# Functional genomics of the parasitic habits of blowflies and botflies

**DOI:** 10.1101/2019.12.13.871723

**Authors:** Gisele Antoniazzi Cardoso, Marina Deszo, Tatiana Teixeira Torres

## Abstract

Infestation by dipterous larvae – myiasis - is a major problem for livestock industries worldwide and can cause severe economic losses. The Oestroidea superfamily is an interesting model to study the evolution of myiasis-causing flies because of the diversity of parasitism strategies among closely-related species. These flies are saprophagous, obligate parasites, or facultative parasites and can be subdivided into cutaneous, nasopharyngeal, traumatic, and furuncular. We expect that closely-related species have genetic differences that lead to the diverse parasitic strategies. To identify genes with such differences, we used gene expression and coding sequence data from five species (*Cochliomyia hominivorax, Chrysomya megacephala, Lucilia cuprina, Dermatobia hominis*, and *Oestrus ovis*). We tested whether 1,287 orthologs have different expression and evolutionary constraints under different scenarios. We found two up-regulated genes; one in species causing cutaneous myiasis that is involved in iron transportation/metabolization (ferritin), and another in species causing traumatic myiasis that responds to reduced oxygen levels (anoxia up-regulated-like). Our evolutionary analysis showed a similar result. In the *Ch. hominivorax* branch, we found one gene with the same function as ferritin that may be evolving under positive selection, spook. This is the first step towards understanding origins and evolution of parasitic strategy diversity in Oestroidea.

## Introduction

Myiasis is a common infestation of vertebrates by dipterous larvae, which feed on living or dead tissue^1^. These species can be defined as obligatory or facultative parasites. The obligate parasites need to feed on a living host to complete the larval stage of their life cycle (primary myiasis), while facultative parasites mostly feed on decomposing organic matter (saprophagous), or even necrotic tissue from a living host (secondary myiasis) and are opportunist parasites^1^.

Many of these parasites are Calyptrate flies, with many belonging to Oestroidea: Calliphoridae (blowflies) and Oestridae (botflies). This group has a wide diversity of contrasting feeding habits that may be exhibited in the adult or larval stage, with parasitism mostly being exhibited during the latter. One of the most interesting features of Oestroidea is the diversity of parasitism strategies among closely related species. Obligatory parasitism, in particular, has evolved on several independent occasions in this group^2,3^. Understanding the different mechanisms that underpin these life strategies can be the key to unlocking the evolution of parasitism.

To begin to understand the evolution of this group, we used RNA sequence (RNA-seq) data of four myiasis-causing and one saprophagous species. The most widely used classification for myiasis was proposed by Zumpt^1^, however, this author did not consider that species could be strictly saprophagous. In the present study we used three classifications: a) obligate parasitism, b) facultative parasitism, and c) saprophagous.

The first obligate parasite chosen for this study was *D. hominis* (Diptera: Oestridae), which parasitizes warm-blooded animals (mostly domestic animals) causing furuncular myiasis. These larvae are capable of entering injured or intact skin (commonly hair follicles) and migrating to the subcutaneous layer where they feed on soft tissue and exudates^4–6^. An important feature of this species is that there is only one larva per infestation site. Another fly of the same family, *O. ovis*, is host-specific and parasitizes only sheep. Females lay first instar larvae in the nose of the host, which then migrate to the inner parts of the paranasal sinuses and feed on the mucous membranes and mucus^7^. Several larvae of this species can be found in the same infestation site.

*Co. hominivorax* (Diptera: Calliphoridae) is another obligate parasite, which commonly infests domestic animals. The larvae of this species are not capable of initiating an infestation, so females lay their eggs in open wounds or mucous membranes. A single wound can be infested by hundreds of larvae at the same time^8^. Infestations by obligate parasites represent a major problem for livestock industries worldwide. There is a loss of productivity and increased management costs, which can be substantial; Grisi and colleagues^9^ estimated a cost of US$720 million annually for managing myiasis-causing Diptera in Brazil.

*Ch. megacephala* (Diptera: Calliphoridae) is a saprophagous species that feeds exclusively on decaying organic matter^8^._Flies of this species can be found in garbage dumps, carcasses, sewers, as well as food displayed in open markets and markets, making them important epidemiologically vectors^10^. They are also important in Forensic Entomology, a science that uses insects as evidence to solve cases of interest to the law, often related to crimes^11^. Lastly, *L. cuprina* (Diptera: Calliphoridae) is a facultative parasite, which is responsible for large economic losses in Australia by causing cutaneous myiasis in sheep. Larvae stay deep in the dermis of the sheep, feeding on tissue fluids, dermal tissue, and blood^12^. They are able to parasitize the host by excreting digestive proteases, as their mouth hooks are poorly developed^13^.

Molecular and genomic information regarding these species are still mostly lacking. Only *L. cuprina* has a well-annotated genome^14^, and some transcriptomic information of Calliphoridae species can be found in public databases such as SRA (Sequence Read Archive). We generated RNA-seq data for *O. ovis* and *D. hominis* and, for *Co. hominivorax* and *Ch. megacephala*, we used public sequences to assemble the transcriptome of all species (excluding *L. cuprina* for which predictions of coding sequences are already available), and used statistical and evolutionary analyses to identify genes that may be involved in parasitism. For *D. hominis* and *O. ovis*, this is the first study to generate large scale transcriptomic information.

## Results

We obtained two biological replicates for each species by *de novo* sequencing or from public resources. On average, more than 56 million reads were obtained per species (Table 1). After quality trimming, we discarded 20% of sequences for *Co. hominivorax*, and less than 3% for the other species. The trimmed reads were used as input for the assemblies.

**Table 1.** RNA-seq data.

| Species | Replicate | Raw reads | Trimmed reads | Contigs | Average length (bp) | CDS |
| --- | --- | --- | --- | --- | --- | --- |
| *Co. hominivorax* | 1 | 13,028,851 | 11,055,926 | 38,749 | 965 | 15,247 |
|  | 2 | 26,807,116 | 26,277,392 |  |  |  |
| *Ch. megacephala* | 1 | 19,549,331 | 19,386,665 | 68,516 | 974 | 18,466 |
|  | 2 | 35,112,471 | 34,108,071 |  |  |  |
| *D. hominis* | 1 | 46,794,071 | 45,397,596 | 166,823 | 1042 | 33,040 |
|  | 2 | 30,006,392 | 29,273,934 |  |  |  |
| *L. cuprina* | 1 | 28,386,962 | 27,827,093 | -- | -- | 18,543 |
|  | 2 | 19,439,280 | 18,987,105 |  |  |  |
| *O. ovis* | 1 | 33,799,262 | 32,827,955 | 103,386 | 1098 | 15,906 |
|  | 2 | 30,643,772 | 29,882,997 |  |  |  |

*Dermatobia hominis and O. ovis* transcriptomes had the highest number of contigs, more than 100,000 (Table 1) with an average length of 1,042 and 1,098 bp, respectively. For *Co. hominivorax*, 38,749 contigs were assembled with an average size of 965 bp. According to our quality evaluation, our assemblies had at least 72% of complete, and less than 10% of fragmented, dipteran orthologs (Figure 1). Furthermore, a large number of duplicated orthologs was obtained for *D. hominis* and *O. ovis*. This resulted from the presence of many alternative isoforms in their transcriptomes (data not shown). From this dataset, we searched for the shared predicted coding sequences (CDS) among the five species with a pipeline that uses a taxonomic grouping method. We identified 1,275 orthologs among the five species.

**Figure 1.**
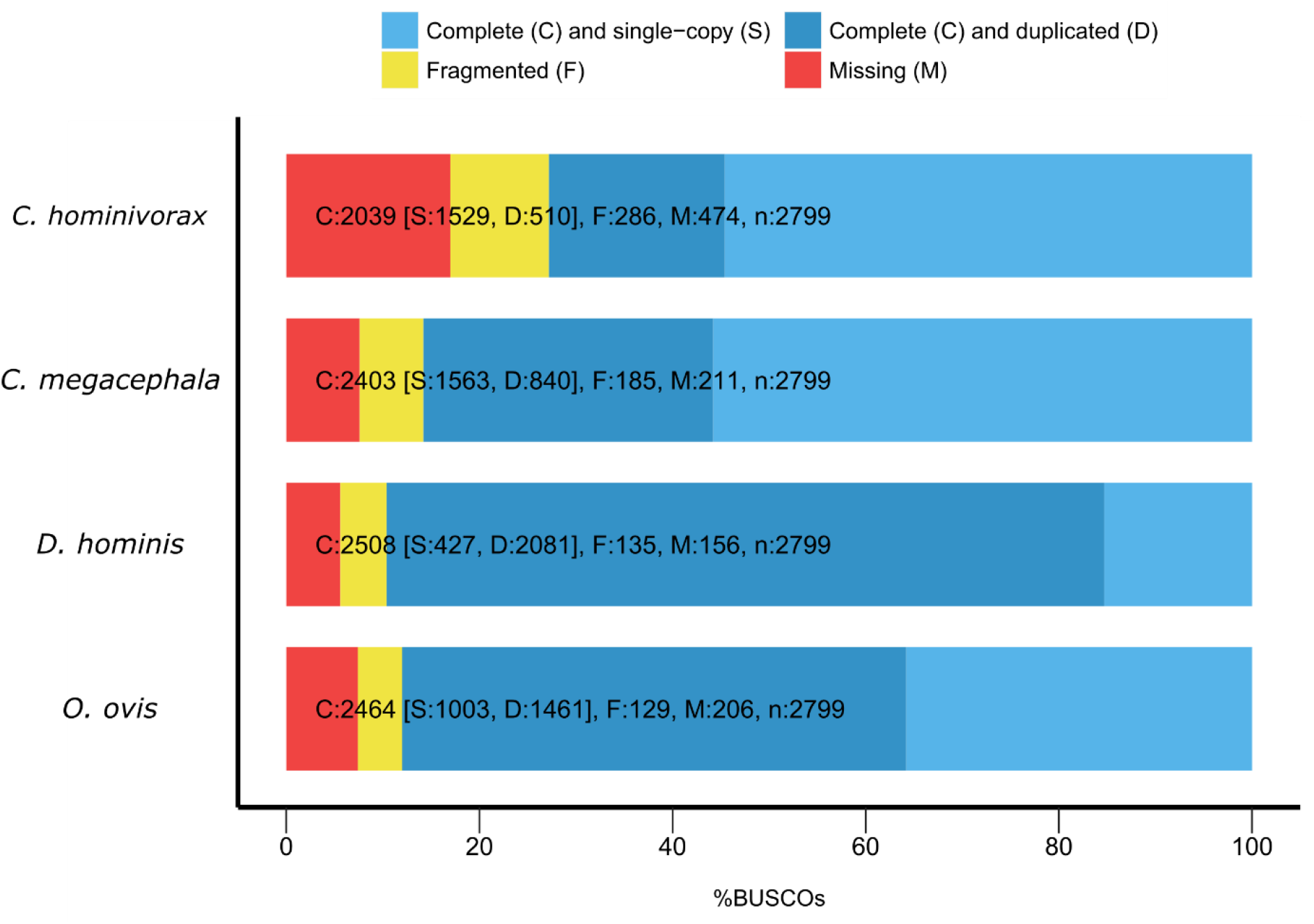
Assembly completeness. The evaluation was performed for the species in which the transcriptome assembly was performed. Bars are the values and proportions of aligned orthologs in each category. n: total number of orthologs in Diptera.

### Gene expression analysis

Reads mapped against the orthologs were counted and used as expression values for each sample in the comparisons among contrasting feeding habits in different scenarios. We used different methods to compute these counts and to infer differently expressed (DE) genes: a) eXpress + edgeR; b) eXpress + DESeq2; c) RSEM + edgeR; and d) RSEM + DESeq2 (Supplementary Tables S1-S4). The intersection of these results was used as the set of DE genes, i.e. only genes that were deemed DE by the four methods were kept for further analyses (Figure 2). Genes identified with all approaches are less dependent on the intrinsic differences of each method. This seems to be a good starting point for the identification of candidate genes, with the drawback being an increased number of false negatives.

**Figure 2.**
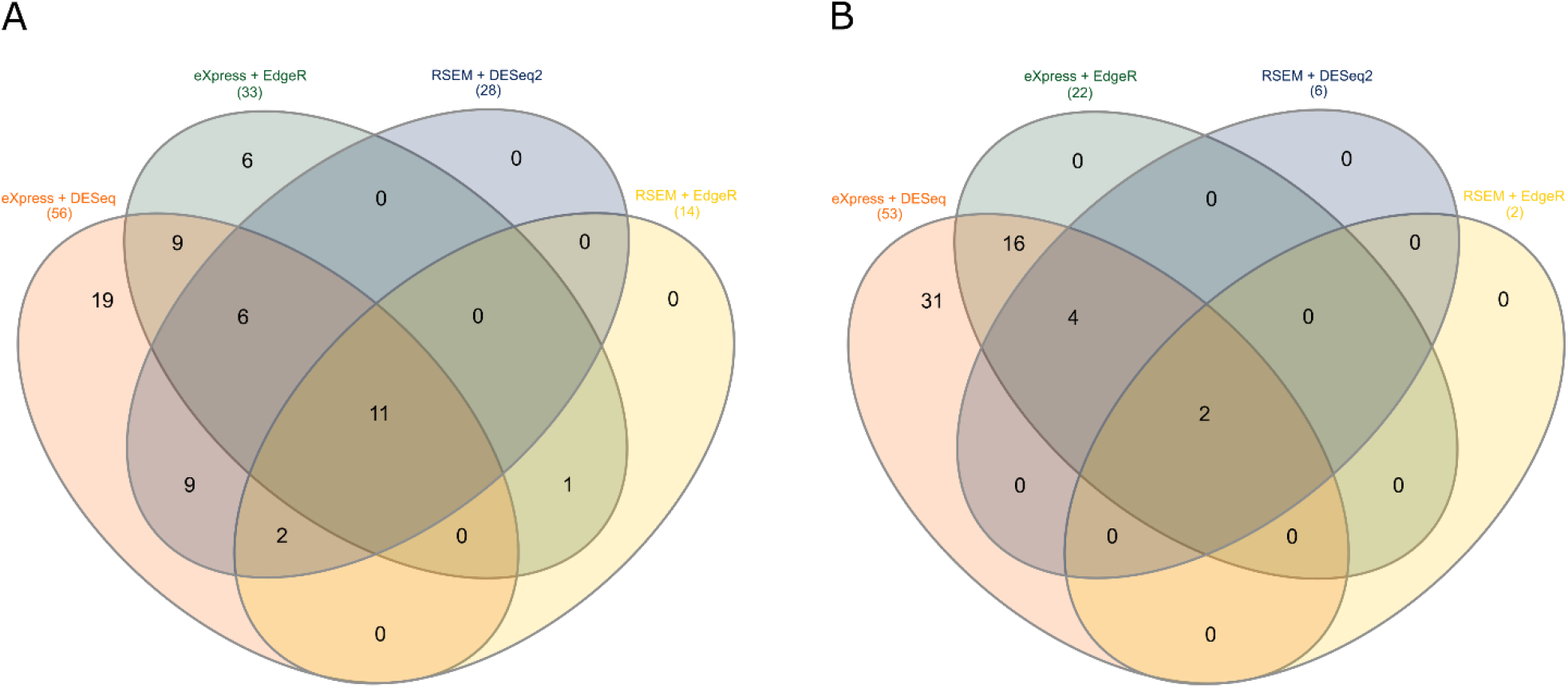
Venn diagrams illustrating the intersection of significant tests in the combination of the different methods. (A) comparison between dermic and nasopharyngeal myiasis. (B) comparison between traumatic/wound and furuncular myiasis. The tests were performed by combining different read counting algorithms (eXpress and RSEM) and the R statistical libraries (edgeR and DESeq2). Only genes with significant differences in all combinations were deemed differentially expressed.Venn diagrams were generated with the InteractiVenn tool^52^.

We tested four different scenarios to search for DE genes that could be linked to the different feeding habits in the species studied. When we compared parasites with saprophagous species and Oestridae with Calliphoridae, we did not find any DE genes. A total of 11 DE genes, however, were detected when we compared cutaneous with nasopharyngeal parasites. Nine genes were up-regulated in cutaneous parasites and two were down-regulated (Figure 2A, Table 2). Almost 45% of the DE genes are ribosomal proteins, the most different among them was the ribosomal protein L8 60S with a mean log2FC = 12.42. Analyzing the function of DE genes, it was possible to identify potential candidates involved in cutaneous parasitism such as the trypsin and ferritin genes. The other hypothesis tested, was that there would be a difference between traumatic and furuncular parasitism. In this test, we found two differently expressed genes; one of them is regulated in hypoxia or in the absence of oxygen (protein anoxia like up-regulated) and the other is a ribosomal protein (Figure 2B, Table 2).

**Table 2.**
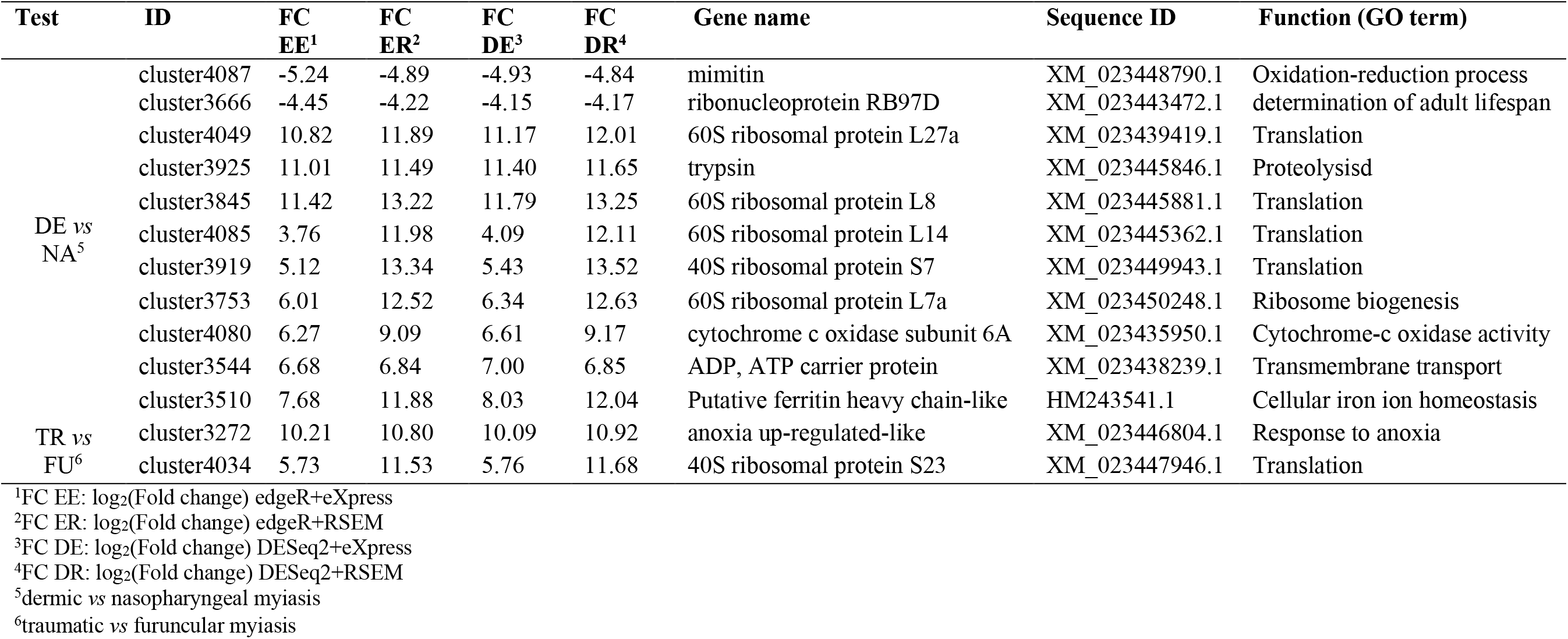
Annotation of the DE genes.

### Evidence of selection in the coding sequences

We searched for evidence of positive selection with codeml using six different models (Supplementary Table S19-S25). We compared the results with the null hypothesis that all sequences were evolving neutrally. In our data, we found ω as high as 999 (a number the program outputs to indicate an infinite number). This ω means that there was a very low rate of non-synonymous substitutions.

In general, there was high constraint of coding sequences in all models tested; at least 80.5% were found to be evolving under a purifying selection regime. Some genes, however, might be evolving under positive selection. When we tested the *Co. hominivorax* branch, we found nine genes under positive selection (Table 3). In agreeance with our expression results, among these genes we found the *spook* gene (*spo*) involved in oxidation-reduction processes and iron binding. For the *D. hominis* branch, we identified four genes with evidence of positive selection; one in common with *Co. hominivorax:* the Samui gene, with a heat shock protein binding function.

**Table 3.**
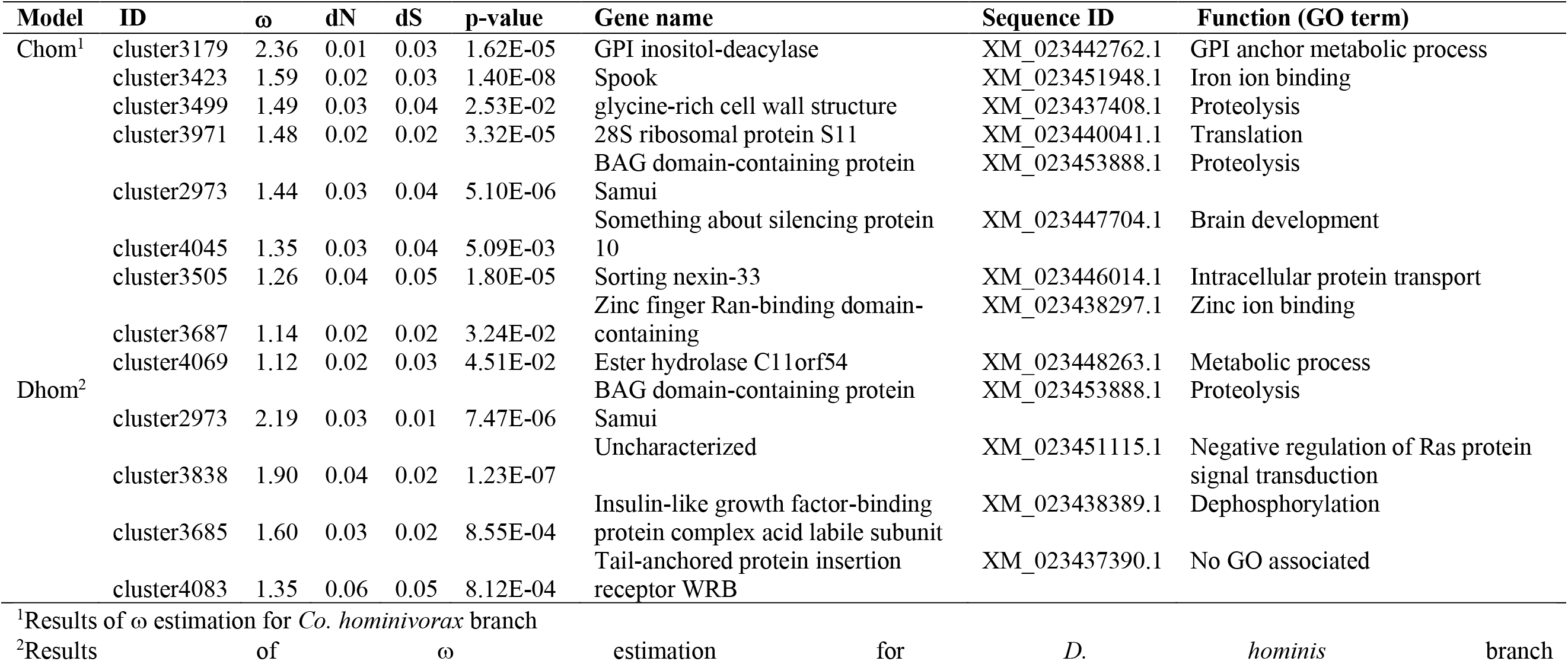
Annotation of positive selected genes.

## Discussion

Herein, we employed a comparative approach to identify DE genes in various scenarios and genes evolving in different selection regimes in specific branches of the cladogram of blowflies (Calliphoridae) and botflies (Oestridae). Understanding the genetic basis of parasitic habit is a crucial step towards identifying the specific genes involved in coding for different parasitism strategies.

Regarding our DE results, we found a great number of ribosomal genes. In the comparison between cutaneous and nasopharyngeal parasites, six ribosomal genes were found to be different. Ribosomal proteins play important roles in all biological processes, translating mRNA molecules into proteins, and it is generally assumed that they have a conserved level of gene expression among species. However, reports on the expression profiles of these genes are scarce, and their expression was compared between different tissues and conditions, but not among different species^15–17^. Also, one previous study investigated the use of ribosomal genes to normalize qPCR data from different Calliphoridae species^18^. The authors showed that a ribosomal gene, *RpS17*, had a dynamic expression among three species in two life stages (larva and adult females). Recent evidence has shown a possible regulatory role for ribosomal proteins, in addition to their constitutive role in translation. Any defects on ribosomal components or losses of central ribosomal proteins may alter the patterns of gene expression^19,20^, which may result in tissue-specific alterations. It was shown that overexpression or silencing of various ribosomal proteins lead to changes in gene transcription, increasing or decreasing gene transcription rates in *Drosophila melanogaster* and *Schistosoma japonicum*, for example^17,21^. Thus, in addition to protein synthesis, dynamic and heterogeneous expression patterns of ribosomal proteins probably play important roles in the regulation of protein translation. The ribosomal genes identified in this study may have a regulatory role on specific transcripts and/or groups of transcripts in Oestroidea species.

The genes adenine nucleotide translocator (ATP/ADP carrier), cytochrome oxidase c subunit 6A (Levy) and ferritin were differently expressed in the comparison of dermic and nasopharyngeal myiasis. All three genes were up-regulated in the dermic parasites. The ATP/ADP carrier and Levy gene products are associated with cellular energy production. The first participates in the regulation of ATP biosynthesis in *D. melanogaster*^22,23^. Mutants of this gene exhibited a reduction in ATP production, lactate accumulation, and changes in the gene expression pattern that are consistent with shifts in ATP production to glycolysis^23^. The second gene, Levy, encodes a subunit of the cytochrome c oxidase complex. *Drosophila melanogaster* mutants for this gene have a decreased rate of assembly of this complex, causing a reduction in lifespan and increase in the neurodegeneration rate due to the decreased production of ATP levels, and higher levels of free radicals^20^. Both genes are involved in the regulation of cellular metabolism in the mitochondrial membrane. Different diets can alter mitochondrial activity^24^. Larvae of *D. melanogaster* reared in diets supplemented with different fatty acids (saturated and unsaturated), for instance, showed differences in cellular metabolism. Individuals fed with a diet supplemented with polyunsaturated fatty acid had higher rate of cellular respiration than those developed in a saturated fatty acid supplementation. This increased activity of the respiratory chain leads to an increase in oxidative stress, decreasing the expression of ADP/ATP carriers^24^. In our study, these genes had a lower level of gene expression in *O. ovis* that feeds on mucus in contrast to those that feed on muscle tissue. This difference may be related to the different diets of the studied species.

The ferritin gene product is widely found among bacteria, plants and animals; and is associated with iron absorption and cellular homeostasis. Ferritin silencing in *D. melanogaster* results in decreased iron uptake and absorption by intestinal cells, and iron deposition in ocular and neuronal cells. This deposition leads to changes in the development of eye and brain tissues, causing malformation of these structures^25^, demonstrating the toxic effects of the iron excess in tissues. To avoid these deleterious effects in a diet rich in iron, usually there is an increase the expression of ferritin gene and protein^26–28^. This mechanism was observed by Georgieva and colleagues^28^ in *D. melanogaster* when individuals fed on media supplement with iron in which increased ferritin mRNA levels was detected in larvae, pupae and imagoes. These previous findings lead us to hypothesize that the up-regulation of ferritin in dermic parasites (*Co. hominivorax*, *D. hominis* and *L. cuprina*) is caused by the high concentrations of iron present in the host blood in which larvae are in constant contact during feeding. This might be the key to the survival of these species in conditions where iron levels are elevated.

In the comparison involving traumatic and furuncular myiasis, we observed an up-regulation of the protein anoxia up-regulated (fau) in the species that causes traumatic wounds (*Co. hominivorax* and *L. cuprina*). Fau is up-regulated in response to low concentrations of oxygen. This response was observed in adults of *D. melanogaster* that were exposed to concentrations of oxygen smaller than 0.02% (anoxia condition)^29^. The up-regulation of fau in our work may be a result of the different parasitism strategies of these species. In both cases, the infestation site can contain hundreds of larvae compared to furuncular myiasis in which there is only one larva per infestation. Because of the large number of larvae, there is an aggregation behavior where the individuals form a larval mass. It has been reported for saprophagous calliphorids the individuals in larval mass go through periods of hypoxia^30^. Thus, species that form traumatic wounds are more susceptible to low concentrations of O_2_.

Besides working with a limited number of genes, we observed that most of them had a conserved (not statistically different; fold change < 2, and/or p > 0.05) expression profile among the scenarios tested. We could assume that this conservation is the result of a constraint in the regulatory regions. Also, we found a similar result for coding sequences in which most of sequences are evolving under negative selection. We detected evidence of positive selection in only nine and four genes in the branch of *Co. hominivorax and D. hominis*, respectively. Among these, we found one gene, spook (spo) on *Co. hominivorax* branch with a similar function to ferritin observed in the differential expression analysis. The spook gene is involved in oxidation and iron binding processes, and it is part of the Halloween family of developmental genes. Spook encodes an enzyme of the P450 cytochrome family that catalyzes the hydroxylation steps in the conversion of cholesterol to 20-hydroxyecysone^31^. It is possible that this gene presents evidence of positive selection in *Co. hominivorax* due to selection in a phenotype that is not necessarily related to the feeding habit. However, automatic annotation of this gene indicates similarity to iron binding proteins (InterPro: IPR001128, InterPro: IPR002401), a biological function also observed in our DE analysis.

The present study is the first to compare different expression profiles of parasites to attempt to understand the underlying mechanisms of parasitism. Genes related to hypoxia and iron transportation/metabolization may play an important role in parasite survival during traumatic myiasis (*Co. hominivorax* and *L. cuprina*), where we observe the aggregation of a large maggot mass, and in the presence of high levels of iron (*Co. hominivorax*, *L. cuprina*, and *D. hominis*). In future studies, we aim to expand the number of samples examined to improve the transcriptomes and find a greater number of orthologs to reveal ther genes involved in mechanisms associated with parasitism.

## Materials and Methods

### Public resources

For *Co. hominivorax, L. cuprina*, and *Ch. megacephala*, we used public RNA-seq data (Table 4). Samples of *Ch. megacephala* and *L. cuprina* consisted of a mix of larvae at different life stages (first instar to pre-pupae), whereas samples of *Co. hominivorax* consisted of feeding third instar larvae.

**Table 4.** Accession numbers of public data.

| **Species** | **NCBI accession** | **Publication** |
| --- | --- | --- |
| *Ch. megacephala* | SRR620248 and SRR1663113 | 49,50 |
| *Co. hominivorax* | SRP044071 | 51 |
| *L. cuprina* | SRR1853100 | 14 |

### RNA isolation and RNA-seq

Live third instar larvae of *D. hominis* and *O. ovis* were collected directly from infested animals. For *D. hominis*, the sampling was carried out manually at cattle breeding farms (Espírito Santo do Pinhal and Guarulhos, both in São Paulo, Brazil). *Oestrus ovis* larvae were donated by Professor Alessandro Francisco Talamini do Amarante (State University of Sao Paulo, Botucatu, Brazil). Larvae were sampled from euthanized sheep. For *L. cuprina*, RNA-seq data from larval samples was publicly available for only one experiment. Thus, we sequenced one sample of third instar larvae to generate a biological replicate. Females of *L. cuprina* were collected in São Paulo, Brazil, with an entomological net using decomposing beef liver as bait. Females were kept in a laboratory colony until the next generation of larvae according to Cardoso *et al* (2016). All samples were conserved at −80°C until RNA extraction was carried out.

Total RNA was extracted with the TRIzol reagent (Thermo Fisher Scientific, USA) according to the manufacturer’s instructions. Samples were quantified with Qubit 2.0 using the Qubit RNA BR kit protocol (Thermo Fischer Scientific, USA). The RNA-seq was outsourced to the sequencing facility at Laboratório de Biotecnologia, ESALQ). The TruSeq mRNA protocol (Illumina, San Diego, USA) was used to prepare the libraries. They were sequenced using HiSeq 2500 equipment using the HiSeq SBS v3 kit (Illumina, San Diego, USA), with paired 100 bp reads. RNA-seq libraries were prepared and sequenced for two individuals of each species.

### Processing of reads and transcriptome assembly

Raw reads were trimmed with trimmomatic version 0.32^32^ using a sliding window of four bases with a 15 base quality cut off. The options “leading” and “trailing” were set to three. After quality trimming, we used an in-house Perl script to maintain only one copy of each duplicated read pair. With this set, the transcriptomes were assembled with the default options of Trinity, version 2.4.0^33^. As the genome of *L. cuprina* is already fully sequenced and publicly available, we used the predicted CDS instead of assembling a transcriptome. The assembled transcriptomes were evaluated for completeness with BUSCO version 3.0^34^ against a Diptera ortholog database.

### Ortholog search

Identification of orthologous sequences among species was based on the protocol developed by Yang and Smith^35^. We used all steps described in the bitbucket repository (https://bitbucket.org/yangya/phylogenomic_dataset_construction) to retain only 1-to-1 orthologs. In brief, this pipeline consists of six steps: 1) predicting functional candidate contigs (CDS) with TransDecoder (https://github.com/TransDecoder/TransDecoder/wiki); 2) removing redundancy of the datasets with cd-hit-est^36^; 3) performing all-by-all blast; 4) filtering blast results by hit fraction length (0.4); 5) Markov-clustering and; 6) retaining 1-to-1 orthologs based on homolog trees.

For each species we retained a set of CDS identified as having an ortholog in all other species. The orthologs were trimmed to the shortest length, ensuring that all comparisons were carried out with same-length sequences.

### Gene expression analysis

We used different methods to analyze gene expression for the purpose of increasing the robustness of our analysis and therefore confidence in our results. To obtain expression counts, we used eXpress version 1.5.0^37^ and RSEM version 1.3.0^38^. The first allows gapped alignments, while the latter does not. Hence, we ran Bowtie2 version 2.2.6^39,40^ twice, with different parameters to map the trimmed reads (not collapsed) to the set of ortholog CDS of each species. For gapped mapping we used: local, very-sensitive-local, maxins 1000, and no-discordant. For ungapped alignments we used: dpad 0, gbar 99999999, mp 1.1, np 1 and, score-min L, 0 −0.1, no-mixed, and no-discordant.

The resulting matrix of expression levels was used as an input to find DE genes with edgeR^41^ and DESeq2^42^, both developed for the R environment. For both analyses, only comparisons with an adjusted p-value < 0.05, and a fold change > 2 were considered DE. Only genes deemed DE in all four different combinations with different read counting and statistical methods were considered for further analyses.

We performed four tests comparing: a) parasites with saprophagous; b) Oestridae species with Calliphoridae species (excluding *Ch. megacephala*); c) dermic myiasis (*Ch. hominivorax*, *D. hominis*, and *L. cuprina*) with nasopharyngeal myiasis (*O. ovis*) and; d) traumatic/wound myiasis (*Co. hominivorax* and *L. cuprina*) with furuncular myiasis (*D. hominis*). DE genes were manually annotated using BLAST^43^.

### Coding sequence analysis

In addition to changes in gene expression, changes in coding regions of orthologous sequences were studied in an attempt to identify changes associated with different food preferences. To achieve this, we estimated the ω (d_N_/d_S_ or K_a_/K_s_) to infer the non-synonymous to synonymous substitution ratio.

For this step, we used the CDS and amino acid sequences predicted previously with TransDecoder for the species with no publicly available genome. First, we aligned the amino acid sequences with muscle^44^. Then the alignment was back-translated to obtain a nucleotide alignment with translatorX^45^. This program also uses Gblocks^46^ to mask poorly and/or very divergent aligned regions.

These alignments were used to estimate the values of d_N_, d_S_, and ω between sequences of orthologs with codeml in PAML version 4 package^47^. We tested six models: (a) a neutral model where ω= 1 (null hypothesis); (b) an average value of ω for the whole tree (model 1); (c) an average ω for *Co. hominivorax* branch and an average value for the other species (model 2); (d) an average value of ω for the branch of dermic parasites (*Co. hominivorax, D. hominis*, *and L. cuprina*) and a different average value of ω for *O. ovis* and *Ch. megacephala* (model 3); (e) an average ω for all parasites and a different ω for the saprophagous species, *Ch. megacephala* (model 4); an ω for *D. hominis* and an average ω for the other species (model 5) and lastly; an ω for *O. ovis* and an average ω for the other species (model 6).

As proposed by Villanueva-Cañas^48^, we discarded branches with d_S_ < 0.01 and d_S_ and d_N_ > 2, as these values can mislead d_N_/d_S_ estimates. Also, we discarded genes with d_N_/d_S_ > 10. The likelihood estimates for each model were compared with the likelihood of our null hypothesis with a χ^2^ test with α set at 0.05. The false discovery rate (FDR) was used for multiple test corrections. For the significant tests we assume that: (i) ω= 1 neutral evolution, (ii) ω < 1 purifying selection and (iii) ω>1 positive selection. Genes with evidences of positive selection were manually annotated using BLAST^43^.

## Data availability statement

All sequences of *D. hominis, O. ovis* and *L. cuprina* are available in the SRA database with the BioSample accession numbers: SAMN11520382, SAMN11520383 and SAMN11655705.

## Acknowledgements

We especially thank Professor Alessandro Francisco Talamini do Amarante, PhD (UNESP, Botucatu campus) for providing *O. ovis* samples. This work was supported by grants to TTT from Fundação de Amparo à Pesquisa do Estado de São Paulo (FAPESP, grant 2014/13933-8 and 2016/09659-3). GAC and MSD were supported by a fellowship from FAPESP (2014/01600-4 and 2012/23200-2, respectively). This study was financed in part by the Coordenação de Aperfeiçoamento de Pessoal de Nível Superior - Brasil (CAPES) - Finance Code 001.

## Authors contributions

MSD carried out all expression analysis including annotation. MSD and GAC performed all evolutionary inference. GAC and TTT wrote the manuscript. TTT supervised the project and coordinated all activities. All authors read and approved the final manuscript.

