## Supplementary information for "Functional genomics of the parasitic habits of blowflies and botflies"

**Supplementary Table 1.** Raw counts eXpress.

| ID | Chom1[1] | Chom2[1] | Dhom1[2] | Dhom2[2] | Oovi1[3] | Oovi2[3] | Lcup1[4] | Lcup2[4] | Cmeg1[5] | Cmeg2[5] |
|---|---|---|---|---|---|---|---|---|---|---|
| cluster1127 | 97 | 287 | 1287 | 9 | 214 | 193 | 262 | 72 | 347 | 384 |
| cluster2853 | 600 | 1381 | 3257 | 6754 | 1939 | 2297 | 3939 | 1164 | 3099 | 7057 |
| cluster2854 | 514 | 1343 | 3982 | 561 | 5122 | 982 | 338 | 184 | 1461 | 2197 |
| cluster2855 | 846 | 1995 | 5075 | 456 | 4374 | 4759 | 2633 | 709 | 3678 | 6455 |
| cluster2856 | 580 | 857 | 3952 | 42 | 5202 | 4534 | 4251 | 454 | 4611 | 3500 |
| cluster2857 | 40624 | 127618 | 221818 | 106992 | 274461 | 271633 | 264094 | 217824 | 246814 | 372542 |
| cluster2858 | 390 | 945 | 1695 | 644 | 785 | 708 | 966 | 110 | 3650 | 2321 |
| cluster2859 | 181 | 447 | 1203 | 216 | 347 | 864 | 561 | 356 | 531 | 993 |
| cluster2860 | 365 | 780 | 3707 | 4215 | 2116 | 1871 | 2365 | 406 | 1181 | 2263 |
| cluster2861 | 238 | 582 | 1372 | 1336 | 1023 | 1445 | 265 | 78 | 1204 | 2755 |
| cluster2862 | 242 | 624 | 471 | 777 | 391 | 496 | 977 | 268 | 675 | 1463 |
| cluster2863 | 289 | 720 | 2538 | 834 | 969 | 2190 | 1124 | 262 | 769 | 1403 |
| cluster2864 | 185 | 384 | 469 | 529 | 1343 | 1195 | 947 | 147 | 721 | 1368 |
| cluster2865 | 268 | 661 | 427 | 799 | 1455 | 2230 | 1080 | 204 | 1111 | 1470 |
| cluster2866 | 99 | 305 | 5074 | 32 | 450 | 646 | 1450 | 225 | 1307 | 2018 |
| cluster2867 | 298 | 680 | 1489 | 1 | 147 | 154 | 330 | 165 | 165 | 410 |
| cluster2869 | 149 | 378 | 875 | 672 | 46 | 53 | 584 | 167 | 645 | 841 |
| cluster2870 | 147 | 401 | 3169 | 97 | 647 | 1084 | 919 | 204 | 820 | 1231 |
| cluster2871 | 100 | 282 | 659 | 147 | 605 | 954 | 1286 | 292 | 925 | 1433 |
| cluster2872 | 405 | 1008 | 3874 | 182 | 1311 | 976 | 823 | 515 | 2235 | 4312 |
| cluster2873 | 174 | 433 | 848 | 1220 | 682 | 632 | 618 | 202 | 576 | 1421 |
| cluster2874 | 308 | 679 | 2061 | 231 | 1610 | 1406 | 1077 | 305 | 1555 | 2585 |
| cluster2875 | 480 | 1079 | 2936 | 4410 | 3398 | 4006 | 7914 | 832 | 4538 | 6902 |
| cluster2876 | 714 | 2058 | 156 | 230 | 14792 | 279 | 1438 | 890 | 801 | 2500 |
| cluster2877 | 296 | 796 | 1468 | 582 | 1994 | 1851 | 896 | 368 | 1753 | 2833 |
| cluster2878 | 225 | 406 | 2861 | 963 | 420 | 1773 | 1836 | 63 | 4296 | 2091 |
| cluster2879 | 5133 | 16178 | 24708 | 5107 | 10635 | 8741 | 6439 | 1920 | 12959 | 20736 |
| cluster2880 | 687 | 1751 | 7625 | 113 | 1246 | 1719 | 1360 | 573 | 2410 | 4283 |
| cluster2881 | 396 | 790 | 1219 | 2505 | 3068 | 2904 | 1003 | 258 | 2122 | 2952 |

| | | | | | | | | | | |
|---|---|---|---|---|---|---|---|---|---|---|
| cluster2882 | 101 | 238 | 693 | 316 | 341 | 513 | 351 | 70 | 416 | 976 |
| cluster2883 | 823 | 2240 | 2063 | 2638 | 2640 | 4717 | 2667 | 1921 | 2614 | 7201 |
| cluster2884 | 333 | 886 | 3981 | 323 | 1243 | 966 | 869 | 232 | 1292 | 1708 |
| cluster2885 | 151 | 364 | 2051 | 13 | 428 | 503 | 776 | 203 | 2230 | 2564 |
| cluster2886 | 260 | 671 | 607 | 1222 | 589 | 844 | 1817 | 128 | 635 | 1511 |
| cluster2887 | 199 | 504 | 830 | 38 | 1688 | 1883 | 948 | 401 | 573 | 1053 |
| cluster2888 | 123 | 356 | 160 | 0 | 116 | 1745 | 262 | 454 | 412 | 270 |
| cluster2889 | 157 | 411 | 1211 | 58 | 599 | 938 | 392 | 192 | 633 | 866 |
| cluster2890 | 306 | 840 | 3504 | 1195 | 799 | 891 | 2728 | 640 | 2154 | 3727 |
| cluster2891 | 506 | 1049 | 4989 | 15 | 1425 | 1253 | 2928 | 887 | 1578 | 3681 |
| cluster2892 | 178 | 505 | 468 | 538 | 1589 | 2213 | 707 | 215 | 481 | 1278 |
| cluster2893 | 1626 | 2794 | 486 | 354 | 7972 | 318 | 1342 | 392 | 320 | 1977 |
| cluster2894 | 1106 | 2524 | 3473 | 234 | 186 | 68 | 2077 | 377 | 3801 | 11039 |
| cluster2895 | 108 | 278 | 1677 | 35 | 224 | 339 | 687 | 185 | 653 | 864 |
| cluster2896 | 432 | 989 | 1793 | 205 | 847 | 1291 | 585 | 330 | 1127 | 1412 |
| cluster2897 | 126 | 294 | 1783 | 47 | 440 | 706 | 1096 | 95 | 1059 | 1195 |
| cluster2898 | 51 | 195 | 105 | 181 | 366 | 151 | 283 | 11 | 179 | 324 |
| cluster2899 | 239 | 791 | 1477 | 2277 | 315 | 379 | 739 | 189 | 916 | 1913 |
| cluster2900 | 82 | 231 | 1040 | 999 | 111 | 194 | 1034 | 166 | 704 | 1090 |
| cluster2901 | 202 | 612 | 1487 | 16 | 357 | 367 | 1852 | 205 | 2150 | 1567 |
| cluster2902 | 84 | 215 | 4433 | 18 | 883 | 1871 | 264 | 91 | 110 | 352 |
| cluster2903 | 89 | 239 | 662 | 942 | 392 | 542 | 3085 | 82 | 769 | 1437 |
| cluster2904 | 345 | 694 | 4853 | 753 | 170 | 218 | 759 | 119 | 4746 | 2230 |
| cluster2905 | 336 | 920 | 2878 | 2628 | 1418 | 1592 | 2322 | 495 | 4496 | 5356 |
| cluster2906 | 218 | 572 | 801 | 1725 | 791 | 970 | 1215 | 199 | 3073 | 2067 |
| cluster2907 | 155 | 434 | 2574 | 65 | 605 | 692 | 1323 | 258 | 874 | 1393 |
| cluster2908 | 229 | 686 | 2776 | 530 | 242 | 443 | 527 | 102 | 1676 | 2439 |
| cluster2909 | 64 | 160 | 638 | 191 | 392 | 548 | 416 | 127 | 393 | 547 |
| cluster2910 | 230 | 698 | 2995 | 150 | 838 | 971 | 2071 | 312 | 1680 | 2769 |
| cluster2911 | 2428 | 6631 | 5950 | 1588 | 4235 | 13774 | 20647 | 3962 | 8924 | 15325 |
| cluster2912 | 212 | 392 | 1732 | 443 | 2544 | 3127 | 956 | 176 | 550 | 1035 |

| | | | | | | | | | | |
|---|---|---|---|---|---|---|---|---|---|---|
| cluster2913 | 277 | 793 | 2588 | 240 | 491 | 669 | 1367 | 342 | 541 | 1687 |
| cluster2914 | 4104 | 9328 | 5457 | 981 | 3189 | 1234 | 4554 | 1315 | 1732 | 11572 |
| cluster2915 | 81 | 237 | 632 | 40 | 50 | 95 | 530 | 68 | 212 | 261 |
| cluster2916 | 78 | 189 | 174 | 419 | 403 | 163 | 332 | 22 | 292 | 619 |
| cluster2917 | 319 | 812 | 1644 | 3 | 262 | 1574 | 565 | 199 | 867 | 1179 |
| cluster2918 | 144 | 388 | 1202 | 59 | 203 | 231 | 338 | 98 | 824 | 1109 |
| cluster2919 | 117 | 411 | 399 | 606 | 760 | 595 | 743 | 117 | 1117 | 1661 |
| cluster2920 | 144 | 403 | 2417 | 86 | 470 | 445 | 1107 | 211 | 825 | 1316 |
| cluster2921 | 138 | 420 | 3379 | 306 | 757 | 765 | 296 | 66 | 910 | 1647 |
| cluster2922 | 178 | 555 | 2568 | 38 | 2844 | 2388 | 2041 | 318 | 1868 | 2345 |
| cluster2923 | 280 | 831 | 4844 | 1059 | 1536 | 1882 | 2401 | 398 | 2822 | 4781 |
| cluster2924 | 116 | 254 | 3247 | 282 | 267 | 3940 | 856 | 185 | 2537 | 6348 |
| cluster2925 | 81 | 244 | 1522 | 49 | 63 | 96 | 262 | 89 | 1113 | 1690 |
| cluster2926 | 222 | 665 | 441 | 1113 | 919 | 1360 | 740 | 178 | 175 | 1180 |
| cluster2927 | 85 | 207 | 789 | 1 | 487 | 996 | 239 | 122 | 277 | 524 |
| cluster2928 | 41 | 90 | 325 | 38 | 716 | 741 | 178 | 133 | 144 | 228 |
| cluster2929 | 214 | 462 | 2281 | 224 | 850 | 755 | 1001 | 166 | 644 | 1308 |
| cluster2930 | 105 | 291 | 2352 | 206 | 1243 | 2499 | 578 | 207 | 814 | 1097 |
| cluster2931 | 124 | 300 | 1275 | 12 | 255 | 340 | 602 | 156 | 687 | 742 |
| cluster2932 | 103 | 279 | 1075 | 64 | 304 | 362 | 959 | 46 | 302 | 479 |
| cluster2933 | 67 | 185 | 775 | 1032 | 1064 | 371 | 843 | 80 | 1862 | 2078 |
| cluster2934 | 129 | 261 | 740 | 113 | 208 | 407 | 674 | 73 | 1003 | 1104 |
| cluster2935 | 732 | 1618 | 1611 | 2720 | 2132 | 1410 | 1102 | 754 | 1086 | 2450 |
| cluster2936 | 141 | 376 | 645 | 0 | 662 | 1149 | 193 | 87 | 239 | 671 |
| cluster2937 | 1367 | 2624 | 2063 | 326 | 1243 | 1562 | 1344 | 421 | 2373 | 5109 |
| cluster2938 | 319 | 741 | 962 | 840 | 738 | 903 | 1016 | 601 | 609 | 1997 |
| cluster2939 | 189 | 348 | 208 | 118 | 338 | 264 | 362 | 102 | 379 | 558 |
| cluster2940 | 471 | 1306 | 2703 | 66 | 1028 | 1196 | 2156 | 662 | 1171 | 3135 |
| cluster2941 | 226 | 852 | 2879 | 543 | 2186 | 2979 | 2069 | 1288 | 3152 | 4118 |
| cluster2942 | 109 | 249 | 1526 | 68 | 1012 | 809 | 389 | 88 | 748 | 1050 |
| cluster2943 | 425 | 1035 | 2482 | 2535 | 2830 | 3320 | 940 | 144 | 3086 | 4072 |

| | | | | | | | | | | |
|---|---|---|---|---|---|---|---|---|---|---|
| cluster2944 | 50 | 161 | 1532 | 40 | 293 | 390 | 934 | 119 | 497 | 821 |
| cluster2945 | 1060 | 2514 | 9412 | 0 | 2016 | 2026 | 4031 | 1192 | 2718 | 4515 |
| cluster2946 | 734 | 2010 | 8096 | 9109 | 5711 | 6964 | 6686 | 1402 | 9598 | 18708 |
| cluster2947 | 42 | 117 | 799 | 208 | 497 | 558 | 267 | 35 | 302 | 439 |
| cluster2948 | 42 | 147 | 883 | 1826 | 423 | 455 | 583 | 101 | 1044 | 1135 |
| cluster2949 | 150 | 505 | 566 | 490 | 747 | 793 | 412 | 211 | 1482 | 1114 |
| cluster2950 | 74 | 262 | 556 | 480 | 363 | 447 | 634 | 112 | 639 | 1341 |
| cluster2951 | 118 | 327 | 803 | 4 | 1064 | 829 | 657 | 196 | 809 | 1170 |
| cluster2952 | 85 | 214 | 918 | 7 | 574 | 514 | 842 | 84 | 592 | 941 |
| cluster2953 | 80 | 209 | 708 | 279 | 270 | 207 | 580 | 113 | 577 | 1129 |
| cluster2954 | 1389 | 2929 | 3370 | 6328 | 1544 | 1779 | 5226 | 1792 | 5795 | 12609 |
| cluster2955 | 274 | 687 | 2723 | 332 | 372 | 1151 | 897 | 343 | 1250 | 1228 |
| cluster2956 | 89 | 239 | 409 | 216 | 508 | 670 | 1591 | 83 | 1023 | 1308 |
| cluster2957 | 59 | 185 | 520 | 6 | 85 | 135 | 641 | 22 | 565 | 840 |
| cluster2958 | 149 | 325 | 1327 | 347 | 563 | 500 | 800 | 243 | 615 | 1178 |
| cluster2959 | 134 | 361 | 189 | 268 | 855 | 828 | 708 | 166 | 1088 | 1938 |
| cluster2960 | 115 | 279 | 673 | 1714 | 279 | 480 | 528 | 149 | 579 | 1195 |
| cluster2961 | 183 | 516 | 592 | 1322 | 890 | 1126 | 447 | 137 | 219 | 439 |
| cluster2962 | 54 | 182 | 254 | 632 | 175 | 473 | 190 | 69 | 87 | 333 |
| cluster2963 | 612 | 1245 | 1523 | 3485 | 2265 | 507 | 1369 | 449 | 2906 | 2642 |
| cluster2964 | 208 | 456 | 790 | 1040 | 1213 | 1947 | 909 | 333 | 1395 | 2949 |
| cluster2965 | 173 | 342 | 4148 | 342 | 890 | 961 | 786 | 180 | 1593 | 2434 |
| cluster2966 | 168 | 463 | 1572 | 16 | 2447 | 1888 | 1122 | 382 | 1456 | 2128 |
| cluster2967 | 62 | 156 | 281 | 613 | 897 | 1040 | 440 | 30 | 1063 | 594 |
| cluster2968 | 250 | 600 | 757 | 521 | 741 | 1358 | 1147 | 231 | 1136 | 2110 |
| cluster2969 | 95 | 381 | 1855 | 41 | 1777 | 789 | 721 | 45 | 494 | 978 |
| cluster2970 | 214 | 476 | 2139 | 326 | 1125 | 1244 | 1073 | 334 | 676 | 1640 |
| cluster2971 | 262 | 782 | 1766 | 926 | 1214 | 1383 | 1627 | 913 | 1282 | 2423 |
| cluster2972 | 1257 | 2606 | 1669 | 3075 | 1263 | 1158 | 3639 | 3763 | 2275 | 10047 |
| cluster2973 | 351 | 915 | 3551 | 22 | 2141 | 2180 | 1241 | 514 | 3641 | 3350 |
| cluster2974 | 132 | 411 | 1080 | 748 | 92 | 331 | 1608 | 183 | 990 | 1975 |

| | | | | | | | | | | |
|---|---|---|---|---|---|---|---|---|---|---|
| cluster2975 | 413 | 1174 | 2408 | 361 | 2954 | 2010 | 997 | 511 | 1531 | 2456 |
| cluster2976 | 327 | 767 | 248 | 122 | 766 | 1599 | 1314 | 298 | 818 | 1319 |
| cluster2977 | 90 | 281 | 1609 | 333 | 842 | 1277 | 600 | 76 | 1505 | 1638 |
| cluster2978 | 247 | 577 | 2351 | 2856 | 1055 | 1276 | 2185 | 369 | 1279 | 3623 |
| cluster2979 | 139 | 419 | 2206 | 135 | 744 | 1005 | 917 | 190 | 646 | 1120 |
| cluster2980 | 452 | 1158 | 355 | 4 | 108 | 163 | 422 | 591 | 322 | 1561 |
| cluster2981 | 80 | 281 | 1633 | 63 | 575 | 789 | 400 | 167 | 354 | 447 |
| cluster2982 | 143 | 495 | 2959 | 223 | 186 | 668 | 998 | 80 | 1896 | 2116 |
| cluster2983 | 208 | 494 | 1275 | 69 | 667 | 1004 | 378 | 212 | 745 | 1509 |
| cluster2984 | 252 | 533 | 1380 | 2292 | 573 | 762 | 452 | 363 | 1065 | 1409 |
| cluster2985 | 184 | 309 | 153 | 371 | 552 | 790 | 680 | 60 | 540 | 521 |
| cluster2986 | 3301 | 8044 | 18123 | 1181 | 19354 | 8812 | 10144 | 4259 | 11180 | 19192 |
| cluster2987 | 90 | 250 | 448 | 188 | 835 | 850 | 375 | 186 | 854 | 1116 |
| cluster2988 | 89 | 206 | 925 | 61 | 533 | 502 | 394 | 125 | 231 | 499 |
| cluster2989 | 36 | 111 | 284 | 391 | 212 | 337 | 245 | 20 | 496 | 429 |
| cluster2990 | 81 | 272 | 412 | 68 | 237 | 1051 | 306 | 171 | 721 | 740 |
| cluster2991 | 54 | 150 | 293 | 22 | 136 | 306 | 130 | 103 | 211 | 389 |
| cluster2992 | 118 | 277 | 1513 | 457 | 200 | 236 | 1426 | 246 | 1185 | 1365 |
| cluster2993 | 164 | 396 | 504 | 1330 | 726 | 1053 | 347 | 87 | 501 | 1399 |
| cluster2994 | 52 | 208 | 1469 | 142 | 528 | 1035 | 658 | 98 | 178 | 247 |
| cluster2995 | 5 | 20 | 93 | 13 | 51 | 169 | 37 | 11 | 12 | 58 |
| cluster2996 | 126 | 282 | 513 | 203 | 323 | 676 | 857 | 25 | 270 | 1276 |
| cluster2997 | 2295 | 5252 | 12819 | 431 | 4403 | 5444 | 6397 | 1728 | 9908 | 16073 |
| cluster2998 | 344 | 811 | 1261 | 419 | 865 | 789 | 1393 | 807 | 435 | 2489 |
| cluster2999 | 517 | 1131 | 1676 | 18 | 1786 | 1629 | 2357 | 1144 | 1308 | 2062 |
| cluster3000 | 16279 | 33621 | 40035 | 2622 | 17588 | 20422 | 82464 | 43758 | 42785 | 106805 |
| cluster3001 | 113 | 301 | 1093 | 476 | 819 | 1029 | 1845 | 291 | 813 | 1878 |
| cluster3002 | 89 | 223 | 1234 | 264 | 318 | 739 | 785 | 92 | 937 | 814 |
| cluster3003 | 114 | 301 | 1638 | 45 | 95 | 80 | 387 | 99 | 522 | 811 |
| cluster3004 | 284 | 841 | 843 | 782 | 396 | 623 | 528 | 111 | 3888 | 3755 |
| cluster3005 | 332 | 1038 | 4468 | 502 | 883 | 1723 | 623 | 403 | 3138 | 4573 |

| | | | | | | | | | | |
|---|---|---|---|---|---|---|---|---|---|---|
| cluster3006 | 299 | 654 | 12839 | 15484 | 4511 | 4337 | 3116 | 558 | 7808 | 6099 |
| cluster3007 | 67 | 226 | 683 | 18 | 292 | 545 | 248 | 106 | 197 | 524 |
| cluster3008 | 61 | 207 | 627 | 46 | 846 | 1445 | 256 | 128 | 153 | 248 |
| cluster3009 | 616 | 1270 | 3026 | 174 | 299 | 429 | 1512 | 1580 | 771 | 2086 |
| cluster3010 | 158 | 507 | 1365 | 46 | 170 | 210 | 810 | 173 | 810 | 1136 |
| cluster3011 | 227 | 708 | 436 | 1819 | 4127 | 1073 | 1763 | 581 | 1084 | 2192 |
| cluster3012 | 104 | 201 | 65 | 4 | 508 | 1429 | 274 | 145 | 106 | 390 |
| cluster3013 | 289 | 840 | 1121 | 1287 | 565 | 2761 | 810 | 301 | 1805 | 2446 |
| cluster3014 | 347 | 862 | 495 | 21 | 259 | 423 | 244 | 104 | 308 | 1454 |
| cluster3015 | 63 | 193 | 1597 | 126 | 110 | 108 | 75 | 20 | 837 | 704 |
| cluster3016 | 106 | 242 | 359 | 0 | 70 | 27 | 230 | 52 | 327 | 360 |
| cluster3017 | 56 | 163 | 3132 | 217 | 103 | 232 | 423 | 107 | 361 | 386 |
| cluster3018 | 222 | 560 | 2233 | 97 | 778 | 1316 | 877 | 380 | 632 | 1288 |
| cluster3019 | 128 | 364 | 3050 | 138 | 992 | 1325 | 568 | 185 | 385 | 454 |
| cluster3020 | 1377 | 3611 | 16713 | 277 | 5305 | 5502 | 6181 | 2927 | 10952 | 15421 |
| cluster3021 | 197 | 401 | 806 | 601 | 490 | 862 | 616 | 136 | 543 | 1115 |
| cluster3022 | 93 | 213 | 456 | 4 | 181 | 247 | 427 | 57 | 446 | 863 |
| cluster3023 | 101 | 280 | 649 | 814 | 523 | 773 | 446 | 130 | 407 | 783 |
| cluster3024 | 67 | 308 | 866 | 531 | 105 | 88 | 613 | 106 | 1958 | 1570 |
| cluster3025 | 222 | 647 | 2591 | 807 | 646 | 1332 | 1487 | 319 | 863 | 1741 |
| cluster3026 | 42 | 79 | 183 | 5 | 68 | 71 | 373 | 17 | 860 | 516 |
| cluster3027 | 145 | 489 | 809 | 1233 | 827 | 1405 | 860 | 262 | 761 | 1734 |
| cluster3028 | 76 | 246 | 648 | 56 | 469 | 677 | 644 | 77 | 447 | 871 |
| cluster3029 | 128 | 269 | 1195 | 137 | 1042 | 1195 | 1117 | 273 | 1009 | 1674 |
| cluster3030 | 135 | 355 | 1736 | 3131 | 407 | 855 | 329 | 121 | 2148 | 3054 |
| cluster3031 | 154 | 318 | 289 | 635 | 559 | 414 | 607 | 167 | 1288 | 1073 |
| cluster3032 | 92 | 203 | 661 | 716 | 554 | 642 | 414 | 118 | 912 | 1419 |
| cluster3033 | 1759 | 5410 | 9030 | 2051 | 3235 | 7983 | 8071 | 2909 | 5369 | 6347 |
| cluster3034 | 64 | 213 | 427 | 27 | 407 | 256 | 746 | 100 | 260 | 328 |
| cluster3035 | 526 | 1366 | 3762 | 1148 | 828 | 695 | 1438 | 409 | 2213 | 3412 |
| cluster3036 | 75 | 194 | 1019 | 26 | 564 | 507 | 1027 | 169 | 860 | 1173 |

| | | | | | | | | | | |
|---|---|---|---|---|---|---|---|---|---|---|
| cluster3037 | 600 | 1281 | 2906 | 152 | 1468 | 723 | 984 | 432 | 5260 | 2149 |
| cluster3038 | 57 | 176 | 765 | 9 | 497 | 1114 | 919 | 124 | 1126 | 989 |
| cluster3039 | 117 | 342 | 543 | 548 | 1066 | 1238 | 1059 | 201 | 688 | 1102 |
| cluster3040 | 65 | 170 | 253 | 293 | 102 | 136 | 316 | 180 | 457 | 694 |
| cluster3041 | 40 | 120 | 298 | 15 | 105 | 127 | 148 | 29 | 75 | 76 |
| cluster3042 | 49 | 146 | 239 | 8 | 336 | 505 | 589 | 52 | 105 | 330 |
| cluster3043 | 27 | 57 | 283 | 63 | 57 | 115 | 181 | 40 | 99 | 175 |
| cluster3044 | 58 | 147 | 919 | 2 | 79 | 696 | 420 | 85 | 436 | 735 |
| cluster3045 | 46 | 144 | 387 | 235 | 651 | 906 | 265 | 20 | 568 | 528 |
| cluster3046 | 65 | 185 | 1234 | 121 | 270 | 312 | 672 | 42 | 1214 | 993 |
| cluster3047 | 2909 | 7244 | 6869 | 6066 | 11527 | 732 | 7061 | 8523 | 14672 | 17440 |
| cluster3048 | 170 | 519 | 699 | 53 | 502 | 594 | 523 | 298 | 290 | 816 |
| cluster3049 | 218 | 510 | 1072 | 1775 | 1116 | 1704 | 1098 | 591 | 1368 | 2888 |
| cluster3050 | 100 | 319 | 1075 | 124 | 114 | 407 | 747 | 98 | 609 | 1210 |
| cluster3051 | 48 | 173 | 217 | 2 | 460 | 475 | 858 | 162 | 163 | 1126 |
| cluster3052 | 76 | 178 | 2780 | 39 | 72 | 108 | 2746 | 140 | 1480 | 1301 |
| cluster3053 | 155 | 371 | 348 | 1248 | 116 | 857 | 232 | 71 | 345 | 762 |
| cluster3054 | 453 | 1499 | 4781 | 543 | 1638 | 2573 | 1807 | 1215 | 2465 | 4576 |
| cluster3055 | 142 | 362 | 608 | 308 | 599 | 619 | 590 | 87 | 759 | 960 |
| cluster3056 | 75 | 228 | 1129 | 230 | 267 | 241 | 872 | 106 | 699 | 605 |
| cluster3057 | 150 | 385 | 226 | 291 | 17 | 132 | 385 | 182 | 548 | 539 |
| cluster3058 | 156 | 351 | 1041 | 336 | 704 | 814 | 1103 | 153 | 545 | 1107 |
| cluster3059 | 63 | 136 | 1342 | 3 | 459 | 657 | 1067 | 79 | 481 | 223 |
| cluster3060 | 266 | 711 | 160 | 5 | 638 | 923 | 333 | 152 | 131 | 557 |
| cluster3061 | 130 | 376 | 1202 | 1577 | 955 | 1119 | 758 | 333 | 634 | 1173 |
| cluster3062 | 56 | 133 | 239 | 68 | 269 | 99 | 543 | 107 | 148 | 463 |
| cluster3063 | 42 | 103 | 456 | 622 | 857 | 1919 | 232 | 151 | 233 | 526 |
| cluster3064 | 55 | 114 | 300 | 77 | 84 | 317 | 85 | 3 | 395 | 420 |
| cluster3065 | 174 | 560 | 1524 | 240 | 151 | 314 | 217 | 146 | 1075 | 1701 |
| cluster3066 | 146 | 524 | 839 | 1507 | 1863 | 1902 | 1065 | 381 | 1262 | 2711 |
| cluster3067 | 132 | 296 | 382 | 502 | 431 | 924 | 397 | 236 | 657 | 1542 |

| | | | | | | | | | | |
|---|---|---|---|---|---|---|---|---|---|---|
| cluster3068 | 140 | 399 | 487 | 229 | 340 | 261 | 730 | 182 | 715 | 1453 |
| cluster3069 | 385 | 1293 | 3987 | 1290 | 944 | 1504 | 1659 | 973 | 2718 | 4371 |
| cluster3070 | 201 | 527 | 743 | 669 | 726 | 1370 | 1278 | 46 | 1744 | 2555 |
| cluster3071 | 107 | 281 | 529 | 821 | 595 | 790 | 725 | 204 | 1211 | 1417 |
| cluster3072 | 794 | 1825 | 9750 | 29 | 1688 | 1897 | 2198 | 598 | 4524 | 4750 |
| cluster3073 | 459 | 1227 | 1633 | 696 | 723 | 1256 | 1086 | 887 | 1413 | 3090 |
| cluster3074 | 712 | 1617 | 5178 | 276 | 1120 | 1177 | 1397 | 759 | 1841 | 3376 |
| cluster3075 | 77 | 205 | 96 | 199 | 172 | 164 | 569 | 273 | 128 | 385 |
| cluster3076 | 22 | 36 | 143 | 14 | 170 | 349 | 66 | 31 | 64 | 157 |
| cluster3077 | 3027 | 6694 | 6727 | 11459 | 5579 | 7019 | 1770 | 2910 | 2548 | 6282 |
| cluster3078 | 124 | 278 | 666 | 100 | 586 | 551 | 298 | 143 | 177 | 536 |
| cluster3079 | 39 | 105 | 178 | 307 | 190 | 187 | 300 | 109 | 155 | 354 |
| cluster3080 | 41 | 138 | 844 | 99 | 136 | 183 | 690 | 76 | 337 | 634 |
| cluster3081 | 626 | 1594 | 900 | 1185 | 714 | 681 | 1521 | 1677 | 660 | 2950 |
| cluster3082 | 384 | 939 | 684 | 0 | 162 | 312 | 1812 | 90 | 281 | 1775 |
| cluster3083 | 85 | 189 | 476 | 300 | 783 | 720 | 397 | 103 | 294 | 559 |
| cluster3084 | 191 | 596 | 2210 | 702 | 406 | 644 | 1940 | 403 | 1378 | 2096 |
| cluster3085 | 52 | 120 | 407 | 2 | 97 | 154 | 557 | 99 | 305 | 667 |
| cluster3086 | 157 | 424 | 1766 | 155 | 243 | 265 | 709 | 143 | 547 | 1095 |
| cluster3087 | 68 | 152 | 485 | 40 | 150 | 645 | 246 | 68 | 204 | 369 |
| cluster3088 | 53 | 131 | 1566 | 52 | 274 | 345 | 377 | 31 | 207 | 326 |
| cluster3089 | 1817 | 4535 | 5904 | 9582 | 9148 | 6914 | 4374 | 1958 | 3797 | 7472 |
| cluster3090 | 132 | 347 | 4115 | 1 | 37 | 288 | 315 | 373 | 1734 | 4127 |
| cluster3091 | 461 | 1269 | 1457 | 1224 | 1875 | 3120 | 2570 | 1272 | 2077 | 5234 |
| cluster3092 | 411 | 1103 | 1500 | 838 | 952 | 1447 | 895 | 777 | 934 | 2283 |
| cluster3093 | 1739 | 3732 | 16959 | 1432 | 30764 | 6926 | 18890 | 10658 | 9937 | 31500 |
| cluster3094 | 23 | 97 | 564 | 0 | 508 | 572 | 244 | 98 | 134 | 356 |
| cluster3095 | 141 | 380 | 265 | 437 | 205 | 390 | 1091 | 653 | 363 | 847 |
| cluster3096 | 220 | 702 | 2395 | 109 | 833 | 1167 | 1101 | 364 | 551 | 1409 |
| cluster3097 | 46 | 106 | 592 | 327 | 1095 | 1175 | 583 | 64 | 776 | 859 |
| cluster3098 | 153 | 322 | 929 | 86 | 349 | 501 | 491 | 149 | 293 | 445 |

| | | | | | | | | | | |
|---|---|---|---|---|---|---|---|---|---|---|
| cluster3099 | 26 | 59 | 596 | 76 | 300 | 300 | 249 | 58 | 567 | 2070 |
| cluster3100 | 244 | 586 | 1774 | 164 | 273 | 6517 | 282 | 8 | 5697 | 2926 |
| cluster3101 | 133 | 409 | 888 | 1043 | 658 | 1320 | 485 | 151 | 343 | 559 |
| cluster3102 | 36 | 97 | 177 | 438 | 127 | 268 | 161 | 39 | 257 | 550 |
| cluster3103 | 79 | 195 | 787 | 7 | 178 | 9 | 551 | 44 | 29 | 96 |
| cluster3104 | 322 | 758 | 1321 | 313 | 1168 | 873 | 1045 | 383 | 994 | 2333 |
| cluster3105 | 7170 | 14025 | 4095 | 400 | 2352 | 1125 | 5072 | 6740 | 6284 | 33344 |
| cluster3106 | 366 | 1055 | 4970 | 2293 | 3478 | 4853 | 3005 | 1556 | 7596 | 11615 |
| cluster3107 | 72 | 201 | 1313 | 152 | 198 | 379 | 249 | 153 | 267 | 342 |
| cluster3108 | 108 | 175 | 1632 | 23 | 564 | 1166 | 1019 | 111 | 2335 | 2217 |
| cluster3109 | 621 | 1401 | 907 | 80 | 2455 | 3772 | 980 | 271 | 284 | 1675 |
| cluster3110 | 127 | 454 | 2855 | 282 | 1341 | 2425 | 963 | 230 | 1140 | 1787 |
| cluster3111 | 391 | 1260 | 1485 | 322 | 94 | 368 | 1504 | 177 | 5382 | 7468 |
| cluster3112 | 169 | 496 | 522 | 1 | 156 | 1376 | 291 | 333 | 1601 | 2152 |
| cluster3113 | 112 | 260 | 1427 | 174 | 402 | 585 | 211 | 150 | 720 | 1475 |
| cluster3114 | 183 | 427 | 1019 | 121 | 247 | 276 | 462 | 148 | 785 | 1081 |
| cluster3115 | 107 | 322 | 2321 | 204 | 591 | 1014 | 763 | 185 | 1129 | 1197 |
| cluster3116 | 97 | 249 | 1024 | 88 | 634 | 381 | 390 | 207 | 401 | 829 |
| cluster3117 | 53 | 171 | 225 | 0 | 221 | 329 | 58 | 10 | 59 | 63 |
| cluster3118 | 35 | 107 | 982 | 144 | 427 | 588 | 285 | 37 | 1144 | 1213 |
| cluster3119 | 58 | 207 | 300 | 593 | 221 | 339 | 252 | 121 | 254 | 574 |
| cluster3120 | 225 | 600 | 1683 | 449 | 1065 | 1296 | 1159 | 257 | 897 | 2046 |
| cluster3121 | 1674 | 4499 | 2429 | 6991 | 3356 | 3294 | 3100 | 1937 | 11315 | 16203 |
| cluster3122 | 991 | 2506 | 1020 | 1651 | 903 | 1532 | 2080 | 1143 | 1936 | 3784 |
| cluster3123 | 81 | 215 | 322 | 132 | 88 | 288 | 287 | 33 | 680 | 883 |
| cluster3124 | 495 | 1037 | 3378 | 340 | 1307 | 1813 | 2221 | 740 | 2340 | 4513 |
| cluster3125 | 524 | 1535 | 909 | 438 | 1819 | 2120 | 1298 | 956 | 930 | 2326 |
| cluster3126 | 221 | 544 | 1799 | 5166 | 1320 | 1241 | 666 | 548 | 2132 | 3654 |
| cluster3127 | 613 | 2193 | 2060 | 821 | 1106 | 1599 | 3387 | 841 | 1155 | 2678 |
| cluster3128 | 150 | 371 | 976 | 265 | 440 | 439 | 509 | 73 | 1993 | 1365 |
| cluster3129 | 395 | 1091 | 374 | 0 | 198 | 20 | 574 | 449 | 454 | 3956 |

| | | | | | | | | | | |
|---|---|---|---|---|---|---|---|---|---|---|
| cluster3130 | 140 | 371 | 354 | 37 | 172 | 259 | 537 | 399 | 366 | 1099 |
| cluster3131 | 982 | 1886 | 5176 | 183 | 1108 | 976 | 2508 | 1104 | 957 | 2569 |
| cluster3132 | 50 | 125 | 280 | 302 | 305 | 439 | 213 | 119 | 281 | 416 |
| cluster3133 | 43 | 145 | 476 | 25 | 252 | 319 | 448 | 129 | 246 | 710 |
| cluster3134 | 78 | 201 | 286 | 55 | 204 | 260 | 447 | 55 | 301 | 472 |
| cluster3135 | 4310 | 11087 | 24609 | 1014 | 2505 | 6459 | 14359 | 8565 | 27319 | 29280 |
| cluster3136 | 25 | 109 | 670 | 19 | 64 | 89 | 248 | 34 | 98 | 137 |
| cluster3137 | 70 | 264 | 3693 | 35 | 668 | 962 | 85 | 40 | 914 | 1584 |
| cluster3138 | 62 | 157 | 353 | 563 | 328 | 415 | 144 | 48 | 300 | 532 |
| cluster3139 | 77 | 217 | 103 | 0 | 49 | 253 | 43 | 8 | 72 | 83 |
| cluster3140 | 114 | 159 | 699 | 125 | 412 | 626 | 343 | 70 | 454 | 368 |
| cluster3141 | 226 | 582 | 1231 | 1013 | 679 | 1095 | 392 | 241 | 790 | 2092 |
| cluster3142 | 63 | 165 | 1281 | 82 | 50 | 103 | 102 | 22 | 153 | 177 |
| cluster3143 | 50 | 176 | 301 | 226 | 440 | 390 | 400 | 28 | 312 | 460 |
| cluster3144 | 420 | 1110 | 563 | 0 | 421 | 717 | 250 | 442 | 220 | 712 |
| cluster3145 | 49 | 95 | 749 | 227 | 588 | 665 | 933 | 300 | 532 | 1024 |
| cluster3146 | 843 | 2072 | 4425 | 2634 | 675 | 2703 | 3493 | 1428 | 9257 | 15426 |
| cluster3147 | 182 | 516 | 3142 | 764 | 1080 | 1183 | 667 | 424 | 634 | 1741 |
| cluster3148 | 182 | 458 | 1874 | 204 | 861 | 607 | 730 | 214 | 823 | 1456 |
| cluster3149 | 160 | 457 | 849 | 837 | 1066 | 1279 | 829 | 343 | 669 | 1359 |
| cluster3150 | 51 | 103 | 734 | 17 | 542 | 110 | 1757 | 61 | 894 | 614 |
| cluster3151 | 395 | 837 | 3091 | 437 | 589 | 617 | 1370 | 381 | 3249 | 3630 |
| cluster3152 | 41 | 113 | 204 | 0 | 133 | 187 | 139 | 19 | 88 | 165 |
| cluster3153 | 627 | 1641 | 4270 | 796 | 2866 | 6838 | 2373 | 661 | 4532 | 5240 |
| cluster3154 | 59 | 169 | 263 | 173 | 457 | 523 | 655 | 87 | 346 | 525 |
| cluster3155 | 48 | 110 | 542 | 1 | 205 | 366 | 120 | 79 | 285 | 506 |
| cluster3156 | 239 | 551 | 645 | 26 | 614 | 700 | 328 | 573 | 198 | 658 |
| cluster3157 | 140 | 398 | 3580 | 1925 | 493 | 1276 | 1323 | 180 | 1591 | 2680 |
| cluster3158 | 182 | 472 | 52 | 60 | 97 | 1537 | 283 | 142 | 256 | 1651 |
| cluster3159 | 93 | 246 | 629 | 70 | 282 | 457 | 235 | 202 | 401 | 890 |
| cluster3160 | 988 | 3359 | 5943 | 1013 | 696 | 1430 | 2906 | 1793 | 779 | 2467 |

| | | | | | | | | | | |
|---|---|---|---|---|---|---|---|---|---|---|
| cluster3161 | 100 | 248 | 1140 | 99 | 109 | 196 | 822 | 356 | 475 | 1094 |
| cluster3162 | 23 | 69 | 197 | 13 | 266 | 575 | 189 | 17 | 173 | 104 |
| cluster3163 | 655 | 1389 | 3367 | 1 | 970 | 884 | 484 | 605 | 382 | 1595 |
| cluster3164 | 187 | 386 | 1127 | 0 | 405 | 529 | 1006 | 596 | 1812 | 2570 |
| cluster3165 | 30 | 125 | 1312 | 8 | 179 | 198 | 125 | 32 | 157 | 303 |
| cluster3166 | 261 | 612 | 941 | 1106 | 704 | 1192 | 607 | 392 | 455 | 920 |
| cluster3167 | 51 | 160 | 649 | 0 | 342 | 280 | 648 | 44 | 1029 | 1030 |
| cluster3168 | 77 | 205 | 254 | 313 | 747 | 914 | 206 | 78 | 256 | 750 |
| cluster3169 | 148 | 414 | 1734 | 11 | 624 | 1776 | 435 | 344 | 1159 | 1003 |
| cluster3170 | 38 | 115 | 501 | 207 | 246 | 310 | 500 | 26 | 390 | 676 |
| cluster3171 | 72 | 175 | 354 | 626 | 117 | 152 | 603 | 127 | 1005 | 1012 |
| cluster3172 | 178 | 515 | 990 | 1224 | 424 | 248 | 2070 | 345 | 981 | 1685 |
| cluster3173 | 60 | 140 | 533 | 0 | 201 | 211 | 188 | 35 | 73 | 224 |
| cluster3174 | 112 | 293 | 1929 | 9 | 185 | 989 | 358 | 48 | 620 | 1029 |
| cluster3175 | 358 | 921 | 292 | 9 | 327 | 152 | 712 | 316 | 95 | 337 |
| cluster3176 | 153 | 394 | 700 | 186 | 1273 | 964 | 5737 | 114 | 2638 | 1817 |
| cluster3177 | 19082 | 47751 | 16643 | 5738 | 12951 | 15995 | 6912 | 64270 | 10007 | 23535 |
| cluster3178 | 39 | 102 | 531 | 1530 | 1479 | 1616 | 276 | 91 | 329 | 666 |
| cluster3179 | 30 | 97 | 447 | 3 | 497 | 1026 | 175 | 53 | 58 | 151 |
| cluster3180 | 89 | 253 | 776 | 99 | 201 | 185 | 411 | 322 | 371 | 631 |
| cluster3181 | 59 | 202 | 664 | 690 | 406 | 589 | 598 | 135 | 476 | 796 |
| cluster3182 | 16116 | 38526 | 14272 | 4262 | 20234 | 8774 | 15392 | 22714 | 4349 | 22990 |
| cluster3183 | 47 | 191 | 666 | 39 | 84 | 670 | 245 | 264 | 253 | 629 |
| cluster3184 | 72 | 199 | 630 | 26 | 284 | 344 | 355 | 92 | 345 | 467 |
| cluster3185 | 1509 | 3213 | 1172 | 159 | 1228 | 754 | 740 | 1463 | 287 | 2767 |
| cluster3186 | 931 | 3080 | 995 | 64 | 1695 | 2681 | 589 | 1121 | 225 | 2226 |
| cluster3187 | 267 | 651 | 540 | 1 | 45 | 178 | 366 | 103 | 725 | 2616 |
| cluster3188 | 76 | 217 | 260 | 814 | 405 | 301 | 537 | 61 | 516 | 594 |
| cluster3189 | 31 | 126 | 566 | 5 | 77 | 121 | 186 | 59 | 265 | 524 |
| cluster3190 | 42 | 107 | 128 | 0 | 332 | 177 | 129 | 4 | 61 | 258 |
| cluster3191 | 286 | 933 | 41 | 47 | 517 | 4 | 1078 | 107 | 29 | 362 |

| | | | | | | | | | | |
|---|---|---|---|---|---|---|---|---|---|---|
| cluster3192 | 167 | 473 | 2099 | 587 | 273 | 431 | 509 | 480 | 172 | 1082 |
| cluster3193 | 375 | 961 | 3859 | 171 | 422 | 771 | 933 | 411 | 1622 | 2371 |
| cluster3194 | 121 | 240 | 889 | 122 | 166 | 233 | 1307 | 47 | 2877 | 2012 |
| cluster3195 | 191 | 603 | 418 | 624 | 653 | 896 | 2366 | 543 | 2663 | 5855 |
| cluster3196 | 186 | 610 | 1247 | 736 | 486 | 574 | 1282 | 380 | 1575 | 1888 |
| cluster3197 | 163 | 331 | 1541 | 1151 | 661 | 411 | 472 | 93 | 1536 | 1258 |
| cluster3198 | 207 | 548 | 1219 | 17 | 1360 | 1531 | 1069 | 443 | 346 | 1020 |
| cluster3199 | 325 | 919 | 2166 | 697 | 603 | 908 | 675 | 407 | 1792 | 2626 |
| cluster3200 | 84 | 271 | 773 | 979 | 1185 | 1089 | 556 | 146 | 415 | 922 |
| cluster3201 | 258 | 497 | 4058 | 767 | 2519 | 2387 | 3403 | 512 | 2765 | 3915 |
| cluster3202 | 49 | 96 | 981 | 163 | 636 | 643 | 727 | 108 | 535 | 612 |
| cluster3203 | 23 | 78 | 204 | 44 | 49 | 73 | 117 | 91 | 73 | 130 |
| cluster3204 | 90 | 213 | 383 | 131 | 148 | 143 | 47 | 30 | 298 | 282 |
| cluster3205 | 50 | 120 | 154 | 104 | 68 | 97 | 191 | 80 | 199 | 377 |
| cluster3206 | 45 | 100 | 155 | 1 | 110 | 399 | 119 | 98 | 117 | 430 |
| cluster3207 | 81 | 212 | 212 | 444 | 1505 | 2045 | 1221 | 174 | 272 | 722 |
| cluster3208 | 811 | 2308 | 3464 | 3469 | 3068 | 3333 | 4049 | 1537 | 4381 | 9062 |
| cluster3209 | 340 | 836 | 1122 | 57 | 243 | 592 | 656 | 165 | 526 | 1241 |
| cluster3210 | 66 | 152 | 333 | 83 | 304 | 473 | 177 | 67 | 243 | 464 |
| cluster3211 | 131 | 371 | 538 | 0 | 507 | 849 | 353 | 106 | 173 | 270 |
| cluster3212 | 2432 | 7023 | 1299 | 3556 | 3633 | 3723 | 4000 | 11658 | 2084 | 4736 |
| cluster3213 | 88 | 226 | 1275 | 901 | 644 | 258 | 2039 | 206 | 360 | 1475 |
| cluster3214 | 46 | 97 | 52 | 82 | 67 | 141 | 78 | 44 | 85 | 180 |
| cluster3215 | 109 | 309 | 1033 | 311 | 481 | 475 | 436 | 174 | 925 | 952 |
| cluster3216 | 38 | 148 | 394 | 7 | 130 | 149 | 111 | 57 | 111 | 283 |
| cluster3217 | 87 | 247 | 356 | 492 | 133 | 275 | 882 | 131 | 623 | 758 |
| cluster3218 | 61 | 249 | 1486 | 2 | 973 | 509 | 515 | 95 | 363 | 492 |
| cluster3219 | 133 | 286 | 431 | 146 | 508 | 716 | 484 | 205 | 209 | 521 |
| cluster3220 | 975 | 2963 | 828 | 978 | 342 | 1775 | 2530 | 1994 | 1544 | 5890 |
| cluster3221 | 103 | 378 | 1040 | 4 | 447 | 527 | 247 | 174 | 476 | 1242 |
| cluster3222 | 266 | 668 | 1095 | 570 | 905 | 1000 | 1317 | 347 | 411 | 1155 |

| | | | | | | | | | | |
|---|---|---|---|---|---|---|---|---|---|---|
| cluster3223 | 42 | 97 | 1025 | 101 | 55 | 75 | 588 | 43 | 253 | 285 |
| cluster3224 | 90 | 238 | 966 | 141 | 613 | 687 | 1650 | 162 | 1371 | 1559 |
| cluster3225 | 63 | 175 | 145 | 0 | 227 | 254 | 175 | 175 | 201 | 518 |
| cluster3226 | 231 | 570 | 462 | 5 | 267 | 267 | 506 | 277 | 273 | 688 |
| cluster3227 | 57 | 170 | 703 | 22 | 96 | 171 | 120 | 53 | 330 | 324 |
| cluster3228 | 129 | 320 | 985 | 19 | 338 | 739 | 226 | 98 | 378 | 766 |
| cluster3229 | 80 | 224 | 481 | 519 | 66 | 106 | 308 | 49 | 468 | 1283 |
| cluster3230 | 38 | 81 | 190 | 277 | 103 | 120 | 423 | 25 | 186 | 599 |
| cluster3231 | 75 | 226 | 6855 | 300 | 535 | 1066 | 2598 | 1109 | 5328 | 20919 |
| cluster3232 | 102 | 237 | 1211 | 28 | 471 | 604 | 293 | 75 | 290 | 769 |
| cluster3233 | 82 | 226 | 2401 | 4321 | 123 | 701 | 118 | 137 | 472 | 612 |
| cluster3234 | 724 | 2107 | 456 | 537 | 985 | 1266 | 996 | 948 | 1323 | 1950 |
| cluster3235 | 187 | 461 | 1374 | 0 | 438 | 900 | 203 | 40 | 33 | 164 |
| cluster3236 | 74 | 209 | 453 | 13 | 402 | 1008 | 148 | 207 | 185 | 271 |
| cluster3237 | 43 | 162 | 2846 | 17 | 180 | 225 | 232 | 15 | 201 | 337 |
| cluster3238 | 105 | 273 | 1005 | 196 | 470 | 501 | 269 | 135 | 448 | 895 |
| cluster3239 | 142 | 442 | 2309 | 743 | 623 | 952 | 958 | 220 | 1316 | 2706 |
| cluster3240 | 37 | 126 | 234 | 453 | 325 | 545 | 269 | 52 | 67 | 229 |
| cluster3241 | 45 | 118 | 459 | 67 | 88 | 257 | 176 | 37 | 270 | 438 |
| cluster3242 | 168 | 501 | 2891 | 64 | 1706 | 2947 | 1104 | 477 | 1182 | 1772 |
| cluster3243 | 462 | 1202 | 2418 | 3255 | 2065 | 1913 | 3859 | 539 | 4789 | 7869 |
| cluster3244 | 49 | 108 | 211 | 241 | 304 | 430 | 179 | 50 | 330 | 708 |
| cluster3245 | 14 | 28 | 356 | 9 | 141 | 107 | 46 | 2 | 188 | 192 |
| cluster3246 | 193 | 478 | 666 | 1156 | 1260 | 1255 | 592 | 161 | 541 | 888 |
| cluster3247 | 83 | 215 | 472 | 68 | 384 | 490 | 107 | 9 | 305 | 456 |
| cluster3248 | 9 | 40 | 445 | 0 | 115 | 96 | 110 | 90 | 74 | 254 |
| cluster3249 | 207 | 563 | 608 | 178 | 365 | 189 | 564 | 147 | 567 | 1630 |
| cluster3250 | 80 | 158 | 740 | 119 | 218 | 4499 | 100 | 53 | 2039 | 3036 |
| cluster3251 | 172 | 474 | 899 | 42 | 450 | 783 | 500 | 234 | 714 | 1385 |
| cluster3252 | 608 | 1544 | 3807 | 1 | 207 | 10548 | 349 | 1284 | 880 | 1138 |
| cluster3253 | 176 | 451 | 657 | 5 | 703 | 868 | 755 | 234 | 625 | 870 |

| | | | | | | | | | | |
|---|---|---|---|---|---|---|---|---|---|---|
| cluster3254 | 348 | 1004 | 329 | 24 | 132 | 119 | 156 | 52 | 260 | 615 |
| cluster3255 | 490 | 1368 | 2043 | 10 | 1680 | 329 | 1017 | 1099 | 2070 | 2899 |
| cluster3256 | 32 | 97 | 271 | 127 | 166 | 181 | 181 | 67 | 71 | 118 |
| cluster3257 | 72 | 209 | 386 | 299 | 542 | 662 | 387 | 141 | 231 | 451 |
| cluster3258 | 440 | 1509 | 8653 | 2158 | 3085 | 3590 | 2759 | 939 | 4185 | 6408 |
| cluster3259 | 293 | 836 | 1017 | 435 | 155 | 893 | 719 | 459 | 1379 | 3633 |
| cluster3260 | 222 | 605 | 312 | 670 | 760 | 707 | 2431 | 558 | 367 | 1115 |
| cluster3261 | 84 | 211 | 574 | 0 | 137 | 171 | 248 | 202 | 292 | 655 |
| cluster3262 | 35 | 73 | 449 | 120 | 95 | 203 | 155 | 69 | 200 | 388 |
| cluster3263 | 39 | 131 | 586 | 87 | 325 | 373 | 194 | 129 | 410 | 900 |
| cluster3264 | 1460 | 4131 | 5946 | 1477 | 2766 | 3951 | 2055 | 2518 | 3192 | 6784 |
| cluster3265 | 177 | 407 | 2515 | 115 | 217 | 396 | 383 | 164 | 337 | 778 |
| cluster3266 | 26 | 92 | 566 | 87 | 247 | 181 | 414 | 49 | 384 | 567 |
| cluster3267 | 43 | 138 | 246 | 238 | 456 | 508 | 298 | 127 | 214 | 763 |
| cluster3268 | 289 | 730 | 764 | 90 | 306 | 1002 | 460 | 344 | 416 | 947 |
| cluster3269 | 454 | 997 | 1145 | 0 | 704 | 796 | 888 | 421 | 1599 | 2764 |
| cluster3270 | 134 | 402 | 604 | 8 | 2151 | 3212 | 1321 | 298 | 619 | 1686 |
| cluster3271 | 269 | 669 | 561 | 466 | 538 | 536 | 530 | 337 | 347 | 1084 |
| cluster3272 | 1795 | 2757 | 0 | 5 | 4183 | 28047 | 8721 | 3612 | 31006 | 15896 |
| cluster3273 | 43 | 84 | 296 | 173 | 353 | 286 | 284 | 65 | 372 | 586 |
| cluster3274 | 19 | 72 | 588 | 8 | 362 | 628 | 473 | 81 | 97 | 221 |
| cluster3275 | 65 | 206 | 340 | 1 | 74 | 43 | 180 | 29 | 65 | 239 |
| cluster3276 | 65 | 217 | 232 | 246 | 352 | 356 | 348 | 65 | 711 | 824 |
| cluster3277 | 19 | 61 | 94 | 1 | 104 | 147 | 59 | 91 | 193 | 165 |
| cluster3278 | 147 | 359 | 2377 | 646 | 1083 | 1136 | 1381 | 528 | 1302 | 1558 |
| cluster3279 | 75 | 176 | 154 | 31 | 581 | 261 | 102 | 81 | 90 | 297 |
| cluster3280 | 107 | 267 | 764 | 141 | 277 | 525 | 130 | 70 | 640 | 937 |
| cluster3281 | 401 | 880 | 980 | 41 | 860 | 1275 | 454 | 115 | 827 | 1595 |
| cluster3282 | 143 | 386 | 624 | 29 | 181 | 245 | 289 | 136 | 537 | 846 |
| cluster3283 | 78 | 198 | 142 | 150 | 281 | 347 | 189 | 85 | 546 | 1061 |
| cluster3284 | 54 | 104 | 451 | 38 | 153 | 189 | 158 | 35 | 169 | 229 |

| | | | | | | | | | | |
|---|---|---|---|---|---|---|---|---|---|---|
| cluster3285 | 52 | 89 | 623 | 76 | 159 | 126 | 447 | 33 | 273 | 328 |
| cluster3286 | 182 | 583 | 2828 | 723 | 838 | 991 | 1476 | 358 | 2432 | 3553 |
| cluster3287 | 17 | 67 | 111 | 48 | 17 | 23 | 35 | 9 | 92 | 148 |
| cluster3288 | 13 | 42 | 161 | 16 | 280 | 402 | 69 | 45 | 140 | 106 |
| cluster3289 | 44 | 129 | 393 | 396 | 320 | 392 | 465 | 137 | 369 | 648 |
| cluster3290 | 110 | 252 | 107 | 119 | 89 | 34 | 928 | 72 | 299 | 761 |
| cluster3291 | 81 | 131 | 165 | 296 | 653 | 293 | 416 | 64 | 181 | 391 |
| cluster3292 | 2291 | 5530 | 17347 | 112 | 1304 | 2066 | 5011 | 6499 | 7486 | 15017 |
| cluster3293 | 56 | 187 | 132 | 145 | 335 | 485 | 271 | 88 | 120 | 386 |
| cluster3294 | 1135 | 2327 | 1375 | 2503 | 2468 | 4021 | 2340 | 1120 | 1203 | 5932 |
| cluster3295 | 143 | 306 | 1130 | 495 | 628 | 873 | 475 | 161 | 414 | 1565 |
| cluster3296 | 2783 | 7867 | 4916 | 270 | 8892 | 7012 | 4843 | 8354 | 1606 | 10650 |
| cluster3297 | 140 | 423 | 1008 | 879 | 478 | 843 | 86 | 16 | 180 | 253 |
| cluster3298 | 85 | 200 | 509 | 310 | 342 | 260 | 1051 | 178 | 342 | 1174 |
| cluster3299 | 87 | 218 | 214 | 6 | 472 | 525 | 155 | 303 | 157 | 207 |
| cluster3300 | 30 | 54 | 290 | 76 | 60 | 276 | 119 | 12 | 200 | 393 |
| cluster3301 | 57 | 191 | 416 | 832 | 314 | 525 | 1304 | 419 | 121 | 298 |
| cluster3302 | 121 | 266 | 501 | 13 | 365 | 393 | 256 | 145 | 768 | 891 |
| cluster3303 | 13 | 40 | 407 | 976 | 292 | 361 | 209 | 42 | 114 | 362 |
| cluster3304 | 374 | 1047 | 3517 | 2856 | 1816 | 3356 | 2229 | 460 | 3120 | 3188 |
| cluster3305 | 78 | 260 | 934 | 14 | 156 | 1720 | 1352 | 144 | 497 | 2712 |
| cluster3306 | 71 | 216 | 141 | 29 | 1515 | 3275 | 72 | 189 | 143 | 497 |
| cluster3307 | 356 | 881 | 2123 | 51 | 689 | 1858 | 606 | 822 | 1447 | 2887 |
| cluster3308 | 112 | 251 | 383 | 437 | 645 | 720 | 902 | 344 | 528 | 1387 |
| cluster3309 | 51 | 129 | 436 | 369 | 313 | 336 | 57 | 21 | 1498 | 827 |
| cluster3310 | 990 | 1726 | 4160 | 1539 | 2368 | 3302 | 4041 | 1209 | 4806 | 8053 |
| cluster3311 | 27 | 81 | 207 | 264 | 394 | 415 | 272 | 53 | 183 | 322 |
| cluster3312 | 71 | 146 | 36 | 0 | 367 | 72 | 344 | 262 | 118 | 485 |
| cluster3313 | 4609 | 12632 | 3330 | 1094 | 7283 | 14949 | 12738 | 9301 | 4084 | 15537 |
| cluster3314 | 27 | 89 | 2029 | 23 | 428 | 510 | 1077 | 103 | 1511 | 1417 |
| cluster3315 | 218 | 647 | 605 | 1007 | 389 | 784 | 422 | 385 | 219 | 714 |

| | | | | | | | | | | |
|---|---|---|---|---|---|---|---|---|---|---|
| cluster3316 | 32 | 96 | 306 | 15 | 440 | 72 | 550 | 34 | 504 | 527 |
| cluster3317 | 41 | 115 | 1120 | 110 | 125 | 141 | 47 | 15 | 263 | 211 |
| cluster3318 | 36 | 96 | 261 | 511 | 107 | 207 | 292 | 135 | 326 | 400 |
| cluster3319 | 351 | 966 | 843 | 22 | 194 | 427 | 656 | 299 | 2384 | 3316 |
| cluster3320 | 42 | 110 | 242 | 11 | 131 | 234 | 560 | 27 | 260 | 187 |
| cluster3321 | 48 | 298 | 1236 | 127 | 280 | 384 | 494 | 81 | 577 | 1016 |
| cluster3322 | 89 | 351 | 352 | 1 | 292 | 906 | 602 | 84 | 311 | 750 |
| cluster3323 | 15 | 58 | 378 | 350 | 180 | 250 | 160 | 11 | 100 | 163 |
| cluster3324 | 125 | 337 | 110 | 150 | 296 | 289 | 202 | 73 | 388 | 554 |
| cluster3325 | 67 | 85 | 662 | 0 | 16 | 9 | 490 | 26 | 80 | 347 |
| cluster3326 | 247 | 855 | 1624 | 543 | 138 | 91 | 522 | 242 | 1303 | 1959 |
| cluster3327 | 44 | 114 | 152 | 253 | 50 | 225 | 187 | 31 | 116 | 359 |
| cluster3328 | 173 | 389 | 585 | 183 | 441 | 484 | 330 | 156 | 880 | 1314 |
| cluster3329 | 24 | 83 | 57 | 0 | 217 | 333 | 474 | 173 | 27 | 97 |
| cluster3330 | 64 | 153 | 1072 | 51 | 879 | 784 | 780 | 105 | 419 | 851 |
| cluster3331 | 101 | 259 | 538 | 75 | 234 | 390 | 664 | 109 | 321 | 937 |
| cluster3332 | 37 | 99 | 583 | 20 | 413 | 527 | 133 | 32 | 209 | 245 |
| cluster3333 | 75 | 199 | 594 | 13 | 223 | 497 | 154 | 138 | 133 | 161 |
| cluster3334 | 15 | 75 | 127 | 61 | 104 | 240 | 100 | 31 | 94 | 160 |
| cluster3335 | 32 | 129 | 97 | 216 | 101 | 348 | 99 | 18 | 152 | 281 |
| cluster3336 | 12 | 91 | 161 | 208 | 123 | 166 | 618 | 53 | 417 | 509 |
| cluster3337 | 36 | 176 | 464 | 493 | 112 | 145 | 213 | 73 | 237 | 238 |
| cluster3338 | 28 | 82 | 158 | 3 | 187 | 282 | 479 | 85 | 121 | 247 |
| cluster3339 | 84 | 209 | 112 | 307 | 179 | 287 | 257 | 116 | 369 | 608 |
| cluster3340 | 214 | 504 | 1285 | 438 | 260 | 762 | 275 | 88 | 551 | 1148 |
| cluster3341 | 95 | 276 | 334 | 17 | 590 | 601 | 1569 | 198 | 570 | 981 |
| cluster3342 | 99 | 347 | 250 | 9 | 889 | 1055 | 403 | 121 | 104 | 385 |
| cluster3343 | 454 | 1249 | 5374 | 622 | 1386 | 3884 | 873 | 1482 | 4162 | 7417 |
| cluster3344 | 17 | 55 | 151 | 132 | 74 | 129 | 381 | 20 | 184 | 212 |
| cluster3345 | 60 | 112 | 214 | 10 | 275 | 400 | 251 | 156 | 141 | 286 |
| cluster3346 | 262 | 656 | 2084 | 3088 | 1042 | 1431 | 784 | 907 | 849 | 1865 |

| | | | | | | | | | | |
|---|---|---|---|---|---|---|---|---|---|---|
| cluster3347 | 69 | 217 | 328 | 213 | 194 | 364 | 235 | 132 | 243 | 445 |
| cluster3348 | 20 | 63 | 476 | 139 | 116 | 180 | 287 | 10 | 399 | 446 |
| cluster3349 | 685 | 1843 | 1600 | 5156 | 15219 | 12514 | 7399 | 2342 | 1692 | 8634 |
| cluster3350 | 181 | 522 | 573 | 20 | 208 | 346 | 279 | 56 | 412 | 1030 |
| cluster3351 | 277 | 899 | 2260 | 280 | 1189 | 1085 | 1061 | 546 | 801 | 1355 |
| cluster3352 | 73 | 237 | 502 | 6 | 149 | 177 | 119 | 30 | 321 | 186 |
| cluster3353 | 54 | 210 | 752 | 132 | 257 | 539 | 301 | 105 | 383 | 613 |
| cluster3354 | 55 | 161 | 1309 | 138 | 506 | 682 | 596 | 209 | 621 | 643 |
| cluster3355 | 47 | 141 | 173 | 7 | 68 | 32 | 117 | 8 | 161 | 226 |
| cluster3356 | 50 | 108 | 80 | 205 | 387 | 467 | 234 | 49 | 53 | 185 |
| cluster3357 | 61 | 240 | 641 | 91 | 634 | 483 | 100 | 53 | 672 | 771 |
| cluster3358 | 24 | 88 | 304 | 386 | 167 | 305 | 469 | 68 | 236 | 458 |
| cluster3359 | 225 | 508 | 803 | 3 | 444 | 474 | 545 | 137 | 126 | 352 |
| cluster3360 | 755 | 2492 | 391 | 452 | 769 | 665 | 677 | 810 | 488 | 2041 |
| cluster3361 | 406 | 883 | 1611 | 64 | 1099 | 1557 | 750 | 1062 | 1050 | 1837 |
| cluster3362 | 865 | 1862 | 2023 | 3931 | 2158 | 3106 | 2356 | 1390 | 1655 | 3317 |
| cluster3363 | 56 | 169 | 380 | 44 | 412 | 566 | 102 | 41 | 131 | 242 |
| cluster3364 | 2390 | 6007 | 15901 | 1250 | 2373 | 9804 | 2445 | 4764 | 2864 | 5807 |
| cluster3365 | 75 | 173 | 575 | 12 | 306 | 1692 | 244 | 55 | 245 | 285 |
| cluster3366 | 39 | 68 | 379 | 0 | 283 | 448 | 89 | 51 | 110 | 145 |
| cluster3367 | 11 | 44 | 66 | 327 | 53 | 101 | 63 | 20 | 62 | 95 |
| cluster3368 | 172 | 558 | 571 | 228 | 386 | 200 | 379 | 232 | 748 | 2049 |
| cluster3369 | 61 | 150 | 1232 | 95 | 475 | 525 | 136 | 109 | 424 | 717 |
| cluster3370 | 43 | 105 | 569 | 11 | 456 | 442 | 534 | 49 | 196 | 400 |
| cluster3371 | 17 | 59 | 4038 | 0 | 3 | 206 | 102 | 41 | 475 | 441 |
| cluster3372 | 457 | 827 | 1620 | 1656 | 1985 | 2406 | 1504 | 769 | 1278 | 2886 |
| cluster3373 | 29 | 71 | 378 | 12 | 206 | 243 | 164 | 32 | 163 | 213 |
| cluster3374 | 51 | 122 | 169 | 1 | 796 | 787 | 328 | 115 | 17 | 36 |
| cluster3375 | 14560 | 48795 | 28872 | 11691 | 31502 | 33458 | 44744 | 36098 | 24749 | 59002 |
| cluster3376 | 196 | 505 | 401 | 434 | 512 | 722 | 609 | 325 | 467 | 944 |
| cluster3377 | 91 | 206 | 118 | 22 | 77 | 129 | 103 | 78 | 241 | 623 |

| | | | | | | | | | | |
|---|---|---|---|---|---|---|---|---|---|---|
| cluster3378 | 34 | 92 | 15 | 115 | 131 | 226 | 165 | 60 | 55 | 138 |
| cluster3379 | 566 | 1393 | 1816 | 407 | 280 | 175 | 726 | 415 | 124 | 356 |
| cluster3380 | 48 | 127 | 539 | 82 | 182 | 355 | 1028 | 135 | 480 | 924 |
| cluster3381 | 753 | 2102 | 1812 | 2406 | 649 | 979 | 959 | 770 | 825 | 4771 |
| cluster3382 | 5372 | 9668 | 25372 | 8468 | 5385 | 8197 | 5802 | 6249 | 8776 | 14275 |
| cluster3383 | 239 | 778 | 881 | 0 | 58 | 73 | 320 | 128 | 281 | 1401 |
| cluster3384 | 517 | 1203 | 571 | 956 | 1845 | 174 | 562 | 413 | 1203 | 2050 |
| cluster3385 | 406 | 1012 | 816 | 10 | 3215 | 616 | 808 | 1237 | 534 | 931 |
| cluster3386 | 624 | 1398 | 660 | 20 | 1545 | 1529 | 657 | 633 | 249 | 904 |
| cluster3387 | 226 | 634 | 863 | 167 | 2216 | 2769 | 1764 | 1048 | 398 | 1752 |
| cluster3388 | 67 | 219 | 537 | 29 | 228 | 530 | 2337 | 128 | 324 | 730 |
| cluster3389 | 224 | 625 | 255 | 80 | 965 | 972 | 289 | 338 | 308 | 1046 |
| cluster3390 | 70 | 61 | 393 | 208 | 326 | 176 | 563 | 68 | 568 | 386 |
| cluster3391 | 88 | 295 | 655 | 662 | 269 | 297 | 357 | 129 | 290 | 493 |
| cluster3392 | 109 | 306 | 64 | 1 | 332 | 226 | 233 | 166 | 19 | 364 |
| cluster3393 | 274 | 683 | 187 | 4 | 91 | 72 | 1556 | 448 | 42 | 896 |
| cluster3394 | 330 | 688 | 537 | 492 | 704 | 1564 | 361 | 614 | 1284 | 2481 |
| cluster3395 | 166 | 482 | 206 | 200 | 109 | 282 | 342 | 290 | 257 | 569 |
| cluster3396 | 27 | 53 | 199 | 227 | 207 | 182 | 528 | 63 | 237 | 340 |
| cluster3397 | 357 | 845 | 988 | 14 | 463 | 613 | 1647 | 1552 | 679 | 6001 |
| cluster3398 | 1441 | 3202 | 6413 | 755 | 1382 | 2252 | 4291 | 2674 | 3101 | 5045 |
| cluster3399 | 12 | 67 | 432 | 242 | 385 | 151 | 416 | 14 | 1004 | 607 |
| cluster3400 | 144 | 364 | 2085 | 689 | 483 | 428 | 287 | 157 | 802 | 2134 |
| cluster3401 | 389 | 1132 | 5482 | 511 | 2879 | 4118 | 1485 | 519 | 2363 | 4677 |
| cluster3402 | 14 | 60 | 439 | 253 | 281 | 368 | 319 | 69 | 732 | 917 |
| cluster3403 | 987 | 2557 | 2899 | 371 | 1799 | 2598 | 1834 | 1072 | 2232 | 5949 |
| cluster3404 | 447 | 1327 | 3471 | 2686 | 1784 | 2305 | 3490 | 1101 | 2137 | 2132 |
| cluster3405 | 83 | 217 | 1357 | 53 | 199 | 225 | 228 | 90 | 727 | 826 |
| cluster3406 | 172 | 496 | 275 | 544 | 842 | 964 | 515 | 655 | 211 | 815 |
| cluster3407 | 191 | 534 | 672 | 1405 | 1068 | 1386 | 1741 | 443 | 657 | 3284 |
| cluster3408 | 28 | 81 | 534 | 109 | 41 | 30 | 99 | 19 | 346 | 517 |

| | | | | | | | | | | |
|---|---|---|---|---|---|---|---|---|---|---|
| cluster3409 | 108 | 294 | 916 | 0 | 540 | 437 | 250 | 237 | 204 | 426 |
| cluster3410 | 51 | 139 | 817 | 1221 | 229 | 349 | 689 | 233 | 256 | 532 |
| cluster3411 | 169 | 443 | 794 | 116 | 1050 | 1173 | 1276 | 421 | 383 | 996 |
| cluster3412 | 41 | 94 | 389 | 10 | 138 | 174 | 263 | 22 | 146 | 140 |
| cluster3413 | 36 | 85 | 746 | 8 | 125 | 68 | 351 | 260 | 59 | 157 |
| cluster3414 | 49 | 233 | 614 | 473 | 178 | 328 | 552 | 84 | 471 | 689 |
| cluster3415 | 64 | 126 | 711 | 27 | 62 | 141 | 195 | 93 | 360 | 361 |
| cluster3416 | 87 | 211 | 37 | 0 | 431 | 568 | 101 | 101 | 114 | 257 |
| cluster3417 | 12 | 29 | 76 | 158 | 76 | 89 | 98 | 34 | 29 | 78 |
| cluster3418 | 41 | 108 | 176 | 10 | 181 | 435 | 94 | 71 | 86 | 278 |
| cluster3419 | 122 | 264 | 1295 | 195 | 159 | 327 | 577 | 136 | 2187 | 2242 |
| cluster3420 | 63 | 226 | 681 | 26 | 181 | 621 | 551 | 46 | 1097 | 810 |
| cluster3421 | 14 | 42 | 1150 | 23 | 393 | 762 | 667 | 40 | 344 | 386 |
| cluster3422 | 17 | 75 | 140 | 262 | 12 | 70 | 436 | 3 | 470 | 309 |
| cluster3423 | 37 | 134 | 184 | 0 | 30 | 195 | 88 | 8 | 64 | 168 |
| cluster3424 | 100 | 283 | 12 | 154 | 46 | 1 | 221 | 48 | 20 | 489 |
| cluster3425 | 94 | 334 | 309 | 37 | 69 | 210 | 718 | 53 | 404 | 703 |
| cluster3426 | 68 | 152 | 452 | 529 | 274 | 265 | 162 | 50 | 383 | 554 |
| cluster3427 | 39 | 51 | 731 | 23 | 79 | 77 | 554 | 21 | 1075 | 573 |
| cluster3428 | 540 | 1311 | 541 | 944 | 964 | 1165 | 1048 | 299 | 5022 | 5562 |
| cluster3429 | 109 | 258 | 430 | 710 | 313 | 580 | 279 | 418 | 417 | 979 |
| cluster3430 | 28 | 100 | 93 | 0 | 446 | 658 | 55 | 11 | 187 | 304 |
| cluster3431 | 44 | 143 | 320 | 34 | 489 | 848 | 126 | 58 | 330 | 842 |
| cluster3432 | 66 | 160 | 168 | 168 | 73 | 101 | 673 | 323 | 227 | 554 |
| cluster3433 | 21 | 70 | 380 | 62 | 119 | 53 | 210 | 14 | 172 | 340 |
| cluster3434 | 55 | 149 | 172 | 218 | 58 | 116 | 960 | 98 | 219 | 657 |
| cluster3435 | 31 | 70 | 491 | 1 | 45 | 180 | 183 | 335 | 232 | 273 |
| cluster3436 | 16 | 48 | 304 | 0 | 773 | 841 | 160 | 51 | 235 | 564 |
| cluster3437 | 68 | 157 | 49 | 82 | 114 | 194 | 194 | 190 | 102 | 468 |
| cluster3438 | 10 | 53 | 61 | 9 | 200 | 182 | 40 | 18 | 72 | 108 |
| cluster3439 | 80 | 227 | 303 | 27 | 83 | 188 | 414 | 106 | 371 | 962 |

| | | | | | | | | | | |
|---|---|---|---|---|---|---|---|---|---|---|
| cluster3440 | 163 | 481 | 1168 | 447 | 605 | 290 | 672 | 255 | 845 | 1504 |
| cluster3441 | 37 | 107 | 419 | 47 | 528 | 366 | 197 | 67 | 260 | 461 |
| cluster3442 | 23 | 82 | 525 | 5 | 1186 | 849 | 417 | 117 | 485 | 720 |
| cluster3443 | 237 | 663 | 1153 | 109 | 5888 | 4333 | 1143 | 920 | 700 | 1596 |
| cluster3444 | 647 | 1598 | 7107 | 1949 | 1672 | 1568 | 2248 | 1105 | 2441 | 5901 |
| cluster3445 | 110 | 248 | 317 | 261 | 352 | 437 | 661 | 280 | 453 | 753 |
| cluster3446 | 171 | 398 | 536 | 72 | 154 | 289 | 229 | 274 | 453 | 843 |
| cluster3447 | 66 | 169 | 395 | 2 | 205 | 309 | 269 | 63 | 99 | 210 |
| cluster3448 | 29 | 83 | 179 | 0 | 103 | 56 | 534 | 47 | 59 | 32 |
| cluster3449 | 111 | 219 | 893 | 644 | 262 | 463 | 503 | 131 | 268 | 538 |
| cluster3450 | 4613 | 11143 | 6434 | 0 | 2515 | 2482 | 2502 | 2611 | 3579 | 6179 |
| cluster3451 | 17 | 53 | 338 | 0 | 2132 | 11 | 711 | 8 | 332 | 2921 |
| cluster3452 | 72 | 213 | 272 | 5 | 273 | 611 | 319 | 378 | 157 | 740 |
| cluster3453 | 1003 | 2536 | 360 | 1208 | 1008 | 705 | 2047 | 2234 | 2531 | 4459 |
| cluster3454 | 625 | 1558 | 1039 | 284 | 512 | 772 | 1431 | 919 | 432 | 1539 |
| cluster3455 | 17 | 40 | 296 | 201 | 66 | 87 | 499 | 33 | 406 | 333 |
| cluster3456 | 34 | 62 | 403 | 116 | 121 | 126 | 200 | 29 | 222 | 265 |
| cluster3457 | 78 | 135 | 1559 | 12 | 61 | 214 | 908 | 90 | 4657 | 5913 |
| cluster3458 | 104 | 332 | 485 | 376 | 476 | 884 | 247 | 163 | 736 | 878 |
| cluster3459 | 192 | 471 | 250 | 0 | 339 | 224 | 775 | 151 | 281 | 671 |
| cluster3460 | 29 | 83 | 424 | 3 | 207 | 309 | 113 | 40 | 229 | 331 |
| cluster3461 | 14 | 68 | 854 | 45 | 49 | 198 | 1789 | 1244 | 64 | 147 |
| cluster3462 | 1579 | 5336 | 3493 | 1219 | 1342 | 4007 | 5753 | 5636 | 1064 | 12323 |
| cluster3463 | 38 | 113 | 151 | 12 | 559 | 443 | 93 | 86 | 417 | 188 |
| cluster3464 | 27 | 76 | 195 | 3 | 114 | 285 | 145 | 35 | 98 | 149 |
| cluster3465 | 107 | 320 | 474 | 803 | 361 | 310 | 430 | 228 | 470 | 903 |
| cluster3466 | 1076 | 3320 | 613 | 642 | 858 | 921 | 975 | 810 | 2243 | 3643 |
| cluster3467 | 68 | 192 | 348 | 28 | 513 | 533 | 471 | 95 | 104 | 376 |
| cluster3468 | 36 | 137 | 566 | 258 | 111 | 139 | 285 | 31 | 328 | 456 |
| cluster3469 | 164 | 413 | 583 | 984 | 141 | 329 | 188 | 132 | 564 | 1162 |
| cluster3470 | 274 | 730 | 956 | 259 | 1006 | 1036 | 524 | 578 | 439 | 1793 |

| | | | | | | | | | | |
|---|---|---|---|---|---|---|---|---|---|---|
| cluster3471 | 194 | 485 | 539 | 36 | 1580 | 2091 | 194 | 330 | 401 | 604 |
| cluster3472 | 209 | 545 | 825 | 1338 | 732 | 1065 | 868 | 478 | 722 | 1822 |
| cluster3473 | 41 | 141 | 56 | 234 | 734 | 268 | 288 | 41 | 53 | 548 |
| cluster3474 | 31 | 86 | 173 | 6 | 758 | 1037 | 146 | 34 | 45 | 178 |
| cluster3475 | 42 | 114 | 80 | 57 | 486 | 264 | 136 | 100 | 164 | 561 |
| cluster3476 | 61 | 168 | 184 | 0 | 258 | 454 | 125 | 45 | 28 | 101 |
| cluster3477 | 22 | 52 | 166 | 192 | 79 | 175 | 402 | 20 | 304 | 500 |
| cluster3478 | 65 | 164 | 832 | 541 | 332 | 394 | 222 | 148 | 681 | 1370 |
| cluster3479 | 129 | 266 | 436 | 0 | 114 | 225 | 160 | 93 | 76 | 257 |
| cluster3480 | 44 | 113 | 1691 | 46 | 290 | 395 | 138 | 90 | 146 | 310 |
| cluster3481 | 52 | 120 | 15 | 29 | 7 | 8 | 154 | 12 | 36 | 54 |
| cluster3482 | 85 | 240 | 517 | 163 | 85 | 172 | 1437 | 274 | 225 | 220 |
| cluster3483 | 571 | 1374 | 2309 | 501 | 1751 | 3410 | 961 | 1200 | 3213 | 4682 |
| cluster3484 | 46 | 97 | 1416 | 11 | 144 | 10 | 450 | 340 | 448 | 725 |
| cluster3485 | 3195 | 8341 | 6695 | 1 | 3484 | 4605 | 7986 | 8974 | 3938 | 9908 |
| cluster3486 | 53 | 149 | 630 | 408 | 93 | 97 | 407 | 48 | 240 | 360 |
| cluster3487 | 1748 | 5584 | 3730 | 411 | 1518 | 1124 | 1599 | 2977 | 749 | 1460 |
| cluster3488 | 209 | 571 | 724 | 137 | 175 | 331 | 524 | 205 | 696 | 1120 |
| cluster3489 | 109 | 314 | 659 | 12 | 1095 | 992 | 385 | 275 | 433 | 853 |
| cluster3490 | 38 | 116 | 189 | 1 | 86 | 101 | 194 | 93 | 42 | 209 |
| cluster3491 | 63 | 184 | 720 | 25 | 493 | 502 | 642 | 129 | 303 | 722 |
| cluster3492 | 49 | 148 | 849 | 24 | 313 | 579 | 614 | 79 | 458 | 696 |
| cluster3493 | 10 | 43 | 646 | 8 | 435 | 128 | 250 | 17 | 189 | 275 |
| cluster3494 | 108 | 322 | 471 | 986 | 396 | 449 | 290 | 177 | 230 | 633 |
| cluster3495 | 130 | 393 | 2939 | 154 | 763 | 1192 | 262 | 359 | 617 | 1125 |
| cluster3496 | 23 | 75 | 139 | 148 | 323 | 324 | 333 | 35 | 404 | 278 |
| cluster3497 | 212 | 559 | 1926 | 1494 | 472 | 748 | 722 | 328 | 440 | 1262 |
| cluster3498 | 46 | 116 | 691 | 2 | 156 | 216 | 136 | 54 | 69 | 243 |
| cluster3499 | 80 | 224 | 466 | 1 | 297 | 306 | 339 | 75 | 176 | 457 |
| cluster3500 | 10614 | 24771 | 9114 | 1709 | 17203 | 8066 | 4638 | 18978 | 8666 | 12011 |
| cluster3501 | 1267 | 3636 | 423 | 1039 | 1254 | 1735 | 1956 | 7432 | 958 | 2472 |

| | | | | | | | | | | |
|---|---|---|---|---|---|---|---|---|---|---|
| cluster3502 | 51 | 113 | 779 | 0 | 541 | 82 | 259 | 92 | 82 | 157 |
| cluster3503 | 67 | 202 | 223 | 24 | 276 | 677 | 181 | 98 | 142 | 396 |
| cluster3504 | 2198 | 3885 | 3940 | 7688 | 779 | 672 | 795 | 2753 | 898 | 2301 |
| cluster3505 | 169 | 450 | 1173 | 116 | 352 | 440 | 160 | 123 | 808 | 1377 |
| cluster3506 | 122 | 389 | 743 | 67 | 362 | 671 | 212 | 155 | 167 | 376 |
| cluster3507 | 83 | 222 | 697 | 42 | 83 | 34 | 92 | 117 | 139 | 339 |
| cluster3508 | 16 | 46 | 206 | 0 | 230 | 333 | 71 | 17 | 74 | 139 |
| cluster3509 | 37 | 121 | 455 | 45 | 209 | 321 | 166 | 64 | 188 | 618 |
| cluster3510 | 4277 | 9463 | 14847 | 782 | 34 | 84 | 4646 | 12053 | 6189 | 14247 |
| cluster3511 | 670 | 1865 | 1257 | 23 | 1781 | 3131 | 987 | 1702 | 686 | 1990 |
| cluster3512 | 26 | 65 | 65 | 25 | 441 | 208 | 385 | 26 | 47 | 198 |
| cluster3513 | 144 | 412 | 1592 | 1250 | 598 | 657 | 1043 | 84 | 2163 | 2567 |
| cluster3514 | 29 | 75 | 677 | 631 | 320 | 182 | 227 | 103 | 381 | 725 |
| cluster3515 | 34 | 90 | 856 | 11 | 61 | 25 | 43 | 9 | 128 | 461 |
| cluster3516 | 373 | 944 | 134 | 5 | 556 | 157 | 134 | 233 | 69 | 334 |
| cluster3517 | 417 | 1034 | 2395 | 817 | 1409 | 1848 | 1475 | 833 | 909 | 1946 |
| cluster3518 | 394 | 1037 | 634 | 4 | 413 | 45 | 477 | 172 | 224 | 1488 |
| cluster3519 | 4 | 44 | 34 | 1 | 6 | 5 | 153 | 2 | 57 | 35 |
| cluster3520 | 21 | 76 | 493 | 24 | 14 | 5 | 14 | 0 | 134 | 126 |
| cluster3521 | 53 | 205 | 568 | 18 | 224 | 233 | 227 | 193 | 381 | 517 |
| cluster3522 | 98 | 259 | 165 | 3 | 139 | 196 | 364 | 186 | 295 | 635 |
| cluster3523 | 108 | 244 | 834 | 0 | 631 | 627 | 509 | 216 | 398 | 528 |
| cluster3524 | 65 | 199 | 799 | 119 | 168 | 315 | 488 | 76 | 254 | 757 |
| cluster3525 | 65 | 162 | 319 | 61 | 152 | 175 | 323 | 275 | 471 | 859 |
| cluster3526 | 46 | 151 | 815 | 117 | 286 | 353 | 236 | 129 | 259 | 440 |
| cluster3527 | 40 | 89 | 135 | 0 | 120 | 154 | 64 | 63 | 102 | 189 |
| cluster3528 | 5385 | 12702 | 19191 | 14192 | 8260 | 12215 | 14136 | 6429 | 18607 | 32944 |
| cluster3529 | 34 | 90 | 726 | 7 | 400 | 335 | 547 | 92 | 1781 | 988 |
| cluster3530 | 73 | 228 | 1148 | 100 | 248 | 308 | 263 | 61 | 323 | 597 |
| cluster3531 | 68 | 212 | 112 | 39 | 303 | 618 | 195 | 310 | 93 | 623 |
| cluster3532 | 26 | 55 | 222 | 473 | 243 | 168 | 875 | 91 | 291 | 494 |

| | | | | | | | | | | |
|---|---|---|---|---|---|---|---|---|---|---|
| cluster3533 | 67 | 173 | 343 | 318 | 96 | 1450 | 702 | 201 | 159 | 707 |
| cluster3534 | 131 | 324 | 312 | 153 | 26 | 97 | 1148 | 253 | 1138 | 1039 |
| cluster3535 | 16 | 59 | 354 | 0 | 42 | 94 | 132 | 24 | 117 | 139 |
| cluster3536 | 83 | 187 | 309 | 285 | 436 | 739 | 201 | 120 | 452 | 812 |
| cluster3537 | 21 | 60 | 368 | 58 | 248 | 297 | 87 | 40 | 95 | 185 |
| cluster3538 | 536 | 1267 | 5814 | 3093 | 266 | 1287 | 1094 | 157 | 2099 | 6076 |
| cluster3539 | 319 | 825 | 832 | 0 | 103 | 1033 | 143 | 251 | 396 | 979 |
| cluster3540 | 21 | 94 | 288 | 0 | 56 | 64 | 269 | 55 | 170 | 351 |
| cluster3541 | 187 | 494 | 877 | 1302 | 1104 | 1831 | 597 | 594 | 1571 | 3395 |
| cluster3542 | 15 | 49 | 82 | 0 | 155 | 153 | 173 | 15 | 75 | 123 |
| cluster3543 | 29 | 69 | 517 | 1 | 25 | 33 | 195 | 33 | 37 | 70 |
| cluster3544 | 11290 | 39743 | 95808 | 19978 | 207 | 905 | 42704 | 31983 | 35597 | 41200 |
| cluster3545 | 24 | 92 | 179 | 16 | 237 | 421 | 270 | 42 | 40 | 96 |
| cluster3546 | 540 | 1607 | 135 | 37 | 1067 | 1140 | 635 | 3400 | 118 | 645 |
| cluster3547 | 255 | 752 | 1501 | 1564 | 1232 | 2069 | 921 | 773 | 283 | 460 |
| cluster3548 | 25 | 78 | 421 | 160 | 220 | 370 | 274 | 82 | 222 | 400 |
| cluster3549 | 328 | 898 | 1317 | 1495 | 1796 | 2730 | 472 | 612 | 1633 | 3852 |
| cluster3550 | 21 | 54 | 155 | 107 | 53 | 50 | 125 | 16 | 180 | 321 |
| cluster3551 | 29 | 91 | 386 | 309 | 98 | 145 | 135 | 19 | 337 | 211 |
| cluster3552 | 335 | 918 | 814 | 639 | 1888 | 3473 | 858 | 356 | 1200 | 2548 |
| cluster3553 | 25 | 42 | 74 | 6 | 82 | 199 | 192 | 27 | 76 | 162 |
| cluster3554 | 29 | 63 | 359 | 16 | 206 | 245 | 372 | 138 | 411 | 807 |
| cluster3555 | 663 | 1831 | 3327 | 1 | 1143 | 5609 | 803 | 1569 | 3157 | 4846 |
| cluster3556 | 31 | 106 | 265 | 8 | 271 | 649 | 65 | 28 | 62 | 49 |
| cluster3557 | 54 | 149 | 277 | 6 | 160 | 232 | 231 | 78 | 122 | 308 |
| cluster3558 | 48 | 146 | 183 | 4 | 144 | 318 | 136 | 59 | 243 | 576 |
| cluster3559 | 29 | 106 | 320 | 33 | 72 | 139 | 124 | 27 | 176 | 562 |
| cluster3560 | 89 | 244 | 426 | 0 | 449 | 665 | 148 | 163 | 291 | 685 |
| cluster3561 | 20 | 55 | 611 | 46 | 97 | 136 | 46 | 6 | 390 | 319 |
| cluster3562 | 49 | 149 | 494 | 4 | 228 | 221 | 200 | 144 | 549 | 601 |
| cluster3563 | 256 | 710 | 254 | 361 | 204 | 403 | 1321 | 860 | 1057 | 2135 |

| | | | | | | | | | | |
|---|---|---|---|---|---|---|---|---|---|---|
| cluster3564 | 21 | 45 | 117 | 585 | 425 | 879 | 321 | 43 | 253 | 448 |
| cluster3565 | 127 | 322 | 150 | 5 | 153 | 301 | 253 | 125 | 232 | 402 |
| cluster3566 | 89 | 230 | 194 | 29 | 112 | 248 | 208 | 61 | 262 | 332 |
| cluster3567 | 77 | 257 | 2254 | 0 | 227 | 240 | 275 | 17 | 2764 | 1016 |
| cluster3568 | 82 | 270 | 367 | 7 | 35 | 26 | 189 | 337 | 627 | 1120 |
| cluster3569 | 80 | 182 | 187 | 6 | 585 | 1403 | 395 | 144 | 300 | 433 |
| cluster3570 | 193 | 622 | 935 | 0 | 2090 | 2440 | 1899 | 353 | 656 | 1258 |
| cluster3571 | 23 | 62 | 56 | 173 | 42 | 74 | 919 | 97 | 45 | 152 |
| cluster3572 | 108 | 219 | 144 | 0 | 2166 | 137 | 396 | 21 | 18 | 567 |
| cluster3573 | 191 | 536 | 1003 | 18 | 208 | 869 | 527 | 633 | 633 | 1939 |
| cluster3574 | 21 | 74 | 243 | 1 | 46 | 59 | 470 | 391 | 110 | 320 |
| cluster3575 | 116 | 323 | 2981 | 103 | 153 | 293 | 121 | 34 | 909 | 1544 |
| cluster3576 | 39 | 81 | 132 | 12 | 276 | 372 | 106 | 55 | 250 | 294 |
| cluster3577 | 300 | 802 | 912 | 1207 | 1676 | 616 | 443 | 279 | 995 | 1872 |
| cluster3578 | 12 | 40 | 74 | 6 | 14 | 40 | 127 | 7 | 90 | 135 |
| cluster3579 | 112 | 359 | 876 | 1041 | 848 | 866 | 1177 | 648 | 550 | 1888 |
| cluster3580 | 19 | 53 | 150 | 0 | 270 | 362 | 58 | 35 | 120 | 186 |
| cluster3581 | 885 | 2205 | 4266 | 194 | 335 | 800 | 1222 | 1971 | 2136 | 3338 |
| cluster3582 | 319 | 938 | 2771 | 1459 | 788 | 1637 | 595 | 424 | 1811 | 2382 |
| cluster3583 | 420 | 1249 | 3268 | 790 | 1589 | 3259 | 1147 | 1305 | 3598 | 6612 |
| cluster3584 | 33 | 61 | 162 | 0 | 285 | 380 | 163 | 21 | 121 | 149 |
| cluster3585 | 40 | 121 | 334 | 88 | 312 | 219 | 639 | 227 | 704 | 862 |
| cluster3586 | 217 | 671 | 1401 | 398 | 966 | 442 | 210 | 377 | 996 | 1072 |
| cluster3587 | 638 | 1752 | 11740 | 1 | 609 | 736 | 840 | 1101 | 694 | 2164 |
| cluster3588 | 44 | 75 | 1733 | 79 | 388 | 301 | 890 | 27 | 581 | 335 |
| cluster3589 | 65 | 162 | 679 | 19 | 213 | 287 | 325 | 72 | 345 | 412 |
| cluster3590 | 24 | 51 | 153 | 2 | 186 | 165 | 129 | 25 | 92 | 252 |
| cluster3591 | 1815 | 4724 | 3494 | 2669 | 2353 | 3604 | 3421 | 2567 | 2312 | 7754 |
| cluster3592 | 78 | 212 | 819 | 17 | 334 | 652 | 357 | 189 | 184 | 539 |
| cluster3593 | 130 | 348 | 166 | 4 | 191 | 307 | 362 | 127 | 106 | 276 |
| cluster3594 | 85 | 275 | 246 | 9 | 37 | 110 | 199 | 38 | 112 | 126 |

| | | | | | | | | | | |
|---|---|---|---|---|---|---|---|---|---|---|
| cluster3595 | 27 | 149 | 712 | 304 | 544 | 1130 | 296 | 122 | 264 | 693 |
| cluster3596 | 7 | 25 | 51 | 3 | 10 | 61 | 39 | 21 | 28 | 79 |
| cluster3597 | 24 | 57 | 465 | 298 | 194 | 298 | 145 | 42 | 398 | 366 |
| cluster3598 | 21 | 59 | 31 | 0 | 6 | 780 | 46 | 27 | 1 | 141 |
| cluster3599 | 257 | 658 | 304 | 5 | 163 | 982 | 1156 | 609 | 438 | 988 |
| cluster3600 | 115 | 309 | 2188 | 227 | 559 | 1278 | 525 | 311 | 1415 | 2001 |
| cluster3601 | 141 | 433 | 709 | 7 | 508 | 895 | 223 | 170 | 366 | 1258 |
| cluster3602 | 8 | 55 | 302 | 93 | 76 | 123 | 144 | 44 | 124 | 154 |
| cluster3603 | 23 | 119 | 570 | 9 | 524 | 336 | 232 | 67 | 317 | 481 |
| cluster3604 | 35 | 53 | 231 | 2 | 180 | 483 | 234 | 5 | 54 | 84 |
| cluster3605 | 27 | 79 | 224 | 4 | 118 | 129 | 106 | 42 | 99 | 175 |
| cluster3606 | 134 | 253 | 584 | 0 | 578 | 732 | 339 | 229 | 198 | 488 |
| cluster3607 | 75 | 181 | 544 | 9 | 592 | 81 | 203 | 89 | 413 | 2261 |
| cluster3608 | 828 | 2139 | 1448 | 1140 | 2390 | 2154 | 1831 | 1928 | 1805 | 4386 |
| cluster3609 | 209 | 616 | 2028 | 0 | 570 | 1859 | 111 | 32 | 2360 | 2539 |
| cluster3610 | 27 | 63 | 55 | 11 | 362 | 462 | 447 | 19 | 113 | 535 |
| cluster3611 | 20 | 82 | 138 | 0 | 451 | 365 | 152 | 25 | 31 | 80 |
| cluster3612 | 44 | 116 | 406 | 0 | 226 | 258 | 143 | 82 | 209 | 512 |
| cluster3613 | 18 | 57 | 364 | 214 | 50 | 786 | 272 | 96 | 470 | 620 |
| cluster3614 | 13 | 43 | 374 | 0 | 216 | 604 | 473 | 18 | 82 | 167 |
| cluster3615 | 13 | 38 | 128 | 206 | 145 | 112 | 100 | 9 | 126 | 207 |
| cluster3616 | 743 | 1819 | 1035 | 3885 | 3635 | 161 | 811 | 2693 | 82 | 364 |
| cluster3617 | 234 | 658 | 1341 | 21 | 78 | 51 | 68 | 77 | 277 | 323 |
| cluster3618 | 14 | 21 | 645 | 0 | 34 | 14 | 255 | 7 | 366 | 116 |
| cluster3619 | 45 | 116 | 376 | 325 | 541 | 745 | 354 | 62 | 130 | 252 |
| cluster3620 | 20 | 62 | 168 | 152 | 214 | 207 | 606 | 62 | 110 | 187 |
| cluster3621 | 39 | 105 | 413 | 7 | 170 | 172 | 61 | 5 | 99 | 137 |
| cluster3622 | 143 | 420 | 1259 | 263 | 255 | 340 | 333 | 265 | 442 | 643 |
| cluster3623 | 196 | 558 | 680 | 50 | 546 | 840 | 192 | 489 | 354 | 659 |
| cluster3624 | 89 | 335 | 720 | 132 | 454 | 1489 | 405 | 267 | 482 | 889 |
| cluster3625 | 31 | 67 | 291 | 39 | 203 | 358 | 129 | 124 | 137 | 244 |

| | | | | | | | | | | |
|---|---|---|---|---|---|---|---|---|---|---|
| cluster3626 | 36 | 104 | 178 | 209 | 118 | 102 | 191 | 97 | 126 | 480 |
| cluster3627 | 38 | 102 | 585 | 48 | 162 | 162 | 341 | 114 | 183 | 348 |
| cluster3628 | 25 | 62 | 351 | 7 | 368 | 443 | 439 | 59 | 120 | 309 |
| cluster3629 | 19 | 57 | 159 | 12 | 501 | 607 | 174 | 26 | 57 | 172 |
| cluster3630 | 192 | 422 | 1876 | 657 | 264 | 786 | 624 | 411 | 734 | 1312 |
| cluster3631 | 18 | 84 | 482 | 989 | 137 | 205 | 201 | 59 | 156 | 182 |
| cluster3632 | 67 | 159 | 140 | 272 | 188 | 257 | 134 | 109 | 186 | 346 |
| cluster3633 | 9834 | 33708 | 24136 | 21432 | 13887 | 14285 | 39015 | 28031 | 27169 | 48186 |
| cluster3634 | 74 | 219 | 457 | 464 | 114 | 224 | 142 | 118 | 347 | 533 |
| cluster3635 | 114 | 490 | 1453 | 430 | 240 | 469 | 563 | 281 | 584 | 1071 |
| cluster3636 | 506 | 1153 | 1500 | 573 | 1005 | 1667 | 2554 | 1243 | 1532 | 3501 |
| cluster3637 | 92 | 254 | 844 | 233 | 492 | 674 | 354 | 227 | 334 | 658 |
| cluster3638 | 281 | 728 | 282 | 22 | 3027 | 215 | 509 | 134 | 298 | 1029 |
| cluster3639 | 31 | 68 | 0 | 46 | 812 | 1851 | 864 | 199 | 12 | 57 |
| cluster3640 | 1352 | 3863 | 98 | 0 | 732 | 4 | 416 | 662 | 68 | 1968 |
| cluster3641 | 45 | 124 | 424 | 316 | 151 | 205 | 587 | 81 | 354 | 584 |
| cluster3642 | 14 | 90 | 438 | 8 | 393 | 623 | 495 | 59 | 134 | 382 |
| cluster3643 | 20 | 81 | 274 | 0 | 214 | 299 | 142 | 31 | 167 | 281 |
| cluster3644 | 170 | 534 | 165 | 141 | 446 | 80 | 300 | 420 | 144 | 726 |
| cluster3645 | 840 | 1904 | 1725 | 0 | 911 | 223 | 2117 | 9974 | 95 | 3316 |
| cluster3646 | 19 | 60 | 231 | 0 | 406 | 492 | 506 | 34 | 103 | 181 |
| cluster3647 | 43 | 93 | 143 | 29 | 198 | 272 | 65 | 56 | 69 | 159 |
| cluster3648 | 479 | 1286 | 821 | 1044 | 1120 | 3356 | 740 | 910 | 2964 | 5915 |
| cluster3649 | 92 | 212 | 455 | 51 | 122 | 211 | 256 | 156 | 385 | 658 |
| cluster3650 | 133 | 366 | 4087 | 147 | 73 | 93 | 316 | 190 | 479 | 555 |
| cluster3651 | 22 | 48 | 6 | 47 | 57 | 5 | 435 | 239 | 5 | 175 |
| cluster3652 | 54 | 165 | 358 | 849 | 1804 | 1619 | 284 | 125 | 98 | 321 |
| cluster3653 | 269 | 650 | 721 | 1171 | 305 | 627 | 308 | 172 | 232 | 1384 |
| cluster3654 | 39 | 122 | 225 | 53 | 323 | 294 | 127 | 220 | 117 | 246 |
| cluster3655 | 32 | 99 | 259 | 6 | 156 | 130 | 207 | 63 | 78 | 412 |
| cluster3656 | 28 | 108 | 136 | 221 | 338 | 418 | 86 | 64 | 213 | 228 |

| | | | | | | | | | | |
|---|---|---|---|---|---|---|---|---|---|---|
| cluster3657 | 126 | 416 | 399 | 68 | 755 | 1384 | 354 | 114 | 185 | 616 |
| cluster3658 | 4348 | 11756 | 24211 | 8634 | 4816 | 17057 | 4253 | 3991 | 35121 | 29267 |
| cluster3659 | 716 | 1500 | 672 | 16 | 956 | 510 | 788 | 955 | 379 | 1438 |
| cluster3660 | 22 | 82 | 28 | 0 | 177 | 80 | 82 | 58 | 9 | 433 |
| cluster3661 | 42 | 99 | 539 | 135 | 435 | 890 | 364 | 60 | 167 | 522 |
| cluster3662 | 19 | 58 | 228 | 1 | 40 | 58 | 124 | 74 | 68 | 156 |
| cluster3663 | 121 | 308 | 786 | 2 | 266 | 493 | 260 | 242 | 192 | 267 |
| cluster3664 | 49 | 97 | 190 | 65 | 76 | 103 | 62 | 20 | 102 | 173 |
| cluster3665 | 84 | 228 | 626 | 0 | 200 | 381 | 142 | 319 | 174 | 496 |
| cluster3666 | 94 | 283 | 538 | 28 | 5066 | 9660 | 457 | 194 | 369 | 630 |
| cluster3667 | 46 | 101 | 488 | 176 | 169 | 405 | 404 | 332 | 163 | 547 |
| cluster3668 | 136 | 345 | 425 | 100 | 370 | 683 | 270 | 195 | 268 | 413 |
| cluster3669 | 953 | 2380 | 1564 | 383 | 2867 | 1453 | 1105 | 1439 | 1718 | 2898 |
| cluster3670 | 1045 | 2796 | 174 | 577 | 243 | 4919 | 963 | 1395 | 368 | 1099 |
| cluster3671 | 469 | 1007 | 1790 | 95 | 2002 | 1527 | 509 | 1036 | 811 | 1292 |
| cluster3672 | 82 | 268 | 140 | 8 | 129 | 196 | 534 | 252 | 187 | 354 |
| cluster3673 | 12 | 38 | 245 | 0 | 208 | 226 | 427 | 38 | 117 | 180 |
| cluster3674 | 205 | 549 | 78 | 0 | 32 | 39 | 74 | 27 | 27 | 340 |
| cluster3675 | 607 | 1311 | 3365 | 1130 | 515 | 1063 | 1459 | 1212 | 2715 | 3765 |
| cluster3676 | 37 | 121 | 702 | 0 | 491 | 777 | 139 | 144 | 301 | 407 |
| cluster3677 | 112 | 341 | 272 | 503 | 1008 | 1572 | 331 | 385 | 342 | 919 |
| cluster3678 | 69 | 152 | 209 | 88 | 472 | 610 | 97 | 125 | 113 | 252 |
| cluster3679 | 249 | 534 | 548 | 509 | 589 | 1367 | 132 | 87 | 507 | 1183 |
| cluster3680 | 57 | 167 | 558 | 20 | 350 | 467 | 282 | 118 | 98 | 642 |
| cluster3681 | 31 | 107 | 177 | 8 | 402 | 724 | 389 | 97 | 39 | 268 |
| cluster3682 | 10 | 42 | 256 | 0 | 67 | 95 | 262 | 37 | 128 | 278 |
| cluster3683 | 76 | 132 | 396 | 78 | 760 | 1354 | 597 | 213 | 448 | 625 |
| cluster3684 | 35 | 98 | 349 | 2 | 75 | 131 | 170 | 26 | 151 | 221 |
| cluster3685 | 190 | 421 | 2003 | 10 | 224 | 118 | 613 | 206 | 638 | 1613 |
| cluster3686 | 57 | 156 | 140 | 0 | 147 | 201 | 84 | 20 | 54 | 86 |
| cluster3687 | 19 | 71 | 196 | 0 | 149 | 149 | 128 | 54 | 98 | 166 |

| | | | | | | | | | | |
|---|---|---|---|---|---|---|---|---|---|---|
| cluster3688 | 239 | 636 | 1434 | 41 | 953 | 1791 | 409 | 466 | 321 | 802 |
| cluster3689 | 42 | 167 | 1 | 25 | 292 | 22 | 1299 | 47 | 4 | 41 |
| cluster3690 | 201 | 523 | 253 | 489 | 214 | 370 | 403 | 333 | 757 | 1154 |
| cluster3691 | 32 | 77 | 515 | 165 | 160 | 313 | 108 | 11 | 188 | 239 |
| cluster3692 | 38 | 103 | 420 | 1 | 85 | 171 | 122 | 76 | 104 | 193 |
| cluster3693 | 127 | 69 | 65 | 0 | 356 | 451 | 85 | 61 | 51 | 95 |
| cluster3694 | 1335 | 2674 | 1942 | 494 | 616 | 1394 | 3027 | 2310 | 1536 | 4068 |
| cluster3695 | 95 | 256 | 86 | 35 | 231 | 346 | 334 | 199 | 181 | 434 |
| cluster3696 | 77 | 291 | 758 | 63 | 153 | 302 | 217 | 161 | 209 | 468 |
| cluster3697 | 335 | 622 | 529 | 1463 | 1702 | 20 | 596 | 587 | 473 | 3348 |
| cluster3698 | 81 | 271 | 878 | 329 | 145 | 191 | 484 | 79 | 688 | 1076 |
| cluster3699 | 67 | 165 | 394 | 0 | 312 | 499 | 111 | 73 | 223 | 236 |
| cluster3700 | 61 | 247 | 201 | 6 | 198 | 398 | 132 | 89 | 147 | 462 |
| cluster3701 | 57 | 128 | 139 | 147 | 105 | 204 | 218 | 60 | 81 | 238 |
| cluster3702 | 33 | 103 | 142 | 11 | 353 | 381 | 221 | 45 | 358 | 508 |
| cluster3703 | 112 | 425 | 857 | 625 | 1277 | 2127 | 979 | 304 | 349 | 1305 |
| cluster3704 | 504 | 1252 | 1185 | 2476 | 3590 | 5496 | 628 | 253 | 452 | 1090 |
| cluster3705 | 12 | 37 | 60 | 175 | 122 | 80 | 65 | 20 | 106 | 74 |
| cluster3706 | 56 | 129 | 239 | 59 | 330 | 295 | 384 | 102 | 258 | 472 |
| cluster3707 | 15 | 32 | 315 | 0 | 81 | 108 | 148 | 14 | 11 | 50 |
| cluster3708 | 4 | 31 | 77 | 0 | 92 | 26 | 135 | 1 | 8 | 80 |
| cluster3709 | 93 | 376 | 3634 | 777 | 753 | 635 | 704 | 359 | 2760 | 3536 |
| cluster3710 | 23 | 63 | 110 | 5 | 141 | 351 | 120 | 42 | 38 | 128 |
| cluster3711 | 745 | 2217 | 6638 | 902 | 1941 | 1259 | 2730 | 1626 | 2531 | 4256 |
| cluster3712 | 22 | 65 | 47 | 100 | 119 | 148 | 62 | 22 | 104 | 84 |
| cluster3713 | 18 | 61 | 260 | 17 | 90 | 142 | 94 | 25 | 23 | 65 |
| cluster3714 | 24 | 60 | 196 | 330 | 62 | 85 | 110 | 49 | 162 | 220 |
| cluster3715 | 41 | 170 | 36 | 10 | 171 | 365 | 48 | 35 | 2 | 66 |
| cluster3716 | 9 | 28 | 47 | 0 | 43 | 28 | 30 | 47 | 18 | 199 |
| cluster3717 | 505 | 1228 | 880 | 743 | 1315 | 1277 | 531 | 914 | 318 | 1755 |
| cluster3718 | 31 | 78 | 180 | 0 | 54 | 67 | 74 | 29 | 28 | 58 |

| | | | | | | | | | | |
|---|---|---|---|---|---|---|---|---|---|---|
| cluster3719 | 13 | 39 | 129 | 8 | 176 | 250 | 51 | 2 | 37 | 62 |
| cluster3720 | 73 | 241 | 520 | 0 | 411 | 621 | 437 | 177 | 257 | 458 |
| cluster3721 | 8 | 30 | 155 | 174 | 41 | 107 | 117 | 15 | 130 | 104 |
| cluster3722 | 39 | 97 | 289 | 1 | 112 | 144 | 69 | 38 | 141 | 176 |
| cluster3723 | 104 | 303 | 791 | 385 | 847 | 1314 | 532 | 255 | 224 | 759 |
| cluster3724 | 24 | 103 | 241 | 4 | 613 | 432 | 257 | 79 | 125 | 226 |
| cluster3725 | 32 | 122 | 392 | 0 | 411 | 477 | 328 | 113 | 145 | 355 |
| cluster3726 | 34 | 114 | 285 | 77 | 75 | 100 | 148 | 49 | 143 | 399 |
| cluster3727 | 39 | 96 | 599 | 0 | 101 | 167 | 185 | 96 | 247 | 342 |
| cluster3728 | 4331 | 11029 | 9874 | 9232 | 10020 | 12863 | 3559 | 8646 | 4244 | 10963 |
| cluster3729 | 175 | 428 | 1581 | 55 | 544 | 808 | 710 | 599 | 670 | 2125 |
| cluster3730 | 76 | 267 | 1049 | 703 | 274 | 429 | 409 | 142 | 528 | 794 |
| cluster3731 | 18 | 38 | 280 | 302 | 204 | 478 | 287 | 37 | 129 | 256 |
| cluster3732 | 93 | 233 | 137 | 36 | 364 | 565 | 442 | 128 | 48 | 153 |
| cluster3733 | 29 | 91 | 17 | 14 | 130 | 40 | 57 | 29 | 75 | 71 |
| cluster3734 | 86 | 270 | 562 | 4 | 1007 | 1128 | 233 | 165 | 282 | 700 |
| cluster3735 | 484 | 1349 | 626 | 1089 | 1117 | 1357 | 1913 | 3283 | 701 | 2933 |
| cluster3736 | 10 | 29 | 47 | 1 | 21 | 43 | 79 | 1 | 63 | 80 |
| cluster3737 | 114 | 373 | 665 | 395 | 476 | 861 | 456 | 183 | 589 | 1395 |
| cluster3738 | 15 | 36 | 507 | 1701 | 310 | 517 | 186 | 73 | 90 | 256 |
| cluster3739 | 825 | 1809 | 6708 | 1730 | 1975 | 2906 | 1821 | 1143 | 2101 | 5046 |
| cluster3740 | 10 | 58 | 184 | 5 | 308 | 431 | 186 | 183 | 147 | 300 |
| cluster3741 | 156 | 508 | 3855 | 1238 | 602 | 1794 | 340 | 124 | 3657 | 4123 |
| cluster3742 | 49 | 138 | 299 | 0 | 530 | 420 | 79 | 54 | 215 | 378 |
| cluster3743 | 73 | 254 | 352 | 52 | 393 | 430 | 263 | 195 | 154 | 357 |
| cluster3744 | 40 | 99 | 480 | 20 | 275 | 312 | 584 | 108 | 378 | 547 |
| cluster3745 | 14 | 49 | 815 | 5 | 399 | 2815 | 156 | 101 | 187 | 1528 |
| cluster3746 | 11 | 47 | 72 | 129 | 145 | 170 | 82 | 26 | 35 | 39 |
| cluster3747 | 120 | 351 | 1558 | 40 | 505 | 833 | 289 | 131 | 234 | 602 |
| cluster3748 | 178 | 596 | 363 | 1193 | 508 | 1443 | 923 | 301 | 315 | 921 |
| cluster3749 | 818 | 1884 | 406 | 2 | 1525 | 1537 | 815 | 1095 | 458 | 1246 |

| | | | | | | | | | | |
|---|---|---|---|---|---|---|---|---|---|---|
| cluster3750 | 8 | 30 | 239 | 1 | 126 | 69 | 917 | 77 | 178 | 203 |
| cluster3751 | 19 | 43 | 215 | 36 | 204 | 197 | 280 | 71 | 120 | 278 |
| cluster3752 | 434 | 1250 | 37 | 7 | 160 | 96 | 869 | 274 | 101 | 554 |
| cluster3753 | 13941 | 36034 | 56664 | 7798 | 517 | 869 | 38239 | 34121 | 35878 | 49754 |
| cluster3754 | 32 | 62 | 176 | 69 | 80 | 332 | 168 | 47 | 33 | 191 |
| cluster3755 | 16 | 37 | 500 | 1733 | 169 | 1056 | 253 | 3 | 31 | 381 |
| cluster3756 | 93 | 268 | 345 | 9 | 185 | 319 | 220 | 286 | 301 | 637 |
| cluster3757 | 19 | 33 | 2212 | 4 | 8 | 199 | 116 | 18 | 283 | 233 |
| cluster3758 | 29 | 92 | 144 | 165 | 299 | 189 | 1101 | 72 | 105 | 231 |
| cluster3759 | 18 | 31 | 112 | 106 | 59 | 86 | 230 | 28 | 161 | 288 |
| cluster3760 | 95 | 229 | 576 | 798 | 546 | 1009 | 319 | 134 | 45 | 281 |
| cluster3761 | 122 | 302 | 1659 | 12 | 41 | 82 | 69 | 124 | 407 | 518 |
| cluster3762 | 20 | 51 | 532 | 378 | 196 | 432 | 677 | 16 | 717 | 907 |
| cluster3763 | 7 | 38 | 202 | 236 | 101 | 91 | 36 | 17 | 159 | 195 |
| cluster3764 | 174 | 480 | 200 | 1 | 363 | 516 | 147 | 437 | 148 | 634 |
| cluster3765 | 339 | 1014 | 352 | 619 | 1004 | 1061 | 462 | 1370 | 645 | 1264 |
| cluster3766 | 2400 | 5700 | 3351 | 440 | 1476 | 1919 | 2357 | 3271 | 3662 | 5988 |
| cluster3767 | 70 | 112 | 137 | 2 | 124 | 317 | 47 | 68 | 106 | 218 |
| cluster3768 | 196 | 410 | 3194 | 5 | 323 | 645 | 1547 | 742 | 1093 | 1852 |
| cluster3769 | 64 | 132 | 341 | 60 | 251 | 433 | 666 | 86 | 670 | 423 |
| cluster3770 | 272 | 673 | 70 | 0 | 2271 | 17 | 184 | 1207 | 7 | 270 |
| cluster3771 | 18 | 51 | 168 | 5 | 229 | 294 | 87 | 28 | 140 | 189 |
| cluster3772 | 38 | 107 | 269 | 34 | 42 | 73 | 504 | 120 | 307 | 374 |
| cluster3773 | 1002 | 3925 | 1896 | 1100 | 2401 | 1294 | 4710 | 2294 | 1317 | 4550 |
| cluster3774 | 47 | 111 | 680 | 256 | 112 | 255 | 467 | 42 | 693 | 608 |
| cluster3775 | 106 | 265 | 566 | 267 | 666 | 1463 | 424 | 711 | 971 | 1271 |
| cluster3776 | 21 | 48 | 36 | 92 | 44 | 36 | 178 | 26 | 105 | 78 |
| cluster3777 | 47 | 134 | 73 | 18 | 66 | 58 | 76 | 40 | 81 | 363 |
| cluster3778 | 792 | 1724 | 3268 | 4576 | 996 | 1380 | 1546 | 712 | 593 | 1546 |
| cluster3779 | 192 | 469 | 227 | 507 | 519 | 699 | 154 | 96 | 102 | 362 |
| cluster3780 | 529 | 1513 | 1187 | 4 | 16 | 10 | 90 | 70 | 34 | 60 |

| | | | | | | | | | | |
|---|---|---|---|---|---|---|---|---|---|---|
| cluster3781 | 22 | 59 | 43 | 50 | 125 | 234 | 108 | 7 | 55 | 119 |
| cluster3782 | 9 | 21 | 226 | 44 | 86 | 130 | 169 | 57 | 347 | 542 |
| cluster3783 | 31 | 84 | 232 | 0 | 469 | 606 | 557 | 66 | 101 | 203 |
| cluster3784 | 6 | 24 | 27 | 41 | 44 | 80 | 38 | 20 | 26 | 48 |
| cluster3785 | 40 | 35 | 654 | 35 | 200 | 155 | 527 | 21 | 757 | 421 |
| cluster3786 | 6 | 26 | 482 | 1043 | 488 | 1815 | 175 | 51 | 392 | 1500 |
| cluster3787 | 157 | 337 | 468 | 3 | 64 | 67 | 299 | 187 | 49 | 160 |
| cluster3788 | 3143 | 12068 | 52995 | 16421 | 7195 | 8471 | 9999 | 4934 | 16096 | 22869 |
| cluster3789 | 14 | 37 | 90 | 102 | 220 | 203 | 474 | 9 | 462 | 112 |
| cluster3790 | 14 | 27 | 190 | 0 | 89 | 122 | 141 | 31 | 21 | 562 |
| cluster3791 | 161 | 498 | 509 | 66 | 142 | 381 | 46 | 31 | 417 | 514 |
| cluster3792 | 279 | 634 | 1003 | 100 | 535 | 790 | 1082 | 729 | 604 | 2484 |
| cluster3793 | 100 | 326 | 385 | 79 | 674 | 664 | 183 | 279 | 121 | 539 |
| cluster3794 | 16 | 49 | 72 | 4 | 63 | 138 | 234 | 46 | 21 | 111 |
| cluster3795 | 138 | 392 | 383 | 196 | 415 | 572 | 234 | 584 | 381 | 815 |
| cluster3796 | 24 | 66 | 98 | 0 | 27 | 7 | 18 | 20 | 64 | 49 |
| cluster3797 | 22 | 51 | 557 | 8 | 125 | 151 | 335 | 30 | 249 | 404 |
| cluster3798 | 16 | 47 | 243 | 5 | 81 | 192 | 189 | 41 | 105 | 213 |
| cluster3799 | 14 | 51 | 60 | 113 | 140 | 175 | 76 | 28 | 138 | 129 |
| cluster3800 | 3952 | 9678 | 9360 | 4593 | 981 | 1568 | 6664 | 8164 | 2210 | 8159 |
| cluster3801 | 13 | 41 | 194 | 0 | 135 | 277 | 91 | 36 | 82 | 118 |
| cluster3802 | 9 | 32 | 65 | 216 | 24 | 186 | 21 | 13 | 4 | 20 |
| cluster3803 | 342 | 1183 | 1621 | 0 | 1398 | 3426 | 528 | 886 | 607 | 3947 |
| cluster3804 | 14 | 65 | 221 | 55 | 480 | 551 | 402 | 40 | 91 | 169 |
| cluster3805 | 13 | 56 | 647 | 57 | 130 | 229 | 66 | 93 | 30 | 144 |
| cluster3806 | 109 | 273 | 441 | 2 | 271 | 449 | 133 | 261 | 208 | 494 |
| cluster3807 | 29 | 116 | 294 | 200 | 78 | 130 | 270 | 112 | 223 | 467 |
| cluster3808 | 51 | 161 | 588 | 0 | 206 | 379 | 183 | 88 | 388 | 407 |
| cluster3809 | 26 | 95 | 447 | 74 | 321 | 397 | 379 | 115 | 200 | 298 |
| cluster3810 | 28 | 91 | 127 | 180 | 283 | 401 | 142 | 84 | 36 | 125 |
| cluster3811 | 212 | 662 | 1758 | 1072 | 472 | 777 | 395 | 345 | 401 | 1899 |

| | | | | | | | | | | |
|---|---|---|---|---|---|---|---|---|---|---|
| cluster3812 | 10 | 31 | 415 | 21 | 19 | 18 | 113 | 12 | 226 | 256 |
| cluster3813 | 20 | 61 | 95 | 0 | 142 | 220 | 80 | 26 | 44 | 97 |
| cluster3814 | 59 | 160 | 346 | 266 | 38 | 163 | 280 | 53 | 432 | 359 |
| cluster3815 | 10 | 28 | 124 | 0 | 271 | 355 | 148 | 40 | 60 | 177 |
| cluster3816 | 162 | 409 | 1512 | 3 | 524 | 548 | 373 | 318 | 625 | 911 |
| cluster3817 | 13 | 44 | 158 | 382 | 40 | 88 | 221 | 53 | 106 | 275 |
| cluster3818 | 17 | 42 | 123 | 389 | 166 | 322 | 66 | 54 | 49 | 87 |
| cluster3819 | 17 | 36 | 213 | 7 | 4 | 396 | 109 | 164 | 18 | 172 |
| cluster3820 | 57 | 177 | 430 | 112 | 236 | 283 | 356 | 194 | 156 | 477 |
| cluster3821 | 103 | 376 | 187 | 206 | 403 | 164 | 500 | 493 | 2406 | 2818 |
| cluster3822 | 11 | 34 | 499 | 0 | 491 | 577 | 105 | 41 | 11 | 21 |
| cluster3823 | 15 | 40 | 258 | 157 | 162 | 283 | 498 | 96 | 132 | 212 |
| cluster3824 | 34 | 72 | 121 | 4 | 739 | 707 | 78 | 149 | 49 | 220 |
| cluster3825 | 36 | 140 | 155 | 38 | 108 | 88 | 399 | 17 | 133 | 282 |
| cluster3826 | 9 | 31 | 100 | 0 | 22 | 52 | 118 | 14 | 12 | 81 |
| cluster3827 | 27 | 97 | 15 | 47 | 151 | 312 | 276 | 205 | 2 | 134 |
| cluster3828 | 12 | 41 | 174 | 2 | 139 | 219 | 151 | 42 | 106 | 125 |
| cluster3829 | 8 | 31 | 243 | 0 | 107 | 160 | 80 | 23 | 66 | 161 |
| cluster3830 | 270 | 708 | 5852 | 1209 | 4008 | 3827 | 1554 | 643 | 1499 | 1967 |
| cluster3831 | 74 | 148 | 107 | 9 | 357 | 483 | 204 | 126 | 83 | 261 |
| cluster3832 | 104 | 253 | 808 | 18 | 578 | 813 | 140 | 144 | 396 | 699 |
| cluster3833 | 46 | 100 | 45 | 3 | 70 | 175 | 78 | 3 | 196 | 309 |
| cluster3834 | 105 | 360 | 466 | 6 | 221 | 197 | 230 | 626 | 88 | 438 |
| cluster3835 | 26 | 97 | 231 | 0 | 71 | 174 | 159 | 42 | 87 | 236 |
| cluster3836 | 64 | 244 | 421 | 28 | 636 | 1069 | 932 | 183 | 215 | 647 |
| cluster3837 | 9 | 29 | 169 | 6 | 199 | 248 | 218 | 65 | 130 | 207 |
| cluster3838 | 26 | 79 | 199 | 1 | 88 | 205 | 124 | 35 | 242 | 433 |
| cluster3839 | 23 | 64 | 55 | 3 | 3 | 10 | 156 | 51 | 127 | 187 |
| cluster3840 | 70 | 168 | 440 | 921 | 713 | 804 | 148 | 71 | 284 | 444 |
| cluster3841 | 150 | 427 | 865 | 3 | 181 | 269 | 226 | 279 | 481 | 585 |
| cluster3842 | 6 | 18 | 66 | 63 | 131 | 81 | 75 | 4 | 249 | 268 |

| | | | | | | | | | | |
|---|---|---|---|---|---|---|---|---|---|---|
| cluster3843 | 39 | 82 | 229 | 90 | 290 | 230 | 182 | 96 | 61 | 148 |
| cluster3844 | 25 | 87 | 8 | 0 | 13 | 35 | 201 | 4 | 2 | 140 |
| cluster3845 | 10205 | 26064 | 28892 | 30081 | 15 | 13 | 22102 | 21820 | 21814 | 38484 |
| cluster3846 | 26 | 132 | 126 | 49 | 105 | 112 | 113 | 64 | 74 | 252 |
| cluster3847 | 231 | 718 | 916 | 127 | 538 | 729 | 144 | 70 | 510 | 737 |
| cluster3848 | 132 | 341 | 1458 | 34 | 72 | 849 | 150 | 462 | 1352 | 1665 |
| cluster3849 | 24 | 47 | 214 | 23 | 160 | 174 | 31 | 19 | 44 | 84 |
| cluster3850 | 4887 | 12992 | 7237 | 1287 | 7942 | 5210 | 4604 | 12266 | 6622 | 15274 |
| cluster3851 | 94 | 212 | 240 | 3 | 116 | 227 | 59 | 24 | 48 | 56 |
| cluster3852 | 17 | 65 | 236 | 467 | 369 | 519 | 73 | 46 | 58 | 139 |
| cluster3853 | 21 | 69 | 253 | 24 | 733 | 1370 | 183 | 76 | 87 | 147 |
| cluster3854 | 100 | 284 | 270 | 95 | 193 | 419 | 303 | 237 | 45 | 336 |
| cluster3855 | 17 | 32 | 60 | 0 | 105 | 118 | 95 | 21 | 12 | 46 |
| cluster3856 | 474 | 1000 | 1508 | 140 | 1880 | 2424 | 529 | 1134 | 889 | 1302 |
| cluster3857 | 11 | 52 | 101 | 5 | 99 | 128 | 167 | 58 | 87 | 182 |
| cluster3858 | 16 | 81 | 278 | 629 | 346 | 649 | 340 | 130 | 36 | 125 |
| cluster3859 | 448 | 1467 | 5668 | 493 | 1045 | 2339 | 915 | 1608 | 2376 | 34002 |
| cluster3860 | 15 | 38 | 138 | 76 | 300 | 58 | 118 | 60 | 211 | 262 |
| cluster3861 | 61 | 211 | 230 | 63 | 148 | 283 | 143 | 114 | 97 | 327 |
| cluster3862 | 126 | 311 | 996 | 286 | 736 | 1201 | 391 | 324 | 576 | 1471 |
| cluster3863 | 14 | 27 | 101 | 141 | 95 | 216 | 38 | 22 | 60 | 126 |
| cluster3864 | 57 | 113 | 193 | 23 | 267 | 311 | 193 | 66 | 71 | 203 |
| cluster3865 | 10 | 34 | 74 | 0 | 178 | 219 | 11 | 9 | 13 | 29 |
| cluster3866 | 16 | 23 | 25 | 0 | 58 | 182 | 54 | 10 | 0 | 36 |
| cluster3867 | 1057 | 2894 | 5426 | 595 | 1216 | 1516 | 2377 | 1778 | 2026 | 3658 |
| cluster3868 | 68 | 236 | 307 | 159 | 305 | 306 | 568 | 170 | 351 | 620 |
| cluster3869 | 9 | 40 | 56 | 0 | 346 | 326 | 65 | 29 | 56 | 107 |
| cluster3870 | 41 | 136 | 152 | 0 | 54 | 16 | 59 | 15 | 56 | 134 |
| cluster3871 | 32 | 76 | 203 | 11 | 280 | 345 | 87 | 46 | 42 | 94 |
| cluster3872 | 13 | 26 | 217 | 3 | 81 | 218 | 95 | 11 | 75 | 132 |
| cluster3873 | 237 | 594 | 1336 | 890 | 772 | 1298 | 341 | 299 | 845 | 1246 |

| | | | | | | | | | | |
|---|---|---|---|---|---|---|---|---|---|---|
| cluster3874 | 21 | 46 | 343 | 73 | 137 | 174 | 339 | 52 | 320 | 307 |
| cluster3875 | 51 | 96 | 196 | 71 | 555 | 831 | 294 | 87 | 218 | 271 |
| cluster3876 | 408 | 1076 | 3026 | 2664 | 1644 | 4340 | 867 | 341 | 2646 | 3032 |
| cluster3877 | 134 | 409 | 276 | 300 | 623 | 585 | 362 | 523 | 294 | 465 |
| cluster3879 | 12 | 66 | 132 | 10 | 137 | 139 | 69 | 41 | 53 | 141 |
| cluster3880 | 40 | 124 | 499 | 0 | 6 | 16 | 57 | 8 | 54 | 42 |
| cluster3881 | 18 | 49 | 152 | 0 | 67 | 148 | 44 | 18 | 43 | 54 |
| cluster3882 | 10 | 21 | 61 | 4 | 29 | 30 | 43 | 0 | 7 | 38 |
| cluster3883 | 10 | 24 | 1519 | 5 | 37 | 510 | 29 | 154 | 466 | 873 |
| cluster3884 | 9 | 33 | 248 | 54 | 206 | 225 | 959 | 15 | 90 | 155 |
| cluster3885 | 303 | 635 | 1645 | 64 | 273 | 512 | 1899 | 525 | 5760 | 5274 |
| cluster3886 | 54 | 143 | 1104 | 8 | 415 | 470 | 286 | 153 | 142 | 243 |
| cluster3887 | 10 | 39 | 19 | 0 | 44 | 57 | 89 | 41 | 15 | 79 |
| cluster3888 | 64 | 198 | 614 | 0 | 198 | 178 | 198 | 142 | 206 | 238 |
| cluster3889 | 4 | 14 | 21 | 0 | 108 | 228 | 70 | 18 | 4 | 17 |
| cluster3890 | 53 | 169 | 478 | 1014 | 365 | 700 | 45 | 114 | 111 | 279 |
| cluster3891 | 1143 | 2638 | 486 | 669 | 3815 | 3991 | 255 | 544 | 336 | 336 |
| cluster3892 | 398 | 1395 | 102 | 240 | 389 | 172 | 1979 | 165 | 56 | 562 |
| cluster3893 | 75 | 260 | 418 | 367 | 418 | 546 | 494 | 226 | 265 | 744 |
| cluster3894 | 47 | 127 | 99 | 25 | 79 | 175 | 163 | 121 | 33 | 248 |
| cluster3895 | 21 | 58 | 145 | 0 | 172 | 283 | 204 | 33 | 95 | 264 |
| cluster3896 | 126 | 406 | 628 | 0 | 306 | 594 | 252 | 150 | 280 | 1058 |
| cluster3897 | 43 | 134 | 120 | 297 | 287 | 549 | 45 | 22 | 96 | 190 |
| cluster3898 | 12 | 21 | 188 | 194 | 197 | 385 | 174 | 6 | 200 | 158 |
| cluster3899 | 29 | 88 | 275 | 15 | 148 | 334 | 50 | 28 | 17 | 103 |
| cluster3900 | 2673 | 7765 | 10144 | 0 | 9443 | 815 | 1018 | 1703 | 4373 | 11417 |
| cluster3901 | 31 | 91 | 154 | 26 | 1181 | 1996 | 283 | 50 | 77 | 133 |
| cluster3902 | 110 | 294 | 557 | 179 | 2049 | 1432 | 679 | 148 | 447 | 427 |
| cluster3903 | 138 | 438 | 1660 | 91 | 472 | 1171 | 486 | 533 | 842 | 1733 |
| cluster3904 | 712 | 2146 | 786 | 285 | 740 | 924 | 1354 | 3336 | 673 | 1514 |
| cluster3905 | 811 | 1725 | 217 | 0 | 186 | 3 | 78 | 159 | 199 | 221 |

| | | | | | | | | | | |
|---|---|---|---|---|---|---|---|---|---|---|
| cluster3906 | 26 | 42 | 113 | 6 | 140 | 180 | 94 | 79 | 37 | 79 |
| cluster3907 | 129 | 251 | 997 | 0 | 337 | 853 | 625 | 666 | 772 | 1381 |
| cluster3908 | 44 | 154 | 160 | 31 | 307 | 344 | 348 | 96 | 112 | 237 |
| cluster3909 | 107 | 253 | 259 | 18 | 605 | 652 | 429 | 161 | 130 | 193 |
| cluster3910 | 58 | 242 | 424 | 0 | 78 | 96 | 55 | 117 | 76 | 410 |
| cluster3911 | 13 | 30 | 453 | 0 | 213 | 316 | 210 | 31 | 170 | 419 |
| cluster3912 | 253 | 583 | 1201 | 316 | 672 | 1124 | 756 | 470 | 551 | 809 |
| cluster3913 | 8 | 28 | 196 | 7 | 8 | 25 | 45 | 36 | 39 | 234 |
| cluster3914 | 384 | 1111 | 853 | 0 | 414 | 5 | 499 | 44 | 227 | 1161 |
| cluster3915 | 150 | 381 | 601 | 3 | 288 | 160 | 102 | 1047 | 64 | 703 |
| cluster3916 | 21 | 49 | 192 | 36 | 87 | 228 | 55 | 27 | 47 | 99 |
| cluster3917 | 8 | 32 | 380 | 11 | 414 | 515 | 348 | 84 | 171 | 349 |
| cluster3918 | 16 | 65 | 410 | 2 | 119 | 238 | 72 | 32 | 71 | 198 |
| cluster3919 | 6086 | 19084 | 36644 | 5817 | 450 | 1014 | 25863 | 17186 | 21490 | 25287 |
| cluster3920 | 11 | 29 | 246 | 40 | 45 | 50 | 509 | 46 | 90 | 128 |
| cluster3921 | 382 | 1154 | 3905 | 0 | 1450 | 3179 | 731 | 864 | 1611 | 4332 |
| cluster3922 | 709 | 2520 | 1295 | 0 | 1952 | 2712 | 1770 | 2253 | 1519 | 2192 |
| cluster3923 | 24 | 43 | 114 | 41 | 41 | 60 | 80 | 40 | 60 | 118 |
| cluster3924 | 5 | 21 | 24 | 3 | 113 | 148 | 64 | 25 | 6 | 52 |
| cluster3925 | 3419 | 9202 | 5948 | 0 | 6 | 0 | 3466 | 5890 | 512 | 10761 |
| cluster3926 | 237 | 610 | 690 | 1 | 2 | 993 | 261 | 198 | 59 | 34 |
| cluster3927 | 33 | 78 | 259 | 7 | 82 | 237 | 111 | 43 | 51 | 90 |
| cluster3928 | 111 | 208 | 148 | 35 | 119 | 233 | 190 | 59 | 77 | 388 |
| cluster3929 | 4 | 31 | 87 | 17 | 130 | 207 | 155 | 121 | 157 | 279 |
| cluster3930 | 25 | 69 | 95 | 99 | 211 | 265 | 105 | 47 | 15 | 107 |
| cluster3931 | 14 | 21 | 1732 | 26 | 102 | 296 | 72 | 79 | 211 | 344 |
| cluster3932 | 18 | 36 | 141 | 66 | 14 | 18 | 1336 | 21 | 73 | 110 |
| cluster3933 | 47 | 86 | 185 | 12 | 37 | 63 | 225 | 100 | 140 | 578 |
| cluster3934 | 506 | 1530 | 3028 | 814 | 2208 | 3660 | 1692 | 439 | 2046 | 4719 |
| cluster3935 | 2 | 22 | 62 | 0 | 34 | 25 | 34 | 14 | 10 | 114 |
| cluster3936 | 34 | 51 | 252 | 2 | 293 | 478 | 162 | 46 | 36 | 99 |

| | | | | | | | | | | |
|---|---|---|---|---|---|---|---|---|---|---|
| cluster3937 | 7 | 20 | 51 | 91 | 55 | 143 | 62 | 20 | 36 | 90 |
| cluster3938 | 19 | 41 | 66 | 0 | 71 | 111 | 45 | 31 | 10 | 60 |
| cluster3939 | 24 | 75 | 133 | 6 | 188 | 187 | 137 | 32 | 74 | 126 |
| cluster3940 | 38 | 125 | 161 | 243 | 633 | 841 | 217 | 74 | 158 | 291 |
| cluster3941 | 73 | 179 | 519 | 222 | 114 | 214 | 278 | 178 | 539 | 743 |
| cluster3942 | 139 | 367 | 712 | 32 | 189 | 132 | 312 | 179 | 642 | 997 |
| cluster3943 | 13 | 72 | 382 | 0 | 244 | 346 | 169 | 54 | 153 | 212 |
| cluster3944 | 38 | 114 | 301 | 11 | 261 | 384 | 114 | 110 | 48 | 220 |
| cluster3945 | 109 | 281 | 1377 | 2 | 130 | 199 | 226 | 169 | 206 | 381 |
| cluster3946 | 120 | 303 | 831 | 1 | 344 | 336 | 379 | 194 | 459 | 877 |
| cluster3947 | 25 | 62 | 121 | 30 | 170 | 255 | 139 | 39 | 120 | 160 |
| cluster3948 | 364 | 1264 | 652 | 77 | 991 | 1368 | 1215 | 876 | 835 | 1456 |
| cluster3949 | 10 | 19 | 212 | 4 | 45 | 144 | 159 | 25 | 175 | 282 |
| cluster3950 | 101 | 280 | 293 | 471 | 887 | 1783 | 615 | 516 | 109 | 511 |
| cluster3951 | 15 | 37 | 180 | 0 | 125 | 240 | 142 | 38 | 55 | 100 |
| cluster3952 | 79 | 159 | 496 | 101 | 655 | 1699 | 169 | 135 | 202 | 386 |
| cluster3953 | 510 | 1253 | 1414 | 327 | 564 | 306 | 971 | 798 | 1244 | 2777 |
| cluster3954 | 8 | 23 | 65 | 0 | 98 | 163 | 286 | 29 | 33 | 164 |
| cluster3955 | 48 | 152 | 52 | 0 | 146 | 252 | 253 | 274 | 85 | 142 |
| cluster3956 | 42 | 88 | 226 | 148 | 403 | 559 | 89 | 78 | 41 | 133 |
| cluster3957 | 9 | 30 | 81 | 11 | 174 | 279 | 174 | 26 | 21 | 74 |
| cluster3958 | 4 | 21 | 7 | 1 | 37 | 8 | 195 | 10 | 20 | 45 |
| cluster3959 | 7 | 23 | 26 | 63 | 445 | 95 | 699 | 102 | 200 | 470 |
| cluster3960 | 18 | 30 | 498 | 11 | 135 | 119 | 949 | 5 | 369 | 179 |
| cluster3961 | 182 | 621 | 1507 | 0 | 751 | 1527 | 420 | 400 | 611 | 1215 |
| cluster3962 | 67 | 163 | 343 | 38 | 226 | 347 | 127 | 63 | 317 | 461 |
| cluster3963 | 264 | 677 | 543 | 0 | 238 | 448 | 164 | 311 | 152 | 711 |
| cluster3964 | 8 | 24 | 140 | 1 | 325 | 242 | 650 | 37 | 310 | 431 |
| cluster3965 | 6 | 16 | 94 | 0 | 18 | 9 | 35 | 12 | 10 | 16 |
| cluster3966 | 44 | 117 | 365 | 132 | 337 | 470 | 240 | 93 | 153 | 277 |
| cluster3967 | 15 | 26 | 56 | 97 | 44 | 59 | 114 | 11 | 80 | 105 |

| | | | | | | | | | | |
|---|---|---|---|---|---|---|---|---|---|---|
| cluster3968 | 14 | 29 | 99 | 3 | 134 | 296 | 43 | 54 | 38 | 114 |
| cluster3969 | 224 | 638 | 1834 | 657 | 465 | 795 | 960 | 427 | 816 | 1346 |
| cluster3970 | 4 | 27 | 97 | 3 | 249 | 385 | 94 | 34 | 56 | 99 |
| cluster3971 | 98 | 275 | 216 | 5 | 551 | 827 | 140 | 67 | 42 | 117 |
| cluster3972 | 27 | 104 | 149 | 48 | 301 | 284 | 436 | 97 | 35 | 180 |
| cluster3973 | 5790 | 21849 | 34171 | 1187 | 14266 | 19434 | 21997 | 21992 | 24977 | 41322 |
| cluster3974 | 5 | 19 | 104 | 0 | 484 | 687 | 439 | 38 | 82 | 247 |
| cluster3975 | 113 | 348 | 235 | 125 | 990 | 1310 | 550 | 155 | 38 | 115 |
| cluster3976 | 963 | 2356 | 973 | 1834 | 1002 | 1716 | 1726 | 1697 | 1097 | 1948 |
| cluster3977 | 719 | 2334 | 868 | 6 | 724 | 470 | 1078 | 460 | 725 | 3948 |
| cluster3978 | 92 | 216 | 414 | 120 | 130 | 237 | 317 | 249 | 61 | 252 |
| cluster3979 | 52 | 101 | 233 | 291 | 409 | 488 | 243 | 112 | 128 | 504 |
| cluster3980 | 49 | 137 | 312 | 95 | 362 | 439 | 257 | 92 | 107 | 452 |
| cluster3981 | 14 | 76 | 108 | 44 | 884 | 1125 | 163 | 55 | 128 | 246 |
| cluster3982 | 37 | 130 | 216 | 24 | 268 | 308 | 253 | 87 | 122 | 213 |
| cluster3983 | 3 | 34 | 60 | 3 | 255 | 260 | 81 | 22 | 18 | 44 |
| cluster3984 | 73 | 233 | 223 | 0 | 349 | 2 | 183 | 193 | 412 | 801 |
| cluster3985 | 10 | 28 | 60 | 56 | 122 | 227 | 66 | 26 | 33 | 94 |
| cluster3986 | 24 | 57 | 157 | 22 | 107 | 167 | 57 | 59 | 84 | 141 |
| cluster3987 | 30 | 45 | 112 | 172 | 302 | 480 | 62 | 46 | 82 | 174 |
| cluster3988 | 47 | 137 | 554 | 130 | 261 | 304 | 380 | 166 | 277 | 481 |
| cluster3989 | 4 | 16 | 47 | 62 | 49 | 62 | 76 | 9 | 108 | 200 |
| cluster3990 | 9 | 24 | 94 | 52 | 69 | 91 | 39 | 16 | 75 | 141 |
| cluster3991 | 23 | 61 | 42 | 0 | 83 | 104 | 38 | 34 | 14 | 82 |
| cluster3992 | 16 | 32 | 214 | 284 | 167 | 307 | 261 | 81 | 142 | 248 |
| cluster3993 | 67 | 153 | 147 | 23 | 144 | 255 | 178 | 68 | 82 | 164 |
| cluster3994 | 243 | 566 | 229 | 3 | 66 | 74 | 169 | 391 | 449 | 531 |
| cluster3995 | 30 | 96 | 126 | 3 | 36 | 13 | 61 | 115 | 120 | 214 |
| cluster3996 | 18 | 89 | 186 | 14 | 106 | 179 | 71 | 39 | 115 | 180 |
| cluster3997 | 39 | 142 | 286 | 0 | 203 | 417 | 69 | 66 | 119 | 220 |
| cluster3998 | 30 | 66 | 28 | 14 | 28 | 247 | 49 | 0 | 7 | 81 |

| | | | | | | | | | | |
|---|---|---|---|---|---|---|---|---|---|---|
| cluster3999 | 7 | 24 | 91 | 0 | 41 | 53 | 133 | 32 | 36 | 74 |
| cluster4000 | 14 | 31 | 102 | 10 | 79 | 118 | 16 | 6 | 23 | 73 |
| cluster4001 | 247 | 735 | 948 | 50 | 802 | 651 | 1003 | 2180 | 311 | 897 |
| cluster4002 | 7 | 37 | 48 | 8 | 219 | 325 | 64 | 12 | 23 | 33 |
| cluster4003 | 113 | 258 | 659 | 0 | 229 | 415 | 55 | 126 | 768 | 698 |
| cluster4004 | 371 | 977 | 3236 | 397 | 947 | 1820 | 930 | 958 | 1913 | 2692 |
| cluster4005 | 27 | 79 | 150 | 2 | 75 | 174 | 73 | 117 | 107 | 325 |
| cluster4006 | 186 | 479 | 1177 | 128 | 456 | 513 | 222 | 350 | 1041 | 1006 |
| cluster4007 | 30 | 108 | 280 | 55 | 97 | 214 | 103 | 57 | 84 | 201 |
| cluster4008 | 16 | 37 | 49 | 0 | 97 | 142 | 34 | 20 | 34 | 62 |
| cluster4009 | 1 | 11 | 23 | 0 | 21 | 38 | 18 | 4 | 43 | 50 |
| cluster4010 | 11 | 46 | 69 | 0 | 126 | 160 | 113 | 16 | 11 | 18 |
| cluster4011 | 2253 | 6749 | 5762 | 459 | 61 | 83 | 4405 | 5621 | 3778 | 8386 |
| cluster4012 | 5 | 17 | 175 | 19 | 80 | 71 | 14 | 4 | 50 | 65 |
| cluster4013 | 7 | 8 | 9 | 0 | 14 | 5 | 518 | 0 | 25 | 18 |
| cluster4014 | 269 | 918 | 1051 | 15 | 1892 | 2710 | 490 | 843 | 329 | 1363 |
| cluster4015 | 5019 | 16095 | 30155 | 6045 | 2798 | 5055 | 10039 | 14620 | 5706 | 25938 |
| cluster4016 | 7 | 48 | 70 | 59 | 313 | 533 | 71 | 50 | 46 | 72 |
| cluster4017 | 12 | 40 | 122 | 0 | 53 | 119 | 71 | 20 | 18 | 86 |
| cluster4018 | 7 | 10 | 52 | 1 | 25 | 64 | 32 | 3 | 51 | 33 |
| cluster4019 | 508 | 1453 | 3827 | 54 | 426 | 6605 | 115 | 188 | 1091 | 3332 |
| cluster4020 | 92 | 163 | 220 | 249 | 197 | 313 | 198 | 138 | 249 | 421 |
| cluster4021 | 4 | 19 | 110 | 0 | 43 | 94 | 30 | 23 | 38 | 75 |
| cluster4022 | 3954 | 12391 | 17225 | 1461 | 10434 | 17361 | 9866 | 11466 | 2507 | 13771 |
| cluster4023 | 8 | 11 | 76 | 1 | 17 | 16 | 71 | 6 | 83 | 52 |
| cluster4024 | 6 | 19 | 58 | 15 | 131 | 145 | 97 | 19 | 25 | 75 |
| cluster4025 | 101 | 251 | 670 | 227 | 379 | 636 | 149 | 287 | 400 | 1032 |
| cluster4026 | 46 | 110 | 189 | 0 | 71 | 128 | 58 | 93 | 63 | 112 |
| cluster4027 | 42 | 100 | 135 | 0 | 254 | 276 | 192 | 33 | 55 | 126 |
| cluster4028 | 11 | 37 | 238 | 21 | 138 | 255 | 139 | 48 | 89 | 255 |
| cluster4029 | 125 | 382 | 1947 | 0 | 36 | 125 | 270 | 342 | 231 | 370 |

| | | | | | | | | | | |
|---|---|---|---|---|---|---|---|---|---|---|
| cluster4030 | 451 | 1471 | 944 | 27 | 2122 | 2975 | 1341 | 1211 | 255 | 1538 |
| cluster4031 | 50 | 180 | 192 | 584 | 128 | 201 | 464 | 184 | 47 | 292 |
| cluster4032 | 4 | 17 | 82 | 10 | 20 | 48 | 58 | 3 | 68 | 137 |
| cluster4033 | 4 | 12 | 30 | 0 | 0 | 22 | 36 | 23 | 7 | 15 |
| cluster4034 | 4574 | 11945 | 956 | 6 | 294 | 330 | 9088 | 9108 | 5261 | 17543 |
| cluster4035 | 11 | 26 | 56 | 92 | 69 | 77 | 56 | 11 | 33 | 84 |
| cluster4036 | 17 | 54 | 98 | 27 | 130 | 175 | 97 | 48 | 75 | 123 |
| cluster4037 | 37 | 107 | 198 | 51 | 285 | 528 | 106 | 87 | 142 | 242 |
| cluster4038 | 44 | 125 | 118 | 0 | 243 | 598 | 176 | 101 | 52 | 170 |
| cluster4039 | 11 | 24 | 361 | 87 | 225 | 129 | 293 | 14 | 307 | 426 |
| cluster4040 | 13 | 45 | 40 | 4 | 54 | 71 | 26 | 25 | 8 | 20 |
| cluster4041 | 1525 | 3909 | 8197 | 88 | 3730 | 3704 | 607 | 2677 | 2631 | 6565 |
| cluster4042 | 490 | 1531 | 1734 | 5 | 609 | 1183 | 1024 | 986 | 244 | 899 |
| cluster4043 | 4 | 15 | 4 | 0 | 10 | 8 | 12 | 3 | 8 | 27 |
| cluster4044 | 35 | 83 | 120 | 9 | 244 | 401 | 71 | 47 | 62 | 130 |
| cluster4045 | 8 | 25 | 95 | 1 | 439 | 567 | 341 | 33 | 58 | 103 |
| cluster4046 | 262 | 159 | 82 | 31 | 12 | 5 | 1321 | 92 | 89 | 313 |
| cluster4047 | 5 | 12 | 60 | 0 | 363 | 426 | 122 | 22 | 78 | 189 |
| cluster4048 | 9 | 21 | 140 | 28 | 331 | 124 | 2021 | 19 | 71 | 130 |
| cluster4049 | 5143 | 12658 | 18317 | 146 | 12 | 6 | 16544 | 19731 | 40496 | 27093 |
| cluster4050 | 119 | 396 | 339 | 2 | 479 | 885 | 282 | 302 | 178 | 464 |
| cluster4051 | 2 | 17 | 66 | 6 | 58 | 198 | 83 | 3 | 22 | 31 |
| cluster4052 | 12 | 26 | 123 | 209 | 212 | 195 | 65 | 17 | 64 | 121 |
| cluster4053 | 18 | 40 | 101 | 11 | 73 | 126 | 178 | 119 | 25 | 157 |
| cluster4054 | 6 | 22 | 63 | 1 | 117 | 234 | 66 | 14 | 26 | 43 |
| cluster4055 | 60 | 244 | 123 | 28 | 107 | 193 | 339 | 263 | 39 | 153 |
| cluster4056 | 164 | 397 | 88 | 64 | 30 | 193 | 204 | 136 | 224 | 416 |
| cluster4057 | 34 | 93 | 155 | 0 | 297 | 539 | 318 | 95 | 43 | 91 |
| cluster4058 | 125 | 284 | 131 | 24 | 310 | 440 | 298 | 82 | 66 | 142 |
| cluster4059 | 841 | 1770 | 1051 | 0 | 2125 | 3825 | 1940 | 1979 | 1529 | 2369 |
| cluster4060 | 3 | 16 | 111 | 4 | 125 | 187 | 15 | 9 | 21 | 43 |

| | | | | | | | | | | |
|---|---|---|---|---|---|---|---|---|---|---|
| cluster4061 | 18 | 51 | 185 | 0 | 86 | 140 | 63 | 27 | 41 | 71 |
| cluster4062 | 2 | 10 | 191 | 3 | 5 | 17 | 203 | 2 | 131 | 144 |
| cluster4063 | 47 | 134 | 603 | 0 | 9 | 590 | 138 | 251 | 171 | 310 |
| cluster4064 | 16 | 44 | 153 | 6 | 182 | 290 | 73 | 17 | 76 | 104 |
| cluster4065 | 17 | 83 | 198 | 26 | 206 | 318 | 162 | 65 | 155 | 206 |
| cluster4066 | 22 | 50 | 47 | 1 | 25 | 103 | 100 | 42 | 26 | 57 |
| cluster4067 | 4 | 29 | 299 | 0 | 147 | 3764 | 108 | 37 | 172 | 232 |
| cluster4068 | 11 | 29 | 37 | 0 | 95 | 202 | 128 | 33 | 24 | 45 |
| cluster4069 | 488 | 1202 | 402 | 117 | 1640 | 3813 | 331 | 863 | 397 | 1183 |
| cluster4070 | 2 | 13 | 68 | 1 | 0 | 2 | 183 | 8 | 19 | 45 |
| cluster4071 | 16 | 72 | 96 | 0 | 382 | 578 | 512 | 46 | 35 | 233 |
| cluster4072 | 146 | 401 | 1608 | 497 | 866 | 1927 | 296 | 444 | 845 | 1107 |
| cluster4073 | 11 | 21 | 220 | 0 | 575 | 916 | 68 | 21 | 105 | 192 |
| cluster4074 | 6 | 24 | 38 | 0 | 136 | 192 | 24 | 2 | 2 | 11 |
| cluster4075 | 6 | 20 | 16 | 12 | 6 | 18 | 1 | 0 | 36 | 27 |
| cluster4076 | 7 | 25 | 186 | 521 | 103 | 206 | 95 | 14 | 97 | 110 |
| cluster4077 | 9 | 18 | 148 | 36 | 164 | 178 | 255 | 25 | 188 | 115 |
| cluster4078 | 152 | 473 | 641 | 1524 | 686 | 1015 | 201 | 165 | 82 | 367 |
| cluster4079 | 24 | 44 | 136 | 175 | 558 | 497 | 85 | 49 | 68 | 150 |
| cluster4080 | 1662 | 3942 | 952 | 791 | 51 | 41 | 2466 | 2718 | 1571 | 2353 |
| cluster4081 | 40 | 110 | 238 | 39 | 306 | 633 | 216 | 79 | 47 | 93 |
| cluster4082 | 4 | 14 | 38 | 0 | 28 | 46 | 35 | 0 | 17 | 19 |
| cluster4083 | 44 | 144 | 94 | 0 | 244 | 360 | 67 | 56 | 92 | 109 |
| cluster4084 | 4326 | 9427 | 7858 | 11197 | 4123 | 5615 | 8143 | 9610 | 5698 | 15786 |
| cluster4085 | 3935 | 15064 | 17583 | 3469 | 918 | 1707 | 16101 | 15811 | 21888 | 24628 |
| cluster4086 | 4 | 29 | 122 | 0 | 43 | 49 | 15 | 15 | 7 | 28 |
| cluster4087 | 16 | 64 | 103 | 6 | 2469 | 4268 | 129 | 82 | 52 | 102 |
| cluster4088 | 4 | 11 | 38 | 3 | 71 | 176 | 177 | 28 | 120 | 71 |
| cluster4089 | 305 | 791 | 61047 | 500 | 2914 | 2803 | 2106 | 5842 | 4530 | 10644 |
| cluster4090 | 2 | 16 | 86 | 0 | 3 | 1 | 73 | 78 | 80 | 214 |
| cluster4091 | 1 | 10 | 105 | 3 | 3 | 5 | 101 | 1 | 449 | 320 |

| | | | | | | | | | | |
|---|---|---|---|---|---|---|---|---|---|---|
| cluster4092 | 1 | 13 | 104 | 0 | 51 | 51 | 71 | 27 | 40 | 159 |
| cluster4093 | 18 | 58 | 244 | 0 | 113 | 1270 | 142 | 32 | 183 | 209 |
| cluster4094 | 3 | 7 | 46 | 5 | 22 | 38 | 21 | 0 | 25 | 29 |
| cluster4095 | 2 | 12 | 18 | 61 | 83 | 86 | 38 | 6 | 15 | 21 |
| cluster4096 | 78 | 219 | 288 | 117 | 301 | 532 | 153 | 264 | 141 | 445 |
| cluster4097 | 4545 | 10584 | 20010 | 869 | 14 | 4 | 9498 | 8028 | 9821 | 15079 |
| cluster4098 | 7 | 12 | 83 | 210 | 183 | 295 | 60 | 20 | 24 | 58 |
| cluster4099 | 23 | 91 | 168 | 41 | 442 | 502 | 96 | 37 | 73 | 225 |
| cluster4100 | 9 | 21 | 48 | 0 | 120 | 140 | 48 | 24 | 18 | 64 |
| cluster4101 | 5 | 8 | 176 | 444 | 11 | 515 | 17 | 14 | 5 | 14 |
| cluster4102 | 7 | 10 | 67 | 2 | 28 | 23 | 20 | 13 | 28 | 64 |
| cluster4103 | 2 | 10 | 126 | 174 | 85 | 178 | 83 | 24 | 169 | 246 |
| cluster4104 | 37 | 121 | 94 | 0 | 263 | 474 | 198 | 74 | 43 | 83 |
| cluster4105 | 67 | 161 | 220 | 79 | 93 | 191 | 15 | 20 | 71 | 148 |
| cluster4106 | 0 | 5 | 105 | 1 | 62 | 19 | 144 | 19 | 18 | 49 |
| cluster4107 | 8 | 18 | 21 | 0 | 134 | 166 | 44 | 4 | 16 | 24 |
| cluster4108 | 2 | 10 | 5 | 36 | 36 | 18 | 24 | 31 | 4 | 11 |
| cluster4109 | 4 | 11 | 134 | 1 | 109 | 176 | 87 | 9 | 31 | 65 |
| cluster4110 | 7 | 38 | 114 | 0 | 107 | 168 | 43 | 40 | 50 | 109 |
| cluster4111 | 3 | 13 | 4 | 0 | 1 | 9 | 95 | 7 | 28 | 130 |
| cluster4112 | 2 | 5 | 14 | 46 | 42 | 86 | 5 | 2 | 5 | 35 |
| cluster4113 | 5608 | 11326 | 20383 | 7122 | 4781 | 5237 | 10479 | 11053 | 4558 | 12753 |
| cluster4114 | 2 | 6 | 57 | 0 | 854 | 544 | 118 | 3 | 240 | 389 |
| cluster4115 | 5 | 7 | 123 | 0 | 19 | 43 | 203 | 9 | 48 | 42 |
| cluster4116 | 1 | 6 | 11 | 44 | 38 | 18 | 645 | 22 | 0 | 11 |
| cluster4117 | 3 | 12 | 6 | 0 | 137 | 27 | 144 | 0 | 12 | 343 |
| cluster4118 | 0 | 7 | 24 | 0 | 24 | 89 | 601 | 104 | 470 | 501 |
| cluster4119 | 23 | 61 | 80 | 0 | 961 | 761 | 101 | 18 | 22 | 80 |
| cluster4120 | 1 | 5 | 32 | 2 | 5 | 1 | 92 | 2 | 77 | 25 |
| cluster4121 | 4 | 9 | 35 | 11 | 60 | 140 | 11 | 12 | 6 | 29 |
| cluster4122 | 2 | 11 | 33 | 1 | 3 | 11 | 213 | 2 | 14 | 45 |

| | | | | | | | | | | |
|---|---|---|---|---|---|---|---|---|---|---|
| cluster4123 | 5 | 11 | 24 | 0 | 6 | 5 | 2 | 3 | 9 | 1 |
| cluster4124 | 1 | 5 | 10 | 0 | 5 | 2 | 11 | 5 | 6 | 17 |
| cluster4125 | 252 | 3 | 117 | 169 | 16 | 0 | 84 | 2 | 9 | 32 |
| cluster4126 | 1 | 5 | 102 | 0 | 8 | 111 | 28 | 35 | 12 | 41 |
| cluster4127 | 1 | 3 | 173 | 0 | 11 | 71 | 286 | 9 | 2 | 7 |
| cluster4128 | 2 | 4 | 15 | 0 | 71 | 149 | 171 | 35 | 17 | 65 |

[1]Chom: Replicates of *C. hominiv*

[2]Dhom: Replicates of *D. hominis*

[3]Oovi: Replicates of *O. ovis*

[4]Lcup: Replicates of *L. cuprina*

[5]Cmeg: Replicates of *C. megace*

**Supplementary Table 2**- Raw counts RSEM.

| ID | Chom1[1] | Chom2[1] | Dhom1[2] | Dhom2[2] | Oovi1[3] | Oovi2[3] | Lcup1[4] | Lcup2[4] | Cmeg1[5] | Cmeg2[5] |
|---|---|---|---|---|---|---|---|---|---|---|
| cluster1127 | 96 | 280 | 1246 | 0 | 208 | 181 | 255 | 69 | 342 | 381 |
| cluster2853 | 597 | 1352 | 3041 | 6540 | 1885 | 2244 | 3890 | 1153 | 3083 | 6974 |
| cluster2854 | 495 | 1247 | 3667 | 15 | 4969 | 963 | 323 | 178 | 1442 | 2132 |
| cluster2855 | 838 | 1935 | 4726 | 0 | 4232 | 4636 | 2605 | 684 | 3655 | 6349 |
| cluster2856 | 573 | 842 | 3601 | 0 | 5071 | 4435 | 4132 | 431 | 4535 | 3344 |
| cluster2857 | 40250 | 124542 | 181146 | 52211 | 268078 | 266748 | 259965 | 213578 | 245236 | 361572 |
| cluster2858 | 385 | 915 | 1508 | 438 | 771 | 696 | 948 | 109 | 3511 | 2226 |
| cluster2859 | 179 | 440 | 1090 | 44 | 334 | 855 | 560 | 345 | 531 | 972 |
| cluster2860 | 364 | 768 | 2619 | 3966 | 2058 | 1831 | 2343 | 402 | 1171 | 2234 |
| cluster2861 | 236 | 563 | 1125 | 1251 | 997 | 1419 | 264 | 76 | 1191 | 2687 |
| cluster2862 | 242 | 608 | 358 | 766 | 379 | 486 | 962 | 262 | 674 | 1434 |
| cluster2863 | 287 | 708 | 2303 | 680 | 932 | 2139 | 1099 | 259 | 758 | 1380 |
| cluster2864 | 184 | 376 | 237 | 514 | 1309 | 1158 | 936 | 143 | 712 | 1349 |
| cluster2865 | 261 | 653 | 299 | 777 | 1431 | 2197 | 1047 | 202 | 1080 | 1441 |
| cluster2866 | 97 | 298 | 4815 | 0 | 443 | 638 | 1418 | 221 | 1288 | 1988 |
| cluster2867 | 284 | 606 | 1361 | 0 | 145 | 150 | 324 | 158 | 164 | 404 |
| cluster2869 | 149 | 368 | 819 | 591 | 46 | 50 | 574 | 159 | 625 | 813 |
| cluster2870 | 138 | 369 | 2950 | 17 | 636 | 1041 | 825 | 184 | 791 | 1113 |
| cluster2871 | 101 | 270 | 575 | 38 | 577 | 921 | 1247 | 281 | 914 | 1395 |
| cluster2872 | 403 | 986 | 3678 | 11 | 1264 | 948 | 812 | 501 | 2057 | 3892 |
| cluster2873 | 172 | 425 | 639 | 1179 | 669 | 630 | 602 | 196 | 560 | 1384 |
| cluster2874 | 307 | 664 | 1873 | 15 | 1562 | 1383 | 1062 | 296 | 1537 | 2541 |
| cluster2875 | 476 | 1047 | 2193 | 4269 | 3301 | 3916 | 7775 | 810 | 4512 | 6758 |
| cluster2876 | 709 | 2025 | 25 | 224 | 14324 | 269 | 1407 | 858 | 773 | 2475 |
| cluster2877 | 297 | 784 | 1339 | 553 | 1956 | 1822 | 851 | 340 | 1736 | 2786 |
| cluster2878 | 218 | 392 | 2095 | 824 | 408 | 1702 | 1714 | 60 | 4186 | 1974 |
| cluster2879 | 5069 | 15727 | 20917 | 3962 | 10399 | 8610 | 6215 | 1715 | 12303 | 20376 |
| cluster2880 | 680 | 1692 | 7431 | 0 | 1190 | 1656 | 1352 | 567 | 2386 | 4190 |

| | | | | | | | | | | |
|---|---|---|---|---|---|---|---|---|---|---|
| cluster2881 | 394 | 777 | 787 | 2469 | 2950 | 2798 | 987 | 253 | 2080 | 2902 |
| cluster2882 | 87 | 236 | 580 | 299 | 330 | 498 | 305 | 66 | 403 | 955 |
| cluster2883 | 815 | 2192 | 1699 | 2551 | 2555 | 4580 | 2634 | 1894 | 2588 | 7054 |
| cluster2884 | 332 | 862 | 3704 | 15 | 1196 | 944 | 833 | 211 | 1285 | 1672 |
| cluster2885 | 150 | 356 | 1999 | 0 | 414 | 474 | 752 | 191 | 2144 | 2430 |
| cluster2886 | 256 | 657 | 524 | 1200 | 578 | 824 | 1793 | 126 | 625 | 1485 |
| cluster2887 | 197 | 491 | 787 | 0 | 1661 | 1855 | 942 | 387 | 566 | 1020 |
| cluster2888 | 122 | 339 | 155 | 0 | 109 | 1688 | 256 | 438 | 404 | 267 |
| cluster2889 | 158 | 406 | 1153 | 0 | 590 | 924 | 377 | 178 | 629 | 857 |
| cluster2890 | 304 | 823 | 2576 | 1150 | 786 | 878 | 2690 | 589 | 2149 | 3670 |
| cluster2891 | 501 | 1009 | 4825 | 0 | 1391 | 1226 | 2885 | 854 | 1540 | 3579 |
| cluster2892 | 177 | 492 | 287 | 530 | 1557 | 2168 | 695 | 207 | 478 | 1259 |
| cluster2893 | 1614 | 2754 | 411 | 0 | 7865 | 316 | 1307 | 379 | 298 | 1892 |
| cluster2894 | 1089 | 2485 | 3433 | 57 | 182 | 68 | 2029 | 371 | 3789 | 10895 |
| cluster2895 | 107 | 270 | 1611 | 0 | 218 | 332 | 667 | 173 | 652 | 848 |
| cluster2896 | 429 | 978 | 1623 | 0 | 834 | 1272 | 573 | 324 | 1128 | 1392 |
| cluster2897 | 125 | 281 | 1691 | 22 | 427 | 694 | 1071 | 91 | 1020 | 1168 |
| cluster2898 | 51 | 193 | 96 | 179 | 361 | 149 | 281 | 8 | 179 | 322 |
| cluster2899 | 236 | 764 | 1134 | 2209 | 297 | 368 | 723 | 176 | 914 | 1872 |
| cluster2900 | 82 | 227 | 343 | 962 | 110 | 192 | 1020 | 163 | 696 | 1070 |
| cluster2901 | 201 | 602 | 1447 | 0 | 352 | 363 | 1827 | 202 | 2037 | 1522 |
| cluster2902 | 84 | 207 | 4268 | 0 | 855 | 1814 | 261 | 90 | 108 | 347 |
| cluster2903 | 85 | 233 | 339 | 908 | 382 | 535 | 3038 | 80 | 768 | 1406 |
| cluster2904 | 339 | 673 | 2325 | 728 | 168 | 214 | 743 | 114 | 4707 | 2177 |
| cluster2905 | 337 | 904 | 1225 | 2490 | 1362 | 1545 | 2289 | 481 | 4463 | 5251 |
| cluster2906 | 207 | 560 | 558 | 1712 | 784 | 949 | 1191 | 197 | 3024 | 2022 |
| cluster2907 | 154 | 425 | 2467 | 0 | 572 | 647 | 1286 | 232 | 853 | 1378 |
| cluster2908 | 224 | 658 | 2472 | 186 | 232 | 427 | 521 | 99 | 1667 | 2410 |
| cluster2909 | 64 | 153 | 547 | 49 | 377 | 532 | 407 | 121 | 390 | 535 |
| cluster2910 | 229 | 678 | 2879 | 0 | 800 | 932 | 2024 | 294 | 1670 | 2696 |
| cluster2911 | 2412 | 6531 | 3123 | 1561 | 4146 | 13635 | 20297 | 3877 | 8695 | 15108 |

| | | | | | | | | | | |
|---|---|---|---|---|---|---|---|---|---|---|
| cluster2912 | 212 | 378 | 1483 | 27 | 2490 | 3066 | 931 | 157 | 539 | 1008 |
| cluster2913 | 270 | 773 | 2477 | 31 | 472 | 643 | 1247 | 309 | 534 | 1630 |
| cluster2914 | 4089 | 9176 | 865 | 963 | 3103 | 1208 | 4504 | 1291 | 1729 | 11407 |
| cluster2915 | 81 | 231 | 590 | 3 | 49 | 95 | 520 | 68 | 205 | 255 |
| cluster2916 | 78 | 182 | 115 | 402 | 392 | 160 | 325 | 20 | 279 | 600 |
| cluster2917 | 311 | 784 | 1547 | 0 | 254 | 1545 | 557 | 194 | 795 | 1116 |
| cluster2918 | 139 | 364 | 1136 | 11 | 198 | 227 | 232 | 72 | 815 | 1093 |
| cluster2919 | 115 | 407 | 226 | 599 | 750 | 584 | 668 | 113 | 1104 | 1639 |
| cluster2920 | 144 | 393 | 2357 | 3 | 458 | 423 | 1089 | 209 | 821 | 1304 |
| cluster2921 | 136 | 409 | 3239 | 46 | 738 | 760 | 291 | 65 | 890 | 1622 |
| cluster2922 | 173 | 537 | 2429 | 1 | 2751 | 2316 | 1990 | 304 | 1852 | 2302 |
| cluster2923 | 276 | 810 | 4217 | 246 | 1450 | 1780 | 2353 | 382 | 2783 | 4673 |
| cluster2924 | 116 | 250 | 2933 | 0 | 262 | 3879 | 849 | 180 | 2521 | 6241 |
| cluster2925 | 79 | 236 | 1393 | 0 | 63 | 93 | 256 | 89 | 1096 | 1663 |
| cluster2926 | 217 | 649 | 420 | 1091 | 896 | 1322 | 724 | 171 | 170 | 1148 |
| cluster2927 | 85 | 198 | 768 | 0 | 459 | 930 | 232 | 112 | 265 | 516 |
| cluster2928 | 41 | 88 | 309 | 3 | 708 | 730 | 173 | 126 | 141 | 221 |
| cluster2929 | 214 | 455 | 2157 | 1 | 828 | 746 | 964 | 159 | 638 | 1289 |
| cluster2930 | 105 | 284 | 2201 | 21 | 1191 | 2447 | 572 | 188 | 807 | 1075 |
| cluster2931 | 124 | 287 | 1139 | 0 | 246 | 333 | 564 | 142 | 678 | 716 |
| cluster2932 | 95 | 263 | 904 | 55 | 295 | 353 | 942 | 38 | 280 | 463 |
| cluster2933 | 64 | 177 | 554 | 1016 | 1024 | 366 | 840 | 79 | 1812 | 2051 |
| cluster2934 | 126 | 256 | 669 | 28 | 206 | 397 | 643 | 65 | 983 | 1073 |
| cluster2935 | 716 | 1551 | 1141 | 2681 | 2090 | 1391 | 1027 | 691 | 997 | 2170 |
| cluster2936 | 142 | 369 | 632 | 0 | 635 | 1123 | 191 | 85 | 234 | 651 |
| cluster2937 | 1351 | 2566 | 1236 | 324 | 1174 | 1494 | 1322 | 410 | 2315 | 4984 |
| cluster2938 | 319 | 724 | 559 | 817 | 727 | 885 | 996 | 578 | 608 | 1968 |
| cluster2939 | 186 | 339 | 38 | 115 | 327 | 258 | 357 | 98 | 377 | 546 |
| cluster2940 | 469 | 1284 | 2649 | 0 | 1006 | 1175 | 2134 | 649 | 1169 | 3110 |
| cluster2941 | 225 | 830 | 1394 | 535 | 2097 | 2912 | 1979 | 1258 | 3005 | 4022 |
| cluster2942 | 108 | 242 | 1432 | 1 | 980 | 791 | 380 | 80 | 737 | 1026 |

| | | | | | | | | | | |
|---|---|---|---|---|---|---|---|---|---|---|
| cluster2943 | 423 | 1020 | 1134 | 2474 | 2778 | 3273 | 927 | 141 | 3084 | 4016 |
| cluster2944 | 50 | 151 | 1466 | 3 | 285 | 382 | 921 | 119 | 490 | 811 |
| cluster2945 | 1046 | 2431 | 9046 | 0 | 1932 | 1966 | 3916 | 1134 | 2623 | 4338 |
| cluster2946 | 652 | 1940 | 5028 | 8838 | 5529 | 6783 | 6572 | 1379 | 9563 | 18397 |
| cluster2947 | 41 | 114 | 667 | 129 | 486 | 546 | 264 | 29 | 293 | 432 |
| cluster2948 | 42 | 143 | 755 | 1797 | 406 | 436 | 526 | 90 | 1035 | 1117 |
| cluster2949 | 148 | 488 | 364 | 479 | 711 | 756 | 399 | 207 | 1463 | 1103 |
| cluster2950 | 73 | 259 | 402 | 452 | 350 | 440 | 619 | 110 | 629 | 1312 |
| cluster2951 | 114 | 319 | 742 | 0 | 1011 | 801 | 645 | 187 | 785 | 1139 |
| cluster2952 | 85 | 210 | 868 | 0 | 558 | 508 | 830 | 84 | 577 | 934 |
| cluster2953 | 79 | 204 | 637 | 268 | 263 | 206 | 570 | 103 | 571 | 1117 |
| cluster2954 | 1378 | 2815 | 2560 | 6103 | 1520 | 1751 | 5058 | 1756 | 5597 | 12288 |
| cluster2955 | 272 | 652 | 2389 | 58 | 368 | 1134 | 825 | 314 | 1225 | 1189 |
| cluster2956 | 89 | 234 | 272 | 207 | 496 | 656 | 1559 | 81 | 1027 | 1301 |
| cluster2957 | 58 | 175 | 504 | 0 | 82 | 131 | 594 | 20 | 536 | 823 |
| cluster2958 | 149 | 319 | 1175 | 21 | 538 | 485 | 792 | 235 | 605 | 1151 |
| cluster2959 | 131 | 344 | 116 | 262 | 826 | 803 | 656 | 143 | 1065 | 1861 |
| cluster2960 | 115 | 269 | 583 | 1682 | 272 | 468 | 523 | 143 | 573 | 1164 |
| cluster2961 | 181 | 505 | 521 | 1287 | 857 | 1106 | 435 | 136 | 219 | 436 |
| cluster2962 | 54 | 173 | 224 | 606 | 166 | 451 | 187 | 69 | 84 | 324 |
| cluster2963 | 601 | 1228 | 1155 | 3418 | 2218 | 502 | 1273 | 411 | 2905 | 2615 |
| cluster2964 | 205 | 446 | 554 | 1004 | 1175 | 1929 | 890 | 311 | 1237 | 2849 |
| cluster2965 | 172 | 336 | 3941 | 0 | 859 | 933 | 777 | 175 | 1543 | 2373 |
| cluster2966 | 165 | 455 | 1475 | 0 | 2367 | 1835 | 1094 | 354 | 1437 | 2091 |
| cluster2967 | 60 | 153 | 179 | 602 | 871 | 1013 | 429 | 30 | 1052 | 581 |
| cluster2968 | 244 | 579 | 567 | 451 | 705 | 1297 | 1120 | 223 | 1119 | 2066 |
| cluster2969 | 92 | 370 | 1782 | 2 | 1746 | 778 | 685 | 41 | 470 | 951 |
| cluster2970 | 215 | 467 | 1977 | 8 | 1092 | 1214 | 1057 | 322 | 671 | 1613 |
| cluster2971 | 258 | 750 | 1269 | 653 | 1190 | 1360 | 1567 | 851 | 1269 | 2355 |
| cluster2972 | 1237 | 2520 | 998 | 3020 | 1222 | 1142 | 3603 | 3709 | 2244 | 9900 |
| cluster2973 | 344 | 878 | 3364 | 0 | 2052 | 2121 | 1234 | 493 | 3583 | 3263 |

| | | | | | | | | | | |
|---|---|---|---|---|---|---|---|---|---|---|
| cluster2974 | 130 | 408 | 852 | 101 | 91 | 327 | 1593 | 179 | 997 | 1961 |
| cluster2975 | 394 | 1134 | 2264 | 293 | 2877 | 1962 | 980 | 486 | 1421 | 2362 |
| cluster2976 | 316 | 734 | 182 | 4 | 722 | 1514 | 1274 | 291 | 775 | 1266 |
| cluster2977 | 90 | 268 | 1373 | 1 | 813 | 1251 | 586 | 73 | 1404 | 1578 |
| cluster2978 | 243 | 558 | 1141 | 2768 | 1030 | 1261 | 2151 | 363 | 1268 | 3569 |
| cluster2979 | 137 | 410 | 2082 | 1 | 721 | 968 | 892 | 179 | 629 | 1079 |
| cluster2980 | 441 | 1103 | 328 | 0 | 100 | 154 | 392 | 561 | 306 | 1501 |
| cluster2981 | 80 | 274 | 1531 | 0 | 561 | 768 | 398 | 159 | 339 | 433 |
| cluster2982 | 143 | 486 | 2844 | 55 | 183 | 664 | 998 | 78 | 1875 | 2083 |
| cluster2983 | 206 | 484 | 1221 | 0 | 638 | 963 | 364 | 209 | 740 | 1481 |
| cluster2984 | 247 | 523 | 980 | 2236 | 557 | 742 | 439 | 347 | 952 | 1379 |
| cluster2985 | 173 | 271 | 147 | 366 | 529 | 764 | 620 | 44 | 510 | 494 |
| cluster2986 | 3287 | 7830 | 17128 | 1 | 19049 | 8689 | 9927 | 3986 | 11085 | 18650 |
| cluster2987 | 89 | 242 | 392 | 136 | 804 | 823 | 345 | 181 | 834 | 1090 |
| cluster2988 | 89 | 196 | 887 | 2 | 514 | 490 | 389 | 122 | 224 | 492 |
| cluster2989 | 35 | 102 | 155 | 385 | 210 | 333 | 239 | 20 | 489 | 418 |
| cluster2990 | 80 | 247 | 383 | 0 | 228 | 1023 | 297 | 165 | 708 | 700 |
| cluster2991 | 54 | 144 | 277 | 0 | 137 | 303 | 129 | 97 | 198 | 383 |
| cluster2992 | 107 | 267 | 1284 | 313 | 198 | 226 | 1360 | 219 | 1148 | 1315 |
| cluster2993 | 163 | 387 | 387 | 1313 | 713 | 1045 | 299 | 80 | 488 | 1377 |
| cluster2994 | 52 | 200 | 1378 | 25 | 520 | 1016 | 640 | 96 | 173 | 244 |
| cluster2995 | 5 | 15 | 82 | 1 | 48 | 158 | 29 | 11 | 12 | 52 |
| cluster2996 | 122 | 278 | 468 | 190 | 312 | 667 | 840 | 26 | 263 | 1243 |
| cluster2997 | 2268 | 5124 | 12337 | 11 | 4265 | 5282 | 6054 | 1654 | 9105 | 15736 |
| cluster2998 | 341 | 789 | 1077 | 0 | 837 | 773 | 1369 | 791 | 413 | 2385 |
| cluster2999 | 511 | 1121 | 1626 | 0 | 1749 | 1593 | 2309 | 1093 | 1270 | 2014 |
| cluster3000 | 16176 | 33060 | 38454 | 157 | 16955 | 19665 | 81542 | 43232 | 42664 | 105956 |
| cluster3001 | 110 | 288 | 849 | 355 | 794 | 1009 | 1809 | 284 | 812 | 1843 |
| cluster3002 | 88 | 216 | 1071 | 173 | 310 | 723 | 772 | 91 | 934 | 806 |
| cluster3003 | 113 | 289 | 1564 | 2 | 94 | 76 | 354 | 88 | 517 | 788 |
| cluster3004 | 281 | 818 | 497 | 764 | 374 | 592 | 522 | 103 | 3860 | 3617 |

| | | | | | | | | | | |
|---|---|---|---|---|---|---|---|---|---|---|
| cluster3005 | 327 | 1006 | 3821 | 76 | 843 | 1648 | 567 | 356 | 3148 | 4483 |
| cluster3006 | 290 | 625 | 12221 | 14985 | 4409 | 4255 | 2977 | 518 | 7105 | 5901 |
| cluster3007 | 66 | 221 | 659 | 0 | 285 | 532 | 248 | 103 | 194 | 515 |
| cluster3008 | 60 | 200 | 598 | 1 | 827 | 1422 | 245 | 111 | 147 | 238 |
| cluster3009 | 612 | 1233 | 2889 | 81 | 293 | 420 | 1479 | 1547 | 758 | 2045 |
| cluster3010 | 158 | 483 | 1284 | 9 | 163 | 196 | 750 | 162 | 759 | 1101 |
| cluster3011 | 226 | 671 | 421 | 1780 | 4034 | 1048 | 1730 | 567 | 1059 | 2164 |
| cluster3012 | 100 | 195 | 59 | 0 | 492 | 1407 | 260 | 135 | 103 | 387 |
| cluster3013 | 286 | 823 | 861 | 1245 | 565 | 2731 | 780 | 272 | 1747 | 2192 |
| cluster3014 | 347 | 845 | 468 | 0 | 249 | 411 | 241 | 102 | 303 | 1431 |
| cluster3015 | 62 | 186 | 1516 | 21 | 110 | 104 | 66 | 20 | 829 | 685 |
| cluster3016 | 105 | 239 | 351 | 0 | 62 | 26 | 224 | 49 | 317 | 349 |
| cluster3017 | 57 | 158 | 2980 | 12 | 101 | 227 | 415 | 103 | 361 | 377 |
| cluster3018 | 218 | 533 | 2067 | 0 | 751 | 1272 | 852 | 360 | 627 | 1254 |
| cluster3019 | 125 | 347 | 2912 | 1 | 917 | 1234 | 532 | 168 | 339 | 422 |
| cluster3020 | 1363 | 3514 | 16240 | 0 | 5149 | 5370 | 6000 | 2822 | 11023 | 15222 |
| cluster3021 | 191 | 388 | 437 | 583 | 463 | 820 | 598 | 132 | 481 | 1087 |
| cluster3022 | 90 | 206 | 339 | 0 | 173 | 243 | 389 | 51 | 444 | 855 |
| cluster3023 | 99 | 270 | 176 | 804 | 515 | 765 | 405 | 118 | 408 | 760 |
| cluster3024 | 66 | 289 | 654 | 170 | 93 | 80 | 596 | 103 | 1882 | 1449 |
| cluster3025 | 222 | 632 | 2354 | 224 | 620 | 1304 | 1419 | 300 | 867 | 1714 |
| cluster3026 | 40 | 75 | 159 | 0 | 67 | 68 | 335 | 17 | 806 | 513 |
| cluster3027 | 145 | 476 | 628 | 1195 | 787 | 1367 | 819 | 246 | 753 | 1698 |
| cluster3028 | 74 | 229 | 598 | 0 | 448 | 653 | 632 | 71 | 436 | 851 |
| cluster3029 | 126 | 263 | 1118 | 1 | 1025 | 1169 | 1103 | 265 | 1007 | 1647 |
| cluster3030 | 136 | 347 | 963 | 3072 | 395 | 846 | 327 | 120 | 2143 | 3014 |
| cluster3031 | 148 | 302 | 263 | 627 | 540 | 406 | 564 | 157 | 1240 | 1043 |
| cluster3032 | 91 | 202 | 467 | 682 | 530 | 622 | 409 | 109 | 900 | 1386 |
| cluster3033 | 1742 | 5259 | 8294 | 501 | 3122 | 7795 | 7951 | 2785 | 5222 | 6250 |
| cluster3034 | 57 | 212 | 389 | 1 | 392 | 253 | 731 | 93 | 253 | 325 |
| cluster3035 | 521 | 1310 | 2742 | 982 | 803 | 676 | 1369 | 384 | 2087 | 3286 |

| | | | | | | | | | | |
|---|---|---|---|---|---|---|---|---|---|---|
| cluster3036 | 75 | 173 | 950 | 0 | 543 | 475 | 932 | 158 | 841 | 1142 |
| cluster3037 | 591 | 1253 | 2810 | 33 | 1433 | 700 | 962 | 424 | 5221 | 2096 |
| cluster3038 | 58 | 168 | 696 | 0 | 467 | 1064 | 899 | 121 | 1075 | 908 |
| cluster3039 | 116 | 336 | 292 | 532 | 1051 | 1227 | 1019 | 194 | 687 | 1086 |
| cluster3040 | 65 | 155 | 136 | 280 | 99 | 131 | 307 | 169 | 444 | 657 |
| cluster3041 | 40 | 117 | 268 | 3 | 101 | 125 | 140 | 28 | 71 | 74 |
| cluster3042 | 49 | 142 | 228 | 0 | 324 | 491 | 578 | 49 | 103 | 323 |
| cluster3043 | 26 | 53 | 235 | 0 | 54 | 114 | 176 | 40 | 97 | 172 |
| cluster3044 | 58 | 139 | 887 | 0 | 78 | 678 | 402 | 80 | 430 | 722 |
| cluster3045 | 45 | 135 | 360 | 202 | 622 | 878 | 260 | 19 | 562 | 524 |
| cluster3046 | 63 | 179 | 1165 | 35 | 264 | 298 | 653 | 36 | 1186 | 981 |
| cluster3047 | 2872 | 7146 | 1376 | 6001 | 11208 | 724 | 6963 | 8367 | 14634 | 17328 |
| cluster3048 | 166 | 495 | 645 | 0 | 492 | 581 | 504 | 287 | 286 | 799 |
| cluster3049 | 216 | 501 | 644 | 1736 | 1080 | 1672 | 1083 | 578 | 1363 | 2837 |
| cluster3050 | 100 | 313 | 1036 | 27 | 112 | 402 | 738 | 97 | 590 | 1171 |
| cluster3051 | 47 | 169 | 210 | 0 | 443 | 465 | 835 | 156 | 161 | 1103 |
| cluster3052 | 72 | 174 | 2664 | 2 | 69 | 105 | 2608 | 123 | 1426 | 1272 |
| cluster3053 | 137 | 353 | 325 | 1215 | 114 | 842 | 219 | 69 | 344 | 744 |
| cluster3054 | 452 | 1469 | 4432 | 44 | 1593 | 2536 | 1778 | 1184 | 2392 | 4488 |
| cluster3055 | 141 | 355 | 515 | 18 | 586 | 613 | 585 | 90 | 751 | 948 |
| cluster3056 | 69 | 216 | 938 | 208 | 265 | 243 | 828 | 104 | 635 | 586 |
| cluster3057 | 150 | 368 | 68 | 288 | 16 | 112 | 379 | 178 | 520 | 522 |
| cluster3058 | 150 | 345 | 400 | 331 | 691 | 794 | 1084 | 149 | 530 | 1086 |
| cluster3059 | 63 | 132 | 1299 | 0 | 452 | 654 | 995 | 72 | 442 | 220 |
| cluster3060 | 264 | 696 | 145 | 0 | 603 | 884 | 326 | 150 | 130 | 546 |
| cluster3061 | 128 | 363 | 962 | 1531 | 921 | 1077 | 744 | 331 | 616 | 1148 |
| cluster3062 | 55 | 131 | 170 | 54 | 260 | 97 | 531 | 104 | 145 | 460 |
| cluster3063 | 40 | 100 | 413 | 521 | 818 | 1827 | 230 | 147 | 222 | 514 |
| cluster3064 | 55 | 113 | 264 | 31 | 82 | 310 | 82 | 3 | 395 | 405 |
| cluster3065 | 172 | 535 | 1322 | 17 | 147 | 311 | 199 | 123 | 1016 | 1635 |
| cluster3066 | 146 | 512 | 622 | 1479 | 1771 | 1813 | 1043 | 366 | 1258 | 2565 |

| | | | | | | | | | | |
|---|---|---|---|---|---|---|---|---|---|---|
| cluster3067 | 131 | 291 | 219 | 489 | 416 | 898 | 394 | 227 | 646 | 1518 |
| cluster3068 | 135 | 369 | 335 | 152 | 332 | 260 | 680 | 174 | 650 | 1400 |
| cluster3069 | 380 | 1244 | 3650 | 568 | 907 | 1462 | 1623 | 948 | 2647 | 4305 |
| cluster3070 | 192 | 491 | 547 | 657 | 692 | 1335 | 1165 | 41 | 1714 | 2449 |
| cluster3071 | 107 | 271 | 345 | 792 | 571 | 766 | 710 | 201 | 1208 | 1391 |
| cluster3072 | 790 | 1765 | 8851 | 0 | 1569 | 1791 | 2037 | 519 | 4262 | 4459 |
| cluster3073 | 451 | 1179 | 1299 | 30 | 693 | 1200 | 1062 | 856 | 1369 | 2944 |
| cluster3074 | 702 | 1531 | 4857 | 0 | 1077 | 1137 | 1349 | 731 | 1784 | 3293 |
| cluster3075 | 77 | 200 | 61 | 196 | 167 | 161 | 555 | 271 | 128 | 376 |
| cluster3076 | 22 | 35 | 124 | 0 | 147 | 329 | 57 | 31 | 64 | 151 |
| cluster3077 | 2999 | 6519 | 4618 | 9995 | 5494 | 6958 | 1746 | 2864 | 2502 | 6013 |
| cluster3078 | 122 | 270 | 644 | 79 | 569 | 533 | 293 | 137 | 178 | 527 |
| cluster3079 | 39 | 104 | 152 | 303 | 189 | 184 | 295 | 99 | 153 | 346 |
| cluster3080 | 40 | 132 | 793 | 10 | 134 | 182 | 682 | 74 | 335 | 617 |
| cluster3081 | 614 | 1587 | 497 | 1163 | 688 | 661 | 1491 | 1636 | 650 | 2894 |
| cluster3082 | 379 | 898 | 660 | 0 | 159 | 311 | 1759 | 82 | 280 | 1756 |
| cluster3083 | 84 | 187 | 198 | 296 | 765 | 705 | 391 | 100 | 299 | 549 |
| cluster3084 | 187 | 570 | 1609 | 546 | 394 | 628 | 1911 | 388 | 1371 | 2049 |
| cluster3085 | 52 | 116 | 367 | 0 | 92 | 151 | 546 | 98 | 295 | 655 |
| cluster3086 | 151 | 408 | 1591 | 19 | 226 | 257 | 668 | 130 | 510 | 1031 |
| cluster3087 | 68 | 151 | 446 | 3 | 147 | 636 | 243 | 65 | 204 | 362 |
| cluster3088 | 51 | 129 | 1540 | 33 | 268 | 342 | 371 | 29 | 201 | 312 |
| cluster3089 | 1758 | 4106 | 3510 | 9138 | 8980 | 6831 | 4140 | 1591 | 3740 | 7349 |
| cluster3090 | 130 | 328 | 3841 | 0 | 37 | 280 | 308 | 358 | 1630 | 3995 |
| cluster3091 | 453 | 1216 | 716 | 1210 | 1789 | 3014 | 2507 | 1211 | 2059 | 5166 |
| cluster3092 | 407 | 1076 | 1204 | 11 | 923 | 1419 | 875 | 746 | 922 | 2239 |
| cluster3093 | 1704 | 3672 | 16230 | 247 | 30136 | 6801 | 18338 | 10232 | 9864 | 31120 |
| cluster3094 | 23 | 91 | 529 | 0 | 495 | 563 | 237 | 93 | 130 | 347 |
| cluster3095 | 140 | 368 | 197 | 417 | 200 | 378 | 1074 | 647 | 363 | 826 |
| cluster3096 | 219 | 690 | 2301 | 19 | 815 | 1144 | 1084 | 358 | 549 | 1385 |
| cluster3097 | 46 | 101 | 404 | 290 | 1076 | 1166 | 574 | 62 | 757 | 843 |

| | | | | | | | | | | |
|---|---|---|---|---|---|---|---|---|---|---|
| cluster3098 | 151 | 296 | 864 | 0 | 317 | 477 | 467 | 144 | 290 | 437 |
| cluster3099 | 25 | 54 | 473 | 57 | 285 | 282 | 245 | 54 | 561 | 1987 |
| cluster3100 | 240 | 566 | 1728 | 2 | 271 | 6395 | 277 | 8 | 5671 | 2890 |
| cluster3101 | 133 | 403 | 713 | 1036 | 621 | 1276 | 477 | 147 | 325 | 518 |
| cluster3102 | 36 | 96 | 133 | 430 | 127 | 266 | 160 | 35 | 254 | 547 |
| cluster3103 | 67 | 181 | 753 | 0 | 175 | 8 | 541 | 46 | 28 | 85 |
| cluster3104 | 314 | 728 | 761 | 236 | 1125 | 851 | 1025 | 378 | 986 | 2298 |
| cluster3105 | 7035 | 13603 | 3734 | 0 | 2286 | 1101 | 4927 | 6385 | 5063 | 30370 |
| cluster3106 | 361 | 1031 | 4072 | 1697 | 3410 | 4790 | 2958 | 1528 | 7325 | 11377 |
| cluster3107 | 72 | 199 | 1224 | 47 | 193 | 374 | 246 | 145 | 252 | 337 |
| cluster3108 | 100 | 167 | 1373 | 0 | 541 | 1146 | 961 | 99 | 2190 | 2168 |
| cluster3109 | 618 | 1368 | 814 | 0 | 2382 | 3655 | 950 | 260 | 280 | 1641 |
| cluster3110 | 127 | 432 | 2688 | 38 | 1281 | 2348 | 928 | 211 | 1103 | 1738 |
| cluster3111 | 387 | 1235 | 1315 | 1 | 89 | 345 | 1504 | 169 | 5199 | 7354 |
| cluster3112 | 166 | 483 | 485 | 0 | 154 | 1353 | 285 | 318 | 1579 | 2067 |
| cluster3113 | 110 | 254 | 1273 | 9 | 393 | 579 | 209 | 146 | 698 | 1418 |
| cluster3114 | 182 | 409 | 931 | 16 | 238 | 268 | 452 | 140 | 786 | 1062 |
| cluster3115 | 106 | 314 | 2165 | 0 | 579 | 987 | 754 | 178 | 1119 | 1160 |
| cluster3116 | 97 | 246 | 939 | 3 | 605 | 361 | 377 | 201 | 382 | 786 |
| cluster3117 | 51 | 168 | 214 | 0 | 217 | 325 | 54 | 10 | 59 | 61 |
| cluster3118 | 35 | 106 | 894 | 91 | 406 | 567 | 271 | 37 | 1085 | 1194 |
| cluster3119 | 57 | 202 | 277 | 583 | 217 | 333 | 245 | 119 | 252 | 550 |
| cluster3120 | 222 | 587 | 1439 | 10 | 1035 | 1270 | 1145 | 252 | 895 | 2017 |
| cluster3121 | 1667 | 4418 | 1995 | 6830 | 3276 | 3227 | 3045 | 1901 | 10977 | 15959 |
| cluster3122 | 982 | 2463 | 688 | 1635 | 859 | 1470 | 2039 | 1096 | 1835 | 3575 |
| cluster3123 | 81 | 209 | 140 | 124 | 85 | 284 | 281 | 32 | 679 | 863 |
| cluster3124 | 486 | 1012 | 3093 | 50 | 1282 | 1784 | 2200 | 699 | 2192 | 4464 |
| cluster3125 | 515 | 1496 | 808 | 49 | 1757 | 2066 | 1277 | 884 | 924 | 2288 |
| cluster3126 | 220 | 524 | 1608 | 5083 | 1223 | 1153 | 662 | 529 | 2089 | 3583 |
| cluster3127 | 605 | 2144 | 1704 | 104 | 1074 | 1556 | 3273 | 815 | 1152 | 2635 |
| cluster3128 | 125 | 352 | 828 | 78 | 423 | 429 | 489 | 70 | 1897 | 1261 |

| | | | | | | | | | | |
|---|---|---|---|---|---|---|---|---|---|---|
| cluster3129 | 386 | 1033 | 348 | 0 | 183 | 20 | 553 | 439 | 444 | 3793 |
| cluster3130 | 136 | 350 | 328 | 0 | 166 | 253 | 531 | 371 | 348 | 1077 |
| cluster3131 | 972 | 1849 | 5053 | 0 | 1010 | 909 | 2459 | 1039 | 916 | 2505 |
| cluster3132 | 49 | 119 | 194 | 301 | 302 | 429 | 212 | 103 | 282 | 414 |
| cluster3133 | 39 | 139 | 453 | 0 | 246 | 313 | 443 | 122 | 241 | 702 |
| cluster3134 | 73 | 190 | 246 | 43 | 189 | 238 | 407 | 45 | 270 | 443 |
| cluster3135 | 4284 | 10952 | 23838 | 3 | 2444 | 6366 | 14036 | 8408 | 27279 | 28991 |
| cluster3136 | 25 | 105 | 633 | 3 | 63 | 87 | 236 | 29 | 98 | 133 |
| cluster3137 | 70 | 262 | 3580 | 0 | 621 | 896 | 82 | 39 | 893 | 1500 |
| cluster3138 | 61 | 153 | 271 | 551 | 321 | 406 | 140 | 48 | 286 | 523 |
| cluster3139 | 71 | 214 | 98 | 0 | 44 | 240 | 40 | 7 | 65 | 78 |
| cluster3140 | 98 | 154 | 633 | 54 | 402 | 621 | 314 | 69 | 452 | 369 |
| cluster3141 | 222 | 555 | 559 | 951 | 635 | 1041 | 380 | 226 | 786 | 2040 |
| cluster3142 | 62 | 160 | 1178 | 31 | 48 | 99 | 96 | 18 | 133 | 167 |
| cluster3143 | 49 | 170 | 131 | 223 | 427 | 384 | 385 | 28 | 309 | 451 |
| cluster3144 | 399 | 1052 | 535 | 0 | 404 | 688 | 243 | 424 | 217 | 699 |
| cluster3145 | 48 | 94 | 636 | 171 | 575 | 650 | 909 | 297 | 526 | 1018 |
| cluster3146 | 814 | 1999 | 1602 | 2545 | 655 | 2644 | 3327 | 1364 | 8547 | 14655 |
| cluster3147 | 179 | 504 | 2720 | 22 | 1052 | 1162 | 652 | 415 | 615 | 1708 |
| cluster3148 | 179 | 445 | 1818 | 54 | 834 | 598 | 704 | 211 | 817 | 1423 |
| cluster3149 | 160 | 434 | 403 | 820 | 1018 | 1237 | 807 | 331 | 665 | 1327 |
| cluster3150 | 48 | 104 | 717 | 6 | 532 | 110 | 1728 | 57 | 886 | 614 |
| cluster3151 | 375 | 733 | 2604 | 75 | 565 | 598 | 1272 | 341 | 2940 | 3271 |
| cluster3152 | 41 | 103 | 177 | 0 | 121 | 176 | 134 | 18 | 85 | 163 |
| cluster3153 | 621 | 1608 | 3722 | 4 | 2817 | 6759 | 2324 | 642 | 4493 | 5077 |
| cluster3154 | 57 | 166 | 218 | 173 | 451 | 496 | 649 | 84 | 347 | 520 |
| cluster3155 | 45 | 106 | 515 | 0 | 200 | 352 | 113 | 76 | 280 | 493 |
| cluster3156 | 234 | 533 | 574 | 0 | 591 | 664 | 319 | 534 | 195 | 640 |
| cluster3157 | 138 | 381 | 2744 | 617 | 475 | 1250 | 1249 | 153 | 1539 | 2641 |
| cluster3158 | 180 | 457 | 27 | 57 | 94 | 1519 | 283 | 138 | 252 | 1600 |
| cluster3159 | 92 | 243 | 603 | 0 | 277 | 454 | 233 | 201 | 400 | 878 |

| | | | | | | | | | | |
|---|---|---|---|---|---|---|---|---|---|---|
| cluster3160 | 968 | 3258 | 5374 | 563 | 667 | 1365 | 2852 | 1759 | 766 | 2379 |
| cluster3161 | 98 | 241 | 1084 | 0 | 105 | 190 | 799 | 346 | 463 | 1038 |
| cluster3162 | 23 | 67 | 184 | 1 | 254 | 559 | 185 | 17 | 170 | 105 |
| cluster3163 | 627 | 1356 | 3270 | 0 | 938 | 861 | 476 | 590 | 375 | 1569 |
| cluster3164 | 184 | 381 | 1095 | 0 | 381 | 519 | 982 | 580 | 1794 | 2537 |
| cluster3165 | 29 | 121 | 1201 | 0 | 169 | 187 | 123 | 32 | 149 | 292 |
| cluster3166 | 259 | 594 | 846 | 1088 | 685 | 1174 | 595 | 380 | 447 | 901 |
| cluster3167 | 50 | 153 | 608 | 0 | 336 | 279 | 637 | 43 | 1005 | 994 |
| cluster3168 | 77 | 195 | 137 | 293 | 716 | 886 | 203 | 73 | 255 | 733 |
| cluster3169 | 147 | 394 | 1675 | 0 | 594 | 1736 | 425 | 322 | 1136 | 978 |
| cluster3170 | 37 | 103 | 462 | 154 | 233 | 298 | 570 | 34 | 379 | 655 |
| cluster3171 | 69 | 164 | 240 | 616 | 105 | 141 | 599 | 121 | 935 | 926 |
| cluster3172 | 178 | 506 | 744 | 1212 | 403 | 239 | 2034 | 329 | 973 | 1664 |
| cluster3173 | 59 | 139 | 522 | 0 | 195 | 207 | 187 | 32 | 73 | 220 |
| cluster3174 | 103 | 230 | 1558 | 0 | 150 | 772 | 304 | 39 | 480 | 817 |
| cluster3175 | 357 | 906 | 286 | 0 | 311 | 150 | 683 | 306 | 84 | 333 |
| cluster3176 | 152 | 380 | 492 | 93 | 1246 | 949 | 5643 | 108 | 2611 | 1779 |
| cluster3177 | 18943 | 46439 | 13506 | 4334 | 12459 | 15482 | 6860 | 63025 | 9699 | 23078 |
| cluster3178 | 38 | 101 | 513 | 1509 | 1421 | 1562 | 275 | 89 | 322 | 655 |
| cluster3179 | 30 | 95 | 437 | 0 | 468 | 977 | 165 | 52 | 57 | 145 |
| cluster3180 | 88 | 237 | 677 | 0 | 189 | 175 | 405 | 308 | 350 | 605 |
| cluster3181 | 58 | 189 | 422 | 662 | 389 | 566 | 588 | 130 | 472 | 786 |
| cluster3182 | 15948 | 37441 | 2272 | 4193 | 19167 | 8406 | 14443 | 21301 | 4360 | 22721 |
| cluster3183 | 47 | 185 | 613 | 4 | 79 | 645 | 247 | 255 | 252 | 621 |
| cluster3184 | 71 | 193 | 609 | 5 | 273 | 340 | 351 | 89 | 339 | 457 |
| cluster3185 | 1493 | 3137 | 1090 | 0 | 1190 | 737 | 730 | 1431 | 288 | 2737 |
| cluster3186 | 910 | 2879 | 921 | 0 | 1638 | 2601 | 575 | 1082 | 224 | 2091 |
| cluster3187 | 263 | 623 | 514 | 0 | 44 | 177 | 353 | 100 | 722 | 2550 |
| cluster3188 | 76 | 215 | 204 | 801 | 394 | 296 | 486 | 60 | 506 | 579 |
| cluster3189 | 31 | 125 | 543 | 0 | 76 | 121 | 167 | 53 | 258 | 517 |
| cluster3190 | 42 | 101 | 123 | 0 | 321 | 157 | 125 | 4 | 59 | 256 |

| | | | | | | | | | | |
|---|---|---|---|---|---|---|---|---|---|---|
| cluster3191 | 279 | 919 | 42 | 47 | 489 | 4 | 1058 | 106 | 29 | 354 |
| cluster3192 | 165 | 450 | 1943 | 573 | 265 | 416 | 487 | 459 | 163 | 1029 |
| cluster3193 | 372 | 933 | 3672 | 14 | 395 | 733 | 891 | 396 | 1608 | 2292 |
| cluster3194 | 119 | 231 | 817 | 120 | 163 | 230 | 1279 | 44 | 2833 | 1869 |
| cluster3195 | 189 | 584 | 258 | 612 | 622 | 869 | 2320 | 524 | 2454 | 5703 |
| cluster3196 | 181 | 591 | 441 | 684 | 464 | 558 | 1225 | 369 | 1552 | 1839 |
| cluster3197 | 152 | 299 | 729 | 1135 | 641 | 395 | 452 | 90 | 1506 | 1184 |
| cluster3198 | 207 | 525 | 1166 | 0 | 1314 | 1488 | 1055 | 431 | 346 | 999 |
| cluster3199 | 320 | 885 | 1696 | 11 | 590 | 879 | 673 | 401 | 1731 | 2541 |
| cluster3200 | 83 | 265 | 485 | 933 | 1159 | 1065 | 541 | 137 | 407 | 907 |
| cluster3201 | 256 | 491 | 3590 | 324 | 2479 | 2336 | 3281 | 475 | 2802 | 3884 |
| cluster3202 | 48 | 95 | 916 | 23 | 616 | 634 | 722 | 103 | 530 | 600 |
| cluster3203 | 23 | 71 | 166 | 0 | 41 | 69 | 116 | 85 | 70 | 119 |
| cluster3204 | 89 | 197 | 222 | 127 | 143 | 135 | 47 | 28 | 286 | 271 |
| cluster3205 | 46 | 105 | 50 | 94 | 66 | 94 | 180 | 73 | 190 | 356 |
| cluster3206 | 45 | 98 | 144 | 0 | 104 | 387 | 108 | 87 | 115 | 407 |
| cluster3207 | 79 | 207 | 155 | 431 | 1466 | 2005 | 1205 | 175 | 269 | 708 |
| cluster3208 | 802 | 2247 | 1727 | 3423 | 2944 | 3271 | 3612 | 1344 | 4371 | 8951 |
| cluster3209 | 337 | 803 | 1064 | 0 | 238 | 587 | 642 | 162 | 515 | 1206 |
| cluster3210 | 66 | 151 | 295 | 1 | 292 | 458 | 162 | 64 | 236 | 459 |
| cluster3211 | 133 | 360 | 504 | 0 | 494 | 836 | 347 | 110 | 172 | 266 |
| cluster3212 | 2405 | 6899 | 943 | 3482 | 3508 | 3612 | 3903 | 11313 | 2072 | 4679 |
| cluster3213 | 82 | 211 | 1176 | 867 | 623 | 254 | 2007 | 192 | 352 | 1404 |
| cluster3214 | 40 | 96 | 24 | 80 | 65 | 135 | 72 | 43 | 71 | 177 |
| cluster3215 | 109 | 301 | 939 | 206 | 464 | 470 | 409 | 167 | 893 | 928 |
| cluster3216 | 38 | 140 | 386 | 0 | 122 | 138 | 111 | 57 | 109 | 273 |
| cluster3217 | 87 | 240 | 198 | 471 | 124 | 259 | 815 | 108 | 617 | 744 |
| cluster3218 | 58 | 243 | 1466 | 0 | 941 | 491 | 505 | 96 | 344 | 484 |
| cluster3219 | 132 | 282 | 369 | 4 | 497 | 700 | 481 | 205 | 210 | 510 |
| cluster3220 | 964 | 2861 | 235 | 950 | 322 | 1715 | 2473 | 1951 | 1506 | 5777 |
| cluster3221 | 100 | 372 | 1006 | 0 | 441 | 520 | 225 | 164 | 469 | 1232 |

| | | | | | | | | | | |
|---|---|---|---|---|---|---|---|---|---|---|
| cluster3222 | 262 | 651 | 831 | 420 | 888 | 982 | 1302 | 338 | 395 | 1135 |
| cluster3223 | 35 | 91 | 982 | 34 | 53 | 72 | 573 | 43 | 240 | 277 |
| cluster3224 | 86 | 228 | 890 | 36 | 597 | 656 | 1584 | 153 | 1357 | 1426 |
| cluster3225 | 62 | 169 | 142 | 0 | 226 | 247 | 172 | 169 | 197 | 498 |
| cluster3226 | 227 | 548 | 446 | 0 | 265 | 259 | 483 | 260 | 266 | 667 |
| cluster3227 | 56 | 164 | 650 | 1 | 95 | 165 | 119 | 52 | 327 | 318 |
| cluster3228 | 129 | 314 | 942 | 0 | 333 | 735 | 222 | 95 | 373 | 742 |
| cluster3229 | 78 | 222 | 384 | 511 | 66 | 104 | 310 | 49 | 453 | 1258 |
| cluster3230 | 38 | 77 | 129 | 271 | 100 | 112 | 416 | 25 | 184 | 587 |
| cluster3231 | 72 | 224 | 6518 | 0 | 511 | 1038 | 2413 | 1050 | 5303 | 20611 |
| cluster3232 | 101 | 227 | 1184 | 0 | 462 | 597 | 273 | 62 | 285 | 754 |
| cluster3233 | 80 | 221 | 2262 | 4113 | 117 | 672 | 102 | 122 | 460 | 592 |
| cluster3234 | 717 | 1996 | 202 | 521 | 932 | 1229 | 947 | 853 | 1307 | 1929 |
| cluster3235 | 184 | 452 | 1339 | 0 | 420 | 870 | 201 | 38 | 33 | 160 |
| cluster3236 | 71 | 203 | 443 | 0 | 402 | 999 | 148 | 202 | 184 | 266 |
| cluster3237 | 43 | 145 | 2608 | 0 | 170 | 222 | 229 | 14 | 199 | 322 |
| cluster3238 | 104 | 271 | 907 | 67 | 467 | 496 | 266 | 129 | 447 | 890 |
| cluster3239 | 139 | 430 | 1997 | 333 | 596 | 924 | 940 | 214 | 1281 | 2671 |
| cluster3240 | 36 | 123 | 184 | 445 | 321 | 536 | 261 | 51 | 63 | 214 |
| cluster3241 | 43 | 116 | 417 | 2 | 86 | 243 | 173 | 37 | 264 | 418 |
| cluster3242 | 169 | 489 | 2641 | 26 | 1664 | 2879 | 1073 | 471 | 1152 | 1749 |
| cluster3243 | 455 | 1144 | 1642 | 3158 | 1952 | 1848 | 3563 | 467 | 4269 | 7499 |
| cluster3244 | 49 | 103 | 184 | 233 | 287 | 415 | 173 | 49 | 320 | 683 |
| cluster3245 | 14 | 26 | 332 | 0 | 136 | 105 | 43 | 2 | 190 | 188 |
| cluster3246 | 191 | 451 | 567 | 1126 | 1230 | 1227 | 571 | 157 | 531 | 863 |
| cluster3247 | 79 | 207 | 427 | 17 | 380 | 487 | 104 | 6 | 293 | 430 |
| cluster3248 | 9 | 37 | 410 | 0 | 110 | 95 | 109 | 82 | 71 | 245 |
| cluster3249 | 204 | 551 | 533 | 13 | 349 | 182 | 552 | 143 | 555 | 1598 |
| cluster3250 | 77 | 157 | 640 | 34 | 215 | 4457 | 95 | 51 | 2003 | 2974 |
| cluster3251 | 171 | 453 | 851 | 0 | 434 | 760 | 468 | 204 | 711 | 1354 |
| cluster3252 | 603 | 1489 | 3574 | 0 | 197 | 10060 | 340 | 1230 | 854 | 1080 |

| | | | | | | | | | | |
|---|---|---|---|---|---|---|---|---|---|---|
| cluster3253 | 173 | 436 | 621 | 0 | 665 | 835 | 742 | 216 | 616 | 841 |
| cluster3254 | 345 | 978 | 297 | 0 | 128 | 118 | 150 | 52 | 253 | 600 |
| cluster3255 | 477 | 1297 | 1873 | 0 | 1623 | 316 | 1004 | 1069 | 2038 | 2829 |
| cluster3256 | 31 | 93 | 217 | 62 | 163 | 171 | 169 | 66 | 68 | 112 |
| cluster3257 | 72 | 206 | 153 | 294 | 511 | 632 | 374 | 136 | 231 | 443 |
| cluster3258 | 435 | 1475 | 7721 | 165 | 2955 | 3489 | 2658 | 898 | 4132 | 6202 |
| cluster3259 | 288 | 810 | 503 | 430 | 148 | 874 | 704 | 447 | 1364 | 3552 |
| cluster3260 | 222 | 588 | 169 | 651 | 736 | 695 | 2372 | 550 | 366 | 1094 |
| cluster3261 | 83 | 200 | 560 | 0 | 130 | 167 | 242 | 196 | 277 | 635 |
| cluster3262 | 34 | 72 | 361 | 1 | 94 | 199 | 154 | 68 | 197 | 380 |
| cluster3263 | 37 | 130 | 525 | 9 | 307 | 363 | 187 | 120 | 411 | 881 |
| cluster3264 | 1450 | 4031 | 5410 | 334 | 2686 | 3883 | 1977 | 2431 | 3108 | 6510 |
| cluster3265 | 175 | 388 | 2326 | 0 | 208 | 380 | 377 | 161 | 333 | 756 |
| cluster3266 | 26 | 89 | 539 | 60 | 243 | 179 | 401 | 49 | 384 | 549 |
| cluster3267 | 43 | 134 | 171 | 226 | 435 | 491 | 285 | 119 | 211 | 751 |
| cluster3268 | 286 | 719 | 694 | 0 | 303 | 981 | 441 | 315 | 386 | 934 |
| cluster3269 | 445 | 980 | 1100 | 0 | 670 | 769 | 866 | 398 | 1592 | 2705 |
| cluster3270 | 134 | 401 | 575 | 0 | 2083 | 3121 | 1297 | 288 | 596 | 1675 |
| cluster3271 | 264 | 647 | 328 | 393 | 527 | 515 | 526 | 330 | 347 | 1073 |
| cluster3272 | 1703 | 2607 | 0 | 4 | 4110 | 27618 | 8562 | 3482 | 29651 | 15477 |
| cluster3273 | 43 | 79 | 272 | 161 | 343 | 283 | 267 | 63 | 366 | 572 |
| cluster3274 | 20 | 68 | 549 | 0 | 339 | 597 | 455 | 73 | 93 | 214 |
| cluster3275 | 65 | 202 | 324 | 0 | 68 | 43 | 172 | 28 | 57 | 233 |
| cluster3276 | 60 | 202 | 107 | 231 | 332 | 333 | 343 | 61 | 698 | 732 |
| cluster3277 | 19 | 55 | 90 | 1 | 95 | 139 | 58 | 90 | 171 | 158 |
| cluster3278 | 144 | 349 | 2057 | 58 | 1063 | 1123 | 1273 | 494 | 1298 | 1537 |
| cluster3279 | 74 | 172 | 134 | 2 | 562 | 245 | 102 | 80 | 90 | 291 |
| cluster3280 | 104 | 261 | 686 | 8 | 275 | 519 | 129 | 68 | 640 | 912 |
| cluster3281 | 393 | 861 | 731 | 0 | 828 | 1237 | 440 | 113 | 820 | 1528 |
| cluster3282 | 142 | 376 | 594 | 1 | 174 | 234 | 296 | 134 | 534 | 821 |
| cluster3283 | 76 | 189 | 131 | 151 | 279 | 336 | 187 | 79 | 535 | 1038 |

| | | | | | | | | | | |
|---|---|---|---|---|---|---|---|---|---|---|
| cluster3284 | 53 | 96 | 417 | 0 | 147 | 183 | 155 | 34 | 154 | 222 |
| cluster3285 | 52 | 88 | 587 | 30 | 156 | 124 | 432 | 32 | 272 | 323 |
| cluster3286 | 181 | 554 | 2382 | 296 | 818 | 970 | 1459 | 353 | 2416 | 3490 |
| cluster3287 | 17 | 66 | 96 | 6 | 17 | 22 | 34 | 7 | 92 | 148 |
| cluster3288 | 13 | 39 | 141 | 0 | 240 | 359 | 61 | 43 | 136 | 97 |
| cluster3289 | 44 | 124 | 211 | 392 | 313 | 388 | 459 | 133 | 370 | 642 |
| cluster3290 | 102 | 243 | 62 | 112 | 87 | 35 | 900 | 65 | 289 | 731 |
| cluster3291 | 78 | 122 | 102 | 292 | 623 | 278 | 392 | 59 | 177 | 372 |
| cluster3292 | 2273 | 5380 | 16879 | 0 | 1233 | 1992 | 4935 | 6321 | 7268 | 14492 |
| cluster3293 | 55 | 184 | 101 | 141 | 322 | 471 | 266 | 88 | 120 | 380 |
| cluster3294 | 1127 | 2280 | 814 | 2444 | 2405 | 3917 | 2318 | 1109 | 1196 | 5852 |
| cluster3295 | 141 | 301 | 931 | 91 | 617 | 855 | 466 | 154 | 407 | 1542 |
| cluster3296 | 2766 | 7699 | 4714 | 0 | 8662 | 6855 | 4717 | 8123 | 1606 | 10283 |
| cluster3297 | 138 | 414 | 554 | 849 | 452 | 794 | 82 | 15 | 163 | 238 |
| cluster3298 | 81 | 191 | 109 | 297 | 330 | 253 | 1016 | 167 | 332 | 1115 |
| cluster3299 | 86 | 214 | 199 | 0 | 462 | 512 | 149 | 294 | 136 | 195 |
| cluster3300 | 29 | 51 | 256 | 18 | 59 | 272 | 118 | 10 | 192 | 385 |
| cluster3301 | 57 | 188 | 279 | 811 | 312 | 528 | 1133 | 400 | 112 | 269 |
| cluster3302 | 122 | 264 | 490 | 1 | 361 | 385 | 243 | 140 | 745 | 872 |
| cluster3303 | 13 | 38 | 391 | 945 | 280 | 349 | 202 | 38 | 107 | 352 |
| cluster3304 | 371 | 1031 | 1653 | 2756 | 1752 | 3268 | 2101 | 430 | 3096 | 3098 |
| cluster3305 | 77 | 246 | 858 | 0 | 149 | 1671 | 1339 | 126 | 479 | 2621 |
| cluster3306 | 69 | 211 | 140 | 0 | 1436 | 3097 | 70 | 174 | 142 | 485 |
| cluster3307 | 347 | 852 | 1976 | 0 | 663 | 1814 | 531 | 780 | 1316 | 2703 |
| cluster3308 | 112 | 250 | 169 | 426 | 631 | 702 | 891 | 335 | 521 | 1373 |
| cluster3309 | 51 | 126 | 341 | 356 | 302 | 324 | 53 | 19 | 1429 | 796 |
| cluster3310 | 987 | 1673 | 2135 | 1392 | 2264 | 3150 | 3956 | 1183 | 4797 | 7922 |
| cluster3311 | 27 | 76 | 126 | 261 | 387 | 410 | 255 | 52 | 181 | 318 |
| cluster3312 | 70 | 140 | 35 | 0 | 359 | 66 | 340 | 262 | 118 | 471 |
| cluster3313 | 4584 | 12446 | 386 | 1078 | 7114 | 14683 | 12370 | 8949 | 4071 | 15447 |
| cluster3314 | 26 | 85 | 1791 | 0 | 410 | 484 | 1028 | 99 | 1440 | 1380 |

| | | | | | | | | | | |
|---|---|---|---|---|---|---|---|---|---|---|
| cluster3315 | 216 | 617 | 424 | 958 | 367 | 752 | 419 | 374 | 216 | 699 |
| cluster3316 | 32 | 94 | 281 | 1 | 418 | 69 | 533 | 33 | 502 | 517 |
| cluster3317 | 41 | 111 | 1078 | 26 | 123 | 136 | 45 | 15 | 259 | 199 |
| cluster3318 | 34 | 88 | 226 | 500 | 106 | 206 | 292 | 132 | 326 | 392 |
| cluster3319 | 344 | 941 | 819 | 0 | 186 | 410 | 654 | 291 | 2321 | 3235 |
| cluster3320 | 40 | 100 | 223 | 0 | 126 | 216 | 517 | 27 | 238 | 168 |
| cluster3321 | 48 | 288 | 1105 | 2 | 271 | 369 | 475 | 75 | 554 | 978 |
| cluster3322 | 89 | 337 | 333 | 0 | 284 | 885 | 591 | 83 | 306 | 728 |
| cluster3323 | 14 | 53 | 364 | 330 | 175 | 229 | 153 | 9 | 101 | 162 |
| cluster3324 | 125 | 323 | 74 | 138 | 280 | 280 | 196 | 68 | 388 | 536 |
| cluster3325 | 66 | 85 | 640 | 0 | 16 | 9 | 481 | 24 | 75 | 344 |
| cluster3326 | 241 | 771 | 1411 | 387 | 135 | 84 | 503 | 234 | 1251 | 1924 |
| cluster3327 | 43 | 109 | 137 | 232 | 47 | 213 | 180 | 29 | 113 | 351 |
| cluster3328 | 172 | 387 | 457 | 71 | 426 | 463 | 318 | 149 | 845 | 1288 |
| cluster3329 | 24 | 78 | 54 | 0 | 208 | 325 | 462 | 169 | 27 | 93 |
| cluster3330 | 64 | 149 | 953 | 21 | 803 | 747 | 728 | 77 | 392 | 821 |
| cluster3331 | 101 | 242 | 486 | 0 | 226 | 375 | 628 | 98 | 317 | 900 |
| cluster3332 | 34 | 95 | 516 | 0 | 395 | 503 | 130 | 31 | 199 | 232 |
| cluster3333 | 75 | 193 | 557 | 0 | 219 | 478 | 154 | 129 | 131 | 152 |
| cluster3334 | 15 | 71 | 99 | 7 | 101 | 238 | 100 | 27 | 91 | 160 |
| cluster3335 | 32 | 123 | 80 | 203 | 99 | 342 | 94 | 18 | 153 | 273 |
| cluster3336 | 12 | 87 | 92 | 203 | 122 | 161 | 595 | 51 | 410 | 482 |
| cluster3337 | 35 | 176 | 250 | 488 | 111 | 143 | 209 | 72 | 236 | 233 |
| cluster3338 | 28 | 78 | 153 | 0 | 178 | 273 | 452 | 80 | 114 | 234 |
| cluster3339 | 84 | 206 | 104 | 296 | 175 | 283 | 255 | 114 | 363 | 597 |
| cluster3340 | 209 | 476 | 1193 | 414 | 230 | 662 | 273 | 89 | 543 | 1099 |
| cluster3341 | 93 | 249 | 318 | 0 | 539 | 548 | 1507 | 178 | 556 | 933 |
| cluster3342 | 99 | 341 | 239 | 0 | 822 | 975 | 393 | 120 | 103 | 376 |
| cluster3343 | 447 | 1204 | 4908 | 0 | 1351 | 3802 | 847 | 1418 | 4088 | 7246 |
| cluster3344 | 17 | 51 | 52 | 123 | 69 | 121 | 372 | 18 | 175 | 204 |
| cluster3345 | 60 | 110 | 204 | 0 | 259 | 373 | 248 | 150 | 140 | 271 |

| | | | | | | | | | | |
|---|---|---|---|---|---|---|---|---|---|---|
| cluster3346 | 259 | 628 | 1138 | 2977 | 1008 | 1395 | 762 | 873 | 837 | 1816 |
| cluster3347 | 68 | 212 | 97 | 207 | 189 | 354 | 195 | 127 | 208 | 424 |
| cluster3348 | 19 | 62 | 401 | 50 | 112 | 170 | 275 | 10 | 367 | 438 |
| cluster3349 | 679 | 1812 | 1190 | 5029 | 14913 | 12301 | 7285 | 2286 | 1672 | 8519 |
| cluster3350 | 177 | 494 | 532 | 0 | 204 | 332 | 272 | 55 | 400 | 1000 |
| cluster3351 | 274 | 861 | 1968 | 8 | 1136 | 1039 | 1027 | 529 | 778 | 1297 |
| cluster3352 | 71 | 236 | 489 | 0 | 147 | 176 | 119 | 29 | 322 | 181 |
| cluster3353 | 54 | 198 | 663 | 8 | 242 | 508 | 288 | 99 | 383 | 598 |
| cluster3354 | 55 | 159 | 1225 | 19 | 493 | 674 | 585 | 204 | 617 | 635 |
| cluster3355 | 46 | 139 | 151 | 0 | 67 | 33 | 115 | 6 | 160 | 222 |
| cluster3356 | 51 | 105 | 55 | 201 | 369 | 459 | 223 | 47 | 54 | 182 |
| cluster3357 | 61 | 231 | 571 | 46 | 625 | 478 | 102 | 50 | 610 | 754 |
| cluster3358 | 24 | 85 | 199 | 372 | 163 | 300 | 459 | 67 | 235 | 451 |
| cluster3359 | 225 | 493 | 736 | 0 | 428 | 463 | 458 | 122 | 126 | 338 |
| cluster3360 | 746 | 2383 | 92 | 446 | 735 | 639 | 655 | 774 | 480 | 1960 |
| cluster3361 | 400 | 856 | 1539 | 0 | 1051 | 1488 | 741 | 1027 | 1038 | 1800 |
| cluster3362 | 857 | 1816 | 1033 | 3884 | 2110 | 3059 | 2323 | 1370 | 1651 | 3290 |
| cluster3363 | 56 | 162 | 342 | 0 | 395 | 546 | 98 | 36 | 122 | 236 |
| cluster3364 | 2356 | 5786 | 15083 | 7 | 2336 | 9646 | 2350 | 4545 | 2826 | 5684 |
| cluster3365 | 72 | 165 | 562 | 0 | 299 | 1650 | 238 | 56 | 227 | 272 |
| cluster3366 | 37 | 67 | 373 | 0 | 276 | 432 | 87 | 49 | 109 | 139 |
| cluster3367 | 11 | 43 | 54 | 319 | 51 | 101 | 57 | 19 | 59 | 90 |
| cluster3368 | 168 | 549 | 318 | 227 | 379 | 195 | 377 | 224 | 739 | 2016 |
| cluster3369 | 61 | 149 | 1136 | 0 | 465 | 517 | 134 | 104 | 421 | 708 |
| cluster3370 | 43 | 103 | 546 | 1 | 435 | 429 | 520 | 46 | 193 | 388 |
| cluster3371 | 17 | 58 | 3955 | 0 | 3 | 203 | 101 | 39 | 464 | 437 |
| cluster3372 | 452 | 806 | 467 | 1632 | 1948 | 2377 | 1467 | 740 | 1273 | 2836 |
| cluster3373 | 28 | 67 | 363 | 0 | 192 | 235 | 155 | 27 | 160 | 210 |
| cluster3374 | 48 | 117 | 156 | 0 | 783 | 766 | 306 | 96 | 17 | 35 |
| cluster3375 | 14234 | 47264 | 10679 | 11408 | 30524 | 32596 | 42906 | 34671 | 24481 | 58424 |
| cluster3376 | 195 | 474 | 261 | 430 | 470 | 672 | 572 | 293 | 460 | 894 |

| | | | | | | | | | | |
|---|---|---|---|---|---|---|---|---|---|---|
| cluster3377 | 88 | 197 | 112 | 0 | 69 | 122 | 98 | 76 | 234 | 614 |
| cluster3378 | 34 | 89 | 14 | 112 | 129 | 217 | 163 | 59 | 55 | 132 |
| cluster3379 | 556 | 1354 | 1646 | 18 | 271 | 172 | 713 | 398 | 119 | 344 |
| cluster3380 | 47 | 122 | 489 | 9 | 176 | 345 | 1009 | 130 | 478 | 913 |
| cluster3381 | 741 | 2018 | 1418 | 2358 | 624 | 951 | 938 | 754 | 819 | 4677 |
| cluster3382 | 5258 | 9328 | 22679 | 2781 | 5129 | 7940 | 5617 | 6015 | 8701 | 13444 |
| cluster3383 | 234 | 754 | 845 | 0 | 55 | 74 | 315 | 128 | 270 | 1356 |
| cluster3384 | 517 | 1181 | 182 | 937 | 1808 | 170 | 543 | 400 | 1189 | 2007 |
| cluster3385 | 403 | 990 | 796 | 0 | 3070 | 596 | 780 | 1190 | 521 | 893 |
| cluster3386 | 621 | 1344 | 609 | 0 | 1516 | 1502 | 650 | 622 | 249 | 865 |
| cluster3387 | 219 | 615 | 812 | 0 | 2179 | 2717 | 1717 | 1013 | 394 | 1698 |
| cluster3388 | 67 | 209 | 502 | 3 | 218 | 524 | 2313 | 126 | 322 | 719 |
| cluster3389 | 219 | 606 | 224 | 0 | 927 | 939 | 284 | 322 | 297 | 1027 |
| cluster3390 | 33 | 55 | 309 | 145 | 318 | 174 | 549 | 64 | 561 | 376 |
| cluster3391 | 86 | 289 | 354 | 80 | 256 | 290 | 354 | 125 | 267 | 486 |
| cluster3392 | 109 | 297 | 59 | 0 | 319 | 220 | 229 | 164 | 19 | 352 |
| cluster3393 | 233 | 669 | 182 | 0 | 86 | 70 | 1506 | 436 | 42 | 876 |
| cluster3394 | 317 | 656 | 247 | 425 | 647 | 1458 | 346 | 589 | 1265 | 2367 |
| cluster3395 | 165 | 458 | 111 | 191 | 104 | 264 | 336 | 276 | 249 | 554 |
| cluster3396 | 27 | 51 | 119 | 216 | 200 | 174 | 515 | 60 | 239 | 336 |
| cluster3397 | 350 | 819 | 965 | 0 | 451 | 599 | 1603 | 1487 | 673 | 5913 |
| cluster3398 | 1411 | 3097 | 5929 | 43 | 1320 | 2157 | 4146 | 2543 | 3055 | 4880 |
| cluster3399 | 11 | 66 | 404 | 232 | 371 | 147 | 403 | 14 | 952 | 597 |
| cluster3400 | 140 | 347 | 1534 | 677 | 456 | 419 | 270 | 152 | 733 | 2063 |
| cluster3401 | 387 | 1116 | 5005 | 12 | 2809 | 4051 | 1442 | 493 | 2360 | 4589 |
| cluster3402 | 13 | 59 | 341 | 218 | 277 | 366 | 310 | 60 | 724 | 907 |
| cluster3403 | 979 | 2487 | 2672 | 0 | 1712 | 2489 | 1649 | 959 | 2214 | 5861 |
| cluster3404 | 436 | 1264 | 3227 | 2404 | 1678 | 2205 | 3404 | 1066 | 2118 | 1986 |
| cluster3405 | 81 | 211 | 1253 | 19 | 152 | 211 | 222 | 85 | 721 | 774 |
| cluster3406 | 171 | 485 | 235 | 540 | 794 | 922 | 509 | 630 | 211 | 804 |
| cluster3407 | 190 | 522 | 613 | 1374 | 1021 | 1326 | 1665 | 424 | 644 | 3250 |

| | | | | | | | | | | |
|---|---|---|---|---|---|---|---|---|---|---|
| cluster3408 | 26 | 79 | 386 | 74 | 40 | 30 | 97 | 19 | 335 | 503 |
| cluster3409 | 108 | 283 | 894 | 0 | 515 | 431 | 244 | 231 | 203 | 415 |
| cluster3410 | 52 | 128 | 713 | 1186 | 225 | 333 | 613 | 216 | 252 | 506 |
| cluster3411 | 166 | 434 | 738 | 0 | 1034 | 1156 | 1244 | 402 | 382 | 973 |
| cluster3412 | 40 | 94 | 376 | 0 | 136 | 169 | 262 | 18 | 138 | 125 |
| cluster3413 | 35 | 80 | 697 | 0 | 119 | 61 | 341 | 238 | 56 | 155 |
| cluster3414 | 47 | 212 | 280 | 440 | 173 | 314 | 449 | 64 | 397 | 660 |
| cluster3415 | 64 | 123 | 668 | 12 | 60 | 135 | 183 | 92 | 347 | 333 |
| cluster3416 | 80 | 196 | 34 | 0 | 397 | 536 | 96 | 97 | 108 | 239 |
| cluster3417 | 12 | 29 | 69 | 149 | 71 | 84 | 98 | 31 | 28 | 75 |
| cluster3418 | 41 | 107 | 167 | 0 | 177 | 420 | 91 | 66 | 85 | 273 |
| cluster3419 | 116 | 245 | 1116 | 83 | 141 | 301 | 563 | 121 | 1931 | 1962 |
| cluster3420 | 58 | 210 | 648 | 0 | 174 | 576 | 509 | 42 | 1083 | 814 |
| cluster3421 | 14 | 41 | 1127 | 2 | 381 | 750 | 583 | 38 | 336 | 369 |
| cluster3422 | 16 | 73 | 125 | 260 | 11 | 70 | 428 | 3 | 464 | 299 |
| cluster3423 | 36 | 126 | 180 | 0 | 27 | 188 | 85 | 8 | 63 | 164 |
| cluster3424 | 99 | 271 | 4 | 144 | 44 | 1 | 214 | 47 | 20 | 480 |
| cluster3425 | 90 | 325 | 293 | 15 | 66 | 207 | 700 | 51 | 400 | 665 |
| cluster3426 | 68 | 146 | 231 | 510 | 262 | 256 | 153 | 48 | 371 | 521 |
| cluster3427 | 37 | 44 | 663 | 9 | 79 | 78 | 532 | 17 | 998 | 523 |
| cluster3428 | 538 | 1272 | 273 | 917 | 941 | 1150 | 1038 | 283 | 4767 | 5388 |
| cluster3429 | 108 | 247 | 279 | 700 | 303 | 566 | 286 | 407 | 414 | 968 |
| cluster3430 | 28 | 93 | 86 | 0 | 437 | 653 | 52 | 9 | 186 | 296 |
| cluster3431 | 43 | 135 | 288 | 0 | 464 | 807 | 122 | 55 | 334 | 832 |
| cluster3432 | 66 | 156 | 63 | 163 | 72 | 98 | 658 | 313 | 225 | 547 |
| cluster3433 | 19 | 64 | 349 | 49 | 118 | 52 | 183 | 12 | 153 | 338 |
| cluster3434 | 55 | 148 | 122 | 211 | 54 | 113 | 939 | 97 | 219 | 647 |
| cluster3435 | 31 | 63 | 455 | 0 | 45 | 177 | 181 | 319 | 231 | 264 |
| cluster3436 | 15 | 45 | 288 | 0 | 745 | 817 | 157 | 51 | 189 | 458 |
| cluster3437 | 66 | 153 | 31 | 79 | 112 | 188 | 187 | 186 | 101 | 451 |
| cluster3438 | 10 | 51 | 55 | 0 | 186 | 175 | 39 | 19 | 69 | 106 |

| | | | | | | | | | | |
|---|---|---|---|---|---|---|---|---|---|---|
| cluster3439 | 79 | 219 | 266 | 0 | 80 | 183 | 406 | 104 | 357 | 939 |
| cluster3440 | 160 | 455 | 1033 | 47 | 576 | 274 | 616 | 231 | 825 | 1442 |
| cluster3441 | 35 | 107 | 392 | 8 | 486 | 336 | 184 | 66 | 246 | 443 |
| cluster3442 | 22 | 81 | 494 | 0 | 1157 | 834 | 409 | 113 | 449 | 712 |
| cluster3443 | 236 | 648 | 1069 | 13 | 5638 | 4199 | 1124 | 892 | 689 | 1539 |
| cluster3444 | 640 | 1558 | 6287 | 442 | 1596 | 1517 | 2208 | 1070 | 2423 | 5771 |
| cluster3445 | 109 | 236 | 126 | 256 | 331 | 421 | 632 | 265 | 450 | 733 |
| cluster3446 | 171 | 380 | 489 | 0 | 150 | 284 | 220 | 263 | 447 | 795 |
| cluster3447 | 66 | 164 | 383 | 0 | 200 | 296 | 264 | 61 | 99 | 203 |
| cluster3448 | 25 | 82 | 163 | 0 | 98 | 54 | 501 | 43 | 57 | 28 |
| cluster3449 | 103 | 205 | 744 | 577 | 247 | 410 | 470 | 120 | 209 | 475 |
| cluster3450 | 4545 | 10741 | 6222 | 0 | 2292 | 2299 | 2442 | 2553 | 3506 | 5974 |
| cluster3451 | 17 | 53 | 330 | 0 | 2079 | 11 | 671 | 6 | 303 | 2761 |
| cluster3452 | 71 | 208 | 258 | 0 | 261 | 584 | 318 | 369 | 148 | 699 |
| cluster3453 | 974 | 2436 | 216 | 1181 | 974 | 690 | 1994 | 2155 | 2189 | 4379 |
| cluster3454 | 601 | 1501 | 825 | 234 | 481 | 740 | 1393 | 885 | 420 | 1517 |
| cluster3455 | 17 | 37 | 134 | 200 | 62 | 86 | 471 | 32 | 406 | 329 |
| cluster3456 | 34 | 60 | 346 | 63 | 118 | 122 | 197 | 29 | 219 | 260 |
| cluster3457 | 76 | 133 | 1493 | 0 | 57 | 210 | 888 | 89 | 4403 | 5776 |
| cluster3458 | 100 | 310 | 265 | 331 | 450 | 837 | 192 | 126 | 728 | 838 |
| cluster3459 | 186 | 462 | 243 | 0 | 330 | 217 | 747 | 146 | 271 | 663 |
| cluster3460 | 29 | 82 | 404 | 0 | 198 | 296 | 110 | 39 | 222 | 325 |
| cluster3461 | 14 | 61 | 797 | 0 | 47 | 186 | 1763 | 1236 | 62 | 139 |
| cluster3462 | 1538 | 4906 | 3137 | 681 | 1256 | 3804 | 4892 | 5122 | 1033 | 11826 |
| cluster3463 | 38 | 110 | 124 | 0 | 523 | 422 | 88 | 84 | 396 | 180 |
| cluster3464 | 26 | 73 | 184 | 0 | 100 | 252 | 142 | 35 | 96 | 146 |
| cluster3465 | 105 | 306 | 313 | 788 | 342 | 297 | 422 | 219 | 460 | 888 |
| cluster3466 | 1048 | 3239 | 203 | 626 | 827 | 880 | 953 | 776 | 2236 | 3595 |
| cluster3467 | 67 | 187 | 332 | 0 | 495 | 517 | 457 | 92 | 104 | 367 |
| cluster3468 | 35 | 128 | 494 | 204 | 101 | 131 | 283 | 27 | 320 | 452 |
| cluster3469 | 160 | 393 | 480 | 768 | 134 | 318 | 188 | 130 | 563 | 1143 |

| | | | | | | | | | | |
|---|---|---|---|---|---|---|---|---|---|---|
| cluster3470 | 272 | 715 | 843 | 32 | 998 | 1020 | 515 | 563 | 436 | 1765 |
| cluster3471 | 191 | 462 | 511 | 0 | 1507 | 1998 | 193 | 311 | 397 | 591 |
| cluster3472 | 203 | 517 | 340 | 1187 | 680 | 984 | 809 | 449 | 679 | 1738 |
| cluster3473 | 39 | 132 | 53 | 227 | 698 | 267 | 282 | 39 | 51 | 536 |
| cluster3474 | 31 | 83 | 162 | 0 | 734 | 1014 | 142 | 32 | 44 | 171 |
| cluster3475 | 41 | 112 | 55 | 0 | 463 | 248 | 129 | 96 | 162 | 545 |
| cluster3476 | 60 | 157 | 170 | 0 | 245 | 437 | 125 | 44 | 27 | 99 |
| cluster3477 | 22 | 50 | 87 | 184 | 72 | 170 | 396 | 20 | 284 | 488 |
| cluster3478 | 65 | 158 | 213 | 516 | 321 | 382 | 215 | 143 | 674 | 1332 |
| cluster3479 | 120 | 251 | 403 | 0 | 103 | 218 | 141 | 87 | 70 | 226 |
| cluster3480 | 42 | 112 | 1636 | 1 | 281 | 387 | 137 | 86 | 144 | 303 |
| cluster3481 | 48 | 111 | 9 | 25 | 5 | 8 | 148 | 12 | 36 | 51 |
| cluster3482 | 84 | 234 | 405 | 33 | 83 | 168 | 1409 | 271 | 224 | 221 |
| cluster3483 | 557 | 1335 | 2070 | 0 | 1697 | 3329 | 941 | 1127 | 3120 | 4579 |
| cluster3484 | 46 | 93 | 1290 | 0 | 140 | 10 | 433 | 332 | 443 | 704 |
| cluster3485 | 3166 | 8190 | 6561 | 0 | 3303 | 4316 | 7844 | 8810 | 3946 | 9844 |
| cluster3486 | 53 | 135 | 251 | 397 | 85 | 93 | 398 | 46 | 237 | 346 |
| cluster3487 | 1702 | 5361 | 3149 | 1 | 1471 | 1102 | 1565 | 2796 | 735 | 1416 |
| cluster3488 | 206 | 550 | 657 | 1 | 164 | 324 | 500 | 186 | 621 | 1062 |
| cluster3489 | 105 | 304 | 615 | 0 | 1030 | 957 | 372 | 258 | 420 | 829 |
| cluster3490 | 38 | 111 | 177 | 0 | 76 | 94 | 169 | 88 | 42 | 205 |
| cluster3491 | 59 | 162 | 647 | 0 | 482 | 495 | 586 | 105 | 290 | 701 |
| cluster3492 | 48 | 144 | 824 | 0 | 304 | 564 | 603 | 77 | 449 | 678 |
| cluster3493 | 10 | 42 | 633 | 0 | 429 | 128 | 247 | 15 | 187 | 271 |
| cluster3494 | 103 | 308 | 386 | 948 | 388 | 440 | 287 | 171 | 225 | 592 |
| cluster3495 | 128 | 375 | 2472 | 0 | 737 | 1174 | 259 | 348 | 599 | 1096 |
| cluster3496 | 22 | 73 | 80 | 145 | 310 | 315 | 322 | 34 | 395 | 275 |
| cluster3497 | 207 | 530 | 1164 | 1414 | 426 | 693 | 684 | 311 | 413 | 1102 |
| cluster3498 | 45 | 111 | 665 | 0 | 149 | 207 | 121 | 46 | 68 | 231 |
| cluster3499 | 77 | 210 | 387 | 0 | 285 | 295 | 311 | 63 | 162 | 408 |
| cluster3500 | 10484 | 24217 | 8138 | 1 | 16792 | 7887 | 4438 | 18041 | 8212 | 11642 |

| | | | | | | | | | | |
|---|---|---|---|---|---|---|---|---|---|---|
| cluster3501 | 1250 | 3553 | 263 | 1022 | 1233 | 1706 | 1914 | 7237 | 956 | 2414 |
| cluster3502 | 51 | 110 | 720 | 0 | 512 | 80 | 251 | 88 | 78 | 154 |
| cluster3503 | 66 | 187 | 164 | 0 | 265 | 660 | 183 | 93 | 142 | 381 |
| cluster3504 | 2186 | 3824 | 3681 | 7525 | 733 | 631 | 782 | 2716 | 893 | 2265 |
| cluster3505 | 165 | 426 | 1099 | 0 | 339 | 422 | 145 | 115 | 784 | 1322 |
| cluster3506 | 120 | 369 | 692 | 0 | 339 | 636 | 202 | 145 | 155 | 358 |
| cluster3507 | 82 | 220 | 680 | 0 | 78 | 33 | 86 | 110 | 138 | 336 |
| cluster3508 | 16 | 42 | 185 | 0 | 222 | 330 | 65 | 15 | 74 | 135 |
| cluster3509 | 37 | 116 | 426 | 16 | 201 | 316 | 155 | 62 | 183 | 593 |
| cluster3510 | 4203 | 9160 | 14061 | 0 | 6 | 0 | 4408 | 11422 | 6017 | 13427 |
| cluster3511 | 657 | 1800 | 1143 | 0 | 1541 | 2708 | 893 | 1587 | 667 | 1861 |
| cluster3512 | 24 | 63 | 58 | 16 | 430 | 205 | 379 | 23 | 47 | 195 |
| cluster3513 | 136 | 357 | 1346 | 990 | 516 | 563 | 952 | 76 | 1993 | 2331 |
| cluster3514 | 28 | 71 | 464 | 616 | 315 | 177 | 224 | 101 | 377 | 713 |
| cluster3515 | 34 | 87 | 832 | 2 | 58 | 24 | 44 | 9 | 128 | 444 |
| cluster3516 | 374 | 925 | 116 | 0 | 549 | 157 | 130 | 228 | 65 | 325 |
| cluster3517 | 417 | 1016 | 2071 | 179 | 1360 | 1811 | 1446 | 822 | 907 | 1914 |
| cluster3518 | 393 | 1003 | 591 | 0 | 399 | 44 | 472 | 169 | 227 | 1462 |
| cluster3519 | 4 | 43 | 33 | 0 | 6 | 5 | 141 | 2 | 55 | 33 |
| cluster3520 | 19 | 71 | 445 | 0 | 13 | 4 | 14 | 0 | 123 | 114 |
| cluster3521 | 49 | 200 | 535 | 0 | 216 | 228 | 221 | 190 | 373 | 489 |
| cluster3522 | 98 | 256 | 157 | 0 | 127 | 185 | 356 | 177 | 275 | 616 |
| cluster3523 | 109 | 222 | 802 | 0 | 579 | 590 | 490 | 198 | 367 | 493 |
| cluster3524 | 65 | 190 | 726 | 7 | 155 | 311 | 480 | 72 | 251 | 730 |
| cluster3525 | 64 | 155 | 303 | 19 | 142 | 172 | 323 | 270 | 450 | 844 |
| cluster3526 | 46 | 150 | 776 | 3 | 275 | 346 | 224 | 120 | 228 | 424 |
| cluster3527 | 39 | 86 | 124 | 0 | 119 | 151 | 64 | 61 | 99 | 186 |
| cluster3528 | 5348 | 12471 | 5646 | 13905 | 7957 | 11756 | 13928 | 6235 | 18558 | 32311 |
| cluster3529 | 30 | 86 | 652 | 0 | 385 | 313 | 491 | 81 | 1706 | 970 |
| cluster3530 | 72 | 216 | 1063 | 1 | 244 | 297 | 267 | 62 | 317 | 595 |
| cluster3531 | 64 | 205 | 82 | 0 | 292 | 582 | 195 | 299 | 82 | 601 |

| | | | | | | | | | | |
|---|---|---|---|---|---|---|---|---|---|---|
| cluster3532 | 26 | 53 | 192 | 471 | 237 | 160 | 858 | 89 | 275 | 479 |
| cluster3533 | 67 | 165 | 177 | 296 | 94 | 1422 | 688 | 197 | 159 | 687 |
| cluster3534 | 129 | 321 | 173 | 149 | 27 | 93 | 1131 | 241 | 1135 | 1022 |
| cluster3535 | 16 | 58 | 333 | 0 | 41 | 92 | 130 | 24 | 117 | 136 |
| cluster3536 | 81 | 177 | 164 | 269 | 426 | 730 | 196 | 119 | 434 | 791 |
| cluster3537 | 21 | 58 | 334 | 0 | 237 | 293 | 81 | 37 | 96 | 180 |
| cluster3538 | 534 | 1235 | 3763 | 3030 | 248 | 1221 | 1046 | 146 | 2096 | 6045 |
| cluster3539 | 302 | 776 | 735 | 0 | 101 | 1030 | 137 | 249 | 360 | 893 |
| cluster3540 | 21 | 85 | 275 | 0 | 54 | 62 | 260 | 43 | 157 | 347 |
| cluster3541 | 186 | 483 | 633 | 1237 | 1072 | 1682 | 586 | 564 | 1552 | 3321 |
| cluster3542 | 15 | 47 | 80 | 0 | 152 | 151 | 170 | 13 | 75 | 123 |
| cluster3543 | 29 | 67 | 485 | 0 | 24 | 29 | 188 | 25 | 36 | 64 |
| cluster3544 | 11133 | 38834 | 41868 | 19528 | 154 | 873 | 42123 | 31469 | 35802 | 40493 |
| cluster3545 | 24 | 89 | 172 | 0 | 231 | 415 | 264 | 42 | 40 | 88 |
| cluster3546 | 532 | 1585 | 114 | 0 | 1031 | 1096 | 584 | 3248 | 117 | 640 |
| cluster3547 | 249 | 702 | 911 | 1496 | 1161 | 1954 | 888 | 737 | 281 | 448 |
| cluster3548 | 22 | 72 | 353 | 151 | 212 | 366 | 264 | 80 | 198 | 378 |
| cluster3549 | 319 | 841 | 916 | 1443 | 1718 | 2646 | 452 | 601 | 1632 | 3750 |
| cluster3550 | 21 | 53 | 131 | 49 | 52 | 49 | 115 | 16 | 176 | 311 |
| cluster3551 | 28 | 87 | 188 | 306 | 93 | 143 | 129 | 19 | 322 | 198 |
| cluster3552 | 329 | 873 | 312 | 621 | 1805 | 3368 | 839 | 347 | 1190 | 2492 |
| cluster3553 | 25 | 38 | 70 | 0 | 81 | 197 | 191 | 27 | 75 | 158 |
| cluster3554 | 29 | 61 | 337 | 0 | 204 | 237 | 336 | 134 | 401 | 756 |
| cluster3555 | 647 | 1795 | 3206 | 0 | 1089 | 5271 | 776 | 1519 | 3108 | 4686 |
| cluster3556 | 30 | 104 | 253 | 0 | 264 | 623 | 60 | 28 | 59 | 48 |
| cluster3557 | 54 | 146 | 256 | 0 | 140 | 207 | 222 | 76 | 122 | 301 |
| cluster3558 | 47 | 135 | 157 | 0 | 138 | 305 | 135 | 57 | 243 | 556 |
| cluster3559 | 27 | 104 | 302 | 0 | 72 | 134 | 123 | 26 | 173 | 546 |
| cluster3560 | 88 | 233 | 407 | 0 | 430 | 637 | 146 | 155 | 290 | 669 |
| cluster3561 | 18 | 52 | 518 | 29 | 94 | 129 | 44 | 6 | 367 | 300 |
| cluster3562 | 48 | 147 | 482 | 0 | 218 | 215 | 196 | 136 | 529 | 581 |

| | | | | | | | | | | |
|---|---|---|---|---|---|---|---|---|---|---|
| cluster3563 | 253 | 680 | 120 | 354 | 195 | 390 | 1221 | 822 | 1048 | 2086 |
| cluster3564 | 21 | 44 | 86 | 574 | 405 | 851 | 313 | 41 | 248 | 429 |
| cluster3565 | 125 | 302 | 148 | 0 | 138 | 283 | 231 | 116 | 232 | 393 |
| cluster3566 | 87 | 221 | 162 | 0 | 101 | 233 | 166 | 44 | 244 | 330 |
| cluster3567 | 75 | 242 | 2156 | 0 | 174 | 230 | 267 | 17 | 2625 | 882 |
| cluster3568 | 81 | 264 | 349 | 0 | 35 | 25 | 182 | 316 | 607 | 1082 |
| cluster3569 | 78 | 167 | 173 | 0 | 544 | 1317 | 385 | 140 | 283 | 415 |
| cluster3570 | 190 | 606 | 898 | 0 | 1896 | 2330 | 1873 | 337 | 649 | 1249 |
| cluster3571 | 22 | 61 | 51 | 162 | 41 | 71 | 906 | 95 | 44 | 150 |
| cluster3572 | 108 | 212 | 140 | 0 | 2120 | 136 | 387 | 20 | 18 | 541 |
| cluster3573 | 188 | 498 | 922 | 0 | 201 | 847 | 476 | 565 | 604 | 1793 |
| cluster3574 | 21 | 70 | 231 | 0 | 47 | 58 | 446 | 385 | 105 | 310 |
| cluster3575 | 114 | 311 | 2771 | 0 | 146 | 279 | 117 | 33 | 889 | 1504 |
| cluster3576 | 39 | 73 | 118 | 1 | 264 | 357 | 106 | 53 | 243 | 287 |
| cluster3577 | 298 | 790 | 670 | 1187 | 1591 | 594 | 436 | 268 | 982 | 1828 |
| cluster3578 | 12 | 37 | 63 | 0 | 14 | 39 | 104 | 7 | 89 | 127 |
| cluster3579 | 112 | 341 | 463 | 992 | 786 | 801 | 1158 | 625 | 534 | 1827 |
| cluster3580 | 19 | 51 | 141 | 0 | 257 | 343 | 52 | 31 | 119 | 180 |
| cluster3581 | 869 | 2120 | 4085 | 9 | 288 | 715 | 1193 | 1925 | 1980 | 3256 |
| cluster3582 | 317 | 911 | 1419 | 1418 | 748 | 1586 | 571 | 406 | 1752 | 2290 |
| cluster3583 | 415 | 1203 | 2802 | 5 | 1556 | 3211 | 1124 | 1252 | 3575 | 6504 |
| cluster3584 | 32 | 61 | 151 | 0 | 265 | 357 | 154 | 18 | 120 | 142 |
| cluster3585 | 39 | 119 | 142 | 87 | 285 | 195 | 627 | 219 | 697 | 841 |
| cluster3586 | 215 | 663 | 1329 | 398 | 948 | 434 | 205 | 371 | 982 | 1040 |
| cluster3587 | 637 | 1722 | 11454 | 0 | 601 | 717 | 814 | 1076 | 643 | 2029 |
| cluster3588 | 43 | 70 | 1643 | 15 | 361 | 286 | 859 | 26 | 555 | 325 |
| cluster3589 | 65 | 149 | 624 | 0 | 205 | 266 | 299 | 67 | 332 | 399 |
| cluster3590 | 23 | 43 | 128 | 0 | 178 | 160 | 123 | 22 | 91 | 242 |
| cluster3591 | 1789 | 4588 | 1328 | 2574 | 2295 | 3539 | 3301 | 2481 | 2240 | 7480 |
| cluster3592 | 78 | 203 | 781 | 0 | 327 | 583 | 347 | 187 | 167 | 506 |
| cluster3593 | 129 | 343 | 155 | 0 | 182 | 288 | 349 | 118 | 104 | 268 |

| | | | | | | | | | | |
|---|---|---|---|---|---|---|---|---|---|---|
| cluster3594 | 84 | 265 | 235 | 0 | 36 | 108 | 199 | 38 | 110 | 123 |
| cluster3595 | 26 | 147 | 162 | 294 | 524 | 1107 | 290 | 116 | 261 | 672 |
| cluster3596 | 7 | 26 | 45 | 0 | 9 | 58 | 37 | 20 | 28 | 70 |
| cluster3597 | 23 | 52 | 425 | 268 | 189 | 287 | 140 | 40 | 395 | 359 |
| cluster3598 | 18 | 50 | 31 | 0 | 6 | 738 | 46 | 24 | 1 | 136 |
| cluster3599 | 252 | 648 | 296 | 0 | 158 | 941 | 1117 | 571 | 429 | 942 |
| cluster3600 | 116 | 309 | 2021 | 9 | 544 | 1262 | 505 | 308 | 1301 | 1979 |
| cluster3601 | 140 | 418 | 667 | 0 | 488 | 865 | 211 | 162 | 346 | 1222 |
| cluster3602 | 8 | 54 | 136 | 92 | 75 | 123 | 142 | 43 | 116 | 146 |
| cluster3603 | 22 | 110 | 543 | 0 | 502 | 324 | 229 | 60 | 315 | 464 |
| cluster3604 | 34 | 51 | 224 | 0 | 177 | 473 | 225 | 5 | 53 | 79 |
| cluster3605 | 26 | 76 | 210 | 0 | 116 | 124 | 103 | 42 | 97 | 170 |
| cluster3606 | 132 | 248 | 556 | 0 | 563 | 707 | 333 | 216 | 196 | 476 |
| cluster3607 | 72 | 174 | 520 | 0 | 572 | 80 | 193 | 87 | 406 | 2224 |
| cluster3608 | 803 | 2063 | 512 | 1092 | 2236 | 2039 | 1757 | 1820 | 1776 | 4206 |
| cluster3609 | 202 | 600 | 1861 | 0 | 538 | 1776 | 103 | 30 | 2336 | 2465 |
| cluster3610 | 27 | 63 | 41 | 0 | 347 | 446 | 442 | 18 | 112 | 524 |
| cluster3611 | 20 | 75 | 117 | 0 | 390 | 311 | 141 | 20 | 32 | 74 |
| cluster3612 | 44 | 112 | 390 | 0 | 218 | 254 | 142 | 77 | 207 | 496 |
| cluster3613 | 17 | 53 | 215 | 212 | 46 | 777 | 271 | 97 | 467 | 609 |
| cluster3614 | 13 | 40 | 367 | 0 | 206 | 582 | 469 | 17 | 77 | 158 |
| cluster3615 | 13 | 35 | 71 | 197 | 139 | 107 | 94 | 9 | 123 | 190 |
| cluster3616 | 736 | 1760 | 535 | 3186 | 3437 | 151 | 782 | 2644 | 61 | 322 |
| cluster3617 | 229 | 647 | 1297 | 0 | 72 | 49 | 40 | 55 | 273 | 319 |
| cluster3618 | 14 | 19 | 620 | 0 | 34 | 14 | 236 | 6 | 356 | 108 |
| cluster3619 | 44 | 116 | 268 | 13 | 515 | 725 | 348 | 61 | 127 | 246 |
| cluster3620 | 20 | 62 | 153 | 93 | 201 | 195 | 593 | 61 | 110 | 186 |
| cluster3621 | 38 | 99 | 389 | 0 | 166 | 167 | 60 | 4 | 91 | 131 |
| cluster3622 | 140 | 391 | 1153 | 1 | 241 | 313 | 322 | 258 | 422 | 615 |
| cluster3623 | 195 | 545 | 628 | 0 | 513 | 806 | 189 | 461 | 352 | 632 |
| cluster3624 | 87 | 319 | 629 | 0 | 426 | 1416 | 393 | 262 | 470 | 851 |

| | | | | | | | | | | |
|---|---|---|---|---|---|---|---|---|---|---|
| cluster3625 | 30 | 64 | 261 | 0 | 187 | 344 | 117 | 113 | 136 | 238 |
| cluster3626 | 36 | 102 | 83 | 198 | 112 | 101 | 188 | 96 | 114 | 459 |
| cluster3627 | 38 | 96 | 540 | 2 | 150 | 151 | 331 | 110 | 182 | 336 |
| cluster3628 | 25 | 56 | 346 | 0 | 356 | 425 | 435 | 58 | 107 | 303 |
| cluster3629 | 19 | 55 | 151 | 0 | 484 | 599 | 171 | 27 | 55 | 171 |
| cluster3630 | 190 | 390 | 1634 | 304 | 247 | 730 | 597 | 389 | 658 | 1261 |
| cluster3631 | 18 | 81 | 438 | 961 | 131 | 198 | 189 | 58 | 153 | 174 |
| cluster3632 | 68 | 153 | 113 | 262 | 181 | 248 | 133 | 107 | 185 | 340 |
| cluster3633 | 9784 | 33126 | 7619 | 21001 | 13377 | 13838 | 38347 | 27555 | 25935 | 47774 |
| cluster3634 | 73 | 213 | 299 | 119 | 105 | 199 | 133 | 115 | 335 | 495 |
| cluster3635 | 111 | 450 | 742 | 406 | 217 | 445 | 542 | 280 | 578 | 1050 |
| cluster3636 | 493 | 1104 | 415 | 553 | 982 | 1615 | 2491 | 1179 | 1503 | 3344 |
| cluster3637 | 90 | 242 | 730 | 81 | 470 | 644 | 346 | 221 | 333 | 628 |
| cluster3638 | 279 | 717 | 244 | 2 | 2957 | 215 | 485 | 131 | 297 | 1008 |
| cluster3639 | 18 | 62 | 0 | 42 | 774 | 1747 | 775 | 188 | 12 | 55 |
| cluster3640 | 1305 | 3685 | 93 | 0 | 713 | 4 | 395 | 646 | 62 | 1924 |
| cluster3641 | 43 | 119 | 237 | 298 | 150 | 199 | 565 | 79 | 350 | 550 |
| cluster3642 | 14 | 84 | 414 | 0 | 356 | 576 | 454 | 58 | 134 | 375 |
| cluster3643 | 20 | 78 | 270 | 0 | 205 | 294 | 135 | 28 | 160 | 273 |
| cluster3644 | 166 | 486 | 145 | 93 | 417 | 76 | 293 | 406 | 142 | 711 |
| cluster3645 | 807 | 1792 | 1667 | 0 | 831 | 202 | 2065 | 9723 | 92 | 3235 |
| cluster3646 | 19 | 57 | 217 | 0 | 391 | 469 | 455 | 32 | 101 | 172 |
| cluster3647 | 42 | 90 | 138 | 0 | 189 | 262 | 62 | 56 | 69 | 156 |
| cluster3648 | 474 | 1249 | 581 | 1004 | 1094 | 3291 | 727 | 892 | 2915 | 5801 |
| cluster3649 | 87 | 204 | 435 | 10 | 118 | 205 | 241 | 138 | 362 | 614 |
| cluster3650 | 131 | 354 | 3840 | 119 | 64 | 79 | 308 | 177 | 449 | 543 |
| cluster3651 | 22 | 45 | 6 | 46 | 57 | 5 | 423 | 237 | 5 | 173 |
| cluster3652 | 54 | 150 | 327 | 832 | 1750 | 1579 | 281 | 118 | 96 | 308 |
| cluster3653 | 267 | 641 | 374 | 1027 | 291 | 602 | 278 | 163 | 228 | 1368 |
| cluster3654 | 39 | 118 | 190 | 1 | 295 | 279 | 116 | 207 | 107 | 239 |
| cluster3655 | 32 | 92 | 246 | 0 | 143 | 120 | 194 | 63 | 76 | 384 |

| | | | | | | | | | | |
|---|---|---|---|---|---|---|---|---|---|---|
| cluster3656 | 28 | 102 | 122 | 214 | 326 | 405 | 77 | 60 | 177 | 221 |
| cluster3657 | 125 | 394 | 366 | 0 | 721 | 1323 | 340 | 112 | 183 | 598 |
| cluster3658 | 4249 | 11265 | 8684 | 8341 | 4270 | 15393 | 3552 | 3267 | 35002 | 26970 |
| cluster3659 | 706 | 1446 | 645 | 0 | 923 | 484 | 752 | 918 | 338 | 1353 |
| cluster3660 | 22 | 80 | 27 | 0 | 175 | 76 | 81 | 55 | 8 | 412 |
| cluster3661 | 41 | 99 | 493 | 100 | 431 | 878 | 354 | 57 | 162 | 506 |
| cluster3662 | 20 | 55 | 225 | 0 | 38 | 57 | 125 | 72 | 69 | 154 |
| cluster3663 | 119 | 299 | 705 | 0 | 253 | 477 | 235 | 222 | 191 | 251 |
| cluster3664 | 48 | 92 | 140 | 2 | 73 | 99 | 57 | 19 | 101 | 166 |
| cluster3665 | 82 | 219 | 599 | 0 | 194 | 372 | 141 | 313 | 170 | 477 |
| cluster3666 | 92 | 262 | 502 | 0 | 4901 | 9357 | 444 | 184 | 367 | 616 |
| cluster3667 | 46 | 99 | 418 | 22 | 165 | 399 | 396 | 327 | 158 | 528 |
| cluster3668 | 132 | 328 | 368 | 0 | 362 | 663 | 263 | 186 | 251 | 397 |
| cluster3669 | 944 | 2312 | 1282 | 1 | 2717 | 1393 | 1083 | 1401 | 1679 | 2806 |
| cluster3670 | 1030 | 2731 | 142 | 567 | 234 | 4690 | 942 | 1296 | 366 | 1079 |
| cluster3671 | 445 | 963 | 1604 | 0 | 1973 | 1501 | 504 | 1022 | 783 | 1268 |
| cluster3672 | 81 | 257 | 128 | 0 | 121 | 190 | 515 | 239 | 176 | 321 |
| cluster3673 | 12 | 36 | 221 | 0 | 199 | 223 | 423 | 36 | 106 | 176 |
| cluster3674 | 204 | 515 | 74 | 0 | 28 | 38 | 70 | 27 | 26 | 329 |
| cluster3675 | 593 | 1248 | 1497 | 1016 | 478 | 979 | 1345 | 1109 | 2673 | 3609 |
| cluster3676 | 37 | 114 | 691 | 0 | 464 | 736 | 136 | 133 | 300 | 401 |
| cluster3677 | 106 | 316 | 130 | 446 | 978 | 1512 | 306 | 357 | 338 | 884 |
| cluster3678 | 67 | 143 | 178 | 59 | 442 | 588 | 94 | 121 | 112 | 245 |
| cluster3679 | 234 | 504 | 255 | 479 | 560 | 1275 | 124 | 79 | 464 | 1118 |
| cluster3680 | 57 | 161 | 525 | 0 | 317 | 434 | 268 | 115 | 95 | 620 |
| cluster3681 | 30 | 103 | 170 | 0 | 387 | 660 | 356 | 94 | 35 | 247 |
| cluster3682 | 9 | 41 | 251 | 0 | 64 | 92 | 253 | 35 | 125 | 274 |
| cluster3683 | 74 | 128 | 340 | 0 | 720 | 1308 | 583 | 207 | 437 | 612 |
| cluster3684 | 35 | 90 | 333 | 0 | 73 | 125 | 167 | 23 | 148 | 212 |
| cluster3685 | 183 | 396 | 1928 | 0 | 213 | 117 | 606 | 204 | 638 | 1584 |
| cluster3686 | 55 | 155 | 136 | 0 | 138 | 192 | 67 | 18 | 54 | 76 |

| | | | | | | | | | | |
|---|---|---|---|---|---|---|---|---|---|---|
| cluster3687 | 18 | 69 | 187 | 0 | 143 | 137 | 120 | 50 | 97 | 161 |
| cluster3688 | 239 | 617 | 1328 | 0 | 930 | 1761 | 408 | 448 | 321 | 778 |
| cluster3689 | 40 | 161 | 0 | 23 | 283 | 19 | 1122 | 43 | 4 | 41 |
| cluster3690 | 197 | 505 | 191 | 464 | 205 | 351 | 388 | 316 | 730 | 1080 |
| cluster3691 | 31 | 75 | 440 | 3 | 155 | 308 | 103 | 11 | 182 | 229 |
| cluster3692 | 37 | 98 | 395 | 0 | 82 | 165 | 114 | 67 | 101 | 183 |
| cluster3693 | 19 | 67 | 60 | 0 | 338 | 435 | 80 | 60 | 50 | 93 |
| cluster3694 | 1318 | 2618 | 431 | 467 | 581 | 1340 | 2979 | 2260 | 1531 | 4003 |
| cluster3695 | 94 | 243 | 72 | 3 | 218 | 331 | 308 | 190 | 179 | 414 |
| cluster3696 | 77 | 277 | 724 | 3 | 150 | 291 | 214 | 157 | 126 | 445 |
| cluster3697 | 330 | 603 | 473 | 1387 | 1639 | 19 | 579 | 512 | 469 | 3300 |
| cluster3698 | 72 | 231 | 315 | 273 | 130 | 167 | 419 | 68 | 509 | 948 |
| cluster3699 | 67 | 156 | 374 | 0 | 305 | 493 | 109 | 71 | 222 | 226 |
| cluster3700 | 61 | 231 | 183 | 0 | 182 | 377 | 128 | 86 | 143 | 449 |
| cluster3701 | 54 | 124 | 54 | 136 | 102 | 199 | 197 | 53 | 80 | 220 |
| cluster3702 | 34 | 92 | 127 | 0 | 319 | 346 | 217 | 43 | 337 | 489 |
| cluster3703 | 105 | 385 | 435 | 583 | 1139 | 1916 | 812 | 266 | 337 | 1176 |
| cluster3704 | 495 | 1161 | 837 | 2361 | 3348 | 5184 | 559 | 229 | 436 | 985 |
| cluster3705 | 12 | 36 | 54 | 173 | 116 | 79 | 63 | 19 | 102 | 71 |
| cluster3706 | 55 | 122 | 225 | 0 | 329 | 292 | 363 | 94 | 246 | 435 |
| cluster3707 | 13 | 29 | 281 | 0 | 67 | 99 | 145 | 10 | 6 | 40 |
| cluster3708 | 3 | 31 | 61 | 0 | 90 | 25 | 133 | 1 | 7 | 78 |
| cluster3709 | 87 | 356 | 3383 | 302 | 738 | 627 | 692 | 345 | 2742 | 3498 |
| cluster3710 | 21 | 63 | 88 | 0 | 138 | 337 | 117 | 38 | 31 | 127 |
| cluster3711 | 739 | 2125 | 6233 | 191 | 1863 | 1240 | 2581 | 1554 | 2511 | 4124 |
| cluster3712 | 22 | 63 | 16 | 100 | 115 | 144 | 60 | 22 | 102 | 81 |
| cluster3713 | 17 | 60 | 234 | 0 | 82 | 128 | 88 | 23 | 19 | 60 |
| cluster3714 | 23 | 58 | 125 | 302 | 61 | 84 | 109 | 45 | 162 | 209 |
| cluster3715 | 40 | 159 | 29 | 4 | 162 | 360 | 46 | 30 | 2 | 57 |
| cluster3716 | 9 | 27 | 48 | 0 | 42 | 28 | 29 | 47 | 12 | 186 |
| cluster3717 | 499 | 1134 | 421 | 689 | 1242 | 1211 | 501 | 865 | 311 | 1692 |

| | | | | | | | | | | |
|---|---|---|---|---|---|---|---|---|---|---|
| cluster3718 | 30 | 77 | 170 | 0 | 53 | 66 | 70 | 29 | 26 | 54 |
| cluster3719 | 13 | 37 | 119 | 2 | 172 | 250 | 49 | 1 | 38 | 59 |
| cluster3720 | 73 | 230 | 497 | 0 | 405 | 598 | 430 | 171 | 254 | 445 |
| cluster3721 | 8 | 30 | 123 | 172 | 40 | 105 | 116 | 14 | 119 | 95 |
| cluster3722 | 39 | 95 | 279 | 0 | 110 | 139 | 65 | 33 | 137 | 171 |
| cluster3723 | 101 | 293 | 634 | 37 | 753 | 1189 | 473 | 213 | 213 | 680 |
| cluster3724 | 22 | 91 | 227 | 0 | 577 | 420 | 245 | 79 | 124 | 213 |
| cluster3725 | 32 | 118 | 371 | 0 | 400 | 463 | 320 | 107 | 144 | 346 |
| cluster3726 | 33 | 112 | 245 | 7 | 70 | 90 | 145 | 48 | 141 | 378 |
| cluster3727 | 39 | 96 | 581 | 0 | 96 | 160 | 175 | 83 | 244 | 342 |
| cluster3728 | 4208 | 10379 | 4486 | 8010 | 9545 | 12210 | 3379 | 8079 | 4067 | 10212 |
| cluster3729 | 172 | 412 | 1510 | 0 | 523 | 778 | 698 | 583 | 667 | 2063 |
| cluster3730 | 70 | 245 | 852 | 663 | 251 | 403 | 393 | 132 | 426 | 747 |
| cluster3731 | 17 | 37 | 164 | 298 | 188 | 459 | 278 | 36 | 128 | 251 |
| cluster3732 | 92 | 222 | 112 | 0 | 352 | 544 | 428 | 126 | 46 | 147 |
| cluster3733 | 28 | 80 | 16 | 14 | 126 | 39 | 57 | 28 | 72 | 68 |
| cluster3734 | 82 | 256 | 496 | 0 | 978 | 1105 | 222 | 160 | 258 | 668 |
| cluster3735 | 482 | 1302 | 365 | 1040 | 1017 | 1265 | 1719 | 2952 | 547 | 2735 |
| cluster3736 | 10 | 28 | 46 | 0 | 17 | 43 | 75 | 1 | 62 | 77 |
| cluster3737 | 113 | 356 | 525 | 111 | 434 | 795 | 413 | 168 | 515 | 1315 |
| cluster3738 | 15 | 34 | 446 | 1594 | 283 | 469 | 185 | 69 | 90 | 251 |
| cluster3739 | 800 | 1744 | 5953 | 591 | 1839 | 2734 | 1764 | 1105 | 2046 | 4807 |
| cluster3740 | 10 | 53 | 179 | 1 | 298 | 414 | 175 | 178 | 146 | 297 |
| cluster3741 | 150 | 494 | 3266 | 90 | 586 | 1718 | 330 | 121 | 3632 | 3961 |
| cluster3742 | 43 | 121 | 235 | 0 | 529 | 424 | 73 | 42 | 202 | 355 |
| cluster3743 | 72 | 247 | 318 | 0 | 369 | 405 | 257 | 191 | 150 | 347 |
| cluster3744 | 37 | 79 | 400 | 0 | 242 | 224 | 570 | 78 | 306 | 468 |
| cluster3745 | 12 | 46 | 765 | 0 | 387 | 2765 | 145 | 96 | 182 | 1490 |
| cluster3746 | 11 | 45 | 54 | 120 | 133 | 162 | 77 | 24 | 35 | 37 |
| cluster3747 | 109 | 314 | 1413 | 10 | 477 | 797 | 258 | 117 | 208 | 493 |
| cluster3748 | 174 | 578 | 286 | 1162 | 488 | 1410 | 905 | 293 | 313 | 876 |

| | | | | | | | | | | |
|---|---|---|---|---|---|---|---|---|---|---|
| cluster3749 | 802 | 1806 | 389 | 0 | 1443 | 1460 | 800 | 1070 | 456 | 1185 |
| cluster3750 | 8 | 30 | 232 | 0 | 118 | 66 | 907 | 71 | 175 | 199 |
| cluster3751 | 18 | 40 | 195 | 1 | 199 | 192 | 277 | 71 | 119 | 268 |
| cluster3752 | 416 | 1200 | 31 | 0 | 155 | 93 | 859 | 271 | 99 | 538 |
| cluster3753 | 13496 | 34137 | 52416 | 319 | 14 | 0 | 37436 | 33057 | 30526 | 46484 |
| cluster3754 | 31 | 61 | 144 | 12 | 76 | 309 | 166 | 46 | 33 | 185 |
| cluster3755 | 16 | 37 | 468 | 1695 | 160 | 1023 | 249 | 2 | 31 | 370 |
| cluster3756 | 90 | 256 | 321 | 0 | 179 | 298 | 204 | 265 | 292 | 602 |
| cluster3757 | 19 | 31 | 2109 | 0 | 7 | 187 | 108 | 19 | 277 | 230 |
| cluster3758 | 29 | 86 | 64 | 157 | 260 | 173 | 1088 | 72 | 97 | 220 |
| cluster3759 | 16 | 30 | 61 | 91 | 59 | 81 | 219 | 25 | 157 | 281 |
| cluster3760 | 92 | 211 | 495 | 752 | 512 | 964 | 303 | 132 | 44 | 264 |
| cluster3761 | 118 | 283 | 1604 | 0 | 37 | 78 | 66 | 98 | 382 | 498 |
| cluster3762 | 20 | 49 | 322 | 366 | 191 | 426 | 672 | 16 | 705 | 874 |
| cluster3763 | 7 | 37 | 182 | 235 | 97 | 90 | 35 | 17 | 149 | 182 |
| cluster3764 | 172 | 454 | 182 | 0 | 345 | 473 | 127 | 418 | 132 | 610 |
| cluster3765 | 334 | 980 | 159 | 604 | 976 | 1036 | 448 | 1310 | 641 | 1230 |
| cluster3766 | 2374 | 5355 | 3097 | 0 | 1324 | 1709 | 2225 | 3117 | 2142 | 5177 |
| cluster3767 | 70 | 107 | 121 | 0 | 124 | 314 | 47 | 67 | 105 | 209 |
| cluster3768 | 196 | 402 | 3031 | 0 | 295 | 603 | 1481 | 706 | 1090 | 1835 |
| cluster3769 | 63 | 126 | 316 | 27 | 231 | 401 | 615 | 86 | 657 | 409 |
| cluster3770 | 247 | 609 | 67 | 0 | 2032 | 13 | 177 | 1147 | 7 | 250 |
| cluster3771 | 18 | 48 | 162 | 0 | 223 | 278 | 83 | 28 | 138 | 184 |
| cluster3772 | 37 | 100 | 234 | 1 | 33 | 66 | 382 | 79 | 307 | 367 |
| cluster3773 | 999 | 3842 | 1344 | 722 | 2229 | 1200 | 4597 | 2230 | 1305 | 4484 |
| cluster3774 | 48 | 107 | 605 | 85 | 102 | 234 | 454 | 37 | 639 | 581 |
| cluster3775 | 105 | 261 | 325 | 0 | 642 | 1448 | 411 | 665 | 989 | 1260 |
| cluster3776 | 20 | 45 | 30 | 91 | 43 | 36 | 163 | 23 | 105 | 75 |
| cluster3777 | 44 | 122 | 65 | 2 | 63 | 55 | 75 | 39 | 76 | 338 |
| cluster3778 | 775 | 1612 | 1616 | 4450 | 947 | 1296 | 1450 | 667 | 336 | 1320 |
| cluster3779 | 186 | 452 | 187 | 484 | 503 | 686 | 151 | 88 | 101 | 347 |

| | | | | | | | | | | |
|---|---|---|---|---|---|---|---|---|---|---|
| cluster3780 | 524 | 1485 | 1150 | 0 | 13 | 10 | 87 | 69 | 34 | 58 |
| cluster3781 | 22 | 58 | 25 | 48 | 120 | 224 | 105 | 7 | 55 | 111 |
| cluster3782 | 8 | 20 | 188 | 3 | 84 | 127 | 167 | 49 | 344 | 536 |
| cluster3783 | 31 | 80 | 207 | 0 | 435 | 578 | 535 | 59 | 100 | 197 |
| cluster3784 | 6 | 23 | 17 | 41 | 41 | 74 | 36 | 19 | 25 | 44 |
| cluster3785 | 25 | 30 | 608 | 2 | 191 | 146 | 393 | 18 | 733 | 396 |
| cluster3786 | 6 | 24 | 393 | 1018 | 476 | 1740 | 161 | 45 | 384 | 1465 |
| cluster3787 | 156 | 331 | 451 | 0 | 61 | 63 | 294 | 188 | 48 | 156 |
| cluster3788 | 3025 | 11413 | 47960 | 10945 | 6742 | 7987 | 9667 | 4785 | 13093 | 21839 |
| cluster3789 | 12 | 33 | 55 | 100 | 213 | 201 | 463 | 9 | 460 | 113 |
| cluster3790 | 14 | 24 | 182 | 0 | 83 | 117 | 141 | 30 | 21 | 552 |
| cluster3791 | 145 | 444 | 443 | 3 | 130 | 344 | 17 | 21 | 405 | 467 |
| cluster3792 | 270 | 602 | 940 | 5 | 520 | 770 | 1055 | 716 | 596 | 2401 |
| cluster3793 | 97 | 294 | 330 | 0 | 661 | 651 | 172 | 269 | 112 | 508 |
| cluster3794 | 16 | 46 | 63 | 0 | 58 | 133 | 228 | 43 | 21 | 110 |
| cluster3795 | 131 | 364 | 322 | 0 | 389 | 545 | 225 | 551 | 361 | 765 |
| cluster3796 | 20 | 57 | 84 | 0 | 23 | 4 | 17 | 19 | 43 | 36 |
| cluster3797 | 21 | 49 | 511 | 0 | 120 | 142 | 321 | 27 | 238 | 396 |
| cluster3798 | 16 | 43 | 229 | 1 | 78 | 185 | 179 | 40 | 96 | 206 |
| cluster3799 | 11 | 48 | 48 | 109 | 130 | 163 | 75 | 28 | 130 | 123 |
| cluster3800 | 3719 | 8245 | 6959 | 463 | 7 | 8 | 6217 | 7719 | 2098 | 6988 |
| cluster3801 | 13 | 38 | 168 | 0 | 126 | 226 | 90 | 36 | 69 | 105 |
| cluster3802 | 9 | 31 | 52 | 202 | 24 | 182 | 19 | 10 | 4 | 17 |
| cluster3803 | 334 | 1146 | 1574 | 0 | 1362 | 3355 | 499 | 847 | 587 | 3814 |
| cluster3804 | 12 | 58 | 188 | 0 | 463 | 526 | 350 | 33 | 87 | 162 |
| cluster3805 | 12 | 55 | 585 | 0 | 126 | 216 | 66 | 87 | 28 | 134 |
| cluster3806 | 109 | 265 | 384 | 0 | 245 | 402 | 132 | 259 | 198 | 461 |
| cluster3807 | 27 | 109 | 100 | 194 | 76 | 125 | 256 | 109 | 217 | 446 |
| cluster3808 | 50 | 141 | 525 | 0 | 195 | 356 | 176 | 79 | 365 | 373 |
| cluster3809 | 26 | 90 | 401 | 0 | 297 | 365 | 365 | 113 | 198 | 282 |
| cluster3810 | 28 | 87 | 79 | 178 | 271 | 384 | 141 | 82 | 36 | 124 |

| | | | | | | | | | | |
|---|---|---|---|---|---|---|---|---|---|---|
| cluster3811 | 194 | 563 | 1303 | 456 | 429 | 655 | 370 | 308 | 368 | 1530 |
| cluster3812 | 10 | 28 | 373 | 4 | 17 | 20 | 111 | 11 | 222 | 248 |
| cluster3813 | 19 | 60 | 91 | 0 | 133 | 201 | 75 | 26 | 42 | 94 |
| cluster3814 | 58 | 154 | 114 | 261 | 37 | 156 | 268 | 47 | 429 | 351 |
| cluster3815 | 10 | 28 | 122 | 0 | 251 | 336 | 140 | 34 | 50 | 153 |
| cluster3816 | 160 | 393 | 1379 | 0 | 486 | 518 | 364 | 292 | 624 | 899 |
| cluster3817 | 10 | 40 | 129 | 335 | 38 | 83 | 204 | 44 | 81 | 245 |
| cluster3818 | 17 | 38 | 115 | 382 | 161 | 315 | 58 | 45 | 49 | 79 |
| cluster3819 | 16 | 32 | 180 | 0 | 4 | 387 | 104 | 158 | 17 | 168 |
| cluster3820 | 55 | 169 | 369 | 0 | 231 | 270 | 343 | 189 | 155 | 464 |
| cluster3821 | 102 | 358 | 25 | 203 | 391 | 164 | 487 | 469 | 2369 | 2725 |
| cluster3822 | 11 | 29 | 462 | 0 | 477 | 567 | 104 | 35 | 10 | 20 |
| cluster3823 | 15 | 39 | 212 | 131 | 154 | 271 | 483 | 90 | 132 | 208 |
| cluster3824 | 34 | 71 | 110 | 0 | 662 | 617 | 69 | 127 | 47 | 214 |
| cluster3825 | 36 | 132 | 133 | 0 | 101 | 85 | 392 | 17 | 131 | 270 |
| cluster3826 | 9 | 28 | 96 | 0 | 22 | 48 | 105 | 13 | 11 | 79 |
| cluster3827 | 26 | 92 | 13 | 47 | 151 | 305 | 267 | 202 | 2 | 132 |
| cluster3828 | 12 | 41 | 158 | 0 | 135 | 211 | 147 | 42 | 105 | 110 |
| cluster3829 | 8 | 29 | 223 | 0 | 105 | 156 | 77 | 23 | 65 | 157 |
| cluster3830 | 268 | 656 | 4881 | 251 | 3737 | 3577 | 1458 | 594 | 1373 | 1837 |
| cluster3831 | 72 | 134 | 101 | 0 | 318 | 433 | 182 | 122 | 80 | 252 |
| cluster3832 | 104 | 248 | 751 | 0 | 523 | 746 | 137 | 138 | 385 | 665 |
| cluster3833 | 46 | 97 | 43 | 0 | 66 | 170 | 74 | 3 | 194 | 299 |
| cluster3834 | 104 | 349 | 436 | 0 | 211 | 187 | 222 | 608 | 87 | 429 |
| cluster3835 | 25 | 92 | 224 | 0 | 66 | 158 | 156 | 40 | 86 | 223 |
| cluster3836 | 63 | 240 | 391 | 0 | 615 | 1052 | 907 | 176 | 197 | 635 |
| cluster3837 | 9 | 27 | 147 | 0 | 192 | 242 | 215 | 62 | 127 | 201 |
| cluster3838 | 26 | 76 | 193 | 0 | 83 | 185 | 120 | 35 | 232 | 416 |
| cluster3839 | 23 | 64 | 52 | 0 | 2 | 9 | 153 | 49 | 127 | 185 |
| cluster3840 | 68 | 155 | 283 | 890 | 684 | 782 | 141 | 61 | 278 | 413 |
| cluster3841 | 149 | 415 | 834 | 0 | 158 | 240 | 220 | 260 | 478 | 568 |

| | | | | | | | | | | |
|---|---|---|---|---|---|---|---|---|---|---|
| cluster3842 | 6 | 19 | 43 | 59 | 129 | 80 | 71 | 4 | 243 | 247 |
| cluster3843 | 38 | 77 | 199 | 10 | 276 | 206 | 165 | 95 | 54 | 143 |
| cluster3844 | 24 | 82 | 7 | 0 | 13 | 35 | 197 | 4 | 2 | 136 |
| cluster3845 | 9953 | 25199 | 12922 | 28298 | 4 | 4 | 21432 | 20927 | 21674 | 36350 |
| cluster3846 | 25 | 124 | 86 | 47 | 95 | 105 | 109 | 61 | 73 | 243 |
| cluster3847 | 227 | 680 | 827 | 0 | 505 | 679 | 127 | 63 | 507 | 716 |
| cluster3848 | 131 | 326 | 1375 | 0 | 67 | 797 | 148 | 443 | 1249 | 1609 |
| cluster3849 | 24 | 48 | 199 | 0 | 155 | 172 | 29 | 18 | 44 | 81 |
| cluster3850 | 4723 | 12338 | 6797 | 0 | 7611 | 5000 | 4235 | 11767 | 5401 | 14852 |
| cluster3851 | 92 | 193 | 199 | 0 | 112 | 216 | 59 | 22 | 43 | 51 |
| cluster3852 | 17 | 64 | 181 | 431 | 352 | 509 | 72 | 44 | 52 | 129 |
| cluster3853 | 21 | 65 | 219 | 0 | 577 | 1326 | 175 | 66 | 69 | 135 |
| cluster3854 | 99 | 266 | 192 | 0 | 175 | 370 | 296 | 195 | 39 | 301 |
| cluster3855 | 16 | 28 | 55 | 0 | 101 | 116 | 93 | 19 | 12 | 46 |
| cluster3856 | 469 | 944 | 1387 | 0 | 1822 | 2392 | 508 | 1103 | 881 | 1238 |
| cluster3857 | 11 | 49 | 97 | 0 | 92 | 126 | 161 | 52 | 87 | 173 |
| cluster3858 | 16 | 78 | 266 | 616 | 341 | 635 | 327 | 125 | 36 | 121 |
| cluster3859 | 434 | 1382 | 5160 | 0 | 996 | 2268 | 869 | 1512 | 2151 | 32416 |
| cluster3860 | 15 | 35 | 115 | 48 | 295 | 57 | 113 | 55 | 208 | 252 |
| cluster3861 | 61 | 200 | 179 | 0 | 140 | 273 | 128 | 100 | 93 | 311 |
| cluster3862 | 123 | 294 | 853 | 0 | 654 | 1066 | 366 | 296 | 552 | 1354 |
| cluster3863 | 14 | 26 | 57 | 135 | 89 | 210 | 37 | 21 | 54 | 121 |
| cluster3864 | 55 | 106 | 180 | 0 | 263 | 305 | 189 | 62 | 69 | 189 |
| cluster3865 | 10 | 31 | 63 | 0 | 168 | 201 | 11 | 9 | 13 | 28 |
| cluster3866 | 16 | 23 | 25 | 0 | 54 | 156 | 50 | 10 | 0 | 34 |
| cluster3867 | 990 | 2487 | 4575 | 413 | 1077 | 1359 | 2168 | 1585 | 1968 | 3428 |
| cluster3868 | 67 | 227 | 237 | 0 | 294 | 289 | 548 | 166 | 326 | 578 |
| cluster3869 | 9 | 37 | 54 | 0 | 331 | 314 | 60 | 28 | 55 | 102 |
| cluster3870 | 41 | 132 | 128 | 0 | 52 | 16 | 56 | 13 | 54 | 131 |
| cluster3871 | 31 | 72 | 188 | 0 | 277 | 336 | 79 | 37 | 39 | 91 |
| cluster3872 | 13 | 24 | 210 | 0 | 81 | 213 | 93 | 10 | 74 | 130 |

| | | | | | | | | | | |
|---|---|---|---|---|---|---|---|---|---|---|
| cluster3873 | 233 | 552 | 1022 | 680 | 685 | 1153 | 325 | 282 | 744 | 1105 |
| cluster3874 | 21 | 42 | 262 | 54 | 133 | 169 | 333 | 44 | 314 | 299 |
| cluster3875 | 51 | 88 | 136 | 5 | 491 | 717 | 268 | 69 | 214 | 235 |
| cluster3876 | 397 | 998 | 1574 | 2004 | 1446 | 3785 | 781 | 284 | 2572 | 2864 |
| cluster3877 | 131 | 381 | 120 | 276 | 581 | 544 | 342 | 477 | 272 | 426 |
| cluster3879 | 12 | 61 | 120 | 0 | 126 | 133 | 65 | 36 | 52 | 135 |
| cluster3880 | 39 | 117 | 471 | 0 | 5 | 15 | 55 | 6 | 46 | 38 |
| cluster3881 | 18 | 48 | 147 | 0 | 63 | 143 | 42 | 18 | 36 | 50 |
| cluster3882 | 10 | 20 | 61 | 0 | 27 | 29 | 37 | 0 | 7 | 37 |
| cluster3883 | 10 | 21 | 1398 | 0 | 36 | 487 | 28 | 148 | 450 | 852 |
| cluster3884 | 8 | 29 | 205 | 0 | 196 | 207 | 927 | 12 | 88 | 145 |
| cluster3885 | 294 | 598 | 1458 | 0 | 258 | 495 | 1834 | 515 | 5728 | 5191 |
| cluster3886 | 52 | 133 | 1052 | 0 | 365 | 423 | 279 | 150 | 142 | 237 |
| cluster3887 | 10 | 38 | 19 | 0 | 44 | 53 | 89 | 36 | 14 | 67 |
| cluster3888 | 62 | 183 | 552 | 0 | 191 | 167 | 176 | 133 | 197 | 224 |
| cluster3889 | 3 | 13 | 22 | 0 | 88 | 174 | 60 | 16 | 3 | 15 |
| cluster3890 | 48 | 156 | 396 | 927 | 327 | 648 | 41 | 109 | 101 | 263 |
| cluster3891 | 1133 | 2558 | 400 | 652 | 3688 | 3894 | 251 | 537 | 335 | 327 |
| cluster3892 | 394 | 1342 | 54 | 67 | 364 | 158 | 1917 | 155 | 56 | 551 |
| cluster3893 | 73 | 250 | 155 | 345 | 404 | 521 | 475 | 221 | 250 | 721 |
| cluster3894 | 45 | 109 | 83 | 0 | 62 | 129 | 156 | 115 | 29 | 234 |
| cluster3895 | 21 | 56 | 136 | 0 | 159 | 264 | 199 | 32 | 94 | 253 |
| cluster3896 | 122 | 352 | 578 | 0 | 275 | 547 | 217 | 133 | 238 | 1003 |
| cluster3897 | 43 | 132 | 91 | 301 | 277 | 536 | 40 | 19 | 95 | 184 |
| cluster3898 | 12 | 20 | 161 | 192 | 193 | 371 | 172 | 5 | 196 | 153 |
| cluster3899 | 29 | 82 | 251 | 0 | 139 | 322 | 49 | 27 | 16 | 96 |
| cluster3900 | 2581 | 7342 | 9656 | 0 | 8627 | 760 | 973 | 1611 | 3019 | 9956 |
| cluster3901 | 31 | 85 | 136 | 0 | 1140 | 1932 | 266 | 46 | 75 | 124 |
| cluster3902 | 109 | 288 | 499 | 0 | 1942 | 1371 | 638 | 138 | 446 | 415 |
| cluster3903 | 133 | 429 | 1524 | 0 | 426 | 1065 | 452 | 500 | 841 | 1659 |
| cluster3904 | 675 | 2025 | 652 | 0 | 672 | 849 | 1301 | 3133 | 665 | 1455 |

| | | | | | | | | | | |
|---|---|---|---|---|---|---|---|---|---|---|
| cluster3905 | 749 | 1476 | 205 | 0 | 134 | 1 | 76 | 152 | 198 | 205 |
| cluster3906 | 24 | 41 | 105 | 0 | 138 | 174 | 86 | 74 | 37 | 75 |
| cluster3907 | 125 | 244 | 990 | 0 | 328 | 839 | 613 | 650 | 765 | 1322 |
| cluster3908 | 44 | 146 | 138 | 1 | 280 | 324 | 333 | 90 | 96 | 219 |
| cluster3909 | 102 | 241 | 247 | 0 | 573 | 624 | 423 | 154 | 130 | 182 |
| cluster3910 | 57 | 224 | 385 | 0 | 67 | 90 | 50 | 109 | 70 | 363 |
| cluster3911 | 12 | 30 | 439 | 0 | 208 | 313 | 209 | 29 | 161 | 398 |
| cluster3912 | 249 | 537 | 565 | 177 | 587 | 986 | 721 | 443 | 500 | 776 |
| cluster3913 | 8 | 24 | 168 | 0 | 7 | 25 | 42 | 32 | 40 | 226 |
| cluster3914 | 375 | 1066 | 782 | 0 | 392 | 6 | 474 | 37 | 214 | 1134 |
| cluster3915 | 147 | 368 | 563 | 0 | 258 | 152 | 94 | 1008 | 64 | 684 |
| cluster3916 | 20 | 49 | 174 | 0 | 63 | 185 | 50 | 24 | 47 | 92 |
| cluster3917 | 8 | 31 | 363 | 0 | 407 | 501 | 331 | 82 | 170 | 341 |
| cluster3918 | 16 | 61 | 392 | 0 | 115 | 226 | 59 | 29 | 70 | 189 |
| cluster3919 | 5987 | 17487 | 31296 | 47 | 4 | 0 | 24265 | 15811 | 15858 | 21893 |
| cluster3920 | 11 | 25 | 228 | 0 | 42 | 49 | 490 | 44 | 90 | 124 |
| cluster3921 | 373 | 1119 | 3764 | 0 | 1416 | 3137 | 705 | 837 | 1586 | 4188 |
| cluster3922 | 689 | 2349 | 1228 | 0 | 1725 | 2404 | 1629 | 2138 | 745 | 1720 |
| cluster3923 | 23 | 41 | 99 | 0 | 38 | 57 | 78 | 39 | 60 | 111 |
| cluster3924 | 5 | 20 | 19 | 0 | 112 | 145 | 56 | 24 | 6 | 50 |
| cluster3925 | 3396 | 8754 | 5493 | 0 | 5 | 0 | 3271 | 5470 | 485 | 9667 |
| cluster3926 | 232 | 581 | 545 | 0 | 2 | 950 | 248 | 182 | 57 | 35 |
| cluster3927 | 32 | 77 | 243 | 0 | 83 | 225 | 109 | 40 | 50 | 86 |
| cluster3928 | 104 | 199 | 125 | 0 | 111 | 216 | 179 | 55 | 75 | 367 |
| cluster3929 | 4 | 29 | 75 | 0 | 116 | 194 | 134 | 108 | 145 | 260 |
| cluster3930 | 25 | 69 | 34 | 96 | 197 | 254 | 101 | 44 | 15 | 103 |
| cluster3931 | 14 | 20 | 1658 | 15 | 100 | 265 | 72 | 76 | 209 | 336 |
| cluster3932 | 18 | 34 | 68 | 60 | 14 | 18 | 1253 | 21 | 66 | 106 |
| cluster3933 | 45 | 83 | 165 | 0 | 33 | 57 | 208 | 97 | 138 | 542 |
| cluster3934 | 494 | 1412 | 2668 | 1 | 2022 | 3402 | 1557 | 392 | 1841 | 4303 |
| cluster3935 | 2 | 21 | 61 | 0 | 32 | 25 | 31 | 14 | 10 | 105 |

| | | | | | | | | | | |
|---|---|---|---|---|---|---|---|---|---|---|
| cluster3936 | 33 | 48 | 240 | 0 | 278 | 460 | 148 | 41 | 35 | 95 |
| cluster3937 | 7 | 20 | 14 | 88 | 54 | 139 | 61 | 15 | 34 | 85 |
| cluster3938 | 17 | 36 | 63 | 0 | 63 | 107 | 43 | 27 | 9 | 54 |
| cluster3939 | 23 | 66 | 122 | 0 | 176 | 166 | 133 | 31 | 74 | 121 |
| cluster3940 | 38 | 121 | 94 | 193 | 602 | 794 | 209 | 70 | 142 | 255 |
| cluster3941 | 67 | 153 | 414 | 79 | 103 | 193 | 260 | 161 | 520 | 657 |
| cluster3942 | 136 | 356 | 633 | 0 | 185 | 127 | 299 | 172 | 630 | 952 |
| cluster3943 | 13 | 67 | 347 | 0 | 224 | 323 | 143 | 44 | 149 | 208 |
| cluster3944 | 38 | 103 | 288 | 0 | 254 | 376 | 110 | 108 | 47 | 216 |
| cluster3945 | 105 | 272 | 1222 | 0 | 111 | 182 | 220 | 156 | 193 | 364 |
| cluster3946 | 119 | 296 | 732 | 0 | 324 | 323 | 322 | 141 | 451 | 829 |
| cluster3947 | 25 | 60 | 113 | 0 | 158 | 237 | 138 | 37 | 118 | 137 |
| cluster3948 | 344 | 1082 | 566 | 0 | 972 | 1353 | 1081 | 757 | 786 | 1237 |
| cluster3949 | 9 | 19 | 188 | 0 | 40 | 123 | 154 | 25 | 172 | 269 |
| cluster3950 | 100 | 267 | 239 | 449 | 843 | 1703 | 579 | 478 | 107 | 481 |
| cluster3951 | 14 | 33 | 169 | 0 | 118 | 219 | 137 | 37 | 51 | 96 |
| cluster3952 | 76 | 150 | 425 | 0 | 541 | 1441 | 167 | 117 | 196 | 365 |
| cluster3953 | 490 | 1204 | 1317 | 1 | 550 | 306 | 906 | 744 | 1214 | 2626 |
| cluster3954 | 8 | 23 | 62 | 0 | 93 | 155 | 282 | 28 | 30 | 155 |
| cluster3955 | 46 | 134 | 48 | 0 | 132 | 230 | 247 | 256 | 80 | 138 |
| cluster3956 | 41 | 86 | 180 | 0 | 367 | 506 | 82 | 65 | 40 | 128 |
| cluster3957 | 9 | 28 | 73 | 0 | 161 | 256 | 166 | 25 | 21 | 71 |
| cluster3958 | 4 | 20 | 5 | 0 | 36 | 8 | 178 | 10 | 20 | 42 |
| cluster3959 | 7 | 21 | 20 | 58 | 424 | 92 | 688 | 99 | 199 | 461 |
| cluster3960 | 12 | 28 | 486 | 8 | 120 | 114 | 858 | 5 | 342 | 173 |
| cluster3961 | 177 | 582 | 1427 | 0 | 679 | 1405 | 377 | 374 | 567 | 1129 |
| cluster3962 | 65 | 153 | 319 | 0 | 207 | 306 | 111 | 52 | 309 | 449 |
| cluster3963 | 258 | 630 | 481 | 0 | 225 | 438 | 154 | 294 | 150 | 680 |
| cluster3964 | 8 | 22 | 130 | 0 | 302 | 234 | 622 | 35 | 301 | 426 |
| cluster3965 | 5 | 16 | 92 | 0 | 17 | 9 | 32 | 11 | 9 | 16 |
| cluster3966 | 44 | 111 | 295 | 0 | 323 | 449 | 239 | 90 | 147 | 259 |

| | | | | | | | | | | |
|---|---|---|---|---|---|---|---|---|---|---|
| cluster3967 | 12 | 21 | 26 | 89 | 39 | 56 | 109 | 10 | 77 | 99 |
| cluster3968 | 14 | 28 | 93 | 1 | 130 | 288 | 43 | 52 | 37 | 109 |
| cluster3969 | 222 | 620 | 1536 | 149 | 445 | 764 | 936 | 406 | 814 | 1314 |
| cluster3970 | 4 | 24 | 96 | 0 | 240 | 384 | 86 | 33 | 56 | 97 |
| cluster3971 | 93 | 247 | 197 | 0 | 480 | 738 | 126 | 63 | 38 | 112 |
| cluster3972 | 27 | 98 | 139 | 0 | 291 | 270 | 413 | 94 | 35 | 174 |
| cluster3973 | 5742 | 21307 | 30608 | 0 | 12794 | 17660 | 21583 | 21397 | 24834 | 40680 |
| cluster3974 | 5 | 18 | 92 | 0 | 454 | 651 | 415 | 36 | 81 | 233 |
| cluster3975 | 111 | 340 | 191 | 0 | 969 | 1288 | 536 | 145 | 36 | 101 |
| cluster3976 | 958 | 2258 | 621 | 1760 | 936 | 1654 | 1609 | 1598 | 1078 | 1806 |
| cluster3977 | 716 | 2266 | 836 | 0 | 709 | 441 | 877 | 381 | 725 | 3908 |
| cluster3978 | 91 | 197 | 327 | 2 | 125 | 234 | 303 | 228 | 61 | 238 |
| cluster3979 | 51 | 100 | 90 | 269 | 397 | 471 | 231 | 105 | 124 | 490 |
| cluster3980 | 48 | 130 | 217 | 88 | 352 | 422 | 241 | 89 | 103 | 441 |
| cluster3981 | 14 | 72 | 75 | 39 | 789 | 964 | 162 | 54 | 127 | 235 |
| cluster3982 | 37 | 125 | 200 | 0 | 239 | 297 | 247 | 83 | 120 | 209 |
| cluster3983 | 3 | 32 | 53 | 0 | 246 | 249 | 80 | 21 | 16 | 42 |
| cluster3984 | 73 | 224 | 211 | 0 | 321 | 2 | 176 | 187 | 363 | 731 |
| cluster3985 | 10 | 21 | 20 | 49 | 107 | 193 | 51 | 24 | 33 | 82 |
| cluster3986 | 24 | 56 | 144 | 0 | 102 | 156 | 55 | 54 | 73 | 126 |
| cluster3987 | 30 | 41 | 41 | 157 | 263 | 423 | 62 | 45 | 80 | 173 |
| cluster3988 | 46 | 131 | 485 | 8 | 244 | 287 | 366 | 161 | 268 | 454 |
| cluster3989 | 4 | 14 | 29 | 56 | 46 | 58 | 69 | 6 | 89 | 191 |
| cluster3990 | 9 | 21 | 75 | 49 | 70 | 85 | 39 | 16 | 73 | 139 |
| cluster3991 | 23 | 56 | 40 | 0 | 81 | 103 | 35 | 33 | 14 | 69 |
| cluster3992 | 15 | 31 | 138 | 268 | 158 | 289 | 265 | 80 | 140 | 243 |
| cluster3993 | 67 | 148 | 133 | 0 | 120 | 221 | 162 | 67 | 79 | 159 |
| cluster3994 | 230 | 526 | 138 | 0 | 64 | 70 | 157 | 374 | 291 | 314 |
| cluster3995 | 29 | 97 | 107 | 0 | 35 | 13 | 58 | 107 | 112 | 209 |
| cluster3996 | 18 | 80 | 146 | 0 | 90 | 160 | 65 | 36 | 111 | 168 |
| cluster3997 | 39 | 140 | 275 | 0 | 194 | 392 | 68 | 62 | 112 | 212 |

| | | | | | | | | | | |
|---|---|---|---|---|---|---|---|---|---|---|
| cluster3998 | 29 | 62 | 16 | 0 | 25 | 233 | 47 | 0 | 6 | 79 |
| cluster3999 | 7 | 23 | 82 | 0 | 38 | 50 | 124 | 31 | 34 | 69 |
| cluster4000 | 13 | 27 | 77 | 6 | 77 | 117 | 15 | 6 | 23 | 70 |
| cluster4001 | 240 | 686 | 803 | 0 | 742 | 600 | 973 | 2031 | 280 | 822 |
| cluster4002 | 7 | 34 | 40 | 0 | 193 | 295 | 61 | 11 | 22 | 30 |
| cluster4003 | 109 | 232 | 597 | 0 | 203 | 363 | 50 | 107 | 630 | 643 |
| cluster4004 | 364 | 919 | 2899 | 0 | 902 | 1775 | 894 | 857 | 1763 | 2274 |
| cluster4005 | 26 | 67 | 144 | 0 | 68 | 167 | 53 | 110 | 93 | 295 |
| cluster4006 | 176 | 447 | 1069 | 0 | 415 | 459 | 214 | 325 | 971 | 846 |
| cluster4007 | 27 | 104 | 245 | 1 | 96 | 199 | 96 | 54 | 81 | 196 |
| cluster4008 | 16 | 31 | 45 | 0 | 92 | 133 | 32 | 19 | 34 | 55 |
| cluster4009 | 1 | 11 | 12 | 0 | 21 | 37 | 18 | 4 | 42 | 41 |
| cluster4010 | 9 | 43 | 51 | 0 | 121 | 148 | 96 | 15 | 9 | 12 |
| cluster4011 | 2203 | 6503 | 5358 | 0 | 16 | 0 | 4104 | 5053 | 3659 | 7890 |
| cluster4012 | 5 | 15 | 146 | 0 | 42 | 29 | 12 | 4 | 48 | 62 |
| cluster4013 | 6 | 8 | 9 | 0 | 12 | 5 | 496 | 0 | 25 | 15 |
| cluster4014 | 267 | 878 | 978 | 0 | 1780 | 2570 | 479 | 808 | 310 | 1306 |
| cluster4015 | 4739 | 14149 | 26342 | 2074 | 27 | 122 | 9297 | 13165 | 5109 | 21909 |
| cluster4016 | 6 | 46 | 39 | 55 | 298 | 508 | 69 | 48 | 43 | 71 |
| cluster4017 | 12 | 35 | 113 | 0 | 47 | 102 | 56 | 16 | 18 | 82 |
| cluster4018 | 3 | 9 | 45 | 1 | 24 | 63 | 31 | 1 | 46 | 31 |
| cluster4019 | 489 | 1314 | 3728 | 0 | 369 | 5764 | 110 | 164 | 1034 | 2800 |
| cluster4020 | 90 | 151 | 94 | 209 | 181 | 291 | 179 | 130 | 247 | 373 |
| cluster4021 | 4 | 19 | 107 | 0 | 42 | 95 | 32 | 25 | 38 | 76 |
| cluster4022 | 3685 | 11280 | 15739 | 0 | 10029 | 16850 | 9094 | 10712 | 2313 | 12400 |
| cluster4023 | 7 | 10 | 71 | 0 | 17 | 16 | 70 | 6 | 79 | 50 |
| cluster4024 | 6 | 17 | 52 | 0 | 109 | 123 | 96 | 19 | 25 | 66 |
| cluster4025 | 97 | 215 | 425 | 19 | 347 | 610 | 135 | 238 | 160 | 738 |
| cluster4026 | 45 | 101 | 167 | 0 | 59 | 113 | 54 | 90 | 57 | 104 |
| cluster4027 | 40 | 88 | 124 | 0 | 224 | 260 | 168 | 30 | 39 | 109 |
| cluster4028 | 11 | 35 | 220 | 0 | 129 | 233 | 124 | 31 | 87 | 234 |

| | | | | | | | | | | |
|---|---|---|---|---|---|---|---|---|---|---|
| cluster4029 | 123 | 356 | 1902 | 0 | 35 | 116 | 262 | 326 | 227 | 359 |
| cluster4030 | 434 | 1307 | 807 | 0 | 1801 | 2490 | 886 | 994 | 245 | 1426 |
| cluster4031 | 49 | 152 | 161 | 550 | 118 | 190 | 430 | 166 | 46 | 276 |
| cluster4032 | 4 | 16 | 79 | 2 | 21 | 48 | 57 | 3 | 65 | 133 |
| cluster4033 | 4 | 12 | 27 | 0 | 0 | 19 | 36 | 22 | 5 | 12 |
| cluster4034 | 4108 | 10339 | 0 | 5 | 6 | 0 | 7432 | 7836 | 4797 | 14554 |
| cluster4035 | 11 | 26 | 43 | 91 | 66 | 74 | 54 | 11 | 33 | 80 |
| cluster4036 | 16 | 47 | 82 | 0 | 115 | 161 | 93 | 39 | 70 | 111 |
| cluster4037 | 37 | 105 | 148 | 0 | 253 | 480 | 104 | 81 | 116 | 206 |
| cluster4038 | 38 | 108 | 110 | 0 | 232 | 565 | 161 | 91 | 52 | 161 |
| cluster4039 | 10 | 23 | 337 | 31 | 219 | 127 | 290 | 13 | 292 | 419 |
| cluster4040 | 13 | 45 | 39 | 1 | 45 | 59 | 24 | 21 | 8 | 20 |
| cluster4041 | 1460 | 3714 | 7820 | 0 | 3661 | 3645 | 603 | 2617 | 2616 | 6435 |
| cluster4042 | 488 | 1504 | 1690 | 0 | 574 | 1131 | 987 | 898 | 232 | 805 |
| cluster4043 | 3 | 13 | 3 | 0 | 8 | 8 | 11 | 3 | 8 | 24 |
| cluster4044 | 34 | 79 | 116 | 0 | 217 | 354 | 68 | 45 | 59 | 117 |
| cluster4045 | 7 | 21 | 87 | 0 | 420 | 545 | 333 | 33 | 46 | 87 |
| cluster4046 | 77 | 121 | 68 | 1 | 5 | 5 | 1278 | 85 | 85 | 296 |
| cluster4047 | 5 | 11 | 54 | 0 | 340 | 405 | 119 | 21 | 76 | 176 |
| cluster4048 | 8 | 19 | 127 | 8 | 326 | 124 | 1902 | 21 | 69 | 127 |
| cluster4049 | 4938 | 11778 | 17197 | 0 | 9 | 0 | 15864 | 17855 | 27447 | 25574 |
| cluster4050 | 117 | 357 | 314 | 0 | 441 | 848 | 235 | 234 | 172 | 431 |
| cluster4051 | 2 | 16 | 65 | 0 | 39 | 161 | 75 | 0 | 16 | 24 |
| cluster4052 | 12 | 25 | 99 | 195 | 204 | 181 | 61 | 15 | 56 | 109 |
| cluster4053 | 17 | 37 | 89 | 0 | 65 | 116 | 158 | 109 | 21 | 122 |
| cluster4054 | 5 | 22 | 60 | 0 | 102 | 197 | 59 | 14 | 25 | 38 |
| cluster4055 | 53 | 210 | 94 | 0 | 98 | 170 | 306 | 234 | 38 | 138 |
| cluster4056 | 149 | 331 | 42 | 0 | 29 | 183 | 167 | 125 | 216 | 402 |
| cluster4057 | 32 | 84 | 151 | 0 | 280 | 492 | 289 | 78 | 43 | 88 |
| cluster4058 | 112 | 223 | 97 | 0 | 295 | 427 | 289 | 74 | 65 | 135 |
| cluster4059 | 798 | 1658 | 986 | 0 | 1905 | 3489 | 1713 | 1705 | 1514 | 1892 |

| | | | | | | | | | | |
|---|---|---|---|---|---|---|---|---|---|---|
| cluster4060 | 3 | 14 | 97 | 0 | 120 | 178 | 13 | 7 | 20 | 42 |
| cluster4061 | 18 | 46 | 173 | 0 | 77 | 131 | 58 | 21 | 38 | 70 |
| cluster4062 | 2 | 10 | 145 | 0 | 6 | 13 | 178 | 1 | 109 | 133 |
| cluster4063 | 44 | 120 | 541 | 0 | 7 | 535 | 134 | 248 | 153 | 306 |
| cluster4064 | 14 | 35 | 129 | 0 | 157 | 251 | 68 | 11 | 72 | 86 |
| cluster4065 | 17 | 73 | 167 | 0 | 184 | 296 | 144 | 61 | 147 | 193 |
| cluster4066 | 21 | 49 | 37 | 0 | 25 | 97 | 99 | 40 | 26 | 55 |
| cluster4067 | 2 | 28 | 289 | 0 | 143 | 3662 | 107 | 34 | 163 | 222 |
| cluster4068 | 7 | 25 | 34 | 0 | 82 | 184 | 109 | 28 | 23 | 41 |
| cluster4069 | 473 | 1122 | 314 | 38 | 1580 | 3707 | 320 | 813 | 396 | 1167 |
| cluster4070 | 1 | 12 | 65 | 1 | 0 | 1 | 144 | 6 | 19 | 42 |
| cluster4071 | 16 | 63 | 92 | 0 | 343 | 512 | 467 | 44 | 25 | 223 |
| cluster4072 | 143 | 378 | 1309 | 0 | 820 | 1850 | 288 | 430 | 828 | 995 |
| cluster4073 | 11 | 17 | 211 | 0 | 553 | 879 | 65 | 20 | 103 | 190 |
| cluster4074 | 6 | 22 | 35 | 0 | 114 | 173 | 21 | 2 | 1 | 7 |
| cluster4075 | 6 | 18 | 7 | 10 | 6 | 17 | 1 | 0 | 31 | 24 |
| cluster4076 | 7 | 24 | 118 | 510 | 101 | 193 | 94 | 14 | 91 | 111 |
| cluster4077 | 9 | 17 | 132 | 0 | 161 | 174 | 255 | 24 | 182 | 112 |
| cluster4078 | 150 | 437 | 313 | 1391 | 660 | 982 | 197 | 158 | 76 | 360 |
| cluster4079 | 22 | 43 | 70 | 162 | 526 | 479 | 78 | 45 | 66 | 146 |
| cluster4080 | 1548 | 3411 | 246 | 731 | 12 | 0 | 2154 | 2435 | 987 | 2063 |
| cluster4081 | 40 | 107 | 201 | 0 | 294 | 610 | 198 | 74 | 43 | 80 |
| cluster4082 | 3 | 13 | 35 | 0 | 28 | 44 | 29 | 0 | 16 | 19 |
| cluster4083 | 44 | 131 | 90 | 0 | 222 | 324 | 57 | 51 | 87 | 107 |
| cluster4084 | 3407 | 7778 | 2889 | 8722 | 3611 | 4926 | 6460 | 7635 | 4573 | 12199 |
| cluster4085 | 3885 | 14797 | 16412 | 779 | 7 | 1 | 13118 | 14667 | 15484 | 21066 |
| cluster4086 | 4 | 27 | 111 | 0 | 41 | 48 | 15 | 13 | 6 | 26 |
| cluster4087 | 16 | 58 | 89 | 0 | 2186 | 3823 | 123 | 81 | 51 | 87 |
| cluster4088 | 4 | 9 | 31 | 0 | 70 | 169 | 158 | 28 | 120 | 68 |
| cluster4089 | 275 | 721 | 56786 | 2 | 2818 | 2730 | 2016 | 5586 | 4492 | 10398 |
| cluster4090 | 2 | 15 | 82 | 0 | 2 | 1 | 71 | 76 | 79 | 212 |

| | | | | | | | | | | |
|---|---|---|---|---|---|---|---|---|---|---|
| cluster4091 | 1 | 7 | 97 | 1 | 3 | 4 | 95 | 0 | 440 | 317 |
| cluster4092 | 1 | 11 | 101 | 0 | 49 | 45 | 70 | 26 | 34 | 147 |
| cluster4093 | 18 | 52 | 208 | 0 | 101 | 1145 | 132 | 30 | 168 | 188 |
| cluster4094 | 3 | 6 | 35 | 2 | 20 | 35 | 18 | 0 | 23 | 27 |
| cluster4095 | 2 | 10 | 9 | 55 | 78 | 77 | 35 | 5 | 15 | 19 |
| cluster4096 | 77 | 207 | 238 | 0 | 272 | 496 | 90 | 177 | 135 | 437 |
| cluster4097 | 4130 | 9106 | 17672 | 0 | 10 | 0 | 8818 | 7180 | 8272 | 11481 |
| cluster4098 | 7 | 10 | 71 | 205 | 174 | 287 | 59 | 18 | 22 | 58 |
| cluster4099 | 19 | 79 | 131 | 0 | 404 | 455 | 93 | 35 | 63 | 185 |
| cluster4100 | 9 | 22 | 42 | 0 | 111 | 131 | 45 | 23 | 16 | 61 |
| cluster4101 | 4 | 6 | 165 | 417 | 10 | 492 | 17 | 13 | 5 | 13 |
| cluster4102 | 7 | 7 | 63 | 0 | 27 | 23 | 16 | 13 | 28 | 63 |
| cluster4103 | 2 | 8 | 90 | 171 | 82 | 175 | 87 | 24 | 162 | 240 |
| cluster4104 | 34 | 107 | 91 | 0 | 219 | 407 | 163 | 63 | 27 | 67 |
| cluster4105 | 65 | 144 | 160 | 0 | 85 | 182 | 8 | 19 | 62 | 140 |
| cluster4106 | 0 | 5 | 95 | 0 | 58 | 17 | 142 | 16 | 18 | 47 |
| cluster4107 | 7 | 15 | 17 | 0 | 112 | 123 | 38 | 3 | 16 | 16 |
| cluster4108 | 2 | 9 | 2 | 35 | 35 | 18 | 24 | 31 | 3 | 10 |
| cluster4109 | 4 | 6 | 104 | 0 | 95 | 157 | 77 | 8 | 28 | 60 |
| cluster4110 | 7 | 35 | 97 | 0 | 80 | 147 | 36 | 35 | 43 | 97 |
| cluster4111 | 3 | 11 | 2 | 0 | 0 | 8 | 83 | 6 | 28 | 118 |
| cluster4112 | 2 | 4 | 12 | 46 | 38 | 85 | 5 | 2 | 4 | 32 |
| cluster4113 | 5562 | 10929 | 16152 | 416 | 4342 | 4822 | 8480 | 9490 | 4331 | 12433 |
| cluster4114 | 2 | 5 | 51 | 0 | 838 | 534 | 118 | 3 | 218 | 354 |
| cluster4115 | 5 | 6 | 120 | 0 | 19 | 39 | 199 | 9 | 48 | 42 |
| cluster4116 | 1 | 5 | 8 | 40 | 33 | 14 | 634 | 21 | 0 | 9 |
| cluster4117 | 3 | 11 | 3 | 0 | 131 | 26 | 142 | 0 | 12 | 332 |
| cluster4118 | 0 | 7 | 23 | 0 | 23 | 88 | 591 | 99 | 468 | 497 |
| cluster4119 | 23 | 56 | 71 | 0 | 853 | 684 | 89 | 17 | 22 | 76 |
| cluster4120 | 1 | 4 | 31 | 1 | 5 | 0 | 89 | 2 | 77 | 22 |
| cluster4121 | 4 | 7 | 33 | 0 | 59 | 133 | 11 | 12 | 6 | 29 |

| | | | | | | | | | | |
|---|---|---|---|---|---|---|---|---|---|---|
| cluster4122 | 2 | 10 | 28 | 0 | 1 | 9 | 181 | 2 | 10 | 35 |
| cluster4123 | 5 | 8 | 19 | 0 | 5 | 5 | 2 | 3 | 6 | 1 |
| cluster4124 | 1 | 3 | 10 | 0 | 5 | 2 | 10 | 5 | 6 | 16 |
| cluster4125 | 2 | 3 | 68 | 79 | 13 | 0 | 76 | 2 | 9 | 31 |
| cluster4126 | 1 | 3 | 98 | 0 | 8 | 103 | 24 | 28 | 12 | 39 |
| cluster4127 | 1 | 2 | 157 | 0 | 11 | 72 | 262 | 9 | 2 | 5 |
| cluster4128 | 2 | 1 | 14 | 0 | 64 | 137 | 149 | 31 | 15 | 63 |

[1]Chom: Replicates of *C. homini*

[2]Dhom: Replicates of *D. homin*

[3]Oovi: Replicates of *O. ovis*

[4]Lcup: Replicates of *L. cuprina*

[5]Cmeg: Replicates of *C. megac*

**Supplementary Table 3.** DESeq2 results with eXpress counts - Parasites vs Saprophagous

| ID | log2FoldChange | Adjusted p value |
|---|---|---|
| cluster3975 | 3.52 | 0.003 |
| cluster3616 | 4.81 | 0.012 |
| cluster3374 | 4.16 | 0.052 |
| cluster3250 | -2.58 | 0.088 |
| cluster3547 | 2.74 | 0.088 |
| cluster3457 | -3.43 | 0.106 |
| cluster3822 | 4.18 | 0.106 |
| cluster3891 | 3.27 | 0.106 |
| cluster3639 | 4.58 | 0.117 |
| cluster4081 | 2.26 | 0.120 |
| cluster3379 | 2.85 | 0.137 |
| cluster3689 | 4.39 | 0.137 |
| cluster3760 | 2.89 | 0.137 |
| cluster3858 | 3.28 | 0.137 |
| cluster3732 | 2.28 | 0.143 |
| cluster3780 | 4.37 | 0.200 |
| cluster3802 | 2.71 | 0.200 |
| cluster4078 | 1.99 | 0.200 |
| cluster4101 | 3.73 | 0.200 |
| cluster4127 | 4.39 | 0.200 |
| cluster4091 | -3.58 | 0.209 |
| cluster3956 | 2.23 | 0.243 |
| cluster4040 | 2.27 | 0.243 |
| cluster4074 | 3.69 | 0.243 |
| cluster3099 | -1.91 | 0.248 |
| cluster3301 | 2.56 | 0.248 |
| cluster3419 | -1.87 | 0.248 |
| cluster3715 | 2.85 | 0.248 |
| cluster3810 | 2.20 | 0.248 |
| cluster3859 | -2.24 | 0.248 |
| cluster3885 | -2.23 | 0.248 |
| cluster3926 | 3.65 | 0.248 |
| cluster4010 | 2.80 | 0.248 |
| cluster4116 | 3.58 | 0.248 |
| cluster3652 | 2.76 | 0.251 |
| cluster3971 | 2.59 | 0.251 |
| cluster3930 | 2.11 | 0.258 |
| cluster3461 | 3.39 | 0.262 |
| cluster3950 | 2.17 | 0.262 |
| cluster3713 | 1.71 | 0.263 |
| cluster3487 | 2.34 | 0.265 |
| cluster3546 | 3.14 | 0.265 |

| | | |
|---|---|---|
| cluster3567 | -2.92 | 0.265 |
| cluster4058 | 2.00 | 0.265 |
| cluster3704 | 2.43 | 0.266 |
| cluster3889 | 3.14 | 0.266 |
| cluster3030 | -1.66 | 0.266 |
| cluster3100 | -2.53 | 0.266 |
| cluster3111 | -2.27 | 0.266 |
| cluster3843 | 1.55 | 0.266 |
| cluster3194 | -2.16 | 0.271 |
| cluster3235 | 2.99 | 0.271 |
| cluster3978 | 1.75 | 0.271 |
| cluster2994 | 1.77 | 0.273 |
| cluster3545 | 1.90 | 0.273 |
| cluster4108 | 2.90 | 0.273 |
| cluster2961 | 2.29 | 0.285 |
| cluster3231 | -2.36 | 0.309 |
| cluster3746 | 2.19 | 0.309 |
| cluster4055 | 2.08 | 0.309 |
| cluster3026 | -2.26 | 0.310 |
| cluster3778 | 2.33 | 0.319 |
| cluster3103 | 2.62 | 0.328 |
| cluster3256 | 1.52 | 0.328 |
| cluster3901 | 2.65 | 0.328 |
| cluster4120 | -2.54 | 0.347 |
| cluster3972 | 1.72 | 0.348 |
| cluster3821 | -1.94 | 0.355 |
| cluster3770 | 3.70 | 0.359 |
| cluster2905 | -1.21 | 0.372 |
| cluster2902 | 2.46 | 0.374 |
| cluster3504 | 2.61 | 0.374 |
| cluster3779 | 2.02 | 0.374 |
| cluster3854 | 1.62 | 0.374 |
| cluster4098 | 2.60 | 0.374 |
| cluster2906 | -1.37 | 0.379 |
| cluster3529 | -2.06 | 0.379 |
| cluster3852 | 2.43 | 0.379 |
| cluster3899 | 1.83 | 0.379 |
| cluster3670 | 2.33 | 0.400 |
| cluster3827 | 2.46 | 0.400 |
| cluster3892 | 2.38 | 0.400 |
| cluster3240 | 2.06 | 0.403 |
| cluster3356 | 1.87 | 0.403 |
| cluster3909 | 1.70 | 0.404 |
| cluster4119 | 2.96 | 0.404 |

| | | |
|---|---|---|
| cluster3871 | 1.69 | 0.421 |
| cluster4031 | 2.20 | 0.421 |
| cluster4002 | 2.17 | 0.429 |
| cluster3556 | 2.02 | 0.442 |
| cluster3008 | 1.72 | 0.453 |
| cluster3160 | 1.61 | 0.453 |
| cluster3782 | -1.56 | 0.453 |
| cluster4022 | 1.54 | 0.454 |
| cluster3297 | 2.27 | 0.463 |
| cluster3851 | 2.04 | 0.463 |
| cluster3476 | 2.18 | 0.466 |
| cluster3182 | 1.90 | 0.473 |
| cluster3175 | 2.18 | 0.473 |
| cluster4057 | 2.19 | 0.473 |
| cluster3004 | -1.71 | 0.477 |
| cluster3128 | -1.39 | 0.477 |
| cluster3171 | -1.43 | 0.477 |
| cluster3176 | -1.39 | 0.477 |
| cluster3212 | 2.00 | 0.477 |
| cluster3315 | 1.74 | 0.477 |
| cluster3329 | 2.39 | 0.477 |
| cluster3619 | 1.66 | 0.477 |
| cluster3707 | 2.14 | 0.477 |
| cluster3936 | 1.93 | 0.477 |
| cluster4016 | 1.95 | 0.477 |
| cluster3077 | 1.85 | 0.488 |
| cluster3805 | 1.68 | 0.488 |
| cluster3853 | 2.02 | 0.488 |
| cluster3957 | 1.75 | 0.488 |
| cluster3983 | 2.14 | 0.488 |
| cluster3728 | 1.54 | 0.493 |
| cluster3787 | 2.12 | 0.493 |
| cluster3243 | -1.12 | 0.529 |
| cluster3179 | 2.01 | 0.534 |
| cluster3177 | 2.06 | 0.534 |
| cluster4104 | 2.04 | 0.534 |
| cluster3019 | 1.40 | 0.537 |
| cluster2885 | -1.63 | 0.548 |
| cluster2893 | 2.08 | 0.548 |
| cluster2925 | -1.71 | 0.548 |
| cluster3108 | -1.61 | 0.548 |
| cluster3219 | 0.97 | 0.548 |
| cluster3222 | 1.06 | 0.548 |
| cluster3371 | -2.11 | 0.548 |

| | | |
|---|---|---|
| cluster3386 | 1.80 | 0.548 |
| cluster3406 | 1.49 | 0.548 |
| cluster3427 | -1.87 | 0.548 |
| cluster3428 | -1.50 | 0.548 |
| cluster3451 | -2.53 | 0.548 |
| cluster3474 | 2.01 | 0.548 |
| cluster3501 | 2.06 | 0.548 |
| cluster3506 | 1.17 | 0.548 |
| cluster3516 | 2.17 | 0.548 |
| cluster3611 | 2.11 | 0.548 |
| cluster3723 | 1.20 | 0.548 |
| cluster3855 | 1.79 | 0.548 |
| cluster3865 | 2.25 | 0.548 |
| cluster3866 | 2.36 | 0.548 |
| cluster4079 | 1.78 | 0.548 |
| cluster4095 | 2.09 | 0.548 |
| cluster4113 | 1.45 | 0.548 |
| cluster4118 | -2.56 | 0.548 |
| cluster4121 | 1.83 | 0.548 |
| cluster4125 | 2.88 | 0.548 |
| cluster3478 | -1.05 | 0.552 |
| cluster3906 | 1.42 | 0.557 |
| cluster3032 | -0.97 | 0.558 |
| cluster3191 | 2.36 | 0.558 |
| cluster3413 | 1.79 | 0.558 |
| cluster3571 | 2.00 | 0.558 |
| cluster4042 | 1.86 | 0.558 |
| cluster3607 | -1.64 | 0.562 |
| cluster3118 | -1.40 | 0.569 |
| cluster4030 | 1.68 | 0.569 |
| cluster2946 | -1.01 | 0.570 |
| cluster3319 | -1.62 | 0.573 |
| cluster3818 | 1.31 | 0.574 |
| cluster3195 | -1.35 | 0.593 |
| cluster3123 | -1.23 | 0.616 |
| cluster3146 | -1.32 | 0.616 |
| cluster3166 | 1.33 | 0.616 |
| cluster3333 | 1.31 | 0.616 |
| cluster3359 | 1.64 | 0.616 |
| cluster3364 | 1.29 | 0.616 |
| cluster3678 | 1.24 | 0.616 |
| cluster3904 | 1.68 | 0.616 |
| cluster3912 | 0.88 | 0.616 |
| cluster3927 | 1.25 | 0.616 |

| | | |
|---|---|---|
| cluster3693 | 2.00 | 0.616 |
| cluster3924 | 1.71 | 0.621 |
| cluster3748 | 1.50 | 0.624 |
| cluster2948 | -1.11 | 0.625 |
| cluster2926 | 1.50 | 0.627 |
| cluster3126 | -0.92 | 0.627 |
| cluster3420 | -1.32 | 0.627 |
| cluster3482 | 1.50 | 0.627 |
| cluster3717 | 1.28 | 0.627 |
| cluster3800 | 1.63 | 0.627 |
| cluster3877 | 1.25 | 0.627 |
| cluster3976 | 1.40 | 0.627 |
| cluster2912 | 1.27 | 0.627 |
| cluster3041 | 1.14 | 0.627 |
| cluster3151 | -1.10 | 0.627 |
| cluster3342 | 1.54 | 0.627 |
| cluster3448 | 1.91 | 0.627 |
| cluster4062 | -1.87 | 0.634 |
| cluster4069 | 1.52 | 0.634 |
| cluster3102 | -0.90 | 0.636 |
| cluster3758 | 1.50 | 0.636 |
| cluster3804 | 1.41 | 0.636 |
| cluster3864 | 1.05 | 0.636 |
| cluster3757 | -1.54 | 0.639 |
| cluster4086 | 1.67 | 0.640 |
| cluster3101 | 0.93 | 0.650 |
| cluster3754 | 1.14 | 0.650 |
| cluster2924 | -1.45 | 0.657 |
| cluster2934 | -1.02 | 0.657 |
| cluster3071 | -0.88 | 0.657 |
| cluster3109 | 1.43 | 0.657 |
| cluster3517 | 0.85 | 0.657 |
| cluster3681 | 1.50 | 0.657 |
| cluster3207 | 1.43 | 0.659 |
| cluster3688 | 1.23 | 0.663 |
| cluster3993 | 1.01 | 0.669 |
| cluster3037 | -1.24 | 0.670 |
| cluster3046 | -1.21 | 0.674 |
| cluster3078 | 0.87 | 0.674 |
| cluster3186 | 1.56 | 0.676 |
| cluster3260 | 1.28 | 0.676 |
| cluster3283 | -1.04 | 0.676 |
| cluster3309 | -1.58 | 0.676 |
| cluster3793 | 1.12 | 0.676 |

| | | |
|---|---|---|
| cluster3944 | 1.21 | 0.677 |
| cluster2858 | -1.15 | 0.679 |
| cluster2928 | 1.24 | 0.679 |
| cluster2995 | 1.26 | 0.679 |
| cluster3024 | -1.46 | 0.679 |
| cluster3063 | 1.48 | 0.679 |
| cluster3089 | 1.25 | 0.679 |
| cluster3117 | 1.69 | 0.679 |
| cluster3156 | 1.27 | 0.679 |
| cluster3178 | 1.73 | 0.679 |
| cluster3192 | 1.23 | 0.679 |
| cluster3306 | 1.78 | 0.679 |
| cluster3378 | 1.33 | 0.679 |
| cluster3387 | 1.16 | 0.679 |
| cluster3443 | 1.39 | 0.679 |
| cluster3454 | 1.18 | 0.679 |
| cluster3467 | 1.06 | 0.679 |
| cluster3500 | 1.49 | 0.679 |
| cluster3559 | -0.97 | 0.679 |
| cluster3604 | 1.60 | 0.679 |
| cluster3618 | -1.87 | 0.679 |
| cluster3743 | 0.91 | 0.679 |
| cluster3824 | 1.69 | 0.679 |
| cluster3842 | -1.42 | 0.679 |
| cluster3849 | 1.12 | 0.679 |
| cluster3886 | 1.29 | 0.679 |
| cluster3890 | 1.16 | 0.679 |
| cluster3897 | 1.53 | 0.679 |
| cluster3902 | 1.19 | 0.679 |
| cluster3916 | 1.00 | 0.679 |
| cluster3938 | 1.47 | 0.679 |
| cluster3952 | 1.21 | 0.679 |
| cluster3966 | 0.89 | 0.679 |
| cluster4068 | 1.61 | 0.679 |
| cluster4107 | 1.81 | 0.679 |
| cluster3417 | 0.93 | 0.688 |
| cluster3005 | -0.97 | 0.695 |
| cluster3718 | 1.38 | 0.695 |
| cluster3940 | 1.26 | 0.695 |
| cluster4032 | -1.14 | 0.700 |
| cluster3543 | 1.57 | 0.704 |
| cluster3257 | 0.96 | 0.712 |
| cluster3274 | 1.32 | 0.716 |
| cluster3362 | 1.19 | 0.716 |

| | | |
|---|---|---|
| cluster3502 | 1.60 | 0.716 |
| cluster3629 | 1.39 | 0.716 |
| cluster3985 | 1.17 | 0.716 |
| cluster4045 | 1.70 | 0.716 |
| cluster3167 | -1.53 | 0.718 |
| cluster3299 | 1.33 | 0.718 |
| cluster3349 | 1.32 | 0.718 |
| cluster3645 | 2.29 | 0.718 |
| cluster3838 | -1.20 | 0.718 |
| cluster2989 | -0.96 | 0.718 |
| cluster3098 | 0.77 | 0.718 |
| cluster3323 | 1.50 | 0.718 |
| cluster3385 | 1.49 | 0.718 |
| cluster3620 | 1.19 | 0.718 |
| cluster3703 | 1.02 | 0.718 |
| cluster3965 | 1.45 | 0.718 |
| cluster3987 | 1.29 | 0.718 |
| cluster4038 | 1.49 | 0.718 |
| cluster2935 | 1.19 | 0.720 |
| cluster3554 | -1.05 | 0.721 |
| cluster3106 | -0.96 | 0.724 |
| cluster3752 | 1.78 | 0.724 |
| cluster3908 | 0.91 | 0.724 |
| cluster2863 | 0.88 | 0.725 |
| cluster3185 | 1.40 | 0.725 |
| cluster3411 | 0.85 | 0.725 |
| cluster3657 | 1.02 | 0.725 |
| cluster3668 | 0.79 | 0.725 |
| cluster3785 | -1.25 | 0.725 |
| cluster3830 | 1.00 | 0.725 |
| cluster3834 | 1.45 | 0.725 |
| cluster3840 | 1.42 | 0.725 |
| cluster4051 | 1.41 | 0.725 |
| cluster4054 | 1.41 | 0.725 |
| cluster3710 | 1.13 | 0.727 |
| cluster4044 | 1.12 | 0.729 |
| cluster3651 | 1.84 | 0.739 |
| cluster3246 | 1.16 | 0.739 |
| cluster3701 | 0.97 | 0.741 |
| cluster3647 | 0.88 | 0.749 |
| cluster3654 | 0.96 | 0.749 |
| cluster2876 | 1.63 | 0.751 |
| cluster2894 | -1.42 | 0.751 |
| cluster2904 | -1.33 | 0.751 |

| | | |
|---|---|---|
| cluster3060 | 1.35 | 0.751 |
| cluster3075 | 1.12 | 0.751 |
| cluster3112 | -1.40 | 0.751 |
| cluster3211 | 1.36 | 0.751 |
| cluster3236 | 1.12 | 0.751 |
| cluster3252 | 1.72 | 0.751 |
| cluster3296 | 1.23 | 0.751 |
| cluster3328 | -0.73 | 0.751 |
| cluster3439 | -0.93 | 0.751 |
| cluster3512 | 1.19 | 0.751 |
| cluster3666 | 1.49 | 0.751 |
| cluster3686 | 1.33 | 0.751 |
| cluster3709 | -1.08 | 0.751 |
| cluster3749 | 1.51 | 0.751 |
| cluster3794 | 1.16 | 0.751 |
| cluster3831 | 1.10 | 0.751 |
| cluster3884 | 1.36 | 0.751 |
| cluster3949 | -1.10 | 0.751 |
| cluster4001 | 0.91 | 0.751 |
| cluster4014 | 1.25 | 0.751 |
| cluster4024 | 1.06 | 0.751 |
| cluster4052 | 1.38 | 0.751 |
| cluster4060 | 1.35 | 0.751 |
| cluster4066 | 1.17 | 0.751 |
| cluster4096 | 0.88 | 0.751 |
| cluster4112 | 1.64 | 0.751 |
| cluster2867 | 1.45 | 0.751 |
| cluster2957 | -1.22 | 0.751 |
| cluster2962 | 0.75 | 0.751 |
| cluster2982 | -1.05 | 0.751 |
| cluster3090 | -1.58 | 0.751 |
| cluster3198 | 1.04 | 0.751 |
| cluster3286 | -0.82 | 0.751 |
| cluster3302 | -0.95 | 0.751 |
| cluster3314 | -1.26 | 0.751 |
| cluster3363 | 0.92 | 0.751 |
| cluster3399 | -1.28 | 0.751 |
| cluster3402 | -1.06 | 0.751 |
| cluster3471 | 1.14 | 0.751 |
| cluster3480 | 1.06 | 0.751 |
| cluster3511 | 1.21 | 0.751 |
| cluster3561 | -1.18 | 0.751 |
| cluster3583 | -0.74 | 0.751 |
| cluster3593 | 1.17 | 0.751 |

| | | |
|---|---|---|
| cluster3663 | 1.17 | 0.751 |
| cluster3719 | 1.25 | 0.751 |
| cluster3735 | 1.18 | 0.751 |
| cluster3763 | -0.90 | 0.751 |
| cluster3765 | 1.07 | 0.751 |
| cluster3773 | 0.94 | 0.751 |
| cluster3812 | -1.27 | 0.751 |
| cluster3856 | 0.95 | 0.751 |
| cluster3955 | 1.45 | 0.751 |
| cluster3998 | 1.47 | 0.751 |
| cluster4009 | -1.20 | 0.751 |
| cluster4027 | 1.25 | 0.751 |
| cluster3127 | 0.77 | 0.752 |
| cluster3351 | 0.62 | 0.752 |
| cluster3981 | 1.29 | 0.757 |
| cluster3131 | 0.95 | 0.760 |
| cluster3404 | 0.92 | 0.760 |
| cluster3585 | -0.87 | 0.760 |
| cluster3012 | 1.30 | 0.764 |
| cluster3343 | -0.81 | 0.764 |
| cluster2872 | -0.78 | 0.764 |
| cluster2918 | -0.87 | 0.764 |
| cluster2959 | -0.84 | 0.764 |
| cluster3187 | -1.40 | 0.764 |
| cluster3200 | 1.05 | 0.764 |
| cluster3263 | -0.70 | 0.764 |
| cluster3291 | 0.99 | 0.764 |
| cluster3293 | 0.82 | 0.764 |
| cluster3313 | 1.00 | 0.764 |
| cluster3368 | -0.82 | 0.764 |
| cluster3481 | 1.39 | 0.764 |
| cluster3562 | -1.00 | 0.764 |
| cluster3671 | 0.92 | 0.764 |
| cluster3738 | 0.96 | 0.764 |
| cluster3741 | -1.13 | 0.764 |
| cluster3783 | 1.32 | 0.764 |
| cluster3784 | 0.98 | 0.764 |
| cluster4053 | 1.03 | 0.764 |
| cluster4071 | 1.42 | 0.764 |
| cluster4084 | 1.03 | 0.764 |
| cluster4099 | 0.99 | 0.764 |
| cluster3819 | 1.37 | 0.765 |
| cluster3224 | -0.82 | 0.767 |
| cluster3259 | -0.87 | 0.767 |

| | | |
|---|---|---|
| cluster3279 | 0.93 | 0.767 |
| cluster3405 | -0.87 | 0.767 |
| cluster3408 | -1.11 | 0.767 |
| cluster3677 | 0.95 | 0.767 |
| cluster3905 | 2.39 | 0.767 |
| cluster3915 | 1.46 | 0.767 |
| cluster3982 | 0.73 | 0.767 |
| cluster4087 | 0.89 | 0.767 |
| cluster2878 | -1.07 | 0.768 |
| cluster3107 | 0.78 | 0.768 |
| cluster3392 | 1.32 | 0.768 |
| cluster3609 | -1.47 | 0.768 |
| cluster3823 | 0.97 | 0.768 |
| cluster3848 | -1.09 | 0.768 |
| cluster4105 | 1.03 | 0.768 |
| cluster2901 | -1.07 | 0.768 |
| cluster3747 | 0.84 | 0.768 |
| cluster3731 | 1.10 | 0.771 |
| cluster2981 | 0.78 | 0.772 |
| cluster4033 | 1.46 | 0.772 |
| cluster3600 | -0.72 | 0.775 |
| cluster3447 | 0.98 | 0.777 |
| cluster4046 | 1.53 | 0.777 |
| cluster3991 | 1.13 | 0.777 |
| cluster3271 | 0.78 | 0.778 |
| cluster3533 | 0.99 | 0.778 |
| cluster3659 | 1.16 | 0.778 |
| cluster3894 | 0.87 | 0.778 |
| cluster3375 | 0.78 | 0.779 |
| cluster3065 | -0.88 | 0.781 |
| cluster3144 | 1.35 | 0.781 |
| cluster4035 | 1.03 | 0.781 |
| cluster2988 | 0.66 | 0.783 |
| cluster3551 | -0.78 | 0.783 |
| cluster3658 | -0.85 | 0.783 |
| cluster3833 | -1.11 | 0.783 |
| cluster3861 | 0.67 | 0.783 |
| cluster3164 | -1.24 | 0.789 |
| cluster2892 | 0.83 | 0.796 |
| cluster2960 | -0.54 | 0.796 |
| cluster3033 | 0.59 | 0.796 |
| cluster3083 | 0.75 | 0.796 |
| cluster3276 | -0.77 | 0.796 |
| cluster3360 | 1.00 | 0.796 |

| | | |
|---|---|---|
| cluster3382 | 0.80 | 0.796 |
| cluster3479 | 1.15 | 0.796 |
| cluster3505 | -0.74 | 0.796 |
| cluster3568 | -1.22 | 0.796 |
| cluster3598 | 1.63 | 0.796 |
| cluster3674 | -1.06 | 0.796 |
| cluster3937 | 1.00 | 0.796 |
| cluster4007 | 0.61 | 0.796 |
| cluster4023 | -1.02 | 0.796 |
| cluster4050 | 1.07 | 0.796 |
| cluster4100 | 1.11 | 0.796 |
| cluster3064 | -0.96 | 0.798 |
| cluster3653 | 1.04 | 0.800 |
| cluster3537 | 0.74 | 0.806 |
| cluster3280 | -0.70 | 0.810 |
| cluster3572 | 1.53 | 0.810 |
| cluster3015 | -1.06 | 0.811 |
| cluster3022 | -0.90 | 0.812 |
| cluster3053 | 1.17 | 0.812 |
| cluster3125 | 0.67 | 0.812 |
| cluster3203 | 0.72 | 0.812 |
| cluster3525 | -0.74 | 0.812 |
| cluster3558 | -0.83 | 0.812 |
| cluster3979 | 0.83 | 0.814 |
| cluster3377 | -0.82 | 0.814 |
| cluster3882 | 1.01 | 0.814 |
| cluster3980 | 0.62 | 0.814 |
| cluster2977 | -0.76 | 0.814 |
| cluster3702 | -0.82 | 0.814 |
| cluster3623 | 0.80 | 0.816 |
| cluster4061 | 0.94 | 0.816 |
| cluster3113 | -0.65 | 0.816 |
| cluster2973 | -0.90 | 0.817 |
| cluster3637 | 0.56 | 0.822 |
| cluster4111 | -1.35 | 0.822 |
| cluster3042 | 0.87 | 0.822 |
| cluster3081 | 0.89 | 0.822 |
| cluster3095 | 0.88 | 0.822 |
| cluster3136 | 0.85 | 0.822 |
| cluster3189 | -0.85 | 0.822 |
| cluster3316 | -0.92 | 0.822 |
| cluster3614 | 1.20 | 0.822 |
| cluster3820 | 0.58 | 0.822 |
| cluster3875 | 0.77 | 0.822 |

| | | |
|---|---|---|
| cluster3942 | -0.78 | 0.822 |
| cluster3587 | 1.26 | 0.823 |
| cluster3863 | 0.96 | 0.823 |
| cluster3393 | 1.28 | 0.825 |
| cluster3135 | -0.83 | 0.828 |
| cluster3594 | 0.92 | 0.828 |
| cluster3836 | 0.79 | 0.828 |
| cluster3880 | 1.33 | 0.831 |
| cluster3020 | -0.73 | 0.831 |
| cluster3163 | 1.11 | 0.831 |
| cluster3357 | -0.73 | 0.831 |
| cluster3365 | 0.93 | 0.831 |
| cluster3426 | -0.56 | 0.831 |
| cluster3449 | 0.52 | 0.831 |
| cluster3578 | -0.83 | 0.831 |
| cluster3648 | -0.73 | 0.831 |
| cluster3724 | 0.93 | 0.831 |
| cluster3970 | 1.02 | 0.831 |
| cluster3644 | 0.90 | 0.833 |
| cluster2997 | -0.67 | 0.836 |
| cluster3992 | 0.91 | 0.836 |
| cluster3142 | 0.94 | 0.836 |
| cluster3147 | 0.66 | 0.836 |
| cluster3230 | -0.71 | 0.836 |
| cluster3646 | 1.10 | 0.836 |
| cluster3968 | 0.90 | 0.836 |
| cluster3989 | -0.87 | 0.836 |
| cluster4083 | 1.01 | 0.838 |
| cluster3790 | -1.02 | 0.846 |
| cluster2908 | -0.79 | 0.846 |
| cluster3011 | 0.89 | 0.846 |
| cluster3038 | -0.86 | 0.846 |
| cluster3068 | -0.59 | 0.846 |
| cluster3139 | 1.13 | 0.846 |
| cluster3667 | 0.71 | 0.846 |
| cluster3881 | 0.88 | 0.846 |
| cluster3933 | -0.79 | 0.846 |
| cluster4003 | -1.04 | 0.846 |
| cluster4015 | 0.82 | 0.846 |
| cluster4026 | 0.94 | 0.846 |
| cluster4114 | -1.32 | 0.846 |
| cluster4128 | 1.20 | 0.846 |
| cluster3534 | -0.93 | 0.847 |
| cluster2860 | 0.52 | 0.847 |

| | | |
|---|---|---|
| cluster3088 | 0.78 | 0.847 |
| cluster3591 | 0.66 | 0.847 |
| cluster3781 | 0.77 | 0.847 |
| cluster3893 | 0.63 | 0.847 |
| cluster4017 | 0.85 | 0.847 |
| cluster4122 | -0.90 | 0.847 |
| cluster3048 | 0.59 | 0.851 |
| cluster3079 | 0.78 | 0.851 |
| cluster3498 | 0.84 | 0.851 |
| cluster3813 | 0.88 | 0.851 |
| cluster3096 | 0.56 | 0.853 |
| cluster2887 | 0.72 | 0.854 |
| cluster3017 | 0.89 | 0.854 |
| cluster3124 | -0.47 | 0.854 |
| cluster3173 | 0.91 | 0.854 |
| cluster3497 | 0.50 | 0.854 |
| cluster3951 | 0.88 | 0.854 |
| cluster4037 | 0.62 | 0.854 |
| cluster4080 | 1.16 | 0.854 |
| cluster4126 | 1.12 | 0.854 |
| cluster3638 | 0.98 | 0.855 |
| cluster3294 | 0.69 | 0.856 |
| cluster3755 | 1.02 | 0.856 |
| cluster3769 | -0.61 | 0.857 |
| cluster4000 | 0.74 | 0.857 |
| cluster3473 | 0.89 | 0.858 |
| cluster3070 | -0.73 | 0.858 |
| cluster3009 | 0.80 | 0.859 |
| cluster3311 | 0.75 | 0.859 |
| cluster3661 | 0.65 | 0.859 |
| cluster3714 | -0.51 | 0.859 |
| cluster2853 | -0.46 | 0.869 |
| cluster2886 | 0.77 | 0.869 |
| cluster3494 | 0.46 | 0.869 |
| cluster3809 | 0.55 | 0.869 |
| cluster3939 | 0.64 | 0.869 |
| cluster4109 | 0.91 | 0.869 |
| cluster3006 | -0.71 | 0.869 |
| cluster3036 | -0.66 | 0.869 |
| cluster3114 | -0.54 | 0.869 |
| cluster3149 | 0.60 | 0.869 |
| cluster3245 | -0.84 | 0.869 |
| cluster3266 | -0.59 | 0.869 |
| cluster3310 | -0.46 | 0.869 |

| | | |
|---|---|---|
| cluster3348 | -0.76 | 0.869 |
| cluster3366 | 0.89 | 0.869 |
| cluster3372 | 0.60 | 0.869 |
| cluster3376 | 0.57 | 0.869 |
| cluster3409 | 0.91 | 0.869 |
| cluster3575 | -0.85 | 0.869 |
| cluster3592 | 0.64 | 0.869 |
| cluster3606 | 0.90 | 0.869 |
| cluster3869 | 0.99 | 0.869 |
| cluster4039 | -0.76 | 0.869 |
| cluster4029 | 1.08 | 0.871 |
| cluster4036 | 0.51 | 0.872 |
| cluster3483 | -0.50 | 0.873 |
| cluster3345 | 0.64 | 0.874 |
| cluster3395 | 0.68 | 0.874 |
| cluster2987 | -0.52 | 0.876 |
| cluster3268 | 0.58 | 0.876 |
| cluster3490 | 0.77 | 0.877 |
| cluster3213 | 0.78 | 0.878 |
| cluster3564 | 0.92 | 0.878 |
| cluster3774 | -0.70 | 0.878 |
| cluster3265 | 0.64 | 0.879 |
| cluster3416 | 0.96 | 0.879 |
| cluster3550 | -0.69 | 0.879 |
| cluster2990 | -0.55 | 0.885 |
| cluster3867 | 0.56 | 0.889 |
| cluster3850 | 0.71 | 0.891 |
| cluster4106 | 0.98 | 0.891 |
| cluster3050 | -0.56 | 0.892 |
| cluster2956 | -0.62 | 0.894 |
| cluster3764 | 0.90 | 0.894 |
| cluster2932 | 0.60 | 0.895 |
| cluster2933 | -0.79 | 0.895 |
| cluster2938 | 0.55 | 0.895 |
| cluster3300 | -0.61 | 0.895 |
| cluster3649 | -0.49 | 0.895 |
| cluster3913 | -0.75 | 0.895 |
| cluster3920 | 0.76 | 0.895 |
| cluster4008 | 0.76 | 0.895 |
| cluster4117 | -1.20 | 0.895 |
| cluster2910 | -0.51 | 0.897 |
| cluster2949 | -0.54 | 0.897 |
| cluster3025 | 0.51 | 0.897 |
| cluster3092 | 0.49 | 0.897 |

| | | |
|---|---|---|
| cluster3241 | -0.48 | 0.897 |
| cluster3282 | -0.57 | 0.897 |
| cluster3305 | -0.77 | 0.897 |
| cluster3410 | 0.52 | 0.897 |
| cluster3632 | 0.64 | 0.897 |
| cluster3811 | 0.66 | 0.897 |
| cluster3963 | 0.91 | 0.897 |
| cluster4090 | -1.10 | 0.897 |
| cluster3640 | 1.67 | 0.898 |
| cluster4073 | 1.02 | 0.899 |
| cluster3321 | -0.51 | 0.902 |
| cluster2913 | 0.48 | 0.902 |
| cluster2970 | 0.39 | 0.902 |
| cluster3034 | 0.55 | 0.902 |
| cluster3389 | 0.59 | 0.902 |
| cluster3486 | 0.73 | 0.902 |
| cluster3503 | 0.54 | 0.902 |
| cluster3513 | -0.70 | 0.902 |
| cluster3595 | 0.62 | 0.902 |
| cluster3617 | 0.85 | 0.902 |
| cluster3736 | -0.73 | 0.902 |
| cluster3887 | 0.78 | 0.902 |
| cluster3974 | 0.99 | 0.902 |
| cluster4013 | -0.89 | 0.902 |
| cluster3076 | 0.59 | 0.903 |
| cluster3622 | 0.50 | 0.903 |
| cluster3694 | 0.63 | 0.903 |
| cluster2897 | -0.57 | 0.903 |
| cluster2953 | -0.51 | 0.903 |
| cluster3272 | -1.50 | 0.903 |
| cluster3625 | 0.48 | 0.903 |
| cluster3698 | -0.59 | 0.903 |
| cluster3948 | 0.57 | 0.903 |
| cluster3962 | -0.44 | 0.903 |
| cluster3883 | -0.93 | 0.903 |
| cluster2965 | -0.54 | 0.907 |
| cluster3018 | 0.45 | 0.907 |
| cluster4123 | 0.95 | 0.909 |
| cluster2874 | -0.38 | 0.910 |
| cluster2923 | -0.48 | 0.910 |
| cluster2941 | -0.43 | 0.910 |
| cluster2993 | 0.71 | 0.910 |
| cluster2998 | 0.47 | 0.910 |
| cluster3318 | -0.45 | 0.910 |

| | | |
|---|---|---|
| cluster3346 | 0.41 | 0.910 |
| cluster3470 | 0.44 | 0.910 |
| cluster3531 | 0.61 | 0.910 |
| cluster3579 | 0.59 | 0.910 |
| cluster3628 | 0.63 | 0.910 |
| cluster3633 | 0.56 | 0.910 |
| cluster3712 | 0.60 | 0.910 |
| cluster3797 | -0.62 | 0.910 |
| cluster3826 | 0.72 | 0.910 |
| cluster3839 | -0.79 | 0.910 |
| cluster3874 | -0.51 | 0.910 |
| cluster3945 | 0.73 | 0.910 |
| cluster3085 | -0.66 | 0.912 |
| cluster3249 | -0.49 | 0.912 |
| cluster3650 | 0.74 | 0.912 |
| cluster3941 | -0.50 | 0.912 |
| cluster4064 | 0.60 | 0.912 |
| cluster3485 | 0.88 | 0.913 |
| cluster3669 | 0.51 | 0.913 |
| cluster3450 | 0.95 | 0.913 |
| cluster3656 | 0.60 | 0.913 |
| cluster2888 | 0.87 | 0.914 |
| cluster3795 | 0.52 | 0.915 |
| cluster3958 | 0.84 | 0.915 |
| cluster3969 | 0.45 | 0.915 |
| cluster2914 | 0.64 | 0.916 |
| cluster3121 | -0.61 | 0.916 |
| cluster3570 | 0.79 | 0.916 |
| cluster3696 | 0.43 | 0.916 |
| cluster3708 | 0.85 | 0.916 |
| cluster3873 | 0.51 | 0.916 |
| cluster3922 | 0.84 | 0.916 |
| cluster3928 | 0.50 | 0.919 |
| cluster2978 | -0.38 | 0.920 |
| cluster3509 | -0.39 | 0.920 |
| cluster3984 | -0.88 | 0.920 |
| cluster3199 | -0.44 | 0.921 |
| cluster3251 | -0.43 | 0.921 |
| cluster3129 | -0.93 | 0.922 |
| cluster3188 | -0.42 | 0.922 |
| cluster3762 | -0.69 | 0.923 |
| cluster4067 | -0.70 | 0.923 |
| cluster3215 | -0.40 | 0.923 |
| cluster3269 | -0.74 | 0.923 |

| | | |
|---|---|---|
| cluster3347 | 0.42 | 0.923 |
| cluster3672 | 0.61 | 0.923 |
| cluster4006 | -0.44 | 0.923 |
| cluster3879 | 0.45 | 0.923 |
| cluster2919 | -0.51 | 0.924 |
| cluster3234 | 0.52 | 0.924 |
| cluster4093 | -0.61 | 0.924 |
| cluster2871 | -0.40 | 0.926 |
| cluster3462 | 0.57 | 0.926 |
| cluster4048 | 0.57 | 0.926 |
| cluster2859 | 0.35 | 0.927 |
| cluster3642 | 0.60 | 0.927 |
| cluster3739 | 0.40 | 0.927 |
| cluster2899 | -0.43 | 0.927 |
| cluster3380 | -0.47 | 0.927 |
| cluster3398 | 0.44 | 0.927 |
| cluster3431 | -0.51 | 0.927 |
| cluster3094 | 0.69 | 0.927 |
| cluster3777 | -0.49 | 0.927 |
| cluster2915 | 0.53 | 0.929 |
| cluster3010 | -0.50 | 0.929 |
| cluster3141 | -0.35 | 0.929 |
| cluster3168 | 0.51 | 0.929 |
| cluster3239 | -0.45 | 0.929 |
| cluster3277 | -0.58 | 0.929 |
| cluster3569 | 0.61 | 0.929 |
| cluster4059 | 0.76 | 0.929 |
| cluster3574 | 0.77 | 0.929 |
| cluster2854 | 0.56 | 0.930 |
| cluster3540 | -0.63 | 0.930 |
| cluster3608 | 0.42 | 0.930 |
| cluster3043 | 0.40 | 0.932 |
| cluster3058 | 0.35 | 0.932 |
| cluster3153 | -0.36 | 0.932 |
| cluster3162 | 0.55 | 0.932 |
| cluster3214 | 0.46 | 0.932 |
| cluster3264 | 0.38 | 0.932 |
| cluster3355 | -0.56 | 0.932 |
| cluster3391 | 0.32 | 0.932 |
| cluster3397 | -0.64 | 0.932 |
| cluster3415 | -0.46 | 0.932 |
| cluster3418 | 0.46 | 0.932 |
| cluster3631 | 0.39 | 0.932 |
| cluster3695 | 0.47 | 0.932 |

| | | |
|---|---|---|
| cluster3815 | 0.67 | 0.932 |
| cluster4020 | 0.44 | 0.932 |
| cluster4072 | 0.44 | 0.932 |
| cluster4075 | -0.72 | 0.932 |
| cluster3021 | 0.41 | 0.936 |
| cluster3029 | -0.35 | 0.936 |
| cluster3031 | -0.49 | 0.936 |
| cluster3039 | 0.41 | 0.936 |
| cluster3072 | -0.55 | 0.936 |
| cluster3132 | 0.45 | 0.936 |
| cluster3159 | -0.33 | 0.936 |
| cluster3218 | 0.59 | 0.936 |
| cluster3226 | 0.56 | 0.936 |
| cluster3326 | -0.54 | 0.936 |
| cluster3334 | 0.38 | 0.936 |
| cluster3338 | 0.52 | 0.936 |
| cluster3381 | 0.55 | 0.936 |
| cluster3477 | -0.55 | 0.936 |
| cluster3532 | 0.59 | 0.936 |
| cluster3599 | 0.60 | 0.936 |
| cluster3621 | 0.54 | 0.936 |
| cluster3716 | -0.61 | 0.936 |
| cluster3789 | -0.60 | 0.936 |
| cluster3986 | 0.35 | 0.936 |
| cluster4092 | -0.61 | 0.936 |
| cluster2930 | 0.42 | 0.939 |
| cluster2985 | 0.47 | 0.939 |
| cluster3155 | -0.52 | 0.939 |
| cluster3273 | -0.38 | 0.939 |
| cluster3275 | 0.58 | 0.939 |
| cluster3287 | -0.49 | 0.939 |
| cluster3327 | 0.52 | 0.939 |
| cluster3358 | 0.51 | 0.939 |
| cluster3412 | 0.43 | 0.939 |
| cluster3425 | -0.46 | 0.939 |
| cluster3433 | -0.43 | 0.939 |
| cluster3495 | 0.41 | 0.939 |
| cluster3508 | 0.59 | 0.939 |
| cluster3526 | 0.33 | 0.939 |
| cluster3725 | 0.57 | 0.939 |
| cluster3788 | 0.46 | 0.939 |
| cluster3923 | 0.36 | 0.939 |
| cluster4049 | -0.89 | 0.939 |
| cluster4082 | 0.62 | 0.939 |

| | | |
|---|---|---|
| cluster3059 | 0.57 | 0.940 |
| cluster3227 | -0.42 | 0.940 |
| cluster2857 | 0.29 | 0.941 |
| cluster2865 | 0.38 | 0.941 |
| cluster2877 | -0.29 | 0.941 |
| cluster2898 | 0.48 | 0.941 |
| cluster2927 | 0.44 | 0.941 |
| cluster2929 | 0.31 | 0.941 |
| cluster2931 | -0.38 | 0.941 |
| cluster2936 | 0.59 | 0.941 |
| cluster2947 | 0.42 | 0.941 |
| cluster2950 | -0.34 | 0.941 |
| cluster2952 | -0.40 | 0.941 |
| cluster2969 | 0.47 | 0.941 |
| cluster2971 | 0.31 | 0.941 |
| cluster2979 | 0.28 | 0.941 |
| cluster2983 | -0.30 | 0.941 |
| cluster2984 | -0.25 | 0.941 |
| cluster2992 | -0.41 | 0.941 |
| cluster2999 | 0.47 | 0.941 |
| cluster3002 | -0.36 | 0.941 |
| cluster3007 | 0.32 | 0.941 |
| cluster3044 | -0.51 | 0.941 |
| cluster3052 | -0.63 | 0.941 |
| cluster3061 | 0.28 | 0.941 |
| cluster3062 | 0.31 | 0.941 |
| cluster3087 | 0.35 | 0.941 |
| cluster3115 | -0.30 | 0.941 |
| cluster3122 | 0.41 | 0.941 |
| cluster3137 | -0.53 | 0.941 |
| cluster3138 | -0.28 | 0.941 |
| cluster3150 | -0.54 | 0.941 |
| cluster3165 | 0.45 | 0.941 |
| cluster3193 | -0.35 | 0.941 |
| cluster3196 | -0.36 | 0.941 |
| cluster3221 | -0.49 | 0.941 |
| cluster3229 | -0.50 | 0.941 |
| cluster3270 | 0.48 | 0.941 |
| cluster3288 | 0.46 | 0.941 |
| cluster3307 | -0.37 | 0.941 |
| cluster3332 | 0.42 | 0.941 |
| cluster3335 | 0.42 | 0.941 |
| cluster3336 | -0.49 | 0.941 |
| cluster3350 | -0.43 | 0.941 |

| | | |
|---|---|---|
| cluster3370 | 0.43 | 0.941 |
| cluster3394 | -0.34 | 0.941 |
| cluster3422 | -0.65 | 0.941 |
| cluster3424 | 0.62 | 0.941 |
| cluster3432 | 0.42 | 0.941 |
| cluster3455 | -0.48 | 0.941 |
| cluster3460 | -0.42 | 0.941 |
| cluster3464 | 0.41 | 0.941 |
| cluster3488 | -0.34 | 0.941 |
| cluster3492 | -0.33 | 0.941 |
| cluster3507 | 0.46 | 0.941 |
| cluster3515 | -0.56 | 0.941 |
| cluster3521 | -0.33 | 0.941 |
| cluster3544 | 0.63 | 0.941 |
| cluster3576 | -0.39 | 0.941 |
| cluster3577 | 0.38 | 0.941 |
| cluster3584 | 0.52 | 0.941 |
| cluster3602 | 0.35 | 0.941 |
| cluster3612 | -0.45 | 0.941 |
| cluster3613 | -0.48 | 0.941 |
| cluster3630 | 0.36 | 0.941 |
| cluster3665 | 0.55 | 0.941 |
| cluster3679 | 0.38 | 0.941 |
| cluster3680 | 0.34 | 0.941 |
| cluster3682 | -0.53 | 0.941 |
| cluster3683 | 0.37 | 0.941 |
| cluster3685 | -0.52 | 0.941 |
| cluster3691 | 0.41 | 0.941 |
| cluster3692 | 0.44 | 0.941 |
| cluster3711 | 0.29 | 0.941 |
| cluster3720 | 0.48 | 0.941 |
| cluster3721 | 0.47 | 0.941 |
| cluster3726 | -0.28 | 0.941 |
| cluster3730 | -0.26 | 0.941 |
| cluster3733 | 0.43 | 0.941 |
| cluster3734 | 0.52 | 0.941 |
| cluster3744 | -0.37 | 0.941 |
| cluster3759 | -0.39 | 0.941 |
| cluster3772 | -0.44 | 0.941 |
| cluster3776 | 0.43 | 0.941 |
| cluster3801 | 0.45 | 0.941 |
| cluster3806 | 0.46 | 0.941 |
| cluster3808 | -0.47 | 0.941 |
| cluster3817 | -0.37 | 0.941 |

| | | |
|---|---|---|
| cluster3844 | 0.64 | 0.941 |
| cluster3845 | 0.70 | 0.941 |
| cluster3846 | 0.28 | 0.941 |
| cluster3860 | -0.37 | 0.941 |
| cluster3888 | 0.48 | 0.941 |
| cluster3935 | -0.44 | 0.941 |
| cluster3947 | 0.27 | 0.941 |
| cluster3954 | 0.52 | 0.941 |
| cluster3964 | -0.61 | 0.941 |
| cluster3997 | 0.51 | 0.941 |
| cluster3999 | 0.48 | 0.941 |
| cluster4018 | -0.45 | 0.941 |
| cluster4063 | 0.61 | 0.941 |
| cluster4070 | 0.67 | 0.941 |
| cluster4089 | -0.46 | 0.941 |
| cluster4102 | -0.35 | 0.941 |
| cluster3003 | -0.39 | 0.945 |
| cluster3016 | -0.50 | 0.945 |
| cluster3152 | 0.42 | 0.945 |
| cluster2855 | -0.25 | 0.948 |
| cluster3069 | -0.28 | 0.948 |
| cluster3170 | -0.33 | 0.948 |
| cluster3244 | -0.33 | 0.948 |
| cluster3388 | -0.26 | 0.948 |
| cluster3673 | 0.46 | 0.948 |
| cluster3766 | 0.38 | 0.948 |
| cluster3929 | -0.38 | 0.948 |
| cluster2866 | -0.40 | 0.952 |
| cluster2881 | 0.36 | 0.952 |
| cluster2903 | 0.42 | 0.952 |
| cluster2951 | -0.38 | 0.952 |
| cluster3040 | -0.34 | 0.952 |
| cluster3133 | -0.27 | 0.952 |
| cluster3201 | -0.27 | 0.952 |
| cluster3223 | 0.38 | 0.952 |
| cluster3361 | 0.31 | 0.952 |
| cluster3465 | -0.22 | 0.952 |
| cluster3557 | 0.31 | 0.952 |
| cluster3627 | 0.26 | 0.952 |
| cluster3664 | 0.28 | 0.952 |
| cluster3767 | 0.37 | 0.952 |
| cluster3847 | 0.31 | 0.952 |
| cluster3898 | 0.44 | 0.952 |
| cluster3911 | -0.44 | 0.952 |

| | | |
|---|---|---|
| cluster3918 | 0.36 | 0.952 |
| cluster4041 | 0.38 | 0.952 |
| cluster3452 | 0.38 | 0.953 |
| cluster3903 | -0.27 | 0.953 |
| cluster2968 | -0.24 | 0.955 |
| cluster3172 | 0.36 | 0.955 |
| cluster3988 | 0.22 | 0.955 |
| cluster2862 | 0.31 | 0.958 |
| cluster2864 | 0.28 | 0.958 |
| cluster3390 | -0.31 | 0.958 |
| cluster3437 | 0.32 | 0.958 |
| cluster3510 | 0.52 | 0.958 |
| cluster3624 | 0.24 | 0.958 |
| cluster3740 | 0.37 | 0.958 |
| cluster4115 | 0.45 | 0.958 |
| cluster2883 | 0.26 | 0.958 |
| cluster2937 | -0.28 | 0.958 |
| cluster2942 | -0.27 | 0.958 |
| cluster2975 | 0.21 | 0.958 |
| cluster3154 | 0.25 | 0.958 |
| cluster3174 | -0.36 | 0.958 |
| cluster3255 | -0.37 | 0.958 |
| cluster3396 | 0.32 | 0.958 |
| cluster3400 | -0.30 | 0.958 |
| cluster3489 | 0.31 | 0.958 |
| cluster3496 | -0.30 | 0.958 |
| cluster3523 | 0.39 | 0.958 |
| cluster3573 | -0.32 | 0.958 |
| cluster3615 | 0.38 | 0.958 |
| cluster3634 | 0.34 | 0.958 |
| cluster3660 | -0.44 | 0.958 |
| cluster3729 | -0.26 | 0.958 |
| cluster3925 | 1.00 | 0.958 |
| cluster4110 | 0.36 | 0.958 |
| cluster2856 | -0.33 | 0.960 |
| cluster2895 | -0.27 | 0.960 |
| cluster2963 | 0.35 | 0.960 |
| cluster3284 | 0.21 | 0.960 |
| cluster3303 | 0.27 | 0.960 |
| cluster3438 | 0.30 | 0.960 |
| cluster3541 | -0.28 | 0.960 |
| cluster3699 | 0.37 | 0.960 |
| cluster3914 | 0.50 | 0.960 |
| cluster3967 | 0.32 | 0.960 |

| | | |
|---|---|---|
| cluster4047 | 0.45 | 0.960 |
| cluster4056 | 0.33 | 0.960 |
| cluster3028 | -0.22 | 0.960 |
| cluster3205 | -0.24 | 0.960 |
| cluster3522 | -0.32 | 0.960 |
| cluster3140 | 0.21 | 0.962 |
| cluster3190 | 0.39 | 0.962 |
| cluster3331 | -0.20 | 0.962 |
| cluster3635 | 0.24 | 0.962 |
| cluster3727 | -0.34 | 0.962 |
| cluster3876 | 0.30 | 0.962 |
| cluster3407 | 0.28 | 0.963 |
| cluster3946 | -0.32 | 0.963 |
| cluster2900 | 0.30 | 0.964 |
| cluster2909 | 0.19 | 0.964 |
| cluster2916 | 0.28 | 0.964 |
| cluster2964 | -0.24 | 0.964 |
| cluster2966 | -0.27 | 0.964 |
| cluster2974 | -0.29 | 0.964 |
| cluster2976 | 0.23 | 0.964 |
| cluster2980 | 0.37 | 0.964 |
| cluster3054 | -0.16 | 0.964 |
| cluster3055 | -0.20 | 0.964 |
| cluster3067 | -0.23 | 0.964 |
| cluster3143 | 0.24 | 0.964 |
| cluster3158 | -0.32 | 0.964 |
| cluster3181 | -0.16 | 0.964 |
| cluster3237 | -0.23 | 0.964 |
| cluster3242 | 0.23 | 0.964 |
| cluster3261 | -0.31 | 0.964 |
| cluster3289 | 0.24 | 0.964 |
| cluster3304 | 0.24 | 0.964 |
| cluster3312 | 0.39 | 0.964 |
| cluster3384 | 0.27 | 0.964 |
| cluster3421 | 0.30 | 0.964 |
| cluster3423 | 0.31 | 0.964 |
| cluster3440 | -0.19 | 0.964 |
| cluster3445 | 0.21 | 0.964 |
| cluster3459 | 0.37 | 0.964 |
| cluster3463 | -0.27 | 0.964 |
| cluster3469 | 0.30 | 0.964 |
| cluster3499 | 0.28 | 0.964 |
| cluster3539 | 0.38 | 0.964 |
| cluster3548 | 0.22 | 0.964 |

| | | |
|---|---|---|
| cluster3552 | 0.22 | 0.964 |
| cluster3555 | -0.34 | 0.964 |
| cluster3589 | -0.20 | 0.964 |
| cluster3603 | -0.28 | 0.964 |
| cluster3675 | -0.22 | 0.964 |
| cluster3697 | 0.33 | 0.964 |
| cluster3750 | 0.38 | 0.964 |
| cluster3751 | 0.19 | 0.964 |
| cluster3917 | 0.31 | 0.964 |
| cluster3995 | -0.31 | 0.964 |
| cluster4005 | -0.28 | 0.964 |
| cluster4011 | 0.39 | 0.964 |
| cluster4065 | 0.19 | 0.964 |
| cluster3325 | 0.38 | 0.964 |
| cluster1127 | 0.18 | 0.965 |
| cluster2870 | 0.15 | 0.965 |
| cluster2873 | -0.13 | 0.965 |
| cluster2879 | 0.16 | 0.965 |
| cluster2880 | -0.21 | 0.965 |
| cluster2884 | 0.16 | 0.965 |
| cluster2890 | -0.19 | 0.965 |
| cluster2896 | 0.16 | 0.965 |
| cluster2920 | -0.15 | 0.965 |
| cluster2921 | -0.19 | 0.965 |
| cluster2922 | -0.18 | 0.965 |
| cluster2945 | 0.27 | 0.965 |
| cluster2958 | 0.14 | 0.965 |
| cluster2972 | 0.23 | 0.965 |
| cluster2991 | -0.17 | 0.965 |
| cluster3023 | 0.13 | 0.965 |
| cluster3027 | -0.14 | 0.965 |
| cluster3035 | -0.18 | 0.965 |
| cluster3049 | 0.20 | 0.965 |
| cluster3066 | 0.19 | 0.965 |
| cluster3080 | -0.18 | 0.965 |
| cluster3104 | -0.12 | 0.965 |
| cluster3105 | -0.28 | 0.965 |
| cluster3119 | -0.12 | 0.965 |
| cluster3130 | -0.16 | 0.965 |
| cluster3134 | -0.17 | 0.965 |
| cluster3169 | -0.23 | 0.965 |
| cluster3180 | 0.15 | 0.965 |
| cluster3197 | -0.20 | 0.965 |
| cluster3209 | 0.16 | 0.965 |

| | | |
|---|---|---|
| cluster3216 | 0.15 | 0.965 |
| cluster3220 | 0.20 | 0.965 |
| cluster3232 | 0.15 | 0.965 |
| cluster3233 | 0.25 | 0.965 |
| cluster3267 | 0.19 | 0.965 |
| cluster3278 | 0.16 | 0.965 |
| cluster3292 | -0.21 | 0.965 |
| cluster3295 | 0.17 | 0.965 |
| cluster3317 | 0.23 | 0.965 |
| cluster3330 | 0.17 | 0.965 |
| cluster3340 | 0.18 | 0.965 |
| cluster3352 | -0.18 | 0.965 |
| cluster3401 | -0.14 | 0.965 |
| cluster3403 | -0.16 | 0.965 |
| cluster3435 | 0.24 | 0.965 |
| cluster3446 | -0.18 | 0.965 |
| cluster3458 | -0.15 | 0.965 |
| cluster3484 | -0.28 | 0.965 |
| cluster3514 | 0.28 | 0.965 |
| cluster3518 | 0.28 | 0.965 |
| cluster3520 | -0.35 | 0.965 |
| cluster3542 | 0.28 | 0.965 |
| cluster3553 | 0.17 | 0.965 |
| cluster3563 | -0.24 | 0.965 |
| cluster3565 | 0.21 | 0.965 |
| cluster3566 | -0.14 | 0.965 |
| cluster3590 | -0.18 | 0.965 |
| cluster3601 | -0.20 | 0.965 |
| cluster3626 | 0.19 | 0.965 |
| cluster3643 | -0.25 | 0.965 |
| cluster3662 | 0.24 | 0.965 |
| cluster3676 | 0.23 | 0.965 |
| cluster3684 | -0.20 | 0.965 |
| cluster3687 | 0.19 | 0.965 |
| cluster3705 | -0.17 | 0.965 |
| cluster3737 | -0.14 | 0.965 |
| cluster3753 | 0.26 | 0.965 |
| cluster3786 | 0.36 | 0.965 |
| cluster3799 | 0.24 | 0.965 |
| cluster3814 | -0.22 | 0.965 |
| cluster3828 | 0.23 | 0.965 |
| cluster3870 | 0.30 | 0.965 |
| cluster3907 | -0.30 | 0.965 |
| cluster3910 | 0.23 | 0.965 |

| | | |
|---|---|---|
| cluster3919 | 0.25 | 0.965 |
| cluster3921 | -0.29 | 0.965 |
| cluster3931 | 0.29 | 0.965 |
| cluster3943 | 0.23 | 0.965 |
| cluster3953 | -0.16 | 0.965 |
| cluster3961 | 0.28 | 0.965 |
| cluster3990 | -0.19 | 0.965 |
| cluster4004 | -0.14 | 0.965 |
| cluster4012 | 0.21 | 0.965 |
| cluster4043 | -0.23 | 0.965 |
| cluster4085 | -0.23 | 0.965 |
| cluster4088 | -0.26 | 0.965 |
| cluster4124 | -0.31 | 0.965 |
| cluster2939 | 0.13 | 0.966 |
| cluster3290 | -0.19 | 0.966 |
| cluster3369 | -0.14 | 0.966 |
| cluster3383 | -0.25 | 0.966 |
| cluster3434 | 0.19 | 0.966 |
| cluster2869 | 0.20 | 0.967 |
| cluster3056 | -0.14 | 0.967 |
| cluster3148 | -0.10 | 0.967 |
| cluster3161 | -0.15 | 0.967 |
| cluster3225 | -0.19 | 0.967 |
| cluster3896 | -0.19 | 0.967 |
| cluster3934 | -0.10 | 0.967 |
| cluster4019 | 0.21 | 0.967 |
| cluster2875 | 0.15 | 0.970 |
| cluster2944 | -0.14 | 0.970 |
| cluster3051 | -0.18 | 0.970 |
| cluster3082 | 0.23 | 0.970 |
| cluster3206 | -0.16 | 0.970 |
| cluster3258 | -0.12 | 0.970 |
| cluster3308 | 0.12 | 0.970 |
| cluster3324 | -0.13 | 0.970 |
| cluster3581 | 0.16 | 0.970 |
| cluster3636 | 0.11 | 0.970 |
| cluster4034 | 0.25 | 0.970 |
| cluster4103 | -0.19 | 0.970 |
| cluster3183 | 0.13 | 0.970 |
| cluster3475 | -0.13 | 0.971 |
| cluster2911 | 0.12 | 0.971 |
| cluster3116 | 0.10 | 0.971 |
| cluster3414 | 0.14 | 0.971 |
| cluster4077 | -0.14 | 0.971 |

| | | |
|---|---|---|
| cluster3960 | -0.19 | 0.972 |
| cluster3084 | -0.11 | 0.973 |
| cluster3208 | -0.11 | 0.973 |
| cluster3444 | 0.11 | 0.973 |
| cluster3655 | -0.12 | 0.973 |
| cluster3932 | 0.14 | 0.973 |
| cluster4076 | 0.12 | 0.973 |
| cluster2967 | -0.15 | 0.974 |
| cluster3580 | 0.16 | 0.974 |
| cluster3597 | -0.14 | 0.974 |
| cluster3792 | -0.11 | 0.974 |
| cluster3791 | -0.14 | 0.975 |
| cluster2954 | -0.13 | 0.976 |
| cluster3491 | 0.10 | 0.976 |
| cluster3605 | 0.10 | 0.976 |
| cluster3868 | 0.09 | 0.976 |
| cluster4094 | -0.13 | 0.976 |
| cluster2861 | -0.12 | 0.977 |
| cluster2882 | -0.08 | 0.977 |
| cluster2955 | 0.08 | 0.977 |
| cluster3045 | -0.11 | 0.977 |
| cluster3047 | -0.12 | 0.977 |
| cluster3110 | 0.08 | 0.977 |
| cluster3120 | 0.06 | 0.977 |
| cluster3157 | 0.12 | 0.977 |
| cluster3184 | -0.09 | 0.977 |
| cluster3202 | 0.09 | 0.977 |
| cluster3248 | 0.14 | 0.977 |
| cluster3262 | -0.09 | 0.977 |
| cluster3337 | 0.07 | 0.977 |
| cluster3344 | 0.10 | 0.977 |
| cluster3453 | 0.12 | 0.977 |
| cluster3456 | -0.09 | 0.977 |
| cluster3519 | -0.16 | 0.977 |
| cluster3535 | -0.14 | 0.977 |
| cluster3536 | -0.10 | 0.977 |
| cluster3582 | 0.09 | 0.977 |
| cluster3586 | 0.08 | 0.977 |
| cluster3596 | -0.11 | 0.977 |
| cluster3690 | -0.10 | 0.977 |
| cluster3775 | -0.09 | 0.977 |
| cluster3872 | 0.11 | 0.977 |
| cluster3977 | -0.13 | 0.977 |
| cluster3994 | 0.16 | 0.977 |

| | | |
|---|---|---|
| cluster3996 | -0.07 | 0.977 |
| cluster3353 | -0.06 | 0.979 |
| cluster3442 | -0.11 | 0.979 |
| cluster3097 | -0.09 | 0.979 |
| cluster3373 | 0.08 | 0.979 |
| cluster3468 | -0.09 | 0.979 |
| cluster4097 | 0.16 | 0.979 |
| cluster3013 | -0.08 | 0.981 |
| cluster3768 | -0.11 | 0.981 |
| cluster3588 | 0.09 | 0.983 |
| cluster3000 | -0.08 | 0.984 |
| cluster3281 | -0.08 | 0.984 |
| cluster3756 | -0.08 | 0.984 |
| cluster2986 | 0.06 | 0.985 |
| cluster3210 | 0.05 | 0.985 |
| cluster3253 | 0.07 | 0.985 |
| cluster3700 | 0.07 | 0.985 |
| cluster3722 | -0.07 | 0.985 |
| cluster3761 | -0.10 | 0.985 |
| cluster3771 | -0.07 | 0.985 |
| cluster3825 | 0.07 | 0.985 |
| cluster4021 | 0.08 | 0.985 |
| cluster4028 | 0.06 | 0.985 |
| cluster3527 | 0.08 | 0.985 |
| cluster3610 | -0.08 | 0.985 |
| cluster3093 | 0.07 | 0.986 |
| cluster3341 | -0.07 | 0.986 |
| cluster3298 | 0.05 | 0.987 |
| cluster3441 | -0.05 | 0.987 |
| cluster3798 | -0.06 | 0.987 |
| cluster3900 | 0.10 | 0.987 |
| cluster3641 | 0.06 | 0.988 |
| cluster3706 | -0.04 | 0.988 |
| cluster2889 | 0.03 | 0.988 |
| cluster2907 | -0.04 | 0.988 |
| cluster2943 | -0.06 | 0.988 |
| cluster2996 | -0.04 | 0.988 |
| cluster3014 | -0.05 | 0.988 |
| cluster3074 | 0.04 | 0.988 |
| cluster3091 | -0.04 | 0.988 |
| cluster3247 | -0.05 | 0.988 |
| cluster3339 | -0.04 | 0.988 |
| cluster3354 | 0.04 | 0.988 |
| cluster3367 | 0.03 | 0.988 |

| | | |
|---|---|---|
| cluster3429 | -0.04 | 0.988 |
| cluster3493 | 0.06 | 0.988 |
| cluster3524 | -0.04 | 0.988 |
| cluster3528 | 0.04 | 0.988 |
| cluster3530 | 0.04 | 0.988 |
| cluster3745 | -0.08 | 0.988 |
| cluster3837 | 0.06 | 0.988 |
| cluster2917 | 0.04 | 0.989 |
| cluster2940 | 0.03 | 0.989 |
| cluster3057 | -0.05 | 0.989 |
| cluster3204 | -0.04 | 0.989 |
| cluster3238 | -0.03 | 0.989 |
| cluster3322 | 0.05 | 0.989 |
| cluster3803 | 0.05 | 0.989 |
| cluster3832 | 0.04 | 0.989 |
| cluster3835 | 0.04 | 0.989 |
| cluster2891 | 0.03 | 0.990 |
| cluster3001 | 0.02 | 0.990 |
| cluster3086 | -0.03 | 0.990 |
| cluster3145 | 0.03 | 0.990 |
| cluster3254 | 0.03 | 0.990 |
| cluster3538 | -0.04 | 0.990 |
| cluster3742 | 0.04 | 0.990 |
| cluster3862 | 0.02 | 0.990 |
| cluster3466 | -0.03 | 0.990 |
| cluster3560 | -0.03 | 0.990 |
| cluster3829 | 0.03 | 0.990 |
| cluster3857 | -0.02 | 0.990 |
| cluster3959 | -0.03 | 0.990 |
| cluster3073 | -0.01 | 0.991 |
| cluster3320 | 0.02 | 0.991 |
| cluster3472 | -0.01 | 0.991 |
| cluster3549 | -0.02 | 0.991 |
| cluster3895 | 0.02 | 0.991 |
| cluster3973 | 0.02 | 0.991 |
| cluster4025 | -0.02 | 0.991 |
| cluster3807 | -0.01 | 0.991 |
| cluster3228 | -0.01 | 0.993 |
| cluster3285 | -0.01 | 0.993 |
| cluster3436 | -0.02 | 0.993 |
| cluster3430 | 0.01 | 0.995 |
| cluster3841 | -0.01 | 0.995 |
| cluster3816 | -0.01 | 0.997 |
| cluster3217 | 0.00 | 0.999 |

| | | |
|---|---|---|
| cluster3796 | 0.00 | 0.999 |

**Supplementary Table 4.** edgeR results with eXpress counts - Parasites vs Saprophagous

| ID | log2FoldChange | Adjusted p value |
|---|---|---|
| cluster3457 | -3.53 | 0.025 |
| cluster4091 | -3.68 | 0.153 |
| cluster3885 | -2.35 | 0.392 |
| cluster3859 | -2.36 | 0.392 |
| cluster3975 | 3.41 | 0.392 |
| cluster3111 | -2.39 | 0.392 |
| cluster3194 | -2.27 | 0.392 |
| cluster3419 | -1.99 | 0.392 |
| cluster3231 | -2.47 | 0.392 |
| cluster3099 | -2.01 | 0.392 |
| cluster3026 | -2.37 | 0.401 |
| cluster3616 | 4.68 | 0.431 |
| cluster3821 | -2.07 | 0.434 |
| cluster3529 | -2.16 | 0.578 |
| cluster3374 | 4.05 | 0.672 |
| cluster3547 | 2.62 | 0.953 |
| cluster3004 | -1.83 | 0.966 |
| cluster3891 | 3.16 | 0.966 |
| cluster3379 | 2.73 | 0.966 |
| cluster3639 | 4.47 | 0.966 |
| cluster3427 | -1.97 | 0.966 |
| cluster4101 | 5.23 | 0.966 |
| cluster3858 | 3.15 | 0.966 |
| cluster3822 | 4.08 | 0.966 |
| cluster3760 | 2.77 | 0.966 |
| cluster2925 | -1.81 | 0.966 |
| cluster2885 | -1.74 | 0.966 |
| cluster3782 | -1.67 | 0.966 |
| cluster3802 | 4.02 | 0.966 |
| cluster3108 | -1.71 | 0.966 |
| cluster3128 | -1.50 | 0.966 |
| cluster3319 | -1.74 | 0.966 |
| cluster3567 | -2.29 | 0.966 |
| cluster3428 | -1.62 | 0.966 |
| cluster3607 | -1.74 | 0.966 |
| cluster4078 | 2.88 | 0.996 |
| cluster4116 | 5.01 | 0.996 |
| cluster4081 | 2.15 | 0.999 |
| cluster3689 | 4.26 | 1.000 |
| cluster3732 | 2.17 | 1.000 |
| cluster4127 | 4.23 | 1.000 |
| cluster3118 | -1.50 | 1.000 |

| | | |
|---|---|---|
| cluster3195 | -1.48 | 1.000 |
| cluster3301 | 2.43 | 1.000 |
| cluster4074 | 3.57 | 1.000 |
| cluster3652 | 2.65 | 1.000 |
| cluster3956 | 2.12 | 1.000 |
| cluster3309 | -1.69 | 1.000 |
| cluster3780 | 4.25 | 1.000 |
| cluster3950 | 2.05 | 1.000 |
| cluster3420 | -1.42 | 1.000 |
| cluster3146 | -1.45 | 1.000 |
| cluster2924 | -1.56 | 1.000 |
| cluster3926 | 3.53 | 1.000 |
| cluster3971 | 2.49 | 1.000 |
| cluster3810 | 2.09 | 1.000 |
| cluster4040 | 2.14 | 1.000 |
| cluster4087 | 3.87 | 1.000 |
| cluster3024 | -1.58 | 1.000 |
| cluster3715 | 2.74 | 1.000 |
| cluster3123 | -1.34 | 1.000 |
| cluster3704 | 2.32 | 1.000 |
| cluster4010 | 2.68 | 1.000 |
| cluster3461 | 3.24 | 1.000 |
| cluster3546 | 2.99 | 1.000 |
| cluster3487 | 2.21 | 1.000 |
| cluster2961 | 2.17 | 1.000 |
| cluster3889 | 3.02 | 1.000 |
| cluster3235 | 2.89 | 1.000 |
| cluster3037 | -1.35 | 1.000 |
| cluster4058 | 1.89 | 1.000 |
| cluster3930 | 1.99 | 1.000 |
| cluster4108 | 2.73 | 1.000 |
| cluster3167 | -1.63 | 1.000 |
| cluster3046 | -1.31 | 1.000 |
| cluster3842 | -1.53 | 1.000 |
| cluster3901 | 2.55 | 1.000 |
| cluster3151 | -1.21 | 1.000 |
| cluster3100 | -1.73 | 1.000 |
| cluster3778 | 2.20 | 1.000 |
| cluster4055 | 1.95 | 1.000 |
| cluster2858 | -1.27 | 1.000 |
| cluster3103 | 2.52 | 1.000 |
| cluster2902 | 2.37 | 1.000 |
| cluster3746 | 2.07 | 1.000 |
| cluster3283 | -1.15 | 1.000 |

| | | |
|---|---|---|
| cluster2994 | 1.66 | 1.000 |
| cluster3112 | -1.52 | 1.000 |
| cluster3978 | 1.62 | 1.000 |
| cluster3779 | 1.91 | 1.000 |
| cluster3892 | 2.26 | 1.000 |
| cluster2894 | -1.54 | 1.000 |
| cluster3785 | -1.36 | 1.000 |
| cluster2934 | -1.13 | 1.000 |
| cluster3852 | 2.31 | 1.000 |
| cluster3545 | 1.79 | 1.000 |
| cluster3818 | 2.35 | 1.000 |
| cluster3090 | -1.68 | 1.000 |
| cluster3240 | 1.94 | 1.000 |
| cluster4031 | 2.07 | 1.000 |
| cluster2904 | -1.44 | 1.000 |
| cluster3504 | 2.48 | 1.000 |
| cluster3670 | 2.20 | 1.000 |
| cluster4032 | -1.25 | 1.000 |
| cluster3399 | -1.39 | 1.000 |
| cluster3827 | 2.33 | 1.000 |
| cluster3250 | -1.51 | 1.000 |
| cluster4098 | 2.48 | 1.000 |
| cluster3738 | 2.86 | 1.000 |
| cluster3838 | -1.30 | 1.000 |
| cluster3713 | 1.60 | 1.000 |
| cluster3554 | -1.17 | 1.000 |
| cluster4125 | 3.94 | 1.000 |
| cluster3356 | 1.76 | 1.000 |
| cluster3314 | -1.36 | 1.000 |
| cluster3709 | -1.20 | 1.000 |
| cluster3972 | 1.61 | 1.000 |
| cluster2957 | -1.33 | 1.000 |
| cluster3770 | 3.58 | 1.000 |
| cluster3187 | -1.52 | 1.000 |
| cluster3812 | -1.37 | 1.000 |
| cluster3005 | -1.08 | 1.000 |
| cluster4119 | 2.87 | 1.000 |
| cluster4009 | -1.29 | 1.000 |
| cluster3909 | 1.59 | 1.000 |
| cluster3890 | 2.26 | 1.000 |
| cluster3297 | 2.15 | 1.000 |
| cluster3008 | 1.62 | 1.000 |
| cluster3854 | 1.50 | 1.000 |
| cluster3899 | 1.73 | 1.000 |

| | | |
|---|---|---|
| cluster3175 | 2.06 | 1.000 |
| cluster4002 | 2.07 | 1.000 |
| cluster3402 | -1.18 | 1.000 |
| cluster2982 | -1.15 | 1.000 |
| cluster3843 | 1.43 | 1.000 |
| cluster3609 | -1.57 | 1.000 |
| cluster3561 | -1.28 | 1.000 |
| cluster3559 | -1.08 | 1.000 |
| cluster3949 | -1.21 | 1.000 |
| cluster3106 | -1.08 | 1.000 |
| cluster3741 | -1.24 | 1.000 |
| cluster3556 | 1.92 | 1.000 |
| cluster3315 | 1.61 | 1.000 |
| cluster3848 | -1.22 | 1.000 |
| cluster3302 | -1.06 | 1.000 |
| cluster3619 | 1.55 | 1.000 |
| cluster3568 | -1.36 | 1.000 |
| cluster3851 | 1.93 | 1.000 |
| cluster3853 | 1.92 | 1.000 |
| cluster3476 | 2.08 | 1.000 |
| cluster3439 | -1.05 | 1.000 |
| cluster2901 | -1.18 | 1.000 |
| cluster3787 | 2.00 | 1.000 |
| cluster3164 | -1.36 | 1.000 |
| cluster4057 | 2.08 | 1.000 |
| cluster3329 | 2.27 | 1.000 |
| cluster3182 | 1.78 | 1.000 |
| cluster3160 | 1.48 | 1.000 |
| cluster4118 | -1.75 | 1.000 |
| cluster3256 | 1.40 | 1.000 |
| cluster2878 | -1.18 | 1.000 |
| cluster3408 | -1.22 | 1.000 |
| cluster3212 | 1.86 | 1.000 |
| cluster3386 | 1.69 | 1.000 |
| cluster3871 | 1.59 | 1.000 |
| cluster3707 | 2.04 | 1.000 |
| cluster3585 | -0.99 | 1.000 |
| cluster3562 | -1.12 | 1.000 |
| cluster3179 | 1.91 | 1.000 |
| cluster3983 | 2.04 | 1.000 |
| cluster4016 | 1.84 | 1.000 |
| cluster4120 | -1.46 | 1.000 |
| cluster3936 | 1.83 | 1.000 |
| cluster3077 | 1.73 | 1.000 |

| | | |
|---|---|---|
| cluster3259 | -0.99 | 1.000 |
| cluster4111 | -1.46 | 1.000 |
| cluster3666 | 2.35 | 1.000 |
| cluster3755 | 2.87 | 1.000 |
| cluster3957 | 1.65 | 1.000 |
| cluster3177 | 1.92 | 1.000 |
| cluster4022 | 1.42 | 1.000 |
| cluster3101 | 1.48 | 1.000 |
| cluster4095 | 1.97 | 1.000 |
| cluster3501 | 1.91 | 1.000 |
| cluster3866 | 2.25 | 1.000 |
| cluster2959 | -0.95 | 1.000 |
| cluster2893 | 1.98 | 1.000 |
| cluster3405 | -0.97 | 1.000 |
| cluster3286 | -0.93 | 1.000 |
| cluster4079 | 1.67 | 1.000 |
| cluster3833 | -1.21 | 1.000 |
| cluster3805 | 1.57 | 1.000 |
| cluster4042 | 1.74 | 1.000 |
| cluster2918 | -0.97 | 1.000 |
| cluster3015 | -1.16 | 1.000 |
| cluster4121 | 1.72 | 1.000 |
| cluster3474 | 1.92 | 1.000 |
| cluster3406 | 1.36 | 1.000 |
| cluster3516 | 2.06 | 1.000 |
| cluster3019 | 1.30 | 1.000 |
| cluster3224 | -0.93 | 1.000 |
| cluster3368 | -0.94 | 1.000 |
| cluster3065 | -0.99 | 1.000 |
| cluster4001 | 1.64 | 1.000 |
| cluster3865 | 2.15 | 1.000 |
| cluster4023 | -1.12 | 1.000 |
| cluster4104 | 1.93 | 1.000 |
| cluster2962 | 1.67 | 1.000 |
| cluster3343 | -0.93 | 1.000 |
| cluster3728 | 1.42 | 1.000 |
| cluster4030 | 1.56 | 1.000 |
| cluster3417 | 1.68 | 1.000 |
| cluster3571 | 1.87 | 1.000 |
| cluster3413 | 1.67 | 1.000 |
| cluster3611 | 2.02 | 1.000 |
| cluster3064 | -1.07 | 1.000 |
| cluster3191 | 2.25 | 1.000 |
| cluster3272 | -1.62 | 1.000 |

| | | |
|---|---|---|
| cluster2872 | -0.89 | 1.000 |
| cluster3855 | 1.68 | 1.000 |
| cluster3583 | -0.86 | 1.000 |
| cluster3022 | -1.01 | 1.000 |
| cluster3328 | -0.84 | 1.000 |
| cluster3658 | -0.97 | 1.000 |
| cluster3359 | 1.53 | 1.000 |
| cluster3276 | -0.89 | 1.000 |
| cluster2973 | -1.00 | 1.000 |
| cluster3316 | -1.02 | 1.000 |
| cluster3748 | 1.38 | 1.000 |
| cluster3631 | 2.00 | 1.000 |
| cluster2926 | 1.38 | 1.000 |
| cluster4048 | 2.14 | 1.000 |
| cluster3534 | -1.06 | 1.000 |
| cluster3904 | 1.54 | 1.000 |
| cluster3600 | -0.83 | 1.000 |
| cluster4003 | -1.15 | 1.000 |
| cluster3942 | -0.90 | 1.000 |
| cluster3525 | -0.87 | 1.000 |
| cluster3342 | 1.44 | 1.000 |
| cluster3377 | -0.94 | 1.000 |
| cluster4113 | 1.32 | 1.000 |
| cluster3558 | -0.94 | 1.000 |
| cluster3924 | 1.60 | 1.000 |
| cluster3482 | 1.37 | 1.000 |
| cluster3263 | -0.81 | 1.000 |
| cluster3233 | 2.34 | 1.000 |
| cluster3166 | 1.21 | 1.000 |
| cluster3702 | -0.92 | 1.000 |
| cluster2977 | -0.87 | 1.000 |
| cluster3505 | -0.85 | 1.000 |
| cluster4069 | 1.40 | 1.000 |
| cluster4062 | -1.24 | 1.000 |
| cluster3189 | -0.96 | 1.000 |
| cluster3135 | -0.95 | 1.000 |
| cluster3038 | -0.97 | 1.000 |
| cluster3693 | 1.90 | 1.000 |
| cluster3578 | -0.94 | 1.000 |
| cluster3790 | -1.12 | 1.000 |
| cluster3906 | 1.30 | 1.000 |
| cluster2908 | -0.90 | 1.000 |
| cluster3506 | 1.06 | 1.000 |
| cluster3989 | -0.99 | 1.000 |

| | | |
|---|---|---|
| cluster3207 | 1.33 | 1.000 |
| cluster3280 | -0.81 | 1.000 |
| cluster3976 | 1.27 | 1.000 |
| cluster3758 | 1.39 | 1.000 |
| cluster3800 | 1.50 | 1.000 |
| cluster2912 | 1.17 | 1.000 |
| cluster3648 | -0.85 | 1.000 |
| cluster3109 | 1.33 | 1.000 |
| cluster3723 | 1.09 | 1.000 |
| cluster3717 | 1.16 | 1.000 |
| cluster3333 | 1.20 | 1.000 |
| cluster3357 | -0.83 | 1.000 |
| cluster3804 | 1.30 | 1.000 |
| cluster3681 | 1.40 | 1.000 |
| cluster3410 | 1.52 | 1.000 |
| cluster3186 | 1.44 | 1.000 |
| cluster3933 | -0.91 | 1.000 |
| cluster3448 | 1.79 | 1.000 |
| cluster3070 | -0.84 | 1.000 |
| cluster3877 | 1.12 | 1.000 |
| cluster3678 | 1.13 | 1.000 |
| cluster3020 | -0.84 | 1.000 |
| cluster3113 | -0.76 | 1.000 |
| cluster3824 | 1.59 | 1.000 |
| cluster3260 | 1.15 | 1.000 |
| cluster3688 | 1.12 | 1.000 |
| cluster3364 | 1.17 | 1.000 |
| cluster4090 | -1.24 | 1.000 |
| cluster4117 | -1.28 | 1.000 |
| cluster3306 | 1.68 | 1.000 |
| cluster3575 | -0.95 | 1.000 |
| cluster4086 | 1.56 | 1.000 |
| cluster3222 | 0.95 | 1.000 |
| cluster3063 | 1.37 | 1.000 |
| cluster3178 | 1.61 | 1.000 |
| cluster3348 | -0.87 | 1.000 |
| cluster3897 | 1.41 | 1.000 |
| cluster2933 | -0.91 | 1.000 |
| cluster3927 | 1.13 | 1.000 |
| cluster3443 | 1.29 | 1.000 |
| cluster3303 | 1.75 | 1.000 |
| cluster2997 | -0.78 | 1.000 |
| cluster3192 | 1.11 | 1.000 |
| cluster3618 | -1.16 | 1.000 |

| | | |
|---|---|---|
| cluster2860 | 1.32 | 1.000 |
| cluster4107 | 1.71 | 1.000 |
| cluster3774 | -0.81 | 1.000 |
| cluster4039 | -0.87 | 1.000 |
| cluster3245 | -0.93 | 1.000 |
| cluster3494 | 1.27 | 1.000 |
| cluster3500 | 1.36 | 1.000 |
| cluster3902 | 1.10 | 1.000 |
| cluster3952 | 1.11 | 1.000 |
| cluster3886 | 1.19 | 1.000 |
| cluster3454 | 1.06 | 1.000 |
| cluster3793 | 1.00 | 1.000 |
| cluster3156 | 1.15 | 1.000 |
| cluster3036 | -0.77 | 1.000 |
| cluster3769 | -0.73 | 1.000 |
| cluster3451 | -1.26 | 1.000 |
| cluster4068 | 1.49 | 1.000 |
| cluster3305 | -0.87 | 1.000 |
| cluster2928 | 1.14 | 1.000 |
| cluster3068 | -0.71 | 1.000 |
| cluster3604 | 1.50 | 1.000 |
| cluster3940 | 1.15 | 1.000 |
| cluster4076 | 1.76 | 1.000 |
| cluster3387 | 1.05 | 1.000 |
| cluster3944 | 1.11 | 1.000 |
| cluster3913 | -0.86 | 1.000 |
| cluster3117 | 1.59 | 1.000 |
| cluster3938 | 1.36 | 1.000 |
| cluster3513 | -0.82 | 1.000 |
| cluster3883 | -1.03 | 1.000 |
| cluster3385 | 1.37 | 1.000 |
| cluster3274 | 1.22 | 1.000 |
| cluster3497 | 1.12 | 1.000 |
| cluster3543 | 1.47 | 1.000 |
| cluster3550 | -0.81 | 1.000 |
| cluster3323 | 1.38 | 1.000 |
| cluster3754 | 1.03 | 1.000 |
| cluster3502 | 1.50 | 1.000 |
| cluster3629 | 1.30 | 1.000 |
| cluster4045 | 1.60 | 1.000 |
| cluster3219 | 0.86 | 1.000 |
| cluster3299 | 1.21 | 1.000 |
| cluster3378 | 1.21 | 1.000 |
| cluster2956 | -0.73 | 1.000 |

| | | |
|---|---|---|
| cluster3834 | 1.31 | 1.000 |
| cluster3346 | 1.23 | 1.000 |
| cluster3089 | 1.13 | 1.000 |
| cluster3367 | 1.58 | 1.000 |
| cluster3041 | 1.03 | 1.000 |
| cluster2995 | 1.15 | 1.000 |
| cluster3129 | -1.05 | 1.000 |
| cluster3839 | -0.92 | 1.000 |
| cluster3266 | -0.70 | 1.000 |
| cluster3752 | 1.66 | 1.000 |
| cluster3349 | 1.21 | 1.000 |
| cluster3840 | 1.30 | 1.000 |
| cluster3987 | 1.18 | 1.000 |
| cluster3905 | 2.27 | 1.000 |
| cluster3282 | -0.68 | 1.000 |
| cluster3185 | 1.28 | 1.000 |
| cluster2990 | -0.66 | 1.000 |
| cluster3984 | -1.00 | 1.000 |
| cluster3114 | -0.66 | 1.000 |
| cluster3736 | -0.83 | 1.000 |
| cluster3362 | 1.06 | 1.000 |
| cluster3864 | 0.94 | 1.000 |
| cluster2949 | -0.66 | 1.000 |
| cluster3467 | 0.96 | 1.000 |
| cluster3449 | 1.03 | 1.000 |
| cluster3050 | -0.67 | 1.000 |
| cluster2935 | 1.06 | 1.000 |
| cluster3620 | 1.07 | 1.000 |
| cluster4038 | 1.38 | 1.000 |
| cluster3698 | -0.71 | 1.000 |
| cluster3246 | 1.04 | 1.000 |
| cluster3718 | 1.27 | 1.000 |
| cluster4051 | 1.31 | 1.000 |
| cluster3849 | 1.02 | 1.000 |
| cluster3085 | -0.77 | 1.000 |
| cluster4054 | 1.30 | 1.000 |
| cluster3985 | 1.06 | 1.000 |
| cluster3965 | 1.33 | 1.000 |
| cluster3703 | 0.91 | 1.000 |
| cluster3300 | -0.72 | 1.000 |
| cluster4014 | 1.14 | 1.000 |
| cluster3932 | 1.64 | 1.000 |
| cluster2905 | -0.71 | 1.000 |
| cluster3060 | 1.25 | 1.000 |

| | | |
|---|---|---|
| cluster3269 | -0.85 | 1.000 |
| cluster2987 | -0.63 | 1.000 |
| cluster3762 | -0.80 | 1.000 |
| cluster2876 | 1.53 | 1.000 |
| cluster3483 | -0.62 | 1.000 |
| cluster3749 | 1.39 | 1.000 |
| cluster3645 | 2.13 | 1.000 |
| cluster2897 | -0.67 | 1.000 |
| cluster2910 | -0.62 | 1.000 |
| cluster3121 | -0.74 | 1.000 |
| cluster2965 | -0.64 | 1.000 |
| cluster3171 | -0.76 | 1.000 |
| cluster3797 | -0.72 | 1.000 |
| cluster4112 | 1.52 | 1.000 |
| cluster3912 | 0.77 | 1.000 |
| cluster3649 | -0.61 | 1.000 |
| cluster3993 | 0.90 | 1.000 |
| cluster3321 | -0.61 | 1.000 |
| cluster2953 | -0.63 | 1.000 |
| cluster3941 | -0.63 | 1.000 |
| cluster3236 | 1.01 | 1.000 |
| cluster3257 | 0.85 | 1.000 |
| cluster3252 | 1.61 | 1.000 |
| cluster3884 | 1.25 | 1.000 |
| cluster3124 | -0.58 | 1.000 |
| cluster3211 | 1.26 | 1.000 |
| cluster4052 | 1.27 | 1.000 |
| cluster3075 | 0.98 | 1.000 |
| cluster3657 | 0.92 | 1.000 |
| cluster3471 | 1.03 | 1.000 |
| cluster2867 | 1.34 | 1.000 |
| cluster3651 | 1.70 | 1.000 |
| cluster3296 | 1.11 | 1.000 |
| cluster3397 | -0.77 | 1.000 |
| cluster3916 | 0.90 | 1.000 |
| cluster3078 | 0.77 | 1.000 |
| cluster3710 | 1.03 | 1.000 |
| cluster3512 | 1.09 | 1.000 |
| cluster3794 | 1.04 | 1.000 |
| cluster3830 | 0.90 | 1.000 |
| cluster2919 | -0.63 | 1.000 |
| cluster3511 | 1.08 | 1.000 |
| cluster4060 | 1.25 | 1.000 |
| cluster2906 | -0.70 | 1.000 |

| | | |
|---|---|---|
| cluster3735 | 1.04 | 1.000 |
| cluster3310 | -0.58 | 1.000 |
| cluster3874 | -0.62 | 1.000 |
| cluster4044 | 1.02 | 1.000 |
| cluster3593 | 1.06 | 1.000 |
| cluster3831 | 0.99 | 1.000 |
| cluster4049 | -1.03 | 1.000 |
| cluster3241 | -0.59 | 1.000 |
| cluster3981 | 1.19 | 1.000 |
| cluster3249 | -0.60 | 1.000 |
| cluster3743 | 0.79 | 1.000 |
| cluster3012 | 1.20 | 1.000 |
| cluster3663 | 1.06 | 1.000 |
| cluster3243 | -0.66 | 1.000 |
| cluster3198 | 0.94 | 1.000 |
| cluster3719 | 1.16 | 1.000 |
| cluster2923 | -0.59 | 1.000 |
| cluster3277 | -0.70 | 1.000 |
| cluster3765 | 0.93 | 1.000 |
| cluster3480 | 0.96 | 1.000 |
| cluster3998 | 1.36 | 1.000 |
| cluster3380 | -0.59 | 1.000 |
| cluster3955 | 1.31 | 1.000 |
| cluster3686 | 1.22 | 1.000 |
| cluster3010 | -0.62 | 1.000 |
| cluster3326 | -0.66 | 1.000 |
| cluster3061 | 0.95 | 1.000 |
| cluster4066 | 1.05 | 1.000 |
| cluster3540 | -0.74 | 1.000 |
| cluster3966 | 0.78 | 1.000 |
| cluster3031 | -0.61 | 1.000 |
| cluster4075 | -0.83 | 1.000 |
| cluster3701 | 0.85 | 1.000 |
| cluster3517 | 0.73 | 1.000 |
| cluster3654 | 0.83 | 1.000 |
| cluster3477 | -0.67 | 1.000 |
| cluster3199 | -0.55 | 1.000 |
| cluster4024 | 0.95 | 1.000 |
| cluster2863 | 0.77 | 1.000 |
| cluster3200 | 0.94 | 1.000 |
| cluster3789 | -0.72 | 1.000 |
| cluster3431 | -0.61 | 1.000 |
| cluster2941 | -0.55 | 1.000 |
| cluster3239 | -0.56 | 1.000 |

| | | |
|---|---|---|
| cluster3908 | 0.80 | 1.000 |
| cluster3391 | 0.93 | 1.000 |
| cluster3962 | -0.54 | 1.000 |
| cluster4006 | -0.56 | 1.000 |
| cluster3716 | -0.73 | 1.000 |
| cluster3355 | -0.67 | 1.000 |
| cluster3411 | 0.74 | 1.000 |
| cluster4092 | -0.71 | 1.000 |
| cluster4071 | 1.33 | 1.000 |
| cluster3783 | 1.22 | 1.000 |
| cluster4027 | 1.15 | 1.000 |
| cluster3052 | -0.74 | 1.000 |
| cluster3856 | 0.83 | 1.000 |
| cluster3291 | 0.88 | 1.000 |
| cluster3251 | -0.54 | 1.000 |
| cluster3072 | -0.65 | 1.000 |
| cluster2871 | -0.52 | 1.000 |
| cluster3422 | -0.77 | 1.000 |
| cluster3915 | 1.31 | 1.000 |
| cluster3777 | -0.60 | 1.000 |
| cluster3392 | 1.20 | 1.000 |
| cluster3659 | 1.04 | 1.000 |
| cluster3481 | 1.27 | 1.000 |
| cluster3415 | -0.58 | 1.000 |
| cluster4033 | 1.31 | 1.000 |
| cluster3677 | 0.83 | 1.000 |
| cluster3363 | 0.82 | 1.000 |
| cluster3131 | 0.83 | 1.000 |
| cluster4096 | 0.76 | 1.000 |
| cluster3819 | 1.24 | 1.000 |
| cluster3478 | -0.58 | 1.000 |
| cluster3215 | -0.51 | 1.000 |
| cluster4099 | 0.89 | 1.000 |
| cluster3731 | 0.98 | 1.000 |
| cluster3425 | -0.58 | 1.000 |
| cluster3155 | -0.63 | 1.000 |
| cluster3773 | 0.82 | 1.000 |
| cluster3671 | 0.81 | 1.000 |
| cluster3964 | -0.71 | 1.000 |
| cluster2874 | -0.49 | 1.000 |
| cluster3336 | -0.61 | 1.000 |
| cluster3144 | 1.23 | 1.000 |
| cluster3685 | -0.63 | 1.000 |
| cluster4053 | 0.90 | 1.000 |

| | | |
|---|---|---|
| cluster3221 | -0.59 | 1.000 |
| cluster4046 | 1.41 | 1.000 |
| cluster3613 | -0.60 | 1.000 |
| cluster3668 | 0.67 | 1.000 |
| cluster3176 | -0.63 | 1.000 |
| cluster3404 | 0.80 | 1.000 |
| cluster3279 | 0.83 | 1.000 |
| cluster3287 | -0.61 | 1.000 |
| cluster3313 | 0.88 | 1.000 |
| cluster3533 | 0.87 | 1.000 |
| cluster4084 | 0.90 | 1.000 |
| cluster3772 | -0.57 | 1.000 |
| cluster3229 | -0.63 | 1.000 |
| cluster3509 | -0.49 | 1.000 |
| cluster3044 | -0.62 | 1.000 |
| cluster3647 | 0.77 | 1.000 |
| cluster3150 | -0.64 | 1.000 |
| cluster3784 | 0.86 | 1.000 |
| cluster3823 | 0.85 | 1.000 |
| cluster4050 | 0.95 | 1.000 |
| cluster4105 | 0.92 | 1.000 |
| cluster3098 | 0.65 | 1.000 |
| cluster3350 | -0.54 | 1.000 |
| cluster3653 | 0.91 | 1.000 |
| cluster3360 | 0.87 | 1.000 |
| cluster4013 | 1.89 | 1.000 |
| cluster2992 | -0.53 | 1.000 |
| cluster3682 | -0.64 | 1.000 |
| cluster3227 | -0.53 | 1.000 |
| cluster3137 | -0.62 | 1.000 |
| cluster4035 | 0.91 | 1.000 |
| cluster3447 | 0.87 | 1.000 |
| cluster3515 | -0.65 | 1.000 |
| cluster3455 | -0.60 | 1.000 |
| cluster3273 | -0.49 | 1.000 |
| cluster3747 | 0.74 | 1.000 |
| cluster3053 | 1.04 | 1.000 |
| cluster3991 | 1.01 | 1.000 |
| cluster3196 | -0.48 | 1.000 |
| cluster3433 | -0.54 | 1.000 |
| cluster3029 | -0.46 | 1.000 |
| cluster3572 | 1.45 | 1.000 |
| cluster3479 | 1.04 | 1.000 |
| cluster3293 | 0.70 | 1.000 |

| | | |
|---|---|---|
| cluster3159 | -0.44 | 1.000 |
| cluster3153 | -0.47 | 1.000 |
| cluster2892 | 0.72 | 1.000 |
| cluster3307 | -0.49 | 1.000 |
| cluster4100 | 1.00 | 1.000 |
| cluster3337 | 0.89 | 1.000 |
| cluster3002 | -0.47 | 1.000 |
| cluster3744 | -0.49 | 1.000 |
| cluster4018 | -0.55 | 1.000 |
| cluster3937 | 0.88 | 1.000 |
| cluster3271 | 0.66 | 1.000 |
| cluster3808 | -0.57 | 1.000 |
| cluster3107 | 0.67 | 1.000 |
| cluster3193 | -0.47 | 1.000 |
| cluster3488 | -0.46 | 1.000 |
| cluster3460 | -0.52 | 1.000 |
| cluster3394 | -0.46 | 1.000 |
| cluster3576 | -0.50 | 1.000 |
| cluster3127 | 0.65 | 1.000 |
| cluster3016 | -0.62 | 1.000 |
| cluster3598 | 1.53 | 1.000 |
| cluster3612 | -0.56 | 1.000 |
| cluster3894 | 0.75 | 1.000 |
| cluster2981 | 0.68 | 1.000 |
| cluster2931 | -0.49 | 1.000 |
| cluster3587 | 1.16 | 1.000 |
| cluster3759 | -0.51 | 1.000 |
| cluster3163 | 1.01 | 1.000 |
| cluster2952 | -0.50 | 1.000 |
| cluster2950 | -0.46 | 1.000 |
| cluster4067 | 1.52 | 1.000 |
| cluster3979 | 0.71 | 1.000 |
| cluster3095 | 0.74 | 1.000 |
| cluster3003 | -0.50 | 1.000 |
| cluster3882 | 0.91 | 1.000 |
| cluster3393 | 1.16 | 1.000 |
| cluster3365 | 0.83 | 1.000 |
| cluster3614 | 1.10 | 1.000 |
| cluster3935 | -0.54 | 1.000 |
| cluster3521 | -0.45 | 1.000 |
| cluster3081 | 0.75 | 1.000 |
| cluster3929 | -0.50 | 1.000 |
| cluster3040 | -0.47 | 1.000 |
| cluster3042 | 0.77 | 1.000 |

| | | |
|---|---|---|
| cluster3032 | -0.46 | 1.000 |
| cluster2866 | -0.50 | 1.000 |
| cluster3860 | -0.48 | 1.000 |
| cluster3724 | 0.83 | 1.000 |
| cluster3623 | 0.68 | 1.000 |
| cluster2877 | -0.40 | 1.000 |
| cluster3115 | -0.41 | 1.000 |
| cluster3644 | 0.77 | 1.000 |
| cluster3083 | 0.64 | 1.000 |
| cluster3492 | -0.43 | 1.000 |
| cluster3023 | 0.71 | 1.000 |
| cluster3170 | -0.44 | 1.000 |
| cluster3880 | 1.22 | 1.000 |
| cluster3911 | -0.54 | 1.000 |
| cluster3244 | -0.44 | 1.000 |
| cluster2951 | -0.48 | 1.000 |
| cluster2983 | -0.40 | 1.000 |
| cluster3836 | 0.69 | 1.000 |
| cluster3142 | 0.83 | 1.000 |
| cluster3382 | 0.68 | 1.000 |
| cluster3992 | 0.79 | 1.000 |
| cluster3982 | 0.62 | 1.000 |
| cluster3594 | 0.80 | 1.000 |
| cluster3863 | 0.84 | 1.000 |
| cluster3646 | 1.00 | 1.000 |
| cluster3970 | 0.93 | 1.000 |
| cluster3875 | 0.66 | 1.000 |
| cluster3573 | -0.45 | 1.000 |
| cluster4102 | -0.45 | 1.000 |
| cluster3255 | -0.49 | 1.000 |
| cluster4061 | 0.84 | 1.000 |
| cluster3011 | 0.77 | 1.000 |
| cluster3069 | -0.40 | 1.000 |
| cluster3375 | 0.65 | 1.000 |
| cluster3136 | 0.74 | 1.000 |
| cluster3638 | 0.89 | 1.000 |
| cluster3071 | -0.43 | 1.000 |
| cluster3660 | -0.54 | 1.000 |
| cluster2988 | 0.55 | 1.000 |
| cluster3390 | -0.42 | 1.000 |
| cluster3903 | -0.38 | 1.000 |
| cluster3968 | 0.79 | 1.000 |
| cluster3174 | -0.46 | 1.000 |
| cluster3017 | 0.79 | 1.000 |

| | | |
|---|---|---|
| cluster3400 | -0.42 | 1.000 |
| cluster3726 | -0.40 | 1.000 |
| cluster3351 | 0.51 | 1.000 |
| cluster4083 | 0.90 | 1.000 |
| cluster3133 | -0.38 | 1.000 |
| cluster3496 | -0.42 | 1.000 |
| cluster4128 | 1.08 | 1.000 |
| cluster3522 | -0.45 | 1.000 |
| cluster3729 | -0.38 | 1.000 |
| cluster3541 | -0.41 | 1.000 |
| cluster3640 | 1.55 | 1.000 |
| cluster3088 | 0.68 | 1.000 |
| cluster2946 | -0.44 | 1.000 |
| cluster3537 | 0.64 | 1.000 |
| cluster3201 | -0.38 | 1.000 |
| cluster3473 | 0.78 | 1.000 |
| cluster4126 | 1.00 | 1.000 |
| cluster3139 | 1.02 | 1.000 |
| cluster2968 | -0.35 | 1.000 |
| cluster3861 | 0.55 | 1.000 |
| cluster3498 | 0.74 | 1.000 |
| cluster2937 | -0.39 | 1.000 |
| cluster3946 | -0.43 | 1.000 |
| cluster3125 | 0.55 | 1.000 |
| cluster2856 | -0.43 | 1.000 |
| cluster4080 | 1.02 | 1.000 |
| cluster2974 | -0.42 | 1.000 |
| cluster3147 | 0.55 | 1.000 |
| cluster3727 | -0.45 | 1.000 |
| cluster3237 | 0.87 | 1.000 |
| cluster2887 | 0.61 | 1.000 |
| cluster4015 | 0.70 | 1.000 |
| cluster3667 | 0.58 | 1.000 |
| cluster2895 | -0.38 | 1.000 |
| cluster2886 | 0.65 | 1.000 |
| cluster2855 | -0.35 | 1.000 |
| cluster3203 | 0.60 | 1.000 |
| cluster3079 | 0.65 | 1.000 |
| cluster3881 | 0.77 | 1.000 |
| cluster3158 | -0.43 | 1.000 |
| cluster3009 | 0.67 | 1.000 |
| cluster3555 | -0.46 | 1.000 |
| cluster4026 | 0.82 | 1.000 |
| cluster3781 | 0.67 | 1.000 |

| | | |
|---|---|---|
| cluster3409 | 0.80 | 1.000 |
| cluster3173 | 0.81 | 1.000 |
| cluster3995 | -0.44 | 1.000 |
| cluster2942 | -0.36 | 1.000 |
| cluster3606 | 0.79 | 1.000 |
| cluster2964 | -0.36 | 1.000 |
| cluster4109 | 0.81 | 1.000 |
| cluster3261 | -0.43 | 1.000 |
| cluster3205 | -0.36 | 1.000 |
| cluster3311 | 0.63 | 1.000 |
| cluster4005 | -0.41 | 1.000 |
| cluster3813 | 0.78 | 1.000 |
| cluster4029 | 0.96 | 1.000 |
| cluster3067 | -0.35 | 1.000 |
| cluster3563 | -0.37 | 1.000 |
| cluster4000 | 0.64 | 1.000 |
| cluster3603 | -0.37 | 1.000 |
| cluster3980 | 0.51 | 1.000 |
| cluster4017 | 0.75 | 1.000 |
| cluster3907 | -0.43 | 1.000 |
| cluster3564 | 0.80 | 1.000 |
| cluster3675 | -0.35 | 1.000 |
| cluster3951 | 0.77 | 1.000 |
| cluster3213 | 0.66 | 1.000 |
| cluster2966 | -0.37 | 1.000 |
| cluster3028 | -0.32 | 1.000 |
| cluster3661 | 0.55 | 1.000 |
| cluster3869 | 0.90 | 1.000 |
| cluster3463 | -0.37 | 1.000 |
| cluster3331 | -0.31 | 1.000 |
| cluster3294 | 0.57 | 1.000 |
| cluster3520 | -0.45 | 1.000 |
| cluster3366 | 0.79 | 1.000 |
| cluster3416 | 0.86 | 1.000 |
| cluster3893 | 0.51 | 1.000 |
| cluster3817 | 0.81 | 1.000 |
| cluster2984 | 0.70 | 1.000 |
| cluster3764 | 0.77 | 1.000 |
| cluster3105 | -0.40 | 1.000 |
| cluster3440 | -0.31 | 1.000 |
| cluster3055 | -0.31 | 1.000 |
| cluster3484 | -0.40 | 1.000 |
| cluster3591 | 0.54 | 1.000 |
| cluster3119 | 0.62 | 1.000 |

| | | |
|---|---|---|
| cluster3674 | 1.07 | 1.000 |
| cluster3921 | -0.40 | 1.000 |
| cluster3169 | -0.34 | 1.000 |
| cluster3592 | 0.53 | 1.000 |
| cluster3395 | 0.55 | 1.000 |
| cluster4088 | -0.37 | 1.000 |
| cluster2890 | -0.31 | 1.000 |
| cluster4085 | -0.37 | 1.000 |
| cluster3589 | -0.31 | 1.000 |
| cluster3820 | 0.46 | 1.000 |
| cluster3814 | -0.34 | 1.000 |
| cluster4007 | 0.49 | 1.000 |
| cluster3446 | -0.30 | 1.000 |
| cluster2880 | -0.32 | 1.000 |
| cluster3149 | 0.49 | 1.000 |
| cluster3637 | 0.45 | 1.000 |
| cluster3265 | 0.53 | 1.000 |
| cluster3033 | 0.47 | 1.000 |
| cluster3048 | 0.47 | 1.000 |
| cluster4037 | 0.50 | 1.000 |
| cluster3811 | 0.54 | 1.000 |
| cluster4106 | 0.88 | 1.000 |
| cluster3963 | 0.79 | 1.000 |
| cluster2991 | -0.29 | 1.000 |
| cluster3096 | 0.45 | 1.000 |
| cluster3617 | 0.74 | 1.000 |
| cluster3643 | -0.35 | 1.000 |
| cluster3345 | 0.53 | 1.000 |
| cluster3490 | 0.65 | 1.000 |
| cluster4073 | 0.94 | 1.000 |
| cluster3035 | -0.30 | 1.000 |
| cluster3197 | -0.32 | 1.000 |
| cluster3054 | -0.27 | 1.000 |
| cluster3376 | 0.45 | 1.000 |
| cluster3486 | 0.61 | 1.000 |
| cluster2932 | 0.50 | 1.000 |
| cluster2960 | 0.68 | 1.000 |
| cluster3134 | -0.28 | 1.000 |
| cluster3939 | 0.54 | 1.000 |
| cluster3850 | 0.58 | 1.000 |
| cluster3292 | -0.33 | 1.000 |
| cluster3130 | -0.29 | 1.000 |
| cluster3372 | 0.48 | 1.000 |
| cluster3953 | -0.29 | 1.000 |

| | | |
|---|---|---|
| cluster3383 | -0.36 | 1.000 |
| cluster3990 | -0.30 | 1.000 |
| cluster3080 | -0.29 | 1.000 |
| cluster3974 | 0.90 | 1.000 |
| cluster3601 | -0.31 | 1.000 |
| cluster3920 | 0.64 | 1.000 |
| cluster3403 | -0.27 | 1.000 |
| cluster3705 | 0.69 | 1.000 |
| cluster2921 | -0.29 | 1.000 |
| cluster3290 | -0.31 | 1.000 |
| cluster3684 | -0.30 | 1.000 |
| cluster3268 | 0.46 | 1.000 |
| cluster3458 | -0.27 | 1.000 |
| cluster4004 | -0.25 | 1.000 |
| cluster4124 | -0.41 | 1.000 |
| cluster3595 | 0.51 | 1.000 |
| cluster2989 | -0.30 | 1.000 |
| cluster3632 | 0.51 | 1.000 |
| cluster2920 | -0.26 | 1.000 |
| cluster3161 | -0.27 | 1.000 |
| cluster3225 | -0.31 | 1.000 |
| cluster2993 | 0.59 | 1.000 |
| cluster4008 | 0.66 | 1.000 |
| cluster3737 | -0.25 | 1.000 |
| cluster3809 | 0.44 | 1.000 |
| cluster3887 | 0.66 | 1.000 |
| cluster2938 | 0.43 | 1.000 |
| cluster3352 | -0.29 | 1.000 |
| cluster2922 | -0.28 | 1.000 |
| cluster3590 | -0.28 | 1.000 |
| cluster3896 | -0.30 | 1.000 |
| cluster3104 | -0.23 | 1.000 |
| cluster3945 | 0.62 | 1.000 |
| cluster2873 | 0.51 | 1.000 |
| cluster3472 | 0.48 | 1.000 |
| cluster3566 | -0.26 | 1.000 |
| cluster4103 | -0.32 | 1.000 |
| cluster3389 | 0.48 | 1.000 |
| cluster3650 | 0.63 | 1.000 |
| cluster3056 | -0.25 | 1.000 |
| cluster3401 | -0.24 | 1.000 |
| cluster3051 | -0.28 | 1.000 |
| cluster3948 | 0.45 | 1.000 |
| cluster3694 | 0.50 | 1.000 |

| | | |
|---|---|---|
| cluster3867 | 0.44 | 1.000 |
| cluster3369 | -0.24 | 1.000 |
| cluster3324 | -0.25 | 1.000 |
| cluster3570 | 0.69 | 1.000 |
| cluster4114 | -0.32 | 1.000 |
| cluster3579 | 0.47 | 1.000 |
| cluster3628 | 0.53 | 1.000 |
| cluster3960 | -0.30 | 1.000 |
| cluster3206 | -0.27 | 1.000 |
| cluster3531 | 0.49 | 1.000 |
| cluster2944 | -0.25 | 1.000 |
| cluster2967 | -0.27 | 1.000 |
| cluster3084 | -0.23 | 1.000 |
| cluster3025 | 0.39 | 1.000 |
| cluster2954 | -0.26 | 1.000 |
| cluster3034 | 0.44 | 1.000 |
| cluster3258 | -0.23 | 1.000 |
| cluster3503 | 0.44 | 1.000 |
| cluster3076 | 0.49 | 1.000 |
| cluster3597 | -0.26 | 1.000 |
| cluster3792 | -0.23 | 1.000 |
| cluster2888 | 0.75 | 1.000 |
| cluster3208 | -0.23 | 1.000 |
| cluster3708 | 0.75 | 1.000 |
| cluster3450 | 0.83 | 1.000 |
| cluster4077 | -0.25 | 1.000 |
| cluster3148 | -0.21 | 1.000 |
| cluster3485 | 0.75 | 1.000 |
| cluster3475 | -0.24 | 1.000 |
| cluster3791 | -0.25 | 1.000 |
| cluster3656 | 0.48 | 1.000 |
| cluster3519 | -0.27 | 1.000 |
| cluster3047 | -0.25 | 1.000 |
| cluster2914 | 0.53 | 1.000 |
| cluster3922 | 0.71 | 1.000 |
| cluster3934 | -0.21 | 1.000 |
| cluster3826 | 0.61 | 1.000 |
| cluster3775 | -0.22 | 1.000 |
| cluster3690 | -0.23 | 1.000 |
| cluster3958 | 0.72 | 1.000 |
| cluster4043 | -0.34 | 1.000 |
| cluster2861 | -0.24 | 1.000 |
| cluster4123 | 0.83 | 1.000 |
| cluster3977 | -0.25 | 1.000 |

| | | |
|---|---|---|
| cluster4064 | 0.50 | 1.000 |
| cluster3655 | -0.23 | 1.000 |
| cluster3536 | -0.21 | 1.000 |
| cluster3535 | -0.24 | 1.000 |
| cluster2913 | 0.37 | 1.000 |
| cluster3092 | 0.37 | 1.000 |
| cluster3642 | 0.50 | 1.000 |
| cluster4094 | -0.23 | 1.000 |
| cluster3262 | -0.21 | 1.000 |
| cluster3712 | 0.48 | 1.000 |
| cluster3622 | 0.38 | 1.000 |
| cluster2899 | 0.56 | 1.000 |
| cluster3045 | -0.22 | 1.000 |
| cluster3569 | 0.50 | 1.000 |
| cluster3873 | 0.39 | 1.000 |
| cluster3768 | -0.22 | 1.000 |
| cluster4036 | 0.39 | 1.000 |
| cluster3094 | 0.59 | 1.000 |
| cluster3672 | 0.48 | 1.000 |
| cluster3184 | -0.19 | 1.000 |
| cluster3000 | -0.21 | 1.000 |
| cluster3429 | 0.44 | 1.000 |
| cluster3013 | -0.20 | 1.000 |
| cluster2882 | -0.19 | 1.000 |
| cluster3756 | -0.20 | 1.000 |
| cluster3633 | 0.43 | 1.000 |
| cluster3456 | -0.20 | 1.000 |
| cluster2998 | 0.35 | 1.000 |
| cluster3468 | -0.21 | 1.000 |
| cluster2854 | 0.46 | 1.000 |
| cluster3669 | 0.39 | 1.000 |
| cluster3596 | -0.22 | 1.000 |
| cluster3795 | 0.38 | 1.000 |
| cluster3442 | -0.21 | 1.000 |
| cluster3097 | -0.20 | 1.000 |
| cluster3018 | 0.34 | 1.000 |
| cluster3138 | 0.47 | 1.000 |
| cluster4059 | 0.64 | 1.000 |
| cluster3353 | -0.17 | 1.000 |
| cluster3030 | -0.21 | 1.000 |
| cluster3928 | 0.39 | 1.000 |
| cluster3996 | -0.18 | 1.000 |
| cluster3714 | 0.53 | 1.000 |
| cluster3168 | 0.40 | 1.000 |

| | | |
|---|---|---|
| cluster3234 | 0.40 | 1.000 |
| cluster3761 | -0.20 | 1.000 |
| cluster3462 | 0.44 | 1.000 |
| cluster2915 | 0.42 | 1.000 |
| cluster3281 | -0.18 | 1.000 |
| cluster3625 | 0.36 | 1.000 |
| cluster3388 | 0.46 | 1.000 |
| cluster3254 | 0.54 | 1.000 |
| cluster3341 | -0.18 | 1.000 |
| cluster3470 | 0.32 | 1.000 |
| cluster3574 | 0.62 | 1.000 |
| cluster4122 | 0.72 | 1.000 |
| cluster3969 | 0.33 | 1.000 |
| cluster3771 | -0.17 | 1.000 |
| cluster3218 | 0.50 | 1.000 |
| cluster3722 | -0.18 | 1.000 |
| cluster3006 | 0.62 | 1.000 |
| cluster3226 | 0.45 | 1.000 |
| cluster2943 | -0.17 | 1.000 |
| cluster3091 | -0.16 | 1.000 |
| cluster3057 | -0.18 | 1.000 |
| cluster3610 | -0.18 | 1.000 |
| cluster3162 | 0.45 | 1.000 |
| cluster3339 | -0.17 | 1.000 |
| cluster3745 | -0.17 | 1.000 |
| cluster4089 | 0.60 | 1.000 |
| cluster3532 | 0.47 | 1.000 |
| cluster3815 | 0.57 | 1.000 |
| cluster3465 | 0.40 | 1.000 |
| cluster3706 | -0.15 | 1.000 |
| cluster3798 | -0.17 | 1.000 |
| cluster3599 | 0.48 | 1.000 |
| cluster3381 | 0.42 | 1.000 |
| cluster3441 | -0.15 | 1.000 |
| cluster2907 | -0.15 | 1.000 |
| cluster3524 | -0.15 | 1.000 |
| cluster3027 | 0.36 | 1.000 |
| cluster3014 | -0.16 | 1.000 |
| cluster3247 | -0.15 | 1.000 |
| cluster2970 | 0.28 | 1.000 |
| cluster3696 | 0.32 | 1.000 |
| cluster3538 | -0.15 | 1.000 |
| cluster3059 | 0.47 | 1.000 |
| cluster2996 | -0.15 | 1.000 |

| | | |
|---|---|---|
| cluster3204 | -0.15 | 1.000 |
| cluster3466 | -0.15 | 1.000 |
| cluster4072 | 0.32 | 1.000 |
| cluster3230 | -0.15 | 1.000 |
| cluster3621 | 0.44 | 1.000 |
| cluster3695 | 0.35 | 1.000 |
| cluster3073 | -0.14 | 1.000 |
| cluster3181 | 0.37 | 1.000 |
| cluster3238 | -0.14 | 1.000 |
| cluster3959 | -0.15 | 1.000 |
| cluster3086 | -0.14 | 1.000 |
| cluster3725 | 0.47 | 1.000 |
| cluster3327 | 0.40 | 1.000 |
| cluster4025 | -0.13 | 1.000 |
| cluster3132 | 0.33 | 1.000 |
| cluster3879 | 0.34 | 1.000 |
| cluster3358 | 0.39 | 1.000 |
| cluster3347 | 0.30 | 1.000 |
| cluster3807 | -0.14 | 1.000 |
| cluster3857 | -0.14 | 1.000 |
| cluster4020 | 0.32 | 1.000 |
| cluster3338 | 0.41 | 1.000 |
| cluster2985 | 0.36 | 1.000 |
| cluster3734 | 0.42 | 1.000 |
| cluster3549 | -0.13 | 1.000 |
| cluster3398 | 0.32 | 1.000 |
| cluster3560 | -0.14 | 1.000 |
| cluster3275 | 0.47 | 1.000 |
| cluster3214 | 0.34 | 1.000 |
| cluster2936 | 0.49 | 1.000 |
| cluster2969 | 0.38 | 1.000 |
| cluster3418 | 0.35 | 1.000 |
| cluster3841 | -0.13 | 1.000 |
| cluster2930 | 0.32 | 1.000 |
| cluster3495 | 0.31 | 1.000 |
| cluster3217 | -0.13 | 1.000 |
| cluster3508 | 0.50 | 1.000 |
| cluster3039 | 0.30 | 1.000 |
| cluster3021 | 0.29 | 1.000 |
| cluster3608 | 0.30 | 1.000 |
| cluster3188 | 0.41 | 1.000 |
| cluster3285 | -0.12 | 1.000 |
| cluster2898 | 0.37 | 1.000 |
| cluster3436 | -0.11 | 1.000 |

| | | |
|---|---|---|
| cluster3816 | -0.12 | 1.000 |
| cluster2853 | 0.40 | 1.000 |
| cluster3228 | -0.11 | 1.000 |
| cluster3796 | -0.12 | 1.000 |
| cluster2947 | 0.32 | 1.000 |
| cluster3739 | 0.28 | 1.000 |
| cluster3973 | -0.11 | 1.000 |
| cluster3788 | 0.35 | 1.000 |
| cluster2999 | 0.35 | 1.000 |
| cluster4082 | 0.52 | 1.000 |
| cluster2978 | 0.36 | 1.000 |
| cluster3370 | 0.34 | 1.000 |
| cluster3430 | -0.08 | 1.000 |
| cluster3507 | 0.35 | 1.000 |
| cluster3332 | 0.32 | 1.000 |
| cluster3126 | 0.45 | 1.000 |
| cluster3412 | 0.32 | 1.000 |
| cluster3862 | -0.09 | 1.000 |
| cluster3320 | -0.09 | 1.000 |
| cluster3318 | 0.36 | 1.000 |
| cluster3665 | 0.42 | 1.000 |
| cluster3145 | -0.09 | 1.000 |
| cluster3001 | -0.09 | 1.000 |
| cluster3058 | 0.24 | 1.000 |
| cluster3288 | 0.36 | 1.000 |
| cluster3043 | 0.28 | 1.000 |
| cluster3895 | -0.08 | 1.000 |
| cluster3544 | 0.50 | 1.000 |
| cluster3270 | 0.38 | 1.000 |
| cluster3528 | -0.08 | 1.000 |
| cluster3264 | 0.26 | 1.000 |
| cluster2859 | 0.23 | 1.000 |
| cluster3829 | -0.07 | 1.000 |
| cluster2891 | -0.08 | 1.000 |
| cluster3997 | 0.40 | 1.000 |
| cluster3803 | -0.06 | 1.000 |
| cluster2940 | -0.08 | 1.000 |
| cluster4063 | 0.48 | 1.000 |
| cluster3165 | 0.36 | 1.000 |
| cluster2865 | 0.27 | 1.000 |
| cluster3334 | 0.27 | 1.000 |
| cluster3730 | 0.30 | 1.000 |
| cluster3584 | 0.42 | 1.000 |
| cluster3720 | 0.37 | 1.000 |

| | | |
|---|---|---|
| cluster2917 | -0.07 | 1.000 |
| cluster3122 | 0.28 | 1.000 |
| cluster2889 | -0.07 | 1.000 |
| cluster3923 | 0.25 | 1.000 |
| cluster3806 | 0.34 | 1.000 |
| cluster3742 | -0.06 | 1.000 |
| cluster3354 | -0.07 | 1.000 |
| cluster3835 | -0.07 | 1.000 |
| cluster3074 | -0.07 | 1.000 |
| cluster3999 | 0.36 | 1.000 |
| cluster3322 | -0.06 | 1.000 |
| cluster3733 | 0.31 | 1.000 |
| cluster3832 | -0.07 | 1.000 |
| cluster2927 | 0.34 | 1.000 |
| cluster3888 | 0.37 | 1.000 |
| cluster3683 | 0.26 | 1.000 |
| cluster3530 | -0.07 | 1.000 |
| cluster3087 | 0.24 | 1.000 |
| cluster3954 | 0.42 | 1.000 |
| cluster3641 | -0.06 | 1.000 |
| cluster3298 | -0.06 | 1.000 |
| cluster3692 | 0.33 | 1.000 |
| cluster3691 | 0.30 | 1.000 |
| cluster3464 | 0.30 | 1.000 |
| cluster3900 | 0.00 | 1.000 |
| cluster3837 | -0.05 | 1.000 |
| cluster4097 | 0.03 | 1.000 |
| cluster3630 | 0.24 | 1.000 |
| cluster3210 | -0.06 | 1.000 |
| cluster3335 | 0.31 | 1.000 |
| cluster3577 | 0.26 | 1.000 |
| cluster3844 | 0.53 | 1.000 |
| cluster3093 | -0.05 | 1.000 |
| cluster3424 | 0.50 | 1.000 |
| cluster2929 | 0.21 | 1.000 |
| cluster3679 | 0.26 | 1.000 |
| cluster3432 | 0.28 | 1.000 |
| cluster4070 | 0.55 | 1.000 |
| cluster3986 | 0.23 | 1.000 |
| cluster2986 | -0.05 | 1.000 |
| cluster3526 | 0.21 | 1.000 |
| cluster3493 | -0.03 | 1.000 |
| cluster4021 | -0.03 | 1.000 |
| cluster4028 | -0.05 | 1.000 |

| | | |
|---|---|---|
| cluster3527 | -0.04 | 1.000 |
| cluster3551 | -0.03 | 1.000 |
| cluster3925 | 0.87 | 1.000 |
| cluster3825 | -0.04 | 1.000 |
| cluster3253 | -0.04 | 1.000 |
| cluster3680 | 0.24 | 1.000 |
| cluster3700 | -0.04 | 1.000 |
| cluster3776 | 0.30 | 1.000 |
| cluster3120 | -0.05 | 1.000 |
| cluster3801 | 0.34 | 1.000 |
| cluster3721 | 0.34 | 1.000 |
| cluster2903 | 0.30 | 1.000 |
| cluster3582 | -0.03 | 1.000 |
| cluster3586 | -0.03 | 1.000 |
| cluster2955 | -0.04 | 1.000 |
| cluster3373 | -0.03 | 1.000 |
| cluster4093 | 0.41 | 1.000 |
| cluster3898 | 0.32 | 1.000 |
| cluster3152 | 0.31 | 1.000 |
| cluster3453 | -0.01 | 1.000 |
| cluster3868 | -0.03 | 1.000 |
| cluster3845 | 0.57 | 1.000 |
| cluster3344 | -0.02 | 1.000 |
| cluster3007 | 0.21 | 1.000 |
| cluster3673 | 0.36 | 1.000 |
| cluster3588 | -0.01 | 1.000 |
| cluster4041 | 0.27 | 1.000 |
| cluster3223 | 0.27 | 1.000 |
| cluster3766 | 0.25 | 1.000 |
| cluster3157 | 0.00 | 1.000 |
| cluster2971 | 0.18 | 1.000 |
| cluster3110 | -0.02 | 1.000 |
| cluster2881 | 0.25 | 1.000 |
| cluster2979 | 0.17 | 1.000 |
| cluster3602 | 0.23 | 1.000 |
| cluster3202 | -0.01 | 1.000 |
| cluster3452 | 0.25 | 1.000 |
| cluster3172 | 0.24 | 1.000 |
| cluster3062 | 0.20 | 1.000 |
| cluster3847 | 0.21 | 1.000 |
| cluster3636 | -0.02 | 1.000 |
| cluster3444 | -0.01 | 1.000 |
| cluster3605 | 0.00 | 1.000 |
| cluster4034 | 0.12 | 1.000 |

| | | |
|---|---|---|
| cluster3918 | 0.27 | 1.000 |
| cluster3116 | -0.01 | 1.000 |
| cluster2911 | 0.00 | 1.000 |
| cluster3767 | 0.26 | 1.000 |
| cluster3361 | 0.19 | 1.000 |
| cluster3491 | 0.00 | 1.000 |
| cluster3523 | 0.28 | 1.000 |
| cluster3711 | 0.17 | 1.000 |
| cluster3557 | 0.20 | 1.000 |
| cluster3740 | 0.25 | 1.000 |
| cluster4115 | 0.35 | 1.000 |
| cluster3489 | 0.21 | 1.000 |
| cluster3308 | 0.00 | 1.000 |
| cluster3634 | 0.21 | 1.000 |
| cluster2862 | 0.18 | 1.000 |
| cluster3437 | 0.19 | 1.000 |
| cluster3396 | 0.20 | 1.000 |
| cluster2864 | 0.17 | 1.000 |
| cluster2875 | 0.03 | 1.000 |
| cluster3581 | 0.03 | 1.000 |
| cluster3846 | 0.17 | 1.000 |
| cluster3615 | 0.26 | 1.000 |
| cluster3664 | 0.17 | 1.000 |
| cluster3699 | 0.26 | 1.000 |
| cluster4047 | 0.37 | 1.000 |
| cluster3510 | 0.39 | 1.000 |
| cluster2963 | 0.23 | 1.000 |
| cluster4110 | 0.25 | 1.000 |
| cluster2857 | 0.16 | 1.000 |
| cluster3914 | 0.39 | 1.000 |
| cluster3947 | 0.16 | 1.000 |
| cluster4056 | 0.21 | 1.000 |
| cluster3190 | 0.30 | 1.000 |
| cluster3627 | 0.15 | 1.000 |
| cluster3154 | 0.14 | 1.000 |
| cluster3459 | 0.25 | 1.000 |
| cluster3082 | 0.12 | 1.000 |
| cluster3876 | 0.18 | 1.000 |
| cluster2948 | 0.26 | 1.000 |
| cluster3917 | 0.21 | 1.000 |
| cluster3967 | 0.20 | 1.000 |
| cluster3438 | 0.20 | 1.000 |
| cluster2958 | 0.02 | 1.000 |
| cluster2879 | 0.05 | 1.000 |

| | | |
|---|---|---|
| cluster3753 | 0.12 | 1.000 |
| cluster3141 | 0.04 | 1.000 |
| cluster3407 | 0.16 | 1.000 |
| cluster3421 | 0.21 | 1.000 |
| cluster3624 | 0.13 | 1.000 |
| cluster3220 | 0.07 | 1.000 |
| cluster2870 | 0.04 | 1.000 |
| cluster3209 | 0.05 | 1.000 |
| cluster3919 | 0.12 | 1.000 |
| cluster4019 | 0.11 | 1.000 |
| cluster3539 | 0.27 | 1.000 |
| cluster3312 | 0.27 | 1.000 |
| cluster2883 | 0.13 | 1.000 |
| cluster3426 | 0.17 | 1.000 |
| cluster3750 | 0.27 | 1.000 |
| cluster3278 | 0.04 | 1.000 |
| cluster3499 | 0.18 | 1.000 |
| cluster2980 | 0.24 | 1.000 |
| cluster3635 | 0.12 | 1.000 |
| cluster3988 | 0.11 | 1.000 |
| cluster3049 | 0.07 | 1.000 |
| cluster3143 | 0.13 | 1.000 |
| cluster2975 | 0.11 | 1.000 |
| cluster2896 | 0.05 | 1.000 |
| cluster2972 | 0.09 | 1.000 |
| cluster2884 | 0.05 | 1.000 |
| cluster3384 | 0.15 | 1.000 |
| cluster3066 | 0.08 | 1.000 |
| cluster2916 | 0.16 | 1.000 |
| cluster3469 | 0.17 | 1.000 |
| cluster3340 | 0.07 | 1.000 |
| cluster3242 | 0.13 | 1.000 |
| cluster3697 | 0.21 | 1.000 |
| cluster2945 | 0.17 | 1.000 |
| cluster3140 | 0.10 | 1.000 |
| cluster2900 | 0.17 | 1.000 |
| cluster2976 | 0.12 | 1.000 |
| cluster3289 | 0.12 | 1.000 |
| cluster4011 | 0.25 | 1.000 |
| cluster3552 | 0.11 | 1.000 |
| cluster3295 | 0.06 | 1.000 |
| cluster3325 | 0.28 | 1.000 |
| cluster3423 | 0.21 | 1.000 |
| cluster3931 | 0.19 | 1.000 |

| | | |
|---|---|---|
| cluster3284 | 0.10 | 1.000 |
| cluster3786 | 0.24 | 1.000 |
| cluster3548 | 0.10 | 1.000 |
| cluster3304 | 0.12 | 1.000 |
| cluster3514 | 0.15 | 1.000 |
| cluster3870 | 0.20 | 1.000 |
| cluster3445 | 0.08 | 1.000 |
| cluster2909 | 0.08 | 1.000 |
| cluster3542 | 0.17 | 1.000 |
| cluster3961 | 0.17 | 1.000 |
| cluster3518 | 0.17 | 1.000 |
| cluster3757 | 0.14 | 1.000 |
| cluster3676 | 0.13 | 1.000 |
| cluster3943 | 0.13 | 1.000 |
| cluster3317 | 0.13 | 1.000 |
| cluster3763 | 0.12 | 1.000 |
| cluster3828 | 0.12 | 1.000 |
| cluster3662 | 0.12 | 1.000 |
| cluster3799 | 0.12 | 1.000 |
| cluster3910 | 0.12 | 1.000 |
| cluster4012 | 0.11 | 1.000 |
| cluster3435 | 0.10 | 1.000 |
| cluster3565 | 0.09 | 1.000 |
| cluster3687 | 0.08 | 1.000 |
| cluster3751 | 0.08 | 1.000 |
| cluster1127 | 0.08 | 1.000 |
| cluster4065 | 0.08 | 1.000 |
| cluster3267 | 0.07 | 1.000 |
| cluster2869 | 0.07 | 1.000 |
| cluster3330 | 0.07 | 1.000 |
| cluster3371 | 0.07 | 1.000 |
| cluster3553 | 0.07 | 1.000 |
| cluster3580 | 0.06 | 1.000 |
| cluster3434 | 0.06 | 1.000 |
| cluster3626 | 0.06 | 1.000 |
| cluster3232 | 0.05 | 1.000 |
| cluster3216 | 0.05 | 1.000 |
| cluster3248 | 0.03 | 1.000 |
| cluster3102 | 0.03 | 1.000 |
| cluster3180 | 0.03 | 1.000 |
| cluster3994 | 0.03 | 1.000 |
| cluster2939 | 0.02 | 1.000 |
| cluster3414 | 0.02 | 1.000 |
| cluster3872 | 0.01 | 1.000 |

| | | |
|---|---|---|
| cluster3183 | 0.01 | 1.000 |

**Supplementary Table 5**. DESeq2 results with RSEM counts - Parasites vs Saprophagous

| ID | log2FoldChange | Adjusted p value |
|---|---|---|
| cluster3616 | 4.83 | 0.014 |
| cluster3547 | 2.62 | 0.081 |
| cluster3374 | 4.32 | 0.117 |
| cluster3822 | 4.52 | 0.117 |
| cluster3891 | 3.40 | 0.117 |
| cluster4101 | 5.05 | 0.117 |
| cluster3760 | 2.76 | 0.117 |
| cluster3639 | 4.66 | 0.128 |
| cluster3858 | 3.10 | 0.128 |
| cluster4074 | 4.45 | 0.173 |
| cluster3975 | 3.69 | 0.195 |
| cluster4127 | 4.85 | 0.195 |
| cluster3250 | -2.57 | 0.197 |
| cluster3457 | -3.53 | 0.197 |
| cluster3802 | 3.03 | 0.197 |
| cluster3689 | 4.33 | 0.206 |
| cluster4078 | 2.70 | 0.210 |
| cluster3301 | 2.50 | 0.239 |
| cluster4010 | 3.29 | 0.239 |
| cluster3652 | 2.70 | 0.285 |
| cluster4116 | 3.55 | 0.285 |
| cluster3778 | 2.46 | 0.295 |
| cluster3926 | 3.85 | 0.324 |
| cluster3950 | 2.20 | 0.324 |
| cluster2961 | 2.11 | 0.337 |
| cluster3379 | 2.88 | 0.337 |
| cluster3704 | 2.39 | 0.337 |
| cluster3709 | -1.65 | 0.337 |
| cluster3715 | 3.12 | 0.337 |
| cluster3810 | 2.14 | 0.337 |
| cluster3235 | 3.26 | 0.337 |
| cluster3889 | 3.29 | 0.344 |
| cluster4108 | 3.02 | 0.376 |
| cluster3030 | -1.58 | 0.533 |
| cluster3461 | 3.64 | 0.533 |
| cluster3746 | 2.05 | 0.533 |
| cluster4091 | -3.43 | 0.533 |
| cluster3004 | -1.80 | 0.548 |
| cluster3015 | -1.97 | 0.548 |
| cluster3099 | -1.83 | 0.548 |
| cluster3103 | 2.95 | 0.548 |
| cluster3194 | -1.97 | 0.548 |

| | | |
|---|---|---|
| cluster3240 | 1.97 | 0.548 |
| cluster3256 | 1.44 | 0.548 |
| cluster3504 | 2.46 | 0.548 |
| cluster3546 | 3.29 | 0.548 |
| cluster3567 | -2.74 | 0.548 |
| cluster3670 | 2.42 | 0.548 |
| cluster3707 | 2.74 | 0.548 |
| cluster3779 | 1.96 | 0.548 |
| cluster3780 | 4.62 | 0.548 |
| cluster3818 | 2.26 | 0.548 |
| cluster3827 | 2.55 | 0.548 |
| cluster3852 | 2.30 | 0.548 |
| cluster3930 | 2.05 | 0.548 |
| cluster3971 | 2.75 | 0.548 |
| cluster4040 | 2.31 | 0.548 |
| cluster4081 | 2.49 | 0.548 |
| cluster4098 | 2.43 | 0.548 |
| cluster4104 | 2.51 | 0.548 |
| cluster4119 | 3.06 | 0.548 |
| cluster3024 | -1.85 | 0.575 |
| cluster2902 | 2.82 | 0.593 |
| cluster3101 | 1.56 | 0.593 |
| cluster3297 | 2.14 | 0.593 |
| cluster3315 | 1.60 | 0.593 |
| cluster3329 | 2.58 | 0.593 |
| cluster3356 | 1.81 | 0.593 |
| cluster3419 | -1.70 | 0.593 |
| cluster3476 | 2.38 | 0.593 |
| cluster3561 | -1.71 | 0.593 |
| cluster3732 | 2.41 | 0.593 |
| cluster3738 | 2.50 | 0.593 |
| cluster3812 | -1.99 | 0.593 |
| cluster3890 | 2.11 | 0.593 |
| cluster3901 | 2.84 | 0.593 |
| cluster4031 | 1.96 | 0.593 |
| cluster3892 | 2.35 | 0.637 |
| cluster4057 | 2.30 | 0.637 |
| cluster2994 | 1.92 | 0.639 |
| cluster3128 | -1.42 | 0.725 |
| cluster3160 | 1.72 | 0.725 |
| cluster3166 | 1.28 | 0.725 |
| cluster3175 | 2.40 | 0.725 |
| cluster3177 | 2.24 | 0.725 |
| cluster3191 | 2.48 | 0.725 |

| | | |
|---|---|---|
| cluster3212 | 2.08 | 0.725 |
| cluster3406 | 1.49 | 0.725 |
| cluster3428 | -1.48 | 0.725 |
| cluster3487 | 2.47 | 0.725 |
| cluster3501 | 2.20 | 0.725 |
| cluster3545 | 2.11 | 0.725 |
| cluster3556 | 2.28 | 0.725 |
| cluster3755 | 2.58 | 0.725 |
| cluster3770 | 3.87 | 0.725 |
| cluster3787 | 2.37 | 0.725 |
| cluster3821 | -1.90 | 0.725 |
| cluster3851 | 2.34 | 0.725 |
| cluster3865 | 2.41 | 0.725 |
| cluster3866 | 2.51 | 0.725 |
| cluster3983 | 2.38 | 0.725 |
| cluster4016 | 2.03 | 0.725 |
| cluster4120 | -2.30 | 0.725 |
| cluster3077 | 1.69 | 0.734 |
| cluster3111 | -2.25 | 0.734 |
| cluster3417 | 1.62 | 0.734 |
| cluster3611 | 2.19 | 0.734 |
| cluster3936 | 2.14 | 0.734 |
| cluster3179 | 2.25 | 0.734 |
| cluster3222 | 1.06 | 0.734 |
| cluster4002 | 2.29 | 0.734 |
| cluster3026 | -2.08 | 0.748 |
| cluster2926 | 1.40 | 0.757 |
| cluster3146 | -1.40 | 0.757 |
| cluster3309 | -1.66 | 0.757 |
| cluster3448 | 2.12 | 0.757 |
| cluster3516 | 2.38 | 0.757 |
| cluster3571 | 1.94 | 0.757 |
| cluster3713 | 1.96 | 0.757 |
| cluster3748 | 1.39 | 0.757 |
| cluster3843 | 1.53 | 0.757 |
| cluster3853 | 2.29 | 0.757 |
| cluster3855 | 1.92 | 0.757 |
| cluster3899 | 2.05 | 0.757 |
| cluster4042 | 2.17 | 0.757 |
| cluster4055 | 2.15 | 0.757 |
| cluster4058 | 2.04 | 0.757 |
| cluster4079 | 1.71 | 0.757 |
| cluster4086 | 2.03 | 0.757 |
| cluster4095 | 1.89 | 0.757 |

| | | |
|---|---|---|
| cluster3885 | -2.07 | 0.803 |
| cluster3474 | 2.22 | 0.815 |
| cluster3195 | -1.31 | 0.820 |
| cluster3956 | 1.98 | 0.820 |
| cluster3976 | 1.41 | 0.820 |
| cluster3008 | 1.93 | 0.824 |
| cluster3123 | -1.28 | 0.825 |
| cluster3176 | -1.32 | 0.832 |
| cluster3182 | 1.95 | 0.832 |
| cluster3207 | 1.47 | 0.832 |
| cluster3386 | 2.01 | 0.832 |
| cluster3449 | 1.21 | 0.832 |
| cluster3451 | -2.31 | 0.832 |
| cluster3728 | 1.52 | 0.832 |
| cluster3871 | 1.89 | 0.832 |
| cluster3909 | 1.87 | 0.832 |
| cluster4118 | -2.35 | 0.832 |
| cluster3782 | -1.63 | 0.841 |
| cluster3877 | 1.32 | 0.857 |
| cluster4045 | 2.13 | 0.857 |
| cluster2904 | -1.52 | 0.861 |
| cluster3117 | 1.93 | 0.861 |
| cluster3192 | 1.34 | 0.861 |
| cluster3359 | 1.83 | 0.861 |
| cluster3371 | -1.92 | 0.861 |
| cluster3413 | 2.03 | 0.861 |
| cluster3529 | -1.84 | 0.861 |
| cluster3678 | 1.31 | 0.861 |
| cluster3717 | 1.27 | 0.861 |
| cluster3718 | 1.72 | 0.861 |
| cluster3741 | -1.50 | 0.861 |
| cluster3758 | 1.52 | 0.861 |
| cluster3938 | 1.71 | 0.861 |
| cluster3978 | 1.70 | 0.861 |
| cluster4069 | 1.63 | 0.861 |
| cluster4107 | 1.96 | 0.861 |
| cluster3502 | 1.89 | 0.861 |
| cluster3957 | 1.83 | 0.866 |
| cluster3859 | -2.10 | 0.873 |
| cluster3965 | 1.78 | 0.873 |
| cluster4033 | 2.02 | 0.875 |
| cluster2935 | 1.18 | 0.877 |
| cluster3231 | -2.17 | 0.877 |
| cluster3842 | -1.44 | 0.877 |

| | | |
|---|---|---|
| cluster3260 | 1.26 | 0.878 |
| cluster3972 | 1.75 | 0.878 |
| cluster2893 | 2.22 | 0.881 |
| cluster3063 | 1.40 | 0.881 |
| cluster3378 | 1.32 | 0.881 |
| cluster3410 | 1.40 | 0.881 |
| cluster3454 | 1.27 | 0.881 |
| cluster3543 | 1.92 | 0.881 |
| cluster3924 | 1.85 | 0.881 |
| cluster3940 | 1.32 | 0.881 |
| cluster4027 | 1.66 | 0.881 |
| cluster3604 | 1.86 | 0.885 |
| cluster3019 | 1.73 | 0.887 |
| cluster3494 | 1.22 | 0.888 |
| cluster4068 | 1.69 | 0.888 |
| cluster4121 | 1.83 | 0.888 |
| cluster2858 | -1.11 | 0.889 |
| cluster3283 | -0.99 | 0.889 |
| cluster3497 | 1.13 | 0.889 |
| cluster3805 | 1.83 | 0.889 |
| cluster2860 | 1.20 | 0.901 |
| cluster3118 | -1.26 | 0.901 |
| cluster3178 | 1.53 | 0.901 |
| cluster3482 | 1.48 | 0.901 |
| cluster3693 | 1.76 | 0.901 |
| cluster3906 | 1.59 | 0.901 |
| cluster2863 | 0.92 | 0.902 |
| cluster3342 | 1.69 | 0.902 |
| cluster3681 | 1.72 | 0.902 |
| cluster3897 | 1.43 | 0.902 |
| cluster3211 | 1.60 | 0.915 |
| cluster3246 | 1.06 | 0.915 |
| cluster3252 | 2.00 | 0.915 |
| cluster3299 | 1.67 | 0.915 |
| cluster3402 | -1.16 | 0.915 |
| cluster3443 | 1.57 | 0.915 |
| cluster3854 | 1.65 | 0.915 |
| cluster4030 | 1.73 | 0.915 |
| cluster4038 | 1.63 | 0.915 |
| cluster3041 | 1.36 | 0.916 |
| cluster3078 | 1.02 | 0.916 |
| cluster2962 | 0.96 | 0.924 |
| cluster3303 | 1.55 | 0.924 |
| cluster3306 | 1.91 | 0.924 |

| | | |
|---|---|---|
| cluster3333 | 1.57 | 0.924 |
| cluster3427 | -1.57 | 0.924 |
| cluster3619 | 1.30 | 0.924 |
| cluster3651 | 1.92 | 0.924 |
| cluster3686 | 1.57 | 0.924 |
| cluster3824 | 1.78 | 0.924 |
| cluster3618 | -1.75 | 0.931 |
| cluster3385 | 1.71 | 0.940 |
| cluster3735 | 1.32 | 0.940 |
| cluster3749 | 1.71 | 0.940 |
| cluster3752 | 1.92 | 0.940 |
| cluster3927 | 1.48 | 0.940 |
| cluster4054 | 1.57 | 0.940 |
| cluster2867 | 1.66 | 0.948 |
| cluster2876 | 1.78 | 0.948 |
| cluster2928 | 1.42 | 0.948 |
| cluster3106 | -0.96 | 0.948 |
| cluster3186 | 1.75 | 0.948 |
| cluster3349 | 1.30 | 0.948 |
| cluster3362 | 1.04 | 0.948 |
| cluster3550 | -0.97 | 0.948 |
| cluster3703 | 0.97 | 0.948 |
| cluster3912 | 0.88 | 0.948 |
| cluster3955 | 1.63 | 0.948 |
| cluster3991 | 1.47 | 0.948 |
| cluster4052 | 1.31 | 0.948 |
| cluster4112 | 1.61 | 0.948 |
| cluster4123 | 1.62 | 0.948 |
| cluster3886 | 1.54 | 0.948 |
| cluster3834 | 1.66 | 0.949 |
| cluster3922 | 1.65 | 0.949 |
| cluster2878 | -1.12 | 0.950 |
| cluster2912 | 1.29 | 0.950 |
| cluster3075 | 1.13 | 0.950 |
| cluster3274 | 1.55 | 0.950 |
| cluster3286 | -0.88 | 0.950 |
| cluster3323 | 1.29 | 0.950 |
| cluster3399 | -1.26 | 0.950 |
| cluster3404 | 0.88 | 0.950 |
| cluster3408 | -1.22 | 0.950 |
| cluster3620 | 1.08 | 0.950 |
| cluster3723 | 1.06 | 0.950 |
| cluster3840 | 1.21 | 0.950 |
| cluster3880 | 1.85 | 0.950 |

| | | |
|---|---|---|
| cluster3904 | 1.77 | 0.950 |
| cluster4071 | 1.60 | 0.950 |
| cluster2948 | -0.92 | 0.951 |
| cluster3089 | 1.06 | 0.951 |
| cluster3109 | 1.58 | 0.951 |
| cluster3479 | 1.46 | 0.951 |
| cluster3629 | 1.57 | 0.951 |
| cluster4051 | 1.73 | 0.951 |
| cluster3512 | 1.28 | 0.952 |
| cluster2995 | 1.34 | 0.954 |
| cluster3060 | 1.52 | 0.954 |
| cluster3061 | 0.92 | 0.954 |
| cluster3144 | 1.52 | 0.954 |
| cluster3156 | 1.44 | 0.954 |
| cluster3257 | 0.89 | 0.954 |
| cluster3392 | 1.50 | 0.954 |
| cluster3506 | 1.35 | 0.954 |
| cluster3572 | 1.73 | 0.954 |
| cluster3587 | 1.73 | 0.954 |
| cluster3607 | -1.43 | 0.954 |
| cluster3614 | 1.50 | 0.954 |
| cluster3663 | 1.41 | 0.954 |
| cluster3666 | 1.67 | 0.954 |
| cluster3688 | 1.45 | 0.954 |
| cluster3710 | 1.34 | 0.954 |
| cluster3719 | 1.43 | 0.954 |
| cluster3747 | 1.23 | 0.954 |
| cluster3754 | 1.05 | 0.954 |
| cluster3757 | -1.42 | 0.954 |
| cluster3765 | 1.12 | 0.954 |
| cluster3783 | 1.48 | 0.954 |
| cluster3784 | 0.99 | 0.954 |
| cluster3830 | 1.02 | 0.954 |
| cluster3881 | 1.32 | 0.954 |
| cluster3915 | 1.66 | 0.954 |
| cluster3944 | 1.41 | 0.954 |
| cluster3981 | 1.33 | 0.954 |
| cluster3987 | 1.11 | 0.954 |
| cluster4014 | 1.47 | 0.954 |
| cluster4022 | 1.76 | 0.954 |
| cluster4053 | 1.38 | 0.954 |
| cluster4060 | 1.52 | 0.954 |
| cluster4062 | -1.59 | 0.954 |
| cluster4066 | 1.33 | 0.954 |

| | | |
|---|---|---|
| cluster4100 | 1.34 | 0.954 |
| cluster4113 | 1.30 | 0.954 |
| cluster4125 | 1.75 | 0.954 |
| cluster3593 | 1.35 | 0.958 |
| cluster2885 | -1.29 | 0.963 |
| cluster2892 | 0.85 | 0.963 |
| cluster2905 | -0.82 | 0.963 |
| cluster2925 | -1.46 | 0.963 |
| cluster2933 | -0.96 | 0.963 |
| cluster2934 | -0.94 | 0.963 |
| cluster2959 | -0.81 | 0.963 |
| cluster3012 | 1.46 | 0.963 |
| cluster3046 | -1.07 | 0.963 |
| cluster3088 | 1.13 | 0.963 |
| cluster3102 | -0.72 | 0.963 |
| cluster3107 | 0.94 | 0.963 |
| cluster3108 | -1.37 | 0.963 |
| cluster3126 | -0.70 | 0.963 |
| cluster3139 | 1.40 | 0.963 |
| cluster3142 | 1.26 | 0.963 |
| cluster3151 | -1.00 | 0.963 |
| cluster3163 | 1.38 | 0.963 |
| cluster3167 | -1.26 | 0.963 |
| cluster3185 | 1.52 | 0.963 |
| cluster3198 | 1.25 | 0.963 |
| cluster3200 | 0.90 | 0.963 |
| cluster3236 | 1.35 | 0.963 |
| cluster3259 | -0.87 | 0.963 |
| cluster3276 | -0.82 | 0.963 |
| cluster3291 | 0.93 | 0.963 |
| cluster3293 | 0.84 | 0.963 |
| cluster3319 | -1.39 | 0.963 |
| cluster3364 | 1.45 | 0.963 |
| cluster3368 | -0.80 | 0.963 |
| cluster3387 | 1.30 | 0.963 |
| cluster3447 | 1.23 | 0.963 |
| cluster3467 | 1.21 | 0.963 |
| cluster3471 | 1.31 | 0.963 |
| cluster3480 | 1.37 | 0.963 |
| cluster3481 | 1.39 | 0.963 |
| cluster3500 | 1.68 | 0.963 |
| cluster3511 | 1.38 | 0.963 |
| cluster3513 | -0.88 | 0.963 |
| cluster3517 | 0.80 | 0.963 |

| | | |
|---|---|---|
| cluster3646 | 1.29 | 0.963 |
| cluster3658 | -1.00 | 0.963 |
| cluster3659 | 1.42 | 0.963 |
| cluster3677 | 0.91 | 0.963 |
| cluster3701 | 0.88 | 0.963 |
| cluster3773 | 0.97 | 0.963 |
| cluster3793 | 1.25 | 0.963 |
| cluster3794 | 1.26 | 0.963 |
| cluster3804 | 1.42 | 0.963 |
| cluster3819 | 1.55 | 0.963 |
| cluster3823 | 0.92 | 0.963 |
| cluster3831 | 1.22 | 0.963 |
| cluster3849 | 1.25 | 0.963 |
| cluster3864 | 1.23 | 0.963 |
| cluster3905 | 2.48 | 0.963 |
| cluster3908 | 1.12 | 0.963 |
| cluster3985 | 1.06 | 0.963 |
| cluster3989 | -0.97 | 0.963 |
| cluster4026 | 1.22 | 0.963 |
| cluster4029 | 1.43 | 0.963 |
| cluster4044 | 1.32 | 0.963 |
| cluster4061 | 1.17 | 0.963 |
| cluster3533 | 0.96 | 0.964 |
| cluster4050 | 1.21 | 0.966 |
| cluster3005 | -0.95 | 0.967 |
| cluster3360 | 1.07 | 0.967 |
| cluster4087 | 1.15 | 0.969 |
| cluster2894 | -1.25 | 0.970 |
| cluster2908 | -0.85 | 0.970 |
| cluster2924 | -1.34 | 0.970 |
| cluster3009 | 1.01 | 0.970 |
| cluster3037 | -1.01 | 0.970 |
| cluster3064 | -0.95 | 0.970 |
| cluster3070 | -0.78 | 0.970 |
| cluster3081 | 0.89 | 0.970 |
| cluster3095 | 0.90 | 0.970 |
| cluster3100 | -1.45 | 0.970 |
| cluster3112 | -1.16 | 0.970 |
| cluster3136 | 1.09 | 0.970 |
| cluster3173 | 1.19 | 0.970 |
| cluster3219 | 0.99 | 0.970 |
| cluster3271 | 0.72 | 0.970 |
| cluster3279 | 1.04 | 0.970 |
| cluster3296 | 1.42 | 0.970 |

| | | |
|---|---|---|
| cluster3328 | -0.70 | 0.970 |
| cluster3348 | -0.86 | 0.970 |
| cluster3365 | 1.24 | 0.970 |
| cluster3366 | 1.17 | 0.970 |
| cluster3393 | 1.38 | 0.970 |
| cluster3409 | 1.19 | 0.970 |
| cluster3420 | -1.17 | 0.970 |
| cluster3478 | -0.77 | 0.970 |
| cluster3498 | 1.15 | 0.970 |
| cluster3585 | -0.84 | 0.970 |
| cluster3606 | 1.12 | 0.970 |
| cluster3650 | 1.11 | 0.970 |
| cluster3654 | 1.05 | 0.970 |
| cluster3657 | 1.13 | 0.970 |
| cluster3724 | 1.13 | 0.970 |
| cluster3731 | 0.92 | 0.970 |
| cluster3774 | -0.83 | 0.970 |
| cluster3785 | -1.21 | 0.970 |
| cluster3800 | 1.55 | 0.970 |
| cluster3813 | 1.09 | 0.970 |
| cluster3869 | 1.20 | 0.970 |
| cluster3884 | 1.37 | 0.970 |
| cluster3887 | 1.10 | 0.970 |
| cluster3902 | 1.23 | 0.970 |
| cluster3951 | 1.13 | 0.970 |
| cluster3952 | 1.19 | 0.970 |
| cluster3968 | 1.14 | 0.970 |
| cluster3970 | 1.22 | 0.970 |
| cluster3993 | 1.10 | 0.970 |
| cluster3998 | 1.48 | 0.970 |
| cluster4001 | 1.13 | 0.970 |
| cluster4024 | 1.09 | 0.970 |
| cluster4032 | -1.00 | 0.970 |
| cluster4035 | 0.90 | 0.970 |
| cluster4080 | 1.54 | 0.970 |
| cluster4083 | 1.18 | 0.970 |
| cluster4084 | 1.00 | 0.970 |
| cluster4099 | 1.18 | 0.970 |
| cluster4126 | 1.31 | 0.970 |
| cluster3313 | 1.11 | 0.973 |
| cluster3644 | 0.94 | 0.973 |
| cluster3980 | 0.68 | 0.973 |
| cluster3011 | 0.82 | 0.973 |
| cluster3187 | -1.20 | 0.978 |

| | | |
|---|---|---|
| cluster3416 | 1.16 | 0.978 |
| cluster4111 | -1.34 | 0.978 |
| cluster2906 | -0.75 | 0.980 |
| cluster2932 | 0.80 | 0.980 |
| cluster2965 | -1.03 | 0.980 |
| cluster3042 | 1.04 | 0.980 |
| cluster3083 | 0.69 | 0.980 |
| cluster3131 | 1.17 | 0.980 |
| cluster3363 | 1.07 | 0.980 |
| cluster3375 | 0.83 | 0.980 |
| cluster3594 | 1.13 | 0.980 |
| cluster3609 | -1.19 | 0.980 |
| cluster3638 | 1.14 | 0.980 |
| cluster3648 | -0.70 | 0.980 |
| cluster3661 | 0.76 | 0.980 |
| cluster3671 | 1.13 | 0.980 |
| cluster3674 | 1.34 | 0.980 |
| cluster3698 | -0.67 | 0.980 |
| cluster3743 | 1.03 | 0.980 |
| cluster3856 | 1.14 | 0.980 |
| cluster3863 | 0.85 | 0.980 |
| cluster3882 | 1.14 | 0.980 |
| cluster4009 | -0.96 | 0.980 |
| cluster4128 | 1.29 | 0.980 |
| cluster3781 | 0.82 | 0.982 |
| cluster3963 | 1.11 | 0.982 |
| cluster4008 | 0.98 | 0.982 |
| cluster4017 | 0.97 | 0.982 |
| cluster3764 | 1.12 | 0.986 |
| cluster2981 | 1.04 | 0.993 |
| cluster3053 | 0.87 | 0.993 |
| cluster3838 | -0.94 | 0.993 |
| cluster1127 | 0.50 | 0.994 |
| cluster2854 | 0.63 | 0.994 |
| cluster2857 | 0.32 | 0.994 |
| cluster2859 | 0.41 | 0.994 |
| cluster2865 | 0.40 | 0.994 |
| cluster2870 | 0.43 | 0.994 |
| cluster2872 | -0.46 | 0.994 |
| cluster2873 | 0.47 | 0.994 |
| cluster2886 | 0.66 | 0.994 |
| cluster2887 | 0.93 | 0.994 |
| cluster2888 | 1.09 | 0.994 |
| cluster2898 | 0.45 | 0.994 |

| | | |
|---|---|---|
| cluster2899 | 0.40 | 0.994 |
| cluster2901 | -0.76 | 0.994 |
| cluster2903 | -0.35 | 0.994 |
| cluster2913 | 0.59 | 0.994 |
| cluster2914 | 0.67 | 0.994 |
| cluster2915 | 0.73 | 0.994 |
| cluster2918 | -0.74 | 0.994 |
| cluster2919 | -0.61 | 0.994 |
| cluster2923 | -0.55 | 0.994 |
| cluster2927 | 0.70 | 0.994 |
| cluster2930 | 0.57 | 0.994 |
| cluster2936 | 0.84 | 0.994 |
| cluster2938 | 0.50 | 0.994 |
| cluster2941 | -0.35 | 0.994 |
| cluster2945 | 0.59 | 0.994 |
| cluster2946 | -0.53 | 0.994 |
| cluster2947 | 0.42 | 0.994 |
| cluster2949 | -0.57 | 0.994 |
| cluster2950 | -0.44 | 0.994 |
| cluster2953 | -0.49 | 0.994 |
| cluster2956 | -0.59 | 0.994 |
| cluster2957 | -0.99 | 0.994 |
| cluster2960 | -0.32 | 0.994 |
| cluster2968 | -0.25 | 0.994 |
| cluster2969 | 0.74 | 0.994 |
| cluster2970 | 0.44 | 0.994 |
| cluster2971 | 0.25 | 0.994 |
| cluster2973 | -0.62 | 0.994 |
| cluster2974 | -0.73 | 0.994 |
| cluster2975 | 0.43 | 0.994 |
| cluster2977 | -0.77 | 0.994 |
| cluster2978 | -0.31 | 0.994 |
| cluster2979 | 0.47 | 0.994 |
| cluster2980 | 0.56 | 0.994 |
| cluster2982 | -0.85 | 0.994 |
| cluster2985 | 0.48 | 0.994 |
| cluster2987 | -0.44 | 0.994 |
| cluster2988 | 0.84 | 0.994 |
| cluster2989 | -0.38 | 0.994 |
| cluster2992 | -0.45 | 0.994 |
| cluster2993 | 0.49 | 0.994 |
| cluster2999 | 0.70 | 0.994 |
| cluster3002 | -0.32 | 0.994 |
| cluster3006 | 0.56 | 0.994 |

| | | |
|---|---|---|
| cluster3007 | 0.55 | 0.994 |
| cluster3018 | 0.64 | 0.994 |
| cluster3021 | 0.38 | 0.994 |
| cluster3022 | -0.79 | 0.994 |
| cluster3023 | 0.58 | 0.994 |
| cluster3025 | 0.38 | 0.994 |
| cluster3027 | 0.34 | 0.994 |
| cluster3031 | -0.56 | 0.994 |
| cluster3032 | -0.50 | 0.994 |
| cluster3033 | 0.67 | 0.994 |
| cluster3034 | 0.70 | 0.994 |
| cluster3036 | -0.48 | 0.994 |
| cluster3038 | -0.59 | 0.994 |
| cluster3039 | 0.38 | 0.994 |
| cluster3040 | -0.43 | 0.994 |
| cluster3048 | 0.74 | 0.994 |
| cluster3050 | -0.41 | 0.994 |
| cluster3058 | 0.31 | 0.994 |
| cluster3059 | 0.94 | 0.994 |
| cluster3062 | 0.36 | 0.994 |
| cluster3065 | -0.87 | 0.994 |
| cluster3067 | -0.29 | 0.994 |
| cluster3068 | -0.57 | 0.994 |
| cluster3069 | -0.32 | 0.994 |
| cluster3071 | -0.45 | 0.994 |
| cluster3076 | 0.68 | 0.994 |
| cluster3079 | 0.66 | 0.994 |
| cluster3087 | 0.50 | 0.994 |
| cluster3090 | -1.16 | 0.994 |
| cluster3094 | 0.96 | 0.994 |
| cluster3096 | 0.77 | 0.994 |
| cluster3098 | 0.87 | 0.994 |
| cluster3113 | -0.57 | 0.994 |
| cluster3114 | -0.46 | 0.994 |
| cluster3119 | 0.59 | 0.994 |
| cluster3121 | -0.72 | 0.994 |
| cluster3122 | 0.46 | 0.994 |
| cluster3125 | 0.69 | 0.994 |
| cluster3127 | 0.72 | 0.994 |
| cluster3129 | -0.74 | 0.994 |
| cluster3132 | 0.35 | 0.994 |
| cluster3135 | -0.61 | 0.994 |
| cluster3138 | 0.42 | 0.994 |
| cluster3147 | 0.52 | 0.994 |

| | | |
|---|---|---|
| cluster3149 | 0.51 | 0.994 |
| cluster3152 | 0.60 | 0.994 |
| cluster3154 | 0.29 | 0.994 |
| cluster3157 | -0.42 | 0.994 |
| cluster3162 | 0.73 | 0.994 |
| cluster3164 | -0.99 | 0.994 |
| cluster3168 | 0.46 | 0.994 |
| cluster3171 | -0.76 | 0.994 |
| cluster3189 | -0.57 | 0.994 |
| cluster3190 | 0.56 | 0.994 |
| cluster3196 | -0.49 | 0.994 |
| cluster3197 | -0.45 | 0.994 |
| cluster3199 | -0.59 | 0.994 |
| cluster3203 | 0.73 | 0.994 |
| cluster3205 | -0.34 | 0.994 |
| cluster3213 | 0.73 | 0.994 |
| cluster3214 | 0.50 | 0.994 |
| cluster3215 | -0.35 | 0.994 |
| cluster3218 | 0.93 | 0.994 |
| cluster3223 | 0.53 | 0.994 |
| cluster3224 | -0.70 | 0.994 |
| cluster3226 | 0.77 | 0.994 |
| cluster3229 | -0.68 | 0.994 |
| cluster3233 | 0.61 | 0.994 |
| cluster3234 | 0.56 | 0.994 |
| cluster3239 | -0.57 | 0.994 |
| cluster3241 | -0.43 | 0.994 |
| cluster3242 | 0.48 | 0.994 |
| cluster3243 | -0.63 | 0.994 |
| cluster3244 | -0.36 | 0.994 |
| cluster3245 | -0.60 | 0.994 |
| cluster3249 | -0.50 | 0.994 |
| cluster3254 | 0.83 | 0.994 |
| cluster3263 | -0.67 | 0.994 |
| cluster3264 | 0.45 | 0.994 |
| cluster3265 | 0.84 | 0.994 |
| cluster3266 | -0.46 | 0.994 |
| cluster3268 | 0.74 | 0.994 |
| cluster3269 | -0.53 | 0.994 |
| cluster3270 | 0.67 | 0.994 |
| cluster3272 | -1.26 | 0.994 |
| cluster3273 | -0.36 | 0.994 |
| cluster3275 | 0.88 | 0.994 |
| cluster3280 | -0.69 | 0.994 |

| | | |
|---|---|---|
| cluster3287 | -0.79 | 0.994 |
| cluster3288 | 0.56 | 0.994 |
| cluster3294 | 0.64 | 0.994 |
| cluster3300 | -0.68 | 0.994 |
| cluster3302 | -0.69 | 0.994 |
| cluster3305 | -0.57 | 0.994 |
| cluster3310 | -0.49 | 0.994 |
| cluster3311 | 0.65 | 0.994 |
| cluster3312 | 0.59 | 0.994 |
| cluster3314 | -0.98 | 0.994 |
| cluster3316 | -0.77 | 0.994 |
| cluster3325 | 0.68 | 0.994 |
| cluster3326 | -0.57 | 0.994 |
| cluster3332 | 0.66 | 0.994 |
| cluster3336 | -0.58 | 0.994 |
| cluster3337 | 0.76 | 0.994 |
| cluster3338 | 0.74 | 0.994 |
| cluster3343 | -0.71 | 0.994 |
| cluster3345 | 0.84 | 0.994 |
| cluster3346 | 0.54 | 0.994 |
| cluster3347 | 0.43 | 0.994 |
| cluster3351 | 0.73 | 0.994 |
| cluster3357 | -0.52 | 0.994 |
| cluster3361 | 0.52 | 0.994 |
| cluster3370 | 0.66 | 0.994 |
| cluster3372 | 0.49 | 0.994 |
| cluster3376 | 0.56 | 0.994 |
| cluster3377 | -0.72 | 0.994 |
| cluster3380 | -0.42 | 0.994 |
| cluster3381 | 0.43 | 0.994 |
| cluster3382 | 0.75 | 0.994 |
| cluster3389 | 0.70 | 0.994 |
| cluster3390 | -0.48 | 0.994 |
| cluster3391 | 0.38 | 0.994 |
| cluster3394 | -0.34 | 0.994 |
| cluster3395 | 0.72 | 0.994 |
| cluster3398 | 0.56 | 0.994 |
| cluster3400 | -0.33 | 0.994 |
| cluster3405 | -0.63 | 0.994 |
| cluster3411 | 0.97 | 0.994 |
| cluster3412 | 0.75 | 0.994 |
| cluster3418 | 0.63 | 0.994 |
| cluster3421 | 0.58 | 0.994 |
| cluster3422 | -0.83 | 0.994 |

| | | |
|---|---|---|
| cluster3423 | 0.54 | 0.994 |
| cluster3429 | 0.45 | 0.994 |
| cluster3432 | 0.42 | 0.994 |
| cluster3437 | 0.40 | 0.994 |
| cluster3439 | -0.79 | 0.994 |
| cluster3440 | -0.37 | 0.994 |
| cluster3450 | 1.17 | 0.994 |
| cluster3452 | 0.63 | 0.994 |
| cluster3455 | -0.69 | 0.994 |
| cluster3459 | 0.55 | 0.994 |
| cluster3462 | 0.61 | 0.994 |
| cluster3464 | 0.58 | 0.994 |
| cluster3465 | 0.35 | 0.994 |
| cluster3470 | 0.49 | 0.994 |
| cluster3472 | 0.41 | 0.994 |
| cluster3473 | 0.83 | 0.994 |
| cluster3477 | -0.67 | 0.994 |
| cluster3485 | 1.08 | 0.994 |
| cluster3486 | 0.44 | 0.994 |
| cluster3489 | 0.52 | 0.994 |
| cluster3490 | 0.92 | 0.994 |
| cluster3495 | 0.58 | 0.994 |
| cluster3499 | 0.55 | 0.994 |
| cluster3503 | 0.66 | 0.994 |
| cluster3505 | -0.61 | 0.994 |
| cluster3507 | 0.64 | 0.994 |
| cluster3508 | 0.82 | 0.994 |
| cluster3523 | 0.69 | 0.994 |
| cluster3525 | -0.59 | 0.994 |
| cluster3526 | 0.47 | 0.994 |
| cluster3531 | 0.75 | 0.994 |
| cluster3532 | 0.48 | 0.994 |
| cluster3534 | -0.88 | 0.994 |
| cluster3537 | 0.75 | 0.994 |
| cluster3539 | 0.69 | 0.994 |
| cluster3541 | -0.40 | 0.994 |
| cluster3544 | 0.58 | 0.994 |
| cluster3548 | 0.33 | 0.994 |
| cluster3554 | -0.84 | 0.994 |
| cluster3557 | 0.47 | 0.994 |
| cluster3558 | -0.66 | 0.994 |
| cluster3559 | -0.87 | 0.994 |
| cluster3562 | -0.71 | 0.994 |
| cluster3564 | 0.74 | 0.994 |

| | | |
|---|---|---|
| cluster3568 | -0.99 | 0.994 |
| cluster3569 | 0.81 | 0.994 |
| cluster3570 | 0.96 | 0.994 |
| cluster3574 | 1.01 | 0.994 |
| cluster3575 | -0.58 | 0.994 |
| cluster3578 | -0.79 | 0.994 |
| cluster3579 | 0.47 | 0.994 |
| cluster3583 | -0.76 | 0.994 |
| cluster3584 | 0.71 | 0.994 |
| cluster3591 | 0.66 | 0.994 |
| cluster3592 | 0.94 | 0.994 |
| cluster3595 | 0.48 | 0.994 |
| cluster3599 | 0.80 | 0.994 |
| cluster3600 | -0.55 | 0.994 |
| cluster3608 | 0.44 | 0.994 |
| cluster3613 | -0.53 | 0.994 |
| cluster3617 | 1.10 | 0.994 |
| cluster3621 | 0.87 | 0.994 |
| cluster3622 | 0.48 | 0.994 |
| cluster3623 | 0.98 | 0.994 |
| cluster3625 | 0.54 | 0.994 |
| cluster3628 | 0.90 | 0.994 |
| cluster3630 | 0.31 | 0.994 |
| cluster3631 | 0.67 | 0.994 |
| cluster3632 | 0.55 | 0.994 |
| cluster3633 | 0.57 | 0.994 |
| cluster3637 | 0.56 | 0.994 |
| cluster3640 | 1.80 | 0.994 |
| cluster3642 | 0.74 | 0.994 |
| cluster3645 | 0.84 | 0.994 |
| cluster3647 | 0.97 | 0.994 |
| cluster3653 | 0.75 | 0.994 |
| cluster3656 | 0.70 | 0.994 |
| cluster3662 | 0.50 | 0.994 |
| cluster3665 | 0.83 | 0.994 |
| cluster3667 | 0.68 | 0.994 |
| cluster3668 | 0.89 | 0.994 |
| cluster3669 | 0.62 | 0.994 |
| cluster3672 | 0.86 | 0.994 |
| cluster3673 | 0.74 | 0.994 |
| cluster3675 | -0.32 | 0.994 |
| cluster3680 | 0.52 | 0.994 |
| cluster3692 | 0.72 | 0.994 |
| cluster3694 | 0.69 | 0.994 |

| | | |
|---|---|---|
| cluster3695 | 0.57 | 0.994 |
| cluster3696 | 0.89 | 0.994 |
| cluster3699 | 0.63 | 0.994 |
| cluster3702 | -0.66 | 0.994 |
| cluster3705 | 0.65 | 0.994 |
| cluster3708 | 1.02 | 0.994 |
| cluster3711 | 0.38 | 0.994 |
| cluster3712 | 0.54 | 0.994 |
| cluster3714 | -0.37 | 0.994 |
| cluster3720 | 0.72 | 0.994 |
| cluster3725 | 0.80 | 0.994 |
| cluster3726 | -0.37 | 0.994 |
| cluster3730 | 0.44 | 0.994 |
| cluster3733 | 0.57 | 0.994 |
| cluster3734 | 0.79 | 0.994 |
| cluster3736 | -0.55 | 0.994 |
| cluster3739 | 0.37 | 0.994 |
| cluster3740 | 0.56 | 0.994 |
| cluster3750 | 0.58 | 0.994 |
| cluster3759 | -0.58 | 0.994 |
| cluster3762 | -0.83 | 0.994 |
| cluster3766 | 0.95 | 0.994 |
| cluster3767 | 0.60 | 0.994 |
| cluster3769 | -0.51 | 0.994 |
| cluster3772 | -0.61 | 0.994 |
| cluster3788 | 0.55 | 0.994 |
| cluster3789 | -0.63 | 0.994 |
| cluster3790 | -0.81 | 0.994 |
| cluster3795 | 0.53 | 0.994 |
| cluster3796 | 0.62 | 0.994 |
| cluster3801 | 0.79 | 0.994 |
| cluster3806 | 0.72 | 0.994 |
| cluster3809 | 0.60 | 0.994 |
| cluster3811 | 0.46 | 0.994 |
| cluster3814 | -0.45 | 0.994 |
| cluster3815 | 1.04 | 0.994 |
| cluster3817 | 0.72 | 0.994 |
| cluster3820 | 0.55 | 0.994 |
| cluster3826 | 0.91 | 0.994 |
| cluster3828 | 0.51 | 0.994 |
| cluster3833 | -0.93 | 0.994 |
| cluster3836 | 0.99 | 0.994 |
| cluster3839 | -0.63 | 0.994 |
| cluster3844 | 0.75 | 0.994 |

| | | |
|---|---|---|
| cluster3846 | 0.32 | 0.994 |
| cluster3848 | -0.78 | 0.994 |
| cluster3850 | 0.95 | 0.994 |
| cluster3860 | -0.38 | 0.994 |
| cluster3861 | 0.67 | 0.994 |
| cluster3867 | 0.65 | 0.994 |
| cluster3870 | 0.52 | 0.994 |
| cluster3873 | 0.51 | 0.994 |
| cluster3874 | -0.48 | 0.994 |
| cluster3875 | 0.76 | 0.994 |
| cluster3879 | 0.56 | 0.994 |
| cluster3888 | 0.76 | 0.994 |
| cluster3893 | 0.56 | 0.994 |
| cluster3894 | 0.91 | 0.994 |
| cluster3910 | 0.57 | 0.994 |
| cluster3913 | -0.63 | 0.994 |
| cluster3914 | 0.69 | 0.994 |
| cluster3916 | 0.92 | 0.994 |
| cluster3917 | 0.52 | 0.994 |
| cluster3918 | 0.65 | 0.994 |
| cluster3920 | 0.76 | 0.994 |
| cluster3925 | 1.28 | 0.994 |
| cluster3928 | 0.56 | 0.994 |
| cluster3931 | -0.70 | 0.994 |
| cluster3933 | -0.63 | 0.994 |
| cluster3937 | 0.84 | 0.994 |
| cluster3939 | 0.77 | 0.994 |
| cluster3941 | -0.65 | 0.994 |
| cluster3942 | -0.58 | 0.994 |
| cluster3945 | 1.04 | 0.994 |
| cluster3948 | 0.77 | 0.994 |
| cluster3949 | -0.92 | 0.994 |
| cluster3954 | 0.76 | 0.994 |
| cluster3958 | 0.92 | 0.994 |
| cluster3961 | 0.56 | 0.994 |
| cluster3966 | 0.83 | 0.994 |
| cluster3974 | 1.16 | 0.994 |
| cluster3979 | 0.70 | 0.994 |
| cluster3982 | 0.84 | 0.994 |
| cluster3986 | 0.59 | 0.994 |
| cluster3992 | 0.74 | 0.994 |
| cluster3994 | 0.95 | 0.994 |
| cluster3997 | 0.79 | 0.994 |
| cluster3999 | 0.73 | 0.994 |

| | | |
|---|---|---|
| cluster4000 | 0.84 | 0.994 |
| cluster4003 | -0.65 | 0.994 |
| cluster4007 | 0.59 | 0.994 |
| cluster4013 | -0.71 | 0.994 |
| cluster4015 | 0.90 | 0.994 |
| cluster4020 | 0.35 | 0.994 |
| cluster4023 | -0.75 | 0.994 |
| cluster4025 | 0.49 | 0.994 |
| cluster4036 | 0.53 | 0.994 |
| cluster4037 | 0.83 | 0.994 |
| cluster4039 | -0.76 | 0.994 |
| cluster4041 | 0.62 | 0.994 |
| cluster4046 | 1.16 | 0.994 |
| cluster4047 | 0.66 | 0.994 |
| cluster4048 | 0.68 | 0.994 |
| cluster4059 | 0.99 | 0.994 |
| cluster4063 | 0.90 | 0.994 |
| cluster4064 | 0.75 | 0.994 |
| cluster4070 | 0.65 | 0.994 |
| cluster4073 | 1.23 | 0.994 |
| cluster4075 | -0.68 | 0.994 |
| cluster4082 | 0.80 | 0.994 |
| cluster4090 | -0.86 | 0.994 |
| cluster4096 | 0.68 | 0.994 |
| cluster4105 | 0.92 | 0.994 |
| cluster4106 | 1.16 | 0.994 |
| cluster4109 | 1.07 | 0.994 |
| cluster4110 | 0.59 | 0.994 |
| cluster4114 | -1.02 | 0.994 |
| cluster4115 | 0.68 | 0.994 |
| cluster4117 | -1.08 | 0.994 |
| cluster4122 | -0.48 | 0.994 |
| cluster3542 | 0.46 | 0.995 |
| cluster3900 | 0.61 | 0.995 |
| cluster2998 | 0.48 | 0.995 |
| cluster3181 | 0.30 | 0.995 |
| cluster3327 | 0.34 | 0.995 |
| cluster3676 | 0.49 | 0.995 |
| cluster3679 | 0.35 | 0.995 |
| cluster3683 | 0.48 | 0.995 |
| cluster3721 | 0.39 | 0.995 |
| cluster3969 | 0.29 | 0.995 |
| cluster3984 | -0.54 | 0.995 |
| cluster2871 | -0.33 | 0.998 |

| | | |
|---|---|---|
| cluster2929 | 0.42 | 0.999 |
| cluster3055 | -0.34 | 0.999 |
| cluster3085 | -0.43 | 0.999 |
| cluster3216 | 0.42 | 0.999 |
| cluster3321 | -0.42 | 0.999 |
| cluster3358 | 0.32 | 0.999 |
| cluster3367 | 0.27 | 0.999 |
| cluster3424 | 0.52 | 0.999 |
| cluster3435 | 0.50 | 0.999 |
| cluster3438 | 0.44 | 0.999 |
| cluster3458 | -0.28 | 0.999 |
| cluster3496 | -0.32 | 0.999 |
| cluster3577 | 0.29 | 0.999 |
| cluster3602 | 0.29 | 0.999 |
| cluster3687 | 0.41 | 0.999 |
| cluster3737 | -0.26 | 0.999 |
| cluster4019 | 0.53 | 0.999 |
| cluster2853 | 0.30 | 0.999 |
| cluster2855 | -0.11 | 0.999 |
| cluster2856 | -0.09 | 0.999 |
| cluster2861 | -0.26 | 0.999 |
| cluster2862 | 0.23 | 0.999 |
| cluster2864 | 0.26 | 0.999 |
| cluster2866 | -0.04 | 0.999 |
| cluster2869 | 0.10 | 0.999 |
| cluster2874 | -0.26 | 0.999 |
| cluster2875 | 0.03 | 0.999 |
| cluster2877 | -0.20 | 0.999 |
| cluster2879 | 0.27 | 0.999 |
| cluster2880 | 0.09 | 0.999 |
| cluster2881 | 0.24 | 0.999 |
| cluster2882 | -0.11 | 0.999 |
| cluster2883 | 0.25 | 0.999 |
| cluster2884 | 0.29 | 0.999 |
| cluster2889 | 0.23 | 0.999 |
| cluster2890 | -0.22 | 0.999 |
| cluster2891 | 0.32 | 0.999 |
| cluster2895 | -0.01 | 0.999 |
| cluster2896 | 0.28 | 0.999 |
| cluster2897 | -0.29 | 0.999 |
| cluster2900 | -0.03 | 0.999 |
| cluster2907 | 0.19 | 0.999 |
| cluster2909 | 0.13 | 0.999 |
| cluster2910 | -0.32 | 0.999 |

| | | |
|---|---|---|
| cluster2911 | 0.23 | 0.999 |
| cluster2916 | 0.13 | 0.999 |
| cluster2917 | 0.38 | 0.999 |
| cluster2920 | 0.08 | 0.999 |
| cluster2921 | -0.04 | 0.999 |
| cluster2922 | 0.04 | 0.999 |
| cluster2931 | -0.14 | 0.999 |
| cluster2937 | -0.16 | 0.999 |
| cluster2939 | 0.17 | 0.999 |
| cluster2940 | 0.25 | 0.999 |
| cluster2942 | -0.06 | 0.999 |
| cluster2943 | -0.23 | 0.999 |
| cluster2944 | 0.11 | 0.999 |
| cluster2951 | -0.14 | 0.999 |
| cluster2952 | -0.15 | 0.999 |
| cluster2954 | -0.27 | 0.999 |
| cluster2955 | 0.15 | 0.999 |
| cluster2958 | 0.06 | 0.999 |
| cluster2963 | 0.14 | 0.999 |
| cluster2964 | -0.23 | 0.999 |
| cluster2966 | -0.05 | 0.999 |
| cluster2967 | -0.25 | 0.999 |
| cluster2972 | 0.19 | 0.999 |
| cluster2976 | 0.34 | 0.999 |
| cluster2983 | -0.11 | 0.999 |
| cluster2984 | 0.01 | 0.999 |
| cluster2986 | 0.24 | 0.999 |
| cluster2990 | -0.42 | 0.999 |
| cluster2991 | 0.04 | 0.999 |
| cluster2996 | 0.03 | 0.999 |
| cluster2997 | -0.40 | 0.999 |
| cluster3000 | 0.09 | 0.999 |
| cluster3003 | -0.13 | 0.999 |
| cluster3010 | -0.26 | 0.999 |
| cluster3013 | -0.07 | 0.999 |
| cluster3014 | 0.13 | 0.999 |
| cluster3016 | -0.23 | 0.999 |
| cluster3017 | -0.24 | 0.999 |
| cluster3020 | -0.48 | 0.999 |
| cluster3028 | -0.09 | 0.999 |
| cluster3029 | -0.23 | 0.999 |
| cluster3035 | -0.17 | 0.999 |
| cluster3043 | 0.29 | 0.999 |
| cluster3044 | -0.23 | 0.999 |

| | | |
|---|---|---|
| cluster3045 | -0.09 | 0.999 |
| cluster3047 | -0.14 | 0.999 |
| cluster3049 | 0.04 | 0.999 |
| cluster3051 | -0.01 | 0.999 |
| cluster3052 | -0.35 | 0.999 |
| cluster3054 | -0.02 | 0.999 |
| cluster3056 | 0.01 | 0.999 |
| cluster3057 | -0.07 | 0.999 |
| cluster3066 | 0.10 | 0.999 |
| cluster3072 | -0.21 | 0.999 |
| cluster3073 | -0.09 | 0.999 |
| cluster3074 | 0.24 | 0.999 |
| cluster3080 | -0.10 | 0.999 |
| cluster3082 | 0.38 | 0.999 |
| cluster3084 | -0.16 | 0.999 |
| cluster3086 | 0.12 | 0.999 |
| cluster3091 | -0.05 | 0.999 |
| cluster3092 | 0.31 | 0.999 |
| cluster3093 | 0.22 | 0.999 |
| cluster3097 | -0.07 | 0.999 |
| cluster3104 | -0.10 | 0.999 |
| cluster3105 | 0.02 | 0.999 |
| cluster3110 | 0.23 | 0.999 |
| cluster3115 | -0.16 | 0.999 |
| cluster3116 | 0.28 | 0.999 |
| cluster3120 | -0.01 | 0.999 |
| cluster3124 | -0.31 | 0.999 |
| cluster3130 | -0.02 | 0.999 |
| cluster3133 | -0.11 | 0.999 |
| cluster3134 | -0.02 | 0.999 |
| cluster3137 | -0.12 | 0.999 |
| cluster3140 | 0.25 | 0.999 |
| cluster3141 | -0.04 | 0.999 |
| cluster3143 | 0.17 | 0.999 |
| cluster3145 | 0.08 | 0.999 |
| cluster3148 | 0.04 | 0.999 |
| cluster3150 | -0.33 | 0.999 |
| cluster3153 | -0.29 | 0.999 |
| cluster3155 | -0.25 | 0.999 |
| cluster3158 | -0.17 | 0.999 |
| cluster3159 | -0.19 | 0.999 |
| cluster3161 | 0.02 | 0.999 |
| cluster3165 | -0.15 | 0.999 |
| cluster3169 | 0.05 | 0.999 |

| | | |
|---|---|---|
| cluster3170 | -0.29 | 0.999 |
| cluster3172 | 0.23 | 0.999 |
| cluster3174 | 0.04 | 0.999 |
| cluster3180 | 0.27 | 0.999 |
| cluster3183 | 0.31 | 0.999 |
| cluster3184 | 0.14 | 0.999 |
| cluster3188 | 0.31 | 0.999 |
| cluster3193 | -0.13 | 0.999 |
| cluster3201 | -0.25 | 0.999 |
| cluster3202 | 0.15 | 0.999 |
| cluster3204 | -0.04 | 0.999 |
| cluster3206 | 0.05 | 0.999 |
| cluster3208 | -0.25 | 0.999 |
| cluster3209 | 0.36 | 0.999 |
| cluster3210 | 0.06 | 0.999 |
| cluster3217 | -0.19 | 0.999 |
| cluster3220 | 0.24 | 0.999 |
| cluster3221 | -0.23 | 0.999 |
| cluster3225 | 0.04 | 0.999 |
| cluster3227 | -0.17 | 0.999 |
| cluster3228 | 0.26 | 0.999 |
| cluster3230 | -0.22 | 0.999 |
| cluster3232 | 0.40 | 0.999 |
| cluster3237 | -0.15 | 0.999 |
| cluster3247 | 0.09 | 0.999 |
| cluster3248 | 0.45 | 0.999 |
| cluster3251 | -0.27 | 0.999 |
| cluster3253 | 0.28 | 0.999 |
| cluster3255 | -0.14 | 0.999 |
| cluster3258 | -0.25 | 0.999 |
| cluster3261 | -0.01 | 0.999 |
| cluster3262 | -0.29 | 0.999 |
| cluster3267 | 0.14 | 0.999 |
| cluster3277 | -0.25 | 0.999 |
| cluster3278 | 0.03 | 0.999 |
| cluster3281 | 0.08 | 0.999 |
| cluster3282 | -0.36 | 0.999 |
| cluster3284 | 0.41 | 0.999 |
| cluster3285 | 0.10 | 0.999 |
| cluster3289 | 0.10 | 0.999 |
| cluster3290 | -0.18 | 0.999 |
| cluster3292 | 0.10 | 0.999 |
| cluster3295 | -0.05 | 0.999 |
| cluster3298 | -0.05 | 0.999 |

| | | |
|---|---|---|
| cluster3304 | 0.05 | 0.999 |
| cluster3307 | -0.08 | 0.999 |
| cluster3308 | 0.09 | 0.999 |
| cluster3317 | 0.39 | 0.999 |
| cluster3318 | 0.29 | 0.999 |
| cluster3320 | 0.25 | 0.999 |
| cluster3322 | 0.26 | 0.999 |
| cluster3324 | -0.11 | 0.999 |
| cluster3330 | 0.34 | 0.999 |
| cluster3331 | -0.13 | 0.999 |
| cluster3334 | 0.25 | 0.999 |
| cluster3335 | 0.28 | 0.999 |
| cluster3339 | -0.10 | 0.999 |
| cluster3340 | 0.22 | 0.999 |
| cluster3341 | 0.07 | 0.999 |
| cluster3344 | -0.03 | 0.999 |
| cluster3350 | -0.24 | 0.999 |
| cluster3352 | 0.10 | 0.999 |
| cluster3353 | -0.09 | 0.999 |
| cluster3354 | 0.17 | 0.999 |
| cluster3355 | -0.40 | 0.999 |
| cluster3373 | 0.27 | 0.999 |
| cluster3383 | 0.01 | 0.999 |
| cluster3384 | 0.23 | 0.999 |
| cluster3388 | -0.07 | 0.999 |
| cluster3396 | 0.18 | 0.999 |
| cluster3397 | -0.47 | 0.999 |
| cluster3401 | -0.02 | 0.999 |
| cluster3403 | -0.07 | 0.999 |
| cluster3407 | 0.15 | 0.999 |
| cluster3414 | -0.03 | 0.999 |
| cluster3415 | -0.15 | 0.999 |
| cluster3425 | -0.31 | 0.999 |
| cluster3426 | 0.07 | 0.999 |
| cluster3430 | 0.22 | 0.999 |
| cluster3431 | -0.42 | 0.999 |
| cluster3433 | -0.32 | 0.999 |
| cluster3434 | 0.13 | 0.999 |
| cluster3436 | 0.48 | 0.999 |
| cluster3441 | 0.09 | 0.999 |
| cluster3442 | 0.16 | 0.999 |
| cluster3444 | -0.02 | 0.999 |
| cluster3445 | 0.18 | 0.999 |
| cluster3446 | -0.04 | 0.999 |

| | | |
|---|---|---|
| cluster3453 | 0.26 | 0.999 |
| cluster3456 | -0.14 | 0.999 |
| cluster3460 | -0.14 | 0.999 |
| cluster3463 | -0.05 | 0.999 |
| cluster3466 | 0.03 | 0.999 |
| cluster3468 | -0.22 | 0.999 |
| cluster3469 | -0.05 | 0.999 |
| cluster3475 | -0.14 | 0.999 |
| cluster3483 | -0.43 | 0.999 |
| cluster3484 | 0.02 | 0.999 |
| cluster3488 | -0.21 | 0.999 |
| cluster3491 | 0.25 | 0.999 |
| cluster3492 | -0.09 | 0.999 |
| cluster3493 | 0.37 | 0.999 |
| cluster3509 | -0.22 | 0.999 |
| cluster3510 | 0.74 | 0.999 |
| cluster3514 | 0.03 | 0.999 |
| cluster3515 | -0.16 | 0.999 |
| cluster3518 | 0.45 | 0.999 |
| cluster3519 | 0.01 | 0.999 |
| cluster3520 | -0.07 | 0.999 |
| cluster3521 | -0.08 | 0.999 |
| cluster3522 | -0.09 | 0.999 |
| cluster3524 | -0.01 | 0.999 |
| cluster3527 | 0.31 | 0.999 |
| cluster3528 | -0.09 | 0.999 |
| cluster3530 | 0.15 | 0.999 |
| cluster3535 | 0.18 | 0.999 |
| cluster3536 | -0.11 | 0.999 |
| cluster3538 | -0.28 | 0.999 |
| cluster3540 | -0.40 | 0.999 |
| cluster3549 | -0.09 | 0.999 |
| cluster3551 | -0.13 | 0.999 |
| cluster3552 | 0.23 | 0.999 |
| cluster3553 | 0.34 | 0.999 |
| cluster3555 | -0.09 | 0.999 |
| cluster3560 | 0.19 | 0.999 |
| cluster3563 | -0.20 | 0.999 |
| cluster3565 | 0.35 | 0.999 |
| cluster3566 | -0.09 | 0.999 |
| cluster3573 | -0.09 | 0.999 |
| cluster3576 | -0.22 | 0.999 |
| cluster3580 | 0.36 | 0.999 |
| cluster3581 | 0.42 | 0.999 |

| | | |
|---|---|---|
| cluster3582 | -0.03 | 0.999 |
| cluster3586 | 0.24 | 0.999 |
| cluster3588 | 0.35 | 0.999 |
| cluster3589 | 0.01 | 0.999 |
| cluster3590 | -0.03 | 0.999 |
| cluster3596 | 0.12 | 0.999 |
| cluster3597 | -0.25 | 0.999 |
| cluster3598 | 0.31 | 0.999 |
| cluster3601 | 0.04 | 0.999 |
| cluster3603 | -0.05 | 0.999 |
| cluster3605 | 0.34 | 0.999 |
| cluster3610 | 0.03 | 0.999 |
| cluster3612 | -0.18 | 0.999 |
| cluster3615 | 0.15 | 0.999 |
| cluster3624 | 0.33 | 0.999 |
| cluster3626 | 0.09 | 0.999 |
| cluster3627 | 0.39 | 0.999 |
| cluster3634 | -0.23 | 0.999 |
| cluster3635 | 0.12 | 0.999 |
| cluster3636 | 0.14 | 0.999 |
| cluster3641 | -0.07 | 0.999 |
| cluster3643 | 0.03 | 0.999 |
| cluster3649 | -0.32 | 0.999 |
| cluster3655 | 0.11 | 0.999 |
| cluster3660 | -0.23 | 0.999 |
| cluster3664 | 0.10 | 0.999 |
| cluster3682 | -0.28 | 0.999 |
| cluster3684 | 0.07 | 0.999 |
| cluster3685 | -0.23 | 0.999 |
| cluster3690 | -0.10 | 0.999 |
| cluster3691 | 0.15 | 0.999 |
| cluster3697 | 0.15 | 0.999 |
| cluster3700 | 0.24 | 0.999 |
| cluster3706 | 0.08 | 0.999 |
| cluster3716 | -0.24 | 0.999 |
| cluster3722 | 0.21 | 0.999 |
| cluster3727 | -0.06 | 0.999 |
| cluster3729 | -0.05 | 0.999 |
| cluster3742 | 0.27 | 0.999 |
| cluster3744 | -0.13 | 0.999 |
| cluster3745 | 0.16 | 0.999 |
| cluster3751 | 0.28 | 0.999 |
| cluster3753 | 0.50 | 0.999 |
| cluster3756 | 0.13 | 0.999 |

| | | |
|---|---|---|
| cluster3761 | 0.28 | 0.999 |
| cluster3763 | 0.12 | 0.999 |
| cluster3768 | 0.16 | 0.999 |
| cluster3771 | 0.14 | 0.999 |
| cluster3775 | -0.18 | 0.999 |
| cluster3776 | 0.30 | 0.999 |
| cluster3777 | -0.38 | 0.999 |
| cluster3786 | 0.12 | 0.999 |
| cluster3791 | -0.10 | 0.999 |
| cluster3792 | 0.05 | 0.999 |
| cluster3797 | -0.37 | 0.999 |
| cluster3798 | 0.21 | 0.999 |
| cluster3799 | 0.19 | 0.999 |
| cluster3803 | 0.29 | 0.999 |
| cluster3807 | -0.17 | 0.999 |
| cluster3808 | -0.17 | 0.999 |
| cluster3816 | 0.22 | 0.999 |
| cluster3825 | 0.09 | 0.999 |
| cluster3829 | 0.31 | 0.999 |
| cluster3832 | 0.27 | 0.999 |
| cluster3835 | 0.29 | 0.999 |
| cluster3837 | 0.25 | 0.999 |
| cluster3841 | 0.24 | 0.999 |
| cluster3845 | 0.58 | 0.999 |
| cluster3847 | 0.37 | 0.999 |
| cluster3857 | 0.13 | 0.999 |
| cluster3862 | -0.03 | 0.999 |
| cluster3868 | 0.05 | 0.999 |
| cluster3872 | 0.37 | 0.999 |
| cluster3876 | -0.01 | 0.999 |
| cluster3883 | -0.54 | 0.999 |
| cluster3895 | 0.22 | 0.999 |
| cluster3896 | 0.04 | 0.999 |
| cluster3898 | 0.38 | 0.999 |
| cluster3903 | -0.11 | 0.999 |
| cluster3907 | -0.02 | 0.999 |
| cluster3911 | -0.09 | 0.999 |
| cluster3919 | 0.56 | 0.999 |
| cluster3923 | 0.26 | 0.999 |
| cluster3929 | -0.28 | 0.999 |
| cluster3932 | 0.01 | 0.999 |
| cluster3934 | -0.07 | 0.999 |
| cluster3935 | -0.13 | 0.999 |
| cluster3943 | 0.42 | 0.999 |

| | | |
|---|---|---|
| cluster3946 | -0.14 | 0.999 |
| cluster3947 | 0.40 | 0.999 |
| cluster3953 | -0.10 | 0.999 |
| cluster3959 | 0.02 | 0.999 |
| cluster3960 | 0.03 | 0.999 |
| cluster3962 | -0.37 | 0.999 |
| cluster3964 | -0.46 | 0.999 |
| cluster3967 | 0.07 | 0.999 |
| cluster3973 | 0.20 | 0.999 |
| cluster3977 | -0.02 | 0.999 |
| cluster3988 | 0.22 | 0.999 |
| cluster3990 | -0.20 | 0.999 |
| cluster3995 | -0.08 | 0.999 |
| cluster3996 | 0.01 | 0.999 |
| cluster4004 | 0.07 | 0.999 |
| cluster4006 | -0.23 | 0.999 |
| cluster4011 | 0.55 | 0.999 |
| cluster4012 | -0.09 | 0.999 |
| cluster4018 | -0.27 | 0.999 |
| cluster4021 | 0.39 | 0.999 |
| cluster4028 | 0.13 | 0.999 |
| cluster4034 | 0.42 | 0.999 |
| cluster4043 | -0.17 | 0.999 |
| cluster4049 | -0.32 | 0.999 |
| cluster4056 | 0.24 | 0.999 |
| cluster4065 | 0.28 | 0.999 |
| cluster4067 | -0.38 | 0.999 |
| cluster4072 | 0.37 | 0.999 |
| cluster4076 | 0.29 | 0.999 |
| cluster4077 | -0.09 | 0.999 |
| cluster4085 | 0.17 | 0.999 |
| cluster4088 | -0.14 | 0.999 |
| cluster4089 | -0.38 | 0.999 |
| cluster4092 | -0.25 | 0.999 |
| cluster4093 | -0.33 | 0.999 |
| cluster4094 | -0.05 | 0.999 |
| cluster4097 | 0.53 | 0.999 |
| cluster4102 | -0.18 | 0.999 |
| cluster4103 | -0.36 | 0.999 |
| cluster4124 | -0.15 | 0.999 |
| cluster3001 | 0.00 | 1.000 |
| cluster3238 | 0.00 | 1.000 |
| cluster3369 | 0.00 | 1.000 |
| cluster3921 | 0.00 | 1.000 |

| | | |
|---|---|---|
| cluster4005 | 0.00 | 1.000 |

**Supplementary Table 6.** edgeR results with RSEM counts - Parasites vs Saprophagous

| ID | log2FoldChan | Adjusted p value |
|---|---|---|
| cluster4091 | -3.78 | 0.829 |
| cluster3457 | -3.45 | 0.829 |
| cluster3099 | -2.12 | 1.000 |
| cluster3194 | -2.28 | 1.000 |
| cluster3419 | -2.04 | 1.000 |
| cluster3024 | -2.15 | 1.000 |
| cluster3616 | 4.64 | 1.000 |
| cluster3821 | -2.17 | 1.000 |
| cluster3004 | -2.01 | 1.000 |
| cluster3111 | -2.57 | 1.000 |
| cluster3026 | -2.40 | 1.000 |
| cluster3885 | -2.41 | 1.000 |
| cluster3128 | -1.75 | 1.000 |
| cluster3782 | -1.96 | 1.000 |
| cluster3374 | 4.03 | 1.000 |
| cluster3802 | 3.83 | 1.000 |
| cluster3231 | -2.55 | 1.000 |
| cluster3741 | -1.88 | 1.000 |
| cluster3529 | -2.20 | 1.000 |
| cluster3859 | -2.43 | 1.000 |
| cluster4101 | 4.94 | 1.000 |
| cluster3858 | 2.96 | 1.000 |
| cluster3822 | 4.17 | 1.000 |
| cluster3309 | -1.87 | 1.000 |
| cluster4116 | 5.19 | 1.000 |
| cluster3891 | 3.11 | 1.000 |
| cluster3639 | 4.40 | 1.000 |
| cluster3547 | 2.41 | 1.000 |
| cluster3760 | 2.61 | 1.000 |
| cluster4074 | 4.11 | 1.000 |
| cluster3427 | -1.94 | 1.000 |
| cluster3428 | -1.70 | 1.000 |
| cluster2904 | -1.82 | 1.000 |
| cluster3567 | -2.17 | 1.000 |
| cluster3118 | -1.59 | 1.000 |
| cluster3975 | 3.41 | 1.000 |
| cluster3123 | -1.52 | 1.000 |
| cluster4127 | 4.48 | 1.000 |
| cluster3689 | 4.13 | 1.000 |
| cluster3146 | -1.61 | 1.000 |
| cluster4078 | 2.59 | 1.000 |
| cluster4010 | 2.97 | 1.000 |

| | | |
|---|---|---|
| cluster3842 | -1.65 | 1.000 |
| cluster3709 | -1.50 | 1.000 |
| cluster3195 | -1.52 | 1.000 |
| cluster3652 | 2.53 | 1.000 |
| cluster3301 | 2.33 | 1.000 |
| cluster3408 | -1.53 | 1.000 |
| cluster2925 | -1.87 | 1.000 |
| cluster2858 | -1.40 | 1.000 |
| cluster3715 | 2.87 | 1.000 |
| cluster3402 | -1.40 | 1.000 |
| cluster3607 | -1.73 | 1.000 |
| cluster3379 | 2.56 | 1.000 |
| cluster3780 | 4.28 | 1.000 |
| cluster3108 | -1.72 | 1.000 |
| cluster3889 | 3.00 | 1.000 |
| cluster4108 | 2.80 | 1.000 |
| cluster3319 | -1.72 | 1.000 |
| cluster3399 | -1.49 | 1.000 |
| cluster3046 | -1.43 | 1.000 |
| cluster3950 | 1.98 | 1.000 |
| cluster3235 | 2.94 | 1.000 |
| cluster3926 | 3.49 | 1.000 |
| cluster3778 | 2.34 | 1.000 |
| cluster2885 | -1.66 | 1.000 |
| cluster3151 | -1.35 | 1.000 |
| cluster3283 | -1.22 | 1.000 |
| cluster3785 | -1.58 | 1.000 |
| cluster4087 | 3.86 | 1.000 |
| cluster2961 | 1.96 | 1.000 |
| cluster3167 | -1.59 | 1.000 |
| cluster2878 | -1.39 | 1.000 |
| cluster3810 | 1.94 | 1.000 |
| cluster3704 | 2.20 | 1.000 |
| cluster3550 | -1.23 | 1.000 |
| cluster3286 | -1.20 | 1.000 |
| cluster3100 | -1.80 | 1.000 |
| cluster2934 | -1.27 | 1.000 |
| cluster3005 | -1.31 | 1.000 |
| cluster3812 | -1.48 | 1.000 |
| cluster2924 | -1.69 | 1.000 |
| cluster3250 | -1.56 | 1.000 |
| cluster3420 | -1.51 | 1.000 |
| cluster4032 | -1.32 | 1.000 |
| cluster3037 | -1.38 | 1.000 |

| | | |
|---|---|---|
| cluster3106 | -1.24 | 1.000 |
| cluster3561 | -1.37 | 1.000 |
| cluster3852 | 2.15 | 1.000 |
| cluster2894 | -1.56 | 1.000 |
| cluster4040 | 2.00 | 1.000 |
| cluster3770 | 3.60 | 1.000 |
| cluster3971 | 2.46 | 1.000 |
| cluster3112 | -1.49 | 1.000 |
| cluster3707 | 2.41 | 1.000 |
| cluster3930 | 1.86 | 1.000 |
| cluster4098 | 2.29 | 1.000 |
| cluster3746 | 1.85 | 1.000 |
| cluster2908 | -1.20 | 1.000 |
| cluster3818 | 2.13 | 1.000 |
| cluster3827 | 2.32 | 1.000 |
| cluster3461 | 3.30 | 1.000 |
| cluster3240 | 1.83 | 1.000 |
| cluster3774 | -1.15 | 1.000 |
| cluster3064 | -1.27 | 1.000 |
| cluster3658 | -1.26 | 1.000 |
| cluster3609 | -1.56 | 1.000 |
| cluster3779 | 1.77 | 1.000 |
| cluster3259 | -1.12 | 1.000 |
| cluster2982 | -1.23 | 1.000 |
| cluster4118 | -1.75 | 1.000 |
| cluster3090 | -1.58 | 1.000 |
| cluster3670 | 2.15 | 1.000 |
| cluster3187 | -1.50 | 1.000 |
| cluster4119 | 2.79 | 1.000 |
| cluster3348 | -1.17 | 1.000 |
| cluster3504 | 2.29 | 1.000 |
| cluster3065 | -1.24 | 1.000 |
| cluster3585 | -1.12 | 1.000 |
| cluster4104 | 2.22 | 1.000 |
| cluster3103 | 2.62 | 1.000 |
| cluster3015 | -1.33 | 1.000 |
| cluster4031 | 1.84 | 1.000 |
| cluster3738 | 2.46 | 1.000 |
| cluster4009 | -1.26 | 1.000 |
| cluster4081 | 2.18 | 1.000 |
| cluster3546 | 2.97 | 1.000 |
| cluster4120 | -1.51 | 1.000 |
| cluster4111 | -1.55 | 1.000 |
| cluster3838 | -1.27 | 1.000 |

| | | |
|---|---|---|
| cluster3890 | 2.00 | 1.000 |
| cluster3297 | 1.96 | 1.000 |
| cluster3368 | -1.04 | 1.000 |
| cluster3164 | -1.33 | 1.000 |
| cluster3329 | 2.30 | 1.000 |
| cluster3732 | 2.13 | 1.000 |
| cluster3892 | 2.13 | 1.000 |
| cluster3513 | -1.08 | 1.000 |
| cluster3276 | -1.03 | 1.000 |
| cluster2959 | -1.04 | 1.000 |
| cluster3476 | 2.09 | 1.000 |
| cluster3989 | -1.14 | 1.000 |
| cluster3356 | 1.62 | 1.000 |
| cluster3949 | -1.26 | 1.000 |
| cluster3901 | 2.55 | 1.000 |
| cluster3328 | -1.00 | 1.000 |
| cluster2957 | -1.31 | 1.000 |
| cluster3287 | -1.11 | 1.000 |
| cluster3314 | -1.35 | 1.000 |
| cluster2933 | -1.11 | 1.000 |
| cluster2902 | 2.44 | 1.000 |
| cluster3941 | -0.96 | 1.000 |
| cluster2918 | -1.11 | 1.000 |
| cluster3559 | -1.20 | 1.000 |
| cluster3534 | -1.15 | 1.000 |
| cluster4057 | 2.01 | 1.000 |
| cluster3568 | -1.34 | 1.000 |
| cluster3272 | -1.58 | 1.000 |
| cluster3866 | 2.25 | 1.000 |
| cluster3224 | -1.01 | 1.000 |
| cluster4016 | 1.77 | 1.000 |
| cluster3755 | 2.57 | 1.000 |
| cluster2974 | -1.01 | 1.000 |
| cluster3070 | -0.99 | 1.000 |
| cluster2994 | 1.57 | 1.000 |
| cluster3263 | -1.01 | 1.000 |
| cluster3554 | -1.16 | 1.000 |
| cluster3280 | -1.03 | 1.000 |
| cluster4039 | -1.06 | 1.000 |
| cluster3191 | 2.26 | 1.000 |
| cluster3405 | -1.02 | 1.000 |
| cluster3833 | -1.23 | 1.000 |
| cluster3315 | 1.42 | 1.000 |
| cluster3865 | 2.11 | 1.000 |

| | | |
|---|---|---|
| cluster3983 | 2.10 | 1.000 |
| cluster3851 | 2.00 | 1.000 |
| cluster3212 | 1.79 | 1.000 |
| cluster3300 | -0.99 | 1.000 |
| cluster2905 | -1.00 | 1.000 |
| cluster3175 | 2.12 | 1.000 |
| cluster3239 | -0.87 | 1.000 |
| cluster3787 | 2.05 | 1.000 |
| cluster2977 | -1.11 | 1.000 |
| cluster3101 | 1.37 | 1.000 |
| cluster4023 | -1.10 | 1.000 |
| cluster3571 | 1.76 | 1.000 |
| cluster3583 | -1.10 | 1.000 |
| cluster3648 | -0.94 | 1.000 |
| cluster3578 | -1.09 | 1.000 |
| cluster3501 | 1.89 | 1.000 |
| cluster3177 | 1.92 | 1.000 |
| cluster3022 | -1.09 | 1.000 |
| cluster4002 | 1.99 | 1.000 |
| cluster3545 | 1.81 | 1.000 |
| cluster3302 | -1.03 | 1.000 |
| cluster3478 | -0.93 | 1.000 |
| cluster3179 | 1.93 | 1.000 |
| cluster3417 | 1.48 | 1.000 |
| cluster3487 | 2.14 | 1.000 |
| cluster3698 | -0.88 | 1.000 |
| cluster3936 | 1.84 | 1.000 |
| cluster3439 | -1.09 | 1.000 |
| cluster3556 | 1.94 | 1.000 |
| cluster3848 | -1.17 | 1.000 |
| cluster3562 | -1.07 | 1.000 |
| cluster3762 | -1.01 | 1.000 |
| cluster3611 | 1.91 | 1.000 |
| cluster3525 | -0.92 | 1.000 |
| cluster4062 | -1.25 | 1.000 |
| cluster3316 | -1.07 | 1.000 |
| cluster2962 | 1.44 | 1.000 |
| cluster4058 | 1.76 | 1.000 |
| cluster4055 | 1.85 | 1.000 |
| cluster3077 | 1.51 | 1.000 |
| cluster3790 | -1.09 | 1.000 |
| cluster3199 | -0.94 | 1.000 |
| cluster2923 | -0.88 | 1.000 |
| cluster3068 | -0.83 | 1.000 |

| | | |
|---|---|---|
| cluster3406 | 1.26 | 1.000 |
| cluster3160 | 1.41 | 1.000 |
| cluster2901 | -1.10 | 1.000 |
| cluster4095 | 1.72 | 1.000 |
| cluster4042 | 1.85 | 1.000 |
| cluster4079 | 1.50 | 1.000 |
| cluster3855 | 1.64 | 1.000 |
| cluster4086 | 1.67 | 1.000 |
| cluster3853 | 1.99 | 1.000 |
| cluster3516 | 2.11 | 1.000 |
| cluster3113 | -0.93 | 1.000 |
| cluster3256 | 1.15 | 1.000 |
| cluster4048 | 2.09 | 1.000 |
| cluster3713 | 1.62 | 1.000 |
| cluster2949 | -0.81 | 1.000 |
| cluster3899 | 1.74 | 1.000 |
| cluster3631 | 1.70 | 1.000 |
| cluster2906 | -0.91 | 1.000 |
| cluster3600 | -0.92 | 1.000 |
| cluster4117 | -1.26 | 1.000 |
| cluster4090 | -1.21 | 1.000 |
| cluster3474 | 1.95 | 1.000 |
| cluster3182 | 1.69 | 1.000 |
| cluster3171 | -0.91 | 1.000 |
| cluster3377 | -1.02 | 1.000 |
| cluster3558 | -0.97 | 1.000 |
| cluster3666 | 2.34 | 1.000 |
| cluster3448 | 1.80 | 1.000 |
| cluster3769 | -0.83 | 1.000 |
| cluster4003 | -1.02 | 1.000 |
| cluster2926 | 1.25 | 1.000 |
| cluster3357 | -0.84 | 1.000 |
| cluster3121 | -0.90 | 1.000 |
| cluster2956 | -0.83 | 1.000 |
| cluster3326 | -0.86 | 1.000 |
| cluster3008 | 1.61 | 1.000 |
| cluster2919 | -0.80 | 1.000 |
| cluster3956 | 1.68 | 1.000 |
| cluster3343 | -1.07 | 1.000 |
| cluster3748 | 1.23 | 1.000 |
| cluster3233 | 1.99 | 1.000 |
| cluster3702 | -0.96 | 1.000 |
| cluster3913 | -0.99 | 1.000 |
| cluster3772 | -0.94 | 1.000 |

| | | |
|---|---|---|
| cluster3386 | 1.72 | 1.000 |
| cluster3843 | 1.23 | 1.000 |
| cluster3455 | -0.88 | 1.000 |
| cluster3909 | 1.56 | 1.000 |
| cluster3874 | -0.78 | 1.000 |
| cluster3505 | -0.97 | 1.000 |
| cluster3266 | -0.77 | 1.000 |
| cluster2953 | -0.74 | 1.000 |
| cluster3189 | -0.93 | 1.000 |
| cluster3422 | -0.98 | 1.000 |
| cluster3477 | -0.83 | 1.000 |
| cluster3871 | 1.58 | 1.000 |
| cluster4045 | 1.86 | 1.000 |
| cluster3978 | 1.38 | 1.000 |
| cluster3618 | -1.08 | 1.000 |
| cluster2973 | -0.98 | 1.000 |
| cluster3451 | -1.17 | 1.000 |
| cluster3207 | 1.26 | 1.000 |
| cluster3166 | 1.07 | 1.000 |
| cluster3031 | -0.76 | 1.000 |
| cluster3758 | 1.33 | 1.000 |
| cluster3933 | -0.94 | 1.000 |
| cluster3245 | -0.98 | 1.000 |
| cluster3114 | -0.80 | 1.000 |
| cluster3942 | -0.92 | 1.000 |
| cluster2987 | -0.72 | 1.000 |
| cluster3413 | 1.67 | 1.000 |
| cluster3728 | 1.28 | 1.000 |
| cluster3924 | 1.59 | 1.000 |
| cluster3229 | -0.82 | 1.000 |
| cluster3938 | 1.42 | 1.000 |
| cluster3310 | -0.74 | 1.000 |
| cluster3359 | 1.53 | 1.000 |
| cluster3957 | 1.56 | 1.000 |
| cluster3575 | -1.01 | 1.000 |
| cluster3196 | -0.72 | 1.000 |
| cluster3243 | -0.79 | 1.000 |
| cluster3038 | -0.92 | 1.000 |
| cluster4107 | 1.66 | 1.000 |
| cluster3759 | -0.77 | 1.000 |
| cluster4033 | 1.67 | 1.000 |
| cluster2992 | -0.73 | 1.000 |
| cluster3117 | 1.60 | 1.000 |
| cluster3789 | -0.86 | 1.000 |

| | | |
|---|---|---|
| cluster3129 | -1.02 | 1.000 |
| cluster3249 | -0.79 | 1.000 |
| cluster3135 | -0.97 | 1.000 |
| cluster3390 | -0.74 | 1.000 |
| cluster4069 | 1.34 | 1.000 |
| cluster3718 | 1.38 | 1.000 |
| cluster3502 | 1.54 | 1.000 |
| cluster2872 | -0.81 | 1.000 |
| cluster3972 | 1.49 | 1.000 |
| cluster3157 | -0.70 | 1.000 |
| cluster4001 | 1.69 | 1.000 |
| cluster3178 | 1.40 | 1.000 |
| cluster3410 | 1.27 | 1.000 |
| cluster4121 | 1.53 | 1.000 |
| cluster3192 | 1.08 | 1.000 |
| cluster3050 | -0.74 | 1.000 |
| cluster3839 | -0.95 | 1.000 |
| cluster3336 | -0.76 | 1.000 |
| cluster3215 | -0.66 | 1.000 |
| cluster4075 | -0.92 | 1.000 |
| cluster3241 | -0.78 | 1.000 |
| cluster3613 | -0.77 | 1.000 |
| cluster3303 | 1.49 | 1.000 |
| cluster3976 | 1.17 | 1.000 |
| cluster3965 | 1.40 | 1.000 |
| cluster3440 | -0.69 | 1.000 |
| cluster3736 | -0.85 | 1.000 |
| cluster3883 | -0.98 | 1.000 |
| cluster3897 | 1.26 | 1.000 |
| cluster3063 | 1.19 | 1.000 |
| cluster2893 | 1.98 | 1.000 |
| cluster3269 | -0.84 | 1.000 |
| cluster3032 | -0.68 | 1.000 |
| cluster3321 | -0.77 | 1.000 |
| cluster3681 | 1.45 | 1.000 |
| cluster3604 | 1.54 | 1.000 |
| cluster3678 | 1.03 | 1.000 |
| cluster4068 | 1.40 | 1.000 |
| cluster4027 | 1.38 | 1.000 |
| cluster3940 | 1.10 | 1.000 |
| cluster3877 | 1.05 | 1.000 |
| cluster3378 | 1.12 | 1.000 |
| cluster3449 | 1.01 | 1.000 |
| cluster3260 | 1.06 | 1.000 |

| | | |
|---|---|---|
| cluster3717 | 1.04 | 1.000 |
| cluster3036 | -0.81 | 1.000 |
| cluster3305 | -0.85 | 1.000 |
| cluster3543 | 1.54 | 1.000 |
| cluster3726 | -0.69 | 1.000 |
| cluster3645 | 2.17 | 1.000 |
| cluster3342 | 1.42 | 1.000 |
| cluster3069 | -0.64 | 1.000 |
| cluster3494 | 1.08 | 1.000 |
| cluster3380 | -0.71 | 1.000 |
| cluster3019 | 1.35 | 1.000 |
| cluster3482 | 1.17 | 1.000 |
| cluster3805 | 1.46 | 1.000 |
| cluster3002 | -0.63 | 1.000 |
| cluster4112 | 1.48 | 1.000 |
| cluster2950 | -0.64 | 1.000 |
| cluster3905 | 2.16 | 1.000 |
| cluster3651 | 1.68 | 1.000 |
| cluster3693 | 1.46 | 1.000 |
| cluster3854 | 1.37 | 1.000 |
| cluster3906 | 1.27 | 1.000 |
| cluster3984 | -0.86 | 1.000 |
| cluster2941 | -0.64 | 1.000 |
| cluster3824 | 1.50 | 1.000 |
| cluster3932 | 1.59 | 1.000 |
| cluster4076 | 1.44 | 1.000 |
| cluster3071 | -0.63 | 1.000 |
| cluster4038 | 1.34 | 1.000 |
| cluster3649 | -0.66 | 1.000 |
| cluster3454 | 1.00 | 1.000 |
| cluster3020 | -0.84 | 1.000 |
| cluster3860 | -0.66 | 1.000 |
| cluster3040 | -0.64 | 1.000 |
| cluster2946 | -0.69 | 1.000 |
| cluster3273 | -0.60 | 1.000 |
| cluster2990 | -0.75 | 1.000 |
| cluster3055 | -0.65 | 1.000 |
| cluster3299 | 1.35 | 1.000 |
| cluster2871 | -0.63 | 1.000 |
| cluster3211 | 1.28 | 1.000 |
| cluster3085 | -0.74 | 1.000 |
| cluster3367 | 1.32 | 1.000 |
| cluster4030 | 1.44 | 1.000 |
| cluster2965 | -0.77 | 1.000 |

| | | |
|---|---|---|
| cluster2997 | -0.74 | 1.000 |
| cluster3497 | 0.96 | 1.000 |
| cluster3443 | 1.28 | 1.000 |
| cluster4052 | 1.16 | 1.000 |
| cluster3282 | -0.70 | 1.000 |
| cluster3306 | 1.64 | 1.000 |
| cluster2897 | -0.64 | 1.000 |
| cluster3124 | -0.63 | 1.000 |
| cluster3962 | -0.71 | 1.000 |
| cluster3394 | -0.60 | 1.000 |
| cluster3540 | -0.73 | 1.000 |
| cluster3814 | -0.64 | 1.000 |
| cluster3483 | -0.76 | 1.000 |
| cluster2935 | 1.01 | 1.000 |
| cluster4125 | 1.64 | 1.000 |
| cluster3752 | 1.66 | 1.000 |
| cluster3222 | 0.81 | 1.000 |
| cluster3991 | 1.18 | 1.000 |
| cluster3797 | -0.72 | 1.000 |
| cluster3541 | -0.60 | 1.000 |
| cluster3197 | -0.62 | 1.000 |
| cluster3433 | -0.62 | 1.000 |
| cluster3431 | -0.72 | 1.000 |
| cluster3252 | 1.64 | 1.000 |
| cluster2860 | 1.05 | 1.000 |
| cluster4054 | 1.26 | 1.000 |
| cluster3777 | -0.66 | 1.000 |
| cluster3619 | 1.01 | 1.000 |
| cluster3176 | -0.69 | 1.000 |
| cluster3349 | 1.12 | 1.000 |
| cluster3686 | 1.26 | 1.000 |
| cluster3397 | -0.76 | 1.000 |
| cluster3333 | 1.21 | 1.000 |
| cluster3323 | 1.11 | 1.000 |
| cluster3675 | -0.59 | 1.000 |
| cluster2876 | 1.54 | 1.000 |
| cluster3355 | -0.71 | 1.000 |
| cluster3385 | 1.40 | 1.000 |
| cluster4071 | 1.37 | 1.000 |
| cluster3186 | 1.47 | 1.000 |
| cluster3737 | -0.54 | 1.000 |
| cluster3244 | -0.56 | 1.000 |
| cluster3041 | 1.02 | 1.000 |
| cluster3262 | -0.63 | 1.000 |

| | | |
|---|---|---|
| cluster3955 | 1.30 | 1.000 |
| cluster3840 | 1.08 | 1.000 |
| cluster3964 | -0.73 | 1.000 |
| cluster3749 | 1.41 | 1.000 |
| cluster3258 | -0.60 | 1.000 |
| cluster3735 | 1.06 | 1.000 |
| cluster2928 | 1.11 | 1.000 |
| cluster3834 | 1.33 | 1.000 |
| cluster3205 | -0.57 | 1.000 |
| cluster3425 | -0.60 | 1.000 |
| cluster3400 | -0.56 | 1.000 |
| cluster3927 | 1.14 | 1.000 |
| cluster2874 | -0.59 | 1.000 |
| cluster3512 | 1.05 | 1.000 |
| cluster3201 | -0.56 | 1.000 |
| cluster3246 | 0.88 | 1.000 |
| cluster3170 | -0.54 | 1.000 |
| cluster3052 | -0.72 | 1.000 |
| cluster3010 | -0.61 | 1.000 |
| cluster3274 | 1.23 | 1.000 |
| cluster4018 | -0.61 | 1.000 |
| cluster3509 | -0.54 | 1.000 |
| cluster2867 | 1.32 | 1.000 |
| cluster2910 | -0.67 | 1.000 |
| cluster3886 | 1.17 | 1.000 |
| cluster3629 | 1.29 | 1.000 |
| cluster3496 | -0.55 | 1.000 |
| cluster3075 | 0.91 | 1.000 |
| cluster3922 | 1.34 | 1.000 |
| cluster3634 | -0.51 | 1.000 |
| cluster3150 | -0.64 | 1.000 |
| cluster3277 | -0.59 | 1.000 |
| cluster2912 | 0.98 | 1.000 |
| cluster3783 | 1.19 | 1.000 |
| cluster3458 | -0.52 | 1.000 |
| cluster2968 | -0.49 | 1.000 |
| cluster3904 | 1.44 | 1.000 |
| cluster3109 | 1.32 | 1.000 |
| cluster3346 | 0.96 | 1.000 |
| cluster3479 | 1.14 | 1.000 |
| cluster3981 | 1.07 | 1.000 |
| cluster3060 | 1.25 | 1.000 |
| cluster4022 | 1.45 | 1.000 |
| cluster3153 | -0.62 | 1.000 |

| | | |
|---|---|---|
| cluster2995 | 1.03 | 1.000 |
| cluster3572 | 1.52 | 1.000 |
| cluster4013 | 1.90 | 1.000 |
| cluster3392 | 1.24 | 1.000 |
| cluster3155 | -0.60 | 1.000 |
| cluster3415 | -0.54 | 1.000 |
| cluster4051 | 1.41 | 1.000 |
| cluster3144 | 1.22 | 1.000 |
| cluster3251 | -0.60 | 1.000 |
| cluster3067 | -0.50 | 1.000 |
| cluster3682 | -0.60 | 1.000 |
| cluster2989 | -0.54 | 1.000 |
| cluster3944 | 1.10 | 1.000 |
| cluster4006 | -0.60 | 1.000 |
| cluster3620 | 0.84 | 1.000 |
| cluster3880 | 1.43 | 1.000 |
| cluster4066 | 1.03 | 1.000 |
| cluster3227 | -0.56 | 1.000 |
| cluster3719 | 1.12 | 1.000 |
| cluster3078 | 0.73 | 1.000 |
| cluster3688 | 1.12 | 1.000 |
| cluster3929 | -0.60 | 1.000 |
| cluster2877 | -0.47 | 1.000 |
| cluster4014 | 1.17 | 1.000 |
| cluster3987 | 0.91 | 1.000 |
| cluster4123 | 1.20 | 1.000 |
| cluster3663 | 1.06 | 1.000 |
| cluster3044 | -0.58 | 1.000 |
| cluster3012 | 1.19 | 1.000 |
| cluster3089 | 0.89 | 1.000 |
| cluster4053 | 1.08 | 1.000 |
| cluster3156 | 1.12 | 1.000 |
| cluster3030 | -0.57 | 1.000 |
| cluster3685 | -0.60 | 1.000 |
| cluster3703 | 0.76 | 1.000 |
| cluster4100 | 1.06 | 1.000 |
| cluster3221 | -0.57 | 1.000 |
| cluster3481 | 1.14 | 1.000 |
| cluster3710 | 1.06 | 1.000 |
| cluster4092 | -0.57 | 1.000 |
| cluster3029 | -0.55 | 1.000 |
| cluster3576 | -0.54 | 1.000 |
| cluster3362 | 0.87 | 1.000 |
| cluster3614 | 1.20 | 1.000 |

| | | |
|---|---|---|
| cluster4113 | 0.98 | 1.000 |
| cluster3747 | 0.88 | 1.000 |
| cluster3915 | 1.33 | 1.000 |
| cluster2890 | -0.47 | 1.000 |
| cluster3881 | 0.97 | 1.000 |
| cluster3593 | 1.06 | 1.000 |
| cluster4060 | 1.20 | 1.000 |
| cluster3072 | -0.60 | 1.000 |
| cluster3597 | -0.48 | 1.000 |
| cluster4102 | -0.53 | 1.000 |
| cluster3765 | 0.84 | 1.000 |
| cluster3587 | 1.33 | 1.000 |
| cluster3506 | 1.02 | 1.000 |
| cluster3193 | -0.51 | 1.000 |
| cluster3488 | -0.53 | 1.000 |
| cluster3016 | -0.57 | 1.000 |
| cluster3468 | -0.47 | 1.000 |
| cluster3808 | -0.53 | 1.000 |
| cluster3985 | 0.84 | 1.000 |
| cluster3350 | -0.55 | 1.000 |
| cluster3515 | -0.57 | 1.000 |
| cluster3456 | -0.45 | 1.000 |
| cluster3598 | 1.50 | 1.000 |
| cluster3804 | 1.14 | 1.000 |
| cluster4067 | 1.58 | 1.000 |
| cluster3723 | 0.77 | 1.000 |
| cluster3159 | -0.52 | 1.000 |
| cluster3115 | -0.54 | 1.000 |
| cluster3084 | -0.43 | 1.000 |
| cluster3612 | -0.51 | 1.000 |
| cluster3500 | 1.35 | 1.000 |
| cluster4049 | -0.68 | 1.000 |
| cluster3754 | 0.78 | 1.000 |
| cluster3716 | -0.53 | 1.000 |
| cluster3003 | -0.53 | 1.000 |
| cluster3035 | -0.44 | 1.000 |
| cluster3794 | 0.99 | 1.000 |
| cluster4103 | -0.52 | 1.000 |
| cluster3061 | 0.75 | 1.000 |
| cluster3364 | 1.09 | 1.000 |
| cluster3990 | -0.44 | 1.000 |
| cluster3208 | -0.45 | 1.000 |
| cluster2964 | -0.43 | 1.000 |
| cluster3236 | 1.02 | 1.000 |

| | | |
|---|---|---|
| cluster4044 | 1.01 | 1.000 |
| cluster3563 | -0.47 | 1.000 |
| cluster3659 | 1.13 | 1.000 |
| cluster3784 | 0.77 | 1.000 |
| cluster2931 | -0.50 | 1.000 |
| cluster3137 | -0.54 | 1.000 |
| cluster2863 | 0.65 | 1.000 |
| cluster3775 | -0.51 | 1.000 |
| cluster2861 | -0.44 | 1.000 |
| cluster3460 | -0.49 | 1.000 |
| cluster3819 | 1.23 | 1.000 |
| cluster3185 | 1.25 | 1.000 |
| cluster2937 | -0.44 | 1.000 |
| cluster3291 | 0.75 | 1.000 |
| cluster3864 | 0.93 | 1.000 |
| cluster3946 | -0.48 | 1.000 |
| cluster3353 | -0.43 | 1.000 |
| cluster4012 | -0.49 | 1.000 |
| cluster3257 | 0.67 | 1.000 |
| cluster2967 | -0.45 | 1.000 |
| cluster2954 | -0.44 | 1.000 |
| cluster3200 | 0.73 | 1.000 |
| cluster3447 | 0.91 | 1.000 |
| cluster3480 | 0.96 | 1.000 |
| cluster3080 | -0.43 | 1.000 |
| cluster3538 | -0.45 | 1.000 |
| cluster3139 | 1.09 | 1.000 |
| cluster3104 | -0.37 | 1.000 |
| cluster3467 | 0.93 | 1.000 |
| cluster3903 | -0.47 | 1.000 |
| cluster2952 | -0.47 | 1.000 |
| cluster2921 | -0.43 | 1.000 |
| cluster3163 | 1.04 | 1.000 |
| cluster2951 | -0.46 | 1.000 |
| cluster3511 | 1.07 | 1.000 |
| cluster3793 | 0.96 | 1.000 |
| cluster3198 | 0.95 | 1.000 |
| cluster4035 | 0.74 | 1.000 |
| cluster3998 | 1.23 | 1.000 |
| cluster3646 | 1.00 | 1.000 |
| cluster3255 | -0.48 | 1.000 |
| cluster3831 | 0.93 | 1.000 |
| cluster3533 | 0.75 | 1.000 |
| cluster3701 | 0.68 | 1.000 |

| | | |
|---|---|---|
| cluster3520 | -0.52 | 1.000 |
| cluster3908 | 0.83 | 1.000 |
| cluster3073 | -0.41 | 1.000 |
| cluster3640 | 1.55 | 1.000 |
| cluster3744 | -0.44 | 1.000 |
| cluster3521 | -0.44 | 1.000 |
| cluster2983 | -0.44 | 1.000 |
| cluster3727 | -0.44 | 1.000 |
| cluster2943 | -0.41 | 1.000 |
| cluster3142 | 0.85 | 1.000 |
| cluster3471 | 1.00 | 1.000 |
| cluster3660 | -0.47 | 1.000 |
| cluster3387 | 1.02 | 1.000 |
| cluster3331 | -0.43 | 1.000 |
| cluster3731 | 0.75 | 1.000 |
| cluster3360 | 0.83 | 1.000 |
| cluster3849 | 0.91 | 1.000 |
| cluster4050 | 0.91 | 1.000 |
| cluster4088 | -0.45 | 1.000 |
| cluster3912 | 0.59 | 1.000 |
| cluster4084 | 0.78 | 1.000 |
| cluster3935 | -0.43 | 1.000 |
| cluster3791 | -0.44 | 1.000 |
| cluster3158 | -0.42 | 1.000 |
| cluster4026 | 0.88 | 1.000 |
| cluster3133 | -0.42 | 1.000 |
| cluster3953 | -0.43 | 1.000 |
| cluster2866 | -0.45 | 1.000 |
| cluster3492 | -0.42 | 1.000 |
| cluster3884 | 1.11 | 1.000 |
| cluster3293 | 0.62 | 1.000 |
| cluster3911 | -0.43 | 1.000 |
| cluster3088 | 0.75 | 1.000 |
| cluster3807 | -0.37 | 1.000 |
| cluster3677 | 0.68 | 1.000 |
| cluster3968 | 0.83 | 1.000 |
| cluster2882 | -0.35 | 1.000 |
| cluster4061 | 0.83 | 1.000 |
| cluster3573 | -0.42 | 1.000 |
| cluster3217 | -0.37 | 1.000 |
| cluster3324 | -0.36 | 1.000 |
| cluster3823 | 0.68 | 1.000 |
| cluster3555 | -0.44 | 1.000 |
| cluster3307 | -0.42 | 1.000 |

| | | |
|---|---|---|
| cluster3295 | -0.34 | 1.000 |
| cluster3830 | 0.69 | 1.000 |
| cluster3296 | 1.13 | 1.000 |
| cluster2942 | -0.40 | 1.000 |
| cluster3995 | -0.43 | 1.000 |
| cluster3393 | 1.13 | 1.000 |
| cluster3475 | -0.41 | 1.000 |
| cluster2855 | -0.43 | 1.000 |
| cluster3290 | -0.39 | 1.000 |
| cluster3773 | 0.71 | 1.000 |
| cluster3047 | -0.40 | 1.000 |
| cluster3173 | 0.86 | 1.000 |
| cluster3444 | -0.35 | 1.000 |
| cluster3566 | -0.40 | 1.000 |
| cluster3536 | -0.34 | 1.000 |
| cluster3028 | -0.40 | 1.000 |
| cluster2892 | 0.63 | 1.000 |
| cluster3522 | -0.40 | 1.000 |
| cluster3054 | -0.36 | 1.000 |
| cluster4077 | -0.41 | 1.000 |
| cluster3045 | -0.35 | 1.000 |
| cluster4094 | -0.37 | 1.000 |
| cluster3404 | 0.64 | 1.000 |
| cluster4029 | 1.03 | 1.000 |
| cluster3409 | 0.84 | 1.000 |
| cluster2856 | -0.41 | 1.000 |
| cluster4083 | 0.87 | 1.000 |
| cluster3313 | 0.83 | 1.000 |
| cluster3053 | 0.76 | 1.000 |
| cluster3365 | 0.91 | 1.000 |
| cluster3934 | -0.38 | 1.000 |
| cluster3729 | -0.38 | 1.000 |
| cluster3095 | 0.65 | 1.000 |
| cluster3238 | -0.32 | 1.000 |
| cluster3230 | -0.35 | 1.000 |
| cluster2895 | -0.39 | 1.000 |
| cluster3603 | -0.38 | 1.000 |
| cluster3951 | 0.81 | 1.000 |
| cluster3887 | 0.82 | 1.000 |
| cluster3902 | 0.93 | 1.000 |
| cluster3690 | -0.33 | 1.000 |
| cluster3463 | -0.38 | 1.000 |
| cluster3952 | 0.88 | 1.000 |
| cluster3800 | 1.24 | 1.000 |

| | | |
|---|---|---|
| cluster4099 | 0.90 | 1.000 |
| cluster3813 | 0.79 | 1.000 |
| cluster3401 | -0.36 | 1.000 |
| cluster3279 | 0.76 | 1.000 |
| cluster3446 | -0.37 | 1.000 |
| cluster3403 | -0.38 | 1.000 |
| cluster3907 | -0.37 | 1.000 |
| cluster4024 | 0.81 | 1.000 |
| cluster3369 | -0.37 | 1.000 |
| cluster3219 | 0.69 | 1.000 |
| cluster3524 | -0.33 | 1.000 |
| cluster3606 | 0.81 | 1.000 |
| cluster4124 | -0.45 | 1.000 |
| cluster3993 | 0.80 | 1.000 |
| cluster2966 | -0.37 | 1.000 |
| cluster3261 | -0.36 | 1.000 |
| cluster3863 | 0.68 | 1.000 |
| cluster3869 | 0.91 | 1.000 |
| cluster3862 | -0.36 | 1.000 |
| cluster3278 | -0.31 | 1.000 |
| cluster4043 | -0.42 | 1.000 |
| cluster4128 | 1.03 | 1.000 |
| cluster3366 | 0.83 | 1.000 |
| cluster3657 | 0.84 | 1.000 |
| cluster4080 | 1.25 | 1.000 |
| cluster3724 | 0.83 | 1.000 |
| cluster3416 | 0.87 | 1.000 |
| cluster3484 | -0.38 | 1.000 |
| cluster3644 | 0.67 | 1.000 |
| cluster3970 | 0.91 | 1.000 |
| cluster3134 | -0.30 | 1.000 |
| cluster3638 | 0.89 | 1.000 |
| cluster3674 | 1.09 | 1.000 |
| cluster3148 | -0.30 | 1.000 |
| cluster3097 | -0.31 | 1.000 |
| cluster3120 | -0.32 | 1.000 |
| cluster3589 | -0.34 | 1.000 |
| cluster3081 | 0.66 | 1.000 |
| cluster3339 | -0.30 | 1.000 |
| cluster3204 | -0.31 | 1.000 |
| cluster3937 | 0.68 | 1.000 |
| cluster4126 | 0.94 | 1.000 |
| cluster3549 | -0.30 | 1.000 |
| cluster3654 | 0.73 | 1.000 |

| | | |
|---|---|---|
| cluster3996 | -0.33 | 1.000 |
| cluster3921 | -0.34 | 1.000 |
| cluster3056 | -0.29 | 1.000 |
| cluster3528 | -0.30 | 1.000 |
| cluster3107 | 0.56 | 1.000 |
| cluster3130 | -0.33 | 1.000 |
| cluster3174 | -0.33 | 1.000 |
| cluster4005 | -0.33 | 1.000 |
| cluster3498 | 0.79 | 1.000 |
| cluster3590 | -0.32 | 1.000 |
| cluster3013 | -0.29 | 1.000 |
| cluster3473 | 0.67 | 1.000 |
| cluster3091 | -0.29 | 1.000 |
| cluster3161 | -0.33 | 1.000 |
| cluster3011 | 0.64 | 1.000 |
| cluster3042 | 0.77 | 1.000 |
| cluster3169 | -0.32 | 1.000 |
| cluster3551 | -0.30 | 1.000 |
| cluster3653 | 0.61 | 1.000 |
| cluster3136 | 0.71 | 1.000 |
| cluster3009 | 0.67 | 1.000 |
| cluster3057 | -0.30 | 1.000 |
| cluster3594 | 0.80 | 1.000 |
| cluster2991 | -0.30 | 1.000 |
| cluster3882 | 0.84 | 1.000 |
| cluster3764 | 0.82 | 1.000 |
| cluster3643 | -0.29 | 1.000 |
| cluster3963 | 0.80 | 1.000 |
| cluster2958 | -0.27 | 1.000 |
| cluster3743 | 0.72 | 1.000 |
| cluster3641 | -0.27 | 1.000 |
| cluster3781 | 0.60 | 1.000 |
| cluster3363 | 0.75 | 1.000 |
| cluster3337 | 0.61 | 1.000 |
| cluster3383 | -0.30 | 1.000 |
| cluster4073 | 0.93 | 1.000 |
| cluster2920 | -0.28 | 1.000 |
| cluster3960 | -0.29 | 1.000 |
| cluster3271 | 0.49 | 1.000 |
| cluster4004 | -0.29 | 1.000 |
| cluster4017 | 0.65 | 1.000 |
| cluster3977 | -0.29 | 1.000 |
| cluster3001 | -0.24 | 1.000 |
| cluster3974 | 0.90 | 1.000 |

| | | |
|---|---|---|
| cluster2880 | -0.29 | 1.000 |
| cluster3237 | 0.91 | 1.000 |
| cluster3671 | 0.80 | 1.000 |
| cluster3131 | 0.84 | 1.000 |
| cluster3856 | 0.80 | 1.000 |
| cluster2922 | -0.28 | 1.000 |
| cluster3582 | -0.25 | 1.000 |
| cluster3684 | -0.28 | 1.000 |
| cluster3352 | -0.27 | 1.000 |
| cluster3896 | -0.27 | 1.000 |
| cluster4008 | 0.67 | 1.000 |
| cluster3225 | -0.27 | 1.000 |
| cluster3601 | -0.27 | 1.000 |
| cluster3519 | -0.28 | 1.000 |
| cluster3792 | -0.26 | 1.000 |
| cluster3210 | -0.26 | 1.000 |
| cluster3105 | -0.27 | 1.000 |
| cluster3051 | -0.26 | 1.000 |
| cluster3650 | 0.69 | 1.000 |
| cluster3292 | -0.27 | 1.000 |
| cluster3979 | 0.52 | 1.000 |
| cluster3298 | -0.24 | 1.000 |
| cluster4109 | 0.75 | 1.000 |
| cluster3815 | 0.75 | 1.000 |
| cluster3517 | 0.49 | 1.000 |
| cluster3992 | 0.56 | 1.000 |
| cluster3285 | -0.24 | 1.000 |
| cluster3206 | -0.26 | 1.000 |
| cluster3868 | -0.25 | 1.000 |
| cluster3213 | 0.55 | 1.000 |
| cluster3661 | 0.49 | 1.000 |
| cluster3836 | 0.71 | 1.000 |
| cluster2944 | -0.24 | 1.000 |
| cluster4106 | 0.86 | 1.000 |
| cluster3141 | -0.22 | 1.000 |
| cluster3466 | -0.24 | 1.000 |
| cluster3086 | -0.23 | 1.000 |
| cluster3079 | 0.49 | 1.000 |
| cluster3441 | -0.22 | 1.000 |
| cluster2886 | 0.50 | 1.000 |
| cluster3647 | 0.66 | 1.000 |
| cluster4122 | 0.91 | 1.000 |
| cluster4000 | 0.54 | 1.000 |
| cluster3876 | -0.22 | 1.000 |

| | | |
|---|---|---|
| cluster3247 | -0.22 | 1.000 |
| cluster3102 | -0.22 | 1.000 |
| cluster2955 | -0.21 | 1.000 |
| cluster4046 | 0.93 | 1.000 |
| cluster3411 | 0.67 | 1.000 |
| cluster3469 | -0.22 | 1.000 |
| cluster3083 | 0.47 | 1.000 |
| cluster3706 | -0.22 | 1.000 |
| cluster3817 | 0.63 | 1.000 |
| cluster3375 | 0.56 | 1.000 |
| cluster2932 | 0.50 | 1.000 |
| cluster3000 | -0.22 | 1.000 |
| cluster3344 | -0.22 | 1.000 |
| cluster3530 | -0.22 | 1.000 |
| cluster3281 | -0.22 | 1.000 |
| cluster2981 | 0.68 | 1.000 |
| cluster3691 | -0.21 | 1.000 |
| cluster3564 | 0.61 | 1.000 |
| cluster3980 | 0.43 | 1.000 |
| cluster3414 | -0.20 | 1.000 |
| cluster3664 | -0.22 | 1.000 |
| cluster3656 | 0.49 | 1.000 |
| cluster3826 | 0.61 | 1.000 |
| cluster3341 | -0.21 | 1.000 |
| cluster3145 | -0.20 | 1.000 |
| cluster3768 | -0.21 | 1.000 |
| cluster3696 | 0.55 | 1.000 |
| cluster4085 | -0.18 | 1.000 |
| cluster3184 | -0.20 | 1.000 |
| cluster3708 | 0.78 | 1.000 |
| cluster3610 | -0.21 | 1.000 |
| cluster2996 | -0.19 | 1.000 |
| cluster3354 | -0.19 | 1.000 |
| cluster3756 | -0.20 | 1.000 |
| cluster4114 | -0.18 | 1.000 |
| cluster3450 | 0.85 | 1.000 |
| cluster3959 | -0.20 | 1.000 |
| cluster2888 | 0.75 | 1.000 |
| cluster3570 | 0.67 | 1.000 |
| cluster2909 | -0.18 | 1.000 |
| cluster3490 | 0.62 | 1.000 |
| cluster3592 | 0.61 | 1.000 |
| cluster3945 | 0.66 | 1.000 |
| cluster3535 | -0.20 | 1.000 |

| | | |
|---|---|---|
| cluster3596 | -0.20 | 1.000 |
| cluster4028 | -0.19 | 1.000 |
| cluster3655 | -0.19 | 1.000 |
| cluster3771 | -0.19 | 1.000 |
| cluster3623 | 0.65 | 1.000 |
| cluster2984 | 0.51 | 1.000 |
| cluster3894 | 0.63 | 1.000 |
| cluster3202 | -0.18 | 1.000 |
| cluster3094 | 0.64 | 1.000 |
| cluster3395 | 0.46 | 1.000 |
| cluster3916 | 0.58 | 1.000 |
| cluster3825 | -0.18 | 1.000 |
| cluster3311 | 0.46 | 1.000 |
| cluster3017 | 0.61 | 1.000 |
| cluster2887 | 0.62 | 1.000 |
| cluster2907 | -0.16 | 1.000 |
| cluster2988 | 0.51 | 1.000 |
| cluster3857 | -0.17 | 1.000 |
| cluster3628 | 0.60 | 1.000 |
| cluster3973 | -0.14 | 1.000 |
| cluster2900 | -0.16 | 1.000 |
| cluster3668 | 0.57 | 1.000 |
| cluster3745 | -0.14 | 1.000 |
| cluster3014 | -0.15 | 1.000 |
| cluster3119 | 0.44 | 1.000 |
| cluster3761 | -0.14 | 1.000 |
| cluster3442 | -0.14 | 1.000 |
| cluster4059 | 0.67 | 1.000 |
| cluster3485 | 0.76 | 1.000 |
| cluster3294 | 0.45 | 1.000 |
| cluster2875 | -0.15 | 1.000 |
| cluster3304 | -0.15 | 1.000 |
| cluster3705 | 0.50 | 1.000 |
| cluster3096 | 0.44 | 1.000 |
| cluster3816 | -0.14 | 1.000 |
| cluster3023 | 0.43 | 1.000 |
| cluster3982 | 0.53 | 1.000 |
| cluster3722 | -0.14 | 1.000 |
| cluster3574 | 0.68 | 1.000 |
| cluster3127 | 0.43 | 1.000 |
| cluster3560 | -0.13 | 1.000 |
| cluster3841 | -0.13 | 1.000 |
| cluster3345 | 0.53 | 1.000 |
| cluster3074 | -0.12 | 1.000 |

| | | |
|---|---|---|
| cluster3049 | -0.13 | 1.000 |
| cluster3635 | -0.13 | 1.000 |
| cluster3617 | 0.73 | 1.000 |
| cluster3591 | 0.43 | 1.000 |
| cluster3958 | 0.68 | 1.000 |
| cluster3308 | -0.13 | 1.000 |
| cluster3098 | 0.53 | 1.000 |
| cluster3218 | 0.58 | 1.000 |
| cluster4105 | 0.60 | 1.000 |
| cluster2889 | -0.11 | 1.000 |
| cluster3382 | 0.42 | 1.000 |
| cluster3110 | -0.12 | 1.000 |
| cluster3514 | -0.12 | 1.000 |
| cluster3636 | -0.12 | 1.000 |
| cluster2869 | -0.12 | 1.000 |
| cluster3275 | 0.57 | 1.000 |
| cluster3966 | 0.51 | 1.000 |
| cluster3875 | 0.46 | 1.000 |
| cluster3798 | -0.11 | 1.000 |
| cluster3988 | -0.11 | 1.000 |
| cluster3893 | 0.36 | 1.000 |
| cluster4037 | 0.52 | 1.000 |
| cluster3672 | 0.55 | 1.000 |
| cluster2986 | -0.08 | 1.000 |
| cluster3059 | 0.57 | 1.000 |
| cluster2936 | 0.54 | 1.000 |
| cluster4034 | 0.12 | 1.000 |
| cluster3430 | -0.07 | 1.000 |
| cluster3376 | 0.33 | 1.000 |
| cluster3180 | -0.08 | 1.000 |
| cluster3725 | 0.49 | 1.000 |
| cluster3228 | -0.08 | 1.000 |
| cluster2940 | -0.07 | 1.000 |
| cluster3632 | 0.35 | 1.000 |
| cluster3093 | -0.08 | 1.000 |
| cluster3125 | 0.39 | 1.000 |
| cluster4056 | -0.06 | 1.000 |
| cluster3994 | 0.61 | 1.000 |
| cluster3832 | -0.07 | 1.000 |
| cluster2939 | -0.09 | 1.000 |
| cluster3373 | -0.08 | 1.000 |
| cluster3939 | 0.46 | 1.000 |
| cluster3837 | -0.07 | 1.000 |
| cluster2884 | -0.08 | 1.000 |

| | | |
|---|---|---|
| cluster2960 | 0.42 | 1.000 |
| cluster3895 | -0.07 | 1.000 |
| cluster3289 | -0.08 | 1.000 |
| cluster2896 | -0.06 | 1.000 |
| cluster3351 | 0.39 | 1.000 |
| cluster3766 | 0.63 | 1.000 |
| cluster3569 | 0.51 | 1.000 |
| cluster3426 | -0.08 | 1.000 |
| cluster3320 | -0.06 | 1.000 |
| cluster3967 | -0.08 | 1.000 |
| cluster3665 | 0.49 | 1.000 |
| cluster3066 | -0.08 | 1.000 |
| cluster3801 | 0.46 | 1.000 |
| cluster3954 | 0.51 | 1.000 |
| cluster3491 | -0.06 | 1.000 |
| cluster3626 | -0.08 | 1.000 |
| cluster3140 | -0.08 | 1.000 |
| cluster2948 | -0.06 | 1.000 |
| cluster3700 | -0.06 | 1.000 |
| cluster3508 | 0.50 | 1.000 |
| cluster3667 | 0.36 | 1.000 |
| cluster3997 | 0.46 | 1.000 |
| cluster3923 | -0.07 | 1.000 |
| cluster3734 | 0.49 | 1.000 |
| cluster3149 | 0.30 | 1.000 |
| cluster3116 | -0.06 | 1.000 |
| cluster2993 | 0.37 | 1.000 |
| cluster3267 | -0.07 | 1.000 |
| cluster3694 | 0.41 | 1.000 |
| cluster3621 | 0.51 | 1.000 |
| cluster3850 | 0.63 | 1.000 |
| cluster3434 | -0.06 | 1.000 |
| cluster4065 | -0.04 | 1.000 |
| cluster3742 | -0.04 | 1.000 |
| cluster3586 | -0.06 | 1.000 |
| cluster2891 | -0.02 | 1.000 |
| cluster3043 | -0.05 | 1.000 |
| cluster3599 | 0.50 | 1.000 |
| cluster3388 | 0.46 | 1.000 |
| cluster3033 | 0.34 | 1.000 |
| cluster3338 | 0.45 | 1.000 |
| cluster3445 | -0.06 | 1.000 |
| cluster3253 | -0.03 | 1.000 |
| cluster2873 | 0.32 | 1.000 |

| | | |
|---|---|---|
| cluster3588 | -0.03 | 1.000 |
| cluster3803 | -0.01 | 1.000 |
| cluster2938 | 0.28 | 1.000 |
| cluster3226 | 0.46 | 1.000 |
| cluster4063 | 0.53 | 1.000 |
| cluster3322 | -0.03 | 1.000 |
| cluster3183 | -0.04 | 1.000 |
| cluster3999 | 0.41 | 1.000 |
| cluster3829 | -0.03 | 1.000 |
| cluster4097 | 0.17 | 1.000 |
| cluster3317 | -0.03 | 1.000 |
| cluster3763 | -0.03 | 1.000 |
| cluster3867 | 0.33 | 1.000 |
| cluster3334 | -0.05 | 1.000 |
| cluster2914 | 0.43 | 1.000 |
| cluster3948 | 0.46 | 1.000 |
| cluster3835 | -0.03 | 1.000 |
| cluster4082 | 0.47 | 1.000 |
| cluster3034 | 0.39 | 1.000 |
| cluster2911 | -0.04 | 1.000 |
| cluster3048 | 0.43 | 1.000 |
| cluster3786 | 0.00 | 1.000 |
| cluster3753 | 0.15 | 1.000 |
| cluster3751 | -0.03 | 1.000 |
| cluster3720 | 0.40 | 1.000 |
| cluster3925 | 0.99 | 1.000 |
| cluster3340 | -0.04 | 1.000 |
| cluster2972 | -0.02 | 1.000 |
| cluster2915 | 0.39 | 1.000 |
| cluster3143 | -0.04 | 1.000 |
| cluster3254 | 0.54 | 1.000 |
| cluster4015 | 0.58 | 1.000 |
| cluster2879 | -0.03 | 1.000 |
| cluster3531 | 0.46 | 1.000 |
| cluster2963 | -0.02 | 1.000 |
| cluster3265 | 0.46 | 1.000 |
| cluster4089 | 0.63 | 1.000 |
| cluster3673 | 0.44 | 1.000 |
| cluster3642 | 0.44 | 1.000 |
| cluster3873 | 0.26 | 1.000 |
| cluster4064 | 0.42 | 1.000 |
| cluster2969 | 0.42 | 1.000 |
| cluster3712 | 0.33 | 1.000 |
| cluster3268 | 0.42 | 1.000 |

| | | |
|---|---|---|
| cluster4072 | 0.01 | 1.000 |
| cluster3579 | 0.29 | 1.000 |
| cluster3220 | 0.00 | 1.000 |
| cluster3969 | -0.03 | 1.000 |
| cluster3527 | -0.01 | 1.000 |
| cluster3888 | 0.39 | 1.000 |
| cluster3799 | -0.02 | 1.000 |
| cluster3162 | 0.41 | 1.000 |
| cluster3697 | 0.01 | 1.000 |
| cluster3537 | 0.41 | 1.000 |
| cluster3532 | 0.34 | 1.000 |
| cluster3453 | 0.00 | 1.000 |
| cluster3919 | 0.21 | 1.000 |
| cluster3412 | 0.41 | 1.000 |
| cluster3372 | 0.29 | 1.000 |
| cluster4011 | 0.22 | 1.000 |
| cluster2916 | -0.01 | 1.000 |
| cluster3637 | 0.25 | 1.000 |
| cluster3214 | 0.29 | 1.000 |
| cluster3203 | 0.39 | 1.000 |
| cluster3462 | 0.34 | 1.000 |
| cluster3092 | 0.00 | 1.000 |
| cluster3806 | 0.39 | 1.000 |
| cluster3584 | 0.40 | 1.000 |
| cluster3389 | 0.42 | 1.000 |
| cluster3552 | -0.01 | 1.000 |
| cluster2917 | 0.04 | 1.000 |
| cluster3006 | 0.43 | 1.000 |
| cluster3396 | -0.01 | 1.000 |
| cluster2971 | -0.02 | 1.000 |
| cluster3844 | 0.55 | 1.000 |
| cluster3209 | 0.04 | 1.000 |
| cluster3861 | 0.37 | 1.000 |
| cluster3920 | 0.45 | 1.000 |
| cluster3733 | 0.29 | 1.000 |
| cluster3407 | 0.00 | 1.000 |
| cluster3633 | 0.32 | 1.000 |
| cluster3692 | 0.36 | 1.000 |
| cluster3503 | 0.37 | 1.000 |
| cluster3370 | 0.35 | 1.000 |
| cluster3234 | 0.29 | 1.000 |
| cluster3384 | 0.02 | 1.000 |
| cluster3165 | 0.42 | 1.000 |
| cluster2927 | 0.37 | 1.000 |

| | | |
|---|---|---|
| cluster3581 | 0.06 | 1.000 |
| cluster4096 | 0.37 | 1.000 |
| cluster3595 | 0.27 | 1.000 |
| cluster3486 | 0.28 | 1.000 |
| cluster2857 | 0.02 | 1.000 |
| cluster3523 | 0.35 | 1.000 |
| cluster4093 | 0.43 | 1.000 |
| cluster3076 | 0.37 | 1.000 |
| cluster3429 | 0.26 | 1.000 |
| cluster3138 | 0.27 | 1.000 |
| cluster3630 | 0.00 | 1.000 |
| cluster3168 | 0.25 | 1.000 |
| cluster2913 | 0.26 | 1.000 |
| cluster2985 | 0.26 | 1.000 |
| cluster3270 | 0.40 | 1.000 |
| cluster3472 | 0.24 | 1.000 |
| cluster2999 | 0.38 | 1.000 |
| cluster3711 | 0.03 | 1.000 |
| cluster2976 | 0.06 | 1.000 |
| cluster3730 | 0.24 | 1.000 |
| cluster3418 | 0.33 | 1.000 |
| cluster2883 | 0.03 | 1.000 |
| cluster3811 | 0.20 | 1.000 |
| cluster2899 | 0.28 | 1.000 |
| cluster3695 | 0.28 | 1.000 |
| cluster2898 | 0.25 | 1.000 |
| cluster3082 | 0.12 | 1.000 |
| cluster3347 | 0.21 | 1.000 |
| cluster3381 | 0.26 | 1.000 |
| cluster3788 | 0.25 | 1.000 |
| cluster2870 | 0.05 | 1.000 |
| cluster2929 | 0.08 | 1.000 |
| cluster3510 | 0.39 | 1.000 |
| cluster3452 | 0.33 | 1.000 |
| cluster3539 | 0.37 | 1.000 |
| cluster2862 | 0.04 | 1.000 |
| cluster2881 | 0.06 | 1.000 |
| cluster3172 | 0.06 | 1.000 |
| cluster3122 | 0.23 | 1.000 |
| cluster2864 | 0.04 | 1.000 |
| cluster3739 | 0.05 | 1.000 |
| cluster4007 | 0.25 | 1.000 |
| cluster3152 | 0.29 | 1.000 |
| cluster3699 | 0.29 | 1.000 |

| | | |
|---|---|---|
| cluster3332 | 0.31 | 1.000 |
| cluster4047 | 0.39 | 1.000 |
| cluster4019 | 0.19 | 1.000 |
| cluster4115 | 0.35 | 1.000 |
| cluster2854 | 0.30 | 1.000 |
| cluster3018 | 0.29 | 1.000 |
| cluster3021 | 0.18 | 1.000 |
| cluster2978 | 0.08 | 1.000 |
| cluster3928 | 0.29 | 1.000 |
| cluster3918 | 0.29 | 1.000 |
| cluster3767 | 0.29 | 1.000 |
| cluster3914 | 0.41 | 1.000 |
| cluster3986 | 0.25 | 1.000 |
| cluster3464 | 0.26 | 1.000 |
| cluster3669 | 0.31 | 1.000 |
| cluster2979 | 0.12 | 1.000 |
| cluster3809 | 0.27 | 1.000 |
| cluster3470 | 0.19 | 1.000 |
| cluster2865 | 0.17 | 1.000 |
| cluster4110 | 0.26 | 1.000 |
| cluster2930 | 0.22 | 1.000 |
| cluster3879 | 0.25 | 1.000 |
| cluster2903 | 0.11 | 1.000 |
| cluster3424 | 0.37 | 1.000 |
| cluster3465 | 0.18 | 1.000 |
| cluster3027 | 0.17 | 1.000 |
| cluster4025 | 0.18 | 1.000 |
| cluster3608 | 0.19 | 1.000 |
| cluster3721 | 0.22 | 1.000 |
| cluster3025 | 0.07 | 1.000 |
| cluster3459 | 0.27 | 1.000 |
| cluster3325 | 0.36 | 1.000 |
| cluster3039 | 0.16 | 1.000 |
| cluster3312 | 0.30 | 1.000 |
| cluster2970 | 0.11 | 1.000 |
| cluster3499 | 0.24 | 1.000 |
| cluster3190 | 0.29 | 1.000 |
| cluster3327 | 0.18 | 1.000 |
| cluster3398 | 0.22 | 1.000 |
| cluster3361 | 0.18 | 1.000 |
| cluster3058 | 0.08 | 1.000 |
| cluster3820 | 0.24 | 1.000 |
| cluster3740 | 0.24 | 1.000 |
| cluster3126 | 0.16 | 1.000 |

| | | |
|---|---|---|
| cluster3845 | 0.37 | 1.000 |
| cluster4036 | 0.21 | 1.000 |
| cluster3147 | 0.17 | 1.000 |
| cluster3577 | 0.10 | 1.000 |
| cluster3714 | 0.19 | 1.000 |
| cluster2998 | 0.19 | 1.000 |
| cluster3358 | 0.16 | 1.000 |
| cluster3507 | 0.27 | 1.000 |
| cluster3132 | 0.14 | 1.000 |
| cluster2945 | 0.24 | 1.000 |
| cluster3796 | 0.25 | 1.000 |
| cluster3007 | 0.22 | 1.000 |
| cluster3188 | 0.18 | 1.000 |
| cluster3264 | 0.13 | 1.000 |
| cluster3680 | 0.22 | 1.000 |
| cluster3495 | 0.20 | 1.000 |
| cluster3900 | 0.29 | 1.000 |
| cluster3432 | 0.17 | 1.000 |
| cluster3931 | 0.24 | 1.000 |
| cluster4070 | 0.37 | 1.000 |
| cluster3242 | 0.14 | 1.000 |
| cluster3961 | 0.22 | 1.000 |
| cluster3087 | 0.17 | 1.000 |
| cluster2975 | 0.13 | 1.000 |
| cluster3750 | 0.29 | 1.000 |
| cluster3437 | 0.15 | 1.000 |
| cluster3544 | 0.28 | 1.000 |
| cluster2980 | 0.26 | 1.000 |
| cluster3423 | 0.22 | 1.000 |
| cluster2853 | 0.17 | 1.000 |
| cluster4041 | 0.28 | 1.000 |
| cluster3625 | 0.20 | 1.000 |
| cluster3910 | 0.23 | 1.000 |
| cluster3757 | 0.24 | 1.000 |
| cluster3371 | 0.23 | 1.000 |
| cluster3288 | 0.22 | 1.000 |
| cluster3421 | 0.22 | 1.000 |
| cluster3489 | 0.21 | 1.000 |
| cluster3870 | 0.20 | 1.000 |
| cluster3917 | 0.20 | 1.000 |
| cluster3795 | 0.20 | 1.000 |
| cluster3436 | 0.19 | 1.000 |
| cluster3828 | 0.18 | 1.000 |
| cluster3518 | 0.18 | 1.000 |

| | | |
|---|---|---|
| cluster3898 | 0.17 | 1.000 |
| cluster3542 | 0.17 | 1.000 |
| cluster3683 | 0.17 | 1.000 |
| cluster3223 | 0.17 | 1.000 |
| cluster3557 | 0.16 | 1.000 |
| cluster3662 | 0.15 | 1.000 |
| cluster3438 | 0.14 | 1.000 |
| cluster3679 | 0.14 | 1.000 |
| cluster3676 | 0.13 | 1.000 |
| cluster3318 | 0.13 | 1.000 |
| cluster3181 | 0.13 | 1.000 |
| cluster2947 | 0.13 | 1.000 |
| cluster4020 | 0.12 | 1.000 |
| cluster3622 | 0.12 | 1.000 |
| cluster1127 | 0.11 | 1.000 |
| cluster3435 | 0.11 | 1.000 |
| cluster3526 | 0.11 | 1.000 |
| cluster3062 | 0.11 | 1.000 |
| cluster3776 | 0.11 | 1.000 |
| cluster3335 | 0.10 | 1.000 |
| cluster3391 | 0.09 | 1.000 |
| cluster3248 | 0.08 | 1.000 |
| cluster2859 | 0.08 | 1.000 |
| cluster3947 | 0.08 | 1.000 |
| cluster3548 | 0.08 | 1.000 |
| cluster3687 | 0.08 | 1.000 |
| cluster3216 | 0.08 | 1.000 |
| cluster3943 | 0.08 | 1.000 |
| cluster3846 | 0.07 | 1.000 |
| cluster3553 | 0.06 | 1.000 |
| cluster3284 | 0.05 | 1.000 |
| cluster3232 | 0.05 | 1.000 |
| cluster3154 | 0.05 | 1.000 |
| cluster3602 | 0.05 | 1.000 |
| cluster3580 | 0.05 | 1.000 |
| cluster3847 | 0.04 | 1.000 |
| cluster3565 | 0.04 | 1.000 |
| cluster4021 | 0.04 | 1.000 |
| cluster3627 | 0.04 | 1.000 |
| cluster3330 | 0.03 | 1.000 |
| cluster3872 | 0.03 | 1.000 |
| cluster3615 | 0.02 | 1.000 |
| cluster3493 | 0.01 | 1.000 |
| cluster3605 | 0.01 | 1.000 |

| | | |
|---|---|---|
| cluster3624 | 0.00 | 1.000 |

**Supplementary Table 7.** DESeq2 results with eXpress counts - Oestridae vs Calliphoridae

| ID | log2FoldChange | Adjusted p value |
|---|---|---|
| cluster3178 | 2.92 | 0.403 |
| cluster3233 | 3.63 | 0.403 |
| cluster3755 | 3.74 | 0.403 |
| cluster3786 | 3.39 | 0.403 |
| cluster4013 | -4.54 | 0.403 |
| cluster4046 | -3.40 | 0.403 |
| cluster4101 | 4.03 | 0.403 |
| cluster3461 | -3.04 | 0.447 |
| cluster3006 | 2.85 | 0.483 |
| cluster3574 | -2.81 | 0.483 |
| cluster3763 | 2.65 | 0.483 |
| cluster3852 | 2.57 | 0.483 |
| cluster4112 | 3.23 | 0.483 |
| cluster4118 | -3.51 | 0.483 |
| cluster3137 | 2.49 | 0.534 |
| cluster3741 | 2.13 | 0.539 |
| cluster4012 | 2.25 | 0.539 |
| cluster3645 | -3.37 | 0.546 |
| cluster4087 | 3.19 | 0.576 |
| cluster3000 | -1.86 | 0.594 |
| cluster3651 | -2.71 | 0.594 |
| cluster3652 | 2.23 | 0.594 |
| cluster3802 | 2.68 | 0.594 |
| cluster3840 | 2.27 | 0.594 |
| cluster3309 | 2.11 | 0.648 |
| cluster3323 | 2.19 | 0.723 |
| cluster3666 | 2.64 | 0.723 |
| cluster3897 | 2.12 | 0.723 |
| cluster4001 | -1.99 | 0.723 |
| cluster4052 | 2.17 | 0.723 |
| cluster4060 | 2.23 | 0.723 |
| cluster4098 | 2.34 | 0.723 |
| cluster4111 | -2.75 | 0.723 |
| cluster3397 | -2.02 | 0.802 |
| cluster2861 | 1.81 | 0.911 |
| cluster2902 | 2.04 | 0.911 |
| cluster2921 | 1.63 | 0.911 |
| cluster2943 | 1.76 | 0.911 |
| cluster2947 | 1.44 | 0.911 |
| cluster2967 | 1.74 | 0.911 |
| cluster2993 | 1.74 | 0.911 |
| cluster3045 | 1.68 | 0.911 |

| | | |
|---|---|---|
| cluster3063 | 1.89 | 0.911 |
| cluster3130 | -1.46 | 0.911 |
| cluster3200 | 1.57 | 0.911 |
| cluster3317 | 1.86 | 0.911 |
| cluster3357 | 1.41 | 0.911 |
| cluster3501 | -1.92 | 0.911 |
| cluster3546 | -2.08 | 0.911 |
| cluster3561 | 1.86 | 0.911 |
| cluster3597 | 1.63 | 0.911 |
| cluster3615 | 1.86 | 0.911 |
| cluster3672 | -1.63 | 0.911 |
| cluster3691 | 1.68 | 0.911 |
| cluster3694 | -1.48 | 0.911 |
| cluster3704 | 1.77 | 0.911 |
| cluster3750 | -2.23 | 0.911 |
| cluster3772 | -1.53 | 0.911 |
| cluster3818 | 2.00 | 0.911 |
| cluster3839 | -2.04 | 0.911 |
| cluster3863 | 1.73 | 0.911 |
| cluster3876 | 1.62 | 0.911 |
| cluster3904 | -1.82 | 0.911 |
| cluster3933 | -1.56 | 0.911 |
| cluster3958 | -2.17 | 0.911 |
| cluster3981 | 1.85 | 0.911 |
| cluster4053 | -1.45 | 0.911 |
| cluster4073 | 2.45 | 0.911 |
| cluster4079 | 1.73 | 0.911 |
| cluster4090 | -2.46 | 0.911 |
| cluster4116 | -2.43 | 0.911 |
| cluster2881 | 1.53 | 0.913 |
| cluster2961 | 1.51 | 0.913 |
| cluster2962 | 1.65 | 0.913 |
| cluster2977 | 1.23 | 0.913 |
| cluster2989 | 1.54 | 0.913 |
| cluster3015 | 1.62 | 0.913 |
| cluster3100 | 1.90 | 0.913 |
| cluster3101 | 1.44 | 0.913 |
| cluster3118 | 1.35 | 0.913 |
| cluster3168 | 1.33 | 0.913 |
| cluster3177 | -1.74 | 0.913 |
| cluster3237 | 1.80 | 0.913 |
| cluster3240 | 1.51 | 0.913 |
| cluster3335 | 1.56 | 0.913 |
| cluster3432 | -1.49 | 0.913 |

| | | |
|---|---|---|
| cluster3482 | -1.49 | 0.913 |
| cluster3525 | -1.19 | 0.913 |
| cluster3534 | -1.57 | 0.913 |
| cluster3551 | 1.70 | 0.913 |
| cluster3564 | 1.92 | 0.913 |
| cluster3595 | 1.40 | 0.913 |
| cluster3656 | 1.45 | 0.913 |
| cluster3719 | 1.64 | 0.913 |
| cluster3745 | 2.05 | 0.913 |
| cluster3760 | 1.43 | 0.913 |
| cluster3865 | 1.92 | 0.913 |
| cluster3885 | -1.33 | 0.913 |
| cluster3898 | 1.74 | 0.913 |
| cluster3901 | 1.84 | 0.913 |
| cluster3915 | -2.04 | 0.913 |
| cluster4055 | -1.39 | 0.913 |
| cluster4070 | -2.29 | 0.913 |
| cluster4074 | 2.03 | 0.913 |
| cluster4095 | 1.87 | 0.913 |
| cluster4099 | 1.40 | 0.913 |
| cluster4103 | 1.81 | 0.913 |
| cluster4114 | 2.67 | 0.913 |
| cluster4119 | 2.16 | 0.913 |
| cluster3705 | 1.64 | 0.919 |
| cluster3245 | 1.62 | 0.920 |
| cluster3940 | 1.38 | 0.920 |
| cluster3987 | 1.47 | 0.920 |
| cluster2930 | 1.22 | 0.925 |
| cluster3009 | -1.41 | 0.925 |
| cluster3053 | 1.70 | 0.925 |
| cluster3176 | -1.50 | 0.925 |
| cluster3197 | 1.47 | 0.925 |
| cluster3430 | 1.89 | 0.925 |
| cluster3474 | 1.67 | 0.925 |
| cluster3514 | 1.61 | 0.925 |
| cluster3537 | 1.16 | 0.925 |
| cluster3563 | -1.38 | 0.925 |
| cluster3792 | -1.13 | 0.925 |
| cluster3830 | 1.18 | 0.925 |
| cluster3834 | -1.61 | 0.925 |
| cluster4002 | 1.56 | 0.925 |
| cluster4016 | 1.48 | 0.925 |
| cluster2854 | 1.38 | 0.925 |
| cluster3157 | 1.48 | 0.925 |

| | | |
|---|---|---|
| cluster3448 | -1.66 | 0.925 |
| cluster3462 | -1.25 | 0.925 |
| cluster3556 | 1.42 | 0.925 |
| cluster3568 | -1.69 | 0.925 |
| cluster3689 | -1.96 | 0.925 |
| cluster3853 | 1.52 | 0.925 |
| cluster4034 | -2.17 | 0.925 |
| cluster3306 | 1.72 | 0.937 |
| cluster3246 | 1.29 | 0.941 |
| cluster3332 | 1.27 | 0.941 |
| cluster3435 | -1.66 | 0.941 |
| cluster3599 | -1.55 | 0.941 |
| cluster3746 | 1.35 | 0.941 |
| cluster2912 | 1.16 | 0.945 |
| cluster3105 | -1.62 | 0.945 |
| cluster3575 | 1.49 | 0.948 |
| cluster2860 | 1.39 | 0.966 |
| cluster3097 | 1.28 | 0.966 |
| cluster3400 | 1.24 | 0.966 |
| cluster3431 | 1.20 | 0.966 |
| cluster3619 | 1.21 | 0.966 |
| cluster3735 | -1.34 | 0.966 |
| cluster3955 | -1.65 | 0.966 |
| cluster4049 | -2.24 | 0.966 |
| cluster3995 | -1.50 | 0.966 |
| cluster2899 | 1.49 | 0.969 |
| cluster2911 | -0.96 | 0.969 |
| cluster2924 | 1.20 | 0.969 |
| cluster2933 | 1.37 | 0.969 |
| cluster2946 | 1.19 | 0.969 |
| cluster2980 | -1.52 | 0.969 |
| cluster2995 | 1.17 | 0.969 |
| cluster3008 | 1.07 | 0.969 |
| cluster3019 | 0.98 | 0.969 |
| cluster3026 | -1.31 | 0.969 |
| cluster3032 | 1.21 | 0.969 |
| cluster3066 | 1.16 | 0.969 |
| cluster3085 | -1.30 | 0.969 |
| cluster3110 | 1.00 | 0.969 |
| cluster3165 | 1.39 | 0.969 |
| cluster3179 | 1.32 | 0.969 |
| cluster3212 | -1.32 | 0.969 |
| cluster3244 | 1.06 | 0.969 |
| cluster3290 | -1.42 | 0.969 |

| | | |
|---|---|---|
| cluster3304 | 1.13 | 0.969 |
| cluster3311 | 1.13 | 0.969 |
| cluster3312 | -1.66 | 0.969 |
| cluster3337 | 1.41 | 0.969 |
| cluster3341 | -1.16 | 0.969 |
| cluster3363 | 1.09 | 0.969 |
| cluster3369 | 1.03 | 0.969 |
| cluster3380 | -0.99 | 0.969 |
| cluster3398 | -0.95 | 0.969 |
| cluster3436 | 1.58 | 0.969 |
| cluster3438 | 1.27 | 0.969 |
| cluster3454 | -1.06 | 0.969 |
| cluster3478 | 1.15 | 0.969 |
| cluster3480 | 1.17 | 0.969 |
| cluster3485 | -1.61 | 0.969 |
| cluster3510 | -1.76 | 0.969 |
| cluster3513 | 1.19 | 0.969 |
| cluster3520 | 1.72 | 0.969 |
| cluster3522 | -1.35 | 0.969 |
| cluster3538 | 1.49 | 0.969 |
| cluster3629 | 1.25 | 0.969 |
| cluster3636 | -0.90 | 0.969 |
| cluster3659 | -1.32 | 0.969 |
| cluster3721 | 1.43 | 0.969 |
| cluster3730 | 1.16 | 0.969 |
| cluster3731 | 1.29 | 0.969 |
| cluster3766 | -1.18 | 0.969 |
| cluster3794 | -1.14 | 0.969 |
| cluster3799 | 1.17 | 0.969 |
| cluster3822 | 1.68 | 0.969 |
| cluster3842 | 1.39 | 0.969 |
| cluster3849 | 1.03 | 0.969 |
| cluster3850 | -1.21 | 0.969 |
| cluster3858 | 1.40 | 0.969 |
| cluster3894 | -0.99 | 0.969 |
| cluster3920 | -1.37 | 0.969 |
| cluster3931 | 1.52 | 0.969 |
| cluster3952 | 1.07 | 0.969 |
| cluster3956 | 1.09 | 0.969 |
| cluster3990 | 1.08 | 0.969 |
| cluster4000 | 1.20 | 0.969 |
| cluster4011 | -1.61 | 0.969 |
| cluster4033 | -1.72 | 0.969 |
| cluster4035 | 1.24 | 0.969 |

| | | |
|---|---|---|
| cluster4085 | -1.36 | 0.969 |
| cluster4121 | 1.37 | 0.969 |
| cluster3150 | -1.33 | 0.970 |
| cluster3161 | -1.05 | 0.970 |
| cluster3247 | 1.10 | 0.970 |
| cluster3313 | -1.07 | 0.970 |
| cluster3401 | 0.86 | 0.970 |
| cluster3571 | -1.37 | 0.970 |
| cluster3582 | 1.02 | 0.970 |
| cluster3585 | -0.94 | 0.970 |
| cluster3787 | -1.33 | 0.970 |
| cluster4066 | -1.17 | 0.970 |
| cluster4080 | -1.57 | 0.970 |
| cluster2892 | 0.93 | 0.971 |
| cluster2926 | 1.00 | 0.971 |
| cluster2942 | 0.92 | 0.971 |
| cluster2965 | 0.91 | 0.971 |
| cluster2969 | 1.15 | 0.971 |
| cluster2999 | -1.03 | 0.971 |
| cluster3013 | 0.97 | 0.971 |
| cluster3017 | 1.11 | 0.971 |
| cluster3027 | 1.05 | 0.971 |
| cluster3062 | -0.87 | 0.971 |
| cluster3075 | -0.99 | 0.971 |
| cluster3082 | -1.38 | 0.971 |
| cluster3083 | 0.84 | 0.971 |
| cluster3089 | 1.03 | 0.971 |
| cluster3095 | -1.05 | 0.971 |
| cluster3135 | -1.04 | 0.971 |
| cluster3141 | 1.03 | 0.971 |
| cluster3142 | 1.12 | 0.971 |
| cluster3147 | 0.83 | 0.971 |
| cluster3164 | -1.34 | 0.971 |
| cluster3175 | -1.27 | 0.971 |
| cluster3181 | 0.99 | 0.971 |
| cluster3182 | -1.10 | 0.971 |
| cluster3185 | -1.17 | 0.971 |
| cluster3188 | 1.22 | 0.971 |
| cluster3226 | -1.11 | 0.971 |
| cluster3258 | 0.87 | 0.971 |
| cluster3288 | 1.08 | 0.971 |
| cluster3292 | -1.18 | 0.971 |
| cluster3296 | -1.04 | 0.971 |
| cluster3329 | -1.35 | 0.971 |

| | | |
|---|---|---|
| cluster3391 | 1.03 | 0.971 |
| cluster3413 | -1.23 | 0.971 |
| cluster3437 | -1.00 | 0.971 |
| cluster3439 | -0.95 | 0.971 |
| cluster3458 | 0.83 | 0.971 |
| cluster3469 | 1.21 | 0.971 |
| cluster3473 | 1.12 | 0.971 |
| cluster3490 | -1.04 | 0.971 |
| cluster3497 | 1.00 | 0.971 |
| cluster3500 | -1.16 | 0.971 |
| cluster3508 | 1.27 | 0.971 |
| cluster3515 | 1.28 | 0.971 |
| cluster3519 | -1.48 | 0.971 |
| cluster3549 | 0.97 | 0.971 |
| cluster3573 | -1.02 | 0.971 |
| cluster3577 | 1.01 | 0.971 |
| cluster3581 | -1.11 | 0.971 |
| cluster3661 | 0.90 | 0.971 |
| cluster3679 | 0.98 | 0.971 |
| cluster3695 | -0.94 | 0.971 |
| cluster3709 | 1.03 | 0.971 |
| cluster3712 | 1.01 | 0.971 |
| cluster3716 | -1.14 | 0.971 |
| cluster3753 | -1.38 | 0.971 |
| cluster3756 | -1.02 | 0.971 |
| cluster3788 | 1.03 | 0.971 |
| cluster3827 | -1.09 | 0.971 |
| cluster3844 | -1.53 | 0.971 |
| cluster3873 | 0.91 | 0.971 |
| cluster3887 | -1.25 | 0.971 |
| cluster3899 | 0.99 | 0.971 |
| cluster3902 | 0.92 | 0.971 |
| cluster3937 | 1.16 | 0.971 |
| cluster3960 | -1.35 | 0.971 |
| cluster3973 | -0.94 | 0.971 |
| cluster3983 | 1.22 | 0.971 |
| cluster3985 | 0.98 | 0.971 |
| cluster3999 | -1.13 | 0.971 |
| cluster4019 | 1.23 | 0.971 |
| cluster4064 | 1.03 | 0.971 |
| cluster4093 | 1.48 | 0.971 |
| cluster4115 | -1.43 | 0.971 |
| cluster3869 | 1.30 | 0.972 |
| cluster3402 | 0.98 | 0.982 |

| | | |
|---|---|---|
| cluster3452 | -1.05 | 0.982 |
| cluster3453 | -1.06 | 0.982 |
| cluster3821 | -0.94 | 0.982 |
| cluster2863 | 0.74 | 0.988 |
| cluster2901 | -1.00 | 0.988 |
| cluster2998 | -0.71 | 0.988 |
| cluster3129 | -1.41 | 0.988 |
| cluster3220 | -0.97 | 0.988 |
| cluster3295 | 0.77 | 0.988 |
| cluster3320 | -0.92 | 0.988 |
| cluster3334 | 0.79 | 0.988 |
| cluster3338 | -1.01 | 0.988 |
| cluster3365 | 1.04 | 0.988 |
| cluster3375 | -0.80 | 0.988 |
| cluster3399 | 1.07 | 0.988 |
| cluster3408 | 1.05 | 0.988 |
| cluster3425 | -0.93 | 0.988 |
| cluster3459 | -1.19 | 0.988 |
| cluster3468 | 0.97 | 0.988 |
| cluster3811 | 0.91 | 0.988 |
| cluster3907 | -1.22 | 0.988 |
| cluster3953 | -0.86 | 0.988 |
| cluster3978 | -0.77 | 0.988 |
| cluster4075 | 1.39 | 0.988 |
| cluster2882 | 0.74 | 0.990 |
| cluster2916 | 1.00 | 0.990 |
| cluster2963 | 1.03 | 0.990 |
| cluster2994 | 0.77 | 0.990 |
| cluster3005 | 0.74 | 0.990 |
| cluster3131 | -0.83 | 0.990 |
| cluster3143 | 0.84 | 0.990 |
| cluster3156 | -0.88 | 0.990 |
| cluster3162 | 0.92 | 0.990 |
| cluster3180 | -0.77 | 0.990 |
| cluster3194 | -0.87 | 0.990 |
| cluster3239 | 0.80 | 0.990 |
| cluster3280 | 0.74 | 0.990 |
| cluster3300 | 0.84 | 0.990 |
| cluster3434 | -0.97 | 0.990 |
| cluster3446 | -0.80 | 0.990 |
| cluster3536 | 0.80 | 0.990 |
| cluster3609 | 1.31 | 0.990 |
| cluster3621 | 1.01 | 0.990 |
| cluster3634 | 0.96 | 0.990 |

| | | |
|---|---|---|
| cluster3653 | 0.98 | 0.990 |
| cluster3662 | -0.98 | 0.990 |
| cluster3703 | 0.78 | 0.990 |
| cluster3778 | 0.99 | 0.990 |
| cluster3810 | 0.85 | 0.990 |
| cluster3817 | 1.17 | 0.990 |
| cluster3916 | 0.77 | 0.990 |
| cluster3919 | -1.12 | 0.990 |
| cluster4127 | -1.42 | 0.990 |
| cluster3554 | -0.81 | 0.991 |
| cluster4056 | -0.97 | 0.993 |
| cluster3441 | 0.74 | 0.994 |
| cluster3487 | -0.92 | 0.994 |
| cluster3552 | 0.74 | 0.994 |
| cluster3948 | -0.82 | 0.994 |
| cluster3970 | 1.06 | 0.994 |
| cluster2865 | 0.69 | 0.995 |
| cluster2871 | -0.56 | 0.995 |
| cluster2877 | 0.59 | 0.995 |
| cluster2884 | 0.60 | 0.995 |
| cluster2898 | 0.78 | 0.995 |
| cluster2904 | 0.78 | 0.995 |
| cluster2908 | 0.86 | 0.995 |
| cluster2914 | -0.71 | 0.995 |
| cluster2915 | -0.72 | 0.995 |
| cluster2919 | 0.73 | 0.995 |
| cluster2923 | 0.56 | 0.995 |
| cluster2928 | 0.67 | 0.995 |
| cluster2940 | -0.68 | 0.995 |
| cluster2949 | 0.67 | 0.995 |
| cluster2957 | -0.86 | 0.995 |
| cluster2964 | 0.78 | 0.995 |
| cluster2972 | -0.84 | 0.995 |
| cluster2981 | 0.62 | 0.995 |
| cluster3004 | 0.69 | 0.995 |
| cluster3011 | 0.86 | 0.995 |
| cluster3016 | -0.93 | 0.995 |
| cluster3034 | -0.64 | 0.995 |
| cluster3049 | 0.75 | 0.995 |
| cluster3051 | -0.92 | 0.995 |
| cluster3064 | 0.88 | 0.995 |
| cluster3071 | 0.76 | 0.995 |
| cluster3076 | 0.74 | 0.995 |
| cluster3077 | 0.80 | 0.995 |

| | | |
|---|---|---|
| cluster3081 | -0.79 | 0.995 |
| cluster3088 | 0.84 | 0.995 |
| cluster3099 | 0.63 | 0.995 |
| cluster3113 | 0.66 | 0.995 |
| cluster3115 | 0.57 | 0.995 |
| cluster3117 | 0.86 | 0.995 |
| cluster3132 | 0.75 | 0.995 |
| cluster3149 | 0.64 | 0.995 |
| cluster3153 | 0.57 | 0.995 |
| cluster3174 | 0.81 | 0.995 |
| cluster3187 | -0.91 | 0.995 |
| cluster3191 | -0.96 | 0.995 |
| cluster3195 | -0.69 | 0.995 |
| cluster3224 | -0.58 | 0.995 |
| cluster3225 | -0.89 | 0.995 |
| cluster3229 | 0.99 | 0.995 |
| cluster3238 | 0.58 | 0.995 |
| cluster3255 | -0.84 | 0.995 |
| cluster3257 | 0.61 | 0.995 |
| cluster3260 | -0.75 | 0.995 |
| cluster3261 | -0.93 | 0.995 |
| cluster3267 | 0.61 | 0.995 |
| cluster3276 | 0.59 | 0.995 |
| cluster3287 | 0.79 | 0.995 |
| cluster3319 | -0.87 | 0.995 |
| cluster3321 | 0.63 | 0.995 |
| cluster3325 | -1.20 | 0.995 |
| cluster3327 | 0.85 | 0.995 |
| cluster3331 | -0.53 | 0.995 |
| cluster3340 | 0.77 | 0.995 |
| cluster3345 | -0.65 | 0.995 |
| cluster3348 | 0.81 | 0.995 |
| cluster3349 | 0.81 | 0.995 |
| cluster3353 | 0.56 | 0.995 |
| cluster3356 | 0.84 | 0.995 |
| cluster3358 | 0.73 | 0.995 |
| cluster3361 | -0.79 | 0.995 |
| cluster3377 | -0.81 | 0.995 |
| cluster3392 | -0.83 | 0.995 |
| cluster3395 | -0.68 | 0.995 |
| cluster3414 | 0.79 | 0.995 |
| cluster3442 | 0.86 | 0.995 |
| cluster3449 | 0.78 | 0.995 |
| cluster3481 | -1.15 | 0.995 |

| | | |
|---|---|---|
| cluster3484 | -0.99 | 0.995 |
| cluster3486 | 0.81 | 0.995 |
| cluster3493 | 0.99 | 0.995 |
| cluster3495 | 0.73 | 0.995 |
| cluster3511 | -0.76 | 0.995 |
| cluster3530 | 0.70 | 0.995 |
| cluster3531 | -0.69 | 0.995 |
| cluster3540 | -0.88 | 0.995 |
| cluster3541 | 0.76 | 0.995 |
| cluster3547 | 0.62 | 0.995 |
| cluster3553 | -0.66 | 0.995 |
| cluster3565 | -0.87 | 0.995 |
| cluster3567 | 1.15 | 0.995 |
| cluster3578 | -0.86 | 0.995 |
| cluster3580 | 1.01 | 0.995 |
| cluster3593 | -0.89 | 0.995 |
| cluster3598 | 1.13 | 0.995 |
| cluster3602 | 0.66 | 0.995 |
| cluster3603 | 0.77 | 0.995 |
| cluster3611 | 1.01 | 0.995 |
| cluster3613 | 0.79 | 0.995 |
| cluster3633 | -0.64 | 0.995 |
| cluster3644 | -0.88 | 0.995 |
| cluster3649 | -0.67 | 0.995 |
| cluster3657 | 0.64 | 0.995 |
| cluster3667 | -0.65 | 0.995 |
| cluster3723 | 0.64 | 0.995 |
| cluster3729 | -0.69 | 0.995 |
| cluster3737 | 0.59 | 0.995 |
| cluster3742 | 0.90 | 0.995 |
| cluster3744 | -0.65 | 0.995 |
| cluster3747 | 0.63 | 0.995 |
| cluster3748 | 0.74 | 0.995 |
| cluster3749 | -0.98 | 0.995 |
| cluster3758 | -0.88 | 0.995 |
| cluster3764 | -1.02 | 0.995 |
| cluster3768 | -0.98 | 0.995 |
| cluster3769 | -0.61 | 0.995 |
| cluster3771 | 0.83 | 0.995 |
| cluster3773 | -0.78 | 0.995 |
| cluster3779 | 0.87 | 0.995 |
| cluster3781 | 0.75 | 0.995 |
| cluster3795 | -0.71 | 0.995 |
| cluster3800 | -0.96 | 0.995 |

| | | |
|---|---|---|
| cluster3805 | 0.75 | 0.995 |
| cluster3819 | -0.92 | 0.995 |
| cluster3826 | -0.81 | 0.995 |
| cluster3854 | -0.61 | 0.995 |
| cluster3857 | -0.70 | 0.995 |
| cluster3867 | -0.61 | 0.995 |
| cluster3892 | -0.92 | 0.995 |
| cluster3918 | 0.92 | 0.995 |
| cluster3922 | -0.99 | 0.995 |
| cluster3928 | -0.66 | 0.995 |
| cluster3929 | -0.81 | 0.995 |
| cluster3930 | 0.76 | 0.995 |
| cluster3934 | 0.59 | 0.995 |
| cluster3942 | -0.65 | 0.995 |
| cluster3954 | -1.04 | 0.995 |
| cluster3959 | -1.00 | 0.995 |
| cluster3964 | -0.90 | 0.995 |
| cluster3966 | 0.53 | 0.995 |
| cluster3979 | 0.75 | 0.995 |
| cluster3989 | 0.88 | 0.995 |
| cluster3992 | 0.79 | 0.995 |
| cluster4005 | -0.88 | 0.995 |
| cluster4015 | -0.70 | 0.995 |
| cluster4022 | -0.65 | 0.995 |
| cluster4023 | -0.93 | 0.995 |
| cluster4026 | -0.80 | 0.995 |
| cluster4042 | -0.89 | 0.995 |
| cluster4047 | 1.26 | 0.995 |
| cluster4051 | 0.91 | 0.995 |
| cluster4054 | 0.89 | 0.995 |
| cluster4059 | -1.03 | 0.995 |
| cluster4072 | 0.74 | 0.995 |
| cluster4086 | 0.85 | 0.995 |
| cluster4094 | 0.99 | 0.995 |
| cluster4097 | -1.36 | 0.995 |
| cluster4106 | -0.96 | 0.995 |
| cluster4107 | 1.15 | 0.995 |
| cluster4113 | -0.72 | 0.995 |
| cluster4124 | -1.11 | 0.995 |
| cluster4128 | -1.07 | 0.995 |
| cluster1127 | 0.27 | 0.997 |
| cluster2855 | 0.31 | 0.997 |
| cluster2857 | -0.47 | 0.997 |
| cluster2858 | 0.36 | 0.997 |

| | | |
|---|---|---|
| cluster2859 | -0.20 | 0.997 |
| cluster2864 | 0.54 | 0.997 |
| cluster2866 | 0.30 | 0.997 |
| cluster2867 | -0.53 | 0.997 |
| cluster2869 | 0.35 | 0.997 |
| cluster2870 | 0.43 | 0.997 |
| cluster2874 | 0.21 | 0.997 |
| cluster2875 | 0.34 | 0.997 |
| cluster2876 | 0.84 | 0.997 |
| cluster2878 | 0.66 | 0.997 |
| cluster2879 | 0.48 | 0.997 |
| cluster2886 | 0.35 | 0.997 |
| cluster2888 | -0.61 | 0.997 |
| cluster2889 | 0.23 | 0.997 |
| cluster2891 | -0.70 | 0.997 |
| cluster2894 | -0.83 | 0.997 |
| cluster2895 | -0.30 | 0.997 |
| cluster2900 | 0.64 | 0.997 |
| cluster2907 | -0.20 | 0.997 |
| cluster2909 | 0.36 | 0.997 |
| cluster2910 | -0.27 | 0.997 |
| cluster2913 | -0.24 | 0.997 |
| cluster2922 | 0.37 | 0.997 |
| cluster2925 | 0.30 | 0.997 |
| cluster2927 | 0.51 | 0.997 |
| cluster2929 | 0.22 | 0.997 |
| cluster2931 | -0.42 | 0.997 |
| cluster2932 | -0.32 | 0.997 |
| cluster2934 | -0.35 | 0.997 |
| cluster2935 | 0.64 | 0.997 |
| cluster2936 | 0.65 | 0.997 |
| cluster2939 | -0.53 | 0.997 |
| cluster2944 | -0.23 | 0.997 |
| cluster2945 | -0.47 | 0.997 |
| cluster2950 | 0.61 | 0.997 |
| cluster2952 | -0.22 | 0.997 |
| cluster2954 | 0.24 | 0.997 |
| cluster2956 | -0.55 | 0.997 |
| cluster2966 | 0.35 | 0.997 |
| cluster2970 | 0.21 | 0.997 |
| cluster2973 | 0.23 | 0.997 |
| cluster2975 | 0.47 | 0.997 |
| cluster2976 | -0.57 | 0.997 |
| cluster2979 | 0.34 | 0.997 |

| | | |
|---|---|---|
| cluster2982 | 0.51 | 0.997 |
| cluster2983 | 0.16 | 0.997 |
| cluster2985 | 0.20 | 0.997 |
| cluster2987 | 0.49 | 0.997 |
| cluster2988 | 0.22 | 0.997 |
| cluster2991 | -0.23 | 0.997 |
| cluster2992 | -0.33 | 0.997 |
| cluster2997 | -0.33 | 0.997 |
| cluster3001 | -0.18 | 0.997 |
| cluster3002 | 0.37 | 0.997 |
| cluster3007 | 0.27 | 0.997 |
| cluster3010 | -0.54 | 0.997 |
| cluster3014 | -0.49 | 0.997 |
| cluster3020 | -0.24 | 0.997 |
| cluster3021 | 0.55 | 0.997 |
| cluster3022 | -0.62 | 0.997 |
| cluster3024 | 0.63 | 0.997 |
| cluster3025 | 0.41 | 0.997 |
| cluster3031 | 0.41 | 0.997 |
| cluster3033 | -0.31 | 0.997 |
| cluster3035 | 0.33 | 0.997 |
| cluster3036 | -0.50 | 0.997 |
| cluster3037 | -0.23 | 0.997 |
| cluster3039 | 0.50 | 0.997 |
| cluster3042 | -0.37 | 0.997 |
| cluster3046 | 0.25 | 0.997 |
| cluster3047 | -0.52 | 0.997 |
| cluster3048 | -0.45 | 0.997 |
| cluster3050 | -0.18 | 0.997 |
| cluster3055 | 0.42 | 0.997 |
| cluster3057 | -0.44 | 0.997 |
| cluster3059 | -0.23 | 0.997 |
| cluster3065 | 0.48 | 0.997 |
| cluster3067 | 0.45 | 0.997 |
| cluster3068 | -0.39 | 0.997 |
| cluster3069 | 0.20 | 0.997 |
| cluster3070 | 0.56 | 0.997 |
| cluster3073 | -0.34 | 0.997 |
| cluster3074 | -0.24 | 0.997 |
| cluster3078 | 0.20 | 0.997 |
| cluster3080 | -0.32 | 0.997 |
| cluster3087 | 0.22 | 0.997 |
| cluster3091 | -0.18 | 0.997 |
| cluster3093 | -0.57 | 0.997 |

| | | |
|---|---|---|
| cluster3094 | 0.63 | 0.997 |
| cluster3098 | -0.21 | 0.997 |
| cluster3103 | -0.64 | 0.997 |
| cluster3104 | -0.16 | 0.997 |
| cluster3106 | 0.49 | 0.997 |
| cluster3107 | 0.48 | 0.997 |
| cluster3109 | 0.29 | 0.997 |
| cluster3111 | -0.44 | 0.997 |
| cluster3112 | -0.40 | 0.997 |
| cluster3114 | -0.27 | 0.997 |
| cluster3120 | 0.32 | 0.997 |
| cluster3121 | 0.58 | 0.997 |
| cluster3122 | -0.34 | 0.997 |
| cluster3124 | -0.40 | 0.997 |
| cluster3125 | -0.31 | 0.997 |
| cluster3127 | -0.45 | 0.997 |
| cluster3128 | 0.42 | 0.997 |
| cluster3133 | -0.48 | 0.997 |
| cluster3134 | -0.45 | 0.997 |
| cluster3140 | 0.32 | 0.997 |
| cluster3144 | -0.78 | 0.997 |
| cluster3145 | -0.31 | 0.997 |
| cluster3148 | 0.25 | 0.997 |
| cluster3151 | -0.17 | 0.997 |
| cluster3155 | 0.26 | 0.997 |
| cluster3159 | -0.18 | 0.997 |
| cluster3160 | -0.45 | 0.997 |
| cluster3163 | -0.40 | 0.997 |
| cluster3166 | 0.61 | 0.997 |
| cluster3167 | -0.33 | 0.997 |
| cluster3169 | 0.26 | 0.997 |
| cluster3170 | 0.52 | 0.997 |
| cluster3183 | -0.35 | 0.997 |
| cluster3186 | -0.38 | 0.997 |
| cluster3190 | 0.54 | 0.997 |
| cluster3192 | 0.19 | 0.997 |
| cluster3198 | -0.22 | 0.997 |
| cluster3199 | 0.34 | 0.997 |
| cluster3202 | 0.35 | 0.997 |
| cluster3203 | -0.57 | 0.997 |
| cluster3204 | 0.63 | 0.997 |
| cluster3205 | -0.39 | 0.997 |
| cluster3206 | -0.43 | 0.997 |
| cluster3207 | 0.65 | 0.997 |

| | | |
|---|---|---|
| cluster3208 | 0.29 | 0.997 |
| cluster3209 | -0.50 | 0.997 |
| cluster3210 | 0.47 | 0.997 |
| cluster3215 | 0.46 | 0.997 |
| cluster3218 | 0.76 | 0.997 |
| cluster3219 | -0.17 | 0.997 |
| cluster3221 | 0.21 | 0.997 |
| cluster3227 | 0.34 | 0.997 |
| cluster3228 | 0.35 | 0.997 |
| cluster3230 | 0.37 | 0.997 |
| cluster3231 | -0.39 | 0.997 |
| cluster3232 | 0.58 | 0.997 |
| cluster3234 | -0.56 | 0.997 |
| cluster3235 | 0.64 | 0.997 |
| cluster3241 | 0.41 | 0.997 |
| cluster3242 | 0.49 | 0.997 |
| cluster3249 | -0.33 | 0.997 |
| cluster3251 | -0.24 | 0.997 |
| cluster3252 | 0.37 | 0.997 |
| cluster3253 | -0.42 | 0.997 |
| cluster3256 | 0.39 | 0.997 |
| cluster3259 | -0.30 | 0.997 |
| cluster3262 | 0.42 | 0.997 |
| cluster3263 | 0.40 | 0.997 |
| cluster3264 | -0.22 | 0.997 |
| cluster3265 | 0.33 | 0.997 |
| cluster3266 | 0.18 | 0.997 |
| cluster3268 | -0.49 | 0.997 |
| cluster3269 | -0.82 | 0.997 |
| cluster3270 | 0.44 | 0.997 |
| cluster3273 | 0.52 | 0.997 |
| cluster3275 | -0.40 | 0.997 |
| cluster3277 | -0.69 | 0.997 |
| cluster3278 | 0.18 | 0.997 |
| cluster3279 | 0.33 | 0.997 |
| cluster3282 | -0.47 | 0.997 |
| cluster3283 | 0.26 | 0.997 |
| cluster3284 | 0.21 | 0.997 |
| cluster3285 | -0.20 | 0.997 |
| cluster3286 | 0.39 | 0.997 |
| cluster3289 | 0.54 | 0.997 |
| cluster3291 | 0.55 | 0.997 |
| cluster3293 | 0.39 | 0.997 |
| cluster3298 | -0.46 | 0.997 |

| | | |
|---|---|---|
| cluster3299 | -0.63 | 0.997 |
| cluster3302 | -0.35 | 0.997 |
| cluster3305 | -0.36 | 0.997 |
| cluster3307 | -0.47 | 0.997 |
| cluster3310 | -0.26 | 0.997 |
| cluster3315 | 0.56 | 0.997 |
| cluster3316 | -0.44 | 0.997 |
| cluster3326 | 0.27 | 0.997 |
| cluster3330 | 0.31 | 0.997 |
| cluster3339 | 0.27 | 0.997 |
| cluster3342 | 0.43 | 0.997 |
| cluster3347 | 0.27 | 0.997 |
| cluster3352 | 0.30 | 0.997 |
| cluster3354 | 0.23 | 0.997 |
| cluster3355 | -0.28 | 0.997 |
| cluster3359 | -0.49 | 0.997 |
| cluster3360 | -0.53 | 0.997 |
| cluster3362 | 0.46 | 0.997 |
| cluster3364 | -0.29 | 0.997 |
| cluster3366 | 0.68 | 0.997 |
| cluster3372 | 0.40 | 0.997 |
| cluster3373 | 0.43 | 0.997 |
| cluster3374 | 0.36 | 0.997 |
| cluster3379 | -0.44 | 0.997 |
| cluster3381 | 0.44 | 0.997 |
| cluster3383 | -0.54 | 0.997 |
| cluster3384 | 0.24 | 0.997 |
| cluster3385 | -0.67 | 0.997 |
| cluster3386 | -0.58 | 0.997 |
| cluster3387 | -0.44 | 0.997 |
| cluster3403 | -0.40 | 0.997 |
| cluster3404 | 0.32 | 0.997 |
| cluster3405 | 0.43 | 0.997 |
| cluster3407 | 0.42 | 0.997 |
| cluster3411 | -0.48 | 0.997 |
| cluster3415 | -0.32 | 0.997 |
| cluster3418 | 0.20 | 0.997 |
| cluster3421 | 0.52 | 0.997 |
| cluster3422 | 0.45 | 0.997 |
| cluster3423 | 0.25 | 0.997 |
| cluster3424 | -0.35 | 0.997 |
| cluster3427 | -0.67 | 0.997 |
| cluster3429 | 0.32 | 0.997 |
| cluster3433 | 0.42 | 0.997 |

| | | |
|---|---|---|
| cluster3440 | 0.21 | 0.997 |
| cluster3443 | 0.66 | 0.997 |
| cluster3444 | 0.35 | 0.997 |
| cluster3445 | -0.50 | 0.997 |
| cluster3447 | -0.23 | 0.997 |
| cluster3450 | -0.89 | 0.997 |
| cluster3456 | 0.50 | 0.997 |
| cluster3457 | -0.66 | 0.997 |
| cluster3460 | 0.67 | 0.997 |
| cluster3463 | 0.68 | 0.997 |
| cluster3466 | -0.46 | 0.997 |
| cluster3470 | -0.24 | 0.997 |
| cluster3471 | 0.55 | 0.997 |
| cluster3475 | 0.27 | 0.997 |
| cluster3476 | 0.31 | 0.997 |
| cluster3477 | 0.23 | 0.997 |
| cluster3479 | -0.65 | 0.997 |
| cluster3488 | -0.51 | 0.997 |
| cluster3491 | -0.20 | 0.997 |
| cluster3496 | 0.60 | 0.997 |
| cluster3498 | 0.34 | 0.997 |
| cluster3499 | -0.23 | 0.997 |
| cluster3503 | 0.21 | 0.997 |
| cluster3504 | 0.29 | 0.997 |
| cluster3505 | 0.39 | 0.997 |
| cluster3506 | 0.25 | 0.997 |
| cluster3507 | -0.23 | 0.997 |
| cluster3509 | 0.47 | 0.997 |
| cluster3516 | -0.77 | 0.997 |
| cluster3518 | -0.78 | 0.997 |
| cluster3521 | -0.43 | 0.997 |
| cluster3523 | -0.26 | 0.997 |
| cluster3526 | 0.46 | 0.997 |
| cluster3527 | -0.46 | 0.997 |
| cluster3539 | -0.38 | 0.997 |
| cluster3543 | -0.38 | 0.997 |
| cluster3544 | -0.60 | 0.997 |
| cluster3545 | 0.19 | 0.997 |
| cluster3548 | 0.57 | 0.997 |
| cluster3550 | 0.59 | 0.997 |
| cluster3555 | -0.27 | 0.997 |
| cluster3557 | -0.47 | 0.997 |
| cluster3559 | 0.37 | 0.997 |
| cluster3562 | -0.39 | 0.997 |

| | | |
|---|---|---|
| cluster3566 | -0.45 | 0.997 |
| cluster3569 | 0.25 | 0.997 |
| cluster3572 | 0.96 | 0.997 |
| cluster3576 | 0.31 | 0.997 |
| cluster3584 | 0.48 | 0.997 |
| cluster3586 | 0.40 | 0.997 |
| cluster3588 | 0.24 | 0.997 |
| cluster3591 | -0.32 | 0.997 |
| cluster3594 | -0.54 | 0.997 |
| cluster3596 | -0.52 | 0.997 |
| cluster3600 | 0.51 | 0.997 |
| cluster3601 | 0.21 | 0.997 |
| cluster3604 | 0.36 | 0.997 |
| cluster3606 | -0.33 | 0.997 |
| cluster3608 | -0.49 | 0.997 |
| cluster3620 | -0.34 | 0.997 |
| cluster3623 | -0.50 | 0.997 |
| cluster3624 | 0.34 | 0.997 |
| cluster3626 | 0.24 | 0.997 |
| cluster3627 | -0.39 | 0.997 |
| cluster3630 | 0.26 | 0.997 |
| cluster3632 | 0.49 | 0.997 |
| cluster3635 | 0.35 | 0.997 |
| cluster3637 | 0.33 | 0.997 |
| cluster3638 | 0.51 | 0.997 |
| cluster3640 | -0.96 | 0.997 |
| cluster3642 | 0.25 | 0.997 |
| cluster3643 | 0.62 | 0.997 |
| cluster3647 | 0.24 | 0.997 |
| cluster3648 | 0.20 | 0.997 |
| cluster3650 | 0.78 | 0.997 |
| cluster3654 | -0.41 | 0.997 |
| cluster3655 | -0.44 | 0.997 |
| cluster3658 | 0.57 | 0.997 |
| cluster3660 | -0.44 | 0.997 |
| cluster3663 | -0.48 | 0.997 |
| cluster3664 | 0.31 | 0.997 |
| cluster3665 | -0.80 | 0.997 |
| cluster3669 | -0.54 | 0.997 |
| cluster3670 | -0.52 | 0.997 |
| cluster3671 | -0.44 | 0.997 |
| cluster3673 | -0.59 | 0.997 |
| cluster3675 | -0.35 | 0.997 |
| cluster3676 | 0.69 | 0.997 |

| | | |
|---|---|---|
| cluster3677 | 0.64 | 0.997 |
| cluster3678 | 0.48 | 0.997 |
| cluster3682 | -0.78 | 0.997 |
| cluster3685 | -0.46 | 0.997 |
| cluster3692 | -0.25 | 0.997 |
| cluster3693 | -0.53 | 0.997 |
| cluster3697 | 0.44 | 0.997 |
| cluster3698 | 0.50 | 0.997 |
| cluster3699 | 0.37 | 0.997 |
| cluster3702 | 0.31 | 0.997 |
| cluster3706 | -0.36 | 0.997 |
| cluster3710 | 0.20 | 0.997 |
| cluster3711 | -0.28 | 0.997 |
| cluster3713 | 0.46 | 0.997 |
| cluster3715 | 0.62 | 0.997 |
| cluster3717 | -0.20 | 0.997 |
| cluster3718 | -0.39 | 0.997 |
| cluster3720 | -0.27 | 0.997 |
| cluster3724 | 0.52 | 0.997 |
| cluster3727 | -0.34 | 0.997 |
| cluster3732 | -0.45 | 0.997 |
| cluster3734 | 0.71 | 0.997 |
| cluster3736 | -0.57 | 0.997 |
| cluster3739 | 0.34 | 0.997 |
| cluster3740 | -0.39 | 0.997 |
| cluster3743 | -0.20 | 0.997 |
| cluster3751 | -0.36 | 0.997 |
| cluster3754 | 0.17 | 0.997 |
| cluster3762 | 0.78 | 0.997 |
| cluster3765 | -0.68 | 0.997 |
| cluster3767 | -0.39 | 0.997 |
| cluster3770 | -1.10 | 0.997 |
| cluster3774 | 0.50 | 0.997 |
| cluster3775 | -0.37 | 0.997 |
| cluster3776 | -0.23 | 0.997 |
| cluster3777 | -0.54 | 0.997 |
| cluster3780 | -0.41 | 0.997 |
| cluster3784 | 0.62 | 0.997 |
| cluster3785 | -0.37 | 0.997 |
| cluster3790 | -0.35 | 0.997 |
| cluster3791 | 0.42 | 0.997 |
| cluster3796 | -0.48 | 0.997 |
| cluster3798 | -0.28 | 0.997 |
| cluster3801 | 0.50 | 0.997 |

| | | |
|---|---|---|
| cluster3804 | 0.58 | 0.997 |
| cluster3806 | -0.64 | 0.997 |
| cluster3812 | 0.48 | 0.997 |
| cluster3813 | 0.35 | 0.997 |
| cluster3814 | 0.47 | 0.997 |
| cluster3815 | 0.53 | 0.997 |
| cluster3816 | -0.25 | 0.997 |
| cluster3820 | -0.39 | 0.997 |
| cluster3823 | -0.22 | 0.997 |
| cluster3824 | 0.61 | 0.997 |
| cluster3825 | -0.63 | 0.997 |
| cluster3829 | 0.66 | 0.997 |
| cluster3831 | -0.35 | 0.997 |
| cluster3832 | 0.51 | 0.997 |
| cluster3835 | -0.27 | 0.997 |
| cluster3836 | -0.27 | 0.997 |
| cluster3837 | -0.25 | 0.997 |
| cluster3841 | -0.69 | 0.997 |
| cluster3845 | -0.36 | 0.997 |
| cluster3847 | 0.62 | 0.997 |
| cluster3848 | -0.45 | 0.997 |
| cluster3855 | -0.29 | 0.997 |
| cluster3856 | -0.40 | 0.997 |
| cluster3860 | 0.45 | 0.997 |
| cluster3862 | 0.38 | 0.997 |
| cluster3868 | -0.37 | 0.997 |
| cluster3871 | 0.62 | 0.997 |
| cluster3872 | 0.64 | 0.997 |
| cluster3875 | 0.55 | 0.997 |
| cluster3877 | -0.39 | 0.997 |
| cluster3879 | 0.35 | 0.997 |
| cluster3880 | 0.43 | 0.997 |
| cluster3881 | 0.43 | 0.997 |
| cluster3883 | 0.92 | 0.997 |
| cluster3886 | 0.25 | 0.997 |
| cluster3888 | -0.39 | 0.997 |
| cluster3889 | 0.59 | 0.997 |
| cluster3891 | 0.36 | 0.997 |
| cluster3893 | 0.25 | 0.997 |
| cluster3903 | -0.23 | 0.997 |
| cluster3905 | -1.13 | 0.997 |
| cluster3906 | -0.58 | 0.997 |
| cluster3908 | -0.30 | 0.997 |
| cluster3909 | -0.26 | 0.997 |

| | | |
|---|---|---|
| cluster3910 | -0.32 | 0.997 |
| cluster3911 | 0.55 | 0.997 |
| cluster3912 | -0.22 | 0.997 |
| cluster3913 | -0.35 | 0.997 |
| cluster3914 | -0.44 | 0.997 |
| cluster3917 | 0.27 | 0.997 |
| cluster3923 | -0.37 | 0.997 |
| cluster3924 | 0.26 | 0.997 |
| cluster3925 | -1.64 | 0.997 |
| cluster3926 | -0.43 | 0.997 |
| cluster3936 | 0.44 | 0.997 |
| cluster3938 | -0.31 | 0.997 |
| cluster3943 | 0.58 | 0.997 |
| cluster3946 | -0.48 | 0.997 |
| cluster3947 | 0.21 | 0.997 |
| cluster3949 | -0.33 | 0.997 |
| cluster3950 | 0.19 | 0.997 |
| cluster3957 | 0.25 | 0.997 |
| cluster3962 | 0.25 | 0.997 |
| cluster3963 | -0.80 | 0.997 |
| cluster3965 | -0.39 | 0.997 |
| cluster3967 | 0.52 | 0.997 |
| cluster3968 | 0.31 | 0.997 |
| cluster3971 | 0.61 | 0.997 |
| cluster3972 | -0.45 | 0.997 |
| cluster3974 | 0.36 | 0.997 |
| cluster3975 | 0.45 | 0.997 |
| cluster3976 | -0.42 | 0.997 |
| cluster3977 | -0.73 | 0.997 |
| cluster3980 | 0.34 | 0.997 |
| cluster3984 | -0.92 | 0.997 |
| cluster3991 | -0.33 | 0.997 |
| cluster3993 | -0.46 | 0.997 |
| cluster3996 | 0.46 | 0.997 |
| cluster3997 | 0.51 | 0.997 |
| cluster3998 | 0.47 | 0.997 |
| cluster4004 | -0.17 | 0.997 |
| cluster4006 | -0.18 | 0.997 |
| cluster4007 | 0.37 | 0.997 |
| cluster4008 | 0.29 | 0.997 |
| cluster4009 | 0.63 | 0.997 |
| cluster4018 | 0.24 | 0.997 |
| cluster4021 | 0.34 | 0.997 |
| cluster4024 | 0.39 | 0.997 |

| | | |
|---|---|---|
| cluster4025 | 0.17 | 0.997 |
| cluster4028 | 0.31 | 0.997 |
| cluster4029 | -0.49 | 0.997 |
| cluster4030 | -0.49 | 0.997 |
| cluster4032 | 0.32 | 0.997 |
| cluster4037 | 0.55 | 0.997 |
| cluster4039 | 0.56 | 0.997 |
| cluster4041 | -0.33 | 0.997 |
| cluster4043 | -0.47 | 0.997 |
| cluster4044 | 0.54 | 0.997 |
| cluster4045 | 0.45 | 0.997 |
| cluster4050 | -0.35 | 0.997 |
| cluster4058 | -0.41 | 0.997 |
| cluster4061 | 0.24 | 0.997 |
| cluster4063 | -0.61 | 0.997 |
| cluster4065 | 0.29 | 0.997 |
| cluster4068 | -0.36 | 0.997 |
| cluster4081 | 0.40 | 0.997 |
| cluster4082 | 0.42 | 0.997 |
| cluster4083 | 0.34 | 0.997 |
| cluster4084 | -0.39 | 0.997 |
| cluster4088 | -0.70 | 0.997 |
| cluster4092 | -0.38 | 0.997 |
| cluster4096 | -0.24 | 0.997 |
| cluster4100 | 0.32 | 0.997 |
| cluster4102 | -0.22 | 0.997 |
| cluster4105 | 0.66 | 0.997 |
| cluster4109 | 0.81 | 0.997 |
| cluster4110 | 0.38 | 0.997 |
| cluster4117 | -0.41 | 0.997 |
| cluster4125 | -1.53 | 0.997 |
| cluster2880 | 0.19 | 0.998 |
| cluster2885 | -0.18 | 0.998 |
| cluster2959 | 0.16 | 0.998 |
| cluster2990 | 0.15 | 0.998 |
| cluster3060 | -0.22 | 0.998 |
| cluster3084 | -0.14 | 0.998 |
| cluster3123 | 0.15 | 0.998 |
| cluster3136 | 0.17 | 0.998 |
| cluster3201 | 0.15 | 0.998 |
| cluster3216 | 0.15 | 0.998 |
| cluster3236 | 0.19 | 0.998 |
| cluster3272 | -0.45 | 0.998 |
| cluster3274 | 0.18 | 0.998 |

| | | |
|---|---|---|
| cluster3294 | 0.16 | 0.998 |
| cluster3308 | -0.17 | 0.998 |
| cluster3350 | -0.17 | 0.998 |
| cluster3368 | -0.15 | 0.998 |
| cluster3394 | -0.15 | 0.998 |
| cluster3406 | -0.16 | 0.998 |
| cluster3412 | -0.15 | 0.998 |
| cluster3483 | -0.13 | 0.998 |
| cluster3489 | 0.17 | 0.998 |
| cluster3529 | -0.18 | 0.998 |
| cluster3533 | 0.17 | 0.998 |
| cluster3641 | 0.18 | 0.998 |
| cluster3690 | -0.19 | 0.998 |
| cluster3726 | 0.16 | 0.998 |
| cluster3733 | -0.19 | 0.998 |
| cluster3782 | -0.18 | 0.998 |
| cluster3797 | -0.18 | 0.998 |
| cluster3859 | -0.14 | 0.998 |
| cluster3870 | -0.23 | 0.998 |
| cluster3874 | -0.17 | 0.998 |
| cluster3969 | 0.14 | 0.998 |
| cluster3982 | -0.14 | 0.998 |
| cluster3988 | -0.14 | 0.998 |
| cluster4010 | 0.20 | 0.998 |
| cluster4057 | -0.23 | 0.998 |
| cluster4126 | -0.25 | 0.998 |
| cluster2920 | -0.14 | 0.998 |
| cluster2941 | -0.13 | 0.998 |
| cluster2968 | 0.12 | 0.998 |
| cluster3012 | 0.18 | 0.998 |
| cluster3248 | -0.19 | 0.998 |
| cluster3390 | -0.15 | 0.998 |
| cluster3528 | 0.14 | 0.998 |
| cluster3579 | 0.15 | 0.998 |
| cluster3708 | -0.21 | 0.998 |
| cluster3861 | -0.12 | 0.998 |
| cluster3864 | -0.13 | 0.998 |
| cluster3945 | -0.18 | 0.998 |
| cluster2856 | 0.05 | 0.999 |
| cluster2862 | 0.06 | 0.999 |
| cluster2872 | 0.10 | 0.999 |
| cluster2883 | 0.12 | 0.999 |
| cluster2887 | 0.02 | 0.999 |
| cluster2890 | -0.04 | 0.999 |

| | | |
|---|---|---|
| cluster2893 | -0.06 | 0.999 |
| cluster2896 | -0.02 | 0.999 |
| cluster2897 | -0.11 | 0.999 |
| cluster2917 | -0.05 | 0.999 |
| cluster2918 | 0.01 | 0.999 |
| cluster2938 | -0.11 | 0.999 |
| cluster2951 | 0.05 | 0.999 |
| cluster2953 | 0.12 | 0.999 |
| cluster2955 | 0.11 | 0.999 |
| cluster2958 | 0.05 | 0.999 |
| cluster2971 | -0.12 | 0.999 |
| cluster2974 | -0.05 | 0.999 |
| cluster2986 | -0.05 | 0.999 |
| cluster2996 | -0.02 | 0.999 |
| cluster3003 | 0.01 | 0.999 |
| cluster3018 | -0.03 | 0.999 |
| cluster3028 | 0.09 | 0.999 |
| cluster3029 | -0.06 | 0.999 |
| cluster3038 | -0.12 | 0.999 |
| cluster3040 | -0.13 | 0.999 |
| cluster3041 | 0.01 | 0.999 |
| cluster3043 | -0.06 | 0.999 |
| cluster3044 | 0.03 | 0.999 |
| cluster3054 | -0.05 | 0.999 |
| cluster3056 | -0.04 | 0.999 |
| cluster3058 | 0.08 | 0.999 |
| cluster3072 | 0.12 | 0.999 |
| cluster3086 | -0.03 | 0.999 |
| cluster3090 | 0.20 | 0.999 |
| cluster3092 | -0.02 | 0.999 |
| cluster3096 | -0.02 | 0.999 |
| cluster3108 | 0.04 | 0.999 |
| cluster3116 | 0.07 | 0.999 |
| cluster3139 | 0.04 | 0.999 |
| cluster3146 | -0.01 | 0.999 |
| cluster3152 | 0.02 | 0.999 |
| cluster3154 | 0.04 | 0.999 |
| cluster3158 | -0.04 | 0.999 |
| cluster3172 | 0.01 | 0.999 |
| cluster3173 | 0.13 | 0.999 |
| cluster3184 | -0.09 | 0.999 |
| cluster3189 | -0.02 | 0.999 |
| cluster3193 | -0.07 | 0.999 |
| cluster3196 | 0.03 | 0.999 |

| | | |
|---|---|---|
| cluster3211 | 0.14 | 0.999 |
| cluster3213 | 0.08 | 0.999 |
| cluster3214 | -0.01 | 0.999 |
| cluster3217 | -0.01 | 0.999 |
| cluster3222 | -0.03 | 0.999 |
| cluster3223 | -0.16 | 0.999 |
| cluster3271 | -0.10 | 0.999 |
| cluster3281 | 0.02 | 0.999 |
| cluster3314 | 0.03 | 0.999 |
| cluster3322 | -0.12 | 0.999 |
| cluster3324 | 0.10 | 0.999 |
| cluster3328 | 0.01 | 0.999 |
| cluster3336 | -0.14 | 0.999 |
| cluster3343 | 0.12 | 0.999 |
| cluster3351 | -0.03 | 0.999 |
| cluster3370 | 0.04 | 0.999 |
| cluster3376 | -0.08 | 0.999 |
| cluster3378 | 0.14 | 0.999 |
| cluster3382 | -0.05 | 0.999 |
| cluster3389 | -0.14 | 0.999 |
| cluster3396 | 0.01 | 0.999 |
| cluster3409 | -0.15 | 0.999 |
| cluster3416 | 0.06 | 0.999 |
| cluster3419 | -0.03 | 0.999 |
| cluster3420 | 0.04 | 0.999 |
| cluster3428 | 0.14 | 0.999 |
| cluster3455 | -0.02 | 0.999 |
| cluster3464 | 0.08 | 0.999 |
| cluster3467 | -0.05 | 0.999 |
| cluster3492 | -0.01 | 0.999 |
| cluster3502 | 0.18 | 0.999 |
| cluster3512 | -0.08 | 0.999 |
| cluster3517 | -0.02 | 0.999 |
| cluster3524 | 0.07 | 0.999 |
| cluster3532 | 0.13 | 0.999 |
| cluster3535 | 0.05 | 0.999 |
| cluster3542 | -0.10 | 0.999 |
| cluster3558 | -0.08 | 0.999 |
| cluster3560 | 0.08 | 0.999 |
| cluster3570 | -0.05 | 0.999 |
| cluster3583 | 0.06 | 0.999 |
| cluster3587 | 0.13 | 0.999 |
| cluster3589 | -0.08 | 0.999 |
| cluster3590 | 0.10 | 0.999 |

| | | |
|---|---|---|
| cluster3592 | -0.11 | 0.999 |
| cluster3605 | -0.08 | 0.999 |
| cluster3607 | 0.14 | 0.999 |
| cluster3610 | -0.10 | 0.999 |
| cluster3614 | 0.14 | 0.999 |
| cluster3616 | 0.14 | 0.999 |
| cluster3617 | 0.01 | 0.999 |
| cluster3618 | -0.08 | 0.999 |
| cluster3622 | 0.09 | 0.999 |
| cluster3625 | -0.07 | 0.999 |
| cluster3628 | -0.05 | 0.999 |
| cluster3639 | 0.07 | 0.999 |
| cluster3646 | -0.03 | 0.999 |
| cluster3668 | -0.11 | 0.999 |
| cluster3680 | 0.07 | 0.999 |
| cluster3681 | 0.05 | 0.999 |
| cluster3683 | 0.11 | 0.999 |
| cluster3684 | -0.14 | 0.999 |
| cluster3686 | 0.08 | 0.999 |
| cluster3687 | -0.15 | 0.999 |
| cluster3688 | 0.04 | 0.999 |
| cluster3696 | -0.03 | 0.999 |
| cluster3700 | 0.04 | 0.999 |
| cluster3701 | 0.04 | 0.999 |
| cluster3707 | 0.10 | 0.999 |
| cluster3722 | 0.09 | 0.999 |
| cluster3725 | 0.02 | 0.999 |
| cluster3728 | 0.01 | 0.999 |
| cluster3759 | -0.06 | 0.999 |
| cluster3761 | 0.18 | 0.999 |
| cluster3783 | -0.14 | 0.999 |
| cluster3789 | -0.09 | 0.999 |
| cluster3793 | 0.03 | 0.999 |
| cluster3803 | -0.01 | 0.999 |
| cluster3807 | 0.10 | 0.999 |
| cluster3808 | 0.16 | 0.999 |
| cluster3809 | 0.05 | 0.999 |
| cluster3828 | -0.01 | 0.999 |
| cluster3833 | -0.16 | 0.999 |
| cluster3838 | -0.05 | 0.999 |
| cluster3843 | 0.11 | 0.999 |
| cluster3851 | -0.09 | 0.999 |
| cluster3866 | 0.12 | 0.999 |
| cluster3882 | 0.03 | 0.999 |

| | | |
|---|---|---|
| cluster3895 | -0.06 | 0.999 |
| cluster3896 | -0.08 | 0.999 |
| cluster3900 | 0.08 | 0.999 |
| cluster3921 | 0.14 | 0.999 |
| cluster3927 | -0.01 | 0.999 |
| cluster3939 | 0.12 | 0.999 |
| cluster3941 | -0.09 | 0.999 |
| cluster3944 | 0.10 | 0.999 |
| cluster3961 | 0.10 | 0.999 |
| cluster3986 | -0.01 | 0.999 |
| cluster4003 | -0.02 | 0.999 |
| cluster4014 | -0.01 | 0.999 |
| cluster4017 | 0.05 | 0.999 |
| cluster4020 | 0.09 | 0.999 |
| cluster4027 | -0.02 | 0.999 |
| cluster4036 | 0.08 | 0.999 |
| cluster4038 | -0.04 | 0.999 |
| cluster4040 | -0.10 | 0.999 |
| cluster4069 | -0.09 | 0.999 |
| cluster4071 | -0.12 | 0.999 |
| cluster4077 | -0.07 | 0.999 |
| cluster4104 | 0.01 | 0.999 |
| cluster4108 | -0.04 | 0.999 |
| cluster4123 | -0.11 | 0.999 |
| cluster3951 | 0.01 | 0.999 |
| cluster3612 | 0.00 | 0.999 |
| cluster3846 | 0.00 | 0.999 |
| cluster3935 | 0.00 | 0.999 |
| cluster3333 | 0.00 | 0.999 |

**Supplementary Table 8.** edgeR results with eXpress counts - Oestridae vs Calliphoridae

| ID | log2FoldChange | Adjusted p value |
|---|---|---|
| cluster4046 | -3.39 | 0.665 |
| cluster4013 | -4.44 | 0.665 |
| cluster3461 | -2.92 | 0.885 |
| cluster3178 | 2.97 | 0.885 |
| cluster3786 | 3.42 | 0.885 |
| cluster4101 | 4.05 | 0.885 |
| cluster3574 | -2.66 | 0.885 |
| cluster3233 | 3.62 | 0.885 |
| cluster3852 | 2.64 | 0.885 |
| cluster4118 | -3.31 | 0.885 |
| cluster3645 | -3.19 | 0.885 |
| cluster3137 | 2.69 | 0.885 |
| cluster4112 | 3.30 | 0.885 |
| cluster3755 | 3.63 | 0.885 |
| cluster3651 | -2.67 | 0.885 |
| cluster3932 | -2.70 | 0.885 |
| cluster3741 | 2.21 | 0.885 |
| cluster3738 | 3.16 | 0.885 |
| cluster4111 | -2.66 | 0.885 |
| cluster4122 | -2.72 | 0.885 |
| cluster3652 | 2.37 | 0.885 |
| cluster3840 | 2.34 | 0.885 |
| cluster3763 | 2.66 | 0.885 |
| cluster3030 | 2.70 | 0.885 |
| cluster4076 | 2.94 | 0.885 |
| cluster4012 | 2.39 | 0.885 |
| cluster4087 | 3.51 | 0.885 |
| cluster3250 | 2.68 | 0.885 |
| cluster3006 | 2.80 | 0.885 |
| cluster3309 | 2.21 | 0.969 |
| cluster3890 | 2.44 | 0.969 |
| cluster3303 | 2.69 | 0.969 |
| cluster3297 | 2.30 | 0.969 |
| cluster3666 | 2.95 | 0.969 |
| cluster3897 | 2.25 | 0.969 |
| cluster3802 | 2.69 | 0.969 |
| cluster3393 | -2.19 | 0.969 |
| cluster3397 | -1.84 | 0.969 |
| cluster4060 | 2.47 | 0.978 |
| cluster3000 | -1.72 | 0.978 |
| cluster4001 | -1.77 | 0.978 |
| cluster3063 | 2.05 | 0.978 |

| | | |
|---|---|---|
| cluster2861 | 1.90 | 0.978 |
| cluster4116 | -2.50 | 0.978 |
| cluster4052 | 2.22 | 0.978 |
| cluster4098 | 2.40 | 0.978 |
| cluster2902 | 2.24 | 0.978 |
| cluster3323 | 2.16 | 0.978 |
| cluster3631 | 2.43 | 0.978 |
| cluster2921 | 1.76 | 0.978 |
| cluster4067 | 3.49 | 0.978 |
| cluster4048 | -2.23 | 0.978 |
| cluster4078 | 1.93 | 0.978 |
| cluster2943 | 1.86 | 0.978 |
| cluster3839 | -1.97 | 0.978 |
| cluster3904 | -1.66 | 0.978 |
| cluster3704 | 1.92 | 0.978 |
| cluster3750 | -2.12 | 0.978 |
| cluster3317 | 1.98 | 0.978 |
| cluster3045 | 1.81 | 0.978 |
| cluster3876 | 1.72 | 0.978 |
| cluster3501 | -1.78 | 0.978 |
| cluster3138 | 1.84 | 0.978 |
| cluster3561 | 1.98 | 0.978 |
| cluster3958 | -2.06 | 0.978 |
| cluster4090 | -2.33 | 0.978 |
| cluster4079 | 1.92 | 0.978 |
| cluster3981 | 2.10 | 0.978 |
| cluster3357 | 1.62 | 0.978 |
| cluster4073 | 2.73 | 0.978 |
| cluster2948 | 2.11 | 0.978 |
| cluster3388 | -1.63 | 0.978 |
| cluster2993 | 1.79 | 0.978 |
| cluster3482 | -1.47 | 0.978 |
| cluster3818 | 2.05 | 0.978 |
| cluster3126 | 2.09 | 0.978 |
| cluster3367 | 2.26 | 0.978 |
| cluster3691 | 1.77 | 0.978 |
| cluster4070 | -2.29 | 0.978 |
| cluster3534 | -1.57 | 0.978 |
| cluster3546 | -1.85 | 0.978 |
| cluster3200 | 1.63 | 0.978 |
| cluster3772 | -1.47 | 0.978 |
| cluster4120 | -1.98 | 0.978 |
| cluster2947 | 1.54 | 0.978 |
| cluster3432 | -1.46 | 0.978 |

| | | |
|---|---|---|
| cluster2967 | 1.80 | 0.978 |
| cluster3915 | -1.82 | 0.978 |
| cluster3745 | 2.34 | 0.978 |
| cluster3694 | -1.36 | 0.978 |
| cluster3597 | 1.67 | 0.978 |
| cluster3933 | -1.43 | 0.978 |
| cluster3100 | 2.16 | 0.978 |
| cluster3901 | 2.09 | 0.978 |
| cluster3672 | -1.48 | 0.978 |
| cluster3863 | 1.82 | 0.978 |
| cluster3101 | 1.50 | 0.978 |
| cluster3595 | 1.54 | 0.978 |
| cluster2881 | 1.62 | 0.978 |
| cluster2961 | 1.57 | 0.978 |
| cluster3130 | -1.31 | 0.978 |
| cluster3564 | 1.94 | 0.978 |
| cluster4099 | 1.63 | 0.978 |
| cluster3306 | 2.07 | 0.978 |
| cluster2962 | 1.67 | 0.978 |
| cluster3656 | 1.58 | 0.978 |
| cluster3615 | 1.85 | 0.978 |
| cluster3118 | 1.47 | 0.978 |
| cluster2930 | 1.42 | 0.978 |
| cluster3563 | -1.31 | 0.978 |
| cluster3760 | 1.50 | 0.978 |
| cluster3177 | -1.57 | 0.978 |
| cluster3168 | 1.50 | 0.978 |
| cluster4119 | 2.44 | 0.978 |
| cluster3426 | 1.58 | 0.978 |
| cluster3551 | 1.69 | 0.978 |
| cluster2854 | 1.61 | 0.978 |
| cluster4095 | 1.92 | 0.978 |
| cluster3940 | 1.54 | 0.978 |
| cluster3015 | 1.73 | 0.978 |
| cluster3853 | 1.79 | 0.978 |
| cluster3865 | 2.22 | 0.978 |
| cluster4114 | 2.87 | 0.978 |
| cluster3474 | 1.94 | 0.978 |
| cluster3719 | 1.85 | 0.978 |
| cluster4074 | 2.27 | 0.978 |
| cluster3987 | 1.64 | 0.978 |
| cluster3240 | 1.54 | 0.978 |
| cluster4034 | -2.07 | 0.978 |
| cluster2977 | 1.35 | 0.978 |

| | | |
|---|---|---|
| cluster3237 | 1.89 | 0.978 |
| cluster3885 | -1.22 | 0.978 |
| cluster3335 | 1.62 | 0.978 |
| cluster3197 | 1.49 | 0.978 |
| cluster3689 | -1.88 | 0.978 |
| cluster3290 | -1.43 | 0.978 |
| cluster4016 | 1.71 | 0.978 |
| cluster3371 | 2.77 | 0.978 |
| cluster3568 | -1.56 | 0.978 |
| cluster3514 | 1.62 | 0.978 |
| cluster3245 | 1.78 | 0.978 |
| cluster3556 | 1.68 | 0.978 |
| cluster3053 | 1.72 | 0.978 |
| cluster3009 | -1.28 | 0.978 |
| cluster3176 | -1.41 | 0.978 |
| cluster4002 | 1.79 | 0.978 |
| cluster3430 | 2.18 | 0.978 |
| cluster3830 | 1.33 | 0.978 |
| cluster3246 | 1.37 | 0.981 |
| cluster3898 | 1.76 | 0.981 |
| cluster3834 | -1.41 | 0.990 |
| cluster2989 | 1.54 | 0.990 |
| cluster3575 | 1.63 | 0.990 |
| cluster3332 | 1.48 | 0.990 |
| cluster4055 | -1.24 | 0.990 |
| cluster2912 | 1.35 | 0.990 |
| cluster3102 | 1.59 | 0.990 |
| cluster3431 | 1.45 | 0.990 |
| cluster3705 | 1.65 | 0.990 |
| cluster4049 | -2.14 | 0.990 |
| cluster2960 | 1.61 | 0.990 |
| cluster4053 | -1.25 | 0.990 |
| cluster3400 | 1.31 | 0.990 |
| cluster4103 | 1.81 | 0.990 |
| cluster3105 | -1.48 | 0.990 |
| cluster3994 | -1.54 | 0.990 |
| cluster3757 | 2.19 | 0.990 |
| cluster3735 | -1.22 | 0.994 |
| cluster3571 | -1.46 | 0.994 |
| cluster3448 | -1.56 | 0.994 |
| cluster3157 | 1.46 | 0.994 |
| cluster3537 | 1.34 | 0.994 |
| cluster3097 | 1.40 | 0.994 |
| cluster3920 | -1.36 | 0.994 |

| | | |
|---|---|---|
| cluster3599 | -1.36 | 0.994 |
| cluster3363 | 1.32 | 0.994 |
| cluster4033 | -1.54 | 0.994 |
| cluster3023 | 1.29 | 0.994 |
| cluster3438 | 1.51 | 0.994 |
| cluster4121 | 1.61 | 0.994 |
| cluster3369 | 1.22 | 0.994 |
| cluster3435 | -1.46 | 0.994 |
| cluster3619 | 1.30 | 0.994 |
| cluster3822 | 1.93 | 0.994 |
| cluster3110 | 1.16 | 0.994 |
| cluster2924 | 1.37 | 0.994 |
| cluster3746 | 1.42 | 0.994 |
| cluster3931 | 1.69 | 0.994 |
| cluster3952 | 1.33 | 0.994 |
| cluster3480 | 1.34 | 0.994 |
| cluster3462 | -1.12 | 0.994 |
| cluster3165 | 1.53 | 0.994 |
| cluster3956 | 1.28 | 0.994 |
| cluster3032 | 1.25 | 0.994 |
| cluster3008 | 1.32 | 0.994 |
| cluster3525 | -1.04 | 0.994 |
| cluster3538 | 1.50 | 0.994 |
| cluster3995 | -1.34 | 0.994 |
| cluster2860 | 1.38 | 0.994 |
| cluster3061 | 1.27 | 0.994 |
| cluster3337 | 1.39 | 0.994 |
| cluster4085 | -1.28 | 0.994 |
| cluster3179 | 1.57 | 0.994 |
| cluster2899 | 1.46 | 0.994 |
| cluster3510 | -1.64 | 0.994 |
| cluster4080 | -1.51 | 0.994 |
| cluster4000 | 1.41 | 0.994 |
| cluster2995 | 1.36 | 0.994 |
| cluster3019 | 1.15 | 0.994 |
| cluster2984 | 1.38 | 0.994 |
| cluster3494 | 1.30 | 0.994 |
| cluster4011 | -1.51 | 0.994 |
| cluster3842 | 1.44 | 0.994 |
| cluster2933 | 1.36 | 0.994 |
| cluster3858 | 1.42 | 0.994 |
| cluster3066 | 1.23 | 0.994 |
| cluster3792 | -0.95 | 0.994 |
| cluster3436 | 1.84 | 0.994 |

| | | |
|---|---|---|
| cluster3212 | -1.20 | 0.994 |
| cluster3955 | -1.41 | 0.994 |
| cluster2980 | -1.38 | 0.994 |
| cluster3026 | -1.22 | 0.994 |
| cluster3478 | 1.21 | 0.994 |
| cluster3629 | 1.48 | 0.994 |
| cluster3247 | 1.30 | 0.994 |
| cluster3141 | 1.11 | 0.994 |
| cluster3522 | -1.18 | 0.994 |
| cluster3304 | 1.17 | 0.994 |
| cluster2969 | 1.32 | 0.994 |
| cluster3519 | -1.48 | 0.994 |
| cluster3849 | 1.25 | 0.994 |
| cluster3730 | 1.18 | 0.994 |
| cluster3312 | -1.41 | 0.994 |
| cluster3582 | 1.11 | 0.994 |
| cluster3085 | -1.18 | 0.994 |
| cluster3244 | 1.16 | 0.994 |
| cluster2892 | 1.10 | 0.994 |
| cluster3485 | -1.43 | 0.994 |
| cluster3150 | -1.23 | 0.994 |
| cluster3799 | 1.26 | 0.994 |
| cluster4019 | 1.51 | 0.994 |
| cluster3549 | 1.12 | 0.994 |
| cluster3095 | -1.01 | 0.994 |
| cluster3401 | 1.05 | 0.994 |
| cluster3731 | 1.30 | 0.994 |
| cluster3027 | 1.10 | 0.994 |
| cluster3288 | 1.34 | 0.994 |
| cluster3311 | 1.20 | 0.994 |
| cluster3709 | 1.10 | 0.994 |
| cluster3990 | 1.18 | 0.994 |
| cluster3513 | 1.17 | 0.994 |
| cluster3787 | -1.20 | 0.994 |
| cluster3844 | -1.46 | 0.994 |
| cluster3454 | -0.93 | 0.994 |
| cluster3679 | 1.15 | 0.994 |
| cluster3873 | 1.02 | 0.994 |
| cluster2942 | 1.12 | 0.994 |
| cluster4093 | 1.74 | 0.994 |
| cluster3161 | -0.97 | 0.994 |
| cluster3659 | -1.12 | 0.994 |
| cluster4035 | 1.27 | 0.994 |
| cluster3721 | 1.38 | 0.994 |

| | | |
|---|---|---|
| cluster3902 | 1.13 | 0.994 |
| cluster3577 | 1.11 | 0.994 |
| cluster3766 | -1.03 | 0.994 |
| cluster3753 | -1.31 | 0.994 |
| cluster4115 | -1.35 | 0.994 |
| cluster2946 | 1.20 | 0.994 |
| cluster3899 | 1.21 | 0.994 |
| cluster3119 | 1.16 | 0.994 |
| cluster2965 | 1.03 | 0.994 |
| cluster3520 | 1.83 | 0.994 |
| cluster3960 | -1.30 | 0.994 |
| cluster2873 | 1.09 | 0.994 |
| cluster3850 | -1.02 | 0.994 |
| cluster3365 | 1.29 | 0.994 |
| cluster3714 | 1.34 | 0.994 |
| cluster3175 | -1.12 | 0.994 |
| cluster2926 | 1.07 | 0.994 |
| cluster3013 | 1.06 | 0.994 |
| cluster3661 | 1.05 | 0.994 |
| cluster4064 | 1.25 | 0.994 |
| cluster3469 | 1.23 | 0.994 |
| cluster3380 | -0.91 | 0.994 |
| cluster3983 | 1.45 | 0.994 |
| cluster3147 | 0.95 | 0.994 |
| cluster3473 | 1.23 | 0.994 |
| cluster3413 | -1.08 | 0.994 |
| cluster3508 | 1.51 | 0.994 |
| cluster3089 | 1.10 | 0.994 |
| cluster3082 | -1.29 | 0.994 |
| cluster3985 | 1.14 | 0.994 |
| cluster3052 | -1.20 | 0.994 |
| cluster3937 | 1.21 | 0.994 |
| cluster3341 | -1.00 | 0.994 |
| cluster3075 | -0.94 | 0.994 |
| cluster3434 | -1.02 | 0.994 |
| cluster3752 | -1.26 | 0.994 |
| cluster3458 | 0.97 | 0.994 |
| cluster3164 | -1.15 | 0.994 |
| cluster3869 | 1.58 | 0.994 |
| cluster3581 | -0.97 | 0.994 |
| cluster3142 | 1.22 | 0.994 |
| cluster3258 | 0.97 | 0.994 |
| cluster3788 | 1.08 | 0.994 |
| cluster3811 | 0.98 | 0.994 |

| | | |
|---|---|---|
| cluster3188 | 1.18 | 0.994 |
| cluster3609 | 1.56 | 0.994 |
| cluster3453 | -0.97 | 0.994 |
| cluster3497 | 1.02 | 0.994 |
| cluster3515 | 1.42 | 0.994 |
| cluster3083 | 0.97 | 0.994 |
| cluster3185 | -0.99 | 0.994 |
| cluster3391 | 1.05 | 0.994 |
| cluster4072 | 0.93 | 0.994 |
| cluster3712 | 1.11 | 0.994 |
| cluster3017 | 1.17 | 0.994 |
| cluster3794 | -0.97 | 0.994 |
| cluster3135 | -0.91 | 0.994 |
| cluster3636 | -0.77 | 0.994 |
| cluster3398 | -0.82 | 0.994 |
| cluster3346 | 1.10 | 0.994 |
| cluster3536 | 0.95 | 0.994 |
| cluster3005 | 0.89 | 0.994 |
| cluster3181 | 1.00 | 0.994 |
| cluster3129 | -1.29 | 0.994 |
| cluster3182 | -0.94 | 0.994 |
| cluster3280 | 0.91 | 0.994 |
| cluster3292 | -1.03 | 0.994 |
| cluster3295 | 0.88 | 0.994 |
| cluster3703 | 0.91 | 0.994 |
| cluster3552 | 0.94 | 0.994 |
| cluster4127 | -1.35 | 0.994 |
| cluster3495 | 0.93 | 0.994 |
| cluster4125 | -1.53 | 0.994 |
| cluster3439 | -0.84 | 0.994 |
| cluster4066 | -0.98 | 0.994 |
| cluster3621 | 1.21 | 0.994 |
| cluster3226 | -0.94 | 0.994 |
| cluster3653 | 1.02 | 0.994 |
| cluster3585 | -0.82 | 0.994 |
| cluster3220 | -0.88 | 0.994 |
| cluster3970 | 1.31 | 0.994 |
| cluster2901 | -0.91 | 0.994 |
| cluster3810 | 0.99 | 0.994 |
| cluster3919 | -1.07 | 0.994 |
| cluster2911 | -0.83 | 0.994 |
| cluster3887 | -1.05 | 0.994 |
| cluster3162 | 1.13 | 0.994 |
| cluster3402 | 1.02 | 0.994 |

| | | |
|---|---|---|
| cluster3481 | -1.13 | 0.994 |
| cluster3500 | -0.96 | 0.994 |
| cluster2863 | 0.85 | 0.994 |
| cluster3441 | 0.95 | 0.994 |
| cluster3313 | -0.88 | 0.994 |
| cluster3634 | 1.00 | 0.994 |
| cluster3437 | -0.86 | 0.994 |
| cluster3779 | 0.99 | 0.994 |
| cluster3329 | -1.12 | 0.994 |
| cluster3239 | 0.85 | 0.994 |
| cluster3425 | -0.86 | 0.994 |
| cluster3916 | 0.94 | 0.994 |
| cluster2964 | 0.88 | 0.994 |
| cluster2906 | 1.00 | 0.994 |
| cluster3194 | -0.84 | 0.994 |
| cluster3821 | -0.82 | 0.994 |
| cluster3884 | -1.02 | 0.994 |
| cluster4075 | 1.52 | 0.994 |
| cluster2908 | 0.93 | 0.994 |
| cluster3894 | -0.82 | 0.994 |
| cluster3334 | 0.92 | 0.994 |
| cluster2963 | 1.06 | 0.994 |
| cluster3340 | 0.88 | 0.994 |
| cluster3918 | 1.12 | 0.994 |
| cluster3459 | -1.03 | 0.994 |
| cluster2978 | 1.00 | 0.994 |
| cluster3953 | -0.74 | 0.994 |
| cluster3800 | -0.90 | 0.994 |
| cluster2882 | 0.83 | 0.994 |
| cluster4047 | 1.50 | 0.994 |
| cluster4107 | 1.39 | 0.994 |
| cluster2994 | 0.89 | 0.994 |
| cluster2916 | 1.02 | 0.994 |
| cluster3143 | 0.91 | 0.994 |
| cluster3062 | -0.77 | 0.994 |
| cluster3468 | 0.96 | 0.994 |
| cluster3999 | -0.98 | 0.994 |
| cluster3113 | 0.83 | 0.994 |
| cluster3408 | 1.08 | 0.994 |
| cluster3827 | -0.90 | 0.994 |
| cluster3356 | 0.96 | 0.994 |
| cluster3573 | -0.82 | 0.994 |
| cluster3580 | 1.29 | 0.994 |
| cluster3771 | 1.06 | 0.994 |

| | | |
|---|---|---|
| cluster3778 | 1.00 | 0.994 |
| cluster4094 | 1.10 | 0.994 |
| cluster3758 | -0.86 | 0.994 |
| cluster3723 | 0.79 | 0.994 |
| cluster3076 | 0.99 | 0.994 |
| cluster3756 | -0.81 | 0.994 |
| cluster3088 | 0.94 | 0.994 |
| cluster3399 | 1.06 | 0.994 |
| cluster4097 | -1.27 | 0.994 |
| cluster3530 | 0.82 | 0.994 |
| cluster3892 | -0.88 | 0.994 |
| cluster3410 | 1.04 | 0.994 |
| cluster2949 | 0.79 | 0.994 |
| cluster4056 | -0.85 | 0.994 |
| cluster3907 | -0.98 | 0.994 |
| cluster2928 | 0.93 | 0.994 |
| cluster3132 | 0.85 | 0.994 |
| cluster3325 | -1.15 | 0.994 |
| cluster3541 | 0.86 | 0.994 |
| cluster3064 | 1.02 | 0.994 |
| cluster3930 | 0.90 | 0.994 |
| cluster2999 | -0.82 | 0.994 |
| cluster3979 | 0.86 | 0.994 |
| cluster3300 | 0.94 | 0.994 |
| cluster3657 | 0.87 | 0.994 |
| cluster3603 | 0.97 | 0.994 |
| cluster2877 | 0.76 | 0.994 |
| cluster2972 | -0.80 | 0.994 |
| cluster3611 | 1.24 | 0.994 |
| cluster3493 | 1.12 | 0.994 |
| cluster3490 | -0.87 | 0.994 |
| cluster3805 | 0.92 | 0.994 |
| cluster4054 | 1.12 | 0.994 |
| cluster3695 | -0.74 | 0.994 |
| cluster3716 | -0.91 | 0.994 |
| cluster2865 | 0.82 | 0.994 |
| cluster3011 | 0.95 | 0.994 |
| cluster3567 | 1.26 | 0.994 |
| cluster3934 | 0.74 | 0.994 |
| cluster3742 | 1.18 | 0.994 |
| cluster3925 | -1.52 | 0.994 |
| cluster3320 | -0.81 | 0.994 |
| cluster3296 | -0.81 | 0.994 |
| cluster2905 | 0.85 | 0.994 |

| | | |
|---|---|---|
| cluster3442 | 1.10 | 0.994 |
| cluster3260 | -0.71 | 0.994 |
| cluster3747 | 0.83 | 0.994 |
| cluster3238 | 0.73 | 0.994 |
| cluster3973 | -0.75 | 0.994 |
| cluster2884 | 0.74 | 0.994 |
| cluster3229 | 0.96 | 0.994 |
| cluster3463 | 0.96 | 0.994 |
| cluster3081 | -0.72 | 0.994 |
| cluster4124 | -0.95 | 0.994 |
| cluster3817 | 1.09 | 0.994 |
| cluster3487 | -0.77 | 0.994 |
| cluster3452 | -0.79 | 0.994 |
| cluster3174 | 0.98 | 0.994 |
| cluster2981 | 0.79 | 0.994 |
| cluster3338 | -0.83 | 0.994 |
| cluster3768 | -0.83 | 0.994 |
| cluster3153 | 0.76 | 0.994 |
| cluster3349 | 0.95 | 0.994 |
| cluster3598 | 1.44 | 0.994 |
| cluster4062 | -1.03 | 0.994 |
| cluster3677 | 0.82 | 0.994 |
| cluster4051 | 1.06 | 0.994 |
| cluster3149 | 0.74 | 0.994 |
| cluster2853 | 0.93 | 0.994 |
| cluster3662 | -0.83 | 0.994 |
| cluster3978 | -0.65 | 0.994 |
| cluster3443 | 0.95 | 0.994 |
| cluster3131 | -0.69 | 0.994 |
| cluster3071 | 0.79 | 0.994 |
| cluster3417 | 0.96 | 0.994 |
| cluster3734 | 0.99 | 0.994 |
| cluster3180 | -0.64 | 0.994 |
| cluster3578 | -0.79 | 0.994 |
| cluster4023 | -0.83 | 0.994 |
| cluster3449 | 0.80 | 0.994 |
| cluster3773 | -0.68 | 0.994 |
| cluster3847 | 0.81 | 0.994 |
| cluster2919 | 0.77 | 0.994 |
| cluster2998 | -0.59 | 0.994 |
| cluster3257 | 0.73 | 0.994 |
| cluster3644 | -0.73 | 0.994 |
| cluster3077 | 0.88 | 0.994 |
| cluster3446 | -0.64 | 0.994 |

| | | |
|---|---|---|
| cluster4086 | 1.05 | 0.994 |
| cluster3195 | -0.65 | 0.994 |
| cluster3049 | 0.80 | 0.994 |
| cluster3319 | -0.72 | 0.994 |
| cluster3353 | 0.69 | 0.994 |
| cluster3737 | 0.70 | 0.994 |
| cluster3871 | 0.87 | 0.994 |
| cluster3781 | 0.89 | 0.994 |
| cluster4037 | 0.79 | 0.994 |
| cluster2957 | -0.79 | 0.994 |
| cluster3115 | 0.70 | 0.994 |
| cluster3748 | 0.78 | 0.994 |
| cluster3267 | 0.73 | 0.994 |
| cluster3375 | -0.65 | 0.994 |
| cluster3613 | 0.89 | 0.994 |
| cluster3099 | 0.76 | 0.994 |
| cluster3471 | 0.86 | 0.994 |
| cluster3232 | 0.77 | 0.994 |
| cluster3484 | -0.86 | 0.994 |
| cluster3321 | 0.71 | 0.994 |
| cluster3004 | 0.75 | 0.994 |
| cluster3959 | -0.87 | 0.994 |
| cluster3989 | 0.88 | 0.994 |
| cluster3117 | 1.10 | 0.994 |
| cluster3414 | 0.77 | 0.994 |
| cluster3327 | 0.85 | 0.994 |
| cluster2975 | 0.68 | 0.994 |
| cluster2876 | 1.13 | 0.994 |
| cluster3600 | 0.67 | 0.994 |
| cluster3547 | 0.72 | 0.994 |
| cluster3166 | 0.69 | 0.994 |
| cluster2898 | 0.84 | 0.994 |
| cluster3875 | 0.77 | 0.994 |
| cluster3948 | -0.62 | 0.994 |
| cluster3883 | 1.14 | 0.994 |
| cluster3593 | -0.71 | 0.994 |
| cluster3460 | 0.88 | 0.994 |
| cluster3156 | -0.64 | 0.994 |
| cluster3465 | 0.76 | 0.994 |
| cluster3218 | 0.93 | 0.994 |
| cluster3971 | 0.89 | 0.994 |
| cluster3954 | -0.85 | 0.994 |
| cluster4113 | -0.63 | 0.994 |
| cluster2915 | -0.66 | 0.994 |

| | | |
|---|---|---|
| cluster4109 | 0.99 | 0.994 |
| cluster2904 | 0.83 | 0.994 |
| cluster3966 | 0.68 | 0.994 |
| cluster2987 | 0.68 | 0.994 |
| cluster3207 | 0.79 | 0.994 |
| cluster3832 | 0.77 | 0.994 |
| cluster3287 | 0.84 | 0.994 |
| cluster3678 | 0.72 | 0.994 |
| cluster3348 | 0.81 | 0.994 |
| cluster3764 | -0.77 | 0.994 |
| cluster3572 | 1.22 | 0.994 |
| cluster3191 | -0.86 | 0.994 |
| cluster3650 | 0.89 | 0.994 |
| cluster3377 | -0.65 | 0.994 |
| cluster3992 | 0.82 | 0.994 |
| cluster2894 | -0.75 | 0.994 |
| cluster3540 | -0.78 | 0.994 |
| cluster3051 | -0.72 | 0.994 |
| cluster3565 | -0.67 | 0.994 |
| cluster3204 | 0.76 | 0.994 |
| cluster4042 | -0.70 | 0.994 |
| cluster3554 | -0.63 | 0.994 |
| cluster2935 | 0.72 | 0.994 |
| cluster4105 | 0.83 | 0.994 |
| cluster3187 | -0.78 | 0.994 |
| cluster3261 | -0.75 | 0.994 |
| cluster3276 | 0.66 | 0.994 |
| cluster2864 | 0.66 | 0.994 |
| cluster4044 | 0.82 | 0.994 |
| cluster3715 | 0.88 | 0.994 |
| cluster3210 | 0.65 | 0.994 |
| cluster3824 | 0.95 | 0.994 |
| cluster3658 | 0.70 | 0.994 |
| cluster3016 | -0.80 | 0.994 |
| cluster3395 | -0.59 | 0.994 |
| cluster3021 | 0.63 | 0.994 |
| cluster3676 | 0.95 | 0.994 |
| cluster2923 | 0.62 | 0.994 |
| cluster3242 | 0.71 | 0.994 |
| cluster3366 | 0.94 | 0.994 |
| cluster3548 | 0.66 | 0.994 |
| cluster3318 | 0.77 | 0.994 |
| cluster4106 | -0.83 | 0.994 |
| cluster3486 | 0.76 | 0.994 |

| | | |
|---|---|---|
| cluster3872 | 0.82 | 0.994 |
| cluster3749 | -0.75 | 0.994 |
| cluster3361 | -0.56 | 0.994 |
| cluster3784 | 0.73 | 0.994 |
| cluster3509 | 0.64 | 0.994 |
| cluster3905 | -0.98 | 0.994 |
| cluster3602 | 0.71 | 0.994 |
| cluster4015 | -0.60 | 0.994 |
| cluster3649 | -0.53 | 0.994 |
| cluster3633 | -0.58 | 0.994 |
| cluster2936 | 0.92 | 0.994 |
| cluster3975 | 0.66 | 0.994 |
| cluster3826 | -0.69 | 0.994 |
| cluster3804 | 0.74 | 0.994 |
| cluster4059 | -0.77 | 0.994 |
| cluster3922 | -0.76 | 0.994 |
| cluster4005 | -0.64 | 0.994 |
| cluster3526 | 0.60 | 0.994 |
| cluster3107 | 0.60 | 0.994 |
| cluster4128 | -0.82 | 0.994 |
| cluster3472 | 0.63 | 0.994 |
| cluster3795 | -0.53 | 0.994 |
| cluster2950 | 0.63 | 0.994 |
| cluster3667 | -0.53 | 0.994 |
| cluster3255 | -0.64 | 0.994 |
| cluster3215 | 0.57 | 0.994 |
| cluster3496 | 0.68 | 0.994 |
| cluster3273 | 0.62 | 0.994 |
| cluster4091 | -0.80 | 0.994 |
| cluster2878 | 0.68 | 0.994 |
| cluster3039 | 0.60 | 0.994 |
| cluster3862 | 0.56 | 0.994 |
| cluster3358 | 0.70 | 0.994 |
| cluster3996 | 0.64 | 0.994 |
| cluster3106 | 0.61 | 0.994 |
| cluster2940 | -0.52 | 0.994 |
| cluster2914 | -0.59 | 0.994 |
| cluster3315 | 0.62 | 0.994 |
| cluster3427 | -0.63 | 0.994 |
| cluster3825 | -0.58 | 0.994 |
| cluster3964 | -0.73 | 0.994 |
| cluster3829 | 0.85 | 0.994 |
| cluster3291 | 0.64 | 0.994 |
| cluster2879 | 0.62 | 0.994 |

| | | |
|---|---|---|
| cluster4081 | 0.62 | 0.994 |
| cluster3729 | -0.49 | 0.994 |
| cluster3724 | 0.76 | 0.994 |
| cluster3094 | 0.86 | 0.994 |
| cluster3263 | 0.57 | 0.994 |
| cluster3713 | 0.61 | 0.994 |
| cluster3070 | 0.61 | 0.994 |
| cluster3586 | 0.56 | 0.994 |
| cluster2927 | 0.76 | 0.994 |
| cluster3225 | -0.64 | 0.994 |
| cluster3867 | -0.48 | 0.994 |
| cluster3765 | -0.52 | 0.994 |
| cluster4009 | 0.80 | 0.994 |
| cluster3942 | -0.50 | 0.994 |
| cluster3682 | -0.66 | 0.994 |
| cluster2870 | 0.57 | 0.994 |
| cluster3451 | 1.19 | 0.994 |
| cluster3235 | 0.86 | 0.994 |
| cluster3762 | 0.73 | 0.994 |
| cluster3034 | -0.50 | 0.994 |
| cluster3643 | 0.82 | 0.994 |
| cluster3457 | -0.59 | 0.994 |
| cluster3769 | -0.48 | 0.994 |
| cluster3505 | 0.56 | 0.994 |
| cluster3224 | -0.48 | 0.994 |
| cluster3638 | 0.78 | 0.994 |
| cluster3819 | -0.66 | 0.994 |
| cluster3067 | 0.56 | 0.994 |
| cluster3065 | 0.59 | 0.994 |
| cluster3928 | -0.49 | 0.994 |
| cluster3518 | -0.63 | 0.994 |
| cluster3929 | -0.58 | 0.994 |
| cluster3392 | -0.61 | 0.994 |
| cluster3770 | -0.79 | 0.994 |
| cluster3624 | 0.54 | 0.994 |
| cluster3269 | -0.62 | 0.994 |
| cluster3854 | -0.45 | 0.994 |
| cluster2891 | -0.54 | 0.994 |
| cluster3289 | 0.58 | 0.994 |
| cluster3744 | -0.50 | 0.994 |
| cluster3342 | 0.68 | 0.994 |
| cluster3243 | 0.60 | 0.994 |
| cluster3121 | 0.63 | 0.994 |
| cluster3984 | -0.70 | 0.994 |

| | | |
|---|---|---|
| cluster3450 | -0.72 | 0.994 |
| cluster3977 | -0.58 | 0.994 |
| cluster2956 | -0.48 | 0.994 |
| cluster3055 | 0.52 | 0.994 |
| cluster3943 | 0.79 | 0.994 |
| cluster3405 | 0.57 | 0.994 |
| cluster3857 | -0.51 | 0.994 |
| cluster3997 | 0.77 | 0.994 |
| cluster4026 | -0.58 | 0.994 |
| cluster3632 | 0.57 | 0.994 |
| cluster3373 | 0.61 | 0.994 |
| cluster3640 | -0.81 | 0.994 |
| cluster3128 | 0.51 | 0.994 |
| cluster2982 | 0.56 | 0.994 |
| cluster3228 | 0.57 | 0.994 |
| cluster3889 | 0.85 | 0.994 |
| cluster3293 | 0.53 | 0.994 |
| cluster3936 | 0.69 | 0.994 |
| cluster3331 | -0.41 | 0.994 |
| cluster3203 | -0.46 | 0.994 |
| cluster3980 | 0.50 | 0.994 |
| cluster2909 | 0.48 | 0.994 |
| cluster4089 | 0.80 | 0.994 |
| cluster3279 | 0.59 | 0.994 |
| cluster3963 | -0.59 | 0.994 |
| cluster4022 | -0.45 | 0.994 |
| cluster3456 | 0.55 | 0.994 |
| cluster3270 | 0.68 | 0.994 |
| cluster3637 | 0.48 | 0.994 |
| cluster2855 | 0.51 | 0.994 |
| cluster3511 | -0.51 | 0.994 |
| cluster3025 | 0.47 | 0.994 |
| cluster3665 | -0.57 | 0.994 |
| cluster3140 | 0.49 | 0.994 |
| cluster3372 | 0.50 | 0.994 |
| cluster2871 | -0.41 | 0.994 |
| cluster3103 | -0.56 | 0.994 |
| cluster3841 | -0.50 | 0.994 |
| cluster3241 | 0.52 | 0.994 |
| cluster3801 | 0.73 | 0.994 |
| cluster3298 | -0.43 | 0.994 |
| cluster3879 | 0.56 | 0.994 |
| cluster2979 | 0.48 | 0.994 |
| cluster3674 | -0.67 | 0.994 |

| | | |
|---|---|---|
| cluster3010 | -0.45 | 0.994 |
| cluster3550 | 0.58 | 0.994 |
| cluster3170 | 0.54 | 0.994 |
| cluster3860 | 0.60 | 0.994 |
| cluster3120 | 0.44 | 0.994 |
| cluster3516 | -0.56 | 0.994 |
| cluster4007 | 0.51 | 0.994 |
| cluster3445 | -0.40 | 0.994 |
| cluster2966 | 0.59 | 0.994 |
| cluster3791 | 0.60 | 0.994 |
| cluster3815 | 0.77 | 0.994 |
| cluster3144 | -0.57 | 0.994 |
| cluster3891 | 0.61 | 0.994 |
| cluster3488 | -0.40 | 0.994 |
| cluster3024 | 0.59 | 0.994 |
| cluster3698 | 0.51 | 0.994 |
| cluster3262 | 0.51 | 0.994 |
| cluster3022 | -0.47 | 0.994 |
| cluster2937 | -0.42 | 0.994 |
| cluster3553 | -0.47 | 0.994 |
| cluster3199 | 0.44 | 0.994 |
| cluster3234 | -0.42 | 0.994 |
| cluster3421 | 0.64 | 0.994 |
| cluster3286 | 0.45 | 0.994 |
| cluster4024 | 0.57 | 0.994 |
| cluster4039 | 0.60 | 0.994 |
| cluster3739 | 0.46 | 0.994 |
| cluster3531 | -0.43 | 0.994 |
| cluster3576 | 0.57 | 0.994 |
| cluster3362 | 0.52 | 0.994 |
| cluster3256 | 0.48 | 0.994 |
| cluster3584 | 0.71 | 0.994 |
| cluster3190 | 0.76 | 0.994 |
| cluster2922 | 0.56 | 0.994 |
| cluster3544 | -0.56 | 0.994 |
| cluster2939 | -0.38 | 0.994 |
| cluster3911 | 0.72 | 0.994 |
| cluster3109 | 0.54 | 0.994 |
| cluster3127 | -0.37 | 0.994 |
| cluster3506 | 0.45 | 0.994 |
| cluster2976 | -0.39 | 0.994 |
| cluster3881 | 0.65 | 0.994 |
| cluster3559 | 0.48 | 0.994 |
| cluster2858 | 0.44 | 0.994 |

| | | |
|---|---|---|
| cluster4088 | -0.50 | 0.994 |
| cluster3774 | 0.49 | 0.994 |
| cluster3345 | -0.41 | 0.994 |
| cluster3111 | -0.40 | 0.994 |
| cluster2900 | 0.56 | 0.994 |
| cluster3968 | 0.61 | 0.994 |
| cluster3202 | 0.46 | 0.994 |
| cluster3374 | 0.63 | 0.994 |
| cluster4065 | 0.49 | 0.994 |
| cluster3702 | 0.53 | 0.994 |
| cluster3777 | -0.39 | 0.994 |
| cluster3002 | 0.44 | 0.994 |
| cluster3007 | 0.47 | 0.994 |
| cluster3594 | -0.43 | 0.994 |
| cluster3057 | -0.43 | 0.994 |
| cluster3360 | -0.40 | 0.994 |
| cluster3475 | 0.50 | 0.994 |
| cluster3227 | 0.49 | 0.994 |
| cluster3699 | 0.63 | 0.994 |
| cluster3962 | 0.45 | 0.994 |
| cluster2889 | 0.44 | 0.994 |
| cluster3444 | 0.43 | 0.994 |
| cluster3635 | 0.42 | 0.994 |
| cluster4110 | 0.64 | 0.994 |
| cluster3330 | 0.48 | 0.994 |
| cluster3277 | -0.43 | 0.994 |
| cluster3608 | -0.34 | 0.994 |
| cluster3998 | 0.68 | 0.994 |
| cluster3479 | -0.46 | 0.994 |
| cluster3265 | 0.46 | 0.994 |
| cluster3736 | -0.45 | 0.994 |
| cluster3160 | -0.36 | 0.994 |
| cluster3031 | 0.45 | 0.994 |
| cluster3381 | 0.48 | 0.994 |
| cluster4028 | 0.49 | 0.994 |
| cluster3047 | -0.40 | 0.994 |
| cluster3169 | 0.50 | 0.994 |
| cluster2874 | 0.39 | 0.994 |
| cluster3148 | 0.39 | 0.994 |
| cluster3647 | 0.47 | 0.994 |
| cluster3078 | 0.39 | 0.994 |
| cluster3407 | 0.45 | 0.994 |
| cluster3209 | -0.35 | 0.994 |
| cluster4045 | 0.67 | 0.994 |

| | | |
|---|---|---|
| cluster2988 | 0.40 | 0.994 |
| cluster2903 | -0.40 | 0.994 |
| cluster3379 | -0.36 | 0.994 |
| cluster3813 | 0.59 | 0.994 |
| cluster3669 | -0.34 | 0.994 |
| cluster3134 | -0.33 | 0.994 |
| cluster3498 | 0.52 | 0.994 |
| cluster3503 | 0.45 | 0.994 |
| cluster3806 | -0.40 | 0.994 |
| cluster3068 | -0.32 | 0.994 |
| cluster3967 | 0.50 | 0.994 |
| cluster3814 | 0.46 | 0.994 |
| cluster4083 | 0.61 | 0.994 |
| cluster3036 | -0.35 | 0.994 |
| cluster3236 | 0.46 | 0.994 |
| cluster3976 | -0.34 | 0.994 |
| cluster3283 | 0.41 | 0.994 |
| cluster2992 | -0.34 | 0.994 |
| cluster2970 | 0.36 | 0.994 |
| cluster3569 | 0.52 | 0.994 |
| cluster3087 | 0.41 | 0.994 |
| cluster3035 | 0.40 | 0.994 |
| cluster4100 | 0.59 | 0.994 |
| cluster3664 | 0.43 | 0.994 |
| cluster4021 | 0.56 | 0.994 |
| cluster3433 | 0.46 | 0.994 |
| cluster3252 | 0.67 | 0.994 |
| cluster2857 | -0.32 | 0.994 |
| cluster3886 | 0.45 | 0.994 |
| cluster3347 | 0.38 | 0.994 |
| cluster3385 | -0.39 | 0.994 |
| cluster3282 | -0.31 | 0.994 |
| cluster3268 | -0.30 | 0.994 |
| cluster3133 | -0.32 | 0.994 |
| cluster4008 | 0.56 | 0.994 |
| cluster4084 | -0.34 | 0.994 |
| cluster3466 | -0.34 | 0.994 |
| cluster3093 | -0.35 | 0.994 |
| cluster3079 | 0.42 | 0.994 |
| cluster3411 | -0.29 | 0.994 |
| cluster3673 | -0.43 | 0.994 |
| cluster2973 | 0.45 | 0.994 |
| cluster2983 | 0.37 | 0.994 |
| cluster3476 | 0.56 | 0.994 |

| | | |
|---|---|---|
| cluster3418 | 0.45 | 0.994 |
| cluster3205 | -0.32 | 0.994 |
| cluster3352 | 0.47 | 0.994 |
| cluster3812 | 0.51 | 0.994 |
| cluster3604 | 0.56 | 0.994 |
| cluster3601 | 0.45 | 0.994 |
| cluster2929 | 0.35 | 0.994 |
| cluster3924 | 0.50 | 0.994 |
| cluster3299 | -0.35 | 0.994 |
| cluster3596 | -0.34 | 0.994 |
| cluster3383 | -0.42 | 0.994 |
| cluster3354 | 0.37 | 0.994 |
| cluster3906 | -0.33 | 0.994 |
| cluster3947 | 0.40 | 0.994 |
| cluster3489 | 0.43 | 0.994 |
| cluster2867 | -0.38 | 0.994 |
| cluster3868 | -0.28 | 0.994 |
| cluster3386 | -0.33 | 0.994 |
| cluster3155 | 0.48 | 0.994 |
| cluster3697 | 0.51 | 0.994 |
| cluster3566 | -0.29 | 0.994 |
| cluster3972 | -0.30 | 0.994 |
| cluster3917 | 0.48 | 0.994 |
| cluster3685 | -0.33 | 0.994 |
| cluster3620 | -0.32 | 0.994 |
| cluster2990 | 0.36 | 0.994 |
| cluster3429 | 0.39 | 0.994 |
| cluster3648 | 0.36 | 0.994 |
| cluster3254 | -0.35 | 0.994 |
| cluster3957 | 0.44 | 0.994 |
| cluster4082 | 0.57 | 0.994 |
| cluster4031 | 0.42 | 0.994 |
| cluster4025 | 0.34 | 0.994 |
| cluster3710 | 0.44 | 0.994 |
| cluster3675 | -0.27 | 0.994 |
| cluster3124 | -0.25 | 0.994 |
| cluster3208 | 0.35 | 0.994 |
| cluster3048 | -0.27 | 0.994 |
| cluster1127 | 0.42 | 0.994 |
| cluster3014 | -0.31 | 0.994 |
| cluster3893 | 0.34 | 0.994 |
| cluster3950 | 0.37 | 0.994 |
| cluster3221 | 0.42 | 0.994 |
| cluster3670 | -0.33 | 0.994 |

| | | |
|---|---|---|
| cluster3080 | -0.28 | 0.994 |
| cluster3820 | -0.26 | 0.994 |
| cluster3404 | 0.35 | 0.994 |
| cluster3627 | -0.27 | 0.994 |
| cluster3171 | 0.41 | 0.994 |
| cluster4063 | -0.38 | 0.994 |
| cluster3630 | 0.33 | 0.994 |
| cluster3012 | 0.48 | 0.994 |
| cluster3557 | -0.28 | 0.994 |
| cluster3923 | -0.27 | 0.994 |
| cluster4032 | 0.39 | 0.994 |
| cluster3993 | -0.27 | 0.994 |
| cluster2934 | -0.25 | 0.994 |
| cluster3284 | 0.35 | 0.994 |
| cluster3359 | -0.30 | 0.994 |
| cluster3122 | -0.27 | 0.994 |
| cluster3545 | 0.38 | 0.994 |
| cluster3231 | -0.30 | 0.994 |
| cluster2925 | 0.39 | 0.994 |
| cluster2931 | -0.27 | 0.994 |
| cluster3316 | -0.30 | 0.994 |
| cluster3623 | -0.26 | 0.994 |
| cluster3642 | 0.42 | 0.994 |
| cluster2880 | 0.35 | 0.994 |
| cluster4061 | 0.45 | 0.994 |
| cluster3343 | 0.32 | 0.994 |
| cluster3785 | -0.29 | 0.994 |
| cluster4029 | -0.34 | 0.994 |
| cluster3796 | -0.32 | 0.994 |
| cluster3655 | -0.27 | 0.994 |
| cluster3946 | -0.29 | 0.994 |
| cluster3521 | -0.25 | 0.994 |
| cluster3974 | 0.57 | 0.994 |
| cluster3732 | -0.25 | 0.994 |
| cluster2866 | 0.39 | 0.994 |
| cluster3307 | -0.24 | 0.994 |
| cluster3073 | -0.22 | 0.994 |
| cluster3424 | -0.34 | 0.994 |
| cluster3384 | 0.36 | 0.994 |
| cluster2886 | 0.34 | 0.994 |
| cluster3683 | 0.34 | 0.994 |
| cluster3403 | -0.22 | 0.994 |
| cluster3249 | -0.23 | 0.994 |
| cluster3543 | -0.30 | 0.994 |

| | | |
|---|---|---|
| cluster2932 | -0.24 | 0.994 |
| cluster3877 | -0.23 | 0.994 |
| cluster2945 | -0.31 | 0.994 |
| cluster3663 | -0.26 | 0.994 |
| cluster3339 | 0.33 | 0.994 |
| cluster3440 | 0.28 | 0.994 |
| cluster3845 | -0.36 | 0.994 |
| cluster3278 | 0.28 | 0.994 |
| cluster3944 | 0.34 | 0.994 |
| cluster2959 | 0.30 | 0.994 |
| cluster3275 | -0.28 | 0.994 |
| cluster2888 | -0.31 | 0.994 |
| cluster4018 | 0.42 | 0.994 |
| cluster3046 | 0.31 | 0.994 |
| cluster4043 | -0.30 | 0.994 |
| cluster3069 | 0.28 | 0.994 |
| cluster3607 | 0.36 | 0.994 |
| cluster3880 | 0.54 | 0.994 |
| cluster3591 | -0.21 | 0.994 |
| cluster2875 | 0.33 | 0.994 |
| cluster2872 | 0.28 | 0.994 |
| cluster3754 | 0.31 | 0.994 |
| cluster4030 | -0.24 | 0.994 |
| cluster3848 | -0.23 | 0.994 |
| cluster3274 | 0.36 | 0.994 |
| cluster3965 | -0.26 | 0.994 |
| cluster3259 | -0.20 | 0.994 |
| cluster3914 | -0.30 | 0.994 |
| cluster3776 | -0.25 | 0.994 |
| cluster3688 | 0.29 | 0.994 |
| cluster3326 | 0.32 | 0.994 |
| cluster3216 | 0.33 | 0.994 |
| cluster3939 | 0.32 | 0.994 |
| cluster3706 | -0.19 | 0.994 |
| cluster4010 | 0.40 | 0.994 |
| cluster4058 | -0.20 | 0.994 |
| cluster3843 | 0.27 | 0.994 |
| cluster3423 | 0.42 | 0.994 |
| cluster3387 | -0.20 | 0.994 |
| cluster3033 | -0.18 | 0.994 |
| cluster3192 | 0.27 | 0.994 |
| cluster3116 | 0.25 | 0.994 |
| cluster2985 | 0.29 | 0.994 |
| cluster3253 | -0.21 | 0.994 |

| | | |
|---|---|---|
| cluster3913 | -0.23 | 0.994 |
| cluster3751 | -0.20 | 0.994 |
| cluster3793 | 0.27 | 0.994 |
| cluster3680 | 0.27 | 0.994 |
| cluster3294 | 0.27 | 0.994 |
| cluster3145 | -0.19 | 0.994 |
| cluster3808 | 0.37 | 0.994 |
| cluster3201 | 0.26 | 0.994 |
| cluster3780 | -0.27 | 0.994 |
| cluster2955 | 0.24 | 0.994 |
| cluster3211 | 0.37 | 0.994 |
| cluster3671 | -0.19 | 0.994 |
| cluster3230 | 0.31 | 0.994 |
| cluster3415 | -0.19 | 0.994 |
| cluster2895 | -0.18 | 0.994 |
| cluster3654 | -0.18 | 0.994 |
| cluster2968 | 0.23 | 0.994 |
| cluster3718 | -0.22 | 0.994 |
| cluster4036 | 0.27 | 0.994 |
| cluster3527 | -0.21 | 0.994 |
| cluster3583 | 0.23 | 0.994 |
| cluster3502 | 0.38 | 0.994 |
| cluster3042 | -0.20 | 0.994 |
| cluster2887 | 0.27 | 0.994 |
| cluster3305 | -0.20 | 0.994 |
| cluster3626 | 0.27 | 0.994 |
| cluster3028 | 0.24 | 0.994 |
| cluster2910 | -0.16 | 0.994 |
| cluster2869 | 0.32 | 0.994 |
| cluster3921 | 0.38 | 0.994 |
| cluster3562 | -0.19 | 0.994 |
| cluster3533 | 0.28 | 0.994 |
| cluster3266 | 0.25 | 0.994 |
| cluster3711 | -0.16 | 0.994 |
| cluster3560 | 0.35 | 0.994 |
| cluster3926 | -0.22 | 0.994 |
| cluster3888 | -0.20 | 0.994 |
| cluster3590 | 0.31 | 0.994 |
| cluster2997 | -0.17 | 0.994 |
| cluster3114 | -0.15 | 0.994 |
| cluster3310 | -0.15 | 0.994 |
| cluster3588 | 0.29 | 0.994 |
| cluster3866 | 0.38 | 0.994 |
| cluster3761 | 0.34 | 0.994 |

| | | |
|---|---|---|
| cluster3336 | -0.19 | 0.994 |
| cluster3969 | 0.21 | 0.994 |
| cluster3823 | -0.17 | 0.994 |
| cluster3775 | -0.16 | 0.994 |
| cluster3167 | -0.20 | 0.994 |
| cluster3961 | 0.34 | 0.994 |
| cluster3163 | -0.19 | 0.994 |
| cluster3726 | 0.24 | 0.994 |
| cluster3949 | -0.19 | 0.994 |
| cluster3660 | -0.20 | 0.994 |
| cluster2883 | 0.23 | 0.994 |
| cluster3324 | 0.24 | 0.994 |
| cluster3681 | 0.28 | 0.994 |
| cluster3693 | -0.20 | 0.994 |
| cluster2913 | -0.14 | 0.994 |
| cluster3428 | 0.23 | 0.994 |
| cluster4092 | -0.20 | 0.994 |
| cluster3464 | 0.28 | 0.994 |
| cluster2951 | 0.28 | 0.994 |
| cluster4117 | -0.24 | 0.994 |
| cluster3700 | 0.27 | 0.994 |
| cluster3072 | 0.27 | 0.994 |
| cluster3504 | 0.32 | 0.994 |
| cluster3206 | -0.17 | 0.994 |
| cluster3123 | 0.23 | 0.994 |
| cluster3281 | 0.24 | 0.994 |
| cluster3416 | 0.35 | 0.994 |
| cluster4014 | 0.27 | 0.994 |
| cluster3186 | -0.16 | 0.994 |
| cluster3790 | -0.17 | 0.994 |
| cluster3722 | 0.29 | 0.994 |
| cluster3856 | -0.14 | 0.994 |
| cluster3183 | -0.15 | 0.994 |
| cluster3223 | -0.16 | 0.994 |
| cluster3727 | -0.17 | 0.994 |
| cluster3622 | 0.20 | 0.994 |
| cluster3355 | -0.16 | 0.994 |
| cluster3173 | 0.31 | 0.994 |
| cluster3090 | 0.35 | 0.994 |
| cluster3125 | -0.13 | 0.994 |
| cluster2944 | -0.14 | 0.994 |
| cluster2856 | 0.26 | 0.994 |
| cluster3285 | -0.14 | 0.994 |
| cluster3084 | -0.12 | 0.994 |

| | | |
|---|---|---|
| cluster3302 | -0.13 | 0.994 |
| cluster3579 | 0.20 | 0.994 |
| cluster3333 | 0.22 | 0.994 |
| cluster3539 | -0.16 | 0.994 |
| cluster3112 | -0.15 | 0.994 |
| cluster3136 | 0.24 | 0.994 |
| cluster3528 | 0.20 | 0.994 |
| cluster3809 | 0.20 | 0.994 |
| cluster3378 | 0.24 | 0.994 |
| cluster3058 | 0.17 | 0.994 |
| cluster3690 | -0.12 | 0.994 |
| cluster2954 | 0.23 | 0.994 |
| cluster3272 | -0.16 | 0.994 |
| cluster3686 | 0.30 | 0.994 |
| cluster3910 | -0.14 | 0.994 |
| cluster3050 | -0.11 | 0.994 |
| cluster3908 | -0.12 | 0.994 |
| cluster3001 | -0.11 | 0.994 |
| cluster4020 | 0.19 | 0.994 |
| cluster3422 | 0.31 | 0.994 |
| cluster3986 | 0.20 | 0.994 |
| cluster3301 | -0.12 | 0.994 |
| cluster3328 | 0.17 | 0.994 |
| cluster3798 | -0.12 | 0.994 |
| cluster2896 | 0.16 | 0.994 |
| cluster3767 | -0.12 | 0.994 |
| cluster3587 | 0.29 | 0.994 |
| cluster4068 | -0.13 | 0.994 |
| cluster3108 | 0.21 | 0.994 |
| cluster3040 | -0.11 | 0.994 |
| cluster3614 | 0.30 | 0.994 |
| cluster3370 | 0.21 | 0.994 |
| cluster3740 | -0.11 | 0.994 |
| cluster3364 | -0.10 | 0.994 |
| cluster3639 | 0.31 | 0.994 |
| cluster2974 | -0.10 | 0.994 |
| cluster4104 | 0.26 | 0.994 |
| cluster3507 | -0.10 | 0.994 |
| cluster3390 | -0.10 | 0.994 |
| cluster4041 | -0.10 | 0.994 |
| cluster3018 | 0.15 | 0.994 |
| cluster3074 | -0.09 | 0.994 |
| cluster3059 | -0.10 | 0.994 |
| cluster3803 | 0.26 | 0.994 |

| | | |
|---|---|---|
| cluster3725 | 0.24 | 0.994 |
| cluster2953 | 0.15 | 0.994 |
| cluster4069 | 0.20 | 0.994 |
| cluster2958 | 0.14 | 0.994 |
| cluster3835 | -0.10 | 0.994 |
| cluster4017 | 0.24 | 0.994 |
| cluster4050 | -0.09 | 0.994 |
| cluster3151 | -0.08 | 0.994 |
| cluster3874 | -0.09 | 0.994 |
| cluster3927 | 0.19 | 0.994 |
| cluster3524 | 0.15 | 0.994 |
| cluster3900 | 0.30 | 0.994 |
| cluster3420 | 0.17 | 0.994 |
| cluster3831 | -0.09 | 0.994 |
| cluster3351 | 0.13 | 0.994 |
| cluster3096 | 0.14 | 0.994 |
| cluster3455 | -0.09 | 0.994 |
| cluster3641 | 0.16 | 0.994 |
| cluster3616 | 0.22 | 0.994 |
| cluster3154 | 0.15 | 0.994 |
| cluster3612 | 0.22 | 0.994 |
| cluster2907 | -0.07 | 0.994 |
| cluster3870 | -0.08 | 0.994 |
| cluster3836 | -0.08 | 0.994 |
| cluster3708 | -0.10 | 0.994 |
| cluster3828 | 0.20 | 0.994 |
| cluster3707 | 0.23 | 0.994 |
| cluster3308 | -0.07 | 0.994 |
| cluster4038 | 0.23 | 0.994 |
| cluster3477 | 0.18 | 0.994 |
| cluster3158 | 0.20 | 0.994 |
| cluster3951 | 0.22 | 0.994 |
| cluster3606 | -0.08 | 0.994 |
| cluster2986 | 0.15 | 0.994 |
| cluster3912 | -0.06 | 0.994 |
| cluster3625 | 0.15 | 0.994 |
| cluster3938 | -0.07 | 0.994 |
| cluster3139 | 0.26 | 0.994 |
| cluster3797 | -0.07 | 0.994 |
| cluster4003 | 0.22 | 0.994 |
| cluster3344 | -0.08 | 0.994 |
| cluster3991 | -0.08 | 0.994 |
| cluster3044 | 0.19 | 0.994 |
| cluster3467 | 0.14 | 0.994 |

| | | |
|---|---|---|
| cluster3152 | 0.21 | 0.994 |
| cluster3728 | 0.15 | 0.994 |
| cluster3264 | -0.06 | 0.994 |
| cluster3098 | -0.06 | 0.994 |
| cluster3692 | -0.07 | 0.994 |
| cluster2952 | -0.06 | 0.994 |
| cluster2918 | 0.14 | 0.994 |
| cluster3091 | -0.06 | 0.994 |
| cluster3020 | -0.06 | 0.994 |
| cluster3855 | -0.06 | 0.994 |
| cluster4027 | 0.21 | 0.994 |
| cluster3720 | -0.06 | 0.994 |
| cluster3759 | -0.07 | 0.994 |
| cluster2859 | -0.05 | 0.994 |
| cluster3041 | 0.15 | 0.994 |
| cluster3517 | 0.11 | 0.994 |
| cluster3037 | -0.05 | 0.994 |
| cluster2893 | 0.19 | 0.994 |
| cluster3782 | -0.07 | 0.994 |
| cluster3251 | -0.05 | 0.994 |
| cluster3618 | -0.05 | 0.994 |
| cluster3499 | -0.05 | 0.994 |
| cluster3789 | -0.06 | 0.994 |
| cluster3470 | -0.05 | 0.994 |
| cluster3492 | 0.13 | 0.994 |
| cluster3054 | 0.11 | 0.994 |
| cluster3447 | -0.05 | 0.994 |
| cluster3029 | 0.11 | 0.994 |
| cluster3717 | -0.04 | 0.994 |
| cluster3523 | -0.04 | 0.994 |
| cluster3816 | -0.04 | 0.994 |
| cluster3696 | 0.11 | 0.994 |
| cluster3846 | 0.12 | 0.994 |
| cluster4096 | -0.04 | 0.994 |
| cluster3368 | -0.04 | 0.994 |
| cluster3837 | -0.04 | 0.994 |
| cluster3172 | -0.04 | 0.994 |
| cluster3217 | -0.04 | 0.994 |
| cluster2917 | 0.15 | 0.994 |
| cluster2920 | -0.04 | 0.994 |
| cluster3092 | 0.10 | 0.994 |
| cluster3491 | -0.04 | 0.994 |
| cluster3807 | 0.12 | 0.994 |
| cluster2885 | -0.03 | 0.994 |

| | | |
|---|---|---|
| cluster3701 | 0.11 | 0.994 |
| cluster3935 | 0.16 | 0.994 |
| cluster3558 | 0.14 | 0.994 |
| cluster3535 | 0.17 | 0.994 |
| cluster3570 | 0.17 | 0.994 |
| cluster3909 | -0.03 | 0.994 |
| cluster3628 | 0.13 | 0.994 |
| cluster3882 | 0.16 | 0.994 |
| cluster3945 | -0.02 | 0.994 |
| cluster3529 | -0.02 | 0.994 |
| cluster3617 | 0.16 | 0.994 |
| cluster2991 | -0.02 | 0.994 |
| cluster3389 | 0.11 | 0.994 |
| cluster3838 | 0.14 | 0.994 |
| cluster3903 | -0.02 | 0.994 |
| cluster2938 | -0.02 | 0.994 |
| cluster3406 | -0.02 | 0.994 |
| cluster3412 | -0.02 | 0.994 |
| cluster4102 | -0.03 | 0.994 |
| cluster3555 | 0.00 | 0.994 |
| cluster3668 | 0.09 | 0.994 |
| cluster3941 | -0.02 | 0.994 |
| cluster3314 | 0.13 | 0.994 |
| cluster3248 | -0.01 | 0.994 |
| cluster2971 | -0.02 | 0.994 |
| cluster3988 | -0.02 | 0.994 |
| cluster3895 | 0.15 | 0.994 |
| cluster4040 | 0.12 | 0.994 |
| cluster3605 | 0.11 | 0.994 |
| cluster2862 | 0.09 | 0.994 |
| cluster3214 | 0.10 | 0.994 |
| cluster3646 | 0.16 | 0.994 |
| cluster3896 | 0.13 | 0.994 |
| cluster3592 | 0.09 | 0.994 |
| cluster3851 | 0.13 | 0.994 |
| cluster3733 | 0.00 | 0.994 |
| cluster2890 | -0.01 | 0.994 |
| cluster3104 | -0.01 | 0.994 |
| cluster3003 | 0.10 | 0.994 |
| cluster3483 | 0.08 | 0.994 |
| cluster3350 | 0.00 | 0.995 |
| cluster3056 | -0.01 | 0.995 |
| cluster3864 | 0.09 | 0.995 |
| cluster3222 | 0.07 | 0.995 |

| | | |
|---|---|---|
| cluster3189 | 0.11 | 0.996 |
| cluster3743 | 0.00 | 0.996 |
| cluster3196 | 0.07 | 0.996 |
| cluster3198 | 0.00 | 0.996 |
| cluster4123 | 0.12 | 0.997 |
| cluster2897 | 0.00 | 0.997 |
| cluster3184 | 0.07 | 0.997 |
| cluster3060 | 0.02 | 0.997 |
| cluster4004 | 0.00 | 0.997 |
| cluster3193 | 0.07 | 0.997 |
| cluster3271 | 0.00 | 0.997 |
| cluster3219 | 0.00 | 0.997 |
| cluster4006 | 0.01 | 0.997 |
| cluster3512 | 0.09 | 0.999 |
| cluster3589 | 0.07 | 0.999 |
| cluster3610 | 0.09 | 1.000 |
| cluster2941 | 0.05 | 1.000 |
| cluster3394 | 0.02 | 1.000 |
| cluster2996 | 0.05 | 1.000 |
| cluster3382 | 0.05 | 1.000 |
| cluster3419 | 0.05 | 1.000 |
| cluster3086 | 0.05 | 1.000 |
| cluster3213 | 0.03 | 1.000 |
| cluster3982 | 0.05 | 1.000 |
| cluster3409 | 0.08 | 1.000 |
| cluster3146 | 0.03 | 1.000 |
| cluster3859 | 0.03 | 1.000 |
| cluster3861 | 0.05 | 1.000 |
| cluster3038 | 0.06 | 1.000 |
| cluster3532 | 0.06 | 1.000 |
| cluster3322 | 0.07 | 1.000 |
| cluster3542 | 0.08 | 1.000 |
| cluster3376 | 0.03 | 1.000 |
| cluster4071 | 0.08 | 1.000 |
| cluster3783 | 0.06 | 1.000 |
| cluster3833 | 0.06 | 1.000 |
| cluster4077 | 0.05 | 1.000 |
| cluster3687 | 0.05 | 1.000 |
| cluster4108 | 0.04 | 1.000 |
| cluster3043 | 0.02 | 1.000 |
| cluster4057 | 0.01 | 1.000 |
| cluster3159 | 0.01 | 1.000 |
| cluster3684 | 0.00 | 1.000 |
| cluster4126 | 0.00 | 1.000 |

| | | |
|---|---|---|
| cluster3396 | 0.00 | 1.000 |

**Supplementary Table 9.** DESeq2 results with RSEM counts - Oestridae vs Calliphoridae

| ID | log2FoldChange | Adjusted p value |
|---|---|---|
| cluster1127 | 0.53 | 0.999 |
| cluster2854 | 1.30 | 0.999 |
| cluster2855 | 0.30 | 0.999 |
| cluster2856 | 0.18 | 0.999 |
| cluster2857 | -0.80 | 0.999 |
| cluster2858 | 0.15 | 0.999 |
| cluster2859 | -0.38 | 0.999 |
| cluster2860 | 0.88 | 0.999 |
| cluster2861 | 1.46 | 0.999 |
| cluster2862 | -0.31 | 0.999 |
| cluster2863 | 0.59 | 0.999 |
| cluster2864 | 0.30 | 0.999 |
| cluster2865 | 0.50 | 0.999 |
| cluster2866 | 0.62 | 0.999 |
| cluster2867 | -0.35 | 0.999 |
| cluster2869 | -0.08 | 0.999 |
| cluster2870 | 0.67 | 0.999 |
| cluster2871 | -0.70 | 0.999 |
| cluster2872 | 0.16 | 0.999 |
| cluster2873 | 0.63 | 0.999 |
| cluster2874 | 0.15 | 0.999 |
| cluster2875 | -0.04 | 0.999 |
| cluster2876 | 0.81 | 0.999 |
| cluster2877 | 0.53 | 0.999 |
| cluster2878 | 0.44 | 0.999 |
| cluster2879 | 0.42 | 0.999 |
| cluster2880 | 0.38 | 0.999 |
| cluster2881 | 1.16 | 0.999 |
| cluster2882 | 0.70 | 0.999 |
| cluster2883 | -0.19 | 0.999 |
| cluster2884 | 0.61 | 0.999 |
| cluster2885 | 0.08 | 0.999 |
| cluster2886 | -0.01 | 0.999 |
| cluster2887 | 0.07 | 0.999 |
| cluster2888 | -0.60 | 0.999 |
| cluster2889 | 0.32 | 0.999 |
| cluster2890 | -0.28 | 0.999 |
| cluster2891 | -0.47 | 0.999 |
| cluster2892 | 0.76 | 0.999 |
| cluster2893 | -0.12 | 0.999 |
| cluster2894 | -0.72 | 0.999 |
| cluster2895 | -0.08 | 0.999 |

| | | |
|---|---|---|
| cluster2896 | -0.09 | 0.999 |
| cluster2897 | 0.09 | 0.999 |
| cluster2898 | 0.61 | 0.999 |
| cluster2899 | 0.97 | 0.999 |
| cluster2900 | -0.09 | 0.999 |
| cluster2901 | -0.78 | 0.999 |
| cluster2902 | 2.27 | 0.999 |
| cluster2904 | 0.30 | 0.999 |
| cluster2905 | 0.29 | 0.999 |
| cluster2906 | 0.57 | 0.999 |
| cluster2907 | 0.00 | 0.999 |
| cluster2908 | 0.62 | 0.999 |
| cluster2909 | 0.05 | 0.999 |
| cluster2910 | -0.17 | 0.999 |
| cluster2911 | -1.05 | 0.999 |
| cluster2912 | 1.02 | 0.999 |
| cluster2913 | -0.18 | 0.999 |
| cluster2914 | -0.93 | 0.999 |
| cluster2915 | -0.67 | 0.999 |
| cluster2916 | 0.60 | 0.999 |
| cluster2917 | 0.12 | 0.999 |
| cluster2918 | 0.32 | 0.999 |
| cluster2919 | 0.47 | 0.999 |
| cluster2920 | 0.01 | 0.999 |
| cluster2921 | 1.63 | 0.999 |
| cluster2922 | 0.51 | 0.999 |
| cluster2923 | 0.26 | 0.999 |
| cluster2924 | 1.17 | 0.999 |
| cluster2925 | 0.42 | 0.999 |
| cluster2926 | 0.69 | 0.999 |
| cluster2927 | 0.65 | 0.999 |
| cluster2928 | 0.66 | 0.999 |
| cluster2929 | 0.20 | 0.999 |
| cluster2930 | 1.24 | 0.999 |
| cluster2931 | -0.21 | 0.999 |
| cluster2932 | -0.20 | 0.999 |
| cluster2933 | 0.96 | 0.999 |
| cluster2934 | -0.40 | 0.999 |
| cluster2935 | 0.28 | 0.999 |
| cluster2936 | 0.75 | 0.999 |
| cluster2937 | -0.67 | 0.999 |
| cluster2938 | -0.46 | 0.999 |
| cluster2939 | -0.75 | 0.999 |
| cluster2940 | -0.54 | 0.999 |

| | | |
|---|---|---|
| cluster2941 | -0.35 | 0.999 |
| cluster2942 | 1.02 | 0.999 |
| cluster2943 | 1.39 | 0.999 |
| cluster2944 | -0.05 | 0.999 |
| cluster2945 | -0.21 | 0.999 |
| cluster2946 | 0.76 | 0.999 |
| cluster2947 | 1.30 | 0.999 |
| cluster2948 | 1.73 | 0.999 |
| cluster2949 | 0.38 | 0.999 |
| cluster2950 | 0.26 | 0.999 |
| cluster2951 | 0.16 | 0.999 |
| cluster2952 | -0.05 | 0.999 |
| cluster2953 | -0.03 | 0.999 |
| cluster2954 | -0.27 | 0.999 |
| cluster2955 | 0.03 | 0.999 |
| cluster2956 | -0.69 | 0.999 |
| cluster2957 | -0.57 | 0.999 |
| cluster2958 | -0.35 | 0.999 |
| cluster2959 | 0.07 | 0.999 |
| cluster2961 | 1.11 | 0.999 |
| cluster2962 | 1.13 | 0.999 |
| cluster2963 | 0.60 | 0.999 |
| cluster2964 | 0.49 | 0.999 |
| cluster2965 | 0.89 | 0.999 |
| cluster2966 | 0.47 | 0.999 |
| cluster2967 | 1.46 | 0.999 |
| cluster2968 | -0.11 | 0.999 |
| cluster2969 | 1.36 | 0.999 |
| cluster2970 | 0.05 | 0.999 |
| cluster2971 | -0.46 | 0.999 |
| cluster2972 | -1.32 | 0.999 |
| cluster2973 | 0.37 | 0.999 |
| cluster2974 | -1.07 | 0.999 |
| cluster2975 | 0.50 | 0.999 |
| cluster2976 | -0.70 | 0.999 |
| cluster2977 | 0.97 | 0.999 |
| cluster2979 | 0.41 | 0.999 |
| cluster2981 | 0.70 | 0.999 |
| cluster2982 | 0.57 | 0.999 |
| cluster2983 | 0.20 | 0.999 |
| cluster2985 | 0.07 | 0.999 |
| cluster2986 | -0.01 | 0.999 |
| cluster2987 | 0.36 | 0.999 |
| cluster2988 | 0.24 | 0.999 |

| | | |
|---|---|---|
| cluster2989 | 1.12 | 0.999 |
| cluster2990 | 0.04 | 0.999 |
| cluster2991 | -0.22 | 0.999 |
| cluster2992 | -0.57 | 0.999 |
| cluster2993 | 1.37 | 0.999 |
| cluster2994 | 0.77 | 0.999 |
| cluster2995 | 1.13 | 0.999 |
| cluster2996 | -0.08 | 0.999 |
| cluster2997 | -0.19 | 0.999 |
| cluster2998 | -1.11 | 0.999 |
| cluster2999 | -0.91 | 0.999 |
| cluster3000 | -1.85 | 0.999 |
| cluster3001 | -0.43 | 0.999 |
| cluster3002 | 0.23 | 0.999 |
| cluster3003 | 0.26 | 0.999 |
| cluster3004 | 0.34 | 0.999 |
| cluster3005 | 0.66 | 0.999 |
| cluster3006 | 2.42 | 0.999 |
| cluster3007 | 0.38 | 0.999 |
| cluster3008 | 1.17 | 0.999 |
| cluster3009 | -1.35 | 0.999 |
| cluster3010 | -0.38 | 0.999 |
| cluster3011 | 0.54 | 0.999 |
| cluster3012 | 0.27 | 0.999 |
| cluster3013 | 0.71 | 0.999 |
| cluster3014 | -0.44 | 0.999 |
| cluster3015 | 1.66 | 0.999 |
| cluster3016 | -0.71 | 0.999 |
| cluster3018 | 0.04 | 0.999 |
| cluster3019 | 1.10 | 0.999 |
| cluster3020 | -0.06 | 0.999 |
| cluster3021 | 0.21 | 0.999 |
| cluster3022 | -0.49 | 0.999 |
| cluster3023 | 0.81 | 0.999 |
| cluster3024 | -0.27 | 0.999 |
| cluster3025 | 0.07 | 0.999 |
| cluster3026 | -1.13 | 0.999 |
| cluster3027 | 0.69 | 0.999 |
| cluster3028 | 0.11 | 0.999 |
| cluster3029 | -0.10 | 0.999 |
| cluster3031 | 0.10 | 0.999 |
| cluster3032 | 0.81 | 0.999 |
| cluster3033 | -0.48 | 0.999 |
| cluster3034 | -0.54 | 0.999 |

| | | |
|---|---|---|
| cluster3035 | 0.07 | 0.999 |
| cluster3036 | -0.33 | 0.999 |
| cluster3037 | -0.15 | 0.999 |
| cluster3038 | -0.02 | 0.999 |
| cluster3039 | 0.27 | 0.999 |
| cluster3040 | -0.65 | 0.999 |
| cluster3041 | 0.08 | 0.999 |
| cluster3042 | -0.27 | 0.999 |
| cluster3043 | -0.48 | 0.999 |
| cluster3044 | 0.28 | 0.999 |
| cluster3045 | 1.55 | 0.999 |
| cluster3046 | 0.30 | 0.999 |
| cluster3047 | -0.95 | 0.999 |
| cluster3048 | -0.45 | 0.999 |
| cluster3049 | 0.28 | 0.999 |
| cluster3050 | -0.21 | 0.999 |
| cluster3051 | -0.81 | 0.999 |
| cluster3053 | 1.30 | 0.999 |
| cluster3054 | -0.15 | 0.999 |
| cluster3055 | -0.04 | 0.999 |
| cluster3056 | -0.10 | 0.999 |
| cluster3057 | -0.94 | 0.999 |
| cluster3058 | -0.20 | 0.999 |
| cluster3059 | 0.10 | 0.999 |
| cluster3060 | -0.20 | 0.999 |
| cluster3061 | 0.78 | 0.999 |
| cluster3062 | -1.03 | 0.999 |
| cluster3063 | 1.56 | 0.999 |
| cluster3064 | 0.70 | 0.999 |
| cluster3065 | 0.29 | 0.999 |
| cluster3066 | 0.80 | 0.999 |
| cluster3067 | 0.10 | 0.999 |
| cluster3068 | -0.67 | 0.999 |
| cluster3069 | -0.16 | 0.999 |
| cluster3070 | 0.41 | 0.999 |
| cluster3071 | 0.32 | 0.999 |
| cluster3072 | 0.37 | 0.999 |
| cluster3073 | -0.85 | 0.999 |
| cluster3074 | -0.20 | 0.999 |
| cluster3075 | -1.35 | 0.999 |
| cluster3076 | 0.66 | 0.999 |
| cluster3077 | 0.28 | 0.999 |
| cluster3078 | 0.21 | 0.999 |
| cluster3079 | 0.03 | 0.999 |

| | | |
|---|---|---|
| cluster3080 | -0.40 | 0.999 |
| cluster3081 | -1.17 | 0.999 |
| cluster3082 | -1.20 | 0.999 |
| cluster3083 | 0.57 | 0.999 |
| cluster3084 | -0.48 | 0.999 |
| cluster3085 | -1.14 | 0.999 |
| cluster3086 | -0.03 | 0.999 |
| cluster3087 | 0.24 | 0.999 |
| cluster3088 | 1.11 | 0.999 |
| cluster3089 | 0.70 | 0.999 |
| cluster3090 | 0.50 | 0.999 |
| cluster3091 | -0.46 | 0.999 |
| cluster3092 | -0.62 | 0.999 |
| cluster3093 | -0.57 | 0.999 |
| cluster3094 | 0.77 | 0.999 |
| cluster3095 | -1.47 | 0.999 |
| cluster3096 | 0.07 | 0.999 |
| cluster3097 | 1.11 | 0.999 |
| cluster3098 | -0.27 | 0.999 |
| cluster3099 | 0.55 | 0.999 |
| cluster3100 | 1.96 | 0.999 |
| cluster3101 | 1.08 | 0.999 |
| cluster3102 | 1.12 | 0.999 |
| cluster3103 | -0.34 | 0.999 |
| cluster3104 | -0.38 | 0.999 |
| cluster3105 | -1.61 | 0.999 |
| cluster3106 | 0.18 | 0.999 |
| cluster3107 | 0.40 | 0.999 |
| cluster3108 | 0.24 | 0.999 |
| cluster3109 | 0.31 | 0.999 |
| cluster3110 | 1.00 | 0.999 |
| cluster3111 | -0.69 | 0.999 |
| cluster3112 | -0.30 | 0.999 |
| cluster3113 | 0.51 | 0.999 |
| cluster3114 | -0.36 | 0.999 |
| cluster3115 | 0.54 | 0.999 |
| cluster3116 | 0.02 | 0.999 |
| cluster3117 | 1.04 | 0.999 |
| cluster3118 | 1.33 | 0.999 |
| cluster3119 | 0.70 | 0.999 |
| cluster3120 | 0.00 | 0.999 |
| cluster3121 | 0.14 | 0.999 |
| cluster3122 | -0.64 | 0.999 |
| cluster3123 | -0.16 | 0.999 |

| | | |
|---|---|---|
| cluster3124 | -0.43 | 0.999 |
| cluster3125 | -0.51 | 0.999 |
| cluster3127 | -0.77 | 0.999 |
| cluster3128 | 0.24 | 0.999 |
| cluster3129 | -1.37 | 0.999 |
| cluster3130 | -1.50 | 0.999 |
| cluster3131 | -0.70 | 0.999 |
| cluster3132 | 0.48 | 0.999 |
| cluster3133 | -0.39 | 0.999 |
| cluster3134 | -0.40 | 0.999 |
| cluster3135 | -0.94 | 0.999 |
| cluster3136 | 0.43 | 0.999 |
| cluster3137 | 2.71 | 0.999 |
| cluster3138 | 1.36 | 0.999 |
| cluster3139 | 0.24 | 0.999 |
| cluster3140 | 0.36 | 0.999 |
| cluster3141 | 0.57 | 0.999 |
| cluster3142 | 1.24 | 0.999 |
| cluster3143 | 0.59 | 0.999 |
| cluster3144 | -0.70 | 0.999 |
| cluster3145 | -0.50 | 0.999 |
| cluster3146 | -0.49 | 0.999 |
| cluster3147 | 0.38 | 0.999 |
| cluster3148 | 0.24 | 0.999 |
| cluster3149 | 0.27 | 0.999 |
| cluster3150 | -1.09 | 0.999 |
| cluster3151 | -0.34 | 0.999 |
| cluster3152 | 0.09 | 0.999 |
| cluster3153 | 0.44 | 0.999 |
| cluster3154 | -0.08 | 0.999 |
| cluster3155 | 0.46 | 0.999 |
| cluster3156 | -0.86 | 0.999 |
| cluster3157 | 0.71 | 0.999 |
| cluster3158 | -0.06 | 0.999 |
| cluster3159 | -0.26 | 0.999 |
| cluster3160 | -0.58 | 0.999 |
| cluster3161 | -1.10 | 0.999 |
| cluster3162 | 0.97 | 0.999 |
| cluster3163 | -0.18 | 0.999 |
| cluster3164 | -1.17 | 0.999 |
| cluster3165 | 1.61 | 0.999 |
| cluster3166 | 0.28 | 0.999 |
| cluster3167 | -0.11 | 0.999 |
| cluster3168 | 1.08 | 0.999 |

| | | |
|---|---|---|
| cluster3169 | 0.42 | 0.999 |
| cluster3170 | 0.10 | 0.999 |
| cluster3171 | -0.09 | 0.999 |
| cluster3172 | -0.44 | 0.999 |
| cluster3173 | 0.38 | 0.999 |
| cluster3174 | 0.96 | 0.999 |
| cluster3175 | -1.20 | 0.999 |
| cluster3176 | -1.58 | 0.999 |
| cluster3177 | -1.89 | 0.999 |
| cluster3178 | 2.50 | 0.999 |
| cluster3179 | 1.42 | 0.999 |
| cluster3180 | -1.00 | 0.999 |
| cluster3181 | 0.55 | 0.999 |
| cluster3182 | -1.27 | 0.999 |
| cluster3183 | -0.37 | 0.999 |
| cluster3184 | 0.02 | 0.999 |
| cluster3185 | -1.21 | 0.999 |
| cluster3186 | -0.40 | 0.999 |
| cluster3187 | -0.76 | 0.999 |
| cluster3188 | 0.79 | 0.999 |
| cluster3189 | 0.30 | 0.999 |
| cluster3190 | 0.61 | 0.999 |
| cluster3191 | -0.93 | 0.999 |
| cluster3192 | 0.04 | 0.999 |
| cluster3193 | 0.04 | 0.999 |
| cluster3194 | -0.79 | 0.999 |
| cluster3195 | -0.99 | 0.999 |
| cluster3196 | -0.45 | 0.999 |
| cluster3197 | 0.99 | 0.999 |
| cluster3198 | -0.15 | 0.999 |
| cluster3199 | -0.23 | 0.999 |
| cluster3200 | 1.21 | 0.999 |
| cluster3201 | 0.05 | 0.999 |
| cluster3202 | 0.22 | 0.999 |
| cluster3203 | -1.01 | 0.999 |
| cluster3204 | 0.34 | 0.999 |
| cluster3205 | -0.81 | 0.999 |
| cluster3206 | -0.29 | 0.999 |
| cluster3207 | 0.50 | 0.999 |
| cluster3208 | 0.00 | 0.999 |
| cluster3209 | -0.43 | 0.999 |
| cluster3210 | 0.28 | 0.999 |
| cluster3211 | 0.21 | 0.999 |
| cluster3212 | -1.62 | 0.999 |

| | | |
|---|---|---|
| cluster3213 | -0.21 | 0.999 |
| cluster3214 | -0.22 | 0.999 |
| cluster3215 | 0.28 | 0.999 |
| cluster3216 | 0.24 | 0.999 |
| cluster3217 | -0.41 | 0.999 |
| cluster3218 | 0.97 | 0.999 |
| cluster3219 | -0.45 | 0.999 |
| cluster3220 | -1.26 | 0.999 |
| cluster3221 | 0.43 | 0.999 |
| cluster3222 | -0.31 | 0.999 |
| cluster3224 | -0.61 | 0.999 |
| cluster3225 | -0.82 | 0.999 |
| cluster3226 | -0.99 | 0.999 |
| cluster3227 | 0.45 | 0.999 |
| cluster3228 | 0.49 | 0.999 |
| cluster3229 | 0.54 | 0.999 |
| cluster3230 | -0.11 | 0.999 |
| cluster3231 | -0.24 | 0.999 |
| cluster3232 | 0.81 | 0.999 |
| cluster3233 | 3.06 | 0.999 |
| cluster3234 | -0.72 | 0.999 |
| cluster3235 | 0.85 | 0.999 |
| cluster3236 | 0.25 | 0.999 |
| cluster3237 | 2.09 | 0.999 |
| cluster3238 | 0.42 | 0.999 |
| cluster3239 | 0.42 | 0.999 |
| cluster3240 | 1.15 | 0.999 |
| cluster3241 | 0.25 | 0.999 |
| cluster3242 | 0.57 | 0.999 |
| cluster3243 | 0.25 | 0.999 |
| cluster3244 | 0.77 | 0.999 |
| cluster3245 | 1.79 | 0.999 |
| cluster3246 | 0.96 | 0.999 |
| cluster3247 | 1.13 | 0.999 |
| cluster3248 | 0.04 | 0.999 |
| cluster3249 | -0.60 | 0.999 |
| cluster3250 | 2.42 | 0.999 |
| cluster3251 | -0.13 | 0.999 |
| cluster3252 | 0.43 | 0.999 |
| cluster3253 | -0.32 | 0.999 |
| cluster3254 | -0.46 | 0.999 |
| cluster3255 | -0.77 | 0.999 |
| cluster3256 | -0.01 | 0.999 |
| cluster3257 | 0.29 | 0.999 |

| | | |
|---|---|---|
| cluster3258 | 0.49 | 0.999 |
| cluster3259 | -0.60 | 0.999 |
| cluster3260 | -1.09 | 0.999 |
| cluster3261 | -0.75 | 0.999 |
| cluster3262 | -0.14 | 0.999 |
| cluster3263 | 0.27 | 0.999 |
| cluster3264 | -0.44 | 0.999 |
| cluster3265 | 0.38 | 0.999 |
| cluster3266 | 0.18 | 0.999 |
| cluster3267 | 0.36 | 0.999 |
| cluster3268 | -0.50 | 0.999 |
| cluster3269 | -0.67 | 0.999 |
| cluster3270 | 0.50 | 0.999 |
| cluster3271 | -0.46 | 0.999 |
| cluster3272 | -0.39 | 0.999 |
| cluster3273 | 0.36 | 0.999 |
| cluster3274 | 0.33 | 0.999 |
| cluster3275 | -0.20 | 0.999 |
| cluster3276 | 0.27 | 0.999 |
| cluster3277 | -0.73 | 0.999 |
| cluster3278 | -0.18 | 0.999 |
| cluster3279 | 0.23 | 0.999 |
| cluster3280 | 0.55 | 0.999 |
| cluster3281 | 0.03 | 0.999 |
| cluster3282 | -0.43 | 0.999 |
| cluster3283 | 0.12 | 0.999 |
| cluster3284 | 0.17 | 0.999 |
| cluster3285 | -0.20 | 0.999 |
| cluster3286 | 0.05 | 0.999 |
| cluster3287 | 0.28 | 0.999 |
| cluster3288 | 0.96 | 0.999 |
| cluster3289 | 0.11 | 0.999 |
| cluster3290 | -1.63 | 0.999 |
| cluster3291 | 0.30 | 0.999 |
| cluster3292 | -0.98 | 0.999 |
| cluster3293 | 0.21 | 0.999 |
| cluster3294 | -0.17 | 0.999 |
| cluster3295 | 0.27 | 0.999 |
| cluster3296 | -1.04 | 0.999 |
| cluster3298 | -0.94 | 0.999 |
| cluster3299 | -0.62 | 0.999 |
| cluster3300 | 0.60 | 0.999 |
| cluster3301 | -0.52 | 0.999 |
| cluster3302 | -0.24 | 0.999 |

| | | |
|---|---|---|
| cluster3303 | 2.24 | 0.999 |
| cluster3304 | 0.72 | 0.999 |
| cluster3305 | -0.21 | 0.999 |
| cluster3306 | 1.71 | 0.999 |
| cluster3307 | -0.36 | 0.999 |
| cluster3308 | -0.52 | 0.999 |
| cluster3309 | 1.79 | 0.999 |
| cluster3310 | -0.59 | 0.999 |
| cluster3311 | 0.84 | 0.999 |
| cluster3312 | -1.69 | 0.999 |
| cluster3313 | -1.15 | 0.999 |
| cluster3314 | 0.26 | 0.999 |
| cluster3315 | 0.09 | 0.999 |
| cluster3316 | -0.36 | 0.999 |
| cluster3317 | 1.82 | 0.999 |
| cluster3318 | 0.24 | 0.999 |
| cluster3319 | -0.77 | 0.999 |
| cluster3320 | -0.80 | 0.999 |
| cluster3321 | 0.56 | 0.999 |
| cluster3322 | -0.01 | 0.999 |
| cluster3323 | 1.91 | 0.999 |
| cluster3324 | -0.10 | 0.999 |
| cluster3325 | -0.78 | 0.999 |
| cluster3326 | -0.03 | 0.999 |
| cluster3327 | 0.43 | 0.999 |
| cluster3328 | -0.23 | 0.999 |
| cluster3329 | -1.32 | 0.999 |
| cluster3330 | 0.47 | 0.999 |
| cluster3331 | -0.60 | 0.999 |
| cluster3332 | 1.36 | 0.999 |
| cluster3333 | 0.10 | 0.999 |
| cluster3334 | 0.45 | 0.999 |
| cluster3335 | 1.21 | 0.999 |
| cluster3336 | -0.52 | 0.999 |
| cluster3337 | 0.86 | 0.999 |
| cluster3338 | -0.88 | 0.999 |
| cluster3339 | -0.12 | 0.999 |
| cluster3340 | 0.60 | 0.999 |
| cluster3341 | -1.12 | 0.999 |
| cluster3342 | 0.42 | 0.999 |
| cluster3343 | -0.01 | 0.999 |
| cluster3344 | -0.55 | 0.999 |
| cluster3345 | -0.66 | 0.999 |
| cluster3346 | 0.46 | 0.999 |

| | | |
|---|---|---|
| cluster3347 | -0.07 | 0.999 |
| cluster3348 | 0.46 | 0.999 |
| cluster3349 | 0.57 | 0.999 |
| cluster3350 | -0.09 | 0.999 |
| cluster3351 | -0.17 | 0.999 |
| cluster3352 | 0.50 | 0.999 |
| cluster3353 | 0.35 | 0.999 |
| cluster3354 | 0.17 | 0.999 |
| cluster3355 | -0.17 | 0.999 |
| cluster3356 | 0.59 | 0.999 |
| cluster3357 | 1.35 | 0.999 |
| cluster3358 | 0.27 | 0.999 |
| cluster3359 | -0.28 | 0.999 |
| cluster3360 | -0.71 | 0.999 |
| cluster3361 | -0.77 | 0.999 |
| cluster3362 | -0.01 | 0.999 |
| cluster3363 | 1.06 | 0.999 |
| cluster3364 | -0.30 | 0.999 |
| cluster3365 | 1.15 | 0.999 |
| cluster3366 | 0.85 | 0.999 |
| cluster3368 | -0.38 | 0.999 |
| cluster3369 | 0.99 | 0.999 |
| cluster3370 | 0.20 | 0.999 |
| cluster3372 | -0.01 | 0.999 |
| cluster3373 | 0.61 | 0.999 |
| cluster3374 | 0.54 | 0.999 |
| cluster3375 | -0.99 | 0.999 |
| cluster3376 | -0.34 | 0.999 |
| cluster3377 | -0.91 | 0.999 |
| cluster3378 | -0.17 | 0.999 |
| cluster3379 | -0.72 | 0.999 |
| cluster3380 | -1.12 | 0.999 |
| cluster3381 | 0.03 | 0.999 |
| cluster3382 | -0.47 | 0.999 |
| cluster3383 | -0.36 | 0.999 |
| cluster3384 | -0.10 | 0.999 |
| cluster3385 | -0.65 | 0.999 |
| cluster3386 | -0.56 | 0.999 |
| cluster3387 | -0.50 | 0.999 |
| cluster3388 | -1.63 | 0.999 |
| cluster3389 | -0.23 | 0.999 |
| cluster3390 | -0.28 | 0.999 |
| cluster3391 | -0.25 | 0.999 |
| cluster3392 | -0.83 | 0.999 |

| | | |
|---|---|---|
| cluster3394 | -0.48 | 0.999 |
| cluster3395 | -0.98 | 0.999 |
| cluster3396 | -0.41 | 0.999 |
| cluster3397 | -1.92 | 0.999 |
| cluster3398 | -1.04 | 0.999 |
| cluster3399 | 0.91 | 0.999 |
| cluster3400 | 0.96 | 0.999 |
| cluster3401 | 0.84 | 0.999 |
| cluster3402 | 0.73 | 0.999 |
| cluster3403 | -0.42 | 0.999 |
| cluster3404 | -0.02 | 0.999 |
| cluster3405 | 0.53 | 0.999 |
| cluster3406 | -0.46 | 0.999 |
| cluster3407 | 0.06 | 0.999 |
| cluster3408 | 0.73 | 0.999 |
| cluster3409 | -0.01 | 0.999 |
| cluster3410 | 0.61 | 0.999 |
| cluster3411 | -0.52 | 0.999 |
| cluster3412 | 0.05 | 0.999 |
| cluster3413 | -1.00 | 0.999 |
| cluster3414 | 0.49 | 0.999 |
| cluster3415 | -0.19 | 0.999 |
| cluster3416 | 0.08 | 0.999 |
| cluster3417 | 0.47 | 0.999 |
| cluster3418 | 0.24 | 0.999 |
| cluster3419 | -0.17 | 0.999 |
| cluster3420 | 0.23 | 0.999 |
| cluster3421 | 0.89 | 0.999 |
| cluster3422 | 0.00 | 0.999 |
| cluster3423 | 0.42 | 0.999 |
| cluster3424 | -0.72 | 0.999 |
| cluster3425 | -0.87 | 0.999 |
| cluster3426 | 1.03 | 0.999 |
| cluster3427 | -0.42 | 0.999 |
| cluster3428 | -0.15 | 0.999 |
| cluster3429 | -0.20 | 0.999 |
| cluster3430 | 2.01 | 0.999 |
| cluster3431 | 1.16 | 0.999 |
| cluster3432 | -1.96 | 0.999 |
| cluster3433 | 0.55 | 0.999 |
| cluster3434 | -1.36 | 0.999 |
| cluster3435 | -1.49 | 0.999 |
| cluster3436 | 1.64 | 0.999 |
| cluster3437 | -1.23 | 0.999 |

| | | |
|---|---|---|
| cluster3438 | 1.14 | 0.999 |
| cluster3439 | -0.99 | 0.999 |
| cluster3440 | -0.26 | 0.999 |
| cluster3441 | 0.70 | 0.999 |
| cluster3442 | 0.95 | 0.999 |
| cluster3443 | 0.62 | 0.999 |
| cluster3444 | -0.09 | 0.999 |
| cluster3445 | -0.84 | 0.999 |
| cluster3446 | -0.93 | 0.999 |
| cluster3447 | -0.08 | 0.999 |
| cluster3448 | -1.40 | 0.999 |
| cluster3449 | 0.43 | 0.999 |
| cluster3450 | -0.79 | 0.999 |
| cluster3451 | 1.20 | 0.999 |
| cluster3452 | -1.04 | 0.999 |
| cluster3453 | -1.36 | 0.999 |
| cluster3454 | -1.16 | 0.999 |
| cluster3455 | -0.53 | 0.999 |
| cluster3456 | 0.18 | 0.999 |
| cluster3457 | -0.32 | 0.999 |
| cluster3458 | 0.68 | 0.999 |
| cluster3459 | -1.06 | 0.999 |
| cluster3460 | 0.81 | 0.999 |
| cluster3461 | -3.07 | 0.999 |
| cluster3462 | -1.38 | 0.999 |
| cluster3463 | 0.61 | 0.999 |
| cluster3464 | 0.11 | 0.999 |
| cluster3465 | 0.24 | 0.999 |
| cluster3466 | -0.61 | 0.999 |
| cluster3467 | -0.02 | 0.999 |
| cluster3468 | 0.65 | 0.999 |
| cluster3469 | 0.53 | 0.999 |
| cluster3470 | -0.45 | 0.999 |
| cluster3471 | 0.55 | 0.999 |
| cluster3472 | 0.03 | 0.999 |
| cluster3473 | 0.91 | 0.999 |
| cluster3474 | 1.73 | 0.999 |
| cluster3475 | 0.01 | 0.999 |
| cluster3476 | 0.36 | 0.999 |
| cluster3477 | -0.25 | 0.999 |
| cluster3478 | 0.50 | 0.999 |
| cluster3479 | -0.45 | 0.999 |
| cluster3480 | 1.36 | 0.999 |
| cluster3481 | -1.34 | 0.999 |

| | | |
|---|---|---|
| cluster3482 | -1.96 | 0.999 |
| cluster3483 | -0.30 | 0.999 |
| cluster3484 | -0.77 | 0.999 |
| cluster3485 | -1.54 | 0.999 |
| cluster3486 | 0.18 | 0.999 |
| cluster3487 | -0.98 | 0.999 |
| cluster3488 | -0.67 | 0.999 |
| cluster3489 | 0.24 | 0.999 |
| cluster3490 | -0.91 | 0.999 |
| cluster3491 | 0.00 | 0.999 |
| cluster3492 | 0.15 | 0.999 |
| cluster3493 | 1.25 | 0.999 |
| cluster3494 | 0.78 | 0.999 |
| cluster3495 | 0.67 | 0.999 |
| cluster3496 | 0.39 | 0.999 |
| cluster3497 | 0.49 | 0.999 |
| cluster3498 | 0.67 | 0.999 |
| cluster3499 | -0.06 | 0.999 |
| cluster3500 | -1.23 | 0.999 |
| cluster3501 | -2.15 | 0.999 |
| cluster3502 | 0.35 | 0.999 |
| cluster3503 | 0.13 | 0.999 |
| cluster3504 | -0.25 | 0.999 |
| cluster3505 | 0.35 | 0.999 |
| cluster3506 | 0.22 | 0.999 |
| cluster3507 | -0.16 | 0.999 |
| cluster3508 | 1.44 | 0.999 |
| cluster3509 | 0.47 | 0.999 |
| cluster3510 | -1.72 | 0.999 |
| cluster3511 | -0.78 | 0.999 |
| cluster3512 | -0.05 | 0.999 |
| cluster3513 | 0.77 | 0.999 |
| cluster3514 | 1.09 | 0.999 |
| cluster3515 | 1.56 | 0.999 |
| cluster3516 | -0.74 | 0.999 |
| cluster3517 | -0.39 | 0.999 |
| cluster3518 | -0.69 | 0.999 |
| cluster3519 | -1.23 | 0.999 |
| cluster3520 | 1.87 | 0.999 |
| cluster3521 | -0.36 | 0.999 |
| cluster3522 | -1.29 | 0.999 |
| cluster3523 | -0.14 | 0.999 |
| cluster3524 | -0.10 | 0.999 |
| cluster3525 | -1.38 | 0.999 |

| | | |
|---|---|---|
| cluster3526 | 0.33 | 0.999 |
| cluster3527 | -0.37 | 0.999 |
| cluster3528 | -0.35 | 0.999 |
| cluster3529 | 0.08 | 0.999 |
| cluster3530 | 0.63 | 0.999 |
| cluster3531 | -0.85 | 0.999 |
| cluster3532 | -0.33 | 0.999 |
| cluster3533 | -0.12 | 0.999 |
| cluster3534 | -1.85 | 0.999 |
| cluster3535 | 0.30 | 0.999 |
| cluster3536 | 0.49 | 0.999 |
| cluster3537 | 1.03 | 0.999 |
| cluster3538 | 1.05 | 0.999 |
| cluster3539 | -0.27 | 0.999 |
| cluster3540 | -0.58 | 0.999 |
| cluster3541 | 0.35 | 0.999 |
| cluster3542 | 0.05 | 0.999 |
| cluster3543 | 0.01 | 0.999 |
| cluster3544 | -1.09 | 0.999 |
| cluster3545 | 0.23 | 0.999 |
| cluster3546 | -2.12 | 0.999 |
| cluster3547 | 0.20 | 0.999 |
| cluster3548 | 0.42 | 0.999 |
| cluster3549 | 0.63 | 0.999 |
| cluster3550 | -0.01 | 0.999 |
| cluster3551 | 1.19 | 0.999 |
| cluster3552 | 0.55 | 0.999 |
| cluster3553 | -0.64 | 0.999 |
| cluster3554 | -0.71 | 0.999 |
| cluster3555 | -0.18 | 0.999 |
| cluster3556 | 1.49 | 0.999 |
| cluster3557 | -0.43 | 0.999 |
| cluster3558 | -0.06 | 0.999 |
| cluster3559 | 0.36 | 0.999 |
| cluster3560 | 0.16 | 0.999 |
| cluster3561 | 1.94 | 0.999 |
| cluster3562 | -0.22 | 0.999 |
| cluster3563 | -1.67 | 0.999 |
| cluster3564 | 1.52 | 0.999 |
| cluster3565 | -0.82 | 0.999 |
| cluster3566 | -0.39 | 0.999 |
| cluster3567 | 1.47 | 0.999 |
| cluster3568 | -1.56 | 0.999 |
| cluster3569 | 0.24 | 0.999 |

| | | |
|---|---|---|
| cluster3570 | 0.01 | 0.999 |
| cluster3571 | -1.87 | 0.999 |
| cluster3572 | 1.03 | 0.999 |
| cluster3573 | -0.88 | 0.999 |
| cluster3574 | -2.66 | 0.999 |
| cluster3575 | 1.64 | 0.999 |
| cluster3576 | 0.26 | 0.999 |
| cluster3577 | 0.66 | 0.999 |
| cluster3578 | -0.74 | 0.999 |
| cluster3579 | -0.36 | 0.999 |
| cluster3580 | 1.13 | 0.999 |
| cluster3581 | -1.08 | 0.999 |
| cluster3582 | 0.59 | 0.999 |
| cluster3583 | -0.25 | 0.999 |
| cluster3584 | 0.61 | 0.999 |
| cluster3585 | -1.26 | 0.999 |
| cluster3586 | 0.31 | 0.999 |
| cluster3587 | 0.44 | 0.999 |
| cluster3588 | 0.41 | 0.999 |
| cluster3589 | 0.05 | 0.999 |
| cluster3590 | 0.20 | 0.999 |
| cluster3591 | -0.65 | 0.999 |
| cluster3592 | -0.05 | 0.999 |
| cluster3593 | -0.82 | 0.999 |
| cluster3594 | -0.47 | 0.999 |
| cluster3595 | 1.04 | 0.999 |
| cluster3596 | -0.50 | 0.999 |
| cluster3597 | 1.33 | 0.999 |
| cluster3598 | 1.25 | 0.999 |
| cluster3599 | -1.45 | 0.999 |
| cluster3600 | 0.40 | 0.999 |
| cluster3601 | 0.30 | 0.999 |
| cluster3602 | 0.23 | 0.999 |
| cluster3603 | 0.93 | 0.999 |
| cluster3604 | 0.55 | 0.999 |
| cluster3605 | 0.03 | 0.999 |
| cluster3606 | -0.21 | 0.999 |
| cluster3607 | 0.27 | 0.999 |
| cluster3608 | -0.78 | 0.999 |
| cluster3609 | 1.50 | 0.999 |
| cluster3610 | -0.09 | 0.999 |
| cluster3611 | 1.03 | 0.999 |
| cluster3612 | 0.16 | 0.999 |
| cluster3613 | 0.46 | 0.999 |

| | | |
|---|---|---|
| cluster3614 | 0.33 | 0.999 |
| cluster3615 | 1.38 | 0.999 |
| cluster3616 | -0.54 | 0.999 |
| cluster3617 | 0.32 | 0.999 |
| cluster3618 | 0.43 | 0.999 |
| cluster3619 | 0.53 | 0.999 |
| cluster3620 | -0.75 | 0.999 |
| cluster3621 | 1.19 | 0.999 |
| cluster3622 | -0.28 | 0.999 |
| cluster3623 | -0.52 | 0.999 |
| cluster3624 | 0.16 | 0.999 |
| cluster3625 | -0.15 | 0.999 |
| cluster3626 | -0.30 | 0.999 |
| cluster3627 | -0.45 | 0.999 |
| cluster3628 | 0.05 | 0.999 |
| cluster3629 | 1.27 | 0.999 |
| cluster3630 | -0.19 | 0.999 |
| cluster3631 | 1.91 | 0.999 |
| cluster3632 | 0.06 | 0.999 |
| cluster3633 | -1.06 | 0.999 |
| cluster3634 | -0.17 | 0.999 |
| cluster3635 | -0.09 | 0.999 |
| cluster3636 | -1.21 | 0.999 |
| cluster3637 | 0.04 | 0.999 |
| cluster3638 | 0.55 | 0.999 |
| cluster3639 | 0.17 | 0.999 |
| cluster3640 | -0.92 | 0.999 |
| cluster3641 | -0.26 | 0.999 |
| cluster3642 | 0.38 | 0.999 |
| cluster3643 | 0.82 | 0.999 |
| cluster3644 | -1.15 | 0.999 |
| cluster3645 | -3.35 | 0.999 |
| cluster3646 | 0.18 | 0.999 |
| cluster3647 | 0.10 | 0.999 |
| cluster3648 | -0.06 | 0.999 |
| cluster3649 | -0.64 | 0.999 |
| cluster3650 | 1.02 | 0.999 |
| cluster3651 | -3.10 | 0.999 |
| cluster3652 | 1.96 | 0.999 |
| cluster3653 | 0.43 | 0.999 |
| cluster3654 | -0.65 | 0.999 |
| cluster3655 | -0.37 | 0.999 |
| cluster3656 | 1.19 | 0.999 |
| cluster3657 | 0.58 | 0.999 |

| | | |
|---|---|---|
| cluster3658 | 0.23 | 0.999 |
| cluster3659 | -1.28 | 0.999 |
| cluster3660 | -0.41 | 0.999 |
| cluster3661 | 0.87 | 0.999 |
| cluster3662 | -0.81 | 0.999 |
| cluster3663 | -0.31 | 0.999 |
| cluster3664 | -0.18 | 0.999 |
| cluster3665 | -0.68 | 0.999 |
| cluster3666 | 2.68 | 0.999 |
| cluster3667 | -1.18 | 0.999 |
| cluster3668 | -0.28 | 0.999 |
| cluster3669 | -0.73 | 0.999 |
| cluster3670 | -0.61 | 0.999 |
| cluster3671 | -0.44 | 0.999 |
| cluster3672 | -1.61 | 0.999 |
| cluster3673 | -0.44 | 0.999 |
| cluster3675 | -0.80 | 0.999 |
| cluster3676 | 0.82 | 0.999 |
| cluster3677 | 0.37 | 0.999 |
| cluster3678 | 0.32 | 0.999 |
| cluster3679 | 0.74 | 0.999 |
| cluster3680 | 0.10 | 0.999 |
| cluster3681 | 0.11 | 0.999 |
| cluster3682 | -0.47 | 0.999 |
| cluster3683 | 0.01 | 0.999 |
| cluster3684 | 0.08 | 0.999 |
| cluster3685 | -0.20 | 0.999 |
| cluster3686 | 0.28 | 0.999 |
| cluster3687 | 0.02 | 0.999 |
| cluster3688 | 0.09 | 0.999 |
| cluster3689 | -1.82 | 0.999 |
| cluster3690 | -0.56 | 0.999 |
| cluster3691 | 1.19 | 0.999 |
| cluster3692 | 0.00 | 0.999 |
| cluster3693 | 0.73 | 0.999 |
| cluster3694 | -1.78 | 0.999 |
| cluster3695 | -1.06 | 0.999 |
| cluster3696 | -0.08 | 0.999 |
| cluster3697 | 0.04 | 0.999 |
| cluster3698 | -0.02 | 0.999 |
| cluster3699 | 0.48 | 0.999 |
| cluster3700 | 0.03 | 0.999 |
| cluster3701 | -0.26 | 0.999 |
| cluster3702 | 0.23 | 0.999 |

| | | |
|---|---|---|
| cluster3703 | 0.61 | 0.999 |
| cluster3704 | 1.49 | 0.999 |
| cluster3705 | 1.19 | 0.999 |
| cluster3706 | -0.46 | 0.999 |
| cluster3707 | 0.37 | 0.999 |
| cluster3708 | -0.04 | 0.999 |
| cluster3709 | 0.76 | 0.999 |
| cluster3710 | 0.28 | 0.999 |
| cluster3711 | -0.39 | 0.999 |
| cluster3712 | 0.65 | 0.999 |
| cluster3713 | 0.50 | 0.999 |
| cluster3715 | 0.66 | 0.999 |
| cluster3716 | -1.08 | 0.999 |
| cluster3717 | -0.52 | 0.999 |
| cluster3718 | -0.19 | 0.999 |
| cluster3719 | 1.77 | 0.999 |
| cluster3720 | -0.16 | 0.999 |
| cluster3721 | 1.01 | 0.999 |
| cluster3722 | 0.31 | 0.999 |
| cluster3723 | 0.27 | 0.999 |
| cluster3724 | 0.56 | 0.999 |
| cluster3725 | 0.15 | 0.999 |
| cluster3726 | -0.25 | 0.999 |
| cluster3727 | 0.00 | 0.999 |
| cluster3728 | -0.38 | 0.999 |
| cluster3729 | -0.63 | 0.999 |
| cluster3730 | 0.81 | 0.999 |
| cluster3731 | 0.91 | 0.999 |
| cluster3732 | -0.54 | 0.999 |
| cluster3733 | -0.28 | 0.999 |
| cluster3734 | 0.78 | 0.999 |
| cluster3735 | -1.64 | 0.999 |
| cluster3736 | -0.41 | 0.999 |
| cluster3737 | 0.12 | 0.999 |
| cluster3738 | 2.51 | 0.999 |
| cluster3739 | 0.02 | 0.999 |
| cluster3740 | -0.37 | 0.999 |
| cluster3741 | 1.52 | 0.999 |
| cluster3742 | 1.15 | 0.999 |
| cluster3743 | -0.34 | 0.999 |
| cluster3744 | -0.66 | 0.999 |
| cluster3745 | 2.20 | 0.999 |
| cluster3746 | 1.01 | 0.999 |
| cluster3747 | 0.83 | 0.999 |

| | | |
|---|---|---|
| cluster3748 | 0.34 | 0.999 |
| cluster3749 | -0.98 | 0.999 |
| cluster3750 | -1.99 | 0.999 |
| cluster3751 | -0.51 | 0.999 |
| cluster3753 | -1.58 | 0.999 |
| cluster3754 | -0.21 | 0.999 |
| cluster3755 | 3.21 | 0.999 |
| cluster3756 | -0.94 | 0.999 |
| cluster3758 | -1.27 | 0.999 |
| cluster3759 | -0.50 | 0.999 |
| cluster3760 | 1.04 | 0.999 |
| cluster3761 | 0.57 | 0.999 |
| cluster3762 | 0.38 | 0.999 |
| cluster3763 | 2.20 | 0.999 |
| cluster3764 | -1.03 | 0.999 |
| cluster3765 | -0.97 | 0.999 |
| cluster3766 | -1.29 | 0.999 |
| cluster3767 | -0.35 | 0.999 |
| cluster3768 | -0.71 | 0.999 |
| cluster3769 | -0.66 | 0.999 |
| cluster3770 | -1.20 | 0.999 |
| cluster3771 | 0.87 | 0.999 |
| cluster3772 | -1.38 | 0.999 |
| cluster3773 | -1.03 | 0.999 |
| cluster3774 | -0.01 | 0.999 |
| cluster3775 | -0.79 | 0.999 |
| cluster3776 | -0.54 | 0.999 |
| cluster3777 | -0.69 | 0.999 |
| cluster3778 | 0.42 | 0.999 |
| cluster3779 | 0.60 | 0.999 |
| cluster3780 | -0.23 | 0.999 |
| cluster3781 | 0.60 | 0.999 |
| cluster3782 | -0.41 | 0.999 |
| cluster3783 | -0.04 | 0.999 |
| cluster3784 | 0.32 | 0.999 |
| cluster3785 | 0.14 | 0.999 |
| cluster3786 | 3.07 | 0.999 |
| cluster3787 | -1.19 | 0.999 |
| cluster3788 | 0.73 | 0.999 |
| cluster3789 | -0.28 | 0.999 |
| cluster3790 | -0.20 | 0.999 |
| cluster3791 | 0.48 | 0.999 |
| cluster3792 | -1.18 | 0.999 |
| cluster3793 | -0.12 | 0.999 |

| | | |
|---|---|---|
| cluster3794 | -1.12 | 0.999 |
| cluster3795 | -1.15 | 0.999 |
| cluster3796 | -0.36 | 0.999 |
| cluster3797 | 0.07 | 0.999 |
| cluster3798 | -0.14 | 0.999 |
| cluster3799 | 0.88 | 0.999 |
| cluster3800 | -1.83 | 0.999 |
| cluster3801 | 0.44 | 0.999 |
| cluster3802 | 2.24 | 0.999 |
| cluster3803 | 0.09 | 0.999 |
| cluster3804 | 0.64 | 0.999 |
| cluster3805 | 0.66 | 0.999 |
| cluster3806 | -0.67 | 0.999 |
| cluster3807 | -0.45 | 0.999 |
| cluster3808 | 0.30 | 0.999 |
| cluster3809 | -0.16 | 0.999 |
| cluster3810 | 0.53 | 0.999 |
| cluster3811 | 0.23 | 0.999 |
| cluster3813 | 0.42 | 0.999 |
| cluster3814 | -0.05 | 0.999 |
| cluster3815 | 0.68 | 0.999 |
| cluster3816 | -0.09 | 0.999 |
| cluster3817 | 0.61 | 0.999 |
| cluster3818 | 1.64 | 0.999 |
| cluster3819 | -0.92 | 0.999 |
| cluster3820 | -0.73 | 0.999 |
| cluster3821 | -1.28 | 0.999 |
| cluster3822 | 1.87 | 0.999 |
| cluster3823 | -0.51 | 0.999 |
| cluster3824 | 0.62 | 0.999 |
| cluster3825 | -0.85 | 0.999 |
| cluster3826 | -0.53 | 0.999 |
| cluster3827 | -1.23 | 0.999 |
| cluster3828 | 0.07 | 0.999 |
| cluster3829 | 0.81 | 0.999 |
| cluster3830 | 0.92 | 0.999 |
| cluster3831 | -0.42 | 0.999 |
| cluster3832 | 0.53 | 0.999 |
| cluster3833 | -0.11 | 0.999 |
| cluster3834 | -1.57 | 0.999 |
| cluster3835 | -0.11 | 0.999 |
| cluster3836 | -0.21 | 0.999 |
| cluster3837 | -0.22 | 0.999 |
| cluster3838 | 0.04 | 0.999 |

| | | |
|---|---|---|
| cluster3839 | -1.98 | 0.999 |
| cluster3840 | 1.87 | 0.999 |
| cluster3841 | -0.53 | 0.999 |
| cluster3842 | 1.16 | 0.999 |
| cluster3843 | -0.28 | 0.999 |
| cluster3844 | -1.44 | 0.999 |
| cluster3845 | -0.96 | 0.999 |
| cluster3846 | -0.19 | 0.999 |
| cluster3847 | 0.54 | 0.999 |
| cluster3848 | -0.36 | 0.999 |
| cluster3849 | 0.98 | 0.999 |
| cluster3850 | -1.32 | 0.999 |
| cluster3851 | -0.05 | 0.999 |
| cluster3852 | 2.10 | 0.999 |
| cluster3853 | 1.49 | 0.999 |
| cluster3854 | -0.91 | 0.999 |
| cluster3855 | -0.18 | 0.999 |
| cluster3856 | -0.46 | 0.999 |
| cluster3857 | -0.61 | 0.999 |
| cluster3858 | 0.99 | 0.999 |
| cluster3859 | -0.22 | 0.999 |
| cluster3860 | 0.20 | 0.999 |
| cluster3861 | -0.34 | 0.999 |
| cluster3862 | 0.02 | 0.999 |
| cluster3863 | 1.29 | 0.999 |
| cluster3864 | -0.13 | 0.999 |
| cluster3865 | 1.87 | 0.999 |
| cluster3866 | 0.10 | 0.999 |
| cluster3867 | -0.66 | 0.999 |
| cluster3868 | -0.87 | 0.999 |
| cluster3869 | 1.35 | 0.999 |
| cluster3870 | -0.10 | 0.999 |
| cluster3871 | 0.78 | 0.999 |
| cluster3872 | 0.84 | 0.999 |
| cluster3873 | 0.48 | 0.999 |
| cluster3874 | -0.32 | 0.999 |
| cluster3875 | 0.38 | 0.999 |
| cluster3876 | 1.12 | 0.999 |
| cluster3877 | -0.69 | 0.999 |
| cluster3879 | 0.37 | 0.999 |
| cluster3880 | 0.77 | 0.999 |
| cluster3881 | 0.56 | 0.999 |
| cluster3882 | 0.20 | 0.999 |
| cluster3883 | 1.11 | 0.999 |

| | | |
|---|---|---|
| cluster3884 | -1.29 | 0.999 |
| cluster3885 | -1.28 | 0.999 |
| cluster3886 | 0.37 | 0.999 |
| cluster3887 | -1.15 | 0.999 |
| cluster3888 | -0.21 | 0.999 |
| cluster3889 | 0.57 | 0.999 |
| cluster3890 | 1.82 | 0.999 |
| cluster3891 | 0.27 | 0.999 |
| cluster3893 | -0.16 | 0.999 |
| cluster3894 | -1.37 | 0.999 |
| cluster3895 | 0.02 | 0.999 |
| cluster3896 | 0.04 | 0.999 |
| cluster3897 | 1.86 | 0.999 |
| cluster3898 | 1.50 | 0.999 |
| cluster3899 | 0.97 | 0.999 |
| cluster3900 | 0.19 | 0.999 |
| cluster3901 | 1.90 | 0.999 |
| cluster3902 | 0.82 | 0.999 |
| cluster3903 | -0.22 | 0.999 |
| cluster3904 | -2.05 | 0.999 |
| cluster3905 | -1.16 | 0.999 |
| cluster3906 | -0.50 | 0.999 |
| cluster3907 | -1.06 | 0.999 |
| cluster3908 | -0.41 | 0.999 |
| cluster3909 | -0.24 | 0.999 |
| cluster3910 | -0.20 | 0.999 |
| cluster3911 | 0.80 | 0.999 |
| cluster3912 | -0.64 | 0.999 |
| cluster3913 | -0.29 | 0.999 |
| cluster3914 | -0.28 | 0.999 |
| cluster3915 | -1.99 | 0.999 |
| cluster3916 | 0.50 | 0.999 |
| cluster3917 | 0.39 | 0.999 |
| cluster3918 | 1.20 | 0.999 |
| cluster3919 | -1.44 | 0.999 |
| cluster3920 | -1.61 | 0.999 |
| cluster3921 | 0.30 | 0.999 |
| cluster3922 | -1.00 | 0.999 |
| cluster3923 | -0.91 | 0.999 |
| cluster3924 | 0.31 | 0.999 |
| cluster3925 | -1.53 | 0.999 |
| cluster3926 | -0.36 | 0.999 |
| cluster3927 | 0.12 | 0.999 |
| cluster3928 | -0.77 | 0.999 |

| | | |
|---|---|---|
| cluster3929 | -0.90 | 0.999 |
| cluster3930 | 0.48 | 0.999 |
| cluster3931 | 1.76 | 0.999 |
| cluster3933 | -1.57 | 0.999 |
| cluster3934 | 0.36 | 0.999 |
| cluster3935 | 0.17 | 0.999 |
| cluster3936 | 0.61 | 0.999 |
| cluster3937 | 0.83 | 0.999 |
| cluster3938 | -0.18 | 0.999 |
| cluster3939 | 0.11 | 0.999 |
| cluster3940 | 1.10 | 0.999 |
| cluster3941 | -0.64 | 0.999 |
| cluster3942 | -0.60 | 0.999 |
| cluster3943 | 0.86 | 0.999 |
| cluster3944 | 0.11 | 0.999 |
| cluster3945 | 0.03 | 0.999 |
| cluster3946 | -0.16 | 0.999 |
| cluster3947 | 0.03 | 0.999 |
| cluster3948 | -0.79 | 0.999 |
| cluster3949 | -0.25 | 0.999 |
| cluster3950 | -0.02 | 0.999 |
| cluster3951 | 0.11 | 0.999 |
| cluster3952 | 0.86 | 0.999 |
| cluster3953 | -1.06 | 0.999 |
| cluster3954 | -0.93 | 0.999 |
| cluster3955 | -1.70 | 0.999 |
| cluster3956 | 0.65 | 0.999 |
| cluster3957 | 0.20 | 0.999 |
| cluster3958 | -2.09 | 0.999 |
| cluster3959 | -1.17 | 0.999 |
| cluster3960 | -0.94 | 0.999 |
| cluster3961 | 0.21 | 0.999 |
| cluster3962 | 0.25 | 0.999 |
| cluster3963 | -0.73 | 0.999 |
| cluster3964 | -0.78 | 0.999 |
| cluster3965 | 0.01 | 0.999 |
| cluster3966 | 0.10 | 0.999 |
| cluster3967 | 0.05 | 0.999 |
| cluster3968 | 0.33 | 0.999 |
| cluster3969 | -0.36 | 0.999 |
| cluster3970 | 1.16 | 0.999 |
| cluster3971 | 0.61 | 0.999 |
| cluster3972 | -0.61 | 0.999 |
| cluster3973 | -0.95 | 0.999 |

| | | |
|---|---|---|
| cluster3974 | 0.46 | 0.999 |
| cluster3975 | 0.33 | 0.999 |
| cluster3976 | -0.77 | 0.999 |
| cluster3977 | -0.55 | 0.999 |
| cluster3978 | -1.20 | 0.999 |
| cluster3979 | 0.38 | 0.999 |
| cluster3980 | 0.22 | 0.999 |
| cluster3981 | 1.67 | 0.999 |
| cluster3982 | -0.20 | 0.999 |
| cluster3983 | 1.23 | 0.999 |
| cluster3984 | -0.89 | 0.999 |
| cluster3985 | 0.69 | 0.999 |
| cluster3986 | -0.12 | 0.999 |
| cluster3987 | 1.01 | 0.999 |
| cluster3988 | -0.48 | 0.999 |
| cluster3989 | 0.59 | 0.999 |
| cluster3990 | 0.79 | 0.999 |
| cluster3991 | -0.29 | 0.999 |
| cluster3992 | 0.28 | 0.999 |
| cluster3993 | -0.61 | 0.999 |
| cluster3994 | -1.74 | 0.999 |
| cluster3995 | -1.41 | 0.999 |
| cluster3996 | 0.32 | 0.999 |
| cluster3997 | 0.61 | 0.999 |
| cluster3998 | 0.33 | 0.999 |
| cluster3999 | -0.98 | 0.999 |
| cluster4000 | 1.18 | 0.999 |
| cluster4001 | -2.04 | 0.999 |
| cluster4002 | 1.50 | 0.999 |
| cluster4003 | 0.12 | 0.999 |
| cluster4004 | -0.27 | 0.999 |
| cluster4005 | -0.80 | 0.999 |
| cluster4006 | -0.29 | 0.999 |
| cluster4007 | 0.17 | 0.999 |
| cluster4008 | 0.29 | 0.999 |
| cluster4009 | 0.57 | 0.999 |
| cluster4010 | 0.40 | 0.999 |
| cluster4011 | -1.59 | 0.999 |
| cluster4012 | 1.71 | 0.999 |
| cluster4013 | -4.38 | 0.999 |
| cluster4014 | -0.01 | 0.999 |
| cluster4015 | -1.12 | 0.999 |
| cluster4016 | 1.34 | 0.999 |
| cluster4017 | 0.29 | 0.999 |

| | | |
|---|---|---|
| cluster4018 | 0.87 | 0.999 |
| cluster4019 | 1.33 | 0.999 |
| cluster4020 | -0.39 | 0.999 |
| cluster4021 | 0.38 | 0.999 |
| cluster4022 | -0.66 | 0.999 |
| cluster4023 | -0.69 | 0.999 |
| cluster4024 | 0.09 | 0.999 |
| cluster4025 | -0.28 | 0.999 |
| cluster4026 | -0.80 | 0.999 |
| cluster4027 | 0.09 | 0.999 |
| cluster4028 | 0.52 | 0.999 |
| cluster4029 | -0.19 | 0.999 |
| cluster4030 | -0.41 | 0.999 |
| cluster4031 | -0.09 | 0.999 |
| cluster4032 | 0.33 | 0.999 |
| cluster4033 | -1.65 | 0.999 |
| cluster4034 | -2.25 | 0.999 |
| cluster4035 | 0.85 | 0.999 |
| cluster4036 | -0.09 | 0.999 |
| cluster4037 | 0.29 | 0.999 |
| cluster4038 | 0.08 | 0.999 |
| cluster4039 | 0.38 | 0.999 |
| cluster4040 | -0.11 | 0.999 |
| cluster4041 | -0.22 | 0.999 |
| cluster4042 | -0.73 | 0.999 |
| cluster4043 | -0.41 | 0.999 |
| cluster4044 | 0.48 | 0.999 |
| cluster4045 | 0.52 | 0.999 |
| cluster4046 | -3.44 | 0.999 |
| cluster4047 | 1.29 | 0.999 |
| cluster4049 | -2.06 | 0.999 |
| cluster4050 | -0.16 | 0.999 |
| cluster4051 | 1.01 | 0.999 |
| cluster4052 | 1.80 | 0.999 |
| cluster4053 | -1.51 | 0.999 |
| cluster4054 | 0.93 | 0.999 |
| cluster4055 | -1.55 | 0.999 |
| cluster4056 | -1.33 | 0.999 |
| cluster4057 | -0.06 | 0.999 |
| cluster4058 | -0.47 | 0.999 |
| cluster4059 | -0.93 | 0.999 |
| cluster4060 | 2.42 | 0.999 |
| cluster4061 | 0.46 | 0.999 |
| cluster4063 | -0.57 | 0.999 |

| | | |
|---|---|---|
| cluster4064 | 1.14 | 0.999 |
| cluster4065 | 0.19 | 0.999 |
| cluster4066 | -1.14 | 0.999 |
| cluster4068 | -0.15 | 0.999 |
| cluster4069 | -0.13 | 0.999 |
| cluster4070 | -1.65 | 0.999 |
| cluster4071 | -0.06 | 0.999 |
| cluster4072 | 0.29 | 0.999 |
| cluster4073 | 2.53 | 0.999 |
| cluster4074 | 2.04 | 0.999 |
| cluster4075 | 0.98 | 0.999 |
| cluster4076 | 2.38 | 0.999 |
| cluster4077 | -0.27 | 0.999 |
| cluster4078 | 1.29 | 0.999 |
| cluster4079 | 1.54 | 0.999 |
| cluster4080 | -1.88 | 0.999 |
| cluster4081 | 0.35 | 0.999 |
| cluster4082 | 0.86 | 0.999 |
| cluster4083 | 0.35 | 0.999 |
| cluster4084 | -0.84 | 0.999 |
| cluster4085 | -1.55 | 0.999 |
| cluster4086 | 1.04 | 0.999 |
| cluster4087 | 3.07 | 0.999 |
| cluster4088 | -0.63 | 0.999 |
| cluster4090 | -2.24 | 0.999 |
| cluster4092 | -0.25 | 0.999 |
| cluster4093 | 1.52 | 0.999 |
| cluster4094 | 0.98 | 0.999 |
| cluster4095 | 1.54 | 0.999 |
| cluster4096 | -0.27 | 0.999 |
| cluster4097 | -1.31 | 0.999 |
| cluster4098 | 1.98 | 0.999 |
| cluster4099 | 1.28 | 0.999 |
| cluster4100 | 0.34 | 0.999 |
| cluster4101 | 3.55 | 0.999 |
| cluster4102 | -0.16 | 0.999 |
| cluster4103 | 1.29 | 0.999 |
| cluster4104 | 0.09 | 0.999 |
| cluster4105 | 0.19 | 0.999 |
| cluster4106 | -0.76 | 0.999 |
| cluster4107 | 1.14 | 0.999 |
| cluster4108 | -0.59 | 0.999 |
| cluster4109 | 0.88 | 0.999 |
| cluster4110 | 0.39 | 0.999 |

| | | |
|---|---|---|
| cluster4111 | -2.81 | 0.999 |
| cluster4112 | 2.85 | 0.999 |
| cluster4113 | -1.21 | 0.999 |
| cluster4114 | 2.75 | 0.999 |
| cluster4115 | -1.13 | 0.999 |
| cluster4116 | -2.99 | 0.999 |
| cluster4117 | -0.37 | 0.999 |
| cluster4118 | -3.39 | 0.999 |
| cluster4119 | 2.15 | 0.999 |
| cluster4121 | 1.11 | 0.999 |
| cluster4123 | -0.24 | 0.999 |
| cluster4124 | -1.11 | 0.999 |
| cluster4125 | 0.67 | 0.999 |
| cluster4126 | 0.10 | 0.999 |
| cluster4127 | -1.03 | 0.999 |
| cluster4128 | -0.97 | 0.999 |

**Supplementary Table 10**. edgeR results with RSEM counts - Oestridae vs Calliphoridae

| ID | log2FoldChange | Adjusted p value |
|---|---|---|
| cluster4013 | 4.53 | 1.000 |
| cluster4046 | 3.59 | 1.000 |
| cluster3932 | 3.11 | 1.000 |
| cluster3651 | 2.99 | 1.000 |
| cluster4118 | 3.48 | 1.000 |
| cluster4101 | -3.71 | 1.000 |
| cluster3461 | 3.18 | 1.000 |
| cluster3645 | 3.34 | 1.000 |
| cluster3786 | -3.15 | 1.000 |
| cluster4116 | 2.96 | 1.000 |
| cluster3574 | 2.72 | 1.000 |
| cluster4111 | 2.88 | 1.000 |
| cluster3233 | -3.27 | 1.000 |
| cluster3178 | -2.61 | 1.000 |
| cluster3482 | 2.06 | 1.000 |
| cluster3755 | -3.28 | 1.000 |
| cluster4122 | 2.79 | 1.000 |
| cluster4112 | -2.91 | 1.000 |
| cluster4048 | 2.45 | 1.000 |
| cluster3738 | -2.69 | 1.000 |
| cluster3852 | -2.22 | 1.000 |
| cluster3432 | 1.86 | 1.000 |
| cluster3763 | -2.31 | 1.000 |
| cluster4076 | -2.51 | 1.000 |
| cluster3534 | 1.85 | 1.000 |
| cluster3802 | -2.40 | 1.000 |
| cluster3501 | 2.04 | 1.000 |
| cluster3303 | -2.36 | 1.000 |
| cluster3006 | -2.51 | 1.000 |
| cluster3958 | 2.20 | 1.000 |
| cluster3652 | -2.04 | 1.000 |
| cluster3250 | -2.41 | 1.000 |
| cluster3750 | 2.18 | 1.000 |
| cluster4067 | -3.32 | 1.000 |
| cluster3393 | 2.19 | 1.000 |
| cluster3571 | 1.81 | 1.000 |
| cluster3840 | -2.00 | 1.000 |
| cluster3030 | -2.22 | 1.000 |
| cluster3694 | 1.74 | 1.000 |
| cluster4090 | 2.33 | 1.000 |
| cluster4087 | -3.09 | 1.000 |
| cluster3897 | -1.96 | 1.000 |

| | | |
|---|---|---|
| cluster3309 | -1.87 | 1.000 |
| cluster3839 | 2.07 | 1.000 |
| cluster3563 | 1.59 | 1.000 |
| cluster4098 | -2.07 | 1.000 |
| cluster3137 | -2.61 | 1.000 |
| cluster3890 | -2.01 | 1.000 |
| cluster3000 | 1.90 | 1.000 |
| cluster4001 | 2.03 | 1.000 |
| cluster3323 | -1.92 | 1.000 |
| cluster3290 | 1.65 | 1.000 |
| cluster3388 | 1.80 | 1.000 |
| cluster4120 | 2.07 | 1.000 |
| cluster3176 | 1.71 | 1.000 |
| cluster4060 | -2.37 | 1.000 |
| cluster4052 | -1.89 | 1.000 |
| cluster3631 | -2.06 | 1.000 |
| cluster3397 | 1.95 | 1.000 |
| cluster3177 | 1.83 | 1.000 |
| cluster3915 | 1.98 | 1.000 |
| cluster4073 | -2.49 | 1.000 |
| cluster3297 | -1.89 | 1.000 |
| cluster3904 | 2.03 | 1.000 |
| cluster3561 | -1.82 | 1.000 |
| cluster3546 | 2.06 | 1.000 |
| cluster3666 | -2.68 | 1.000 |
| cluster3525 | 1.40 | 1.000 |
| cluster3818 | -1.80 | 1.000 |
| cluster3735 | 1.51 | 1.000 |
| cluster3920 | 1.81 | 1.000 |
| cluster2948 | -1.85 | 1.000 |
| cluster3095 | 1.37 | 1.000 |
| cluster3689 | 1.91 | 1.000 |
| cluster3367 | -1.91 | 1.000 |
| cluster3672 | 1.65 | 1.000 |
| cluster3371 | -2.92 | 1.000 |
| cluster3933 | 1.65 | 1.000 |
| cluster3955 | 1.68 | 1.000 |
| cluster3063 | -1.63 | 1.000 |
| cluster2861 | -1.54 | 1.000 |
| cluster4033 | 1.70 | 1.000 |
| cluster4114 | -2.63 | 1.000 |
| cluster3585 | 1.25 | 1.000 |
| cluster3772 | 1.51 | 1.000 |
| cluster3045 | -1.52 | 1.000 |

| | | |
|---|---|---|
| cluster4053 | 1.54 | 1.000 |
| cluster3312 | 1.67 | 1.000 |
| cluster3212 | 1.50 | 1.000 |
| cluster2902 | -2.15 | 1.000 |
| cluster4079 | -1.61 | 1.000 |
| cluster4055 | 1.57 | 1.000 |
| cluster3434 | 1.33 | 1.000 |
| cluster4070 | 1.89 | 1.000 |
| cluster3009 | 1.41 | 1.000 |
| cluster3994 | 1.73 | 1.000 |
| cluster3317 | -1.71 | 1.000 |
| cluster3462 | 1.36 | 1.000 |
| cluster3448 | 1.55 | 1.000 |
| cluster3130 | 1.53 | 1.000 |
| cluster3075 | 1.26 | 1.000 |
| cluster4119 | -2.12 | 1.000 |
| cluster3981 | -1.66 | 1.000 |
| cluster3138 | -1.47 | 1.000 |
| cluster2993 | -1.50 | 1.000 |
| cluster3704 | -1.58 | 1.000 |
| cluster4074 | -1.97 | 1.000 |
| cluster3834 | 1.57 | 1.000 |
| cluster3745 | -2.18 | 1.000 |
| cluster3758 | 1.29 | 1.000 |
| cluster3126 | -1.67 | 1.000 |
| cluster3435 | 1.55 | 1.000 |
| cluster3741 | -1.44 | 1.000 |
| cluster3430 | -1.98 | 1.000 |
| cluster4078 | -1.45 | 1.000 |
| cluster3800 | 1.85 | 1.000 |
| cluster3568 | 1.60 | 1.000 |
| cluster3667 | 1.21 | 1.000 |
| cluster3865 | -1.87 | 1.000 |
| cluster2921 | -1.51 | 1.000 |
| cluster4095 | -1.58 | 1.000 |
| cluster3380 | 1.26 | 1.000 |
| cluster3597 | -1.37 | 1.000 |
| cluster2943 | -1.46 | 1.000 |
| cluster3636 | 1.17 | 1.000 |
| cluster3564 | -1.59 | 1.000 |
| cluster2967 | -1.49 | 1.000 |
| cluster3757 | -2.29 | 1.000 |
| cluster3894 | 1.39 | 1.000 |
| cluster3599 | 1.48 | 1.000 |

| | | |
|---|---|---|
| cluster2974 | 1.19 | 1.000 |
| cluster3357 | -1.33 | 1.000 |
| cluster4080 | 1.82 | 1.000 |
| cluster3844 | 1.55 | 1.000 |
| cluster3719 | -1.68 | 1.000 |
| cluster3437 | 1.15 | 1.000 |
| cluster3454 | 1.15 | 1.000 |
| cluster3978 | 1.24 | 1.000 |
| cluster3863 | -1.40 | 1.000 |
| cluster3995 | 1.43 | 1.000 |
| cluster3615 | -1.46 | 1.000 |
| cluster3656 | -1.29 | 1.000 |
| cluster3792 | 1.22 | 1.000 |
| cluster3898 | -1.49 | 1.000 |
| cluster3453 | 1.25 | 1.000 |
| cluster3822 | -1.80 | 1.000 |
| cluster3885 | 1.39 | 1.000 |
| cluster3053 | -1.45 | 1.000 |
| cluster3062 | 1.06 | 1.000 |
| cluster3200 | -1.28 | 1.000 |
| cluster3522 | 1.33 | 1.000 |
| cluster3931 | -1.65 | 1.000 |
| cluster3100 | -1.91 | 1.000 |
| cluster2947 | -1.23 | 1.000 |
| cluster4113 | 1.25 | 1.000 |
| cluster3827 | 1.18 | 1.000 |
| cluster3485 | 1.56 | 1.000 |
| cluster3015 | -1.54 | 1.000 |
| cluster2980 | 1.48 | 1.000 |
| cluster3329 | 1.35 | 1.000 |
| cluster3481 | 1.35 | 1.000 |
| cluster4049 | 2.13 | 1.000 |
| cluster3884 | 1.48 | 1.000 |
| cluster3118 | -1.23 | 1.000 |
| cluster3821 | 1.18 | 1.000 |
| cluster3237 | -1.88 | 1.000 |
| cluster2972 | 1.17 | 1.000 |
| cluster3085 | 1.28 | 1.000 |
| cluster2961 | -1.22 | 1.000 |
| cluster4034 | 2.25 | 1.000 |
| cluster3892 | 1.24 | 1.000 |
| cluster3335 | -1.28 | 1.000 |
| cluster3150 | 1.28 | 1.000 |
| cluster3105 | 1.62 | 1.000 |

| | | |
|---|---|---|
| cluster3901 | -1.85 | 1.000 |
| cluster3220 | 1.19 | 1.000 |
| cluster4016 | -1.39 | 1.000 |
| cluster3474 | -1.70 | 1.000 |
| cluster2962 | -1.26 | 1.000 |
| cluster3260 | 1.03 | 1.000 |
| cluster3026 | 1.27 | 1.000 |
| cluster4115 | 1.35 | 1.000 |
| cluster3101 | -1.16 | 1.000 |
| cluster3240 | -1.23 | 1.000 |
| cluster3705 | -1.31 | 1.000 |
| cluster3644 | 1.10 | 1.000 |
| cluster3551 | -1.28 | 1.000 |
| cluster3519 | 1.42 | 1.000 |
| cluster3876 | -1.20 | 1.000 |
| cluster3794 | 1.20 | 1.000 |
| cluster4056 | 1.34 | 1.000 |
| cluster4066 | 1.18 | 1.000 |
| cluster3887 | 1.18 | 1.000 |
| cluster3182 | 1.23 | 1.000 |
| cluster2911 | 1.08 | 1.000 |
| cluster4012 | -1.59 | 1.000 |
| cluster2881 | -1.25 | 1.000 |
| cluster3164 | 1.24 | 1.000 |
| cluster3245 | -1.65 | 1.000 |
| cluster3081 | 1.05 | 1.000 |
| cluster3102 | -1.24 | 1.000 |
| cluster3129 | 1.39 | 1.000 |
| cluster3082 | 1.32 | 1.000 |
| cluster3306 | -1.76 | 1.000 |
| cluster3787 | 1.26 | 1.000 |
| cluster3760 | -1.14 | 1.000 |
| cluster3195 | 0.96 | 1.000 |
| cluster3959 | 1.19 | 1.000 |
| cluster3436 | -1.60 | 1.000 |
| cluster3341 | 1.23 | 1.000 |
| cluster3168 | -1.13 | 1.000 |
| cluster4103 | -1.37 | 1.000 |
| cluster3659 | 1.28 | 1.000 |
| cluster3773 | 1.02 | 1.000 |
| cluster3695 | 1.07 | 1.000 |
| cluster4085 | 1.59 | 1.000 |
| cluster3398 | 1.11 | 1.000 |
| cluster2989 | -1.18 | 1.000 |

| | | |
|---|---|---|
| cluster3514 | -1.21 | 1.000 |
| cluster3752 | 1.37 | 1.000 |
| cluster3426 | -1.15 | 1.000 |
| cluster3175 | 1.23 | 1.000 |
| cluster2960 | -1.22 | 1.000 |
| cluster3940 | -1.13 | 1.000 |
| cluster3556 | -1.47 | 1.000 |
| cluster3161 | 1.22 | 1.000 |
| cluster3313 | 1.13 | 1.000 |
| cluster3999 | 1.09 | 1.000 |
| cluster2998 | 1.15 | 1.000 |
| cluster3960 | 1.18 | 1.000 |
| cluster3425 | 0.99 | 1.000 |
| cluster3753 | 1.66 | 1.000 |
| cluster4124 | 1.15 | 1.000 |
| cluster3716 | 1.08 | 1.000 |
| cluster3766 | 1.30 | 1.000 |
| cluster3842 | -1.15 | 1.000 |
| cluster3246 | -1.04 | 1.000 |
| cluster3165 | -1.47 | 1.000 |
| cluster3953 | 1.10 | 1.000 |
| cluster4002 | -1.44 | 1.000 |
| cluster3581 | 1.13 | 1.000 |
| cluster3459 | 1.12 | 1.000 |
| cluster3510 | 1.77 | 1.000 |
| cluster3595 | -1.08 | 1.000 |
| cluster3439 | 1.08 | 1.000 |
| cluster3203 | 1.07 | 1.000 |
| cluster2930 | -1.17 | 1.000 |
| cluster3746 | -1.08 | 1.000 |
| cluster3194 | 0.93 | 1.000 |
| cluster3395 | 0.90 | 1.000 |
| cluster3795 | 1.13 | 1.000 |
| cluster3850 | 1.30 | 1.000 |
| cluster3197 | -1.09 | 1.000 |
| cluster3853 | -1.46 | 1.000 |
| cluster3298 | 0.89 | 1.000 |
| cluster3508 | -1.37 | 1.000 |
| cluster4011 | 1.63 | 1.000 |
| cluster3907 | 1.09 | 1.000 |
| cluster4000 | -1.15 | 1.000 |
| cluster3400 | -1.00 | 1.000 |
| cluster3179 | -1.36 | 1.000 |
| cluster3097 | -1.08 | 1.000 |

| | | |
|---|---|---|
| cluster4127 | 1.27 | 1.000 |
| cluster4093 | -1.46 | 1.000 |
| cluster3180 | 1.06 | 1.000 |
| cluster3987 | -1.11 | 1.000 |
| cluster3413 | 1.10 | 1.000 |
| cluster3185 | 1.21 | 1.000 |
| cluster3375 | 0.95 | 1.000 |
| cluster3515 | -1.41 | 1.000 |
| cluster3633 | 0.96 | 1.000 |
| cluster3858 | -1.10 | 1.000 |
| cluster3609 | -1.45 | 1.000 |
| cluster3923 | 0.97 | 1.000 |
| cluster3919 | 1.54 | 1.000 |
| cluster2854 | -1.28 | 1.000 |
| cluster3452 | 1.03 | 1.000 |
| cluster3575 | -1.50 | 1.000 |
| cluster3954 | 1.05 | 1.000 |
| cluster3073 | 0.88 | 1.000 |
| cluster3332 | -1.28 | 1.000 |
| cluster3500 | 1.20 | 1.000 |
| cluster3480 | -1.26 | 1.000 |
| cluster3226 | 1.03 | 1.000 |
| cluster2899 | -1.10 | 1.000 |
| cluster3925 | 1.56 | 1.000 |
| cluster3869 | -1.34 | 1.000 |
| cluster3490 | 0.97 | 1.000 |
| cluster2969 | -1.24 | 1.000 |
| cluster3538 | -1.12 | 1.000 |
| cluster3127 | 0.84 | 1.000 |
| cluster3721 | -1.06 | 1.000 |
| cluster3338 | 0.97 | 1.000 |
| cluster4015 | 1.15 | 1.000 |
| cluster4099 | -1.26 | 1.000 |
| cluster3247 | -1.08 | 1.000 |
| cluster3756 | 0.95 | 1.000 |
| cluster3825 | 0.99 | 1.000 |
| cluster3765 | 0.85 | 1.000 |
| cluster3520 | -1.72 | 1.000 |
| cluster3337 | -0.99 | 1.000 |
| cluster3620 | 0.80 | 1.000 |
| cluster3799 | -0.96 | 1.000 |
| cluster2933 | -1.01 | 1.000 |
| cluster3445 | 0.77 | 1.000 |
| cluster3868 | 0.94 | 1.000 |

| | | |
|---|---|---|
| cluster3854 | 0.93 | 1.000 |
| cluster3052 | 1.06 | 1.000 |
| cluster2914 | 0.93 | 1.000 |
| cluster3142 | -1.11 | 1.000 |
| cluster3764 | 1.00 | 1.000 |
| cluster3446 | 0.96 | 1.000 |
| cluster3691 | -1.08 | 1.000 |
| cluster3544 | 1.05 | 1.000 |
| cluster3320 | 0.95 | 1.000 |
| cluster2871 | 0.77 | 1.000 |
| cluster3494 | -0.93 | 1.000 |
| cluster3377 | 0.93 | 1.000 |
| cluster3769 | 0.75 | 1.000 |
| cluster3135 | 1.03 | 1.000 |
| cluster3032 | -0.89 | 1.000 |
| cluster2984 | -0.99 | 1.000 |
| cluster4035 | -0.93 | 1.000 |
| cluster3023 | -0.93 | 1.000 |
| cluster3008 | -1.14 | 1.000 |
| cluster3061 | -0.89 | 1.000 |
| cluster3191 | 0.98 | 1.000 |
| cluster3662 | 0.90 | 1.000 |
| cluster3770 | 1.13 | 1.000 |
| cluster2995 | -1.07 | 1.000 |
| cluster3205 | 0.74 | 1.000 |
| cluster2999 | 0.95 | 1.000 |
| cluster3731 | -0.95 | 1.000 |
| cluster2956 | 0.75 | 1.000 |
| cluster4062 | 1.08 | 1.000 |
| cluster3473 | -0.94 | 1.000 |
| cluster3922 | 0.99 | 1.000 |
| cluster2860 | -0.96 | 1.000 |
| cluster4097 | 1.40 | 1.000 |
| cluster3567 | -1.29 | 1.000 |
| cluster3487 | 0.99 | 1.000 |
| cluster3311 | -0.89 | 1.000 |
| cluster3629 | -1.21 | 1.000 |
| cluster2901 | 0.93 | 1.000 |
| cluster3675 | 0.73 | 1.000 |
| cluster3578 | 0.88 | 1.000 |
| cluster4128 | 1.03 | 1.000 |
| cluster3573 | 0.91 | 1.000 |
| cluster3730 | -0.88 | 1.000 |
| cluster3057 | 0.83 | 1.000 |

| | | |
|---|---|---|
| cluster3068 | 0.68 | 1.000 |
| cluster3244 | -0.83 | 1.000 |
| cluster3292 | 1.05 | 1.000 |
| cluster3431 | -1.14 | 1.000 |
| cluster2857 | 0.77 | 1.000 |
| cluster3438 | -1.12 | 1.000 |
| cluster3051 | 0.89 | 1.000 |
| cluster3742 | -1.14 | 1.000 |
| cluster3296 | 1.02 | 1.000 |
| cluster3905 | 1.17 | 1.000 |
| cluster3749 | 0.96 | 1.000 |
| cluster3983 | -1.18 | 1.000 |
| cluster3918 | -1.12 | 1.000 |
| cluster2915 | 0.82 | 1.000 |
| cluster3224 | 0.74 | 1.000 |
| cluster3110 | -0.91 | 1.000 |
| cluster3066 | -0.88 | 1.000 |
| cluster3593 | 0.86 | 1.000 |
| cluster3325 | 1.00 | 1.000 |
| cluster3088 | -0.94 | 1.000 |
| cluster2903 | 0.78 | 1.000 |
| cluster4023 | 0.86 | 1.000 |
| cluster3929 | 0.90 | 1.000 |
| cluster3580 | -1.11 | 1.000 |
| cluster3047 | 0.83 | 1.000 |
| cluster2912 | -0.96 | 1.000 |
| cluster4059 | 0.92 | 1.000 |
| cluster3565 | 0.84 | 1.000 |
| cluster2939 | 0.72 | 1.000 |
| cluster3941 | 0.65 | 1.000 |
| cluster3990 | -0.82 | 1.000 |
| cluster4064 | -1.07 | 1.000 |
| cluster3937 | -0.91 | 1.000 |
| cluster4047 | -1.23 | 1.000 |
| cluster3964 | 0.92 | 1.000 |
| cluster3156 | 0.85 | 1.000 |
| cluster3661 | -0.81 | 1.000 |
| cluster3830 | -0.87 | 1.000 |
| cluster3379 | 0.78 | 1.000 |
| cluster4019 | -1.30 | 1.000 |
| cluster3225 | 0.81 | 1.000 |
| cluster3649 | 0.69 | 1.000 |
| cluster3188 | -0.89 | 1.000 |
| cluster2892 | -0.80 | 1.000 |

| | | |
|---|---|---|
| cluster3973 | 0.98 | 1.000 |
| cluster3912 | 0.63 | 1.000 |
| cluster4026 | 0.81 | 1.000 |
| cluster3819 | 0.92 | 1.000 |
| cluster3119 | -0.83 | 1.000 |
| cluster3261 | 0.81 | 1.000 |
| cluster3027 | -0.78 | 1.000 |
| cluster3392 | 0.83 | 1.000 |
| cluster3928 | 0.80 | 1.000 |
| cluster4121 | -1.12 | 1.000 |
| cluster3608 | 0.69 | 1.000 |
| cluster3867 | 0.69 | 1.000 |
| cluster3714 | -0.89 | 1.000 |
| cluster4106 | 0.93 | 1.000 |
| cluster3365 | -1.09 | 1.000 |
| cluster3970 | -1.12 | 1.000 |
| cluster3820 | 0.79 | 1.000 |
| cluster2926 | -0.77 | 1.000 |
| cluster4005 | 0.79 | 1.000 |
| cluster3777 | 0.72 | 1.000 |
| cluster3531 | 0.82 | 1.000 |
| cluster3621 | -1.10 | 1.000 |
| cluster2894 | 0.82 | 1.000 |
| cluster3984 | 0.89 | 1.000 |
| cluster3319 | 0.82 | 1.000 |
| cluster3744 | 0.79 | 1.000 |
| cluster3363 | -1.02 | 1.000 |
| cluster3679 | -0.82 | 1.000 |
| cluster3768 | 0.83 | 1.000 |
| cluster3598 | -1.25 | 1.000 |
| cluster3554 | 0.78 | 1.000 |
| cluster3019 | -0.99 | 1.000 |
| cluster3458 | -0.76 | 1.000 |
| cluster3111 | 0.81 | 1.000 |
| cluster3484 | 0.90 | 1.000 |
| cluster2992 | 0.64 | 1.000 |
| cluster4086 | -0.98 | 1.000 |
| cluster3277 | 0.72 | 1.000 |
| cluster2976 | 0.74 | 1.000 |
| cluster2937 | 0.67 | 1.000 |
| cluster3013 | -0.78 | 1.000 |
| cluster3948 | 0.80 | 1.000 |
| cluster4107 | -1.07 | 1.000 |
| cluster3513 | -0.78 | 1.000 |

| | | |
|---|---|---|
| cluster3187 | 0.83 | 1.000 |
| cluster3399 | -0.85 | 1.000 |
| cluster4084 | 0.70 | 1.000 |
| cluster2882 | -0.71 | 1.000 |
| cluster3577 | -0.76 | 1.000 |
| cluster3016 | 0.80 | 1.000 |
| cluster3234 | 0.66 | 1.000 |
| cluster3304 | -0.79 | 1.000 |
| cluster3976 | 0.66 | 1.000 |
| cluster3249 | 0.67 | 1.000 |
| cluster3488 | 0.74 | 1.000 |
| cluster2873 | -0.74 | 1.000 |
| cluster2946 | -0.83 | 1.000 |
| cluster3553 | 0.72 | 1.000 |
| cluster3537 | -0.97 | 1.000 |
| cluster3361 | 0.78 | 1.000 |
| cluster3493 | -1.07 | 1.000 |
| cluster3117 | -0.99 | 1.000 |
| cluster2942 | -0.93 | 1.000 |
| cluster3255 | 0.80 | 1.000 |
| cluster3017 | -0.96 | 1.000 |
| cluster3089 | -0.81 | 1.000 |
| cluster3775 | 0.76 | 1.000 |
| cluster2924 | -1.06 | 1.000 |
| cluster3549 | -0.75 | 1.000 |
| cluster4042 | 0.77 | 1.000 |
| cluster3540 | 0.73 | 1.000 |
| cluster2957 | 0.75 | 1.000 |
| cluster3131 | 0.79 | 1.000 |
| cluster3331 | 0.71 | 1.000 |
| cluster3402 | -0.73 | 1.000 |
| cluster3611 | -0.96 | 1.000 |
| cluster3849 | -0.94 | 1.000 |
| cluster3650 | -0.90 | 1.000 |
| cluster3877 | 0.58 | 1.000 |
| cluster3511 | 0.77 | 1.000 |
| cluster3269 | 0.72 | 1.000 |
| cluster3963 | 0.74 | 1.000 |
| cluster3899 | -0.93 | 1.000 |
| cluster3092 | 0.63 | 1.000 |
| cluster3160 | 0.62 | 1.000 |
| cluster3857 | 0.68 | 1.000 |
| cluster3162 | -0.88 | 1.000 |
| cluster3288 | -0.95 | 1.000 |

| | | |
|---|---|---|
| cluster3034 | 0.66 | 1.000 |
| cluster3360 | 0.65 | 1.000 |
| cluster3310 | 0.57 | 1.000 |
| cluster3345 | 0.68 | 1.000 |
| cluster4094 | -0.86 | 1.000 |
| cluster3826 | 0.67 | 1.000 |
| cluster3141 | -0.67 | 1.000 |
| cluster3640 | 0.91 | 1.000 |
| cluster3369 | -0.93 | 1.000 |
| cluster3674 | 0.81 | 1.000 |
| cluster3450 | 0.82 | 1.000 |
| cluster3665 | 0.70 | 1.000 |
| cluster3669 | 0.72 | 1.000 |
| cluster3712 | -0.74 | 1.000 |
| cluster3972 | 0.68 | 1.000 |
| cluster3259 | 0.55 | 1.000 |
| cluster3709 | -0.69 | 1.000 |
| cluster3518 | 0.74 | 1.000 |
| cluster3985 | -0.74 | 1.000 |
| cluster3806 | 0.67 | 1.000 |
| cluster3582 | -0.68 | 1.000 |
| cluster3427 | 0.65 | 1.000 |
| cluster4088 | 0.71 | 1.000 |
| cluster3729 | 0.68 | 1.000 |
| cluster2977 | -0.86 | 1.000 |
| cluster3654 | 0.62 | 1.000 |
| cluster3993 | 0.64 | 1.000 |
| cluster3144 | 0.70 | 1.000 |
| cluster3942 | 0.67 | 1.000 |
| cluster3747 | -0.76 | 1.000 |
| cluster3410 | -0.75 | 1.000 |
| cluster3703 | -0.64 | 1.000 |
| cluster3084 | 0.51 | 1.000 |
| cluster3516 | 0.73 | 1.000 |
| cluster3779 | -0.70 | 1.000 |
| cluster4054 | -0.87 | 1.000 |
| cluster3080 | 0.58 | 1.000 |
| cluster3591 | 0.57 | 1.000 |
| cluster3988 | 0.55 | 1.000 |
| cluster4091 | 0.77 | 1.000 |
| cluster3122 | 0.55 | 1.000 |
| cluster3145 | 0.52 | 1.000 |
| cluster3157 | -0.63 | 1.000 |
| cluster2963 | -0.74 | 1.000 |

| | | |
|---|---|---|
| cluster3040 | 0.52 | 1.000 |
| cluster3883 | -1.06 | 1.000 |
| cluster3682 | 0.64 | 1.000 |
| cluster4075 | -1.01 | 1.000 |
| cluster3751 | 0.59 | 1.000 |
| cluster2906 | -0.67 | 1.000 |
| cluster3823 | 0.52 | 1.000 |
| cluster3788 | -0.72 | 1.000 |
| cluster3603 | -0.84 | 1.000 |
| cluster3033 | 0.54 | 1.000 |
| cluster3442 | -0.89 | 1.000 |
| cluster3043 | 0.60 | 1.000 |
| cluster2934 | 0.51 | 1.000 |
| cluster3771 | -0.83 | 1.000 |
| cluster3627 | 0.56 | 1.000 |
| cluster3401 | -0.77 | 1.000 |
| cluster3478 | -0.64 | 1.000 |
| cluster3466 | 0.56 | 1.000 |
| cluster3845 | 0.81 | 1.000 |
| cluster3952 | -0.86 | 1.000 |
| cluster3001 | 0.47 | 1.000 |
| cluster3356 | -0.64 | 1.000 |
| cluster3022 | 0.59 | 1.000 |
| cluster3218 | -0.85 | 1.000 |
| cluster2940 | 0.63 | 1.000 |
| cluster3093 | 0.59 | 1.000 |
| cluster3572 | -0.99 | 1.000 |
| cluster3817 | -0.74 | 1.000 |
| cluster3340 | -0.61 | 1.000 |
| cluster3469 | -0.67 | 1.000 |
| cluster3366 | -0.81 | 1.000 |
| cluster3083 | -0.61 | 1.000 |
| cluster3344 | 0.53 | 1.000 |
| cluster2916 | -0.66 | 1.000 |
| cluster3810 | -0.61 | 1.000 |
| cluster3124 | 0.51 | 1.000 |
| cluster3134 | 0.47 | 1.000 |
| cluster3181 | -0.62 | 1.000 |
| cluster3497 | -0.60 | 1.000 |
| cluster3408 | -0.67 | 1.000 |
| cluster3732 | 0.58 | 1.000 |
| cluster4018 | -0.74 | 1.000 |
| cluster4022 | 0.67 | 1.000 |
| cluster3424 | 0.64 | 1.000 |

| | | |
|---|---|---|
| cluster3441 | -0.65 | 1.000 |
| cluster3596 | 0.55 | 1.000 |
| cluster3299 | 0.58 | 1.000 |
| cluster3468 | -0.61 | 1.000 |
| cluster3125 | 0.51 | 1.000 |
| cluster3676 | -0.80 | 1.000 |
| cluster3943 | -0.78 | 1.000 |
| cluster3132 | -0.57 | 1.000 |
| cluster3385 | 0.62 | 1.000 |
| cluster3670 | 0.55 | 1.000 |
| cluster3336 | 0.51 | 1.000 |
| cluster3841 | 0.58 | 1.000 |
| cluster3411 | 0.57 | 1.000 |
| cluster3174 | -0.82 | 1.000 |
| cluster3143 | -0.58 | 1.000 |
| cluster2863 | -0.56 | 1.000 |
| cluster2994 | -0.64 | 1.000 |
| cluster3873 | -0.55 | 1.000 |
| cluster3536 | -0.57 | 1.000 |
| cluster3782 | 0.51 | 1.000 |
| cluster3673 | 0.58 | 1.000 |
| cluster3451 | -1.06 | 1.000 |
| cluster3346 | -0.64 | 1.000 |
| cluster3308 | 0.45 | 1.000 |
| cluster3989 | -0.62 | 1.000 |
| cluster2964 | -0.56 | 1.000 |
| cluster3871 | -0.75 | 1.000 |
| cluster2891 | 0.58 | 1.000 |
| cluster4063 | 0.60 | 1.000 |
| cluster3382 | 0.48 | 1.000 |
| cluster3460 | -0.73 | 1.000 |
| cluster3010 | 0.50 | 1.000 |
| cluster3736 | 0.55 | 1.000 |
| cluster3643 | -0.73 | 1.000 |
| cluster2877 | -0.55 | 1.000 |
| cluster3151 | 0.45 | 1.000 |
| cluster3470 | 0.45 | 1.000 |
| cluster3064 | -0.65 | 1.000 |
| cluster3977 | 0.59 | 1.000 |
| cluster3229 | -0.63 | 1.000 |
| cluster3759 | 0.46 | 1.000 |
| cluster3690 | 0.45 | 1.000 |
| cluster3455 | 0.49 | 1.000 |
| cluster3219 | 0.48 | 1.000 |

| | | |
|---|---|---|
| cluster3734 | -0.77 | 1.000 |
| cluster3552 | -0.59 | 1.000 |
| cluster3706 | 0.52 | 1.000 |
| cluster3969 | 0.41 | 1.000 |
| cluster3594 | 0.55 | 1.000 |
| cluster2971 | 0.43 | 1.000 |
| cluster3011 | -0.62 | 1.000 |
| cluster2898 | -0.61 | 1.000 |
| cluster3902 | -0.77 | 1.000 |
| cluster3776 | 0.47 | 1.000 |
| cluster3282 | 0.50 | 1.000 |
| cluster3906 | 0.51 | 1.000 |
| cluster3829 | -0.72 | 1.000 |
| cluster2859 | 0.43 | 1.000 |
| cluster3235 | -0.78 | 1.000 |
| cluster3268 | 0.53 | 1.000 |
| cluster2888 | 0.56 | 1.000 |
| cluster2958 | 0.44 | 1.000 |
| cluster3711 | 0.46 | 1.000 |
| cluster4109 | -0.76 | 1.000 |
| cluster3005 | -0.59 | 1.000 |
| cluster3557 | 0.50 | 1.000 |
| cluster3386 | 0.55 | 1.000 |
| cluster2876 | -0.85 | 1.000 |
| cluster3114 | 0.45 | 1.000 |
| cluster2928 | -0.66 | 1.000 |
| cluster3517 | 0.42 | 1.000 |
| cluster3417 | -0.59 | 1.000 |
| cluster3781 | -0.59 | 1.000 |
| cluster3264 | 0.45 | 1.000 |
| cluster3104 | 0.39 | 1.000 |
| cluster3715 | -0.67 | 1.000 |
| cluster3693 | -0.74 | 1.000 |
| cluster3414 | -0.54 | 1.000 |
| cluster4058 | 0.50 | 1.000 |
| cluster4051 | -0.83 | 1.000 |
| cluster3717 | 0.42 | 1.000 |
| cluster3232 | -0.72 | 1.000 |
| cluster3872 | -0.71 | 1.000 |
| cluster3623 | 0.51 | 1.000 |
| cluster3479 | 0.50 | 1.000 |
| cluster3316 | 0.50 | 1.000 |
| cluster2965 | -0.75 | 1.000 |
| cluster2865 | -0.53 | 1.000 |

| | | |
|---|---|---|
| cluster3094 | -0.71 | 1.000 |
| cluster3209 | 0.51 | 1.000 |
| cluster3653 | -0.56 | 1.000 |
| cluster3457 | 0.54 | 1.000 |
| cluster3271 | 0.39 | 1.000 |
| cluster2919 | -0.53 | 1.000 |
| cluster3349 | -0.62 | 1.000 |
| cluster3196 | 0.39 | 1.000 |
| cluster3908 | 0.46 | 1.000 |
| cluster3133 | 0.48 | 1.000 |
| cluster2936 | -0.72 | 1.000 |
| cluster3048 | 0.49 | 1.000 |
| cluster3930 | -0.54 | 1.000 |
| cluster2938 | 0.38 | 1.000 |
| cluster3421 | -0.70 | 1.000 |
| cluster3874 | 0.40 | 1.000 |
| cluster3091 | 0.40 | 1.000 |
| cluster2978 | -0.56 | 1.000 |
| cluster3103 | 0.52 | 1.000 |
| cluster3394 | 0.39 | 1.000 |
| cluster3387 | 0.50 | 1.000 |
| cluster3655 | 0.46 | 1.000 |
| cluster3146 | 0.40 | 1.000 |
| cluster3778 | -0.58 | 1.000 |
| cluster4082 | -0.71 | 1.000 |
| cluster2908 | -0.53 | 1.000 |
| cluster3301 | 0.40 | 1.000 |
| cluster3443 | -0.64 | 1.000 |
| cluster3566 | 0.44 | 1.000 |
| cluster3956 | -0.66 | 1.000 |
| cluster3014 | 0.47 | 1.000 |
| cluster2975 | -0.49 | 1.000 |
| cluster3911 | -0.66 | 1.000 |
| cluster2853 | -0.56 | 1.000 |
| cluster3036 | 0.45 | 1.000 |
| cluster2870 | -0.54 | 1.000 |
| cluster3449 | -0.48 | 1.000 |
| cluster3254 | 0.49 | 1.000 |
| cluster3183 | 0.40 | 1.000 |
| cluster3327 | -0.51 | 1.000 |
| cluster3076 | -0.64 | 1.000 |
| cluster4043 | 0.45 | 1.000 |
| cluster2949 | -0.45 | 1.000 |
| cluster3396 | 0.37 | 1.000 |

| | | |
|---|---|---|
| cluster3099 | -0.47 | 1.000 |
| cluster3406 | 0.35 | 1.000 |
| cluster3172 | 0.37 | 1.000 |
| cluster3403 | 0.45 | 1.000 |
| cluster3541 | -0.47 | 1.000 |
| cluster3831 | 0.41 | 1.000 |
| cluster3979 | -0.46 | 1.000 |
| cluster3824 | -0.66 | 1.000 |
| cluster3856 | 0.44 | 1.000 |
| cluster3300 | -0.50 | 1.000 |
| cluster3463 | -0.63 | 1.000 |
| cluster2867 | 0.44 | 1.000 |
| cluster3671 | 0.43 | 1.000 |
| cluster3521 | 0.41 | 1.000 |
| cluster3677 | -0.48 | 1.000 |
| cluster3285 | 0.35 | 1.000 |
| cluster3368 | 0.33 | 1.000 |
| cluster2932 | 0.35 | 1.000 |
| cluster3222 | 0.31 | 1.000 |
| cluster3050 | 0.35 | 1.000 |
| cluster3207 | -0.49 | 1.000 |
| cluster3807 | 0.35 | 1.000 |
| cluster3280 | -0.50 | 1.000 |
| cluster3997 | -0.59 | 1.000 |
| cluster2927 | -0.60 | 1.000 |
| cluster4089 | -0.80 | 1.000 |
| cluster3383 | 0.44 | 1.000 |
| cluster2941 | 0.33 | 1.000 |
| cluster3613 | -0.50 | 1.000 |
| cluster2981 | -0.61 | 1.000 |
| cluster3242 | -0.51 | 1.000 |
| cluster3971 | -0.60 | 1.000 |
| cluster3660 | 0.40 | 1.000 |
| cluster3527 | 0.38 | 1.000 |
| cluster3949 | 0.40 | 1.000 |
| cluster4125 | -0.69 | 1.000 |
| cluster3815 | -0.61 | 1.000 |
| cluster3217 | 0.34 | 1.000 |
| cluster3495 | -0.62 | 1.000 |
| cluster4117 | 0.49 | 1.000 |
| cluster3440 | 0.32 | 1.000 |
| cluster2884 | -0.51 | 1.000 |
| cluster3619 | -0.47 | 1.000 |
| cluster3726 | 0.33 | 1.000 |

| | | |
|---|---|---|
| cluster3548 | -0.42 | 1.000 |
| cluster3024 | 0.33 | 1.000 |
| cluster3796 | 0.40 | 1.000 |
| cluster3748 | -0.44 | 1.000 |
| cluster3042 | 0.38 | 1.000 |
| cluster3805 | -0.60 | 1.000 |
| cluster3913 | 0.40 | 1.000 |
| cluster4077 | 0.39 | 1.000 |
| cluster4108 | 0.41 | 1.000 |
| cluster3267 | -0.42 | 1.000 |
| cluster3253 | 0.37 | 1.000 |
| cluster4030 | 0.40 | 1.000 |
| cluster3848 | 0.40 | 1.000 |
| cluster3390 | 0.31 | 1.000 |
| cluster3307 | 0.38 | 1.000 |
| cluster3861 | 0.36 | 1.000 |
| cluster3498 | -0.56 | 1.000 |
| cluster3740 | 0.36 | 1.000 |
| cluster3743 | 0.36 | 1.000 |
| cluster3098 | 0.36 | 1.000 |
| cluster3936 | -0.55 | 1.000 |
| cluster3622 | 0.35 | 1.000 |
| cluster3071 | -0.41 | 1.000 |
| cluster3663 | 0.35 | 1.000 |
| cluster3616 | 0.35 | 1.000 |
| cluster3530 | -0.51 | 1.000 |
| cluster3186 | 0.38 | 1.000 |
| cluster2890 | 0.29 | 1.000 |
| cluster3767 | 0.35 | 1.000 |
| cluster4020 | 0.29 | 1.000 |
| cluster3433 | -0.44 | 1.000 |
| cluster3789 | 0.33 | 1.000 |
| cluster3391 | 0.27 | 1.000 |
| cluster3843 | 0.30 | 1.000 |
| cluster3359 | 0.35 | 1.000 |
| cluster3584 | -0.54 | 1.000 |
| cluster3231 | 0.39 | 1.000 |
| cluster3509 | -0.41 | 1.000 |
| cluster3471 | -0.57 | 1.000 |
| cluster3657 | -0.55 | 1.000 |
| cluster3004 | -0.41 | 1.000 |
| cluster3926 | 0.39 | 1.000 |
| cluster3166 | -0.38 | 1.000 |
| cluster3070 | -0.40 | 1.000 |

| | | |
|---|---|---|
| cluster2913 | 0.29 | 1.000 |
| cluster4009 | -0.50 | 1.000 |
| cluster2931 | 0.33 | 1.000 |
| cluster3376 | 0.27 | 1.000 |
| cluster3238 | -0.38 | 1.000 |
| cluster3113 | -0.44 | 1.000 |
| cluster3419 | 0.27 | 1.000 |
| cluster3305 | 0.34 | 1.000 |
| cluster4092 | 0.34 | 1.000 |
| cluster3199 | 0.29 | 1.000 |
| cluster3638 | -0.54 | 1.000 |
| cluster3273 | -0.37 | 1.000 |
| cluster3415 | 0.29 | 1.000 |
| cluster3790 | 0.32 | 1.000 |
| cluster3724 | -0.52 | 1.000 |
| cluster3364 | 0.33 | 1.000 |
| cluster3239 | -0.36 | 1.000 |
| cluster3881 | -0.50 | 1.000 |
| cluster3049 | -0.40 | 1.000 |
| cluster3223 | 0.30 | 1.000 |
| cluster3272 | 0.39 | 1.000 |
| cluster3190 | -0.54 | 1.000 |
| cluster2935 | -0.41 | 1.000 |
| cluster3784 | -0.40 | 1.000 |
| cluster4006 | 0.32 | 1.000 |
| cluster3685 | 0.33 | 1.000 |
| cluster3373 | -0.50 | 1.000 |
| cluster3405 | -0.42 | 1.000 |
| cluster4025 | 0.27 | 1.000 |
| cluster3804 | -0.54 | 1.000 |
| cluster3832 | -0.51 | 1.000 |
| cluster3914 | 0.35 | 1.000 |
| cluster3206 | 0.30 | 1.000 |
| cluster2987 | -0.37 | 1.000 |
| cluster3334 | -0.40 | 1.000 |
| cluster3889 | -0.52 | 1.000 |
| cluster3278 | 0.26 | 1.000 |
| cluster3159 | 0.30 | 1.000 |
| cluster2910 | 0.31 | 1.000 |
| cluster3328 | 0.25 | 1.000 |
| cluster3496 | -0.38 | 1.000 |
| cluster4004 | 0.32 | 1.000 |
| cluster2945 | 0.33 | 1.000 |
| cluster3880 | -0.62 | 1.000 |

| | | |
|---|---|---|
| cluster3275 | 0.31 | 1.000 |
| cluster3374 | -0.51 | 1.000 |
| cluster3668 | 0.30 | 1.000 |
| cluster3112 | 0.31 | 1.000 |
| cluster2878 | -0.38 | 1.000 |
| cluster3077 | -0.42 | 1.000 |
| cluster3754 | 0.26 | 1.000 |
| cluster3302 | 0.28 | 1.000 |
| cluster3258 | -0.40 | 1.000 |
| cluster3586 | -0.35 | 1.000 |
| cluster3728 | 0.27 | 1.000 |
| cluster3733 | 0.26 | 1.000 |
| cluster3149 | -0.35 | 1.000 |
| cluster3483 | 0.31 | 1.000 |
| cluster3465 | -0.38 | 1.000 |
| cluster3204 | -0.39 | 1.000 |
| cluster3579 | 0.24 | 1.000 |
| cluster3678 | -0.36 | 1.000 |
| cluster2905 | -0.38 | 1.000 |
| cluster3318 | -0.39 | 1.000 |
| cluster3991 | 0.28 | 1.000 |
| cluster2879 | -0.40 | 1.000 |
| cluster3532 | 0.27 | 1.000 |
| cluster3257 | -0.35 | 1.000 |
| cluster3562 | 0.28 | 1.000 |
| cluster3836 | 0.29 | 1.000 |
| cluster3837 | 0.29 | 1.000 |
| cluster3583 | 0.27 | 1.000 |
| cluster2982 | -0.40 | 1.000 |
| cluster4044 | -0.47 | 1.000 |
| cluster2862 | 0.23 | 1.000 |
| cluster3888 | 0.27 | 1.000 |
| cluster3847 | -0.50 | 1.000 |
| cluster3718 | 0.27 | 1.000 |
| cluster2997 | 0.28 | 1.000 |
| cluster3909 | 0.27 | 1.000 |
| cluster3528 | 0.25 | 1.000 |
| cluster4123 | 0.29 | 1.000 |
| cluster3539 | 0.28 | 1.000 |
| cluster3321 | -0.42 | 1.000 |
| cluster3699 | -0.45 | 1.000 |
| cluster3916 | -0.44 | 1.000 |
| cluster3524 | 0.24 | 1.000 |
| cluster3992 | -0.37 | 1.000 |

| | | |
|---|---|---|
| cluster4029 | 0.29 | 1.000 |
| cluster3074 | 0.28 | 1.000 |
| cluster3058 | 0.21 | 1.000 |
| cluster3167 | 0.26 | 1.000 |
| cluster3982 | 0.26 | 1.000 |
| cluster3641 | 0.22 | 1.000 |
| cluster3664 | 0.23 | 1.000 |
| cluster3798 | 0.25 | 1.000 |
| cluster2991 | 0.25 | 1.000 |
| cluster3291 | -0.35 | 1.000 |
| cluster3355 | 0.27 | 1.000 |
| cluster3903 | 0.26 | 1.000 |
| cluster2864 | -0.33 | 1.000 |
| cluster3351 | 0.23 | 1.000 |
| cluster4096 | 0.25 | 1.000 |
| cluster3213 | 0.22 | 1.000 |
| cluster3630 | 0.20 | 1.000 |
| cluster4028 | -0.43 | 1.000 |
| cluster3276 | -0.31 | 1.000 |
| cluster3859 | 0.26 | 1.000 |
| cluster3780 | 0.29 | 1.000 |
| cluster3791 | -0.45 | 1.000 |
| cluster3809 | 0.24 | 1.000 |
| cluster3477 | 0.23 | 1.000 |
| cluster3606 | 0.24 | 1.000 |
| cluster3855 | 0.23 | 1.000 |
| cluster3262 | 0.22 | 1.000 |
| cluster2944 | 0.23 | 1.000 |
| cluster3228 | -0.43 | 1.000 |
| cluster3910 | 0.24 | 1.000 |
| cluster3330 | -0.36 | 1.000 |
| cluster3946 | 0.23 | 1.000 |
| cluster4102 | 0.22 | 1.000 |
| cluster3107 | -0.32 | 1.000 |
| cluster3547 | -0.32 | 1.000 |
| cluster3270 | -0.45 | 1.000 |
| cluster3358 | -0.32 | 1.000 |
| cluster3507 | 0.23 | 1.000 |
| cluster3037 | 0.21 | 1.000 |
| cluster3054 | 0.20 | 1.000 |
| cluster3069 | 0.19 | 1.000 |
| cluster3713 | -0.39 | 1.000 |
| cluster3762 | -0.35 | 1.000 |
| cluster3163 | 0.23 | 1.000 |

| | | |
|---|---|---|
| cluster3720 | 0.22 | 1.000 |
| cluster3140 | -0.30 | 1.000 |
| cluster3352 | -0.41 | 1.000 |
| cluster3701 | 0.19 | 1.000 |
| cluster3155 | -0.40 | 1.000 |
| cluster3348 | -0.32 | 1.000 |
| cluster3604 | -0.43 | 1.000 |
| cluster3115 | -0.42 | 1.000 |
| cluster2895 | 0.22 | 1.000 |
| cluster3039 | -0.29 | 1.000 |
| cluster4041 | 0.23 | 1.000 |
| cluster2950 | -0.29 | 1.000 |
| cluster2966 | -0.42 | 1.000 |
| cluster3056 | 0.18 | 1.000 |
| cluster3389 | 0.21 | 1.000 |
| cluster4061 | -0.38 | 1.000 |
| cluster3835 | 0.21 | 1.000 |
| cluster2922 | -0.40 | 1.000 |
| cluster3600 | -0.34 | 1.000 |
| cluster3875 | -0.34 | 1.000 |
| cluster3801 | -0.39 | 1.000 |
| cluster3938 | 0.20 | 1.000 |
| cluster3761 | -0.47 | 1.000 |
| cluster3147 | -0.32 | 1.000 |
| cluster3813 | -0.38 | 1.000 |
| cluster3626 | 0.18 | 1.000 |
| cluster3243 | -0.31 | 1.000 |
| cluster1127 | -0.40 | 1.000 |
| cluster4110 | -0.38 | 1.000 |
| cluster3523 | 0.20 | 1.000 |
| cluster3060 | 0.20 | 1.000 |
| cluster3153 | -0.38 | 1.000 |
| cluster3198 | 0.20 | 1.000 |
| cluster3342 | -0.39 | 1.000 |
| cluster3811 | -0.27 | 1.000 |
| cluster3846 | 0.16 | 1.000 |
| cluster4068 | 0.20 | 1.000 |
| cluster3029 | 0.18 | 1.000 |
| cluster3221 | -0.38 | 1.000 |
| cluster3251 | 0.19 | 1.000 |
| cluster3021 | -0.26 | 1.000 |
| cluster2866 | -0.42 | 1.000 |
| cluster3215 | -0.26 | 1.000 |
| cluster4045 | -0.43 | 1.000 |

| | | |
|---|---|---|
| cluster3227 | -0.35 | 1.000 |
| cluster3658 | -0.31 | 1.000 |
| cluster2952 | 0.18 | 1.000 |
| cluster3891 | -0.33 | 1.000 |
| cluster3610 | 0.19 | 1.000 |
| cluster3708 | 0.21 | 1.000 |
| cluster3123 | 0.15 | 1.000 |
| cluster3968 | -0.34 | 1.000 |
| cluster3086 | 0.16 | 1.000 |
| cluster3169 | -0.37 | 1.000 |
| cluster3555 | 0.18 | 1.000 |
| cluster3543 | 0.18 | 1.000 |
| cluster3864 | 0.17 | 1.000 |
| cluster3447 | 0.17 | 1.000 |
| cluster3812 | -0.36 | 1.000 |
| cluster3625 | 0.17 | 1.000 |
| cluster3444 | 0.15 | 1.000 |
| cluster2904 | -0.29 | 1.000 |
| cluster4083 | -0.36 | 1.000 |
| cluster3252 | -0.45 | 1.000 |
| cluster3879 | -0.33 | 1.000 |
| cluster3870 | 0.17 | 1.000 |
| cluster4050 | 0.16 | 1.000 |
| cluster3602 | -0.25 | 1.000 |
| cluster2996 | 0.14 | 1.000 |
| cluster4071 | 0.16 | 1.000 |
| cluster3634 | 0.13 | 1.000 |
| cluster3833 | 0.15 | 1.000 |
| cluster2954 | 0.14 | 1.000 |
| cluster3350 | 0.15 | 1.000 |
| cluster3696 | 0.14 | 1.000 |
| cluster3353 | -0.27 | 1.000 |
| cluster4010 | -0.33 | 1.000 |
| cluster3986 | 0.14 | 1.000 |
| cluster3816 | 0.15 | 1.000 |
| cluster3293 | -0.24 | 1.000 |
| cluster3476 | -0.33 | 1.000 |
| cluster2920 | 0.14 | 1.000 |
| cluster4100 | -0.33 | 1.000 |
| cluster3214 | 0.13 | 1.000 |
| cluster2907 | 0.15 | 1.000 |
| cluster3499 | 0.14 | 1.000 |
| cluster3020 | 0.15 | 1.000 |
| cluster3121 | -0.27 | 1.000 |

| | | |
|---|---|---|
| cluster3980 | -0.22 | 1.000 |
| cluster4021 | -0.33 | 1.000 |
| cluster4081 | -0.32 | 1.000 |
| cluster2896 | 0.14 | 1.000 |
| cluster3038 | 0.14 | 1.000 |
| cluster3723 | -0.24 | 1.000 |
| cluster3512 | 0.13 | 1.000 |
| cluster3007 | -0.31 | 1.000 |
| cluster4036 | 0.13 | 1.000 |
| cluster3774 | 0.12 | 1.000 |
| cluster3783 | 0.13 | 1.000 |
| cluster3090 | -0.38 | 1.000 |
| cluster3295 | -0.22 | 1.000 |
| cluster3423 | -0.33 | 1.000 |
| cluster3486 | -0.25 | 1.000 |
| cluster3315 | -0.22 | 1.000 |
| cluster2979 | -0.29 | 1.000 |
| cluster4040 | 0.12 | 1.000 |
| cluster3526 | -0.26 | 1.000 |
| cluster3592 | 0.12 | 1.000 |
| cluster3055 | 0.11 | 1.000 |
| cluster2883 | 0.11 | 1.000 |
| cluster2893 | 0.12 | 1.000 |
| cluster2968 | 0.10 | 1.000 |
| cluster4057 | 0.11 | 1.000 |
| cluster3974 | -0.35 | 1.000 |
| cluster3154 | 0.10 | 1.000 |
| cluster3797 | 0.12 | 1.000 |
| cluster3996 | -0.28 | 1.000 |
| cluster3934 | -0.29 | 1.000 |
| cluster3491 | 0.11 | 1.000 |
| cluster4037 | -0.29 | 1.000 |
| cluster3727 | 0.11 | 1.000 |
| cluster3793 | 0.10 | 1.000 |
| cluster4008 | -0.28 | 1.000 |
| cluster3263 | -0.23 | 1.000 |
| cluster3558 | 0.10 | 1.000 |
| cluster3860 | -0.22 | 1.000 |
| cluster3924 | -0.30 | 1.000 |
| cluster3975 | -0.30 | 1.000 |
| cluster3886 | -0.29 | 1.000 |
| cluster3917 | -0.30 | 1.000 |
| cluster3505 | -0.29 | 1.000 |
| cluster2973 | -0.30 | 1.000 |

| | | |
|---|---|---|
| cluster3322 | 0.10 | 1.000 |
| cluster4039 | -0.23 | 1.000 |
| cluster3965 | 0.11 | 1.000 |
| cluster4069 | 0.09 | 1.000 |
| cluster3428 | 0.09 | 1.000 |
| cluster3106 | -0.21 | 1.000 |
| cluster3078 | -0.19 | 1.000 |
| cluster3533 | 0.09 | 1.000 |
| cluster3412 | 0.09 | 1.000 |
| cluster3467 | 0.09 | 1.000 |
| cluster3601 | -0.28 | 1.000 |
| cluster3378 | 0.09 | 1.000 |
| cluster3173 | -0.27 | 1.000 |
| cluster3587 | -0.34 | 1.000 |
| cluster3067 | -0.19 | 1.000 |
| cluster4072 | -0.28 | 1.000 |
| cluster3851 | 0.08 | 1.000 |
| cluster3289 | -0.19 | 1.000 |
| cluster2918 | -0.23 | 1.000 |
| cluster3210 | -0.24 | 1.000 |
| cluster3576 | -0.24 | 1.000 |
| cluster3120 | 0.08 | 1.000 |
| cluster3136 | -0.25 | 1.000 |
| cluster3893 | 0.08 | 1.000 |
| cluster3294 | 0.08 | 1.000 |
| cluster3184 | 0.08 | 1.000 |
| cluster3031 | -0.19 | 1.000 |
| cluster2889 | -0.26 | 1.000 |
| cluster3692 | 0.08 | 1.000 |
| cluster3279 | -0.24 | 1.000 |
| cluster3109 | -0.29 | 1.000 |
| cluster3632 | -0.19 | 1.000 |
| cluster3059 | 0.07 | 1.000 |
| cluster3945 | 0.07 | 1.000 |
| cluster3236 | -0.26 | 1.000 |
| cluster2897 | 0.07 | 1.000 |
| cluster3193 | 0.07 | 1.000 |
| cluster3502 | -0.26 | 1.000 |
| cluster2925 | -0.28 | 1.000 |
| cluster3065 | -0.21 | 1.000 |
| cluster3283 | -0.17 | 1.000 |
| cluster3895 | 0.07 | 1.000 |
| cluster2986 | 0.06 | 1.000 |
| cluster3710 | -0.24 | 1.000 |

| | | |
|---|---|---|
| cluster3589 | 0.06 | 1.000 |
| cluster3570 | 0.06 | 1.000 |
| cluster3265 | -0.27 | 1.000 |
| cluster3921 | -0.27 | 1.000 |
| cluster3012 | -0.27 | 1.000 |
| cluster3642 | -0.25 | 1.000 |
| cluster3722 | -0.24 | 1.000 |
| cluster2880 | -0.28 | 1.000 |
| cluster2955 | 0.06 | 1.000 |
| cluster3559 | -0.24 | 1.000 |
| cluster3686 | -0.25 | 1.000 |
| cluster3628 | 0.06 | 1.000 |
| cluster3504 | 0.06 | 1.000 |
| cluster3324 | 0.06 | 1.000 |
| cluster3128 | -0.17 | 1.000 |
| cluster3808 | -0.23 | 1.000 |
| cluster3684 | 0.06 | 1.000 |
| cluster3409 | 0.05 | 1.000 |
| cluster3418 | -0.23 | 1.000 |
| cluster3785 | 0.05 | 1.000 |
| cluster3158 | 0.05 | 1.000 |
| cluster3542 | 0.05 | 1.000 |
| cluster2885 | 0.05 | 1.000 |
| cluster3635 | 0.05 | 1.000 |
| cluster3248 | 0.05 | 1.000 |
| cluster3230 | 0.06 | 1.000 |
| cluster3529 | 0.05 | 1.000 |
| cluster3472 | -0.16 | 1.000 |
| cluster3343 | 0.04 | 1.000 |
| cluster3687 | 0.04 | 1.000 |
| cluster3605 | 0.04 | 1.000 |
| cluster3201 | 0.05 | 1.000 |
| cluster3116 | 0.04 | 1.000 |
| cluster3018 | 0.04 | 1.000 |
| cluster3287 | -0.20 | 1.000 |
| cluster3489 | -0.23 | 1.000 |
| cluster2953 | 0.05 | 1.000 |
| cluster3148 | -0.16 | 1.000 |
| cluster3072 | -0.25 | 1.000 |
| cluster3683 | 0.03 | 1.000 |
| cluster3326 | 0.04 | 1.000 |
| cluster3838 | 0.03 | 1.000 |
| cluster3607 | -0.22 | 1.000 |
| cluster3569 | -0.22 | 1.000 |

| | | |
|---|---|---|
| cluster2970 | 0.03 | 1.000 |
| cluster4017 | -0.20 | 1.000 |
| cluster3550 | 0.04 | 1.000 |
| cluster2855 | -0.24 | 1.000 |
| cluster3998 | -0.28 | 1.000 |
| cluster3697 | -0.19 | 1.000 |
| cluster3588 | -0.19 | 1.000 |
| cluster3274 | -0.21 | 1.000 |
| cluster3002 | -0.14 | 1.000 |
| cluster3962 | -0.20 | 1.000 |
| cluster3947 | 0.02 | 1.000 |
| cluster3429 | 0.03 | 1.000 |
| cluster4032 | -0.17 | 1.000 |
| cluster3407 | -0.14 | 1.000 |
| cluster2923 | -0.14 | 1.000 |
| cluster3087 | -0.16 | 1.000 |
| cluster2869 | 0.02 | 1.000 |
| cluster3286 | 0.03 | 1.000 |
| cluster3096 | 0.02 | 1.000 |
| cluster3381 | -0.15 | 1.000 |
| cluster3041 | 0.02 | 1.000 |
| cluster3281 | 0.00 | 1.000 |
| cluster3862 | 0.00 | 1.000 |
| cluster3079 | -0.13 | 1.000 |
| cluster3707 | -0.19 | 1.000 |
| cluster3506 | -0.18 | 1.000 |
| cluster3896 | 0.00 | 1.000 |
| cluster3025 | 0.02 | 1.000 |
| cluster3028 | 0.00 | 1.000 |
| cluster3617 | -0.24 | 1.000 |
| cluster2988 | -0.16 | 1.000 |
| cluster3961 | -0.19 | 1.000 |
| cluster3739 | 0.01 | 1.000 |
| cluster4014 | -0.01 | 1.000 |
| cluster2875 | 0.01 | 1.000 |
| cluster3828 | 0.00 | 1.000 |
| cluster3339 | 0.01 | 1.000 |
| cluster2858 | -0.12 | 1.000 |
| cluster3702 | -0.17 | 1.000 |
| cluster3256 | 0.02 | 1.000 |
| cluster3139 | -0.21 | 1.000 |
| cluster2909 | 0.01 | 1.000 |
| cluster3189 | -0.17 | 1.000 |
| cluster3211 | -0.17 | 1.000 |

| | | |
|---|---|---|
| cluster3456 | -0.11 | 1.000 |
| cluster3046 | -0.13 | 1.000 |
| cluster3216 | -0.16 | 1.000 |
| cluster3384 | -0.01 | 1.000 |
| cluster3967 | -0.13 | 1.000 |
| cluster3241 | -0.14 | 1.000 |
| cluster3362 | -0.12 | 1.000 |
| cluster3737 | -0.10 | 1.000 |
| cluster3560 | -0.15 | 1.000 |
| cluster2983 | -0.15 | 1.000 |
| cluster3044 | -0.16 | 1.000 |
| cluster3614 | -0.16 | 1.000 |
| cluster3535 | -0.15 | 1.000 |
| cluster4105 | -0.18 | 1.000 |
| cluster4065 | -0.15 | 1.000 |
| cluster2887 | -0.04 | 1.000 |
| cluster2900 | -0.02 | 1.000 |
| cluster3545 | -0.14 | 1.000 |
| cluster2985 | -0.10 | 1.000 |
| cluster3314 | -0.05 | 1.000 |
| cluster3404 | -0.02 | 1.000 |
| cluster3202 | -0.10 | 1.000 |
| cluster4007 | -0.12 | 1.000 |
| cluster2917 | -0.05 | 1.000 |
| cluster3372 | -0.09 | 1.000 |
| cluster2959 | -0.09 | 1.000 |
| cluster3624 | -0.13 | 1.000 |
| cluster3648 | -0.03 | 1.000 |
| cluster3590 | -0.12 | 1.000 |
| cluster3503 | -0.12 | 1.000 |
| cluster2886 | -0.04 | 1.000 |
| cluster3900 | -0.17 | 1.000 |
| cluster3618 | -0.17 | 1.000 |
| cluster3208 | -0.08 | 1.000 |
| cluster2856 | -0.09 | 1.000 |
| cluster3192 | -0.07 | 1.000 |
| cluster3957 | -0.12 | 1.000 |
| cluster3266 | -0.07 | 1.000 |
| cluster3003 | -0.11 | 1.000 |
| cluster3688 | -0.08 | 1.000 |
| cluster3035 | -0.07 | 1.000 |
| cluster2872 | -0.09 | 1.000 |
| cluster2874 | -0.08 | 1.000 |
| cluster3639 | -0.14 | 1.000 |

| | | |
|---|---|---|
| cluster3612 | -0.10 | 1.000 |
| cluster3803 | -0.10 | 1.000 |
| cluster3354 | -0.07 | 1.000 |
| cluster3944 | -0.10 | 1.000 |
| cluster2951 | -0.10 | 1.000 |
| cluster3950 | -0.06 | 1.000 |
| cluster3647 | -0.10 | 1.000 |
| cluster3416 | -0.11 | 1.000 |
| cluster3108 | -0.10 | 1.000 |
| cluster3935 | -0.10 | 1.000 |
| cluster4003 | -0.09 | 1.000 |
| cluster3725 | -0.09 | 1.000 |
| cluster2929 | -0.08 | 1.000 |
| cluster3420 | -0.08 | 1.000 |
| cluster4038 | -0.07 | 1.000 |
| cluster3370 | -0.07 | 1.000 |
| cluster3284 | -0.07 | 1.000 |
| cluster3882 | -0.07 | 1.000 |
| cluster3866 | -0.07 | 1.000 |
| cluster4104 | -0.06 | 1.000 |
| cluster3646 | -0.06 | 1.000 |
| cluster4031 | -0.06 | 1.000 |
| cluster3681 | -0.06 | 1.000 |
| cluster3333 | -0.06 | 1.000 |
| cluster4126 | -0.06 | 1.000 |
| cluster3966 | -0.05 | 1.000 |
| cluster3927 | -0.05 | 1.000 |
| cluster3939 | -0.05 | 1.000 |
| cluster3680 | -0.04 | 1.000 |
| cluster3814 | -0.04 | 1.000 |
| cluster4027 | -0.03 | 1.000 |
| cluster3464 | -0.03 | 1.000 |
| cluster3951 | -0.03 | 1.000 |
| cluster3422 | -0.03 | 1.000 |
| cluster3171 | -0.03 | 1.000 |
| cluster3475 | -0.02 | 1.000 |
| cluster3637 | -0.02 | 1.000 |
| cluster3170 | -0.02 | 1.000 |
| cluster3347 | -0.02 | 1.000 |
| cluster2990 | -0.02 | 1.000 |
| cluster3700 | -0.02 | 1.000 |
| cluster4024 | -0.01 | 1.000 |
| cluster3492 | -0.01 | 1.000 |
| cluster3152 | -0.01 | 1.000 |

| | | |
|---|---|---|
| cluster3698 | -0.01 | 1.000 |

**Supplementary Table 11.** DESeq2 results with eXpress counts - Dermic vs nasopharyngeal

| ID | log2FoldChange | Adjusted p value |
|---|---|---|
| cluster3845 | -11.79 | 5.20E-43 |
| cluster4097 | -10.56 | 3.02E-28 |
| cluster3544 | -7.00 | 1.21E-22 |
| cluster4049 | -11.17 | 8.83E-20 |
| cluster3753 | -6.34 | 1.89E-18 |
| cluster3919 | -5.44 | 3.76E-16 |
| cluster3510 | -8.03 | 9.62E-16 |
| cluster4011 | -6.80 | 5.83E-13 |
| cluster4080 | -6.61 | 2.90E-11 |
| cluster3925 | -11.40 | 2.08E-09 |
| cluster4087 | 4.93 | 3.00E-07 |
| cluster3666 | 4.15 | 1.22E-06 |
| cluster4085 | -4.09 | 1.23E-06 |
| cluster2869 | -4.18 | 1.99E-05 |
| cluster4046 | -5.90 | 3.30E-05 |
| cluster3800 | -3.54 | 3.77E-05 |
| cluster3901 | 3.31 | 6.09E-05 |
| cluster2894 | -4.25 | 6.29E-05 |
| cluster3780 | -6.20 | 0.0003 |
| cluster3326 | -3.26 | 0.0003 |
| cluster3981 | 3.00 | 0.0003 |
| cluster3534 | -3.47 | 0.0005 |
| cluster3853 | 2.78 | 0.0016 |
| cluster4015 | -2.66 | 0.0039 |
| cluster3474 | 2.97 | 0.0044 |
| cluster3379 | -2.74 | 0.0052 |
| cluster4034 | -5.35 | 0.0052 |
| cluster3650 | -3.48 | 0.0055 |
| cluster3839 | -3.95 | 0.0055 |
| cluster4002 | 2.62 | 0.0058 |
| cluster3481 | -4.03 | 0.0070 |
| cluster4119 | 3.67 | 0.0094 |
| cluster3306 | 3.21 | 0.0100 |
| cluster3160 | -2.08 | 0.0113 |
| cluster2904 | -3.11 | 0.0119 |
| cluster3111 | -2.66 | 0.0120 |
| cluster3932 | -4.34 | 0.0120 |
| cluster3057 | -3.09 | 0.0121 |
| cluster3504 | -3.63 | 0.0131 |
| cluster3568 | -3.58 | 0.0135 |
| cluster3629 | 2.47 | 0.0136 |
| cluster3009 | -2.68 | 0.0151 |

| | | |
|---|---|---|
| cluster4070 | -5.76 | 0.0151 |
| cluster4099 | 1.97 | 0.0197 |
| cluster3024 | -2.84 | 0.0204 |
| cluster4073 | 3.53 | 0.0204 |
| cluster4067 | 4.35 | 0.0214 |
| cluster3325 | -4.25 | 0.0219 |
| cluster4090 | -4.97 | 0.0219 |
| cluster3052 | -3.51 | 0.0290 |
| cluster2915 | -2.32 | 0.0327 |
| cluster4113 | -2.07 | 0.0327 |
| cluster3482 | -2.44 | 0.0347 |
| cluster3161 | -2.14 | 0.0432 |
| cluster4125 | -5.11 | 0.0432 |
| cluster3003 | -2.50 | 0.0487 |
| cluster2900 | -2.89 | 0.0513 |
| cluster2974 | -2.55 | 0.0513 |
| cluster3288 | 2.00 | 0.0513 |
| cluster3431 | 1.85 | 0.0513 |
| cluster2925 | -2.45 | 0.0530 |
| cluster2992 | -2.17 | 0.0540 |
| cluster3432 | -2.49 | 0.0540 |
| cluster4074 | 2.93 | 0.0540 |
| cluster3581 | -2.50 | 0.0541 |
| cluster3223 | -2.50 | 0.0594 |
| cluster3902 | 1.79 | 0.0594 |
| cluster3008 | 1.84 | 0.0606 |
| cluster3393 | -3.42 | 0.0607 |
| cluster3290 | -2.70 | 0.0620 |
| cluster3983 | 2.50 | 0.0620 |
| cluster3675 | -1.84 | 0.0631 |
| cluster3229 | -2.74 | 0.0638 |
| cluster3772 | -2.19 | 0.0668 |
| cluster3869 | 2.78 | 0.0668 |
| cluster3952 | 1.82 | 0.0668 |
| cluster3250 | 2.76 | 0.0668 |
| cluster3617 | -3.10 | 0.0668 |
| cluster3880 | -3.72 | 0.0687 |
| cluster4047 | 3.08 | 0.0705 |
| cluster3438 | 1.95 | 0.0716 |
| cluster3970 | 2.44 | 0.0839 |
| cluster3761 | -2.93 | 0.0846 |
| cluster2972 | -2.31 | 0.0857 |
| cluster3105 | -2.92 | 0.0857 |
| cluster3408 | -2.43 | 0.0857 |

| | | |
|---|---|---|
| cluster3424 | -3.50 | 0.0857 |
| cluster3434 | -2.41 | 0.0857 |
| cluster3443 | 2.14 | 0.0857 |
| cluster3507 | -2.27 | 0.0857 |
| cluster4107 | 2.69 | 0.0857 |
| cluster2928 | 1.80 | 0.0864 |
| cluster3430 | 2.87 | 0.1000 |
| cluster3633 | -1.88 | 0.1095 |
| cluster3875 | 1.62 | 0.1095 |
| cluster4060 | 2.34 | 0.1095 |
| cluster3436 | 2.83 | 0.1098 |
| cluster3953 | -1.83 | 0.1109 |
| cluster3694 | -2.02 | 0.1146 |
| cluster3865 | 2.63 | 0.1146 |
| cluster3994 | -3.05 | 0.1146 |
| cluster3076 | 1.68 | 0.1162 |
| cluster3486 | -2.36 | 0.1162 |
| cluster3745 | 2.85 | 0.1162 |
| cluster3812 | -2.48 | 0.1162 |
| cluster2954 | -2.22 | 0.1163 |
| cluster3787 | -2.63 | 0.1163 |
| cluster3383 | -3.23 | 0.1170 |
| cluster4016 | 2.08 | 0.1173 |
| cluster3520 | -3.46 | 0.1195 |
| cluster3135 | -1.95 | 0.1215 |
| cluster3461 | -3.23 | 0.1233 |
| cluster4121 | 2.03 | 0.1236 |
| cluster3172 | -2.22 | 0.1236 |
| cluster2980 | -2.99 | 0.1249 |
| cluster3292 | -2.50 | 0.1272 |
| cluster3563 | -2.12 | 0.1274 |
| cluster3171 | -2.35 | 0.1304 |
| cluster4054 | 2.17 | 0.1322 |
| cluster3398 | -1.55 | 0.1343 |
| cluster4084 | -2.01 | 0.1343 |
| cluster3162 | 1.88 | 0.1344 |
| cluster3081 | -1.97 | 0.1353 |
| cluster3192 | -1.77 | 0.1353 |
| cluster3571 | -2.63 | 0.1376 |
| cluster3010 | -1.83 | 0.1433 |
| cluster3463 | 1.87 | 0.1433 |
| cluster4029 | -3.10 | 0.1469 |
| cluster4064 | 1.77 | 0.1504 |
| cluster3040 | -1.90 | 0.1532 |

| | | |
|---|---|---|
| cluster3287 | -2.06 | 0.1532 |
| cluster3381 | -2.05 | 0.1532 |
| cluster3519 | -3.09 | 0.1532 |
| cluster3556 | 1.94 | 0.1532 |
| cluster3867 | -1.50 | 0.1532 |
| cluster4091 | -3.23 | 0.1532 |
| cluster3179 | 2.19 | 0.1560 |
| cluster3824 | 2.32 | 0.1575 |
| cluster3151 | -1.37 | 0.1584 |
| cluster3488 | -1.45 | 0.1620 |
| cluster3084 | -1.56 | 0.1630 |
| cluster3422 | -2.92 | 0.1630 |
| cluster3698 | -1.74 | 0.1644 |
| cluster3933 | -1.84 | 0.1648 |
| cluster3086 | -1.51 | 0.1661 |
| cluster3719 | 2.04 | 0.1661 |
| cluster3913 | -2.11 | 0.1663 |
| cluster3035 | -1.59 | 0.1674 |
| cluster2912 | 1.52 | 0.1686 |
| cluster4093 | 2.63 | 0.1783 |
| cluster3814 | -1.96 | 0.1793 |
| cluster3017 | -2.16 | 0.1829 |
| cluster3889 | 2.40 | 0.1829 |
| cluster3771 | 1.71 | 0.1841 |
| cluster3065 | -1.68 | 0.1872 |
| cluster3543 | -2.42 | 0.1872 |
| cluster3674 | -3.25 | 0.1872 |
| cluster3142 | -2.14 | 0.1932 |
| cluster3442 | 2.17 | 0.2022 |
| cluster3538 | -2.32 | 0.2022 |
| cluster3752 | -2.83 | 0.2022 |
| cluster3995 | -2.31 | 0.2022 |
| cluster2908 | -1.79 | 0.2063 |
| cluster3487 | -1.93 | 0.2101 |
| cluster4044 | 1.61 | 0.2101 |
| cluster3220 | -1.90 | 0.2135 |
| cluster2890 | -1.48 | 0.2144 |
| cluster3454 | -1.57 | 0.2166 |
| cluster3205 | -1.44 | 0.2215 |
| cluster3419 | -1.36 | 0.2215 |
| cluster3651 | -2.82 | 0.2215 |
| cluster4081 | 1.27 | 0.2215 |
| cluster3453 | -2.03 | 0.2216 |
| cluster3395 | -1.69 | 0.2218 |

| | | |
|---|---|---|
| cluster3574 | -2.68 | 0.2244 |
| cluster3095 | -1.80 | 0.2248 |
| cluster3146 | -1.56 | 0.2248 |
| cluster3365 | 1.87 | 0.2248 |
| cluster3778 | -2.03 | 0.2248 |
| cluster3788 | -1.72 | 0.2248 |
| cluster3817 | -2.43 | 0.2248 |
| cluster3871 | 1.47 | 0.2248 |
| cluster3885 | -1.68 | 0.2248 |
| cluster3941 | -1.44 | 0.2248 |
| cluster3180 | -1.38 | 0.2270 |
| cluster3382 | -1.49 | 0.2270 |
| cluster3217 | -1.80 | 0.2288 |
| cluster3580 | 2.09 | 0.2288 |
| cluster3645 | -3.38 | 0.2288 |
| cluster3611 | 2.18 | 0.2289 |
| cluster3129 | -2.95 | 0.2323 |
| cluster3469 | -2.14 | 0.2323 |
| cluster3766 | -1.83 | 0.2323 |
| cluster3484 | -2.45 | 0.2368 |
| cluster4120 | -2.93 | 0.2400 |
| cluster3576 | 1.44 | 0.2426 |
| cluster3254 | -2.32 | 0.2440 |
| cluster3635 | -1.37 | 0.2440 |
| cluster3807 | -1.60 | 0.2440 |
| cluster3924 | 1.79 | 0.2440 |
| cluster3203 | -1.35 | 0.2452 |
| cluster3090 | -2.66 | 0.2505 |
| cluster3187 | -2.43 | 0.2505 |
| cluster3270 | 2.00 | 0.2505 |
| cluster3622 | -1.27 | 0.2505 |
| cluster3905 | -3.45 | 0.2505 |
| cluster3683 | 1.39 | 0.2513 |
| cluster3975 | 1.34 | 0.2513 |
| cluster3177 | -2.17 | 0.2555 |
| cluster3471 | 1.71 | 0.2555 |
| cluster3657 | 1.33 | 0.2555 |
| cluster4062 | -2.68 | 0.2555 |
| cluster3920 | -1.92 | 0.2580 |
| cluster3685 | -1.97 | 0.2599 |
| cluster3892 | -2.06 | 0.2599 |
| cluster3904 | -1.96 | 0.2599 |
| cluster4056 | -1.91 | 0.2599 |
| cluster4031 | -1.96 | 0.2638 |

| | | |
|---|---|---|
| cluster2914 | -1.79 | 0.2655 |
| cluster3462 | -1.58 | 0.2674 |
| cluster3978 | -1.25 | 0.2676 |
| cluster3000 | -1.73 | 0.2696 |
| cluster2913 | -1.15 | 0.2697 |
| cluster3114 | -1.19 | 0.2697 |
| cluster3724 | 1.76 | 0.2697 |
| cluster4037 | 1.21 | 0.2697 |
| cluster3957 | 1.57 | 0.2717 |
| cluster3122 | -1.53 | 0.2779 |
| cluster3131 | -1.47 | 0.2779 |
| cluster3501 | -2.14 | 0.2779 |
| cluster3508 | 1.98 | 0.2779 |
| cluster3569 | 1.79 | 0.2779 |
| cluster3598 | 2.60 | 0.2779 |
| cluster3702 | 1.43 | 0.2779 |
| cluster3711 | -1.16 | 0.2779 |
| cluster3804 | 1.54 | 0.2782 |
| cluster3815 | 2.04 | 0.2788 |
| cluster3441 | 1.20 | 0.2793 |
| cluster3363 | 1.30 | 0.2806 |
| cluster2867 | -2.25 | 0.2810 |
| cluster3374 | 1.97 | 0.2829 |
| cluster3415 | -1.44 | 0.2840 |
| cluster3016 | -2.28 | 0.2878 |
| cluster3196 | -1.29 | 0.2878 |
| cluster3194 | -1.57 | 0.2899 |
| cluster3455 | -1.89 | 0.2899 |
| cluster3457 | -1.94 | 0.2905 |
| cluster3690 | -1.57 | 0.2905 |
| cluster4024 | 1.41 | 0.2908 |
| cluster3968 | 1.65 | 0.2944 |
| cluster3193 | -1.34 | 0.2946 |
| cluster3822 | 2.20 | 0.2977 |
| cluster3942 | -1.43 | 0.2977 |
| cluster3410 | -2.00 | 0.3057 |
| cluster4100 | 1.77 | 0.3072 |
| cluster3594 | -1.71 | 0.3078 |
| cluster3634 | -1.69 | 0.3095 |
| cluster4051 | 1.79 | 0.3217 |
| cluster3572 | 2.48 | 0.3276 |
| cluster3444 | -1.22 | 0.3311 |
| cluster3734 | 1.71 | 0.3311 |
| cluster3936 | 1.63 | 0.3379 |

| | | |
|---|---|---|
| cluster3726 | -1.10 | 0.3386 |
| cluster3776 | -1.67 | 0.3397 |
| cluster3259 | -1.30 | 0.3411 |
| cluster3332 | 1.41 | 0.3411 |
| cluster3976 | -1.48 | 0.3411 |
| cluster3015 | -1.82 | 0.3417 |
| cluster3425 | -1.50 | 0.3417 |
| cluster2876 | 2.18 | 0.3422 |
| cluster3069 | -1.14 | 0.3422 |
| cluster3100 | 2.08 | 0.3422 |
| cluster3466 | -1.60 | 0.3422 |
| cluster2995 | 1.35 | 0.3430 |
| cluster3969 | -1.11 | 0.3430 |
| cluster3127 | -1.06 | 0.3431 |
| cluster3528 | -1.24 | 0.3431 |
| cluster3618 | -2.46 | 0.3431 |
| cluster2862 | -1.40 | 0.3479 |
| cluster3265 | -1.35 | 0.3519 |
| cluster3342 | 1.55 | 0.3574 |
| cluster4111 | -2.55 | 0.3578 |
| cluster3446 | -1.26 | 0.3612 |
| cluster3012 | 1.81 | 0.3641 |
| cluster3319 | -1.59 | 0.3641 |
| cluster3497 | -1.52 | 0.3641 |
| cluster3681 | 1.50 | 0.3641 |
| cluster2930 | 1.24 | 0.3645 |
| cluster3075 | -1.47 | 0.3645 |
| cluster3212 | -1.73 | 0.3645 |
| cluster3630 | -1.26 | 0.3645 |
| cluster3710 | 1.35 | 0.3645 |
| cluster3318 | -1.73 | 0.3661 |
| cluster3080 | -1.29 | 0.3687 |
| cluster3249 | -1.16 | 0.3725 |
| cluster3427 | -1.67 | 0.3725 |
| cluster3468 | -1.49 | 0.3725 |
| cluster3130 | -1.39 | 0.3757 |
| cluster3413 | -1.70 | 0.3851 |
| cluster3451 | 2.67 | 0.3870 |
| cluster3056 | -1.19 | 0.3872 |
| cluster3050 | -1.16 | 0.3885 |
| cluster3136 | -1.41 | 0.3885 |
| cluster3515 | -1.89 | 0.3885 |
| cluster3550 | -1.43 | 0.3885 |
| cluster3587 | -2.18 | 0.3885 |

| | | |
|---|---|---|
| cluster3773 | -1.24 | 0.3885 |
| cluster3584 | 1.73 | 0.3921 |
| cluster3439 | -1.20 | 0.3928 |
| cluster3649 | -1.03 | 0.3942 |
| cluster3591 | -1.15 | 0.3987 |
| cluster2918 | -1.20 | 0.3996 |
| cluster4079 | 1.52 | 0.4032 |
| cluster4109 | 1.69 | 0.4032 |
| cluster3678 | 1.13 | 0.4052 |
| cluster3866 | 1.67 | 0.4052 |
| cluster4116 | -2.36 | 0.4052 |
| cluster3971 | 1.54 | 0.4056 |
| cluster3360 | -1.49 | 0.4112 |
| cluster3545 | 1.17 | 0.4146 |
| cluster3068 | -1.00 | 0.4151 |
| cluster3368 | -1.09 | 0.4213 |
| cluster3641 | -1.34 | 0.4213 |
| cluster3653 | -1.53 | 0.4273 |
| cluster3626 | -1.31 | 0.4275 |
| cluster3082 | -2.10 | 0.4283 |
| cluster2966 | 1.43 | 0.4320 |
| cluster3175 | -1.71 | 0.4341 |
| cluster3661 | 1.11 | 0.4341 |
| cluster3073 | -0.98 | 0.4361 |
| cluster2887 | 1.23 | 0.4373 |
| cluster3230 | -1.53 | 0.4375 |
| cluster3742 | 1.71 | 0.4377 |
| cluster3923 | -1.01 | 0.4381 |
| cluster3947 | 0.92 | 0.4381 |
| cluster2953 | -1.09 | 0.4406 |
| cluster4089 | -1.95 | 0.4406 |
| cluster3185 | -1.56 | 0.4435 |
| cluster3475 | 1.16 | 0.4461 |
| cluster3525 | -1.12 | 0.4461 |
| cluster3735 | -1.44 | 0.4486 |
| cluster2987 | 0.91 | 0.4513 |
| cluster3199 | -0.99 | 0.4513 |
| cluster3275 | -1.64 | 0.4513 |
| cluster3279 | 1.19 | 0.4513 |
| cluster3397 | -1.63 | 0.4513 |
| cluster3097 | 1.31 | 0.4529 |
| cluster3940 | 1.29 | 0.4529 |
| cluster2892 | 1.11 | 0.4737 |
| cluster3182 | -1.38 | 0.4737 |

| | | |
|---|---|---|
| cluster4008 | 1.44 | 0.4737 |
| cluster3917 | 1.46 | 0.4741 |
| cluster3074 | -1.03 | 0.4760 |
| cluster2982 | -1.25 | 0.4786 |
| cluster3575 | -1.55 | 0.4786 |
| cluster3207 | 1.32 | 0.4821 |
| cluster3045 | 1.29 | 0.4884 |
| cluster4065 | 0.93 | 0.4928 |
| cluster3414 | -1.27 | 0.4955 |
| cluster3552 | 1.00 | 0.4961 |
| cluster3811 | -1.22 | 0.4961 |
| cluster2903 | -1.54 | 0.4984 |
| cluster3213 | -1.39 | 0.4984 |
| cluster3231 | -1.45 | 0.4984 |
| cluster3512 | 1.31 | 0.4984 |
| cluster3638 | 1.61 | 0.4984 |
| cluster3639 | 2.01 | 0.4984 |
| cluster4000 | 1.18 | 0.4984 |
| cluster3153 | 0.81 | 0.5063 |
| cluster3418 | 1.09 | 0.5063 |
| cluster3796 | -1.70 | 0.5063 |
| cluster2922 | 1.26 | 0.5088 |
| cluster3109 | 1.23 | 0.5088 |
| cluster3121 | -1.32 | 0.5088 |
| cluster3489 | 1.24 | 0.5088 |
| cluster3503 | 1.01 | 0.5088 |
| cluster3956 | 1.10 | 0.5088 |
| cluster3004 | -1.21 | 0.5104 |
| cluster2901 | -1.34 | 0.5108 |
| cluster3103 | -1.61 | 0.5108 |
| cluster3301 | -1.38 | 0.5108 |
| cluster3899 | 1.06 | 0.5184 |
| cluster3774 | -1.19 | 0.5218 |
| cluster3043 | -0.93 | 0.5224 |
| cluster3915 | -1.78 | 0.5224 |
| cluster3730 | -1.24 | 0.5224 |
| cluster3813 | 1.35 | 0.5346 |
| cluster2998 | -0.84 | 0.5357 |
| cluster2957 | -1.39 | 0.5377 |
| cluster3518 | -1.66 | 0.5377 |
| cluster3603 | 1.19 | 0.5377 |
| cluster3604 | 1.47 | 0.5377 |
| cluster3985 | 1.10 | 0.5377 |
| cluster3357 | 1.02 | 0.5377 |

| | | |
|---|---|---|
| cluster3945 | -1.46 | 0.5406 |
| cluster2969 | 1.27 | 0.5416 |
| cluster3242 | 1.08 | 0.5416 |
| cluster2895 | -1.09 | 0.5432 |
| cluster3298 | -1.02 | 0.5432 |
| cluster3449 | -1.15 | 0.5441 |
| cluster3236 | 1.16 | 0.5495 |
| cluster3440 | -0.91 | 0.5505 |
| cluster4055 | -1.17 | 0.5505 |
| cluster2978 | -1.31 | 0.5524 |
| cluster3209 | -1.09 | 0.5529 |
| cluster3551 | -1.37 | 0.5529 |
| cluster3315 | -1.17 | 0.5544 |
| cluster3578 | -1.20 | 0.5544 |
| cluster3642 | 1.26 | 0.5593 |
| cluster3844 | -1.82 | 0.5593 |
| cluster3849 | 0.99 | 0.5593 |
| cluster4071 | 1.61 | 0.5593 |
| cluster3366 | 1.42 | 0.5594 |
| cluster3768 | -1.44 | 0.5594 |
| cluster3801 | 1.35 | 0.5594 |
| cluster3834 | -1.43 | 0.5594 |
| cluster3157 | -1.33 | 0.5600 |
| cluster3282 | -0.99 | 0.5601 |
| cluster4010 | 1.32 | 0.5601 |
| cluster3346 | -1.31 | 0.5661 |
| cluster2856 | 1.23 | 0.5719 |
| cluster3647 | 0.85 | 0.5731 |
| cluster3537 | 0.91 | 0.5812 |
| cluster2938 | -0.89 | 0.5869 |
| cluster3662 | -1.26 | 0.5869 |
| cluster3850 | -1.18 | 0.5878 |
| cluster3939 | 0.96 | 0.5878 |
| cluster2942 | 0.96 | 0.5900 |
| cluster3094 | 1.43 | 0.5900 |
| cluster3168 | 1.04 | 0.5900 |
| cluster3624 | 0.80 | 0.5900 |
| cluster3715 | 1.20 | 0.5900 |
| cluster3400 | -1.09 | 0.5957 |
| cluster3821 | -1.09 | 0.5961 |
| cluster2955 | -0.79 | 0.5978 |
| cluster3377 | -1.03 | 0.5995 |
| cluster3435 | -1.50 | 0.5995 |
| cluster3110 | 0.86 | 0.5998 |

| | | |
|---|---|---|
| cluster3330 | 0.96 | 0.5998 |
| cluster3355 | -1.20 | 0.5998 |
| cluster3450 | -1.82 | 0.5998 |
| cluster3467 | 0.87 | 0.5998 |
| cluster3977 | -1.44 | 0.5998 |
| cluster3636 | -0.79 | 0.6030 |
| cluster3672 | -1.21 | 0.6030 |
| cluster2886 | -1.14 | 0.6045 |
| cluster2975 | 0.74 | 0.6045 |
| cluster3532 | -1.33 | 0.6045 |
| cluster4104 | 1.33 | 0.6045 |
| cluster4110 | 1.26 | 0.6045 |
| cluster3646 | 1.46 | 0.6053 |
| cluster3759 | -1.08 | 0.6053 |
| cluster3628 | 1.11 | 0.6088 |
| cluster3696 | -0.79 | 0.6088 |
| cluster3590 | 1.06 | 0.6095 |
| cluster3271 | -0.85 | 0.6151 |
| cluster3375 | -0.87 | 0.6151 |
| cluster3777 | -0.96 | 0.6151 |
| cluster3870 | -1.44 | 0.6151 |
| cluster3404 | -0.92 | 0.6152 |
| cluster3783 | 1.40 | 0.6158 |
| cluster4038 | 1.32 | 0.6183 |
| cluster3540 | -1.31 | 0.6190 |
| cluster3676 | 1.38 | 0.6190 |
| cluster3190 | 1.43 | 0.6224 |
| cluster3327 | -1.14 | 0.6224 |
| cluster4068 | 1.27 | 0.6224 |
| cluster3285 | -0.92 | 0.6229 |
| cluster3832 | 0.98 | 0.6249 |
| cluster2905 | -1.01 | 0.6307 |
| cluster3349 | 1.09 | 0.6307 |
| cluster3405 | -0.91 | 0.6307 |
| cluster3513 | -1.12 | 0.6307 |
| cluster3910 | -1.38 | 0.6307 |
| cluster4001 | -1.23 | 0.6307 |
| cluster4009 | 1.22 | 0.6307 |
| cluster4083 | 1.29 | 0.6307 |
| cluster3401 | 0.74 | 0.6313 |
| cluster3524 | -0.74 | 0.6315 |
| cluster2920 | -0.87 | 0.6318 |
| cluster3644 | -1.10 | 0.6318 |
| cluster3836 | 0.95 | 0.6318 |

| | | |
|---|---|---|
| cluster3226 | -1.17 | 0.6343 |
| cluster3643 | 1.20 | 0.6343 |
| cluster3841 | -1.20 | 0.6343 |
| cluster3872 | 1.07 | 0.6343 |
| cluster3980 | 0.64 | 0.6343 |
| cluster3210 | 0.65 | 0.6349 |
| cluster3026 | -1.11 | 0.6370 |
| cluster2963 | -1.19 | 0.6388 |
| cluster2855 | 0.69 | 0.6391 |
| cluster3077 | -1.10 | 0.6391 |
| cluster3260 | -0.96 | 0.6391 |
| cluster3336 | -1.11 | 0.6391 |
| cluster3387 | 0.88 | 0.6391 |
| cluster3721 | -1.22 | 0.6391 |
| cluster3831 | 0.99 | 0.6391 |
| cluster3837 | 1.04 | 0.6391 |
| cluster4028 | 0.82 | 0.6391 |
| cluster4057 | 1.25 | 0.6391 |
| cluster2927 | 1.18 | 0.6404 |
| cluster3781 | 0.96 | 0.6404 |
| cluster3864 | 0.72 | 0.6404 |
| cluster3879 | 0.79 | 0.6404 |
| cluster3197 | -1.10 | 0.6438 |
| cluster3914 | -1.72 | 0.6438 |
| cluster2858 | -0.83 | 0.6468 |
| cluster2936 | 1.29 | 0.6468 |
| cluster2937 | -0.90 | 0.6468 |
| cluster3006 | -1.41 | 0.6468 |
| cluster3053 | -1.30 | 0.6468 |
| cluster3085 | -1.12 | 0.6468 |
| cluster3107 | -0.76 | 0.6468 |
| cluster3234 | -0.92 | 0.6468 |
| cluster3274 | 1.06 | 0.6468 |
| cluster3344 | -1.02 | 0.6468 |
| cluster3416 | 1.36 | 0.6468 |
| cluster3429 | -1.02 | 0.6468 |
| cluster3485 | -1.48 | 0.6468 |
| cluster3640 | -2.53 | 0.6468 |
| cluster3709 | -0.94 | 0.6468 |
| cluster3740 | 1.11 | 0.6468 |
| cluster3895 | 1.14 | 0.6468 |
| cluster3909 | 0.89 | 0.6468 |
| cluster3944 | 0.86 | 0.6468 |
| cluster3965 | -1.19 | 0.6468 |

| | | |
|---|---|---|
| cluster3967 | -1.07 | 0.6468 |
| cluster4033 | -1.51 | 0.6468 |
| cluster4036 | 0.67 | 0.6468 |
| cluster4014 | 1.08 | 0.6499 |
| cluster3195 | -0.87 | 0.6530 |
| cluster3189 | -0.97 | 0.6532 |
| cluster3828 | 0.97 | 0.6540 |
| cluster4069 | 1.02 | 0.6540 |
| cluster3264 | -0.70 | 0.6542 |
| cluster2990 | 0.72 | 0.6608 |
| cluster3227 | -0.85 | 0.6608 |
| cluster3514 | -1.15 | 0.6608 |
| cluster3595 | 0.88 | 0.6611 |
| cluster3627 | -0.72 | 0.6611 |
| cluster2854 | 0.99 | 0.6635 |
| cluster2866 | -1.08 | 0.6635 |
| cluster3079 | -0.94 | 0.6635 |
| cluster3123 | -0.78 | 0.6635 |
| cluster3286 | -0.71 | 0.6635 |
| cluster3476 | 1.14 | 0.6635 |
| cluster3542 | 1.11 | 0.6635 |
| cluster3570 | 1.29 | 0.6635 |
| cluster3825 | -0.87 | 0.6635 |
| cluster3943 | 1.15 | 0.6635 |
| cluster4027 | 1.10 | 0.6635 |
| cluster2934 | -0.68 | 0.6644 |
| cluster2951 | 1.01 | 0.6644 |
| cluster3693 | 1.28 | 0.6644 |
| cluster2906 | -0.99 | 0.6646 |
| cluster2924 | 0.97 | 0.6646 |
| cluster3025 | -0.70 | 0.6646 |
| cluster3033 | -0.57 | 0.6646 |
| cluster3237 | -1.24 | 0.6646 |
| cluster3243 | -0.92 | 0.6646 |
| cluster3317 | -1.06 | 0.6646 |
| cluster3362 | -0.89 | 0.6646 |
| cluster3614 | 1.29 | 0.6646 |
| cluster3703 | 0.75 | 0.6646 |
| cluster3739 | -0.66 | 0.6646 |
| cluster3792 | -0.75 | 0.6646 |
| cluster3855 | 1.02 | 0.6646 |
| cluster4023 | -1.05 | 0.6646 |
| cluster3477 | -1.00 | 0.6682 |
| cluster3029 | 0.64 | 0.6719 |

| | | |
|---|---|---|
| cluster3047 | -0.99 | 0.6744 |
| cluster4019 | 1.17 | 0.6785 |
| cluster2877 | 0.60 | 0.6814 |
| cluster3204 | -0.88 | 0.6831 |
| cluster3437 | -0.86 | 0.6831 |
| cluster3339 | -0.86 | 0.6837 |
| cluster3951 | 1.03 | 0.6843 |
| cluster3083 | 0.71 | 0.6870 |
| cluster3188 | -1.04 | 0.6870 |
| cluster3616 | -1.17 | 0.6870 |
| cluster3460 | 0.89 | 0.6878 |
| cluster3659 | -1.07 | 0.6878 |
| cluster3701 | -0.73 | 0.6878 |
| cluster2958 | -0.60 | 0.6878 |
| cluster1127 | -0.91 | 0.6899 |
| cluster2935 | -0.86 | 0.6899 |
| cluster3014 | -0.98 | 0.6899 |
| cluster3038 | 0.92 | 0.6899 |
| cluster3191 | -1.29 | 0.6899 |
| cluster3247 | 0.85 | 0.6899 |
| cluster3261 | -1.10 | 0.6899 |
| cluster3262 | -0.69 | 0.6899 |
| cluster3370 | 0.85 | 0.6899 |
| cluster3433 | -0.81 | 0.6899 |
| cluster3456 | -0.74 | 0.6899 |
| cluster3472 | -0.84 | 0.6899 |
| cluster3500 | -0.99 | 0.6899 |
| cluster3546 | -1.24 | 0.6899 |
| cluster3658 | -0.75 | 0.6899 |
| cluster3793 | 0.69 | 0.6899 |
| cluster3830 | 0.73 | 0.6899 |
| cluster3859 | -0.71 | 0.6899 |
| cluster3973 | -0.82 | 0.6899 |
| cluster4018 | 0.92 | 0.6899 |
| cluster4088 | 1.04 | 0.6899 |
| cluster3092 | -0.62 | 0.6953 |
| cluster3464 | 0.80 | 0.6953 |
| cluster3582 | -0.78 | 0.6953 |
| cluster3725 | 1.04 | 0.6953 |
| cluster3997 | 1.03 | 0.6953 |
| cluster4118 | -1.60 | 0.6953 |
| cluster3916 | 0.64 | 0.6964 |
| cluster3473 | 0.93 | 0.6981 |
| cluster3750 | -1.22 | 0.6981 |

| | | |
|---|---|---|
| cluster2879 | -0.71 | 0.7008 |
| cluster3522 | -0.94 | 0.7008 |
| cluster3982 | 0.61 | 0.7008 |
| cluster3448 | -1.11 | 0.7021 |
| cluster3664 | -0.70 | 0.7021 |
| cluster3373 | 0.70 | 0.7055 |
| cluster3380 | -0.72 | 0.7055 |
| cluster3493 | 1.00 | 0.7055 |
| cluster3667 | -0.73 | 0.7055 |
| cluster3699 | 1.01 | 0.7055 |
| cluster3208 | -0.68 | 0.7089 |
| cluster3723 | 0.60 | 0.7090 |
| cluster2931 | -0.80 | 0.7099 |
| cluster3117 | 1.08 | 0.7104 |
| cluster3911 | 1.07 | 0.7113 |
| cluster3340 | -0.74 | 0.7123 |
| cluster3358 | -0.88 | 0.7123 |
| cluster3930 | 0.68 | 0.7136 |
| cluster2880 | -0.76 | 0.7295 |
| cluster3677 | 0.75 | 0.7295 |
| cluster3966 | 0.55 | 0.7315 |
| cluster3042 | 0.80 | 0.7321 |
| cluster3239 | -0.65 | 0.7397 |
| cluster3252 | 1.22 | 0.7397 |
| cluster3478 | -0.78 | 0.7397 |
| cluster3998 | 1.03 | 0.7405 |
| cluster3602 | -0.71 | 0.7473 |
| cluster3198 | 0.76 | 0.7475 |
| cluster3829 | 0.92 | 0.7475 |
| cluster3625 | 0.59 | 0.7510 |
| cluster3445 | -0.59 | 0.7523 |
| cluster4042 | -0.93 | 0.7523 |
| cluster3310 | -0.48 | 0.7553 |
| cluster3388 | -0.87 | 0.7553 |
| cluster3428 | -0.69 | 0.7623 |
| cluster3763 | -0.99 | 0.7623 |
| cluster3908 | 0.58 | 0.7623 |
| cluster2950 | -0.67 | 0.7626 |
| cluster2940 | -0.66 | 0.7643 |
| cluster2859 | -0.47 | 0.7673 |
| cluster2977 | 0.62 | 0.7673 |
| cluster3028 | 0.56 | 0.7673 |
| cluster3046 | -0.65 | 0.7673 |
| cluster3364 | -0.65 | 0.7673 |

| | | |
|---|---|---|
| cluster3560 | 0.93 | 0.7673 |
| cluster3579 | -0.71 | 0.7673 |
| cluster3820 | -0.49 | 0.7673 |
| cluster4020 | -0.63 | 0.7673 |
| cluster4082 | 1.01 | 0.7673 |
| cluster4124 | -1.08 | 0.7673 |
| cluster2971 | -0.54 | 0.7689 |
| cluster3619 | 0.69 | 0.7707 |
| cluster3218 | 0.87 | 0.7726 |
| cluster3356 | 0.71 | 0.7726 |
| cluster3490 | -0.79 | 0.7726 |
| cluster3688 | 0.65 | 0.7726 |
| cluster3704 | 0.80 | 0.7726 |
| cluster4115 | -1.03 | 0.7726 |
| cluster2864 | 0.61 | 0.7747 |
| cluster3087 | 0.55 | 0.7747 |
| cluster3680 | 0.57 | 0.7747 |
| cluster3421 | 0.85 | 0.7760 |
| cluster4123 | -1.01 | 0.7760 |
| cluster3960 | -0.98 | 0.7822 |
| cluster3389 | 0.64 | 0.7862 |
| cluster3906 | 0.64 | 0.7862 |
| cluster2874 | 0.45 | 0.7864 |
| cluster2910 | -0.54 | 0.7864 |
| cluster3031 | -0.67 | 0.7864 |
| cluster3078 | 0.44 | 0.7864 |
| cluster3166 | -0.63 | 0.7864 |
| cluster3169 | 0.73 | 0.7864 |
| cluster3331 | -0.48 | 0.7864 |
| cluster3728 | -0.64 | 0.7864 |
| cluster3894 | -0.59 | 0.7864 |
| cluster3950 | 0.65 | 0.7864 |
| cluster4012 | 0.76 | 0.7864 |
| cluster4077 | 0.68 | 0.7864 |
| cluster4094 | 0.80 | 0.7864 |
| cluster4128 | 1.04 | 0.7871 |
| cluster2932 | -0.61 | 0.7903 |
| cluster3118 | 0.62 | 0.7908 |
| cluster3350 | -0.68 | 0.7908 |
| cluster3958 | -0.99 | 0.7909 |
| cluster3573 | -0.72 | 0.7916 |
| cluster3632 | -0.65 | 0.7916 |
| cluster2947 | 0.61 | 0.7934 |
| cluster3062 | -0.56 | 0.7934 |

| | | |
|---|---|---|
| cluster3334 | 0.51 | 0.7934 |
| cluster3559 | -0.50 | 0.7934 |
| cluster3765 | -0.69 | 0.7934 |
| cluster3891 | 0.76 | 0.7934 |
| cluster4021 | 0.78 | 0.7934 |
| cluster3732 | 0.56 | 0.7950 |
| cluster2865 | 0.60 | 0.7956 |
| cluster2959 | 0.55 | 0.7956 |
| cluster3479 | -0.84 | 0.7983 |
| cluster3881 | 0.74 | 0.7983 |
| cluster4006 | -0.51 | 0.7983 |
| cluster2994 | 0.54 | 0.8001 |
| cluster3158 | 0.77 | 0.8001 |
| cluster3347 | -0.49 | 0.8001 |
| cluster3862 | 0.45 | 0.8001 |
| cluster3868 | -0.46 | 0.8001 |
| cluster3293 | 0.51 | 0.8006 |
| cluster3963 | -0.91 | 0.8006 |
| cluster2893 | 0.84 | 0.8023 |
| cluster2967 | 0.75 | 0.8025 |
| cluster3211 | 0.82 | 0.8025 |
| cluster3396 | -0.66 | 0.8025 |
| cluster3934 | 0.42 | 0.8025 |
| cluster3164 | -0.86 | 0.8037 |
| cluster3214 | -0.58 | 0.8037 |
| cluster3245 | 0.75 | 0.8037 |
| cluster3321 | -0.51 | 0.8037 |
| cluster3496 | 0.60 | 0.8037 |
| cluster3599 | -0.80 | 0.8037 |
| cluster3601 | 0.66 | 0.8037 |
| cluster3846 | -0.45 | 0.8037 |
| cluster3854 | -0.47 | 0.8037 |
| cluster3996 | 0.47 | 0.8037 |
| cluster2945 | -0.92 | 0.8039 |
| cluster3072 | -0.71 | 0.8039 |
| cluster3567 | -0.95 | 0.8039 |
| cluster3608 | -0.51 | 0.8045 |
| cluster3656 | 0.63 | 0.8045 |
| cluster4127 | -1.05 | 0.8045 |
| cluster3553 | 0.58 | 0.8051 |
| cluster3093 | 0.62 | 0.8061 |
| cluster3181 | -0.63 | 0.8061 |
| cluster3530 | -0.49 | 0.8061 |
| cluster3938 | 0.68 | 0.8061 |

| | | |
|---|---|---|
| cluster2891 | -0.69 | 0.8073 |
| cluster2939 | -0.51 | 0.8073 |
| cluster2973 | 0.63 | 0.8073 |
| cluster3376 | -0.48 | 0.8073 |
| cluster3348 | -0.64 | 0.8080 |
| cluster3140 | 0.43 | 0.8087 |
| cluster3345 | 0.56 | 0.8087 |
| cluster3918 | 0.66 | 0.8087 |
| cluster2944 | -0.59 | 0.8089 |
| cluster3860 | 0.59 | 0.8089 |
| cluster3206 | 0.68 | 0.8094 |
| cluster2907 | -0.52 | 0.8097 |
| cluster3128 | -0.46 | 0.8097 |
| cluster3842 | 0.68 | 0.8097 |
| cluster3962 | 0.42 | 0.8097 |
| cluster4004 | -0.42 | 0.8097 |
| cluster3609 | 0.91 | 0.8104 |
| cluster3124 | -0.38 | 0.8105 |
| cluster3697 | -0.80 | 0.8109 |
| cluster3202 | 0.51 | 0.8131 |
| cluster3163 | -0.80 | 0.8141 |
| cluster3516 | -0.82 | 0.8141 |
| cluster2997 | -0.49 | 0.8189 |
| cluster3803 | 0.83 | 0.8189 |
| cluster3810 | 0.54 | 0.8189 |
| cluster2878 | -0.61 | 0.8193 |
| cluster3459 | -0.79 | 0.8193 |
| cluster3809 | 0.44 | 0.8193 |
| cluster3689 | -0.92 | 0.8198 |
| cluster3928 | -0.49 | 0.8204 |
| cluster3141 | -0.54 | 0.8206 |
| cluster3272 | 1.44 | 0.8206 |
| cluster2885 | -0.58 | 0.8216 |
| cluster2988 | 0.40 | 0.8216 |
| cluster3222 | -0.38 | 0.8216 |
| cluster3322 | 0.68 | 0.8216 |
| cluster3411 | 0.42 | 0.8216 |
| cluster3505 | -0.47 | 0.8216 |
| cluster3535 | -0.70 | 0.8216 |
| cluster3536 | 0.48 | 0.8216 |
| cluster3558 | 0.54 | 0.8216 |
| cluster3795 | -0.51 | 0.8216 |
| cluster3848 | -0.64 | 0.8216 |
| cluster4095 | 0.71 | 0.8216 |

| | | |
|---|---|---|
| cluster3299 | 0.62 | 0.8225 |
| cluster3660 | 0.71 | 0.8225 |
| cluster3756 | -0.57 | 0.8225 |
| cluster3791 | -0.62 | 0.8225 |
| cluster3758 | -0.60 | 0.8245 |
| cluster3037 | -0.47 | 0.8253 |
| cluster3019 | 0.47 | 0.8264 |
| cluster3215 | -0.39 | 0.8264 |
| cluster3566 | -0.45 | 0.8264 |
| cluster3718 | -0.62 | 0.8264 |
| cluster3826 | -0.64 | 0.8264 |
| cluster3972 | 0.45 | 0.8264 |
| cluster4072 | 0.46 | 0.8264 |
| cluster3108 | 0.55 | 0.8283 |
| cluster3267 | 0.46 | 0.8283 |
| cluster3509 | 0.36 | 0.8283 |
| cluster3607 | 0.58 | 0.8283 |
| cluster3748 | -0.56 | 0.8283 |
| cluster3856 | 0.49 | 0.8283 |
| cluster4030 | 0.58 | 0.8283 |
| cluster3099 | 0.44 | 0.8295 |
| cluster2888 | 0.80 | 0.8305 |
| cluster2889 | 0.40 | 0.8305 |
| cluster3606 | 0.68 | 0.8305 |
| cluster2875 | -0.54 | 0.8312 |
| cluster2923 | -0.40 | 0.8312 |
| cluster2986 | 0.41 | 0.8312 |
| cluster3005 | -0.41 | 0.8312 |
| cluster3007 | 0.44 | 0.8312 |
| cluster3039 | 0.44 | 0.8312 |
| cluster3144 | -0.76 | 0.8312 |
| cluster3289 | -0.48 | 0.8312 |
| cluster3305 | 0.59 | 0.8312 |
| cluster3369 | 0.45 | 0.8312 |
| cluster3390 | -0.48 | 0.8312 |
| cluster3403 | -0.39 | 0.8312 |
| cluster3517 | -0.32 | 0.8312 |
| cluster3585 | -0.45 | 0.8312 |
| cluster3717 | -0.44 | 0.8312 |
| cluster3727 | -0.65 | 0.8312 |
| cluster3986 | 0.38 | 0.8312 |
| cluster3988 | -0.36 | 0.8312 |
| cluster3991 | 0.60 | 0.8312 |
| cluster4050 | 0.61 | 0.8312 |

| | | |
|---|---|---|
| cluster4061 | 0.58 | 0.8312 |
| cluster3232 | 0.45 | 0.8317 |
| cluster3491 | 0.43 | 0.8317 |
| cluster3767 | 0.57 | 0.8317 |
| cluster3407 | -0.50 | 0.8323 |
| cluster3021 | -0.42 | 0.8348 |
| cluster3049 | -0.50 | 0.8356 |
| cluster4040 | 0.45 | 0.8356 |
| cluster4075 | -0.73 | 0.8388 |
| cluster3843 | 0.35 | 0.8391 |
| cluster4108 | -0.64 | 0.8430 |
| cluster3241 | -0.37 | 0.8435 |
| cluster3255 | -0.60 | 0.8435 |
| cluster3700 | 0.48 | 0.8460 |
| cluster4017 | 0.53 | 0.8460 |
| cluster2872 | -0.38 | 0.8472 |
| cluster2916 | -0.56 | 0.8472 |
| cluster3098 | -0.32 | 0.8472 |
| cluster3155 | 0.54 | 0.8472 |
| cluster3931 | -0.64 | 0.8472 |
| cluster3268 | -0.40 | 0.8479 |
| cluster3898 | 0.60 | 0.8479 |
| cluster3384 | -0.52 | 0.8519 |
| cluster3521 | -0.41 | 0.8519 |
| cluster3673 | 0.62 | 0.8519 |
| cluster3729 | -0.41 | 0.8519 |
| cluster3751 | 0.38 | 0.8519 |
| cluster3385 | 0.59 | 0.8523 |
| cluster3054 | -0.30 | 0.8531 |
| cluster3154 | 0.37 | 0.8531 |
| cluster2981 | 0.38 | 0.8542 |
| cluster3311 | 0.46 | 0.8544 |
| cluster3071 | -0.44 | 0.8553 |
| cluster3219 | 0.28 | 0.8553 |
| cluster3256 | -0.34 | 0.8553 |
| cluster3257 | 0.36 | 0.8553 |
| cluster3313 | -0.45 | 0.8553 |
| cluster3565 | -0.49 | 0.8553 |
| cluster3687 | 0.51 | 0.8553 |
| cluster3706 | 0.32 | 0.8553 |
| cluster3720 | 0.56 | 0.8553 |
| cluster3762 | -0.56 | 0.8553 |
| cluster3897 | 0.54 | 0.8553 |
| cluster3929 | 0.49 | 0.8553 |

| | | |
|---|---|---|
| cluster3954 | 0.60 | 0.8553 |
| cluster4026 | -0.55 | 0.8553 |
| cluster3323 | -0.56 | 0.8554 |
| cluster3531 | 0.44 | 0.8554 |
| cluster3684 | -0.47 | 0.8554 |
| cluster4053 | -0.44 | 0.8554 |
| cluster2941 | 0.32 | 0.8586 |
| cluster3263 | 0.31 | 0.8592 |
| cluster3041 | -0.35 | 0.8595 |
| cluster2983 | 0.33 | 0.8606 |
| cluster3741 | -0.48 | 0.8619 |
| cluster3888 | -0.54 | 0.8643 |
| cluster3593 | -0.47 | 0.8674 |
| cluster2911 | -0.36 | 0.8675 |
| cluster2989 | -0.48 | 0.8675 |
| cluster3316 | 0.46 | 0.8675 |
| cluster3523 | 0.54 | 0.8675 |
| cluster3586 | -0.36 | 0.8675 |
| cluster3621 | 0.47 | 0.8675 |
| cluster3743 | 0.30 | 0.8675 |
| cluster3022 | -0.43 | 0.8678 |
| cluster3874 | -0.35 | 0.8678 |
| cluster2884 | -0.32 | 0.8694 |
| cluster3011 | 0.46 | 0.8694 |
| cluster3051 | 0.49 | 0.8694 |
| cluster3060 | 0.50 | 0.8694 |
| cluster3134 | -0.30 | 0.8694 |
| cluster3253 | 0.43 | 0.8694 |
| cluster3294 | -0.36 | 0.8694 |
| cluster3304 | -0.39 | 0.8694 |
| cluster3747 | 0.36 | 0.8694 |
| cluster3838 | 0.42 | 0.8694 |
| cluster3961 | 0.55 | 0.8694 |
| cluster3269 | -0.55 | 0.8695 |
| cluster3352 | -0.42 | 0.8695 |
| cluster3529 | 0.42 | 0.8695 |
| cluster3682 | -0.51 | 0.8695 |
| cluster3775 | 0.36 | 0.8695 |
| cluster3147 | -0.33 | 0.8699 |
| cluster3654 | 0.34 | 0.8699 |
| cluster3228 | 0.37 | 0.8701 |
| cluster3833 | 0.48 | 0.8701 |
| cluster3912 | -0.24 | 0.8701 |
| cluster3754 | 0.30 | 0.8754 |

| | | |
|---|---|---|
| cluster3492 | 0.35 | 0.8764 |
| cluster3692 | -0.42 | 0.8778 |
| cluster2883 | -0.31 | 0.8818 |
| cluster2952 | 0.39 | 0.8818 |
| cluster3089 | -0.37 | 0.8818 |
| cluster3165 | -0.44 | 0.8818 |
| cluster3278 | -0.29 | 0.8818 |
| cluster3281 | 0.37 | 0.8818 |
| cluster3547 | -0.34 | 0.8818 |
| cluster3588 | -0.42 | 0.8818 |
| cluster3592 | 0.33 | 0.8818 |
| cluster3612 | 0.45 | 0.8818 |
| cluster3620 | -0.38 | 0.8818 |
| cluster3785 | -0.40 | 0.8818 |
| cluster3886 | 0.38 | 0.8818 |
| cluster3893 | -0.30 | 0.8818 |
| cluster4058 | 0.35 | 0.8818 |
| cluster3150 | -0.46 | 0.8821 |
| cluster3291 | 0.36 | 0.8821 |
| cluster3686 | 0.45 | 0.8821 |
| cluster3823 | -0.37 | 0.8821 |
| cluster4105 | -0.39 | 0.8821 |
| cluster3297 | -0.47 | 0.8822 |
| cluster4126 | 0.53 | 0.8856 |
| cluster3002 | -0.28 | 0.8876 |
| cluster3143 | 0.33 | 0.8876 |
| cluster3244 | 0.33 | 0.8876 |
| cluster3502 | 0.47 | 0.8876 |
| cluster3708 | 0.51 | 0.8876 |
| cluster3861 | -0.25 | 0.8876 |
| cluster3927 | 0.31 | 0.8876 |
| cluster3328 | -0.23 | 0.8925 |
| cluster2965 | -0.30 | 0.8935 |
| cluster3152 | 0.41 | 0.8935 |
| cluster3597 | -0.37 | 0.8935 |
| cluster3992 | -0.38 | 0.8935 |
| cluster3999 | -0.40 | 0.8935 |
| cluster4022 | -0.28 | 0.8935 |
| cluster2882 | -0.27 | 0.8953 |
| cluster2976 | 0.29 | 0.8953 |
| cluster3020 | -0.29 | 0.8953 |
| cluster3170 | -0.31 | 0.8953 |
| cluster3258 | -0.26 | 0.8953 |
| cluster3273 | 0.27 | 0.8953 |

| | | |
|---|---|---|
| cluster3277 | 0.36 | 0.8953 |
| cluster3351 | -0.20 | 0.8953 |
| cluster3483 | 0.23 | 0.8953 |
| cluster3539 | -0.48 | 0.8953 |
| cluster3665 | -0.43 | 0.8953 |
| cluster3671 | 0.30 | 0.8953 |
| cluster3790 | 0.37 | 0.8953 |
| cluster3797 | -0.35 | 0.8953 |
| cluster3799 | 0.33 | 0.8953 |
| cluster3873 | -0.29 | 0.8953 |
| cluster3921 | 0.46 | 0.8953 |
| cluster3959 | 0.43 | 0.8953 |
| cluster3979 | 0.31 | 0.8953 |
| cluster3333 | 0.29 | 0.8974 |
| cluster3533 | 0.32 | 0.8974 |
| cluster3555 | 0.43 | 0.9007 |
| cluster3201 | 0.24 | 0.9082 |
| cluster3716 | -0.37 | 0.9082 |
| cluster3613 | 0.33 | 0.9095 |
| cluster3955 | -0.42 | 0.9095 |
| cluster4086 | 0.36 | 0.9095 |
| cluster2933 | -0.35 | 0.9102 |
| cluster3877 | -0.26 | 0.9103 |
| cluster3324 | -0.27 | 0.9104 |
| cluster4066 | -0.32 | 0.9108 |
| cluster3235 | 0.41 | 0.9118 |
| cluster2929 | -0.21 | 0.9143 |
| cluster4035 | -0.31 | 0.9143 |
| cluster3088 | -0.28 | 0.9149 |
| cluster3216 | -0.26 | 0.9149 |
| cluster3668 | 0.19 | 0.9149 |
| cluster3679 | 0.29 | 0.9155 |
| cluster3266 | -0.23 | 0.9173 |
| cluster3946 | -0.32 | 0.9181 |
| cluster3112 | 0.36 | 0.9192 |
| cluster3561 | -0.30 | 0.9196 |
| cluster2964 | 0.24 | 0.9247 |
| cluster3338 | 0.29 | 0.9247 |
| cluster3506 | 0.19 | 0.9247 |
| cluster3808 | 0.32 | 0.9247 |
| cluster3857 | 0.25 | 0.9247 |
| cluster4041 | -0.30 | 0.9253 |
| cluster3359 | -0.29 | 0.9301 |
| cluster3458 | 0.21 | 0.9301 |

| | | |
|---|---|---|
| cluster3577 | -0.26 | 0.9301 |
| cluster3798 | 0.24 | 0.9301 |
| cluster3926 | -0.40 | 0.9301 |
| cluster3984 | -0.40 | 0.9301 |
| cluster3989 | -0.28 | 0.9301 |
| cluster3948 | -0.23 | 0.9303 |
| cluster3329 | 0.34 | 0.9315 |
| cluster3527 | 0.29 | 0.9315 |
| cluster2896 | -0.18 | 0.9319 |
| cluster3296 | -0.26 | 0.9319 |
| cluster3907 | -0.34 | 0.9319 |
| cluster2897 | -0.22 | 0.9327 |
| cluster2909 | 0.16 | 0.9327 |
| cluster3669 | -0.21 | 0.9327 |
| cluster3746 | 0.25 | 0.9327 |
| cluster3782 | -0.21 | 0.9346 |
| cluster3248 | -0.31 | 0.9364 |
| cluster3789 | 0.27 | 0.9367 |
| cluster3549 | 0.22 | 0.9378 |
| cluster3713 | 0.17 | 0.9378 |
| cluster3320 | -0.23 | 0.9386 |
| cluster3048 | -0.17 | 0.9407 |
| cluster3420 | 0.20 | 0.9407 |
| cluster3670 | -0.28 | 0.9407 |
| cluster2871 | 0.16 | 0.9420 |
| cluster3001 | -0.18 | 0.9420 |
| cluster3308 | -0.18 | 0.9420 |
| cluster3770 | 0.40 | 0.9420 |
| cluster2970 | 0.13 | 0.9428 |
| cluster3104 | -0.13 | 0.9428 |
| cluster3284 | -0.16 | 0.9428 |
| cluster3695 | -0.20 | 0.9428 |
| cluster3990 | 0.20 | 0.9428 |
| cluster4092 | 0.27 | 0.9428 |
| cluster2921 | -0.21 | 0.9444 |
| cluster2943 | 0.22 | 0.9444 |
| cluster3116 | 0.14 | 0.9444 |
| cluster3148 | -0.13 | 0.9444 |
| cluster3354 | 0.15 | 0.9444 |
| cluster3589 | -0.16 | 0.9444 |
| cluster3596 | -0.20 | 0.9444 |
| cluster3605 | 0.18 | 0.9444 |
| cluster3663 | -0.23 | 0.9444 |
| cluster3733 | 0.21 | 0.9444 |

| | | |
|---|---|---|
| cluster3736 | -0.24 | 0.9444 |
| cluster3764 | -0.26 | 0.9444 |
| cluster3816 | -0.23 | 0.9444 |
| cluster4007 | -0.13 | 0.9444 |
| cluster2919 | -0.18 | 0.9449 |
| cluster3156 | -0.19 | 0.9451 |
| cluster4043 | -0.26 | 0.9451 |
| cluster3113 | -0.15 | 0.9465 |
| cluster3406 | -0.18 | 0.9466 |
| cluster2857 | -0.14 | 0.9473 |
| cluster2926 | -0.19 | 0.9473 |
| cluster3059 | 0.23 | 0.9473 |
| cluster3447 | 0.19 | 0.9473 |
| cluster3858 | -0.22 | 0.9473 |
| cluster3409 | 0.23 | 0.9497 |
| cluster3749 | -0.25 | 0.9497 |
| cluster3386 | 0.20 | 0.9499 |
| cluster3922 | -0.26 | 0.9522 |
| cluster4103 | -0.23 | 0.9522 |
| cluster3884 | -0.22 | 0.9525 |
| cluster3314 | -0.20 | 0.9532 |
| cluster3159 | -0.12 | 0.9541 |
| cluster3526 | -0.12 | 0.9545 |
| cluster3096 | -0.12 | 0.9557 |
| cluster3302 | 0.15 | 0.9557 |
| cluster3495 | 0.14 | 0.9557 |
| cluster3091 | -0.12 | 0.9579 |
| cluster2861 | -0.18 | 0.9580 |
| cluster3784 | 0.15 | 0.9580 |
| cluster3221 | 0.17 | 0.9583 |
| cluster3712 | 0.15 | 0.9583 |
| cluster3819 | 0.21 | 0.9583 |
| cluster3600 | 0.12 | 0.9586 |
| cluster3805 | -0.15 | 0.9598 |
| cluster3779 | -0.16 | 0.9616 |
| cluster3106 | 0.12 | 0.9630 |
| cluster3499 | 0.16 | 0.9630 |
| cluster4032 | -0.14 | 0.9630 |
| cluster3312 | -0.21 | 0.9646 |
| cluster2870 | -0.12 | 0.9647 |
| cluster3044 | 0.17 | 0.9647 |
| cluster3067 | -0.12 | 0.9647 |
| cluster3372 | -0.12 | 0.9647 |
| cluster3883 | -0.21 | 0.9647 |

| | | |
|---|---|---|
| cluster4039 | 0.15 | 0.9647 |
| cluster4096 | 0.11 | 0.9647 |
| cluster3309 | -0.16 | 0.9648 |
| cluster3283 | 0.11 | 0.9651 |
| cluster4059 | 0.19 | 0.9665 |
| cluster2956 | -0.10 | 0.9671 |
| cluster2979 | 0.09 | 0.9671 |
| cluster2985 | 0.13 | 0.9671 |
| cluster2996 | -0.10 | 0.9671 |
| cluster3036 | 0.11 | 0.9671 |
| cluster3058 | -0.08 | 0.9671 |
| cluster3064 | 0.13 | 0.9671 |
| cluster3120 | 0.07 | 0.9671 |
| cluster3125 | 0.08 | 0.9671 |
| cluster3174 | 0.14 | 0.9671 |
| cluster3186 | -0.12 | 0.9671 |
| cluster3224 | -0.09 | 0.9671 |
| cluster3300 | 0.10 | 0.9671 |
| cluster3307 | -0.10 | 0.9671 |
| cluster3361 | -0.11 | 0.9671 |
| cluster3452 | 0.14 | 0.9671 |
| cluster3498 | -0.12 | 0.9671 |
| cluster3511 | 0.14 | 0.9671 |
| cluster3541 | -0.11 | 0.9671 |
| cluster3637 | 0.07 | 0.9671 |
| cluster3851 | -0.12 | 0.9671 |
| cluster3887 | -0.15 | 0.9671 |
| cluster3896 | 0.13 | 0.9671 |
| cluster3949 | 0.13 | 0.9671 |
| cluster4005 | -0.13 | 0.9671 |
| cluster4106 | -0.16 | 0.9671 |
| cluster3013 | -0.10 | 0.9673 |
| cluster2991 | 0.07 | 0.9674 |
| cluster2999 | -0.10 | 0.9674 |
| cluster3145 | 0.08 | 0.9674 |
| cluster3167 | 0.12 | 0.9674 |
| cluster3183 | -0.09 | 0.9674 |
| cluster3295 | -0.07 | 0.9674 |
| cluster3335 | -0.10 | 0.9674 |
| cluster3341 | 0.10 | 0.9674 |
| cluster3394 | -0.08 | 0.9674 |
| cluster3412 | -0.09 | 0.9674 |
| cluster3548 | 0.08 | 0.9674 |
| cluster3562 | -0.09 | 0.9674 |

| | | |
|---|---|---|
| cluster3935 | 0.11 | 0.9674 |
| cluster4025 | -0.08 | 0.9674 |
| cluster4063 | -0.15 | 0.9674 |
| cluster3769 | 0.07 | 0.9693 |
| cluster3648 | 0.08 | 0.9700 |
| cluster3731 | 0.10 | 0.9701 |
| cluster2968 | -0.06 | 0.9709 |
| cluster3018 | 0.06 | 0.9709 |
| cluster3744 | 0.07 | 0.9709 |
| cluster3903 | -0.07 | 0.9709 |
| cluster3993 | 0.06 | 0.9724 |
| cluster3225 | 0.09 | 0.9738 |
| cluster3353 | 0.05 | 0.9741 |
| cluster3276 | -0.06 | 0.9768 |
| cluster3900 | -0.11 | 0.9768 |
| cluster3173 | 0.08 | 0.9778 |
| cluster3722 | 0.07 | 0.9778 |
| cluster2898 | 0.07 | 0.9799 |
| cluster3655 | -0.05 | 0.9799 |
| cluster4102 | -0.06 | 0.9808 |
| cluster3737 | -0.04 | 0.9814 |
| cluster3835 | -0.06 | 0.9814 |
| cluster3343 | -0.04 | 0.9820 |
| cluster3399 | -0.06 | 0.9820 |
| cluster3238 | -0.03 | 0.9838 |
| cluster3184 | 0.04 | 0.9838 |
| cluster3794 | -0.05 | 0.9838 |
| cluster3132 | -0.04 | 0.9839 |
| cluster3937 | 0.05 | 0.9849 |
| cluster3423 | -0.05 | 0.9887 |
| cluster3847 | 0.04 | 0.9887 |
| cluster3882 | -0.04 | 0.9897 |
| cluster3055 | 0.03 | 0.9902 |
| cluster3707 | 0.04 | 0.9902 |
| cluster2863 | -0.02 | 0.9908 |
| cluster2917 | -0.03 | 0.9908 |
| cluster3149 | 0.02 | 0.9908 |
| cluster3251 | 0.02 | 0.9908 |
| cluster3378 | -0.03 | 0.9908 |
| cluster4003 | 0.04 | 0.9908 |
| cluster3034 | -0.02 | 0.9929 |
| cluster3070 | -0.02 | 0.9929 |
| cluster3115 | -0.02 | 0.9929 |
| cluster3139 | -0.02 | 0.9929 |

| | | |
|---|---|---|
| cluster3280 | 0.01 | 0.9929 |
| cluster3402 | 0.02 | 0.9929 |
| cluster3470 | -0.01 | 0.9929 |
| cluster3554 | 0.02 | 0.9929 |
| cluster3557 | -0.02 | 0.9929 |
| cluster3806 | -0.03 | 0.9929 |
| cluster3876 | -0.02 | 0.9929 |
| cluster2949 | -0.01 | 0.9951 |
| cluster3392 | -0.01 | 0.9951 |
| cluster3583 | -0.01 | 0.9951 |
| cluster3623 | 0.01 | 0.9978 |
| cluster3691 | 0.00 | 0.9991 |
| cluster3133 | 0.00 | 0.9999 |
| cluster3827 | 0.00 | 0.9999 |

**Supplementary Table 12**. edgeR results with eXpress counts - Dermic vs nasopharyngeal

| ID | log2FoldChange | Adjusted p value |
|---|---|---|
| cluster3845 | -11.42 | 2.43E-09 |
| cluster4087 | 5.24 | 9.60E-08 |
| cluster4097 | -10.23 | 2.40E-07 |
| cluster3666 | 4.45 | 1.09E-06 |
| cluster4049 | -10.83 | 1.93E-05 |
| cluster3544 | -6.68 | 2.60E-05 |
| cluster3510 | -7.68 | 9.36E-05 |
| cluster3753 | -6.01 | 0.0001 |
| cluster3901 | 3.60 | 0.0002 |
| cluster4011 | -6.47 | 0.0004 |
| cluster3981 | 3.30 | 0.0006 |
| cluster4080 | -6.27 | 0.0006 |
| cluster3919 | -5.12 | 0.0006 |
| cluster3306 | 3.55 | 0.0010 |
| cluster3853 | 3.08 | 0.0017 |
| cluster4114 | 4.69 | 0.0019 |
| cluster4119 | 3.95 | 0.0021 |
| cluster3474 | 3.26 | 0.0024 |
| cluster4067 | 4.61 | 0.0033 |
| cluster4073 | 3.80 | 0.0056 |
| cluster3925 | -11.01 | 0.0059 |
| cluster4002 | 2.91 | 0.0109 |
| cluster3250 | 3.08 | 0.0198 |
| cluster3629 | 2.75 | 0.0201 |
| cluster4085 | -3.76 | 0.0311 |
| cluster4074 | 3.20 | 0.0353 |
| cluster4046 | -5.59 | 0.0361 |
| cluster2869 | -3.83 | 0.0414 |
| cluster4047 | 3.34 | 0.0435 |
| cluster2894 | -3.95 | 0.0435 |
| cluster3443 | 2.46 | 0.0452 |
| cluster3869 | 3.06 | 0.0483 |
| cluster3983 | 2.77 | 0.0485 |
| cluster3745 | 3.12 | 0.0503 |
| cluster3430 | 3.16 | 0.0503 |
| cluster3436 | 3.10 | 0.0516 |
| cluster4107 | 2.95 | 0.0595 |
| cluster3970 | 2.72 | 0.0595 |
| cluster4099 | 2.28 | 0.0615 |
| cluster3800 | -3.20 | 0.0682 |
| cluster4060 | 2.62 | 0.0682 |
| cluster3865 | 2.91 | 0.0697 |

| | | |
|---|---|---|
| cluster3780 | -5.88 | 0.0707 |
| cluster3824 | 2.66 | 0.0746 |
| cluster3288 | 2.31 | 0.0805 |
| cluster3431 | 2.15 | 0.0807 |
| cluster3952 | 2.15 | 0.0807 |
| cluster4016 | 2.41 | 0.0807 |
| cluster3902 | 2.09 | 0.0811 |
| cluster3008 | 2.14 | 0.0858 |
| cluster3438 | 2.25 | 0.0910 |
| cluster2928 | 2.11 | 0.0910 |
| cluster3534 | -3.17 | 0.0981 |
| cluster4093 | 2.90 | 0.0998 |
| cluster3179 | 2.46 | 0.0998 |
| cluster4121 | 2.34 | 0.1031 |
| cluster3326 | -2.93 | 0.1044 |
| cluster4054 | 2.44 | 0.1064 |
| cluster3463 | 2.18 | 0.1145 |
| cluster3442 | 2.44 | 0.1201 |
| cluster3889 | 2.67 | 0.1303 |
| cluster3556 | 2.23 | 0.1346 |
| cluster3162 | 2.16 | 0.1469 |
| cluster3719 | 2.32 | 0.1475 |
| cluster3875 | 1.93 | 0.1475 |
| cluster3365 | 2.16 | 0.1681 |
| cluster4070 | -5.33 | 0.1681 |
| cluster3839 | -3.62 | 0.1681 |
| cluster3650 | -3.21 | 0.1681 |
| cluster3270 | 2.28 | 0.1681 |
| cluster3076 | 1.98 | 0.1681 |
| cluster4034 | -5.01 | 0.1724 |
| cluster4064 | 2.06 | 0.1724 |
| cluster3471 | 2.04 | 0.1724 |
| cluster3481 | -3.69 | 0.1752 |
| cluster2912 | 1.83 | 0.1858 |
| cluster3611 | 2.45 | 0.1880 |
| cluster4045 | 2.55 | 0.1880 |
| cluster3598 | 2.91 | 0.1885 |
| cluster3580 | 2.38 | 0.2054 |
| cluster3569 | 2.09 | 0.2054 |
| cluster3374 | 2.27 | 0.2054 |
| cluster3504 | -3.25 | 0.2054 |
| cluster2904 | -2.80 | 0.2083 |
| cluster3771 | 1.99 | 0.2083 |
| cluster3057 | -2.74 | 0.2083 |

| | | |
|---|---|---|
| cluster3932 | -4.09 | 0.2083 |
| cluster3924 | 2.08 | 0.2146 |
| cluster2876 | 2.50 | 0.2146 |
| cluster3572 | 2.76 | 0.2173 |
| cluster3379 | -2.41 | 0.2222 |
| cluster4044 | 1.92 | 0.2222 |
| cluster3568 | -3.25 | 0.2227 |
| cluster3100 | 2.38 | 0.2299 |
| cluster3724 | 2.04 | 0.2299 |
| cluster3822 | 2.46 | 0.2312 |
| cluster4015 | -2.33 | 0.2409 |
| cluster3815 | 2.31 | 0.2576 |
| cluster3012 | 2.13 | 0.2577 |
| cluster3508 | 2.25 | 0.2577 |
| cluster3734 | 2.01 | 0.2613 |
| cluster4090 | -4.56 | 0.2696 |
| cluster3968 | 1.97 | 0.2696 |
| cluster3052 | -3.27 | 0.2696 |
| cluster3111 | -2.36 | 0.2696 |
| cluster3009 | -2.35 | 0.2696 |
| cluster3974 | 2.54 | 0.2696 |
| cluster3683 | 1.69 | 0.2776 |
| cluster3804 | 1.82 | 0.2776 |
| cluster3975 | 1.65 | 0.2776 |
| cluster3871 | 1.77 | 0.2776 |
| cluster3024 | -2.51 | 0.2776 |
| cluster3325 | -3.99 | 0.2811 |
| cluster3576 | 1.75 | 0.2811 |
| cluster3957 | 1.84 | 0.2811 |
| cluster4051 | 2.04 | 0.2811 |
| cluster4100 | 2.06 | 0.2841 |
| cluster4079 | 1.88 | 0.2854 |
| cluster3451 | 2.88 | 0.2907 |
| cluster3657 | 1.64 | 0.2907 |
| cluster3342 | 1.85 | 0.2907 |
| cluster3936 | 1.92 | 0.2996 |
| cluster4081 | 1.57 | 0.3064 |
| cluster3702 | 1.72 | 0.3150 |
| cluster3363 | 1.60 | 0.3279 |
| cluster2900 | -2.55 | 0.3279 |
| cluster3971 | 1.85 | 0.3279 |
| cluster4024 | 1.70 | 0.3279 |
| cluster3681 | 1.78 | 0.3279 |
| cluster3332 | 1.69 | 0.3412 |

| | | |
|---|---|---|
| cluster3940 | 1.64 | 0.3607 |
| cluster4037 | 1.53 | 0.3607 |
| cluster3393 | -3.12 | 0.3607 |
| cluster4125 | -4.71 | 0.3620 |
| cluster2966 | 1.72 | 0.3620 |
| cluster3639 | 2.30 | 0.3620 |
| cluster3617 | -2.80 | 0.3620 |
| cluster3229 | -2.39 | 0.3620 |
| cluster2995 | 1.64 | 0.3620 |
| cluster3742 | 2.00 | 0.3620 |
| cluster3584 | 2.00 | 0.3620 |
| cluster3482 | -2.15 | 0.3620 |
| cluster3581 | -2.17 | 0.3620 |
| cluster2974 | -2.23 | 0.3620 |
| cluster2930 | 1.53 | 0.3620 |
| cluster3003 | -2.22 | 0.3620 |
| cluster3441 | 1.50 | 0.3620 |
| cluster4109 | 1.93 | 0.3643 |
| cluster3866 | 1.96 | 0.3657 |
| cluster3638 | 1.92 | 0.3763 |
| cluster3097 | 1.63 | 0.3763 |
| cluster3207 | 1.64 | 0.3763 |
| cluster3290 | -2.41 | 0.3828 |
| cluster3045 | 1.62 | 0.3840 |
| cluster3710 | 1.64 | 0.3840 |
| cluster3678 | 1.47 | 0.3854 |
| cluster3432 | -2.17 | 0.3854 |
| cluster2887 | 1.53 | 0.3869 |
| cluster3610 | 1.82 | 0.3869 |
| cluster2925 | -2.17 | 0.4004 |
| cluster2892 | 1.45 | 0.4037 |
| cluster3223 | -2.23 | 0.4037 |
| cluster3160 | -1.77 | 0.4037 |
| cluster3105 | -2.58 | 0.4037 |
| cluster3761 | -2.64 | 0.4044 |
| cluster3279 | 1.52 | 0.4044 |
| cluster2915 | -2.04 | 0.4044 |
| cluster3880 | -3.43 | 0.4044 |
| cluster3109 | 1.54 | 0.4113 |
| cluster3917 | 1.72 | 0.4136 |
| cluster3475 | 1.50 | 0.4136 |
| cluster3661 | 1.41 | 0.4136 |
| cluster3424 | -3.16 | 0.4136 |
| cluster3489 | 1.54 | 0.4455 |

| | | |
|---|---|---|
| cluster3956 | 1.45 | 0.4455 |
| cluster4008 | 1.75 | 0.4455 |
| cluster3552 | 1.34 | 0.4455 |
| cluster3161 | -1.84 | 0.4455 |
| cluster3994 | -2.71 | 0.4455 |
| cluster2992 | -1.87 | 0.4497 |
| cluster2972 | -1.95 | 0.4600 |
| cluster3236 | 1.48 | 0.4605 |
| cluster2922 | 1.52 | 0.4605 |
| cluster4113 | -1.73 | 0.4605 |
| cluster3545 | 1.45 | 0.4605 |
| cluster3434 | -2.11 | 0.4605 |
| cluster2899 | -2.26 | 0.4605 |
| cluster3383 | -2.93 | 0.4605 |
| cluster3461 | -2.91 | 0.4726 |
| cluster3512 | 1.58 | 0.4780 |
| cluster2980 | -2.65 | 0.4805 |
| cluster3652 | 1.61 | 0.4895 |
| cluster4000 | 1.48 | 0.4895 |
| cluster3787 | -2.31 | 0.4895 |
| cluster2969 | 1.53 | 0.4895 |
| cluster3520 | -3.17 | 0.4895 |
| cluster3168 | 1.39 | 0.4895 |
| cluster3486 | -2.02 | 0.4895 |
| cluster4091 | -2.96 | 0.4895 |
| cluster3604 | 1.73 | 0.4895 |
| cluster3985 | 1.44 | 0.4895 |
| cluster3292 | -2.19 | 0.4895 |
| cluster4071 | 1.85 | 0.4895 |
| cluster3507 | -1.96 | 0.4895 |
| cluster3772 | -1.90 | 0.4895 |
| cluster3813 | 1.64 | 0.4895 |
| cluster2856 | 1.50 | 0.4895 |
| cluster3408 | -2.12 | 0.4895 |
| cluster3366 | 1.71 | 0.4895 |
| cluster3418 | 1.41 | 0.4895 |
| cluster4029 | -2.80 | 0.4895 |
| cluster3357 | 1.32 | 0.4895 |
| cluster3172 | -1.89 | 0.4895 |
| cluster3503 | 1.32 | 0.4895 |
| cluster3603 | 1.46 | 0.4895 |
| cluster3242 | 1.37 | 0.4895 |
| cluster3063 | 1.52 | 0.4895 |
| cluster3812 | -2.21 | 0.4895 |

| | | |
|---|---|---|
| cluster2954 | -1.87 | 0.4895 |
| cluster3171 | -2.00 | 0.4895 |
| cluster2987 | 1.23 | 0.4944 |
| cluster3571 | -2.34 | 0.4944 |
| cluster3094 | 1.70 | 0.5027 |
| cluster3349 | 1.44 | 0.5053 |
| cluster3642 | 1.51 | 0.5082 |
| cluster3694 | -1.69 | 0.5108 |
| cluster3801 | 1.63 | 0.5138 |
| cluster3715 | 1.52 | 0.5143 |
| cluster3563 | -1.78 | 0.5190 |
| cluster3899 | 1.36 | 0.5190 |
| cluster3519 | -2.83 | 0.5238 |
| cluster3675 | -1.51 | 0.5238 |
| cluster3676 | 1.67 | 0.5241 |
| cluster3987 | 1.44 | 0.5241 |
| cluster4010 | 1.59 | 0.5241 |
| cluster4104 | 1.63 | 0.5319 |
| cluster4038 | 1.63 | 0.5357 |
| cluster3081 | -1.62 | 0.5414 |
| cluster3646 | 1.70 | 0.5414 |
| cluster4069 | 1.37 | 0.5414 |
| cluster3783 | 1.65 | 0.5414 |
| cluster3422 | -2.59 | 0.5414 |
| cluster3381 | -1.69 | 0.5414 |
| cluster3849 | 1.30 | 0.5414 |
| cluster3947 | 1.22 | 0.5414 |
| cluster4110 | 1.55 | 0.5414 |
| cluster3153 | 1.12 | 0.5414 |
| cluster4009 | 1.48 | 0.5414 |
| cluster4084 | -1.65 | 0.5414 |
| cluster3416 | 1.69 | 0.5414 |
| cluster3832 | 1.29 | 0.5414 |
| cluster3135 | -1.64 | 0.5414 |
| cluster4065 | 1.23 | 0.5414 |
| cluster3190 | 1.70 | 0.5437 |
| cluster4083 | 1.60 | 0.5456 |
| cluster4014 | 1.40 | 0.5515 |
| cluster3953 | -1.51 | 0.5515 |
| cluster2927 | 1.47 | 0.5515 |
| cluster2942 | 1.24 | 0.5515 |
| cluster3633 | -1.54 | 0.5515 |
| cluster2936 | 1.59 | 0.5515 |
| cluster3752 | -2.52 | 0.5515 |

| | | |
|---|---|---|
| cluster3628 | 1.37 | 0.5515 |
| cluster3831 | 1.31 | 0.5515 |
| cluster4068 | 1.55 | 0.5515 |
| cluster3538 | -1.97 | 0.5515 |
| cluster3674 | -2.92 | 0.5525 |
| cluster3387 | 1.19 | 0.5538 |
| cluster3110 | 1.15 | 0.5642 |
| cluster3017 | -1.88 | 0.5642 |
| cluster3537 | 1.21 | 0.5642 |
| cluster4057 | 1.53 | 0.5642 |
| cluster2854 | 1.30 | 0.5642 |
| cluster3693 | 1.63 | 0.5642 |
| cluster3330 | 1.23 | 0.5651 |
| cluster3740 | 1.43 | 0.5684 |
| cluster3624 | 1.12 | 0.5684 |
| cluster3590 | 1.33 | 0.5686 |
| cluster3939 | 1.25 | 0.5686 |
| cluster3467 | 1.15 | 0.5736 |
| cluster3909 | 1.19 | 0.5736 |
| cluster3647 | 1.17 | 0.5762 |
| cluster3142 | -1.85 | 0.5775 |
| cluster3570 | 1.57 | 0.5784 |
| cluster3595 | 1.21 | 0.5784 |
| cluster2975 | 1.05 | 0.5784 |
| cluster3543 | -2.15 | 0.5784 |
| cluster3836 | 1.22 | 0.5784 |
| cluster3781 | 1.28 | 0.5784 |
| cluster3643 | 1.47 | 0.5784 |
| cluster3453 | -1.68 | 0.5784 |
| cluster3287 | -1.73 | 0.5784 |
| cluster4019 | 1.47 | 0.5784 |
| cluster3574 | -2.36 | 0.5784 |
| cluster3040 | -1.55 | 0.5784 |
| cluster3872 | 1.33 | 0.5784 |
| cluster3010 | -1.54 | 0.5784 |
| cluster3129 | -2.62 | 0.5784 |
| cluster3192 | -1.43 | 0.5784 |
| cluster3995 | -1.98 | 0.5784 |
| cluster3274 | 1.31 | 0.5784 |
| cluster3814 | -1.62 | 0.5784 |
| cluster3233 | -2.64 | 0.5784 |
| cluster3837 | 1.32 | 0.5784 |
| cluster3913 | -1.81 | 0.5784 |
| cluster3651 | -2.50 | 0.5784 |

| | | |
|---|---|---|
| cluster3220 | -1.57 | 0.5784 |
| cluster3484 | -2.16 | 0.5784 |
| cluster3487 | -1.60 | 0.5784 |
| cluster2924 | 1.25 | 0.5784 |
| cluster3476 | 1.44 | 0.5784 |
| cluster3778 | -1.66 | 0.5784 |
| cluster2951 | 1.29 | 0.5784 |
| cluster3817 | -2.06 | 0.5784 |
| cluster3944 | 1.17 | 0.5784 |
| cluster3254 | -2.00 | 0.5784 |
| cluster3645 | -3.02 | 0.5784 |
| cluster3090 | -2.38 | 0.5784 |
| cluster3703 | 1.08 | 0.5784 |
| cluster3401 | 1.03 | 0.5784 |
| cluster3187 | -2.12 | 0.5784 |
| cluster3469 | -1.76 | 0.5812 |
| cluster3698 | -1.42 | 0.5877 |
| cluster3895 | 1.41 | 0.5877 |
| cluster3473 | 1.28 | 0.5877 |
| cluster4120 | -2.66 | 0.5877 |
| cluster4062 | -2.45 | 0.5877 |
| cluster4122 | -2.63 | 0.5877 |
| cluster2990 | 1.03 | 0.5995 |
| cluster3943 | 1.42 | 0.6007 |
| cluster2908 | -1.48 | 0.6019 |
| cluster4027 | 1.39 | 0.6019 |
| cluster3828 | 1.25 | 0.6019 |
| cluster3677 | 1.12 | 0.6019 |
| cluster4028 | 1.12 | 0.6063 |
| cluster3830 | 1.05 | 0.6063 |
| cluster3614 | 1.52 | 0.6063 |
| cluster3879 | 1.09 | 0.6090 |
| cluster3892 | -1.77 | 0.6090 |
| cluster3177 | -1.81 | 0.6095 |
| cluster3542 | 1.37 | 0.6095 |
| cluster2855 | 0.99 | 0.6095 |
| cluster3083 | 1.04 | 0.6106 |
| cluster3247 | 1.16 | 0.6106 |
| cluster3904 | -1.61 | 0.6106 |
| cluster3095 | -1.46 | 0.6106 |
| cluster3933 | -1.54 | 0.6106 |
| cluster3766 | -1.49 | 0.6106 |
| cluster3905 | -3.11 | 0.6106 |
| cluster3855 | 1.30 | 0.6106 |

| | | |
|---|---|---|
| cluster4117 | 1.73 | 0.6106 |
| cluster3793 | 1.02 | 0.6106 |
| cluster3864 | 1.03 | 0.6106 |
| cluster3252 | 1.54 | 0.6106 |
| cluster3704 | 1.17 | 0.6106 |
| cluster3035 | -1.26 | 0.6106 |
| cluster3210 | 0.97 | 0.6106 |
| cluster4031 | -1.59 | 0.6106 |
| cluster3997 | 1.33 | 0.6106 |
| cluster3685 | -1.69 | 0.6106 |
| cluster3725 | 1.32 | 0.6106 |
| cluster3217 | -1.47 | 0.6106 |
| cluster3337 | -1.60 | 0.6106 |
| cluster3951 | 1.30 | 0.6106 |
| cluster4088 | 1.29 | 0.6106 |
| cluster3699 | 1.31 | 0.6106 |
| cluster3501 | -1.78 | 0.6106 |
| cluster3885 | -1.39 | 0.6106 |
| cluster3065 | -1.36 | 0.6106 |
| cluster3038 | 1.17 | 0.6106 |
| cluster3788 | -1.39 | 0.6106 |
| cluster3084 | -1.25 | 0.6106 |
| cluster3980 | 0.96 | 0.6106 |
| cluster3395 | -1.35 | 0.6106 |
| cluster3398 | -1.23 | 0.6106 |
| cluster2867 | -1.95 | 0.6143 |
| cluster4056 | -1.57 | 0.6211 |
| cluster3460 | 1.17 | 0.6215 |
| cluster3457 | -1.69 | 0.6215 |
| cluster4018 | 1.19 | 0.6215 |
| cluster3272 | 1.76 | 0.6215 |
| cluster3117 | 1.37 | 0.6225 |
| cluster2877 | 0.93 | 0.6248 |
| cluster2914 | -1.47 | 0.6298 |
| cluster4036 | 0.98 | 0.6303 |
| cluster3370 | 1.11 | 0.6319 |
| cluster3723 | 0.93 | 0.6340 |
| cluster3867 | -1.18 | 0.6340 |
| cluster3029 | 0.93 | 0.6378 |
| cluster3998 | 1.35 | 0.6381 |
| cluster3198 | 1.05 | 0.6406 |
| cluster3619 | 1.03 | 0.6406 |
| cluster3016 | -1.98 | 0.6426 |
| cluster3930 | 1.02 | 0.6426 |

| | | |
|---|---|---|
| cluster3454 | -1.24 | 0.6426 |
| cluster3000 | -1.41 | 0.6427 |
| cluster3920 | -1.66 | 0.6427 |
| cluster3356 | 1.05 | 0.6439 |
| cluster3410 | -1.63 | 0.6451 |
| cluster3950 | 1.00 | 0.6453 |
| cluster3493 | 1.23 | 0.6453 |
| cluster3891 | 1.10 | 0.6461 |
| cluster3560 | 1.24 | 0.6461 |
| cluster3146 | -1.22 | 0.6461 |
| cluster3911 | 1.31 | 0.6461 |
| cluster3464 | 1.08 | 0.6464 |
| cluster4111 | -2.27 | 0.6482 |
| cluster3714 | -1.65 | 0.6503 |
| cluster2890 | -1.17 | 0.6508 |
| cluster3086 | -1.22 | 0.6521 |
| cluster3389 | 0.98 | 0.6541 |
| cluster2864 | 0.94 | 0.6552 |
| cluster3042 | 1.06 | 0.6616 |
| cluster3688 | 0.97 | 0.6616 |
| cluster4082 | 1.24 | 0.6616 |
| cluster3455 | -1.58 | 0.6616 |
| cluster3916 | 0.95 | 0.6616 |
| cluster2865 | 0.94 | 0.6673 |
| cluster2967 | 1.11 | 0.6673 |
| cluster3618 | -2.23 | 0.6748 |
| cluster2977 | 0.93 | 0.6748 |
| cluster3488 | -1.14 | 0.6748 |
| cluster3829 | 1.18 | 0.6748 |
| cluster3462 | -1.24 | 0.6748 |
| cluster3634 | -1.33 | 0.6748 |
| cluster3194 | -1.30 | 0.6748 |
| cluster3807 | -1.26 | 0.6748 |
| cluster3373 | 0.98 | 0.6748 |
| cluster2893 | 1.17 | 0.6748 |
| cluster3169 | 1.03 | 0.6748 |
| cluster3382 | -1.15 | 0.6761 |
| cluster3158 | 1.10 | 0.6761 |
| cluster3122 | -1.19 | 0.6761 |
| cluster3966 | 0.87 | 0.6761 |
| cluster3151 | -1.07 | 0.6761 |
| cluster3656 | 1.00 | 0.6761 |
| cluster3625 | 0.91 | 0.6761 |
| cluster4013 | -3.05 | 0.6761 |

| | | |
|---|---|---|
| cluster3982 | 0.91 | 0.6761 |
| cluster3015 | -1.53 | 0.6761 |
| cluster2960 | -1.57 | 0.6774 |
| cluster3218 | 1.13 | 0.6787 |
| cluster2947 | 0.93 | 0.6805 |
| cluster4112 | 1.29 | 0.6805 |
| cluster3131 | -1.16 | 0.6805 |
| cluster3587 | -1.90 | 0.6808 |
| cluster4094 | 1.06 | 0.6894 |
| cluster3466 | -1.27 | 0.6898 |
| cluster3421 | 1.08 | 0.6926 |
| cluster3690 | -1.21 | 0.6926 |
| cluster4012 | 1.05 | 0.6926 |
| cluster3906 | 0.96 | 0.6926 |
| cluster2959 | 0.88 | 0.6926 |
| cluster3594 | -1.41 | 0.6926 |
| cluster3908 | 0.87 | 0.6926 |
| cluster3118 | 0.91 | 0.6926 |
| cluster3180 | -1.07 | 0.6958 |
| cluster3631 | -1.76 | 0.6958 |
| cluster4128 | 1.30 | 0.6958 |
| cluster3941 | -1.10 | 0.6960 |
| cluster3211 | 1.11 | 0.7020 |
| cluster3028 | 0.83 | 0.7038 |
| cluster3601 | 0.96 | 0.7038 |
| cluster3419 | -1.06 | 0.7048 |
| cluster3087 | 0.85 | 0.7048 |
| cluster3680 | 0.86 | 0.7048 |
| cluster3318 | -1.36 | 0.7048 |
| cluster3319 | -1.28 | 0.7048 |
| cluster3212 | -1.36 | 0.7048 |
| cluster3496 | 0.91 | 0.7048 |
| cluster3732 | 0.87 | 0.7048 |
| cluster3293 | 0.84 | 0.7048 |
| cluster3810 | 0.89 | 0.7048 |
| cluster3093 | 0.92 | 0.7048 |
| cluster4077 | 0.95 | 0.7048 |
| cluster4021 | 1.06 | 0.7056 |
| cluster3205 | -1.11 | 0.7056 |
| cluster3862 | 0.78 | 0.7056 |
| cluster3635 | -1.04 | 0.7056 |
| cluster3803 | 1.15 | 0.7056 |
| cluster3776 | -1.34 | 0.7056 |
| cluster3976 | -1.12 | 0.7056 |

| | | |
|---|---|---|
| cluster3536 | 0.83 | 0.7056 |
| cluster2973 | 0.91 | 0.7056 |
| cluster3082 | -1.82 | 0.7056 |
| cluster3427 | -1.42 | 0.7056 |
| cluster3425 | -1.23 | 0.7056 |
| cluster3515 | -1.63 | 0.7056 |
| cluster4116 | -2.11 | 0.7056 |
| cluster3842 | 1.01 | 0.7056 |
| cluster3942 | -1.12 | 0.7056 |
| cluster3299 | 0.96 | 0.7056 |
| cluster3413 | -1.40 | 0.7056 |
| cluster2902 | 1.06 | 0.7060 |
| cluster3497 | -1.17 | 0.7060 |
| cluster3345 | 0.87 | 0.7060 |
| cluster3078 | 0.75 | 0.7060 |
| cluster3860 | 0.92 | 0.7060 |
| cluster3881 | 1.02 | 0.7060 |
| cluster3609 | 1.19 | 0.7060 |
| cluster4095 | 1.07 | 0.7060 |
| cluster3334 | 0.83 | 0.7060 |
| cluster3856 | 0.82 | 0.7070 |
| cluster3193 | -1.05 | 0.7070 |
| cluster2874 | 0.75 | 0.7070 |
| cluster3206 | 0.99 | 0.7087 |
| cluster4072 | 0.80 | 0.7087 |
| cluster3245 | 1.01 | 0.7096 |
| cluster4089 | -1.68 | 0.7096 |
| cluster4030 | 0.90 | 0.7096 |
| cluster3934 | 0.73 | 0.7096 |
| cluster3415 | -1.14 | 0.7096 |
| cluster3267 | 0.80 | 0.7096 |
| cluster3030 | -1.59 | 0.7096 |
| cluster3938 | 0.98 | 0.7098 |
| cluster2984 | -1.30 | 0.7098 |
| cluster2994 | 0.82 | 0.7124 |
| cluster3075 | -1.14 | 0.7127 |
| cluster2888 | 1.14 | 0.7193 |
| cluster2862 | -1.06 | 0.7211 |
| cluster3175 | -1.39 | 0.7230 |
| cluster3140 | 0.73 | 0.7233 |
| cluster2853 | -1.25 | 0.7255 |
| cluster3360 | -1.15 | 0.7255 |
| cluster3202 | 0.79 | 0.7255 |
| cluster3918 | 0.93 | 0.7255 |

| | | |
|---|---|---|
| cluster3468 | -1.16 | 0.7255 |
| cluster3322 | 0.96 | 0.7255 |
| cluster3964 | 1.09 | 0.7255 |
| cluster3371 | -2.37 | 0.7255 |
| cluster3553 | 0.86 | 0.7255 |
| cluster4050 | 0.93 | 0.7255 |
| cluster3606 | 0.99 | 0.7255 |
| cluster3039 | 0.77 | 0.7255 |
| cluster3196 | -0.96 | 0.7275 |
| cluster3265 | -1.06 | 0.7275 |
| cluster3203 | -1.03 | 0.7275 |
| cluster3660 | 1.02 | 0.7275 |
| cluster3185 | -1.22 | 0.7275 |
| cluster3397 | -1.31 | 0.7287 |
| cluster3130 | -1.07 | 0.7287 |
| cluster3996 | 0.77 | 0.7287 |
| cluster3653 | -1.16 | 0.7310 |
| cluster3622 | -0.95 | 0.7310 |
| cluster3607 | 0.87 | 0.7345 |
| cluster3411 | 0.72 | 0.7364 |
| cluster3897 | 0.92 | 0.7386 |
| cluster3259 | -0.97 | 0.7408 |
| cluster3369 | 0.75 | 0.7411 |
| cluster3558 | 0.84 | 0.7411 |
| cluster3962 | 0.73 | 0.7411 |
| cluster3809 | 0.73 | 0.7440 |
| cluster3311 | 0.81 | 0.7472 |
| cluster3385 | 0.92 | 0.7472 |
| cluster3230 | -1.20 | 0.7472 |
| cluster3767 | 0.90 | 0.7472 |
| cluster3898 | 0.95 | 0.7472 |
| cluster3019 | 0.74 | 0.7472 |
| cluster3575 | -1.28 | 0.7476 |
| cluster2988 | 0.69 | 0.7476 |
| cluster3136 | -1.14 | 0.7476 |
| cluster3080 | -1.01 | 0.7476 |
| cluster3978 | -0.92 | 0.7476 |
| cluster3446 | -0.93 | 0.7476 |
| cluster2986 | 0.71 | 0.7476 |
| cluster3735 | -1.09 | 0.7476 |
| cluster3108 | 0.81 | 0.7476 |
| cluster3444 | -0.90 | 0.7476 |
| cluster3972 | 0.74 | 0.7476 |
| cluster3531 | 0.78 | 0.7476 |

| | | |
|---|---|---|
| cluster3275 | -1.35 | 0.7476 |
| cluster3011 | 0.81 | 0.7476 |
| cluster3991 | 0.92 | 0.7476 |
| cluster2889 | 0.70 | 0.7476 |
| cluster3550 | -1.10 | 0.7476 |
| cluster3099 | 0.73 | 0.7476 |
| cluster3257 | 0.69 | 0.7476 |
| cluster2903 | -1.24 | 0.7476 |
| cluster3232 | 0.73 | 0.7476 |
| cluster3114 | -0.88 | 0.7476 |
| cluster3231 | -1.17 | 0.7476 |
| cluster3007 | 0.73 | 0.7476 |
| cluster3641 | -1.02 | 0.7492 |
| cluster3630 | -0.92 | 0.7511 |
| cluster3711 | -0.85 | 0.7511 |
| cluster3775 | 0.71 | 0.7511 |
| cluster3305 | 0.84 | 0.7514 |
| cluster3700 | 0.78 | 0.7527 |
| cluster3491 | 0.71 | 0.7534 |
| cluster3773 | -0.92 | 0.7534 |
| cluster3103 | -1.35 | 0.7534 |
| cluster4061 | 0.86 | 0.7534 |
| cluster3986 | 0.70 | 0.7534 |
| cluster3528 | -0.90 | 0.7534 |
| cluster2913 | -0.86 | 0.7534 |
| cluster3640 | -2.20 | 0.7534 |
| cluster3843 | 0.68 | 0.7545 |
| cluster3154 | 0.68 | 0.7545 |
| cluster3391 | -1.03 | 0.7546 |
| cluster4040 | 0.76 | 0.7564 |
| cluster3929 | 0.81 | 0.7585 |
| cluster3213 | -1.07 | 0.7607 |
| cluster3155 | 0.83 | 0.7607 |
| cluster2941 | 0.64 | 0.7607 |
| cluster3056 | -0.90 | 0.7607 |
| cluster2918 | -0.90 | 0.7607 |
| cluster3182 | -1.04 | 0.7608 |
| cluster3509 | 0.66 | 0.7615 |
| cluster4098 | 0.91 | 0.7615 |
| cluster3720 | 0.85 | 0.7615 |
| cluster3465 | -1.00 | 0.7649 |
| cluster3915 | -1.42 | 0.7649 |
| cluster3796 | -1.38 | 0.7652 |
| cluster2901 | -1.08 | 0.7661 |

| | | |
|---|---|---|
| cluster3518 | -1.35 | 0.7661 |
| cluster3069 | -0.81 | 0.7661 |
| cluster3654 | 0.68 | 0.7661 |
| cluster2982 | -0.97 | 0.7688 |
| cluster4017 | 0.81 | 0.7700 |
| cluster2983 | 0.64 | 0.7714 |
| cluster3060 | 0.82 | 0.7714 |
| cluster3291 | 0.70 | 0.7714 |
| cluster3050 | -0.88 | 0.7714 |
| cluster3178 | 0.86 | 0.7738 |
| cluster2981 | 0.66 | 0.7740 |
| cluster2948 | -1.30 | 0.7740 |
| cluster3945 | -1.17 | 0.7740 |
| cluster3626 | -0.96 | 0.7740 |
| cluster3523 | 0.83 | 0.7740 |
| cluster3301 | -1.04 | 0.7740 |
| cluster3219 | 0.60 | 0.7740 |
| cluster3961 | 0.85 | 0.7740 |
| cluster3439 | -0.90 | 0.7740 |
| cluster3244 | 0.68 | 0.7740 |
| cluster3743 | 0.62 | 0.7740 |
| cluster3263 | 0.62 | 0.7740 |
| cluster3687 | 0.80 | 0.7740 |
| cluster3751 | 0.67 | 0.7740 |
| cluster3706 | 0.62 | 0.7740 |
| cluster3979 | 0.66 | 0.7740 |
| cluster3414 | -0.94 | 0.7740 |
| cluster3253 | 0.73 | 0.7740 |
| cluster3768 | -1.16 | 0.7740 |
| cluster3249 | -0.84 | 0.7740 |
| cluster2957 | -1.14 | 0.7740 |
| cluster3844 | -1.55 | 0.7740 |
| cluster3786 | 0.91 | 0.7748 |
| cluster3673 | 0.86 | 0.7756 |
| cluster3228 | 0.66 | 0.7756 |
| cluster3126 | -1.22 | 0.7760 |
| cluster3281 | 0.68 | 0.7767 |
| cluster3969 | -0.78 | 0.7767 |
| cluster3564 | 0.80 | 0.7767 |
| cluster3747 | 0.64 | 0.7767 |
| cluster3954 | 0.84 | 0.7767 |
| cluster3591 | -0.81 | 0.7767 |
| cluster4058 | 0.66 | 0.7767 |
| cluster3121 | -0.95 | 0.7767 |

| | | |
|---|---|---|
| cluster3143 | 0.65 | 0.7767 |
| cluster3051 | 0.75 | 0.7767 |
| cluster2881 | 0.68 | 0.7767 |
| cluster3811 | -0.87 | 0.7767 |
| cluster3757 | -1.58 | 0.7767 |
| cluster3127 | -0.75 | 0.7767 |
| cluster3621 | 0.75 | 0.7767 |
| cluster3592 | 0.63 | 0.7767 |
| cluster3833 | 0.78 | 0.7767 |
| cluster3483 | 0.57 | 0.7767 |
| cluster3671 | 0.63 | 0.7814 |
| cluster3799 | 0.69 | 0.7814 |
| cluster3367 | -1.31 | 0.7814 |
| cluster3838 | 0.71 | 0.7825 |
| cluster3834 | -1.09 | 0.7825 |
| cluster3754 | 0.62 | 0.7835 |
| cluster3316 | 0.71 | 0.7835 |
| cluster2978 | -0.95 | 0.7855 |
| cluster3886 | 0.66 | 0.7870 |
| cluster3004 | -0.86 | 0.7870 |
| cluster3157 | -0.99 | 0.7870 |
| cluster3679 | 0.64 | 0.7870 |
| cluster3529 | 0.68 | 0.7882 |
| cluster3612 | 0.74 | 0.7896 |
| cluster3533 | 0.65 | 0.7896 |
| cluster3450 | -1.50 | 0.7896 |
| cluster2964 | 0.59 | 0.7896 |
| cluster3730 | -0.89 | 0.7898 |
| cluster3273 | 0.59 | 0.7901 |
| cluster2953 | -0.77 | 0.7903 |
| cluster2976 | 0.60 | 0.7903 |
| cluster4126 | 0.83 | 0.7903 |
| cluster3492 | 0.61 | 0.7903 |
| cluster3774 | -0.88 | 0.7914 |
| cluster3977 | -1.13 | 0.7953 |
| cluster3551 | -1.01 | 0.7953 |
| cluster3686 | 0.75 | 0.7953 |
| cluster2952 | 0.65 | 0.7962 |
| cluster3346 | -0.93 | 0.7962 |
| cluster3613 | 0.67 | 0.7962 |
| cluster3368 | -0.76 | 0.7962 |
| cluster3525 | -0.79 | 0.7962 |
| cluster3555 | 0.75 | 0.7967 |
| cluster3549 | 0.58 | 0.7967 |

| | | |
|---|---|---|
| cluster3277 | 0.69 | 0.7967 |
| cluster3726 | -0.78 | 0.7974 |
| cluster3921 | 0.76 | 0.7981 |
| cluster3066 | 0.60 | 0.8033 |
| cluster3502 | 0.75 | 0.8041 |
| cluster3333 | 0.59 | 0.8042 |
| cluster3458 | 0.55 | 0.8042 |
| cluster3927 | 0.60 | 0.8042 |
| cluster3074 | -0.73 | 0.8042 |
| cluster3435 | -1.18 | 0.8042 |
| cluster3668 | 0.52 | 0.8042 |
| cluster3200 | 0.60 | 0.8050 |
| cluster2895 | -0.81 | 0.8063 |
| cluster3532 | -1.00 | 0.8093 |
| cluster3201 | 0.53 | 0.8114 |
| cluster3315 | -0.80 | 0.8118 |
| cluster3152 | 0.69 | 0.8118 |
| cluster3449 | -0.80 | 0.8118 |
| cluster4055 | -0.85 | 0.8118 |
| cluster3578 | -0.93 | 0.8165 |
| cluster3649 | -0.71 | 0.8165 |
| cluster3209 | -0.78 | 0.8165 |
| cluster3708 | 0.73 | 0.8202 |
| cluster3112 | 0.68 | 0.8202 |
| cluster3959 | 0.70 | 0.8202 |
| cluster3850 | -0.83 | 0.8202 |
| cluster3746 | 0.61 | 0.8202 |
| cluster3506 | 0.50 | 0.8202 |
| cluster3662 | -0.96 | 0.8202 |
| cluster2943 | 0.58 | 0.8202 |
| cluster4033 | -1.19 | 0.8202 |
| cluster3068 | -0.69 | 0.8209 |
| cluster3199 | -0.67 | 0.8234 |
| cluster3672 | -0.90 | 0.8234 |
| cluster3870 | -1.14 | 0.8234 |
| cluster3235 | 0.69 | 0.8283 |
| cluster4086 | 0.63 | 0.8285 |
| cluster2909 | 0.48 | 0.8285 |
| cluster3790 | 0.64 | 0.8308 |
| cluster3073 | -0.65 | 0.8308 |
| cluster2886 | -0.80 | 0.8326 |
| cluster3914 | -1.42 | 0.8326 |
| cluster3852 | 0.63 | 0.8348 |
| cluster3540 | -1.05 | 0.8348 |

| | | |
|---|---|---|
| cluster4052 | 0.63 | 0.8348 |
| cluster3355 | -0.91 | 0.8398 |
| cluster3400 | -0.76 | 0.8399 |
| cluster3841 | -0.88 | 0.8421 |
| cluster3298 | -0.71 | 0.8421 |
| cluster3910 | -1.07 | 0.8421 |
| cluster3923 | -0.68 | 0.8433 |
| cluster4001 | -0.88 | 0.8433 |
| cluster3137 | 0.62 | 0.8433 |
| cluster3527 | 0.60 | 0.8440 |
| cluster3990 | 0.53 | 0.8440 |
| cluster3857 | 0.54 | 0.8442 |
| cluster3808 | 0.60 | 0.8454 |
| cluster3821 | -0.75 | 0.8486 |
| cluster3770 | 0.76 | 0.8486 |
| cluster3226 | -0.85 | 0.8486 |
| cluster3485 | -1.15 | 0.8514 |
| cluster2970 | 0.44 | 0.8538 |
| cluster3338 | 0.56 | 0.8538 |
| cluster3329 | 0.63 | 0.8538 |
| cluster2871 | 0.45 | 0.8538 |
| cluster3513 | -0.78 | 0.8540 |
| cluster3116 | 0.44 | 0.8540 |
| cluster3282 | -0.69 | 0.8540 |
| cluster2963 | -0.82 | 0.8540 |
| cluster3863 | 0.56 | 0.8540 |
| cluster4096 | 0.46 | 0.8543 |
| cluster3106 | 0.45 | 0.8543 |
| cluster3798 | 0.51 | 0.8552 |
| cluster3237 | -1.00 | 0.8552 |
| cluster3386 | 0.53 | 0.8552 |
| cluster3733 | 0.53 | 0.8554 |
| cluster3713 | 0.46 | 0.8554 |
| cluster3494 | -0.78 | 0.8579 |
| cluster3789 | 0.55 | 0.8579 |
| cluster3712 | 0.51 | 0.8582 |
| cluster3354 | 0.45 | 0.8582 |
| cluster3006 | -1.04 | 0.8582 |
| cluster3644 | -0.76 | 0.8582 |
| cluster3759 | -0.76 | 0.8582 |
| cluster3053 | -0.91 | 0.8582 |
| cluster3336 | -0.81 | 0.8582 |
| cluster3327 | -0.79 | 0.8582 |
| cluster3283 | 0.45 | 0.8582 |

| | | |
|---|---|---|
| cluster2866 | -0.84 | 0.8583 |
| cluster3784 | 0.50 | 0.8601 |
| cluster3026 | -0.85 | 0.8601 |
| cluster3420 | 0.47 | 0.8601 |
| cluster3085 | -0.85 | 0.8613 |
| cluster3125 | 0.41 | 0.8613 |
| cluster3738 | -1.13 | 0.8613 |
| cluster3648 | 0.43 | 0.8613 |
| cluster3119 | -0.73 | 0.8613 |
| cluster3600 | 0.42 | 0.8613 |
| cluster3302 | 0.45 | 0.8613 |
| cluster3197 | -0.75 | 0.8613 |
| cluster3495 | 0.44 | 0.8613 |
| cluster2905 | -0.67 | 0.8613 |
| cluster3965 | -0.91 | 0.8613 |
| cluster3605 | 0.46 | 0.8613 |
| cluster3721 | -0.86 | 0.8613 |
| cluster2985 | 0.46 | 0.8615 |
| cluster3377 | -0.71 | 0.8648 |
| cluster3409 | 0.53 | 0.8648 |
| cluster3077 | -0.72 | 0.8648 |
| cluster3637 | 0.40 | 0.8658 |
| cluster3440 | -0.59 | 0.8663 |
| cluster3511 | 0.48 | 0.8717 |
| cluster3447 | 0.48 | 0.8717 |
| cluster3260 | -0.65 | 0.8727 |
| cluster3514 | -0.79 | 0.8745 |
| cluster4092 | 0.54 | 0.8745 |
| cluster3819 | 0.54 | 0.8745 |
| cluster3120 | 0.38 | 0.8745 |
| cluster3221 | 0.46 | 0.8762 |
| cluster3452 | 0.46 | 0.8762 |
| cluster3043 | -0.63 | 0.8762 |
| cluster3285 | -0.64 | 0.8812 |
| cluster3246 | 0.42 | 0.8812 |
| cluster3404 | -0.58 | 0.8812 |
| cluster3405 | -0.62 | 0.8812 |
| cluster2938 | -0.55 | 0.8834 |
| cluster2920 | -0.60 | 0.8834 |
| cluster3548 | 0.40 | 0.8834 |
| cluster4059 | 0.52 | 0.8834 |
| cluster3709 | -0.62 | 0.8835 |
| cluster3429 | -0.64 | 0.8835 |
| cluster3059 | 0.47 | 0.8835 |

| | | |
|---|---|---|
| cluster3064 | 0.45 | 0.8835 |
| cluster4118 | -1.35 | 0.8835 |
| cluster3967 | -0.72 | 0.8835 |
| cluster3777 | -0.64 | 0.8835 |
| cluster3750 | -0.99 | 0.8835 |
| cluster3191 | -1.00 | 0.8835 |
| cluster3317 | -0.76 | 0.8835 |
| cluster2998 | -0.51 | 0.8835 |
| cluster2979 | 0.37 | 0.8835 |
| cluster3344 | -0.71 | 0.8835 |
| cluster3731 | 0.44 | 0.8835 |
| cluster3499 | 0.44 | 0.8835 |
| cluster3145 | 0.39 | 0.8835 |
| cluster3149 | 0.37 | 0.8835 |
| cluster2991 | 0.38 | 0.8835 |
| cluster4039 | 0.43 | 0.8835 |
| cluster3300 | 0.42 | 0.8835 |
| cluster3353 | 0.35 | 0.8835 |
| cluster3189 | -0.70 | 0.8875 |
| cluster3546 | -0.88 | 0.8875 |
| cluster2937 | -0.58 | 0.8877 |
| cluster3018 | 0.35 | 0.8880 |
| cluster3659 | -0.74 | 0.8880 |
| cluster3993 | 0.37 | 0.8897 |
| cluster3234 | -0.58 | 0.8897 |
| cluster3616 | -0.80 | 0.8919 |
| cluster3375 | -0.54 | 0.8930 |
| cluster4023 | -0.78 | 0.8934 |
| cluster3044 | 0.43 | 0.8934 |
| cluster3896 | 0.43 | 0.8944 |
| cluster3055 | 0.35 | 0.8944 |
| cluster3261 | -0.80 | 0.8944 |
| cluster3036 | 0.37 | 0.8944 |
| cluster3583 | 0.32 | 0.8944 |
| cluster3769 | 0.35 | 0.8944 |
| cluster2906 | -0.63 | 0.8944 |
| cluster3195 | -0.55 | 0.8944 |
| cluster3047 | -0.64 | 0.8944 |
| cluster3271 | -0.51 | 0.8944 |
| cluster2898 | 0.39 | 0.8944 |
| cluster3477 | -0.68 | 0.8944 |
| cluster3448 | -0.86 | 0.8944 |
| cluster3470 | 0.32 | 0.8944 |
| cluster3174 | 0.40 | 0.8944 |

| | | |
|---|---|---|
| cluster2949 | 0.33 | 0.8944 |
| cluster3937 | 0.41 | 0.8944 |
| cluster3243 | -0.58 | 0.8944 |
| cluster4076 | -0.91 | 0.8944 |
| cluster3280 | 0.33 | 0.8944 |
| cluster3949 | 0.39 | 0.8944 |
| cluster3014 | -0.66 | 0.8944 |
| cluster3188 | -0.68 | 0.8947 |
| cluster2955 | -0.49 | 0.8947 |
| cluster3847 | 0.35 | 0.8947 |
| cluster3225 | 0.42 | 0.8947 |
| cluster3079 | -0.58 | 0.8947 |
| cluster2860 | -0.63 | 0.8947 |
| cluster3636 | -0.47 | 0.8947 |
| cluster1127 | -0.64 | 0.8947 |
| cluster3840 | 0.39 | 0.8947 |
| cluster3251 | 0.33 | 0.8963 |
| cluster3184 | 0.32 | 0.8963 |
| cluster3876 | 0.34 | 0.8963 |
| cluster2873 | -0.57 | 0.8963 |
| cluster3623 | 0.34 | 0.8963 |
| cluster2858 | -0.51 | 0.8963 |
| cluster3500 | -0.64 | 0.8963 |
| cluster3744 | 0.34 | 0.8963 |
| cluster3825 | -0.60 | 0.8967 |
| cluster3696 | -0.49 | 0.8981 |
| cluster3341 | 0.36 | 0.8981 |
| cluster3402 | 0.35 | 0.8982 |
| cluster3227 | -0.56 | 0.8998 |
| cluster3722 | 0.36 | 0.9014 |
| cluster3362 | -0.53 | 0.9025 |
| cluster2863 | 0.30 | 0.9025 |
| cluster3935 | 0.38 | 0.9025 |
| cluster3240 | 0.34 | 0.9031 |
| cluster3737 | 0.29 | 0.9083 |
| cluster3132 | 0.31 | 0.9083 |
| cluster3522 | -0.63 | 0.9087 |
| cluster3167 | 0.37 | 0.9120 |
| cluster3238 | 0.28 | 0.9120 |
| cluster3102 | -0.63 | 0.9148 |
| cluster3070 | 0.31 | 0.9148 |
| cluster3107 | -0.46 | 0.9148 |
| cluster3173 | 0.36 | 0.9148 |
| cluster3343 | 0.28 | 0.9148 |

| | | |
|---|---|---|
| cluster3204 | -0.54 | 0.9148 |
| cluster3388 | -0.63 | 0.9148 |
| cluster3691 | 0.32 | 0.9148 |
| cluster3827 | 0.33 | 0.9148 |
| cluster3554 | 0.30 | 0.9148 |
| cluster4124 | -0.80 | 0.9148 |
| cluster3524 | -0.45 | 0.9148 |
| cluster3973 | -0.51 | 0.9148 |
| cluster4003 | 0.35 | 0.9148 |
| cluster3133 | 0.28 | 0.9148 |
| cluster3339 | -0.51 | 0.9148 |
| cluster2935 | -0.50 | 0.9148 |
| cluster2968 | 0.26 | 0.9148 |
| cluster3378 | 0.32 | 0.9148 |
| cluster2931 | -0.52 | 0.9148 |
| cluster3115 | 0.27 | 0.9148 |
| cluster3437 | -0.51 | 0.9150 |
| cluster3394 | 0.27 | 0.9150 |
| cluster3960 | -0.76 | 0.9150 |
| cluster3276 | 0.28 | 0.9152 |
| cluster4115 | -0.81 | 0.9156 |
| cluster3123 | -0.46 | 0.9160 |
| cluster4025 | 0.27 | 0.9160 |
| cluster3295 | 0.26 | 0.9160 |
| cluster4042 | -0.61 | 0.9160 |
| cluster3472 | -0.47 | 0.9160 |
| cluster3760 | 0.29 | 0.9165 |
| cluster3426 | -0.54 | 0.9191 |
| cluster3358 | -0.54 | 0.9191 |
| cluster3792 | -0.44 | 0.9191 |
| cluster3557 | 0.28 | 0.9204 |
| cluster3806 | 0.30 | 0.9204 |
| cluster3392 | 0.31 | 0.9204 |
| cluster2880 | -0.48 | 0.9204 |
| cluster3303 | -0.73 | 0.9240 |
| cluster3433 | -0.52 | 0.9251 |
| cluster3541 | 0.26 | 0.9255 |
| cluster3380 | -0.45 | 0.9260 |
| cluster3903 | 0.25 | 0.9263 |
| cluster3013 | 0.25 | 0.9263 |
| cluster3034 | 0.25 | 0.9282 |
| cluster3707 | 0.29 | 0.9282 |
| cluster3582 | -0.43 | 0.9282 |
| cluster3372 | 0.24 | 0.9282 |

| | | |
|---|---|---|
| cluster3286 | -0.40 | 0.9285 |
| cluster3058 | 0.23 | 0.9285 |
| cluster3958 | -0.76 | 0.9285 |
| cluster3627 | -0.43 | 0.9285 |
| cluster3067 | 0.24 | 0.9316 |
| cluster3658 | -0.42 | 0.9354 |
| cluster2934 | -0.39 | 0.9354 |
| cluster3139 | 0.29 | 0.9354 |
| cluster3025 | -0.39 | 0.9355 |
| cluster2917 | 0.26 | 0.9355 |
| cluster3335 | 0.26 | 0.9357 |
| cluster3091 | 0.21 | 0.9367 |
| cluster3859 | -0.39 | 0.9389 |
| cluster4123 | -0.70 | 0.9389 |
| cluster3399 | 0.25 | 0.9389 |
| cluster3264 | -0.37 | 0.9389 |
| cluster3417 | -0.52 | 0.9389 |
| cluster3763 | -0.61 | 0.9389 |
| cluster3667 | -0.40 | 0.9389 |
| cluster2879 | -0.39 | 0.9389 |
| cluster3655 | 0.23 | 0.9389 |
| cluster3361 | 0.22 | 0.9389 |
| cluster3183 | 0.22 | 0.9389 |
| cluster3567 | -0.70 | 0.9389 |
| cluster3340 | -0.41 | 0.9389 |
| cluster2962 | -0.50 | 0.9389 |
| cluster3307 | 0.22 | 0.9389 |
| cluster3456 | -0.43 | 0.9389 |
| cluster3159 | 0.20 | 0.9389 |
| cluster3794 | 0.23 | 0.9417 |
| cluster4127 | -0.84 | 0.9417 |
| cluster3478 | -0.43 | 0.9439 |
| cluster3701 | -0.39 | 0.9439 |
| cluster3423 | 0.23 | 0.9439 |
| cluster2945 | -0.64 | 0.9439 |
| cluster3104 | 0.18 | 0.9439 |
| cluster2857 | 0.19 | 0.9439 |
| cluster4102 | 0.24 | 0.9439 |
| cluster3779 | 0.21 | 0.9439 |
| cluster2999 | 0.21 | 0.9439 |
| cluster3739 | -0.34 | 0.9439 |
| cluster3526 | 0.19 | 0.9439 |
| cluster2996 | 0.20 | 0.9439 |
| cluster3186 | 0.21 | 0.9439 |

| | | |
|---|---|---|
| cluster3835 | 0.21 | 0.9439 |
| cluster3562 | 0.20 | 0.9439 |
| cluster3309 | 0.21 | 0.9463 |
| cluster3882 | 0.23 | 0.9463 |
| cluster3490 | -0.49 | 0.9463 |
| cluster2940 | -0.37 | 0.9463 |
| cluster4007 | 0.18 | 0.9463 |
| cluster3046 | -0.37 | 0.9463 |
| cluster3262 | -0.37 | 0.9463 |
| cluster3963 | -0.59 | 0.9463 |
| cluster4005 | 0.20 | 0.9463 |
| cluster3406 | 0.18 | 0.9463 |
| cluster3164 | -0.55 | 0.9463 |
| cluster3479 | -0.53 | 0.9463 |
| cluster2956 | 0.18 | 0.9463 |
| cluster3224 | 0.17 | 0.9463 |
| cluster2861 | 0.19 | 0.9463 |
| cluster3900 | 0.19 | 0.9463 |
| cluster3096 | 0.17 | 0.9463 |
| cluster3148 | 0.16 | 0.9463 |
| cluster3480 | 0.18 | 0.9463 |
| cluster3208 | -0.34 | 0.9463 |
| cluster3664 | -0.37 | 0.9463 |
| cluster3851 | 0.19 | 0.9463 |
| cluster2926 | 0.17 | 0.9463 |
| cluster3579 | -0.36 | 0.9463 |
| cluster3113 | 0.16 | 0.9463 |
| cluster3072 | -0.44 | 0.9463 |
| cluster3689 | -0.69 | 0.9463 |
| cluster3412 | 0.17 | 0.9463 |
| cluster3599 | -0.49 | 0.9482 |
| cluster3602 | -0.39 | 0.9489 |
| cluster3239 | -0.34 | 0.9493 |
| cluster3350 | -0.38 | 0.9512 |
| cluster3101 | 0.16 | 0.9512 |
| cluster3573 | -0.40 | 0.9512 |
| cluster2919 | 0.16 | 0.9512 |
| cluster3428 | -0.34 | 0.9512 |
| cluster2870 | 0.15 | 0.9512 |
| cluster4063 | 0.17 | 0.9512 |
| cluster3308 | 0.15 | 0.9512 |
| cluster2932 | -0.35 | 0.9512 |
| cluster2891 | -0.41 | 0.9530 |
| cluster3805 | 0.16 | 0.9530 |

| | | |
|---|---|---|
| cluster2950 | -0.34 | 0.9558 |
| cluster3498 | 0.15 | 0.9558 |
| cluster2896 | 0.14 | 0.9558 |
| cluster3048 | 0.14 | 0.9558 |
| cluster3858 | 0.15 | 0.9558 |
| cluster2958 | -0.28 | 0.9558 |
| cluster3284 | 0.14 | 0.9558 |
| cluster3156 | 0.14 | 0.9558 |
| cluster3163 | -0.49 | 0.9558 |
| cluster3364 | -0.32 | 0.9558 |
| cluster3092 | -0.28 | 0.9558 |
| cluster3459 | -0.50 | 0.9558 |
| cluster3887 | 0.15 | 0.9609 |
| cluster3755 | -0.66 | 0.9613 |
| cluster3001 | 0.12 | 0.9631 |
| cluster4103 | 0.14 | 0.9631 |
| cluster3669 | 0.12 | 0.9647 |
| cluster3695 | 0.13 | 0.9647 |
| cluster3516 | -0.49 | 0.9647 |
| cluster3312 | 0.11 | 0.9662 |
| cluster2944 | -0.33 | 0.9662 |
| cluster3589 | 0.12 | 0.9662 |
| cluster4032 | 0.13 | 0.9662 |
| cluster3031 | -0.31 | 0.9676 |
| cluster3396 | -0.34 | 0.9676 |
| cluster3765 | -0.33 | 0.9676 |
| cluster3033 | -0.26 | 0.9681 |
| cluster3351 | 0.10 | 0.9715 |
| cluster3535 | -0.44 | 0.9715 |
| cluster2910 | -0.27 | 0.9715 |
| cluster3697 | -0.43 | 0.9715 |
| cluster3577 | 0.10 | 0.9715 |
| cluster4020 | -0.28 | 0.9747 |
| cluster4106 | 0.07 | 0.9747 |
| cluster3749 | 0.09 | 0.9747 |
| cluster3328 | 0.10 | 0.9747 |
| cluster3728 | -0.28 | 0.9747 |
| cluster3445 | -0.26 | 0.9747 |
| cluster3348 | -0.34 | 0.9747 |
| cluster3883 | 0.07 | 0.9747 |
| cluster4048 | -0.52 | 0.9747 |
| cluster3877 | 0.09 | 0.9747 |
| cluster3782 | 0.09 | 0.9747 |
| cluster3166 | -0.27 | 0.9747 |

| | | |
|---|---|---|
| cluster3922 | 0.07 | 0.9747 |
| cluster3764 | 0.08 | 0.9747 |
| cluster3632 | -0.29 | 0.9747 |
| cluster3596 | 0.10 | 0.9747 |
| cluster3948 | 0.08 | 0.9747 |
| cluster2921 | 0.08 | 0.9747 |
| cluster3912 | 0.08 | 0.9747 |
| cluster2885 | -0.31 | 0.9747 |
| cluster2929 | 0.08 | 0.9747 |
| cluster3062 | -0.27 | 0.9749 |
| cluster3670 | 0.07 | 0.9749 |
| cluster3663 | 0.08 | 0.9749 |
| cluster3816 | 0.07 | 0.9749 |
| cluster3758 | -0.33 | 0.9749 |
| cluster3296 | 0.07 | 0.9749 |
| cluster2878 | -0.30 | 0.9754 |
| cluster3861 | 0.08 | 0.9771 |
| cluster3144 | -0.43 | 0.9774 |
| cluster3324 | 0.07 | 0.9774 |
| cluster3181 | -0.28 | 0.9774 |
| cluster2907 | -0.26 | 0.9774 |
| cluster4101 | 0.01 | 0.9774 |
| cluster3848 | -0.32 | 0.9774 |
| cluster3826 | -0.38 | 0.9774 |
| cluster3873 | 0.07 | 0.9774 |
| cluster3894 | -0.26 | 0.9807 |
| cluster3258 | 0.06 | 0.9814 |
| cluster3727 | -0.37 | 0.9814 |
| cluster2882 | 0.06 | 0.9814 |
| cluster3791 | -0.30 | 0.9814 |
| cluster2971 | -0.21 | 0.9839 |
| cluster3931 | -0.38 | 0.9841 |
| cluster3321 | -0.23 | 0.9841 |
| cluster3314 | 0.04 | 0.9841 |
| cluster3027 | 0.05 | 0.9848 |
| cluster3266 | 0.05 | 0.9848 |
| cluster3989 | 0.06 | 0.9848 |
| cluster4078 | -0.30 | 0.9866 |
| cluster2897 | 0.04 | 0.9868 |
| cluster4035 | 0.05 | 0.9868 |
| cluster3736 | 0.02 | 0.9868 |
| cluster4022 | 0.04 | 0.9868 |
| cluster3884 | 0.02 | 0.9868 |
| cluster2883 | 0.04 | 0.9870 |

| | | |
|---|---|---|
| cluster3718 | -0.33 | 0.9870 |
| cluster3893 | 0.04 | 0.9876 |
| cluster4041 | 0.02 | 0.9876 |
| cluster3216 | 0.03 | 0.9876 |
| cluster4075 | -0.38 | 0.9876 |
| cluster3320 | 0.02 | 0.9876 |
| cluster3214 | -0.23 | 0.9876 |
| cluster3802 | -0.38 | 0.9876 |
| cluster3818 | 0.01 | 0.9876 |
| cluster2961 | 0.02 | 0.9878 |
| cluster3756 | -0.25 | 0.9888 |
| cluster3331 | -0.19 | 0.9897 |
| cluster3530 | -0.20 | 0.9897 |
| cluster3061 | -0.24 | 0.9897 |
| cluster3359 | 0.01 | 0.9897 |
| cluster3032 | 0.02 | 0.9897 |
| cluster3255 | -0.28 | 0.9897 |
| cluster4006 | -0.19 | 0.9901 |
| cluster3278 | 0.02 | 0.9901 |
| cluster2933 | 0.00 | 0.9902 |
| cluster3547 | 0.02 | 0.9912 |
| cluster2997 | -0.19 | 0.9914 |
| cluster3559 | -0.20 | 0.9914 |
| cluster2939 | -0.18 | 0.9914 |
| cluster3820 | -0.17 | 0.9914 |
| cluster3002 | 0.01 | 0.9925 |
| cluster3310 | -0.16 | 0.9934 |
| cluster2875 | -0.21 | 0.9973 |
| cluster3020 | 0.00 | 0.9973 |
| cluster2859 | -0.15 | 0.9973 |
| cluster3089 | 0.00 | 0.9973 |
| cluster3608 | -0.16 | 0.9978 |
| cluster3054 | 0.01 | 0.9981 |
| cluster3141 | -0.18 | 0.9981 |
| cluster3748 | -0.20 | 0.9991 |
| cluster2946 | -0.21 | 1.0000 |
| cluster3517 | 0.01 | 1.0000 |
| cluster3147 | 0.00 | 1.0000 |
| cluster3868 | -0.15 | 1.0000 |
| cluster4043 | 0.03 | 1.0000 |
| cluster3037 | -0.17 | 1.0000 |
| cluster2965 | -0.01 | 1.0000 |
| cluster3294 | -0.01 | 1.0000 |
| cluster3347 | -0.15 | 1.0000 |

| | | |
|---|---|---|
| cluster3762 | -0.24 | 1.0000 |
| cluster3890 | -0.26 | 1.0000 |
| cluster3928 | -0.17 | 1.0000 |
| cluster3505 | -0.16 | 1.0000 |
| cluster3854 | -0.14 | 1.0000 |
| cluster2916 | -0.20 | 1.0000 |
| cluster3128 | -0.14 | 1.0000 |
| cluster3376 | -0.14 | 1.0000 |
| cluster4108 | -0.27 | 1.0000 |
| cluster3390 | -0.17 | 1.0000 |
| cluster3795 | -0.15 | 1.0000 |
| cluster3150 | -0.24 | 1.0000 |
| cluster3888 | -0.24 | 1.0000 |
| cluster2993 | -0.06 | 1.0000 |
| cluster3586 | -0.03 | 1.0000 |
| cluster2884 | -0.03 | 1.0000 |
| cluster3684 | -0.20 | 1.0000 |
| cluster3269 | -0.24 | 1.0000 |
| cluster4026 | -0.23 | 1.0000 |
| cluster3682 | -0.26 | 1.0000 |
| cluster3846 | -0.13 | 1.0000 |
| cluster3585 | -0.14 | 1.0000 |
| cluster3407 | -0.15 | 1.0000 |
| cluster3304 | -0.05 | 1.0000 |
| cluster3384 | -0.17 | 1.0000 |
| cluster3289 | -0.14 | 1.0000 |
| cluster3565 | -0.17 | 1.0000 |
| cluster3566 | -0.14 | 1.0000 |
| cluster3023 | -0.06 | 1.0000 |
| cluster3323 | -0.20 | 1.0000 |
| cluster3588 | -0.18 | 1.0000 |
| cluster3049 | -0.14 | 1.0000 |
| cluster3297 | -0.10 | 1.0000 |
| cluster4004 | -0.10 | 1.0000 |
| cluster3593 | -0.16 | 1.0000 |
| cluster3165 | -0.19 | 1.0000 |
| cluster3022 | -0.16 | 1.0000 |
| cluster2911 | -0.06 | 1.0000 |
| cluster3741 | -0.15 | 1.0000 |
| cluster3176 | -0.11 | 1.0000 |
| cluster2923 | -0.10 | 1.0000 |
| cluster3005 | -0.10 | 1.0000 |
| cluster3313 | -0.12 | 1.0000 |
| cluster3222 | -0.06 | 1.0000 |

| | | |
|---|---|---|
| cluster3785 | -0.14 | 1.0000 |
| cluster3521 | -0.11 | 1.0000 |
| cluster3071 | -0.09 | 1.0000 |
| cluster3717 | -0.09 | 1.0000 |
| cluster3729 | -0.10 | 1.0000 |
| cluster3124 | -0.07 | 1.0000 |
| cluster3403 | -0.07 | 1.0000 |
| cluster3352 | -0.13 | 1.0000 |
| cluster2872 | -0.08 | 1.0000 |
| cluster3539 | -0.16 | 1.0000 |
| cluster3705 | -0.14 | 1.0000 |
| cluster3692 | -0.13 | 1.0000 |
| cluster3999 | -0.13 | 1.0000 |
| cluster2989 | -0.13 | 1.0000 |
| cluster4053 | -0.12 | 1.0000 |
| cluster3665 | -0.11 | 1.0000 |
| cluster3138 | -0.10 | 1.0000 |
| cluster3797 | -0.10 | 1.0000 |
| cluster3955 | -0.09 | 1.0000 |
| cluster3984 | -0.09 | 1.0000 |
| cluster3021 | -0.08 | 1.0000 |
| cluster3926 | -0.08 | 1.0000 |
| cluster3620 | -0.08 | 1.0000 |
| cluster3268 | -0.08 | 1.0000 |
| cluster3615 | -0.07 | 1.0000 |
| cluster3241 | -0.07 | 1.0000 |
| cluster3215 | -0.06 | 1.0000 |
| cluster3041 | -0.06 | 1.0000 |
| cluster3874 | -0.06 | 1.0000 |
| cluster3823 | -0.06 | 1.0000 |
| cluster4105 | -0.05 | 1.0000 |
| cluster3988 | -0.04 | 1.0000 |
| cluster3716 | -0.04 | 1.0000 |
| cluster3946 | -0.03 | 1.0000 |
| cluster3992 | -0.03 | 1.0000 |
| cluster3088 | -0.03 | 1.0000 |
| cluster3248 | -0.03 | 1.0000 |
| cluster3561 | -0.02 | 1.0000 |
| cluster3907 | -0.02 | 1.0000 |
| cluster3597 | -0.02 | 1.0000 |
| cluster3256 | -0.02 | 1.0000 |
| cluster4066 | -0.01 | 1.0000 |
| cluster3098 | -0.01 | 1.0000 |
| cluster3134 | -0.01 | 1.0000 |

| | | |
|---|---|---|
| cluster3170 | 0.00 | 1.0000 |

**Supplementary Table 13.** DESeq2 results with RSEM counts - Dermic vs nasopharyngeal

| ID | log2FoldChange | Adjusted p value |
|---|---|---|
| cluster3845 | -13.25 | 4.04E-36 |
| cluster4085 | -12.11 | 3.37E-20 |
| cluster3753 | -12.63 | 5.92E-18 |
| cluster3800 | -10.53 | 5.92E-18 |
| cluster3919 | -13.52 | 1.39E-16 |
| cluster3544 | -6.85 | 8.55E-14 |
| cluster4015 | -8.20 | 8.55E-14 |
| cluster4080 | -9.17 | 4.03E-10 |
| cluster4049 | -12.01 | 1.45E-09 |
| cluster3510 | -12.04 | 2.01E-09 |
| cluster3925 | -11.65 | 5.78E-09 |
| cluster4097 | -11.20 | 1.13E-08 |
| cluster4011 | -9.67 | 1.81E-06 |
| cluster2869 | -3.86 | 1.94E-05 |
| cluster4046 | -6.10 | 0.001 |
| cluster3780 | -6.43 | 0.002 |
| cluster4087 | 4.84 | 0.002 |
| cluster4034 | -11.64 | 0.003 |
| cluster3326 | -3.04 | 0.005 |
| cluster2894 | -4.25 | 0.006 |
| cluster3981 | 3.02 | 0.009 |
| cluster3534 | -3.31 | 0.015 |
| cluster3650 | -3.80 | 0.021 |
| cluster3666 | 4.17 | 0.034 |
| cluster3481 | -4.02 | 0.035 |
| cluster4090 | -5.46 | 0.038 |
| cluster4070 | -6.40 | 0.045 |
| cluster3504 | -3.35 | 0.049 |
| cluster2904 | -2.67 | 0.055 |
| cluster4119 | 3.58 | 0.061 |
| cluster3325 | -4.33 | 0.065 |
| cluster3839 | -4.15 | 0.072 |
| cluster3057 | -2.98 | 0.093 |
| cluster3901 | 3.49 | 0.099 |
| cluster3024 | -2.30 | 0.101 |
| cluster4067 | 4.22 | 0.101 |
| cluster3250 | 3.03 | 0.163 |
| cluster4073 | 3.38 | 0.192 |
| cluster3009 | -2.70 | 0.195 |
| cluster3880 | -3.97 | 0.195 |
| cluster3568 | -3.60 | 0.201 |
| cluster3052 | -3.56 | 0.219 |

| | | |
|---|---|---|
| cluster4016 | 2.37 | 0.234 |
| cluster3229 | -2.32 | 0.234 |
| cluster3474 | 2.97 | 0.274 |
| cluster3306 | 3.25 | 0.274 |
| cluster2900 | -2.31 | 0.282 |
| cluster3393 | -3.39 | 0.310 |
| cluster4047 | 3.04 | 0.313 |
| cluster3383 | -3.29 | 0.319 |
| cluster3424 | -3.29 | 0.334 |
| cluster2992 | -1.89 | 0.351 |
| cluster3290 | -2.47 | 0.351 |
| cluster3430 | 2.89 | 0.351 |
| cluster3869 | 2.74 | 0.351 |
| cluster4002 | 2.72 | 0.351 |
| cluster4029 | -3.31 | 0.351 |
| cluster4074 | 2.78 | 0.351 |
| cluster2899 | -2.15 | 0.363 |
| cluster3160 | -2.02 | 0.363 |
| cluster3172 | -1.92 | 0.363 |
| cluster3379 | -2.55 | 0.363 |
| cluster3432 | -2.30 | 0.363 |
| cluster3434 | -2.19 | 0.363 |
| cluster3853 | 2.81 | 0.363 |
| cluster4107 | 2.60 | 0.403 |
| cluster3223 | -2.43 | 0.420 |
| cluster3171 | -2.00 | 0.431 |
| cluster4079 | 2.02 | 0.436 |
| cluster3674 | -3.31 | 0.436 |
| cluster3905 | -3.78 | 0.436 |
| cluster2972 | -2.10 | 0.445 |
| cluster2980 | -3.03 | 0.445 |
| cluster3436 | 2.70 | 0.445 |
| cluster3482 | -2.21 | 0.445 |
| cluster3571 | -2.39 | 0.445 |
| cluster3617 | -3.22 | 0.445 |
| cluster3761 | -3.12 | 0.445 |
| cluster3408 | -2.08 | 0.445 |
| cluster3581 | -2.66 | 0.445 |
| cluster3940 | 1.75 | 0.445 |
| cluster3994 | -3.02 | 0.457 |
| cluster2954 | -1.83 | 0.459 |
| cluster3129 | -3.03 | 0.459 |
| cluster3486 | -1.88 | 0.459 |
| cluster3629 | 2.52 | 0.459 |

| | | |
|---|---|---|
| cluster3675 | -1.59 | 0.459 |
| cluster3787 | -2.74 | 0.459 |
| cluster2974 | -1.88 | 0.459 |
| cluster3381 | -1.75 | 0.459 |
| cluster3461 | -3.29 | 0.459 |
| cluster3645 | -3.54 | 0.459 |
| cluster3865 | 2.52 | 0.459 |
| cluster3983 | 2.53 | 0.459 |
| cluster4091 | -3.31 | 0.459 |
| cluster3192 | -1.65 | 0.462 |
| cluster3003 | -2.56 | 0.464 |
| cluster3097 | 1.72 | 0.464 |
| cluster3519 | -2.97 | 0.464 |
| cluster3520 | -3.55 | 0.465 |
| cluster3563 | -1.97 | 0.476 |
| cluster4121 | 2.37 | 0.476 |
| cluster3422 | -2.47 | 0.494 |
| cluster3081 | -1.81 | 0.502 |
| cluster3651 | -2.73 | 0.503 |
| cluster3970 | 2.46 | 0.504 |
| cluster2915 | -2.27 | 0.515 |
| cluster4093 | 2.52 | 0.527 |
| cluster2912 | 1.92 | 0.528 |
| cluster3111 | -2.51 | 0.528 |
| cluster3812 | -2.44 | 0.528 |
| cluster4120 | -3.04 | 0.528 |
| cluster4122 | -3.10 | 0.528 |
| cluster3543 | -2.60 | 0.531 |
| cluster3090 | -2.85 | 0.534 |
| cluster4099 | 2.27 | 0.534 |
| cluster3745 | 2.72 | 0.535 |
| cluster3443 | 2.14 | 0.553 |
| cluster3538 | -1.91 | 0.569 |
| cluster3574 | -2.69 | 0.569 |
| cluster3598 | 2.68 | 0.569 |
| cluster3752 | -2.80 | 0.570 |
| cluster4060 | 2.40 | 0.570 |
| cluster3016 | -2.46 | 0.570 |
| cluster4125 | -2.82 | 0.570 |
| cluster2892 | 1.41 | 0.570 |
| cluster2925 | -2.49 | 0.580 |
| cluster3035 | -1.33 | 0.580 |
| cluster3040 | -1.55 | 0.580 |
| cluster3045 | 1.60 | 0.580 |

| | | |
|---|---|---|
| cluster3142 | -2.15 | 0.580 |
| cluster3177 | -2.17 | 0.580 |
| cluster3207 | 1.62 | 0.580 |
| cluster3633 | -1.68 | 0.580 |
| cluster3694 | -1.93 | 0.580 |
| cluster3875 | 1.88 | 0.580 |
| cluster3889 | 2.22 | 0.580 |
| cluster3063 | 1.66 | 0.589 |
| cluster3693 | 2.18 | 0.589 |
| cluster3084 | -1.29 | 0.599 |
| cluster3095 | -1.63 | 0.599 |
| cluster3453 | -1.89 | 0.599 |
| cluster3233 | -2.38 | 0.603 |
| cluster3105 | -2.90 | 0.611 |
| cluster3187 | -2.42 | 0.611 |
| cluster3824 | 2.25 | 0.613 |
| cluster3719 | 2.12 | 0.625 |
| cluster4045 | 2.34 | 0.625 |
| cluster2867 | -2.28 | 0.629 |
| cluster2890 | -1.23 | 0.629 |
| cluster2928 | 1.89 | 0.629 |
| cluster3010 | -1.88 | 0.629 |
| cluster3017 | -2.13 | 0.629 |
| cluster3162 | 1.92 | 0.629 |
| cluster3168 | 1.44 | 0.629 |
| cluster3292 | -2.64 | 0.629 |
| cluster3374 | 2.05 | 0.629 |
| cluster3395 | -1.56 | 0.629 |
| cluster3438 | 2.00 | 0.629 |
| cluster3454 | -1.52 | 0.629 |
| cluster3469 | -1.57 | 0.629 |
| cluster3484 | -2.58 | 0.629 |
| cluster3507 | -2.34 | 0.629 |
| cluster3572 | 2.47 | 0.629 |
| cluster3580 | 2.00 | 0.629 |
| cluster3595 | 1.46 | 0.629 |
| cluster3611 | 2.05 | 0.629 |
| cluster3618 | -2.62 | 0.629 |
| cluster3619 | 1.63 | 0.629 |
| cluster3772 | -2.01 | 0.629 |
| cluster3778 | -1.55 | 0.629 |
| cluster3867 | -1.49 | 0.629 |
| cluster3974 | 2.33 | 0.629 |
| cluster3995 | -2.31 | 0.629 |

| | | |
|---|---|---|
| cluster4054 | 2.00 | 0.629 |
| cluster4062 | -2.64 | 0.629 |
| cluster2908 | -1.55 | 0.632 |
| cluster3008 | 1.92 | 0.632 |
| cluster3217 | -1.43 | 0.632 |
| cluster3220 | -1.77 | 0.632 |
| cluster3288 | 1.99 | 0.632 |
| cluster3501 | -2.09 | 0.632 |
| cluster3698 | -1.27 | 0.632 |
| cluster3788 | -1.48 | 0.632 |
| cluster3814 | -1.49 | 0.632 |
| cluster3817 | -1.79 | 0.632 |
| cluster3902 | 2.07 | 0.632 |
| cluster4084 | -1.59 | 0.632 |
| cluster4111 | -2.62 | 0.632 |
| cluster4031 | -1.52 | 0.637 |
| cluster3892 | -1.94 | 0.638 |
| cluster3179 | 2.02 | 0.638 |
| cluster3442 | 2.10 | 0.638 |
| cluster3451 | 2.65 | 0.638 |
| cluster3815 | 1.97 | 0.638 |
| cluster3924 | 1.95 | 0.638 |
| cluster3985 | 1.49 | 0.638 |
| cluster4113 | -1.79 | 0.638 |
| cluster3552 | 1.28 | 0.658 |
| cluster3161 | -2.10 | 0.661 |
| cluster3431 | 1.94 | 0.661 |
| cluster3661 | 1.31 | 0.661 |
| cluster3194 | -1.49 | 0.663 |
| cluster3508 | 1.93 | 0.663 |
| cluster3587 | -2.39 | 0.663 |
| cluster3685 | -2.12 | 0.663 |
| cluster3254 | -2.29 | 0.664 |
| cluster2876 | 2.21 | 0.665 |
| cluster3122 | -1.35 | 0.678 |
| cluster3690 | -1.33 | 0.678 |
| cluster3913 | -2.06 | 0.678 |
| cluster3515 | -2.16 | 0.679 |
| cluster4116 | -2.36 | 0.679 |
| cluster3270 | 1.98 | 0.683 |
| cluster3337 | -1.43 | 0.687 |
| cluster3639 | 2.21 | 0.687 |
| cluster3804 | 1.96 | 0.687 |
| cluster3822 | 2.06 | 0.687 |

| | | |
|---|---|---|
| cluster2987 | 1.10 | 0.688 |
| cluster3410 | -1.50 | 0.688 |
| cluster3463 | 1.86 | 0.688 |
| cluster3556 | 1.89 | 0.688 |
| cluster3635 | -1.09 | 0.688 |
| cluster3678 | 1.28 | 0.688 |
| cluster3742 | 1.86 | 0.688 |
| cluster3933 | -1.90 | 0.688 |
| cluster3952 | 1.91 | 0.688 |
| cluster3987 | 1.46 | 0.688 |
| cluster3082 | -2.06 | 0.691 |
| cluster3100 | 2.17 | 0.691 |
| cluster4064 | 1.81 | 0.695 |
| cluster3349 | 1.45 | 0.715 |
| cluster3457 | -2.06 | 0.717 |
| cluster3703 | 1.09 | 0.732 |
| cluster3287 | -1.55 | 0.741 |
| cluster3956 | 1.76 | 0.743 |
| cluster4100 | 1.67 | 0.743 |
| cluster3415 | -1.52 | 0.744 |
| cluster2862 | -1.11 | 0.750 |
| cluster3000 | -1.75 | 0.750 |
| cluster3012 | 1.88 | 0.750 |
| cluster2914 | -1.65 | 0.750 |
| cluster2995 | 1.60 | 0.750 |
| cluster3015 | -1.76 | 0.750 |
| cluster3076 | 1.70 | 0.750 |
| cluster3083 | 1.09 | 0.750 |
| cluster3212 | -1.64 | 0.750 |
| cluster3365 | 1.81 | 0.750 |
| cluster3419 | -1.26 | 0.750 |
| cluster3455 | -1.45 | 0.750 |
| cluster3462 | -1.46 | 0.750 |
| cluster3466 | -1.49 | 0.750 |
| cluster3497 | -1.13 | 0.750 |
| cluster3512 | 1.47 | 0.750 |
| cluster3569 | 1.76 | 0.750 |
| cluster3714 | -1.40 | 0.750 |
| cluster3796 | -1.91 | 0.750 |
| cluster3968 | 1.60 | 0.753 |
| cluster3135 | -1.97 | 0.758 |
| cluster3146 | -1.16 | 0.758 |
| cluster3724 | 1.72 | 0.758 |
| cluster3976 | -1.27 | 0.758 |

| | | |
|---|---|---|
| cluster3086 | -1.44 | 0.767 |
| cluster3771 | 1.64 | 0.771 |
| cluster3584 | 1.66 | 0.777 |
| cluster3807 | -1.19 | 0.780 |
| cluster4000 | 1.39 | 0.780 |
| cluster3075 | -1.30 | 0.782 |
| cluster3275 | -1.76 | 0.783 |
| cluster3425 | -1.42 | 0.783 |
| cluster3468 | -1.20 | 0.783 |
| cluster3473 | 1.30 | 0.783 |
| cluster3734 | 1.68 | 0.783 |
| cluster3776 | -1.29 | 0.783 |
| cluster3941 | -1.09 | 0.783 |
| cluster3413 | -1.82 | 0.789 |
| cluster3427 | -1.61 | 0.789 |
| cluster3631 | -1.55 | 0.789 |
| cluster3957 | 1.65 | 0.789 |
| cluster3576 | 1.47 | 0.791 |
| cluster3915 | -1.94 | 0.803 |
| cluster3318 | -1.25 | 0.804 |
| cluster3382 | -1.21 | 0.804 |
| cluster3398 | -1.49 | 0.804 |
| cluster3471 | 1.70 | 0.804 |
| cluster3487 | -1.87 | 0.804 |
| cluster3594 | -1.70 | 0.804 |
| cluster3704 | 1.24 | 0.804 |
| cluster3866 | 1.53 | 0.804 |
| cluster3871 | 1.56 | 0.804 |
| cluster3904 | -1.96 | 0.804 |
| cluster3936 | 1.59 | 0.804 |
| cluster3953 | -1.65 | 0.804 |
| cluster3975 | 1.63 | 0.804 |
| cluster4089 | -2.18 | 0.804 |
| cluster4109 | 1.69 | 0.804 |
| cluster2967 | 1.27 | 0.806 |
| cluster3065 | -1.40 | 0.806 |
| cluster3781 | 1.23 | 0.806 |
| cluster3175 | -1.74 | 0.808 |
| cluster3259 | -1.08 | 0.808 |
| cluster3735 | -1.33 | 0.808 |
| cluster3885 | -1.67 | 0.808 |
| cluster3677 | 1.14 | 0.810 |
| cluster3360 | -1.39 | 0.810 |
| cluster3945 | -1.69 | 0.813 |

| | | |
|---|---|---|
| cluster3193 | -1.42 | 0.814 |
| cluster3610 | 1.68 | 0.819 |
| cluster4044 | 1.51 | 0.819 |
| cluster3920 | -1.74 | 0.822 |
| cluster2984 | -1.16 | 0.829 |
| cluster3319 | -1.63 | 0.829 |
| cluster3638 | 1.63 | 0.829 |
| cluster3773 | -1.18 | 0.829 |
| cluster2853 | -1.15 | 0.830 |
| cluster2930 | 1.32 | 0.830 |
| cluster3056 | -0.99 | 0.830 |
| cluster3136 | -1.45 | 0.830 |
| cluster3151 | -1.14 | 0.830 |
| cluster3196 | -0.92 | 0.830 |
| cluster3205 | -1.06 | 0.830 |
| cluster3356 | 1.06 | 0.830 |
| cluster3357 | 1.15 | 0.830 |
| cluster3397 | -1.65 | 0.830 |
| cluster3441 | 1.21 | 0.830 |
| cluster3450 | -1.94 | 0.830 |
| cluster3518 | -1.71 | 0.830 |
| cluster3630 | -0.96 | 0.830 |
| cluster3681 | 1.49 | 0.830 |
| cluster3683 | 1.53 | 0.830 |
| cluster3723 | 1.08 | 0.830 |
| cluster3766 | -1.89 | 0.830 |
| cluster4008 | 1.42 | 0.830 |
| cluster4009 | 1.38 | 0.830 |
| cluster4010 | 1.44 | 0.830 |
| cluster4024 | 1.45 | 0.830 |
| cluster4071 | 1.58 | 0.830 |
| cluster4081 | 1.45 | 0.830 |
| cluster3710 | 1.43 | 0.830 |
| cluster3230 | -1.15 | 0.830 |
| cluster3768 | -1.62 | 0.831 |
| cluster4033 | -1.70 | 0.838 |
| cluster4112 | 1.48 | 0.839 |
| cluster2864 | 0.93 | 0.845 |
| cluster2865 | 0.94 | 0.845 |
| cluster2877 | 0.77 | 0.845 |
| cluster2953 | -0.89 | 0.845 |
| cluster3069 | -0.87 | 0.845 |
| cluster3103 | -1.67 | 0.845 |
| cluster3131 | -1.59 | 0.845 |

| | | |
|---|---|---|
| cluster3213 | -1.10 | 0.845 |
| cluster3342 | 1.46 | 0.845 |
| cluster3363 | 1.42 | 0.845 |
| cluster3465 | -0.97 | 0.845 |
| cluster3475 | 1.44 | 0.845 |
| cluster3634 | -0.91 | 0.845 |
| cluster3646 | 1.48 | 0.845 |
| cluster3653 | -1.04 | 0.845 |
| cluster3656 | 1.02 | 0.845 |
| cluster3657 | 1.43 | 0.845 |
| cluster3702 | 1.40 | 0.845 |
| cluster3715 | 1.32 | 0.845 |
| cluster3830 | 1.06 | 0.845 |
| cluster3842 | 1.15 | 0.845 |
| cluster3844 | -1.70 | 0.845 |
| cluster3910 | -1.55 | 0.845 |
| cluster3914 | -1.78 | 0.845 |
| cluster3930 | 1.01 | 0.845 |
| cluster3971 | 1.46 | 0.845 |
| cluster3980 | 0.83 | 0.845 |
| cluster4037 | 1.38 | 0.845 |
| cluster4038 | 1.38 | 0.845 |
| cluster4051 | 1.69 | 0.845 |
| cluster4056 | -1.60 | 0.845 |
| cluster4068 | 1.38 | 0.845 |
| cluster4095 | 1.31 | 0.845 |
| cluster2982 | -1.21 | 0.851 |
| cluster3182 | -1.31 | 0.851 |
| cluster3368 | -0.91 | 0.859 |
| cluster3435 | -1.55 | 0.859 |
| cluster3834 | -1.52 | 0.859 |
| cluster3575 | -1.66 | 0.860 |
| cluster3279 | 1.27 | 0.861 |
| cluster3334 | 1.05 | 0.861 |
| cluster3662 | -1.39 | 0.861 |
| cluster3942 | -1.42 | 0.861 |
| cluster3405 | -1.17 | 0.862 |
| cluster3604 | 1.45 | 0.862 |
| cluster3626 | -0.95 | 0.862 |
| cluster3641 | -0.92 | 0.862 |
| cluster3711 | -1.07 | 0.862 |
| cluster3783 | 1.38 | 0.862 |
| cluster3870 | -1.47 | 0.862 |
| cluster2903 | -1.14 | 0.863 |

| | | |
|---|---|---|
| cluster2913 | -1.09 | 0.863 |
| cluster3114 | -1.09 | 0.863 |
| cluster3332 | 1.39 | 0.863 |
| cluster3525 | -1.08 | 0.863 |
| cluster3536 | 0.86 | 0.863 |
| cluster3841 | -1.42 | 0.863 |
| cluster3488 | -1.29 | 0.870 |
| cluster2966 | 1.37 | 0.871 |
| cluster3004 | -0.90 | 0.871 |
| cluster3730 | -0.90 | 0.871 |
| cluster3917 | 1.43 | 0.871 |
| cluster2901 | -1.38 | 0.872 |
| cluster2947 | 0.95 | 0.872 |
| cluster2957 | -1.40 | 0.872 |
| cluster3050 | -1.06 | 0.872 |
| cluster3496 | 0.94 | 0.872 |
| cluster3540 | -1.33 | 0.872 |
| cluster3640 | -2.51 | 0.872 |
| cluster3813 | 1.23 | 0.872 |
| cluster4104 | 1.27 | 0.872 |
| cluster3185 | -1.55 | 0.873 |
| cluster3311 | 0.92 | 0.873 |
| cluster3950 | 0.89 | 0.873 |
| cluster3965 | -1.31 | 0.873 |
| cluster4057 | 1.27 | 0.875 |
| cluster3039 | 0.80 | 0.876 |
| cluster3094 | 1.32 | 0.876 |
| cluster3449 | -0.83 | 0.876 |
| cluster3550 | -0.86 | 0.876 |
| cluster3080 | -1.13 | 0.876 |
| cluster3265 | -1.39 | 0.876 |
| cluster3444 | -0.90 | 0.876 |
| cluster3528 | -0.89 | 0.876 |
| cluster3121 | -0.95 | 0.878 |
| cluster3130 | -1.33 | 0.878 |
| cluster3190 | 1.35 | 0.878 |
| cluster3366 | 1.27 | 0.878 |
| cluster3416 | 1.36 | 0.878 |
| cluster3485 | -1.58 | 0.878 |
| cluster3545 | 1.24 | 0.878 |
| cluster3977 | -1.42 | 0.878 |
| cluster4018 | 1.18 | 0.878 |
| cluster2918 | -1.11 | 0.879 |
| cluster3180 | -1.31 | 0.879 |

| | | |
|---|---|---|
| cluster3810 | 0.86 | 0.879 |
| cluster3860 | 0.95 | 0.883 |
| cluster2959 | 0.82 | 0.893 |
| cluster2975 | 0.79 | 0.893 |
| cluster3301 | -0.93 | 0.893 |
| cluster3315 | -0.82 | 0.893 |
| cluster3591 | -0.90 | 0.893 |
| cluster4069 | 1.11 | 0.893 |
| cluster3231 | -1.48 | 0.896 |
| cluster3644 | -1.03 | 0.899 |
| cluster2887 | 1.24 | 0.900 |
| cluster2948 | -1.08 | 0.900 |
| cluster3267 | 0.76 | 0.900 |
| cluster4083 | 1.21 | 0.900 |
| cluster3242 | 1.01 | 0.903 |
| cluster3203 | -1.21 | 0.904 |
| cluster3897 | 1.02 | 0.911 |
| cluster3261 | -1.21 | 0.922 |
| cluster2854 | 1.19 | 0.923 |
| cluster2886 | -0.77 | 0.923 |
| cluster2922 | 1.19 | 0.923 |
| cluster2936 | 1.18 | 0.923 |
| cluster2969 | 1.19 | 0.923 |
| cluster3068 | -0.71 | 0.923 |
| cluster3085 | -1.16 | 0.923 |
| cluster3109 | 1.27 | 0.923 |
| cluster3110 | 0.95 | 0.923 |
| cluster3191 | -1.30 | 0.923 |
| cluster3226 | -1.16 | 0.923 |
| cluster3247 | 1.06 | 0.923 |
| cluster3249 | -0.97 | 0.923 |
| cluster3293 | 0.72 | 0.923 |
| cluster3330 | 0.97 | 0.923 |
| cluster3400 | -0.81 | 0.923 |
| cluster3418 | 1.10 | 0.923 |
| cluster3439 | -1.14 | 0.923 |
| cluster3446 | -1.17 | 0.923 |
| cluster3489 | 1.17 | 0.923 |
| cluster3503 | 1.13 | 0.923 |
| cluster3532 | -0.95 | 0.923 |
| cluster3537 | 1.18 | 0.923 |
| cluster3542 | 1.12 | 0.923 |
| cluster3564 | 1.03 | 0.923 |
| cluster3567 | -1.38 | 0.923 |

| | | |
|---|---|---|
| cluster3570 | 1.20 | 0.923 |
| cluster3590 | 1.11 | 0.923 |
| cluster3642 | 1.17 | 0.923 |
| cluster3649 | -0.93 | 0.923 |
| cluster3672 | -1.21 | 0.923 |
| cluster3676 | 1.18 | 0.923 |
| cluster3801 | 1.10 | 0.923 |
| cluster3811 | -0.70 | 0.923 |
| cluster3855 | 1.05 | 0.923 |
| cluster3898 | 1.05 | 0.923 |
| cluster3969 | -0.73 | 0.923 |
| cluster4001 | -1.28 | 0.923 |
| cluster4027 | 1.08 | 0.923 |
| cluster3011 | 0.84 | 0.923 |
| cluster3236 | 1.14 | 0.923 |
| cluster3476 | 1.09 | 0.923 |
| cluster3643 | 1.09 | 0.923 |
| cluster3947 | 1.06 | 0.923 |
| cluster3979 | 0.77 | 0.923 |
| cluster4088 | 1.19 | 0.923 |
| cluster2895 | -1.16 | 0.923 |
| cluster3774 | -0.80 | 0.923 |
| cluster2866 | -1.25 | 0.925 |
| cluster2937 | -0.85 | 0.925 |
| cluster3153 | 1.07 | 0.925 |
| cluster3200 | 0.76 | 0.925 |
| cluster3257 | 0.68 | 0.925 |
| cluster3282 | -1.04 | 0.925 |
| cluster3404 | -0.65 | 0.925 |
| cluster3513 | -0.77 | 0.925 |
| cluster3622 | -1.07 | 0.925 |
| cluster3750 | -1.27 | 0.925 |
| cluster3849 | 1.10 | 0.925 |
| cluster4065 | 1.05 | 0.925 |
| cluster4118 | -1.54 | 0.925 |
| cluster3614 | 1.19 | 0.925 |
| cluster3895 | 1.04 | 0.925 |
| cluster4094 | 1.00 | 0.925 |
| cluster2856 | 1.21 | 0.925 |
| cluster3726 | -0.85 | 0.925 |
| cluster1127 | -1.09 | 0.926 |
| cluster2881 | 0.78 | 0.926 |
| cluster2920 | -0.96 | 0.926 |
| cluster2927 | 1.01 | 0.926 |

| | | |
|---|---|---|
| cluster2938 | -0.61 | 0.926 |
| cluster2943 | 0.77 | 0.926 |
| cluster2964 | 0.63 | 0.926 |
| cluster2977 | 1.05 | 0.926 |
| cluster2978 | -0.75 | 0.926 |
| cluster3127 | -0.80 | 0.926 |
| cluster3143 | 0.72 | 0.926 |
| cluster3189 | -1.04 | 0.926 |
| cluster3209 | -1.05 | 0.926 |
| cluster3210 | 0.92 | 0.926 |
| cluster3234 | -0.78 | 0.926 |
| cluster3244 | 0.65 | 0.926 |
| cluster3260 | -0.74 | 0.926 |
| cluster3272 | 1.51 | 0.926 |
| cluster3285 | -0.81 | 0.926 |
| cluster3291 | 0.72 | 0.926 |
| cluster3317 | -0.99 | 0.926 |
| cluster3327 | -0.74 | 0.926 |
| cluster3340 | -0.72 | 0.926 |
| cluster3346 | -0.79 | 0.926 |
| cluster3355 | -1.14 | 0.926 |
| cluster3375 | -0.74 | 0.926 |
| cluster3377 | -1.02 | 0.926 |
| cluster3414 | -0.69 | 0.926 |
| cluster3448 | -1.08 | 0.926 |
| cluster3458 | 0.64 | 0.926 |
| cluster3522 | -1.02 | 0.926 |
| cluster3546 | -1.26 | 0.926 |
| cluster3551 | -0.81 | 0.926 |
| cluster3603 | 1.09 | 0.926 |
| cluster3628 | 1.05 | 0.926 |
| cluster3659 | -1.10 | 0.926 |
| cluster3740 | 1.06 | 0.926 |
| cluster3821 | -0.91 | 0.926 |
| cluster3837 | 1.06 | 0.926 |
| cluster3891 | 0.87 | 0.926 |
| cluster3899 | 1.07 | 0.926 |
| cluster3943 | 1.06 | 0.926 |
| cluster3966 | 0.98 | 0.926 |
| cluster3978 | -0.95 | 0.926 |
| cluster3998 | 1.29 | 0.926 |
| cluster4023 | -1.01 | 0.926 |
| cluster4055 | -1.09 | 0.926 |
| cluster4082 | 1.10 | 0.926 |

| | | |
|---|---|---|
| cluster4110 | 1.03 | 0.926 |
| cluster4115 | -1.13 | 0.926 |
| cluster4123 | -1.04 | 0.926 |
| cluster4124 | -1.00 | 0.926 |
| cluster3026 | -1.01 | 0.927 |
| cluster3117 | 1.03 | 0.927 |
| cluster3578 | -0.97 | 0.927 |
| cluster3647 | 0.96 | 0.927 |
| cluster3725 | 0.97 | 0.927 |
| cluster3836 | 1.00 | 0.927 |
| cluster3939 | 0.93 | 0.927 |
| cluster4028 | 0.98 | 0.927 |
| cluster4036 | 0.92 | 0.927 |
| cluster2924 | 1.11 | 0.928 |
| cluster3140 | 0.66 | 0.932 |
| cluster3195 | -0.66 | 0.932 |
| cluster3872 | 0.99 | 0.932 |
| cluster2942 | 0.93 | 0.932 |
| cluster2951 | 0.94 | 0.932 |
| cluster2963 | -0.72 | 0.932 |
| cluster3014 | -1.00 | 0.932 |
| cluster3066 | 0.63 | 0.932 |
| cluster3074 | -1.04 | 0.932 |
| cluster3118 | 0.73 | 0.932 |
| cluster3119 | -0.66 | 0.932 |
| cluster3158 | 0.88 | 0.932 |
| cluster3164 | -0.95 | 0.932 |
| cluster3227 | -0.90 | 0.932 |
| cluster3271 | -0.59 | 0.932 |
| cluster3274 | 0.98 | 0.932 |
| cluster3336 | -0.73 | 0.932 |
| cluster3401 | 0.85 | 0.932 |
| cluster3429 | -0.66 | 0.932 |
| cluster3437 | -0.73 | 0.932 |
| cluster3467 | 0.91 | 0.932 |
| cluster3490 | -0.90 | 0.932 |
| cluster3494 | -0.66 | 0.932 |
| cluster3624 | 0.95 | 0.932 |
| cluster3699 | 0.93 | 0.932 |
| cluster3746 | 0.68 | 0.932 |
| cluster3777 | -0.83 | 0.932 |
| cluster3799 | 0.70 | 0.932 |
| cluster3828 | 0.91 | 0.932 |
| cluster3831 | 0.93 | 0.932 |

| | | |
|---|---|---|
| cluster3951 | 0.91 | 0.932 |
| cluster3960 | -1.00 | 0.932 |
| cluster4014 | 1.00 | 0.932 |
| cluster4042 | -1.00 | 0.932 |
| cluster4077 | 0.98 | 0.932 |
| cluster4128 | 1.15 | 0.932 |
| cluster3387 | 0.97 | 0.934 |
| cluster2945 | -1.05 | 0.935 |
| cluster3863 | 0.73 | 0.938 |
| cluster2858 | -0.58 | 0.939 |
| cluster3909 | 0.89 | 0.939 |
| cluster3850 | -1.14 | 0.940 |
| cluster3298 | -0.65 | 0.942 |
| cluster3997 | 0.89 | 0.942 |
| cluster3154 | 0.56 | 0.944 |
| cluster3252 | 1.11 | 0.944 |
| cluster3721 | -0.74 | 0.950 |
| cluster3864 | 0.84 | 0.952 |
| cluster3202 | 0.72 | 0.953 |
| cluster3479 | -0.89 | 0.953 |
| cluster3613 | 0.72 | 0.953 |
| cluster4072 | 0.91 | 0.953 |
| cluster3879 | 0.83 | 0.953 |
| cluster2990 | 0.86 | 0.956 |
| cluster3533 | 0.67 | 0.956 |
| cluster3549 | 0.57 | 0.956 |
| cluster3679 | 0.65 | 0.956 |
| cluster3709 | -0.64 | 0.956 |
| cluster3963 | -0.92 | 0.956 |
| cluster3273 | 0.52 | 0.957 |
| cluster3911 | 0.91 | 0.957 |
| cluster3843 | 0.65 | 0.958 |
| cluster3006 | -0.85 | 0.958 |
| cluster3038 | 0.86 | 0.958 |
| cluster3107 | -0.66 | 0.958 |
| cluster3163 | -0.91 | 0.958 |
| cluster3560 | 0.84 | 0.958 |
| cluster3616 | -0.81 | 0.958 |
| cluster3696 | -0.77 | 0.958 |
| cluster3754 | 0.63 | 0.958 |
| cluster3759 | -0.62 | 0.958 |
| cluster3793 | 0.86 | 0.958 |
| cluster3944 | 0.83 | 0.958 |
| cluster3636 | -0.58 | 0.961 |

| | | |
|---|---|---|
| cluster2931 | -0.84 | 0.962 |
| cluster3047 | -0.77 | 0.962 |
| cluster3053 | -0.70 | 0.962 |
| cluster3073 | -0.67 | 0.962 |
| cluster3535 | -0.84 | 0.962 |
| cluster2880 | -0.92 | 0.965 |
| cluster2905 | -0.52 | 0.965 |
| cluster3042 | 0.81 | 0.965 |
| cluster3077 | -0.63 | 0.965 |
| cluster3204 | -0.61 | 0.965 |
| cluster3339 | -0.54 | 0.965 |
| cluster3370 | 0.78 | 0.965 |
| cluster3388 | -0.81 | 0.965 |
| cluster3079 | -0.54 | 0.967 |
| cluster2941 | 0.51 | 0.968 |
| cluster2955 | -0.60 | 0.968 |
| cluster2994 | 0.65 | 0.968 |
| cluster3029 | 0.75 | 0.968 |
| cluster3099 | 0.56 | 0.968 |
| cluster3295 | 0.50 | 0.968 |
| cluster3344 | -0.63 | 0.968 |
| cluster3433 | -0.63 | 0.968 |
| cluster3514 | -0.63 | 0.968 |
| cluster3627 | -0.72 | 0.968 |
| cluster3727 | -0.80 | 0.968 |
| cluster3792 | -0.71 | 0.968 |
| cluster3829 | 0.79 | 0.968 |
| cluster3832 | 0.82 | 0.968 |
| cluster3876 | 0.53 | 0.968 |
| cluster3894 | -0.78 | 0.968 |
| cluster3937 | 0.65 | 0.968 |
| cluster3964 | 0.88 | 0.968 |
| cluster4019 | 0.95 | 0.968 |
| cluster3973 | -0.95 | 0.969 |
| cluster2855 | 0.75 | 0.970 |
| cluster2873 | -0.48 | 0.970 |
| cluster2874 | 0.55 | 0.970 |
| cluster2879 | -0.57 | 0.970 |
| cluster2885 | -0.74 | 0.970 |
| cluster2888 | 0.78 | 0.970 |
| cluster2891 | -0.79 | 0.970 |
| cluster2893 | 0.97 | 0.970 |
| cluster2906 | -0.51 | 0.970 |
| cluster2909 | 0.50 | 0.970 |

| | | |
|---|---|---|
| cluster2932 | -0.55 | 0.970 |
| cluster2934 | -0.50 | 0.970 |
| cluster2935 | -0.47 | 0.970 |
| cluster2940 | -0.71 | 0.970 |
| cluster2944 | -0.66 | 0.970 |
| cluster2985 | 0.47 | 0.970 |
| cluster3046 | -0.57 | 0.970 |
| cluster3055 | 0.50 | 0.970 |
| cluster3072 | -0.87 | 0.970 |
| cluster3078 | 0.46 | 0.970 |
| cluster3087 | 0.63 | 0.970 |
| cluster3093 | 0.64 | 0.970 |
| cluster3106 | 0.44 | 0.970 |
| cluster3123 | -0.46 | 0.970 |
| cluster3144 | -0.78 | 0.970 |
| cluster3157 | -0.45 | 0.970 |
| cluster3188 | -0.52 | 0.970 |
| cluster3197 | -0.55 | 0.970 |
| cluster3198 | 0.70 | 0.970 |
| cluster3199 | -0.55 | 0.970 |
| cluster3201 | 0.47 | 0.970 |
| cluster3206 | 0.68 | 0.970 |
| cluster3211 | 0.75 | 0.970 |
| cluster3240 | 0.48 | 0.970 |
| cluster3243 | -0.50 | 0.970 |
| cluster3246 | 0.49 | 0.970 |
| cluster3264 | -0.53 | 0.970 |
| cluster3300 | 0.53 | 0.970 |
| cluster3322 | 0.67 | 0.970 |
| cluster3350 | -0.68 | 0.970 |
| cluster3362 | -0.49 | 0.970 |
| cluster3373 | 0.65 | 0.970 |
| cluster3380 | -0.58 | 0.970 |
| cluster3389 | 0.73 | 0.970 |
| cluster3421 | 0.79 | 0.970 |
| cluster3440 | -0.50 | 0.970 |
| cluster3459 | -0.79 | 0.970 |
| cluster3460 | 0.73 | 0.970 |
| cluster3464 | 0.62 | 0.970 |
| cluster3477 | -0.61 | 0.970 |
| cluster3493 | 0.84 | 0.970 |
| cluster3500 | -0.95 | 0.970 |
| cluster3516 | -0.82 | 0.970 |
| cluster3524 | -0.56 | 0.970 |

| | | |
|---|---|---|
| cluster3553 | 0.66 | 0.970 |
| cluster3573 | -0.69 | 0.970 |
| cluster3599 | -0.80 | 0.970 |
| cluster3609 | 0.76 | 0.970 |
| cluster3625 | 0.72 | 0.970 |
| cluster3660 | 0.70 | 0.970 |
| cluster3691 | 0.70 | 0.970 |
| cluster3712 | 0.55 | 0.970 |
| cluster3718 | -0.69 | 0.970 |
| cluster3731 | 0.57 | 0.970 |
| cluster3732 | 0.67 | 0.970 |
| cluster3737 | 0.41 | 0.970 |
| cluster3739 | -0.44 | 0.970 |
| cluster3765 | -0.53 | 0.970 |
| cluster3775 | 0.74 | 0.970 |
| cluster3789 | 0.61 | 0.970 |
| cluster3803 | 0.77 | 0.970 |
| cluster3825 | -0.70 | 0.970 |
| cluster3826 | -0.66 | 0.970 |
| cluster3848 | -0.76 | 0.970 |
| cluster3862 | 0.71 | 0.970 |
| cluster3906 | 0.69 | 0.970 |
| cluster3908 | 0.65 | 0.970 |
| cluster3923 | -0.70 | 0.970 |
| cluster3934 | 0.71 | 0.970 |
| cluster3938 | 0.66 | 0.970 |
| cluster3958 | -0.83 | 0.970 |
| cluster3967 | -0.57 | 0.970 |
| cluster3972 | 0.65 | 0.970 |
| cluster3990 | 0.50 | 0.970 |
| cluster4026 | -0.76 | 0.970 |
| cluster4050 | 0.69 | 0.970 |
| cluster4127 | -0.96 | 0.970 |
| cluster3149 | 0.39 | 0.972 |
| cluster2860 | -0.47 | 0.973 |
| cluster3286 | -0.39 | 0.973 |
| cluster3402 | 0.46 | 0.973 |
| cluster3456 | -0.43 | 0.973 |
| cluster3916 | 0.62 | 0.973 |
| cluster3982 | 0.61 | 0.973 |
| cluster3028 | 0.64 | 0.973 |
| cluster3218 | 0.67 | 0.974 |
| cluster3991 | 0.61 | 0.974 |
| cluster2872 | -0.43 | 0.982 |

| | | |
|---|---|---|
| cluster2898 | 0.37 | 0.982 |
| cluster2907 | -0.63 | 0.982 |
| cluster2910 | -0.57 | 0.982 |
| cluster2950 | -0.31 | 0.982 |
| cluster2973 | 0.51 | 0.982 |
| cluster2997 | -0.52 | 0.982 |
| cluster2998 | -0.61 | 0.982 |
| cluster3013 | 0.30 | 0.982 |
| cluster3025 | -0.33 | 0.982 |
| cluster3031 | -0.30 | 0.982 |
| cluster3033 | -0.40 | 0.982 |
| cluster3037 | -0.50 | 0.982 |
| cluster3043 | -0.56 | 0.982 |
| cluster3051 | 0.49 | 0.982 |
| cluster3062 | -0.43 | 0.982 |
| cluster3064 | 0.41 | 0.982 |
| cluster3070 | 0.34 | 0.982 |
| cluster3088 | -0.42 | 0.982 |
| cluster3108 | 0.58 | 0.982 |
| cluster3120 | 0.41 | 0.982 |
| cluster3132 | 0.36 | 0.982 |
| cluster3165 | -0.67 | 0.982 |
| cluster3166 | -0.31 | 0.982 |
| cluster3169 | 0.62 | 0.982 |
| cluster3208 | -0.29 | 0.982 |
| cluster3219 | 0.52 | 0.982 |
| cluster3239 | -0.28 | 0.982 |
| cluster3245 | 0.65 | 0.982 |
| cluster3255 | -0.67 | 0.982 |
| cluster3263 | 0.48 | 0.982 |
| cluster3269 | -0.61 | 0.982 |
| cluster3283 | 0.33 | 0.982 |
| cluster3299 | 0.59 | 0.982 |
| cluster3303 | -0.48 | 0.982 |
| cluster3305 | 0.60 | 0.982 |
| cluster3316 | 0.46 | 0.982 |
| cluster3335 | 0.38 | 0.982 |
| cluster3345 | 0.50 | 0.982 |
| cluster3352 | -0.50 | 0.982 |
| cluster3358 | -0.41 | 0.982 |
| cluster3364 | -0.59 | 0.982 |
| cluster3369 | 0.52 | 0.982 |
| cluster3372 | 0.30 | 0.982 |
| cluster3376 | -0.29 | 0.982 |

| | | |
|---|---|---|
| cluster3385 | 0.52 | 0.982 |
| cluster3391 | -0.33 | 0.982 |
| cluster3411 | 0.53 | 0.982 |
| cluster3417 | -0.47 | 0.982 |
| cluster3426 | -0.37 | 0.982 |
| cluster3428 | -0.40 | 0.982 |
| cluster3445 | -0.36 | 0.982 |
| cluster3472 | -0.39 | 0.982 |
| cluster3491 | 0.54 | 0.982 |
| cluster3509 | 0.43 | 0.982 |
| cluster3521 | -0.46 | 0.982 |
| cluster3531 | 0.51 | 0.982 |
| cluster3548 | 0.33 | 0.982 |
| cluster3558 | 0.54 | 0.982 |
| cluster3565 | -0.52 | 0.982 |
| cluster3579 | -0.41 | 0.982 |
| cluster3582 | -0.38 | 0.982 |
| cluster3585 | -0.37 | 0.982 |
| cluster3588 | -0.51 | 0.982 |
| cluster3593 | -0.50 | 0.982 |
| cluster3601 | 0.59 | 0.982 |
| cluster3606 | 0.61 | 0.982 |
| cluster3607 | 0.49 | 0.982 |
| cluster3608 | -0.35 | 0.982 |
| cluster3632 | -0.33 | 0.982 |
| cluster3637 | 0.32 | 0.982 |
| cluster3648 | 0.31 | 0.982 |
| cluster3654 | 0.44 | 0.982 |
| cluster3658 | -0.44 | 0.982 |
| cluster3665 | -0.53 | 0.982 |
| cluster3667 | -0.43 | 0.982 |
| cluster3673 | 0.59 | 0.982 |
| cluster3680 | 0.46 | 0.982 |
| cluster3682 | -0.60 | 0.982 |
| cluster3684 | -0.56 | 0.982 |
| cluster3688 | 0.62 | 0.982 |
| cluster3689 | -0.72 | 0.982 |
| cluster3692 | -0.51 | 0.982 |
| cluster3697 | -0.43 | 0.982 |
| cluster3701 | -0.33 | 0.982 |
| cluster3706 | 0.56 | 0.982 |
| cluster3708 | 0.55 | 0.982 |
| cluster3716 | -0.45 | 0.982 |
| cluster3720 | 0.49 | 0.982 |

| | | |
|---|---|---|
| cluster3728 | -0.35 | 0.982 |
| cluster3729 | -0.47 | 0.982 |
| cluster3751 | 0.54 | 0.982 |
| cluster3756 | -0.60 | 0.982 |
| cluster3758 | -0.49 | 0.982 |
| cluster3760 | 0.35 | 0.982 |
| cluster3763 | -0.50 | 0.982 |
| cluster3767 | 0.58 | 0.982 |
| cluster3784 | 0.42 | 0.982 |
| cluster3809 | 0.56 | 0.982 |
| cluster3833 | 0.50 | 0.982 |
| cluster3846 | -0.36 | 0.982 |
| cluster3856 | 0.52 | 0.982 |
| cluster3859 | -0.64 | 0.982 |
| cluster3881 | 0.59 | 0.982 |
| cluster3888 | -0.61 | 0.982 |
| cluster3918 | 0.51 | 0.982 |
| cluster3929 | 0.61 | 0.982 |
| cluster3954 | 0.57 | 0.982 |
| cluster3959 | 0.56 | 0.982 |
| cluster3962 | 0.46 | 0.982 |
| cluster3984 | -0.55 | 0.982 |
| cluster3986 | 0.47 | 0.982 |
| cluster3996 | 0.48 | 0.982 |
| cluster3999 | -0.46 | 0.982 |
| cluster4006 | -0.50 | 0.982 |
| cluster4021 | 0.60 | 0.982 |
| cluster4025 | 0.43 | 0.982 |
| cluster4030 | 0.56 | 0.982 |
| cluster4039 | 0.46 | 0.982 |
| cluster4058 | 0.57 | 0.982 |
| cluster4061 | 0.47 | 0.982 |
| cluster4096 | 0.56 | 0.982 |
| cluster3150 | -0.46 | 0.983 |
| cluster3058 | 0.26 | 0.984 |
| cluster2976 | 0.42 | 0.986 |
| cluster2988 | 0.40 | 0.986 |
| cluster3060 | 0.47 | 0.986 |
| cluster3276 | 0.29 | 0.986 |
| cluster3791 | -0.47 | 0.986 |
| cluster3955 | -0.48 | 0.986 |
| cluster4017 | 0.42 | 0.986 |
| cluster2857 | 0.07 | 0.988 |
| cluster2859 | -0.31 | 0.988 |

| | | |
|---|---|---|
| cluster2861 | 0.27 | 0.988 |
| cluster2863 | 0.18 | 0.988 |
| cluster2870 | -0.15 | 0.988 |
| cluster2871 | 0.31 | 0.988 |
| cluster2875 | -0.15 | 0.988 |
| cluster2878 | -0.26 | 0.988 |
| cluster2883 | -0.08 | 0.988 |
| cluster2884 | -0.26 | 0.988 |
| cluster2889 | 0.40 | 0.988 |
| cluster2897 | -0.28 | 0.988 |
| cluster2911 | -0.22 | 0.988 |
| cluster2916 | -0.10 | 0.988 |
| cluster2919 | 0.23 | 0.988 |
| cluster2921 | -0.13 | 0.988 |
| cluster2923 | -0.12 | 0.988 |
| cluster2926 | 0.20 | 0.988 |
| cluster2929 | -0.10 | 0.988 |
| cluster2933 | 0.13 | 0.988 |
| cluster2939 | -0.32 | 0.988 |
| cluster2949 | 0.25 | 0.988 |
| cluster2952 | 0.32 | 0.988 |
| cluster2956 | 0.11 | 0.988 |
| cluster2958 | -0.29 | 0.988 |
| cluster2961 | 0.11 | 0.988 |
| cluster2965 | -0.24 | 0.988 |
| cluster2968 | 0.18 | 0.988 |
| cluster2970 | 0.31 | 0.988 |
| cluster2971 | -0.22 | 0.988 |
| cluster2979 | 0.10 | 0.988 |
| cluster2981 | 0.33 | 0.988 |
| cluster2983 | 0.30 | 0.988 |
| cluster2986 | 0.44 | 0.988 |
| cluster2989 | 0.09 | 0.988 |
| cluster2991 | 0.13 | 0.988 |
| cluster2993 | 0.12 | 0.988 |
| cluster2996 | 0.07 | 0.988 |
| cluster2999 | -0.13 | 0.988 |
| cluster3001 | 0.09 | 0.988 |
| cluster3002 | -0.08 | 0.988 |
| cluster3005 | -0.24 | 0.988 |
| cluster3007 | 0.37 | 0.988 |
| cluster3019 | 0.37 | 0.988 |
| cluster3020 | -0.39 | 0.988 |
| cluster3021 | -0.09 | 0.988 |

| | | |
|---|---|---|
| cluster3022 | -0.36 | 0.988 |
| cluster3023 | 0.18 | 0.988 |
| cluster3027 | 0.11 | 0.988 |
| cluster3032 | 0.11 | 0.988 |
| cluster3036 | 0.10 | 0.988 |
| cluster3041 | -0.33 | 0.988 |
| cluster3048 | -0.12 | 0.988 |
| cluster3049 | -0.07 | 0.988 |
| cluster3054 | -0.21 | 0.988 |
| cluster3059 | 0.16 | 0.988 |
| cluster3061 | -0.17 | 0.988 |
| cluster3067 | 0.22 | 0.988 |
| cluster3089 | 0.15 | 0.988 |
| cluster3091 | 0.11 | 0.988 |
| cluster3092 | -0.18 | 0.988 |
| cluster3096 | -0.16 | 0.988 |
| cluster3098 | -0.29 | 0.988 |
| cluster3101 | 0.19 | 0.988 |
| cluster3104 | 0.06 | 0.988 |
| cluster3112 | 0.35 | 0.988 |
| cluster3116 | 0.16 | 0.988 |
| cluster3124 | -0.28 | 0.988 |
| cluster3125 | 0.28 | 0.988 |
| cluster3128 | -0.09 | 0.988 |
| cluster3134 | -0.21 | 0.988 |
| cluster3141 | -0.10 | 0.988 |
| cluster3145 | 0.26 | 0.988 |
| cluster3147 | 0.11 | 0.988 |
| cluster3148 | -0.08 | 0.988 |
| cluster3152 | 0.33 | 0.988 |
| cluster3155 | 0.42 | 0.988 |
| cluster3156 | -0.21 | 0.988 |
| cluster3170 | -0.12 | 0.988 |
| cluster3174 | -0.11 | 0.988 |
| cluster3181 | -0.18 | 0.988 |
| cluster3183 | -0.11 | 0.988 |
| cluster3214 | -0.28 | 0.988 |
| cluster3215 | -0.16 | 0.988 |
| cluster3216 | -0.42 | 0.988 |
| cluster3222 | -0.11 | 0.988 |
| cluster3228 | 0.31 | 0.988 |
| cluster3232 | 0.39 | 0.988 |
| cluster3235 | 0.25 | 0.988 |
| cluster3238 | 0.19 | 0.988 |

| | | |
|---|---|---|
| cluster3241 | -0.19 | 0.988 |
| cluster3248 | -0.44 | 0.988 |
| cluster3253 | 0.39 | 0.988 |
| cluster3258 | 0.15 | 0.988 |
| cluster3262 | -0.20 | 0.988 |
| cluster3266 | -0.14 | 0.988 |
| cluster3268 | -0.30 | 0.988 |
| cluster3277 | 0.23 | 0.988 |
| cluster3278 | 0.13 | 0.988 |
| cluster3280 | 0.29 | 0.988 |
| cluster3281 | 0.42 | 0.988 |
| cluster3289 | -0.07 | 0.988 |
| cluster3294 | -0.08 | 0.988 |
| cluster3296 | -0.29 | 0.988 |
| cluster3302 | 0.11 | 0.988 |
| cluster3304 | 0.11 | 0.988 |
| cluster3307 | -0.13 | 0.988 |
| cluster3308 | 0.10 | 0.988 |
| cluster3309 | 0.28 | 0.988 |
| cluster3310 | -0.25 | 0.988 |
| cluster3312 | -0.26 | 0.988 |
| cluster3313 | -0.37 | 0.988 |
| cluster3314 | -0.29 | 0.988 |
| cluster3320 | -0.18 | 0.988 |
| cluster3321 | -0.36 | 0.988 |
| cluster3323 | -0.13 | 0.988 |
| cluster3324 | -0.08 | 0.988 |
| cluster3329 | 0.34 | 0.988 |
| cluster3331 | -0.32 | 0.988 |
| cluster3333 | 0.22 | 0.988 |
| cluster3338 | 0.31 | 0.988 |
| cluster3347 | -0.11 | 0.988 |
| cluster3348 | -0.22 | 0.988 |
| cluster3351 | -0.10 | 0.988 |
| cluster3353 | 0.27 | 0.988 |
| cluster3354 | 0.25 | 0.988 |
| cluster3359 | -0.29 | 0.988 |
| cluster3361 | -0.17 | 0.988 |
| cluster3378 | 0.25 | 0.988 |
| cluster3384 | -0.24 | 0.988 |
| cluster3386 | 0.20 | 0.988 |
| cluster3394 | 0.11 | 0.988 |
| cluster3396 | -0.30 | 0.988 |
| cluster3399 | 0.22 | 0.988 |

| | | |
|---|---|---|
| cluster3403 | -0.31 | 0.988 |
| cluster3407 | -0.16 | 0.988 |
| cluster3412 | -0.13 | 0.988 |
| cluster3420 | 0.20 | 0.988 |
| cluster3423 | -0.13 | 0.988 |
| cluster3447 | 0.10 | 0.988 |
| cluster3470 | 0.17 | 0.988 |
| cluster3478 | -0.17 | 0.988 |
| cluster3480 | -0.25 | 0.988 |
| cluster3483 | 0.42 | 0.988 |
| cluster3492 | 0.30 | 0.988 |
| cluster3495 | 0.20 | 0.988 |
| cluster3498 | -0.28 | 0.988 |
| cluster3499 | 0.19 | 0.988 |
| cluster3502 | 0.29 | 0.988 |
| cluster3505 | -0.38 | 0.988 |
| cluster3506 | 0.22 | 0.988 |
| cluster3523 | 0.40 | 0.988 |
| cluster3527 | 0.26 | 0.988 |
| cluster3529 | 0.40 | 0.988 |
| cluster3530 | -0.41 | 0.988 |
| cluster3539 | -0.45 | 0.988 |
| cluster3541 | 0.24 | 0.988 |
| cluster3555 | 0.31 | 0.988 |
| cluster3557 | -0.18 | 0.988 |
| cluster3559 | -0.38 | 0.988 |
| cluster3561 | -0.25 | 0.988 |
| cluster3562 | -0.21 | 0.988 |
| cluster3566 | -0.30 | 0.988 |
| cluster3583 | 0.28 | 0.988 |
| cluster3586 | -0.29 | 0.988 |
| cluster3589 | -0.21 | 0.988 |
| cluster3592 | 0.19 | 0.988 |
| cluster3596 | -0.21 | 0.988 |
| cluster3600 | 0.23 | 0.988 |
| cluster3602 | -0.27 | 0.988 |
| cluster3605 | 0.12 | 0.988 |
| cluster3612 | 0.34 | 0.988 |
| cluster3615 | 0.16 | 0.988 |
| cluster3620 | -0.08 | 0.988 |
| cluster3621 | 0.38 | 0.988 |
| cluster3655 | -0.17 | 0.988 |
| cluster3663 | -0.28 | 0.988 |
| cluster3664 | -0.25 | 0.988 |

| | | |
|---|---|---|
| cluster3668 | 0.39 | 0.988 |
| cluster3669 | -0.15 | 0.988 |
| cluster3670 | -0.19 | 0.988 |
| cluster3671 | 0.31 | 0.988 |
| cluster3686 | 0.40 | 0.988 |
| cluster3687 | 0.41 | 0.988 |
| cluster3695 | -0.11 | 0.988 |
| cluster3700 | 0.43 | 0.988 |
| cluster3707 | -0.16 | 0.988 |
| cluster3713 | 0.16 | 0.988 |
| cluster3717 | -0.25 | 0.988 |
| cluster3733 | 0.31 | 0.988 |
| cluster3736 | -0.29 | 0.988 |
| cluster3741 | 0.25 | 0.988 |
| cluster3743 | 0.34 | 0.988 |
| cluster3747 | 0.33 | 0.988 |
| cluster3748 | -0.15 | 0.988 |
| cluster3749 | -0.30 | 0.988 |
| cluster3755 | -0.34 | 0.988 |
| cluster3764 | -0.33 | 0.988 |
| cluster3769 | 0.13 | 0.988 |
| cluster3770 | 0.28 | 0.988 |
| cluster3779 | 0.21 | 0.988 |
| cluster3782 | 0.14 | 0.988 |
| cluster3785 | -0.19 | 0.988 |
| cluster3790 | 0.23 | 0.988 |
| cluster3795 | -0.28 | 0.988 |
| cluster3797 | -0.43 | 0.988 |
| cluster3798 | 0.20 | 0.988 |
| cluster3806 | -0.20 | 0.988 |
| cluster3808 | 0.23 | 0.988 |
| cluster3816 | -0.37 | 0.988 |
| cluster3819 | 0.24 | 0.988 |
| cluster3820 | -0.24 | 0.988 |
| cluster3823 | -0.09 | 0.988 |
| cluster3827 | 0.10 | 0.988 |
| cluster3835 | -0.22 | 0.988 |
| cluster3838 | 0.26 | 0.988 |
| cluster3847 | 0.13 | 0.988 |
| cluster3851 | -0.11 | 0.988 |
| cluster3854 | -0.29 | 0.988 |
| cluster3857 | 0.28 | 0.988 |
| cluster3858 | 0.28 | 0.988 |
| cluster3868 | -0.12 | 0.988 |

| | | |
|---|---|---|
| cluster3874 | -0.11 | 0.988 |
| cluster3877 | -0.07 | 0.988 |
| cluster3883 | -0.45 | 0.988 |
| cluster3886 | 0.11 | 0.988 |
| cluster3887 | -0.13 | 0.988 |
| cluster3890 | -0.09 | 0.988 |
| cluster3900 | -0.28 | 0.988 |
| cluster3903 | -0.14 | 0.988 |
| cluster3907 | -0.42 | 0.988 |
| cluster3912 | -0.08 | 0.988 |
| cluster3921 | 0.35 | 0.988 |
| cluster3922 | -0.39 | 0.988 |
| cluster3926 | -0.39 | 0.988 |
| cluster3927 | 0.27 | 0.988 |
| cluster3928 | -0.37 | 0.988 |
| cluster3946 | -0.32 | 0.988 |
| cluster3961 | 0.39 | 0.988 |
| cluster3988 | -0.13 | 0.988 |
| cluster3989 | 0.24 | 0.988 |
| cluster4003 | -0.12 | 0.988 |
| cluster4004 | -0.27 | 0.988 |
| cluster4005 | -0.15 | 0.988 |
| cluster4007 | 0.12 | 0.988 |
| cluster4020 | -0.28 | 0.988 |
| cluster4022 | -0.21 | 0.988 |
| cluster4035 | 0.10 | 0.988 |
| cluster4040 | 0.28 | 0.988 |
| cluster4041 | -0.36 | 0.988 |
| cluster4043 | -0.21 | 0.988 |
| cluster4053 | -0.41 | 0.988 |
| cluster4059 | 0.18 | 0.988 |
| cluster4063 | -0.33 | 0.988 |
| cluster4066 | -0.30 | 0.988 |
| cluster4075 | -0.30 | 0.988 |
| cluster4086 | 0.24 | 0.988 |
| cluster4101 | 0.34 | 0.988 |
| cluster4103 | 0.31 | 0.988 |
| cluster4106 | -0.24 | 0.988 |
| cluster4108 | -0.32 | 0.988 |
| cluster4126 | 0.48 | 0.988 |
| cluster2882 | 0.03 | 0.990 |
| cluster2896 | -0.08 | 0.990 |
| cluster2917 | -0.09 | 0.990 |
| cluster2946 | -0.06 | 0.990 |

| | | |
|---|---|---|
| cluster3018 | 0.05 | 0.990 |
| cluster3034 | 0.04 | 0.990 |
| cluster3044 | 0.06 | 0.990 |
| cluster3071 | -0.03 | 0.990 |
| cluster3113 | 0.04 | 0.990 |
| cluster3115 | 0.08 | 0.990 |
| cluster3138 | 0.04 | 0.990 |
| cluster3139 | -0.05 | 0.990 |
| cluster3159 | -0.04 | 0.990 |
| cluster3167 | 0.07 | 0.990 |
| cluster3173 | -0.06 | 0.990 |
| cluster3186 | -0.10 | 0.990 |
| cluster3221 | 0.08 | 0.990 |
| cluster3224 | 0.05 | 0.990 |
| cluster3225 | 0.07 | 0.990 |
| cluster3251 | 0.05 | 0.990 |
| cluster3256 | 0.04 | 0.990 |
| cluster3284 | -0.08 | 0.990 |
| cluster3341 | 0.09 | 0.990 |
| cluster3343 | 0.09 | 0.990 |
| cluster3392 | -0.04 | 0.990 |
| cluster3409 | 0.09 | 0.990 |
| cluster3452 | 0.07 | 0.990 |
| cluster3517 | -0.04 | 0.990 |
| cluster3526 | 0.06 | 0.990 |
| cluster3554 | 0.05 | 0.990 |
| cluster3577 | 0.06 | 0.990 |
| cluster3597 | 0.03 | 0.990 |
| cluster3762 | -0.08 | 0.990 |
| cluster3805 | -0.06 | 0.990 |
| cluster3893 | 0.05 | 0.990 |
| cluster3896 | 0.07 | 0.990 |
| cluster3948 | -0.05 | 0.990 |
| cluster3949 | -0.05 | 0.990 |
| cluster3992 | 0.04 | 0.990 |
| cluster4032 | 0.05 | 0.990 |
| cluster4078 | -0.05 | 0.990 |
| cluster4092 | 0.09 | 0.990 |
| cluster4102 | -0.04 | 0.990 |
| cluster4105 | 0.07 | 0.990 |
| cluster3744 | -0.04 | 0.990 |
| cluster4012 | 0.04 | 0.991 |
| cluster3133 | 0.03 | 0.992 |
| cluster3297 | 0.03 | 0.992 |

| | | |
|---|---|---|
| cluster3993 | -0.03 | 0.992 |
| cluster3705 | -0.02 | 0.994 |
| cluster3722 | -0.03 | 0.994 |
| cluster3328 | -0.01 | 0.995 |
| cluster3390 | 0.02 | 0.995 |
| cluster3406 | -0.01 | 0.995 |
| cluster3511 | -0.02 | 0.995 |
| cluster3547 | 0.01 | 0.995 |
| cluster3623 | -0.01 | 0.995 |
| cluster3794 | -0.01 | 0.995 |
| cluster3861 | 0.01 | 0.995 |
| cluster3882 | -0.02 | 0.995 |
| cluster3884 | 0.01 | 0.995 |
| cluster3935 | -0.01 | 0.996 |
| cluster3873 | 0.00 | 0.998 |
| cluster3184 | 0.00 | 0.999 |

**Supplementary Table 14.** edgeR results with RSEM counts - Dermic vs nasopharyngeal

| ID | log2FoldChange | Adjusted p value |
|---|---|---|
| cluster3845 | -13.22 | 1.16E-05 |
| cluster4085 | -11.98 | 0.0004 |
| cluster3800 | -10.44 | 0.0008 |
| cluster3919 | -13.34 | 0.0018 |
| cluster3753 | -12.52 | 0.0020 |
| cluster4015 | -8.14 | 0.0049 |
| cluster4087 | 4.89 | 0.0057 |
| cluster3544 | -6.84 | 0.0147 |
| cluster4080 | -9.09 | 0.0150 |
| cluster3666 | 4.22 | 0.0471 |
| cluster4114 | 4.44 | 0.0498 |
| cluster3510 | -11.88 | 0.0498 |
| cluster4049 | -11.89 | 0.0504 |
| cluster3925 | -11.49 | 0.0505 |
| cluster4097 | -11.08 | 0.0596 |
| cluster3981 | 3.01 | 0.0623 |
| cluster4119 | 3.61 | 0.0770 |
| cluster4067 | 4.29 | 0.0889 |
| cluster4011 | -9.57 | 0.1448 |
| cluster3901 | 3.50 | 0.1508 |
| cluster4073 | 3.47 | 0.1508 |
| cluster3250 | 3.10 | 0.1653 |
| cluster4034 | -11.53 | 0.1696 |
| cluster3306 | 3.32 | 0.2879 |
| cluster2869 | -3.90 | 0.2892 |
| cluster3474 | 3.02 | 0.3036 |
| cluster4046 | -6.11 | 0.3154 |
| cluster4047 | 3.04 | 0.3855 |
| cluster4074 | 2.82 | 0.3960 |
| cluster4070 | -6.05 | 0.3960 |
| cluster3430 | 2.94 | 0.4041 |
| cluster4002 | 2.74 | 0.4041 |
| cluster3853 | 2.86 | 0.4041 |
| cluster3869 | 2.78 | 0.4041 |
| cluster3780 | -6.34 | 0.4351 |
| cluster2894 | -4.19 | 0.4351 |
| cluster4016 | 2.33 | 0.4909 |
| cluster3436 | 2.77 | 0.4998 |
| cluster4107 | 2.59 | 0.5428 |
| cluster3865 | 2.60 | 0.5620 |
| cluster3534 | -3.34 | 0.6028 |
| cluster3650 | -3.68 | 0.6028 |

| | | |
|---|---|---|
| cluster3326 | -3.01 | 0.6028 |
| cluster3481 | -4.01 | 0.6028 |
| cluster3983 | 2.56 | 0.6028 |
| cluster3629 | 2.55 | 0.6180 |
| cluster4121 | 2.43 | 0.6180 |
| cluster4060 | 2.51 | 0.6590 |
| cluster3745 | 2.83 | 0.6590 |
| cluster3970 | 2.50 | 0.6794 |
| cluster4093 | 2.57 | 0.6794 |
| cluster4090 | -5.24 | 0.6794 |
| cluster3504 | -3.40 | 0.7574 |
| cluster3598 | 2.71 | 0.7574 |
| cluster3932 | -4.15 | 0.7574 |
| cluster4099 | 2.33 | 0.7574 |
| cluster3839 | -4.10 | 0.7574 |
| cluster3443 | 2.20 | 0.8014 |
| cluster3325 | -4.27 | 0.8641 |
| cluster3057 | -3.03 | 0.8812 |
| cluster3824 | 2.33 | 0.8812 |
| cluster3889 | 2.22 | 0.8812 |
| cluster3719 | 2.17 | 0.8812 |
| cluster4079 | 1.93 | 0.8812 |
| cluster2912 | 1.97 | 0.8812 |
| cluster3693 | 2.23 | 0.8812 |
| cluster2904 | -2.67 | 0.8950 |
| cluster3572 | 2.49 | 0.9773 |
| cluster2928 | 1.96 | 0.9870 |
| cluster3875 | 1.90 | 0.9870 |
| cluster3438 | 2.05 | 0.9870 |
| cluster4054 | 2.03 | 0.9870 |
| cluster4045 | 2.30 | 0.9870 |
| cluster3580 | 2.07 | 0.9870 |
| cluster3611 | 2.08 | 0.9870 |
| cluster3179 | 2.10 | 0.9870 |
| cluster3288 | 2.06 | 0.9870 |
| cluster3451 | 2.63 | 0.9870 |
| cluster3374 | 2.08 | 0.9870 |
| cluster3974 | 2.28 | 0.9870 |
| cluster3162 | 1.94 | 0.9870 |
| cluster3940 | 1.68 | 0.9870 |
| cluster3442 | 2.15 | 0.9870 |
| cluster3822 | 2.17 | 0.9870 |
| cluster3902 | 2.10 | 0.9870 |
| cluster3008 | 1.99 | 0.9870 |

| | | |
|---|---|---|
| cluster3009 | -2.63 | 0.9870 |
| cluster2876 | 2.25 | 0.9870 |
| cluster3024 | -2.30 | 0.9870 |
| cluster3568 | -3.51 | 0.9870 |
| cluster3880 | -3.84 | 0.9870 |
| cluster3508 | 2.01 | 0.9870 |
| cluster3431 | 2.01 | 0.9907 |
| cluster3924 | 1.96 | 0.9907 |
| cluster3815 | 2.00 | 0.9991 |
| cluster3052 | -3.53 | 1.0000 |
| cluster3100 | 2.25 | 1.0000 |
| cluster3619 | 1.65 | 1.0000 |
| cluster3097 | 1.65 | 1.0000 |
| cluster3229 | -2.41 | 1.0000 |
| cluster3952 | 1.98 | 1.0000 |
| cluster3556 | 1.97 | 1.0000 |
| cluster3742 | 1.94 | 1.0000 |
| cluster3463 | 1.92 | 1.0000 |
| cluster4064 | 1.86 | 1.0000 |
| cluster3639 | 2.16 | 1.0000 |
| cluster2900 | -2.44 | 1.0000 |
| cluster3290 | -2.52 | 1.0000 |
| cluster3270 | 1.99 | 1.0000 |
| cluster3804 | 1.95 | 1.0000 |
| cluster3424 | -3.31 | 1.0000 |
| cluster3045 | 1.56 | 1.0000 |
| cluster3393 | -3.38 | 1.0000 |
| cluster3434 | -2.27 | 1.0000 |
| cluster3383 | -3.22 | 1.0000 |
| cluster3956 | 1.83 | 1.0000 |
| cluster2995 | 1.67 | 1.0000 |
| cluster3063 | 1.58 | 1.0000 |
| cluster3432 | -2.32 | 1.0000 |
| cluster3076 | 1.77 | 1.0000 |
| cluster3652 | 1.62 | 1.0000 |
| cluster3012 | 1.91 | 1.0000 |
| cluster3365 | 1.88 | 1.0000 |
| cluster3379 | -2.49 | 1.0000 |
| cluster4100 | 1.72 | 1.0000 |
| cluster3968 | 1.68 | 1.0000 |
| cluster3207 | 1.53 | 1.0000 |
| cluster3223 | -2.38 | 1.0000 |
| cluster4029 | -3.20 | 1.0000 |
| cluster3571 | -2.49 | 1.0000 |

| | | |
|---|---|---|
| cluster3724 | 1.76 | 1.0000 |
| cluster2899 | -2.27 | 1.0000 |
| cluster3569 | 1.79 | 1.0000 |
| cluster3771 | 1.71 | 1.0000 |
| cluster3734 | 1.76 | 1.0000 |
| cluster3171 | -2.11 | 1.0000 |
| cluster3584 | 1.69 | 1.0000 |
| cluster4109 | 1.73 | 1.0000 |
| cluster3482 | -2.23 | 1.0000 |
| cluster3674 | -3.26 | 1.0000 |
| cluster3581 | -2.58 | 1.0000 |
| cluster2892 | 1.35 | 1.0000 |
| cluster3160 | -1.97 | 1.0000 |
| cluster3471 | 1.78 | 1.0000 |
| cluster3172 | -2.00 | 1.0000 |
| cluster4000 | 1.44 | 1.0000 |
| cluster3408 | -2.04 | 1.0000 |
| cluster2992 | -1.90 | 1.0000 |
| cluster3985 | 1.42 | 1.0000 |
| cluster2972 | -2.12 | 1.0000 |
| cluster3168 | 1.37 | 1.0000 |
| cluster3936 | 1.64 | 1.0000 |
| cluster3871 | 1.63 | 1.0000 |
| cluster3422 | -2.61 | 1.0000 |
| cluster3787 | -2.67 | 1.0000 |
| cluster3595 | 1.39 | 1.0000 |
| cluster4091 | -3.23 | 1.0000 |
| cluster3761 | -3.00 | 1.0000 |
| cluster3957 | 1.64 | 1.0000 |
| cluster3576 | 1.52 | 1.0000 |
| cluster2980 | -2.97 | 1.0000 |
| cluster3661 | 1.32 | 1.0000 |
| cluster3617 | -3.12 | 1.0000 |
| cluster3678 | 1.31 | 1.0000 |
| cluster3994 | -2.95 | 1.0000 |
| cluster3486 | -1.98 | 1.0000 |
| cluster3519 | -2.97 | 1.0000 |
| cluster3003 | -2.46 | 1.0000 |
| cluster4044 | 1.57 | 1.0000 |
| cluster3638 | 1.67 | 1.0000 |
| cluster3975 | 1.65 | 1.0000 |
| cluster2915 | -2.23 | 1.0000 |
| cluster3129 | -2.97 | 1.0000 |
| cluster3512 | 1.42 | 1.0000 |

| | | |
|---|---|---|
| cluster3812 | -2.34 | 1.0000 |
| cluster3866 | 1.53 | 1.0000 |
| cluster3651 | -2.72 | 1.0000 |
| cluster3905 | -3.72 | 1.0000 |
| cluster2930 | 1.41 | 1.0000 |
| cluster3563 | -1.98 | 1.0000 |
| cluster4125 | -2.91 | 1.0000 |
| cluster3645 | -3.45 | 1.0000 |
| cluster2954 | -1.91 | 1.0000 |
| cluster3987 | 1.37 | 1.0000 |
| cluster2974 | -1.89 | 1.0000 |
| cluster3111 | -2.49 | 1.0000 |
| cluster3461 | -3.25 | 1.0000 |
| cluster3349 | 1.36 | 1.0000 |
| cluster4008 | 1.47 | 1.0000 |
| cluster4081 | 1.49 | 1.0000 |
| cluster3381 | -1.80 | 1.0000 |
| cluster3552 | 1.25 | 1.0000 |
| cluster3610 | 1.62 | 1.0000 |
| cluster3543 | -2.51 | 1.0000 |
| cluster4051 | 1.69 | 1.0000 |
| cluster3681 | 1.51 | 1.0000 |
| cluster3683 | 1.56 | 1.0000 |
| cluster3081 | -1.81 | 1.0000 |
| cluster4024 | 1.46 | 1.0000 |
| cluster3971 | 1.51 | 1.0000 |
| cluster4010 | 1.45 | 1.0000 |
| cluster3363 | 1.50 | 1.0000 |
| cluster3538 | -1.98 | 1.0000 |
| cluster3441 | 1.28 | 1.0000 |
| cluster4122 | -3.14 | 1.0000 |
| cluster3710 | 1.46 | 1.0000 |
| cluster4009 | 1.38 | 1.0000 |
| cluster3520 | -3.39 | 1.0000 |
| cluster4120 | -3.00 | 1.0000 |
| cluster4071 | 1.54 | 1.0000 |
| cluster3357 | 1.22 | 1.0000 |
| cluster3342 | 1.49 | 1.0000 |
| cluster3574 | -2.63 | 1.0000 |
| cluster3332 | 1.48 | 1.0000 |
| cluster3475 | 1.49 | 1.0000 |
| cluster3142 | -2.03 | 1.0000 |
| cluster3016 | -2.40 | 1.0000 |
| cluster4037 | 1.44 | 1.0000 |

| | | |
|---|---|---|
| cluster3233 | -2.55 | 1.0000 |
| cluster3702 | 1.42 | 1.0000 |
| cluster3192 | -1.63 | 1.0000 |
| cluster3694 | -1.91 | 1.0000 |
| cluster3090 | -2.72 | 1.0000 |
| cluster3657 | 1.48 | 1.0000 |
| cluster3715 | 1.35 | 1.0000 |
| cluster3675 | -1.59 | 1.0000 |
| cluster4038 | 1.42 | 1.0000 |
| cluster3646 | 1.47 | 1.0000 |
| cluster3752 | -2.78 | 1.0000 |
| cluster3279 | 1.32 | 1.0000 |
| cluster2925 | -2.39 | 1.0000 |
| cluster3453 | -1.88 | 1.0000 |
| cluster3604 | 1.47 | 1.0000 |
| cluster3917 | 1.48 | 1.0000 |
| cluster2987 | 1.11 | 1.0000 |
| cluster4068 | 1.38 | 1.0000 |
| cluster3040 | -1.61 | 1.0000 |
| cluster3473 | 1.20 | 1.0000 |
| cluster3094 | 1.40 | 1.0000 |
| cluster2966 | 1.43 | 1.0000 |
| cluster3187 | -2.37 | 1.0000 |
| cluster3177 | -2.10 | 1.0000 |
| cluster3817 | -1.94 | 1.0000 |
| cluster3095 | -1.65 | 1.0000 |
| cluster3723 | 1.13 | 1.0000 |
| cluster3830 | 1.14 | 1.0000 |
| cluster3366 | 1.36 | 1.0000 |
| cluster3772 | -1.99 | 1.0000 |
| cluster3105 | -2.84 | 1.0000 |
| cluster3781 | 1.16 | 1.0000 |
| cluster4111 | -2.64 | 1.0000 |
| cluster3813 | 1.28 | 1.0000 |
| cluster3416 | 1.40 | 1.0000 |
| cluster4013 | -3.37 | 1.0000 |
| cluster3783 | 1.37 | 1.0000 |
| cluster3995 | -2.22 | 1.0000 |
| cluster3501 | -2.04 | 1.0000 |
| cluster3010 | -1.82 | 1.0000 |
| cluster4062 | -2.62 | 1.0000 |
| cluster4104 | 1.30 | 1.0000 |
| cluster3190 | 1.38 | 1.0000 |
| cluster3507 | -2.23 | 1.0000 |

| | | |
|---|---|---|
| cluster3633 | -1.70 | 1.0000 |
| cluster3469 | -1.65 | 1.0000 |
| cluster2854 | 1.29 | 1.0000 |
| cluster3292 | -2.55 | 1.0000 |
| cluster3677 | 1.10 | 1.0000 |
| cluster3220 | -1.77 | 1.0000 |
| cluster4018 | 1.21 | 1.0000 |
| cluster3814 | -1.59 | 1.0000 |
| cluster3778 | -1.65 | 1.0000 |
| cluster4031 | -1.62 | 1.0000 |
| cluster4057 | 1.29 | 1.0000 |
| cluster3892 | -1.95 | 1.0000 |
| cluster3334 | 1.08 | 1.0000 |
| cluster3484 | -2.47 | 1.0000 |
| cluster3704 | 1.14 | 1.0000 |
| cluster4112 | 1.31 | 1.0000 |
| cluster3618 | -2.51 | 1.0000 |
| cluster3395 | -1.56 | 1.0000 |
| cluster3703 | 1.03 | 1.0000 |
| cluster3017 | -2.01 | 1.0000 |
| cluster2867 | -2.19 | 1.0000 |
| cluster3537 | 1.27 | 1.0000 |
| cluster3545 | 1.26 | 1.0000 |
| cluster3083 | 1.03 | 1.0000 |
| cluster4084 | -1.62 | 1.0000 |
| cluster3676 | 1.29 | 1.0000 |
| cluster4083 | 1.26 | 1.0000 |
| cluster2969 | 1.28 | 1.0000 |
| cluster3217 | -1.52 | 1.0000 |
| cluster4113 | -1.72 | 1.0000 |
| cluster2887 | 1.28 | 1.0000 |
| cluster2967 | 1.13 | 1.0000 |
| cluster4069 | 1.17 | 1.0000 |
| cluster3454 | -1.49 | 1.0000 |
| cluster3337 | -1.54 | 1.0000 |
| cluster4116 | -2.48 | 1.0000 |
| cluster3867 | -1.44 | 1.0000 |
| cluster3247 | 1.12 | 1.0000 |
| cluster2922 | 1.24 | 1.0000 |
| cluster3109 | 1.31 | 1.0000 |
| cluster4117 | 1.49 | 1.0000 |
| cluster3489 | 1.24 | 1.0000 |
| cluster3410 | -1.61 | 1.0000 |
| cluster3242 | 1.09 | 1.0000 |

| | | |
|---|---|---|
| cluster3849 | 1.20 | 1.0000 |
| cluster2908 | -1.48 | 1.0000 |
| cluster3801 | 1.17 | 1.0000 |
| cluster2936 | 1.25 | 1.0000 |
| cluster3236 | 1.22 | 1.0000 |
| cluster3194 | -1.51 | 1.0000 |
| cluster4095 | 1.15 | 1.0000 |
| cluster3913 | -1.95 | 1.0000 |
| cluster3161 | -2.04 | 1.0000 |
| cluster3084 | -1.31 | 1.0000 |
| cluster3698 | -1.32 | 1.0000 |
| cluster3503 | 1.18 | 1.0000 |
| cluster3590 | 1.15 | 1.0000 |
| cluster3035 | -1.33 | 1.0000 |
| cluster2947 | 0.97 | 1.0000 |
| cluster3418 | 1.16 | 1.0000 |
| cluster3272 | 1.52 | 1.0000 |
| cluster3788 | -1.45 | 1.0000 |
| cluster3587 | -2.27 | 1.0000 |
| cluster3110 | 1.03 | 1.0000 |
| cluster3643 | 1.16 | 1.0000 |
| cluster3842 | 1.05 | 1.0000 |
| cluster3515 | -2.01 | 1.0000 |
| cluster3685 | -2.02 | 1.0000 |
| cluster3254 | -2.24 | 1.0000 |
| cluster3642 | 1.20 | 1.0000 |
| cluster3603 | 1.18 | 1.0000 |
| cluster3933 | -1.85 | 1.0000 |
| cluster3570 | 1.21 | 1.0000 |
| cluster3356 | 0.97 | 1.0000 |
| cluster3714 | -1.52 | 1.0000 |
| cluster3455 | -1.56 | 1.0000 |
| cluster3287 | -1.49 | 1.0000 |
| cluster3476 | 1.14 | 1.0000 |
| cluster3899 | 1.15 | 1.0000 |
| cluster4110 | 1.11 | 1.0000 |
| cluster3082 | -2.06 | 1.0000 |
| cluster3153 | 1.12 | 1.0000 |
| cluster3943 | 1.14 | 1.0000 |
| cluster2856 | 1.24 | 1.0000 |
| cluster3740 | 1.13 | 1.0000 |
| cluster2960 | -1.54 | 1.0000 |
| cluster3542 | 1.12 | 1.0000 |
| cluster3930 | 0.95 | 1.0000 |

| | | |
|---|---|---|
| cluster3656 | 0.95 | 1.0000 |
| cluster3371 | -2.79 | 1.0000 |
| cluster3122 | -1.37 | 1.0000 |
| cluster4027 | 1.11 | 1.0000 |
| cluster4065 | 1.11 | 1.0000 |
| cluster2977 | 1.12 | 1.0000 |
| cluster3330 | 1.01 | 1.0000 |
| cluster3690 | -1.36 | 1.0000 |
| cluster3947 | 1.09 | 1.0000 |
| cluster3855 | 1.07 | 1.0000 |
| cluster3860 | 0.96 | 1.0000 |
| cluster3631 | -1.71 | 1.0000 |
| cluster3212 | -1.61 | 1.0000 |
| cluster3457 | -2.00 | 1.0000 |
| cluster4088 | 1.16 | 1.0000 |
| cluster3998 | 1.30 | 1.0000 |
| cluster3614 | 1.19 | 1.0000 |
| cluster2890 | -1.25 | 1.0000 |
| cluster4082 | 1.13 | 1.0000 |
| cluster2914 | -1.63 | 1.0000 |
| cluster4094 | 1.03 | 1.0000 |
| cluster3415 | -1.43 | 1.0000 |
| cluster2924 | 1.20 | 1.0000 |
| cluster3776 | -1.37 | 1.0000 |
| cluster3466 | -1.48 | 1.0000 |
| cluster3462 | -1.42 | 1.0000 |
| cluster3980 | 0.83 | 1.0000 |
| cluster2927 | 1.09 | 1.0000 |
| cluster3895 | 1.06 | 1.0000 |
| cluster3837 | 1.09 | 1.0000 |
| cluster3117 | 1.10 | 1.0000 |
| cluster3135 | -1.91 | 1.0000 |
| cluster3015 | -1.63 | 1.0000 |
| cluster3425 | -1.42 | 1.0000 |
| cluster3796 | -1.81 | 1.0000 |
| cluster4028 | 1.05 | 1.0000 |
| cluster3872 | 1.06 | 1.0000 |
| cluster3628 | 1.07 | 1.0000 |
| cluster3966 | 1.03 | 1.0000 |
| cluster3000 | -1.71 | 1.0000 |
| cluster3318 | -1.36 | 1.0000 |
| cluster3210 | 0.98 | 1.0000 |
| cluster2865 | 0.86 | 1.0000 |
| cluster2864 | 0.86 | 1.0000 |

| | | |
|---|---|---|
| cluster2975 | 0.83 | 1.0000 |
| cluster3075 | -1.32 | 1.0000 |
| cluster3647 | 1.02 | 1.0000 |
| cluster3950 | 0.86 | 1.0000 |
| cluster3427 | -1.57 | 1.0000 |
| cluster3252 | 1.19 | 1.0000 |
| cluster2942 | 1.02 | 1.0000 |
| cluster3725 | 1.02 | 1.0000 |
| cluster3699 | 1.01 | 1.0000 |
| cluster3419 | -1.21 | 1.0000 |
| cluster3275 | -1.70 | 1.0000 |
| cluster3976 | -1.29 | 1.0000 |
| cluster4014 | 1.07 | 1.0000 |
| cluster3807 | -1.25 | 1.0000 |
| cluster3497 | -1.21 | 1.0000 |
| cluster3468 | -1.23 | 1.0000 |
| cluster3086 | -1.37 | 1.0000 |
| cluster3496 | 0.85 | 1.0000 |
| cluster3536 | 0.81 | 1.0000 |
| cluster4036 | 0.96 | 1.0000 |
| cluster3401 | 0.94 | 1.0000 |
| cluster3640 | -2.46 | 1.0000 |
| cluster3939 | 0.97 | 1.0000 |
| cluster3594 | -1.66 | 1.0000 |
| cluster3146 | -1.21 | 1.0000 |
| cluster3413 | -1.73 | 1.0000 |
| cluster3624 | 1.01 | 1.0000 |
| cluster3398 | -1.43 | 1.0000 |
| cluster3274 | 1.02 | 1.0000 |
| cluster4098 | 0.96 | 1.0000 |
| cluster2862 | -1.17 | 1.0000 |
| cluster3897 | 0.90 | 1.0000 |
| cluster3487 | -1.80 | 1.0000 |
| cluster4128 | 1.11 | 1.0000 |
| cluster3831 | 0.98 | 1.0000 |
| cluster3898 | 0.92 | 1.0000 |
| cluster3828 | 0.95 | 1.0000 |
| cluster3635 | -1.11 | 1.0000 |
| cluster3997 | 0.97 | 1.0000 |
| cluster3836 | 1.00 | 1.0000 |
| cluster2951 | 0.99 | 1.0000 |
| cluster2877 | 0.77 | 1.0000 |
| cluster3951 | 0.95 | 1.0000 |
| cluster3311 | 0.82 | 1.0000 |

| | | |
|---|---|---|
| cluster4072 | 1.00 | 1.0000 |
| cluster3953 | -1.59 | 1.0000 |
| cluster3891 | 0.88 | 1.0000 |
| cluster3810 | 0.80 | 1.0000 |
| cluster3786 | 1.01 | 1.0000 |
| cluster3387 | 1.01 | 1.0000 |
| cluster3920 | -1.75 | 1.0000 |
| cluster3911 | 0.98 | 1.0000 |
| cluster3230 | -1.26 | 1.0000 |
| cluster3735 | -1.31 | 1.0000 |
| cluster2853 | -1.27 | 1.0000 |
| cluster3360 | -1.37 | 1.0000 |
| cluster3904 | -1.88 | 1.0000 |
| cluster3467 | 0.93 | 1.0000 |
| cluster4077 | 0.97 | 1.0000 |
| cluster2984 | -1.27 | 1.0000 |
| cluster3175 | -1.69 | 1.0000 |
| cluster3885 | -1.64 | 1.0000 |
| cluster3178 | 0.90 | 1.0000 |
| cluster3879 | 0.90 | 1.0000 |
| cluster3915 | -1.84 | 1.0000 |
| cluster2959 | 0.77 | 1.0000 |
| cluster3065 | -1.32 | 1.0000 |
| cluster3909 | 0.93 | 1.0000 |
| cluster4033 | -1.65 | 1.0000 |
| cluster3118 | 0.78 | 1.0000 |
| cluster3941 | -1.05 | 1.0000 |
| cluster3944 | 0.91 | 1.0000 |
| cluster3030 | -1.45 | 1.0000 |
| cluster3193 | -1.33 | 1.0000 |
| cluster4019 | 1.05 | 1.0000 |
| cluster3793 | 0.93 | 1.0000 |
| cluster3560 | 0.91 | 1.0000 |
| cluster3158 | 0.88 | 1.0000 |
| cluster2990 | 0.92 | 1.0000 |
| cluster3945 | -1.59 | 1.0000 |
| cluster3766 | -1.83 | 1.0000 |
| cluster3382 | -1.15 | 1.0000 |
| cluster3319 | -1.58 | 1.0000 |
| cluster3864 | 0.88 | 1.0000 |
| cluster3136 | -1.37 | 1.0000 |
| cluster3832 | 0.91 | 1.0000 |
| cluster3450 | -1.88 | 1.0000 |
| cluster4089 | -2.02 | 1.0000 |

| | | |
|---|---|---|
| cluster3205 | -1.10 | 1.0000 |
| cluster3259 | -1.09 | 1.0000 |
| cluster3829 | 0.88 | 1.0000 |
| cluster3773 | -1.16 | 1.0000 |
| cluster3267 | 0.71 | 1.0000 |
| cluster3039 | 0.73 | 1.0000 |
| cluster3564 | 0.85 | 1.0000 |
| cluster3844 | -1.74 | 1.0000 |
| cluster3213 | -1.17 | 1.0000 |
| cluster3140 | 0.71 | 1.0000 |
| cluster2903 | -1.27 | 1.0000 |
| cluster3757 | -2.03 | 1.0000 |
| cluster3493 | 0.93 | 1.0000 |
| cluster2893 | 1.01 | 1.0000 |
| cluster3011 | 0.75 | 1.0000 |
| cluster3202 | 0.76 | 1.0000 |
| cluster3518 | -1.65 | 1.0000 |
| cluster3370 | 0.82 | 1.0000 |
| cluster3151 | -1.08 | 1.0000 |
| cluster3038 | 0.87 | 1.0000 |
| cluster3397 | -1.60 | 1.0000 |
| cluster3768 | -1.55 | 1.0000 |
| cluster3293 | 0.68 | 1.0000 |
| cluster3691 | 0.79 | 1.0000 |
| cluster3653 | -1.12 | 1.0000 |
| cluster3979 | 0.69 | 1.0000 |
| cluster3843 | 0.71 | 1.0000 |
| cluster3460 | 0.82 | 1.0000 |
| cluster3421 | 0.85 | 1.0000 |
| cluster3910 | -1.45 | 1.0000 |
| cluster4056 | -1.56 | 1.0000 |
| cluster3103 | -1.61 | 1.0000 |
| cluster3465 | -1.06 | 1.0000 |
| cluster3029 | 0.79 | 1.0000 |
| cluster3803 | 0.85 | 1.0000 |
| cluster3196 | -0.97 | 1.0000 |
| cluster3056 | -0.99 | 1.0000 |
| cluster3641 | -1.01 | 1.0000 |
| cluster3211 | 0.80 | 1.0000 |
| cluster2994 | 0.71 | 1.0000 |
| cluster3042 | 0.80 | 1.0000 |
| cluster3625 | 0.79 | 1.0000 |
| cluster3852 | 0.73 | 1.0000 |
| cluster3126 | -1.24 | 1.0000 |

| | | |
|---|---|---|
| cluster3131 | -1.52 | 1.0000 |
| cluster3609 | 0.86 | 1.0000 |
| cluster3182 | -1.27 | 1.0000 |
| cluster2948 | -1.25 | 1.0000 |
| cluster2982 | -1.14 | 1.0000 |
| cluster3662 | -1.31 | 1.0000 |
| cluster3626 | -1.01 | 1.0000 |
| cluster3862 | 0.79 | 1.0000 |
| cluster3934 | 0.77 | 1.0000 |
| cluster3435 | -1.46 | 1.0000 |
| cluster3775 | 0.81 | 1.0000 |
| cluster2888 | 0.83 | 1.0000 |
| cluster3870 | -1.40 | 1.0000 |
| cluster3143 | 0.64 | 1.0000 |
| cluster3942 | -1.35 | 1.0000 |
| cluster3914 | -1.72 | 1.0000 |
| cluster3200 | 0.66 | 1.0000 |
| cluster3389 | 0.79 | 1.0000 |
| cluster2855 | 0.81 | 1.0000 |
| cluster3257 | 0.62 | 1.0000 |
| cluster3906 | 0.75 | 1.0000 |
| cluster2881 | 0.66 | 1.0000 |
| cluster3754 | 0.65 | 1.0000 |
| cluster3405 | -1.07 | 1.0000 |
| cluster3964 | 0.82 | 1.0000 |
| cluster3834 | -1.44 | 1.0000 |
| cluster3630 | -0.92 | 1.0000 |
| cluster3525 | -1.03 | 1.0000 |
| cluster2943 | 0.66 | 1.0000 |
| cluster2957 | -1.39 | 1.0000 |
| cluster3938 | 0.72 | 1.0000 |
| cluster3634 | -0.89 | 1.0000 |
| cluster3087 | 0.68 | 1.0000 |
| cluster2953 | -0.91 | 1.0000 |
| cluster3540 | -1.29 | 1.0000 |
| cluster3198 | 0.75 | 1.0000 |
| cluster3488 | -1.24 | 1.0000 |
| cluster4050 | 0.75 | 1.0000 |
| cluster3080 | -1.10 | 1.0000 |
| cluster3206 | 0.73 | 1.0000 |
| cluster3841 | -1.33 | 1.0000 |
| cluster3660 | 0.74 | 1.0000 |
| cluster3121 | -1.02 | 1.0000 |
| cluster3050 | -1.02 | 1.0000 |

| | | |
|---|---|---|
| cluster3301 | -1.02 | 1.0000 |
| cluster3114 | -1.03 | 1.0000 |
| cluster4052 | 0.68 | 1.0000 |
| cluster3291 | 0.63 | 1.0000 |
| cluster2913 | -1.02 | 1.0000 |
| cluster3458 | 0.60 | 1.0000 |
| cluster3218 | 0.76 | 1.0000 |
| cluster3711 | -0.99 | 1.0000 |
| cluster3730 | -0.96 | 1.0000 |
| cluster3004 | -0.96 | 1.0000 |
| cluster3373 | 0.72 | 1.0000 |
| cluster3916 | 0.70 | 1.0000 |
| cluster3093 | 0.70 | 1.0000 |
| cluster3245 | 0.77 | 1.0000 |
| cluster3367 | -1.21 | 1.0000 |
| cluster3613 | 0.65 | 1.0000 |
| cluster2901 | -1.35 | 1.0000 |
| cluster3099 | 0.60 | 1.0000 |
| cluster3965 | -1.21 | 1.0000 |
| cluster3908 | 0.67 | 1.0000 |
| cluster3532 | -1.08 | 1.0000 |
| cluster4021 | 0.69 | 1.0000 |
| cluster3575 | -1.52 | 1.0000 |
| cluster3244 | 0.58 | 1.0000 |
| cluster3130 | -1.28 | 1.0000 |
| cluster3185 | -1.49 | 1.0000 |
| cluster3799 | 0.61 | 1.0000 |
| cluster3368 | -0.90 | 1.0000 |
| cluster3550 | -0.88 | 1.0000 |
| cluster3679 | 0.60 | 1.0000 |
| cluster3169 | 0.71 | 1.0000 |
| cluster3881 | 0.66 | 1.0000 |
| cluster3732 | 0.69 | 1.0000 |
| cluster3485 | -1.52 | 1.0000 |
| cluster3688 | 0.70 | 1.0000 |
| cluster3528 | -0.94 | 1.0000 |
| cluster3464 | 0.67 | 1.0000 |
| cluster2918 | -1.02 | 1.0000 |
| cluster2874 | 0.61 | 1.0000 |
| cluster3069 | -0.83 | 1.0000 |
| cluster3449 | -0.89 | 1.0000 |
| cluster3180 | -1.23 | 1.0000 |
| cluster3991 | 0.66 | 1.0000 |
| cluster3322 | 0.68 | 1.0000 |

| | | |
|---|---|---|
| cluster2964 | 0.56 | 1.0000 |
| cluster3606 | 0.67 | 1.0000 |
| cluster3315 | -0.89 | 1.0000 |
| cluster3863 | 0.61 | 1.0000 |
| cluster3533 | 0.60 | 1.0000 |
| cluster3028 | 0.67 | 1.0000 |
| cluster3977 | -1.37 | 1.0000 |
| cluster3295 | 0.54 | 1.0000 |
| cluster3982 | 0.65 | 1.0000 |
| cluster3553 | 0.66 | 1.0000 |
| cluster3746 | 0.58 | 1.0000 |
| cluster3591 | -0.91 | 1.0000 |
| cluster3265 | -1.29 | 1.0000 |
| cluster2909 | 0.53 | 1.0000 |
| cluster3972 | 0.65 | 1.0000 |
| cluster3299 | 0.67 | 1.0000 |
| cluster3601 | 0.66 | 1.0000 |
| cluster2941 | 0.53 | 1.0000 |
| cluster3203 | -1.14 | 1.0000 |
| cluster3191 | -1.30 | 1.0000 |
| cluster3644 | -0.99 | 1.0000 |
| cluster3300 | 0.57 | 1.0000 |
| cluster3263 | 0.56 | 1.0000 |
| cluster3929 | 0.67 | 1.0000 |
| cluster3767 | 0.64 | 1.0000 |
| cluster2886 | -0.87 | 1.0000 |
| cluster3369 | 0.64 | 1.0000 |
| cluster3078 | 0.51 | 1.0000 |
| cluster4096 | 0.63 | 1.0000 |
| cluster3672 | -1.18 | 1.0000 |
| cluster3444 | -0.83 | 1.0000 |
| cluster3261 | -1.14 | 1.0000 |
| cluster3066 | 0.53 | 1.0000 |
| cluster3154 | 0.50 | 1.0000 |
| cluster3549 | 0.52 | 1.0000 |
| cluster3809 | 0.61 | 1.0000 |
| cluster3085 | -1.14 | 1.0000 |
| cluster3231 | -1.39 | 1.0000 |
| cluster2902 | 0.67 | 1.0000 |
| cluster3918 | 0.62 | 1.0000 |
| cluster3219 | 0.56 | 1.0000 |
| cluster3108 | 0.63 | 1.0000 |
| cluster3055 | 0.53 | 1.0000 |
| cluster3346 | -0.89 | 1.0000 |

| | | |
|---|---|---|
| cluster2978 | -0.88 | 1.0000 |
| cluster3273 | 0.49 | 1.0000 |
| cluster3551 | -0.92 | 1.0000 |
| cluster3439 | -1.11 | 1.0000 |
| cluster3649 | -0.87 | 1.0000 |
| cluster3706 | 0.59 | 1.0000 |
| cluster3400 | -0.82 | 1.0000 |
| cluster3751 | 0.57 | 1.0000 |
| cluster3558 | 0.59 | 1.0000 |
| cluster4030 | 0.62 | 1.0000 |
| cluster4058 | 0.59 | 1.0000 |
| cluster3491 | 0.59 | 1.0000 |
| cluster3249 | -0.92 | 1.0000 |
| cluster3513 | -0.84 | 1.0000 |
| cluster4025 | 0.51 | 1.0000 |
| cluster3201 | 0.49 | 1.0000 |
| cluster3750 | -1.31 | 1.0000 |
| cluster3509 | 0.49 | 1.0000 |
| cluster3856 | 0.60 | 1.0000 |
| cluster2973 | 0.59 | 1.0000 |
| cluster3673 | 0.59 | 1.0000 |
| cluster3996 | 0.55 | 1.0000 |
| cluster3305 | 0.60 | 1.0000 |
| cluster3607 | 0.58 | 1.0000 |
| cluster3774 | -0.79 | 1.0000 |
| cluster3446 | -1.10 | 1.0000 |
| cluster3327 | -0.83 | 1.0000 |
| cluster4061 | 0.54 | 1.0000 |
| cluster3411 | 0.57 | 1.0000 |
| cluster4118 | -1.59 | 1.0000 |
| cluster3385 | 0.59 | 1.0000 |
| cluster3937 | 0.51 | 1.0000 |
| cluster3414 | -0.79 | 1.0000 |
| cluster3990 | 0.47 | 1.0000 |
| cluster3986 | 0.54 | 1.0000 |
| cluster3567 | -1.25 | 1.0000 |
| cluster3531 | 0.57 | 1.0000 |
| cluster3720 | 0.55 | 1.0000 |
| cluster3282 | -0.98 | 1.0000 |
| cluster3345 | 0.54 | 1.0000 |
| cluster3226 | -1.11 | 1.0000 |
| cluster2937 | -0.82 | 1.0000 |
| cluster3237 | -1.32 | 1.0000 |
| cluster3654 | 0.52 | 1.0000 |

| | | |
|---|---|---|
| cluster3680 | 0.53 | 1.0000 |
| cluster3448 | -1.09 | 1.0000 |
| cluster3962 | 0.53 | 1.0000 |
| cluster4126 | 0.58 | 1.0000 |
| cluster3260 | -0.80 | 1.0000 |
| cluster3355 | -1.09 | 1.0000 |
| cluster3876 | 0.45 | 1.0000 |
| cluster3068 | -0.71 | 1.0000 |
| cluster4001 | -1.20 | 1.0000 |
| cluster3336 | -0.85 | 1.0000 |
| cluster3726 | -0.80 | 1.0000 |
| cluster3954 | 0.53 | 1.0000 |
| cluster2895 | -1.09 | 1.0000 |
| cluster3737 | 0.43 | 1.0000 |
| cluster3106 | 0.43 | 1.0000 |
| cluster3127 | -0.78 | 1.0000 |
| cluster4115 | -1.14 | 1.0000 |
| cluster4039 | 0.47 | 1.0000 |
| cluster3789 | 0.49 | 1.0000 |
| cluster3155 | 0.51 | 1.0000 |
| cluster3821 | -0.90 | 1.0000 |
| cluster3840 | 0.48 | 1.0000 |
| cluster3622 | -0.98 | 1.0000 |
| cluster3978 | -0.89 | 1.0000 |
| cluster3712 | 0.46 | 1.0000 |
| cluster3578 | -0.96 | 1.0000 |
| cluster4055 | -1.05 | 1.0000 |
| cluster3833 | 0.52 | 1.0000 |
| cluster2988 | 0.48 | 1.0000 |
| cluster3969 | -0.69 | 1.0000 |
| cluster3120 | 0.46 | 1.0000 |
| cluster3708 | 0.53 | 1.0000 |
| cluster4023 | -0.98 | 1.0000 |
| cluster3721 | -0.86 | 1.0000 |
| cluster3119 | -0.76 | 1.0000 |
| cluster2866 | -1.15 | 1.0000 |
| cluster3738 | -1.06 | 1.0000 |
| cluster2963 | -0.82 | 1.0000 |
| cluster3494 | -0.77 | 1.0000 |
| cluster3811 | -0.68 | 1.0000 |
| cluster3404 | -0.70 | 1.0000 |
| cluster3285 | -0.77 | 1.0000 |
| cluster3340 | -0.72 | 1.0000 |
| cluster3687 | 0.48 | 1.0000 |

| | | |
|---|---|---|
| cluster4017 | 0.48 | 1.0000 |
| cluster3700 | 0.48 | 1.0000 |
| cluster3060 | 0.50 | 1.0000 |
| cluster3731 | 0.44 | 1.0000 |
| cluster3026 | -1.01 | 1.0000 |
| cluster2986 | 0.51 | 1.0000 |
| cluster4123 | -0.94 | 1.0000 |
| cluster3522 | -0.98 | 1.0000 |
| cluster3959 | 0.48 | 1.0000 |
| cluster2889 | 0.48 | 1.0000 |
| cluster3232 | 0.49 | 1.0000 |
| cluster3019 | 0.48 | 1.0000 |
| cluster3064 | 0.45 | 1.0000 |
| cluster3195 | -0.72 | 1.0000 |
| cluster3051 | 0.48 | 1.0000 |
| cluster3483 | 0.49 | 1.0000 |
| cluster3006 | -0.98 | 1.0000 |
| cluster3234 | -0.77 | 1.0000 |
| cluster3053 | -0.85 | 1.0000 |
| cluster3377 | -0.97 | 1.0000 |
| cluster3961 | 0.47 | 1.0000 |
| cluster3523 | 0.47 | 1.0000 |
| cluster3402 | 0.41 | 1.0000 |
| cluster3621 | 0.48 | 1.0000 |
| cluster3209 | -1.00 | 1.0000 |
| cluster3316 | 0.46 | 1.0000 |
| cluster3246 | 0.40 | 1.0000 |
| cluster2920 | -0.89 | 1.0000 |
| cluster3747 | 0.42 | 1.0000 |
| cluster3960 | -1.02 | 1.0000 |
| cluster1127 | -0.99 | 1.0000 |
| cluster3529 | 0.46 | 1.0000 |
| cluster3007 | 0.45 | 1.0000 |
| cluster3189 | -0.95 | 1.0000 |
| cluster3281 | 0.46 | 1.0000 |
| cluster3686 | 0.46 | 1.0000 |
| cluster3429 | -0.72 | 1.0000 |
| cluster3668 | 0.45 | 1.0000 |
| cluster3298 | -0.72 | 1.0000 |
| cluster2985 | 0.39 | 1.0000 |
| cluster2938 | -0.64 | 1.0000 |
| cluster3317 | -0.86 | 1.0000 |
| cluster3637 | 0.37 | 1.0000 |
| cluster3759 | -0.71 | 1.0000 |

| | | |
|---|---|---|
| cluster2976 | 0.42 | 1.0000 |
| cluster3437 | -0.72 | 1.0000 |
| cluster3777 | -0.79 | 1.0000 |
| cluster3921 | 0.44 | 1.0000 |
| cluster3659 | -1.04 | 1.0000 |
| cluster3253 | 0.43 | 1.0000 |
| cluster3612 | 0.42 | 1.0000 |
| cluster3546 | -1.18 | 1.0000 |
| cluster2981 | 0.42 | 1.0000 |
| cluster3344 | -0.74 | 1.0000 |
| cluster3280 | 0.37 | 1.0000 |
| cluster3375 | -0.72 | 1.0000 |
| cluster3477 | -0.73 | 1.0000 |
| cluster3743 | 0.40 | 1.0000 |
| cluster2970 | 0.37 | 1.0000 |
| cluster3240 | 0.36 | 1.0000 |
| cluster3514 | -0.73 | 1.0000 |
| cluster3112 | 0.40 | 1.0000 |
| cluster3074 | -0.95 | 1.0000 |
| cluster3784 | 0.35 | 1.0000 |
| cluster3616 | -0.84 | 1.0000 |
| cluster3228 | 0.39 | 1.0000 |
| cluster3490 | -0.84 | 1.0000 |
| cluster3671 | 0.40 | 1.0000 |
| cluster3741 | 0.35 | 1.0000 |
| cluster3271 | -0.60 | 1.0000 |
| cluster4042 | -0.93 | 1.0000 |
| cluster3502 | 0.39 | 1.0000 |
| cluster3164 | -0.90 | 1.0000 |
| cluster3014 | -0.95 | 1.0000 |
| cluster3353 | 0.34 | 1.0000 |
| cluster3077 | -0.69 | 1.0000 |
| cluster3149 | 0.32 | 1.0000 |
| cluster3079 | -0.63 | 1.0000 |
| cluster3555 | 0.39 | 1.0000 |
| cluster2945 | -0.98 | 1.0000 |
| cluster2983 | 0.37 | 1.0000 |
| cluster3339 | -0.61 | 1.0000 |
| cluster3227 | -0.80 | 1.0000 |
| cluster3152 | 0.37 | 1.0000 |
| cluster3388 | -0.86 | 1.0000 |
| cluster3583 | 0.36 | 1.0000 |
| cluster2905 | -0.62 | 1.0000 |
| cluster3967 | -0.70 | 1.0000 |

| | | |
|---|---|---|
| cluster3047 | -0.77 | 1.0000 |
| cluster3548 | 0.31 | 1.0000 |
| cluster2952 | 0.36 | 1.0000 |
| cluster3125 | 0.32 | 1.0000 |
| cluster3354 | 0.33 | 1.0000 |
| cluster3850 | -1.07 | 1.0000 |
| cluster3600 | 0.33 | 1.0000 |
| cluster2871 | 0.31 | 1.0000 |
| cluster3770 | 0.37 | 1.0000 |
| cluster3197 | -0.65 | 1.0000 |
| cluster3479 | -0.82 | 1.0000 |
| cluster2858 | -0.57 | 1.0000 |
| cluster4086 | 0.35 | 1.0000 |
| cluster3492 | 0.34 | 1.0000 |
| cluster2906 | -0.63 | 1.0000 |
| cluster3283 | 0.30 | 1.0000 |
| cluster3733 | 0.33 | 1.0000 |
| cluster3927 | 0.34 | 1.0000 |
| cluster4040 | 0.34 | 1.0000 |
| cluster3329 | 0.35 | 1.0000 |
| cluster3648 | 0.30 | 1.0000 |
| cluster3204 | -0.61 | 1.0000 |
| cluster3696 | -0.68 | 1.0000 |
| cluster3235 | 0.34 | 1.0000 |
| cluster3132 | 0.29 | 1.0000 |
| cluster3188 | -0.66 | 1.0000 |
| cluster3527 | 0.33 | 1.0000 |
| cluster3636 | -0.58 | 1.0000 |
| cluster3243 | -0.60 | 1.0000 |
| cluster3808 | 0.31 | 1.0000 |
| cluster3495 | 0.32 | 1.0000 |
| cluster3963 | -0.86 | 1.0000 |
| cluster3277 | 0.31 | 1.0000 |
| cluster3838 | 0.31 | 1.0000 |
| cluster2898 | 0.29 | 1.0000 |
| cluster3931 | -0.82 | 1.0000 |
| cluster3333 | 0.31 | 1.0000 |
| cluster2873 | -0.57 | 1.0000 |
| cluster3338 | 0.30 | 1.0000 |
| cluster4076 | -0.81 | 1.0000 |
| cluster3506 | 0.30 | 1.0000 |
| cluster3535 | -0.77 | 1.0000 |
| cluster3857 | 0.30 | 1.0000 |
| cluster3107 | -0.57 | 1.0000 |

| | | |
|---|---|---|
| cluster3819 | 0.31 | 1.0000 |
| cluster2931 | -0.77 | 1.0000 |
| cluster3335 | 0.27 | 1.0000 |
| cluster3709 | -0.57 | 1.0000 |
| cluster3073 | -0.61 | 1.0000 |
| cluster3238 | 0.26 | 1.0000 |
| cluster3790 | 0.28 | 1.0000 |
| cluster3362 | -0.57 | 1.0000 |
| cluster3760 | 0.26 | 1.0000 |
| cluster3433 | -0.60 | 1.0000 |
| cluster3792 | -0.66 | 1.0000 |
| cluster3145 | 0.26 | 1.0000 |
| cluster3070 | 0.25 | 1.0000 |
| cluster3627 | -0.65 | 1.0000 |
| cluster4127 | -0.98 | 1.0000 |
| cluster2860 | -0.58 | 1.0000 |
| cluster3163 | -0.82 | 1.0000 |
| cluster3459 | -0.77 | 1.0000 |
| cluster3894 | -0.74 | 1.0000 |
| cluster3592 | 0.27 | 1.0000 |
| cluster3417 | -0.57 | 1.0000 |
| cluster4124 | -0.92 | 1.0000 |
| cluster3958 | -0.90 | 1.0000 |
| cluster3372 | 0.24 | 1.0000 |
| cluster2880 | -0.82 | 1.0000 |
| cluster3386 | 0.25 | 1.0000 |
| cluster3470 | 0.24 | 1.0000 |
| cluster3013 | 0.23 | 1.0000 |
| cluster3116 | 0.24 | 1.0000 |
| cluster3258 | 0.23 | 1.0000 |
| cluster3727 | -0.71 | 1.0000 |
| cluster2935 | -0.53 | 1.0000 |
| cluster3798 | 0.24 | 1.0000 |
| cluster3123 | -0.50 | 1.0000 |
| cluster2955 | -0.53 | 1.0000 |
| cluster3102 | -0.57 | 1.0000 |
| cluster3499 | 0.24 | 1.0000 |
| cluster3758 | -0.59 | 1.0000 |
| cluster3420 | 0.24 | 1.0000 |
| cluster2932 | -0.54 | 1.0000 |
| cluster2879 | -0.53 | 1.0000 |
| cluster2949 | 0.21 | 1.0000 |
| cluster3599 | -0.77 | 1.0000 |
| cluster3276 | 0.22 | 1.0000 |

| | | |
|---|---|---|
| cluster3058 | 0.21 | 1.0000 |
| cluster3713 | 0.23 | 1.0000 |
| cluster3358 | -0.53 | 1.0000 |
| cluster4059 | 0.23 | 1.0000 |
| cluster3973 | -0.89 | 1.0000 |
| cluster3923 | -0.64 | 1.0000 |
| cluster3825 | -0.71 | 1.0000 |
| cluster4026 | -0.68 | 1.0000 |
| cluster3380 | -0.58 | 1.0000 |
| cluster3500 | -0.88 | 1.0000 |
| cluster3826 | -0.64 | 1.0000 |
| cluster3147 | 0.21 | 1.0000 |
| cluster3763 | -0.62 | 1.0000 |
| cluster3886 | 0.21 | 1.0000 |
| cluster3847 | 0.21 | 1.0000 |
| cluster3059 | 0.21 | 1.0000 |
| cluster3072 | -0.77 | 1.0000 |
| cluster3309 | 0.20 | 1.0000 |
| cluster2891 | -0.72 | 1.0000 |
| cluster3278 | 0.19 | 1.0000 |
| cluster2861 | 0.20 | 1.0000 |
| cluster3303 | -0.66 | 1.0000 |
| cluster3472 | -0.48 | 1.0000 |
| cluster3718 | -0.62 | 1.0000 |
| cluster3579 | -0.48 | 1.0000 |
| cluster2991 | 0.20 | 1.0000 |
| cluster2863 | 0.18 | 1.0000 |
| cluster3137 | 0.18 | 1.0000 |
| cluster3144 | -0.72 | 1.0000 |
| cluster3343 | 0.18 | 1.0000 |
| cluster3046 | -0.51 | 1.0000 |
| cluster2962 | -0.51 | 1.0000 |
| cluster4007 | 0.20 | 1.0000 |
| cluster3605 | 0.19 | 1.0000 |
| cluster2934 | -0.48 | 1.0000 |
| cluster3541 | 0.18 | 1.0000 |
| cluster3409 | 0.18 | 1.0000 |
| cluster2944 | -0.60 | 1.0000 |
| cluster2979 | 0.18 | 1.0000 |
| cluster3782 | 0.18 | 1.0000 |
| cluster3157 | -0.45 | 1.0000 |
| cluster4012 | 0.17 | 1.0000 |
| cluster2885 | -0.66 | 1.0000 |
| cluster3378 | 0.18 | 1.0000 |

| | | |
|---|---|---|
| cluster3524 | -0.51 | 1.0000 |
| cluster3302 | 0.17 | 1.0000 |
| cluster4103 | 0.17 | 1.0000 |
| cluster3765 | -0.51 | 1.0000 |
| cluster3221 | 0.17 | 1.0000 |
| cluster3115 | 0.17 | 1.0000 |
| cluster3516 | -0.76 | 1.0000 |
| cluster3689 | -0.79 | 1.0000 |
| cluster3426 | -0.48 | 1.0000 |
| cluster2940 | -0.66 | 1.0000 |
| cluster3067 | 0.16 | 1.0000 |
| cluster3264 | -0.46 | 1.0000 |
| cluster4101 | 0.14 | 1.0000 |
| cluster3062 | -0.44 | 1.0000 |
| cluster3658 | -0.46 | 1.0000 |
| cluster3447 | 0.16 | 1.0000 |
| cluster3440 | -0.44 | 1.0000 |
| cluster3399 | 0.15 | 1.0000 |
| cluster3582 | -0.43 | 1.0000 |
| cluster3858 | 0.15 | 1.0000 |
| cluster3350 | -0.62 | 1.0000 |
| cluster3036 | 0.14 | 1.0000 |
| cluster4092 | 0.16 | 1.0000 |
| cluster3526 | 0.15 | 1.0000 |
| cluster3428 | -0.45 | 1.0000 |
| cluster3456 | -0.42 | 1.0000 |
| cluster3779 | 0.15 | 1.0000 |
| cluster4105 | 0.14 | 1.0000 |
| cluster3113 | 0.14 | 1.0000 |
| cluster2968 | 0.14 | 1.0000 |
| cluster3896 | 0.14 | 1.0000 |
| cluster2919 | 0.14 | 1.0000 |
| cluster3199 | -0.48 | 1.0000 |
| cluster3573 | -0.62 | 1.0000 |
| cluster3452 | 0.13 | 1.0000 |
| cluster3225 | 0.13 | 1.0000 |
| cluster3018 | 0.12 | 1.0000 |
| cluster3044 | 0.12 | 1.0000 |
| cluster3769 | 0.13 | 1.0000 |
| cluster3848 | -0.66 | 1.0000 |
| cluster3251 | 0.11 | 1.0000 |
| cluster3632 | -0.39 | 1.0000 |
| cluster3445 | -0.39 | 1.0000 |
| cluster3739 | -0.38 | 1.0000 |

| | | |
|---|---|---|
| cluster3682 | -0.57 | 1.0000 |
| cluster3286 | -0.36 | 1.0000 |
| cluster3394 | 0.11 | 1.0000 |
| cluster3167 | 0.10 | 1.0000 |
| cluster3827 | 0.11 | 1.0000 |
| cluster3554 | 0.10 | 1.0000 |
| cluster2950 | -0.38 | 1.0000 |
| cluster3701 | -0.39 | 1.0000 |
| cluster3031 | -0.38 | 1.0000 |
| cluster2926 | 0.11 | 1.0000 |
| cluster3269 | -0.56 | 1.0000 |
| cluster3166 | -0.36 | 1.0000 |
| cluster3888 | -0.53 | 1.0000 |
| cluster3989 | 0.12 | 1.0000 |
| cluster2998 | -0.56 | 1.0000 |
| cluster2857 | 0.10 | 1.0000 |
| cluster3585 | -0.39 | 1.0000 |
| cluster3756 | -0.53 | 1.0000 |
| cluster3043 | -0.52 | 1.0000 |
| cluster3396 | -0.40 | 1.0000 |
| cluster3805 | 0.07 | 1.0000 |
| cluster3846 | -0.35 | 1.0000 |
| cluster2907 | -0.57 | 1.0000 |
| cluster3037 | -0.43 | 1.0000 |
| cluster3101 | 0.09 | 1.0000 |
| cluster3341 | 0.07 | 1.0000 |
| cluster3133 | 0.08 | 1.0000 |
| cluster3255 | -0.59 | 1.0000 |
| cluster3623 | 0.06 | 1.0000 |
| cluster3697 | -0.48 | 1.0000 |
| cluster4048 | -0.64 | 1.0000 |
| cluster4032 | 0.09 | 1.0000 |
| cluster3511 | 0.05 | 1.0000 |
| cluster3861 | 0.07 | 1.0000 |
| cluster3818 | 0.08 | 1.0000 |
| cluster3208 | -0.36 | 1.0000 |
| cluster3091 | 0.08 | 1.0000 |
| cluster3722 | 0.06 | 1.0000 |
| cluster3033 | -0.37 | 1.0000 |
| cluster3684 | -0.49 | 1.0000 |
| cluster3104 | 0.07 | 1.0000 |
| cluster3034 | 0.05 | 1.0000 |
| cluster3935 | 0.06 | 1.0000 |
| cluster3667 | -0.38 | 1.0000 |

| | | |
|---|---|---|
| cluster3184 | 0.06 | 1.0000 |
| cluster3391 | -0.31 | 1.0000 |
| cluster3376 | -0.32 | 1.0000 |
| cluster2910 | -0.52 | 1.0000 |
| cluster3089 | 0.06 | 1.0000 |
| cluster3023 | 0.06 | 1.0000 |
| cluster3882 | 0.03 | 1.0000 |
| cluster3165 | -0.54 | 1.0000 |
| cluster3565 | -0.48 | 1.0000 |
| cluster3364 | -0.51 | 1.0000 |
| cluster3159 | 0.03 | 1.0000 |
| cluster3593 | -0.47 | 1.0000 |
| cluster3859 | -0.54 | 1.0000 |
| cluster3150 | -0.48 | 1.0000 |
| cluster2997 | -0.46 | 1.0000 |
| cluster4020 | -0.32 | 1.0000 |
| cluster3608 | -0.34 | 1.0000 |
| cluster3001 | 0.05 | 1.0000 |
| cluster3224 | 0.04 | 1.0000 |
| cluster3173 | 0.02 | 1.0000 |
| cluster3728 | -0.35 | 1.0000 |
| cluster3256 | 0.06 | 1.0000 |
| cluster3308 | 0.05 | 1.0000 |
| cluster3588 | -0.44 | 1.0000 |
| cluster2956 | 0.04 | 1.0000 |
| cluster3025 | -0.29 | 1.0000 |
| cluster3999 | -0.44 | 1.0000 |
| cluster3948 | 0.00 | 1.0000 |
| cluster3602 | -0.31 | 1.0000 |
| cluster3214 | -0.32 | 1.0000 |
| cluster2996 | 0.03 | 1.0000 |
| cluster2896 | -0.01 | 1.0000 |
| cluster4102 | 0.05 | 1.0000 |
| cluster3993 | 0.01 | 1.0000 |
| cluster3755 | -0.56 | 1.0000 |
| cluster3692 | -0.42 | 1.0000 |
| cluster3186 | -0.04 | 1.0000 |
| cluster2917 | -0.03 | 1.0000 |
| cluster2939 | -0.33 | 1.0000 |
| cluster3615 | 0.01 | 1.0000 |
| cluster3665 | -0.44 | 1.0000 |
| cluster3328 | 0.02 | 1.0000 |
| cluster3304 | 0.01 | 1.0000 |
| cluster2933 | 0.00 | 1.0000 |

| | | |
|---|---|---|
| cluster3310 | -0.27 | 1.0000 |
| cluster3032 | 0.01 | 1.0000 |
| cluster3307 | -0.04 | 1.0000 |
| cluster3027 | 0.01 | 1.0000 |
| cluster2878 | -0.30 | 1.0000 |
| cluster2921 | -0.01 | 1.0000 |
| cluster2929 | -0.03 | 1.0000 |
| cluster3352 | -0.42 | 1.0000 |
| cluster2961 | 0.00 | 1.0000 |
| cluster3517 | 0.01 | 1.0000 |
| cluster3577 | 0.00 | 1.0000 |
| cluster2993 | -0.01 | 1.0000 |
| cluster3239 | -0.25 | 1.0000 |
| cluster3088 | -0.33 | 1.0000 |
| cluster3181 | -0.28 | 1.0000 |
| cluster2872 | -0.34 | 1.0000 |
| cluster3148 | -0.01 | 1.0000 |
| cluster3955 | -0.43 | 1.0000 |
| cluster4108 | -0.37 | 1.0000 |
| cluster2882 | 0.00 | 1.0000 |
| cluster3351 | -0.03 | 1.0000 |
| cluster4006 | -0.41 | 1.0000 |
| cluster3586 | -0.25 | 1.0000 |
| cluster3716 | -0.37 | 1.0000 |
| cluster3521 | -0.38 | 1.0000 |
| cluster3061 | -0.27 | 1.0000 |
| cluster2999 | -0.08 | 1.0000 |
| cluster3669 | -0.08 | 1.0000 |
| cluster3478 | -0.27 | 1.0000 |
| cluster3313 | -0.35 | 1.0000 |
| cluster3791 | -0.40 | 1.0000 |
| cluster2859 | -0.25 | 1.0000 |
| cluster3361 | -0.09 | 1.0000 |
| cluster3984 | -0.47 | 1.0000 |
| cluster3729 | -0.39 | 1.0000 |
| cluster3873 | -0.02 | 1.0000 |
| cluster3406 | -0.03 | 1.0000 |
| cluster4022 | -0.14 | 1.0000 |
| cluster3407 | -0.25 | 1.0000 |
| cluster2870 | -0.05 | 1.0000 |
| cluster4053 | -0.36 | 1.0000 |
| cluster3748 | -0.25 | 1.0000 |
| cluster3797 | -0.37 | 1.0000 |
| cluster2971 | -0.22 | 1.0000 |

| | | |
|---|---|---|
| cluster2875 | -0.26 | 1.0000 |
| cluster3717 | -0.24 | 1.0000 |
| cluster3041 | -0.27 | 1.0000 |
| cluster3216 | -0.34 | 1.0000 |
| cluster2965 | -0.13 | 1.0000 |
| cluster3900 | -0.18 | 1.0000 |
| cluster3530 | -0.31 | 1.0000 |
| cluster3022 | -0.33 | 1.0000 |
| cluster3928 | -0.34 | 1.0000 |
| cluster3539 | -0.39 | 1.0000 |
| cluster3547 | -0.05 | 1.0000 |
| cluster2911 | -0.24 | 1.0000 |
| cluster3134 | -0.21 | 1.0000 |
| cluster3907 | -0.35 | 1.0000 |
| cluster3124 | -0.23 | 1.0000 |
| cluster3321 | -0.29 | 1.0000 |
| cluster3384 | -0.27 | 1.0000 |
| cluster2958 | -0.23 | 1.0000 |
| cluster3559 | -0.31 | 1.0000 |
| cluster3248 | -0.33 | 1.0000 |
| cluster3912 | -0.06 | 1.0000 |
| cluster4066 | -0.28 | 1.0000 |
| cluster3096 | -0.09 | 1.0000 |
| cluster2923 | -0.07 | 1.0000 |
| cluster3331 | -0.30 | 1.0000 |
| cluster3296 | -0.22 | 1.0000 |
| cluster2897 | -0.23 | 1.0000 |
| cluster3092 | -0.12 | 1.0000 |
| cluster3176 | -0.26 | 1.0000 |
| cluster4004 | -0.18 | 1.0000 |
| cluster3922 | -0.33 | 1.0000 |
| cluster3348 | -0.21 | 1.0000 |
| cluster3141 | -0.17 | 1.0000 |
| cluster2916 | -0.21 | 1.0000 |
| cluster4075 | -0.33 | 1.0000 |
| cluster3323 | -0.23 | 1.0000 |
| cluster3505 | -0.29 | 1.0000 |
| cluster3736 | -0.29 | 1.0000 |
| cluster3816 | -0.28 | 1.0000 |
| cluster3566 | -0.27 | 1.0000 |
| cluster2883 | -0.11 | 1.0000 |
| cluster3890 | -0.22 | 1.0000 |
| cluster3021 | -0.15 | 1.0000 |
| cluster3054 | -0.13 | 1.0000 |

| | | |
|---|---|---|
| cluster3762 | -0.21 | 1.0000 |
| cluster3170 | -0.16 | 1.0000 |
| cluster3946 | -0.24 | 1.0000 |
| cluster3749 | -0.24 | 1.0000 |
| cluster3289 | -0.15 | 1.0000 |
| cluster3854 | -0.24 | 1.0000 |
| cluster3049 | -0.16 | 1.0000 |
| cluster3347 | -0.15 | 1.0000 |
| cluster4041 | -0.26 | 1.0000 |
| cluster3670 | -0.18 | 1.0000 |
| cluster3020 | -0.31 | 1.0000 |
| cluster2946 | -0.17 | 1.0000 |
| cluster3215 | -0.14 | 1.0000 |
| cluster4078 | -0.18 | 1.0000 |
| cluster3268 | -0.25 | 1.0000 |
| cluster3294 | -0.13 | 1.0000 |
| cluster2884 | -0.17 | 1.0000 |
| cluster3222 | -0.12 | 1.0000 |
| cluster3664 | -0.19 | 1.0000 |
| cluster3926 | -0.33 | 1.0000 |
| cluster3005 | -0.15 | 1.0000 |
| cluster3359 | -0.24 | 1.0000 |
| cluster3403 | -0.25 | 1.0000 |
| cluster3802 | -0.22 | 1.0000 |
| cluster3764 | -0.26 | 1.0000 |
| cluster3098 | -0.22 | 1.0000 |
| cluster3823 | -0.15 | 1.0000 |
| cluster3883 | -0.30 | 1.0000 |
| cluster4063 | -0.24 | 1.0000 |
| cluster4106 | -0.23 | 1.0000 |
| cluster3314 | -0.22 | 1.0000 |
| cluster3312 | -0.21 | 1.0000 |
| cluster3663 | -0.21 | 1.0000 |
| cluster3795 | -0.20 | 1.0000 |
| cluster4043 | -0.19 | 1.0000 |
| cluster3320 | -0.19 | 1.0000 |
| cluster3820 | -0.18 | 1.0000 |
| cluster3498 | -0.17 | 1.0000 |
| cluster3835 | -0.17 | 1.0000 |
| cluster3596 | -0.16 | 1.0000 |
| cluster3705 | -0.15 | 1.0000 |
| cluster3561 | -0.15 | 1.0000 |
| cluster3620 | -0.14 | 1.0000 |
| cluster3589 | -0.14 | 1.0000 |

| | | |
|---|---|---|
| cluster3156 | -0.14 | 1.0000 |
| cluster3557 | -0.13 | 1.0000 |
| cluster3785 | -0.13 | 1.0000 |
| cluster3806 | -0.13 | 1.0000 |
| cluster3262 | -0.13 | 1.0000 |
| cluster3562 | -0.12 | 1.0000 |
| cluster3071 | -0.12 | 1.0000 |
| cluster3874 | -0.12 | 1.0000 |
| cluster3241 | -0.12 | 1.0000 |
| cluster3266 | -0.11 | 1.0000 |
| cluster3887 | -0.11 | 1.0000 |
| cluster3655 | -0.11 | 1.0000 |
| cluster3480 | -0.11 | 1.0000 |
| cluster3707 | -0.10 | 1.0000 |
| cluster3324 | -0.10 | 1.0000 |
| cluster3868 | -0.09 | 1.0000 |
| cluster3412 | -0.08 | 1.0000 |
| cluster3138 | -0.08 | 1.0000 |
| cluster3423 | -0.08 | 1.0000 |
| cluster3695 | -0.08 | 1.0000 |
| cluster3988 | -0.07 | 1.0000 |
| cluster3297 | -0.07 | 1.0000 |
| cluster3877 | -0.07 | 1.0000 |
| cluster3002 | -0.07 | 1.0000 |
| cluster3048 | -0.07 | 1.0000 |
| cluster4005 | -0.06 | 1.0000 |
| cluster3992 | -0.06 | 1.0000 |
| cluster3884 | -0.06 | 1.0000 |
| cluster3903 | -0.06 | 1.0000 |
| cluster3128 | -0.05 | 1.0000 |
| cluster3851 | -0.05 | 1.0000 |
| cluster2989 | -0.05 | 1.0000 |
| cluster3183 | -0.03 | 1.0000 |
| cluster4003 | -0.03 | 1.0000 |
| cluster3390 | -0.03 | 1.0000 |
| cluster3597 | -0.02 | 1.0000 |
| cluster3794 | -0.02 | 1.0000 |
| cluster3744 | -0.02 | 1.0000 |
| cluster3174 | -0.01 | 1.0000 |
| cluster3949 | -0.01 | 1.0000 |
| cluster4035 | -0.01 | 1.0000 |
| cluster3139 | -0.01 | 1.0000 |
| cluster3284 | 0.00 | 1.0000 |
| cluster3893 | 0.00 | 1.0000 |

| | | |
|---|---|---|
| cluster3392 | 0.00 | 1.0000 |

**Supplementary Table 15.** DESeq2 results with eXpress counts - Traumatic vs furuncular

| ID | log2FoldChange | Adjusted p value |
|---|---|---|
| cluster3272 | 10.09 | 5.89E-15 |
| cluster3752 | 5.77 | 1.88E-08 |
| cluster3640 | 6.67 | 1.35E-05 |
| cluster4034 | 5.76 | 1.35E-05 |
| cluster3233 | -4.82 | 0.0001 |
| cluster3546 | 5.06 | 0.0001 |
| cluster4101 | -5.26 | 0.0001 |
| cluster3770 | 5.84 | 0.0002 |
| cluster3006 | -4.18 | 0.0007 |
| cluster3312 | 4.87 | 0.0007 |
| cluster3393 | 4.13 | 0.0007 |
| cluster3738 | -4.64 | 0.0012 |
| cluster3755 | -5.04 | 0.0024 |
| cluster3392 | 4.03 | 0.0033 |
| cluster3012 | 3.56 | 0.0036 |
| cluster3030 | -3.77 | 0.0036 |
| cluster3786 | -4.38 | 0.0036 |
| cluster4076 | -4.26 | 0.0036 |
| cluster3303 | -4.06 | 0.0051 |
| cluster3516 | 4.12 | 0.0053 |
| cluster3416 | 4.27 | 0.0054 |
| cluster3631 | -3.73 | 0.0065 |
| cluster3749 | 4.18 | 0.0065 |
| cluster3955 | 4.25 | 0.0065 |
| cluster3178 | -3.75 | 0.0068 |
| cluster3060 | 3.54 | 0.0111 |
| cluster3599 | 3.46 | 0.0111 |
| cluster3105 | 3.19 | 0.0112 |
| cluster3175 | 3.29 | 0.0112 |
| cluster3763 | -3.48 | 0.0112 |
| cluster3191 | 3.41 | 0.0159 |
| cluster2948 | -3.46 | 0.0162 |
| cluster3313 | 2.67 | 0.0168 |
| cluster3672 | 3.00 | 0.0186 |
| cluster2980 | 3.49 | 0.0240 |
| cluster3126 | -3.30 | 0.0240 |
| cluster3741 | -2.70 | 0.0240 |
| cluster3852 | -3.27 | 0.0240 |
| cluster3329 | 3.85 | 0.0274 |
| cluster3905 | 4.49 | 0.0294 |
| cluster4046 | 3.17 | 0.0294 |
| cluster3659 | 3.10 | 0.0318 |

| | | |
|---|---|---|
| cluster3802 | -3.48 | 0.0341 |
| cluster3323 | -3.09 | 0.0394 |
| cluster3674 | 4.09 | 0.0394 |
| cluster3514 | -2.83 | 0.0430 |
| cluster3367 | -3.35 | 0.0440 |
| cluster3840 | -2.98 | 0.0458 |
| cluster3157 | -2.59 | 0.0475 |
| cluster3890 | -3.21 | 0.0475 |
| cluster2888 | 3.55 | 0.0483 |
| cluster3994 | 3.23 | 0.0489 |
| cluster4098 | -3.27 | 0.0498 |
| cluster4052 | -3.00 | 0.0513 |
| cluster3597 | -2.59 | 0.0518 |
| cluster3593 | 2.81 | 0.0529 |
| cluster3948 | 2.33 | 0.0529 |
| cluster3386 | 2.80 | 0.0543 |
| cluster3565 | 2.75 | 0.0550 |
| cluster3186 | 2.79 | 0.0567 |
| cluster3764 | 3.33 | 0.0576 |
| cluster2960 | -2.80 | 0.0591 |
| cluster3296 | 2.61 | 0.0591 |
| cluster3638 | 2.57 | 0.0591 |
| cluster3818 | -3.05 | 0.0591 |
| cluster3522 | 2.76 | 0.0623 |
| cluster4030 | 2.60 | 0.0687 |
| cluster3129 | 3.35 | 0.0752 |
| cluster3185 | 2.58 | 0.0752 |
| cluster3551 | -2.57 | 0.0752 |
| cluster3371 | -3.84 | 0.0761 |
| cluster3757 | -3.33 | 0.0794 |
| cluster2860 | -2.47 | 0.0822 |
| cluster3197 | -2.40 | 0.0825 |
| cluster3831 | 2.37 | 0.0857 |
| cluster3017 | -2.24 | 0.0890 |
| cluster3695 | 2.23 | 0.0924 |
| cluster3693 | 3.27 | 0.0930 |
| cluster3904 | 2.46 | 0.0940 |
| cluster3977 | 2.90 | 0.0942 |
| cluster3500 | 2.53 | 0.0942 |
| cluster2876 | 2.47 | 0.0962 |
| cluster3714 | -2.55 | 0.0972 |
| cluster3615 | -2.74 | 0.0988 |
| cluster3000 | 2.17 | 0.0996 |
| cluster3309 | -2.59 | 0.0996 |

| | | |
|---|---|---|
| cluster3138 | -2.50 | 0.0997 |
| cluster3385 | 2.68 | 0.0997 |
| cluster3426 | -2.40 | 0.0997 |
| cluster3459 | 3.00 | 0.0997 |
| cluster3511 | 2.62 | 0.0997 |
| cluster3538 | -2.36 | 0.0997 |
| cluster3652 | -2.54 | 0.0997 |
| cluster3689 | 3.44 | 0.0997 |
| cluster4055 | 2.23 | 0.0997 |
| cluster4103 | -2.98 | 0.0997 |
| cluster2984 | -2.43 | 0.1026 |
| cluster4059 | 3.28 | 0.1026 |
| cluster3397 | 2.61 | 0.1045 |
| cluster3102 | -2.54 | 0.1092 |
| cluster2899 | -2.36 | 0.1130 |
| cluster3337 | -2.28 | 0.1130 |
| cluster3400 | -2.02 | 0.1130 |
| cluster4058 | 2.11 | 0.1130 |
| cluster3564 | -2.83 | 0.1151 |
| cluster3721 | -2.56 | 0.1151 |
| cluster3317 | -2.38 | 0.1169 |
| cluster3341 | 2.37 | 0.1169 |
| cluster3410 | -2.46 | 0.1169 |
| cluster4069 | 2.28 | 0.1169 |
| cluster4078 | -2.51 | 0.1169 |
| cluster2962 | -2.42 | 0.1184 |
| cluster3063 | -2.34 | 0.1184 |
| cluster3478 | -2.09 | 0.1184 |
| cluster3709 | -1.95 | 0.1184 |
| cluster3730 | -2.05 | 0.1184 |
| cluster3931 | -2.72 | 0.1184 |
| cluster4001 | 2.49 | 0.1184 |
| cluster3691 | -2.23 | 0.1193 |
| cluster2976 | 1.99 | 0.1199 |
| cluster3569 | 2.24 | 0.1237 |
| cluster3876 | -2.13 | 0.1237 |
| cluster3200 | -2.19 | 0.1251 |
| cluster3182 | 2.07 | 0.1265 |
| cluster3061 | -2.18 | 0.1284 |
| cluster3817 | -2.55 | 0.1284 |
| cluster2946 | -2.16 | 0.1308 |
| cluster3922 | 3.10 | 0.1322 |
| cluster2933 | -2.34 | 0.1343 |
| cluster2993 | -2.41 | 0.1353 |

| | | |
|---|---|---|
| cluster3705 | -2.51 | 0.1357 |
| cluster2978 | -2.17 | 0.1383 |
| cluster3760 | -2.14 | 0.1386 |
| cluster3177 | 2.43 | 0.1395 |
| cluster3766 | 2.00 | 0.1395 |
| cluster3130 | 1.92 | 0.1395 |
| cluster2989 | -2.32 | 0.1421 |
| cluster3053 | -2.48 | 0.1421 |
| cluster3497 | -1.95 | 0.1421 |
| cluster3850 | 2.08 | 0.1421 |
| cluster3051 | 2.52 | 0.1456 |
| cluster3299 | 2.33 | 0.1495 |
| cluster3032 | -2.04 | 0.1498 |
| cluster3137 | -2.53 | 0.1498 |
| cluster3513 | -2.11 | 0.1498 |
| cluster3732 | 1.83 | 0.1498 |
| cluster3342 | 2.07 | 0.1503 |
| cluster3346 | -2.19 | 0.1503 |
| cluster3788 | -1.75 | 0.1503 |
| cluster3858 | -2.42 | 0.1503 |
| cluster3898 | -2.54 | 0.1503 |
| cluster2943 | -2.16 | 0.1506 |
| cluster2861 | -2.24 | 0.1513 |
| cluster3158 | 2.16 | 0.1513 |
| cluster3101 | -2.02 | 0.1516 |
| cluster2921 | -1.96 | 0.1551 |
| cluster2853 | -2.13 | 0.1577 |
| cluster3240 | -2.19 | 0.1577 |
| cluster3408 | -1.90 | 0.1608 |
| cluster3225 | 2.57 | 0.1615 |
| cluster3226 | 2.24 | 0.1615 |
| cluster3254 | 2.44 | 0.1615 |
| cluster3304 | -1.89 | 0.1615 |
| cluster3452 | 2.29 | 0.1615 |
| cluster3023 | -1.98 | 0.1623 |
| cluster3494 | -2.13 | 0.1623 |
| cluster3909 | 1.84 | 0.1623 |
| cluster4011 | 1.86 | 0.1623 |
| cluster3731 | -2.24 | 0.1637 |
| cluster3297 | -2.45 | 0.1670 |
| cluster3015 | -2.12 | 0.1676 |
| cluster2947 | -1.80 | 0.1770 |
| cluster3237 | -2.47 | 0.1770 |
| cluster3572 | 2.66 | 0.1770 |

| | | |
|---|---|---|
| cluster3531 | 1.98 | 0.1787 |
| cluster2881 | -2.06 | 0.1809 |
| cluster2893 | 2.12 | 0.1809 |
| cluster3109 | 1.87 | 0.1809 |
| cluster3694 | 1.66 | 0.1809 |
| cluster3486 | -1.89 | 0.1867 |
| cluster2902 | -2.33 | 0.1870 |
| cluster2961 | -2.07 | 0.1890 |
| cluster3778 | -1.94 | 0.1890 |
| cluster2873 | -1.91 | 0.1902 |
| cluster2967 | -2.18 | 0.1902 |
| cluster3144 | 2.71 | 0.1902 |
| cluster3389 | 1.80 | 0.1913 |
| cluster3391 | -1.84 | 0.1913 |
| cluster3449 | -1.77 | 0.1915 |
| cluster3181 | -1.90 | 0.1936 |
| cluster2900 | -1.93 | 0.1941 |
| cluster3188 | -2.15 | 0.1941 |
| cluster4071 | 2.62 | 0.1941 |
| cluster3229 | -1.93 | 0.1943 |
| cluster3119 | -1.99 | 0.1944 |
| cluster4050 | 2.23 | 0.1981 |
| cluster2965 | -1.65 | 0.1992 |
| cluster3469 | -2.03 | 0.1992 |
| cluster3830 | -1.55 | 0.1992 |
| cluster3485 | 2.72 | 0.2003 |
| cluster4038 | 2.34 | 0.2003 |
| cluster3118 | -1.72 | 0.2008 |
| cluster2905 | -1.77 | 0.2009 |
| cluster3318 | -2.03 | 0.2012 |
| cluster3468 | -1.80 | 0.2050 |
| cluster3518 | 2.26 | 0.2070 |
| cluster2904 | -1.70 | 0.2173 |
| cluster3897 | -2.18 | 0.2173 |
| cluster4049 | 2.13 | 0.2173 |
| cluster2999 | 1.99 | 0.2220 |
| cluster3239 | -1.54 | 0.2220 |
| cluster3258 | -1.49 | 0.2220 |
| cluster3374 | 2.13 | 0.2231 |
| cluster3014 | 2.03 | 0.2254 |
| cluster3417 | -2.07 | 0.2259 |
| cluster4080 | 1.75 | 0.2267 |
| cluster2963 | -1.90 | 0.2312 |
| cluster3147 | -1.49 | 0.2312 |

| | | |
|---|---|---|
| cluster3634 | -1.79 | 0.2312 |
| cluster2906 | -1.89 | 0.2324 |
| cluster3650 | -1.73 | 0.2329 |
| cluster3156 | 1.84 | 0.2334 |
| cluster3582 | -1.62 | 0.2364 |
| cluster3834 | 2.13 | 0.2364 |
| cluster3487 | 1.71 | 0.2368 |
| cluster3595 | -1.66 | 0.2368 |
| cluster3270 | 1.94 | 0.2374 |
| cluster3575 | -1.97 | 0.2374 |
| cluster3480 | -1.80 | 0.2381 |
| cluster4079 | -1.86 | 0.2407 |
| cluster3082 | 2.49 | 0.2427 |
| cluster3246 | -1.80 | 0.2427 |
| cluster3319 | 1.79 | 0.2444 |
| cluster3402 | -1.83 | 0.2469 |
| cluster3377 | 1.70 | 0.2482 |
| cluster3574 | 2.37 | 0.2482 |
| cluster3338 | 2.01 | 0.2497 |
| cluster3501 | 2.08 | 0.2497 |
| cluster4056 | 1.74 | 0.2497 |
| cluster3027 | -1.72 | 0.2504 |
| cluster3811 | -1.60 | 0.2556 |
| cluster2911 | 1.45 | 0.2664 |
| cluster3915 | 2.34 | 0.2676 |
| cluster3142 | -1.75 | 0.2683 |
| cluster3925 | 2.60 | 0.2683 |
| cluster3984 | 2.18 | 0.2683 |
| cluster3066 | -1.73 | 0.2706 |
| cluster3454 | 1.41 | 0.2709 |
| cluster3992 | -1.92 | 0.2709 |
| cluster2924 | -1.49 | 0.2717 |
| cluster3987 | -1.82 | 0.2717 |
| cluster3141 | -1.59 | 0.2804 |
| cluster3756 | 1.77 | 0.2804 |
| cluster3619 | -1.63 | 0.2812 |
| cluster2977 | -1.46 | 0.2817 |
| cluster3024 | -1.55 | 0.2817 |
| cluster3387 | 1.50 | 0.2817 |
| cluster3704 | -1.82 | 0.2829 |
| cluster3894 | 1.54 | 0.2829 |
| cluster3971 | 1.82 | 0.2834 |
| cluster3414 | -1.61 | 0.2839 |
| cluster3358 | -1.79 | 0.2871 |

| | | |
|---|---|---|
| cluster3462 | 1.45 | 0.2899 |
| cluster4014 | 1.82 | 0.2943 |
| cluster3465 | -1.65 | 0.2944 |
| cluster3049 | -1.65 | 0.2950 |
| cluster3089 | -1.62 | 0.2957 |
| cluster3311 | -1.69 | 0.2974 |
| cluster3681 | 1.68 | 0.2983 |
| cluster3345 | 1.61 | 0.3021 |
| cluster3399 | -1.89 | 0.3021 |
| cluster3892 | 1.79 | 0.3059 |
| cluster3171 | -1.67 | 0.3065 |
| cluster3306 | 1.64 | 0.3074 |
| cluster3568 | 1.78 | 0.3103 |
| cluster3700 | 1.68 | 0.3103 |
| cluster4057 | 2.03 | 0.3128 |
| cluster4118 | 4.76 | 0.3128 |
| cluster3071 | -1.57 | 0.3247 |
| cluster3669 | 1.41 | 0.3247 |
| cluster3656 | -1.66 | 0.3248 |
| cluster3792 | 1.36 | 0.3257 |
| cluster3450 | 2.43 | 0.3309 |
| cluster2914 | 1.49 | 0.3311 |
| cluster3295 | -1.32 | 0.3311 |
| cluster3327 | -1.64 | 0.3311 |
| cluster3335 | -1.76 | 0.3311 |
| cluster3348 | -1.68 | 0.3311 |
| cluster3639 | 2.37 | 0.3311 |
| cluster3653 | -1.65 | 0.3311 |
| cluster3774 | -1.50 | 0.3311 |
| cluster3112 | 1.92 | 0.3364 |
| cluster2887 | 1.41 | 0.3376 |
| cluster3045 | -1.55 | 0.3386 |
| cluster3322 | 1.86 | 0.3386 |
| cluster3963 | 2.12 | 0.3386 |
| cluster2908 | -1.39 | 0.3400 |
| cluster3244 | -1.51 | 0.3400 |
| cluster2869 | -1.38 | 0.3425 |
| cluster2878 | -1.56 | 0.3500 |
| cluster3243 | -1.51 | 0.3500 |
| cluster3042 | 1.62 | 0.3539 |
| cluster3187 | 1.92 | 0.3539 |
| cluster3836 | 1.45 | 0.3539 |
| cluster3165 | -1.76 | 0.3597 |
| cluster3456 | -1.38 | 0.3665 |

| | | |
|---|---|---|
| cluster3515 | -1.81 | 0.3680 |
| cluster3873 | -1.39 | 0.3682 |
| cluster4042 | 1.79 | 0.3682 |
| cluster3613 | -1.50 | 0.3688 |
| cluster3097 | -1.46 | 0.3696 |
| cluster3253 | 1.63 | 0.3696 |
| cluster3471 | 1.45 | 0.3696 |
| cluster3369 | -1.33 | 0.3707 |
| cluster3670 | 1.66 | 0.3720 |
| cluster3602 | -1.45 | 0.3742 |
| cluster3928 | 1.36 | 0.3742 |
| cluster3510 | 1.47 | 0.3758 |
| cluster3425 | 1.40 | 0.3758 |
| cluster3443 | 1.30 | 0.3760 |
| cluster2923 | -1.22 | 0.3777 |
| cluster3993 | 1.30 | 0.3777 |
| cluster2916 | -1.64 | 0.3804 |
| cluster3132 | -1.40 | 0.3804 |
| cluster3262 | -1.25 | 0.3804 |
| cluster3289 | -1.43 | 0.3804 |
| cluster3444 | -1.15 | 0.3804 |
| cluster3472 | -1.43 | 0.3804 |
| cluster3651 | 1.92 | 0.3804 |
| cluster3964 | 2.06 | 0.3804 |
| cluster4031 | -1.58 | 0.3804 |
| cluster3375 | 1.22 | 0.3813 |
| cluster3541 | -1.45 | 0.3813 |
| cluster3908 | 1.27 | 0.3813 |
| cluster3448 | 1.91 | 0.3839 |
| cluster3698 | -1.25 | 0.3839 |
| cluster3973 | 1.38 | 0.3839 |
| cluster3088 | -1.47 | 0.3885 |
| cluster3787 | 1.61 | 0.3885 |
| cluster3805 | -1.46 | 0.3885 |
| cluster3814 | -1.33 | 0.3885 |
| cluster3013 | -1.34 | 0.3888 |
| cluster3827 | 1.68 | 0.3888 |
| cluster3570 | 1.89 | 0.3970 |
| cluster3361 | 1.37 | 0.4014 |
| cluster2930 | -1.17 | 0.4017 |
| cluster3439 | 1.22 | 0.4017 |
| cluster2863 | -1.15 | 0.4027 |
| cluster2882 | -1.23 | 0.4034 |
| cluster2950 | -1.30 | 0.4042 |

| | | |
|---|---|---|
| cluster3461 | 1.88 | 0.4042 |
| cluster3566 | 1.25 | 0.4042 |
| cluster3577 | -1.38 | 0.4042 |
| cluster3594 | 1.45 | 0.4042 |
| cluster3883 | -1.97 | 0.4042 |
| cluster3360 | 1.43 | 0.4051 |
| cluster3630 | -1.15 | 0.4073 |
| cluster4022 | 1.21 | 0.4073 |
| cluster3547 | -1.30 | 0.4131 |
| cluster3933 | 1.30 | 0.4131 |
| cluster2964 | -1.30 | 0.4132 |
| cluster3268 | 1.18 | 0.4132 |
| cluster3269 | 1.84 | 0.4132 |
| cluster3979 | -1.34 | 0.4132 |
| cluster3107 | -1.14 | 0.4138 |
| cluster3623 | 1.35 | 0.4138 |
| cluster3490 | 1.57 | 0.4149 |
| cluster3019 | -1.21 | 0.4166 |
| cluster2937 | 1.29 | 0.4203 |
| cluster3166 | -1.28 | 0.4203 |
| cluster3230 | -1.49 | 0.4203 |
| cluster3255 | 1.55 | 0.4203 |
| cluster3286 | -1.10 | 0.4203 |
| cluster3644 | 1.31 | 0.4203 |
| cluster2926 | -1.40 | 0.4204 |
| cluster3077 | -1.45 | 0.4204 |
| cluster3549 | -1.32 | 0.4208 |
| cluster3573 | 1.40 | 0.4208 |
| cluster3079 | -1.37 | 0.4264 |
| cluster3856 | 1.26 | 0.4264 |
| cluster3025 | -1.08 | 0.4274 |
| cluster3548 | -1.21 | 0.4360 |
| cluster3558 | 1.38 | 0.4368 |
| cluster3048 | 1.12 | 0.4371 |
| cluster3748 | -1.37 | 0.4374 |
| cluster3750 | 1.88 | 0.4374 |
| cluster3940 | -1.31 | 0.4374 |
| cluster3503 | 1.15 | 0.4384 |
| cluster3404 | -1.17 | 0.4402 |
| cluster3168 | -1.25 | 0.4502 |
| cluster3192 | -1.09 | 0.4525 |
| cluster2954 | -1.25 | 0.4541 |
| cluster2935 | -1.28 | 0.4575 |
| cluster3362 | -1.26 | 0.4631 |

| | | |
|---|---|---|
| cluster3422 | -1.67 | 0.4631 |
| cluster3476 | 1.61 | 0.4631 |
| cluster3106 | -1.12 | 0.4647 |
| cluster3149 | -1.15 | 0.4647 |
| cluster3250 | -1.15 | 0.4647 |
| cluster3740 | 1.52 | 0.4647 |
| cluster3783 | 1.67 | 0.4647 |
| cluster3113 | -1.07 | 0.4654 |
| cluster3806 | 1.50 | 0.4698 |
| cluster3466 | 1.27 | 0.4708 |
| cluster2875 | -1.33 | 0.4715 |
| cluster3110 | -1.03 | 0.4718 |
| cluster3411 | 1.03 | 0.4718 |
| cluster3641 | -1.22 | 0.4718 |
| cluster3773 | 1.05 | 0.4718 |
| cluster3914 | 1.79 | 0.4718 |
| cluster3926 | 1.56 | 0.4718 |
| cluster4085 | 1.05 | 0.4718 |
| cluster3021 | -1.12 | 0.4773 |
| cluster3099 | -1.12 | 0.4773 |
| cluster3429 | -1.29 | 0.4800 |
| cluster3115 | -1.03 | 0.4822 |
| cluster3213 | -1.27 | 0.4868 |
| cluster3315 | -1.21 | 0.4868 |
| cluster4039 | -1.38 | 0.4868 |
| cluster3281 | 1.25 | 0.4870 |
| cluster3011 | -1.30 | 0.4889 |
| cluster3170 | -1.21 | 0.4889 |
| cluster3234 | 1.18 | 0.4889 |
| cluster3279 | 1.10 | 0.4889 |
| cluster3340 | -1.12 | 0.4889 |
| cluster3403 | 0.99 | 0.4889 |
| cluster3407 | -1.23 | 0.4889 |
| cluster3532 | -1.45 | 0.4889 |
| cluster3671 | 1.17 | 0.4889 |
| cluster3697 | -1.26 | 0.4889 |
| cluster3005 | -1.00 | 0.4960 |
| cluster3530 | -1.04 | 0.4972 |
| cluster3956 | -1.12 | 0.4976 |
| cluster3504 | -1.33 | 0.5002 |
| cluster3026 | 1.34 | 0.5016 |
| cluster3083 | -1.04 | 0.5016 |
| cluster3121 | -1.22 | 0.5016 |
| cluster3215 | -0.94 | 0.5016 |

| | | |
|---|---|---|
| cluster3320 | 1.28 | 0.5016 |
| cluster3885 | 1.08 | 0.5016 |
| cluster2982 | -1.09 | 0.5032 |
| cluster3125 | 0.99 | 0.5032 |
| cluster3198 | 1.19 | 0.5032 |
| cluster3539 | 1.63 | 0.5032 |
| cluster3626 | -1.14 | 0.5032 |
| cluster3635 | -0.95 | 0.5032 |
| cluster3810 | -1.18 | 0.5032 |
| cluster4072 | -1.04 | 0.5032 |
| cluster3455 | -1.31 | 0.5066 |
| cluster3034 | 1.09 | 0.5129 |
| cluster3477 | -1.30 | 0.5139 |
| cluster3273 | -1.05 | 0.5147 |
| cluster2919 | -1.12 | 0.5162 |
| cluster3002 | -0.99 | 0.5162 |
| cluster3172 | -1.12 | 0.5162 |
| cluster3220 | 1.07 | 0.5162 |
| cluster3495 | -1.06 | 0.5162 |
| cluster3600 | -0.95 | 0.5162 |
| cluster3739 | -0.91 | 0.5162 |
| cluster3780 | 1.59 | 0.5162 |
| cluster4097 | 1.03 | 0.5162 |
| cluster3035 | -0.92 | 0.5185 |
| cluster3031 | -1.13 | 0.5205 |
| cluster2994 | -1.01 | 0.5214 |
| cluster3536 | -1.04 | 0.5214 |
| cluster3753 | 0.92 | 0.5214 |
| cluster3372 | -1.02 | 0.5273 |
| cluster3467 | 0.98 | 0.5276 |
| cluster3666 | 0.98 | 0.5341 |
| cluster3278 | -0.94 | 0.5378 |
| cluster3359 | 1.27 | 0.5378 |
| cluster3458 | -1.00 | 0.5378 |
| cluster3534 | 0.96 | 0.5378 |
| cluster3557 | 1.11 | 0.5378 |
| cluster3803 | 1.52 | 0.5378 |
| cluster3807 | -1.02 | 0.5378 |
| cluster3632 | -1.12 | 0.5380 |
| cluster3009 | 0.98 | 0.5382 |
| cluster3433 | -1.07 | 0.5382 |
| cluster3489 | 1.10 | 0.5382 |
| cluster3526 | -0.91 | 0.5382 |
| cluster3579 | -1.09 | 0.5382 |

| | | |
|---|---|---|
| cluster3350 | 1.14 | 0.5389 |
| cluster2866 | -1.24 | 0.5395 |
| cluster3661 | -0.95 | 0.5395 |
| cluster2912 | -0.88 | 0.5410 |
| cluster2974 | -1.02 | 0.5435 |
| cluster3209 | 1.01 | 0.5439 |
| cluster3069 | -0.86 | 0.5484 |
| cluster3256 | -0.94 | 0.5484 |
| cluster2886 | -1.12 | 0.5496 |
| cluster2890 | -0.88 | 0.5496 |
| cluster2953 | -0.90 | 0.5506 |
| cluster3164 | 1.41 | 0.5506 |
| cluster3238 | -0.86 | 0.5506 |
| cluster3302 | 1.04 | 0.5506 |
| cluster3563 | 1.03 | 0.5506 |
| cluster3851 | 1.29 | 0.5506 |
| cluster3143 | -1.04 | 0.5596 |
| cluster3446 | 0.92 | 0.5596 |
| cluster3567 | -1.53 | 0.5596 |
| cluster3381 | -1.03 | 0.5608 |
| cluster3208 | -0.93 | 0.5629 |
| cluster2940 | 0.97 | 0.5665 |
| cluster3588 | -1.17 | 0.5665 |
| cluster3657 | 0.87 | 0.5665 |
| cluster3772 | 0.95 | 0.5665 |
| cluster3864 | 0.90 | 0.5665 |
| cluster3067 | -0.97 | 0.5698 |
| cluster2925 | -0.96 | 0.5711 |
| cluster3223 | -1.00 | 0.5711 |
| cluster3265 | -0.92 | 0.5711 |
| cluster3703 | -0.91 | 0.5711 |
| cluster3016 | 1.30 | 0.5722 |
| cluster3290 | 1.03 | 0.5722 |
| cluster3525 | 0.90 | 0.5722 |
| cluster3581 | 0.96 | 0.5722 |
| cluster3982 | 0.89 | 0.5722 |
| cluster3211 | 1.29 | 0.5735 |
| cluster3257 | -0.90 | 0.5735 |
| cluster3606 | 1.29 | 0.5735 |
| cluster3737 | -0.86 | 0.5735 |
| cluster3004 | -0.99 | 0.5753 |
| cluster3969 | -0.80 | 0.5753 |
| cluster3267 | -0.94 | 0.5754 |
| cluster3291 | -1.01 | 0.5754 |

| | | |
|---|---|---|
| cluster3301 | -1.10 | 0.5754 |
| cluster3321 | -0.88 | 0.5754 |
| cluster3356 | -1.04 | 0.5754 |
| cluster3401 | -0.79 | 0.5754 |
| cluster3405 | -0.92 | 0.5754 |
| cluster3473 | -1.11 | 0.5754 |
| cluster3743 | 0.84 | 0.5754 |
| cluster3135 | 0.87 | 0.5763 |
| cluster3217 | -0.98 | 0.5775 |
| cluster3907 | 1.34 | 0.5829 |
| cluster3085 | 1.13 | 0.5855 |
| cluster3212 | 1.09 | 0.5855 |
| cluster3724 | 1.05 | 0.5855 |
| cluster3953 | 0.79 | 0.5855 |
| cluster3493 | -1.17 | 0.5862 |
| cluster2884 | -0.83 | 0.5866 |
| cluster3723 | -0.81 | 0.5866 |
| cluster3759 | -1.03 | 0.5866 |
| cluster3854 | 0.82 | 0.5866 |
| cluster2894 | 0.89 | 0.5901 |
| cluster3396 | -1.04 | 0.5908 |
| cluster3093 | 0.90 | 0.5914 |
| cluster3560 | 1.23 | 0.5914 |
| cluster3646 | 1.31 | 0.5914 |
| cluster3383 | 1.29 | 0.5936 |
| cluster3658 | -0.85 | 0.5981 |
| cluster3146 | -0.78 | 0.5982 |
| cluster3437 | 0.97 | 0.5990 |
| cluster3479 | 1.22 | 0.5990 |
| cluster3555 | 1.27 | 0.5990 |
| cluster3720 | 1.20 | 0.5990 |
| cluster2901 | 1.00 | 0.5998 |
| cluster2909 | -0.77 | 0.5998 |
| cluster3199 | -0.77 | 0.5998 |
| cluster3353 | -0.76 | 0.5998 |
| cluster3972 | 0.88 | 0.6011 |
| cluster3046 | -0.89 | 0.6018 |
| cluster3022 | 1.02 | 0.6023 |
| cluster3236 | 0.94 | 0.6023 |
| cluster3440 | -0.74 | 0.6023 |
| cluster2939 | 0.84 | 0.6036 |
| cluster3901 | 0.84 | 0.6050 |
| cluster2870 | -0.85 | 0.6067 |
| cluster3349 | -0.97 | 0.6087 |

| | | |
|---|---|---|
| cluster3419 | -0.73 | 0.6141 |
| cluster3282 | 0.83 | 0.6146 |
| cluster3793 | 0.81 | 0.6157 |
| cluster3779 | -0.98 | 0.6200 |
| cluster2951 | 0.99 | 0.6207 |
| cluster2992 | -0.79 | 0.6207 |
| cluster3251 | 0.79 | 0.6207 |
| cluster3326 | -0.75 | 0.6207 |
| cluster3398 | 0.71 | 0.6207 |
| cluster3616 | -1.02 | 0.6207 |
| cluster2949 | -0.80 | 0.6215 |
| cluster3196 | -0.75 | 0.6219 |
| cluster3202 | -0.83 | 0.6219 |
| cluster3276 | -0.84 | 0.6219 |
| cluster3496 | -0.92 | 0.6219 |
| cluster3528 | -0.76 | 0.6219 |
| cluster3601 | 0.96 | 0.6219 |
| cluster3679 | -0.94 | 0.6219 |
| cluster2854 | -0.86 | 0.6224 |
| cluster3975 | 0.75 | 0.6224 |
| cluster2958 | -0.70 | 0.6226 |
| cluster3090 | -1.18 | 0.6227 |
| cluster3636 | 0.71 | 0.6227 |
| cluster3056 | -0.76 | 0.6236 |
| cluster3040 | -0.83 | 0.6236 |
| cluster3084 | -0.72 | 0.6263 |
| cluster3891 | 0.99 | 0.6263 |
| cluster3941 | -0.74 | 0.6263 |
| cluster3959 | 1.15 | 0.6263 |
| cluster3241 | -0.74 | 0.6301 |
| cluster3896 | 1.11 | 0.6301 |
| cluster2942 | -0.77 | 0.6321 |
| cluster3070 | -0.89 | 0.6321 |
| cluster3280 | -0.76 | 0.6321 |
| cluster3339 | -0.88 | 0.6321 |
| cluster3357 | -0.78 | 0.6321 |
| cluster3039 | -0.79 | 0.6323 |
| cluster3893 | -0.77 | 0.6333 |
| cluster3065 | -0.75 | 0.6359 |
| cluster3307 | 0.84 | 0.6359 |
| cluster3966 | -0.71 | 0.6361 |
| cluster3128 | -0.72 | 0.6380 |
| cluster3745 | -0.99 | 0.6477 |
| cluster3043 | -0.73 | 0.6495 |

| | | |
|---|---|---|
| cluster3332 | -0.80 | 0.6500 |
| cluster3344 | -0.91 | 0.6500 |
| cluster3453 | 0.86 | 0.6500 |
| cluster3734 | 0.90 | 0.6500 |
| cluster3266 | -0.76 | 0.6507 |
| cluster3354 | -0.72 | 0.6507 |
| cluster3305 | 0.88 | 0.6554 |
| cluster3390 | -0.80 | 0.6554 |
| cluster3841 | 0.95 | 0.6554 |
| cluster3064 | -0.91 | 0.6565 |
| cluster3499 | 0.95 | 0.6565 |
| cluster3637 | -0.65 | 0.6565 |
| cluster3942 | 0.73 | 0.6586 |
| cluster3316 | 0.85 | 0.6618 |
| cluster3134 | 0.69 | 0.6632 |
| cluster3062 | 0.69 | 0.6701 |
| cluster4020 | -0.75 | 0.6701 |
| cluster3201 | -0.69 | 0.6727 |
| cluster3821 | 0.76 | 0.6727 |
| cluster3825 | 0.79 | 0.6727 |
| cluster3174 | -0.90 | 0.6753 |
| cluster3665 | 1.03 | 0.6782 |
| cluster3735 | 0.82 | 0.6782 |
| cluster2981 | -0.69 | 0.6830 |
| cluster3683 | 0.66 | 0.6830 |
| cluster3435 | 1.00 | 0.6847 |
| cluster3688 | 0.73 | 0.6851 |
| cluster2862 | -0.72 | 0.6876 |
| cluster3231 | -0.84 | 0.6876 |
| cluster3622 | -0.61 | 0.6876 |
| cluster3347 | -0.66 | 0.6895 |
| cluster3655 | 0.76 | 0.6895 |
| cluster3447 | 0.84 | 0.6903 |
| cluster3204 | -0.77 | 0.6933 |
| cluster3261 | 0.94 | 0.7006 |
| cluster3482 | 0.67 | 0.7006 |
| cluster2991 | 0.66 | 0.7009 |
| cluster3136 | -0.74 | 0.7015 |
| cluster3382 | -0.63 | 0.7017 |
| cluster3769 | 0.65 | 0.7024 |
| cluster2864 | -0.67 | 0.7048 |
| cluster2865 | -0.70 | 0.7048 |
| cluster2871 | 0.63 | 0.7048 |
| cluster2955 | -0.56 | 0.7048 |

| | | |
|---|---|---|
| cluster2990 | 0.58 | 0.7048 |
| cluster3263 | -0.61 | 0.7048 |
| cluster3275 | 0.87 | 0.7048 |
| cluster3668 | 0.59 | 0.7048 |
| cluster3726 | -0.61 | 0.7048 |
| cluster3900 | 1.10 | 0.7048 |
| cluster2929 | -0.59 | 0.7052 |
| cluster3654 | 0.69 | 0.7052 |
| cluster3663 | 0.83 | 0.7052 |
| cluster4041 | 0.82 | 0.7052 |
| cluster2892 | -0.64 | 0.7090 |
| cluster3292 | 0.78 | 0.7090 |
| cluster3336 | -0.80 | 0.7090 |
| cluster3488 | 0.56 | 0.7090 |
| cluster3677 | -0.71 | 0.7090 |
| cluster3706 | 0.59 | 0.7090 |
| cluster3795 | 0.68 | 0.7090 |
| cluster2986 | 0.57 | 0.7098 |
| cluster3131 | 0.62 | 0.7117 |
| cluster3314 | -0.83 | 0.7117 |
| cluster2858 | -0.63 | 0.7133 |
| cluster3554 | 0.70 | 0.7133 |
| cluster2856 | 0.74 | 0.7166 |
| cluster3003 | -0.64 | 0.7175 |
| cluster3524 | -0.57 | 0.7175 |
| cluster2936 | 0.89 | 0.7179 |
| cluster3725 | 0.86 | 0.7179 |
| cluster2898 | -0.76 | 0.7200 |
| cluster3120 | -0.53 | 0.7200 |
| cluster3395 | 0.62 | 0.7200 |
| cluster2966 | 0.70 | 0.7200 |
| cluster3055 | -0.58 | 0.7280 |
| cluster3862 | -0.54 | 0.7280 |
| cluster3919 | 0.51 | 0.7306 |
| cluster2970 | -0.50 | 0.7332 |
| cluster3845 | -0.60 | 0.7332 |
| cluster3585 | 0.61 | 0.7349 |
| cluster3451 | 0.97 | 0.7370 |
| cluster3475 | 0.60 | 0.7371 |
| cluster3823 | -0.70 | 0.7378 |
| cluster2957 | 0.74 | 0.7379 |
| cluster2979 | -0.54 | 0.7379 |
| cluster3038 | 0.70 | 0.7379 |
| cluster3285 | -0.61 | 0.7379 |

| | | |
|---|---|---|
| cluster3649 | 0.53 | 0.7379 |
| cluster3944 | 0.63 | 0.7379 |
| cluster3331 | 0.52 | 0.7419 |
| cluster3148 | -0.50 | 0.7441 |
| cluster3761 | -0.73 | 0.7441 |
| cluster3867 | 0.50 | 0.7442 |
| cluster3140 | -0.52 | 0.7455 |
| cluster3946 | 0.75 | 0.7455 |
| cluster3086 | -0.51 | 0.7463 |
| cluster3729 | 0.58 | 0.7463 |
| cluster3744 | 0.62 | 0.7468 |
| cluster2917 | 0.74 | 0.7486 |
| cluster3133 | 0.57 | 0.7487 |
| cluster3523 | 0.79 | 0.7487 |
| cluster3874 | -0.59 | 0.7517 |
| cluster3470 | 0.50 | 0.7517 |
| cluster3533 | -0.57 | 0.7517 |
| cluster3586 | -0.55 | 0.7517 |
| cluster3910 | 0.80 | 0.7551 |
| cluster3701 | -0.56 | 0.7572 |
| cluster3432 | 0.58 | 0.7692 |
| cluster2877 | -0.46 | 0.7708 |
| cluster2997 | 0.52 | 0.7724 |
| cluster3252 | 0.79 | 0.7749 |
| cluster3127 | 0.46 | 0.7750 |
| cluster2883 | -0.52 | 0.7804 |
| cluster2969 | -0.59 | 0.7804 |
| cluster2998 | 0.45 | 0.7804 |
| cluster3100 | -0.65 | 0.7804 |
| cluster3628 | 0.62 | 0.7804 |
| cluster4025 | -0.51 | 0.7816 |
| cluster3521 | 0.54 | 0.7828 |
| cluster3861 | 0.47 | 0.7876 |
| cluster2938 | -0.48 | 0.7897 |
| cluster3058 | -0.46 | 0.7897 |
| cluster3151 | -0.42 | 0.7897 |
| cluster3587 | -0.76 | 0.7897 |
| cluster3765 | 0.58 | 0.7897 |
| cluster2971 | -0.46 | 0.7907 |
| cluster3608 | 0.48 | 0.7907 |
| cluster3080 | -0.50 | 0.7927 |
| cluster4007 | -0.46 | 0.7927 |
| cluster3227 | -0.52 | 0.7966 |
| cluster3699 | 0.67 | 0.7982 |

| | | |
|---|---|---|
| cluster3562 | 0.57 | 0.8025 |
| cluster3294 | -0.49 | 0.8085 |
| cluster3509 | -0.44 | 0.8085 |
| cluster3961 | 0.69 | 0.8085 |
| cluster3232 | -0.49 | 0.8123 |
| cluster3675 | -0.41 | 0.8123 |
| cluster3747 | -0.48 | 0.8123 |
| cluster3800 | 0.44 | 0.8123 |
| cluster3934 | -0.40 | 0.8123 |
| cluster3284 | -0.44 | 0.8176 |
| cluster3380 | 0.48 | 0.8180 |
| cluster3483 | 0.41 | 0.8180 |
| cluster3754 | -0.44 | 0.8180 |
| cluster3207 | -0.52 | 0.8183 |
| cluster2867 | 0.63 | 0.8190 |
| cluster3037 | 0.43 | 0.8226 |
| cluster3163 | 0.65 | 0.8238 |
| cluster3249 | 0.40 | 0.8238 |
| cluster3988 | -0.41 | 0.8238 |
| cluster3111 | 0.40 | 0.8382 |
| cluster3491 | 0.44 | 0.8404 |
| cluster4081 | 0.38 | 0.8447 |
| cluster3001 | -0.42 | 0.8449 |
| cluster3343 | -0.40 | 0.8449 |
| cluster3219 | 0.36 | 0.8473 |
| cluster4063 | 0.60 | 0.8473 |
| cluster3036 | 0.45 | 0.8496 |
| cluster3498 | -0.52 | 0.8496 |
| cluster4105 | -0.50 | 0.8545 |
| cluster2920 | -0.39 | 0.8596 |
| cluster3409 | 0.56 | 0.8620 |
| cluster3308 | -0.40 | 0.8628 |
| cluster4084 | -0.41 | 0.8628 |
| cluster2944 | -0.43 | 0.8645 |
| cluster3247 | -0.45 | 0.8645 |
| cluster3517 | -0.33 | 0.8645 |
| cluster3785 | -0.47 | 0.8646 |
| cluster2952 | 0.46 | 0.8650 |
| cluster3441 | -0.34 | 0.8695 |
| cluster3791 | 0.46 | 0.8695 |
| cluster1127 | -0.45 | 0.8705 |
| cluster3218 | -0.50 | 0.8705 |
| cluster2896 | 0.33 | 0.8772 |
| cluster2985 | -0.42 | 0.8772 |

| | | |
|---|---|---|
| cluster3123 | -0.36 | 0.8772 |
| cluster3298 | -0.37 | 0.8772 |
| cluster3843 | -0.34 | 0.8772 |
| cluster3620 | -0.42 | 0.8782 |
| cluster3797 | -0.44 | 0.8782 |
| cluster3888 | 0.50 | 0.8782 |
| cluster4003 | 0.53 | 0.8782 |
| cluster4096 | 0.36 | 0.8782 |
| cluster3607 | 0.39 | 0.8794 |
| cluster3384 | -0.38 | 0.8798 |
| cluster3010 | 0.35 | 0.8800 |
| cluster2891 | 0.43 | 0.8804 |
| cluster2915 | 0.34 | 0.8804 |
| cluster3167 | 0.50 | 0.8804 |
| cluster3886 | -0.41 | 0.8804 |
| cluster3333 | 0.37 | 0.8814 |
| cluster2880 | -0.36 | 0.8831 |
| cluster2968 | -0.31 | 0.8831 |
| cluster3033 | 0.27 | 0.8831 |
| cluster3075 | 0.37 | 0.8831 |
| cluster3153 | -0.27 | 0.8831 |
| cluster3193 | -0.31 | 0.8831 |
| cluster3222 | -0.30 | 0.8831 |
| cluster3378 | -0.42 | 0.8831 |
| cluster3415 | -0.33 | 0.8831 |
| cluster3427 | -0.41 | 0.8831 |
| cluster3505 | -0.32 | 0.8831 |
| cluster3543 | -0.45 | 0.8831 |
| cluster3642 | 0.41 | 0.8831 |
| cluster3950 | -0.36 | 0.8831 |
| cluster3952 | -0.30 | 0.8831 |
| cluster3978 | 0.30 | 0.8831 |
| cluster3072 | -0.40 | 0.8852 |
| cluster3690 | -0.34 | 0.8860 |
| cluster3028 | 0.31 | 0.8861 |
| cluster3224 | 0.32 | 0.8888 |
| cluster3183 | 0.32 | 0.8900 |
| cluster3092 | -0.29 | 0.8908 |
| cluster3592 | 0.33 | 0.8919 |
| cluster3442 | 0.38 | 0.8931 |
| cluster3612 | 0.42 | 0.8931 |
| cluster3680 | 0.31 | 0.8931 |
| cluster3379 | 0.28 | 0.8957 |
| cluster3728 | -0.32 | 0.8957 |

| | | |
|---|---|---|
| cluster2956 | 0.32 | 0.9022 |
| cluster2928 | 0.28 | 0.9055 |
| cluster3457 | -0.36 | 0.9055 |
| cluster3413 | 0.35 | 0.9113 |
| cluster3050 | -0.25 | 0.9121 |
| cluster3145 | -0.30 | 0.9121 |
| cluster3775 | 0.29 | 0.9121 |
| cluster3104 | 0.22 | 0.9159 |
| cluster3180 | 0.25 | 0.9159 |
| cluster3506 | 0.26 | 0.9159 |
| cluster3552 | -0.26 | 0.9159 |
| cluster2922 | 0.29 | 0.9198 |
| cluster3583 | -0.24 | 0.9198 |
| cluster3875 | 0.25 | 0.9198 |
| cluster3210 | -0.23 | 0.9202 |
| cluster3877 | 0.27 | 0.9202 |
| cluster2983 | 0.24 | 0.9209 |
| cluster3324 | 0.28 | 0.9209 |
| cluster3609 | -0.41 | 0.9209 |
| cluster2857 | 0.23 | 0.9215 |
| cluster2872 | -0.23 | 0.9215 |
| cluster2879 | -0.24 | 0.9215 |
| cluster2895 | -0.26 | 0.9215 |
| cluster2987 | -0.21 | 0.9215 |
| cluster2988 | -0.22 | 0.9215 |
| cluster3124 | 0.20 | 0.9215 |
| cluster3159 | 0.22 | 0.9215 |
| cluster3161 | 0.23 | 0.9215 |
| cluster3184 | 0.24 | 0.9215 |
| cluster3221 | 0.29 | 0.9215 |
| cluster3264 | 0.23 | 0.9215 |
| cluster3310 | -0.20 | 0.9215 |
| cluster3352 | 0.29 | 0.9215 |
| cluster3370 | 0.27 | 0.9215 |
| cluster3428 | -0.25 | 0.9215 |
| cluster3809 | -0.23 | 0.9215 |
| cluster3816 | 0.32 | 0.9215 |
| cluster3846 | 0.25 | 0.9215 |
| cluster3980 | -0.22 | 0.9215 |
| cluster4015 | 0.22 | 0.9215 |
| cluster3081 | 0.24 | 0.9225 |
| cluster3095 | 0.25 | 0.9225 |
| cluster3160 | 0.19 | 0.9225 |
| cluster3859 | -0.22 | 0.9225 |

| | | |
|---|---|---|
| cluster3420 | 0.24 | 0.9227 |
| cluster3283 | -0.23 | 0.9241 |
| cluster2907 | -0.21 | 0.9266 |
| cluster2945 | 0.32 | 0.9266 |
| cluster3116 | -0.18 | 0.9266 |
| cluster3189 | -0.24 | 0.9266 |
| cluster3205 | -0.21 | 0.9266 |
| cluster3214 | -0.24 | 0.9266 |
| cluster3363 | -0.20 | 0.9266 |
| cluster3364 | 0.21 | 0.9266 |
| cluster3376 | -0.20 | 0.9266 |
| cluster3603 | -0.25 | 0.9266 |
| cluster3627 | -0.21 | 0.9266 |
| cluster3633 | 0.20 | 0.9266 |
| cluster3648 | -0.22 | 0.9266 |
| cluster3920 | 0.26 | 0.9266 |
| cluster3921 | 0.32 | 0.9266 |
| cluster3962 | 0.19 | 0.9266 |
| cluster3103 | 0.24 | 0.9272 |
| cluster3751 | 0.21 | 0.9309 |
| cluster2855 | 0.17 | 0.9315 |
| cluster2897 | -0.21 | 0.9317 |
| cluster3544 | -0.16 | 0.9317 |
| cluster3293 | -0.19 | 0.9328 |
| cluster2927 | 0.23 | 0.9381 |
| cluster3194 | 0.19 | 0.9381 |
| cluster3195 | 0.18 | 0.9381 |
| cluster3260 | 0.19 | 0.9381 |
| cluster3330 | -0.18 | 0.9381 |
| cluster3365 | 0.19 | 0.9381 |
| cluster3617 | 0.23 | 0.9381 |
| cluster3667 | -0.18 | 0.9381 |
| cluster3711 | -0.14 | 0.9381 |
| cluster3758 | 0.22 | 0.9381 |
| cluster3768 | 0.23 | 0.9381 |
| cluster3847 | 0.19 | 0.9381 |
| cluster3945 | -0.23 | 0.9381 |
| cluster2918 | -0.16 | 0.9399 |
| cluster2973 | 0.18 | 0.9399 |
| cluster3007 | 0.16 | 0.9399 |
| cluster3029 | 0.14 | 0.9399 |
| cluster3098 | 0.14 | 0.9399 |
| cluster3108 | -0.19 | 0.9399 |
| cluster3216 | 0.17 | 0.9399 |

| | | |
|---|---|---|
| cluster3271 | -0.15 | 0.9399 |
| cluster3368 | 0.14 | 0.9399 |
| cluster3484 | -0.19 | 0.9399 |
| cluster3507 | -0.16 | 0.9399 |
| cluster3832 | 0.17 | 0.9399 |
| cluster3848 | 0.19 | 0.9399 |
| cluster3868 | 0.15 | 0.9399 |
| cluster3903 | 0.16 | 0.9399 |
| cluster2913 | -0.13 | 0.9404 |
| cluster2931 | 0.17 | 0.9404 |
| cluster3502 | -0.21 | 0.9404 |
| cluster3057 | -0.15 | 0.9422 |
| cluster3154 | -0.15 | 0.9422 |
| cluster3054 | -0.12 | 0.9435 |
| cluster2885 | -0.16 | 0.9436 |
| cluster2996 | -0.15 | 0.9436 |
| cluster3242 | -0.15 | 0.9436 |
| cluster3692 | 0.18 | 0.9436 |
| cluster3717 | 0.14 | 0.9436 |
| cluster3917 | 0.17 | 0.9436 |
| cluster2889 | 0.13 | 0.9469 |
| cluster4006 | 0.13 | 0.9469 |
| cluster3068 | 0.11 | 0.9492 |
| cluster3406 | -0.14 | 0.9495 |
| cluster3591 | 0.12 | 0.9509 |
| cluster3808 | 0.17 | 0.9522 |
| cluster3624 | 0.10 | 0.9537 |
| cluster3169 | 0.14 | 0.9540 |
| cluster3424 | 0.15 | 0.9545 |
| cluster4004 | -0.10 | 0.9545 |
| cluster3044 | -0.15 | 0.9550 |
| cluster4029 | -0.16 | 0.9561 |
| cluster3678 | 0.11 | 0.9578 |
| cluster2959 | 0.11 | 0.9593 |
| cluster3394 | 0.11 | 0.9593 |
| cluster4037 | 0.10 | 0.9593 |
| cluster3412 | 0.11 | 0.9615 |
| cluster4019 | -0.13 | 0.9615 |
| cluster3325 | 0.14 | 0.9648 |
| cluster4113 | 0.08 | 0.9648 |
| cluster3091 | -0.08 | 0.9705 |
| cluster3571 | 0.11 | 0.9722 |
| cluster3912 | 0.07 | 0.9722 |
| cluster3625 | 0.08 | 0.9747 |

| | | |
|---|---|---|
| cluster2859 | -0.06 | 0.9757 |
| cluster3020 | 0.08 | 0.9757 |
| cluster3676 | -0.11 | 0.9757 |
| cluster3228 | 0.07 | 0.9786 |
| cluster3529 | 0.08 | 0.9786 |
| cluster3073 | 0.06 | 0.9821 |
| cluster3114 | 0.05 | 0.9821 |
| cluster2932 | 0.05 | 0.9825 |
| cluster2975 | 0.05 | 0.9825 |
| cluster3047 | 0.06 | 0.9825 |
| cluster3078 | 0.04 | 0.9825 |
| cluster3096 | 0.05 | 0.9825 |
| cluster3173 | 0.07 | 0.9825 |
| cluster3179 | -0.06 | 0.9825 |
| cluster3274 | 0.07 | 0.9825 |
| cluster3431 | 0.05 | 0.9825 |
| cluster3685 | -0.06 | 0.9825 |
| cluster3727 | -0.08 | 0.9825 |
| cluster3820 | 0.05 | 0.9825 |
| cluster2874 | -0.04 | 0.9834 |
| cluster2941 | 0.04 | 0.9834 |
| cluster3155 | -0.06 | 0.9834 |
| cluster3351 | 0.04 | 0.9834 |
| cluster3434 | -0.05 | 0.9834 |
| cluster2934 | 0.03 | 0.9855 |
| cluster2972 | -0.04 | 0.9855 |
| cluster3087 | 0.03 | 0.9855 |
| cluster3122 | 0.03 | 0.9855 |
| cluster3259 | 0.03 | 0.9855 |
| cluster3492 | -0.04 | 0.9855 |
| cluster3059 | -0.04 | 0.9868 |
| cluster3696 | -0.03 | 0.9873 |
| cluster3008 | 0.02 | 0.9895 |
| cluster2910 | -0.02 | 0.9902 |
| cluster3018 | 0.02 | 0.9902 |
| cluster3074 | 0.02 | 0.9902 |
| cluster3328 | 0.02 | 0.9920 |
| cluster3445 | 0.02 | 0.9920 |
| cluster3235 | -0.02 | 0.9951 |
| cluster3976 | -0.01 | 0.9955 |
| cluster3203 | 0.01 | 0.9967 |
| cluster3902 | 0.01 | 0.9967 |
| cluster3804 | -0.01 | 0.9970 |
| cluster3094 | 0.01 | 0.9976 |

| | | |
|---|---|---|
| cluster3589 | 0.00 | 0.9984 |

**Supplementary Table 16.** edgeR results with eXpress counts - Traumatic vs furuncular

| ID | log2FoldChange | Adjusted p value |
|---|---|---|
| cluster3272 | 10.21 | 6.85E-07 |
| cluster3233 | -4.75 | 0.0004 |
| cluster3752 | 5.79 | 0.0008 |
| cluster4101 | -5.17 | 0.0008 |
| cluster3006 | -4.09 | 0.0023 |
| cluster3738 | -4.55 | 0.0028 |
| cluster3755 | -4.91 | 0.0082 |
| cluster3030 | -3.69 | 0.0084 |
| cluster3786 | -4.28 | 0.0085 |
| cluster4076 | -4.15 | 0.0122 |
| cluster4034 | 5.73 | 0.0122 |
| cluster3303 | -3.96 | 0.0122 |
| cluster3178 | -3.66 | 0.0124 |
| cluster3546 | 5.07 | 0.0126 |
| cluster3631 | -3.64 | 0.0127 |
| cluster3640 | 6.64 | 0.0142 |
| cluster2948 | -3.36 | 0.0281 |
| cluster3770 | 5.78 | 0.0378 |
| cluster3126 | -3.22 | 0.0378 |
| cluster3393 | 4.12 | 0.0378 |
| cluster3763 | -3.40 | 0.0410 |
| cluster3312 | 4.84 | 0.0504 |
| cluster3852 | -3.19 | 0.0576 |
| cluster3741 | -2.65 | 0.0603 |
| cluster3890 | -3.14 | 0.0779 |
| cluster3392 | 4.01 | 0.0871 |
| cluster3840 | -2.89 | 0.0871 |
| cluster3514 | -2.76 | 0.0871 |
| cluster3802 | -3.38 | 0.0928 |
| cluster3323 | -2.99 | 0.0957 |
| cluster3157 | -2.51 | 0.0976 |
| cluster3012 | 3.57 | 0.1011 |
| cluster3367 | -3.25 | 0.1011 |
| cluster2960 | -2.71 | 0.1021 |
| cluster3516 | 4.10 | 0.1021 |
| cluster3749 | 4.14 | 0.1115 |
| cluster4098 | -3.17 | 0.1162 |
| cluster3818 | -2.97 | 0.1162 |
| cluster3599 | 3.44 | 0.1162 |
| cluster3105 | 3.19 | 0.1162 |
| cluster3175 | 3.28 | 0.1162 |
| cluster4112 | -3.55 | 0.1162 |

| | | |
|---|---|---|
| cluster3844 | 5.02 | 0.1162 |
| cluster3060 | 3.53 | 0.1162 |
| cluster3371 | -3.86 | 0.1162 |
| cluster4052 | -2.91 | 0.1162 |
| cluster3955 | 4.21 | 0.1176 |
| cluster3416 | 4.25 | 0.1176 |
| cluster3191 | 3.50 | 0.1176 |
| cluster3757 | -3.33 | 0.1176 |
| cluster3597 | -2.51 | 0.1176 |
| cluster2860 | -2.38 | 0.1258 |
| cluster3197 | -2.32 | 0.1316 |
| cluster3313 | 2.71 | 0.1383 |
| cluster3551 | -2.48 | 0.1589 |
| cluster2980 | 3.46 | 0.1640 |
| cluster2984 | -2.35 | 0.1640 |
| cluster3538 | -2.29 | 0.1641 |
| cluster3017 | -2.23 | 0.1680 |
| cluster3652 | -2.45 | 0.1688 |
| cluster4046 | 3.25 | 0.1688 |
| cluster3672 | 3.00 | 0.1727 |
| cluster3138 | -2.42 | 0.1727 |
| cluster3659 | 3.08 | 0.1761 |
| cluster4078 | -2.43 | 0.1761 |
| cluster3714 | -2.47 | 0.1761 |
| cluster2899 | -2.28 | 0.1769 |
| cluster3309 | -2.51 | 0.1773 |
| cluster3410 | -2.37 | 0.1791 |
| cluster3426 | -2.31 | 0.1791 |
| cluster4111 | 4.26 | 0.1803 |
| cluster3931 | -2.76 | 0.1817 |
| cluster4103 | -2.89 | 0.1845 |
| cluster3615 | -2.64 | 0.1894 |
| cluster3564 | -2.71 | 0.1915 |
| cluster3102 | -2.44 | 0.1958 |
| cluster3063 | -2.27 | 0.1988 |
| cluster2962 | -2.34 | 0.1988 |
| cluster3876 | -2.06 | 0.1988 |
| cluster3337 | -2.20 | 0.1988 |
| cluster3386 | 2.79 | 0.1988 |
| cluster2993 | -2.31 | 0.1988 |
| cluster3400 | -1.97 | 0.1988 |
| cluster3317 | -2.36 | 0.1988 |
| cluster3709 | -1.92 | 0.1988 |
| cluster3200 | -2.10 | 0.1988 |

| | | |
|---|---|---|
| cluster3329 | 3.83 | 0.1988 |
| cluster3186 | 2.79 | 0.1988 |
| cluster3061 | -2.10 | 0.1988 |
| cluster3994 | 3.20 | 0.1988 |
| cluster3053 | -2.39 | 0.1988 |
| cluster3948 | 2.34 | 0.1988 |
| cluster3137 | -2.55 | 0.1988 |
| cluster2933 | -2.23 | 0.1988 |
| cluster2946 | -2.07 | 0.1988 |
| cluster3730 | -1.97 | 0.1988 |
| cluster2978 | -2.08 | 0.1988 |
| cluster3958 | 4.02 | 0.1988 |
| cluster3346 | -2.12 | 0.2008 |
| cluster3721 | -2.46 | 0.2008 |
| cluster3296 | 2.59 | 0.2011 |
| cluster3478 | -2.03 | 0.2011 |
| cluster3905 | 4.46 | 0.2011 |
| cluster3817 | -2.45 | 0.2011 |
| cluster3185 | 2.60 | 0.2054 |
| cluster3760 | -2.06 | 0.2055 |
| cluster3660 | 3.40 | 0.2055 |
| cluster3863 | -2.42 | 0.2088 |
| cluster2861 | -2.16 | 0.2088 |
| cluster3593 | 2.80 | 0.2088 |
| cluster2943 | -2.07 | 0.2108 |
| cluster4030 | 2.58 | 0.2108 |
| cluster3858 | -2.34 | 0.2108 |
| cluster3674 | 4.07 | 0.2108 |
| cluster3638 | 2.59 | 0.2108 |
| cluster2888 | 3.50 | 0.2108 |
| cluster3513 | -2.02 | 0.2108 |
| cluster3497 | -1.88 | 0.2108 |
| cluster3904 | 2.49 | 0.2108 |
| cluster3764 | 3.29 | 0.2108 |
| cluster3565 | 2.74 | 0.2108 |
| cluster2876 | 2.55 | 0.2108 |
| cluster2853 | -2.04 | 0.2141 |
| cluster3887 | 3.31 | 0.2141 |
| cluster3691 | -2.17 | 0.2156 |
| cluster2921 | -1.94 | 0.2156 |
| cluster3101 | -1.94 | 0.2156 |
| cluster2989 | -2.22 | 0.2156 |
| cluster3297 | -2.38 | 0.2156 |
| cluster3705 | -2.42 | 0.2156 |

| | | |
|---|---|---|
| cluster3494 | -2.05 | 0.2156 |
| cluster3032 | -1.95 | 0.2156 |
| cluster4012 | -2.32 | 0.2156 |
| cluster3237 | -2.47 | 0.2156 |
| cluster3522 | 2.75 | 0.2156 |
| cluster2902 | -2.35 | 0.2165 |
| cluster3500 | 2.55 | 0.2165 |
| cluster3561 | -2.28 | 0.2290 |
| cluster3898 | -2.43 | 0.2326 |
| cluster3304 | -1.80 | 0.2347 |
| cluster3240 | -2.09 | 0.2347 |
| cluster3023 | -1.90 | 0.2350 |
| cluster2881 | -1.97 | 0.2360 |
| cluster3015 | -2.11 | 0.2360 |
| cluster3977 | 2.88 | 0.2393 |
| cluster3129 | 3.32 | 0.2393 |
| cluster3788 | -1.70 | 0.2393 |
| cluster2961 | -1.98 | 0.2393 |
| cluster3511 | 2.59 | 0.2394 |
| cluster3731 | -2.14 | 0.2458 |
| cluster3778 | -1.86 | 0.2458 |
| cluster3000 | 2.17 | 0.2479 |
| cluster3385 | 2.64 | 0.2479 |
| cluster4013 | 5.30 | 0.2501 |
| cluster4069 | 2.31 | 0.2504 |
| cluster2873 | -1.83 | 0.2516 |
| cluster3397 | 2.58 | 0.2611 |
| cluster2967 | -2.08 | 0.2611 |
| cluster3469 | -1.95 | 0.2630 |
| cluster3695 | 2.28 | 0.2699 |
| cluster3188 | -2.05 | 0.2708 |
| cluster2900 | -1.84 | 0.2708 |
| cluster3119 | -1.91 | 0.2744 |
| cluster4001 | 2.47 | 0.2805 |
| cluster3831 | 2.38 | 0.2812 |
| cluster3181 | -1.81 | 0.2820 |
| cluster2976 | 2.06 | 0.2820 |
| cluster3182 | 2.10 | 0.2820 |
| cluster4055 | 2.26 | 0.2820 |
| cluster3229 | -1.84 | 0.2820 |
| cluster3341 | 2.39 | 0.2820 |
| cluster3689 | 3.57 | 0.2820 |
| cluster3391 | -1.76 | 0.2820 |
| cluster2905 | -1.69 | 0.2826 |

| | | |
|---|---|---|
| cluster3486 | -1.81 | 0.2826 |
| cluster3318 | -1.95 | 0.2826 |
| cluster3459 | 2.99 | 0.2831 |
| cluster3449 | -1.70 | 0.2835 |
| cluster3693 | 3.24 | 0.2835 |
| cluster2965 | -1.63 | 0.2847 |
| cluster4118 | 4.67 | 0.2863 |
| cluster3408 | -1.86 | 0.2869 |
| cluster2963 | -1.81 | 0.2869 |
| cluster3645 | 3.89 | 0.2869 |
| cluster4059 | 3.24 | 0.2869 |
| cluster3177 | 2.45 | 0.2869 |
| cluster3650 | -1.74 | 0.2869 |
| cluster3575 | -1.97 | 0.2869 |
| cluster2947 | -1.74 | 0.2871 |
| cluster4043 | 2.91 | 0.2881 |
| cluster2904 | -1.67 | 0.2881 |
| cluster3766 | 2.01 | 0.2926 |
| cluster3897 | -2.10 | 0.2926 |
| cluster4058 | 2.15 | 0.2926 |
| cluster4095 | -2.30 | 0.2949 |
| cluster2906 | -1.79 | 0.2949 |
| cluster3830 | -1.52 | 0.2949 |
| cluster4128 | 3.58 | 0.2955 |
| cluster3850 | 2.09 | 0.2987 |
| cluster4117 | 4.11 | 0.3093 |
| cluster3634 | -1.72 | 0.3111 |
| cluster3246 | -1.71 | 0.3111 |
| cluster3569 | 2.23 | 0.3180 |
| cluster3480 | -1.81 | 0.3196 |
| cluster3468 | -1.72 | 0.3245 |
| cluster3582 | -1.56 | 0.3259 |
| cluster2893 | 2.19 | 0.3275 |
| cluster3118 | -1.68 | 0.3275 |
| cluster3158 | 2.24 | 0.3275 |
| cluster3130 | 1.93 | 0.3275 |
| cluster3598 | 2.65 | 0.3275 |
| cluster3027 | -1.64 | 0.3275 |
| cluster3239 | -1.48 | 0.3275 |
| cluster3417 | -1.98 | 0.3308 |
| cluster3258 | -1.44 | 0.3308 |
| cluster3147 | -1.46 | 0.3337 |
| cluster3811 | -1.54 | 0.3337 |
| cluster3922 | 3.06 | 0.3368 |

| | | |
|---|---|---|
| cluster3051 | 2.51 | 0.3382 |
| cluster4011 | 1.86 | 0.3472 |
| cluster3704 | -1.73 | 0.3479 |
| cluster3299 | 2.30 | 0.3483 |
| cluster3066 | -1.64 | 0.3494 |
| cluster3402 | -1.75 | 0.3507 |
| cluster3595 | -1.60 | 0.3507 |
| cluster3839 | 2.32 | 0.3507 |
| cluster4079 | -1.78 | 0.3537 |
| cluster3109 | 1.89 | 0.3537 |
| cluster3142 | -1.74 | 0.3537 |
| cluster4035 | -1.95 | 0.3537 |
| cluster4089 | -2.10 | 0.3537 |
| cluster3254 | 2.45 | 0.3537 |
| cluster3226 | 2.22 | 0.3537 |
| cluster3141 | -1.52 | 0.3543 |
| cluster3342 | 2.07 | 0.3543 |
| cluster3992 | -1.83 | 0.3546 |
| cluster3694 | 1.69 | 0.3592 |
| cluster3732 | 1.88 | 0.3655 |
| cluster2924 | -1.47 | 0.3692 |
| cluster3049 | -1.57 | 0.3717 |
| cluster3452 | 2.26 | 0.3728 |
| cluster3715 | 2.34 | 0.3745 |
| cluster3465 | -1.57 | 0.3745 |
| cluster4066 | 2.35 | 0.3745 |
| cluster3089 | -1.54 | 0.3761 |
| cluster4080 | 1.81 | 0.3763 |
| cluster3909 | 1.85 | 0.3832 |
| cluster3531 | 2.00 | 0.3832 |
| cluster3937 | -1.88 | 0.3832 |
| cluster3389 | 1.83 | 0.3863 |
| cluster3358 | -1.69 | 0.3883 |
| cluster3987 | -1.75 | 0.3945 |
| cluster3414 | -1.52 | 0.3966 |
| cluster3746 | -1.77 | 0.4037 |
| cluster3653 | -1.57 | 0.4037 |
| cluster3024 | -1.47 | 0.4062 |
| cluster3399 | -1.78 | 0.4066 |
| cluster3619 | -1.55 | 0.4102 |
| cluster3225 | 2.54 | 0.4137 |
| cluster4104 | 2.48 | 0.4137 |
| cluster3171 | -1.57 | 0.4137 |
| cluster3311 | -1.60 | 0.4175 |

| | | |
|---|---|---|
| cluster2977 | -1.41 | 0.4222 |
| cluster3071 | -1.49 | 0.4222 |
| cluster3518 | 2.24 | 0.4222 |
| cluster2999 | 1.97 | 0.4222 |
| cluster4068 | 2.50 | 0.4242 |
| cluster3610 | 2.31 | 0.4262 |
| cluster4050 | 2.20 | 0.4262 |
| cluster3572 | 2.65 | 0.4262 |
| cluster3487 | 1.72 | 0.4262 |
| cluster3762 | -1.80 | 0.4262 |
| cluster3501 | 2.14 | 0.4262 |
| cluster3144 | 2.67 | 0.4262 |
| cluster3165 | -1.78 | 0.4320 |
| cluster3991 | 2.36 | 0.4341 |
| cluster4049 | 2.10 | 0.4341 |
| cluster3014 | 2.03 | 0.4348 |
| cluster2908 | -1.35 | 0.4439 |
| cluster3842 | -1.78 | 0.4449 |
| cluster3156 | 1.82 | 0.4474 |
| cluster3656 | -1.59 | 0.4474 |
| cluster3515 | -1.82 | 0.4474 |
| cluster2878 | -1.48 | 0.4488 |
| cluster4071 | 2.63 | 0.4508 |
| cluster3243 | -1.41 | 0.4508 |
| cluster3485 | 2.68 | 0.4518 |
| cluster3319 | 1.78 | 0.4519 |
| cluster3327 | -1.54 | 0.4574 |
| cluster3335 | -1.67 | 0.4574 |
| cluster3834 | 2.09 | 0.4574 |
| cluster3270 | 1.93 | 0.4574 |
| cluster3883 | -2.03 | 0.4574 |
| cluster3866 | 2.25 | 0.4574 |
| cluster3348 | -1.61 | 0.4587 |
| cluster3873 | -1.32 | 0.4587 |
| cluster2869 | -1.31 | 0.4587 |
| cluster2911 | 1.50 | 0.4587 |
| cluster4038 | 2.32 | 0.4587 |
| cluster3774 | -1.42 | 0.4587 |
| cluster3892 | 1.89 | 0.4587 |
| cluster3245 | -1.81 | 0.4587 |
| cluster4056 | 1.81 | 0.4587 |
| cluster3387 | 1.52 | 0.4587 |
| cluster3833 | 2.38 | 0.4610 |
| cluster3045 | -1.46 | 0.4610 |

| | | |
|---|---|---|
| cluster3990 | -1.56 | 0.4651 |
| cluster3295 | -1.26 | 0.4651 |
| cluster3454 | 1.45 | 0.4701 |
| cluster3541 | -1.38 | 0.4727 |
| cluster3462 | 1.49 | 0.4727 |
| cluster3374 | 2.11 | 0.4727 |
| cluster3244 | -1.43 | 0.4745 |
| cluster3472 | -1.35 | 0.4754 |
| cluster4031 | -1.49 | 0.4754 |
| cluster3613 | -1.43 | 0.4819 |
| cluster3082 | 2.48 | 0.4825 |
| cluster3794 | 1.99 | 0.4876 |
| cluster3989 | -1.66 | 0.4897 |
| cluster2916 | -1.54 | 0.4897 |
| cluster3889 | 2.28 | 0.4898 |
| cluster3097 | -1.37 | 0.4903 |
| cluster3369 | -1.33 | 0.4903 |
| cluster3088 | -1.45 | 0.4903 |
| cluster3812 | -1.54 | 0.4951 |
| cluster3338 | 2.02 | 0.4951 |
| cluster3574 | 2.33 | 0.4998 |
| cluster4014 | 1.79 | 0.5011 |
| cluster3289 | -1.34 | 0.5011 |
| cluster3805 | -1.46 | 0.5011 |
| cluster3669 | 1.45 | 0.5011 |
| cluster3377 | 1.73 | 0.5011 |
| cluster3577 | -1.31 | 0.5027 |
| cluster3013 | -1.26 | 0.5027 |
| cluster3915 | 2.28 | 0.5027 |
| cluster2923 | -1.17 | 0.5090 |
| cluster3132 | -1.33 | 0.5135 |
| cluster2914 | 1.53 | 0.5137 |
| cluster3520 | -1.91 | 0.5137 |
| cluster4083 | 2.23 | 0.5157 |
| cluster3602 | -1.39 | 0.5157 |
| cluster3547 | -1.23 | 0.5157 |
| cluster3756 | 1.74 | 0.5157 |
| cluster3077 | -1.37 | 0.5201 |
| cluster3388 | 1.88 | 0.5201 |
| cluster3444 | -1.10 | 0.5201 |
| cluster3019 | -1.21 | 0.5201 |
| cluster3670 | 1.74 | 0.5207 |
| cluster3799 | -1.49 | 0.5207 |
| cluster3639 | 2.48 | 0.5217 |

| | | |
|---|---|---|
| cluster3549 | -1.25 | 0.5217 |
| cluster3984 | 2.15 | 0.5217 |
| cluster2964 | -1.21 | 0.5217 |
| cluster2926 | -1.30 | 0.5217 |
| cluster3971 | 1.81 | 0.5217 |
| cluster3456 | -1.31 | 0.5217 |
| cluster3698 | -1.18 | 0.5217 |
| cluster3925 | 2.56 | 0.5217 |
| cluster4048 | 2.58 | 0.5217 |
| cluster2930 | -1.16 | 0.5218 |
| cluster3550 | -1.39 | 0.5236 |
| cluster3166 | -1.20 | 0.5245 |
| cluster3792 | 1.36 | 0.5266 |
| cluster3824 | 1.89 | 0.5284 |
| cluster3630 | -1.10 | 0.5323 |
| cluster3748 | -1.28 | 0.5347 |
| cluster3306 | 1.65 | 0.5354 |
| cluster3814 | -1.25 | 0.5384 |
| cluster3568 | 1.75 | 0.5388 |
| cluster3894 | 1.56 | 0.5388 |
| cluster2950 | -1.22 | 0.5388 |
| cluster3681 | 1.69 | 0.5437 |
| cluster3967 | -1.48 | 0.5490 |
| cluster2887 | 1.41 | 0.5490 |
| cluster3345 | 1.60 | 0.5490 |
| cluster3979 | -1.27 | 0.5490 |
| cluster3262 | -1.21 | 0.5490 |
| cluster2863 | -1.09 | 0.5490 |
| cluster2882 | -1.16 | 0.5490 |
| cluster3230 | -1.38 | 0.5490 |
| cluster3733 | 1.84 | 0.5499 |
| cluster2935 | -1.20 | 0.5505 |
| cluster3924 | 1.79 | 0.5567 |
| cluster3112 | 1.88 | 0.5567 |
| cluster3107 | -1.12 | 0.5567 |
| cluster3362 | -1.18 | 0.5576 |
| cluster3079 | -1.28 | 0.5576 |
| cluster3700 | 1.67 | 0.5576 |
| cluster2954 | -1.17 | 0.5581 |
| cluster3176 | 1.80 | 0.5581 |
| cluster4088 | 2.00 | 0.5581 |
| cluster3360 | 1.50 | 0.5581 |
| cluster3836 | 1.47 | 0.5581 |
| cluster3537 | -1.22 | 0.5581 |

| | | |
|---|---|---|
| cluster3404 | -1.09 | 0.5581 |
| cluster3954 | 2.27 | 0.5581 |
| cluster4057 | 2.01 | 0.5581 |
| cluster3519 | 2.17 | 0.5581 |
| cluster3443 | 1.30 | 0.5581 |
| cluster3322 | 1.85 | 0.5581 |
| cluster3300 | -1.28 | 0.5581 |
| cluster3375 | 1.26 | 0.5581 |
| cluster3192 | -1.05 | 0.5581 |
| cluster3286 | -1.05 | 0.5586 |
| cluster3471 | 1.44 | 0.5588 |
| cluster3429 | -1.22 | 0.5588 |
| cluster3187 | 1.90 | 0.5588 |
| cluster3940 | -1.22 | 0.5588 |
| cluster3510 | 1.45 | 0.5588 |
| cluster3106 | -1.07 | 0.5588 |
| cluster3139 | 2.33 | 0.5588 |
| cluster4042 | 1.76 | 0.5588 |
| cluster3253 | 1.61 | 0.5588 |
| cluster3422 | -1.54 | 0.5639 |
| cluster3963 | 2.08 | 0.5639 |
| cluster2875 | -1.22 | 0.5639 |
| cluster3042 | 1.63 | 0.5639 |
| cluster3450 | 2.40 | 0.5665 |
| cluster3149 | -1.08 | 0.5665 |
| cluster3315 | -1.14 | 0.5665 |
| cluster3168 | -1.17 | 0.5665 |
| cluster3025 | -1.03 | 0.5665 |
| cluster3548 | -1.15 | 0.5665 |
| cluster3425 | 1.44 | 0.5665 |
| cluster3697 | -1.19 | 0.5689 |
| cluster3504 | -1.26 | 0.5689 |
| cluster3553 | 1.63 | 0.5689 |
| cluster3011 | -1.21 | 0.5702 |
| cluster3973 | 1.36 | 0.5702 |
| cluster3113 | -1.05 | 0.5736 |
| cluster3361 | 1.35 | 0.5759 |
| cluster3213 | -1.17 | 0.5759 |
| cluster2937 | 1.32 | 0.5759 |
| cluster3651 | 2.00 | 0.5760 |
| cluster3110 | -1.01 | 0.5760 |
| cluster3995 | 1.71 | 0.5760 |
| cluster3250 | -1.12 | 0.5760 |
| cluster3407 | -1.14 | 0.5763 |

| | | |
|---|---|---|
| cluster3767 | 1.76 | 0.5763 |
| cluster3532 | -1.33 | 0.5796 |
| cluster3277 | 1.77 | 0.5824 |
| cluster3495 | -1.08 | 0.5834 |
| cluster3340 | -1.07 | 0.5834 |
| cluster3644 | 1.37 | 0.5834 |
| cluster4107 | 1.84 | 0.5834 |
| cluster3121 | -1.14 | 0.5834 |
| cluster3021 | -1.04 | 0.5834 |
| cluster3115 | -1.01 | 0.5834 |
| cluster3641 | -1.13 | 0.5834 |
| cluster3928 | 1.41 | 0.5834 |
| cluster3827 | 1.75 | 0.5845 |
| cluster3787 | 1.59 | 0.5845 |
| cluster3005 | -0.99 | 0.5845 |
| cluster4053 | 1.59 | 0.5845 |
| cluster2982 | -1.06 | 0.5881 |
| cluster4039 | -1.30 | 0.5881 |
| cluster3466 | 1.35 | 0.5885 |
| cluster3206 | 1.70 | 0.5885 |
| cluster4072 | -1.01 | 0.5911 |
| cluster2866 | -1.24 | 0.5924 |
| cluster3964 | 2.08 | 0.5943 |
| cluster3268 | 1.19 | 0.5943 |
| cluster4022 | 1.21 | 0.5949 |
| cluster3993 | 1.33 | 0.5984 |
| cluster4122 | 2.24 | 0.5984 |
| cluster3461 | 1.86 | 0.5984 |
| cluster3570 | 1.88 | 0.5984 |
| cluster3623 | 1.34 | 0.6034 |
| cluster3170 | -1.13 | 0.6047 |
| cluster3856 | 1.25 | 0.6047 |
| cluster3777 | 1.44 | 0.6047 |
| cluster3099 | -1.09 | 0.6061 |
| cluster3573 | 1.37 | 0.6061 |
| cluster3448 | 1.91 | 0.6061 |
| cluster3439 | 1.24 | 0.6061 |
| cluster3255 | 1.52 | 0.6061 |
| cluster3716 | 1.77 | 0.6061 |
| cluster4045 | 1.89 | 0.6061 |
| cluster4087 | 1.50 | 0.6061 |
| cluster3594 | 1.45 | 0.6061 |
| cluster3773 | 1.12 | 0.6061 |
| cluster3567 | -1.54 | 0.6061 |

| | | |
|---|---|---|
| cluster3530 | -1.02 | 0.6061 |
| cluster3600 | -0.94 | 0.6061 |
| cluster3234 | 1.25 | 0.6061 |
| cluster3908 | 1.31 | 0.6099 |
| cluster3269 | 1.82 | 0.6122 |
| cluster3626 | -1.07 | 0.6137 |
| cluster3455 | -1.20 | 0.6148 |
| cluster3172 | -1.02 | 0.6156 |
| cluster3031 | -1.04 | 0.6168 |
| cluster2919 | -1.03 | 0.6173 |
| cluster3566 | 1.28 | 0.6180 |
| cluster3579 | -1.02 | 0.6203 |
| cluster3810 | -1.10 | 0.6204 |
| cluster3635 | -0.90 | 0.6204 |
| cluster3855 | 1.67 | 0.6208 |
| cluster3933 | 1.31 | 0.6208 |
| cluster2994 | -0.97 | 0.6208 |
| cluster3372 | -0.94 | 0.6208 |
| cluster3083 | -0.97 | 0.6208 |
| cluster3712 | -1.18 | 0.6208 |
| cluster3956 | -1.06 | 0.6208 |
| cluster3477 | -1.19 | 0.6208 |
| cluster3739 | -0.87 | 0.6208 |
| cluster3490 | 1.55 | 0.6230 |
| cluster3974 | 2.07 | 0.6282 |
| cluster4060 | -1.22 | 0.6289 |
| cluster3048 | 1.13 | 0.6311 |
| cluster4027 | 1.72 | 0.6353 |
| cluster3278 | -0.90 | 0.6353 |
| cluster3215 | -0.90 | 0.6353 |
| cluster3035 | -0.87 | 0.6368 |
| cluster3220 | 1.14 | 0.6368 |
| cluster4085 | 1.07 | 0.6368 |
| cluster3632 | -1.05 | 0.6368 |
| cluster3536 | -0.97 | 0.6368 |
| cluster3002 | -0.93 | 0.6368 |
| cluster3512 | 1.45 | 0.6405 |
| cluster3588 | -1.14 | 0.6406 |
| cluster3411 | 1.06 | 0.6406 |
| cluster3273 | -0.98 | 0.6406 |
| cluster3458 | -0.93 | 0.6406 |
| cluster3558 | 1.37 | 0.6406 |
| cluster3281 | 1.25 | 0.6406 |
| cluster3750 | 1.89 | 0.6419 |

| | | |
|---|---|---|
| cluster3381 | -0.95 | 0.6419 |
| cluster4040 | 1.38 | 0.6419 |
| cluster3125 | 1.04 | 0.6419 |
| cluster4008 | 1.56 | 0.6419 |
| cluster3403 | 1.01 | 0.6419 |
| cluster2974 | -0.93 | 0.6419 |
| cluster2886 | -1.02 | 0.6419 |
| cluster3702 | 1.27 | 0.6457 |
| cluster3265 | -0.92 | 0.6458 |
| cluster3857 | 1.39 | 0.6485 |
| cluster3671 | 1.15 | 0.6485 |
| cluster2925 | -0.96 | 0.6485 |
| cluster3869 | 1.59 | 0.6485 |
| cluster4119 | 1.60 | 0.6485 |
| cluster3807 | -0.96 | 0.6485 |
| cluster3069 | -0.81 | 0.6521 |
| cluster3433 | -1.02 | 0.6533 |
| cluster4005 | 1.48 | 0.6533 |
| cluster4033 | 1.56 | 0.6533 |
| cluster3926 | 1.53 | 0.6533 |
| cluster3806 | 1.46 | 0.6537 |
| cluster3526 | -0.89 | 0.6537 |
| cluster3481 | 1.53 | 0.6537 |
| cluster3503 | 1.17 | 0.6537 |
| cluster3885 | 1.08 | 0.6537 |
| cluster2912 | -0.83 | 0.6552 |
| cluster3938 | 1.51 | 0.6552 |
| cluster2890 | -0.82 | 0.6552 |
| cluster3686 | 1.67 | 0.6552 |
| cluster3208 | -0.85 | 0.6585 |
| cluster3740 | 1.49 | 0.6587 |
| cluster3301 | -1.01 | 0.6589 |
| cluster3542 | 1.63 | 0.6589 |
| cluster3783 | 1.67 | 0.6589 |
| cluster3661 | -0.89 | 0.6589 |
| cluster3914 | 1.78 | 0.6589 |
| cluster3004 | -0.91 | 0.6589 |
| cluster3493 | -1.16 | 0.6589 |
| cluster3405 | -0.92 | 0.6589 |
| cluster3223 | -0.96 | 0.6589 |
| cluster3563 | 1.10 | 0.6589 |
| cluster3198 | 1.17 | 0.6591 |
| cluster3090 | -1.22 | 0.6603 |
| cluster3143 | -0.95 | 0.6605 |

| | | |
|---|---|---|
| cluster3476 | 1.59 | 0.6609 |
| cluster3067 | -0.90 | 0.6609 |
| cluster3906 | 1.28 | 0.6609 |
| cluster2953 | -0.83 | 0.6664 |
| cluster4010 | 1.50 | 0.6676 |
| cluster3703 | -0.83 | 0.6676 |
| cluster3238 | -0.82 | 0.6676 |
| cluster3463 | 1.19 | 0.6706 |
| cluster2884 | -0.81 | 0.6737 |
| cluster3401 | -0.78 | 0.6737 |
| cluster3534 | 1.03 | 0.6796 |
| cluster3026 | 1.36 | 0.6797 |
| cluster3985 | -0.99 | 0.6797 |
| cluster3320 | 1.31 | 0.6797 |
| cluster3710 | 1.20 | 0.6797 |
| cluster3321 | -0.85 | 0.6797 |
| cluster4097 | 1.02 | 0.6797 |
| cluster2870 | -0.85 | 0.6797 |
| cluster3753 | 0.93 | 0.6797 |
| cluster3034 | 1.11 | 0.6797 |
| cluster3473 | -1.00 | 0.6797 |
| cluster3279 | 1.13 | 0.6818 |
| cluster3256 | -0.88 | 0.6819 |
| cluster3969 | -0.75 | 0.6859 |
| cluster3527 | 1.51 | 0.6859 |
| cluster3291 | -0.91 | 0.6859 |
| cluster3356 | -0.95 | 0.6859 |
| cluster3616 | -0.96 | 0.6862 |
| cluster3217 | -0.88 | 0.6862 |
| cluster3267 | -0.86 | 0.6869 |
| cluster3257 | -0.82 | 0.6869 |
| cluster3737 | -0.79 | 0.6874 |
| cluster3658 | -0.80 | 0.6909 |
| cluster3009 | 0.97 | 0.6935 |
| cluster3539 | 1.60 | 0.6953 |
| cluster3209 | 1.02 | 0.6953 |
| cluster3287 | -0.97 | 0.6953 |
| cluster2854 | -0.84 | 0.6953 |
| cluster3780 | 1.56 | 0.6973 |
| cluster3396 | -0.94 | 0.6973 |
| cluster3359 | 1.26 | 0.6973 |
| cluster3349 | -0.88 | 0.6973 |
| cluster3290 | 1.13 | 0.6973 |
| cluster3046 | -0.84 | 0.6986 |

| | | |
|---|---|---|
| cluster4100 | 1.38 | 0.7022 |
| cluster3759 | -0.94 | 0.7022 |
| cluster3350 | 1.14 | 0.7022 |
| cluster3467 | 1.01 | 0.7022 |
| cluster3146 | -0.72 | 0.7060 |
| cluster3666 | 0.99 | 0.7060 |
| cluster3489 | 1.08 | 0.7068 |
| cluster3723 | -0.75 | 0.7068 |
| cluster3779 | -0.90 | 0.7068 |
| cluster3212 | 1.15 | 0.7068 |
| cluster3784 | -0.95 | 0.7080 |
| cluster3199 | -0.72 | 0.7080 |
| cluster3745 | -1.02 | 0.7080 |
| cluster3679 | -0.87 | 0.7103 |
| cluster3430 | 1.45 | 0.7170 |
| cluster2940 | 0.96 | 0.7170 |
| cluster3803 | 1.48 | 0.7170 |
| cluster3576 | 1.06 | 0.7170 |
| cluster3446 | 0.93 | 0.7203 |
| cluster3618 | -1.21 | 0.7203 |
| cluster3581 | 0.94 | 0.7203 |
| cluster4044 | 1.06 | 0.7258 |
| cluster3822 | -1.10 | 0.7258 |
| cluster3302 | 1.02 | 0.7258 |
| cluster3557 | 1.10 | 0.7258 |
| cluster3440 | -0.69 | 0.7258 |
| cluster2942 | -0.77 | 0.7258 |
| cluster3326 | -0.70 | 0.7258 |
| cluster3202 | -0.78 | 0.7261 |
| cluster2992 | -0.72 | 0.7296 |
| cluster3657 | 0.90 | 0.7296 |
| cluster2949 | -0.73 | 0.7296 |
| cluster3353 | -0.72 | 0.7296 |
| cluster4094 | -1.04 | 0.7296 |
| cluster2909 | -0.72 | 0.7296 |
| cluster3164 | 1.38 | 0.7296 |
| cluster3174 | -0.91 | 0.7297 |
| cluster3772 | 0.99 | 0.7300 |
| cluster2894 | 0.91 | 0.7301 |
| cluster3196 | -0.68 | 0.7313 |
| cluster3851 | 1.27 | 0.7313 |
| cluster3070 | -0.79 | 0.7313 |
| cluster3953 | 0.82 | 0.7313 |
| cluster3231 | -0.84 | 0.7313 |

| | | |
|---|---|---|
| cluster3813 | 1.30 | 0.7317 |
| cluster3065 | -0.72 | 0.7317 |
| cluster3419 | -0.69 | 0.7317 |
| cluster3040 | -0.76 | 0.7317 |
| cluster3929 | 1.24 | 0.7317 |
| cluster3525 | 0.92 | 0.7317 |
| cluster3528 | -0.69 | 0.7317 |
| cluster4047 | 1.40 | 0.7317 |
| cluster4026 | 1.35 | 0.7328 |
| cluster3339 | -0.79 | 0.7328 |
| cluster3357 | -0.75 | 0.7328 |
| cluster3093 | 0.91 | 0.7328 |
| cluster3437 | 1.04 | 0.7328 |
| cluster3056 | -0.71 | 0.7328 |
| cluster3276 | -0.75 | 0.7328 |
| cluster3280 | -0.73 | 0.7328 |
| cluster3930 | -0.80 | 0.7328 |
| cluster3190 | 1.45 | 0.7328 |
| cluster3932 | 1.35 | 0.7328 |
| cluster3135 | 0.86 | 0.7328 |
| cluster2958 | -0.65 | 0.7328 |
| cluster3418 | 1.01 | 0.7328 |
| cluster3332 | -0.80 | 0.7328 |
| cluster2939 | 0.91 | 0.7328 |
| cluster3496 | -0.82 | 0.7328 |
| cluster3354 | -0.70 | 0.7328 |
| cluster3893 | -0.70 | 0.7349 |
| cluster3039 | -0.70 | 0.7349 |
| cluster3864 | 0.93 | 0.7349 |
| cluster3854 | 0.86 | 0.7349 |
| cluster3084 | -0.66 | 0.7349 |
| cluster3941 | -0.69 | 0.7349 |
| cluster3891 | 1.07 | 0.7380 |
| cluster3916 | -0.77 | 0.7381 |
| cluster3849 | -0.80 | 0.7386 |
| cluster2995 | -0.81 | 0.7386 |
| cluster3743 | 0.86 | 0.7386 |
| cluster3606 | 1.26 | 0.7386 |
| cluster3211 | 1.27 | 0.7386 |
| cluster2901 | 1.01 | 0.7386 |
| cluster3064 | -0.86 | 0.7386 |
| cluster2981 | -0.69 | 0.7386 |
| cluster3016 | 1.28 | 0.7397 |
| cluster3085 | 1.13 | 0.7403 |

| | | |
|---|---|---|
| cluster3982 | 0.91 | 0.7403 |
| cluster3421 | -0.88 | 0.7406 |
| cluster3453 | 0.94 | 0.7415 |
| cluster3636 | 0.76 | 0.7415 |
| cluster3128 | -0.67 | 0.7425 |
| cluster3860 | -0.76 | 0.7446 |
| cluster3241 | -0.70 | 0.7476 |
| cluster3266 | -0.71 | 0.7476 |
| cluster3895 | 1.25 | 0.7489 |
| cluster3724 | 1.04 | 0.7513 |
| cluster3939 | 0.99 | 0.7513 |
| cluster3344 | -0.80 | 0.7524 |
| cluster3201 | -0.64 | 0.7524 |
| cluster3907 | 1.30 | 0.7524 |
| cluster3975 | 0.81 | 0.7524 |
| cluster3736 | 1.22 | 0.7524 |
| cluster3966 | -0.66 | 0.7528 |
| cluster3972 | 0.94 | 0.7528 |
| cluster3390 | -0.71 | 0.7528 |
| cluster3637 | -0.61 | 0.7544 |
| cluster3398 | 0.72 | 0.7564 |
| cluster3314 | -0.82 | 0.7568 |
| cluster3793 | 0.83 | 0.7568 |
| cluster3383 | 1.26 | 0.7582 |
| cluster3022 | 1.02 | 0.7591 |
| cluster3646 | 1.32 | 0.7591 |
| cluster3555 | 1.23 | 0.7591 |
| cluster3236 | 0.92 | 0.7591 |
| cluster3043 | -0.68 | 0.7594 |
| cluster3560 | 1.19 | 0.7594 |
| cluster4020 | -0.68 | 0.7594 |
| cluster3959 | 1.26 | 0.7594 |
| cluster3282 | 0.83 | 0.7594 |
| cluster3479 | 1.19 | 0.7594 |
| cluster3720 | 1.17 | 0.7594 |
| cluster3150 | 1.09 | 0.7594 |
| cluster4090 | 1.38 | 0.7594 |
| cluster3204 | -0.72 | 0.7632 |
| cluster3901 | 0.88 | 0.7634 |
| cluster3136 | -0.73 | 0.7634 |
| cluster3251 | 0.79 | 0.7634 |
| cluster3611 | 1.18 | 0.7634 |
| cluster2951 | 0.97 | 0.7634 |
| cluster2862 | -0.63 | 0.7634 |

| | | |
|---|---|---|
| cluster2903 | -0.76 | 0.7658 |
| cluster3735 | 0.88 | 0.7658 |
| cluster3918 | -0.81 | 0.7661 |
| cluster3003 | -0.65 | 0.7678 |
| cluster3622 | -0.58 | 0.7683 |
| cluster3761 | -0.76 | 0.7683 |
| cluster3382 | -0.58 | 0.7684 |
| cluster3601 | 0.94 | 0.7699 |
| cluster3307 | 0.81 | 0.7749 |
| cluster4108 | -0.87 | 0.7782 |
| cluster3821 | 0.82 | 0.7782 |
| cluster3870 | 1.20 | 0.7792 |
| cluster3999 | 1.18 | 0.7792 |
| cluster4070 | 1.25 | 0.7793 |
| cluster3677 | -0.64 | 0.7793 |
| cluster2955 | -0.54 | 0.7827 |
| cluster2865 | -0.61 | 0.7827 |
| cluster3545 | 0.86 | 0.7827 |
| cluster2929 | -0.56 | 0.7832 |
| cluster3983 | 1.00 | 0.7844 |
| cluster3347 | -0.59 | 0.7844 |
| cluster3263 | -0.59 | 0.7891 |
| cluster3355 | 0.97 | 0.7893 |
| cluster3305 | 0.89 | 0.7893 |
| cluster2858 | -0.57 | 0.7897 |
| cluster3334 | -0.63 | 0.7923 |
| cluster4124 | 1.12 | 0.7927 |
| cluster2864 | -0.58 | 0.7927 |
| cluster3474 | 0.86 | 0.7927 |
| cluster3819 | 0.98 | 0.7934 |
| cluster3336 | -0.69 | 0.7941 |
| cluster3896 | 1.08 | 0.7978 |
| cluster3587 | -0.80 | 0.8003 |
| cluster3062 | 0.75 | 0.8005 |
| cluster2892 | -0.55 | 0.8005 |
| cluster3825 | 0.86 | 0.8037 |
| cluster3482 | 0.74 | 0.8070 |
| cluster3134 | 0.74 | 0.8070 |
| cluster2979 | -0.53 | 0.8096 |
| cluster3316 | 0.88 | 0.8111 |
| cluster3942 | 0.72 | 0.8111 |
| cluster3734 | 0.87 | 0.8111 |
| cluster3841 | 0.91 | 0.8119 |
| cluster3726 | -0.56 | 0.8119 |

| | | |
|---|---|---|
| cluster3845 | -0.53 | 0.8119 |
| cluster2898 | -0.65 | 0.8119 |
| cluster3957 | 0.88 | 0.8119 |
| cluster3596 | 0.83 | 0.8119 |
| cluster3683 | 0.70 | 0.8119 |
| cluster4121 | -0.69 | 0.8119 |
| cluster3578 | 0.90 | 0.8119 |
| cluster3499 | 0.94 | 0.8119 |
| cluster3524 | -0.53 | 0.8165 |
| cluster3688 | 0.71 | 0.8165 |
| cluster4016 | -0.62 | 0.8165 |
| cluster3862 | -0.51 | 0.8165 |
| cluster2871 | 0.68 | 0.8165 |
| cluster3823 | -0.61 | 0.8165 |
| cluster3586 | -0.51 | 0.8165 |
| cluster3968 | 0.87 | 0.8165 |
| cluster3120 | -0.47 | 0.8165 |
| cluster3998 | 1.12 | 0.8165 |
| cluster2970 | -0.47 | 0.8165 |
| cluster3776 | -0.63 | 0.8165 |
| cluster3285 | -0.56 | 0.8165 |
| cluster2969 | -0.58 | 0.8165 |
| cluster3584 | 1.03 | 0.8165 |
| cluster3100 | -0.63 | 0.8165 |
| cluster3395 | 0.69 | 0.8193 |
| cluster3769 | 0.71 | 0.8193 |
| cluster4032 | -0.64 | 0.8193 |
| cluster3795 | 0.72 | 0.8193 |
| cluster3148 | -0.47 | 0.8199 |
| cluster3055 | -0.50 | 0.8199 |
| cluster3086 | -0.49 | 0.8213 |
| cluster4116 | 1.31 | 0.8223 |
| cluster4075 | -0.79 | 0.8242 |
| cluster3796 | 0.99 | 0.8286 |
| cluster3782 | -0.58 | 0.8339 |
| cluster3668 | 0.62 | 0.8339 |
| cluster3488 | 0.60 | 0.8389 |
| cluster3815 | 1.00 | 0.8394 |
| cluster3131 | 0.61 | 0.8394 |
| cluster3655 | 0.76 | 0.8394 |
| cluster3140 | -0.48 | 0.8408 |
| cluster3533 | -0.49 | 0.8408 |
| cluster2856 | 0.74 | 0.8408 |
| cluster3665 | 0.99 | 0.8408 |

| | | |
|---|---|---|
| cluster3874 | -0.53 | 0.8408 |
| cluster3447 | 0.83 | 0.8408 |
| cluster3435 | 0.95 | 0.8408 |
| cluster4041 | 0.78 | 0.8408 |
| cluster4025 | -0.48 | 0.8408 |
| cluster2986 | 0.57 | 0.8408 |
| cluster3654 | 0.70 | 0.8415 |
| cluster3664 | -0.53 | 0.8423 |
| cluster3706 | 0.64 | 0.8423 |
| cluster2990 | 0.60 | 0.8423 |
| cluster3747 | -0.49 | 0.8423 |
| cluster4000 | -0.58 | 0.8423 |
| cluster3152 | 0.95 | 0.8423 |
| cluster3292 | 0.74 | 0.8423 |
| cluster3227 | -0.52 | 0.8423 |
| cluster3837 | 0.82 | 0.8426 |
| cluster2991 | 0.66 | 0.8454 |
| cluster2883 | -0.45 | 0.8454 |
| cluster3585 | 0.65 | 0.8454 |
| cluster3621 | -0.60 | 0.8454 |
| cluster3919 | 0.53 | 0.8454 |
| cluster3261 | 0.91 | 0.8454 |
| cluster3663 | 0.80 | 0.8454 |
| cluster2966 | 0.68 | 0.8454 |
| cluster3232 | -0.50 | 0.8454 |
| cluster3275 | 0.86 | 0.8461 |
| cluster4054 | 0.78 | 0.8482 |
| cluster2877 | -0.41 | 0.8482 |
| cluster3701 | -0.48 | 0.8485 |
| cluster4120 | 0.99 | 0.8485 |
| cluster3127 | 0.52 | 0.8513 |
| cluster3554 | 0.70 | 0.8515 |
| cluster3475 | 0.66 | 0.8525 |
| cluster3432 | 0.65 | 0.8525 |
| cluster3470 | 0.54 | 0.8532 |
| cluster3913 | -0.53 | 0.8558 |
| cluster3080 | -0.46 | 0.8558 |
| cluster3900 | 1.07 | 0.8558 |
| cluster3590 | 0.75 | 0.8558 |
| cluster3867 | 0.51 | 0.8558 |
| cluster2971 | -0.40 | 0.8558 |
| cluster3331 | 0.56 | 0.8558 |
| cluster3911 | -0.61 | 0.8558 |
| cluster2938 | -0.41 | 0.8558 |

| | | |
|---|---|---|
| cluster3151 | -0.39 | 0.8558 |
| cluster3608 | 0.54 | 0.8558 |
| cluster3038 | 0.71 | 0.8558 |
| cluster3880 | -0.63 | 0.8558 |
| cluster3713 | -0.48 | 0.8558 |
| cluster3673 | 0.97 | 0.8569 |
| cluster3498 | -0.54 | 0.8570 |
| cluster3464 | 0.71 | 0.8570 |
| cluster3765 | 0.64 | 0.8600 |
| cluster3343 | -0.40 | 0.8606 |
| cluster2936 | 0.87 | 0.8610 |
| cluster2957 | 0.75 | 0.8610 |
| cluster3058 | -0.39 | 0.8610 |
| cluster4007 | -0.43 | 0.8610 |
| cluster3729 | 0.56 | 0.8624 |
| cluster3294 | -0.41 | 0.8626 |
| cluster3725 | 0.84 | 0.8631 |
| cluster3423 | 0.91 | 0.8631 |
| cluster3675 | -0.37 | 0.8631 |
| cluster3649 | 0.54 | 0.8631 |
| cluster2998 | 0.49 | 0.8633 |
| cluster3744 | 0.63 | 0.8636 |
| cluster4067 | -0.58 | 0.8636 |
| cluster3708 | 0.96 | 0.8636 |
| cluster3218 | -0.51 | 0.8636 |
| cluster3829 | -0.56 | 0.8636 |
| cluster3872 | -0.51 | 0.8636 |
| cluster3944 | 0.61 | 0.8636 |
| cluster3826 | 0.82 | 0.8636 |
| cluster3284 | -0.42 | 0.8636 |
| cluster3133 | 0.58 | 0.8636 |
| cluster3509 | -0.42 | 0.8650 |
| cluster1127 | -0.46 | 0.8659 |
| cluster4073 | -0.56 | 0.8659 |
| cluster2917 | 0.72 | 0.8659 |
| cluster3451 | 0.99 | 0.8663 |
| cluster4002 | 0.66 | 0.8663 |
| cluster2997 | 0.52 | 0.8663 |
| cluster3117 | 0.89 | 0.8663 |
| cluster2920 | -0.39 | 0.8663 |
| cluster3934 | -0.35 | 0.8663 |
| cluster3946 | 0.72 | 0.8665 |
| cluster3559 | -0.42 | 0.8665 |
| cluster2944 | -0.42 | 0.8665 |

| | | |
|---|---|---|
| cluster3687 | 0.78 | 0.8710 |
| cluster3207 | -0.41 | 0.8725 |
| cluster3886 | -0.44 | 0.8725 |
| cluster3800 | 0.49 | 0.8731 |
| cluster3970 | 0.67 | 0.8743 |
| cluster3523 | 0.77 | 0.8756 |
| cluster3988 | -0.37 | 0.8762 |
| cluster4105 | -0.45 | 0.8762 |
| cluster3838 | 0.69 | 0.8786 |
| cluster4074 | 0.73 | 0.8786 |
| cluster3072 | -0.42 | 0.8786 |
| cluster2880 | -0.38 | 0.8786 |
| cluster4086 | -0.53 | 0.8786 |
| cluster3785 | -0.43 | 0.8786 |
| cluster3754 | -0.38 | 0.8789 |
| cluster3835 | 0.75 | 0.8808 |
| cluster3247 | -0.41 | 0.8808 |
| cluster3861 | 0.51 | 0.8834 |
| cluster4114 | 0.90 | 0.8834 |
| cluster3707 | -0.50 | 0.8834 |
| cluster3797 | -0.43 | 0.8835 |
| cluster3543 | -0.45 | 0.8835 |
| cluster3001 | -0.34 | 0.8854 |
| cluster3628 | 0.63 | 0.8862 |
| cluster3483 | 0.43 | 0.8884 |
| cluster3380 | 0.53 | 0.8893 |
| cluster3111 | 0.45 | 0.8893 |
| cluster4084 | -0.34 | 0.8893 |
| cluster3540 | 0.74 | 0.8893 |
| cluster3521 | 0.52 | 0.8904 |
| cluster3910 | 0.76 | 0.8908 |
| cluster3249 | 0.45 | 0.8910 |
| cluster3252 | 0.75 | 0.8922 |
| cluster3438 | 0.58 | 0.8922 |
| cluster3718 | 0.70 | 0.8931 |
| cluster3193 | -0.31 | 0.8957 |
| cluster3308 | -0.32 | 0.8970 |
| cluster3884 | 0.72 | 0.8970 |
| cluster3427 | -0.38 | 0.8970 |
| cluster3719 | -0.40 | 0.9028 |
| cluster3517 | -0.28 | 0.9040 |
| cluster3415 | -0.33 | 0.9050 |
| cluster3505 | -0.31 | 0.9050 |
| cluster3441 | -0.32 | 0.9053 |

| | | |
|---|---|---|
| cluster3248 | -0.43 | 0.9053 |
| cluster3609 | -0.44 | 0.9053 |
| cluster3037 | 0.43 | 0.9054 |
| cluster3535 | -0.42 | 0.9058 |
| cluster3662 | 0.62 | 0.9074 |
| cluster3997 | 0.68 | 0.9090 |
| cluster3457 | -0.35 | 0.9107 |
| cluster3219 | 0.41 | 0.9107 |
| cluster3647 | 0.48 | 0.9107 |
| cluster3828 | 0.59 | 0.9107 |
| cluster3384 | -0.30 | 0.9107 |
| cluster3562 | 0.54 | 0.9127 |
| cluster2985 | -0.32 | 0.9154 |
| cluster3843 | -0.29 | 0.9169 |
| cluster3123 | -0.29 | 0.9169 |
| cluster3742 | 0.65 | 0.9169 |
| cluster3298 | -0.28 | 0.9169 |
| cluster3629 | 0.50 | 0.9169 |
| cluster3620 | -0.32 | 0.9169 |
| cluster3961 | 0.65 | 0.9171 |
| cluster3699 | 0.65 | 0.9171 |
| cluster3950 | -0.29 | 0.9176 |
| cluster3460 | -0.34 | 0.9191 |
| cluster3952 | -0.27 | 0.9195 |
| cluster2867 | 0.60 | 0.9195 |
| cluster3491 | 0.45 | 0.9195 |
| cluster3378 | -0.32 | 0.9198 |
| cluster3036 | 0.45 | 0.9229 |
| cluster4109 | -0.35 | 0.9242 |
| cluster4028 | -0.30 | 0.9242 |
| cluster3899 | -0.31 | 0.9242 |
| cluster3690 | -0.27 | 0.9242 |
| cluster4081 | 0.41 | 0.9242 |
| cluster3075 | 0.45 | 0.9242 |
| cluster3580 | 0.59 | 0.9242 |
| cluster3163 | 0.61 | 0.9242 |
| cluster3153 | -0.23 | 0.9242 |
| cluster3728 | -0.26 | 0.9252 |
| cluster3222 | -0.23 | 0.9264 |
| cluster3373 | -0.29 | 0.9275 |
| cluster2895 | -0.26 | 0.9282 |
| cluster3092 | -0.23 | 0.9290 |
| cluster4110 | 0.58 | 0.9290 |
| cluster3789 | -0.30 | 0.9290 |

| | | |
|---|---|---|
| cluster3936 | 0.51 | 0.9290 |
| cluster2896 | 0.35 | 0.9300 |
| cluster2968 | -0.23 | 0.9300 |
| cluster2872 | -0.24 | 0.9300 |
| cluster3791 | 0.49 | 0.9325 |
| cluster4096 | 0.39 | 0.9334 |
| cluster3859 | -0.23 | 0.9334 |
| cluster3951 | 0.57 | 0.9349 |
| cluster3781 | -0.27 | 0.9349 |
| cluster3145 | -0.24 | 0.9349 |
| cluster2956 | 0.42 | 0.9353 |
| cluster2952 | 0.47 | 0.9353 |
| cluster4091 | -0.33 | 0.9353 |
| cluster3033 | 0.31 | 0.9353 |
| cluster4115 | 0.66 | 0.9367 |
| cluster3604 | 0.56 | 0.9367 |
| cluster3583 | -0.21 | 0.9367 |
| cluster3050 | -0.22 | 0.9376 |
| cluster3603 | -0.26 | 0.9377 |
| cluster3224 | 0.37 | 0.9377 |
| cluster3605 | 0.44 | 0.9390 |
| cluster3923 | -0.25 | 0.9390 |
| cluster3871 | 0.40 | 0.9390 |
| cluster4009 | 0.56 | 0.9391 |
| cluster3945 | -0.26 | 0.9392 |
| cluster3978 | 0.34 | 0.9394 |
| cluster3879 | 0.43 | 0.9395 |
| cluster3189 | -0.25 | 0.9398 |
| cluster2915 | 0.36 | 0.9398 |
| cluster3960 | 0.59 | 0.9398 |
| cluster4063 | 0.55 | 0.9398 |
| cluster2988 | -0.21 | 0.9398 |
| cluster2907 | -0.21 | 0.9398 |
| cluster3010 | 0.35 | 0.9400 |
| cluster3379 | 0.31 | 0.9416 |
| cluster3409 | 0.52 | 0.9419 |
| cluster4017 | 0.49 | 0.9430 |
| cluster2879 | -0.20 | 0.9430 |
| cluster3484 | -0.23 | 0.9455 |
| cluster3552 | -0.18 | 0.9479 |
| cluster2891 | 0.41 | 0.9482 |
| cluster3028 | 0.34 | 0.9494 |
| cluster3607 | 0.37 | 0.9496 |
| cluster3210 | -0.19 | 0.9496 |

| | | |
|---|---|---|
| cluster3809 | -0.19 | 0.9496 |
| cluster3502 | -0.23 | 0.9504 |
| cluster2897 | -0.20 | 0.9504 |
| cluster3627 | -0.19 | 0.9525 |
| cluster3116 | -0.18 | 0.9525 |
| cluster3877 | 0.33 | 0.9525 |
| cluster3775 | 0.33 | 0.9525 |
| cluster3642 | 0.42 | 0.9525 |
| cluster3167 | 0.51 | 0.9525 |
| cluster3324 | 0.36 | 0.9529 |
| cluster3104 | 0.27 | 0.9532 |
| cluster3333 | 0.35 | 0.9539 |
| cluster4099 | -0.19 | 0.9539 |
| cluster3081 | 0.30 | 0.9539 |
| cluster3108 | -0.19 | 0.9539 |
| cluster3980 | -0.17 | 0.9539 |
| cluster3363 | -0.18 | 0.9539 |
| cluster3095 | 0.32 | 0.9539 |
| cluster4029 | -0.20 | 0.9539 |
| cluster2987 | -0.16 | 0.9539 |
| cluster3888 | 0.46 | 0.9539 |
| cluster3428 | -0.16 | 0.9539 |
| cluster4062 | -0.20 | 0.9539 |
| cluster4003 | 0.49 | 0.9539 |
| cluster2885 | -0.18 | 0.9539 |
| cluster3310 | -0.15 | 0.9539 |
| cluster2857 | 0.28 | 0.9539 |
| cluster3330 | -0.17 | 0.9539 |
| cluster3242 | -0.16 | 0.9539 |
| cluster3264 | 0.26 | 0.9539 |
| cluster3648 | -0.15 | 0.9552 |
| cluster3183 | 0.31 | 0.9552 |
| cluster3507 | -0.16 | 0.9552 |
| cluster2918 | -0.15 | 0.9552 |
| cluster3680 | 0.30 | 0.9552 |
| cluster3592 | 0.31 | 0.9552 |
| cluster3214 | -0.16 | 0.9552 |
| cluster3283 | -0.15 | 0.9552 |
| cluster3544 | -0.14 | 0.9552 |
| cluster3875 | 0.30 | 0.9556 |
| cluster3633 | 0.26 | 0.9565 |
| cluster3667 | -0.14 | 0.9565 |
| cluster3947 | 0.32 | 0.9565 |
| cluster3180 | 0.26 | 0.9565 |

| | | |
|---|---|---|
| cluster3442 | 0.37 | 0.9565 |
| cluster3205 | -0.14 | 0.9565 |
| cluster3711 | -0.13 | 0.9565 |
| cluster4019 | -0.15 | 0.9565 |
| cluster3044 | -0.16 | 0.9565 |
| cluster2928 | 0.29 | 0.9565 |
| cluster3506 | 0.26 | 0.9565 |
| cluster3376 | -0.13 | 0.9565 |
| cluster3162 | 0.33 | 0.9565 |
| cluster4065 | 0.30 | 0.9565 |
| cluster4082 | 0.47 | 0.9565 |
| cluster3260 | 0.29 | 0.9565 |
| cluster3195 | 0.27 | 0.9565 |
| cluster2922 | 0.29 | 0.9565 |
| cluster3682 | 0.45 | 0.9567 |
| cluster4036 | 0.31 | 0.9567 |
| cluster3846 | 0.29 | 0.9567 |
| cluster3160 | 0.22 | 0.9567 |
| cluster4015 | 0.24 | 0.9568 |
| cluster4123 | 0.34 | 0.9568 |
| cluster3124 | 0.22 | 0.9568 |
| cluster4125 | -0.14 | 0.9568 |
| cluster3676 | -0.14 | 0.9568 |
| cluster3612 | 0.39 | 0.9572 |
| cluster3054 | -0.11 | 0.9572 |
| cluster3758 | 0.33 | 0.9595 |
| cluster2913 | -0.11 | 0.9595 |
| cluster2983 | 0.24 | 0.9595 |
| cluster3161 | 0.24 | 0.9595 |
| cluster3935 | 0.39 | 0.9595 |
| cluster4077 | -0.12 | 0.9595 |
| cluster3293 | -0.11 | 0.9595 |
| cluster3413 | 0.32 | 0.9605 |
| cluster3508 | -0.14 | 0.9608 |
| cluster3920 | 0.32 | 0.9629 |
| cluster3159 | 0.23 | 0.9633 |
| cluster3184 | 0.25 | 0.9633 |
| cluster4021 | -0.15 | 0.9653 |
| cluster4004 | -0.09 | 0.9653 |
| cluster3927 | 0.28 | 0.9653 |
| cluster3727 | -0.11 | 0.9657 |
| cluster3965 | -0.16 | 0.9663 |
| cluster3943 | -0.10 | 0.9663 |
| cluster3420 | 0.26 | 0.9673 |

| | | |
|---|---|---|
| cluster3370 | 0.28 | 0.9673 |
| cluster2855 | 0.19 | 0.9673 |
| cluster4023 | 0.31 | 0.9673 |
| cluster3364 | 0.21 | 0.9673 |
| cluster3271 | -0.08 | 0.9673 |
| cluster4106 | 0.47 | 0.9673 |
| cluster3868 | 0.22 | 0.9673 |
| cluster3194 | 0.24 | 0.9673 |
| cluster3771 | 0.27 | 0.9673 |
| cluster3816 | 0.29 | 0.9673 |
| cluster3685 | -0.08 | 0.9673 |
| cluster3155 | -0.08 | 0.9673 |
| cluster3865 | 0.30 | 0.9673 |
| cluster3406 | -0.07 | 0.9673 |
| cluster3052 | -0.07 | 0.9673 |
| cluster3798 | 0.28 | 0.9673 |
| cluster3368 | 0.19 | 0.9673 |
| cluster2996 | -0.07 | 0.9673 |
| cluster3751 | 0.25 | 0.9673 |
| cluster3057 | -0.07 | 0.9673 |
| cluster3352 | 0.27 | 0.9673 |
| cluster4126 | -0.09 | 0.9673 |
| cluster3717 | 0.20 | 0.9673 |
| cluster3790 | 0.33 | 0.9673 |
| cluster3179 | -0.08 | 0.9673 |
| cluster3221 | 0.27 | 0.9673 |
| cluster4102 | -0.13 | 0.9673 |
| cluster3614 | 0.38 | 0.9673 |
| cluster3591 | 0.18 | 0.9673 |
| cluster4127 | 0.49 | 0.9673 |
| cluster3962 | 0.21 | 0.9674 |
| cluster3154 | -0.06 | 0.9677 |
| cluster3366 | -0.07 | 0.9684 |
| cluster3068 | 0.18 | 0.9684 |
| cluster3847 | 0.22 | 0.9684 |
| cluster3853 | 0.22 | 0.9690 |
| cluster3288 | -0.07 | 0.9690 |
| cluster3235 | -0.04 | 0.9690 |
| cluster3103 | 0.25 | 0.9700 |
| cluster2959 | 0.19 | 0.9707 |
| cluster3029 | 0.17 | 0.9707 |
| cluster3041 | 0.21 | 0.9707 |
| cluster3722 | 0.27 | 0.9707 |
| cluster3059 | -0.03 | 0.9731 |

| | | |
|---|---|---|
| cluster3098 | 0.16 | 0.9736 |
| cluster3949 | -0.06 | 0.9739 |
| cluster3094 | -0.02 | 0.9810 |
| cluster2859 | -0.04 | 0.9810 |
| cluster3424 | 0.25 | 0.9810 |
| cluster3394 | 0.17 | 0.9810 |
| cluster3492 | -0.03 | 0.9810 |
| cluster4024 | 0.24 | 0.9830 |
| cluster2945 | 0.30 | 0.9843 |
| cluster3365 | 0.18 | 0.9846 |
| cluster3903 | 0.15 | 0.9858 |
| cluster3921 | 0.28 | 0.9867 |
| cluster3617 | 0.21 | 0.9867 |
| cluster3696 | -0.02 | 0.9867 |
| cluster4093 | 0.25 | 0.9894 |
| cluster3571 | 0.22 | 0.9894 |
| cluster3007 | 0.15 | 0.9894 |
| cluster3981 | 0.19 | 0.9894 |
| cluster3624 | 0.13 | 0.9894 |
| cluster3091 | -0.01 | 0.9894 |
| cluster2874 | -0.01 | 0.9894 |
| cluster4113 | 0.12 | 0.9894 |
| cluster3768 | 0.20 | 0.9894 |
| cluster3986 | 0.16 | 0.9894 |
| cluster2927 | 0.21 | 0.9894 |
| cluster2910 | 0.00 | 0.9894 |
| cluster2931 | 0.16 | 0.9894 |
| cluster4006 | 0.13 | 0.9894 |
| cluster3678 | 0.15 | 0.9894 |
| cluster3996 | 0.17 | 0.9894 |
| cluster2973 | 0.16 | 0.9894 |
| cluster2889 | 0.12 | 0.9894 |
| cluster3912 | 0.11 | 0.9894 |
| cluster3832 | 0.15 | 0.9894 |
| cluster3643 | 0.23 | 0.9894 |
| cluster3881 | 0.21 | 0.9894 |
| cluster3848 | 0.15 | 0.9902 |
| cluster4092 | 0.24 | 0.9902 |
| cluster3216 | 0.16 | 0.9919 |
| cluster3073 | 0.10 | 0.9919 |
| cluster3917 | 0.18 | 0.9922 |
| cluster4037 | 0.13 | 0.9922 |
| cluster3047 | 0.11 | 0.9922 |
| cluster3122 | 0.11 | 0.9938 |

| | | |
|---|---|---|
| cluster4061 | 0.19 | 0.9947 |
| cluster3801 | 0.20 | 0.9963 |
| cluster3018 | 0.02 | 0.9966 |
| cluster3074 | 0.02 | 0.9967 |
| cluster2972 | 0.03 | 1.0000 |
| cluster3692 | 0.15 | 1.0000 |
| cluster3684 | 0.15 | 1.0000 |
| cluster3259 | 0.08 | 1.0000 |
| cluster3412 | 0.12 | 1.0000 |
| cluster2975 | 0.08 | 1.0000 |
| cluster3114 | 0.08 | 1.0000 |
| cluster3445 | 0.09 | 1.0000 |
| cluster3169 | 0.11 | 1.0000 |
| cluster3820 | 0.08 | 1.0000 |
| cluster3625 | 0.09 | 1.0000 |
| cluster2934 | 0.08 | 1.0000 |
| cluster3078 | 0.07 | 1.0000 |
| cluster3020 | 0.06 | 1.0000 |
| cluster3808 | 0.14 | 1.0000 |
| cluster2941 | 0.06 | 1.0000 |
| cluster3976 | 0.06 | 1.0000 |
| cluster3902 | 0.07 | 1.0000 |
| cluster3351 | 0.06 | 1.0000 |
| cluster3096 | 0.05 | 1.0000 |
| cluster3328 | 0.06 | 1.0000 |
| cluster2932 | 0.08 | 1.0000 |
| cluster4018 | 0.08 | 1.0000 |
| cluster3325 | 0.14 | 1.0000 |
| cluster3882 | 0.10 | 1.0000 |
| cluster3436 | 0.09 | 1.0000 |
| cluster3529 | 0.08 | 1.0000 |
| cluster3274 | 0.08 | 1.0000 |
| cluster3431 | 0.06 | 1.0000 |
| cluster3556 | 0.06 | 1.0000 |
| cluster3228 | 0.06 | 1.0000 |
| cluster3804 | 0.06 | 1.0000 |
| cluster3173 | 0.06 | 1.0000 |
| cluster3434 | 0.06 | 1.0000 |
| cluster3087 | 0.05 | 1.0000 |
| cluster3076 | 0.04 | 1.0000 |
| cluster3203 | 0.03 | 1.0000 |
| cluster3008 | 0.03 | 1.0000 |
| cluster4064 | 0.03 | 1.0000 |
| cluster4051 | 0.02 | 1.0000 |

| | | |
|---|---|---|
| cluster3589 | 0.00 | 1.0000 |

**Supplementary Table 17.** DESeq2 results with RSEM counts - Traumatic vs furuncular

| ID | log2FoldChange | Adjusted p value |
|---|---|---|
| cluster4034 | 11.68 | 9.85E-13 |
| cluster3272 | 10.92 | 1.44E-11 |
| cluster3752 | 6.40 | 0.001 |
| cluster3640 | 6.22 | 0.026 |
| cluster3233 | -4.25 | 0.031 |
| cluster3546 | 6.06 | 0.031 |
| cluster3006 | -3.75 | 0.053 |
| cluster3892 | 3.96 | 0.053 |
| cluster3191 | 3.82 | 0.056 |
| cluster3313 | 3.86 | 0.056 |
| cluster3770 | 5.40 | 0.056 |
| cluster3738 | -3.95 | 0.090 |
| cluster3755 | -4.47 | 0.098 |
| cluster3786 | -3.87 | 0.113 |
| cluster3303 | -3.52 | 0.146 |
| cluster2976 | 3.60 | 0.193 |
| cluster3631 | -3.15 | 0.221 |
| cluster3178 | -3.14 | 0.239 |
| cluster3182 | 3.20 | 0.240 |
| cluster2876 | 3.34 | 0.281 |
| cluster3030 | -3.06 | 0.281 |
| cluster3694 | 2.77 | 0.281 |
| cluster2948 | -2.97 | 0.290 |
| cluster3516 | 4.08 | 0.328 |
| cluster4069 | 3.06 | 0.328 |
| cluster3371 | -4.37 | 0.347 |
| cluster3393 | 3.73 | 0.356 |
| cluster2893 | 4.07 | 0.370 |
| cluster2914 | 2.79 | 0.397 |
| cluster3126 | -2.64 | 0.483 |
| cluster3177 | 2.96 | 0.483 |
| cluster3749 | 3.74 | 0.483 |
| cluster3905 | 3.91 | 0.528 |
| cluster3994 | 3.50 | 0.528 |
| cluster3060 | 3.38 | 0.538 |
| cluster3904 | 3.70 | 0.538 |
| cluster4080 | 2.57 | 0.538 |
| cluster3501 | 2.82 | 0.605 |
| cluster3840 | -2.35 | 0.692 |
| cluster3890 | -2.53 | 0.692 |
| cluster2888 | 3.09 | 0.731 |
| cluster2911 | 2.00 | 0.731 |

| | | |
|---|---|---|
| cluster2960 | -2.18 | 0.731 |
| cluster2980 | 3.13 | 0.731 |
| cluster3125 | 2.22 | 0.731 |
| cluster3175 | 3.00 | 0.731 |
| cluster3375 | 2.01 | 0.731 |
| cluster3599 | 3.05 | 0.731 |
| cluster3638 | 2.70 | 0.731 |
| cluster3644 | 2.12 | 0.731 |
| cluster3773 | 1.90 | 0.731 |
| cluster3105 | 3.36 | 0.753 |
| cluster3670 | 2.33 | 0.753 |
| cluster3764 | 2.97 | 0.783 |
| cluster3129 | 2.90 | 0.817 |
| cluster3389 | 2.82 | 0.823 |
| cluster3500 | 3.14 | 0.823 |
| cluster3360 | 2.26 | 0.835 |
| cluster3462 | 1.92 | 0.835 |
| cluster2860 | -1.84 | 0.854 |
| cluster3410 | -1.98 | 0.854 |
| cluster3454 | 1.76 | 0.854 |
| cluster3652 | -1.95 | 0.854 |
| cluster2899 | -1.76 | 0.860 |
| cluster2902 | -2.81 | 0.860 |
| cluster2984 | -1.78 | 0.860 |
| cluster3017 | -2.32 | 0.860 |
| cluster3053 | -1.90 | 0.860 |
| cluster3137 | -2.95 | 0.860 |
| cluster3176 | 2.32 | 0.860 |
| cluster3185 | 2.93 | 0.860 |
| cluster3186 | 2.76 | 0.860 |
| cluster3197 | -1.79 | 0.860 |
| cluster3220 | 1.90 | 0.860 |
| cluster3234 | 1.89 | 0.860 |
| cluster3237 | -2.94 | 0.860 |
| cluster3386 | 2.60 | 0.860 |
| cluster3400 | -1.67 | 0.860 |
| cluster3459 | 2.47 | 0.860 |
| cluster3466 | 2.07 | 0.860 |
| cluster3538 | -1.84 | 0.860 |
| cluster3650 | -2.06 | 0.860 |
| cluster3659 | 2.78 | 0.860 |
| cluster3730 | -1.63 | 0.860 |
| cluster3800 | 1.88 | 0.860 |
| cluster3948 | 2.53 | 0.860 |

| | | |
|---|---|---|
| cluster4059 | 2.71 | 0.860 |
| cluster2933 | -1.78 | 0.867 |
| cluster2853 | -1.53 | 0.884 |
| cluster2861 | -1.66 | 0.884 |
| cluster2921 | -1.95 | 0.884 |
| cluster2937 | 1.69 | 0.884 |
| cluster2946 | -1.56 | 0.884 |
| cluster2993 | -1.79 | 0.884 |
| cluster3000 | 2.09 | 0.884 |
| cluster3061 | -1.58 | 0.884 |
| cluster3101 | -1.47 | 0.884 |
| cluster3127 | 1.60 | 0.884 |
| cluster3157 | -1.51 | 0.884 |
| cluster3200 | -1.54 | 0.884 |
| cluster3212 | 1.81 | 0.884 |
| cluster3254 | 2.51 | 0.884 |
| cluster3296 | 2.51 | 0.884 |
| cluster3513 | -1.60 | 0.884 |
| cluster3534 | 1.54 | 0.884 |
| cluster3563 | 1.72 | 0.884 |
| cluster3636 | 1.61 | 0.884 |
| cluster3669 | 2.35 | 0.884 |
| cluster3709 | -1.60 | 0.884 |
| cluster3760 | -1.57 | 0.884 |
| cluster3775 | 2.33 | 0.884 |
| cluster3850 | 2.56 | 0.884 |
| cluster3922 | 2.59 | 0.884 |
| cluster3977 | 2.39 | 0.884 |
| cluster4001 | 2.52 | 0.884 |
| cluster4078 | -1.71 | 0.884 |
| cluster2978 | -1.49 | 0.888 |
| cluster3032 | -1.43 | 0.888 |
| cluster3130 | 2.08 | 0.888 |
| cluster3144 | 2.21 | 0.888 |
| cluster3188 | -1.55 | 0.888 |
| cluster3297 | -1.78 | 0.888 |
| cluster3341 | 2.12 | 0.888 |
| cluster3385 | 2.27 | 0.888 |
| cluster3449 | -1.33 | 0.888 |
| cluster3482 | 1.60 | 0.888 |
| cluster3494 | -1.49 | 0.888 |
| cluster3511 | 2.32 | 0.888 |
| cluster3741 | -1.70 | 0.888 |
| cluster3788 | -1.37 | 0.888 |

| | | |
|---|---|---|
| cluster3795 | 2.24 | 0.888 |
| cluster4030 | 2.28 | 0.888 |
| cluster3397 | 2.20 | 0.892 |
| cluster3497 | -1.34 | 0.892 |
| cluster3766 | 2.30 | 0.892 |
| cluster2961 | -1.44 | 0.894 |
| cluster3119 | -1.40 | 0.894 |
| cluster3229 | -1.38 | 0.894 |
| cluster3453 | 1.58 | 0.894 |
| cluster3821 | 1.65 | 0.894 |
| cluster2873 | -1.30 | 0.929 |
| cluster2881 | -1.39 | 0.929 |
| cluster2943 | -1.44 | 0.929 |
| cluster3346 | -1.46 | 0.929 |
| cluster3387 | 2.03 | 0.929 |
| cluster3485 | 2.22 | 0.929 |
| cluster3487 | 2.05 | 0.929 |
| cluster3575 | -2.20 | 0.929 |
| cluster3891 | 1.59 | 0.929 |
| cluster3109 | 2.01 | 0.930 |
| cluster3181 | -1.30 | 0.930 |
| cluster3395 | 1.32 | 0.930 |
| cluster3735 | 1.50 | 0.930 |
| cluster3925 | 2.15 | 0.939 |
| cluster3567 | -2.08 | 0.945 |
| cluster3876 | -1.26 | 0.945 |
| cluster4085 | 1.43 | 0.945 |
| cluster3073 | 1.45 | 0.969 |
| cluster3082 | 1.92 | 0.969 |
| cluster3304 | -1.21 | 0.969 |
| cluster2906 | -1.27 | 0.971 |
| cluster2963 | -1.28 | 0.971 |
| cluster3226 | 1.80 | 0.971 |
| cluster3246 | -1.22 | 0.971 |
| cluster3518 | 1.86 | 0.971 |
| cluster3608 | 1.29 | 0.971 |
| cluster3778 | -1.26 | 0.974 |
| cluster2854 | -0.56 | 0.975 |
| cluster2857 | 1.11 | 0.975 |
| cluster2859 | 0.37 | 0.975 |
| cluster2863 | -0.84 | 0.975 |
| cluster2866 | -1.73 | 0.975 |
| cluster2869 | -0.89 | 0.975 |
| cluster2870 | -1.24 | 0.975 |

| | | |
|---|---|---|
| cluster2871 | 1.14 | 0.975 |
| cluster2875 | -0.75 | 0.975 |
| cluster2878 | -1.20 | 0.975 |
| cluster2880 | -0.77 | 0.975 |
| cluster2884 | -0.81 | 0.975 |
| cluster2885 | -0.67 | 0.975 |
| cluster2886 | -0.55 | 0.975 |
| cluster2887 | 1.24 | 0.975 |
| cluster2890 | -0.54 | 0.975 |
| cluster2894 | 0.73 | 0.975 |
| cluster2895 | -0.65 | 0.975 |
| cluster2896 | 0.61 | 0.975 |
| cluster2897 | -0.61 | 0.975 |
| cluster2900 | -1.15 | 0.975 |
| cluster2901 | 0.53 | 0.975 |
| cluster2904 | -1.11 | 0.975 |
| cluster2905 | -1.01 | 0.975 |
| cluster2907 | -0.59 | 0.975 |
| cluster2908 | -1.02 | 0.975 |
| cluster2918 | -0.51 | 0.975 |
| cluster2919 | -0.51 | 0.975 |
| cluster2920 | -0.69 | 0.975 |
| cluster2923 | -0.69 | 0.975 |
| cluster2924 | -1.38 | 0.975 |
| cluster2925 | -1.17 | 0.975 |
| cluster2926 | -0.80 | 0.975 |
| cluster2929 | -0.50 | 0.975 |
| cluster2930 | -1.15 | 0.975 |
| cluster2935 | -0.68 | 0.975 |
| cluster2940 | 0.61 | 0.975 |
| cluster2941 | 0.65 | 0.975 |
| cluster2942 | -0.97 | 0.975 |
| cluster2944 | -0.80 | 0.975 |
| cluster2950 | -0.73 | 0.975 |
| cluster2951 | 0.56 | 0.975 |
| cluster2953 | -0.67 | 0.975 |
| cluster2954 | -0.61 | 0.975 |
| cluster2956 | 0.79 | 0.975 |
| cluster2959 | 0.68 | 0.975 |
| cluster2964 | -0.68 | 0.975 |
| cluster2965 | -1.63 | 0.975 |
| cluster2969 | -1.01 | 0.975 |
| cluster2972 | 0.68 | 0.975 |
| cluster2974 | 0.61 | 0.975 |

| | | |
|---|---|---|
| cluster2977 | -0.78 | 0.975 |
| cluster2979 | -0.66 | 0.975 |
| cluster2981 | -0.89 | 0.975 |
| cluster2982 | -1.19 | 0.975 |
| cluster2992 | -0.46 | 0.975 |
| cluster2994 | -0.99 | 0.975 |
| cluster2998 | 1.58 | 0.975 |
| cluster2999 | 1.55 | 0.975 |
| cluster3002 | -0.74 | 0.975 |
| cluster3003 | -1.01 | 0.975 |
| cluster3004 | -0.36 | 0.975 |
| cluster3005 | -0.77 | 0.975 |
| cluster3009 | 0.81 | 0.975 |
| cluster3011 | -0.66 | 0.975 |
| cluster3013 | -0.78 | 0.975 |
| cluster3014 | 1.86 | 0.975 |
| cluster3019 | -1.46 | 0.975 |
| cluster3021 | -0.49 | 0.975 |
| cluster3023 | -1.25 | 0.975 |
| cluster3024 | -0.32 | 0.975 |
| cluster3025 | -0.42 | 0.975 |
| cluster3027 | -1.14 | 0.975 |
| cluster3031 | -0.60 | 0.975 |
| cluster3033 | 0.77 | 0.975 |
| cluster3035 | -0.51 | 0.975 |
| cluster3047 | 0.96 | 0.975 |
| cluster3048 | 1.15 | 0.975 |
| cluster3049 | -0.96 | 0.975 |
| cluster3056 | -0.67 | 0.975 |
| cluster3057 | 0.64 | 0.975 |
| cluster3059 | -0.66 | 0.975 |
| cluster3066 | -1.10 | 0.975 |
| cluster3068 | 0.83 | 0.975 |
| cluster3070 | -0.46 | 0.975 |
| cluster3071 | -0.91 | 0.975 |
| cluster3072 | -0.86 | 0.975 |
| cluster3077 | -0.63 | 0.975 |
| cluster3081 | 0.95 | 0.975 |
| cluster3086 | -0.48 | 0.975 |
| cluster3089 | -1.02 | 0.975 |
| cluster3090 | -1.64 | 0.975 |
| cluster3091 | 0.58 | 0.975 |
| cluster3092 | 1.42 | 0.975 |
| cluster3093 | 0.88 | 0.975 |

| | | |
|---|---|---|
| cluster3095 | 0.93 | 0.975 |
| cluster3100 | -0.63 | 0.975 |
| cluster3104 | 0.87 | 0.975 |
| cluster3106 | -0.54 | 0.975 |
| cluster3107 | -1.02 | 0.975 |
| cluster3108 | -0.53 | 0.975 |
| cluster3110 | -1.00 | 0.975 |
| cluster3111 | 1.09 | 0.975 |
| cluster3112 | 1.49 | 0.975 |
| cluster3113 | -0.76 | 0.975 |
| cluster3115 | -0.96 | 0.975 |
| cluster3121 | -0.54 | 0.975 |
| cluster3122 | 0.68 | 0.975 |
| cluster3135 | 0.65 | 0.975 |
| cluster3141 | -0.85 | 0.975 |
| cluster3147 | -0.68 | 0.975 |
| cluster3148 | -0.47 | 0.975 |
| cluster3149 | -0.46 | 0.975 |
| cluster3156 | 1.72 | 0.975 |
| cluster3160 | 0.41 | 0.975 |
| cluster3164 | 0.90 | 0.975 |
| cluster3166 | -0.75 | 0.975 |
| cluster3171 | -1.07 | 0.975 |
| cluster3172 | -0.58 | 0.975 |
| cluster3174 | -1.35 | 0.975 |
| cluster3180 | 0.62 | 0.975 |
| cluster3187 | 1.44 | 0.975 |
| cluster3192 | -0.95 | 0.975 |
| cluster3193 | -0.52 | 0.975 |
| cluster3195 | 0.81 | 0.975 |
| cluster3198 | 0.80 | 0.975 |
| cluster3199 | 0.44 | 0.975 |
| cluster3201 | -0.42 | 0.975 |
| cluster3209 | 0.86 | 0.975 |
| cluster3213 | -0.94 | 0.975 |
| cluster3215 | -0.58 | 0.975 |
| cluster3217 | -0.38 | 0.975 |
| cluster3218 | -1.05 | 0.975 |
| cluster3222 | 0.35 | 0.975 |
| cluster3224 | 0.48 | 0.975 |
| cluster3231 | -1.09 | 0.975 |
| cluster3235 | -0.54 | 0.975 |
| cluster3239 | -0.94 | 0.975 |
| cluster3242 | -0.44 | 0.975 |

| | | |
|---|---|---|
| cluster3243 | -0.98 | 0.975 |
| cluster3249 | 1.28 | 0.975 |
| cluster3251 | 0.57 | 0.975 |
| cluster3253 | 1.18 | 0.975 |
| cluster3255 | 1.16 | 0.975 |
| cluster3258 | -0.76 | 0.975 |
| cluster3259 | 0.68 | 0.975 |
| cluster3260 | 0.88 | 0.975 |
| cluster3264 | 0.85 | 0.975 |
| cluster3265 | -1.06 | 0.975 |
| cluster3268 | 1.44 | 0.975 |
| cluster3269 | 1.33 | 0.975 |
| cluster3270 | 1.53 | 0.975 |
| cluster3271 | 0.65 | 0.975 |
| cluster3281 | 1.40 | 0.975 |
| cluster3286 | -0.55 | 0.975 |
| cluster3298 | 0.50 | 0.975 |
| cluster3301 | -0.52 | 0.975 |
| cluster3307 | 0.54 | 0.975 |
| cluster3310 | 0.47 | 0.975 |
| cluster3314 | -1.24 | 0.975 |
| cluster3315 | -0.52 | 0.975 |
| cluster3319 | 1.46 | 0.975 |
| cluster3326 | -0.37 | 0.975 |
| cluster3328 | 0.80 | 0.975 |
| cluster3340 | -0.92 | 0.975 |
| cluster3354 | -0.60 | 0.975 |
| cluster3359 | 0.81 | 0.975 |
| cluster3361 | 1.17 | 0.975 |
| cluster3362 | -0.54 | 0.975 |
| cluster3368 | 0.72 | 0.975 |
| cluster3376 | 0.35 | 0.975 |
| cluster3379 | 0.94 | 0.975 |
| cluster3381 | -0.43 | 0.975 |
| cluster3383 | 0.79 | 0.975 |
| cluster3384 | 0.41 | 0.975 |
| cluster3388 | 1.65 | 0.975 |
| cluster3394 | 0.96 | 0.975 |
| cluster3398 | 0.92 | 0.975 |
| cluster3401 | -0.66 | 0.975 |
| cluster3403 | 1.25 | 0.975 |
| cluster3404 | -0.70 | 0.975 |
| cluster3406 | 0.48 | 0.975 |
| cluster3407 | -0.66 | 0.975 |

| | | |
|---|---|---|
| cluster3411 | 1.37 | 0.975 |
| cluster3419 | -0.53 | 0.975 |
| cluster3428 | 0.47 | 0.975 |
| cluster3429 | -0.60 | 0.975 |
| cluster3443 | 1.41 | 0.975 |
| cluster3444 | -0.45 | 0.975 |
| cluster3445 | 0.73 | 0.975 |
| cluster3446 | 1.27 | 0.975 |
| cluster3450 | 1.92 | 0.975 |
| cluster3457 | -0.86 | 0.975 |
| cluster3461 | 1.80 | 0.975 |
| cluster3465 | -1.00 | 0.975 |
| cluster3469 | -1.17 | 0.975 |
| cluster3470 | 1.31 | 0.975 |
| cluster3471 | 1.40 | 0.975 |
| cluster3472 | -0.60 | 0.975 |
| cluster3483 | 1.10 | 0.975 |
| cluster3484 | -0.55 | 0.975 |
| cluster3488 | 1.10 | 0.975 |
| cluster3489 | 0.76 | 0.975 |
| cluster3495 | -1.01 | 0.975 |
| cluster3504 | -0.70 | 0.975 |
| cluster3510 | 1.35 | 0.975 |
| cluster3517 | 0.57 | 0.975 |
| cluster3539 | 1.23 | 0.975 |
| cluster3541 | -0.78 | 0.975 |
| cluster3544 | 0.55 | 0.975 |
| cluster3547 | -0.65 | 0.975 |
| cluster3549 | -0.69 | 0.975 |
| cluster3552 | 0.50 | 0.975 |
| cluster3555 | 0.79 | 0.975 |
| cluster3570 | 1.35 | 0.975 |
| cluster3573 | 1.04 | 0.975 |
| cluster3577 | -0.77 | 0.975 |
| cluster3579 | -0.37 | 0.975 |
| cluster3581 | 0.79 | 0.975 |
| cluster3582 | -0.96 | 0.975 |
| cluster3583 | 0.61 | 0.975 |
| cluster3586 | -0.40 | 0.975 |
| cluster3587 | -1.26 | 0.975 |
| cluster3588 | -1.50 | 0.975 |
| cluster3591 | 0.91 | 0.975 |
| cluster3600 | -0.76 | 0.975 |
| cluster3601 | 0.57 | 0.975 |

| | | |
|---|---|---|
| cluster3609 | -0.87 | 0.975 |
| cluster3623 | 1.39 | 0.975 |
| cluster3624 | 0.71 | 0.975 |
| cluster3630 | -0.52 | 0.975 |
| cluster3633 | 1.03 | 0.975 |
| cluster3635 | -0.33 | 0.975 |
| cluster3648 | 0.44 | 0.975 |
| cluster3653 | -0.81 | 0.975 |
| cluster3671 | 1.14 | 0.975 |
| cluster3675 | 0.32 | 0.975 |
| cluster3685 | -0.56 | 0.975 |
| cluster3697 | -0.63 | 0.975 |
| cluster3703 | -0.35 | 0.975 |
| cluster3704 | -1.18 | 0.975 |
| cluster3717 | 0.89 | 0.975 |
| cluster3723 | 0.49 | 0.975 |
| cluster3728 | 0.53 | 0.975 |
| cluster3737 | 0.36 | 0.975 |
| cluster3739 | -0.36 | 0.975 |
| cluster3747 | -0.86 | 0.975 |
| cluster3748 | -0.70 | 0.975 |
| cluster3753 | 1.23 | 0.975 |
| cluster3761 | -1.27 | 0.975 |
| cluster3765 | 1.34 | 0.975 |
| cluster3780 | 1.11 | 0.975 |
| cluster3792 | 1.48 | 0.975 |
| cluster3803 | 1.01 | 0.975 |
| cluster3811 | -0.59 | 0.975 |
| cluster3830 | -1.01 | 0.975 |
| cluster3834 | 1.81 | 0.975 |
| cluster3847 | 0.54 | 0.975 |
| cluster3856 | 1.41 | 0.975 |
| cluster3867 | 0.47 | 0.975 |
| cluster3873 | -0.70 | 0.975 |
| cluster3877 | 1.04 | 0.975 |
| cluster3885 | 0.92 | 0.975 |
| cluster3900 | 0.58 | 0.975 |
| cluster3907 | 0.81 | 0.975 |
| cluster3912 | 1.07 | 0.975 |
| cluster3914 | 1.33 | 0.975 |
| cluster3915 | 1.96 | 0.975 |
| cluster3919 | 1.00 | 0.975 |
| cluster3926 | 1.33 | 0.975 |
| cluster3945 | -0.63 | 0.975 |

| | | |
|---|---|---|
| cluster3953 | 1.44 | 0.975 |
| cluster3963 | 1.72 | 0.975 |
| cluster3973 | 1.24 | 0.975 |
| cluster3976 | 0.68 | 0.975 |
| cluster4011 | 1.88 | 0.975 |
| cluster4014 | 1.49 | 0.975 |
| cluster4015 | 0.58 | 0.975 |
| cluster4019 | -0.62 | 0.975 |
| cluster4022 | 1.26 | 0.975 |
| cluster4029 | -0.70 | 0.975 |
| cluster4031 | -0.98 | 0.975 |
| cluster4042 | 1.28 | 0.975 |
| cluster4049 | 1.67 | 0.975 |
| cluster4084 | 0.38 | 0.975 |
| cluster4097 | 0.83 | 0.975 |
| cluster4113 | 1.20 | 0.975 |
| cluster3208 | -0.32 | 0.978 |
| cluster3688 | 0.51 | 0.978 |
| cluster2986 | 0.50 | 0.981 |
| cluster3308 | 0.33 | 0.981 |
| cluster3690 | 0.33 | 0.981 |
| cluster2872 | -0.40 | 0.984 |
| cluster2955 | -0.34 | 0.984 |
| cluster2968 | 0.27 | 0.984 |
| cluster3305 | 0.47 | 0.984 |
| cluster3779 | -0.34 | 0.984 |
| cluster3841 | 0.47 | 0.984 |
| cluster3120 | 0.36 | 0.986 |
| cluster3351 | 0.38 | 0.986 |
| cluster3058 | 0.27 | 0.987 |
| cluster3067 | -0.28 | 0.987 |
| cluster3069 | -0.27 | 0.987 |
| cluster3934 | 0.41 | 0.987 |
| cluster2855 | 0.26 | 0.991 |
| cluster2856 | 0.33 | 0.991 |
| cluster2858 | -0.17 | 0.991 |
| cluster2862 | -0.11 | 0.991 |
| cluster2867 | 0.14 | 0.991 |
| cluster2874 | 0.17 | 0.991 |
| cluster2877 | -0.21 | 0.991 |
| cluster2883 | 0.09 | 0.991 |
| cluster2910 | -0.23 | 0.991 |
| cluster2912 | -0.10 | 0.991 |
| cluster2913 | -0.16 | 0.991 |

| | | |
|---|---|---|
| cluster2917 | 0.27 | 0.991 |
| cluster2922 | -0.14 | 0.991 |
| cluster2931 | -0.21 | 0.991 |
| cluster2932 | -0.15 | 0.991 |
| cluster2938 | 0.17 | 0.991 |
| cluster2945 | -0.21 | 0.991 |
| cluster2949 | -0.20 | 0.991 |
| cluster2958 | 0.10 | 0.991 |
| cluster2966 | 0.27 | 0.991 |
| cluster2970 | -0.12 | 0.991 |
| cluster2971 | 0.25 | 0.991 |
| cluster2973 | -0.27 | 0.991 |
| cluster2983 | 0.12 | 0.991 |
| cluster2996 | 0.07 | 0.991 |
| cluster2997 | 0.22 | 0.991 |
| cluster3001 | 0.12 | 0.991 |
| cluster3018 | -0.16 | 0.991 |
| cluster3020 | -0.33 | 0.991 |
| cluster3029 | 0.33 | 0.991 |
| cluster3037 | 0.23 | 0.991 |
| cluster3039 | -0.15 | 0.991 |
| cluster3050 | -0.18 | 0.991 |
| cluster3054 | 0.09 | 0.991 |
| cluster3065 | -0.32 | 0.991 |
| cluster3074 | -0.11 | 0.991 |
| cluster3084 | -0.24 | 0.991 |
| cluster3096 | -0.16 | 0.991 |
| cluster3098 | 0.22 | 0.991 |
| cluster3114 | 0.25 | 0.991 |
| cluster3116 | -0.09 | 0.991 |
| cluster3124 | 0.31 | 0.991 |
| cluster3128 | -0.16 | 0.991 |
| cluster3131 | 0.33 | 0.991 |
| cluster3145 | 0.07 | 0.991 |
| cluster3151 | -0.13 | 0.991 |
| cluster3153 | 0.34 | 0.991 |
| cluster3161 | 0.27 | 0.991 |
| cluster3163 | 0.13 | 0.991 |
| cluster3169 | -0.34 | 0.991 |
| cluster3196 | 0.07 | 0.991 |
| cluster3207 | 0.11 | 0.991 |
| cluster3221 | -0.20 | 0.991 |
| cluster3252 | 0.34 | 0.991 |
| cluster3278 | -0.17 | 0.991 |

| | | |
|---|---|---|
| cluster3292 | 0.33 | 0.991 |
| cluster3294 | 0.21 | 0.991 |
| cluster3295 | -0.17 | 0.991 |
| cluster3343 | -0.13 | 0.991 |
| cluster3349 | -0.29 | 0.991 |
| cluster3364 | 0.24 | 0.991 |
| cluster3372 | -0.21 | 0.991 |
| cluster3382 | 0.10 | 0.991 |
| cluster3440 | 0.26 | 0.991 |
| cluster3505 | -0.22 | 0.991 |
| cluster3523 | 0.24 | 0.991 |
| cluster3528 | 0.08 | 0.991 |
| cluster3533 | 0.14 | 0.991 |
| cluster3617 | -0.21 | 0.991 |
| cluster3658 | -0.10 | 0.991 |
| cluster3677 | 0.10 | 0.991 |
| cluster3679 | -0.24 | 0.991 |
| cluster3729 | 0.34 | 0.991 |
| cluster3768 | -0.27 | 0.991 |
| cluster3816 | -0.11 | 0.991 |
| cluster3845 | 0.18 | 0.991 |
| cluster3859 | -0.11 | 0.991 |
| cluster3862 | 0.40 | 0.991 |
| cluster3921 | -0.18 | 0.991 |
| cluster3950 | 0.24 | 0.991 |
| cluster3961 | 0.17 | 0.991 |
| cluster3969 | 0.13 | 0.991 |
| cluster4004 | 0.17 | 0.991 |
| cluster4006 | 0.32 | 0.991 |
| cluster4041 | 0.42 | 0.991 |
| cluster4072 | 0.10 | 0.991 |
| cluster2865 | -0.06 | 0.991 |
| cluster3848 | -0.09 | 0.991 |
| cluster2891 | -0.08 | 0.993 |
| cluster3711 | 0.05 | 0.993 |
| cluster3903 | 0.08 | 0.993 |
| cluster3194 | 0.05 | 0.994 |
| cluster2889 | -0.04 | 0.995 |
| cluster2975 | 0.04 | 0.995 |
| cluster3010 | 0.03 | 0.995 |
| cluster3036 | 0.05 | 0.995 |
| cluster3409 | 0.05 | 0.995 |
| cluster3637 | -0.03 | 0.995 |
| cluster3893 | 0.03 | 0.995 |

| | | |
|---|---|---|
| cluster2864 | 0.01 | 0.997 |
| cluster2879 | -0.01 | 0.997 |
| cluster2892 | -0.01 | 0.997 |
| cluster2952 | 0.01 | 0.997 |
| cluster2985 | 0.02 | 0.997 |
| cluster3146 | -0.02 | 0.997 |
| cluster3616 | -0.01 | 0.997 |
| cluster3622 | 0.01 | 0.997 |

**Supplementary Table 18.** edgeR results with RSEM counts - Traumatic vs furuncular

| ID | log2FoldChange | Adjusted p value |
|---|---|---|
| cluster4034 | 11.53 | 0.005 |
| cluster3272 | 10.80 | 0.005 |
| cluster3233 | -4.31 | 0.206 |
| cluster4101 | -4.75 | 0.206 |
| cluster3752 | 6.66 | 0.413 |
| cluster3738 | -3.98 | 0.488 |
| cluster3755 | -4.41 | 0.488 |
| cluster3006 | -3.69 | 0.488 |
| cluster3786 | -3.87 | 0.625 |
| cluster3640 | 6.48 | 0.744 |
| cluster3546 | 6.27 | 0.765 |
| cluster4076 | -3.59 | 0.765 |
| cluster3303 | -3.50 | 0.765 |
| cluster3178 | -3.16 | 0.914 |
| cluster3030 | -3.11 | 0.914 |
| cluster3631 | -3.16 | 0.948 |
| cluster3770 | 5.60 | 1.000 |
| cluster3371 | -4.06 | 1.000 |
| cluster3892 | 4.00 | 1.000 |
| cluster2948 | -2.95 | 1.000 |
| cluster3312 | 4.76 | 1.000 |
| cluster3763 | -2.97 | 1.000 |
| cluster4056 | 4.57 | 1.000 |
| cluster3844 | 5.09 | 1.000 |
| cluster3191 | 3.85 | 1.000 |
| cluster4112 | -3.14 | 1.000 |
| cluster3126 | -2.72 | 1.000 |
| cluster3313 | 3.79 | 1.000 |
| cluster2976 | 3.89 | 1.000 |
| cluster3958 | 4.75 | 1.000 |
| cluster3695 | 3.63 | 1.000 |
| cluster3757 | -3.51 | 1.000 |
| cluster3802 | -2.89 | 1.000 |
| cluster4046 | 4.18 | 1.000 |
| cluster4111 | 4.86 | 1.000 |
| cluster3323 | -2.65 | 1.000 |
| cluster3852 | -2.60 | 1.000 |
| cluster3367 | -2.75 | 1.000 |
| cluster3392 | 4.06 | 1.000 |
| cluster3416 | 4.15 | 1.000 |
| cluster3931 | -2.86 | 1.000 |
| cluster3890 | -2.62 | 1.000 |

| | | |
|---|---|---|
| cluster3818 | -2.60 | 1.000 |
| cluster4098 | -2.72 | 1.000 |
| cluster3393 | 4.04 | 1.000 |
| cluster3955 | 4.10 | 1.000 |
| cluster4069 | 3.22 | 1.000 |
| cluster3516 | 4.33 | 1.000 |
| cluster2893 | 4.36 | 1.000 |
| cluster3840 | -2.41 | 1.000 |
| cluster3012 | 3.82 | 1.000 |
| cluster3182 | 3.14 | 1.000 |
| cluster3749 | 4.00 | 1.000 |
| cluster2876 | 3.25 | 1.000 |
| cluster4052 | -2.41 | 1.000 |
| cluster2960 | -2.21 | 1.000 |
| cluster3514 | -2.25 | 1.000 |
| cluster3689 | 4.07 | 1.000 |
| cluster4117 | 4.86 | 1.000 |
| cluster3694 | 2.76 | 1.000 |
| cluster3531 | 3.59 | 1.000 |
| cluster3597 | -2.11 | 1.000 |
| cluster3329 | 3.73 | 1.000 |
| cluster3905 | 4.18 | 1.000 |
| cluster3904 | 3.93 | 1.000 |
| cluster3994 | 3.74 | 1.000 |
| cluster3060 | 3.66 | 1.000 |
| cluster3674 | 3.91 | 1.000 |
| cluster3177 | 3.01 | 1.000 |
| cluster3715 | 3.05 | 1.000 |
| cluster3158 | 2.84 | 1.000 |
| cluster2914 | 2.80 | 1.000 |
| cluster3660 | 3.31 | 1.000 |
| cluster4103 | -2.34 | 1.000 |
| cluster3887 | 3.17 | 1.000 |
| cluster4055 | 3.31 | 1.000 |
| cluster3309 | -2.05 | 1.000 |
| cluster4013 | 5.14 | 1.000 |
| cluster3564 | -2.18 | 1.000 |
| cluster3599 | 3.34 | 1.000 |
| cluster3410 | -1.97 | 1.000 |
| cluster3652 | -1.99 | 1.000 |
| cluster3137 | -2.67 | 1.000 |
| cluster3672 | 3.23 | 1.000 |
| cluster3638 | 2.98 | 1.000 |
| cluster2980 | 3.39 | 1.000 |

| | | |
|---|---|---|
| cluster3561 | -2.12 | 1.000 |
| cluster3175 | 3.29 | 1.000 |
| cluster3615 | -2.09 | 1.000 |
| cluster3610 | 3.43 | 1.000 |
| cluster4118 | 4.56 | 1.000 |
| cluster2888 | 3.33 | 1.000 |
| cluster3138 | -1.92 | 1.000 |
| cluster3732 | 3.08 | 1.000 |
| cluster3125 | 2.45 | 1.000 |
| cluster3053 | -1.96 | 1.000 |
| cluster3102 | -1.93 | 1.000 |
| cluster2902 | -2.50 | 1.000 |
| cluster3237 | -2.59 | 1.000 |
| cluster4043 | 3.04 | 1.000 |
| cluster3551 | -1.92 | 1.000 |
| cluster2984 | -1.84 | 1.000 |
| cluster3721 | -2.00 | 1.000 |
| cluster3317 | -2.08 | 1.000 |
| cluster2860 | -1.81 | 1.000 |
| cluster3475 | 3.05 | 1.000 |
| cluster4058 | 3.08 | 1.000 |
| cluster4125 | -2.19 | 1.000 |
| cluster3129 | 3.17 | 1.000 |
| cluster3197 | -1.78 | 1.000 |
| cluster4080 | 2.49 | 1.000 |
| cluster3817 | -1.96 | 1.000 |
| cluster3538 | -1.79 | 1.000 |
| cluster2899 | -1.78 | 1.000 |
| cluster3714 | -1.88 | 1.000 |
| cluster2993 | -1.85 | 1.000 |
| cluster3105 | 3.63 | 1.000 |
| cluster3501 | 2.69 | 1.000 |
| cluster3898 | -2.01 | 1.000 |
| cluster3705 | -1.94 | 1.000 |
| cluster3645 | 3.71 | 1.000 |
| cluster3389 | 3.08 | 1.000 |
| cluster3650 | -1.80 | 1.000 |
| cluster3017 | -1.99 | 1.000 |
| cluster3764 | 3.21 | 1.000 |
| cluster2962 | -1.81 | 1.000 |
| cluster3858 | -1.87 | 1.000 |
| cluster3176 | 2.64 | 1.000 |
| cluster3500 | 3.39 | 1.000 |
| cluster4078 | -1.80 | 1.000 |

| | | |
|---|---|---|
| cluster3593 | 2.83 | 1.000 |
| cluster3659 | 3.05 | 1.000 |
| cluster3863 | -1.84 | 1.000 |
| cluster2933 | -1.73 | 1.000 |
| cluster3185 | 3.19 | 1.000 |
| cluster3426 | -1.73 | 1.000 |
| cluster3015 | -1.89 | 1.000 |
| cluster3061 | -1.60 | 1.000 |
| cluster3400 | -1.59 | 1.000 |
| cluster2911 | 2.12 | 1.000 |
| cluster4059 | 2.98 | 1.000 |
| cluster2861 | -1.66 | 1.000 |
| cluster3730 | -1.58 | 1.000 |
| cluster4128 | 3.38 | 1.000 |
| cluster3522 | 2.73 | 1.000 |
| cluster3948 | 2.81 | 1.000 |
| cluster3297 | -1.80 | 1.000 |
| cluster3565 | 2.71 | 1.000 |
| cluster3186 | 3.02 | 1.000 |
| cluster3644 | 2.14 | 1.000 |
| cluster4066 | 2.62 | 1.000 |
| cluster3386 | 2.87 | 1.000 |
| cluster3063 | -1.64 | 1.000 |
| cluster2989 | -1.69 | 1.000 |
| cluster3760 | -1.58 | 1.000 |
| cluster3337 | -1.63 | 1.000 |
| cluster3639 | 2.99 | 1.000 |
| cluster4048 | 3.15 | 1.000 |
| cluster3459 | 2.79 | 1.000 |
| cluster3670 | 2.23 | 1.000 |
| cluster3998 | 3.29 | 1.000 |
| cluster3975 | 2.68 | 1.000 |
| cluster3462 | 2.04 | 1.000 |
| cluster3831 | 2.66 | 1.000 |
| cluster3773 | 1.97 | 1.000 |
| cluster3240 | -1.60 | 1.000 |
| cluster3200 | -1.54 | 1.000 |
| cluster3515 | -1.93 | 1.000 |
| cluster3922 | 2.85 | 1.000 |
| cluster4089 | -2.21 | 1.000 |
| cluster3800 | 2.09 | 1.000 |
| cluster3894 | 2.64 | 1.000 |
| cluster3454 | 1.88 | 1.000 |
| cluster3669 | 2.61 | 1.000 |

| | | |
|---|---|---|
| cluster2921 | -1.66 | 1.000 |
| cluster2853 | -1.54 | 1.000 |
| cluster3494 | -1.54 | 1.000 |
| cluster3375 | 2.00 | 1.000 |
| cluster2946 | -1.54 | 1.000 |
| cluster4095 | -1.77 | 1.000 |
| cluster3346 | -1.54 | 1.000 |
| cluster3360 | 2.16 | 1.000 |
| cluster3924 | 2.44 | 1.000 |
| cluster3651 | 2.54 | 1.000 |
| cluster3101 | -1.47 | 1.000 |
| cluster3513 | -1.50 | 1.000 |
| cluster3777 | 2.29 | 1.000 |
| cluster3000 | 2.39 | 1.000 |
| cluster3731 | -1.62 | 1.000 |
| cluster3883 | -2.14 | 1.000 |
| cluster3480 | -1.84 | 1.000 |
| cluster3188 | -1.56 | 1.000 |
| cluster3839 | 2.53 | 1.000 |
| cluster3897 | -1.64 | 1.000 |
| cluster2978 | -1.47 | 1.000 |
| cluster3142 | -1.61 | 1.000 |
| cluster2961 | -1.48 | 1.000 |
| cluster4001 | 2.75 | 1.000 |
| cluster3977 | 2.68 | 1.000 |
| cluster3733 | 2.10 | 1.000 |
| cluster3827 | 2.23 | 1.000 |
| cluster3254 | 2.79 | 1.000 |
| cluster2943 | -1.46 | 1.000 |
| cluster2967 | -1.55 | 1.000 |
| cluster3598 | 2.37 | 1.000 |
| cluster3854 | 2.44 | 1.000 |
| cluster3032 | -1.42 | 1.000 |
| cluster2881 | -1.43 | 1.000 |
| cluster3127 | 1.86 | 1.000 |
| cluster3466 | 2.00 | 1.000 |
| cluster3088 | -1.55 | 1.000 |
| cluster3775 | 2.57 | 1.000 |
| cluster3165 | -1.85 | 1.000 |
| cluster3377 | 2.45 | 1.000 |
| cluster4030 | 2.55 | 1.000 |
| cluster3709 | -1.38 | 1.000 |
| cluster3318 | -1.46 | 1.000 |
| cluster3511 | 2.57 | 1.000 |

| | | |
|---|---|---|
| cluster3928 | 2.43 | 1.000 |
| cluster3497 | -1.34 | 1.000 |
| cluster3575 | -1.90 | 1.000 |
| cluster3119 | -1.43 | 1.000 |
| cluster3385 | 2.52 | 1.000 |
| cluster3908 | 2.16 | 1.000 |
| cluster2937 | 1.83 | 1.000 |
| cluster3118 | -1.39 | 1.000 |
| cluster3741 | -1.43 | 1.000 |
| cluster3296 | 2.76 | 1.000 |
| cluster3833 | 2.61 | 1.000 |
| cluster4012 | -1.77 | 1.000 |
| cluster3481 | 2.16 | 1.000 |
| cluster3341 | 2.49 | 1.000 |
| cluster3482 | 1.89 | 1.000 |
| cluster3051 | 2.40 | 1.000 |
| cluster3234 | 1.82 | 1.000 |
| cluster3572 | 2.47 | 1.000 |
| cluster4035 | -1.45 | 1.000 |
| cluster3157 | -1.30 | 1.000 |
| cluster3397 | 2.46 | 1.000 |
| cluster3569 | 2.31 | 1.000 |
| cluster3850 | 2.81 | 1.000 |
| cluster3144 | 2.47 | 1.000 |
| cluster3795 | 2.47 | 1.000 |
| cluster3225 | 2.35 | 1.000 |
| cluster3512 | 2.01 | 1.000 |
| cluster3299 | 2.37 | 1.000 |
| cluster3417 | -1.46 | 1.000 |
| cluster3229 | -1.36 | 1.000 |
| cluster2873 | -1.31 | 1.000 |
| cluster3245 | -1.81 | 1.000 |
| cluster3130 | 2.36 | 1.000 |
| cluster3399 | -1.45 | 1.000 |
| cluster3306 | 2.37 | 1.000 |
| cluster3220 | 1.82 | 1.000 |
| cluster2963 | -1.32 | 1.000 |
| cluster3991 | 2.23 | 1.000 |
| cluster3449 | -1.27 | 1.000 |
| cluster3766 | 2.56 | 1.000 |
| cluster3279 | 2.01 | 1.000 |
| cluster3778 | -1.32 | 1.000 |
| cluster3876 | -1.26 | 1.000 |
| cluster4068 | 2.21 | 1.000 |

| | | |
|---|---|---|
| cluster4071 | 2.39 | 1.000 |
| cluster3387 | 2.32 | 1.000 |
| cluster4088 | 2.42 | 1.000 |
| cluster3866 | 2.11 | 1.000 |
| cluster3794 | 2.28 | 1.000 |
| cluster3567 | -1.74 | 1.000 |
| cluster3109 | 2.31 | 1.000 |
| cluster3408 | -1.29 | 1.000 |
| cluster2906 | -1.28 | 1.000 |
| cluster3181 | -1.27 | 1.000 |
| cluster3788 | -1.22 | 1.000 |
| cluster3636 | 1.61 | 1.000 |
| cluster3023 | -1.30 | 1.000 |
| cluster3487 | 2.31 | 1.000 |
| cluster3693 | 2.14 | 1.000 |
| cluster3452 | 2.25 | 1.000 |
| cluster3342 | 2.14 | 1.000 |
| cluster3246 | -1.24 | 1.000 |
| cluster3468 | -1.24 | 1.000 |
| cluster4104 | 2.15 | 1.000 |
| cluster3563 | 1.66 | 1.000 |
| cluster3868 | 2.14 | 1.000 |
| cluster3485 | 2.48 | 1.000 |
| cluster3304 | -1.20 | 1.000 |
| cluster2939 | 1.75 | 1.000 |
| cluster3534 | 1.61 | 1.000 |
| cluster3875 | 1.77 | 1.000 |
| cluster3073 | 1.70 | 1.000 |
| cluster3909 | 2.08 | 1.000 |
| cluster3082 | 2.27 | 1.000 |
| cluster3704 | -1.23 | 1.000 |
| cluster3388 | 2.06 | 1.000 |
| cluster2947 | -1.21 | 1.000 |
| cluster3425 | 1.72 | 1.000 |
| cluster3402 | -1.25 | 1.000 |
| cluster3469 | -1.20 | 1.000 |
| cluster3478 | -1.23 | 1.000 |
| cluster4038 | 2.07 | 1.000 |
| cluster3925 | 2.40 | 1.000 |
| cluster3972 | 2.10 | 1.000 |
| cluster4085 | 1.65 | 1.000 |
| cluster3226 | 2.09 | 1.000 |
| cluster3812 | -1.38 | 1.000 |
| cluster3027 | -1.15 | 1.000 |

| | | |
|---|---|---|
| cluster3290 | 1.68 | 1.000 |
| cluster3937 | -1.32 | 1.000 |
| cluster3518 | 2.15 | 1.000 |
| cluster3932 | 2.00 | 1.000 |
| cluster3520 | -1.78 | 1.000 |
| cluster3574 | 2.21 | 1.000 |
| cluster3746 | -1.24 | 1.000 |
| cluster3219 | 1.76 | 1.000 |
| cluster3993 | 2.02 | 1.000 |
| cluster3092 | 1.69 | 1.000 |
| cluster3066 | -1.13 | 1.000 |
| cluster3014 | 2.15 | 1.000 |
| cluster3443 | 1.67 | 1.000 |
| cluster3792 | 1.76 | 1.000 |
| cluster3486 | -1.16 | 1.000 |
| cluster4079 | -1.21 | 1.000 |
| cluster3212 | 1.69 | 1.000 |
| cluster3984 | 2.00 | 1.000 |
| cluster3992 | -1.23 | 1.000 |
| cluster3954 | 2.14 | 1.000 |
| cluster3503 | 1.95 | 1.000 |
| cluster3656 | -1.15 | 1.000 |
| cluster3842 | -1.22 | 1.000 |
| cluster3470 | 1.54 | 1.000 |
| cluster3374 | 1.95 | 1.000 |
| cluster4083 | 2.01 | 1.000 |
| cluster3762 | -1.24 | 1.000 |
| cluster2900 | -1.14 | 1.000 |
| cluster4107 | 1.79 | 1.000 |
| cluster4053 | 2.04 | 1.000 |
| cluster3358 | -1.14 | 1.000 |
| cluster3990 | -1.13 | 1.000 |
| cluster4122 | 2.26 | 1.000 |
| cluster3437 | 1.59 | 1.000 |
| cluster3553 | 1.95 | 1.000 |
| cluster3566 | 1.95 | 1.000 |
| cluster3889 | 1.92 | 1.000 |
| cluster3834 | 2.04 | 1.000 |
| cluster3156 | 1.98 | 1.000 |
| cluster4050 | 1.95 | 1.000 |
| cluster3915 | 2.17 | 1.000 |
| cluster3989 | -1.19 | 1.000 |
| cluster3139 | 2.13 | 1.000 |
| cluster3519 | 2.14 | 1.000 |

| | | |
|---|---|---|
| cluster3249 | 1.57 | 1.000 |
| cluster3618 | -1.47 | 1.000 |
| cluster3963 | 1.98 | 1.000 |
| cluster3335 | -1.13 | 1.000 |
| cluster3525 | 1.48 | 1.000 |
| cluster4011 | 2.14 | 1.000 |
| cluster3311 | -1.10 | 1.000 |
| cluster3465 | -1.05 | 1.000 |
| cluster3090 | -1.37 | 1.000 |
| cluster3338 | 1.93 | 1.000 |
| cluster3891 | 1.55 | 1.000 |
| cluster4040 | 1.59 | 1.000 |
| cluster3089 | -1.06 | 1.000 |
| cluster3450 | 2.20 | 1.000 |
| cluster2866 | -1.36 | 1.000 |
| cluster3824 | 1.92 | 1.000 |
| cluster3959 | 1.86 | 1.000 |
| cluster3978 | 1.62 | 1.000 |
| cluster3825 | 1.94 | 1.000 |
| cluster3171 | -1.07 | 1.000 |
| cluster3929 | 2.04 | 1.000 |
| cluster3654 | 1.74 | 1.000 |
| cluster2905 | -1.01 | 1.000 |
| cluster2998 | 1.87 | 1.000 |
| cluster3270 | 1.87 | 1.000 |
| cluster3453 | 1.48 | 1.000 |
| cluster2871 | 1.43 | 1.000 |
| cluster3414 | -1.04 | 1.000 |
| cluster3045 | -1.03 | 1.000 |
| cluster2904 | -0.97 | 1.000 |
| cluster3049 | -1.01 | 1.000 |
| cluster2878 | -1.01 | 1.000 |
| cluster3821 | 1.54 | 1.000 |
| cluster3995 | 1.86 | 1.000 |
| cluster4087 | 1.82 | 1.000 |
| cluster2965 | -1.30 | 1.000 |
| cluster3953 | 1.72 | 1.000 |
| cluster3822 | -1.31 | 1.000 |
| cluster3327 | -1.03 | 1.000 |
| cluster2999 | 1.85 | 1.000 |
| cluster3767 | 1.83 | 1.000 |
| cluster3585 | 1.38 | 1.000 |
| cluster3582 | -0.96 | 1.000 |
| cluster3861 | 1.77 | 1.000 |

| | | |
|---|---|---|
| cluster4113 | 1.45 | 1.000 |
| cluster3277 | 1.65 | 1.000 |
| cluster4031 | -1.02 | 1.000 |
| cluster3971 | 1.78 | 1.000 |
| cluster3493 | -1.29 | 1.000 |
| cluster3700 | 1.75 | 1.000 |
| cluster3799 | -1.03 | 1.000 |
| cluster3244 | -0.98 | 1.000 |
| cluster3345 | 1.74 | 1.000 |
| cluster3681 | 1.75 | 1.000 |
| cluster3062 | 1.31 | 1.000 |
| cluster3756 | 1.74 | 1.000 |
| cluster3439 | 1.70 | 1.000 |
| cluster3735 | 1.40 | 1.000 |
| cluster3793 | 1.76 | 1.000 |
| cluster3243 | -0.95 | 1.000 |
| cluster3987 | -1.04 | 1.000 |
| cluster3019 | -1.15 | 1.000 |
| cluster3319 | 1.75 | 1.000 |
| cluster3322 | 1.72 | 1.000 |
| cluster3112 | 1.76 | 1.000 |
| cluster3964 | 1.98 | 1.000 |
| cluster3753 | 1.50 | 1.000 |
| cluster3463 | 1.71 | 1.000 |
| cluster3461 | 2.08 | 1.000 |
| cluster3974 | 1.99 | 1.000 |
| cluster4060 | -1.20 | 1.000 |
| cluster3071 | -0.93 | 1.000 |
| cluster3422 | -1.09 | 1.000 |
| cluster4057 | 1.69 | 1.000 |
| cluster3836 | 1.74 | 1.000 |
| cluster3268 | 1.72 | 1.000 |
| cluster3448 | 1.75 | 1.000 |
| cluster2916 | -0.98 | 1.000 |
| cluster4045 | 1.85 | 1.000 |
| cluster4014 | 1.74 | 1.000 |
| cluster3411 | 1.69 | 1.000 |
| cluster3568 | 1.69 | 1.000 |
| cluster3187 | 1.73 | 1.000 |
| cluster3745 | -1.18 | 1.000 |
| cluster3912 | 1.18 | 1.000 |
| cluster3588 | -1.08 | 1.000 |
| cluster3132 | -0.90 | 1.000 |
| cluster3570 | 1.70 | 1.000 |

| | | |
|---|---|---|
| cluster3395 | 1.28 | 1.000 |
| cluster4096 | 1.68 | 1.000 |
| cluster3668 | 1.65 | 1.000 |
| cluster3683 | 1.64 | 1.000 |
| cluster3141 | -0.88 | 1.000 |
| cluster3702 | 1.63 | 1.000 |
| cluster2882 | -0.88 | 1.000 |
| cluster3192 | -0.85 | 1.000 |
| cluster4049 | 1.93 | 1.000 |
| cluster2857 | 1.22 | 1.000 |
| cluster3042 | 1.69 | 1.000 |
| cluster3716 | 1.60 | 1.000 |
| cluster3657 | 1.63 | 1.000 |
| cluster3608 | 1.22 | 1.000 |
| cluster3855 | 1.54 | 1.000 |
| cluster3433 | -0.93 | 1.000 |
| cluster4033 | 1.56 | 1.000 |
| cluster3901 | 1.62 | 1.000 |
| cluster3421 | -1.09 | 1.000 |
| cluster3281 | 1.69 | 1.000 |
| cluster3933 | 1.60 | 1.000 |
| cluster3576 | 1.45 | 1.000 |
| cluster3269 | 1.62 | 1.000 |
| cluster4116 | 1.94 | 1.000 |
| cluster4010 | 1.57 | 1.000 |
| cluster3432 | 1.31 | 1.000 |
| cluster3471 | 1.66 | 1.000 |
| cluster2869 | -0.83 | 1.000 |
| cluster3967 | -0.96 | 1.000 |
| cluster3623 | 1.64 | 1.000 |
| cluster3541 | -0.84 | 1.000 |
| cluster3206 | 1.58 | 1.000 |
| cluster3706 | 1.56 | 1.000 |
| cluster3653 | -0.87 | 1.000 |
| cluster3856 | 1.66 | 1.000 |
| cluster3743 | 1.54 | 1.000 |
| cluster3613 | -0.88 | 1.000 |
| cluster3710 | 1.54 | 1.000 |
| cluster2870 | -0.91 | 1.000 |
| cluster4119 | 1.49 | 1.000 |
| cluster3926 | 1.60 | 1.000 |
| cluster2926 | -0.83 | 1.000 |
| cluster3805 | -1.07 | 1.000 |
| cluster3577 | -0.81 | 1.000 |

| | | |
|---|---|---|
| cluster3340 | -0.81 | 1.000 |
| cluster2887 | 1.54 | 1.000 |
| cluster3548 | -0.82 | 1.000 |
| cluster3079 | -0.83 | 1.000 |
| cluster3918 | -1.04 | 1.000 |
| cluster3013 | -0.79 | 1.000 |
| cluster2924 | -1.04 | 1.000 |
| cluster3405 | -0.87 | 1.000 |
| cluster3783 | 1.57 | 1.000 |
| cluster2982 | -0.86 | 1.000 |
| cluster4008 | 1.44 | 1.000 |
| cluster3166 | -0.77 | 1.000 |
| cluster3230 | -0.86 | 1.000 |
| cluster3558 | 1.49 | 1.000 |
| cluster3765 | 1.22 | 1.000 |
| cluster4042 | 1.55 | 1.000 |
| cluster3594 | 1.52 | 1.000 |
| cluster3174 | -1.02 | 1.000 |
| cluster3488 | 1.39 | 1.000 |
| cluster3213 | -0.80 | 1.000 |
| cluster3750 | 1.74 | 1.000 |
| cluster3111 | 1.42 | 1.000 |
| cluster4002 | 1.43 | 1.000 |
| cluster3446 | 1.53 | 1.000 |
| cluster2930 | -0.85 | 1.000 |
| cluster3869 | 1.42 | 1.000 |
| cluster3253 | 1.49 | 1.000 |
| cluster4027 | 1.50 | 1.000 |
| cluster3914 | 1.63 | 1.000 |
| cluster3097 | -0.79 | 1.000 |
| cluster3369 | -0.99 | 1.000 |
| cluster3740 | 1.45 | 1.000 |
| cluster4037 | 1.45 | 1.000 |
| cluster3379 | 1.22 | 1.000 |
| cluster3034 | 1.35 | 1.000 |
| cluster3532 | -0.86 | 1.000 |
| cluster3476 | 1.46 | 1.000 |
| cluster2908 | -0.77 | 1.000 |
| cluster3806 | 1.48 | 1.000 |
| cluster3761 | -1.00 | 1.000 |
| cluster3957 | 1.46 | 1.000 |
| cluster4100 | 1.40 | 1.000 |
| cluster3403 | 1.54 | 1.000 |
| cluster3857 | 1.48 | 1.000 |

| | | |
|---|---|---|
| cluster3787 | 1.47 | 1.000 |
| cluster3104 | 1.00 | 1.000 |
| cluster3542 | 1.45 | 1.000 |
| cluster3830 | -0.75 | 1.000 |
| cluster3289 | -0.78 | 1.000 |
| cluster3691 | -0.89 | 1.000 |
| cluster3055 | 1.16 | 1.000 |
| cluster3239 | -0.72 | 1.000 |
| cluster3769 | 1.10 | 1.000 |
| cluster4036 | 1.38 | 1.000 |
| cluster3587 | -0.99 | 1.000 |
| cluster3906 | 1.43 | 1.000 |
| cluster3549 | -0.74 | 1.000 |
| cluster4009 | 1.32 | 1.000 |
| cluster3748 | -0.75 | 1.000 |
| cluster3048 | 1.44 | 1.000 |
| cluster3099 | -0.76 | 1.000 |
| cluster3490 | 1.40 | 1.000 |
| cluster3398 | 1.20 | 1.000 |
| cluster3107 | -0.77 | 1.000 |
| cluster3864 | 1.41 | 1.000 |
| cluster3539 | 1.50 | 1.000 |
| cluster3348 | -0.80 | 1.000 |
| cluster3772 | 1.30 | 1.000 |
| cluster3982 | 1.39 | 1.000 |
| cluster3884 | 1.74 | 1.000 |
| cluster2863 | -0.69 | 1.000 |
| cluster2935 | -0.73 | 1.000 |
| cluster3504 | -0.77 | 1.000 |
| cluster3264 | 1.06 | 1.000 |
| cluster3527 | 1.41 | 1.000 |
| cluster3033 | 1.03 | 1.000 |
| cluster2964 | -0.70 | 1.000 |
| cluster3919 | 1.28 | 1.000 |
| cluster3686 | 1.44 | 1.000 |
| cluster3026 | 1.43 | 1.000 |
| cluster3255 | 1.42 | 1.000 |
| cluster3361 | 1.43 | 1.000 |
| cluster3223 | -0.76 | 1.000 |
| cluster3667 | 1.11 | 1.000 |
| cluster3595 | -0.72 | 1.000 |
| cluster3547 | -0.69 | 1.000 |
| cluster3093 | 1.16 | 1.000 |
| cluster3438 | 1.26 | 1.000 |

| | | |
|---|---|---|
| cluster3009 | 1.05 | 1.000 |
| cluster3697 | -0.70 | 1.000 |
| cluster3873 | -0.67 | 1.000 |
| cluster3011 | -0.70 | 1.000 |
| cluster3510 | 1.60 | 1.000 |
| cluster3320 | 1.41 | 1.000 |
| cluster2925 | -0.87 | 1.000 |
| cluster2950 | -0.69 | 1.000 |
| cluster3938 | 1.26 | 1.000 |
| cluster3328 | 0.98 | 1.000 |
| cluster2875 | -0.70 | 1.000 |
| cluster3602 | -0.71 | 1.000 |
| cluster3077 | -0.70 | 1.000 |
| cluster4044 | 1.30 | 1.000 |
| cluster3671 | 1.39 | 1.000 |
| cluster3483 | 1.36 | 1.000 |
| cluster3136 | -0.77 | 1.000 |
| cluster3407 | -0.67 | 1.000 |
| cluster3467 | 1.30 | 1.000 |
| cluster3429 | -0.68 | 1.000 |
| cluster3380 | 1.14 | 1.000 |
| cluster3472 | -0.66 | 1.000 |
| cluster3947 | 1.29 | 1.000 |
| cluster4022 | 1.52 | 1.000 |
| cluster2990 | 1.28 | 1.000 |
| cluster3456 | -0.67 | 1.000 |
| cluster4024 | 1.28 | 1.000 |
| cluster3404 | -0.63 | 1.000 |
| cluster4047 | 1.34 | 1.000 |
| cluster3068 | 0.94 | 1.000 |
| cluster3110 | -0.69 | 1.000 |
| cluster3134 | 0.98 | 1.000 |
| cluster3983 | 1.25 | 1.000 |
| cluster3902 | 1.30 | 1.000 |
| cluster3430 | 1.30 | 1.000 |
| cluster4025 | 1.00 | 1.000 |
| cluster3573 | 1.30 | 1.000 |
| cluster2954 | -0.64 | 1.000 |
| cluster3973 | 1.51 | 1.000 |
| cluster3851 | 1.34 | 1.000 |
| cluster3880 | -0.83 | 1.000 |
| cluster3633 | 0.96 | 1.000 |
| cluster3314 | -0.83 | 1.000 |
| cluster4067 | -0.83 | 1.000 |

| | | |
|---|---|---|
| cluster3911 | -0.81 | 1.000 |
| cluster2894 | 1.02 | 1.000 |
| cluster3498 | -0.78 | 1.000 |
| cluster3075 | 1.00 | 1.000 |
| cluster4073 | -0.78 | 1.000 |
| cluster4026 | 1.27 | 1.000 |
| cluster3877 | 0.96 | 1.000 |
| cluster3647 | 1.27 | 1.000 |
| cluster3031 | -0.62 | 1.000 |
| cluster3820 | 1.22 | 1.000 |
| cluster3885 | 1.25 | 1.000 |
| cluster4005 | 1.30 | 1.000 |
| cluster3265 | -0.77 | 1.000 |
| cluster3273 | -0.61 | 1.000 |
| cluster4081 | 1.21 | 1.000 |
| cluster3578 | 1.22 | 1.000 |
| cluster4039 | -0.68 | 1.000 |
| cluster3803 | 1.27 | 1.000 |
| cluster3596 | 1.15 | 1.000 |
| cluster2994 | -0.65 | 1.000 |
| cluster3362 | -0.61 | 1.000 |
| cluster3324 | 0.94 | 1.000 |
| cluster3121 | -0.61 | 1.000 |
| cluster3843 | 0.97 | 1.000 |
| cluster3495 | -0.75 | 1.000 |
| cluster3003 | -0.70 | 1.000 |
| cluster2974 | 0.89 | 1.000 |
| cluster3022 | 1.21 | 1.000 |
| cluster3418 | 1.20 | 1.000 |
| cluster3814 | -0.60 | 1.000 |
| cluster3231 | -0.77 | 1.000 |
| cluster3315 | -0.59 | 1.000 |
| cluster3545 | 1.20 | 1.000 |
| cluster3780 | 1.39 | 1.000 |
| cluster3394 | 0.90 | 1.000 |
| cluster3813 | 1.14 | 1.000 |
| cluster2956 | 0.93 | 1.000 |
| cluster3455 | -0.64 | 1.000 |
| cluster3979 | -0.59 | 1.000 |
| cluster3190 | 1.27 | 1.000 |
| cluster3081 | 0.87 | 1.000 |
| cluster3218 | -0.72 | 1.000 |
| cluster3095 | 0.88 | 1.000 |
| cluster3164 | 1.19 | 1.000 |

| | | |
|---|---|---|
| cluster3591 | 0.86 | 1.000 |
| cluster3557 | 1.13 | 1.000 |
| cluster3331 | 1.16 | 1.000 |
| cluster3581 | 1.04 | 1.000 |
| cluster3168 | -0.57 | 1.000 |
| cluster3923 | 1.13 | 1.000 |
| cluster2969 | -0.66 | 1.000 |
| cluster3758 | 0.99 | 1.000 |
| cluster3350 | 1.20 | 1.000 |
| cluster3211 | 1.15 | 1.000 |
| cluster2942 | -0.65 | 1.000 |
| cluster3666 | 1.14 | 1.000 |
| cluster3332 | -0.68 | 1.000 |
| cluster3810 | -0.57 | 1.000 |
| cluster3870 | 1.20 | 1.000 |
| cluster3785 | -0.66 | 1.000 |
| cluster3260 | 0.87 | 1.000 |
| cluster3046 | -0.59 | 1.000 |
| cluster3530 | -0.64 | 1.000 |
| cluster3172 | -0.54 | 1.000 |
| cluster3829 | -0.67 | 1.000 |
| cluster4090 | 1.24 | 1.000 |
| cluster3619 | 0.89 | 1.000 |
| cluster3736 | 1.17 | 1.000 |
| cluster2953 | -0.54 | 1.000 |
| cluster3209 | 1.16 | 1.000 |
| cluster3641 | -0.54 | 1.000 |
| cluster3477 | -0.58 | 1.000 |
| cluster3939 | 1.12 | 1.000 |
| cluster3517 | 0.79 | 1.000 |
| cluster3712 | -0.60 | 1.000 |
| cluster3717 | 0.83 | 1.000 |
| cluster3661 | -0.53 | 1.000 |
| cluster3895 | 1.12 | 1.000 |
| cluster4086 | -0.68 | 1.000 |
| cluster3707 | -0.67 | 1.000 |
| cluster3543 | -0.65 | 1.000 |
| cluster3811 | -0.50 | 1.000 |
| cluster3754 | 0.88 | 1.000 |
| cluster2886 | -0.53 | 1.000 |
| cluster3302 | 0.99 | 1.000 |
| cluster3747 | -0.58 | 1.000 |
| cluster3301 | -0.54 | 1.000 |
| cluster3334 | 0.89 | 1.000 |

| | | |
|---|---|---|
| cluster3195 | 0.82 | 1.000 |
| cluster2919 | -0.52 | 1.000 |
| cluster3819 | 1.19 | 1.000 |
| cluster3115 | -0.63 | 1.000 |
| cluster3621 | -0.67 | 1.000 |
| cluster3047 | 0.86 | 1.000 |
| cluster3198 | 1.10 | 1.000 |
| cluster3611 | 1.11 | 1.000 |
| cluster3316 | 1.05 | 1.000 |
| cluster3250 | -0.54 | 1.000 |
| cluster3999 | 1.09 | 1.000 |
| cluster3355 | 1.18 | 1.000 |
| cluster3359 | 1.11 | 1.000 |
| cluster3664 | 0.96 | 1.000 |
| cluster3085 | 1.09 | 1.000 |
| cluster3632 | -0.51 | 1.000 |
| cluster3606 | 1.08 | 1.000 |
| cluster3016 | 1.08 | 1.000 |
| cluster3203 | 1.08 | 1.000 |
| cluster3872 | -0.62 | 1.000 |
| cluster3774 | -0.50 | 1.000 |
| cluster2941 | 0.74 | 1.000 |
| cluster1127 | -0.61 | 1.000 |
| cluster3232 | -0.61 | 1.000 |
| cluster3149 | -0.49 | 1.000 |
| cluster3005 | -0.51 | 1.000 |
| cluster3649 | 0.83 | 1.000 |
| cluster3170 | -0.50 | 1.000 |
| cluster3002 | -0.49 | 1.000 |
| cluster3368 | 0.75 | 1.000 |
| cluster4094 | -0.66 | 1.000 |
| cluster3473 | -0.51 | 1.000 |
| cluster3751 | 0.98 | 1.000 |
| cluster3383 | 1.08 | 1.000 |
| cluster3021 | -0.47 | 1.000 |
| cluster2981 | -0.59 | 1.000 |
| cluster3224 | 0.83 | 1.000 |
| cluster3381 | -0.47 | 1.000 |
| cluster3282 | 0.97 | 1.000 |
| cluster4015 | 0.77 | 1.000 |
| cluster3907 | 1.07 | 1.000 |
| cluster3391 | 0.72 | 1.000 |
| cluster3150 | 1.01 | 1.000 |
| cluster3489 | 1.04 | 1.000 |

| | | |
|---|---|---|
| cluster3248 | -0.59 | 1.000 |
| cluster3886 | -0.57 | 1.000 |
| cluster3266 | -0.48 | 1.000 |
| cluster2884 | -0.51 | 1.000 |
| cluster3560 | 1.01 | 1.000 |
| cluster3646 | 1.09 | 1.000 |
| cluster3258 | -0.47 | 1.000 |
| cluster3968 | 0.90 | 1.000 |
| cluster3678 | 0.76 | 1.000 |
| cluster3720 | 1.00 | 1.000 |
| cluster3609 | -0.58 | 1.000 |
| cluster3106 | -0.44 | 1.000 |
| cluster3259 | 0.69 | 1.000 |
| cluster3479 | 0.99 | 1.000 |
| cluster3784 | -0.47 | 1.000 |
| cluster3056 | -0.44 | 1.000 |
| cluster4124 | 0.91 | 1.000 |
| cluster3535 | -0.55 | 1.000 |
| cluster3537 | -0.54 | 1.000 |
| cluster3113 | -0.49 | 1.000 |
| cluster3072 | -0.55 | 1.000 |
| cluster3583 | 0.87 | 1.000 |
| cluster3600 | -0.48 | 1.000 |
| cluster3300 | -0.47 | 1.000 |
| cluster3723 | 0.74 | 1.000 |
| cluster3624 | 0.99 | 1.000 |
| cluster3846 | 0.74 | 1.000 |
| cluster3291 | -0.44 | 1.000 |
| cluster4105 | 1.12 | 1.000 |
| cluster2890 | -0.41 | 1.000 |
| cluster3956 | 0.98 | 1.000 |
| cluster3215 | -0.42 | 1.000 |
| cluster3698 | -0.42 | 1.000 |
| cluster3555 | 1.04 | 1.000 |
| cluster3724 | 0.97 | 1.000 |
| cluster3867 | 0.68 | 1.000 |
| cluster3837 | 1.02 | 1.000 |
| cluster4097 | 1.10 | 1.000 |
| cluster2991 | 0.94 | 1.000 |
| cluster3940 | -0.42 | 1.000 |
| cluster4120 | 1.02 | 1.000 |
| cluster3804 | 1.00 | 1.000 |
| cluster3579 | -0.41 | 1.000 |
| cluster3035 | -0.39 | 1.000 |

| | | |
|---|---|---|
| cluster3210 | 0.85 | 1.000 |
| cluster3981 | 0.73 | 1.000 |
| cluster3356 | -0.42 | 1.000 |
| cluster3797 | -0.49 | 1.000 |
| cluster3445 | 0.71 | 1.000 |
| cluster3458 | -0.40 | 1.000 |
| cluster3424 | 0.86 | 1.000 |
| cluster4126 | -0.51 | 1.000 |
| cluster3626 | -0.41 | 1.000 |
| cluster2923 | -0.39 | 1.000 |
| cluster3474 | 0.94 | 1.000 |
| cluster4099 | 0.92 | 1.000 |
| cluster3708 | 1.06 | 1.000 |
| cluster3227 | -0.45 | 1.000 |
| cluster2880 | -0.48 | 1.000 |
| cluster3135 | 0.94 | 1.000 |
| cluster3457 | -0.48 | 1.000 |
| cluster3791 | 0.94 | 1.000 |
| cluster3070 | -0.40 | 1.000 |
| cluster3942 | 0.92 | 1.000 |
| cluster3571 | 0.79 | 1.000 |
| cluster3236 | 0.90 | 1.000 |
| cluster3779 | -0.40 | 1.000 |
| cluster2940 | 0.92 | 1.000 |
| cluster3499 | 0.90 | 1.000 |
| cluster2977 | -0.44 | 1.000 |
| cluster3147 | -0.40 | 1.000 |
| cluster4065 | 0.91 | 1.000 |
| cluster3630 | -0.36 | 1.000 |
| cluster3271 | 0.64 | 1.000 |
| cluster2944 | -0.43 | 1.000 |
| cluster3460 | -0.45 | 1.000 |
| cluster3544 | 0.62 | 1.000 |
| cluster3920 | 0.99 | 1.000 |
| cluster4029 | -0.44 | 1.000 |
| cluster3180 | 0.89 | 1.000 |
| cluster3004 | -0.37 | 1.000 |
| cluster3536 | -0.36 | 1.000 |
| cluster3199 | 0.72 | 1.000 |
| cluster2896 | 0.90 | 1.000 |
| cluster2959 | 0.67 | 1.000 |
| cluster3160 | 0.61 | 1.000 |
| cluster3966 | 0.86 | 1.000 |
| cluster3357 | -0.38 | 1.000 |

| | | |
|---|---|---|
| cluster3217 | -0.36 | 1.000 |
| cluster3122 | 0.63 | 1.000 |
| cluster3152 | 0.89 | 1.000 |
| cluster3427 | -0.39 | 1.000 |
| cluster3251 | 0.87 | 1.000 |
| cluster3396 | -0.37 | 1.000 |
| cluster3189 | -0.42 | 1.000 |
| cluster2901 | 0.89 | 1.000 |
| cluster3267 | -0.35 | 1.000 |
| cluster3208 | -0.34 | 1.000 |
| cluster3703 | -0.34 | 1.000 |
| cluster2951 | 0.88 | 1.000 |
| cluster3965 | -0.42 | 1.000 |
| cluster3796 | 0.83 | 1.000 |
| cluster3985 | -0.37 | 1.000 |
| cluster3734 | 0.84 | 1.000 |
| cluster3584 | 0.87 | 1.000 |
| cluster4077 | 0.92 | 1.000 |
| cluster3896 | 0.84 | 1.000 |
| cluster3143 | -0.34 | 1.000 |
| cluster2859 | 0.62 | 1.000 |
| cluster3401 | -0.36 | 1.000 |
| cluster2928 | 0.72 | 1.000 |
| cluster3321 | -0.38 | 1.000 |
| cluster3550 | -0.35 | 1.000 |
| cluster3676 | -0.39 | 1.000 |
| cluster3120 | 0.68 | 1.000 |
| cluster4091 | -0.44 | 1.000 |
| cluster3067 | -0.33 | 1.000 |
| cluster3286 | -0.31 | 1.000 |
| cluster3601 | 0.84 | 1.000 |
| cluster3913 | -0.41 | 1.000 |
| cluster2920 | -0.36 | 1.000 |
| cluster3349 | -0.33 | 1.000 |
| cluster3976 | 0.60 | 1.000 |
| cluster3305 | 0.85 | 1.000 |
| cluster3673 | 0.89 | 1.000 |
| cluster3083 | -0.31 | 1.000 |
| cluster3586 | -0.31 | 1.000 |
| cluster3351 | 0.66 | 1.000 |
| cluster3603 | -0.37 | 1.000 |
| cluster3502 | -0.36 | 1.000 |
| cluster3310 | 0.55 | 1.000 |
| cluster3124 | 0.61 | 1.000 |

| | | |
|---|---|---|
| cluster2995 | -0.38 | 1.000 |
| cluster3091 | 0.56 | 1.000 |
| cluster3526 | -0.33 | 1.000 |
| cluster3044 | -0.36 | 1.000 |
| cluster3943 | -0.36 | 1.000 |
| cluster3945 | -0.35 | 1.000 |
| cluster3727 | -0.35 | 1.000 |
| cluster2885 | -0.35 | 1.000 |
| cluster2972 | 0.59 | 1.000 |
| cluster3665 | 0.82 | 1.000 |
| cluster3629 | 0.80 | 1.000 |
| cluster4070 | 0.82 | 1.000 |
| cluster2979 | -0.33 | 1.000 |
| cluster3590 | 0.78 | 1.000 |
| cluster4019 | -0.34 | 1.000 |
| cluster3415 | -0.31 | 1.000 |
| cluster3554 | 0.80 | 1.000 |
| cluster3307 | 0.80 | 1.000 |
| cluster3435 | 0.83 | 1.000 |
| cluster3373 | -0.33 | 1.000 |
| cluster4109 | -0.35 | 1.000 |
| cluster2895 | -0.32 | 1.000 |
| cluster3354 | -0.30 | 1.000 |
| cluster3100 | -0.32 | 1.000 |
| cluster2986 | 0.80 | 1.000 |
| cluster3635 | -0.27 | 1.000 |
| cluster3372 | -0.27 | 1.000 |
| cluster3934 | 0.72 | 1.000 |
| cluster3807 | -0.28 | 1.000 |
| cluster3688 | 0.77 | 1.000 |
| cluster3737 | 0.53 | 1.000 |
| cluster3986 | 0.76 | 1.000 |
| cluster3496 | -0.28 | 1.000 |
| cluster3759 | -0.29 | 1.000 |
| cluster3679 | -0.28 | 1.000 |
| cluster3847 | 0.82 | 1.000 |
| cluster3205 | 0.57 | 1.000 |
| cluster2934 | 0.58 | 1.000 |
| cluster3988 | 0.59 | 1.000 |
| cluster3744 | 0.78 | 1.000 |
| cluster3434 | 0.59 | 1.000 |
| cluster3655 | 0.74 | 1.000 |
| cluster3849 | -0.32 | 1.000 |
| cluster2854 | -0.28 | 1.000 |

| | | |
|---|---|---|
| cluster3419 | -0.26 | 1.000 |
| cluster3339 | -0.26 | 1.000 |
| cluster4114 | 0.87 | 1.000 |
| cluster3815 | 0.74 | 1.000 |
| cluster2992 | -0.25 | 1.000 |
| cluster3153 | 0.65 | 1.000 |
| cluster3366 | -0.30 | 1.000 |
| cluster3057 | 0.57 | 1.000 |
| cluster3440 | 0.52 | 1.000 |
| cluster3900 | 0.85 | 1.000 |
| cluster4108 | -0.27 | 1.000 |
| cluster3879 | 0.70 | 1.000 |
| cluster3029 | 0.66 | 1.000 |
| cluster3326 | -0.24 | 1.000 |
| cluster4032 | -0.28 | 1.000 |
| cluster3133 | 0.71 | 1.000 |
| cluster3484 | -0.27 | 1.000 |
| cluster3508 | -0.29 | 1.000 |
| cluster3155 | -0.27 | 1.000 |
| cluster3261 | 0.70 | 1.000 |
| cluster2897 | -0.25 | 1.000 |
| cluster3841 | 0.74 | 1.000 |
| cluster3285 | -0.25 | 1.000 |
| cluster3275 | 0.71 | 1.000 |
| cluster3059 | -0.26 | 1.000 |
| cluster3620 | 0.56 | 1.000 |
| cluster3159 | 0.70 | 1.000 |
| cluster3222 | 0.45 | 1.000 |
| cluster2915 | 0.59 | 1.000 |
| cluster2907 | -0.25 | 1.000 |
| cluster3685 | -0.25 | 1.000 |
| cluster4054 | 0.69 | 1.000 |
| cluster3238 | -0.23 | 1.000 |
| cluster3447 | 0.68 | 1.000 |
| cluster3235 | -0.25 | 1.000 |
| cluster3390 | -0.22 | 1.000 |
| cluster3663 | 0.70 | 1.000 |
| cluster3028 | 0.68 | 1.000 |
| cluster3114 | 0.54 | 1.000 |
| cluster3193 | -0.23 | 1.000 |
| cluster3552 | 0.47 | 1.000 |
| cluster3464 | 0.67 | 1.000 |
| cluster3725 | 0.68 | 1.000 |
| cluster3444 | -0.21 | 1.000 |

| | | |
|---|---|---|
| cluster2949 | -0.21 | 1.000 |
| cluster3179 | -0.24 | 1.000 |
| cluster3944 | 0.67 | 1.000 |
| cluster3776 | -0.23 | 1.000 |
| cluster2918 | -0.22 | 1.000 |
| cluster3298 | 0.52 | 1.000 |
| cluster4000 | -0.24 | 1.000 |
| cluster3719 | -0.26 | 1.000 |
| cluster2936 | 0.67 | 1.000 |
| cluster3037 | 0.52 | 1.000 |
| cluster3038 | 0.69 | 1.000 |
| cluster3862 | 0.67 | 1.000 |
| cluster3625 | 0.65 | 1.000 |
| cluster3853 | 0.66 | 1.000 |
| cluster3052 | -0.21 | 1.000 |
| cluster3117 | 0.71 | 1.000 |
| cluster2856 | 0.69 | 1.000 |
| cluster3204 | -0.20 | 1.000 |
| cluster3713 | -0.22 | 1.000 |
| cluster4018 | -0.29 | 1.000 |
| cluster3996 | 0.65 | 1.000 |
| cluster2898 | -0.20 | 1.000 |
| cluster3040 | -0.19 | 1.000 |
| cluster3183 | 0.53 | 1.000 |
| cluster3423 | 0.68 | 1.000 |
| cluster3726 | 0.50 | 1.000 |
| cluster3148 | -0.18 | 1.000 |
| cluster3952 | 0.63 | 1.000 |
| cluster3451 | 0.75 | 1.000 |
| cluster3257 | -0.18 | 1.000 |
| cluster3970 | 0.61 | 1.000 |
| cluster3094 | -0.18 | 1.000 |
| cluster3025 | -0.17 | 1.000 |
| cluster3162 | 0.57 | 1.000 |
| cluster3728 | 0.45 | 1.000 |
| cluster3506 | 0.62 | 1.000 |
| cluster3729 | 0.62 | 1.000 |
| cluster2874 | 0.48 | 1.000 |
| cluster3086 | -0.17 | 1.000 |
| cluster2929 | -0.17 | 1.000 |
| cluster3509 | -0.17 | 1.000 |
| cluster3202 | -0.16 | 1.000 |
| cluster2903 | -0.16 | 1.000 |
| cluster3962 | 0.60 | 1.000 |

| | | |
|---|---|---|
| cluster3828 | 0.60 | 1.000 |
| cluster4021 | -0.21 | 1.000 |
| cluster3910 | 0.61 | 1.000 |
| cluster3108 | -0.15 | 1.000 |
| cluster3064 | -0.16 | 1.000 |
| cluster3826 | 0.57 | 1.000 |
| cluster3739 | -0.15 | 1.000 |
| cluster3131 | 0.62 | 1.000 |
| cluster3242 | -0.15 | 1.000 |
| cluster3336 | -0.15 | 1.000 |
| cluster4074 | 0.60 | 1.000 |
| cluster3521 | 0.57 | 1.000 |
| cluster3173 | -0.15 | 1.000 |
| cluster3123 | 0.43 | 1.000 |
| cluster2966 | 0.59 | 1.000 |
| cluster3276 | -0.14 | 1.000 |
| cluster2997 | 0.53 | 1.000 |
| cluster3024 | -0.14 | 1.000 |
| cluster3742 | 0.58 | 1.000 |
| cluster3616 | -0.14 | 1.000 |
| cluster3428 | 0.42 | 1.000 |
| cluster4041 | 0.66 | 1.000 |
| cluster3687 | 0.58 | 1.000 |
| cluster3930 | -0.15 | 1.000 |
| cluster4028 | -0.15 | 1.000 |
| cluster4006 | 0.58 | 1.000 |
| cluster3406 | 0.41 | 1.000 |
| cluster3161 | 0.58 | 1.000 |
| cluster3039 | -0.13 | 1.000 |
| cluster3648 | 0.39 | 1.000 |
| cluster3076 | 0.55 | 1.000 |
| cluster2987 | 0.38 | 1.000 |
| cluster3718 | 0.55 | 1.000 |
| cluster2862 | -0.12 | 1.000 |
| cluster3809 | 0.56 | 1.000 |
| cluster2917 | 0.57 | 1.000 |
| cluster3431 | 0.54 | 1.000 |
| cluster3838 | 0.54 | 1.000 |
| cluster2957 | 0.58 | 1.000 |
| cluster3330 | -0.12 | 1.000 |
| cluster3201 | -0.12 | 1.000 |
| cluster3835 | 0.55 | 1.000 |
| cluster2872 | -0.11 | 1.000 |
| cluster3252 | 0.59 | 1.000 |

| | | |
|---|---|---|
| cluster3523 | 0.54 | 1.000 |
| cluster3675 | 0.35 | 1.000 |
| cluster2855 | 0.57 | 1.000 |
| cluster3293 | 0.38 | 1.000 |
| cluster2958 | 0.40 | 1.000 |
| cluster3969 | 0.36 | 1.000 |
| cluster4062 | -0.09 | 1.000 |
| cluster3628 | 0.56 | 1.000 |
| cluster2877 | -0.11 | 1.000 |
| cluster3946 | 0.53 | 1.000 |
| cluster4102 | -0.16 | 1.000 |
| cluster3658 | -0.10 | 1.000 |
| cluster3058 | 0.35 | 1.000 |
| cluster2968 | 0.34 | 1.000 |
| cluster3344 | -0.10 | 1.000 |
| cluster3098 | 0.53 | 1.000 |
| cluster3899 | -0.10 | 1.000 |
| cluster3069 | -0.10 | 1.000 |
| cluster3140 | -0.10 | 1.000 |
| cluster3284 | -0.08 | 1.000 |
| cluster3871 | 0.50 | 1.000 |
| cluster3292 | 0.59 | 1.000 |
| cluster2971 | 0.33 | 1.000 |
| cluster3364 | 0.49 | 1.000 |
| cluster3997 | 0.50 | 1.000 |
| cluster3169 | -0.06 | 1.000 |
| cluster2865 | -0.08 | 1.000 |
| cluster3084 | -0.08 | 1.000 |
| cluster3823 | -0.08 | 1.000 |
| cluster3874 | -0.08 | 1.000 |
| cluster3436 | -0.07 | 1.000 |
| cluster3325 | -0.05 | 1.000 |
| cluster3529 | -0.05 | 1.000 |
| cluster4110 | 0.48 | 1.000 |
| cluster3376 | 0.33 | 1.000 |
| cluster3699 | 0.48 | 1.000 |
| cluster3020 | -0.03 | 1.000 |
| cluster4123 | 0.40 | 1.000 |
| cluster3241 | -0.06 | 1.000 |
| cluster4061 | -0.06 | 1.000 |
| cluster3087 | 0.40 | 1.000 |
| cluster2955 | -0.06 | 1.000 |
| cluster3492 | -0.04 | 1.000 |
| cluster3054 | 0.36 | 1.000 |

| | | |
|---|---|---|
| cluster3692 | -0.04 | 1.000 |
| cluster3540 | 0.48 | 1.000 |
| cluster3154 | 0.33 | 1.000 |
| cluster3065 | -0.05 | 1.000 |
| cluster4075 | -0.04 | 1.000 |
| cluster3194 | 0.34 | 1.000 |
| cluster3041 | 0.38 | 1.000 |
| cluster3662 | 0.47 | 1.000 |
| cluster3491 | 0.45 | 1.000 |
| cluster4063 | 0.46 | 1.000 |
| cluster4007 | 0.40 | 1.000 |
| cluster3559 | -0.01 | 1.000 |
| cluster3274 | -0.01 | 1.000 |
| cluster3711 | 0.31 | 1.000 |
| cluster3214 | 0.36 | 1.000 |
| cluster3384 | 0.33 | 1.000 |
| cluster3507 | -0.01 | 1.000 |
| cluster3010 | 0.35 | 1.000 |
| cluster3860 | -0.05 | 1.000 |
| cluster2927 | -0.01 | 1.000 |
| cluster4016 | -0.04 | 1.000 |
| cluster3808 | -0.01 | 1.000 |
| cluster3308 | 0.31 | 1.000 |
| cluster3643 | -0.02 | 1.000 |
| cluster3684 | -0.01 | 1.000 |
| cluster3287 | 0.34 | 1.000 |
| cluster3001 | 0.28 | 1.000 |
| cluster3941 | 0.28 | 1.000 |
| cluster3605 | 0.43 | 1.000 |
| cluster3961 | 0.43 | 1.000 |
| cluster3146 | -0.03 | 1.000 |
| cluster3580 | 0.40 | 1.000 |
| cluster3980 | 0.28 | 1.000 |
| cluster4004 | 0.44 | 1.000 |
| cluster2892 | -0.02 | 1.000 |
| cluster3080 | -0.01 | 1.000 |
| cluster2867 | 0.43 | 1.000 |
| cluster4084 | 0.30 | 1.000 |
| cluster2973 | 0.03 | 1.000 |
| cluster3382 | 0.28 | 1.000 |
| cluster3768 | 0.03 | 1.000 |
| cluster3288 | 0.39 | 1.000 |
| cluster2983 | 0.41 | 1.000 |
| cluster4115 | 0.47 | 1.000 |

| | | |
|---|---|---|
| cluster2858 | -0.01 | 1.000 |
| cluster3893 | 0.00 | 1.000 |
| cluster3562 | 0.39 | 1.000 |
| cluster3690 | 0.27 | 1.000 |
| cluster3951 | 0.40 | 1.000 |
| cluster4121 | 0.39 | 1.000 |
| cluster3680 | 0.38 | 1.000 |
| cluster3036 | 0.40 | 1.000 |
| cluster3163 | 0.40 | 1.000 |
| cluster2975 | 0.23 | 1.000 |
| cluster3184 | 0.29 | 1.000 |
| cluster3043 | 0.38 | 1.000 |
| cluster3604 | 0.41 | 1.000 |
| cluster3921 | 0.08 | 1.000 |
| cluster3781 | 0.30 | 1.000 |
| cluster2952 | 0.38 | 1.000 |
| cluster2945 | 0.10 | 1.000 |
| cluster3333 | 0.35 | 1.000 |
| cluster3145 | 0.25 | 1.000 |
| cluster2996 | 0.25 | 1.000 |
| cluster3167 | 0.36 | 1.000 |
| cluster4072 | 0.35 | 1.000 |
| cluster3528 | 0.04 | 1.000 |
| cluster3078 | 0.22 | 1.000 |
| cluster3363 | 0.34 | 1.000 |
| cluster3696 | 0.27 | 1.000 |
| cluster2909 | 0.21 | 1.000 |
| cluster3903 | 0.34 | 1.000 |
| cluster3343 | 0.12 | 1.000 |
| cluster2910 | 0.12 | 1.000 |
| cluster3936 | 0.33 | 1.000 |
| cluster3283 | 0.21 | 1.000 |
| cluster3409 | 0.32 | 1.000 |
| cluster3888 | 0.31 | 1.000 |
| cluster3859 | 0.13 | 1.000 |
| cluster3622 | 0.28 | 1.000 |
| cluster2883 | 0.05 | 1.000 |
| cluster2912 | 0.22 | 1.000 |
| cluster3642 | 0.33 | 1.000 |
| cluster3927 | 0.30 | 1.000 |
| cluster3865 | 0.30 | 1.000 |
| cluster3960 | 0.33 | 1.000 |
| cluster3018 | 0.14 | 1.000 |
| cluster3607 | 0.29 | 1.000 |

| | | |
|---|---|---|
| cluster4106 | 0.41 | 1.000 |
| cluster3950 | 0.19 | 1.000 |
| cluster3592 | 0.29 | 1.000 |
| cluster3782 | 0.26 | 1.000 |
| cluster3196 | 0.07 | 1.000 |
| cluster3378 | 0.22 | 1.000 |
| cluster3637 | 0.17 | 1.000 |
| cluster4003 | 0.29 | 1.000 |
| cluster3771 | 0.28 | 1.000 |
| cluster3442 | 0.29 | 1.000 |
| cluster3008 | 0.23 | 1.000 |
| cluster4023 | 0.25 | 1.000 |
| cluster3074 | 0.18 | 1.000 |
| cluster2938 | 0.15 | 1.000 |
| cluster2879 | 0.16 | 1.000 |
| cluster3294 | 0.16 | 1.000 |
| cluster3151 | 0.15 | 1.000 |
| cluster2970 | 0.19 | 1.000 |
| cluster2932 | 0.18 | 1.000 |
| cluster3420 | 0.25 | 1.000 |
| cluster3116 | 0.20 | 1.000 |
| cluster3798 | 0.23 | 1.000 |
| cluster3278 | 0.12 | 1.000 |
| cluster2889 | 0.25 | 1.000 |
| cluster3845 | 0.11 | 1.000 |
| cluster2922 | 0.22 | 1.000 |
| cluster3533 | 0.16 | 1.000 |
| cluster3370 | 0.22 | 1.000 |
| cluster3096 | 0.14 | 1.000 |
| cluster2891 | 0.24 | 1.000 |
| cluster4127 | 0.31 | 1.000 |
| cluster3682 | 0.22 | 1.000 |
| cluster3612 | 0.21 | 1.000 |
| cluster3413 | 0.20 | 1.000 |
| cluster3789 | 0.20 | 1.000 |
| cluster4017 | 0.20 | 1.000 |
| cluster3935 | 0.19 | 1.000 |
| cluster3801 | 0.19 | 1.000 |
| cluster3882 | 0.19 | 1.000 |
| cluster4093 | 0.19 | 1.000 |
| cluster3917 | 0.18 | 1.000 |
| cluster4082 | 0.18 | 1.000 |
| cluster3614 | 0.18 | 1.000 |
| cluster3790 | 0.17 | 1.000 |

| | | |
|---|---|---|
| cluster3832 | 0.17 | 1.000 |
| cluster3816 | 0.17 | 1.000 |
| cluster3352 | 0.16 | 1.000 |
| cluster3007 | 0.15 | 1.000 |
| cluster3701 | 0.15 | 1.000 |
| cluster3848 | 0.14 | 1.000 |
| cluster3050 | 0.14 | 1.000 |
| cluster2913 | 0.14 | 1.000 |
| cluster3627 | 0.14 | 1.000 |
| cluster3207 | 0.13 | 1.000 |
| cluster4051 | 0.13 | 1.000 |
| cluster3634 | 0.11 | 1.000 |
| cluster2931 | 0.11 | 1.000 |
| cluster3365 | 0.11 | 1.000 |
| cluster3347 | 0.10 | 1.000 |
| cluster3524 | 0.10 | 1.000 |
| cluster3412 | 0.10 | 1.000 |
| cluster3128 | 0.09 | 1.000 |
| cluster3556 | 0.08 | 1.000 |
| cluster3216 | 0.08 | 1.000 |
| cluster4020 | 0.08 | 1.000 |
| cluster3722 | 0.08 | 1.000 |
| cluster3103 | 0.07 | 1.000 |
| cluster3221 | 0.07 | 1.000 |
| cluster3617 | 0.07 | 1.000 |
| cluster3505 | 0.06 | 1.000 |
| cluster3295 | 0.06 | 1.000 |
| cluster4064 | 0.06 | 1.000 |
| cluster3441 | 0.06 | 1.000 |
| cluster3262 | 0.05 | 1.000 |
| cluster3881 | 0.05 | 1.000 |
| cluster3280 | 0.05 | 1.000 |
| cluster3247 | 0.04 | 1.000 |
| cluster4092 | 0.04 | 1.000 |
| cluster2985 | 0.03 | 1.000 |
| cluster3228 | 0.02 | 1.000 |
| cluster3353 | 0.02 | 1.000 |
| cluster2864 | 0.02 | 1.000 |
| cluster3256 | 0.02 | 1.000 |
| cluster3589 | 0.01 | 1.000 |
| cluster3263 | 0.01 | 1.000 |
| cluster3677 | 0.01 | 1.000 |
| cluster3949 | -0.01 | 1.000 |
| cluster2988 | 0.00 | 1.000 |

| | | |
|---|---|---|
| cluster3916 | 0.00 | 1.000 |

**Supplementary Table 19**. codeml results - neutral evolution

| ID | Likelihood | ω |
|---|---|---|
| cluster1127 | -1524.314086 | 1 |
| cluster2853 | -26297.98509 | 1 |
| cluster2854 | -1823.396454 | 1 |
| cluster2855 | -28264.89144 | 1 |
| cluster2856 | -16358.6295 | 1 |
| cluster2857 | -6489.531419 | 1 |
| cluster2858 | -9849.038871 | 1 |
| cluster2859 | -6339.28655 | 1 |
| cluster2860 | -20633.4996 | 1 |
| cluster2861 | -10259.3456 | 1 |
| cluster2862 | -22986.43971 | 1 |
| cluster2863 | -17640.95872 | 1 |
| cluster2864 | -16338.84605 | 1 |
| cluster2865 | -16282.73794 | 1 |
| cluster2866 | -20206.90772 | 1 |
| cluster2867 | -16862.41354 | 1 |
| cluster2869 | -9699.066574 | 1 |
| cluster2870 | -4295.481897 | 1 |
| cluster2871 | -3526.338088 | 1 |
| cluster2872 | -13322.46144 | 1 |
| cluster2873 | -16751.01282 | 1 |
| cluster2874 | -11861.6533 | 1 |
| cluster2875 | -16633.57696 | 1 |
| cluster2876 | -4590.941261 | 1 |
| cluster2877 | -12088.1886 | 1 |
| cluster2878 | -7925.783223 | 1 |
| cluster2879 | -10442.65959 | 1 |
| cluster2880 | -11960.30565 | 1 |
| cluster2881 | -12951.93757 | 1 |
| cluster2882 | -12337.02688 | 1 |
| cluster2883 | -7398.725577 | 1 |
| cluster2884 | -14073.68015 | 1 |
| cluster2885 | -17972.278 | 1 |
| cluster2886 | -13699.40506 | 1 |
| cluster2887 | -10554.73157 | 1 |
| cluster2888 | -8490.220832 | 1 |
| cluster2889 | -8763.937974 | 1 |
| cluster2890 | -2816.941236 | 1 |
| cluster2891 | -13787.68498 | 1 |
| cluster2892 | -9571.776506 | 1 |
| cluster2893 | -9550.898584 | 1 |
| cluster2894 | -6457.004582 | 1 |

| | | |
|---|---|---|
| cluster2895 | -9683.067648 | 1 |
| cluster2896 | -12951.70093 | 1 |
| cluster2897 | -10564.81856 | 1 |
| cluster2898 | -5278.578869 | 1 |
| cluster2899 | -9538.990375 | 1 |
| cluster2900 | -11911.58061 | 1 |
| cluster2901 | -10584.97814 | 1 |
| cluster2902 | -4376.693279 | 1 |
| cluster2903 | -10898.00686 | 1 |
| cluster2904 | -6887.37304 | 1 |
| cluster2905 | -10148.25 | 1 |
| cluster2906 | -15082.86919 | 1 |
| cluster2907 | -13847.50695 | 1 |
| cluster2908 | -6497.386787 | 1 |
| cluster2909 | -3244.478368 | 1 |
| cluster2910 | -10611.70225 | 1 |
| cluster2911 | -4602.305096 | 1 |
| cluster2912 | -10197.08489 | 1 |
| cluster2913 | -7932.213273 | 1 |
| cluster2914 | -9469.138523 | 1 |
| cluster2915 | -5092.104513 | 1 |
| cluster2916 | -11360.9011 | 1 |
| cluster2917 | -10659.60257 | 1 |
| cluster2918 | -4853.372182 | 1 |
| cluster2919 | -10627.12727 | 1 |
| cluster2920 | -8406.11369 | 1 |
| cluster2921 | -4834.192903 | 1 |
| cluster2922 | -6960.304228 | 1 |
| cluster2923 | -6070.824911 | 1 |
| cluster2924 | -12865.3995 | 1 |
| cluster2925 | -6653.496777 | 1 |
| cluster2926 | -13368.98387 | 1 |
| cluster2927 | -5609.237027 | 1 |
| cluster2928 | -8501.208246 | 1 |
| cluster2929 | -10097.22358 | 1 |
| cluster2930 | -11937.44951 | 1 |
| cluster2931 | -7645.200583 | 1 |
| cluster2932 | -10731.74598 | 1 |
| cluster2933 | -9802.593284 | 1 |
| cluster2934 | -11582.45636 | 1 |
| cluster2935 | -5756.102101 | 1 |
| cluster2936 | -10997.72482 | 1 |
| cluster2937 | -5322.614479 | 1 |
| cluster2938 | -8690.749489 | 1 |

| | | |
|---|---|---|
| cluster2939 | -3199.876577 | 1 |
| cluster2940 | -12616.58933 | 1 |
| cluster2941 | -3427.723594 | 1 |
| cluster2942 | -12684.63677 | 1 |
| cluster2943 | -5017.718835 | 1 |
| cluster2944 | -9094.485131 | 1 |
| cluster2945 | -3895.800741 | 1 |
| cluster2946 | -7721.365859 | 1 |
| cluster2947 | -11053.09848 | 1 |
| cluster2948 | -10555.59127 | 1 |
| cluster2949 | -8060.924528 | 1 |
| cluster2950 | -8478.240869 | 1 |
| cluster2951 | -10004.1563 | 1 |
| cluster2952 | -12409.38104 | 1 |
| cluster2953 | -14615.89207 | 1 |
| cluster2954 | -8397.526022 | 1 |
| cluster2955 | -6965.016748 | 1 |
| cluster2956 | -12101.39389 | 1 |
| cluster2957 | -9013.348525 | 1 |
| cluster2958 | -11866.32407 | 1 |
| cluster2959 | -5807.896094 | 1 |
| cluster2960 | -11289.75259 | 1 |
| cluster2961 | -10408.24816 | 1 |
| cluster2962 | -8723.055805 | 1 |
| cluster2963 | -8200.012854 | 1 |
| cluster2964 | -6061.820096 | 1 |
| cluster2965 | -8935.050802 | 1 |
| cluster2966 | -6569.391078 | 1 |
| cluster2967 | -11544.87057 | 1 |
| cluster2968 | -6534.59103 | 1 |
| cluster2969 | -8982.292625 | 1 |
| cluster2970 | -5385.141039 | 1 |
| cluster2971 | -3986.279095 | 1 |
| cluster2972 | -3335.841683 | 1 |
| cluster2973 | -6869.156621 | 1 |
| cluster2974 | -4978.618334 | 1 |
| cluster2975 | -11291.44498 | 1 |
| cluster2976 | -1508.183727 | 1 |
| cluster2977 | -4500.489787 | 1 |
| cluster2978 | -10429.36493 | 1 |
| cluster2979 | -10091.35084 | 1 |
| cluster2980 | -1803.0107 | 1 |
| cluster2981 | -6984.666567 | 1 |
| cluster2982 | -8584.306347 | 1 |

| | | |
|---|---|---|
| cluster2983 | -8913.905385 | 1 |
| cluster2984 | -5574.247474 | 1 |
| cluster2985 | -4079.73646 | 1 |
| cluster2986 | -3719.60801 | 1 |
| cluster2987 | -6950.198048 | 1 |
| cluster2988 | -6746.204106 | 1 |
| cluster2989 | -7474.86903 | 1 |
| cluster2990 | -8492.687771 | 1 |
| cluster2991 | -6937.363446 | 1 |
| cluster2992 | -4851.00895 | 1 |
| cluster2993 | -6353.563307 | 1 |
| cluster2994 | -2716.734209 | 1 |
| cluster2995 | -805.323526 | 1 |
| cluster2996 | -8356.320249 | 1 |
| cluster2997 | -9054.854379 | 1 |
| cluster2998 | -8918.855105 | 1 |
| cluster2999 | -7759.797318 | 1 |
| cluster3000 | -7375.848378 | 1 |
| cluster3001 | -7012.178794 | 1 |
| cluster3002 | -8946.860776 | 1 |
| cluster3003 | -8431.829304 | 1 |
| cluster3004 | -6434.019323 | 1 |
| cluster3005 | -4488.631037 | 1 |
| cluster3006 | -9528.082136 | 1 |
| cluster3007 | -9613.651911 | 1 |
| cluster3008 | -6923.582684 | 1 |
| cluster3009 | -6613.509501 | 1 |
| cluster3010 | -8420.901357 | 1 |
| cluster3011 | -4110.966422 | 1 |
| cluster3012 | -5764.609369 | 1 |
| cluster3013 | -7319.959785 | 1 |
| cluster3014 | -4506.468906 | 1 |
| cluster3015 | -2682.68679 | 1 |
| cluster3016 | -5139.43897 | 1 |
| cluster3017 | -7052.586815 | 1 |
| cluster3018 | -7041.265231 | 1 |
| cluster3019 | -8454.456615 | 1 |
| cluster3020 | -7126.410687 | 1 |
| cluster3021 | -3553.013181 | 1 |
| cluster3022 | -3750.693708 | 1 |
| cluster3023 | -5102.629715 | 1 |
| cluster3024 | -2106.839224 | 1 |
| cluster3025 | -5426.141466 | 1 |
| cluster3026 | -4246.589327 | 1 |

| | | |
|---|---|---|
| cluster3027 | -4486.128295 | 1 |
| cluster3028 | -7664.55557 | 1 |
| cluster3029 | -7629.186813 | 1 |
| cluster3030 | -5864.596816 | 1 |
| cluster3031 | -9486.159636 | 1 |
| cluster3032 | -7046.325226 | 1 |
| cluster3033 | -3951.878082 | 1 |
| cluster3034 | -5254.641512 | 1 |
| cluster3035 | -9010.226121 | 1 |
| cluster3036 | -9750.088071 | 1 |
| cluster3037 | -2936.356417 | 1 |
| cluster3038 | -4971.716149 | 1 |
| cluster3039 | -7331.085478 | 1 |
| cluster3040 | -5635.433392 | 1 |
| cluster3041 | -9532.091041 | 1 |
| cluster3042 | -5095.154742 | 1 |
| cluster3043 | -3387.421312 | 1 |
| cluster3044 | -7428.669819 | 1 |
| cluster3045 | -6891.909769 | 1 |
| cluster3046 | -7560.127205 | 1 |
| cluster3047 | -2523.885758 | 1 |
| cluster3048 | -6347.562164 | 1 |
| cluster3049 | -8550.100404 | 1 |
| cluster3050 | -8560.538111 | 1 |
| cluster3051 | -8490.956568 | 1 |
| cluster3052 | -7745.06137 | 1 |
| cluster3053 | -10124.83853 | 1 |
| cluster3054 | -8387.044354 | 1 |
| cluster3055 | -4511.113371 | 1 |
| cluster3056 | -6437.193528 | 1 |
| cluster3057 | -1006.262388 | 1 |
| cluster3058 | -2856.165439 | 1 |
| cluster3059 | -9363.315499 | 1 |
| cluster3060 | -6070.252315 | 1 |
| cluster3061 | -6714.593606 | 1 |
| cluster3062 | -6385.619472 | 1 |
| cluster3063 | -7130.376809 | 1 |
| cluster3064 | -2189.990897 | 1 |
| cluster3065 | -2464.517539 | 1 |
| cluster3066 | -6824.823806 | 1 |
| cluster3067 | -5607.36759 | 1 |
| cluster3068 | -5457.463746 | 1 |
| cluster3069 | -1830.64844 | 1 |
| cluster3070 | -5986.017108 | 1 |

| | | |
|---|---|---|
| cluster3071 | -6840.682114 | 1 |
| cluster3072 | -8031.261306 | 1 |
| cluster3073 | -4276.334332 | 1 |
| cluster3074 | -8621.707211 | 1 |
| cluster3075 | -6222.478876 | 1 |
| cluster3076 | -2014.409734 | 1 |
| cluster3077 | -5210.448303 | 1 |
| cluster3078 | -5907.381979 | 1 |
| cluster3079 | -8825.051906 | 1 |
| cluster3080 | -6719.06214 | 1 |
| cluster3081 | -4702.336821 | 1 |
| cluster3082 | -7972.898114 | 1 |
| cluster3083 | -4803.143697 | 1 |
| cluster3084 | -3450.551115 | 1 |
| cluster3085 | -8542.186592 | 1 |
| cluster3086 | -3658.598737 | 1 |
| cluster3087 | -7702.724722 | 1 |
| cluster3088 | -7079.624598 | 1 |
| cluster3089 | -4027.291695 | 1 |
| cluster3090 | -6144.374922 | 1 |
| cluster3091 | -6493.840648 | 1 |
| cluster3092 | -5015.416936 | 1 |
| cluster3093 | -5372.692041 | 1 |
| cluster3094 | -9121.622161 | 1 |
| cluster3095 | -6017.970565 | 1 |
| cluster3096 | -7035.320615 | 1 |
| cluster3097 | -8201.887874 | 1 |
| cluster3098 | -1805.960324 | 1 |
| cluster3099 | -4845.468746 | 1 |
| cluster3100 | -4993.778362 | 1 |
| cluster3101 | -3924.232302 | 1 |
| cluster3102 | -2741.204068 | 1 |
| cluster3103 | -5864.329889 | 1 |
| cluster3104 | -5113.58973 | 1 |
| cluster3105 | -9086.806043 | 1 |
| cluster3106 | -6220.730719 | 1 |
| cluster3107 | -3485.346697 | 1 |
| cluster3108 | -4568.798026 | 1 |
| cluster3109 | -4058.579833 | 1 |
| cluster3110 | -6804.484982 | 1 |
| cluster3111 | -5597.4291 | 1 |
| cluster3112 | -6869.290448 | 1 |
| cluster3113 | -3600.399459 | 1 |
| cluster3114 | -5333.773222 | 1 |

| | | |
|---|---|---|
| cluster3115 | -4483.655797 | 1 |
| cluster3116 | -3099.441415 | 1 |
| cluster3117 | -8418.97713 | 1 |
| cluster3118 | -1721.732241 | 1 |
| cluster3119 | -3349.953979 | 1 |
| cluster3120 | -5770.038166 | 1 |
| cluster3121 | -7909.705323 | 1 |
| cluster3122 | -4128.579121 | 1 |
| cluster3123 | -4559.263142 | 1 |
| cluster3124 | -7284.255593 | 1 |
| cluster3125 | -5219.908249 | 1 |
| cluster3126 | -8600.332465 | 1 |
| cluster3127 | -3063.196922 | 1 |
| cluster3128 | -2685.137735 | 1 |
| cluster3129 | -3640.31009 | 1 |
| cluster3130 | -6668.26099 | 1 |
| cluster3131 | -2171.286404 | 1 |
| cluster3132 | -6169.163773 | 1 |
| cluster3133 | -9705.431169 | 1 |
| cluster3134 | -3649.132522 | 1 |
| cluster3135 | -5092.30801 | 1 |
| cluster3136 | -4514.272345 | 1 |
| cluster3137 | -3661.404925 | 1 |
| cluster3138 | -5332.680285 | 1 |
| cluster3139 | -6270.138607 | 1 |
| cluster3140 | -3941.287265 | 1 |
| cluster3141 | -2822.874234 | 1 |
| cluster3142 | -1717.814577 | 1 |
| cluster3143 | -3816.321373 | 1 |
| cluster3144 | -2571.230287 | 1 |
| cluster3145 | -8398.113501 | 1 |
| cluster3146 | -2214.427834 | 1 |
| cluster3147 | -5578.256111 | 1 |
| cluster3148 | -6908.821399 | 1 |
| cluster3149 | -2823.981858 | 1 |
| cluster3150 | -4912.08475 | 1 |
| cluster3151 | -4401.488172 | 1 |
| cluster3152 | -9774.103617 | 1 |
| cluster3153 | -5092.893022 | 1 |
| cluster3154 | -6558.327398 | 1 |
| cluster3155 | -3725.544075 | 1 |
| cluster3156 | -7806.631511 | 1 |
| cluster3157 | -3480.402154 | 1 |
| cluster3158 | -4088.246854 | 1 |

| | | |
|---|---|---|
| cluster3159 | -6532.827667 | 1 |
| cluster3160 | -1175.296784 | 1 |
| cluster3161 | -5676.001035 | 1 |
| cluster3162 | -4229.849973 | 1 |
| cluster3163 | -3920.39354 | 1 |
| cluster3164 | -6872.347191 | 1 |
| cluster3165 | -8923.484175 | 1 |
| cluster3166 | -5472.199551 | 1 |
| cluster3167 | -5733.97602 | 1 |
| cluster3168 | -8546.318604 | 1 |
| cluster3169 | -5843.559145 | 1 |
| cluster3170 | -1759.355814 | 1 |
| cluster3171 | -6765.938426 | 1 |
| cluster3172 | -7264.272082 | 1 |
| cluster3173 | -5408.418495 | 1 |
| cluster3174 | -1625.802562 | 1 |
| cluster3175 | -2263.134561 | 1 |
| cluster3176 | -4367.135086 | 1 |
| cluster3177 | -3811.110862 | 1 |
| cluster3178 | -8435.835718 | 1 |
| cluster3179 | -5049.734478 | 1 |
| cluster3180 | -1325.572378 | 1 |
| cluster3181 | -7970.134474 | 1 |
| cluster3182 | -2978.268973 | 1 |
| cluster3183 | -4396.922109 | 1 |
| cluster3184 | -4766.651626 | 1 |
| cluster3185 | -4333.342558 | 1 |
| cluster3186 | -2620.591782 | 1 |
| cluster3187 | -6347.605921 | 1 |
| cluster3188 | -4784.025151 | 1 |
| cluster3189 | -6991.3721 | 1 |
| cluster3190 | -6228.71686 | 1 |
| cluster3191 | -3140.714589 | 1 |
| cluster3192 | -6603.335163 | 1 |
| cluster3193 | -3884.86505 | 1 |
| cluster3194 | -7713.031477 | 1 |
| cluster3195 | -4786.425558 | 1 |
| cluster3196 | -2564.150691 | 1 |
| cluster3197 | -6026.412608 | 1 |
| cluster3198 | -7563.261365 | 1 |
| cluster3199 | -3539.683917 | 1 |
| cluster3200 | -5174.374312 | 1 |
| cluster3201 | -4767.17141 | 1 |
| cluster3202 | -6829.769706 | 1 |

| | | |
|---|---|---|
| cluster3203 | -3328.27846 | 1 |
| cluster3204 | -2070.514326 | 1 |
| cluster3205 | -2289.367188 | 1 |
| cluster3206 | -4489.317032 | 1 |
| cluster3207 | -6691.743393 | 1 |
| cluster3208 | -3808.121479 | 1 |
| cluster3209 | -6570.415508 | 1 |
| cluster3210 | -4225.778458 | 1 |
| cluster3211 | -3973.650398 | 1 |
| cluster3212 | -3384.829628 | 1 |
| cluster3213 | -4998.9965 | 1 |
| cluster3214 | -6848.037884 | 1 |
| cluster3215 | -2534.847971 | 1 |
| cluster3216 | -5187.676554 | 1 |
| cluster3217 | -4315.975345 | 1 |
| cluster3218 | -5897.915078 | 1 |
| cluster3219 | -6578.759216 | 1 |
| cluster3220 | -3295.409625 | 1 |
| cluster3221 | -6593.597741 | 1 |
| cluster3222 | -2962.562011 | 1 |
| cluster3223 | -3730.669798 | 1 |
| cluster3224 | -1864.565801 | 1 |
| cluster3225 | -7095.676315 | 1 |
| cluster3226 | -4238.782786 | 1 |
| cluster3227 | -4850.995277 | 1 |
| cluster3228 | -4185.532623 | 1 |
| cluster3229 | -4419.031496 | 1 |
| cluster3230 | -4173.809368 | 1 |
| cluster3231 | -5210.977388 | 1 |
| cluster3232 | -6003.860555 | 1 |
| cluster3233 | -3917.292195 | 1 |
| cluster3234 | -4291.237962 | 1 |
| cluster3235 | -8152.062066 | 1 |
| cluster3236 | -4617.85971 | 1 |
| cluster3237 | -3596.299426 | 1 |
| cluster3238 | -3012.79538 | 1 |
| cluster3239 | -3066.903532 | 1 |
| cluster3240 | -8094.038409 | 1 |
| cluster3241 | -6352.19888 | 1 |
| cluster3242 | -7905.37256 | 1 |
| cluster3243 | -6261.744429 | 1 |
| cluster3244 | -7237.293599 | 1 |
| cluster3245 | -3341.1364 | 1 |
| cluster3246 | -5850.237697 | 1 |

| | | |
|---|---|---|
| cluster3247 | -938.36095 | 1 |
| cluster3248 | -4124.739161 | 1 |
| cluster3249 | -4155.284507 | 1 |
| cluster3250 | -3777.699492 | 1 |
| cluster3251 | -4992.855528 | 1 |
| cluster3252 | -8115.39408 | 1 |
| cluster3253 | -4729.73221 | 1 |
| cluster3254 | -3597.349108 | 1 |
| cluster3255 | -6615.346463 | 1 |
| cluster3256 | -6953.821427 | 1 |
| cluster3257 | -6386.907103 | 1 |
| cluster3258 | -2284.898721 | 1 |
| cluster3259 | -3941.050231 | 1 |
| cluster3260 | -5376.568098 | 1 |
| cluster3261 | -5386.667119 | 1 |
| cluster3262 | -6915.507681 | 1 |
| cluster3263 | -2163.265129 | 1 |
| cluster3264 | -2104.246771 | 1 |
| cluster3265 | -5276.696636 | 1 |
| cluster3266 | -3160.204567 | 1 |
| cluster3267 | -4751.834456 | 1 |
| cluster3268 | -3203.509542 | 1 |
| cluster3269 | -3919.90752 | 1 |
| cluster3270 | -2502.425356 | 1 |
| cluster3271 | -4904.456114 | 1 |
| cluster3272 | -954.696178 | 1 |
| cluster3273 | -4273.077035 | 1 |
| cluster3274 | -2320.944888 | 1 |
| cluster3275 | -5123.690146 | 1 |
| cluster3276 | -2654.037954 | 1 |
| cluster3277 | -3566.798961 | 1 |
| cluster3278 | -5446.451009 | 1 |
| cluster3279 | -4421.908491 | 1 |
| cluster3280 | -4405.641691 | 1 |
| cluster3281 | -2328.591702 | 1 |
| cluster3282 | -2522.070317 | 1 |
| cluster3283 | -6040.873129 | 1 |
| cluster3284 | -6886.325885 | 1 |
| cluster3285 | -2735.693069 | 1 |
| cluster3286 | -2437.347322 | 1 |
| cluster3287 | -1643.625195 | 1 |
| cluster3288 | -1549.941093 | 1 |
| cluster3289 | -6492.886359 | 1 |
| cluster3290 | -1423.608181 | 1 |

| | | |
|---|---|---|
| cluster3291 | -5505.11104 | 1 |
| cluster3292 | -3153.341863 | 1 |
| cluster3293 | -3969.528496 | 1 |
| cluster3294 | -3405.442257 | 1 |
| cluster3295 | -2703.096663 | 1 |
| cluster3296 | -2211.245572 | 1 |
| cluster3297 | -2452.292101 | 1 |
| cluster3298 | -3751.102749 | 1 |
| cluster3299 | -8541.73761 | 1 |
| cluster3300 | -2828.777469 | 1 |
| cluster3301 | -2011.176052 | 1 |
| cluster3302 | -4973.986293 | 1 |
| cluster3303 | -5159.752439 | 1 |
| cluster3304 | -3252.728891 | 1 |
| cluster3305 | -4182.373127 | 1 |
| cluster3306 | -5486.775332 | 1 |
| cluster3307 | -2109.91393 | 1 |
| cluster3308 | -5276.370332 | 1 |
| cluster3309 | -3585.321781 | 1 |
| cluster3310 | -4951.165525 | 1 |
| cluster3311 | -6881.139017 | 1 |
| cluster3312 | -2307.10966 | 1 |
| cluster3313 | -2899.037634 | 1 |
| cluster3314 | -7957.907889 | 1 |
| cluster3315 | -4616.272984 | 1 |
| cluster3316 | -6462.192198 | 1 |
| cluster3317 | -1390.037044 | 1 |
| cluster3318 | -3632.263112 | 1 |
| cluster3319 | -4346.638127 | 1 |
| cluster3320 | -3117.292377 | 1 |
| cluster3321 | -3406.472756 | 1 |
| cluster3322 | -3724.981588 | 1 |
| cluster3323 | -6200.01207 | 1 |
| cluster3324 | -1580.906332 | 1 |
| cluster3325 | -2437.164425 | 1 |
| cluster3326 | -1814.621311 | 1 |
| cluster3327 | -4542.685351 | 1 |
| cluster3328 | -2978.04502 | 1 |
| cluster3329 | -3497.13588 | 1 |
| cluster3330 | -5160.797517 | 1 |
| cluster3331 | -3112.158093 | 1 |
| cluster3332 | -6876.253698 | 1 |
| cluster3333 | -4178.133382 | 1 |
| cluster3334 | -1491.275058 | 1 |

| | | |
|---|---|---|
| cluster3335 | -3328.393388 | 1 |
| cluster3336 | -3638.615461 | 1 |
| cluster3337 | -2950.585632 | 1 |
| cluster3338 | -7132.678106 | 1 |
| cluster3339 | -2167.504304 | 1 |
| cluster3340 | -1668.758376 | 1 |
| cluster3341 | -1917.041507 | 1 |
| cluster3342 | -3007.65802 | 1 |
| cluster3343 | -3774.57432 | 1 |
| cluster3344 | -3611.072008 | 1 |
| cluster3345 | -3444.867971 | 1 |
| cluster3346 | -4525.019081 | 1 |
| cluster3347 | -4676.597589 | 1 |
| cluster3348 | -5039.745381 | 1 |
| cluster3349 | -5052.074642 | 1 |
| cluster3350 | -5371.639413 | 1 |
| cluster3351 | -3696.92221 | 1 |
| cluster3352 | -4492.565901 | 1 |
| cluster3353 | -1843.504688 | 1 |
| cluster3354 | -2461.351974 | 1 |
| cluster3355 | -4844.761535 | 1 |
| cluster3356 | -5198.979209 | 1 |
| cluster3357 | -1486.892748 | 1 |
| cluster3358 | -5215.868037 | 1 |
| cluster3359 | -6897.80825 | 1 |
| cluster3360 | -2802.760001 | 1 |
| cluster3361 | -5648.921548 | 1 |
| cluster3362 | -4431.550841 | 1 |
| cluster3363 | -4891.587942 | 1 |
| cluster3364 | -4997.621427 | 1 |
| cluster3365 | -6203.699668 | 1 |
| cluster3366 | -3572.204667 | 1 |
| cluster3367 | -3916.59437 | 1 |
| cluster3368 | -1846.231115 | 1 |
| cluster3369 | -4540.527708 | 1 |
| cluster3370 | -6854.113849 | 1 |
| cluster3371 | -3490.157757 | 1 |
| cluster3372 | -4657.183724 | 1 |
| cluster3373 | -6639.246827 | 1 |
| cluster3374 | -5592.949525 | 1 |
| cluster3375 | -3546.846797 | 1 |
| cluster3376 | -1671.661213 | 1 |
| cluster3377 | -2859.429136 | 1 |
| cluster3378 | -7338.046124 | 1 |

| | | |
|---|---|---|
| cluster3379 | -1787.370322 | 1 |
| cluster3380 | -3029.556263 | 1 |
| cluster3381 | -2582.585428 | 1 |
| cluster3382 | -1001.52788 | 1 |
| cluster3383 | -5692.490156 | 1 |
| cluster3384 | -2235.739212 | 1 |
| cluster3385 | -4450.450002 | 1 |
| cluster3386 | -5067.292897 | 1 |
| cluster3387 | -3443.750849 | 1 |
| cluster3388 | -3290.24606 | 1 |
| cluster3389 | -2168.86349 | 1 |
| cluster3390 | -5628.899409 | 1 |
| cluster3391 | -2915.928064 | 1 |
| cluster3392 | -3069.174911 | 1 |
| cluster3393 | -3494.361701 | 1 |
| cluster3394 | -909.821363 | 1 |
| cluster3395 | -3286.701882 | 1 |
| cluster3396 | -5741.318503 | 1 |
| cluster3397 | -5450.658603 | 1 |
| cluster3398 | -4246.672025 | 1 |
| cluster3399 | -3665.947496 | 1 |
| cluster3400 | -2042.310742 | 1 |
| cluster3401 | -3331.691818 | 1 |
| cluster3402 | -5952.050351 | 1 |
| cluster3403 | -4278.402863 | 1 |
| cluster3404 | -778.880522 | 1 |
| cluster3405 | -1644.366325 | 1 |
| cluster3406 | -4533.981219 | 1 |
| cluster3407 | -6895.024268 | 1 |
| cluster3408 | -2201.065962 | 1 |
| cluster3409 | -5715.427243 | 1 |
| cluster3410 | -1640.686076 | 1 |
| cluster3411 | -3194.890246 | 1 |
| cluster3412 | -3674.43689 | 1 |
| cluster3413 | -2882.860133 | 1 |
| cluster3414 | -2632.768539 | 1 |
| cluster3415 | -3860.672238 | 1 |
| cluster3416 | -677.089105 | 1 |
| cluster3417 | -1554.479328 | 1 |
| cluster3418 | -2809.285065 | 1 |
| cluster3419 | -2809.149219 | 1 |
| cluster3420 | -1837.309704 | 1 |
| cluster3421 | -5130.583656 | 1 |
| cluster3422 | -6229.906099 | 1 |

| | | |
|---|---|---|
| cluster3423 | -4784.256861 | 1 |
| cluster3424 | -3062.690981 | 1 |
| cluster3425 | -2444.245065 | 1 |
| cluster3426 | -4121.832627 | 1 |
| cluster3427 | -4092.972691 | 1 |
| cluster3428 | -2957.960079 | 1 |
| cluster3429 | -4845.141209 | 1 |
| cluster3430 | -5424.374543 | 1 |
| cluster3431 | -5382.898891 | 1 |
| cluster3432 | -4290.934754 | 1 |
| cluster3433 | -5630.364774 | 1 |
| cluster3434 | -3727.157664 | 1 |
| cluster3435 | -4861.307042 | 1 |
| cluster3436 | -3721.93349 | 1 |
| cluster3437 | -3450.459117 | 1 |
| cluster3438 | -4354.34239 | 1 |
| cluster3439 | -2261.029795 | 1 |
| cluster3440 | -2638.118919 | 1 |
| cluster3441 | -3104.870165 | 1 |
| cluster3442 | -3359.054419 | 1 |
| cluster3443 | -4175.940783 | 1 |
| cluster3444 | -4061.674064 | 1 |
| cluster3445 | -3513.956517 | 1 |
| cluster3446 | -4727.989508 | 1 |
| cluster3447 | -6694.235626 | 1 |
| cluster3448 | -2770.864358 | 1 |
| cluster3449 | -687.785076 | 1 |
| cluster3450 | -2977.329469 | 1 |
| cluster3451 | -4165.456852 | 1 |
| cluster3452 | -4431.253385 | 1 |
| cluster3453 | -2078.019566 | 1 |
| cluster3454 | -4438.470063 | 1 |
| cluster3455 | -4014.306695 | 1 |
| cluster3456 | -4773.822238 | 1 |
| cluster3457 | -4212.90792 | 1 |
| cluster3458 | -2032.356782 | 1 |
| cluster3459 | -2968.815908 | 1 |
| cluster3460 | -6372.273822 | 1 |
| cluster3461 | -3390.735145 | 1 |
| cluster3462 | -875.913409 | 1 |
| cluster3463 | -5351.58204 | 1 |
| cluster3464 | -6282.638183 | 1 |
| cluster3465 | -4793.862817 | 1 |
| cluster3466 | -3220.619357 | 1 |

| | | |
|---|---|---|
| cluster3467 | -4650.258016 | 1 |
| cluster3468 | -2289.761398 | 1 |
| cluster3469 | -2489.851776 | 1 |
| cluster3470 | -4657.706088 | 1 |
| cluster3471 | -4726.393985 | 1 |
| cluster3472 | -4457.052059 | 1 |
| cluster3473 | -4054.958498 | 1 |
| cluster3474 | -6082.088159 | 1 |
| cluster3475 | -2759.653022 | 1 |
| cluster3476 | -5847.264051 | 1 |
| cluster3477 | -3735.230022 | 1 |
| cluster3478 | -4241.801722 | 1 |
| cluster3479 | -3213.102988 | 1 |
| cluster3480 | -5162.145449 | 1 |
| cluster3481 | -1235.597465 | 1 |
| cluster3482 | -2097.804323 | 1 |
| cluster3483 | -4478.875696 | 1 |
| cluster3484 | -3225.863775 | 1 |
| cluster3485 | -3887.676289 | 1 |
| cluster3486 | -3502.628248 | 1 |
| cluster3487 | -2708.876956 | 1 |
| cluster3488 | -1691.975043 | 1 |
| cluster3489 | -4574.48861 | 1 |
| cluster3490 | -4038.763319 | 1 |
| cluster3491 | -4223.77799 | 1 |
| cluster3492 | -3860.214292 | 1 |
| cluster3493 | -5080.50543 | 1 |
| cluster3494 | -3388.59821 | 1 |
| cluster3495 | -3287.275142 | 1 |
| cluster3496 | -3794.980448 | 1 |
| cluster3497 | -1315.649511 | 1 |
| cluster3498 | -4611.688246 | 1 |
| cluster3499 | -3905.321338 | 1 |
| cluster3500 | -3918.432916 | 1 |
| cluster3501 | -2699.309182 | 1 |
| cluster3502 | -2603.526303 | 1 |
| cluster3503 | -2162.368625 | 1 |
| cluster3504 | -1420.625787 | 1 |
| cluster3505 | -2715.862249 | 1 |
| cluster3506 | -3049.000407 | 1 |
| cluster3507 | -5627.517594 | 1 |
| cluster3508 | -6363.336872 | 1 |
| cluster3509 | -2674.056013 | 1 |
| cluster3510 | -884.084676 | 1 |

| | | |
|---|---|---|
| cluster3511 | -1129.225375 | 1 |
| cluster3512 | -2653.560059 | 1 |
| cluster3513 | -996.342645 | 1 |
| cluster3514 | -5918.005447 | 1 |
| cluster3515 | -3137.058405 | 1 |
| cluster3516 | -4802.04333 | 1 |
| cluster3517 | -2874.183263 | 1 |
| cluster3518 | -5200.46946 | 1 |
| cluster3519 | -2414.448942 | 1 |
| cluster3520 | -1249.123227 | 1 |
| cluster3521 | -4798.127534 | 1 |
| cluster3522 | -5411.692241 | 1 |
| cluster3523 | -1908.688005 | 1 |
| cluster3524 | -2269.432123 | 1 |
| cluster3525 | -3127.001192 | 1 |
| cluster3526 | -3857.840558 | 1 |
| cluster3527 | -5527.315027 | 1 |
| cluster3528 | -2322.862267 | 1 |
| cluster3529 | -4309.55884 | 1 |
| cluster3530 | -2230.582828 | 1 |
| cluster3531 | -3282.916761 | 1 |
| cluster3532 | -6133.979496 | 1 |
| cluster3533 | -3043.564152 | 1 |
| cluster3534 | -2749.261706 | 1 |
| cluster3535 | -3834.043666 | 1 |
| cluster3536 | -4077.612964 | 1 |
| cluster3537 | -4620.740492 | 1 |
| cluster3538 | -1543.132085 | 1 |
| cluster3539 | -3316.674335 | 1 |
| cluster3540 | -5757.924244 | 1 |
| cluster3541 | -4496.527304 | 1 |
| cluster3542 | -5954.027559 | 1 |
| cluster3543 | -4825.31229 | 1 |
| cluster3544 | -1226.482378 | 1 |
| cluster3545 | -5810.657949 | 1 |
| cluster3546 | -3093.847809 | 1 |
| cluster3547 | -1254.010514 | 1 |
| cluster3548 | -1403.460504 | 1 |
| cluster3549 | -3711.992792 | 1 |
| cluster3550 | -1378.494835 | 1 |
| cluster3551 | -3055.657677 | 1 |
| cluster3552 | -2783.261463 | 1 |
| cluster3553 | -2792.076758 | 1 |
| cluster3554 | -4403.497313 | 1 |

| | | |
|---|---|---|
| cluster3555 | -1931.413553 | 1 |
| cluster3556 | -4943.27839 | 1 |
| cluster3557 | -1461.773039 | 1 |
| cluster3558 | -5291.691371 | 1 |
| cluster3559 | -2870.857463 | 1 |
| cluster3560 | -5659.89408 | 1 |
| cluster3561 | -2015.358357 | 1 |
| cluster3562 | -4908.573423 | 1 |
| cluster3563 | -2332.638622 | 1 |
| cluster3564 | -5551.351901 | 1 |
| cluster3565 | -3840.637555 | 1 |
| cluster3566 | -2128.982404 | 1 |
| cluster3567 | -2603.733327 | 1 |
| cluster3568 | -3613.349401 | 1 |
| cluster3569 | -3974.008006 | 1 |
| cluster3570 | -2770.214584 | 1 |
| cluster3571 | -2379.850532 | 1 |
| cluster3572 | -3856.610822 | 1 |
| cluster3573 | -3825.169563 | 1 |
| cluster3574 | -5615.966646 | 1 |
| cluster3575 | -4365.792603 | 1 |
| cluster3576 | -3706.279853 | 1 |
| cluster3577 | -3077.56445 | 1 |
| cluster3578 | -3860.675259 | 1 |
| cluster3579 | -3369.796867 | 1 |
| cluster3580 | -2934.106452 | 1 |
| cluster3581 | -1523.156552 | 1 |
| cluster3582 | -1224.487839 | 1 |
| cluster3583 | -3213.646822 | 1 |
| cluster3584 | -5515.320056 | 1 |
| cluster3585 | -2287.646132 | 1 |
| cluster3586 | -5718.122704 | 1 |
| cluster3587 | -3829.905502 | 1 |
| cluster3588 | -3814.686139 | 1 |
| cluster3589 | -3419.548369 | 1 |
| cluster3590 | -4541.308336 | 1 |
| cluster3591 | -3396.039896 | 1 |
| cluster3592 | -5358.220299 | 1 |
| cluster3593 | -3880.749839 | 1 |
| cluster3594 | -3491.323991 | 1 |
| cluster3595 | -2517.900114 | 1 |
| cluster3596 | -2668.605497 | 1 |
| cluster3597 | -4855.43119 | 1 |
| cluster3598 | -4775.34061 | 1 |

| | | |
|---|---|---|
| cluster3599 | -2642.59101 | 1 |
| cluster3600 | -3293.804338 | 1 |
| cluster3601 | -4061.784336 | 1 |
| cluster3602 | -3699.563333 | 1 |
| cluster3603 | -4779.747973 | 1 |
| cluster3604 | -1113.55334 | 1 |
| cluster3605 | -3473.745032 | 1 |
| cluster3606 | -5041.561365 | 1 |
| cluster3607 | -2958.794437 | 1 |
| cluster3608 | -3457.289264 | 1 |
| cluster3609 | -4766.74025 | 1 |
| cluster3610 | -1615.150115 | 1 |
| cluster3611 | -2643.836685 | 1 |
| cluster3612 | -3240.244773 | 1 |
| cluster3613 | -4858.281314 | 1 |
| cluster3614 | -4894.063386 | 1 |
| cluster3615 | -1811.783256 | 1 |
| cluster3616 | -891.736599 | 1 |
| cluster3617 | -2154.468784 | 1 |
| cluster3618 | -1961.333239 | 1 |
| cluster3619 | -3184.672529 | 1 |
| cluster3620 | -4074.53872 | 1 |
| cluster3621 | -5502.770522 | 1 |
| cluster3622 | -3215.989232 | 1 |
| cluster3623 | -2698.722606 | 1 |
| cluster3624 | -3429.961691 | 1 |
| cluster3625 | -4055.737858 | 1 |
| cluster3626 | -2308.616755 | 1 |
| cluster3627 | -3767.513023 | 1 |
| cluster3628 | -3480.93176 | 1 |
| cluster3629 | -5135.449289 | 1 |
| cluster3630 | -1381.591497 | 1 |
| cluster3631 | -4294.029828 | 1 |
| cluster3632 | -2837.830573 | 1 |
| cluster3633 | -2856.652562 | 1 |
| cluster3634 | -1074.004657 | 1 |
| cluster3635 | -1757.624914 | 1 |
| cluster3636 | -3012.41062 | 1 |
| cluster3637 | -1185.933039 | 1 |
| cluster3638 | -3091.779349 | 1 |
| cluster3639 | -1438.420868 | 1 |
| cluster3640 | -5162.040154 | 1 |
| cluster3641 | -2001.38979 | 1 |
| cluster3642 | -3820.162283 | 1 |

| | | |
|---|---|---|
| cluster3643 | -3794.444165 | 1 |
| cluster3644 | -2167.42982 | 1 |
| cluster3645 | -2104.19114 | 1 |
| cluster3646 | -3055.402991 | 1 |
| cluster3647 | -4362.114017 | 1 |
| cluster3648 | -2293.777563 | 1 |
| cluster3649 | -1929.996679 | 1 |
| cluster3650 | -1436.934179 | 1 |
| cluster3651 | -4448.570897 | 1 |
| cluster3652 | -3904.111832 | 1 |
| cluster3653 | -1764.985972 | 1 |
| cluster3654 | -3509.57842 | 1 |
| cluster3655 | -2592.759535 | 1 |
| cluster3656 | -3653.586181 | 1 |
| cluster3657 | -3770.069082 | 1 |
| cluster3658 | -797.469243 | 1 |
| cluster3659 | -2565.701943 | 1 |
| cluster3660 | -3570.290053 | 1 |
| cluster3661 | -1984.722431 | 1 |
| cluster3662 | -3209.894813 | 1 |
| cluster3663 | -3495.195436 | 1 |
| cluster3664 | -2761.765103 | 1 |
| cluster3665 | -4307.747421 | 1 |
| cluster3666 | -3092.227028 | 1 |
| cluster3667 | -1769.156256 | 1 |
| cluster3668 | -3547.714103 | 1 |
| cluster3669 | -2950.605381 | 1 |
| cluster3670 | -4579.258098 | 1 |
| cluster3671 | -4599.145472 | 1 |
| cluster3672 | -2819.48664 | 1 |
| cluster3673 | -4818.623774 | 1 |
| cluster3674 | -2298.260652 | 1 |
| cluster3675 | -756.410946 | 1 |
| cluster3676 | -3928.440711 | 1 |
| cluster3677 | -924.670284 | 1 |
| cluster3678 | -3788.214562 | 1 |
| cluster3679 | -1121.458774 | 1 |
| cluster3680 | -2865.491655 | 1 |
| cluster3681 | -3487.590546 | 1 |
| cluster3682 | -5055.197081 | 1 |
| cluster3683 | -2983.105976 | 1 |
| cluster3684 | -3475.90809 | 1 |
| cluster3685 | -3964.06114 | 1 |
| cluster3686 | -1791.901926 | 1 |

| | | |
|---|---|---|
| cluster3687 | -3036.083488 | 1 |
| cluster3688 | -2503.603271 | 1 |
| cluster3689 | -3002.75353 | 1 |
| cluster3690 | -1697.163757 | 1 |
| cluster3691 | -817.968958 | 1 |
| cluster3692 | -1770.412784 | 1 |
| cluster3693 | -3112.851544 | 1 |
| cluster3694 | -2302.019414 | 1 |
| cluster3695 | -2034.864992 | 1 |
| cluster3696 | -946.990973 | 1 |
| cluster3697 | -3910.236412 | 1 |
| cluster3698 | -1297.334758 | 1 |
| cluster3699 | -5051.04228 | 1 |
| cluster3700 | -3244.126822 | 1 |
| cluster3701 | -1368.272492 | 1 |
| cluster3702 | -4457.424285 | 1 |
| cluster3703 | -1745.916737 | 1 |
| cluster3704 | -2809.022275 | 1 |
| cluster3705 | -2334.097902 | 1 |
| cluster3706 | -2451.014148 | 1 |
| cluster3707 | -2163.329842 | 1 |
| cluster3708 | -2901.168193 | 1 |
| cluster3709 | -1701.001309 | 1 |
| cluster3710 | -2296.396628 | 1 |
| cluster3711 | -1558.334672 | 1 |
| cluster3712 | -3959.680902 | 1 |
| cluster3713 | -4499.888533 | 1 |
| cluster3714 | -1960.246504 | 1 |
| cluster3715 | -1525.125445 | 1 |
| cluster3716 | -4128.524189 | 1 |
| cluster3717 | -1594.359257 | 1 |
| cluster3718 | -3544.685494 | 1 |
| cluster3719 | -3697.85769 | 1 |
| cluster3720 | -3168.131405 | 1 |
| cluster3721 | -1229.434569 | 1 |
| cluster3722 | -1344.619184 | 1 |
| cluster3723 | -1192.808897 | 1 |
| cluster3724 | -3575.676996 | 1 |
| cluster3725 | -2266.043754 | 1 |
| cluster3726 | -2176.264212 | 1 |
| cluster3727 | -2736.371965 | 1 |
| cluster3728 | -2452.633493 | 1 |
| cluster3729 | -3514.870765 | 1 |
| cluster3730 | -1924.990564 | 1 |

| | | |
|---|---|---|
| cluster3731 | -3467.691639 | 1 |
| cluster3732 | -3301.484129 | 1 |
| cluster3733 | -3357.732908 | 1 |
| cluster3734 | -3454.94247 | 1 |
| cluster3735 | -3523.298063 | 1 |
| cluster3736 | -3985.410803 | 1 |
| cluster3737 | -1411.789184 | 1 |
| cluster3738 | -4047.099394 | 1 |
| cluster3739 | -1317.47915 | 1 |
| cluster3740 | -4004.714869 | 1 |
| cluster3741 | -1527.750585 | 1 |
| cluster3742 | -1806.607219 | 1 |
| cluster3743 | -3481.783348 | 1 |
| cluster3744 | -1430.846803 | 1 |
| cluster3745 | -3022.893703 | 1 |
| cluster3746 | -4303.90285 | 1 |
| cluster3747 | -1310.086906 | 1 |
| cluster3748 | -2568.160223 | 1 |
| cluster3749 | -3719.063882 | 1 |
| cluster3750 | -1579.627335 | 1 |
| cluster3751 | -3780.353738 | 1 |
| cluster3752 | -3249.528113 | 1 |
| cluster3753 | -1093.025573 | 1 |
| cluster3754 | -995.036534 | 1 |
| cluster3755 | -3020.499655 | 1 |
| cluster3756 | -1947.325022 | 1 |
| cluster3757 | -2350.725996 | 1 |
| cluster3758 | -1713.561814 | 1 |
| cluster3759 | -3355.443705 | 1 |
| cluster3760 | -3020.043232 | 1 |
| cluster3761 | -951.811414 | 1 |
| cluster3762 | -2233.72656 | 1 |
| cluster3763 | -2185.692226 | 1 |
| cluster3764 | -1873.59022 | 1 |
| cluster3765 | -3127.771737 | 1 |
| cluster3766 | -2137.219793 | 1 |
| cluster3767 | -1014.173969 | 1 |
| cluster3768 | -2396.000336 | 1 |
| cluster3769 | -2021.125925 | 1 |
| cluster3770 | -1965.643593 | 1 |
| cluster3771 | -2329.22799 | 1 |
| cluster3772 | -826.950653 | 1 |
| cluster3773 | -1143.609279 | 1 |
| cluster3774 | -1571.36834 | 1 |

| | | |
|---|---|---|
| cluster3775 | -1049.761758 | 1 |
| cluster3776 | -2761.226789 | 1 |
| cluster3777 | -3420.577098 | 1 |
| cluster3778 | -1214.049397 | 1 |
| cluster3779 | -3501.84663 | 1 |
| cluster3780 | -2066.222847 | 1 |
| cluster3781 | -2977.429944 | 1 |
| cluster3782 | -2600.328446 | 1 |
| cluster3783 | -2636.871245 | 1 |
| cluster3784 | -3326.503476 | 1 |
| cluster3785 | -3132.25544 | 1 |
| cluster3786 | -2711.472412 | 1 |
| cluster3787 | -1759.873851 | 1 |
| cluster3788 | -1240.077032 | 1 |
| cluster3789 | -3579.127326 | 1 |
| cluster3790 | -2577.043071 | 1 |
| cluster3791 | -489.869885 | 1 |
| cluster3792 | -2231.809117 | 1 |
| cluster3793 | -2150.01731 | 1 |
| cluster3794 | -2614.23646 | 1 |
| cluster3795 | -1982.956145 | 1 |
| cluster3796 | -1325.744881 | 1 |
| cluster3797 | -4270.721018 | 1 |
| cluster3798 | -4102.882201 | 1 |
| cluster3799 | -2613.885053 | 1 |
| cluster3800 | -449.669291 | 1 |
| cluster3801 | -2761.674782 | 1 |
| cluster3802 | -3015.88075 | 1 |
| cluster3803 | -2558.734476 | 1 |
| cluster3804 | -3002.495651 | 1 |
| cluster3805 | -3123.805801 | 1 |
| cluster3806 | -4084.284172 | 1 |
| cluster3807 | -1689.249744 | 1 |
| cluster3808 | -2107.366524 | 1 |
| cluster3809 | -2868.259106 | 1 |
| cluster3810 | -3019.571731 | 1 |
| cluster3811 | -835.316477 | 1 |
| cluster3812 | -3422.846666 | 1 |
| cluster3813 | -3828.834011 | 1 |
| cluster3814 | -1702.171086 | 1 |
| cluster3815 | -2029.798018 | 1 |
| cluster3816 | -2966.272168 | 1 |
| cluster3817 | -1152.061057 | 1 |
| cluster3818 | -2187.413959 | 1 |

| | | |
|---|---|---|
| cluster3819 | -3904.747448 | 1 |
| cluster3820 | -2575.716513 | 1 |
| cluster3821 | -2621.97838 | 1 |
| cluster3822 | -3942.377103 | 1 |
| cluster3823 | -4084.61773 | 1 |
| cluster3824 | -2755.221452 | 1 |
| cluster3825 | -1706.766405 | 1 |
| cluster3826 | -3405.929306 | 1 |
| cluster3827 | -2803.285214 | 1 |
| cluster3828 | -2671.340635 | 1 |
| cluster3829 | -2612.834144 | 1 |
| cluster3830 | -1810.126706 | 1 |
| cluster3831 | -2696.020227 | 1 |
| cluster3832 | -2185.860036 | 1 |
| cluster3833 | -2550.942465 | 1 |
| cluster3834 | -2464.58891 | 1 |
| cluster3835 | -2420.833333 | 1 |
| cluster3836 | -2240.265876 | 1 |
| cluster3837 | -3025.039795 | 1 |
| cluster3838 | -2672.98309 | 1 |
| cluster3839 | -1791.627893 | 1 |
| cluster3840 | -2315.954774 | 1 |
| cluster3841 | -2470.548465 | 1 |
| cluster3842 | -1850.506678 | 1 |
| cluster3843 | -1724.507183 | 1 |
| cluster3844 | -1148.47472 | 1 |
| cluster3845 | -702.020057 | 1 |
| cluster3846 | -1507.569931 | 1 |
| cluster3847 | -1535.29905 | 1 |
| cluster3848 | -2909.600402 | 1 |
| cluster3849 | -3051.240078 | 1 |
| cluster3850 | -1894.699488 | 1 |
| cluster3851 | -1977.461931 | 1 |
| cluster3852 | -2348.108588 | 1 |
| cluster3853 | -1386.054746 | 1 |
| cluster3854 | -1449.704521 | 1 |
| cluster3855 | -3619.446665 | 1 |
| cluster3856 | -2045.436684 | 1 |
| cluster3857 | -3685.348645 | 1 |
| cluster3858 | -2716.77727 | 1 |
| cluster3859 | -2356.872893 | 1 |
| cluster3860 | -2841.846476 | 1 |
| cluster3861 | -1547.034141 | 1 |
| cluster3862 | -1154.19757 | 1 |

| | | |
|---|---|---|
| cluster3863 | -1849.90542 | 1 |
| cluster3864 | -2875.960096 | 1 |
| cluster3865 | -3364.959969 | 1 |
| cluster3866 | -1928.488801 | 1 |
| cluster3867 | -646.226259 | 1 |
| cluster3868 | -1586.657248 | 1 |
| cluster3869 | -3362.563962 | 1 |
| cluster3870 | -2835.675907 | 1 |
| cluster3871 | -2672.806851 | 1 |
| cluster3872 | -3213.800726 | 1 |
| cluster3873 | -2059.273072 | 1 |
| cluster3874 | -2408.441488 | 1 |
| cluster3875 | -923.047634 | 1 |
| cluster3876 | -870.753545 | 1 |
| cluster3877 | -1676.229871 | 1 |
| cluster3879 | -2232.236782 | 1 |
| cluster3880 | -1559.759248 | 1 |
| cluster3881 | -3355.200155 | 1 |
| cluster3882 | -3142.042053 | 1 |
| cluster3883 | -480.225008 | 1 |
| cluster3884 | -1970.969906 | 1 |
| cluster3885 | -3486.823369 | 1 |
| cluster3886 | -1757.109957 | 1 |
| cluster3887 | -1954.718652 | 1 |
| cluster3888 | -1424.777638 | 1 |
| cluster3889 | -1158.730888 | 1 |
| cluster3890 | -2203.50646 | 1 |
| cluster3891 | -2647.9807 | 1 |
| cluster3892 | -1564.768479 | 1 |
| cluster3893 | -1690.002048 | 1 |
| cluster3894 | -1151.991737 | 1 |
| cluster3895 | -2369.819739 | 1 |
| cluster3896 | -1015.426673 | 1 |
| cluster3897 | -1712.259777 | 1 |
| cluster3898 | -2719.587173 | 1 |
| cluster3899 | -3475.610941 | 1 |
| cluster3900 | -2501.93937 | 1 |
| cluster3901 | -2017.259979 | 1 |
| cluster3902 | -3066.513323 | 1 |
| cluster3903 | -1933.894826 | 1 |
| cluster3904 | -1063.886805 | 1 |
| cluster3905 | -1687.244318 | 1 |
| cluster3906 | -1683.702646 | 1 |
| cluster3907 | -2240.824598 | 1 |

| | | |
|---|---|---|
| cluster3908 | -2158.229009 | 1 |
| cluster3909 | -2857.963946 | 1 |
| cluster3910 | -1350.624218 | 1 |
| cluster3911 | -2151.881598 | 1 |
| cluster3912 | -1130.446823 | 1 |
| cluster3913 | -2885.315599 | 1 |
| cluster3914 | -2441.367747 | 1 |
| cluster3915 | -2565.341763 | 1 |
| cluster3916 | -980.408639 | 1 |
| cluster3917 | -3056.62691 | 1 |
| cluster3918 | -3193.189113 | 1 |
| cluster3919 | -627.993438 | 1 |
| cluster3920 | -743.213252 | 1 |
| cluster3921 | -2513.481765 | 1 |
| cluster3922 | -1376.888407 | 1 |
| cluster3923 | -1543.912565 | 1 |
| cluster3924 | -1988.85432 | 1 |
| cluster3925 | -1212.92873 | 1 |
| cluster3926 | -2644.749529 | 1 |
| cluster3927 | -1444.902945 | 1 |
| cluster3928 | -1319.502681 | 1 |
| cluster3929 | -2021.261171 | 1 |
| cluster3930 | -2407.527595 | 1 |
| cluster3931 | -1899.362001 | 1 |
| cluster3932 | -1252.62753 | 1 |
| cluster3933 | -1233.591461 | 1 |
| cluster3934 | -844.62932 | 1 |
| cluster3935 | -3404.008666 | 1 |
| cluster3936 | -2518.065737 | 1 |
| cluster3937 | -2312.504611 | 1 |
| cluster3938 | -2044.079355 | 1 |
| cluster3939 | -2465.301554 | 1 |
| cluster3940 | -1547.228078 | 1 |
| cluster3941 | -1063.062654 | 1 |
| cluster3942 | -2301.678808 | 1 |
| cluster3943 | -2713.820668 | 1 |
| cluster3944 | -1805.995363 | 1 |
| cluster3945 | -1902.848464 | 1 |
| cluster3946 | -2097.868602 | 1 |
| cluster3947 | -2665.897623 | 1 |
| cluster3948 | -1157.466806 | 1 |
| cluster3949 | -1024.505774 | 1 |
| cluster3950 | -2106.844393 | 1 |
| cluster3951 | -3253.119576 | 1 |

| | | |
|---|---|---|
| cluster3952 | -1878.281431 | 1 |
| cluster3953 | -1575.0686 | 1 |
| cluster3954 | -2065.590741 | 1 |
| cluster3955 | -1376.318622 | 1 |
| cluster3956 | -2411.520598 | 1 |
| cluster3957 | -2386.664407 | 1 |
| cluster3958 | -1281.582502 | 1 |
| cluster3959 | -3086.200358 | 1 |
| cluster3960 | -2640.663732 | 1 |
| cluster3961 | -1799.005924 | 1 |
| cluster3962 | -1726.194723 | 1 |
| cluster3963 | -2377.660672 | 1 |
| cluster3964 | -3167.28568 | 1 |
| cluster3965 | -1730.035617 | 1 |
| cluster3966 | -1657.457626 | 1 |
| cluster3967 | -1888.168862 | 1 |
| cluster3968 | -1377.926515 | 1 |
| cluster3969 | -1056.933956 | 1 |
| cluster3970 | -2762.807273 | 1 |
| cluster3971 | -1936.115055 | 1 |
| cluster3972 | -1185.962617 | 1 |
| cluster3973 | -1617.980316 | 1 |
| cluster3974 | -2157.091791 | 1 |
| cluster3975 | -1449.606371 | 1 |
| cluster3976 | -1870.990163 | 1 |
| cluster3977 | -2389.54653 | 1 |
| cluster3978 | -1048.924242 | 1 |
| cluster3979 | -2067.319962 | 1 |
| cluster3980 | -1042.189379 | 1 |
| cluster3981 | -2306.342487 | 1 |
| cluster3982 | -2054.240366 | 1 |
| cluster3983 | -3039.485015 | 1 |
| cluster3984 | -1506.649839 | 1 |
| cluster3985 | -1161.402817 | 1 |
| cluster3986 | -2065.668771 | 1 |
| cluster3987 | -2517.6064 | 1 |
| cluster3988 | -1484.632178 | 1 |
| cluster3989 | -2031.008847 | 1 |
| cluster3990 | -2585.627238 | 1 |
| cluster3991 | -1803.671949 | 1 |
| cluster3992 | -2127.958434 | 1 |
| cluster3993 | -1903.258832 | 1 |
| cluster3994 | -2651.345512 | 1 |
| cluster3995 | -1839.26104 | 1 |

| | | |
|---|---|---|
| cluster3996 | -2125.070454 | 1 |
| cluster3997 | -2803.801558 | 1 |
| cluster3998 | -1581.159586 | 1 |
| cluster3999 | -2869.283776 | 1 |
| cluster4000 | -2605.928485 | 1 |
| cluster4001 | -2051.039319 | 1 |
| cluster4002 | -2266.194598 | 1 |
| cluster4003 | -1559.164011 | 1 |
| cluster4004 | -1775.524328 | 1 |
| cluster4005 | -2310.929933 | 1 |
| cluster4006 | -1246.605471 | 1 |
| cluster4007 | -1806.457299 | 1 |
| cluster4008 | -1681.532177 | 1 |
| cluster4009 | -1807.523605 | 1 |
| cluster4010 | -1786.210631 | 1 |
| cluster4011 | -1127.997821 | 1 |
| cluster4012 | -2782.92619 | 1 |
| cluster4013 | -979.714002 | 1 |
| cluster4014 | -1374.70825 | 1 |
| cluster4015 | -644.580496 | 1 |
| cluster4016 | -2175.773098 | 1 |
| cluster4017 | -1510.544764 | 1 |
| cluster4018 | -1383.733133 | 1 |
| cluster4019 | -1993.336979 | 1 |
| cluster4020 | -1135.568795 | 1 |
| cluster4021 | -1838.151003 | 1 |
| cluster4022 | -1110.687151 | 1 |
| cluster4023 | -2032.608632 | 1 |
| cluster4024 | -2671.076356 | 1 |
| cluster4025 | -782.018996 | 1 |
| cluster4026 | -1339.840667 | 1 |
| cluster4027 | -1159.883962 | 1 |
| cluster4028 | -2313.43971 | 1 |
| cluster4029 | -2464.250064 | 1 |
| cluster4030 | -1270.952633 | 1 |
| cluster4031 | -1234.59534 | 1 |
| cluster4032 | -1756.567203 | 1 |
| cluster4033 | -2419.686262 | 1 |
| cluster4034 | -580.179924 | 1 |
| cluster4035 | -2352.578031 | 1 |
| cluster4036 | -1292.726225 | 1 |
| cluster4037 | -1898.527709 | 1 |
| cluster4038 | -1080.421198 | 1 |
| cluster4039 | -1643.383775 | 1 |

| | | |
|---|---|---|
| cluster4040 | -1265.191104 | 1 |
| cluster4041 | -1760.340371 | 1 |
| cluster4042 | -1022.415368 | 1 |
| cluster4043 | -1244.969628 | 1 |
| cluster4044 | -1277.941716 | 1 |
| cluster4045 | -2200.112285 | 1 |
| cluster4046 | -1052.493825 | 1 |
| cluster4047 | -2493.121422 | 1 |
| cluster4048 | -1739.398458 | 1 |
| cluster4049 | -628.320607 | 1 |
| cluster4050 | -1456.374562 | 1 |
| cluster4051 | -1632.311801 | 1 |
| cluster4052 | -1529.466512 | 1 |
| cluster4053 | -1925.65143 | 1 |
| cluster4054 | -2208.823427 | 1 |
| cluster4055 | -903.775078 | 1 |
| cluster4056 | -1071.810879 | 1 |
| cluster4057 | -1793.844419 | 1 |
| cluster4058 | -1870.764954 | 1 |
| cluster4059 | -813.768994 | 1 |
| cluster4060 | -2173.110233 | 1 |
| cluster4061 | -1500.373206 | 1 |
| cluster4062 | -1468.696938 | 1 |
| cluster4063 | -1951.806202 | 1 |
| cluster4064 | -931.711636 | 1 |
| cluster4065 | -1169.186283 | 1 |
| cluster4066 | -1806.964513 | 1 |
| cluster4067 | -462.024263 | 1 |
| cluster4068 | -1552.511239 | 1 |
| cluster4069 | -1157.449868 | 1 |
| cluster4070 | -847.653175 | 1 |
| cluster4071 | -1033.836614 | 1 |
| cluster4072 | -1127.570876 | 1 |
| cluster4073 | -2187.812395 | 1 |
| cluster4074 | -1439.740442 | 1 |
| cluster4075 | -710.500896 | 1 |
| cluster4076 | -1787.161471 | 1 |
| cluster4077 | -1861.27818 | 1 |
| cluster4078 | -1444.206323 | 1 |
| cluster4079 | -1550.822628 | 1 |
| cluster4080 | -837.115064 | 1 |
| cluster4081 | -1325.394149 | 1 |
| cluster4082 | -835.146691 | 1 |
| cluster4083 | -1898.204715 | 1 |

| | | |
|---|---|---|
| cluster4084 | -807.175221 | 1 |
| cluster4085 | -920.219861 | 1 |
| cluster4086 | -1514.011459 | 1 |
| cluster4087 | -1441.425767 | 1 |
| cluster4088 | -1432.290933 | 1 |
| cluster4089 | -1826.073196 | 1 |
| cluster4090 | -852.451306 | 1 |
| cluster4091 | -1073.783914 | 1 |
| cluster4092 | -1629.576758 | 1 |
| cluster4093 | -1187.830638 | 1 |
| cluster4094 | -409.832599 | 1 |
| cluster4095 | -1527.835084 | 1 |
| cluster4096 | -1413.918567 | 1 |
| cluster4097 | -730.967827 | 1 |
| cluster4098 | -2146.890217 | 1 |
| cluster4099 | -1023.254346 | 1 |
| cluster4100 | -1308.380975 | 1 |
| cluster4101 | -2137.636334 | 1 |
| cluster4102 | -2003.582807 | 1 |
| cluster4103 | -1217.760217 | 1 |
| cluster4104 | -1273.419289 | 1 |
| cluster4105 | -850.233607 | 1 |
| cluster4106 | -1223.339208 | 1 |
| cluster4107 | -1000.991062 | 1 |
| cluster4108 | -969.110354 | 1 |
| cluster4109 | -1798.080253 | 1 |
| cluster4110 | -1360.955837 | 1 |
| cluster4111 | -1014.400775 | 1 |
| cluster4112 | -1326.74667 | 1 |
| cluster4113 | -861.086711 | 1 |
| cluster4114 | -1202.928318 | 1 |
| cluster4115 | -1402.31419 | 1 |
| cluster4116 | -693.824065 | 1 |
| cluster4117 | -762.391986 | 1 |
| cluster4118 | -1496.173419 | 1 |
| cluster4119 | -1092.355024 | 1 |
| cluster4120 | -1194.582351 | 1 |
| cluster4121 | -1532.695276 | 1 |
| cluster4122 | -546.711557 | 1 |
| cluster4123 | -546.310355 | 1 |
| cluster4124 | -1141.340446 | 1 |
| cluster4125 | -1153.316588 | 1 |
| cluster4126 | -814.002994 | 1 |
| cluster4127 | -854.537285 | 1 |

| | | |
|---|---|---|
| cluster4128 | -1096.61137 | 1 |
| cluster4128 | -1096.61137 | 1 |

**Supplementary Table 20.** codeml results - w for the whole tree

| ID | Likelihood | ω | dN | dS | Adjusted p value |
|---|---|---|---|---|---|
| cluster1127 | -1409.0635 | 0.0572 | 0.4157 | 7.2625 | 0.04 |
| cluster2853 | -22474.442 | 0.0281 | 0.1570 | 5.5945 | 7.95E-65 |
| cluster2854 | -1615.3656 | 0.0354 | 0.0598 | 1.6902 | 0.0005 |
| cluster2855 | -23555.854 | 0.0204 | 0.1527 | 7.4781 | 1.96E-92 |
| cluster2856 | -15426.646 | 0.1130 | 0.6471 | 5.7258 | 3.31E-07 |
| cluster2857 | -6020.4421 | 0.0122 | 0.0205 | 1.6970 | 3.92E-05 |
| cluster2858 | -7755.8194 | 0.0040 | 0.0162 | 4.0719 | 5.27E-54 |
| cluster2859 | -5679.2784 | 0.0414 | 0.0749 | 1.8124 | 4.48E-09 |
| cluster2860 | -16044.757 | 0.0017 | 0.0067 | 3.8455 | 9.24E-124 |
| cluster2861 | -8375.3851 | 0.0105 | 0.0433 | 4.1202 | 1.60E-41 |
| cluster2862 | -18857.655 | 0.0152 | 0.0669 | 4.3850 | 4.33E-88 |
| cluster2863 | -15075.873 | 0.0314 | 0.1610 | 5.1281 | 8.03E-44 |
| cluster2864 | -12551.751 | 0.0037 | 0.0184 | 4.9937 | 3.63E-107 |
| cluster2865 | -15005.038 | 0.0980 | 0.2690 | 2.7462 | 2.19E-12 |
| cluster2866 | -17312.539 | 0.0279 | 0.1643 | 5.8962 | 1.59E-48 |
| cluster2867 | -14864.912 | 0.0495 | 0.2689 | 5.4310 | 9.04E-28 |
| cluster2869 | -8917.0516 | 0.0919 | 0.2591 | 2.8205 | 2.27E-08 |
| cluster2870 | -3881.1185 | 0.0549 | 0.1737 | 3.1646 | 7.29E-06 |
| cluster2871 | -3139.982 | 0.0265 | 0.0396 | 1.4979 | 3.70E-06 |
| cluster2872 | -12260.947 | 0.0833 | 0.4011 | 4.8193 | 1.07E-10 |
| cluster2873 | -13852.572 | 0.0154 | 0.0569 | 3.6930 | 1.88E-59 |
| cluster2874 | -10388.461 | 0.0384 | 0.1084 | 2.8261 | 7.83E-22 |
| cluster2875 | -13637.18 | 0.0159 | 0.1008 | 6.3439 | 3.30E-64 |
| cluster2876 | -3971.74 | 0.0346 | 0.1208 | 3.4955 | 7.59E-11 |
| cluster2877 | -10638.032 | 0.0523 | 0.1845 | 3.5266 | 8.80E-21 |
| cluster2878 | -7059.7819 | 0.0395 | 0.0708 | 1.7893 | 6.20E-12 |
| cluster2879 | -10036.423 | 0.1902 | 0.3010 | 1.5834 | 0.005 |
| cluster2880 | -10642.395 | 0.0496 | 0.2699 | 5.4450 | 1.67E-17 |
| cluster2881 | -11523.3 | 0.0522 | 0.1964 | 3.7599 | 7.04E-19 |
| cluster2882 | -10650.117 | 0.0308 | 0.1392 | 4.5253 | 2.34E-27 |
| cluster2883 | -6765.0394 | 0.0704 | 0.1936 | 2.7489 | 2.00E-07 |
| cluster2884 | -12071.568 | 0.0260 | 0.0742 | 2.8596 | 1.30E-33 |
| cluster2885 | -15741.54 | 0.0440 | 0.2753 | 6.2611 | 2.11E-32 |
| cluster2886 | -12004.79 | 0.0397 | 0.1272 | 3.2050 | 8.33E-25 |
| cluster2887 | -8368.9918 | 0.0065 | 0.0388 | 5.9411 | 9.64E-55 |
| cluster2888 | -7786.5723 | 0.0656 | 0.4405 | 6.7128 | 7.40E-08 |
| cluster2889 | -6931.167 | 0.0085 | 0.0469 | 5.5279 | 1.37E-46 |
| cluster2890 | -2681.1915 | 0.0884 | 0.0649 | 0.7350 | 0.07 |
| cluster2891 | -12098.567 | 0.0388 | 0.2271 | 5.8464 | 1.65E-24 |
| cluster2892 | -7913.3434 | 0.0149 | 0.0504 | 3.3702 | 1.94E-34 |
| cluster2893 | -8469.8212 | 0.0360 | 0.2909 | 8.0795 | 5.17E-15 |
| cluster2894 | -5513.0096 | 0.0169 | 0.0684 | 4.0485 | 5.30E-17 |

| | | | | | |
|---|---|---|---|---|---|
| cluster2895 | -8448.9876 | 0.0363 | 0.1942 | 5.3432 | 5.24E-19 |
| cluster2896 | -10287.9 | 0.0102 | 0.0618 | 6.0313 | 5.79E-66 |
| cluster2897 | -9159.4752 | 0.0343 | 0.1175 | 3.4294 | 2.25E-22 |
| cluster2898 | -4501.103 | 0.0261 | 0.0716 | 2.7435 | 2.21E-14 |
| cluster2899 | -8006.9819 | 0.0193 | 0.1190 | 6.1686 | 1.71E-29 |
| cluster2900 | -9924.1834 | 0.0130 | 0.0430 | 3.3147 | 1.59E-39 |
| cluster2901 | -9241.1257 | 0.0371 | 0.2096 | 5.6424 | 1.74E-20 |
| cluster2902 | -3696.7104 | 0.0264 | 0.1508 | 5.7094 | 1.89E-13 |
| cluster2903 | -8864.182 | 0.0104 | 0.0489 | 4.6791 | 1.38E-45 |
| cluster2904 | -6415.5177 | 0.0905 | 0.2362 | 2.6097 | 5.98E-05 |
| cluster2905 | -8880.6593 | 0.0292 | 0.0704 | 2.4100 | 4.28E-19 |
| cluster2906 | -13667.93 | 0.0733 | 0.1747 | 2.3854 | 5.26E-16 |
| cluster2907 | -12740.73 | 0.0739 | 0.4695 | 6.3491 | 4.09E-11 |
| cluster2908 | -5645.6948 | 0.0124 | 0.0216 | 1.7364 | 5.84E-14 |
| cluster2909 | -2925.4123 | 0.0383 | 0.0626 | 1.6358 | 6.74E-05 |
| cluster2910 | -9271.5007 | 0.0433 | 0.1733 | 4.0029 | 2.52E-20 |
| cluster2911 | -4008.8477 | 0.0046 | 0.0055 | 1.1884 | 4.79E-10 |
| cluster2912 | -8769.1577 | 0.0262 | 0.1468 | 5.5969 | 6.11E-24 |
| cluster2913 | -6297.9977 | 0.0026 | 0.0101 | 3.8036 | 5.67E-41 |
| cluster2914 | -8063.931 | 0.0160 | 0.0412 | 2.5830 | 5.46E-25 |
| cluster2915 | -4207.4058 | 0.0170 | 0.0720 | 4.2415 | 4.84E-19 |
| cluster2916 | -9770.5319 | 0.0317 | 0.1064 | 3.3608 | 1.68E-26 |
| cluster2917 | -9027.3133 | 0.0200 | 0.0965 | 4.8392 | 8.08E-30 |
| cluster2918 | -4475.6107 | 0.0735 | 0.1600 | 2.1775 | 0.0001 |
| cluster2919 | -9565.1892 | 0.0670 | 0.1618 | 2.4135 | 3.73E-13 |
| cluster2920 | -6957.4191 | 0.0156 | 0.1305 | 8.3594 | 4.47E-30 |
| cluster2921 | -3809.2559 | 0.0040 | 0.0213 | 5.3097 | 7.02E-27 |
| cluster2922 | -6244.7533 | 0.0566 | 0.2935 | 5.1877 | 1.31E-09 |
| cluster2923 | -5867.7019 | 0.1795 | 0.5747 | 3.2022 | 0.07 |
| cluster2924 | -10977.144 | 0.0219 | 0.1415 | 6.4686 | 8.43E-33 |
| cluster2925 | -5771.6113 | 0.0332 | 0.1727 | 5.1957 | 1.52E-14 |
| cluster2926 | -11220.042 | 0.0212 | 0.0906 | 4.2699 | 5.56E-41 |
| cluster2927 | -4904.4904 | 0.0311 | 0.2022 | 6.5164 | 2.36E-11 |
| cluster2928 | -7181.0395 | 0.0216 | 0.1115 | 5.1650 | 1.07E-24 |
| cluster2929 | -8011.2774 | 0.0058 | 0.0263 | 4.5691 | 3.63E-52 |
| cluster2930 | -10989.016 | 0.0844 | 0.2681 | 3.1768 | 1.06E-09 |
| cluster2931 | -6866.5858 | 0.0525 | 0.3438 | 6.5456 | 3.14E-10 |
| cluster2932 | -9694.1603 | 0.0613 | 0.2951 | 4.8156 | 1.70E-12 |
| cluster2933 | -8737.174 | 0.0469 | 0.1000 | 2.1297 | 2.91E-14 |
| cluster2934 | -10531.84 | 0.0635 | 0.1583 | 2.4946 | 6.55E-12 |
| cluster2935 | -4952.4939 | 0.0282 | 0.0998 | 3.5386 | 4.47E-14 |
| cluster2936 | -9511.6547 | 0.0365 | 0.2569 | 7.0428 | 5.76E-24 |
| cluster2937 | -4595.6658 | 0.0279 | 0.0676 | 2.4255 | 1.30E-12 |
| cluster2938 | -7241.4465 | 0.0154 | 0.0513 | 3.3364 | 4.41E-29 |

| | | | | | |
|---|---|---|---|---|---|
| cluster2939 | -2817.9719 | 0.0233 | 0.0383 | 1.6407 | 1.49E-06 |
| cluster2940 | -11378.563 | 0.0671 | 0.2934 | 4.3709 | 8.29E-15 |
| cluster2941 | -3030.9524 | 0.0017 | 0.0016 | 0.8924 | 1.40E-06 |
| cluster2942 | -11359.598 | 0.0541 | 0.2883 | 5.3329 | 1.06E-16 |
| cluster2943 | -4040.4456 | 0.0108 | 0.0449 | 4.1645 | 1.67E-23 |
| cluster2944 | -8493.9711 | 0.1037 | 0.4214 | 4.0656 | 9.26E-06 |
| cluster2945 | -3085.051 | 0.0023 | 0.0219 | 9.6240 | 4.04E-21 |
| cluster2946 | -6742.6308 | 0.0095 | 0.0152 | 1.6148 | 2.68E-15 |
| cluster2947 | -9660.0228 | 0.0371 | 0.0985 | 2.6530 | 5.06E-21 |
| cluster2948 | -9376.9031 | 0.0530 | 0.1997 | 3.7702 | 5.15E-16 |
| cluster2949 | -6816.1306 | 0.0173 | 0.0488 | 2.8223 | 3.06E-23 |
| cluster2950 | -7114.9451 | 0.0168 | 0.0630 | 3.7429 | 2.13E-26 |
| cluster2951 | -9094.5785 | 0.0623 | 0.3515 | 5.6433 | 1.51E-10 |
| cluster2952 | -10895.607 | 0.0431 | 0.2145 | 4.9743 | 5.39E-22 |
| cluster2953 | -13290.999 | 0.0709 | 0.4176 | 5.8863 | 1.32E-14 |
| cluster2954 | -7274.0786 | 0.0197 | 0.1169 | 5.9279 | 2.80E-18 |
| cluster2955 | -6030.3797 | 0.0246 | 0.0962 | 3.9190 | 1.67E-15 |
| cluster2956 | -10134.502 | 0.0135 | 0.1451 | 10.7611 | 4.90E-38 |
| cluster2957 | -8388.2594 | 0.0935 | 0.4587 | 4.9079 | 3.59E-06 |
| cluster2958 | -9691.379 | 0.0119 | 0.0563 | 4.7478 | 1.02E-47 |
| cluster2959 | -5169.4772 | 0.0528 | 0.1514 | 2.8717 | 2.96E-09 |
| cluster2960 | -9442.3883 | 0.0237 | 0.1001 | 4.2269 | 5.26E-36 |
| cluster2961 | -8398.2071 | 0.0095 | 0.0979 | 10.3356 | 9.66E-47 |
| cluster2962 | -7495.6031 | 0.0221 | 0.1742 | 7.8740 | 7.48E-21 |
| cluster2963 | -7223.9045 | 0.0358 | 0.0652 | 1.8174 | 2.22E-14 |
| cluster2964 | -5248.8738 | 0.0313 | 0.0735 | 2.3456 | 1.09E-13 |
| cluster2965 | -7391.9295 | 0.0113 | 0.0421 | 3.7378 | 4.84E-32 |
| cluster2966 | -5587.8477 | 0.0227 | 0.1546 | 6.8156 | 5.18E-18 |
| cluster2967 | -9579.4573 | 0.0171 | 0.0493 | 2.8824 | 6.11E-40 |
| cluster2968 | -5456.9461 | 0.0185 | 0.0660 | 3.5801 | 1.09E-21 |
| cluster2969 | -7910.6215 | 0.0425 | 0.1671 | 3.9271 | 1.16E-15 |
| cluster2970 | -4236.3195 | 0.0038 | 0.0211 | 5.6294 | 3.44E-30 |
| cluster2971 | -3608.8835 | 0.0312 | 0.0375 | 1.2050 | 2.21E-05 |
| cluster2972 | -2670.0984 | 0.0062 | 0.0215 | 3.4402 | 7.22E-17 |
| cluster2973 | -6360.5299 | 0.0927 | 0.4092 | 4.4165 | 1.48E-05 |
| cluster2974 | -4423.4039 | 0.0303 | 0.0805 | 2.6535 | 2.33E-08 |
| cluster2975 | -10302.199 | 0.0723 | 0.3338 | 4.6137 | 5.92E-11 |
| cluster2976 | -1469.394 | 0.1964 | 0.3982 | 2.0280 | 0.48 |
| cluster2977 | -3393.0748 | 0.0047 | 0.0468 | 9.8911 | 4.80E-34 |
| cluster2978 | -8215.6155 | 0.0014 | 0.0048 | 3.3426 | 5.84E-57 |
| cluster2979 | -8840.0434 | 0.0397 | 0.1374 | 3.4586 | 9.79E-19 |
| cluster2980 | -1554.107 | 0.0219 | 0.2335 | 10.6816 | 2.56E-05 |
| cluster2981 | -6213.5358 | 0.0565 | 0.2398 | 4.2435 | 6.68E-11 |
| cluster2982 | -8059.248 | 0.1089 | 0.2362 | 2.1672 | 6.52E-05 |

| | | | | | |
|---|---|---|---|---|---|
| cluster2983 | -7519.9609 | 0.0237 | 0.1558 | 6.5725 | 3.39E-26 |
| cluster2984 | -4831.8675 | 0.0319 | 0.0694 | 2.1783 | 1.51E-12 |
| cluster2985 | -3972.6154 | 0.2375 | 0.4109 | 1.7298 | 0.24 |
| cluster2986 | -3497.2496 | 0.1081 | 0.2664 | 2.4635 | 0.01 |
| cluster2987 | -6078.1184 | 0.0303 | 0.0804 | 2.6491 | 1.11E-13 |
| cluster2988 | -5703.4482 | 0.0243 | 0.1248 | 5.1321 | 1.07E-19 |
| cluster2989 | -6589.7126 | 0.0391 | 0.0823 | 2.0996 | 3.99E-13 |
| cluster2990 | -7511.9915 | 0.0447 | 0.2083 | 4.6653 | 4.91E-14 |
| cluster2991 | -6308.8806 | 0.0600 | 0.3475 | 5.7917 | 9.92E-08 |
| cluster2992 | -4178.9613 | 0.0106 | 0.0187 | 1.7641 | 6.28E-12 |
| cluster2993 | -5423.5642 | 0.0324 | 0.1225 | 3.7812 | 8.60E-17 |
| cluster2994 | -2260.2683 | 0.0037 | 0.0088 | 2.3486 | 3.01E-10 |
| cluster2995 | -740.22868 | 0.0203 | 0.1012 | 4.9821 | 0.10 |
| cluster2996 | -7310.9378 | 0.0433 | 0.1361 | 3.1459 | 5.05E-16 |
| cluster2997 | -8172.0123 | 0.0404 | 0.1553 | 3.8387 | 6.13E-11 |
| cluster2998 | -7504.1864 | 0.0188 | 0.0971 | 5.1641 | 6.08E-27 |
| cluster2999 | -6764.2018 | 0.0309 | 0.1960 | 6.3505 | 1.02E-15 |
| cluster3000 | -6334.6519 | 0.0118 | 0.0270 | 2.2911 | 5.62E-18 |
| cluster3001 | -6059.573 | 0.0212 | 0.0657 | 3.1023 | 5.77E-16 |
| cluster3002 | -7669.4529 | 0.0266 | 0.0817 | 3.0699 | 5.67E-22 |
| cluster3003 | -7811.181 | 0.0900 | 0.2171 | 2.4124 | 1.92E-06 |
| cluster3004 | -5965.3045 | 0.0940 | 0.1746 | 1.8578 | 3.69E-05 |
| cluster3005 | -4102.904 | 0.0586 | 0.0835 | 1.4267 | 4.48E-05 |
| cluster3006 | -8672.0211 | 0.0572 | 0.2251 | 3.9383 | 6.50E-10 |
| cluster3007 | -8244.048 | 0.0269 | 0.2070 | 7.6984 | 2.01E-23 |
| cluster3008 | -5954.3444 | 0.0318 | 0.1482 | 4.6625 | 1.08E-16 |
| cluster3009 | -5913.3985 | 0.0434 | 0.1098 | 2.5270 | 1.10E-09 |
| cluster3010 | -8057.4747 | 0.1501 | 0.3388 | 2.2578 | 0.005 |
| cluster3011 | -3372.0484 | 0.0095 | 0.0730 | 7.7052 | 1.08E-16 |
| cluster3012 | -5210.2293 | 0.0585 | 0.3484 | 5.9553 | 2.37E-07 |
| cluster3013 | -6735.1785 | 0.0825 | 0.1512 | 1.8325 | 1.42E-06 |
| cluster3014 | -3561.8607 | 0.0081 | 0.0496 | 6.1100 | 1.39E-24 |
| cluster3015 | -2410.5303 | 0.0417 | 0.0785 | 1.8835 | 0.0002 |
| cluster3016 | -4556.4244 | 0.0393 | 0.0845 | 2.1520 | 7.97E-09 |
| cluster3017 | -5972.1963 | 0.0267 | 0.2010 | 7.5406 | 3.48E-20 |
| cluster3018 | -5679.2016 | 0.0066 | 0.0290 | 4.4170 | 4.84E-32 |
| cluster3019 | -7570.9311 | 0.0533 | 0.2842 | 5.3293 | 1.16E-11 |
| cluster3020 | -6195.2308 | 0.0343 | 0.1219 | 3.5578 | 4.61E-15 |
| cluster3021 | -3024.8757 | 0.0165 | 0.0381 | 2.3177 | 2.35E-10 |
| cluster3022 | -3431.5338 | 0.0601 | 0.3803 | 6.3299 | 0.0002 |
| cluster3023 | -4204.6702 | 0.0100 | 0.0279 | 2.7837 | 1.49E-19 |
| cluster3024 | -1959.8649 | 0.0878 | 0.1398 | 1.5929 | 0.02 |
| cluster3025 | -4691.8613 | 0.0101 | 0.0177 | 1.7521 | 1.30E-12 |
| cluster3026 | -4111.1941 | 0.1713 | 0.4792 | 2.7975 | 0.14 |

| | | | | | |
|---|---|---|---|---|---|
| cluster3027 | -3436.6337 | 0.0023 | 0.0101 | 4.4069 | 1.39E-30 |
| cluster3028 | -6804.0297 | 0.0407 | 0.2614 | 6.4299 | 3.61E-12 |
| cluster3029 | -6675.714 | 0.0288 | 0.0818 | 2.8432 | 9.67E-15 |
| cluster3030 | -5268.4411 | 0.0538 | 0.0966 | 1.7946 | 3.53E-08 |
| cluster3031 | -8797.2709 | 0.0987 | 0.3392 | 3.4344 | 6.44E-07 |
| cluster3032 | -5916.387 | 0.0231 | 0.0911 | 3.9381 | 5.01E-22 |
| cluster3033 | -3824.1384 | 0.1583 | 0.1370 | 0.8658 | 0.15 |
| cluster3034 | -4967.0442 | 0.0339 | 1.2755 | 37.5756 | 0.005 |
| cluster3035 | -7644.6418 | 0.0219 | 0.0742 | 3.3909 | 7.47E-25 |
| cluster3036 | -8982.3954 | 0.0711 | 0.3331 | 4.6874 | 4.43E-08 |
| cluster3037 | -2566.6164 | 0.0185 | 0.0369 | 1.9983 | 1.11E-06 |
| cluster3038 | -4471.4357 | 0.0594 | 0.2423 | 4.0766 | 4.96E-07 |
| cluster3039 | -6528.9869 | 0.0331 | 0.0529 | 1.6003 | 3.45E-11 |
| cluster3040 | -4844.1627 | 0.0190 | 0.0354 | 1.8614 | 5.84E-14 |
| cluster3041 | -8448.2259 | 0.0442 | 0.2364 | 5.3487 | 4.11E-15 |
| cluster3042 | -4404.5545 | 0.0340 | 0.1476 | 4.3472 | 5.76E-12 |
| cluster3043 | -2917.7548 | 0.0242 | 0.2322 | 9.5778 | 8.23E-09 |
| cluster3044 | -6428.3317 | 0.0299 | 0.1432 | 4.7876 | 1.53E-16 |
| cluster3045 | -6128.8607 | 0.0529 | 0.1257 | 2.3785 | 7.81E-11 |
| cluster3046 | -7278.3055 | 0.1755 | 0.4922 | 2.8048 | 0.02 |
| cluster3047 | -2123.7219 | 0.0034 | 0.0110 | 3.2335 | 1.05E-08 |
| cluster3048 | -5407.654 | 0.0220 | 0.0693 | 3.1560 | 3.76E-17 |
| cluster3049 | -7064.6565 | 0.0168 | 0.0554 | 3.3105 | 4.62E-31 |
| cluster3050 | -7562.4745 | 0.0402 | 0.1291 | 3.2076 | 2.22E-14 |
| cluster3051 | -7513.2988 | 0.0389 | 0.2399 | 6.1656 | 5.86E-14 |
| cluster3052 | -6827.3881 | 0.0481 | 0.2437 | 5.0688 | 1.47E-13 |
| cluster3053 | -9371.1202 | 0.0924 | 0.3099 | 3.3541 | 1.37E-07 |
| cluster3054 | -6880.5805 | 0.0094 | 0.0344 | 3.6772 | 1.29E-32 |
| cluster3055 | -3994.9713 | 0.0296 | 0.0946 | 3.2028 | 4.73E-08 |
| cluster3056 | -5995.1868 | 0.1007 | 0.1373 | 1.3626 | 0.0001 |
| cluster3057 | -919.87617 | 0.0526 | 0.0749 | 1.4231 | 0.05 |
| cluster3058 | -2536.0652 | 0.0266 | 0.0425 | 1.5963 | 1.95E-05 |
| cluster3059 | -8713.6377 | 0.0688 | 0.6036 | 8.7695 | 2.29E-06 |
| cluster3060 | -5326.766 | 0.0342 | 0.1592 | 4.6532 | 1.28E-11 |
| cluster3061 | -5740.9543 | 0.0160 | 0.0319 | 1.9993 | 2.41E-17 |
| cluster3062 | -5691.1676 | 0.0367 | 0.0722 | 1.9660 | 7.57E-10 |
| cluster3063 | -6148.0658 | 0.0320 | 0.1137 | 3.5584 | 1.23E-16 |
| cluster3064 | -1919.2507 | 0.0119 | 0.0197 | 1.6513 | 3.47E-05 |
| cluster3065 | -2300.6159 | 0.0940 | 0.3330 | 3.5408 | 0.02 |
| cluster3066 | -6187.6153 | 0.0693 | 0.1818 | 2.6256 | 5.00E-08 |
| cluster3067 | -4920.6966 | 0.0409 | 0.1288 | 3.1475 | 7.58E-11 |
| cluster3068 | -5143.4918 | 0.1100 | 0.1461 | 1.3291 | 0.003 |
| cluster3069 | -1632.4045 | 0.0014 | 0.0014 | 0.9245 | 0.001 |
| cluster3070 | -5192.4983 | 0.0366 | 0.1331 | 3.6355 | 2.97E-13 |

| | | | | | |
|---|---|---|---|---|---|
| cluster3071 | -5857.6636 | 0.0223 | 0.0487 | 2.1799 | 2.41E-17 |
| cluster3072 | -7373.5535 | 0.0849 | 0.3347 | 3.9417 | 2.28E-07 |
| cluster3073 | -3570.6462 | 0.0076 | 0.0215 | 2.8294 | 1.01E-14 |
| cluster3074 | -7651.2147 | 0.0296 | 0.1993 | 6.7458 | 1.41E-13 |
| cluster3075 | -5428.0848 | 0.0370 | 0.1237 | 3.3478 | 8.49E-13 |
| cluster3076 | -1814.4122 | 0.0322 | 0.1984 | 6.1714 | 0.001 |
| cluster3077 | -4385.0912 | 0.0233 | 0.0856 | 3.6817 | 2.77E-16 |
| cluster3078 | -5027.3277 | 0.0227 | 0.0813 | 3.5802 | 2.98E-16 |
| cluster3079 | -7622.0427 | 0.0335 | 0.1114 | 3.3285 | 7.76E-20 |
| cluster3080 | -5650.091 | 0.0162 | 0.0587 | 3.6203 | 9.40E-21 |
| cluster3081 | -3875.9268 | 0.0100 | 0.0280 | 2.7916 | 4.33E-18 |
| cluster3082 | -7187.8188 | 0.0587 | 0.2334 | 3.9732 | 5.38E-10 |
| cluster3083 | -4161.0009 | 0.0262 | 0.0582 | 2.2254 | 4.23E-11 |
| cluster3084 | -3198.0219 | 0.0372 | 0.0312 | 0.8381 | 0.002 |
| cluster3085 | -7344.5362 | 0.0221 | 0.2080 | 9.4149 | 2.55E-20 |
| cluster3086 | -3405.3795 | 0.0930 | 0.2335 | 2.5099 | 0.003 |
| cluster3087 | -6567.9918 | 0.0191 | 0.0498 | 2.6130 | 2.60E-20 |
| cluster3088 | -6393.3756 | 0.0653 | 0.1802 | 2.7607 | 8.33E-09 |
| cluster3089 | -3685.6616 | 0.0726 | 0.1864 | 2.5672 | 0.0001 |
| cluster3090 | -4961.243 | 0.0085 | 0.0386 | 4.5108 | 7.99E-28 |
| cluster3091 | -5911.3896 | 0.0545 | 0.0976 | 1.7889 | 3.28E-07 |
| cluster3092 | -4265.5375 | 0.0174 | 0.0587 | 3.3714 | 3.92E-14 |
| cluster3093 | -4546.7757 | 0.0064 | 0.0326 | 5.0624 | 7.94E-16 |
| cluster3094 | -7866.104 | 0.0301 | 0.2339 | 7.7792 | 7.80E-21 |
| cluster3095 | -4870.8136 | 0.0063 | 0.0243 | 3.8613 | 9.54E-27 |
| cluster3096 | -6132.6258 | 0.0335 | 0.1263 | 3.7711 | 2.13E-14 |
| cluster3097 | -7077.6425 | 0.0210 | 0.0421 | 2.0056 | 9.79E-19 |
| cluster3098 | -1418.7892 | 0.0072 | 0.0424 | 5.9317 | 3.36E-11 |
| cluster3099 | -4273.551 | 0.0405 | 0.1320 | 3.2611 | 5.35E-09 |
| cluster3100 | -4394.3538 | 0.0423 | 0.1566 | 3.7003 | 1.68E-09 |
| cluster3101 | -3436.1543 | 0.0478 | 0.1483 | 3.1057 | 2.85E-08 |
| cluster3102 | -2231.6988 | 0.0146 | 0.0578 | 3.9696 | 2.17E-12 |
| cluster3103 | -5216.6585 | 0.0365 | 0.2747 | 7.5161 | 2.06E-09 |
| cluster3104 | -4327.8651 | 0.0216 | 0.0796 | 3.6891 | 4.02E-15 |
| cluster3105 | -7972.7299 | 0.0280 | 0.2166 | 7.7377 | 1.19E-16 |
| cluster3106 | -5132.2566 | 0.0028 | 0.0063 | 2.3077 | 2.80E-23 |
| cluster3107 | -3137.8962 | 0.0413 | 0.0802 | 1.9450 | 2.86E-05 |
| cluster3108 | -4331.8829 | 0.1406 | 0.3789 | 2.6944 | 0.01 |
| cluster3109 | -3462.7813 | 0.0281 | 0.1277 | 4.5502 | 2.41E-11 |
| cluster3110 | -6381.2098 | 0.1084 | 0.2470 | 2.2799 | 0.0003 |
| cluster3111 | -4889.452 | 0.0326 | 0.0973 | 2.9871 | 1.80E-11 |
| cluster3112 | -6145.7317 | 0.0419 | 0.2802 | 6.6851 | 6.61E-10 |
| cluster3113 | -3068.717 | 0.0253 | 0.1080 | 4.2741 | 2.36E-10 |
| cluster3114 | -4690.338 | 0.0327 | 0.0797 | 2.4376 | 3.95E-10 |

| | | | | | |
|---|---|---|---|---|---|
| cluster3115 | -3777.6541 | 0.0181 | 0.0620 | 3.4264 | 4.42E-14 |
| cluster3116 | -2724.4694 | 0.0379 | 0.1288 | 3.3984 | 1.56E-06 |
| cluster3117 | -7499.9637 | 0.0477 | 0.3423 | 7.1761 | 1.45E-12 |
| cluster3118 | -1576.1741 | 0.0331 | 0.0312 | 0.9459 | 0.01 |
| cluster3119 | -2982.7749 | 0.0394 | 0.1833 | 4.6526 | 6.33E-06 |
| cluster3120 | -4910.8118 | 0.0228 | 0.0726 | 3.1829 | 6.68E-16 |
| cluster3121 | -6821.3303 | 0.0270 | 0.1043 | 3.8647 | 2.94E-18 |
| cluster3122 | -3558.4319 | 0.0209 | 0.0439 | 2.1044 | 2.48E-10 |
| cluster3123 | -3831.671 | 0.0144 | 0.0344 | 2.3839 | 1.21E-14 |
| cluster3124 | -6377.5959 | 0.0391 | 0.1411 | 3.6053 | 4.74E-14 |
| cluster3125 | -4803.2344 | 0.0663 | 0.1784 | 2.6914 | 4.47E-05 |
| cluster3126 | -7553.0047 | 0.0423 | 0.1406 | 3.3237 | 1.20E-15 |
| cluster3127 | -2607.5707 | 0.0081 | 0.0361 | 4.4429 | 3.80E-09 |
| cluster3128 | -2238.3573 | 0.0021 | 0.0044 | 2.0697 | 5.68E-10 |
| cluster3129 | -3264.2144 | 0.0371 | 0.3646 | 9.8345 | 9.32E-06 |
| cluster3130 | -5799.4408 | 0.0359 | 0.1872 | 5.2166 | 3.99E-14 |
| cluster3131 | -1764.779 | 0.0081 | 0.0476 | 5.8478 | 2.84E-10 |
| cluster3132 | -5290.1361 | 0.0247 | 0.0704 | 2.8422 | 1.51E-15 |
| cluster3133 | -8925.5828 | 0.0611 | 0.4682 | 7.6656 | 2.50E-08 |
| cluster3134 | -3291.0444 | 0.0583 | 0.1074 | 1.8409 | 2.53E-05 |
| cluster3135 | -4400.5157 | 0.0109 | 0.0352 | 3.2273 | 5.21E-12 |
| cluster3136 | -3954.0917 | 0.0379 | 0.1859 | 4.9116 | 3.05E-09 |
| cluster3137 | -3090.9939 | 0.0248 | 0.1274 | 5.1413 | 1.49E-11 |
| cluster3138 | -4743.0435 | 0.0531 | 0.2258 | 4.2544 | 1.03E-08 |
| cluster3139 | -5753.88 | 0.0683 | 0.4685 | 6.8591 | 4.12E-06 |
| cluster3140 | -3560.405 | 0.0429 | 0.0659 | 1.5353 | 1.65E-05 |
| cluster3141 | -2415.3379 | 0.0101 | 0.0168 | 1.6624 | 3.94E-08 |
| cluster3142 | -1642.5948 | 0.1019 | 0.0573 | 0.5623 | 0.20 |
| cluster3143 | -3228.3455 | 0.0142 | 0.0325 | 2.2896 | 9.73E-12 |
| cluster3144 | -2088.7696 | 0.0048 | 0.1396 | 29.1979 | 6.45E-12 |
| cluster3145 | -6530.7948 | 0.0054 | 0.0333 | 6.1698 | 9.77E-51 |
| cluster3146 | -1742.1753 | 0.0024 | 0.0471 | 19.9901 | 2.93E-13 |
| cluster3147 | -4473.1553 | 0.0063 | 0.0249 | 3.9836 | 7.02E-27 |
| cluster3148 | -6023.3584 | 0.0290 | 0.0754 | 2.6009 | 3.86E-14 |
| cluster3149 | -2372.924 | 0.0028 | 0.0060 | 2.1442 | 1.10E-09 |
| cluster3150 | -4524.5946 | 0.0670 | 0.0994 | 1.4837 | 9.06E-05 |
| cluster3151 | -3975.3841 | 0.0510 | 0.1285 | 2.5169 | 5.23E-06 |
| cluster3152 | -8746.0796 | 0.0486 | 0.3573 | 7.3446 | 2.09E-13 |
| cluster3153 | -4517.0901 | 0.0430 | 0.1025 | 2.3831 | 1.03E-08 |
| cluster3154 | -5344.1975 | 0.0149 | 0.0657 | 4.4239 | 2.32E-27 |
| cluster3155 | -3014.951 | 0.0087 | 0.0369 | 4.2158 | 4.22E-17 |
| cluster3156 | -6789.2316 | 0.0264 | 0.1653 | 6.2547 | 2.86E-16 |
| cluster3157 | -3281.0113 | 0.0987 | 0.1191 | 1.2071 | 0.02 |
| cluster3158 | -3373.1154 | 0.0152 | 0.0698 | 4.5964 | 8.86E-16 |

| | | | | | |
|---|---|---|---|---|---|
| cluster3159 | -5813.5245 | 0.0481 | 0.3018 | 6.2738 | 2.95E-10 |
| cluster3160 | -1099.8366 | 0.0520 | 0.0409 | 0.7874 | 0.12 |
| cluster3161 | -4982.4473 | 0.0322 | 0.1942 | 6.0249 | 6.45E-11 |
| cluster3162 | -3639.6665 | 0.0091 | 0.0169 | 1.8471 | 9.70E-11 |
| cluster3163 | -3430.2637 | 0.0364 | 0.2003 | 5.5061 | 2.46E-08 |
| cluster3164 | -6180.6921 | 0.0576 | 0.2768 | 4.8050 | 3.73E-09 |
| cluster3165 | -8050.093 | 0.0559 | 0.3239 | 5.7905 | 7.06E-11 |
| cluster3166 | -4633.6964 | 0.0251 | 0.0761 | 3.0376 | 5.40E-16 |
| cluster3167 | -5313.6683 | 0.0933 | 0.1817 | 1.9479 | 8.71E-05 |
| cluster3168 | -7090.792 | 0.0178 | 0.1049 | 5.8938 | 7.68E-30 |
| cluster3169 | -5029.8687 | 0.0301 | 0.1812 | 6.0190 | 3.39E-14 |
| cluster3170 | -1623.243 | 0.0133 | 0.0096 | 0.7155 | 0.02 |
| cluster3171 | -6159.8172 | 0.0616 | 0.2017 | 3.2720 | 1.94E-07 |
| cluster3172 | -6262.2336 | 0.0280 | 0.0944 | 3.3750 | 5.87E-17 |
| cluster3173 | -4457.7883 | 0.0150 | 0.0666 | 4.4469 | 1.38E-20 |
| cluster3174 | -1301.3189 | 0.0020 | 0.0067 | 3.3395 | 5.10E-09 |
| cluster3175 | -1938.2118 | 0.0159 | 0.0428 | 2.6876 | 9.71E-07 |
| cluster3176 | -3888.08 | 0.0245 | 0.0533 | 2.1830 | 2.63E-07 |
| cluster3177 | -3506.9954 | 0.0530 | 0.1159 | 2.1862 | 0.0005 |
| cluster3178 | -6745.3697 | 0.0112 | 0.0608 | 5.4389 | 4.68E-41 |
| cluster3179 | -4630.6594 | 0.0662 | 0.4936 | 7.4594 | 2.98E-05 |
| cluster3180 | -1071.2516 | 0.0033 | 0.0305 | 9.3180 | 3.92E-07 |
| cluster3181 | -6693.4678 | 0.0215 | 0.0957 | 4.4596 | 1.07E-24 |
| cluster3182 | -2530.6341 | 0.0067 | 0.0152 | 2.2708 | 4.23E-09 |
| cluster3183 | -4042.5891 | 0.0825 | 0.3132 | 3.7939 | 0.0002 |
| cluster3184 | -4207.6217 | 0.0320 | 0.0695 | 2.1738 | 8.84E-09 |
| cluster3185 | -3546.7573 | 0.0168 | 0.1040 | 6.1856 | 7.39E-18 |
| cluster3186 | -2290.5937 | 0.0285 | 0.0882 | 3.0970 | 4.12E-06 |
| cluster3187 | -5393.6291 | 0.0207 | 0.0812 | 3.9218 | 1.20E-17 |
| cluster3188 | -4445.3655 | 0.0901 | 0.2064 | 2.2900 | 0.0005 |
| cluster3189 | -6229.5154 | 0.0426 | 0.2545 | 5.9710 | 1.14E-10 |
| cluster3190 | -5398.2093 | 0.0311 | 0.2289 | 7.3493 | 7.06E-14 |
| cluster3191 | -2831.571 | 0.0466 | 0.1450 | 3.1109 | 8.66E-05 |
| cluster3192 | -5887.2122 | 0.0518 | 0.1284 | 2.4786 | 4.45E-10 |
| cluster3193 | -3456.9371 | 0.0363 | 0.0722 | 1.9887 | 1.06E-06 |
| cluster3194 | -6981.9414 | 0.0599 | 0.3296 | 5.5033 | 4.23E-09 |
| cluster3195 | -3960.8499 | 0.0181 | 0.0802 | 4.4456 | 9.86E-18 |
| cluster3196 | -2244.7307 | 0.0045 | 0.0042 | 0.9559 | 6.56E-06 |
| cluster3197 | -5447.2492 | 0.0517 | 0.1093 | 2.1117 | 1.31E-07 |
| cluster3198 | -6432.0722 | 0.0245 | 0.1725 | 7.0522 | 1.49E-20 |
| cluster3199 | -2979.921 | 0.0111 | 0.0309 | 2.7854 | 1.59E-11 |
| cluster3200 | -4274.995 | 0.0117 | 0.0665 | 5.6925 | 2.46E-19 |
| cluster3201 | -4476.7045 | 0.1109 | 0.1401 | 1.2625 | 0.003 |
| cluster3202 | -6071.4804 | 0.0427 | 0.1186 | 2.7801 | 8.38E-11 |

| | | | | | |
|---|---|---|---|---|---|
| cluster3203 | -3017.4434 | 0.0495 | 0.3131 | 6.3276 | 0.0001 |
| cluster3204 | -1878.2925 | 0.0388 | 0.0428 | 1.1081 | 0.003 |
| cluster3205 | -2015.8027 | 0.0061 | 0.0063 | 1.0250 | 4.34E-05 |
| cluster3206 | -3914.8429 | 0.0342 | 0.1525 | 4.4552 | 1.04E-09 |
| cluster3207 | -5500.7496 | 0.0139 | 0.0661 | 4.7623 | 8.66E-26 |
| cluster3208 | -3286.7206 | 0.0050 | 0.0114 | 2.3062 | 1.65E-09 |
| cluster3209 | -5552.231 | 0.0235 | 0.1302 | 5.5344 | 2.55E-19 |
| cluster3210 | -3378.0098 | 0.0030 | 0.0483 | 15.9349 | 3.40E-21 |
| cluster3211 | -3435.1328 | 0.0315 | 0.1523 | 4.8379 | 1.10E-09 |
| cluster3212 | -2811.7606 | 0.0023 | 0.0049 | 2.1189 | 1.49E-12 |
| cluster3213 | -4615.1846 | 0.0933 | 0.3401 | 3.6446 | 0.0001 |
| cluster3214 | -6052.0021 | 0.0472 | 0.1399 | 2.9634 | 9.59E-12 |
| cluster3215 | -2279.3451 | 0.0023 | 0.0018 | 0.8050 | 0.0003 |
| cluster3216 | -4538.6292 | 0.0297 | 0.2976 | 10.0354 | 1.51E-10 |
| cluster3217 | -3824.998 | 0.0422 | 0.1017 | 2.4095 | 1.09E-07 |
| cluster3218 | -5084.3091 | 0.0244 | 0.1613 | 6.6045 | 4.52E-14 |
| cluster3219 | -5449.835 | 0.0123 | 0.0589 | 4.7815 | 1.30E-23 |
| cluster3220 | -2924.5994 | 0.0305 | 0.0651 | 2.1338 | 4.23E-06 |
| cluster3221 | -5736.3138 | 0.0327 | 0.2286 | 6.9919 | 6.19E-14 |
| cluster3222 | -2622.057 | 0.0401 | 0.0981 | 2.4484 | 8.17E-06 |
| cluster3223 | -3467.4743 | 0.0953 | 0.3066 | 3.2184 | 0.002 |
| cluster3224 | -1597.2435 | 0.0001 | 0.0001 | 1.3059 | 8.64E-06 |
| cluster3225 | -6105.9071 | 0.0320 | 0.1562 | 4.8802 | 5.91E-17 |
| cluster3226 | -3727.5489 | 0.0403 | 0.1484 | 3.6825 | 2.32E-08 |
| cluster3227 | -4398.6833 | 0.0691 | 0.3514 | 5.0844 | 4.21E-06 |
| cluster3228 | -3625.7919 | 0.0321 | 0.2024 | 6.3011 | 6.96E-10 |
| cluster3229 | -3870.6885 | 0.0294 | 0.0968 | 3.2950 | 4.43E-09 |
| cluster3230 | -3482.4259 | 0.0110 | 0.0271 | 2.4655 | 1.64E-14 |
| cluster3231 | -4529.0274 | 0.0244 | 0.0847 | 3.4811 | 1.84E-11 |
| cluster3232 | -5054.0095 | 0.0226 | 0.1261 | 5.5814 | 1.67E-18 |
| cluster3233 | -3436.5209 | 0.0412 | 0.1954 | 4.7446 | 4.47E-08 |
| cluster3234 | -3821.3413 | 0.0508 | 0.1630 | 3.2077 | 3.49E-07 |
| cluster3235 | -7271.9722 | 0.0377 | 0.3143 | 8.3436 | 5.74E-12 |
| cluster3236 | -3956.7151 | 0.0330 | 0.1749 | 5.2925 | 3.97E-12 |
| cluster3237 | -3244.7876 | 0.0585 | 0.2366 | 4.0446 | 3.10E-05 |
| cluster3238 | -2785.1486 | 0.0586 | 0.0857 | 1.4618 | 0.003 |
| cluster3239 | -2740.7946 | 0.0226 | 0.0437 | 1.9287 | 2.74E-05 |
| cluster3240 | -6962.1389 | 0.0319 | 0.1305 | 4.0937 | 3.13E-19 |
| cluster3241 | -5756.5255 | 0.0646 | 0.2683 | 4.1509 | 1.27E-07 |
| cluster3242 | -7163.5212 | 0.0443 | 0.3327 | 7.5155 | 3.96E-09 |
| cluster3243 | -5984.2992 | 0.1521 | 0.2340 | 1.5388 | 0.01 |
| cluster3244 | -6473.4941 | 0.0598 | 0.1904 | 3.1834 | 2.19E-10 |
| cluster3245 | -2585.8836 | 0.0029 | 0.0129 | 4.4745 | 1.46E-21 |
| cluster3246 | -5135.0283 | 0.0425 | 0.2237 | 5.2620 | 3.24E-11 |

| | | | | | |
|---|---|---|---|---|---|
| cluster3247 | -849.26146 | 0.0636 | 0.1773 | 2.7873 | 0.04 |
| cluster3248 | -3485.0894 | 0.0214 | 0.0913 | 4.2745 | 1.03E-12 |
| cluster3249 | -3483.2175 | 0.0167 | 0.0716 | 4.2900 | 7.71E-14 |
| cluster3250 | -3552.7387 | 0.1109 | 0.1876 | 1.6908 | 0.01 |
| cluster3251 | -4467.6505 | 0.0577 | 0.1747 | 3.0266 | 1.36E-07 |
| cluster3252 | -7422.9807 | 0.0637 | 0.4254 | 6.6752 | 5.96E-08 |
| cluster3253 | -4317.7487 | 0.0708 | 0.2338 | 3.3034 | 2.20E-05 |
| cluster3254 | -3110.4846 | 0.0347 | 0.1366 | 3.9363 | 6.95E-09 |
| cluster3255 | -5570.8422 | 0.0178 | 0.1487 | 8.3740 | 3.61E-20 |
| cluster3256 | -6101.377 | 0.0411 | 0.1659 | 4.0417 | 4.07E-13 |
| cluster3257 | -5144.8496 | 0.0064 | 0.0255 | 3.9932 | 1.71E-29 |
| cluster3258 | -1919.2434 | 0.0014 | 0.0029 | 2.0785 | 3.75E-08 |
| cluster3259 | -3556.5266 | 0.0360 | 0.0461 | 1.2838 | 1.37E-05 |
| cluster3260 | -4482.5218 | 0.0137 | 0.0407 | 2.9701 | 2.15E-18 |
| cluster3261 | -4948.4176 | 0.0694 | 0.4291 | 6.1843 | 2.40E-05 |
| cluster3262 | -5791.5304 | 0.0185 | 0.1241 | 6.7103 | 3.18E-22 |
| cluster3263 | -2057.4741 | 0.1245 | 0.2693 | 2.1630 | 0.11 |
| cluster3264 | -1835.3272 | 0.0103 | 0.0159 | 1.5565 | 2.68E-05 |
| cluster3265 | -4128.947 | 0.0030 | 0.0116 | 3.9034 | 8.46E-31 |
| cluster3266 | -2856.8814 | 0.0296 | 0.0437 | 1.4759 | 0.0001 |
| cluster3267 | -4125.0833 | 0.0316 | 0.1010 | 3.1927 | 9.72E-11 |
| cluster3268 | -2807.4256 | 0.0374 | 0.1242 | 3.3206 | 6.02E-07 |
| cluster3269 | -3538.4028 | 0.0575 | 0.2072 | 3.6079 | 1.51E-05 |
| cluster3270 | -2185.3925 | 0.0261 | 0.1400 | 5.3661 | 5.88E-06 |
| cluster3271 | -4112.8943 | 0.0124 | 0.0331 | 2.6859 | 5.61E-16 |
| cluster3272 | -939.60973 | 0.3125 | 0.1056 | 0.3380 | 0.73 |
| cluster3273 | -4005.7489 | 0.1157 | 0.4778 | 4.1286 | 0.004 |
| cluster3274 | -2033.335 | 0.0323 | 0.1989 | 6.1551 | 1.95E-05 |
| cluster3275 | -4551.2077 | 0.0484 | 0.2322 | 4.7934 | 1.38E-08 |
| cluster3276 | -2485.6801 | 0.0899 | 0.0832 | 0.9262 | 0.02 |
| cluster3277 | -3228.02 | 0.0499 | 0.2708 | 5.4269 | 5.53E-05 |
| cluster3278 | -4577.4933 | 0.0032 | 0.0058 | 1.7879 | 3.48E-17 |
| cluster3279 | -3733.0686 | 0.0167 | 0.0842 | 5.0435 | 1.22E-13 |
| cluster3280 | -3788.6395 | 0.0222 | 0.0582 | 2.6240 | 3.39E-11 |
| cluster3281 | -1967.4158 | 0.0208 | 0.1110 | 5.3407 | 7.59E-08 |
| cluster3282 | -2049.1961 | 0.0056 | 0.0170 | 3.0329 | 1.05E-11 |
| cluster3283 | -5464.9013 | 0.0645 | 0.1979 | 3.0707 | 1.59E-07 |
| cluster3284 | -5980.3706 | 0.0345 | 0.1400 | 4.0596 | 8.62E-15 |
| cluster3285 | -2362.9261 | 0.0074 | 0.0140 | 1.8948 | 3.43E-07 |
| cluster3286 | -2209.1763 | 0.0302 | 0.0479 | 1.5844 | 0.001 |
| cluster3287 | -1377.4229 | 0.0129 | 0.0377 | 2.9190 | 2.17E-06 |
| cluster3288 | -1241.6146 | 0.0064 | 0.0471 | 7.3323 | 1.26E-08 |
| cluster3289 | -5798.8476 | 0.0478 | 0.0829 | 1.7372 | 1.08E-09 |
| cluster3290 | -1253.0705 | 0.0211 | 0.0651 | 3.0851 | 0.001 |

| | | | | | |
|---|---|---|---|---|---|
| cluster3291 | -4787.8336 | 0.0397 | 0.1475 | 3.7116 | 6.36E-12 |
| cluster3292 | -2774.4252 | 0.0264 | 0.0689 | 2.6071 | 1.50E-06 |
| cluster3293 | -3184.8326 | 0.0095 | 0.0405 | 4.2576 | 1.97E-19 |
| cluster3294 | -2883.2729 | 0.0134 | 0.0520 | 3.8693 | 1.48E-10 |
| cluster3295 | -2241.2247 | 0.0016 | 0.0033 | 2.0402 | 1.63E-10 |
| cluster3296 | -1799.7807 | 0.0043 | 0.0233 | 5.4364 | 2.56E-10 |
| cluster3297 | -2355.5226 | 0.1643 | 0.3746 | 2.2808 | 0.17 |
| cluster3298 | -2944.7741 | 0.0040 | 0.0162 | 4.0732 | 9.67E-22 |
| cluster3299 | -7908.376 | 0.0486 | 0.5573 | 11.4762 | 1.39E-06 |
| cluster3300 | -2252.9511 | 0.0048 | 0.0212 | 4.4175 | 4.28E-15 |
| cluster3301 | -1804.4129 | 0.0491 | 0.0930 | 1.8947 | 0.001 |
| cluster3302 | -4560.683 | 0.0875 | 0.2537 | 2.9001 | 3.34E-05 |
| cluster3303 | -4480.6953 | 0.0355 | 0.2019 | 5.6764 | 1.77E-11 |
| cluster3304 | -2861.0003 | 0.0243 | 0.0459 | 1.8879 | 9.84E-07 |
| cluster3305 | -3805.3559 | 0.0636 | 0.2526 | 3.9720 | 3.54E-05 |
| cluster3306 | -5021.6065 | 0.0716 | 0.3105 | 4.3400 | 8.86E-06 |
| cluster3307 | -1671.3744 | 0.0045 | 0.0201 | 4.4473 | 4.20E-12 |
| cluster3308 | -4388.0274 | 0.0183 | 0.0939 | 5.1415 | 1.63E-18 |
| cluster3309 | -3385.2676 | 0.1189 | 0.2053 | 1.7282 | 0.02 |
| cluster3310 | -4175.2766 | 0.0158 | 0.0529 | 3.3429 | 3.04E-15 |
| cluster3311 | -6158.1264 | 0.0548 | 0.2050 | 3.7375 | 7.02E-10 |
| cluster3312 | -1931.8756 | 0.0268 | 0.1712 | 6.3938 | 1.91E-08 |
| cluster3313 | -2508.8927 | 0.0113 | 0.0202 | 1.7795 | 2.21E-07 |
| cluster3314 | -7142.1498 | 0.0426 | 0.3983 | 9.3416 | 1.06E-10 |
| cluster3315 | -3723.1006 | 0.0058 | 0.0194 | 3.3337 | 1.93E-21 |
| cluster3316 | -5974.5042 | 0.0853 | 0.3707 | 4.3473 | 1.83E-05 |
| cluster3317 | -1142.6546 | 0.0032 | 0.0097 | 3.1076 | 1.54E-06 |
| cluster3318 | -3441.7891 | 0.1079 | 0.5350 | 4.9569 | 0.03 |
| cluster3319 | -3769.6326 | 0.0355 | 0.2012 | 5.6749 | 4.52E-10 |
| cluster3320 | -2814.032 | 0.0549 | 0.2390 | 4.3569 | 0.0001 |
| cluster3321 | -2917.5628 | 0.0027 | 0.0069 | 2.5699 | 2.12E-09 |
| cluster3322 | -3279.181 | 0.0415 | 0.2447 | 5.8951 | 1.98E-07 |
| cluster3323 | -5520.6379 | 0.0503 | 0.2020 | 4.0142 | 9.87E-10 |
| cluster3324 | -1400.704 | 0.0225 | 0.1539 | 6.8427 | 0.001 |
| cluster3325 | -2061.16 | 0.0187 | 0.0887 | 4.7385 | 4.47E-08 |
| cluster3326 | -1704.4545 | 0.0736 | 0.0745 | 1.0128 | 0.07 |
| cluster3327 | -3865.001 | 0.0253 | 0.0864 | 3.4215 | 6.82E-13 |
| cluster3328 | -2608.0927 | 0.0114 | 0.0691 | 6.0644 | 1.32E-06 |
| cluster3329 | -3154.8694 | 0.0630 | 0.4119 | 6.5412 | 3.88E-05 |
| cluster3330 | -4710.9755 | 0.0474 | 0.2611 | 5.5043 | 9.28E-06 |
| cluster3331 | -2869.1642 | 0.0738 | 0.1356 | 1.8361 | 0.002 |
| cluster3332 | -6191.9466 | 0.0627 | 0.3719 | 5.9348 | 5.44E-09 |
| cluster3333 | -3576.0428 | 0.0239 | 0.1262 | 5.2712 | 2.98E-11 |
| cluster3334 | -1380.9122 | 0.0735 | 0.1501 | 2.0426 | 0.04 |

| | | | | | |
|---|---|---|---|---|---|
| cluster3335 | -2858.4299 | 0.0288 | 0.1377 | 4.7844 | 5.81E-09 |
| cluster3336 | -3028.0816 | 0.0038 | 0.0076 | 2.0106 | 3.62E-13 |
| cluster3337 | -2620.428 | 0.0308 | 0.0416 | 1.3487 | 1.48E-05 |
| cluster3338 | -6303.2886 | 0.0388 | 0.2898 | 7.4565 | 3.59E-12 |
| cluster3339 | -1910.627 | 0.0461 | 0.2089 | 4.5340 | 7.92E-05 |
| cluster3340 | -1627.5066 | 0.1675 | 0.1261 | 0.7534 | 0.48 |
| cluster3341 | -1577.0131 | 0.0110 | 0.0729 | 6.6542 | 2.02E-08 |
| cluster3342 | -2556.42 | 0.0214 | 0.1278 | 5.9631 | 3.82E-09 |
| cluster3343 | -2926.5396 | 0.0004 | 0.0019 | 4.2570 | 6.62E-24 |
| cluster3344 | -3149.3637 | 0.0266 | 0.0488 | 1.8345 | 4.28E-08 |
| cluster3345 | -2795.8665 | 0.0129 | 0.0865 | 6.6838 | 1.47E-15 |
| cluster3346 | -3610.3001 | 0.0035 | 0.0125 | 3.5284 | 5.84E-23 |
| cluster3347 | -4176.1333 | 0.0468 | 0.0996 | 2.1288 | 2.06E-07 |
| cluster3348 | -4560.0115 | 0.0500 | 0.1069 | 2.1406 | 1.70E-06 |
| cluster3349 | -4215.0767 | 0.0089 | 0.0385 | 4.3224 | 3.06E-17 |
| cluster3350 | -4778.9189 | 0.0501 | 0.2366 | 4.7284 | 9.81E-09 |
| cluster3351 | -3134.228 | 0.0161 | 0.0661 | 4.0958 | 3.56E-11 |
| cluster3352 | -4085.2769 | 0.0567 | 0.1530 | 2.6986 | 1.65E-05 |
| cluster3353 | -1586.1509 | 0.0108 | 0.0236 | 2.1808 | 1.67E-05 |
| cluster3354 | -2120.9362 | 0.0056 | 0.0094 | 1.7027 | 8.94E-07 |
| cluster3355 | -4288.0535 | 0.0431 | 0.1751 | 4.0650 | 1.34E-08 |
| cluster3356 | -4596.1283 | 0.0406 | 0.1025 | 2.5230 | 2.99E-09 |
| cluster3357 | -1290.149 | 0.0278 | 0.0768 | 2.7670 | 0.0002 |
| cluster3358 | -4171.919 | 0.0016 | 0.0037 | 2.3850 | 9.66E-26 |
| cluster3359 | -6043.5624 | 0.0380 | 0.2361 | 6.2179 | 2.91E-13 |
| cluster3360 | -2471.6558 | 0.0379 | 0.1535 | 4.0515 | 8.07E-06 |
| cluster3361 | -5035.5033 | 0.0371 | 0.2542 | 6.8492 | 7.38E-09 |
| cluster3362 | -3692.8134 | 0.0162 | 0.0544 | 3.3657 | 1.70E-15 |
| cluster3363 | -3721.1665 | 0.0023 | 0.0284 | 12.1496 | 8.30E-35 |
| cluster3364 | -4356.3214 | 0.0238 | 0.1664 | 6.9954 | 1.10E-10 |
| cluster3365 | -5566.0145 | 0.0623 | 0.2447 | 3.9245 | 1.01E-08 |
| cluster3366 | -3149.5658 | 0.0311 | 0.2732 | 8.7934 | 4.79E-07 |
| cluster3367 | -3694.0321 | 0.1339 | 0.3653 | 2.7286 | 0.01 |
| cluster3368 | -1654.7904 | 0.0266 | 0.0263 | 0.9873 | 0.001 |
| cluster3369 | -3766.3322 | 0.0154 | 0.0847 | 5.5151 | 1.57E-16 |
| cluster3370 | -6387.9186 | 0.0994 | 0.3539 | 3.5635 | 7.04E-05 |
| cluster3371 | -2934.5056 | 0.0062 | 0.0121 | 1.9640 | 1.59E-11 |
| cluster3372 | -3841.5257 | 0.0087 | 0.0462 | 5.3247 | 8.26E-18 |
| cluster3373 | -5277.5405 | 0.0084 | 0.0568 | 6.7237 | 4.41E-34 |
| cluster3374 | -5125.5212 | 0.0711 | 0.5310 | 7.4678 | 9.80E-06 |
| cluster3375 | -3052.2342 | 0.0054 | 0.0097 | 1.8344 | 3.03E-09 |
| cluster3376 | -1531.6801 | 0.0369 | 0.0488 | 1.3245 | 0.01 |
| cluster3377 | -2274.0822 | 0.0117 | 0.0720 | 6.1320 | 2.09E-15 |
| cluster3378 | -6508.1657 | 0.0479 | 0.2184 | 4.5606 | 7.09E-12 |

| | | | | | |
|---|---|---|---|---|---|
| cluster3379 | -1568.548 | 0.0338 | 0.0941 | 2.7864 | 0.0002 |
| cluster3380 | -2653.5263 | 0.0126 | 0.0799 | 6.3724 | 1.09E-06 |
| cluster3381 | -2268.0122 | 0.0336 | 0.1036 | 3.0792 | 9.73E-06 |
| cluster3382 | -885.76594 | 0.0065 | 0.0126 | 1.9458 | 0.01 |
| cluster3383 | -5151.7312 | 0.0599 | 0.3547 | 5.9224 | 3.94E-07 |
| cluster3384 | -1972.3235 | 0.0337 | 0.0981 | 2.9153 | 6.77E-05 |
| cluster3385 | -3771.6906 | 0.0220 | 0.1185 | 5.3794 | 3.43E-13 |
| cluster3386 | -4155.9089 | 0.0145 | 0.0782 | 5.3999 | 2.81E-20 |
| cluster3387 | -3031.3044 | 0.0428 | 0.1353 | 3.1653 | 5.53E-07 |
| cluster3388 | -2809.4357 | 0.0254 | 0.0841 | 3.3169 | 2.03E-09 |
| cluster3389 | -1788.9375 | 0.0099 | 0.0578 | 5.8313 | 4.04E-09 |
| cluster3390 | -5117.2051 | 0.0641 | 0.1803 | 2.8135 | 1.40E-06 |
| cluster3391 | -2416.4054 | 0.0084 | 0.0205 | 2.4404 | 2.92E-11 |
| cluster3392 | -2528.4497 | 0.0102 | 0.0604 | 5.9340 | 2.15E-12 |
| cluster3393 | -3179.5408 | 0.0576 | 0.2076 | 3.6031 | 0.0002 |
| cluster3394 | -713.6658 | 0.0036 | 0.0138 | 3.8266 | 1.93E-06 |
| cluster3395 | -2714.583 | 0.0150 | 0.0505 | 3.3674 | 7.09E-13 |
| cluster3396 | -4985.7009 | 0.0362 | 0.0960 | 2.6519 | 1.34E-12 |
| cluster3397 | -4642.0984 | 0.0281 | 0.1665 | 5.9259 | 5.33E-15 |
| cluster3398 | -3833.7546 | 0.0556 | 0.1355 | 2.4374 | 6.93E-06 |
| cluster3399 | -3230.7134 | 0.0442 | 0.1778 | 4.0216 | 3.09E-07 |
| cluster3400 | -1955.0251 | 0.1015 | 0.1024 | 1.0083 | 0.17 |
| cluster3401 | -2870.6706 | 0.0164 | 0.0660 | 4.0128 | 1.14E-08 |
| cluster3402 | -5115.3569 | 0.0228 | 0.0759 | 3.3225 | 1.11E-14 |
| cluster3403 | -3669.6508 | 0.0222 | 0.0760 | 3.4174 | 3.16E-11 |
| cluster3404 | -716.69613 | 0.0001 | 0.0000 | 0.4812 | 0.11 |
| cluster3405 | -1541.3321 | 0.1207 | 0.2248 | 1.8613 | 0.07 |
| cluster3406 | -3800.6507 | 0.0188 | 0.0867 | 4.6109 | 6.03E-15 |
| cluster3407 | -6244.3717 | 0.0605 | 0.2867 | 4.7378 | 3.06E-08 |
| cluster3408 | -1896.6956 | 0.0062 | 0.0086 | 1.3821 | 3.33E-06 |
| cluster3409 | -4909.5257 | 0.0233 | 0.1532 | 6.5779 | 3.03E-14 |
| cluster3410 | -1443.1283 | 0.0240 | 0.0357 | 1.4829 | 0.0005 |
| cluster3411 | -2582.9267 | 0.0003 | 0.0011 | 3.1748 | 5.24E-15 |
| cluster3412 | -3392.2366 | 0.0836 | 0.3041 | 3.6406 | 0.001 |
| cluster3413 | -2560.892 | 0.0464 | 0.1857 | 4.0081 | 1.91E-05 |
| cluster3414 | -2522.9666 | 0.1358 | 0.1502 | 1.1056 | 0.13 |
| cluster3415 | -3493.4086 | 0.0632 | 0.1476 | 2.3364 | 2.70E-05 |
| cluster3416 | -604.01786 | 0.0143 | 1.2849 | 89.7124 | 0.04 |
| cluster3417 | -1215.5877 | 0.0012 | 0.0747 | 61.5260 | 3.65E-10 |
| cluster3418 | -2534.288 | 0.0553 | 0.2122 | 3.8404 | 0.0002 |
| cluster3419 | -2673.1985 | 0.0984 | 0.1408 | 1.4313 | 0.07 |
| cluster3420 | -1786.1686 | 0.2544 | 0.5178 | 2.0348 | 0.40 |
| cluster3421 | -4207.7736 | 0.0113 | 0.0670 | 5.9596 | 1.66E-20 |
| cluster3422 | -5399.3755 | 0.0350 | 0.1057 | 3.0244 | 7.06E-14 |

| | | | | | |
|---|---|---|---|---|---|
| cluster3423 | -4251.2562 | 0.0471 | 0.3639 | 7.7284 | 4.46E-08 |
| cluster3424 | -2856.575 | 0.1021 | 0.2942 | 2.8819 | 0.01 |
| cluster3425 | -2232.6686 | 0.0440 | 0.0717 | 1.6270 | 0.002 |
| cluster3426 | -3411.3248 | 0.0181 | 0.0974 | 5.3802 | 1.83E-15 |
| cluster3427 | -3796.8009 | 0.0925 | 0.2338 | 2.5270 | 0.001 |
| cluster3428 | -2685.9856 | 0.0643 | 0.1307 | 2.0320 | 0.0004 |
| cluster3429 | -4145.7928 | 0.0286 | 0.1100 | 3.8549 | 7.60E-13 |
| cluster3430 | -4824.7322 | 0.0437 | 0.3205 | 7.3403 | 7.87E-09 |
| cluster3431 | -4739.2311 | 0.0425 | 0.2642 | 6.2222 | 4.72E-10 |
| cluster3432 | -3782.7767 | 0.0369 | 0.0819 | 2.2224 | 3.44E-08 |
| cluster3433 | -5393.5588 | 0.1700 | 0.3109 | 1.8282 | 0.03 |
| cluster3434 | -3295.1682 | 0.0365 | 0.0678 | 1.8604 | 4.79E-07 |
| cluster3435 | -4093.3889 | 0.0213 | 0.0948 | 4.4541 | 3.19E-15 |
| cluster3436 | -3135.3385 | 0.0234 | 0.1427 | 6.0869 | 5.81E-12 |
| cluster3437 | -2816.4584 | 0.0129 | 0.0606 | 4.6935 | 7.29E-15 |
| cluster3438 | -3825.024 | 0.0382 | 0.1314 | 3.4445 | 1.15E-08 |
| cluster3439 | -2056.4977 | 0.0525 | 0.1968 | 3.7456 | 0.002 |
| cluster3440 | -2314.4941 | 0.0259 | 0.0506 | 1.9547 | 6.77E-06 |
| cluster3441 | -2756.889 | 0.0376 | 0.2142 | 5.6968 | 8.60E-06 |
| cluster3442 | -2923.8857 | 0.0190 | 0.2372 | 12.4755 | 8.47E-08 |
| cluster3443 | -3532.5264 | 0.0147 | 0.0815 | 5.5490 | 1.06E-12 |
| cluster3444 | -3386.8303 | 0.0101 | 0.0333 | 3.2811 | 3.03E-14 |
| cluster3445 | -2965.094 | 0.0142 | 0.0293 | 2.0711 | 3.35E-11 |
| cluster3446 | -4062.1359 | 0.0307 | 0.1738 | 5.6551 | 5.16E-12 |
| cluster3447 | -5927.1987 | 0.0351 | 0.3318 | 9.4420 | 3.17E-11 |
| cluster3448 | -2378.8926 | 0.0229 | 0.1456 | 6.3606 | 9.86E-08 |
| cluster3449 | -613.68004 | 0.0034 | 0.0025 | 0.7345 | 0.04 |
| cluster3450 | -2518.6721 | 0.0201 | 0.1375 | 6.8332 | 1.68E-09 |
| cluster3451 | -3594.2495 | 0.0274 | 0.1477 | 5.3881 | 2.77E-10 |
| cluster3452 | -3964.8812 | 0.0474 | 0.2294 | 4.8381 | 6.58E-07 |
| cluster3453 | -1849.6933 | 0.0385 | 0.0896 | 2.3286 | 0.0003 |
| cluster3454 | -3695.992 | 0.0108 | 0.0931 | 8.5936 | 1.28E-15 |
| cluster3455 | -3649.3252 | 0.0436 | 0.0549 | 1.2564 | 4.34E-05 |
| cluster3456 | -4231.5565 | 0.0366 | 0.0760 | 2.0740 | 2.49E-08 |
| cluster3457 | -3649.4877 | 0.0307 | 0.1481 | 4.8318 | 6.18E-10 |
| cluster3458 | -1806.6119 | 0.0341 | 0.0832 | 2.4389 | 0.0003 |
| cluster3459 | -2640.8916 | 0.0443 | 0.2999 | 6.7721 | 1.79E-05 |
| cluster3460 | -5580.6164 | 0.0321 | 0.2594 | 8.0715 | 1.95E-12 |
| cluster3461 | -3102.9271 | 0.0660 | 0.2921 | 4.4258 | 0.0004 |
| cluster3462 | -824.35104 | 0.0361 | 0.0171 | 0.4735 | 0.22 |
| cluster3463 | -4771.416 | 0.0475 | 0.2189 | 4.6054 | 1.88E-08 |
| cluster3464 | -5608.2815 | 0.0457 | 0.3287 | 7.1911 | 1.71E-09 |
| cluster3465 | -3956.0017 | 0.0098 | 0.0900 | 9.2026 | 3.20E-18 |
| cluster3466 | -2644.3002 | 0.0125 | 0.0561 | 4.5067 | 2.58E-13 |

| | | | | | |
|---|---|---|---|---|---|
| cluster3467 | -4061.1708 | 0.0330 | 0.1218 | 3.6901 | 7.92E-10 |
| cluster3468 | -1849.9386 | 0.0077 | 0.0674 | 8.8149 | 3.01E-11 |
| cluster3469 | -2105.9078 | 0.0226 | 0.0926 | 4.0913 | 3.28E-08 |
| cluster3470 | -3884.7839 | 0.0135 | 0.0693 | 5.1273 | 4.63E-16 |
| cluster3471 | -4173.3987 | 0.0428 | 0.1814 | 4.2346 | 1.09E-08 |
| cluster3472 | -3722.0736 | 0.0178 | 0.0542 | 3.0365 | 2.93E-15 |
| cluster3473 | -3470.9192 | 0.0273 | 0.0984 | 3.6057 | 5.92E-11 |
| cluster3474 | -5366.9017 | 0.0310 | 0.3024 | 9.7533 | 7.95E-11 |
| cluster3475 | -2340.9421 | 0.0123 | 0.0714 | 5.7832 | 1.09E-08 |
| cluster3476 | -5111.8509 | 0.0296 | 0.3038 | 10.2600 | 8.62E-12 |
| cluster3477 | -3277.3802 | 0.0155 | 0.0195 | 1.2642 | 9.58E-08 |
| cluster3478 | -3582.1278 | 0.0195 | 0.0619 | 3.1810 | 4.14E-13 |
| cluster3479 | -2916.4547 | 0.0481 | 0.3747 | 7.7865 | 0.0002 |
| cluster3480 | -4509.1582 | 0.0333 | 0.1737 | 5.2194 | 1.05E-10 |
| cluster3481 | -1072.3981 | 0.0068 | 0.1288 | 18.9884 | 0.001 |
| cluster3482 | -1967.528 | 0.0768 | 0.0963 | 1.2545 | 0.04 |
| cluster3483 | -3660.4904 | 0.0101 | 0.0433 | 4.2984 | 1.20E-18 |
| cluster3484 | -2769.5142 | 0.0306 | 0.1603 | 5.2410 | 9.23E-09 |
| cluster3485 | -3495.3233 | 0.0412 | 0.1588 | 3.8534 | 7.83E-06 |
| cluster3486 | -3175.1244 | 0.0505 | 0.0871 | 1.7236 | 8.41E-05 |
| cluster3487 | -2348.3355 | 0.0243 | 0.0933 | 3.8407 | 7.27E-07 |
| cluster3488 | -1483.4208 | 0.0278 | 0.0737 | 2.6431 | 0.0003 |
| cluster3489 | -4061.696 | 0.0447 | 0.2498 | 5.5892 | 7.30E-08 |
| cluster3490 | -3433.1788 | 0.0217 | 0.1654 | 7.6395 | 9.73E-12 |
| cluster3491 | -3687.206 | 0.0316 | 0.1271 | 4.0210 | 4.20E-09 |
| cluster3492 | -3372.4835 | 0.0375 | 0.2235 | 5.9650 | 2.23E-08 |
| cluster3493 | -4433.3787 | 0.0345 | 0.1238 | 3.5923 | 1.08E-10 |
| cluster3494 | -2865.9829 | 0.0138 | 0.1349 | 9.7459 | 1.27E-10 |
| cluster3495 | -2843.0887 | 0.0294 | 0.1081 | 3.6813 | 3.11E-08 |
| cluster3496 | -3231.788 | 0.0222 | 0.0621 | 2.8016 | 6.43E-11 |
| cluster3497 | -1194.8717 | 0.0135 | 0.0116 | 0.8520 | 0.02 |
| cluster3498 | -4134.1604 | 0.0483 | 0.3697 | 7.6606 | 6.09E-07 |
| cluster3499 | -3724.8658 | 0.0910 | 0.4720 | 5.1854 | 0.04 |
| cluster3500 | -3424.4386 | 0.0206 | 0.1050 | 5.0914 | 1.89E-08 |
| cluster3501 | -2359.7169 | 0.0269 | 0.0927 | 3.4528 | 3.03E-06 |
| cluster3502 | -2169.5073 | 0.0145 | 0.0780 | 5.3796 | 9.47E-10 |
| cluster3503 | -1952.3476 | 0.0464 | 0.2506 | 5.4049 | 0.001 |
| cluster3504 | -1142.0856 | 0.0078 | 0.0655 | 8.4304 | 7.92E-08 |
| cluster3505 | -2418.2936 | 0.0530 | 0.3663 | 6.9142 | 4.63E-05 |
| cluster3506 | -2550.0888 | 0.0235 | 0.1189 | 5.0573 | 8.85E-11 |
| cluster3507 | -5020.0705 | 0.0542 | 0.2633 | 4.8578 | 9.60E-09 |
| cluster3508 | -5634.2164 | 0.0295 | 0.3659 | 12.4132 | 9.48E-11 |
| cluster3509 | -2402.0262 | 0.0012 | 0.1073 | 90.4179 | 0.0002 |
| cluster3510 | -844.34131 | 0.1070 | 0.2106 | 1.9677 | 0.34 |

| | | | | | |
|---|---|---|---|---|---|
| cluster3511 | -1019.1598 | 0.0337 | 0.1021 | 3.0288 | 0.02 |
| cluster3512 | -2356.9804 | 0.0329 | 0.0699 | 2.1255 | 3.98E-05 |
| cluster3513 | -959.36347 | 0.0711 | 0.0289 | 0.4066 | 0.41 |
| cluster3514 | -5158.9193 | 0.0365 | 0.1183 | 3.2363 | 2.37E-12 |
| cluster3515 | -2605.3412 | 0.0134 | 0.0526 | 3.9282 | 9.11E-12 |
| cluster3516 | -4130.6213 | 0.0273 | 0.1190 | 4.3580 | 5.12E-12 |
| cluster3517 | -2410.3471 | 0.0025 | 0.0055 | 2.1983 | 5.25E-10 |
| cluster3518 | -4784.7089 | 0.0760 | 0.3595 | 4.7313 | 4.49E-05 |
| cluster3519 | -2127.2128 | 0.0418 | 0.1675 | 4.0093 | 2.95E-05 |
| cluster3520 | -1193.995 | 0.1493 | 0.3063 | 2.0515 | 0.27 |
| cluster3521 | -4224.3574 | 0.0376 | 0.2248 | 5.9776 | 4.02E-09 |
| cluster3522 | -4754.0622 | 0.0331 | 0.2447 | 7.3858 | 2.20E-10 |
| cluster3523 | -1657.3417 | 0.0298 | 0.1645 | 5.5265 | 3.69E-05 |
| cluster3524 | -1963.5727 | 0.0126 | 0.0220 | 1.7367 | 4.24E-06 |
| cluster3525 | -2817.1898 | 0.0458 | 0.1142 | 2.4958 | 8.05E-05 |
| cluster3526 | -2999.2043 | 0.0023 | 0.0175 | 7.5572 | 6.11E-24 |
| cluster3527 | -4937.5702 | 0.0474 | 0.3505 | 7.3867 | 1.89E-08 |
| cluster3528 | -1968.4071 | 0.0108 | 0.0259 | 2.4125 | 1.28E-07 |
| cluster3529 | -3742.861 | 0.0311 | 0.2230 | 7.1648 | 7.75E-10 |
| cluster3530 | -1954.2993 | 0.0300 | 0.1960 | 6.5277 | 2.84E-05 |
| cluster3531 | -2920.4097 | 0.0500 | 0.1708 | 3.4178 | 6.65E-06 |
| cluster3532 | -5557.7864 | 0.0621 | 0.2795 | 4.5003 | 1.97E-07 |
| cluster3533 | -2655.2642 | 0.0246 | 0.0505 | 2.0504 | 5.06E-07 |
| cluster3534 | -2560.5428 | 0.0647 | 0.0814 | 1.2585 | 0.01 |
| cluster3535 | -3289.9454 | 0.0218 | 0.2195 | 10.0563 | 3.49E-10 |
| cluster3536 | -3557.2465 | 0.0363 | 0.1078 | 2.9678 | 6.53E-09 |
| cluster3537 | -3667.4536 | 0.0071 | 0.0395 | 5.6014 | 2.17E-24 |
| cluster3538 | -1389.1437 | 0.0374 | 0.0616 | 1.6481 | 0.005 |
| cluster3539 | -2945.6814 | 0.0374 | 0.4145 | 11.0720 | 4.51E-06 |
| cluster3540 | -5364.1978 | 0.0909 | 0.5626 | 6.1898 | 0.0002 |
| cluster3541 | -3877.8657 | 0.0273 | 0.0891 | 3.2632 | 4.89E-11 |
| cluster3542 | -5427.6137 | 0.0602 | 0.4717 | 7.8362 | 1.41E-06 |
| cluster3543 | -4247.7525 | 0.0350 | 0.2932 | 8.3782 | 3.53E-09 |
| cluster3544 | -1088.813 | 0.0139 | 0.0225 | 1.6227 | 0.005 |
| cluster3545 | -5147.2057 | 0.0458 | 0.2886 | 6.3009 | 6.78E-10 |
| cluster3546 | -2680.0269 | 0.0226 | 0.1760 | 7.7878 | 1.07E-07 |
| cluster3547 | -1117.3584 | 0.0087 | 0.0098 | 1.1428 | 0.01 |
| cluster3548 | -1308.9736 | 0.0569 | 0.0465 | 0.8189 | 0.07 |
| cluster3549 | -3233.2799 | 0.0379 | 0.1201 | 3.1727 | 2.12E-08 |
| cluster3550 | -1224.136 | 0.0083 | 0.0094 | 1.1188 | 0.003 |
| cluster3551 | -2798.8758 | 0.0505 | 0.0763 | 1.5103 | 0.001 |
| cluster3552 | -2386.1276 | 0.0231 | 0.1457 | 6.3175 | 7.09E-08 |
| cluster3553 | -2405.5779 | 0.0328 | 0.1081 | 3.2928 | 1.66E-07 |
| cluster3554 | -3824.3981 | 0.0265 | 0.0781 | 2.9428 | 5.14E-10 |

| | | | | | |
|---|---|---|---|---|---|
| cluster3555 | -1633.639 | 0.0205 | 0.1328 | 6.4745 | 1.09E-06 |
| cluster3556 | -4206.1336 | 0.0253 | 0.1560 | 6.1775 | 6.88E-14 |
| cluster3557 | -1368.352 | 0.0608 | 0.5976 | 9.8352 | 0.08 |
| cluster3558 | -4777.2397 | 0.0666 | 0.3240 | 4.8669 | 5.51E-07 |
| cluster3559 | -2458.1321 | 0.0177 | 0.0935 | 5.2913 | 3.47E-08 |
| cluster3560 | -5096.7822 | 0.0588 | 0.3750 | 6.3747 | 1.16E-07 |
| cluster3561 | -1802.2589 | 0.0313 | 0.0548 | 1.7513 | 0.001 |
| cluster3562 | -4278.4323 | 0.0381 | 0.1982 | 5.2048 | 1.57E-10 |
| cluster3563 | -1982.7831 | 0.0148 | 0.0472 | 3.1794 | 1.99E-07 |
| cluster3564 | -5024.1543 | 0.0704 | 0.1796 | 2.5495 | 5.51E-07 |
| cluster3565 | -3351.4846 | 0.0364 | 0.2218 | 6.0998 | 1.87E-08 |
| cluster3566 | -1890.4123 | 0.0440 | 0.1547 | 3.5143 | 0.0002 |
| cluster3567 | -2403.9754 | 0.0669 | 0.2639 | 3.9438 | 0.005 |
| cluster3568 | -3168.28 | 0.0285 | 0.1917 | 6.7340 | 1.33E-07 |
| cluster3569 | -3551.6338 | 0.0405 | 0.2938 | 7.2495 | 1.93E-06 |
| cluster3570 | -2401.0568 | 0.0196 | 0.2227 | 11.3486 | 5.34E-07 |
| cluster3571 | -2036.9029 | 0.0177 | 0.1001 | 5.6713 | 4.67E-07 |
| cluster3572 | -3453.9091 | 0.0512 | 0.2670 | 5.2194 | 4.10E-06 |
| cluster3573 | -3244.3942 | 0.0116 | 0.0913 | 7.8846 | 1.87E-11 |
| cluster3574 | -5131.8072 | 0.0767 | 0.3878 | 5.0578 | 4.89E-06 |
| cluster3575 | -3831.1917 | 0.0397 | 0.1922 | 4.8434 | 8.67E-09 |
| cluster3576 | -3275.0948 | 0.0367 | 0.2055 | 5.6025 | 4.67E-07 |
| cluster3577 | -2731.4253 | 0.0393 | 0.0919 | 2.3441 | 8.71E-06 |
| cluster3578 | -3391.4835 | 0.0338 | 0.1524 | 4.5122 | 7.61E-08 |
| cluster3579 | -2924.2332 | 0.0261 | 0.1390 | 5.3233 | 4.27E-08 |
| cluster3580 | -2417.9393 | 0.0103 | 0.0906 | 8.7705 | 6.96E-12 |
| cluster3581 | -1247.0855 | 0.0077 | 0.0580 | 7.5637 | 3.03E-07 |
| cluster3582 | -1108.0896 | 0.0234 | 0.0209 | 0.8884 | 0.02 |
| cluster3583 | -2533.4362 | 0.0001 | 0.0005 | 4.5593 | 2.82E-18 |
| cluster3584 | -4869.8726 | 0.0308 | 0.2995 | 9.7298 | 6.96E-10 |
| cluster3585 | -2094.4591 | 0.0629 | 0.1054 | 1.6767 | 0.004 |
| cluster3586 | -5224.9747 | 0.0717 | 0.4001 | 5.5764 | 4.02E-06 |
| cluster3587 | -3470.5274 | 0.0522 | 0.4180 | 8.0094 | 3.73E-05 |
| cluster3588 | -3361.2893 | 0.0273 | 0.0507 | 1.8531 | 1.74E-07 |
| cluster3589 | -3018.0656 | 0.0434 | 0.1735 | 3.9976 | 1.01E-06 |
| cluster3590 | -4182.1755 | 0.0609 | 0.4881 | 8.0106 | 0.0002 |
| cluster3591 | -2874.4182 | 0.0120 | 0.0275 | 2.2954 | 1.46E-10 |
| cluster3592 | -4706.153 | 0.0389 | 0.2692 | 6.9210 | 2.53E-10 |
| cluster3593 | -3419.6968 | 0.0478 | 0.3199 | 6.6934 | 1.38E-07 |
| cluster3594 | -3177.8806 | 0.0609 | 0.1920 | 3.1525 | 0.0002 |
| cluster3595 | -2123.6496 | 0.0120 | 0.0288 | 2.3957 | 1.67E-08 |
| cluster3596 | -2316.2729 | 0.0331 | 0.1268 | 3.8342 | 1.08E-06 |
| cluster3597 | -4194.7778 | 0.0296 | 0.1094 | 3.7030 | 1.47E-11 |
| cluster3598 | -4253.8355 | 0.0231 | 0.4837 | 20.9792 | 8.55E-08 |

| | | | | | |
|---|---|---|---|---|---|
| cluster3599 | -2254.0603 | 0.0246 | 0.1148 | 4.6648 | 6.03E-08 |
| cluster3600 | -2861.3 | 0.0253 | 0.0747 | 2.9528 | 7.50E-08 |
| cluster3601 | -3284.9635 | 0.0031 | 0.0576 | 18.3317 | 1.28E-18 |
| cluster3602 | -3065.9593 | 0.0033 | 0.0069 | 2.0808 | 7.03E-14 |
| cluster3603 | -4224.6413 | 0.0526 | 0.2498 | 4.7542 | 1.16E-08 |
| cluster3604 | -1058.292 | 0.0631 | 0.0322 | 0.5111 | 0.24 |
| cluster3605 | -3057.8823 | 0.0380 | 0.1705 | 4.4808 | 5.02E-07 |
| cluster3606 | -4323.6371 | 0.0278 | 0.1901 | 6.8474 | 5.55E-13 |
| cluster3607 | -2597.1671 | 0.0365 | 0.2100 | 5.7552 | 2.09E-06 |
| cluster3608 | -2888.2067 | 0.0108 | 0.0353 | 3.2807 | 3.87E-12 |
| cluster3609 | -4182.2681 | 0.0297 | 0.2558 | 8.6137 | 1.77E-09 |
| cluster3610 | -1395.5598 | 0.0077 | 0.1083 | 14.0116 | 8.61E-05 |
| cluster3611 | -2340.884 | 0.0263 | 0.3183 | 12.0976 | 2.61E-05 |
| cluster3612 | -2831.634 | 0.0343 | 0.2819 | 8.2248 | 3.05E-07 |
| cluster3613 | -4059.3361 | 0.0183 | 0.0594 | 3.2486 | 2.09E-16 |
| cluster3614 | -4308.2333 | 0.0249 | 0.2896 | 11.6159 | 2.72E-09 |
| cluster3615 | -1663.3299 | 0.0470 | 0.0470 | 0.9979 | 0.01 |
| cluster3616 | -774.82574 | 0.0144 | 0.0476 | 3.3082 | 0.005 |
| cluster3617 | -1931.5757 | 0.0442 | 0.2168 | 4.9031 | 0.001 |
| cluster3618 | -1899.3801 | 0.1715 | 0.4927 | 2.8725 | 0.32 |
| cluster3619 | -2552.6021 | 0.0045 | 0.0226 | 5.0085 | 5.26E-16 |
| cluster3620 | -3605.2881 | 0.0514 | 0.1880 | 3.6563 | 1.72E-07 |
| cluster3621 | -4765.4964 | 0.0339 | 0.2856 | 8.4168 | 1.53E-12 |
| cluster3622 | -2724.2596 | 0.0126 | 0.0296 | 2.3501 | 5.25E-10 |
| cluster3623 | -2271.7601 | 0.0092 | 0.1243 | 13.5292 | 3.69E-09 |
| cluster3624 | -2909.9631 | 0.0138 | 0.1306 | 9.4572 | 2.09E-10 |
| cluster3625 | -3308.9197 | 0.0064 | 0.0580 | 9.0077 | 2.88E-17 |
| cluster3626 | -1983.3284 | 0.0293 | 0.0941 | 3.2088 | 1.22E-06 |
| cluster3627 | -3001.0824 | 0.0072 | 0.0350 | 4.8675 | 1.39E-19 |
| cluster3628 | -3023.1107 | 0.0278 | 0.2414 | 8.6870 | 3.06E-08 |
| cluster3629 | -4165.475 | 0.0119 | 0.1327 | 11.1853 | 1.36E-22 |
| cluster3630 | -1210.3912 | 0.0150 | 0.0236 | 1.5777 | 0.001 |
| cluster3631 | -3595.076 | 0.0152 | 0.1019 | 6.6869 | 2.10E-14 |
| cluster3632 | -2460.0197 | 0.0307 | 0.2279 | 7.4274 | 3.98E-07 |
| cluster3633 | -2505.3256 | 0.0080 | 0.0625 | 7.8381 | 2.70E-06 |
| cluster3634 | -939.33847 | 0.0154 | 0.0340 | 2.2046 | 0.003 |
| cluster3635 | -1621.136 | 0.0421 | 0.0358 | 0.8486 | 0.02 |
| cluster3636 | -2532.3699 | 0.0145 | 0.0677 | 4.6943 | 3.49E-10 |
| cluster3637 | -1062.5126 | 0.0160 | 0.0206 | 1.2910 | 0.01 |
| cluster3638 | -2732.7281 | 0.0320 | 0.0882 | 2.7583 | 4.18E-06 |
| cluster3639 | -1272.5784 | 0.0355 | 0.0853 | 2.4009 | 0.002 |
| cluster3640 | -4766.5751 | 0.0743 | 0.4632 | 6.2328 | 9.84E-05 |
| cluster3641 | -1769.1067 | 0.0264 | 0.0333 | 1.2605 | 0.0002 |
| cluster3642 | -3357.8126 | 0.0316 | 0.2133 | 6.7415 | 1.01E-07 |

| | | | | | |
|---|---|---|---|---|---|
| cluster3643 | -3538.3673 | 0.1016 | 0.4704 | 4.6286 | 0.003 |
| cluster3644 | -1913.1111 | 0.0289 | 0.0512 | 1.7703 | 9.35E-05 |
| cluster3645 | -1807.8985 | 0.0307 | 0.1669 | 5.4328 | 3.62E-06 |
| cluster3646 | -2636.2113 | 0.0215 | 0.2549 | 11.8420 | 5.92E-08 |
| cluster3647 | -3430.0126 | 0.0042 | 0.0809 | 19.4638 | 9.17E-25 |
| cluster3648 | -1759.5923 | 0.0014 | 0.0065 | 4.8223 | 3.68E-16 |
| cluster3649 | -1640.6161 | 0.0099 | 0.0307 | 3.1163 | 2.18E-06 |
| cluster3650 | -1258.004 | 0.0139 | 0.2354 | 16.9622 | 0.001 |
| cluster3651 | -3971.3629 | 0.0478 | 0.1770 | 3.7067 | 3.80E-07 |
| cluster3652 | -3310.7348 | 0.0251 | 0.0905 | 3.6090 | 1.11E-11 |
| cluster3653 | -1426.9852 | 0.0039 | 0.0109 | 2.8169 | 5.55E-09 |
| cluster3654 | -2838.0258 | 0.0095 | 0.0623 | 6.5513 | 2.71E-16 |
| cluster3655 | -2366.6571 | 0.0662 | 0.2776 | 4.1972 | 0.002 |
| cluster3656 | -3108.7937 | 0.0255 | 0.1036 | 4.0630 | 1.14E-10 |
| cluster3657 | -3194.4491 | 0.0150 | 0.1573 | 10.4782 | 2.00E-11 |
| cluster3658 | -753.7423 | 0.0867 | 0.1029 | 1.1867 | 0.27 |
| cluster3659 | -2269.8169 | 0.0357 | 0.0949 | 2.6637 | 3.03E-05 |
| cluster3660 | -3017.8106 | 0.0191 | 0.2256 | 11.8227 | 3.59E-11 |
| cluster3661 | -1819.0827 | 0.0019 | 0.0010 | 0.5697 | 0.01 |
| cluster3662 | -2804.2947 | 0.0363 | 0.1995 | 5.5021 | 3.28E-07 |
| cluster3663 | -3134.1494 | 0.0488 | 0.1944 | 3.9870 | 1.41E-05 |
| cluster3664 | -2449.4978 | 0.0397 | 0.3033 | 7.6481 | 2.25E-05 |
| cluster3665 | -3603.4135 | 0.0206 | 0.1434 | 6.9588 | 1.44E-14 |
| cluster3666 | -2711.4958 | 0.0349 | 0.1611 | 4.6175 | 1.04E-06 |
| cluster3667 | -1570.8251 | 0.0211 | 0.0313 | 1.4775 | 0.001 |
| cluster3668 | -2983.8432 | 0.0212 | 0.1060 | 5.0104 | 1.18E-11 |
| cluster3669 | -2601.9726 | 0.0368 | 0.1223 | 3.3259 | 4.71E-06 |
| cluster3670 | -4034.931 | 0.0460 | 0.1986 | 4.3192 | 1.08E-08 |
| cluster3671 | -3996.6827 | 0.0299 | 0.1912 | 6.4004 | 2.54E-10 |
| cluster3672 | -2500.7693 | 0.0449 | 0.1987 | 4.4279 | 1.86E-05 |
| cluster3673 | -4410.3565 | 0.0415 | 0.5507 | 13.2762 | 3.10E-05 |
| cluster3674 | -2025.4369 | 0.0276 | 0.2052 | 7.4457 | 4.70E-05 |
| cluster3675 | -660.30852 | 0.0001 | 0.0000 | 0.7115 | 0.01 |
| cluster3676 | -3526.7623 | 0.0534 | 0.2687 | 5.0370 | 5.31E-06 |
| cluster3677 | -812.08983 | 0.0221 | 0.0550 | 2.4871 | 0.01 |
| cluster3678 | -3307.6017 | 0.0346 | 0.1494 | 4.3129 | 2.61E-08 |
| cluster3679 | -944.67422 | 0.0084 | 0.0166 | 1.9882 | 0.0001 |
| cluster3680 | -2314.4553 | 0.0094 | 0.1050 | 11.1824 | 9.93E-14 |
| cluster3681 | -2959.2588 | 0.0196 | 0.1576 | 8.0642 | 1.52E-10 |
| cluster3682 | -4365.7462 | 0.0343 | 0.2473 | 7.2117 | 5.13E-12 |
| cluster3683 | -2805.3766 | 0.1021 | 0.4948 | 4.8456 | 0.02 |
| cluster3684 | -2993.2098 | 0.0348 | 0.1764 | 5.0723 | 5.00E-09 |
| cluster3685 | -3674.6318 | 0.0833 | 0.5190 | 6.2326 | 0.001 |
| cluster3686 | -1628.7042 | 0.0345 | 0.3824 | 11.0685 | 0.01 |

| | | | | | |
|---|---|---|---|---|---|
| cluster3687 | -2877.6706 | 0.0808 | 0.6804 | 8.4186 | 0.04 |
| cluster3688 | -2051.3421 | 0.0083 | 0.0749 | 9.0475 | 7.17E-11 |
| cluster3689 | -2642.1112 | 0.0391 | 0.1084 | 2.7732 | 2.68E-06 |
| cluster3690 | -1454.9317 | 0.0233 | 0.0864 | 3.7069 | 2.36E-05 |
| cluster3691 | -670.36032 | 0.0019 | 0.0219 | 11.2803 | 0.0002 |
| cluster3692 | -1435.3731 | 0.0110 | 0.0630 | 5.7472 | 8.12E-09 |
| cluster3693 | -2465.0545 | 0.0091 | 0.0935 | 10.2822 | 3.63E-17 |
| cluster3694 | -2092.3146 | 0.0303 | 0.0502 | 1.6588 | 0.002 |
| cluster3695 | -1737.1066 | 0.0247 | 0.1168 | 4.7274 | 2.12E-06 |
| cluster3696 | -817.96765 | 0.0105 | 0.1541 | 14.7304 | 0.003 |
| cluster3697 | -3584.1727 | 0.0723 | 0.2039 | 2.8217 | 0.0002 |
| cluster3698 | -1099.5724 | 0.0001 | 0.0000 | 1.4138 | 7.40E-05 |
| cluster3699 | -4441.8097 | 0.0468 | 0.3304 | 7.0519 | 1.12E-09 |
| cluster3700 | -2819.0929 | 0.0322 | 0.1528 | 4.7367 | 1.00E-07 |
| cluster3701 | -1128.5885 | 0.0011 | 0.0022 | 1.9473 | 2.72E-06 |
| cluster3702 | -3864.568 | 0.0262 | 0.2477 | 9.4744 | 2.54E-10 |
| cluster3703 | -1512.7048 | 0.0167 | 0.0293 | 1.7631 | 6.12E-05 |
| cluster3704 | -2230.317 | 0.0046 | 0.0555 | 12.0501 | 2.40E-15 |
| cluster3705 | -1945.2254 | 0.0136 | 0.0415 | 3.0552 | 6.95E-09 |
| cluster3706 | -2094.656 | 0.0209 | 0.0650 | 3.1115 | 2.45E-07 |
| cluster3707 | -1921.4939 | 0.0416 | 0.2986 | 7.1852 | 0.0002 |
| cluster3708 | -2636.53 | 0.0470 | 0.1761 | 3.7481 | 0.0005 |
| cluster3709 | -1580.4694 | 0.0787 | 0.0977 | 1.2410 | 0.04 |
| cluster3710 | -1924.8677 | 0.0145 | 0.1197 | 8.2624 | 2.44E-08 |
| cluster3711 | -1439.633 | 0.0291 | 0.0335 | 1.1521 | 0.03 |
| cluster3712 | -3402.6566 | 0.0271 | 0.0872 | 3.2144 | 2.64E-10 |
| cluster3713 | -4029.8902 | 0.0529 | 0.3786 | 7.1638 | 6.60E-07 |
| cluster3714 | -1775.8603 | 0.0463 | 0.0854 | 1.8442 | 0.003 |
| cluster3715 | -1286.9457 | 0.0134 | 0.2125 | 15.8817 | 1.09E-05 |
| cluster3716 | -3658.1424 | 0.0464 | 0.2775 | 5.9842 | 1.95E-07 |
| cluster3717 | -1403.9643 | 0.0142 | 0.0183 | 1.2923 | 0.001 |
| cluster3718 | -2990.8424 | 0.0213 | 0.1591 | 7.4780 | 2.74E-11 |
| cluster3719 | -3295.0357 | 0.0447 | 0.1684 | 3.7666 | 2.47E-06 |
| cluster3720 | -2758.9361 | 0.0263 | 0.0918 | 3.4874 | 2.10E-07 |
| cluster3721 | -1098.0568 | 0.0051 | 0.0039 | 0.7603 | 0.01 |
| cluster3722 | -1224.5824 | 0.0398 | 0.4572 | 11.4794 | 0.02 |
| cluster3723 | -989.94266 | 0.0001 | 0.0002 | 2.0506 | 2.10E-05 |
| cluster3724 | -3083.6075 | 0.0336 | 0.1942 | 5.7897 | 4.25E-09 |
| cluster3725 | -1839.95 | 0.0104 | 0.0793 | 7.6431 | 9.87E-11 |
| cluster3726 | -1936.6261 | 0.0486 | 0.1361 | 2.8027 | 0.0002 |
| cluster3727 | -2469.0453 | 0.0539 | 0.3486 | 6.4706 | 0.0003 |
| cluster3728 | -2157.0913 | 0.0210 | 0.0697 | 3.3216 | 2.00E-05 |
| cluster3729 | -2947.3377 | 0.0132 | 0.0909 | 6.8779 | 6.77E-12 |
| cluster3730 | -1782.056 | 0.0822 | 0.1086 | 1.3209 | 0.02 |

| | | | | | |
|---|---|---|---|---|---|
| cluster3731 | -2865.5328 | 0.0090 | 0.0275 | 3.0733 | 1.90E-13 |
| cluster3732 | -2822.9553 | 0.0228 | 0.0889 | 3.9045 | 2.60E-09 |
| cluster3733 | -3072.8629 | 0.0798 | 0.1706 | 2.1397 | 0.0005 |
| cluster3734 | -3043.4571 | 0.0434 | 0.2136 | 4.9264 | 6.13E-07 |
| cluster3735 | -3250.3703 | 0.0807 | 0.1849 | 2.2908 | 0.001 |
| cluster3736 | -3538.2993 | 0.0461 | 0.1340 | 2.9074 | 4.80E-07 |
| cluster3737 | -1209.5778 | 0.0030 | 0.0041 | 1.3457 | 0.0001 |
| cluster3738 | -3259.9057 | 0.0089 | 0.0349 | 3.9098 | 3.59E-19 |
| cluster3739 | -1156.2535 | 0.0001 | 0.0001 | 1.2770 | 0.001 |
| cluster3740 | -3450.4368 | 0.0301 | 0.1610 | 5.3589 | 4.13E-10 |
| cluster3741 | -1312.7963 | 0.0166 | 0.0351 | 2.1220 | 7.58E-05 |
| cluster3742 | -1615.8355 | 0.0564 | 0.2405 | 4.2641 | 0.001 |
| cluster3743 | -2980.39 | 0.0238 | 0.1306 | 5.4950 | 1.23E-09 |
| cluster3744 | -1356.1217 | 0.1064 | 0.2442 | 2.2949 | 0.16 |
| cluster3745 | -2585.3345 | 0.0251 | 0.1562 | 6.2300 | 1.21E-08 |
| cluster3746 | -3786.2235 | 0.0381 | 0.1775 | 4.6565 | 1.99E-08 |
| cluster3747 | -1121.3006 | 0.0274 | 0.0930 | 3.3971 | 0.0002 |
| cluster3748 | -2166.698 | 0.0142 | 0.0362 | 2.5503 | 1.27E-08 |
| cluster3749 | -3071.6507 | 0.0176 | 0.1079 | 6.1409 | 2.27E-14 |
| cluster3750 | -1355.4384 | 0.0243 | 0.2288 | 9.4019 | 4.91E-05 |
| cluster3751 | -3012.0054 | 0.0046 | 0.0216 | 4.6808 | 1.31E-19 |
| cluster3752 | -2663.4966 | 0.0079 | 0.0660 | 8.4185 | 1.28E-13 |
| cluster3753 | -1023.3745 | 0.0195 | 0.0106 | 0.5457 | 0.13 |
| cluster3754 | -881.97724 | 0.0001 | 0.0000 | 0.8180 | 0.01 |
| cluster3755 | -2499.8083 | 0.0158 | 0.0636 | 4.0394 | 9.60E-12 |
| cluster3756 | -1689.7931 | 0.0357 | 0.1513 | 4.2390 | 2.85E-05 |
| cluster3757 | -2091.3825 | 0.0455 | 0.0992 | 2.1801 | 0.0001 |
| cluster3758 | -1511.7918 | 0.0290 | 0.0755 | 2.6017 | 0.0005 |
| cluster3759 | -3066.4698 | 0.0682 | 0.1922 | 2.8166 | 0.0004 |
| cluster3760 | -2600.8759 | 0.0226 | 0.1185 | 5.2541 | 4.87E-08 |
| cluster3761 | -767.99285 | 0.0043 | 0.1177 | 27.6443 | 1.42E-05 |
| cluster3762 | -1968.0081 | 0.0166 | 0.0243 | 1.4658 | 5.78E-05 |
| cluster3763 | -2005.5732 | 0.0557 | 0.0735 | 1.3207 | 0.01 |
| cluster3764 | -1629.8247 | 0.0302 | 0.2133 | 7.0648 | 5.34E-05 |
| cluster3765 | -2670.9464 | 0.0187 | 0.0762 | 4.0884 | 5.06E-09 |
| cluster3766 | -1795.6306 | 0.0115 | 0.0610 | 5.2954 | 1.06E-07 |
| cluster3767 | -862.69539 | 0.0242 | 0.1694 | 6.9996 | 0.001 |
| cluster3768 | -2219.4703 | 0.0421 | 0.2915 | 6.9235 | 0.01 |
| cluster3769 | -1765.3215 | 0.0071 | 0.0103 | 1.4456 | 4.55E-05 |
| cluster3770 | -1775.1472 | 0.0531 | 0.1965 | 3.6999 | 0.002 |
| cluster3771 | -2139.0494 | 0.0703 | 0.3094 | 4.4034 | 0.005 |
| cluster3772 | -725.3054 | 0.0009 | 0.0881 | 95.9994 | 0.01 |
| cluster3773 | -1118.2878 | 0.1920 | 0.1802 | 0.9379 | 0.60 |
| cluster3774 | -1409.1588 | 0.0135 | 0.0139 | 1.0271 | 0.003 |

| | | | | | |
|---|---|---|---|---|---|
| cluster3775 | -906.73253 | 0.0128 | 0.1002 | 7.8614 | 0.001 |
| cluster3776 | -2518.6646 | 0.0780 | 0.1517 | 1.9447 | 0.001 |
| cluster3777 | -3067.0273 | 0.0589 | 0.3120 | 5.2991 | 1.71E-05 |
| cluster3778 | -980.75092 | 0.0012 | 0.0049 | 3.9807 | 1.13E-06 |
| cluster3779 | -2995.1667 | 0.0221 | 0.1566 | 7.0775 | 9.26E-10 |
| cluster3780 | -1685.5833 | 0.0121 | 0.1108 | 9.1674 | 1.42E-09 |
| cluster3781 | -2686.3977 | 0.0573 | 0.1793 | 3.1283 | 0.0001 |
| cluster3782 | -2290.3928 | 0.0430 | 0.1316 | 3.0637 | 1.42E-05 |
| cluster3783 | -2108.275 | 0.0072 | 0.1412 | 19.5996 | 8.71E-14 |
| cluster3784 | -2951.8023 | 0.0484 | 0.2239 | 4.6215 | 3.75E-06 |
| cluster3785 | -2689.595 | 0.0166 | 0.0499 | 3.0147 | 1.54E-08 |
| cluster3786 | -2267.0149 | 0.0086 | 0.1228 | 14.3144 | 8.52E-10 |
| cluster3787 | -1522.4369 | 0.0275 | 0.1030 | 3.7480 | 4.79E-05 |
| cluster3788 | -1160.4015 | 0.0001 | 0.0000 | 0.4121 | 0.11 |
| cluster3789 | -3041.047 | 0.0112 | 0.0261 | 2.3517 | 1.23E-10 |
| cluster3790 | -2177.5489 | 0.0136 | 0.1318 | 9.6679 | 1.61E-08 |
| cluster3791 | -420.104 | 0.0132 | 0.0242 | 1.8305 | 0.02 |
| cluster3792 | -1863.3952 | 0.0060 | 0.0149 | 2.4745 | 1.97E-08 |
| cluster3793 | -1853.4001 | 0.0275 | 0.0999 | 3.6366 | 4.51E-06 |
| cluster3794 | -2189.5782 | 0.0159 | 0.0946 | 5.9346 | 2.41E-09 |
| cluster3795 | -1701.6617 | 0.0195 | 0.0875 | 4.4918 | 5.78E-06 |
| cluster3796 | -1251.9353 | 0.1268 | 0.3717 | 2.9301 | 0.15 |
| cluster3797 | -3913.385 | 0.0549 | 0.5882 | 10.7141 | 0.0001 |
| cluster3798 | -3447.1277 | 0.0242 | 0.1323 | 5.4771 | 2.20E-13 |
| cluster3799 | -2341.5452 | 0.0467 | 0.1637 | 3.5072 | 0.0001 |
| cluster3800 | -431.36581 | 0.0109 | 0.0045 | 0.4136 | 0.54 |
| cluster3801 | -2690.6244 | 0.2061 | 0.4238 | 2.0555 | 0.34 |
| cluster3802 | -2649.9452 | 0.0340 | 0.2088 | 6.1345 | 2.00E-06 |
| cluster3803 | -2239.9663 | 0.0355 | 0.1351 | 3.8094 | 6.68E-06 |
| cluster3804 | -2635.6016 | 0.0320 | 0.1344 | 4.2037 | 1.78E-06 |
| cluster3805 | -2655.9729 | 0.0237 | 0.1300 | 5.4907 | 2.06E-09 |
| cluster3806 | -3567.0081 | 0.0386 | 0.2077 | 5.3870 | 8.27E-09 |
| cluster3807 | -1441.9244 | 0.0075 | 0.0118 | 1.5629 | 1.49E-05 |
| cluster3808 | -1826.0208 | 0.0322 | 0.1865 | 5.7960 | 1.12E-05 |
| cluster3809 | -2361.389 | 0.0093 | 0.0393 | 4.2224 | 9.59E-12 |
| cluster3810 | -2464.596 | 0.0096 | 0.0483 | 5.0214 | 3.23E-13 |
| cluster3811 | -761.71599 | 0.0250 | 0.0229 | 0.9166 | 0.07 |
| cluster3812 | -3000.8509 | 0.0302 | 0.0989 | 3.2748 | 2.73E-07 |
| cluster3813 | -3413.7363 | 0.0350 | 0.3920 | 11.2037 | 1.85E-06 |
| cluster3814 | -1609.0325 | 0.0889 | 0.0889 | 1.0004 | 0.11 |
| cluster3815 | -1708.6144 | 0.0190 | 0.1695 | 8.9184 | 2.91E-07 |
| cluster3816 | -2526.7472 | 0.0267 | 0.2030 | 7.5926 | 7.55E-09 |
| cluster3817 | -992.0692 | 0.0063 | 0.0094 | 1.4945 | 0.001 |
| cluster3818 | -1979.1024 | 0.0626 | 0.3301 | 5.2752 | 0.002 |

| | | | | | |
|---|---|---|---|---|---|
| cluster3819 | -3347.2182 | 0.0275 | 0.1339 | 4.8693 | 1.88E-10 |
| cluster3820 | -2225.8035 | 0.0299 | 0.1293 | 4.3242 | 8.04E-07 |
| cluster3821 | -2243.7508 | 0.0162 | 0.0608 | 3.7560 | 1.21E-07 |
| cluster3822 | -3399.9441 | 0.0286 | 0.1896 | 6.6261 | 7.06E-10 |
| cluster3823 | -3408.8796 | 0.0198 | 0.1304 | 6.5796 | 3.33E-14 |
| cluster3824 | -2354.2077 | 0.0247 | 0.1974 | 7.9766 | 4.41E-08 |
| cluster3825 | -1388.908 | 0.0065 | 0.0249 | 3.8092 | 2.50E-08 |
| cluster3826 | -3155.184 | 0.0740 | 0.4325 | 5.8442 | 0.002 |
| cluster3827 | -2403.7031 | 0.0278 | 0.1259 | 4.5209 | 6.56E-08 |
| cluster3828 | -2365.2484 | 0.0478 | 0.1831 | 3.8283 | 2.38E-05 |
| cluster3829 | -2257.3749 | 0.0237 | 0.1421 | 5.9975 | 6.47E-07 |
| cluster3830 | -1629.3662 | 0.0203 | 0.0348 | 1.7125 | 0.002 |
| cluster3831 | -2348.5286 | 0.0315 | 0.2379 | 7.5500 | 1.73E-06 |
| cluster3832 | -1768.6816 | 0.0044 | 0.0158 | 3.5843 | 1.08E-10 |
| cluster3833 | -2130.5021 | 0.0229 | 0.1069 | 4.6607 | 2.16E-09 |
| cluster3834 | -2059.3846 | 0.0099 | 0.1056 | 10.6840 | 4.38E-09 |
| cluster3835 | -2118.3109 | 0.0344 | 0.1331 | 3.8698 | 1.09E-05 |
| cluster3836 | -1895.0445 | 0.0173 | 0.0718 | 4.1560 | 1.60E-07 |
| cluster3837 | -2639.6853 | 0.0410 | 0.1830 | 4.4598 | 5.68E-07 |
| cluster3838 | -2313.0197 | 0.0385 | 0.1352 | 3.5119 | 6.35E-07 |
| cluster3839 | -1500.5511 | 0.0193 | 0.1087 | 5.6422 | 7.16E-07 |
| cluster3840 | -2024.6286 | 0.0045 | 0.3183 | 70.5400 | 1.48E-05 |
| cluster3841 | -2293.1704 | 0.0755 | 0.4638 | 6.1386 | 0.01 |
| cluster3842 | -1606.9851 | 0.0035 | 0.0041 | 1.1437 | 4.85E-05 |
| cluster3843 | -1441.3701 | 0.0117 | 0.0328 | 2.8235 | 8.81E-07 |
| cluster3844 | -1078.532 | 0.0662 | 0.1636 | 2.4731 | 0.14 |
| cluster3845 | -604.91636 | 0.0001 | 0.0000 | 0.9374 | 0.01 |
| cluster3846 | -1273.9258 | 0.0173 | 0.0598 | 3.4496 | 1.43E-05 |
| cluster3847 | -1156.4518 | 0.0011 | 0.0073 | 6.8218 | 1.29E-12 |
| cluster3848 | -2517.7349 | 0.0317 | 0.2014 | 6.3521 | 2.06E-07 |
| cluster3849 | -2665.1246 | 0.0151 | 0.2520 | 16.6919 | 6.07E-07 |
| cluster3850 | -1678.0715 | 0.0216 | 0.0663 | 3.0709 | 0.0004 |
| cluster3851 | -1618.1867 | 0.0098 | 0.0489 | 5.0017 | 5.28E-09 |
| cluster3852 | -1856.843 | 0.0066 | 0.0550 | 8.2830 | 1.78E-13 |
| cluster3853 | -1101.7646 | 0.0048 | 0.0502 | 10.4091 | 2.69E-08 |
| cluster3854 | -1358.1683 | 0.1001 | 0.2456 | 2.4547 | 0.09 |
| cluster3855 | -2930.9408 | 0.0103 | 0.1496 | 14.5380 | 1.42E-16 |
| cluster3856 | -1696.502 | 0.0185 | 0.1075 | 5.8216 | 2.62E-08 |
| cluster3857 | -3038.0536 | 0.0167 | 0.0989 | 5.9307 | 1.73E-14 |
| cluster3858 | -2416.7328 | 0.0487 | 0.1837 | 3.7724 | 4.02E-05 |
| cluster3859 | -1977.6195 | 0.0131 | 0.0514 | 3.9131 | 1.94E-08 |
| cluster3860 | -2625.4713 | 0.0821 | 0.1851 | 2.2556 | 0.004 |
| cluster3861 | -1301.0629 | 0.0171 | 0.0884 | 5.1562 | 6.48E-06 |
| cluster3862 | -922.33901 | 0.0001 | 0.0003 | 3.1963 | 6.62E-07 |

| | | | | | |
|---|---|---|---|---|---|
| cluster3863 | -1483.3861 | 0.0011 | 0.0037 | 3.3475 | 6.40E-10 |
| cluster3864 | -2220.0883 | 0.0038 | 0.0364 | 9.5409 | 4.31E-19 |
| cluster3865 | -3004.3258 | 0.0452 | 0.3108 | 6.8816 | 9.64E-06 |
| cluster3866 | -1633.4289 | 0.0180 | 0.1403 | 7.7794 | 1.35E-06 |
| cluster3867 | -608.47114 | 0.0001 | 0.0000 | 0.3149 | 0.29 |
| cluster3868 | -1393.7676 | 0.0279 | 0.0630 | 2.2583 | 0.0005 |
| cluster3869 | -2975.6186 | 0.0312 | 0.3724 | 11.9276 | 2.00E-06 |
| cluster3870 | -2462.7396 | 0.0402 | 0.1762 | 4.3858 | 5.59E-07 |
| cluster3871 | -2152.8545 | 0.0075 | 0.0379 | 5.0420 | 3.44E-13 |
| cluster3872 | -2649.6537 | 0.0120 | 0.1125 | 9.3445 | 8.04E-13 |
| cluster3873 | -1788.1612 | 0.0288 | 0.0980 | 3.4014 | 1.85E-05 |
| cluster3874 | -2102.0954 | 0.0135 | 0.0190 | 1.4122 | 8.00E-06 |
| cluster3875 | -770.52529 | 0.0028 | 0.0071 | 2.5690 | 0.0003 |
| cluster3876 | -745.95459 | 0.0109 | 0.0236 | 2.1611 | 0.002 |
| cluster3877 | -1436.1233 | 0.0141 | 0.0313 | 2.2196 | 2.47E-05 |
| cluster3879 | -1881.4762 | 0.0178 | 0.0742 | 4.1685 | 9.52E-08 |
| cluster3880 | -1357.7125 | 0.0205 | 0.1150 | 5.6203 | 0.0002 |
| cluster3881 | -3017.8381 | 0.0475 | 0.3515 | 7.3942 | 3.43E-05 |
| cluster3882 | -2690.5422 | 0.0247 | 0.1390 | 5.6386 | 8.33E-09 |
| cluster3883 | -423.32636 | 0.0051 | 0.4782 | 94.3822 | 0.06 |
| cluster3884 | -1563.7138 | 0.0004 | 0.0691 | 167.8099 | 2.75E-11 |
| cluster3885 | -2858.3022 | 0.0139 | 0.0752 | 5.4238 | 1.78E-14 |
| cluster3886 | -1395.0378 | 0.0033 | 0.0554 | 16.8037 | 3.45E-10 |
| cluster3887 | -1628.4082 | 0.0089 | 0.1299 | 14.6629 | 1.04E-07 |
| cluster3888 | -1182.6967 | 0.0107 | 0.1916 | 17.8395 | 3.56E-06 |
| cluster3889 | -1004.6838 | 0.0239 | 0.1812 | 7.5929 | 0.001 |
| cluster3890 | -1871.9575 | 0.0172 | 0.1280 | 7.4600 | 3.94E-07 |
| cluster3891 | -2184.7662 | 0.0141 | 0.0579 | 4.0978 | 9.09E-11 |
| cluster3892 | -1374.4793 | 0.0146 | 0.0337 | 2.3034 | 0.001 |
| cluster3893 | -1458.9619 | 0.0251 | 0.0705 | 2.8095 | 5.37E-05 |
| cluster3894 | -966.60253 | 0.0079 | 0.0809 | 10.2574 | 7.75E-05 |
| cluster3895 | -1982.8364 | 0.0093 | 0.0684 | 7.3398 | 1.08E-08 |
| cluster3896 | -820.69134 | 0.0032 | 0.0238 | 7.4860 | 8.61E-06 |
| cluster3897 | -1452.62 | 0.0060 | 0.2283 | 38.3673 | 6.30E-06 |
| cluster3898 | -2390.4346 | 0.0246 | 0.0372 | 1.5151 | 6.60E-06 |
| cluster3899 | -3003.2615 | 0.0275 | 0.2427 | 8.8303 | 1.06E-08 |
| cluster3900 | -2364.8813 | 0.0709 | 0.5694 | 8.0305 | 0.05 |
| cluster3901 | -1544.2318 | 0.0023 | 0.0150 | 6.5731 | 1.37E-14 |
| cluster3902 | -2496.4033 | 0.0114 | 0.0873 | 7.6758 | 1.09E-13 |
| cluster3903 | -1623.5663 | 0.0105 | 0.1515 | 14.4578 | 3.76E-07 |
| cluster3904 | -905.32905 | 0.0053 | 0.0719 | 13.5669 | 0.0004 |
| cluster3905 | -1594.9338 | 0.1304 | 0.4029 | 3.0893 | 0.11 |
| cluster3906 | -1453.7143 | 0.0211 | 0.0774 | 3.6767 | 5.62E-05 |
| cluster3907 | -1945.1137 | 0.0412 | 0.1612 | 3.9093 | 7.69E-06 |

| | | | | | |
|---|---|---|---|---|---|
| cluster3908 | -1848.2107 | 0.0141 | 0.0778 | 5.5326 | 1.68E-06 |
| cluster3909 | -2465.7354 | 0.0241 | 0.2016 | 8.3765 | 1.54E-07 |
| cluster3910 | -1193.4564 | 0.0361 | 0.2421 | 6.7007 | 0.002 |
| cluster3911 | -1749.9458 | 0.0085 | 0.0636 | 7.4879 | 3.75E-10 |
| cluster3912 | -951.76626 | 0.0001 | 0.0000 | 1.5462 | 0.0001 |
| cluster3913 | -2544.9987 | 0.0386 | 0.2117 | 5.4849 | 6.23E-06 |
| cluster3914 | -2239.1115 | 0.0569 | 0.3811 | 6.6929 | 0.004 |
| cluster3915 | -2227.4502 | 0.0345 | 0.1905 | 5.5298 | 1.83E-06 |
| cluster3916 | -902.6965 | 0.0795 | 0.2586 | 3.2541 | 0.08 |
| cluster3917 | -2583.1928 | 0.0125 | 0.0813 | 6.4895 | 8.27E-10 |
| cluster3918 | -2759.0807 | 0.0176 | 0.2635 | 14.9838 | 4.03E-08 |
| cluster3919 | -570.0868 | 0.0019 | 0.0031 | 1.6999 | 0.10 |
| cluster3920 | -657.92818 | 0.0109 | 0.4816 | 44.0217 | 0.02 |
| cluster3921 | -2190.7369 | 0.0254 | 0.1270 | 5.0093 | 4.17E-06 |
| cluster3922 | -1240.3656 | 0.0519 | 0.1914 | 3.6893 | 0.01 |
| cluster3923 | -1150.4881 | 0.0004 | 0.0047 | 12.1407 | 1.75E-13 |
| cluster3924 | -1696.2972 | 0.0197 | 0.1020 | 5.1751 | 2.44E-06 |
| cluster3925 | -1093.5828 | 0.0494 | 0.1904 | 3.8532 | 0.01 |
| cluster3926 | -2437.4396 | 0.0451 | 0.4152 | 9.2134 | 0.004 |
| cluster3927 | -1188.1979 | 0.0107 | 0.0649 | 6.0661 | 1.02E-06 |
| cluster3928 | -1159.9465 | 0.0291 | 0.0708 | 2.4308 | 0.002 |
| cluster3929 | -1608.2226 | 0.0060 | 0.0437 | 7.2263 | 2.65E-11 |
| cluster3930 | -2017.8275 | 0.0156 | 0.0423 | 2.7101 | 1.12E-08 |
| cluster3931 | -1830.2079 | 0.1364 | 0.1698 | 1.2448 | 0.26 |
| cluster3932 | -1117.9005 | 0.0110 | 0.0138 | 1.2501 | 0.01 |
| cluster3933 | -1158.492 | 0.0865 | 0.1490 | 1.7216 | 0.13 |
| cluster3934 | -717.10761 | 0.0001 | 0.0002 | 2.3620 | 0.002 |
| cluster3935 | -3108.9888 | 0.0590 | 0.4103 | 6.9593 | 0.0003 |
| cluster3936 | -2135.467 | 0.0172 | 0.1302 | 7.5855 | 4.43E-08 |
| cluster3937 | -1939.0909 | 0.0200 | 0.0571 | 2.8514 | 2.33E-08 |
| cluster3938 | -1782.3333 | 0.0324 | 0.0963 | 2.9730 | 3.36E-05 |
| cluster3939 | -2056.004 | 0.0156 | 0.0874 | 5.6116 | 3.05E-09 |
| cluster3940 | -1338.5646 | 0.0178 | 0.0375 | 2.1051 | 0.0001 |
| cluster3941 | -970.19299 | 0.0301 | 0.0462 | 1.5382 | 0.04 |
| cluster3942 | -2009.6116 | 0.0314 | 0.0876 | 2.7846 | 1.33E-05 |
| cluster3943 | -2268.5583 | 0.0159 | 0.1664 | 10.4380 | 8.08E-10 |
| cluster3944 | -1593.4194 | 0.0458 | 0.2078 | 4.5373 | 0.0003 |
| cluster3945 | -1676.6894 | 0.0380 | 0.2026 | 5.3267 | 0.0002 |
| cluster3946 | -1806.1475 | 0.0257 | 0.1328 | 5.1621 | 4.92E-06 |
| cluster3947 | -2286.0571 | 0.0283 | 0.1764 | 6.2390 | 1.37E-07 |
| cluster3948 | -1015.9267 | 0.0305 | 0.0874 | 2.8665 | 0.003 |
| cluster3949 | -878.24883 | 0.0093 | 0.0922 | 9.9049 | 0.001 |
| cluster3950 | -1743.4762 | 0.0014 | 0.1192 | 86.3360 | 1.13E-08 |
| cluster3951 | -2834.2857 | 0.0392 | 0.2201 | 5.6133 | 1.60E-07 |

| | | | | | |
|---|---|---|---|---|---|
| cluster3952 | -1474.9503 | 0.0031 | 0.0280 | 9.0728 | 1.28E-11 |
| cluster3953 | -1363.2426 | 0.0060 | 0.0545 | 9.1713 | 0.0001 |
| cluster3954 | -1806.6903 | 0.0280 | 0.1022 | 3.6528 | 4.49E-05 |
| cluster3955 | -1192.2205 | 0.0252 | 0.1592 | 6.3221 | 0.0004 |
| cluster3956 | -1995.7935 | 0.0144 | 0.1120 | 7.8000 | 1.07E-09 |
| cluster3957 | -1959.4803 | 0.0123 | 0.1011 | 8.2193 | 2.84E-10 |
| cluster3958 | -1104.8632 | 0.0229 | 0.1107 | 4.8261 | 0.0004 |
| cluster3959 | -2686.1877 | 0.0340 | 0.1110 | 3.2701 | 2.73E-07 |
| cluster3960 | -2424.5799 | 0.0576 | 0.3029 | 5.2565 | 0.003 |
| cluster3961 | -1594.9815 | 0.0378 | 0.1990 | 5.2617 | 0.001 |
| cluster3962 | -1533.3978 | 0.0415 | 0.1769 | 4.2625 | 0.001 |
| cluster3963 | -2078.2859 | 0.0403 | 0.1963 | 4.8757 | 1.12E-05 |
| cluster3964 | -2999.5551 | 0.0621 | 0.8362 | 13.4730 | 0.03 |
| cluster3965 | -1516.852 | 0.0342 | 0.2707 | 7.9082 | 0.0002 |
| cluster3966 | -1433.0606 | 0.0239 | 0.0749 | 3.1312 | 7.49E-05 |
| cluster3967 | -1753.9046 | 0.0872 | 0.1652 | 1.8951 | 0.03 |
| cluster3968 | -1167.7163 | 0.0174 | 0.0756 | 4.3398 | 4.39E-05 |
| cluster3969 | -899.56818 | 0.0001 | 0.0001 | 1.3668 | 0.0005 |
| cluster3970 | -2484.4824 | 0.0455 | 0.2671 | 5.8725 | 0.0002 |
| cluster3971 | -1690.2917 | 0.0365 | 0.1911 | 5.2390 | 6.20E-05 |
| cluster3972 | -990.23328 | 0.0139 | 0.0739 | 5.3121 | 3.89E-05 |
| cluster3973 | -1420.1051 | 0.0094 | 0.0336 | 3.5784 | 0.0004 |
| cluster3974 | -1832.8513 | 0.0180 | 0.0801 | 4.4568 | 5.34E-07 |
| cluster3975 | -1271.1908 | 0.0269 | 0.1129 | 4.2005 | 0.001 |
| cluster3976 | -1581.4275 | 0.0107 | 0.0624 | 5.8282 | 1.45E-06 |
| cluster3977 | -2230.1594 | 0.0974 | 0.3722 | 3.8200 | 0.02 |
| cluster3978 | -887.87911 | 0.0061 | 0.0151 | 2.5044 | 0.0003 |
| cluster3979 | -1621.1723 | 0.0021 | 0.0089 | 4.2242 | 9.37E-13 |
| cluster3980 | -922.24867 | 0.0001 | 0.0000 | 0.9224 | 0.01 |
| cluster3981 | -1981.0157 | 0.0203 | 0.1048 | 5.1633 | 1.20E-06 |
| cluster3982 | -1829.9174 | 0.0493 | 0.2136 | 4.3361 | 0.0004 |
| cluster3983 | -2677.4219 | 0.0339 | 0.2937 | 8.6601 | 2.82E-06 |
| cluster3984 | -1382.3379 | 0.0429 | 0.3446 | 8.0384 | 0.02 |
| cluster3985 | -979.26464 | 0.0120 | 0.0248 | 2.0617 | 0.0001 |
| cluster3986 | -1734.3032 | 0.0168 | 0.1533 | 9.1027 | 1.54E-07 |
| cluster3987 | -2087.2294 | 0.0122 | 0.0654 | 5.3548 | 6.61E-10 |
| cluster3988 | -1245.8627 | 0.0094 | 0.0327 | 3.4841 | 7.72E-06 |
| cluster3989 | -1835.0553 | 0.0550 | 0.2479 | 4.5066 | 0.002 |
| cluster3990 | -2208.6202 | 0.0248 | 0.2025 | 8.1692 | 1.07E-07 |
| cluster3991 | -1522.0949 | 0.0207 | 0.1315 | 6.3498 | 1.81E-06 |
| cluster3992 | -1889.1843 | 0.0368 | 0.0887 | 2.4118 | 0.0002 |
| cluster3993 | -1675.0258 | 0.0344 | 0.1928 | 5.6055 | 0.0002 |
| cluster3994 | -2395.1223 | 0.0580 | 0.2757 | 4.7519 | 0.0004 |
| cluster3995 | -1619.4788 | 0.0365 | 0.1613 | 4.4233 | 0.0002 |

| | | | | | |
|---|---|---|---|---|---|
| cluster3996 | -1708.9928 | 0.0064 | 0.0696 | 10.8597 | 6.40E-11 |
| cluster3997 | -2518.9408 | 0.0538 | 0.2750 | 5.1152 | 0.0001 |
| cluster3998 | -1416.8555 | 0.0540 | 0.1856 | 3.4394 | 0.003 |
| cluster3999 | -2525.0438 | 0.0317 | 0.2255 | 7.1231 | 4.52E-06 |
| cluster4000 | -2290.4198 | 0.0470 | 0.1851 | 3.9397 | 1.01E-05 |
| cluster4001 | -1739.6001 | 0.0190 | 0.0951 | 5.0150 | 7.69E-07 |
| cluster4002 | -1802.1705 | 0.0042 | 0.0691 | 16.6692 | 1.53E-12 |
| cluster4003 | -1421.8275 | 0.0752 | 0.3145 | 4.1823 | 0.01 |
| cluster4004 | -1442.9096 | 0.0080 | 0.0315 | 3.9330 | 1.09E-08 |
| cluster4005 | -2068.3079 | 0.0465 | 0.3446 | 7.4067 | 0.0003 |
| cluster4006 | -1089.5732 | 0.0320 | 0.1046 | 3.2686 | 0.001 |
| cluster4007 | -1425.9581 | 0.0026 | 0.0096 | 3.7062 | 7.58E-11 |
| cluster4008 | -1355.8221 | 0.0078 | 0.0794 | 10.2346 | 8.41E-09 |
| cluster4009 | -1546.5411 | 0.0133 | 0.1372 | 10.3058 | 1.02E-05 |
| cluster4010 | -1627.7451 | 0.0556 | 0.4005 | 7.2024 | 0.01 |
| cluster4011 | -1061.4987 | 0.0693 | 0.0581 | 0.8371 | 0.16 |
| cluster4012 | -2446.933 | 0.0151 | 0.5214 | 34.6538 | 5.46E-06 |
| cluster4013 | -846.68108 | 0.0235 | 0.0526 | 2.2371 | 0.002 |
| cluster4014 | -1138.0748 | 0.0063 | 0.0355 | 5.6705 | 3.91E-06 |
| cluster4015 | -618.62931 | 0.0273 | 0.0071 | 0.2576 | 0.47 |
| cluster4016 | -1815.7295 | 0.0190 | 0.0879 | 4.6257 | 2.70E-08 |
| cluster4017 | -1342.2874 | 0.0419 | 0.2230 | 5.3243 | 0.002 |
| cluster4018 | -1325.8329 | 0.1430 | 0.6281 | 4.3920 | 0.27 |
| cluster4019 | -1826.2203 | 0.0668 | 0.3682 | 5.5101 | 0.01 |
| cluster4020 | -1005.3964 | 0.0361 | 0.0731 | 2.0308 | 0.01 |
| cluster4021 | -1593.3152 | 0.0268 | 0.0649 | 2.4177 | 4.16E-05 |
| cluster4022 | -970.31533 | 0.0092 | 0.0278 | 3.0300 | 0.002 |
| cluster4023 | -1844.5292 | 0.0654 | 0.2344 | 3.5812 | 0.003 |
| cluster4024 | -2359.8226 | 0.0299 | 0.2808 | 9.3865 | 1.72E-05 |
| cluster4025 | -624.40744 | 0.0006 | 0.0074 | 11.8520 | 3.78E-05 |
| cluster4026 | -1088.9947 | 0.0026 | 0.0345 | 13.0745 | 6.58E-07 |
| cluster4027 | -980.16405 | 0.0052 | 0.0645 | 12.3573 | 0.0001 |
| cluster4028 | -1847.2593 | 0.0064 | 0.0707 | 11.1075 | 2.15E-12 |
| cluster4029 | -2251.3568 | 0.0697 | 0.4011 | 5.7531 | 0.002 |
| cluster4030 | -1125.1118 | 0.0372 | 0.1125 | 3.0232 | 0.003 |
| cluster4031 | -1097.6742 | 0.0248 | 0.2024 | 8.1550 | 0.005 |
| cluster4032 | -1475.4081 | 0.0011 | 0.0020 | 1.7127 | 1.34E-06 |
| cluster4033 | -2215.2701 | 0.0705 | 0.4245 | 6.0213 | 0.003 |
| cluster4034 | -493.6726 | 0.0001 | 0.0004 | 3.9353 | 0.01 |
| cluster4035 | -2115.6496 | 0.0602 | 0.1612 | 2.6794 | 0.0005 |
| cluster4036 | -1077.3476 | 0.0088 | 0.0530 | 6.0066 | 1.47E-05 |
| cluster4037 | -1626.0237 | 0.0192 | 0.1372 | 7.1390 | 7.03E-06 |
| cluster4038 | -902.57463 | 0.0166 | 0.1608 | 9.6769 | 8.88E-05 |
| cluster4039 | -1508.327 | 0.0611 | 0.1007 | 1.6492 | 0.02 |

| | | | | | |
|---|---|---|---|---|---|
| cluster4040 | -1126.6264 | 0.0307 | 0.1278 | 4.1634 | 0.005 |
| cluster4041 | -1475.5164 | 0.0138 | 0.0997 | 7.2069 | 9.98E-07 |
| cluster4042 | -899.70109 | 0.0320 | 0.1072 | 3.3505 | 0.01 |
| cluster4043 | -1138.6049 | 0.0447 | 0.5051 | 11.3158 | 0.03 |
| cluster4044 | -958.06892 | 0.0014 | 0.0109 | 7.7057 | 4.77E-11 |
| cluster4045 | -2023.8482 | 0.0654 | 0.3412 | 5.2149 | 0.01 |
| cluster4046 | -889.95957 | 0.0027 | 0.0055 | 2.0802 | 0.0003 |
| cluster4047 | -2069.4508 | 0.0179 | 0.1537 | 8.6058 | 1.00E-09 |
| cluster4048 | -1470.4343 | 0.0084 | 0.0189 | 2.2385 | 3.45E-06 |
| cluster4049 | -573.69745 | 0.0011 | 0.0187 | 17.1847 | 0.12 |
| cluster4050 | -1217.6531 | 0.0089 | 0.0315 | 3.5326 | 6.27E-06 |
| cluster4051 | -1404.3679 | 0.0086 | 0.0131 | 1.5224 | 4.95E-05 |
| cluster4052 | -1288.191 | 0.0192 | 0.0506 | 2.6405 | 8.60E-06 |
| cluster4053 | -1644.3531 | 0.0126 | 0.1502 | 11.9405 | 4.12E-06 |
| cluster4054 | -1860.9335 | 0.0185 | 0.1649 | 8.8938 | 1.03E-07 |
| cluster4055 | -719.97804 | 0.0023 | 0.0733 | 31.7929 | 8.00E-06 |
| cluster4056 | -934.73143 | 0.0351 | 0.1353 | 3.8559 | 0.003 |
| cluster4057 | -1565.1573 | 0.0347 | 0.1847 | 5.3231 | 0.0001 |
| cluster4058 | -1681.4822 | 0.0419 | 0.2259 | 5.3887 | 0.002 |
| cluster4059 | -723.11768 | 0.0124 | 0.1112 | 8.9397 | 0.02 |
| cluster4060 | -1875.8831 | 0.0282 | 0.1240 | 4.4018 | 4.88E-06 |
| cluster4061 | -1188.5742 | 0.0059 | 0.0562 | 9.4925 | 4.68E-09 |
| cluster4062 | -1212.6649 | 0.0137 | 0.0644 | 4.6971 | 1.35E-06 |
| cluster4063 | -1723.0171 | 0.0252 | 0.2730 | 10.8496 | 0.0002 |
| cluster4064 | -783.14254 | 0.0160 | 0.0768 | 4.8013 | 0.0004 |
| cluster4065 | -989.56052 | 0.0083 | 0.0241 | 2.8868 | 0.0001 |
| cluster4066 | -1616.9438 | 0.0460 | 0.2126 | 4.6243 | 0.001 |
| cluster4067 | -418.64638 | 0.0201 | 0.3386 | 16.8235 | 0.15 |
| cluster4068 | -1354.0161 | 0.0183 | 0.2641 | 14.4034 | 0.0003 |
| cluster4069 | -1065.1355 | 0.0620 | 0.1583 | 2.5530 | 0.05 |
| cluster4070 | -696.52304 | 0.0048 | 0.1319 | 27.7411 | 0.0002 |
| cluster4071 | -826.84886 | 0.0021 | 0.0195 | 9.1777 | 2.70E-06 |
| cluster4072 | -988.18633 | 0.0254 | 0.1109 | 4.3654 | 0.003 |
| cluster4073 | -1889.85 | 0.0307 | 0.2057 | 6.6959 | 4.98E-06 |
| cluster4074 | -1257.2498 | 0.0323 | 0.2779 | 8.5955 | 0.0005 |
| cluster4075 | -710.45233 | 0.8159 | 17.4125 | 21.3430 | 1.00 |
| cluster4076 | -1650.3443 | 0.0890 | 0.2283 | 2.5652 | 0.02 |
| cluster4077 | -1680.0443 | 0.0437 | 0.2035 | 4.6542 | 0.003 |
| cluster4078 | -1232.5178 | 0.0219 | 0.1161 | 5.3020 | 5.97E-05 |
| cluster4079 | -1207.5574 | 0.0010 | 0.0048 | 4.5998 | 2.02E-10 |
| cluster4080 | -737.50374 | 0.0182 | 0.0954 | 5.2394 | 0.01 |
| cluster4081 | -1131.3344 | 0.0187 | 0.0671 | 3.5839 | 0.0001 |
| cluster4082 | -743.4072 | 0.0095 | 0.3129 | 33.0818 | 0.02 |
| cluster4083 | -1706.5275 | 0.0586 | 0.3924 | 6.6952 | 0.002 |

| | | | | | |
|---|---|---|---|---|---|
| cluster4084 | -716.07404 | 0.0153 | 0.0376 | 2.4525 | 0.02 |
| cluster4085 | -859.67662 | 0.0281 | 0.0458 | 1.6299 | 0.15 |
| cluster4086 | -1314.8171 | 0.0302 | 0.2123 | 7.0213 | 0.0002 |
| cluster4087 | -1206.0256 | 0.0122 | 0.1317 | 10.8395 | 7.54E-06 |
| cluster4088 | -1314.3205 | 0.0574 | 0.0998 | 1.7402 | 0.03 |
| cluster4089 | -1511.3173 | 0.0062 | 0.0787 | 12.6674 | 1.05E-07 |
| cluster4090 | -755.73188 | 0.0364 | 0.1046 | 2.8723 | 0.02 |
| cluster4091 | -954.10595 | 0.0315 | 0.0526 | 1.6711 | 0.01 |
| cluster4092 | -1425.0216 | 0.0267 | 0.2086 | 7.8111 | 0.0003 |
| cluster4093 | -1006.9591 | 0.0181 | 0.1132 | 6.2525 | 0.0001 |
| cluster4094 | -375.50209 | 0.0086 | 0.0061 | 0.7162 | 0.22 |
| cluster4095 | -1404.7048 | 0.0437 | 0.3069 | 7.0292 | 0.02 |
| cluster4096 | -1213.1169 | 0.0234 | 0.1768 | 7.5534 | 0.0001 |
| cluster4097 | -705.33283 | 0.0964 | 0.0549 | 0.5699 | 0.50 |
| cluster4098 | -1868.7308 | 0.0335 | 0.1934 | 5.7698 | 1.71E-05 |
| cluster4099 | -811.01508 | 0.0026 | 0.0158 | 6.0017 | 1.28E-06 |
| cluster4100 | -1048.7831 | 0.0080 | 0.0441 | 5.5298 | 1.79E-07 |
| cluster4101 | -2089.378 | 0.2405 | 0.6157 | 2.5598 | 0.46 |
| cluster4102 | -1856.9184 | 0.0471 | 0.7040 | 14.9429 | 0.02 |
| cluster4103 | -1107.3743 | 0.0422 | 0.0518 | 1.2268 | 0.02 |
| cluster4104 | -1018.0602 | 0.0074 | 0.0558 | 7.5566 | 1.91E-07 |
| cluster4105 | -735.70163 | 0.0092 | 0.2378 | 25.8613 | 0.005 |
| cluster4106 | -1000.9781 | 0.0078 | 0.0602 | 7.6778 | 4.02E-06 |
| cluster4107 | -786.85262 | 0.0024 | 0.0145 | 6.0800 | 7.47E-07 |
| cluster4108 | -783.54259 | 0.0113 | 0.0832 | 7.3807 | 1.42E-05 |
| cluster4109 | -1651.8951 | 0.0672 | 0.4468 | 6.6455 | 0.01 |
| cluster4110 | -1246.4892 | 0.0630 | 0.3160 | 5.0214 | 0.03 |
| cluster4111 | -925.20938 | 0.0483 | 0.3081 | 6.3858 | 0.05 |
| cluster4112 | -1113.3942 | 0.0162 | 0.0584 | 3.5959 | 2.32E-05 |
| cluster4113 | -773.39303 | 0.0060 | 0.0300 | 4.9864 | 0.03 |
| cluster4114 | -1075.0215 | 0.0430 | 0.1210 | 2.8184 | 0.01 |
| cluster4115 | -1156.5382 | 0.0110 | 0.0795 | 7.2080 | 2.02E-06 |
| cluster4116 | -636.70294 | 0.0292 | 0.2866 | 9.8240 | 0.12 |
| cluster4117 | -660.63193 | 0.0129 | 0.0334 | 2.5917 | 0.01 |
| cluster4118 | -1288.4941 | 0.0300 | 0.2073 | 6.9039 | 0.0001 |
| cluster4119 | -926.42559 | 0.0191 | 0.1340 | 7.0112 | 0.0003 |
| cluster4120 | -968.37896 | 0.0045 | 0.0154 | 3.4073 | 1.98E-06 |
| cluster4121 | -1217.6375 | 0.0043 | 0.1241 | 28.6403 | 4.83E-09 |
| cluster4122 | -527.68051 | 0.0343 | 0.0109 | 0.3139 | 0.56 |
| cluster4123 | -468.66831 | 0.0071 | 0.0232 | 3.2344 | 0.02 |
| cluster4124 | -905.64758 | 0.0044 | 0.0684 | 15.6560 | 3.63E-07 |
| cluster4125 | -1065.915 | 0.0770 | 0.3269 | 4.2482 | 0.07 |
| cluster4126 | -763.08084 | 0.0836 | 0.1208 | 1.4451 | 0.20 |
| cluster4127 | -766.46498 | 0.0571 | 0.1741 | 3.0492 | 0.03 |

| | | | | | |
|---|---|---|---|---|---|
| cluster4128 | -989.65672 | 0.0353 | 0.4426 | 12.5362 | 0.02 |

**Supplementary Table 21.** codeml results - w for *C. hominivora* x/other species

| ID | Likelihood | ϖ other species | dN | dS | ϖ *C. hominivorax* | dN |
|---|---|---|---|---|---|---|
| cluster1127rr | -1408.1518 | 0.0501 | 0.0985 | 1.9672 | 0.1341 | 0.0178 |
| cluster2853rr | -22470.892 | 0.0303 | 0.0352 | 1.1628 | 0.0197 | 0.0156 |
| cluster2854rr | -1614.855 | 0.0330 | 0.0131 | 0.3968 | 0.0637 | 0.0074 |
| cluster2855rr | -23534.714 | 0.0183 | 0.0356 | 1.9432 | 0.0904 | 0.0084 |
| cluster2856rr | -15419.568 | 0.1024 | 0.1528 | 1.4922 | 0.2182 | 0.0309 |
| cluster2857rr | -6020.0198 | 0.0127 | 0.0048 | 0.3768 | 0.0079 | 0.0015 |
| cluster2858rr | -7755.8174 | 0.0039 | 0.0038 | 0.9759 | 0.0041 | 0.0007 |
| cluster2859rr | -5669.5914 | 0.0356 | 0.0161 | 0.4516 | 0.2923 | 0.0093 |
| cluster2860rr | -16042.841 | 0.0019 | 0.0016 | 0.8180 | 0.0005 | 0.0003 |
| cluster2861rr | -8374.8991 | 0.0101 | 0.0096 | 0.9540 | 0.0145 | 0.0047 |
| cluster2862rr | -18853.557 | 0.0165 | 0.0149 | 0.9026 | 0.0095 | 0.0070 |
| cluster2863rr | -15063.766 | 0.0284 | 0.0368 | 1.2951 | 0.1002 | 0.0120 |
| cluster2864rr | -12550.571 | 0.0040 | 0.0041 | 1.0400 | 0.0022 | 0.0018 |
| cluster2865rr | -15005.003 | 0.0986 | 0.0537 | 0.5449 | 0.0953 | 0.0539 |
| cluster2866rr | -17304.629 | 0.0252 | 0.0384 | 1.5217 | 0.0713 | 0.0092 |
| cluster2867rr | -14814.455 | 0.0394 | 0.0555 | 1.4091 | 0.7198 | 0.0305 |
| cluster2869rr | -8907.7315 | 0.0817 | 0.0577 | 0.7059 | 0.2411 | 0.0253 |
| cluster2870rr | -3879.1591 | 0.0571 | 0.0438 | 0.7668 | 0.0001 | 0.0000 |
| cluster2871rr | -3139.9399 | 0.0269 | 0.0094 | 0.3505 | 0.0215 | 0.0020 |
| cluster2872rr | -12254.583 | 0.0765 | 0.0931 | 1.2172 | 0.1688 | 0.0251 |
| cluster2873rr | -13848.078 | 0.0168 | 0.0129 | 0.7694 | 0.0081 | 0.0046 |
| cluster2874rr | -10373.676 | 0.0327 | 0.0226 | 0.6923 | 0.1430 | 0.0144 |
| cluster2875rr | -13632.324 | 0.0176 | 0.0229 | 1.2985 | 0.0089 | 0.0088 |
| cluster2876rr | -3967.251 | 0.0405 | 0.0275 | 0.6793 | 0.0133 | 0.0094 |
| cluster2877rr | -10635.66 | 0.0487 | 0.0374 | 0.7683 | 0.0719 | 0.0352 |
| cluster2878rr | -7056.5537 | 0.0430 | 0.0166 | 0.3862 | 0.0181 | 0.0043 |
| cluster2879rr | -10036.291 | 0.1925 | 0.0680 | 0.3535 | 0.1729 | 0.0289 |
| cluster2880rr | -10622.696 | 0.0426 | 0.0619 | 1.4533 | 0.2863 | 0.0197 |
| cluster2881rr | -11523.044 | 0.0511 | 0.0400 | 0.7836 | 0.0576 | 0.0367 |
| cluster2882rr | -10649.726 | 0.0300 | 0.0324 | 1.0788 | 0.0385 | 0.0097 |
| cluster2883rr | -6761.6587 | 0.0777 | 0.0423 | 0.5441 | 0.0423 | 0.0235 |
| cluster2884rr | -12070.596 | 0.0252 | 0.0174 | 0.6909 | 0.0403 | 0.0042 |
| cluster2885rr | -15711.441 | 0.0368 | 0.0622 | 1.6917 | 0.2460 | 0.0219 |
| cluster2886rr | -12003.19 | 0.0416 | 0.0283 | 0.6796 | 0.0290 | 0.0134 |
| cluster2887rr | -8366.7074 | 0.0057 | 0.0087 | 1.5092 | 0.0151 | 0.0038 |
| cluster2888rr | -7779.9395 | 0.0571 | 0.0996 | 1.7464 | 0.1692 | 0.0340 |
| cluster2889rr | -6920.1193 | 0.0071 | 0.0100 | 1.4119 | 0.0525 | 0.0065 |
| cluster2890rr | -2676.1053 | 0.0969 | 0.0162 | 0.1668 | 0.0001 | 0.0000 |
| cluster2891rr | -12081.88 | 0.0335 | 0.0526 | 1.5695 | 0.1636 | 0.0144 |
| cluster2892rr | -7913.1288 | 0.0154 | 0.0108 | 0.7011 | 0.0128 | 0.0071 |
| cluster2893rr | -8455.2909 | 0.0285 | 0.0674 | 2.3630 | 0.2021 | 0.0173 |
| cluster2894rr | -5507.8383 | 0.0111 | 0.0154 | 1.3925 | 0.0623 | 0.0060 |

| | | | | | | |
|---|---|---|---|---|---|---|
| cluster2895rr | -8442.5767 | 0.0326 | 0.0445 | 1.3676 | 0.1086 | 0.0135 |
| cluster2896rr | -10285.148 | 0.0096 | 0.0142 | 1.4837 | 0.0249 | 0.0042 |
| cluster2897rr | -9158.4277 | 0.0333 | 0.0280 | 0.8420 | 0.0636 | 0.0046 |
| cluster2898rr | -4497.7596 | 0.0292 | 0.0165 | 0.5649 | 0.0094 | 0.0045 |
| cluster2899rr | -8006.6375 | 0.0201 | 0.0231 | 1.1508 | 0.0167 | 0.0261 |
| cluster2900rr | -9924.1481 | 0.0128 | 0.0089 | 0.6913 | 0.0137 | 0.0076 |
| cluster2901rr | -9238.7414 | 0.0348 | 0.0490 | 1.4090 | 0.0670 | 0.0122 |
| cluster2902rr | -3690.4703 | 0.0219 | 0.0336 | 1.5340 | 0.1518 | 0.0134 |
| cluster2903rr | -8864.1596 | 0.0103 | 0.0102 | 0.9858 | 0.0110 | 0.0082 |
| cluster2904rr | -6415.2761 | 0.0924 | 0.0551 | 0.5960 | 0.0756 | 0.0161 |
| cluster2905rr | -8877.1523 | 0.0264 | 0.0138 | 0.5229 | 0.0476 | 0.0159 |
| cluster2906rr | -13666.382 | 0.0768 | 0.0371 | 0.4837 | 0.0575 | 0.0257 |
| cluster2907rr | -12720.927 | 0.0629 | 0.1076 | 1.7108 | 0.2569 | 0.0311 |
| cluster2908rr | -5644.7613 | 0.0131 | 0.0053 | 0.4056 | 0.0040 | 0.0004 |
| cluster2909rr | -2925.2791 | 0.0374 | 0.0146 | 0.3911 | 0.0518 | 0.0040 |
| cluster2910rr | -9268.5489 | 0.0412 | 0.0407 | 0.9891 | 0.1077 | 0.0081 |
| cluster2911rr | -4008.0678 | 0.0050 | 0.0013 | 0.2666 | 0.0001 | 0.0000 |
| cluster2912rr | -8764.768 | 0.0243 | 0.0341 | 1.4009 | 0.0615 | 0.0095 |
| cluster2913rr | -6296.8537 | 0.0028 | 0.0025 | 0.9012 | 0.0001 | 0.0000 |
| cluster2914rr | -8063.58 | 0.0154 | 0.0086 | 0.5600 | 0.0198 | 0.0069 |
| cluster2915rr | -4206.3323 | 0.0159 | 0.0163 | 1.0278 | 0.0356 | 0.0058 |
| cluster2916rr | -9765.0968 | 0.0350 | 0.0247 | 0.7068 | 0.0142 | 0.0070 |
| cluster2917rr | -9026.8308 | 0.0205 | 0.0235 | 1.1472 | 0.0133 | 0.0026 |
| cluster2918rr | -4474.804 | 0.0701 | 0.0366 | 0.5226 | 0.1213 | 0.0118 |
| cluster2919rr | -9564.6002 | 0.0691 | 0.0355 | 0.5138 | 0.0553 | 0.0195 |
| cluster2920rr | -6956.1708 | 0.0146 | 0.0307 | 2.1115 | 0.0328 | 0.0061 |
| cluster2921rr | -3809.2554 | 0.0040 | 0.0052 | 1.2944 | 0.0039 | 0.0005 |
| cluster2922rr | -6244.0286 | 0.0547 | 0.0712 | 1.3012 | 0.0964 | 0.0084 |
| cluster2923rr | -5865.446 | 0.1887 | 0.1426 | 0.7560 | 0.0820 | 0.0071 |
| cluster2924rr | -10971.469 | 0.0196 | 0.0325 | 1.6605 | 0.0553 | 0.0098 |
| cluster2925rr | -5768.9107 | 0.0304 | 0.0389 | 1.2803 | 0.0750 | 0.0139 |
| cluster2926rr | -11211.86 | 0.0243 | 0.0207 | 0.8517 | 0.0090 | 0.0075 |
| cluster2927rr | -4901.5762 | 0.0278 | 0.0456 | 1.6406 | 0.0673 | 0.0176 |
| cluster2928rr | -7178.2081 | 0.0201 | 0.0257 | 1.2795 | 0.0515 | 0.0077 |
| cluster2929rr | -8038.8479 | 0.0031 | 0.0041 | 1.3094 | 0.0239 | 0.0014 |
| cluster2930rr | -10988.15 | 0.0823 | 0.0637 | 0.7745 | 0.1180 | 0.0122 |
| cluster2931rr | -6860.9502 | 0.0452 | 0.0796 | 1.7632 | 0.1467 | 0.0221 |
| cluster2932rr | -9688.3352 | 0.0558 | 0.0672 | 1.2047 | 0.1470 | 0.0234 |
| cluster2933rr | -8731.3363 | 0.0516 | 0.0234 | 0.4537 | 0.0194 | 0.0059 |
| cluster2934rr | -10530.809 | 0.0618 | 0.0374 | 0.6062 | 0.0967 | 0.0079 |
| cluster2935rr | -4952.1413 | 0.0294 | 0.0220 | 0.7488 | 0.0224 | 0.0114 |
| cluster2936rr | -9481.4672 | 0.0287 | 0.0557 | 1.9407 | 0.5383 | 0.0260 |
| cluster2937rr | -4595.2725 | 0.0290 | 0.0152 | 0.5238 | 0.0201 | 0.0066 |
| cluster2938rr | -7239.6019 | 0.0165 | 0.0119 | 0.7192 | 0.0078 | 0.0034 |

| | | | | | | |
|---|---|---|---|---|---|---|
| cluster2939rr | -2817.9719 | 0.0233 | 0.0086 | 0.3676 | 0.0232 | 0.0039 |
| cluster2940rr | -11347.371 | 0.0554 | 0.0642 | 1.1590 | 0.3448 | 0.0307 |
| cluster2941rr | -3030.7479 | 0.0019 | 0.0004 | 0.1959 | 0.0001 | 0.0000 |
| cluster2942rr | -11356.754 | 0.0511 | 0.0679 | 1.3277 | 0.1012 | 0.0146 |
| cluster2943rr | -4037.7666 | 0.0091 | 0.0083 | 0.9125 | 0.0224 | 0.0122 |
| cluster2944rr | -8473.7824 | 0.0815 | 0.0596 | 0.7316 | 0.2769 | 0.1932 |
| cluster2945rr | -3084.1609 | 0.0021 | 0.0050 | 2.4249 | 0.0103 | 0.0016 |
| cluster2946rr | -6736.9947 | 0.0109 | 0.0037 | 0.3430 | 0.0001 | 0.0000 |
| cluster2947rr | -9657.4384 | 0.0349 | 0.0224 | 0.6412 | 0.0722 | 0.0082 |
| cluster2948rr | -9374.6823 | 0.0568 | 0.0467 | 0.8225 | 0.0334 | 0.0127 |
| cluster2949rr | -6814.0319 | 0.0189 | 0.0109 | 0.5752 | 0.0095 | 0.0051 |
| cluster2950rr | -7094.8639 | 0.0208 | 0.0156 | 0.7492 | 0.0001 | 0.0001 |
| cluster2951rr | -9094.4841 | 0.0635 | 0.0827 | 1.3034 | 0.0568 | 0.0211 |
| cluster2952rr | -10887.482 | 0.0380 | 0.0497 | 1.3078 | 0.1275 | 0.0136 |
| cluster2953rr | -13284.205 | 0.0628 | 0.0943 | 1.5014 | 0.1352 | 0.0385 |
| cluster2954rr | -7271.9522 | 0.0178 | 0.0246 | 1.3800 | 0.0317 | 0.0191 |
| cluster2955rr | -6029.9212 | 0.0238 | 0.0226 | 0.9481 | 0.0349 | 0.0056 |
| cluster2956rr | -10114.883 | 0.0135 | 0.0145 | 1.0713 | 0.0802 | 0.0922 |
| cluster2957rr | -8387.4427 | 0.0904 | 0.1110 | 1.2281 | 0.1403 | 0.0132 |
| cluster2958rr | -9681.6095 | 0.0104 | 0.0123 | 1.1842 | 0.0497 | 0.0065 |
| cluster2959rr | -5168.9263 | 0.0550 | 0.0330 | 0.5999 | 0.0414 | 0.0189 |
| cluster2960rr | -9442.3022 | 0.0240 | 0.0210 | 0.8739 | 0.0219 | 0.0160 |
| cluster2961rr | -8397.8143 | 0.0100 | 0.0232 | 2.3141 | 0.0069 | 0.0050 |
| cluster2962rr | -7495.358 | 0.0213 | 0.0372 | 1.7480 | 0.0261 | 0.0257 |
| cluster2963rr | -7221.2214 | 0.0385 | 0.0152 | 0.3935 | 0.0168 | 0.0040 |
| cluster2964rr | -5243.3817 | 0.0356 | 0.0170 | 0.4773 | 0.0095 | 0.0041 |
| cluster2965rr | -7391.4684 | 0.0110 | 0.0100 | 0.9101 | 0.0188 | 0.0021 |
| cluster2966rr | -5587.4167 | 0.0219 | 0.0362 | 1.6561 | 0.0341 | 0.0086 |
| cluster2967rr | -9577.4178 | 0.0184 | 0.0112 | 0.6083 | 0.0097 | 0.0043 |
| cluster2968rr | -5456.6125 | 0.0177 | 0.0140 | 0.7912 | 0.0234 | 0.0103 |
| cluster2969rr | -7909.9152 | 0.0412 | 0.0398 | 0.9660 | 0.0651 | 0.0071 |
| cluster2970rr | -4236.3158 | 0.0038 | 0.0051 | 1.3676 | 0.0034 | 0.0005 |
| cluster2971rr | -3608.877 | 0.0310 | 0.0088 | 0.2846 | 0.0334 | 0.0022 |
| cluster2972rr | -2667.1296 | 0.0081 | 0.0048 | 0.5887 | 0.0012 | 0.0013 |
| cluster2973rr | -6333.7713 | 0.0722 | 0.0893 | 1.2376 | 1.4406 | 0.0408 |
| cluster2974rr | -4423.2914 | 0.0307 | 0.0198 | 0.6457 | 0.0216 | 0.0013 |
| cluster2975rr | -10300.199 | 0.0679 | 0.0774 | 1.1393 | 0.1092 | 0.0232 |
| cluster2976rr | -1469.3575 | 0.1996 | 0.0978 | 0.4902 | 0.1572 | 0.0080 |
| cluster2977rr | -3390.7836 | 0.0044 | 0.0108 | 2.4888 | 0.0266 | 0.0030 |
| cluster2978rr | -8215.3844 | 0.0015 | 0.0011 | 0.7259 | 0.0008 | 0.0003 |
| cluster2979rr | -8821.4663 | 0.0340 | 0.0296 | 0.8726 | 0.1781 | 0.0165 |
| cluster2980rr | -1551.0743 | 0.0171 | 0.0540 | 3.1591 | 0.0967 | 0.0159 |
| cluster2981rr | -6210.1365 | 0.0518 | 0.0554 | 1.0704 | 0.1261 | 0.0165 |
| cluster2982rr | -8044.7287 | 0.0916 | 0.0480 | 0.5241 | 0.2964 | 0.0440 |

| | | | | | | |
|---|---|---|---|---|---|---|
| cluster2983rr | -7499.0262 | 0.0186 | 0.0338 | 1.8198 | 0.1578 | 0.0175 |
| cluster2984rr | -4826.1073 | 0.0365 | 0.0164 | 0.4503 | 0.0083 | 0.0031 |
| cluster2985rr | -3972.6035 | 0.2360 | 0.0899 | 0.3809 | 0.2479 | 0.0513 |
| cluster2986rr | -3496.4645 | 0.1036 | 0.0626 | 0.6037 | 0.1763 | 0.0148 |
| cluster2987rr | -6077.5747 | 0.0292 | 0.0180 | 0.6174 | 0.0415 | 0.0081 |
| cluster2988rr | -5703.3315 | 0.0240 | 0.0303 | 1.2584 | 0.0309 | 0.0038 |
| cluster2989rr | -6588.9965 | 0.0409 | 0.0179 | 0.4381 | 0.0290 | 0.0101 |
| cluster2990rr | -7510.8818 | 0.0425 | 0.0484 | 1.1397 | 0.0684 | 0.0136 |
| cluster2991rr | -6303.484 | 0.0521 | 0.0803 | 1.5410 | 0.1571 | 0.0214 |
| cluster2992rr | -4177.4874 | 0.0092 | 0.0036 | 0.3942 | 0.0230 | 0.0044 |
| cluster2993rr | -5421.3965 | 0.0355 | 0.0275 | 0.7746 | 0.0183 | 0.0117 |
| cluster2994rr | -2259.8762 | 0.0040 | 0.0022 | 0.5554 | 0.0001 | 0.0000 |
| cluster2995rr | -740.22901 | 0.0203 | 0.0253 | 1.2455 | 4.4656 | 0.0000 |
| cluster2996rr | -7310.3719 | 0.0450 | 0.0291 | 0.6459 | 0.0350 | 0.0195 |
| cluster2997rr | -8168.4741 | 0.0376 | 0.0355 | 0.9443 | 0.0817 | 0.0122 |
| cluster2998rr | -7502.2911 | 0.0176 | 0.0227 | 1.2877 | 0.0394 | 0.0056 |
| cluster2999rr | -6764.1194 | 0.0305 | 0.0472 | 1.5481 | 0.0355 | 0.0072 |
| cluster3000rr | -6334.646 | 0.0118 | 0.0063 | 0.5365 | 0.0111 | 0.0016 |
| cluster3001rr | -6054.598 | 0.0188 | 0.0140 | 0.7489 | 0.0642 | 0.0091 |
| cluster3002rr | -7667.6739 | 0.0248 | 0.0182 | 0.7338 | 0.0449 | 0.0086 |
| cluster3003rr | -7809.9123 | 0.0862 | 0.0500 | 0.5798 | 0.1351 | 0.0158 |
| cluster3004rr | -5958.3123 | 0.1054 | 0.0405 | 0.3846 | 0.0360 | 0.0107 |
| cluster3005rr | -4102.8217 | 0.0579 | 0.0199 | 0.3428 | 0.0721 | 0.0042 |
| cluster3006rr | -8670.3245 | 0.0603 | 0.0513 | 0.8514 | 0.0393 | 0.0195 |
| cluster3007rr | -8242.9637 | 0.0256 | 0.0498 | 1.9477 | 0.0442 | 0.0074 |
| cluster3008rr | -5954.3297 | 0.0316 | 0.0356 | 1.1259 | 0.0342 | 0.0056 |
| cluster3009rr | -5909.9548 | 0.0394 | 0.0239 | 0.6060 | 0.0861 | 0.0140 |
| cluster3010rr | -8050.9424 | 0.1356 | 0.0767 | 0.5657 | 0.3331 | 0.0301 |
| cluster3011rr | -3371.1432 | 0.0109 | 0.0172 | 1.5758 | 0.0049 | 0.0036 |
| cluster3012rr | -5208.7879 | 0.0620 | 0.0864 | 1.3927 | 0.0254 | 0.0063 |
| cluster3013rr | -6735.1754 | 0.0823 | 0.0337 | 0.4096 | 0.0841 | 0.0164 |
| cluster3014rr | -3561.6942 | 0.0083 | 0.0122 | 1.4684 | 0.0048 | 0.0009 |
| cluster3015rr | -2410.1194 | 0.0400 | 0.0186 | 0.4654 | 0.0919 | 0.0036 |
| cluster3016rr | -4553.4929 | 0.0361 | 0.0189 | 0.5238 | 0.1043 | 0.0084 |
| cluster3017rr | -5971.6728 | 0.0276 | 0.0502 | 1.8193 | 0.0091 | 0.0009 |
| cluster3018rr | -5679.1851 | 0.0065 | 0.0069 | 1.0549 | 0.0074 | 0.0015 |
| cluster3019rr | -7570.9288 | 0.0532 | 0.0633 | 1.1912 | 0.0541 | 0.0308 |
| cluster3020rr | -6175.053 | 0.0274 | 0.0250 | 0.9110 | 0.2336 | 0.0181 |
| cluster3021rr | -3022.6853 | 0.0188 | 0.0092 | 0.4900 | 0.0037 | 0.0012 |
| cluster3022rr | -3430.8631 | 0.0562 | 0.0918 | 1.6332 | 0.1016 | 0.0120 |
| cluster3023rr | -4202.7556 | 0.0109 | 0.0067 | 0.6170 | 0.0022 | 0.0007 |
| cluster3024rr | -1959.2633 | 0.0836 | 0.0326 | 0.3900 | 0.2095 | 0.0079 |
| cluster3025rr | -4690.8864 | 0.0092 | 0.0037 | 0.4076 | 0.0205 | 0.0027 |
| cluster3026rr | -4110.3918 | 0.1631 | 0.1129 | 0.6923 | 0.2962 | 0.0242 |

| | | | | | | |
|---|---|---|---|---|---|---|
| cluster3027rr | -3436.2028 | 0.0025 | 0.0024 | 0.9297 | 0.0007 | 0.0005 |
| cluster3028rr | -6798.6606 | 0.0355 | 0.0607 | 1.7092 | 0.1265 | 0.0146 |
| cluster3029rr | -6670.6407 | 0.0258 | 0.0177 | 0.6876 | 0.0730 | 0.0102 |
| cluster3030rr | -5268.4411 | 0.0538 | 0.0212 | 0.3932 | 0.0537 | 0.0119 |
| cluster3031rr | -8795.1607 | 0.1061 | 0.0742 | 0.6991 | 0.0684 | 0.0415 |
| cluster3032rr | -5916.3546 | 0.0234 | 0.0190 | 0.8120 | 0.0218 | 0.0150 |
| cluster3033rr | -3822.0889 | 0.1681 | 0.0338 | 0.2013 | 0.0507 | 0.0027 |
| cluster3034rr | -4963.3391 | 0.0152 | 0.0383 | 2.5208 | 0.0346 | 1.1430 |
| cluster3035rr | -7643.4956 | 0.0229 | 0.0174 | 0.7571 | 0.0136 | 0.0046 |
| cluster3036rr | -8982.0122 | 0.0696 | 0.0797 | 1.1465 | 0.0925 | 0.0131 |
| cluster3037rr | -2566.5954 | 0.0187 | 0.0089 | 0.4733 | 0.0154 | 0.0016 |
| cluster3038rr | -4470.5752 | 0.0569 | 0.0580 | 1.0194 | 0.1044 | 0.0094 |
| cluster3039rr | -6527.1089 | 0.0352 | 0.0125 | 0.3564 | 0.0152 | 0.0026 |
| cluster3040rr | -4838.6663 | 0.0218 | 0.0086 | 0.3931 | 0.0021 | 0.0006 |
| cluster3041rr | -8447.1959 | 0.0429 | 0.0565 | 1.3187 | 0.0771 | 0.0089 |
| cluster3042rr | -4396.0632 | 0.0292 | 0.0328 | 1.1254 | 0.1381 | 0.0155 |
| cluster3043rr | -2915.1371 | 0.0197 | 0.0543 | 2.7607 | 0.4311 | 0.0104 |
| cluster3044rr | -6428.187 | 0.0293 | 0.0338 | 1.1548 | 0.0359 | 0.0077 |
| cluster3045rr | -6125.8697 | 0.0572 | 0.0305 | 0.5326 | 0.0206 | 0.0043 |
| cluster3046rr | -7251.4351 | 0.1282 | 0.0491 | 0.3827 | 0.4261 | 0.3184 |
| cluster3047rr | -2123.234 | 0.0036 | 0.0027 | 0.7636 | 0.0001 | 0.0000 |
| cluster3048rr | -5407.5664 | 0.0217 | 0.0166 | 0.7642 | 0.0275 | 0.0029 |
| cluster3049rr | -7060.9741 | 0.0187 | 0.0127 | 0.6781 | 0.0075 | 0.0043 |
| cluster3050rr | -7560.893 | 0.0382 | 0.0301 | 0.7891 | 0.0713 | 0.0078 |
| cluster3051rr | -7508.5905 | 0.0349 | 0.0558 | 1.5960 | 0.1043 | 0.0144 |
| cluster3052rr | -6822.4367 | 0.0437 | 0.0563 | 1.2878 | 0.1658 | 0.0143 |
| cluster3053rr | -9369.1658 | 0.0984 | 0.0689 | 0.6998 | 0.0650 | 0.0340 |
| cluster3054rr | -6879.6766 | 0.0090 | 0.0081 | 0.9014 | 0.0210 | 0.0020 |
| cluster3055rr | -3993.5925 | 0.0307 | 0.0235 | 0.7658 | 0.0074 | 0.0007 |
| cluster3056rr | -5994.6684 | 0.1041 | 0.0311 | 0.2991 | 0.0767 | 0.0125 |
| cluster3057rr | -919.31618 | 0.0498 | 0.0178 | 0.3576 | 999.0000 | 0.0034 |
| cluster3058rr | -2534.6585 | 0.0289 | 0.0103 | 0.3558 | 0.0070 | 0.0011 |
| cluster3059rr | -8709.6709 | 0.0588 | 0.1463 | 2.4876 | 0.2492 | 0.0147 |
| cluster3060rr | -5324.9765 | 0.0315 | 0.0367 | 1.1658 | 0.0634 | 0.0119 |
| cluster3061rr | -5740.3391 | 0.0169 | 0.0069 | 0.4070 | 0.0108 | 0.0041 |
| cluster3062rr | -5687.9668 | 0.0397 | 0.0180 | 0.4530 | 0.0058 | 0.0008 |
| cluster3063rr | -6147.4358 | 0.0332 | 0.0272 | 0.8202 | 0.0207 | 0.0049 |
| cluster3064rr | -1920.7107 | 0.0104 | 0.0044 | 0.4249 | 0.0125 | 0.0012 |
| cluster3065rr | -2295.8551 | 0.0793 | 0.0721 | 0.9094 | 0.3311 | 0.0434 |
| cluster3066rr | -6187.6151 | 0.0692 | 0.0382 | 0.5513 | 0.0695 | 0.0292 |
| cluster3067rr | -4918.3728 | 0.0444 | 0.0300 | 0.6753 | 0.0208 | 0.0086 |
| cluster3068rr | -5143.4898 | 0.1097 | 0.0332 | 0.3027 | 0.1120 | 0.0133 |
| cluster3069rr | -1632.352 | 0.0015 | 0.0003 | 0.2184 | 0.0001 | 0.0000 |
| cluster3070rr | -5192.1412 | 0.0379 | 0.0290 | 0.7639 | 0.0297 | 0.0170 |

| | | | | | | |
|---|---|---|---|---|---|---|
| cluster3071rr | -5856.4396 | 0.0239 | 0.0111 | 0.4629 | 0.0129 | 0.0041 |
| cluster3072rr | -7363.0657 | 0.0723 | 0.0718 | 0.9935 | 0.2124 | 0.0433 |
| cluster3073rr | -3570.6042 | 0.0075 | 0.0052 | 0.6896 | 0.0104 | 0.0008 |
| cluster3074rr | -7651.0033 | 0.0290 | 0.0479 | 1.6566 | 0.0369 | 0.0072 |
| cluster3075rr | -5426.478 | 0.0394 | 0.0288 | 0.7313 | 0.0211 | 0.0082 |
| cluster3076rr | -1814.397 | 0.0334 | 0.0465 | 1.3956 | 0.0300 | 0.0123 |
| cluster3077rr | -4383.9391 | 0.0251 | 0.0195 | 0.7757 | 0.0136 | 0.0073 |
| cluster3078rr | -5019.1801 | 0.0176 | 0.0156 | 0.8849 | 0.0663 | 0.0186 |
| cluster3079rr | -7615.6036 | 0.0381 | 0.0245 | 0.6428 | 0.0158 | 0.0121 |
| cluster3080rr | -5649.7973 | 0.0157 | 0.0136 | 0.8657 | 0.0233 | 0.0039 |
| cluster3081rr | -3874.657 | 0.0108 | 0.0067 | 0.6207 | 0.0028 | 0.0008 |
| cluster3082rr | -7187.4995 | 0.0600 | 0.0566 | 0.9443 | 0.0447 | 0.0074 |
| cluster3083rr | -4160.3112 | 0.0243 | 0.0111 | 0.4565 | 0.0357 | 0.0141 |
| cluster3084rr | -3197.3645 | 0.0395 | 0.0075 | 0.1901 | 0.0145 | 0.0011 |
| cluster3085rr | -7342.2775 | 0.0204 | 0.0495 | 2.4238 | 0.0813 | 0.0072 |
| cluster3086rr | -3401.2974 | 0.1073 | 0.0563 | 0.5250 | 0.0356 | 0.0094 |
| cluster3087rr | -6567.4953 | 0.0195 | 0.0123 | 0.6277 | 0.0092 | 0.0009 |
| cluster3088rr | -6373.6821 | 0.0508 | 0.0351 | 0.6907 | 0.2127 | 0.0397 |
| cluster3089rr | -3685.2768 | 0.0691 | 0.0371 | 0.5371 | 0.0876 | 0.0384 |
| cluster3090rr | -4960.0975 | 0.0090 | 0.0095 | 1.0527 | 0.0022 | 0.0005 |
| cluster3091rr | -5911.0997 | 0.0560 | 0.0221 | 0.3951 | 0.0438 | 0.0089 |
| cluster3092rr | -4264.2091 | 0.0162 | 0.0132 | 0.8162 | 0.0386 | 0.0053 |
| cluster3093rr | -4536.5062 | 0.0047 | 0.0064 | 1.3662 | 0.0333 | 0.0070 |
| cluster3094rr | -7861.3004 | 0.0270 | 0.0537 | 1.9906 | 0.0682 | 0.0171 |
| cluster3095rr | -4869.2445 | 0.0070 | 0.0057 | 0.8159 | 0.0023 | 0.0012 |
| cluster3096rr | -6131.4792 | 0.0319 | 0.0296 | 0.9272 | 0.0601 | 0.0071 |
| cluster3097rr | -7077.2525 | 0.0201 | 0.0088 | 0.4361 | 0.0267 | 0.0071 |
| cluster3098rr | -1415.2574 | 0.0053 | 0.0081 | 1.5242 | 0.0520 | 0.0088 |
| cluster3099rr | -4266.4159 | 0.0324 | 0.0228 | 0.7032 | 0.0872 | 0.0429 |
| cluster3100rr | -4390.1394 | 0.0384 | 0.0362 | 0.9423 | 0.1484 | 0.0113 |
| cluster3101rr | -3432.2695 | 0.0552 | 0.0328 | 0.5952 | 0.0214 | 0.0160 |
| cluster3102rr | -2230.6148 | 0.0164 | 0.0128 | 0.7834 | 0.0071 | 0.0058 |
| cluster3103rr | -5216.6239 | 0.0371 | 0.0660 | 1.7781 | 0.0335 | 0.0108 |
| cluster3104rr | -4326.6516 | 0.0230 | 0.0189 | 0.8213 | 0.0110 | 0.0039 |
| cluster3105rr | -7968.5413 | 0.0243 | 0.0499 | 2.0575 | 0.0651 | 0.0146 |
| cluster3106rr | -5131.4345 | 0.0031 | 0.0016 | 0.5121 | 0.0001 | 0.0000 |
| cluster3107rr | -3137.8714 | 0.0416 | 0.0195 | 0.4689 | 0.0351 | 0.0023 |
| cluster3108rr | -4331.8584 | 0.1395 | 0.0900 | 0.6453 | 0.1533 | 0.0186 |
| cluster3109rr | -3422.2594 | 0.0151 | 0.0228 | 1.5091 | 87.3110 | 0.0371 |
| cluster3110rr | -6379.8629 | 0.1040 | 0.0581 | 0.5588 | 0.1831 | 0.0134 |
| cluster3111rr | -4889.4354 | 0.0324 | 0.0228 | 0.7039 | 0.0350 | 0.0061 |
| cluster3112rr | -6144.8759 | 0.0391 | 0.0672 | 1.7188 | 0.0717 | 0.0103 |
| cluster3113rr | -3068.5633 | 0.0258 | 0.0264 | 1.0256 | 0.0166 | 0.0026 |
| cluster3114rr | -4680.6482 | 0.0255 | 0.0143 | 0.5598 | 0.0839 | 0.0229 |

| | | | | | | |
|---|---|---|---|---|---|---|
| cluster3115rr | -3776.8792 | 0.0188 | 0.0153 | 0.8162 | 0.0062 | 0.0008 |
| cluster3116rr | -2722.4032 | 0.0405 | 0.0320 | 0.7922 | 0.0074 | 0.0012 |
| cluster3117rr | -7470.4339 | 0.0362 | 0.0762 | 2.1055 | 0.6344 | 0.0296 |
| cluster3118rr | -1575.8422 | 0.0337 | 0.0078 | 0.2320 | 0.0001 | 0.0000 |
| cluster3119rr | -2980.1028 | 0.0454 | 0.0435 | 0.9584 | 0.0148 | 0.0076 |
| cluster3120rr | -4901.6498 | 0.0187 | 0.0150 | 0.8011 | 0.0870 | 0.0120 |
| cluster3121rr | -6820.9994 | 0.0258 | 0.0220 | 0.8515 | 0.0323 | 0.0165 |
| cluster3122rr | -3558.2031 | 0.0217 | 0.0098 | 0.4539 | 0.0155 | 0.0044 |
| cluster3123rr | -3830.0935 | 0.0158 | 0.0081 | 0.5124 | 0.0050 | 0.0017 |
| cluster3124rr | -6377.252 | 0.0401 | 0.0340 | 0.8472 | 0.0288 | 0.0058 |
| cluster3125rr | -4802.4516 | 0.0691 | 0.0431 | 0.6242 | 0.0421 | 0.0064 |
| cluster3126rr | -7549.6822 | 0.0460 | 0.0317 | 0.6889 | 0.0234 | 0.0129 |
| cluster3127rr | -2606.3444 | 0.0074 | 0.0082 | 1.1125 | 0.0247 | 0.0032 |
| cluster3128rr | -2238.1404 | 0.0023 | 0.0011 | 0.4835 | 0.0001 | 0.0000 |
| cluster3129rr | -3252.8069 | 0.0229 | 0.0815 | 3.5578 | 0.2701 | 0.0327 |
| cluster3130rr | -5794.996 | 0.0319 | 0.0420 | 1.3170 | 0.1270 | 0.0147 |
| cluster3131rr | -1764.4053 | 0.0075 | 0.0103 | 1.3756 | 0.0139 | 0.0060 |
| cluster3132rr | -5288.941 | 0.0266 | 0.0153 | 0.5752 | 0.0160 | 0.0086 |
| cluster3133rr | -8917.0153 | 0.0526 | 0.1095 | 2.0822 | 0.1793 | 0.0246 |
| cluster3134rr | -3290.1447 | 0.0620 | 0.0251 | 0.4055 | 0.0328 | 0.0069 |
| cluster3135rr | -4396.5337 | 0.0092 | 0.0072 | 0.7813 | 0.0336 | 0.0058 |
| cluster3136rr | -3953.6516 | 0.0368 | 0.0442 | 1.2012 | 0.0585 | 0.0084 |
| cluster3137rr | -3087.9356 | 0.0225 | 0.0292 | 1.3010 | 0.0808 | 0.0100 |
| cluster3138rr | -4740.903 | 0.0592 | 0.0487 | 0.8228 | 0.0319 | 0.0314 |
| cluster3139rr | -5753.8375 | 0.0673 | 0.1116 | 1.6587 | 0.0750 | 0.0213 |
| cluster3140rr | -3560.007 | 0.0411 | 0.0152 | 0.3699 | 0.0740 | 0.0043 |
| cluster3141rr | -2415.2431 | 0.0096 | 0.0034 | 0.3550 | 0.0132 | 0.0032 |
| cluster3142rr | -1633.2804 | 0.0644 | 0.0091 | 0.1408 | 1.2984 | 0.0187 |
| cluster3143rr | -3223.85 | 0.0163 | 0.0080 | 0.4893 | 0.0001 | 0.0000 |
| cluster3144rr | -2087.5661 | 0.0038 | 0.0339 | 9.0223 | 0.0571 | 0.0033 |
| cluster3145rr | -6530.7158 | 0.0053 | 0.0077 | 1.4558 | 0.0065 | 0.0024 |
| cluster3146rr | -1738.4981 | 0.0039 | 0.0104 | 2.6658 | 0.0003 | 0.0051 |
| cluster3147rr | -4471.5212 | 0.0057 | 0.0055 | 0.9723 | 0.0185 | 0.0027 |
| cluster3148rr | -6017.6258 | 0.0261 | 0.0167 | 0.6404 | 0.1048 | 0.0077 |
| cluster3149rr | -2371.9875 | 0.0034 | 0.0015 | 0.4352 | 0.0001 | 0.0000 |
| cluster3150rr | -4524.0271 | 0.0691 | 0.0247 | 0.3572 | 0.0278 | 0.0015 |
| cluster3151rr | -3969.564 | 0.0538 | 0.0323 | 0.6005 | 0.0001 | 0.0000 |
| cluster3152rr | -8737.9555 | 0.0422 | 0.0825 | 1.9557 | 0.1410 | 0.0241 |
| cluster3153rr | -4516.2302 | 0.0410 | 0.0236 | 0.5763 | 0.0744 | 0.0071 |
| cluster3154rr | -5343.172 | 0.0138 | 0.0149 | 1.0785 | 0.0287 | 0.0060 |
| cluster3155rr | -3013.4341 | 0.0079 | 0.0083 | 1.0446 | 0.0442 | 0.0030 |
| cluster3156rr | -6786.0375 | 0.0239 | 0.0383 | 1.6010 | 0.0595 | 0.0107 |
| cluster3157rr | -3280.9767 | 0.0996 | 0.0287 | 0.2883 | 0.0850 | 0.0044 |
| cluster3158rr | -3372.8509 | 0.0161 | 0.0142 | 0.8815 | 0.0122 | 0.0128 |

| | | | | | | |
|---|---|---|---|---|---|---|
| cluster3159rr | -5801.7696 | 0.0383 | 0.0672 | 1.7557 | 0.1901 | 0.0284 |
| cluster3160rr | -1099.804 | 0.0538 | 0.0091 | 0.1696 | 0.0425 | 0.0046 |
| cluster3161rr | -4981.5489 | 0.0305 | 0.0463 | 1.5170 | 0.0545 | 0.0082 |
| cluster3162rr | -3639.5187 | 0.0089 | 0.0040 | 0.4499 | 0.0174 | 0.0008 |
| cluster3163rr | -3430.0933 | 0.0353 | 0.0476 | 1.3472 | 0.0485 | 0.0091 |
| cluster3164rr | -6176.3606 | 0.0528 | 0.0638 | 1.2095 | 0.1600 | 0.0176 |
| cluster3165rr | -8047.5491 | 0.0515 | 0.0751 | 1.4571 | 0.0999 | 0.0211 |
| cluster3166rr | -4633.1464 | 0.0237 | 0.0147 | 0.6232 | 0.0324 | 0.0174 |
| cluster3167rr | -5313.1468 | 0.0907 | 0.0432 | 0.4767 | 0.1529 | 0.0074 |
| cluster3168rr | -7088.571 | 0.0195 | 0.0232 | 1.1899 | 0.0108 | 0.0114 |
| cluster3169rr | -5029.1256 | 0.0286 | 0.0425 | 1.4871 | 0.0483 | 0.0100 |
| cluster3170rr | -1623.083 | 0.0136 | 0.0024 | 0.1751 | 0.0001 | 0.0000 |
| cluster3171rr | -6157.6772 | 0.0662 | 0.0478 | 0.7231 | 0.0344 | 0.0098 |
| cluster3172rr | -6262.224 | 0.0278 | 0.0214 | 0.7703 | 0.0293 | 0.0087 |
| cluster3173rr | -4457.6382 | 0.0152 | 0.0164 | 1.0781 | 0.0094 | 0.0011 |
| cluster3174rr | -1301.2255 | 0.0021 | 0.0017 | 0.8069 | 0.0001 | 0.0000 |
| cluster3175rr | -1937.1279 | 0.0169 | 0.0107 | 0.6338 | 0.0001 | 0.0000 |
| cluster3176rr | -3887.5773 | 0.0256 | 0.0128 | 0.4984 | 0.0142 | 0.0025 |
| cluster3177rr | -3506.7455 | 0.0513 | 0.0250 | 0.4880 | 0.0657 | 0.0161 |
| cluster3178rr | -6745.2006 | 0.0116 | 0.0128 | 1.1047 | 0.0096 | 0.0093 |
| cluster3179rr | -4617.1274 | 0.0453 | 0.1125 | 2.4837 | 2.3577 | 0.0341 |
| cluster3180rr | -1069.3655 | 0.0022 | 0.0069 | 3.0882 | 0.0533 | 0.0026 |
| cluster3181rr | -6693.137 | 0.0223 | 0.0201 | 0.9042 | 0.0177 | 0.0149 |
| cluster3182rr | -2530.5184 | 0.0069 | 0.0035 | 0.5097 | 0.0043 | 0.0010 |
| cluster3183rr | -4042.0067 | 0.0779 | 0.0687 | 0.8816 | 0.1139 | 0.0381 |
| cluster3184rr | -4207.4929 | 0.0325 | 0.0168 | 0.5158 | 0.0242 | 0.0026 |
| cluster3185rr | -3544.2331 | 0.0153 | 0.0239 | 1.5621 | 0.0705 | 0.0066 |
| cluster3186rr | -2289.3322 | 0.0255 | 0.0197 | 0.7721 | 0.0664 | 0.0085 |
| cluster3187rr | -5390.2918 | 0.0186 | 0.0183 | 0.9875 | 0.0577 | 0.0071 |
| cluster3188rr | -4445.3539 | 0.0895 | 0.0422 | 0.4716 | 0.0931 | 0.0375 |
| cluster3189rr | -6228.0426 | 0.0402 | 0.0605 | 1.5061 | 0.0795 | 0.0107 |
| cluster3190rr | -5398.1915 | 0.0309 | 0.0554 | 1.7920 | 0.0340 | 0.0072 |
| cluster3191rr | -2827.8663 | 0.0393 | 0.0288 | 0.7339 | 0.1395 | 0.0296 |
| cluster3192rr | -5886.9431 | 0.0532 | 0.0282 | 0.5311 | 0.0430 | 0.0151 |
| cluster3193rr | -3456.8856 | 0.0359 | 0.0172 | 0.4787 | 0.0434 | 0.0034 |
| cluster3194rr | -6981.449 | 0.0571 | 0.0764 | 1.3396 | 0.0795 | 0.0230 |
| cluster3195rr | -3959.9816 | 0.0195 | 0.0189 | 0.9684 | 0.0092 | 0.0048 |
| cluster3196rr | -2244.4742 | 0.0049 | 0.0011 | 0.2196 | 0.0001 | 0.0000 |
| cluster3197rr | -5440.6921 | 0.0441 | 0.0201 | 0.4574 | 0.1128 | 0.0299 |
| cluster3198rr | -6428.4515 | 0.0219 | 0.0398 | 1.8198 | 0.0619 | 0.0116 |
| cluster3199rr | -2970.7099 | 0.0076 | 0.0052 | 0.6882 | 0.0723 | 0.0092 |
| cluster3200rr | -4268.9634 | 0.0138 | 0.0165 | 1.1908 | 0.0001 | 0.0000 |
| cluster3201rr | -4475.4858 | 0.1048 | 0.0315 | 0.3010 | 0.1874 | 0.0132 |
| cluster3202rr | -6070.422 | 0.0410 | 0.0277 | 0.6754 | 0.0739 | 0.0071 |

| | | | | | | |
|---|---|---|---|---|---|---|
| cluster3203rr | -3017.433 | 0.0492 | 0.0765 | 1.5566 | 0.0540 | 0.0067 |
| cluster3204rr | -1875.6927 | 0.0430 | 0.0107 | 0.2482 | 0.0001 | 0.0000 |
| cluster3205rr | -2015.4128 | 0.0066 | 0.0015 | 0.2334 | 0.0001 | 0.0000 |
| cluster3206rr | -3913.537 | 0.0319 | 0.0349 | 1.0931 | 0.0699 | 0.0112 |
| cluster3207rr | -5500.6036 | 0.0143 | 0.0152 | 1.0627 | 0.0111 | 0.0053 |
| cluster3208rr | -3284.7968 | 0.0056 | 0.0028 | 0.5041 | 0.0001 | 0.0000 |
| cluster3209rr | -5549.5686 | 0.0219 | 0.0304 | 1.3919 | 0.0687 | 0.0075 |
| cluster3210rr | -3377.9059 | 0.0029 | 0.0118 | 4.0285 | 0.0050 | 0.0010 |
| cluster3211rr | -3434.2494 | 0.0298 | 0.0362 | 1.2145 | 0.0607 | 0.0074 |
| cluster3212rr | -2811.2351 | 0.0026 | 0.0012 | 0.4712 | 0.0001 | 0.0000 |
| cluster3213rr | -4613.1135 | 0.1030 | 0.0758 | 0.7366 | 0.0560 | 0.0363 |
| cluster3214rr | -6050.3651 | 0.0503 | 0.0315 | 0.6256 | 0.0305 | 0.0136 |
| cluster3215rr | -2279.2466 | 0.0024 | 0.0005 | 0.1909 | 0.0001 | 0.0000 |
| cluster3216rr | -4525.4827 | 0.0181 | 0.0667 | 3.6928 | 0.1377 | 0.0275 |
| cluster3217rr | -3822.9994 | 0.0458 | 0.0237 | 0.5176 | 0.0203 | 0.0066 |
| cluster3218rr | -5083.1483 | 0.0231 | 0.0387 | 1.6774 | 0.0606 | 0.0054 |
| cluster3219rr | -5449.4152 | 0.0118 | 0.0139 | 1.1749 | 0.0209 | 0.0030 |
| cluster3220rr | -2923.5613 | 0.0333 | 0.0146 | 0.4390 | 0.0165 | 0.0060 |
| cluster3221rr | -5714.5024 | 0.0239 | 0.0495 | 2.0723 | 0.3927 | 0.0265 |
| cluster3222rr | -2618.7333 | 0.0456 | 0.0241 | 0.5282 | 0.0066 | 0.0018 |
| cluster3223rr | -3465.8052 | 0.0989 | 0.0769 | 0.7778 | 0.0001 | 0.0000 |
| cluster3224rr | -1597.2435 | 0.0001 | 0.0000 | 0.3098 | 0.0001 | 0.0000 |
| cluster3225rr | -6093.5293 | 0.0270 | 0.0349 | 1.2941 | 0.1923 | 0.0146 |
| cluster3226rr | -3709.2008 | 0.0284 | 0.0285 | 1.0040 | 0.2090 | 0.0325 |
| cluster3227rr | -4398.6825 | 0.0690 | 0.0845 | 1.2247 | 0.0704 | 0.0133 |
| cluster3228rr | -3619.8244 | 0.0269 | 0.0444 | 1.6474 | 0.1299 | 0.0203 |
| cluster3229rr | -3868.1281 | 0.0327 | 0.0231 | 0.7058 | 0.0104 | 0.0039 |
| cluster3230rr | -3477.7308 | 0.0129 | 0.0066 | 0.5107 | 0.0001 | 0.0000 |
| cluster3231rr | -4528.1842 | 0.0232 | 0.0196 | 0.8454 | 0.0432 | 0.0059 |
| cluster3232rr | -5050.9781 | 0.0202 | 0.0279 | 1.3786 | 0.0638 | 0.0111 |
| cluster3233rr | -3436.5167 | 0.0410 | 0.0435 | 1.0628 | 0.0426 | 0.0214 |
| cluster3234rr | -3819.0382 | 0.0568 | 0.0365 | 0.6425 | 0.0287 | 0.0168 |
| cluster3235rr | -7270.1732 | 0.0346 | 0.0740 | 2.1427 | 0.0669 | 0.0158 |
| cluster3236rr | -3953.3549 | 0.0290 | 0.0390 | 1.3464 | 0.0775 | 0.0176 |
| cluster3237rr | -3243.7937 | 0.0551 | 0.0543 | 0.9858 | 0.1020 | 0.0178 |
| cluster3238rr | -2781.5634 | 0.0628 | 0.0215 | 0.3425 | 0.0001 | 0.0000 |
| cluster3239rr | -2740.6504 | 0.0219 | 0.0099 | 0.4512 | 0.0308 | 0.0040 |
| cluster3240rr | -6960.843 | 0.0341 | 0.0292 | 0.8549 | 0.0222 | 0.0136 |
| cluster3241rr | -5756.287 | 0.0630 | 0.0635 | 1.0075 | 0.0823 | 0.0134 |
| cluster3242rr | -7155.8137 | 0.0382 | 0.0774 | 2.0291 | 0.1999 | 0.0185 |
| cluster3243rr | -5984.0205 | 0.1484 | 0.0510 | 0.3437 | 0.1792 | 0.0304 |
| cluster3244rr | -6473.3167 | 0.0611 | 0.0410 | 0.6709 | 0.0526 | 0.0262 |
| cluster3245rr | -2585.5379 | 0.0027 | 0.0030 | 1.0862 | 0.0076 | 0.0011 |
| cluster3246rr | -5134.5341 | 0.0447 | 0.0467 | 1.0443 | 0.0343 | 0.0363 |

| | | | | | | |
|---|---|---|---|---|---|---|
| cluster3247rr | -849.10166 | 0.0659 | 0.0434 | 0.6577 | 0.0363 | 0.0040 |
| cluster3248rr | -3481.2408 | 0.0184 | 0.0198 | 1.0755 | 0.0949 | 0.0103 |
| cluster3249rr | -3479.7624 | 0.0150 | 0.0161 | 1.0714 | 0.0682 | 0.0067 |
| cluster3250rr | -3547.9895 | 0.1229 | 0.0458 | 0.3721 | 0.0210 | 0.0038 |
| cluster3251rr | -4463.9804 | 0.0521 | 0.0387 | 0.7435 | 0.1528 | 0.0167 |
| cluster3252rr | -7422.2473 | 0.0608 | 0.0978 | 1.6077 | 0.0945 | 0.0275 |
| cluster3253rr | -4317.7481 | 0.0709 | 0.0557 | 0.7851 | 0.0700 | 0.0113 |
| cluster3254rr | -3104.9247 | 0.0290 | 0.0292 | 1.0075 | 0.1560 | 0.0163 |
| cluster3255rr | -5562.2071 | 0.0137 | 0.0320 | 2.3382 | 0.0541 | 0.0193 |
| cluster3256rr | -6096.4157 | 0.0363 | 0.0356 | 0.9811 | 0.0890 | 0.0231 |
| cluster3257rr | -5143.1168 | 0.0072 | 0.0059 | 0.8185 | 0.0026 | 0.0018 |
| cluster3258rr | -1919.1321 | 0.0015 | 0.0007 | 0.4894 | 0.0001 | 0.0000 |
| cluster3259rr | -3556.4357 | 0.0368 | 0.0102 | 0.2766 | 0.0300 | 0.0053 |
| cluster3260rr | -4480.9445 | 0.0150 | 0.0094 | 0.6288 | 0.0062 | 0.0027 |
| cluster3261rr | -4920.1793 | 0.0458 | 0.0894 | 1.9509 | 0.7909 | 0.0563 |
| cluster3262rr | -5790.1119 | 0.0175 | 0.0294 | 1.6807 | 0.0455 | 0.0054 |
| cluster3263rr | -2047.025 | 0.0863 | 0.0520 | 0.6029 | 0.5071 | 0.0640 |
| cluster3264rr | -1834.9466 | 0.0106 | 0.0040 | 0.3755 | 0.0001 | 0.0000 |
| cluster3265rr | -4147.1191 | 0.0027 | 0.0027 | 1.0139 | 0.5905 | 0.0000 |
| cluster3266rr | -2853.6835 | 0.0322 | 0.0110 | 0.3405 | 0.0001 | 0.0000 |
| cluster3267rr | -4124.8476 | 0.0327 | 0.0229 | 0.7002 | 0.0244 | 0.0091 |
| cluster3268rr | -2804.2311 | 0.0321 | 0.0256 | 0.7965 | 0.0912 | 0.0208 |
| cluster3269rr | -3538.3944 | 0.0571 | 0.0502 | 0.8784 | 0.0618 | 0.0064 |
| cluster3270rr | -2185.2993 | 0.0257 | 0.0340 | 1.3251 | 0.0371 | 0.0038 |
| cluster3271rr | -4112.8714 | 0.0122 | 0.0071 | 0.5870 | 0.0135 | 0.0046 |
| cluster3272rr | -939.31394 | 0.3460 | 0.0193 | 0.0559 | 0.2052 | 0.0251 |
| cluster3273rr | -4002.9761 | 0.1037 | 0.1098 | 1.0591 | 0.2354 | 0.0353 |
| cluster3274rr | -2030.5522 | 0.0254 | 0.0442 | 1.7395 | 0.0873 | 0.0222 |
| cluster3275rr | -4551.1739 | 0.0480 | 0.0558 | 1.1613 | 0.0546 | 0.0088 |
| cluster3276rr | -2484.9318 | 0.0830 | 0.0163 | 0.1962 | 0.1337 | 0.0188 |
| cluster3277rr | -3226.8449 | 0.0466 | 0.0642 | 1.3785 | 0.1003 | 0.0124 |
| cluster3278rr | -4577.2235 | 0.0033 | 0.0014 | 0.4332 | 0.0001 | 0.0000 |
| cluster3279rr | -3733.0686 | 0.0167 | 0.0198 | 1.1850 | 0.0168 | 0.0051 |
| cluster3280rr | -3787.1914 | 0.0230 | 0.0146 | 0.6358 | 0.0001 | 0.0000 |
| cluster3281rr | -1964.2724 | 0.0173 | 0.0240 | 1.3904 | 0.1054 | 0.0120 |
| cluster3282rr | -2049.165 | 0.0054 | 0.0040 | 0.7409 | 0.0106 | 0.0007 |
| cluster3283rr | -5463.5367 | 0.0594 | 0.0382 | 0.6432 | 0.0886 | 0.0457 |
| cluster3284rr | -5979.8176 | 0.0336 | 0.0335 | 0.9957 | 0.0595 | 0.0052 |
| cluster3285rr | -2362.2346 | 0.0078 | 0.0035 | 0.4471 | 0.0001 | 0.0000 |
| cluster3286rr | -2209.2334 | 0.0319 | 0.0118 | 0.3693 | 0.0216 | 0.0017 |
| cluster3287rr | -1377.2634 | 0.0124 | 0.0089 | 0.7166 | 0.0293 | 0.0020 |
| cluster3288rr | -1239.4741 | 0.0058 | 0.0112 | 1.9222 | 999.0000 | 0.0025 |
| cluster3289rr | -5793.5533 | 0.0410 | 0.0148 | 0.3600 | 0.0903 | 0.0258 |
| cluster3290rr | -1252.5731 | 0.0221 | 0.0162 | 0.7359 | 0.0001 | 0.0000 |

| | | | | | | |
|---|---|---|---|---|---|---|
| cluster3291rr | -4787.6359 | 0.0410 | 0.0319 | 0.7788 | 0.0340 | 0.0196 |
| cluster3292rr | -2772.6531 | 0.0233 | 0.0148 | 0.6327 | 0.0662 | 0.0091 |
| cluster3293rr | -3183.5311 | 0.0106 | 0.0092 | 0.8664 | 0.0039 | 0.0030 |
| cluster3294rr | -2882.5872 | 0.0144 | 0.0124 | 0.8568 | 0.0062 | 0.0021 |
| cluster3295rr | -2241.1436 | 0.0017 | 0.0008 | 0.4954 | 0.0001 | 0.0000 |
| cluster3296rr | -1799.1212 | 0.0040 | 0.0055 | 1.3659 | 0.0225 | 0.0015 |
| cluster3297rr | -2352.9098 | 0.1871 | 0.0880 | 0.4705 | 0.0731 | 0.0219 |
| cluster3298rr | -2944.6499 | 0.0042 | 0.0036 | 0.8566 | 0.0029 | 0.0018 |
| cluster3299rr | -7904.4223 | 0.0195 | 0.1340 | 6.8641 | 0.8653 | 0.0141 |
| cluster3300rr | -2252.1018 | 0.0051 | 0.0053 | 1.0411 | 0.0001 | 0.0000 |
| cluster3301rr | -1798.0836 | 0.0650 | 0.0214 | 0.3287 | 0.0101 | 0.0061 |
| cluster3302rr | -4535.3754 | 0.0589 | 0.0411 | 0.6976 | 0.3047 | 0.0916 |
| cluster3303rr | -4480.4758 | 0.0338 | 0.0422 | 1.2495 | 0.0415 | 0.0328 |
| cluster3304rr | -2860.5438 | 0.0254 | 0.0109 | 0.4282 | 0.0129 | 0.0022 |
| cluster3305rr | -3802.5533 | 0.0573 | 0.0573 | 1.0006 | 0.1370 | 0.0216 |
| cluster3306rr | -5012.8438 | 0.0619 | 0.0702 | 1.1347 | 0.3403 | 0.0247 |
| cluster3307rr | -1668.2629 | 0.0031 | 0.0035 | 1.1304 | 0.0524 | 0.0051 |
| cluster3308rr | -4382.9492 | 0.0219 | 0.0220 | 1.0034 | 0.0059 | 0.0047 |
| cluster3309rr | -3384.6284 | 0.1245 | 0.0467 | 0.3755 | 0.0819 | 0.0186 |
| cluster3310rr | -4171.8052 | 0.0134 | 0.0107 | 0.7928 | 0.0394 | 0.0104 |
| cluster3311rr | -6158.1203 | 0.0546 | 0.0440 | 0.8054 | 0.0561 | 0.0291 |
| cluster3312rr | -1923.2136 | 0.0189 | 0.0361 | 1.9050 | 999.0000 | 0.0202 |
| cluster3313rr | -2508.5471 | 0.0106 | 0.0043 | 0.4061 | 0.0197 | 0.0031 |
| cluster3314rr | -7138.3241 | 0.0371 | 0.0953 | 2.5676 | 0.1226 | 0.0142 |
| cluster3315rr | -3723.0939 | 0.0059 | 0.0041 | 0.6897 | 0.0055 | 0.0032 |
| cluster3316rr | -5974.4673 | 0.0844 | 0.0890 | 1.0538 | 0.0928 | 0.0145 |
| cluster3317rr | -1142.4909 | 0.0033 | 0.0025 | 0.7491 | 0.0001 | 0.0000 |
| cluster3318rr | -3441.538 | 0.1022 | 0.1140 | 1.1154 | 0.1387 | 0.0796 |
| cluster3319rr | -3763.4809 | 0.0297 | 0.0450 | 1.5184 | 0.1543 | 0.0175 |
| cluster3320rr | -2812.8534 | 0.0573 | 0.0596 | 1.0413 | 0.0134 | 0.0014 |
| cluster3321rr | -2918.9503 | 0.0011 | 0.0009 | 0.8570 | 0.0499 | 0.0025 |
| cluster3322rr | -3278.1366 | 0.0379 | 0.0566 | 1.4921 | 0.0754 | 0.0168 |
| cluster3323rr | -5518.6423 | 0.0467 | 0.0463 | 0.9921 | 0.1014 | 0.0161 |
| cluster3324rr | -1397.6751 | 0.0354 | 0.0380 | 1.0714 | 0.0001 | 0.0000 |
| cluster3325rr | -2061.1568 | 0.0186 | 0.0207 | 1.1146 | 0.0197 | 0.0056 |
| cluster3326rr | -1703.5044 | 0.0757 | 0.0187 | 0.2465 | 0.0001 | 0.0000 |
| cluster3327rr | -3864.9383 | 0.0248 | 0.0187 | 0.7558 | 0.0285 | 0.0115 |
| cluster3328rr | -2623.8382 | 0.0102 | 0.0170 | 1.6719 | 6.9268 | 0.0000 |
| cluster3329rr | -3148.7083 | 0.0513 | 0.0915 | 1.7823 | 0.3623 | 0.0473 |
| cluster3330rr | -4710.6475 | 0.0457 | 0.0626 | 1.3692 | 0.0648 | 0.0101 |
| cluster3331rr | -2869.1606 | 0.0736 | 0.0322 | 0.4371 | 0.0770 | 0.0068 |
| cluster3332rr | -6186.9349 | 0.0551 | 0.0855 | 1.5533 | 0.1670 | 0.0263 |
| cluster3333rr | -3567.2629 | 0.0173 | 0.0277 | 1.5990 | 0.2108 | 0.0139 |
| cluster3334rr | -1375.8791 | 0.0565 | 0.0293 | 0.5185 | 0.3245 | 0.0318 |

| | | | | | | |
|---|---|---|---|---|---|---|
| cluster3335rr | -2855.3126 | 0.0343 | 0.0325 | 0.9475 | 0.0096 | 0.0061 |
| cluster3336rr | -3026.9845 | 0.0044 | 0.0019 | 0.4253 | 0.0001 | 0.0000 |
| cluster3337rr | -2619.9759 | 0.0288 | 0.0085 | 0.2959 | 0.0482 | 0.0077 |
| cluster3338rr | -6299.0993 | 0.0343 | 0.0672 | 1.9602 | 0.1009 | 0.0181 |
| cluster3339rr | -1907.2377 | 0.0564 | 0.0494 | 0.8752 | 0.0135 | 0.0105 |
| cluster3340rr | -1624.9375 | 0.2186 | 0.0226 | 0.1033 | 0.0935 | 0.0339 |
| cluster3341rr | -1576.9999 | 0.0108 | 0.0177 | 1.6338 | 0.0131 | 0.0020 |
| cluster3342rr | -2555.4477 | 0.0191 | 0.0300 | 1.5708 | 0.0532 | 0.0069 |
| cluster3343rr | -2926.4924 | 0.0005 | 0.0005 | 1.0325 | 0.0001 | 0.0000 |
| cluster3344rr | -3148.0069 | 0.0291 | 0.0113 | 0.3884 | 0.0116 | 0.0032 |
| cluster3345rr | -2788.8956 | 0.0101 | 0.0183 | 1.8157 | 0.0927 | 0.0115 |
| cluster3346rr | -3607.1407 | 0.0044 | 0.0031 | 0.7008 | 0.0001 | 0.0001 |
| cluster3347rr | -4175.3144 | 0.0495 | 0.0225 | 0.4545 | 0.0313 | 0.0095 |
| cluster3348rr | -4559.8206 | 0.0488 | 0.0248 | 0.5076 | 0.0636 | 0.0075 |
| cluster3349rr | -4213.1317 | 0.0067 | 0.0084 | 1.2523 | 0.0186 | 0.0060 |
| cluster3350rr | -4778.1113 | 0.0476 | 0.0555 | 1.1654 | 0.0849 | 0.0128 |
| cluster3351rr | -3131.0348 | 0.0138 | 0.0149 | 1.0789 | 0.0639 | 0.0065 |
| cluster3352rr | -4084.9629 | 0.0549 | 0.0356 | 0.6483 | 0.0767 | 0.0100 |
| cluster3353rr | -1586.004 | 0.0110 | 0.0059 | 0.5377 | 0.0001 | 0.0000 |
| cluster3354rr | -2117.6995 | 0.0038 | 0.0016 | 0.4227 | 0.0891 | 0.0025 |
| cluster3355rr | -4267.8354 | 0.0316 | 0.0326 | 1.0310 | 0.2295 | 0.0478 |
| cluster3356rr | -4590.152 | 0.0463 | 0.0245 | 0.5295 | 0.0114 | 0.0041 |
| cluster3357rr | -1290.063 | 0.0285 | 0.0189 | 0.6631 | 0.0158 | 0.0016 |
| cluster3358rr | -4171.9183 | 0.0016 | 0.0008 | 0.4925 | 0.0017 | 0.0007 |
| cluster3359rr | -6031.0669 | 0.0305 | 0.0525 | 1.7224 | 0.2046 | 0.0213 |
| cluster3360rr | -2471.5944 | 0.0370 | 0.0337 | 0.9096 | 0.0437 | 0.0190 |
| cluster3361rr | -5030.1654 | 0.0322 | 0.0588 | 1.8256 | 0.1083 | 0.0170 |
| cluster3362rr | -3690.9974 | 0.0180 | 0.0126 | 0.6966 | 0.0069 | 0.0038 |
| cluster3363rr | -3719.3165 | 0.0019 | 0.0065 | 3.4650 | 0.0155 | 0.0021 |
| cluster3364rr | -4356.1514 | 0.0231 | 0.0394 | 1.7039 | 0.0303 | 0.0083 |
| cluster3365rr | -5563.7598 | 0.0588 | 0.0570 | 0.9692 | 0.1286 | 0.0148 |
| cluster3366rr | -3145.5898 | 0.0257 | 0.0648 | 2.5274 | 0.2108 | 0.0119 |
| cluster3367rr | -3693.7543 | 0.1388 | 0.0714 | 0.5143 | 0.1151 | 0.0792 |
| cluster3368rr | -1652.1421 | 0.0313 | 0.0064 | 0.2048 | 0.0001 | 0.0000 |
| cluster3369rr | -3760.9751 | 0.0129 | 0.0181 | 1.4065 | 0.0663 | 0.0097 |
| cluster3370rr | -6384.7176 | 0.0922 | 0.0826 | 0.8968 | 0.2208 | 0.0210 |
| cluster3371rr | -2933.7179 | 0.0058 | 0.0028 | 0.4851 | 0.0346 | 0.0009 |
| cluster3372rr | -3840.6381 | 0.0094 | 0.0109 | 1.1685 | 0.0044 | 0.0023 |
| cluster3373rr | -5277.5316 | 0.0085 | 0.0139 | 1.6350 | 0.0076 | 0.0013 |
| cluster3374rr | -5121.4659 | 0.0623 | 0.1266 | 2.0339 | 0.2721 | 0.0206 |
| cluster3375rr | -3051.705 | 0.0056 | 0.0024 | 0.4342 | 0.0001 | 0.0000 |
| cluster3376rr | -1529.8379 | 0.0400 | 0.0122 | 0.3053 | 0.0001 | 0.0000 |
| cluster3377rr | -2273.0624 | 0.0107 | 0.0163 | 1.5208 | 0.0375 | 0.0054 |
| cluster3378rr | -6508.1598 | 0.0477 | 0.0452 | 0.9482 | 0.0491 | 0.0377 |

| | | | | | | |
|---|---|---|---|---|---|---|
| cluster3379rr | -1568.5083 | 0.0333 | 0.0223 | 0.6679 | 0.0425 | 0.0050 |
| cluster3380rr | -2653.3256 | 0.0161 | 0.0197 | 1.2248 | 0.0067 | 0.0012 |
| cluster3381rr | -2267.7991 | 0.0351 | 0.0223 | 0.6353 | 0.0264 | 0.0140 |
| cluster3382rr | -885.62258 | 0.0067 | 0.0031 | 0.4707 | 0.0001 | 0.0000 |
| cluster3383rr | -5125.7808 | 0.0394 | 0.0715 | 1.8141 | 0.3170 | 0.0613 |
| cluster3384rr | -1972.3176 | 0.0339 | 0.0225 | 0.6656 | 0.0318 | 0.0079 |
| cluster3385rr | -3770.8625 | 0.0207 | 0.0274 | 1.3229 | 0.0420 | 0.0076 |
| cluster3386rr | -4154.3187 | 0.0134 | 0.0178 | 1.3310 | 0.0334 | 0.0065 |
| cluster3387rr | -3023.4514 | 0.0329 | 0.0255 | 0.7762 | 0.1366 | 0.0335 |
| cluster3388rr | -2809.2204 | 0.0245 | 0.0195 | 0.7946 | 0.0370 | 0.0059 |
| cluster3389rr | -1787.2217 | 0.0082 | 0.0131 | 1.5945 | 0.0443 | 0.0053 |
| cluster3390rr | -5117.0389 | 0.0627 | 0.0387 | 0.6176 | 0.0735 | 0.0256 |
| cluster3391rr | -2414.1855 | 0.0070 | 0.0042 | 0.5967 | 0.0423 | 0.0035 |
| cluster3392rr | -2527.2271 | 0.0072 | 0.0135 | 1.8867 | 0.0309 | 0.0056 |
| cluster3393rr | -3175.9786 | 0.0491 | 0.0465 | 0.9483 | 0.1498 | 0.0206 |
| cluster3394rr | -712.64912 | 0.0021 | 0.0019 | 0.8918 | 0.0147 | 0.0070 |
| cluster3395rr | -2714.5609 | 0.0147 | 0.0100 | 0.6841 | 0.0161 | 0.0102 |
| cluster3396rr | -4983.7846 | 0.0392 | 0.0217 | 0.5553 | 0.0202 | 0.0087 |
| cluster3397rr | -4637.2242 | 0.0247 | 0.0376 | 1.5248 | 0.1050 | 0.0129 |
| cluster3398rr | -3832.5204 | 0.0530 | 0.0320 | 0.6052 | 0.1381 | 0.0066 |
| cluster3399rr | -3230.6195 | 0.0430 | 0.0395 | 0.9184 | 0.0511 | 0.0198 |
| cluster3400rr | -1954.9796 | 0.1003 | 0.0245 | 0.2442 | 0.1283 | 0.0044 |
| cluster3401rr | -2870.6663 | 0.0165 | 0.0157 | 0.9533 | 0.0156 | 0.0030 |
| cluster3402rr | -5114.3627 | 0.0242 | 0.0178 | 0.7332 | 0.0136 | 0.0050 |
| cluster3403rr | -3664.6663 | 0.0188 | 0.0161 | 0.8546 | 0.0822 | 0.0104 |
| cluster3404rr | -716.69613 | 0.0001 | 0.0000 | 0.1155 | 0.0001 | 0.0000 |
| cluster3405rr | -1541.2642 | 0.1225 | 0.0545 | 0.4451 | 0.0922 | 0.0068 |
| cluster3406rr | -3797.0026 | 0.0222 | 0.0191 | 0.8618 | 0.0078 | 0.0087 |
| cluster3407rr | -6244.1164 | 0.0628 | 0.0622 | 0.9900 | 0.0504 | 0.0373 |
| cluster3408rr | -1896.6123 | 0.0058 | 0.0018 | 0.3084 | 0.0095 | 0.0014 |
| cluster3409rr | -4907.5138 | 0.0208 | 0.0345 | 1.6620 | 0.0436 | 0.0142 |
| cluster3410rr | -1442.2328 | 0.0254 | 0.0089 | 0.3491 | 0.0001 | 0.0000 |
| cluster3411rr | -2582.8993 | 0.0004 | 0.0003 | 0.7628 | 0.0001 | 0.0000 |
| cluster3412rr | -3390.1987 | 0.0752 | 0.0704 | 0.9372 | 0.1874 | 0.0187 |
| cluster3413rr | -2553.8058 | 0.0353 | 0.0360 | 1.0187 | 0.1626 | 0.0365 |
| cluster3414rr | -2521.1764 | 0.1484 | 0.0348 | 0.2348 | 0.0634 | 0.0102 |
| cluster3415rr | -3488.9861 | 0.0579 | 0.0340 | 0.5877 | 0.3250 | 0.0101 |
| cluster3416rr | -603.53788 | 0.0134 | 0.2947 | 21.9562 | 0.0266 | 0.0654 |
| cluster3417rr | -1214.6275 | 0.0011 | 0.0170 | 15.5739 | 0.0112 | 0.0085 |
| cluster3418rr | -2531.8295 | 0.0471 | 0.0472 | 1.0029 | 0.1417 | 0.0216 |
| cluster3419rr | -2668.0723 | 0.0857 | 0.0308 | 0.3597 | 0.6014 | 0.0168 |
| cluster3420rr | -1786.1063 | 0.2588 | 0.1267 | 0.4896 | 0.1899 | 0.0119 |
| cluster3421rr | -4207.7595 | 0.0114 | 0.0159 | 1.4006 | 0.0103 | 0.0034 |
| cluster3422rr | -5398.9947 | 0.0364 | 0.0236 | 0.6476 | 0.0268 | 0.0113 |

| | | | | | | |
|---|---|---|---|---|---|---|
| cluster3423rr | -4232.0979 | 0.0320 | 0.0800 | 2.4976 | 1.5852 | 0.0348 |
| cluster3424rr | -2856.4999 | 0.1040 | 0.0687 | 0.6607 | 0.0849 | 0.0194 |
| cluster3425rr | -2232.0719 | 0.0456 | 0.0181 | 0.3965 | 0.0001 | 0.0000 |
| cluster3426rr | -3409.8649 | 0.0202 | 0.0220 | 1.0868 | 0.0097 | 0.0093 |
| cluster3427rr | -3789.6108 | 0.0800 | 0.0505 | 0.6310 | 0.2983 | 0.0299 |
| cluster3428rr | -2685.1341 | 0.0688 | 0.0298 | 0.4335 | 0.0383 | 0.0110 |
| cluster3429rr | -4143.7994 | 0.0314 | 0.0251 | 0.8003 | 0.0153 | 0.0095 |
| cluster3430rr | -4823.3181 | 0.0380 | 0.0755 | 1.9865 | 0.0905 | 0.0170 |
| cluster3431rr | -4736.5368 | 0.0368 | 0.0597 | 1.6213 | 0.0848 | 0.0235 |
| cluster3432rr | -3782.7605 | 0.0366 | 0.0181 | 0.4944 | 0.0393 | 0.0097 |
| cluster3433rr | -5388.7217 | 0.1498 | 0.0628 | 0.4190 | 0.3226 | 0.0611 |
| cluster3434rr | -3295.0475 | 0.0375 | 0.0150 | 0.4007 | 0.0294 | 0.0075 |
| cluster3435rr | -4092.5011 | 0.0198 | 0.0215 | 1.0828 | 0.0396 | 0.0078 |
| cluster3436rr | -3133.3191 | 0.0214 | 0.0331 | 1.5503 | 0.0597 | 0.0098 |
| cluster3437rr | -2816.0319 | 0.0120 | 0.0133 | 1.1176 | 0.0206 | 0.0074 |
| cluster3438rr | -3823.4111 | 0.0353 | 0.0297 | 0.8419 | 0.0830 | 0.0114 |
| cluster3439rr | -2056.1165 | 0.0504 | 0.0471 | 0.9338 | 0.0945 | 0.0076 |
| cluster3440rr | -2314.0282 | 0.0249 | 0.0118 | 0.4751 | 0.0528 | 0.0032 |
| cluster3441rr | -2753.1319 | 0.0322 | 0.0481 | 1.4924 | 0.1348 | 0.0177 |
| cluster3442rr | -2922.4251 | 0.0132 | 0.0569 | 4.3208 | 0.0835 | 0.0091 |
| cluster3443rr | -3518.7274 | 0.0107 | 0.0163 | 1.5200 | 0.1462 | 0.0137 |
| cluster3444rr | -3385.5759 | 0.0106 | 0.0083 | 0.7823 | 0.0001 | 0.0000 |
| cluster3445rr | -2962.9524 | 0.0162 | 0.0070 | 0.4302 | 0.0030 | 0.0010 |
| cluster3446rr | -4061.0629 | 0.0288 | 0.0400 | 1.3883 | 0.0547 | 0.0128 |
| cluster3447rr | -5902.288 | 0.0262 | 0.0714 | 2.7259 | 0.5393 | 0.0325 |
| cluster3448rr | -2377.2154 | 0.0238 | 0.0365 | 1.5352 | 0.0001 | 0.0000 |
| cluster3449rr | -613.53865 | 0.0039 | 0.0006 | 0.1505 | 0.0001 | 0.0000 |
| cluster3450rr | -2512.5312 | 0.0154 | 0.0294 | 1.9143 | 0.1162 | 0.0148 |
| cluster3451rr | -3593.0539 | 0.0251 | 0.0347 | 1.3833 | 0.0628 | 0.0075 |
| cluster3452rr | -3964.8413 | 0.0469 | 0.0554 | 1.1821 | 0.0544 | 0.0074 |
| cluster3453rr | -1845.8418 | 0.0457 | 0.0209 | 0.4577 | 0.0084 | 0.0041 |
| cluster3454rr | -3695.7426 | 0.0115 | 0.0221 | 1.9303 | 0.0078 | 0.0044 |
| cluster3455rr | -3649.1173 | 0.0451 | 0.0123 | 0.2735 | 0.0338 | 0.0053 |
| cluster3456rr | -4229.9614 | 0.0389 | 0.0186 | 0.4791 | 0.0123 | 0.0017 |
| cluster3457rr | -3648.9246 | 0.0291 | 0.0342 | 1.1741 | 0.0464 | 0.0108 |
| cluster3458rr | -1806.0262 | 0.0318 | 0.0180 | 0.5654 | 0.0592 | 0.0117 |
| cluster3459rr | -2639.2485 | 0.0412 | 0.0717 | 1.7413 | 0.1500 | 0.0104 |
| cluster3460rr | -5579.2762 | 0.0299 | 0.0615 | 2.0553 | 0.0648 | 0.0109 |
| cluster3461rr | -3102.8864 | 0.0671 | 0.0687 | 1.0252 | 0.0598 | 0.0175 |
| cluster3462rr | -823.72779 | 0.0309 | 0.0035 | 0.1146 | 0.1610 | 0.0030 |
| cluster3463rr | -4771.3437 | 0.0484 | 0.0528 | 1.0905 | 0.0399 | 0.0080 |
| cluster3464rr | -5597.4135 | 0.0375 | 0.0752 | 2.0075 | 0.2591 | 0.0244 |
| cluster3465rr | -3953.9333 | 0.0113 | 0.0212 | 1.8822 | 0.0041 | 0.0046 |
| cluster3466rr | -2644.1902 | 0.0119 | 0.0123 | 1.0331 | 0.0158 | 0.0071 |

| | | | | | | |
|---|---|---|---|---|---|---|
| cluster3467rr | -4058.6841 | 0.0304 | 0.0277 | 0.9131 | 0.0800 | 0.0100 |
| cluster3468rr | -1845.4686 | 0.0053 | 0.0141 | 2.6565 | 0.0388 | 0.0102 |
| cluster3469rr | -2105.2635 | 0.0235 | 0.0233 | 0.9906 | 0.0001 | 0.0000 |
| cluster3470rr | -3884.4751 | 0.0127 | 0.0163 | 1.2836 | 0.0234 | 0.0034 |
| cluster3471rr | -4170.8059 | 0.0397 | 0.0423 | 1.0667 | 0.1180 | 0.0112 |
| cluster3472rr | -3721.7579 | 0.0187 | 0.0121 | 0.6489 | 0.0128 | 0.0055 |
| cluster3473rr | -3470.9163 | 0.0274 | 0.0207 | 0.7552 | 0.0267 | 0.0155 |
| cluster3474rr | -5362.592 | 0.0260 | 0.0722 | 2.7748 | 0.1444 | 0.0115 |
| cluster3475rr | -2340.0395 | 0.0111 | 0.0159 | 1.4366 | 0.0257 | 0.0074 |
| cluster3476rr | -5095.152 | 0.0203 | 0.0681 | 3.3478 | 0.2555 | 0.0251 |
| cluster3477rr | -3275.7472 | 0.0132 | 0.0037 | 0.2796 | 0.0356 | 0.0052 |
| cluster3478rr | -3581.0765 | 0.0211 | 0.0140 | 0.6655 | 0.0113 | 0.0056 |
| cluster3479rr | -2910.3958 | 0.0343 | 0.0845 | 2.4632 | 0.2022 | 0.0308 |
| cluster3480rr | -4508.8579 | 0.0324 | 0.0415 | 1.2841 | 0.0485 | 0.0068 |
| cluster3481rr | -1071.7406 | 0.0083 | 0.0322 | 3.8612 | 0.0001 | 0.0000 |
| cluster3482rr | -1966.1587 | 0.0804 | 0.0243 | 0.3024 | 0.0001 | 0.0000 |
| cluster3483rr | -3656.3474 | 0.0084 | 0.0091 | 1.0768 | 0.0360 | 0.0068 |
| cluster3484rr | -2759.9302 | 0.0236 | 0.0334 | 1.4140 | 0.1967 | 0.0227 |
| cluster3485rr | -3494.6831 | 0.0396 | 0.0379 | 0.9590 | 0.0703 | 0.0071 |
| cluster3486rr | -3175.0451 | 0.0494 | 0.0177 | 0.3588 | 0.0570 | 0.0164 |
| cluster3487rr | -2346.6405 | 0.0209 | 0.0200 | 0.9595 | 0.0548 | 0.0131 |
| cluster3488rr | -1482.9499 | 0.0260 | 0.0166 | 0.6385 | 0.0520 | 0.0071 |
| cluster3489rr | -4050.3841 | 0.0352 | 0.0519 | 1.4727 | 0.3020 | 0.0319 |
| cluster3490rr | -3431.2101 | 0.0191 | 0.0385 | 2.0144 | 0.0518 | 0.0102 |
| cluster3491rr | -3685.5371 | 0.0290 | 0.0293 | 1.0109 | 0.0750 | 0.0090 |
| cluster3492rr | -3372.4835 | 0.0375 | 0.0548 | 1.4632 | 0.0377 | 0.0043 |
| cluster3493rr | -4432.8899 | 0.0356 | 0.0305 | 0.8563 | 0.0190 | 0.0025 |
| cluster3494rr | -2865.1759 | 0.0178 | 0.0302 | 1.7005 | 0.0086 | 0.0132 |
| cluster3495rr | -2836.3182 | 0.0237 | 0.0229 | 0.9653 | 0.2049 | 0.0132 |
| cluster3496rr | -3231.6561 | 0.0229 | 0.0129 | 0.5604 | 0.0188 | 0.0105 |
| cluster3497rr | -1194.4396 | 0.0112 | 0.0021 | 0.1882 | 0.0317 | 0.0032 |
| cluster3498rr | -4133.5406 | 0.0462 | 0.0892 | 1.9312 | 0.0872 | 0.0105 |
| cluster3499rr | -3707.9932 | 0.0626 | 0.1053 | 1.6817 | 1.4877 | 0.0400 |
| cluster3500rr | -3421.6923 | 0.0177 | 0.0236 | 1.3347 | 0.0655 | 0.0089 |
| cluster3501rr | -2355.4874 | 0.0319 | 0.0226 | 0.7071 | 0.0036 | 0.0014 |
| cluster3502rr | -2168.6798 | 0.0132 | 0.0175 | 1.3317 | 0.0292 | 0.0079 |
| cluster3503rr | -1950.3838 | 0.0401 | 0.0582 | 1.4540 | 0.1371 | 0.0158 |
| cluster3504rr | -1142.0833 | 0.0076 | 0.0154 | 2.0164 | 0.0085 | 0.0038 |
| cluster3505rr | -2402.8941 | 0.0391 | 0.0760 | 1.9427 | 1.2577 | 0.0477 |
| cluster3506rr | -2544.4207 | 0.0193 | 0.0261 | 1.3517 | 0.1213 | 0.0133 |
| cluster3507rr | -5009.4 | 0.0441 | 0.0581 | 1.3170 | 0.3597 | 0.0254 |
| cluster3508rr | -5628.0597 | 0.0236 | 0.0858 | 3.6302 | 0.1774 | 0.0192 |
| cluster3509rr | -2401.973 | 0.0012 | 0.0268 | 22.5888 | 0.0001 | 0.0000 |
| cluster3510rr | -843.97507 | 0.1159 | 0.0508 | 0.4388 | 0.0514 | 0.0081 |

| | | | | | | |
|---|---|---|---|---|---|---|
| cluster3511rr | -1015.2642 | 0.0266 | 0.0211 | 0.7933 | 0.2442 | 0.0152 |
| cluster3512rr | -2356.7153 | 0.0338 | 0.0172 | 0.5089 | 0.0166 | 0.0013 |
| cluster3513rr | -958.0126 | 0.0807 | 0.0072 | 0.0897 | 0.0001 | 0.0000 |
| cluster3514rr | -5157.3676 | 0.0395 | 0.0263 | 0.6644 | 0.0224 | 0.0124 |
| cluster3515rr | -2604.5316 | 0.0122 | 0.0117 | 0.9608 | 0.0300 | 0.0050 |
| cluster3516rr | -4128.2861 | 0.0252 | 0.0271 | 1.0776 | 0.0649 | 0.0095 |
| cluster3517rr | -2410.1871 | 0.0026 | 0.0014 | 0.5305 | 0.0001 | 0.0000 |
| cluster3518rr | -4784.5514 | 0.0743 | 0.0845 | 1.1376 | 0.0904 | 0.0205 |
| cluster3519rr | -2126.8892 | 0.0441 | 0.0390 | 0.8860 | 0.0295 | 0.0116 |
| cluster3520rr | -1193.9854 | 0.1477 | 0.0723 | 0.4894 | 0.1629 | 0.0169 |
| cluster3521rr | -4217.766 | 0.0300 | 0.0498 | 1.6583 | 0.1183 | 0.0230 |
| cluster3522rr | -4751.1723 | 0.0293 | 0.0576 | 1.9689 | 0.0934 | 0.0120 |
| cluster3523rr | -1650.5713 | 0.0234 | 0.0352 | 1.5043 | 999.0000 | 0.0180 |
| cluster3524rr | -1963.4267 | 0.0121 | 0.0051 | 0.4205 | 0.0217 | 0.0014 |
| cluster3525rr | -2817.1817 | 0.0460 | 0.0268 | 0.5829 | 0.0432 | 0.0069 |
| cluster3526rr | -2997.0841 | 0.0017 | 0.0037 | 2.1845 | 0.0188 | 0.0025 |
| cluster3527rr | -4935.0996 | 0.0409 | 0.0822 | 2.0103 | 0.1132 | 0.0190 |
| cluster3528rr | -1968.2261 | 0.0113 | 0.0061 | 0.5424 | 0.0062 | 0.0015 |
| cluster3529rr | -3740.7266 | 0.0284 | 0.0519 | 1.8284 | 0.0990 | 0.0114 |
| cluster3530rr | -1951.8795 | 0.0242 | 0.0441 | 1.8232 | 0.0802 | 0.0175 |
| cluster3531rr | -2917.7683 | 0.0456 | 0.0393 | 0.8604 | 0.1982 | 0.0106 |
| cluster3532rr | -5557.7116 | 0.0634 | 0.0613 | 0.9681 | 0.0564 | 0.0340 |
| cluster3533rr | -2655.1512 | 0.0239 | 0.0108 | 0.4524 | 0.0305 | 0.0073 |
| cluster3534rr | -2560.5178 | 0.0640 | 0.0186 | 0.2912 | 0.0725 | 0.0069 |
| cluster3535rr | -3289.3624 | 0.0205 | 0.0523 | 2.5576 | 0.0374 | 0.0089 |
| cluster3536rr | -3557.1812 | 0.0356 | 0.0228 | 0.6423 | 0.0409 | 0.0167 |
| cluster3537rr | -3665.2824 | 0.0065 | 0.0090 | 1.3895 | 0.0141 | 0.0019 |
| cluster3538rr | -1387.6173 | 0.0399 | 0.0154 | 0.3861 | 0.0001 | 0.0000 |
| cluster3539rr | -2942.7991 | 0.0281 | 0.0950 | 3.3813 | 0.1171 | 0.0334 |
| cluster3540rr | -5363.5555 | 0.0862 | 0.1345 | 1.5599 | 0.1437 | 0.0213 |
| cluster3541rr | -3875.7863 | 0.0304 | 0.0200 | 0.6577 | 0.0139 | 0.0082 |
| cluster3542rr | -5425.0673 | 0.0539 | 0.1126 | 2.0887 | 0.1494 | 0.0176 |
| cluster3543rr | -4245.9441 | 0.0310 | 0.0700 | 2.2576 | 0.0898 | 0.0120 |
| cluster3544rr | -1087.8216 | 0.0120 | 0.0047 | 0.3889 | 0.0478 | 0.0044 |
| cluster3545rr | -5146.4459 | 0.0440 | 0.0692 | 1.5749 | 0.0879 | 0.0095 |
| cluster3546rr | -2669.9241 | 0.0158 | 0.0374 | 2.3691 | 0.1240 | 0.0239 |
| cluster3547rr | -1117.017 | 0.0093 | 0.0025 | 0.2640 | 0.0001 | 0.0000 |
| cluster3548rr | -1308.9721 | 0.0567 | 0.0106 | 0.1876 | 0.0593 | 0.0041 |
| cluster3549rr | -3232.7212 | 0.0352 | 0.0243 | 0.6916 | 0.0512 | 0.0234 |
| cluster3550rr | -1223.742 | 0.0091 | 0.0023 | 0.2567 | 0.0001 | 0.0000 |
| cluster3551rr | -2798.2704 | 0.0536 | 0.0172 | 0.3209 | 0.0311 | 0.0071 |
| cluster3552rr | -2384.1763 | 0.0282 | 0.0329 | 1.1685 | 0.0112 | 0.0132 |
| cluster3553rr | -2403.539 | 0.0295 | 0.0239 | 0.8097 | 0.0843 | 0.0119 |
| cluster3554rr | -3823.9137 | 0.0257 | 0.0186 | 0.7240 | 0.0492 | 0.0035 |

| | | | | | | |
|---|---|---|---|---|---|---|
| cluster3555rr | -1628.1039 | 0.0144 | 0.0255 | 1.7763 | 0.4051 | 0.0189 |
| cluster3556rr | -4203.3581 | 0.0228 | 0.0363 | 1.5886 | 0.1026 | 0.0081 |
| cluster3557rr | -1366.054 | 0.0734 | 0.1505 | 2.0501 | 0.0001 | 0.0000 |
| cluster3558rr | -4776.0861 | 0.0623 | 0.0738 | 1.1833 | 0.1106 | 0.0262 |
| cluster3559rr | -2457.658 | 0.0187 | 0.0231 | 1.2348 | 0.0075 | 0.0013 |
| cluster3560rr | -5081.7603 | 0.0447 | 0.0819 | 1.8329 | 0.3640 | 0.0394 |
| cluster3561rr | -1800.9038 | 0.0329 | 0.0137 | 0.4169 | 0.0001 | 0.0000 |
| cluster3562rr | -4269.6282 | 0.0323 | 0.0441 | 1.3637 | 0.1880 | 0.0193 |
| cluster3563rr | -1977.1087 | 0.0190 | 0.0116 | 0.6086 | 0.0001 | 0.0001 |
| cluster3564rr | -5021.5678 | 0.0766 | 0.0411 | 0.5371 | 0.0369 | 0.0142 |
| cluster3565rr | -3330.9669 | 0.0239 | 0.0442 | 1.8526 | 0.4731 | 0.0362 |
| cluster3566rr | -1890.3471 | 0.0431 | 0.0371 | 0.8595 | 0.0571 | 0.0065 |
| cluster3567rr | -2402.7897 | 0.0616 | 0.0614 | 0.9980 | 0.1554 | 0.0159 |
| cluster3568rr | -3168.2766 | 0.0283 | 0.0461 | 1.6277 | 0.0297 | 0.0071 |
| cluster3569rr | -3551.6267 | 0.0402 | 0.0716 | 1.7811 | 0.0435 | 0.0071 |
| cluster3570rr | -2385.2988 | 0.0118 | 0.0482 | 4.0811 | 0.2527 | 0.0268 |
| cluster3571rr | -2036.9029 | 0.0177 | 0.0231 | 1.3075 | 0.0176 | 0.0078 |
| cluster3572rr | -3449.8145 | 0.0428 | 0.0586 | 1.3684 | 0.1253 | 0.0303 |
| cluster3573rr | -3242.7753 | 0.0083 | 0.0215 | 2.6026 | 0.0342 | 0.0045 |
| cluster3574rr | -5130.6571 | 0.0728 | 0.0917 | 1.2592 | 0.2036 | 0.0150 |
| cluster3575rr | -3831.1903 | 0.0396 | 0.0454 | 1.1474 | 0.0406 | 0.0104 |
| cluster3576rr | -3274.7764 | 0.0354 | 0.0495 | 1.3997 | 0.0572 | 0.0071 |
| cluster3577rr | -2730.3297 | 0.0424 | 0.0210 | 0.4954 | 0.0204 | 0.0071 |
| cluster3578rr | -3391.4741 | 0.0335 | 0.0353 | 1.0539 | 0.0355 | 0.0110 |
| cluster3579rr | -2923.4008 | 0.0284 | 0.0309 | 1.0885 | 0.0162 | 0.0148 |
| cluster3580rr | -2417.7904 | 0.0100 | 0.0212 | 2.1259 | 0.0160 | 0.0049 |
| cluster3581rr | -1246.9356 | 0.0072 | 0.0139 | 1.9341 | 0.0139 | 0.0022 |
| cluster3582rr | -1106.9093 | 0.0266 | 0.0051 | 0.1931 | 0.0001 | 0.0000 |
| cluster3583rr | -2533.4362 | 0.0001 | 0.0001 | 1.0875 | 0.0001 | 0.0000 |
| cluster3584rr | -4868.473 | 0.0281 | 0.0717 | 2.5497 | 0.0640 | 0.0116 |
| cluster3585rr | -2092.262 | 0.0697 | 0.0251 | 0.3603 | 0.0201 | 0.0044 |
| cluster3586rr | -5218.4081 | 0.0605 | 0.0888 | 1.4678 | 0.2899 | 0.0417 |
| cluster3587rr | -3454.529 | 0.0331 | 0.0902 | 2.7225 | 0.5162 | 0.0472 |
| cluster3588rr | -3338.6217 | 0.0124 | 0.0053 | 0.4246 | 0.1258 | 0.0309 |
| cluster3589rr | -3012.8868 | 0.0364 | 0.0362 | 0.9944 | 0.2102 | 0.0205 |
| cluster3590rr | -4177.8593 | 0.0510 | 0.1098 | 2.1514 | 0.1368 | 0.0433 |
| cluster3591rr | -2874.38 | 0.0117 | 0.0060 | 0.5146 | 0.0141 | 0.0034 |
| cluster3592rr | -4700.5286 | 0.0323 | 0.0621 | 1.9257 | 0.1961 | 0.0170 |
| cluster3593rr | -3400.4475 | 0.0337 | 0.0699 | 2.0767 | 999.0000 | 0.0327 |
| cluster3594rr | -3173.4089 | 0.0521 | 0.0419 | 0.8045 | 0.1596 | 0.0221 |
| cluster3595rr | -2123.4976 | 0.0112 | 0.0052 | 0.4670 | 0.0153 | 0.0081 |
| cluster3596rr | -2315.478 | 0.0306 | 0.0288 | 0.9426 | 0.0627 | 0.0104 |
| cluster3597rr | -4193.9191 | 0.0278 | 0.0229 | 0.8246 | 0.0441 | 0.0187 |
| cluster3598rr | -4253.806 | 0.0240 | 0.1197 | 4.9793 | 0.0197 | 0.0054 |

| | | | | | | |
|---|---|---|---|---|---|---|
| cluster3599rr | -2248.9201 | 0.0205 | 0.0249 | 1.2179 | 0.1334 | 0.0137 |
| cluster3600rr | -2861.2813 | 0.0256 | 0.0182 | 0.7109 | 0.0213 | 0.0022 |
| cluster3601rr | -3284.7975 | 0.0057 | 0.0143 | 2.4960 | 0.0018 | 0.0005 |
| cluster3602rr | -3064.9906 | 0.0037 | 0.0017 | 0.4501 | 0.0001 | 0.0000 |
| cluster3603rr | -4217.4513 | 0.0454 | 0.0568 | 1.2504 | 0.2783 | 0.0188 |
| cluster3604rr | -1058.288 | 0.0638 | 0.0072 | 0.1125 | 0.0580 | 0.0035 |
| cluster3605rr | -3057.8793 | 0.0379 | 0.0406 | 1.0725 | 0.0398 | 0.0078 |
| cluster3606rr | -4321.6754 | 0.0250 | 0.0447 | 1.7861 | 0.0728 | 0.0100 |
| cluster3607rr | -2596.3813 | 0.0338 | 0.0481 | 1.4216 | 0.0734 | 0.0140 |
| cluster3608rr | -2888.184 | 0.0106 | 0.0075 | 0.7126 | 0.0120 | 0.0052 |
| cluster3609rr | -4178.0518 | 0.0245 | 0.0578 | 2.3566 | 0.0956 | 0.0206 |
| cluster3610rr | -1395.0984 | 0.0084 | 0.0271 | 3.2272 | 0.0001 | 0.0000 |
| cluster3611rr | -2340.8508 | 0.0268 | 0.0772 | 2.8786 | 0.0229 | 0.0101 |
| cluster3612rr | -2824.1915 | 0.0265 | 0.0627 | 2.3702 | 0.5284 | 0.0255 |
| cluster3613rr | -4058.5985 | 0.0195 | 0.0137 | 0.7028 | 0.0110 | 0.0046 |
| cluster3614rr | -4298.9181 | 0.0200 | 0.0685 | 3.4324 | 0.2750 | 0.0148 |
| cluster3615rr | -1662.5649 | 0.0513 | 0.0107 | 0.2078 | 0.0213 | 0.0036 |
| cluster3616rr | -774.82099 | 0.0143 | 0.0111 | 0.7761 | 0.0160 | 0.0035 |
| cluster3617rr | -1924.3897 | 0.0319 | 0.0461 | 1.4479 | 0.3086 | 0.0293 |
| cluster3618rr | -1899.354 | 0.1749 | 0.1184 | 0.6771 | 0.1541 | 0.0195 |
| cluster3619rr | -2552.2602 | 0.0047 | 0.0057 | 1.2006 | 0.0001 | 0.0000 |
| cluster3620rr | -3604.9378 | 0.0491 | 0.0424 | 0.8628 | 0.0689 | 0.0187 |
| cluster3621rr | -4762.2408 | 0.0297 | 0.0677 | 2.2791 | 0.0927 | 0.0139 |
| cluster3622rr | -2723.1845 | 0.0115 | 0.0066 | 0.5723 | 0.0396 | 0.0030 |
| cluster3623rr | -2271.7416 | 0.0090 | 0.0307 | 3.3961 | 0.0114 | 0.0014 |
| cluster3624rr | -2909.7934 | 0.0133 | 0.0319 | 2.3979 | 0.0220 | 0.0028 |
| cluster3625rr | -3308.3404 | 0.0058 | 0.0138 | 2.3964 | 0.0145 | 0.0026 |
| cluster3626rr | -1982.0228 | 0.0328 | 0.0206 | 0.6281 | 0.0149 | 0.0105 |
| cluster3627rr | -3001.0166 | 0.0074 | 0.0086 | 1.1659 | 0.0043 | 0.0009 |
| cluster3628rr | -3022.4061 | 0.0259 | 0.0575 | 2.2183 | 0.0570 | 0.0096 |
| cluster3629rr | -4158.4503 | 0.0095 | 0.0286 | 3.0184 | 0.0746 | 0.0133 |
| cluster3630rr | -1209.852 | 0.0158 | 0.0059 | 0.3731 | 0.0001 | 0.0000 |
| cluster3631rr | -3594.2194 | 0.0177 | 0.0240 | 1.3598 | 0.0086 | 0.0053 |
| cluster3632rr | -2459.1386 | 0.0277 | 0.0485 | 1.7518 | 0.0516 | 0.0339 |
| cluster3633rr | -2504.7034 | 0.0070 | 0.0152 | 2.1667 | 0.0200 | 0.0025 |
| cluster3634rr | -939.05231 | 0.0159 | 0.0085 | 0.5361 | 0.0001 | 0.0000 |
| cluster3635rr | -1621.07 | 0.0410 | 0.0078 | 0.1902 | 0.0533 | 0.0047 |
| cluster3636rr | -2531.2383 | 0.0162 | 0.0148 | 0.9178 | 0.0076 | 0.0076 |
| cluster3637rr | -1059.6254 | 0.0095 | 0.0030 | 0.3163 | 0.1500 | 0.0064 |
| cluster3638rr | -2731.8549 | 0.0296 | 0.0200 | 0.6762 | 0.0602 | 0.0076 |
| cluster3639rr | -1272.2297 | 0.0384 | 0.0195 | 0.5095 | 0.0217 | 0.0072 |
| cluster3640rr | -4764.8964 | 0.0674 | 0.1087 | 1.6141 | 0.1341 | 0.0252 |
| cluster3641rr | -1767.8038 | 0.0231 | 0.0065 | 0.2837 | 0.0647 | 0.0077 |
| cluster3642rr | -3355.558 | 0.0293 | 0.0506 | 1.7306 | 0.1915 | 0.0085 |

| | | | | | | |
|---|---|---|---|---|---|---|
| cluster3643rr | -3531.6989 | 0.0850 | 0.1081 | 1.2717 | 0.3981 | 0.0328 |
| cluster3644rr | -1913.1096 | 0.0288 | 0.0116 | 0.4042 | 0.0299 | 0.0046 |
| cluster3645rr | -1803.0318 | 0.0236 | 0.0346 | 1.4646 | 0.1109 | 0.0273 |
| cluster3646rr | -2628.7497 | 0.0118 | 0.0582 | 4.9512 | 0.1576 | 0.0217 |
| cluster3647rr | -3428.4801 | 0.0037 | 0.0191 | 5.1512 | 0.0494 | 0.0033 |
| cluster3648rr | -1759.1554 | 0.0015 | 0.0016 | 1.0451 | 0.0001 | 0.0001 |
| cluster3649rr | -1639.0359 | 0.0087 | 0.0067 | 0.7777 | 0.0680 | 0.0032 |
| cluster3650rr | -1253.7424 | 0.0075 | 0.0552 | 7.3428 | 999.0000 | 0.0142 |
| cluster3651rr | -3971.2282 | 0.0490 | 0.0401 | 0.8185 | 0.0402 | 0.0163 |
| cluster3652rr | -3310.2539 | 0.0266 | 0.0196 | 0.7341 | 0.0179 | 0.0118 |
| cluster3653rr | -1426.4184 | 0.0043 | 0.0027 | 0.6358 | 0.0001 | 0.0000 |
| cluster3654rr | -2837.9899 | 0.0096 | 0.0152 | 1.5780 | 0.0073 | 0.0015 |
| cluster3655rr | -2366.5159 | 0.0678 | 0.0676 | 0.9973 | 0.0493 | 0.0075 |
| cluster3656rr | -3108.3044 | 0.0269 | 0.0242 | 0.8991 | 0.0159 | 0.0066 |
| cluster3657rr | -3193.4457 | 0.0138 | 0.0377 | 2.7298 | 0.0318 | 0.0058 |
| cluster3658rr | -753.02419 | 0.0904 | 0.0258 | 0.2858 | 0.0001 | 0.0000 |
| cluster3659rr | -2269.803 | 0.0353 | 0.0218 | 0.6185 | 0.0390 | 0.0075 |
| cluster3660rr | -3016.7066 | 0.0166 | 0.0545 | 3.2873 | 0.1160 | 0.0058 |
| cluster3661rr | -1818.9705 | 0.0022 | 0.0003 | 0.1264 | 0.0001 | 0.0000 |
| cluster3662rr | -2791.9515 | 0.0270 | 0.0422 | 1.5624 | 4.0166 | 0.0224 |
| cluster3663rr | -3133.0792 | 0.0454 | 0.0448 | 0.9883 | 0.0932 | 0.0140 |
| cluster3664rr | -2449.0785 | 0.0375 | 0.0728 | 1.9418 | 0.0711 | 0.0113 |
| cluster3665rr | -3603.1509 | 0.0198 | 0.0342 | 1.7282 | 0.0301 | 0.0061 |
| cluster3666rr | -2710.5195 | 0.0327 | 0.0382 | 1.1700 | 0.0791 | 0.0074 |
| cluster3667rr | -1570.8087 | 0.0209 | 0.0074 | 0.3561 | 0.0263 | 0.0014 |
| cluster3668rr | -2978.6996 | 0.0171 | 0.0225 | 1.3186 | 0.0712 | 0.0151 |
| cluster3669rr | -2601.0088 | 0.0344 | 0.0287 | 0.8337 | 0.0939 | 0.0067 |
| cluster3670rr | -4034.8187 | 0.0472 | 0.0433 | 0.9180 | 0.0403 | 0.0253 |
| cluster3671rr | -3996.6669 | 0.0296 | 0.0459 | 1.5494 | 0.0327 | 0.0074 |
| cluster3672rr | -2495.6425 | 0.0372 | 0.0442 | 1.1903 | 0.1824 | 0.0188 |
| cluster3673rr | -4395.2122 | 0.0297 | 0.1242 | 4.1780 | 0.3335 | 0.0488 |
| cluster3674rr | -2025.1768 | 0.0261 | 0.0494 | 1.8953 | 0.0462 | 0.0068 |
| cluster3675rr | -660.30852 | 0.0001 | 0.0000 | 0.1625 | 0.0001 | 0.0000 |
| cluster3676rr | -3512.4189 | 0.0407 | 0.0600 | 1.4743 | 3.4687 | 0.0265 |
| cluster3677rr | -811.95238 | 0.0242 | 0.0128 | 0.5269 | 0.0136 | 0.0038 |
| cluster3678rr | -3306.5509 | 0.0321 | 0.0338 | 1.0539 | 0.0631 | 0.0141 |
| cluster3679rr | -944.13356 | 0.0091 | 0.0041 | 0.4472 | 0.0001 | 0.0000 |
| cluster3680rr | -2314.0834 | 0.0088 | 0.0250 | 2.8428 | 0.0178 | 0.0046 |
| cluster3681rr | -2959.2499 | 0.0195 | 0.0381 | 1.9582 | 0.0212 | 0.0053 |
| cluster3682rr | -4337.6238 | 0.0241 | 0.0517 | 2.1423 | 0.7821 | 0.0342 |
| cluster3683rr | -2805.2152 | 0.1078 | 0.1102 | 1.0219 | 0.0879 | 0.0551 |
| cluster3684rr | -2988.5857 | 0.0292 | 0.0383 | 1.3115 | 0.1274 | 0.0190 |
| cluster3685rr | -3672.3245 | 0.0756 | 0.1236 | 1.6356 | 0.2335 | 0.0212 |
| cluster3686rr | -1628.4302 | 0.0371 | 0.0969 | 2.6082 | 0.0001 | 0.0000 |

| | | | | | | |
|---|---|---|---|---|---|---|
| cluster3687rr | -2869.8673 | 0.0638 | 0.1613 | 2.5299 | 1.1428 | 0.0231 |
| cluster3688rr | -2042.1982 | 0.0041 | 0.0155 | 3.7349 | 0.0631 | 0.0130 |
| cluster3689rr | -2638.5389 | 0.0445 | 0.0268 | 0.6031 | 0.0033 | 0.0009 |
| cluster3690rr | -1453.7117 | 0.0271 | 0.0201 | 0.7443 | 0.0086 | 0.0056 |
| cluster3691rr | -669.76828 | 0.0029 | 0.0031 | 1.0597 | 0.0255 | 0.0040 |
| cluster3692rr | -1435.8748 | 0.0112 | 0.0162 | 1.4497 | 0.0001 | 0.0000 |
| cluster3693rr | -2463.4347 | 0.0079 | 0.0215 | 2.7208 | 0.0380 | 0.0064 |
| cluster3694rr | -2090.3352 | 0.0334 | 0.0124 | 0.3700 | 0.0025 | 0.0004 |
| cluster3695rr | -1734.5621 | 0.0216 | 0.0264 | 1.2208 | 0.1129 | 0.0101 |
| cluster3696rr | -817.41977 | 0.0122 | 0.0385 | 3.1602 | 0.0001 | 0.0000 |
| cluster3697rr | -3583.7306 | 0.0758 | 0.0448 | 0.5907 | 0.0559 | 0.0244 |
| cluster3698rr | -1099.5724 | 0.0001 | 0.0000 | 0.3097 | 0.0001 | 0.0000 |
| cluster3699rr | -4429.2867 | 0.0374 | 0.0746 | 1.9960 | 0.6057 | 0.0260 |
| cluster3700rr | -2819.0892 | 0.0324 | 0.0367 | 1.1325 | 0.0307 | 0.0060 |
| cluster3701rr | -1128.4346 | 0.0013 | 0.0005 | 0.4068 | 0.0001 | 0.0000 |
| cluster3702rr | -3860.3828 | 0.0208 | 0.0591 | 2.8412 | 0.5334 | 0.0097 |
| cluster3703rr | -1512.7 | 0.0168 | 0.0068 | 0.4042 | 0.0151 | 0.0022 |
| cluster3704rr | -2227.7294 | 0.0077 | 0.0139 | 1.8047 | 0.0010 | 0.0014 |
| cluster3705rr | -1944.3763 | 0.0150 | 0.0097 | 0.6496 | 0.0045 | 0.0020 |
| cluster3706rr | -2093.9412 | 0.0196 | 0.0151 | 0.7700 | 0.0524 | 0.0042 |
| cluster3707rr | -1919.6593 | 0.0355 | 0.0688 | 1.9381 | 0.1695 | 0.0192 |
| cluster3708rr | -2636.4306 | 0.0458 | 0.0419 | 0.9163 | 0.0592 | 0.0081 |
| cluster3709rr | -1579.7113 | 0.0816 | 0.0246 | 0.3017 | 0.0001 | 0.0000 |
| cluster3710rr | -1924.8149 | 0.0142 | 0.0287 | 2.0265 | 0.0193 | 0.0047 |
| cluster3711rr | -1438.6941 | 0.0266 | 0.0076 | 0.2838 | 0.1211 | 0.0030 |
| cluster3712rr | -3402.6326 | 0.0276 | 0.0191 | 0.6918 | 0.0247 | 0.0109 |
| cluster3713rr | -4020.1349 | 0.0388 | 0.0853 | 2.1957 | 0.2096 | 0.0341 |
| cluster3714rr | -1773.411 | 0.0389 | 0.0157 | 0.4051 | 0.1086 | 0.0239 |
| cluster3715rr | -1283.7761 | 0.0115 | 0.0483 | 4.2108 | 999.0000 | 0.0119 |
| cluster3716rr | -3658.0746 | 0.0456 | 0.0663 | 1.4549 | 0.0593 | 0.0107 |
| cluster3717rr | -1401.7415 | 0.0096 | 0.0027 | 0.2827 | 0.0470 | 0.0080 |
| cluster3718rr | -2989.4349 | 0.0190 | 0.0376 | 1.9840 | 0.0539 | 0.0076 |
| cluster3719rr | -3291.0604 | 0.0385 | 0.0381 | 0.9910 | 0.2164 | 0.0133 |
| cluster3720rr | -2757.6688 | 0.0243 | 0.0213 | 0.8759 | 0.0692 | 0.0063 |
| cluster3721rr | -1097.8138 | 0.0057 | 0.0009 | 0.1647 | 0.0001 | 0.0000 |
| cluster3722rr | -1224.2785 | 0.0420 | 0.1138 | 2.7076 | 0.0163 | 0.0037 |
| cluster3723rr | -989.94266 | 0.0001 | 0.0000 | 0.4734 | 0.0001 | 0.0000 |
| cluster3724rr | -3076.2934 | 0.0272 | 0.0424 | 1.5614 | 0.1541 | 0.0208 |
| cluster3725rr | -1839.9498 | 0.0104 | 0.0192 | 1.8465 | 0.0102 | 0.0025 |
| cluster3726rr | -1936.3798 | 0.0502 | 0.0335 | 0.6688 | 0.0253 | 0.0024 |
| cluster3727rr | -2464.5655 | 0.0451 | 0.0815 | 1.8050 | 0.2151 | 0.0199 |
| cluster3728rr | -2157.0851 | 0.0212 | 0.0152 | 0.7169 | 0.0201 | 0.0090 |
| cluster3729rr | -2947.1276 | 0.0128 | 0.0217 | 1.7050 | 0.0201 | 0.0038 |
| cluster3730rr | -1781.561 | 0.0769 | 0.0225 | 0.2932 | 0.1272 | 0.0190 |

| | | | | | | |
|---|---|---|---|---|---|---|
| cluster3731rr | -2864.8692 | 0.0082 | 0.0057 | 0.6927 | 0.0163 | 0.0050 |
| cluster3732rr | -2822.6833 | 0.0218 | 0.0207 | 0.9461 | 0.0371 | 0.0055 |
| cluster3733rr | -3072.861 | 0.0800 | 0.0376 | 0.4695 | 0.0782 | 0.0204 |
| cluster3734rr | -3041.7549 | 0.0394 | 0.0480 | 1.2180 | 0.0882 | 0.0198 |
| cluster3735rr | -3245.3084 | 0.0690 | 0.0362 | 0.5253 | 0.2008 | 0.0428 |
| cluster3736rr | -3537.6093 | 0.0439 | 0.0305 | 0.6957 | 0.0738 | 0.0113 |
| cluster3737rr | -1209.4004 | 0.0033 | 0.0010 | 0.3058 | 0.0001 | 0.0000 |
| cluster3738rr | -3257.6245 | 0.0102 | 0.0084 | 0.8260 | 0.0018 | 0.0010 |
| cluster3739rr | -1156.2535 | 0.0001 | 0.0000 | 0.3065 | 0.0001 | 0.0000 |
| cluster3740rr | -3448.0006 | 0.0267 | 0.0363 | 1.3593 | 0.0790 | 0.0132 |
| cluster3741rr | -1312.358 | 0.0149 | 0.0073 | 0.4880 | 0.0319 | 0.0058 |
| cluster3742rr | -1615.744 | 0.0548 | 0.0558 | 1.0190 | 0.0716 | 0.0164 |
| cluster3743rr | -2979.3391 | 0.0224 | 0.0311 | 1.3876 | 0.0625 | 0.0058 |
| cluster3744rr | -1354.3334 | 0.0938 | 0.0570 | 0.6076 | 0.4670 | 0.0136 |
| cluster3745rr | -2582.6625 | 0.0219 | 0.0337 | 1.5405 | 0.1050 | 0.0163 |
| cluster3746rr | -3783.4819 | 0.0335 | 0.0344 | 1.0289 | 0.0782 | 0.0403 |
| cluster3747rr | -1120.342 | 0.0286 | 0.0234 | 0.8186 | 0.0001 | 0.0000 |
| cluster3748rr | -2164.2734 | 0.0165 | 0.0089 | 0.5404 | 0.0001 | 0.0000 |
| cluster3749rr | -3060.9694 | 0.0119 | 0.0227 | 1.9042 | 0.1419 | 0.0152 |
| cluster3750rr | -1348.7576 | 0.0179 | 0.0500 | 2.7931 | 999.0000 | 0.0231 |
| cluster3751rr | -3010.6071 | 0.0041 | 0.0047 | 1.1505 | 0.0148 | 0.0029 |
| cluster3752rr | -2660.7339 | 0.0068 | 0.0151 | 2.2336 | 0.0619 | 0.0047 |
| cluster3753rr | -1021.7275 | 0.0277 | 0.0027 | 0.0964 | 0.0001 | 0.0000 |
| cluster3754rr | -881.97724 | 0.0001 | 0.0000 | 0.1854 | 0.0001 | 0.0000 |
| cluster3755rr | -2499.5521 | 0.0168 | 0.0144 | 0.8573 | 0.0101 | 0.0060 |
| cluster3756rr | -1689.5556 | 0.0375 | 0.0363 | 0.9673 | 0.0235 | 0.0068 |
| cluster3757rr | -2090.0516 | 0.0419 | 0.0224 | 0.5345 | 0.1298 | 0.0084 |
| cluster3758rr | -1509.5386 | 0.0340 | 0.0188 | 0.5529 | 0.0001 | 0.0000 |
| cluster3759rr | -3065.3892 | 0.0735 | 0.0437 | 0.5941 | 0.0422 | 0.0174 |
| cluster3760rr | -2599.4112 | 0.0269 | 0.0269 | 1.0005 | 0.0113 | 0.0100 |
| cluster3761rr | -767.58093 | 0.0035 | 0.0284 | 8.0284 | 0.0188 | 0.0038 |
| cluster3762rr | -1966.3731 | 0.0183 | 0.0061 | 0.3330 | 0.0001 | 0.0000 |
| cluster3763rr | -2005.5162 | 0.0570 | 0.0168 | 0.2948 | 0.0455 | 0.0065 |
| cluster3764rr | -1629.7722 | 0.0294 | 0.0524 | 1.7789 | 0.0457 | 0.0035 |
| cluster3765rr | -2667.1913 | 0.0227 | 0.0185 | 0.8146 | 0.0027 | 0.0013 |
| cluster3766rr | -1795.3255 | 0.0107 | 0.0143 | 1.3371 | 0.0212 | 0.0035 |
| cluster3767rr | -859.9419 | 0.0064 | 0.0354 | 5.5180 | 727.7810 | 0.0219 |
| cluster3768rr | -2216.9206 | 0.0279 | 0.0705 | 2.5264 | 999.0000 | 0.0082 |
| cluster3769rr | -1764.8087 | 0.0077 | 0.0026 | 0.3348 | 0.0001 | 0.0000 |
| cluster3770rr | -1771.3201 | 0.0419 | 0.0418 | 0.9964 | 0.2051 | 0.0269 |
| cluster3771rr | -2138.4346 | 0.0761 | 0.0745 | 0.9789 | 0.0430 | 0.0122 |
| cluster3772rr | -725.30121 | 0.0009 | 0.0220 | 23.9983 | 0.0001 | 0.0000 |
| cluster3773rr | -1116.9818 | 0.2132 | 0.0446 | 0.2092 | 0.0458 | 0.0027 |
| cluster3774rr | -1409.0234 | 0.0127 | 0.0030 | 0.2387 | 0.0240 | 0.0017 |

| | | | | | | |
|---|---|---|---|---|---|---|
| cluster3775rr | -906.40414 | 0.0135 | 0.0252 | 1.8654 | 0.0001 | 0.0000 |
| cluster3776rr | -2517.1438 | 0.0860 | 0.0334 | 0.3886 | 0.0404 | 0.0165 |
| cluster3777rr | -3066.5066 | 0.0543 | 0.0709 | 1.3047 | 0.0929 | 0.0282 |
| cluster3778rr | -980.5449 | 0.0014 | 0.0012 | 0.8722 | 0.0001 | 0.0000 |
| cluster3779rr | -2995.1116 | 0.0229 | 0.0351 | 1.5337 | 0.0194 | 0.0161 |
| cluster3780rr | -1683.6556 | 0.0099 | 0.0251 | 2.5463 | 0.0530 | 0.0099 |
| cluster3781rr | -2686.1781 | 0.0550 | 0.0359 | 0.6530 | 0.0697 | 0.0362 |
| cluster3782rr | -2289.8738 | 0.0401 | 0.0283 | 0.7058 | 0.0652 | 0.0170 |
| cluster3783rr | -2106.9326 | 0.0063 | 0.0340 | 5.3772 | 0.0242 | 0.0053 |
| cluster3784rr | -2948.9697 | 0.0557 | 0.0502 | 0.9010 | 0.0226 | 0.0225 |
| cluster3785rr | -2688.2734 | 0.0153 | 0.0113 | 0.7402 | 0.0475 | 0.0043 |
| cluster3786rr | -2267.0149 | 0.0086 | 0.0287 | 3.3468 | 0.0086 | 0.0079 |
| cluster3787rr | -1521.1444 | 0.0240 | 0.0221 | 0.9218 | 0.0617 | 0.0146 |
| cluster3788rr | -1160.4015 | 0.0001 | 0.0000 | 0.0984 | 0.0001 | 0.0000 |
| cluster3789rr | -3040.9159 | 0.0116 | 0.0060 | 0.5194 | 0.0074 | 0.0019 |
| cluster3790rr | -2174.775 | 0.0108 | 0.0302 | 2.8010 | 0.0511 | 0.0100 |
| cluster3791rr | -418.9737 | 0.0076 | 0.0030 | 0.3974 | 0.0449 | 0.0119 |
| cluster3792rr | -1862.9345 | 0.0054 | 0.0030 | 0.5701 | 0.0128 | 0.0027 |
| cluster3793rr | -1852.0726 | 0.0240 | 0.0213 | 0.8881 | 0.0713 | 0.0122 |
| cluster3794rr | -2186.2926 | 0.0139 | 0.0220 | 1.5813 | 0.2043 | 0.0059 |
| cluster3795rr | -1701.6593 | 0.0194 | 0.0214 | 1.1011 | 0.0210 | 0.0020 |
| cluster3796rr | -1246.73 | 0.1005 | 0.0791 | 0.7871 | 1.3046 | 0.0420 |
| cluster3797rr | -3907.9104 | 0.0461 | 0.1398 | 3.0345 | 0.1754 | 0.0244 |
| cluster3798rr | -3441.8062 | 0.0199 | 0.0289 | 1.4498 | 0.0884 | 0.0175 |
| cluster3799rr | -2339.1553 | 0.0394 | 0.0331 | 0.8406 | 0.1115 | 0.0307 |
| cluster3800rr | -430.57052 | 0.0254 | 0.0011 | 0.0434 | 0.0001 | 0.0000 |
| cluster3801rr | -2690.1883 | 0.1989 | 0.1015 | 0.5102 | 0.3367 | 0.0160 |
| cluster3802rr | -2649.715 | 0.0356 | 0.0453 | 1.2732 | 0.0267 | 0.0273 |
| cluster3803rr | -2238.7534 | 0.0326 | 0.0308 | 0.9437 | 0.1304 | 0.0085 |
| cluster3804rr | -2634.9625 | 0.0304 | 0.0316 | 1.0394 | 0.0613 | 0.0070 |
| cluster3805rr | -2654.6865 | 0.0213 | 0.0296 | 1.3914 | 0.0522 | 0.0107 |
| cluster3806rr | -3565.1302 | 0.0359 | 0.0485 | 1.3502 | 0.1324 | 0.0108 |
| cluster3807rr | -1440.9893 | 0.0059 | 0.0021 | 0.3514 | 0.0226 | 0.0037 |
| cluster3808rr | -1825.427 | 0.0305 | 0.0440 | 1.4447 | 0.0672 | 0.0094 |
| cluster3809rr | -2360.1046 | 0.0100 | 0.0099 | 0.9874 | 0.0001 | 0.0000 |
| cluster3810rr | -2463.5508 | 0.0082 | 0.0091 | 1.1050 | 0.0167 | 0.0129 |
| cluster3811rr | -761.30254 | 0.0218 | 0.0048 | 0.2229 | 0.0945 | 0.0029 |
| cluster3812rr | -3000.7218 | 0.0295 | 0.0235 | 0.7979 | 0.0400 | 0.0048 |
| cluster3813rr | -3411.2984 | 0.0303 | 0.0916 | 3.0212 | 0.0986 | 0.0207 |
| cluster3814rr | -1609.0289 | 0.0895 | 0.0178 | 0.1986 | 0.0861 | 0.0178 |
| cluster3815rr | -1706.8057 | 0.0162 | 0.0394 | 2.4328 | 0.1367 | 0.0096 |
| cluster3816rr | -2521.629 | 0.0212 | 0.0470 | 2.2127 | 0.3850 | 0.0128 |
| cluster3817rr | -991.40391 | 0.0050 | 0.0018 | 0.3514 | 0.0254 | 0.0025 |
| cluster3818rr | -1978.8595 | 0.0660 | 0.0680 | 1.0301 | 0.0493 | 0.0571 |

| | | | | | | |
|---|---|---|---|---|---|---|
| cluster3819rr | -3343.6023 | 0.0235 | 0.0295 | 1.2570 | 0.0975 | 0.0131 |
| cluster3820rr | -2224.426 | 0.0273 | 0.0296 | 1.0844 | 0.0731 | 0.0098 |
| cluster3821rr | -2243.5257 | 0.0171 | 0.0134 | 0.7845 | 0.0114 | 0.0064 |
| cluster3822rr | -3396.5485 | 0.0241 | 0.0429 | 1.7799 | 0.0925 | 0.0157 |
| cluster3823rr | -3408.781 | 0.0193 | 0.0268 | 1.3908 | 0.0231 | 0.0236 |
| cluster3824rr | -2336.6844 | 0.0150 | 0.0399 | 2.6670 | 0.3443 | 0.0356 |
| cluster3825rr | -1388.7941 | 0.0062 | 0.0056 | 0.9096 | 0.0108 | 0.0022 |
| cluster3826rr | -3154.9609 | 0.0708 | 0.1033 | 1.4587 | 0.0980 | 0.0185 |
| cluster3827rr | -2403.264 | 0.0304 | 0.0263 | 0.8642 | 0.0197 | 0.0202 |
| cluster3828rr | -2361.1859 | 0.0506 | 0.0464 | 0.9165 | 0.0001 | 0.0000 |
| cluster3829rr | -2250.5849 | 0.0191 | 0.0302 | 1.5801 | 0.1332 | 0.0194 |
| cluster3830rr | -1627.8396 | 0.0183 | 0.0078 | 0.4264 | 0.1471 | 0.0031 |
| cluster3831rr | -2348.3867 | 0.0304 | 0.0575 | 1.8953 | 0.0458 | 0.0070 |
| cluster3832rr | -1767.9935 | 0.0048 | 0.0040 | 0.8297 | 0.0001 | 0.0000 |
| cluster3833rr | -2130.1074 | 0.0221 | 0.0253 | 1.1439 | 0.0469 | 0.0056 |
| cluster3834rr | -2054.4449 | 0.0046 | 0.0237 | 5.1196 | 0.0532 | 0.0107 |
| cluster3835rr | -2118.0356 | 0.0333 | 0.0319 | 0.9577 | 0.0670 | 0.0047 |
| cluster3836rr | -1894.321 | 0.0161 | 0.0167 | 1.0371 | 0.0452 | 0.0051 |
| cluster3837rr | -2639.6473 | 0.0416 | 0.0432 | 1.0383 | 0.0365 | 0.0105 |
| cluster3838rr | -2277.9261 | 0.0182 | 0.0205 | 1.1307 | 999.0000 | 0.0494 |
| cluster3839rr | -1481.1629 | 0.0105 | 0.0189 | 1.8006 | 999.0000 | 0.0279 |
| cluster3840rr | -2024.4672 | 0.0102 | 0.0799 | 7.8396 | 0.0001 | 0.0000 |
| cluster3841rr | -2290.2468 | 0.0651 | 0.1074 | 1.6509 | 0.2017 | 0.0298 |
| cluster3842rr | -1604.0559 | 0.0013 | 0.0003 | 0.2578 | 0.0267 | 0.0031 |
| cluster3843rr | -1440.972 | 0.0120 | 0.0082 | 0.6867 | 0.0001 | 0.0000 |
| cluster3844rr | -1078.2424 | 0.0624 | 0.0393 | 0.6309 | 0.1725 | 0.0055 |
| cluster3845rr | -604.91636 | 0.0001 | 0.0000 | 0.1483 | 0.0001 | 0.0000 |
| cluster3846rr | -1273.1932 | 0.0195 | 0.0141 | 0.7211 | 0.0061 | 0.0025 |
| cluster3847rr | -1156.4097 | 0.0011 | 0.0018 | 1.6738 | 0.0001 | 0.0000 |
| cluster3848rr | -2515.2769 | 0.0277 | 0.0450 | 1.6253 | 0.1247 | 0.0162 |
| cluster3849rr | -2655.9956 | 0.0083 | 0.0602 | 7.2741 | 999.0000 | 0.0106 |
| cluster3850rr | -1677.0783 | 0.0237 | 0.0164 | 0.6925 | 0.0054 | 0.0012 |
| cluster3851rr | -1617.8573 | 0.0090 | 0.0112 | 1.2514 | 0.0186 | 0.0038 |
| cluster3852rr | -1856.8249 | 0.0068 | 0.0126 | 1.8589 | 0.0059 | 0.0046 |
| cluster3853rr | -1101.3636 | 0.0052 | 0.0125 | 2.4216 | 0.0001 | 0.0000 |
| cluster3854rr | -1356.8615 | 0.0854 | 0.0539 | 0.6318 | 0.2562 | 0.0230 |
| cluster3855rr | -2920.0033 | 0.0072 | 0.0340 | 4.7573 | 36.2927 | 0.0111 |
| cluster3856rr | -1693.4551 | 0.0150 | 0.0232 | 1.5518 | 0.0748 | 0.0144 |
| cluster3857rr | -3035.9144 | 0.0148 | 0.0221 | 1.4939 | 0.0473 | 0.0092 |
| cluster3858rr | -2416.7313 | 0.0489 | 0.0354 | 0.7243 | 0.0479 | 0.0419 |
| cluster3859rr | -1977.604 | 0.0133 | 0.0123 | 0.9281 | 0.0111 | 0.0022 |
| cluster3860rr | -2623.5891 | 0.0905 | 0.0416 | 0.4595 | 0.0417 | 0.0173 |
| cluster3861rr | -1299.8898 | 0.0152 | 0.0196 | 1.2932 | 0.0456 | 0.0092 |
| cluster3862rr | -922.33901 | 0.0001 | 0.0001 | 0.7844 | 0.0001 | 0.0000 |

| | | | | | | |
|---|---|---|---|---|---|---|
| cluster3863rr | -1483.0515 | 0.0013 | 0.0009 | 0.6947 | 0.0001 | 0.0001 |
| cluster3864rr | -2220.0539 | 0.0041 | 0.0092 | 2.2522 | 0.0007 | 0.0001 |
| cluster3865rr | -3001.9098 | 0.0404 | 0.0718 | 1.7786 | 0.1251 | 0.0199 |
| cluster3866rr | -1633.013 | 0.0196 | 0.0352 | 1.7967 | 0.0001 | 0.0000 |
| cluster3867rr | -608.47114 | 0.0001 | 0.0000 | 0.0675 | 0.0001 | 0.0000 |
| cluster3868rr | -1393.3957 | 0.0289 | 0.0158 | 0.5484 | 0.0001 | 0.0000 |
| cluster3869rr | -2963.3286 | 0.0190 | 0.0854 | 4.4997 | 0.9414 | 0.0265 |
| cluster3870rr | -2459.7348 | 0.0343 | 0.0376 | 1.0978 | 0.0947 | 0.0246 |
| cluster3871rr | -2152.7171 | 0.0073 | 0.0089 | 1.2214 | 0.0129 | 0.0021 |
| cluster3872rr | -2649.6049 | 0.0123 | 0.0278 | 2.2632 | 0.0087 | 0.0016 |
| cluster3873rr | -1788.1612 | 0.0288 | 0.0216 | 0.7517 | 0.0289 | 0.0114 |
| cluster3874rr | -2100.8141 | 0.0146 | 0.0047 | 0.3224 | 0.0001 | 0.0000 |
| cluster3875rr | -770.40263 | 0.0030 | 0.0018 | 0.6012 | 0.0001 | 0.0000 |
| cluster3876rr | -745.22562 | 0.0125 | 0.0058 | 0.4654 | 0.0001 | 0.0000 |
| cluster3877rr | -1435.9876 | 0.0147 | 0.0073 | 0.4964 | 0.0087 | 0.0019 |
| cluster3879rr | -1880.9633 | 0.0167 | 0.0171 | 1.0244 | 0.0371 | 0.0055 |
| cluster3880rr | -1356.0145 | 0.0229 | 0.0288 | 1.2579 | 0.0001 | 0.0000 |
| cluster3881rr | -3015.7212 | 0.0398 | 0.0833 | 2.0914 | 2.2075 | 0.0129 |
| cluster3882rr | -2687.4518 | 0.0211 | 0.0304 | 1.4402 | 0.0859 | 0.0140 |
| cluster3883rr | -423.83591 | 0.0051 | 0.1101 | 21.4267 | 999.0000 | 0.0412 |
| cluster3884rr | -1560.5414 | 0.0004 | 0.0151 | 39.7173 | 677.3629 | 0.0051 |
| cluster3885rr | -2849.5951 | 0.0092 | 0.0140 | 1.5290 | 0.0646 | 0.0166 |
| cluster3886rr | -1394.964 | 0.0034 | 0.0139 | 4.1188 | 0.0001 | 0.0000 |
| cluster3887rr | -1627.5269 | 0.0065 | 0.0307 | 4.7302 | 0.0229 | 0.0064 |
| cluster3888rr | -1182.098 | 0.0059 | 0.0456 | 7.7205 | 999.0000 | 0.0085 |
| cluster3889rr | -1004.6717 | 0.0235 | 0.0432 | 1.8406 | 0.0271 | 0.0081 |
| cluster3890rr | -1871.7003 | 0.0163 | 0.0280 | 1.7160 | 0.0261 | 0.0164 |
| cluster3891rr | -2183.2967 | 0.0163 | 0.0137 | 0.8410 | 0.0044 | 0.0027 |
| cluster3892rr | -1372.8089 | 0.0164 | 0.0084 | 0.5130 | 0.0001 | 0.0000 |
| cluster3893rr | -1458.8435 | 0.0261 | 0.0160 | 0.6144 | 0.0189 | 0.0060 |
| cluster3894rr | -966.3824 | 0.0069 | 0.0194 | 2.8243 | 0.0160 | 0.0031 |
| cluster3895rr | -1980.6266 | 0.0073 | 0.0155 | 2.1203 | 0.0419 | 0.0057 |
| cluster3896rr | -820.61353 | 0.0033 | 0.0059 | 1.8171 | 0.0001 | 0.0000 |
| cluster3897rr | -1451.0637 | 0.0076 | 0.0570 | 7.4937 | 0.0001 | 0.0001 |
| cluster3898rr | -2390.3802 | 0.0252 | 0.0084 | 0.3327 | 0.0203 | 0.0037 |
| cluster3899rr | -2997.1837 | 0.0201 | 0.0540 | 2.6856 | 0.1804 | 0.0226 |
| cluster3900rr | -2358.9101 | 0.0434 | 0.1245 | 2.8706 | 0.7350 | 0.0599 |
| cluster3901rr | -1540.6861 | 0.0015 | 0.0026 | 1.7337 | 0.0409 | 0.0041 |
| cluster3902rr | -2495.0388 | 0.0104 | 0.0195 | 1.8738 | 0.0463 | 0.0066 |
| cluster3903rr | -1623.1553 | 0.0093 | 0.0366 | 3.9443 | 0.0230 | 0.0047 |
| cluster3904rr | -905.11743 | 0.0056 | 0.0180 | 3.2285 | 0.0001 | 0.0000 |
| cluster3905rr | -1590.1704 | 0.0979 | 0.0856 | 0.8744 | 0.5806 | 0.0549 |
| cluster3906rr | -1453.394 | 0.0224 | 0.0189 | 0.8450 | 0.0100 | 0.0021 |
| cluster3907rr | -1944.3392 | 0.0382 | 0.0363 | 0.9511 | 0.0755 | 0.0144 |

| | | | | | | |
|---|---|---|---|---|---|---|
| cluster3908rr | -1848.1275 | 0.0137 | 0.0185 | 1.3550 | 0.0194 | 0.0035 |
| cluster3909rr | -2462.7405 | 0.0201 | 0.0468 | 2.3335 | 0.0849 | 0.0132 |
| cluster3910rr | -1192.1268 | 0.0299 | 0.0545 | 1.8238 | 0.1075 | 0.0195 |
| cluster3911rr | -1741.1914 | 0.0039 | 0.0126 | 3.2704 | 0.0686 | 0.0134 |
| cluster3912rr | -951.76626 | 0.0001 | 0.0000 | 0.3167 | 0.0001 | 0.0000 |
| cluster3913rr | -2544.9975 | 0.0387 | 0.0517 | 1.3384 | 0.0372 | 0.0047 |
| cluster3914rr | -2238.9468 | 0.0520 | 0.0921 | 1.7711 | 0.0907 | 0.0112 |
| cluster3915rr | -2226.9727 | 0.0322 | 0.0442 | 1.3744 | 0.0601 | 0.0122 |
| cluster3916rr | -902.19813 | 0.0728 | 0.0598 | 0.8218 | 0.1782 | 0.0178 |
| cluster3917rr | -2583.1911 | 0.0126 | 0.0199 | 1.5877 | 0.0118 | 0.0015 |
| cluster3918rr | -2747.6243 | 0.0112 | 0.0606 | 5.4034 | 0.2186 | 0.0206 |
| cluster3919rr | -569.99271 | 0.0021 | 0.0008 | 0.3823 | 0.0001 | 0.0000 |
| cluster3920rr | -657.51675 | 0.0089 | 0.1184 | 13.2599 | 999.0000 | 0.0056 |
| cluster3921rr | -2190.091 | 0.0235 | 0.0286 | 1.2189 | 0.0441 | 0.0116 |
| cluster3922rr | -1237.2135 | 0.0395 | 0.0421 | 1.0680 | 3.5386 | 0.0172 |
| cluster3923rr | -1152.1647 | 0.0004 | 0.0011 | 3.1883 | 0.2843 | 0.0000 |
| cluster3924rr | -1692.7756 | 0.0169 | 0.0231 | 1.3635 | 0.1717 | 0.0082 |
| cluster3925rr | -1089.3362 | 0.0363 | 0.0374 | 1.0283 | 0.2908 | 0.0311 |
| cluster3926rr | -2437.4262 | 0.0463 | 0.1009 | 2.1812 | 0.0410 | 0.0118 |
| cluster3927rr | -1188.1436 | 0.0104 | 0.0156 | 1.5035 | 0.0153 | 0.0026 |
| cluster3928rr | -1159.8934 | 0.0299 | 0.0172 | 0.5775 | 0.0192 | 0.0020 |
| cluster3929rr | -1606.7176 | 0.0055 | 0.0105 | 1.9153 | 999.0000 | 0.0019 |
| cluster3930rr | -2015.4593 | 0.0174 | 0.0105 | 0.6040 | 0.0001 | 0.0000 |
| cluster3931rr | -1826.4985 | 0.1514 | 0.0429 | 0.2833 | 0.0001 | 0.0000 |
| cluster3932rr | -1117.7387 | 0.0113 | 0.0034 | 0.3041 | 0.0001 | 0.0000 |
| cluster3933rr | -1158.4283 | 0.0848 | 0.0358 | 0.4224 | 0.1174 | 0.0055 |
| cluster3934rr | -717.10761 | 0.0001 | 0.0001 | 0.5840 | 0.0001 | 0.0000 |
| cluster3935rr | -3094.8269 | 0.0391 | 0.0889 | 2.2708 | 0.4620 | 0.0494 |
| cluster3936rr | -2124.7263 | 0.0123 | 0.0259 | 2.1141 | 0.1635 | 0.0230 |
| cluster3937rr | -1938.2741 | 0.0222 | 0.0132 | 0.5931 | 0.0093 | 0.0043 |
| cluster3938rr | -1780.4203 | 0.0283 | 0.0213 | 0.7528 | 0.1057 | 0.0105 |
| cluster3939rr | -2054.8706 | 0.0144 | 0.0206 | 1.4293 | 0.0482 | 0.0050 |
| cluster3940rr | -1337.0052 | 0.0199 | 0.0092 | 0.4639 | 0.0001 | 0.0000 |
| cluster3941rr | -970.1816 | 0.0297 | 0.0109 | 0.3686 | 0.0354 | 0.0025 |
| cluster3942rr | -2009.0501 | 0.0297 | 0.0205 | 0.6899 | 0.0744 | 0.0049 |
| cluster3943rr | -2265.093 | 0.0119 | 0.0388 | 3.2694 | 9.2442 | 0.0095 |
| cluster3944rr | -1593.1403 | 0.0436 | 0.0497 | 1.1416 | 0.1031 | 0.0073 |
| cluster3945rr | -1675.8262 | 0.0354 | 0.0487 | 1.3751 | 0.1982 | 0.0066 |
| cluster3946rr | -1796.2319 | 0.0190 | 0.0289 | 1.5191 | 0.3879 | 0.0161 |
| cluster3947rr | -2285.7898 | 0.0271 | 0.0413 | 1.5234 | 0.0432 | 0.0101 |
| cluster3948rr | -1015.8119 | 0.0294 | 0.0203 | 0.6896 | 0.0453 | 0.0063 |
| cluster3949rr | -878.13103 | 0.0097 | 0.0230 | 2.3846 | 0.0001 | 0.0000 |
| cluster3950rr | -1741.042 | 0.0134 | 0.0267 | 1.9995 | 0.0080 | 0.0102 |
| cluster3951rr | -2833.0397 | 0.0365 | 0.0509 | 1.3952 | 0.0926 | 0.0143 |

| | | | | | | |
|---|---|---|---|---|---|---|
| cluster3952rr | -1474.8213 | 0.0029 | 0.0065 | 2.2149 | 0.0074 | 0.0016 |
| cluster3953rr | -1361.5093 | 0.0043 | 0.0127 | 2.9240 | 0.0302 | 0.0038 |
| cluster3954rr | -1805.3143 | 0.0255 | 0.0228 | 0.8948 | 0.0801 | 0.0095 |
| cluster3955rr | -1191.4777 | 0.0224 | 0.0375 | 1.6700 | 0.0636 | 0.0091 |
| cluster3956rr | -1995.3289 | 0.0137 | 0.0267 | 1.9585 | 0.0325 | 0.0044 |
| cluster3957rr | -1957.0783 | 0.0097 | 0.0217 | 2.2326 | 0.0446 | 0.0112 |
| cluster3958rr | -1103.3529 | 0.0263 | 0.0276 | 1.0489 | 0.0001 | 0.0000 |
| cluster3959rr | -2685.8019 | 0.0359 | 0.0233 | 0.6491 | 0.0251 | 0.0174 |
| cluster3960rr | -2424.5365 | 0.0569 | 0.0743 | 1.3070 | 0.0703 | 0.0055 |
| cluster3961rr | -1594.9149 | 0.0371 | 0.0482 | 1.3012 | 0.0516 | 0.0057 |
| cluster3962rr | -1533.2058 | 0.0429 | 0.0433 | 1.0092 | 0.0260 | 0.0042 |
| cluster3963rr | -2070.7503 | 0.0305 | 0.0411 | 1.3480 | 0.2346 | 0.0272 |
| cluster3964rr | -2988.6998 | 0.0379 | 0.1971 | 5.1962 | 999.0000 | 0.0240 |
| cluster3965rr | -1515.7792 | 0.0296 | 0.0635 | 2.1478 | 0.1489 | 0.0128 |
| cluster3966rr | -1432.9638 | 0.0233 | 0.0176 | 0.7565 | 0.0337 | 0.0044 |
| cluster3967rr | -1751.114 | 0.1005 | 0.0390 | 0.3875 | 0.0275 | 0.0089 |
| cluster3968rr | -1167.6359 | 0.0168 | 0.0176 | 1.0462 | 0.0268 | 0.0046 |
| cluster3969rr | -899.56818 | 0.0001 | 0.0000 | 0.3287 | 0.0001 | 0.0000 |
| cluster3970rr | -2478.9402 | 0.0379 | 0.0607 | 1.5996 | 0.2084 | 0.0221 |
| cluster3971rr | -1681.6227 | 0.0238 | 0.0412 | 1.7307 | 1.4765 | 0.0233 |
| cluster3972rr | -989.67746 | 0.0128 | 0.0170 | 1.3313 | 0.0333 | 0.0058 |
| cluster3973rr | -1419.7956 | 0.0084 | 0.0080 | 0.9424 | 0.0229 | 0.0018 |
| cluster3974rr | -1830.8066 | 0.0151 | 0.0163 | 1.0801 | 0.0458 | 0.0145 |
| cluster3975rr | -1270.8806 | 0.0251 | 0.0269 | 1.0699 | 0.0562 | 0.0051 |
| cluster3976rr | -1578.0326 | 0.0051 | 0.0132 | 2.5877 | 0.1230 | 0.0110 |
| cluster3977rr | -2229.2747 | 0.0876 | 0.0864 | 0.9869 | 0.1941 | 0.0231 |
| cluster3978rr | -879.53125 | 0.0011 | 0.0008 | 0.6887 | 0.0885 | 0.0122 |
| cluster3979rr | -1620.8315 | 0.0018 | 0.0018 | 0.9821 | 0.0053 | 0.0019 |
| cluster3980rr | -922.24867 | 0.0001 | 0.0000 | 0.2139 | 0.0001 | 0.0000 |
| cluster3981rr | -1980.0849 | 0.0230 | 0.0249 | 1.0845 | 0.0091 | 0.0051 |
| cluster3982rr | -1829.7695 | 0.0478 | 0.0501 | 1.0478 | 0.0685 | 0.0125 |
| cluster3983rr | -2676.6123 | 0.0319 | 0.0702 | 2.1994 | 0.0665 | 0.0113 |
| cluster3984rr | -1373.7174 | 0.0119 | 0.0748 | 6.2939 | 0.4027 | 0.0420 |
| cluster3985rr | -978.69493 | 0.0129 | 0.0062 | 0.4784 | 0.0001 | 0.0000 |
| cluster3986rr | -1731.5109 | 0.0115 | 0.0343 | 2.9855 | 0.0775 | 0.0141 |
| cluster3987rr | -2086.6297 | 0.0107 | 0.0138 | 1.2865 | 0.0200 | 0.0104 |
| cluster3988rr | -1238.345 | 0.0045 | 0.0039 | 0.8783 | 0.1097 | 0.0125 |
| cluster3989rr | -1834.4632 | 0.0508 | 0.0477 | 0.9390 | 0.0815 | 0.0585 |
| cluster3990rr | -2205.9729 | 0.0211 | 0.0469 | 2.2262 | 0.1292 | 0.0128 |
| cluster3991rr | -1522.0606 | 0.0209 | 0.0321 | 1.5361 | 0.0155 | 0.0031 |
| cluster3992rr | -1889.07 | 0.0355 | 0.0177 | 0.4996 | 0.0445 | 0.0185 |
| cluster3993rr | -1672.9724 | 0.0299 | 0.0416 | 1.3906 | 0.2307 | 0.0176 |
| cluster3994rr | -2392.4239 | 0.0674 | 0.0629 | 0.9340 | 0.0259 | 0.0242 |
| cluster3995rr | -1617.8273 | 0.0322 | 0.0365 | 1.1340 | 0.1155 | 0.0135 |

| | | | | | | |
|---|---|---|---|---|---|---|
| cluster3996rr | -1708.986 | 0.0064 | 0.0168 | 2.6412 | 0.0072 | 0.0025 |
| cluster3997rr | -2518.1728 | 0.0503 | 0.0640 | 1.2724 | 0.1088 | 0.0164 |
| cluster3998rr | -1411.3445 | 0.0414 | 0.0378 | 0.9134 | 0.3155 | 0.0337 |
| cluster3999rr | -2524.832 | 0.0303 | 0.0534 | 1.7615 | 0.0455 | 0.0117 |
| cluster4000rr | -2286.7458 | 0.0392 | 0.0396 | 1.0115 | 0.1546 | 0.0253 |
| cluster4001rr | -1737.3809 | 0.0139 | 0.0174 | 1.2492 | 0.0415 | 0.0261 |
| cluster4002rr | -1800.6521 | 0.0028 | 0.0167 | 6.0659 | 999.0000 | 0.0020 |
| cluster4003rr | -1414.9395 | 0.0497 | 0.0624 | 1.2557 | 0.3365 | 0.0594 |
| cluster4004rr | -1442.6554 | 0.0076 | 0.0074 | 0.9674 | 0.0187 | 0.0020 |
| cluster4005rr | -2062.8672 | 0.0339 | 0.0788 | 2.3221 | 999.0000 | 0.0215 |
| cluster4006rr | -1089.5507 | 0.0324 | 0.0255 | 0.7864 | 0.0256 | 0.0027 |
| cluster4007rr | -1424.2913 | 0.0020 | 0.0019 | 0.9382 | 0.0397 | 0.0020 |
| cluster4008rr | -1355.6159 | 0.0082 | 0.0200 | 2.4484 | 0.0001 | 0.0000 |
| cluster4009rr | -1546.5376 | 0.0131 | 0.0333 | 2.5367 | 0.0144 | 0.0041 |
| cluster4010rr | -1626.0971 | 0.0513 | 0.0975 | 1.9012 | 999.0000 | 0.0082 |
| cluster4011rr | -1061.3733 | 0.0727 | 0.0134 | 0.1839 | 0.0470 | 0.0046 |
| cluster4012rr | -2437.8996 | 0.0077 | 0.1256 | 16.3027 | 0.2215 | 0.0204 |
| cluster4013rr | -846.38487 | 0.0212 | 0.0110 | 0.5179 | 0.0435 | 0.0078 |
| cluster4014rr | -1137.3027 | 0.0069 | 0.0089 | 1.2879 | 0.0001 | 0.0000 |
| cluster4015rr | -617.75203 | 0.0434 | 0.0018 | 0.0407 | 0.0001 | 0.0000 |
| cluster4016rr | -1815.7162 | 0.0187 | 0.0186 | 0.9930 | 0.0205 | 0.0134 |
| cluster4017rr | -1342.1763 | 0.0432 | 0.0551 | 1.2756 | 0.0246 | 0.0043 |
| cluster4018rr | -1325.6416 | 0.1331 | 0.1521 | 1.1434 | 0.2317 | 0.0182 |
| cluster4019rr | -1823.9746 | 0.0566 | 0.0848 | 1.4970 | 0.3036 | 0.0251 |
| cluster4020rr | -1004.8938 | 0.0328 | 0.0152 | 0.4635 | 0.0686 | 0.0131 |
| cluster4021rr | -1590.3118 | 0.0218 | 0.0132 | 0.6076 | 0.1209 | 0.0111 |
| cluster4022rr | -970.06149 | 0.0080 | 0.0063 | 0.7918 | 0.0202 | 0.0027 |
| cluster4023rr | -1844.4302 | 0.0668 | 0.0569 | 0.8518 | 0.0494 | 0.0071 |
| cluster4024rr | -2359.8134 | 0.0296 | 0.0666 | 2.2480 | 0.0316 | 0.0145 |
| cluster4025rr | -624.38865 | 0.0006 | 0.0019 | 2.8925 | 0.0001 | 0.0000 |
| cluster4026rr | -1088.9002 | 0.0028 | 0.0086 | 3.1114 | 0.0001 | 0.0000 |
| cluster4027rr | -983.16605 | 0.0051 | 0.0162 | 3.1741 | 2.1480 | 0.0000 |
| cluster4028rr | -1845.8232 | 0.0050 | 0.0159 | 3.1846 | 0.0269 | 0.0062 |
| cluster4029rr | -2241.1366 | 0.0501 | 0.0823 | 1.6433 | 0.7554 | 0.0540 |
| cluster4030rr | -1124.8986 | 0.0357 | 0.0268 | 0.7510 | 0.0764 | 0.0046 |
| cluster4031rr | -1097.6206 | 0.0235 | 0.0483 | 2.0577 | 0.0311 | 0.0099 |
| cluster4032rr | -1475.3307 | 0.0012 | 0.0005 | 0.3930 | 0.0001 | 0.0000 |
| cluster4033rr | -2213.4177 | 0.0608 | 0.0957 | 1.5726 | 0.1807 | 0.0354 |
| cluster4034rr | -493.6726 | 0.0001 | 0.0001 | 0.9153 | 0.0001 | 0.0000 |
| cluster4035rr | -2115.5558 | 0.0619 | 0.0354 | 0.5711 | 0.0503 | 0.0198 |
| cluster4036rr | -1077.1411 | 0.0092 | 0.0132 | 1.4478 | 0.0001 | 0.0000 |
| cluster4037rr | -1625.8552 | 0.0200 | 0.0333 | 1.6674 | 0.0127 | 0.0042 |
| cluster4038rr | -901.66639 | 0.0171 | 0.0355 | 2.0717 | 999.0000 | 0.0100 |
| cluster4039rr | -1507.4682 | 0.0646 | 0.0255 | 0.3946 | 0.0001 | 0.0000 |

| | | | | | | |
|---|---|---|---|---|---|---|
| cluster4040rr | -1126.4622 | 0.0345 | 0.0301 | 0.8725 | 0.0170 | 0.0085 |
| cluster4041rr | -1474.9381 | 0.0150 | 0.0245 | 1.6364 | 0.0054 | 0.0024 |
| cluster4042rr | -899.67193 | 0.0314 | 0.0259 | 0.8256 | 0.0488 | 0.0030 |
| cluster4043rr | -1136.926 | 0.0315 | 0.1197 | 3.8034 | 0.2117 | 0.0218 |
| cluster4044rr | -957.98381 | 0.0015 | 0.0027 | 1.8707 | 0.0001 | 0.0000 |
| cluster4045rr | -2015.4829 | 0.0461 | 0.0765 | 1.6622 | 1.3500 | 0.0353 |
| cluster4046rr | -889.88447 | 0.0028 | 0.0014 | 0.4989 | 0.0001 | 0.0000 |
| cluster4047rr | -2062.7326 | 0.0124 | 0.0327 | 2.6337 | 0.4028 | 0.0192 |
| cluster4048rr | -1470.311 | 0.0080 | 0.0043 | 0.5351 | 0.0152 | 0.0016 |
| cluster4049rr | -573.68806 | 0.0011 | 0.0040 | 3.7384 | 0.0013 | 0.0029 |
| cluster4050rr | -1216.6888 | 0.0096 | 0.0079 | 0.8172 | 0.0001 | 0.0000 |
| cluster4051rr | -1403.7863 | 0.0094 | 0.0033 | 0.3478 | 0.0001 | 0.0000 |
| cluster4052rr | -1288.1388 | 0.0198 | 0.0106 | 0.5325 | 0.0159 | 0.0081 |
| cluster4053rr | -1642.7771 | 0.0097 | 0.0360 | 3.7264 | 0.0764 | 0.0066 |
| cluster4054rr | -1854.0578 | 0.0125 | 0.0364 | 2.9108 | 999.0000 | 0.0170 |
| cluster4055rr | -717.92043 | 0.0019 | 0.0158 | 8.4219 | 999.0000 | 0.0060 |
| cluster4056rr | -932.40754 | 0.0274 | 0.0275 | 1.0041 | 0.1202 | 0.0239 |
| cluster4057rr | -1565.0006 | 0.0334 | 0.0435 | 1.3007 | 0.0488 | 0.0104 |
| cluster4058rr | -1677.9137 | 0.0313 | 0.0507 | 1.6209 | 0.1721 | 0.0218 |
| cluster4059rr | -722.86921 | 0.0137 | 0.0277 | 2.0290 | 0.0001 | 0.0000 |
| cluster4060rr | -1875.8598 | 0.0286 | 0.0295 | 1.0313 | 0.0243 | 0.0060 |
| cluster4061rr | -1186.3151 | 0.0047 | 0.0126 | 2.7052 | 0.1131 | 0.0055 |
| cluster4062rr | -1188.0268 | 0.0022 | 0.0073 | 3.3042 | 26.1086 | 0.0346 |
| cluster4063rr | -1718.0893 | 0.0216 | 0.0625 | 2.8894 | 999.0000 | 0.0156 |
| cluster4064rr | -782.72456 | 0.0175 | 0.0194 | 1.1116 | 0.0001 | 0.0000 |
| cluster4065rr | -989.31178 | 0.0087 | 0.0060 | 0.6913 | 0.0001 | 0.0000 |
| cluster4066rr | -1616.8696 | 0.0469 | 0.0515 | 1.0973 | 0.0347 | 0.0067 |
| cluster4067rr | -418.63845 | 0.0189 | 0.0822 | 4.3475 | 0.0225 | 0.0098 |
| cluster4068rr | -1352.3768 | 0.0152 | 0.0614 | 4.0409 | 0.0976 | 0.0139 |
| cluster4069rr | -1062.0617 | 0.0496 | 0.0323 | 0.6512 | 1.1206 | 0.0252 |
| cluster4070rr | -696.52332 | 0.0048 | 0.0330 | 6.9357 | 8.0501 | 0.0000 |
| cluster4071rr | -826.7776 | 0.0022 | 0.0049 | 2.2473 | 0.0001 | 0.0000 |
| cluster4072rr | -987.67795 | 0.0230 | 0.0256 | 1.1132 | 0.0809 | 0.0065 |
| cluster4073rr | -1877.9081 | 0.0207 | 0.0430 | 2.0739 | 0.5391 | 0.0306 |
| cluster4074rr | -1256.1468 | 0.0289 | 0.0631 | 2.1795 | 0.1043 | 0.0192 |
| cluster4075rr | -709.437 | 0.8709 | 1.4343 | 1.6469 | 0.0758 | 3.6829 |
| cluster4076rr | -1650.0791 | 0.0843 | 0.0465 | 0.5516 | 0.1157 | 0.0425 |
| cluster4077rr | -1676.0967 | 0.0382 | 0.0468 | 1.2261 | 0.5011 | 0.0124 |
| cluster4078rr | -1231.4682 | 0.0253 | 0.0281 | 1.1088 | 0.0065 | 0.0039 |
| cluster4079rr | -1207.4119 | 0.0011 | 0.0012 | 1.0512 | 0.0001 | 0.0000 |
| cluster4080rr | -735.28114 | 0.0042 | 0.0201 | 4.7716 | 0.0440 | 0.0146 |
| cluster4081rr | -1130.2911 | 0.0160 | 0.0139 | 0.8705 | 0.0433 | 0.0118 |
| cluster4082rr | -742.67007 | 0.0108 | 0.0088 | 0.8200 | 0.1489 | 0.2983 |
| cluster4083rr | -1702.3278 | 0.0459 | 0.0861 | 1.8772 | 0.2805 | 0.0402 |

| | | | | | | |
|---|---|---|---|---|---|---|
| cluster4084rr | -715.0729 | 0.0126 | 0.0085 | 0.6723 | 0.0965 | 0.0050 |
| cluster4085rr | -858.81121 | 0.0310 | 0.0115 | 0.3706 | 0.0001 | 0.0000 |
| cluster4086rr | -1314.4881 | 0.0278 | 0.0499 | 1.7944 | 0.0540 | 0.0109 |
| cluster4087rr | -1206.0256 | 0.0122 | 0.0324 | 2.6679 | 0.0122 | 0.0021 |
| cluster4088rr | -1313.1938 | 0.0608 | 0.0250 | 0.4119 | 0.0001 | 0.0000 |
| cluster4089rr | -1507.4876 | 0.0040 | 0.0179 | 4.5197 | 999.0000 | 0.0064 |
| cluster4090rr | -753.87364 | 0.0303 | 0.0223 | 0.7341 | 0.2180 | 0.0132 |
| cluster4091rr | -953.8629 | 0.0302 | 0.0124 | 0.4113 | 0.0778 | 0.0029 |
| cluster4092rr | -1419.818 | 0.0198 | 0.0454 | 2.2984 | 0.8487 | 0.0216 |
| cluster4093rr | -1006.9591 | 0.0181 | 0.0268 | 1.4770 | 0.0181 | 0.0062 |
| cluster4094rr | -375.3961 | 0.0096 | 0.0015 | 0.1605 | 0.0001 | 0.0000 |
| cluster4095rr | -1403.6574 | 0.0475 | 0.0657 | 1.3845 | 0.0160 | 0.0428 |
| cluster4096rr | -1212.7656 | 0.0205 | 0.0410 | 1.9982 | 0.0433 | 0.0119 |
| cluster4097rr | -705.04819 | 0.0808 | 0.0075 | 0.0933 | 0.1313 | 0.0248 |
| cluster4098rr | -1868.6233 | 0.0318 | 0.0420 | 1.3184 | 0.0407 | 0.0260 |
| cluster4099rr | -810.85973 | 0.0028 | 0.0040 | 1.4383 | 0.0001 | 0.0000 |
| cluster4100rr | -1047.952 | 0.0087 | 0.0110 | 1.2657 | 0.0001 | 0.0000 |
| cluster4101rr | -2088.901 | 0.2589 | 0.1397 | 0.5396 | 0.1651 | 0.0576 |
| cluster4102rr | -1856.2727 | 0.0379 | 0.1715 | 4.5251 | 0.1491 | 0.0151 |
| cluster4103rr | -1106.9415 | 0.0398 | 0.0118 | 0.2962 | 0.1035 | 0.0048 |
| cluster4104rr | -1017.8595 | 0.0077 | 0.0139 | 1.8136 | 0.0001 | 0.0000 |
| cluster4105rr | -735.60571 | 0.0084 | 0.0588 | 7.0305 | 999.0000 | 0.0021 |
| cluster4106rr | -1000.5548 | 0.0082 | 0.0151 | 1.8375 | 0.0001 | 0.0000 |
| cluster4107rr | -786.7717 | 0.0025 | 0.0036 | 1.4827 | 0.0001 | 0.0000 |
| cluster4108rr | -782.8522 | 0.0137 | 0.0206 | 1.5078 | 0.0001 | 0.0001 |
| cluster4109rr | -1651.8546 | 0.0659 | 0.1067 | 1.6181 | 0.0783 | 0.0193 |
| cluster4110rr | -1243.9782 | 0.0510 | 0.0721 | 1.4140 | 0.2490 | 0.0240 |
| cluster4111rr | -924.98775 | 0.0459 | 0.0757 | 1.6477 | 0.1186 | 0.0048 |
| cluster4112rr | -1113.3414 | 0.0169 | 0.0138 | 0.8168 | 0.0113 | 0.0030 |
| cluster4113rr | -773.32637 | 0.0061 | 0.0075 | 1.2279 | 0.0001 | 0.0000 |
| cluster4114rr | -1073.3446 | 0.0473 | 0.0310 | 0.6548 | 0.0001 | 0.0000 |
| cluster4115rr | -1155.247 | 0.0088 | 0.0184 | 2.1039 | 0.0544 | 0.0056 |
| cluster4116rr | -634.55051 | 0.0253 | 0.0370 | 1.4612 | 0.2245 | 0.1462 |
| cluster4117rr | -660.32856 | 0.0138 | 0.0083 | 0.6031 | 0.0001 | 0.0000 |
| cluster4118rr | -1288.093 | 0.0277 | 0.0475 | 1.7156 | 0.0884 | 0.0128 |
| cluster4119rr | -922.75144 | 0.0134 | 0.0296 | 2.2054 | 0.2197 | 0.0147 |
| cluster4120rr | -965.76102 | 0.0020 | 0.0017 | 0.8423 | 0.0582 | 0.0065 |
| cluster4121rr | -1217.0348 | 0.0038 | 0.0301 | 8.0094 | 0.0191 | 0.0034 |
| cluster4122rr | -527.68051 | 0.0343 | 0.0027 | 0.0784 | 0.3577 | 0.0000 |
| cluster4123rr | -468.58017 | 0.0074 | 0.0058 | 0.7869 | 0.0001 | 0.0000 |
| cluster4124rr | -905.35774 | 0.0047 | 0.0171 | 3.6257 | 0.0001 | 0.0000 |
| cluster4125rr | -1064.3673 | 0.0982 | 0.0780 | 0.7941 | 0.0283 | 0.0158 |
| cluster4126rr | -762.27556 | 0.0897 | 0.0307 | 0.3429 | 0.0001 | 0.0000 |
| cluster4127rr | -766.41339 | 0.0558 | 0.0411 | 0.7375 | 0.0757 | 0.0092 |

| | | | | | | | | |
|---|---|---|---|---|---|---|---|---|
| cluster4128rr | -987.82585 | | 0.0271 | 0.1038 | 3.8306 | | 0.3266 | 0.0185 |

| dS | Adjusted p value |
|---|---|
| 0.1327 | 0.03 |
| 0.7934 | 5.94E-65 |
| 0.1160 | 0.0005 |
| 0.0930 | 2.61E-93 |
| 0.1418 | 2.66E-07 |
| 0.1842 | 3.83E-05 |
| 0.1697 | 5.27E-54 |
| 0.0319 | 2.63E-09 |
| 0.5653 | 7.16E-124 |
| 0.3284 | 1.52E-41 |
| 0.7357 | 2.83E-88 |
| 0.1194 | 2.97E-44 |
| 0.8147 | 3.08E-107 |
| 0.5655 | 2.17E-12 |
| 0.1296 | 8.39E-49 |
| 0.0424 | 3.31E-29 |
| 0.1051 | 1.52E-08 |
| 0.0629 | 6.51E-06 |
| 0.0942 | 3.65E-06 |
| 0.1486 | 8.20E-11 |
| 0.5679 | 1.20E-59 |
| 0.1007 | 2.90E-22 |
| 0.9871 | 1.99E-64 |
| 0.7075 | 5.47E-11 |
| 0.4905 | 7.51E-21 |
| 0.2373 | 5.07E-12 |
| 0.1671 | 0.005 |
| 0.0689 | 5.19E-18 |
| 0.6367 | 6.88E-19 |
| 0.2518 | 2.27E-27 |
| 0.5551 | 1.70E-07 |
| 0.1045 | 1.21E-33 |
| 0.0890 | 2.78E-33 |
| 0.4639 | 7.23E-25 |
| 0.2522 | 7.29E-55 |
| 0.2012 | 5.45E-08 |
| 0.1232 | 3.67E-47 |
| 0.0649 | 0.06 |
| 0.0882 | 5.78E-25 |
| 0.5602 | 1.90E-34 |
| 0.0854 | 2.12E-15 |
| 0.0961 | 3.47E-17 |

| | |
|---|---|
| 0.1247 | 3.34E-19 |
| 0.1698 | 4.14E-66 |
| 0.0726 | 2.08E-22 |
| 0.4765 | 1.66E-14 |
| 1.5692 | 1.65E-29 |
| 0.5511 | 1.58E-39 |
| 0.1828 | 1.48E-20 |
| 0.0879 | 1.10E-13 |
| 0.7417 | 1.37E-45 |
| 0.2122 | 5.91E-05 |
| 0.3337 | 3.33E-19 |
| 0.4468 | 4.81E-16 |
| 0.1210 | 1.79E-11 |
| 0.1107 | 5.41E-14 |
| 0.0769 | 6.68E-05 |
| 0.0756 | 2.02E-20 |
| 0.1200 | 4.52E-10 |
| 0.1544 | 4.38E-24 |
| 0.1839 | 4.93E-41 |
| 0.3459 | 5.30E-25 |
| 0.1631 | 4.27E-19 |
| 0.4972 | 1.06E-26 |
| 0.1958 | 7.84E-30 |
| 0.0977 | 0.0001 |
| 0.3518 | 3.57E-13 |
| 0.1864 | 4.05E-30 |
| 0.1306 | 6.64E-27 |
| 0.0872 | 1.25E-09 |
| 0.0870 | 0.06 |
| 0.1768 | 5.11E-33 |
| 0.1848 | 1.23E-14 |
| 0.8247 | 2.74E-41 |
| 0.2615 | 1.89E-11 |
| 0.1506 | 8.19E-25 |
| 0.0588 | 9.66E-51 |
| 0.1035 | 1.00E-09 |
| 0.1507 | 2.31E-10 |
| 0.1589 | 1.25E-12 |
| 0.3041 | 2.04E-14 |
| 0.0816 | 6.19E-12 |
| 0.5097 | 4.29E-14 |
| 0.0482 | 6.34E-25 |
| 0.3266 | 1.24E-12 |
| 0.4327 | 3.61E-29 |

| | |
|---|---|
| 0.1703 | 1.48E-06 |
| 0.0892 | 1.63E-15 |
| 0.1086 | 1.37E-06 |
| 0.1444 | 8.91E-17 |
| 0.5448 | 1.23E-23 |
| 0.6979 | 4.57E-06 |
| 0.1532 | 3.61E-21 |
| 0.2415 | 1.79E-15 |
| 0.1136 | 4.22E-21 |
| 0.3805 | 4.44E-16 |
| 0.5346 | 2.54E-23 |
| 0.5920 | 3.62E-27 |
| 0.3711 | 1.49E-10 |
| 0.1065 | 3.10E-22 |
| 0.2852 | 9.44E-15 |
| 0.6030 | 2.39E-18 |
| 0.1613 | 1.58E-15 |
| 1.1497 | 7.84E-39 |
| 0.0944 | 3.45E-06 |
| 0.1308 | 3.61E-48 |
| 0.4577 | 2.82E-09 |
| 0.7288 | 5.21E-36 |
| 0.7148 | 8.70E-47 |
| 0.9837 | 7.26E-21 |
| 0.2381 | 1.87E-14 |
| 0.4291 | 7.23E-14 |
| 0.1113 | 4.75E-32 |
| 0.2511 | 4.99E-18 |
| 0.4411 | 5.00E-40 |
| 0.4393 | 1.05E-21 |
| 0.1088 | 1.10E-15 |
| 0.1535 | 3.53E-30 |
| 0.0671 | 2.18E-05 |
| 1.1166 | 5.08E-17 |
| 0.0284 | 5.10E-06 |
| 0.0621 | 2.28E-08 |
| 0.2122 | 5.35E-11 |
| 0.0506 | 0.48 |
| 0.1130 | 3.51E-34 |
| 0.4372 | 5.67E-57 |
| 0.0925 | 2.82E-19 |
| 0.1644 | 2.00E-05 |
| 0.1311 | 5.45E-11 |
| 0.1483 | 4.05E-05 |

| | |
|---|---|
| 0.1108 | 5.54E-27 |
| 0.3723 | 9.89E-13 |
| 0.2067 | 0.24 |
| 0.0839 | 0.01 |
| 0.1959 | 1.06E-13 |
| 0.1229 | 1.06E-19 |
| 0.3488 | 3.75E-13 |
| 0.1986 | 4.54E-14 |
| 0.1360 | 7.54E-08 |
| 0.1902 | 5.55E-12 |
| 0.6411 | 7.13E-17 |
| 0.1142 | 2.87E-10 |
| 0.0000 | 0.10 |
| 0.5566 | 4.74E-16 |
| 0.1488 | 5.08E-11 |
| 0.1430 | 5.03E-27 |
| 0.2039 | 1.01E-15 |
| 0.1441 | 5.58E-18 |
| 0.1420 | 4.05E-16 |
| 0.1910 | 4.85E-22 |
| 0.1171 | 1.82E-06 |
| 0.2976 | 2.78E-05 |
| 0.0579 | 4.42E-05 |
| 0.4964 | 5.93E-10 |
| 0.1684 | 1.84E-23 |
| 0.1637 | 1.06E-16 |
| 0.1623 | 9.07E-10 |
| 0.0903 | 0.005 |
| 0.7309 | 9.63E-17 |
| 0.2494 | 2.17E-07 |
| 0.1946 | 1.42E-06 |
| 0.1902 | 1.32E-24 |
| 0.0395 | 0.0002 |
| 0.0807 | 6.54E-09 |
| 0.1044 | 3.29E-20 |
| 0.2008 | 4.83E-32 |
| 0.5706 | 1.14E-11 |
| 0.0774 | 1.11E-15 |
| 0.3404 | 1.94E-10 |
| 0.1181 | 0.0002 |
| 0.3066 | 1.23E-19 |
| 0.0379 | 0.02 |
| 0.1302 | 1.19E-12 |
| 0.0817 | 0.14 |

| | |
|---|---|
| 0.6828 | 1.34E-30 |
| 0.1154 | 2.57E-12 |
| 0.1391 | 6.84E-15 |
| 0.2218 | 3.50E-08 |
| 0.6071 | 5.89E-07 |
| 0.6890 | 4.85E-22 |
| 0.0529 | 0.14 |
| 33.0379 | 0.005 |
| 0.3372 | 6.56E-25 |
| 0.1421 | 4.30E-08 |
| 0.1029 | 1.10E-06 |
| 0.0901 | 4.66E-07 |
| 0.1725 | 3.06E-11 |
| 0.2855 | 3.83E-14 |
| 0.1149 | 3.78E-15 |
| 0.1125 | 3.02E-12 |
| 0.0241 | 6.63E-09 |
| 0.2152 | 1.50E-16 |
| 0.2075 | 6.43E-11 |
| 0.7472 | 0.01 |
| 0.1638 | 9.86E-09 |
| 0.1039 | 3.66E-17 |
| 0.5792 | 3.19E-31 |
| 0.1097 | 1.98E-14 |
| 0.1381 | 4.37E-14 |
| 0.0864 | 1.07E-13 |
| 0.5223 | 1.25E-07 |
| 0.0930 | 1.14E-32 |
| 0.1010 | 4.30E-08 |
| 0.1627 | 9.81E-05 |
| 0.0000 | 0.05 |
| 0.1603 | 1.77E-05 |
| 0.0592 | 1.97E-06 |
| 0.1874 | 1.13E-11 |
| 0.3781 | 2.28E-17 |
| 0.1400 | 6.23E-10 |
| 0.2357 | 1.16E-16 |
| 0.0956 | 3.80E-05 |
| 0.1311 | 0.02 |
| 0.4207 | 4.95E-08 |
| 0.4140 | 6.39E-11 |
| 0.1185 | 0.003 |
| 0.0508 | 0.001 |
| 0.5721 | 2.87E-13 |

| | |
|---|---|
| 0.3179 | 2.19E-17 |
| 0.2037 | 1.44E-07 |
| 0.0731 | 9.84E-15 |
| 0.1950 | 1.37E-13 |
| 0.3877 | 7.48E-13 |
| 0.4087 | 0.001 |
| 0.5334 | 2.48E-16 |
| 0.2804 | 1.52E-16 |
| 0.7656 | 4.73E-20 |
| 0.1690 | 9.15E-21 |
| 0.2905 | 3.81E-18 |
| 0.1664 | 5.26E-10 |
| 0.3963 | 4.00E-11 |
| 0.0756 | 0.002 |
| 0.0889 | 2.11E-20 |
| 0.2635 | 0.003 |
| 0.0955 | 2.45E-20 |
| 0.1865 | 3.02E-09 |
| 0.4389 | 0.0001 |
| 0.2343 | 6.88E-28 |
| 0.2040 | 3.20E-07 |
| 0.1368 | 3.47E-14 |
| 0.2107 | 3.37E-16 |
| 0.2503 | 5.43E-21 |
| 0.5368 | 7.72E-27 |
| 0.1176 | 1.93E-14 |
| 0.2643 | 9.36E-19 |
| 0.1683 | 2.14E-11 |
| 0.4918 | 3.40E-09 |
| 0.0760 | 1.27E-09 |
| 0.7478 | 2.18E-08 |
| 0.8180 | 1.92E-12 |
| 0.3222 | 2.05E-09 |
| 0.3539 | 3.56E-15 |
| 0.2245 | 8.96E-17 |
| 0.2713 | 2.54E-23 |
| 0.0653 | 2.83E-05 |
| 0.1216 | 0.01 |
| 0.0004 | 8.51E-13 |
| 0.0730 | 0.0003 |
| 0.1754 | 1.77E-11 |
| 0.1433 | 6.21E-10 |
| 0.1540 | 2.30E-10 |
| 0.2731 | 2.11E-10 |

| | |
|---|---|
| 0.1335 | 4.09E-14 |
| 0.1615 | 1.36E-06 |
| 0.0467 | 2.56E-13 |
| 0.0167 | 0.01 |
| 0.5156 | 5.36E-06 |
| 0.1376 | 3.22E-16 |
| 0.5124 | 2.84E-18 |
| 0.2865 | 2.39E-10 |
| 0.3280 | 1.02E-14 |
| 0.1997 | 4.55E-14 |
| 0.1522 | 4.30E-05 |
| 0.5529 | 9.65E-16 |
| 0.1295 | 3.38E-09 |
| 0.1342 | 5.51E-10 |
| 0.1212 | 4.91E-06 |
| 0.1160 | 2.90E-14 |
| 0.4317 | 2.70E-10 |
| 0.5368 | 1.36E-15 |
| 0.1371 | 1.73E-08 |
| 0.2092 | 2.38E-05 |
| 0.1736 | 3.82E-12 |
| 0.1438 | 2.92E-09 |
| 0.1232 | 1.13E-11 |
| 0.9837 | 8.91E-09 |
| 0.2843 | 4.06E-06 |
| 0.0587 | 1.61E-05 |
| 0.2460 | 3.86E-08 |
| 0.0144 | 0.15 |
| 0.3206 | 6.65E-12 |
| 0.0580 | 5.62E-12 |
| 0.3721 | 9.66E-51 |
| 16.5395 | 1.83E-13 |
| 0.1463 | 5.69E-27 |
| 0.0737 | 2.57E-14 |
| 0.4008 | 1.00E-09 |
| 0.0544 | 8.83E-05 |
| 0.1047 | 3.85E-06 |
| 0.1708 | 1.32E-13 |
| 0.0958 | 9.59E-09 |
| 0.2096 | 2.08E-27 |
| 0.0667 | 3.53E-17 |
| 0.1793 | 2.28E-16 |
| 0.0518 | 0.02 |
| 1.0550 | 8.57E-16 |

| | |
|---|---|
| 0.1495 | 1.49E-10 |
| 0.1076 | 0.12 |
| 0.1514 | 6.03E-11 |
| 0.0489 | 9.46E-11 |
| 0.1879 | 2.39E-08 |
| 0.1097 | 2.91E-09 |
| 0.2112 | 6.11E-11 |
| 0.5364 | 5.06E-16 |
| 0.0486 | 8.50E-05 |
| 1.0529 | 6.32E-30 |
| 0.2080 | 3.17E-14 |
| 0.0147 | 0.02 |
| 0.2837 | 1.72E-07 |
| 0.2965 | 5.75E-17 |
| 0.1184 | 1.35E-20 |
| 0.1071 | 4.96E-09 |
| 0.1159 | 8.83E-07 |
| 0.1778 | 2.52E-07 |
| 0.2454 | 0.0005 |
| 0.9702 | 4.38E-41 |
| 0.0144 | 1.62E-05 |
| 0.0494 | 3.15E-07 |
| 0.8399 | 9.98E-25 |
| 0.2284 | 4.10E-09 |
| 0.3347 | 0.0001 |
| 0.1060 | 8.64E-09 |
| 0.0935 | 5.61E-18 |
| 0.1274 | 3.74E-06 |
| 0.1229 | 8.97E-18 |
| 0.4029 | 0.0005 |
| 0.1343 | 1.04E-10 |
| 0.2105 | 6.99E-14 |
| 0.2124 | 7.08E-05 |
| 0.3521 | 4.36E-10 |
| 0.0776 | 1.05E-06 |
| 0.2898 | 4.05E-09 |
| 0.5165 | 8.97E-18 |
| 0.0770 | 6.37E-06 |
| 0.2654 | 9.28E-08 |
| 0.1868 | 1.11E-20 |
| 0.1275 | 7.03E-12 |
| 0.4450 | 1.35E-19 |
| 0.0706 | 0.003 |
| 0.0966 | 7.77E-11 |

| | |
|---|---|
| 0.1250 | 0.0001 |
| 0.1132 | 0.002 |
| 0.0913 | 4.20E-05 |
| 0.1606 | 9.38E-10 |
| 0.4728 | 8.53E-26 |
| 0.2823 | 1.41E-09 |
| 0.1090 | 2.01E-19 |
| 0.1921 | 3.36E-21 |
| 0.1213 | 1.02E-09 |
| 0.2274 | 1.39E-12 |
| 0.6477 | 0.0001 |
| 0.4459 | 8.47E-12 |
| 0.0410 | 0.0003 |
| 0.1997 | 6.26E-11 |
| 0.3239 | 9.53E-08 |
| 0.0884 | 4.11E-14 |
| 0.1439 | 1.23E-23 |
| 0.3642 | 3.93E-06 |
| 0.0675 | 1.36E-14 |
| 0.2729 | 6.57E-06 |
| 0.0473 | 0.002 |
| 0.0669 | 8.53E-06 |
| 0.0761 | 2.34E-17 |
| 0.1554 | 6.97E-09 |
| 0.1889 | 4.16E-06 |
| 0.1559 | 4.54E-10 |
| 0.3729 | 3.67E-09 |
| 0.4086 | 1.05E-14 |
| 0.1359 | 1.72E-11 |
| 0.1732 | 1.27E-18 |
| 0.5028 | 4.41E-08 |
| 0.5859 | 3.02E-07 |
| 0.2354 | 5.07E-12 |
| 0.2276 | 2.99E-12 |
| 0.1744 | 2.91E-05 |
| 0.0714 | 0.003 |
| 0.1309 | 2.70E-05 |
| 0.6103 | 2.80E-19 |
| 0.1627 | 1.23E-07 |
| 0.0924 | 2.68E-09 |
| 0.1695 | 0.01 |
| 0.4978 | 2.13E-10 |
| 0.1467 | 1.39E-21 |
| 1.0575 | 3.07E-11 |

| | |
|---|---|
| 0.1106 | 0.04 |
| 0.1080 | 7.32E-13 |
| 0.0989 | 5.65E-14 |
| 0.1800 | 0.01 |
| 0.1091 | 1.09E-07 |
| 0.2910 | 5.68E-08 |
| 0.1621 | 2.17E-05 |
| 0.1046 | 4.52E-09 |
| 0.3562 | 1.73E-20 |
| 0.2601 | 2.94E-13 |
| 0.6953 | 1.42E-29 |
| 0.1198 | 3.68E-08 |
| 0.1775 | 1.35E-05 |
| 0.4410 | 1.85E-18 |
| 0.0712 | 6.79E-06 |
| 0.1193 | 2.79E-22 |
| 0.1262 | 0.08 |
| 0.0509 | 2.58E-05 |
| 0.0000 | 7.84E-30 |
| 0.1011 | 0.0001 |
| 0.3751 | 9.46E-11 |
| 0.2278 | 4.79E-07 |
| 0.1027 | 1.50E-05 |
| 0.1019 | 5.77E-06 |
| 0.3400 | 5.51E-16 |
| 0.1223 | 0.72 |
| 0.1499 | 0.003 |
| 0.2539 | 1.61E-05 |
| 0.1614 | 1.36E-08 |
| 0.1409 | 0.02 |
| 0.1239 | 5.19E-05 |
| 0.0543 | 3.35E-17 |
| 0.3039 | 1.20E-13 |
| 0.0680 | 3.01E-11 |
| 0.1134 | 5.68E-08 |
| 0.0675 | 1.04E-11 |
| 0.5164 | 1.46E-07 |
| 0.0879 | 8.15E-15 |
| 0.1022 | 3.21E-07 |
| 0.0764 | 0.001 |
| 0.0669 | 2.13E-06 |
| 0.0000 | 9.81E-09 |
| 0.2855 | 7.97E-10 |
| 0.0778 | 0.001 |

| | |
|---|---|
| 0.5771 | 6.19E-12 |
| 0.1375 | 1.33E-06 |
| 0.7594 | 1.67E-19 |
| 0.3386 | 1.39E-10 |
| 0.0579 | 1.61E-10 |
| 0.0671 | 2.36E-10 |
| 0.2995 | 0.15 |
| 0.6388 | 9.52E-22 |
| 0.0163 | 1.19E-06 |
| 0.2199 | 3.78E-15 |
| 0.6073 | 0.001 |
| 0.3005 | 1.05E-05 |
| 0.7909 | 1.72E-11 |
| 0.1707 | 9.47E-07 |
| 0.1578 | 3.05E-05 |
| 0.0727 | 5.95E-06 |
| 0.0967 | 2.84E-12 |
| 0.8071 | 9.99E-19 |
| 0.2268 | 0.02 |
| 0.2648 | 2.20E-15 |
| 0.5196 | 6.90E-10 |
| 0.0000 | 8.64E-09 |
| 0.1590 | 2.13E-07 |
| 0.1155 | 8.56E-11 |
| 0.5731 | 1.93E-21 |
| 0.1561 | 1.81E-05 |
| 0.1043 | 1.51E-06 |
| 0.5739 | 0.02 |
| 0.1136 | 2.88E-10 |
| 0.1067 | 0.0001 |
| 0.0511 | 2.36E-09 |
| 0.2233 | 1.83E-07 |
| 0.1589 | 8.70E-10 |
| 0.3312 | 0.001 |
| 0.2831 | 4.41E-08 |
| 0.0234 | 0.06 |
| 0.4039 | 6.69E-13 |
| 0.0000 | 3.65E-06 |
| 0.1305 | 2.78E-05 |
| 0.1559 | 9.05E-06 |
| 0.0890 | 0.002 |
| 0.1578 | 4.13E-09 |
| 0.0657 | 1.48E-11 |
| 0.0979 | 0.03 |

| | |
|---|---|
| 0.6351 | 4.50E-09 |
| 0.3102 | 3.23E-13 |
| 0.1588 | 1.43E-05 |
| 0.1797 | 2.71E-12 |
| 0.7811 | 6.31E-05 |
| 0.3620 | 0.45 |
| 0.1545 | 1.98E-08 |
| 0.1304 | 3.46E-09 |
| 0.1265 | 6.58E-24 |
| 0.2732 | 3.86E-08 |
| 0.1246 | 6.92E-16 |
| 0.6977 | 4.00E-23 |
| 0.3027 | 1.93E-07 |
| 0.1181 | 1.67E-06 |
| 0.3247 | 2.54E-17 |
| 0.1510 | 9.14E-09 |
| 0.1021 | 2.72E-11 |
| 0.1310 | 1.61E-05 |
| 0.0251 | 1.64E-05 |
| 0.0279 | 6.91E-07 |
| 0.2082 | 3.85E-09 |
| 0.3628 | 2.04E-09 |
| 0.1034 | 0.0002 |
| 0.4148 | 9.66E-26 |
| 0.1043 | 1.25E-13 |
| 0.4354 | 7.95E-06 |
| 0.1566 | 5.29E-09 |
| 0.5554 | 1.42E-15 |
| 0.1377 | 6.34E-35 |
| 0.2735 | 1.08E-10 |
| 0.1153 | 8.77E-09 |
| 0.0566 | 3.67E-07 |
| 0.6884 | 0.01 |
| 0.1678 | 0.001 |
| 0.1468 | 9.44E-17 |
| 0.0953 | 6.25E-05 |
| 0.0271 | 1.47E-11 |
| 0.5227 | 7.49E-18 |
| 0.1745 | 4.26E-34 |
| 0.0756 | 8.12E-06 |
| 0.0857 | 2.87E-09 |
| 0.0871 | 0.01 |
| 0.1430 | 1.80E-15 |
| 0.7683 | 7.00E-12 |

| | |
|---|---|
| 0.1177 | 0.0002 |
| 0.1834 | 1.07E-06 |
| 0.5299 | 9.51E-06 |
| 0.0563 | 0.01 |
| 0.1933 | 1.04E-07 |
| 0.2494 | 6.75E-05 |
| 0.1800 | 3.17E-13 |
| 0.1946 | 2.38E-20 |
| 0.2452 | 3.28E-07 |
| 0.1582 | 1.98E-09 |
| 0.1200 | 3.38E-09 |
| 0.3488 | 1.38E-06 |
| 0.0816 | 2.31E-11 |
| 0.1824 | 1.88E-12 |
| 0.1374 | 0.0001 |
| 0.4779 | 1.70E-06 |
| 0.6364 | 6.97E-13 |
| 0.4295 | 1.15E-12 |
| 0.1232 | 3.56E-15 |
| 0.0476 | 6.44E-06 |
| 0.3885 | 3.04E-07 |
| 0.0340 | 0.17 |
| 0.1933 | 1.12E-08 |
| 0.3691 | 1.01E-14 |
| 0.1269 | 2.11E-11 |
| 0.0191 | 0.11 |
| 0.0737 | 0.07 |
| 1.1117 | 4.29E-15 |
| 0.7404 | 2.99E-08 |
| 0.1493 | 3.28E-06 |
| 0.3257 | 2.57E-14 |
| 0.0800 | 0.0004 |
| 0.1205 | 5.15E-15 |
| 0.0997 | 0.001 |
| 0.2243 | 1.24E-05 |
| 0.1616 | 0.12 |
| 0.0312 | 2.14E-05 |
| 2.4603 | 0.04 |
| 0.7565 | 3.21E-10 |
| 0.1522 | 0.0002 |
| 0.0279 | 0.06 |
| 0.0625 | 0.40 |
| 0.3326 | 1.65E-20 |
| 0.4224 | 6.86E-14 |

| | |
|---|---|
| 0.0219 | 1.40E-08 |
| 0.2292 | 0.01 |
| 0.0332 | 0.002 |
| 0.9562 | 1.58E-15 |
| 0.1001 | 0.001 |
| 0.2864 | 0.0004 |
| 0.6175 | 6.42E-13 |
| 0.1875 | 7.08E-09 |
| 0.2773 | 3.95E-10 |
| 0.2464 | 3.40E-08 |
| 0.1893 | 0.02 |
| 0.2562 | 4.67E-07 |
| 0.1965 | 2.90E-15 |
| 0.1636 | 4.82E-12 |
| 0.3601 | 6.89E-15 |
| 0.1377 | 1.02E-08 |
| 0.0805 | 0.002 |
| 0.0615 | 6.50E-06 |
| 0.1315 | 6.77E-06 |
| 0.1087 | 7.53E-08 |
| 0.0937 | 3.22E-13 |
| 0.1155 | 2.65E-14 |
| 0.3482 | 2.74E-11 |
| 0.2333 | 4.67E-12 |
| 0.0604 | 6.84E-12 |
| 0.1377 | 8.50E-08 |
| 0.1333 | 0.04 |
| 0.1274 | 9.86E-10 |
| 0.1196 | 2.50E-10 |
| 0.1362 | 6.52E-07 |
| 0.4927 | 0.0003 |
| 0.5633 | 1.24E-15 |
| 0.1574 | 4.27E-05 |
| 0.1412 | 2.23E-08 |
| 0.2322 | 5.86E-10 |
| 0.1983 | 0.0003 |
| 0.0691 | 1.61E-05 |
| 0.1679 | 1.77E-12 |
| 0.2927 | 0.0004 |
| 0.0184 | 0.21 |
| 0.2013 | 1.84E-08 |
| 0.0942 | 9.07E-10 |
| 1.1334 | 2.63E-18 |
| 0.4465 | 2.53E-13 |

| | |
|---|---|
| 0.1246 | 6.60E-10 |
| 0.2619 | 1.81E-11 |
| 0.0772 | 3.08E-08 |
| 0.1471 | 4.42E-16 |
| 0.0953 | 9.19E-09 |
| 0.4287 | 2.78E-15 |
| 0.5812 | 5.82E-11 |
| 0.0798 | 6.07E-11 |
| 0.2877 | 9.92E-09 |
| 0.0981 | 2.75E-12 |
| 0.1462 | 8.45E-08 |
| 0.4944 | 3.75E-13 |
| 0.1521 | 0.0001 |
| 0.1407 | 1.02E-10 |
| 0.2891 | 0.001 |
| 0.0409 | 0.04 |
| 0.1881 | 7.77E-19 |
| 0.1152 | 4.35E-09 |
| 0.1008 | 7.46E-06 |
| 0.2873 | 8.34E-05 |
| 0.2386 | 6.39E-07 |
| 0.1358 | 0.0003 |
| 0.1055 | 3.64E-08 |
| 0.1964 | 8.19E-12 |
| 0.1198 | 3.67E-09 |
| 0.1131 | 2.19E-08 |
| 0.1299 | 1.04E-10 |
| 1.5423 | 1.18E-10 |
| 0.0642 | 1.88E-08 |
| 0.5578 | 6.24E-11 |
| 0.1017 | 0.02 |
| 0.1205 | 5.82E-07 |
| 0.0269 | 0.03 |
| 0.1354 | 1.55E-08 |
| 0.3965 | 2.24E-06 |
| 0.2696 | 8.69E-10 |
| 0.1156 | 0.001 |
| 0.4508 | 7.80E-08 |
| 0.0379 | 1.80E-05 |
| 0.1100 | 5.33E-11 |
| 0.0707 | 5.12E-09 |
| 0.1085 | 6.42E-11 |
| 0.0591 | 0.0002 |
| 0.1581 | 0.34 |

| | |
|---|---|
| 0.0620 | 0.02 |
| 0.0806 | 3.90E-05 |
| 0.0451 | 0.39 |
| 0.5532 | 2.11E-12 |
| 0.1660 | 8.24E-12 |
| 0.1466 | 4.18E-12 |
| 0.0718 | 5.13E-10 |
| 0.2269 | 4.42E-05 |
| 0.3914 | 2.85E-05 |
| 0.1041 | 0.27 |
| 0.1944 | 2.62E-09 |
| 0.1285 | 1.80E-10 |
| 0.0000 | 2.21E-05 |
| 0.0651 | 4.15E-06 |
| 0.1603 | 8.04E-05 |
| 0.1317 | 4.57E-24 |
| 0.1677 | 1.62E-08 |
| 0.2352 | 1.24E-07 |
| 0.1153 | 6.58E-10 |
| 0.2187 | 2.38E-05 |
| 0.0537 | 5.66E-06 |
| 0.6030 | 1.92E-07 |
| 0.2392 | 4.93E-07 |
| 0.0947 | 0.01 |
| 0.2375 | 3.30E-10 |
| 0.4076 | 6.39E-09 |
| 0.1378 | 1.64E-24 |
| 0.0890 | 0.005 |
| 0.2854 | 3.77E-06 |
| 0.1484 | 0.0002 |
| 0.5876 | 4.15E-11 |
| 0.1175 | 1.25E-06 |
| 0.1333 | 3.08E-09 |
| 0.0926 | 0.005 |
| 0.1086 | 6.36E-10 |
| 0.1924 | 5.00E-08 |
| 0.0826 | 0.01 |
| 0.0688 | 0.07 |
| 0.4577 | 2.00E-08 |
| 0.0880 | 0.003 |
| 0.2278 | 0.001 |
| 1.1782 | 5.96E-08 |
| 0.1414 | 1.40E-07 |
| 0.0703 | 4.93E-10 |

| | |
|---|---|
| 0.0466 | 6.72E-07 |
| 0.0793 | 5.43E-14 |
| 0.1496 | 0.07 |
| 0.2367 | 5.10E-07 |
| 0.1679 | 3.31E-08 |
| 0.1083 | 5.16E-08 |
| 0.0734 | 0.001 |
| 0.1024 | 8.60E-11 |
| 0.6249 | 1.23E-07 |
| 0.3843 | 4.77E-07 |
| 0.0765 | 4.44E-09 |
| 0.1137 | 0.0002 |
| 0.1026 | 0.005 |
| 0.2391 | 1.30E-07 |
| 0.1628 | 1.91E-06 |
| 0.1062 | 1.64E-07 |
| 0.4402 | 4.59E-07 |
| 0.2415 | 3.24E-06 |
| 0.1326 | 1.62E-11 |
| 0.0736 | 4.59E-06 |
| 0.2571 | 8.55E-09 |
| 0.1233 | 4.51E-07 |
| 0.3472 | 8.08E-06 |
| 0.3107 | 7.52E-08 |
| 0.9132 | 3.95E-08 |
| 0.3083 | 6.79E-12 |
| 0.1601 | 2.95E-07 |
| 0.1138 | 0.02 |
| 0.2093 | 2.80E-18 |
| 0.1813 | 6.28E-10 |
| 0.2207 | 0.004 |
| 0.1438 | 2.95E-06 |
| 0.0914 | 1.63E-05 |
| 0.2459 | 3.92E-08 |
| 0.0977 | 7.19E-07 |
| 0.3168 | 0.0001 |
| 0.2445 | 1.44E-10 |
| 0.0869 | 1.74E-10 |
| 0.0000 | 3.92E-08 |
| 0.1383 | 0.0001 |
| 0.5309 | 1.63E-08 |
| 0.1655 | 1.01E-06 |
| 0.4237 | 1.36E-11 |
| 0.2748 | 8.42E-08 |

| | |
|---|---|
| 0.1029 | 3.92E-08 |
| 0.1041 | 7.39E-08 |
| 0.2635 | 1.25E-18 |
| 0.2773 | 6.36E-14 |
| 0.0675 | 7.46E-09 |
| 0.0606 | 0.24 |
| 0.1959 | 4.93E-07 |
| 0.1373 | 4.70E-13 |
| 0.1909 | 1.97E-06 |
| 0.4305 | 3.78E-12 |
| 0.2152 | 1.32E-09 |
| 0.1679 | 8.28E-05 |
| 0.4406 | 2.58E-05 |
| 0.0482 | 1.80E-07 |
| 0.4192 | 1.93E-16 |
| 0.0537 | 1.47E-09 |
| 0.1686 | 0.01 |
| 0.2186 | 0.005 |
| 0.0950 | 0.0004 |
| 0.1267 | 0.32 |
| 0.0972 | 4.97E-16 |
| 0.2707 | 1.66E-07 |
| 0.1502 | 1.20E-12 |
| 0.0753 | 4.79E-10 |
| 0.1233 | 3.58E-09 |
| 0.1264 | 2.03E-10 |
| 0.1788 | 2.67E-17 |
| 0.7089 | 1.10E-06 |
| 0.2011 | 1.35E-19 |
| 0.1691 | 2.87E-08 |
| 0.1787 | 6.20E-23 |
| 0.0757 | 0.001 |
| 0.6160 | 1.90E-14 |
| 0.6561 | 3.70E-07 |
| 0.1244 | 2.58E-06 |
| 0.0474 | 0.003 |
| 0.0884 | 0.02 |
| 0.9973 | 3.13E-10 |
| 0.0427 | 0.01 |
| 0.1260 | 3.92E-06 |
| 0.3331 | 0.002 |
| 0.1880 | 9.15E-05 |
| 0.1195 | 0.0002 |
| 0.0442 | 8.55E-08 |

| | |
|---|---|
| 0.0824 | 0.003 |
| 0.1542 | 9.32E-05 |
| 0.2461 | 2.44E-06 |
| 0.1375 | 3.34E-08 |
| 0.0671 | 7.31E-25 |
| 0.5584 | 3.42E-16 |
| 0.0464 | 1.90E-06 |
| 0.0000 | 0.0005 |
| 0.4060 | 3.71E-07 |
| 0.6596 | 1.05E-11 |
| 0.2693 | 5.12E-09 |
| 0.2070 | 2.65E-16 |
| 0.1531 | 0.002 |
| 0.4160 | 1.09E-10 |
| 0.1825 | 1.81E-11 |
| 0.0462 | 0.26 |
| 0.1931 | 3.00E-05 |
| 0.0502 | 3.23E-11 |
| 0.0638 | 0.01 |
| 0.0056 | 1.37E-07 |
| 0.1507 | 1.32E-05 |
| 0.1586 | 2.17E-05 |
| 0.2015 | 1.37E-14 |
| 0.0938 | 9.71E-07 |
| 0.0545 | 0.001 |
| 0.2127 | 7.43E-12 |
| 0.0718 | 4.38E-06 |
| 0.6286 | 1.04E-08 |
| 0.2262 | 2.50E-10 |
| 0.1031 | 1.35E-05 |
| 0.1462 | 1.55E-05 |
| 0.1480 | 4.59E-05 |
| 0.0616 | 0.01 |
| 0.0076 | 2.38E-06 |
| 0.2802 | 0.01 |
| 0.2233 | 2.39E-08 |
| 0.1867 | 0.0001 |
| 0.2560 | 9.44E-14 |
| 0.2518 | 1.50E-10 |
| 0.0438 | 6.13E-13 |
| 0.6261 | 0.02 |
| 0.1492 | 3.49E-09 |
| 0.0910 | 0.001 |
| 0.2050 | 0.01 |

| | |
|---|---|
| 0.0202 | 0.03 |
| 0.2054 | 2.79E-11 |
| 0.2751 | 2.10E-06 |
| 0.6431 | 2.12E-05 |
| 0.1567 | 0.0002 |
| 0.1849 | 8.40E-09 |
| 0.1673 | 2.97E-17 |
| 0.1779 | 0.002 |
| 0.0895 | 1.73E-06 |
| 0.1880 | 0.002 |
| 0.4354 | 0.0002 |
| 0.1751 | 7.38E-05 |
| 0.0430 | 5.05E-10 |
| 0.1956 | 9.83E-08 |
| 0.3204 | 2.66E-06 |
| 0.0183 | 1.87E-10 |
| 0.1450 | 6.10E-05 |
| 1.3845 | 1.74E-15 |
| 0.4355 | 6.32E-09 |
| 0.0796 | 2.29E-07 |
| 0.1130 | 0.0002 |
| 0.1367 | 0.0005 |
| 0.0308 | 0.04 |
| 0.2425 | 2.39E-08 |
| 0.0244 | 0.03 |
| 0.4432 | 2.60E-10 |
| 0.1629 | 3.79E-07 |
| 0.2204 | 0.003 |
| 0.0000 | 8.16E-06 |
| 0.1807 | 1.90E-07 |
| 0.1692 | 0.0005 |
| 0.1412 | 2.36E-11 |
| 0.0616 | 1.94E-06 |
| 0.0916 | 1.90E-07 |
| 0.1016 | 0.01 |
| 0.2293 | 0.02 |
| 0.1571 | 2.08E-05 |
| 0.1351 | 2.46E-09 |
| 0.2495 | 9.77E-11 |
| 0.0952 | 0.0002 |
| 0.0924 | 0.0002 |
| 0.4495 | 1.98E-05 |
| 0.1887 | 6.60E-12 |
| 0.1490 | 0.02 |

| | |
|---|---|
| 0.3057 | 1.77E-13 |
| 0.1487 | 2.52E-09 |
| 0.2608 | 0.0005 |
| 0.2240 | 5.44E-07 |
| 0.2133 | 0.001 |
| 0.1526 | 4.54E-07 |
| 0.1210 | 0.0001 |
| 0.5457 | 2.79E-19 |
| 0.0509 | 0.001 |
| 0.1675 | 3.39E-10 |
| 0.1827 | 7.31E-05 |
| 0.2293 | 0.001 |
| 0.0933 | 1.13E-09 |
| 0.0291 | 0.15 |
| 0.1549 | 9.74E-09 |
| 0.5155 | 1.64E-08 |
| 0.1140 | 0.0002 |
| 0.4094 | 1.02E-08 |
| 0.1072 | 8.15E-15 |
| 0.0000 | 2.86E-05 |
| 0.1975 | 1.11E-19 |
| 0.0767 | 9.62E-14 |
| 0.1621 | 0.12 |
| 0.0765 | 0.01 |
| 0.5950 | 9.19E-12 |
| 0.2875 | 2.78E-05 |
| 0.0645 | 0.0001 |
| 0.3062 | 0.0004 |
| 0.4114 | 0.0004 |
| 0.8822 | 4.30E-08 |
| 0.1999 | 1.35E-05 |
| 0.1299 | 5.17E-05 |
| 0.1423 | 0.01 |
| 0.0776 | 5.29E-05 |
| 0.4851 | 3.70E-09 |
| 0.1648 | 1.02E-07 |
| 0.0000 | 0.0005 |
| 0.0000 | 0.01 |
| 0.1046 | 4.36E-05 |
| 0.1312 | 0.002 |
| 0.2846 | 0.005 |
| 0.0185 | 0.01 |
| 0.0596 | 0.58 |
| 0.0729 | 0.003 |

| | |
|---|---|
| 0.1158 | 0.001 |
| 0.4089 | 0.001 |
| 0.3034 | 1.64E-05 |
| 0.4741 | 1.10E-06 |
| 0.8274 | 9.07E-10 |
| 0.1859 | 1.15E-09 |
| 0.5201 | 0.0001 |
| 0.2609 | 1.36E-05 |
| 0.2179 | 7.39E-14 |
| 0.9948 | 3.13E-06 |
| 0.0903 | 1.37E-08 |
| 0.9256 | 8.47E-10 |
| 0.2369 | 4.33E-05 |
| 0.0184 | 0.11 |
| 0.2599 | 1.21E-10 |
| 0.1963 | 1.25E-08 |
| 0.2657 | 0.02 |
| 0.2082 | 1.86E-08 |
| 0.1715 | 4.04E-06 |
| 0.0287 | 1.78E-09 |
| 0.0964 | 5.72E-06 |
| 0.0322 | 0.12 |
| 0.1390 | 8.25E-05 |
| 0.1979 | 1.36E-13 |
| 0.2751 | 0.0001 |
| 0.2463 | 0.52 |
| 0.0476 | 0.34 |
| 1.0223 | 1.95E-06 |
| 0.0654 | 6.11E-06 |
| 0.1144 | 1.70E-06 |
| 0.2042 | 1.84E-09 |
| 0.0819 | 7.13E-09 |
| 0.1622 | 1.38E-05 |
| 0.1405 | 1.06E-05 |
| 0.1782 | 8.24E-12 |
| 0.7734 | 2.87E-13 |
| 0.0309 | 0.07 |
| 0.1213 | 2.67E-07 |
| 0.2103 | 1.61E-06 |
| 0.2062 | 0.11 |
| 0.0699 | 2.45E-07 |
| 0.0334 | 4.88E-09 |
| 0.0974 | 0.001 |
| 1.1581 | 0.001 |

| | |
|---|---|
| 0.1343 | 1.41E-10 |
| 0.1347 | 7.19E-07 |
| 0.5612 | 1.17E-07 |
| 0.1693 | 5.49E-10 |
| 1.0228 | 3.24E-14 |
| 0.1033 | 1.05E-08 |
| 0.2049 | 2.43E-08 |
| 0.1888 | 0.002 |
| 1.0276 | 6.22E-08 |
| 0.1829 | 1.82E-05 |
| 0.1455 | 3.85E-07 |
| 0.0211 | 0.002 |
| 0.1532 | 1.70E-06 |
| 0.2442 | 1.00E-10 |
| 0.1186 | 2.07E-09 |
| 0.2011 | 2.78E-09 |
| 0.0698 | 1.07E-05 |
| 0.1128 | 1.48E-07 |
| 0.2883 | 5.60E-07 |
| 0.0000 | 4.24E-08 |
| 0.0000 | 1.16E-07 |
| 0.3236 | 1.46E-05 |
| 0.1477 | 0.01 |
| 0.1148 | 3.91E-05 |
| 0.0686 | 8.42E-07 |
| 0.0318 | 0.14 |
| 0.3440 | 0.01 |
| 0.4079 | 1.34E-05 |
| 0.1052 | 1.24E-12 |
| 0.1300 | 1.70E-07 |
| 0.0000 | 3.20E-07 |
| 0.2248 | 0.0003 |
| 0.2063 | 5.01E-09 |
| 0.7871 | 1.76E-13 |
| 0.2378 | 2.53E-08 |
| 0.0899 | 0.08 |
| 0.0003 | 4.35E-17 |
| 0.1926 | 1.94E-08 |
| 0.1950 | 1.37E-14 |
| 0.8746 | 3.99E-05 |
| 0.1931 | 1.90E-08 |
| 0.4147 | 0.004 |
| 0.2018 | 5.77E-06 |
| 0.0588 | 6.58E-07 |

| | |
|---|---|
| 0.5634 | 6.07E-10 |
| 0.1929 | 4.22E-19 |
| 0.1589 | 8.30E-06 |
| 0.1593 | 1.30E-06 |
| 0.0448 | 0.29 |
| 0.0450 | 0.0005 |
| 0.0281 | 9.32E-07 |
| 0.2598 | 4.47E-07 |
| 0.1650 | 3.33E-13 |
| 0.1880 | 7.86E-13 |
| 0.3950 | 1.83E-05 |
| 0.1145 | 7.22E-06 |
| 0.1600 | 0.0003 |
| 0.2378 | 0.002 |
| 0.2181 | 2.41E-05 |
| 0.1471 | 8.91E-08 |
| 0.2291 | 0.0002 |
| 0.0059 | 3.02E-05 |
| 0.1627 | 6.46E-09 |
| 0.0000 | 0.06 |
| 0.0000 | 1.84E-11 |
| 0.2575 | 7.39E-15 |
| 0.0768 | 3.37E-10 |
| 0.2804 | 9.42E-08 |
| 0.0000 | 3.32E-06 |
| 0.2984 | 0.001 |
| 0.6294 | 3.79E-07 |
| 0.6077 | 7.77E-11 |
| 0.2086 | 0.0005 |
| 0.3161 | 5.29E-05 |
| 0.1940 | 7.57E-05 |
| 0.1369 | 8.72E-09 |
| 0.0792 | 8.42E-06 |
| 0.6291 | 5.48E-06 |
| 0.1838 | 6.50E-06 |
| 0.1251 | 6.72E-09 |
| 0.0815 | 0.04 |
| 0.0995 | 8.26E-15 |
| 0.1428 | 9.38E-14 |
| 0.2036 | 3.57E-07 |
| 0.1375 | 0.0004 |
| 0.0945 | 0.09 |
| 0.2083 | 5.45E-05 |
| 0.1901 | 7.17E-06 |

| | |
|---|---|
| 0.1813 | 1.67E-06 |
| 0.1557 | 1.22E-07 |
| 0.1811 | 0.002 |
| 0.1951 | 1.49E-10 |
| 0.2793 | 0.0001 |
| 0.1260 | 6.14E-06 |
| 0.1238 | 0.004 |
| 0.2032 | 1.75E-06 |
| 0.0998 | 0.07 |
| 0.1255 | 8.22E-10 |
| 0.0943 | 1.70E-08 |
| 0.1613 | 0.10 |
| 0.0000 | 0.02 |
| 0.2618 | 3.93E-06 |
| 0.0049 | 0.01 |
| 0.0000 | 2.22E-13 |
| 0.0475 | 1.82E-06 |
| 0.1070 | 0.01 |
| 0.2880 | 0.004 |
| 0.1719 | 1.01E-06 |
| 0.1061 | 0.002 |
| 0.0000 | 2.16E-11 |
| 0.2631 | 8.94E-09 |
| 0.0609 | 0.24 |
| 0.0335 | 0.01 |
| 0.0471 | 0.13 |
| 0.0260 | 0.001 |
| 0.1068 | 0.0002 |
| 0.1404 | 1.78E-08 |
| 0.4642 | 2.13E-08 |
| 0.0996 | 2.90E-05 |
| 0.1028 | 2.73E-09 |
| 0.2186 | 0.0001 |
| 0.0707 | 0.04 |
| 0.0662 | 1.27E-05 |
| 0.0010 | 5.90E-10 |
| 0.0703 | 0.0003 |
| 0.0333 | 0.0002 |
| 0.0415 | 2.26E-06 |
| 0.2327 | 1.32E-07 |
| 0.1398 | 0.003 |
| 0.0450 | 0.001 |
| 1.2720 | 8.88E-09 |
| 0.1545 | 1.44E-07 |

| | |
|---|---|
| 0.2149 | 1.24E-11 |
| 0.1247 | 0.0001 |
| 0.1190 | 4.06E-05 |
| 0.1424 | 0.0003 |
| 0.1369 | 1.00E-09 |
| 0.2513 | 2.22E-10 |
| 0.2291 | 0.0003 |
| 0.6921 | 2.63E-07 |
| 0.0788 | 0.003 |
| 0.1104 | 0.001 |
| 0.1626 | 0.001 |
| 0.1161 | 6.59E-06 |
| 0.0000 | 0.02 |
| 0.0859 | 0.0002 |
| 0.1302 | 7.40E-05 |
| 0.3230 | 0.02 |
| 0.1720 | 4.33E-05 |
| 0.0521 | 0.0005 |
| 0.1061 | 0.0001 |
| 0.0158 | 3.32E-05 |
| 0.1749 | 3.67E-05 |
| 0.0793 | 0.0004 |
| 0.3158 | 4.46E-07 |
| 0.0904 | 0.001 |
| 0.0891 | 1.08E-06 |
| 0.1191 | 0.02 |
| 0.1378 | 0.0002 |
| 0.3538 | 8.80E-13 |
| 0.0670 | 0.01 |
| 0.5570 | 1.11E-06 |
| 0.1822 | 0.0004 |
| 0.1699 | 2.66E-06 |
| 0.1042 | 0.01 |
| 0.1395 | 0.0001 |
| 0.1825 | 1.19E-07 |
| 0.5192 | 6.17E-10 |
| 0.1143 | 3.86E-06 |
| 0.7182 | 0.002 |
| 0.0987 | 8.55E-08 |
| 0.2004 | 1.79E-06 |
| 0.4149 | 0.0002 |
| 0.0762 | 0.0002 |
| 0.9345 | 0.0003 |
| 0.1168 | 0.0002 |

| | |
|---|---|
| 0.3436 | 6.24E-11 |
| 0.1507 | 0.0001 |
| 0.1069 | 0.002 |
| 0.2566 | 4.42E-06 |
| 0.1638 | 7.84E-06 |
| 0.6283 | 6.35E-07 |
| 0.0000 | 1.26E-12 |
| 0.1765 | 0.01 |
| 0.1085 | 1.04E-08 |
| 0.0000 | 0.0002 |
| 0.1073 | 0.001 |
| 0.0491 | 6.11E-11 |
| 0.1217 | 8.12E-09 |
| 0.2846 | 1.01E-05 |
| 0.0000 | 0.01 |
| 0.0987 | 0.16 |
| 0.0920 | 3.00E-06 |
| 0.1803 | 0.002 |
| 0.2733 | 3.59E-06 |
| 0.0953 | 0.45 |
| 0.6561 | 2.66E-08 |
| 0.1761 | 0.002 |
| 0.0784 | 0.27 |
| 0.0827 | 0.01 |
| 0.1910 | 0.005 |
| 0.0918 | 3.30E-05 |
| 0.1319 | 0.002 |
| 0.1434 | 0.003 |
| 0.4575 | 1.70E-05 |
| 0.1289 | 3.75E-05 |
| 0.1133 | 6.47E-07 |
| 0.0000 | 0.0002 |
| 0.2313 | 1.80E-12 |
| 0.0715 | 0.001 |
| 0.0596 | 0.003 |
| 0.3172 | 0.005 |
| 0.1395 | 1.32E-06 |
| 0.1960 | 0.003 |
| 0.2743 | 0.01 |
| 0.3938 | 0.0005 |
| 0.0700 | 1.43E-05 |
| 0.3293 | 6.84E-06 |
| 0.0000 | 8.14E-05 |
| 0.0551 | 0.02 |

| | |
|---|---|
| 0.5033 | 0.005 |
| 0.4565 | 9.40E-07 |
| 0.0625 | 0.01 |
| 0.1028 | 0.03 |
| 0.1823 | 4.68E-11 |
| 0.0261 | 0.005 |
| 0.0824 | 0.0003 |
| 0.0476 | 5.33E-10 |
| 0.1035 | 3.38E-06 |
| 2.2724 | 0.12 |
| 0.2426 | 5.68E-06 |
| 0.1298 | 4.70E-05 |
| 0.5096 | 8.42E-06 |
| 0.0866 | 3.61E-06 |
| 0.0000 | 5.52E-08 |
| 0.0000 | 6.23E-06 |
| 0.1992 | 0.002 |
| 0.2139 | 0.0001 |
| 0.1266 | 0.001 |
| 0.0740 | 0.02 |
| 0.2478 | 4.82E-06 |
| 0.0484 | 3.53E-09 |
| 0.0013 | 1.05E-07 |
| 0.0000 | 0.0002 |
| 0.1755 | 0.0004 |
| 0.0993 | 0.0001 |
| 0.1936 | 0.001 |
| 0.4371 | 0.15 |
| 0.1420 | 0.0003 |
| 0.0225 | 0.05 |
| 0.0000 | 0.0002 |
| 0.1124 | 2.66E-06 |
| 0.0802 | 0.003 |
| 0.0567 | 1.99E-06 |
| 0.1841 | 0.0005 |
| 48.6159 | 0.98 |
| 0.3675 | 0.02 |
| 0.0246 | 0.002 |
| 0.5968 | 5.44E-05 |
| 0.3711 | 1.95E-10 |
| 0.3313 | 0.01 |
| 0.2718 | 0.0001 |
| 2.0034 | 0.02 |
| 0.1433 | 0.001 |

| | |
|---|---|
| 0.0514 | 0.02 |
| 0.0959 | 0.15 |
| 0.2016 | 0.0002 |
| 0.1724 | 7.43E-06 |
| 0.0560 | 0.02 |
| 0.0000 | 7.20E-08 |
| 0.0604 | 0.02 |
| 0.0377 | 0.01 |
| 0.0254 | 0.0002 |
| 0.3441 | 0.0001 |
| 0.0733 | 0.22 |
| 2.6826 | 0.02 |
| 0.2762 | 0.0001 |
| 0.1889 | 0.50 |
| 0.6400 | 1.68E-05 |
| 0.1907 | 1.25E-06 |
| 0.3027 | 1.60E-07 |
| 0.3492 | 0.46 |
| 0.1011 | 0.02 |
| 0.0466 | 0.02 |
| 0.0936 | 1.83E-07 |
| 0.0000 | 0.005 |
| 0.1605 | 3.79E-06 |
| 0.1297 | 7.33E-07 |
| 0.7776 | 1.31E-05 |
| 0.2468 | 0.01 |
| 0.0964 | 0.02 |
| 0.0403 | 0.04 |
| 0.2652 | 2.28E-05 |
| 0.0380 | 0.03 |
| 0.1898 | 0.01 |
| 0.1024 | 1.77E-06 |
| 0.6512 | 0.11 |
| 0.0992 | 0.01 |
| 0.1448 | 0.0001 |
| 0.0670 | 0.0002 |
| 0.1119 | 1.48E-06 |
| 0.1768 | 4.42E-09 |
| 0.0000 | 0.56 |
| 0.0592 | 0.02 |
| 0.2745 | 3.46E-07 |
| 0.5592 | 0.06 |
| 0.0639 | 0.20 |
| 0.1212 | 0.03 |

| | |
|---|---|
| 0.0567 | 0.02 |

**Supplementary Table 22.** codeml results - w for dermic parasites/other species

| ID | Likelihood | ω other species | dN | dS | ω dermic | dN | dS |
|---|---|---|---|---|---|---|---|
| cluster1127 | -1409.04882 | 0.0553 | 0.1400 | 2.5315 | 0.0600 | 0.0451 | 0.7518 |
| cluster2853 | -22473.62650 | 0.0258 | 0.0320 | 1.2403 | 0.0307 | 0.0311 | 1.0135 |
| cluster2854 | -1615.32284 | 0.0371 | 0.0196 | 0.5285 | 0.0328 | 0.0069 | 0.2093 |
| cluster2855 | -23551.90658 | 0.0176 | 0.0538 | 3.0547 | 0.0242 | 0.0149 | 0.6161 |
| cluster2856 | -15420.96309 | 0.0919 | 0.2123 | 2.3116 | 0.1391 | 0.0731 | 0.5256 |
| cluster2857 | -5977.81050 | 0.0027 | 0.0022 | 0.7953 | 0.0529 | 0.0058 | 0.1094 |
| cluster2858 | -7755.51270 | 0.0043 | 0.0061 | 1.4253 | 0.0032 | 0.0013 | 0.3987 |
| cluster2859 | -5673.66765 | 0.0311 | 0.0208 | 0.6675 | 0.0645 | 0.0109 | 0.1684 |
| cluster2860 | -16040.51960 | 0.0026 | 0.0024 | 0.9342 | 0.0009 | 0.0006 | 0.6588 |
| cluster2861 | -8370.11503 | 0.0073 | 0.0092 | 1.2727 | 0.0149 | 0.0084 | 0.5594 |
| cluster2862 | -18857.55791 | 0.0148 | 0.0130 | 0.8835 | 0.0157 | 0.0136 | 0.8689 |
| cluster2863 | -15073.42285 | 0.0277 | 0.0514 | 1.8559 | 0.0364 | 0.0193 | 0.5291 |
| cluster2864 | -12548.54929 | 0.0050 | 0.0050 | 1.0125 | 0.0025 | 0.0026 | 1.0100 |
| cluster2865 | -15004.90358 | 0.1012 | 0.0603 | 0.5958 | 0.0952 | 0.0494 | 0.5187 |
| cluster2866 | -17311.14704 | 0.0246 | 0.0596 | 2.4241 | 0.0310 | 0.0149 | 0.4812 |
| cluster2867 | -14849.34706 | 0.0353 | 0.0730 | 2.0681 | 0.0730 | 0.0383 | 0.5240 |
| cluster2869 | -8915.43303 | 0.0819 | 0.0846 | 1.0327 | 0.1056 | 0.0296 | 0.2799 |
| cluster2870 | -3880.55240 | 0.0488 | 0.0542 | 1.1101 | 0.0635 | 0.0217 | 0.3414 |
| cluster2871 | -3139.48795 | 0.0227 | 0.0101 | 0.4449 | 0.0316 | 0.0065 | 0.2055 |
| cluster2872 | -12260.94647 | 0.0834 | 0.1326 | 1.5902 | 0.0831 | 0.0453 | 0.5454 |
| cluster2873 | -13851.48632 | 0.0174 | 0.0141 | 0.8108 | 0.0135 | 0.0094 | 0.6959 |
| cluster2874 | -10381.06761 | 0.0295 | 0.0292 | 0.9892 | 0.0530 | 0.0159 | 0.3009 |
| cluster2875 | -13636.38464 | 0.0177 | 0.0273 | 1.5476 | 0.0143 | 0.0153 | 1.0710 |
| cluster2876 | -3970.35841 | 0.0432 | 0.0289 | 0.6699 | 0.0272 | 0.0204 | 0.7488 |
| cluster2877 | -10636.95406 | 0.0467 | 0.0465 | 0.9948 | 0.0577 | 0.0307 | 0.5317 |
| cluster2878 | -7056.59855 | 0.0491 | 0.0223 | 0.4541 | 0.0294 | 0.0085 | 0.2905 |
| cluster2879 | -10035.35655 | 0.1742 | 0.0791 | 0.4542 | 0.2106 | 0.0479 | 0.2274 |
| cluster2880 | -10642.17146 | 0.0476 | 0.0916 | 1.9239 | 0.0522 | 0.0288 | 0.5520 |
| cluster2881 | -11523.15648 | 0.0500 | 0.0329 | 0.6569 | 0.0541 | 0.0437 | 0.8081 |
| cluster2882 | -10647.56303 | 0.0261 | 0.0387 | 1.4819 | 0.0364 | 0.0208 | 0.5708 |
| cluster2883 | -6764.08154 | 0.0799 | 0.0407 | 0.5099 | 0.0625 | 0.0370 | 0.5908 |
| cluster2884 | -12070.44795 | 0.0239 | 0.0234 | 0.9824 | 0.0299 | 0.0091 | 0.3037 |
| cluster2885 | -15732.13007 | 0.0334 | 0.0869 | 2.5995 | 0.0575 | 0.0333 | 0.5799 |
| cluster2886 | -12003.78043 | 0.0442 | 0.0275 | 0.6235 | 0.0356 | 0.0237 | 0.6656 |
| cluster2887 | -8362.70402 | 0.0028 | 0.0117 | 4.1390 | 0.0100 | 0.0050 | 0.5062 |
| cluster2888 | -7783.36882 | 0.0511 | 0.1237 | 2.4205 | 0.0806 | 0.0631 | 0.7832 |
| cluster2889 | -6930.65486 | 0.0077 | 0.0153 | 1.9872 | 0.0100 | 0.0055 | 0.5489 |
| cluster2890 | -2680.657563 | 0.0713 | 0.0082 | 0.1151 | 0.1003 | 0.0166 | 0.1658 |
| cluster2891 | -12095.00807 | 0.0311 | 0.0775 | 2.4880 | 0.0465 | 0.0238 | 0.5123 |
| cluster2892 | -7912.72031 | 0.0130 | 0.0086 | 0.6613 | 0.0166 | 0.0112 | 0.6746 |
| cluster2893 | -8580.60264 | 0.0339 | 0.0746 | 2.2021 | 0.0403 | 0.0284 | 0.7052 |
| cluster2894 | -5509.36630 | 0.0051 | 0.0243 | 4.7347 | 0.0193 | 0.0063 | 0.3235 |

| | | | | | | | |
|---|---|---|---|---|---|---|---|
| cluster2895 | -8448.55871 | 0.0336 | 0.0661 | 1.9647 | 0.0394 | 0.0205 | 0.5207 |
| cluster2896 | -10286.29221 | 0.0089 | 0.0185 | 2.0883 | 0.0125 | 0.0082 | 0.6529 |
| cluster2897 | -9157.52134 | 0.0302 | 0.0370 | 1.2238 | 0.0414 | 0.0144 | 0.3481 |
| cluster2898 | -4501.02855 | 0.0249 | 0.0156 | 0.6289 | 0.0276 | 0.0136 | 0.4925 |
| cluster2899 | -8006.61107 | 0.0173 | 0.0216 | 1.2512 | 0.0209 | 0.0255 | 1.2173 |
| cluster2900 | -9923.86854 | 0.0141 | 0.0086 | 0.6079 | 0.0121 | 0.0085 | 0.7056 |
| cluster2901 | -9241.04896 | 0.0383 | 0.0765 | 1.9987 | 0.0359 | 0.0189 | 0.5260 |
| cluster2902 | -3696.66965 | 0.0274 | 0.0529 | 1.9313 | 0.0252 | 0.0150 | 0.5962 |
| cluster2903 | -8864.10097 | 0.0099 | 0.0077 | 0.7739 | 0.0109 | 0.0113 | 1.0332 |
| cluster2904 | -6403.00097 | 0.0529 | 0.0244 | 0.4608 | 0.1365 | 0.0650 | 0.4763 |
| cluster2905 | -8874.64054 | 0.0203 | 0.0101 | 0.4982 | 0.0381 | 0.0173 | 0.4526 |
| cluster2906 | -13667.91923 | 0.0726 | 0.0380 | 0.5233 | 0.0740 | 0.0330 | 0.4457 |
| cluster2907 | -12733.33766 | 0.0580 | 0.1474 | 2.5421 | 0.0940 | 0.0569 | 0.6056 |
| cluster2908 | -5645.65731 | 0.0128 | 0.0072 | 0.5590 | 0.0117 | 0.0024 | 0.2058 |
| cluster2909 | -2924.80584 | 0.0335 | 0.0185 | 0.5531 | 0.0467 | 0.0085 | 0.1813 |
| cluster2910 | -9271.21778 | 0.0415 | 0.0551 | 1.3294 | 0.0464 | 0.0209 | 0.4507 |
| cluster2911 | -4006.28657 | 0.0071 | 0.0022 | 0.3052 | 0.0010 | 0.0002 | 0.1946 |
| cluster2912 | -8762.62883 | 0.0202 | 0.0444 | 2.1965 | 0.0368 | 0.0192 | 0.5207 |
| cluster2913 | -6297.30199 | 0.0031 | 0.0040 | 1.2792 | 0.0017 | 0.0007 | 0.4047 |
| cluster2914 | -8063.42014 | 0.0141 | 0.0078 | 0.5527 | 0.0178 | 0.0087 | 0.4879 |
| cluster2915 | -4206.82416 | 0.0149 | 0.0205 | 1.3795 | 0.0200 | 0.0102 | 0.5090 |
| cluster2916 | -9770.40742 | 0.0303 | 0.0211 | 0.6981 | 0.0330 | 0.0215 | 0.6513 |
| cluster2917 | -9025.45150 | 0.0235 | 0.0366 | 1.5596 | 0.0164 | 0.0079 | 0.4811 |
| cluster2918 | -4474.43706 | 0.0632 | 0.0404 | 0.6405 | 0.0868 | 0.0260 | 0.2995 |
| cluster2919 | -9564.68784 | 0.0720 | 0.0385 | 0.5339 | 0.0620 | 0.0281 | 0.4527 |
| cluster2920 | -6956.88136 | 0.0138 | 0.0441 | 3.1895 | 0.0177 | 0.0139 | 0.7888 |
| cluster2921 | -3808.28907 | 0.0031 | 0.0056 | 1.8275 | 0.0056 | 0.0034 | 0.6032 |
| cluster2922 | -6243.87308 | 0.0507 | 0.0973 | 1.9181 | 0.0641 | 0.0328 | 0.5125 |
| cluster2923 | -5849.44785 | 0.0697 | 0.1435 | 2.0597 | 0.3043 | 0.0967 | 0.3178 |
| cluster2924 | -10970.66338 | 0.0155 | 0.0445 | 2.8736 | 0.0294 | 0.0172 | 0.5859 |
| cluster2925 | -5771.60047 | 0.0337 | 0.0573 | 1.7013 | 0.0327 | 0.0194 | 0.5938 |
| cluster2926 | -11219.97818 | 0.0219 | 0.0192 | 0.8772 | 0.0206 | 0.0173 | 0.8415 |
| cluster2927 | -4902.94126 | 0.0258 | 0.0596 | 2.3111 | 0.0389 | 0.0274 | 0.7047 |
| cluster2928 | -7180.71873 | 0.0202 | 0.0375 | 1.8533 | 0.0237 | 0.0121 | 0.5108 |
| cluster2929 | -8011.25880 | 0.0056 | 0.0087 | 1.5372 | 0.0060 | 0.0030 | 0.5016 |
| cluster2930 | -10989.01477 | 0.0846 | 0.0938 | 1.1091 | 0.0841 | 0.0268 | 0.3189 |
| cluster2931 | -6864.15970 | 0.0429 | 0.1067 | 2.4873 | 0.0644 | 0.0433 | 0.6722 |
| cluster2932 | -9687.06215 | 0.0476 | 0.0812 | 1.7065 | 0.0805 | 0.0439 | 0.5448 |
| cluster2933 | -8733.59755 | 0.0582 | 0.0238 | 0.4082 | 0.0374 | 0.0168 | 0.4500 |
| cluster2934 | -10531.72447 | 0.0650 | 0.0546 | 0.8398 | 0.0610 | 0.0164 | 0.2695 |
| cluster2935 | -4952.31198 | 0.0304 | 0.0221 | 0.7256 | 0.0262 | 0.0184 | 0.7039 |
| cluster2936 | -9505.97460 | 0.0280 | 0.0805 | 2.8780 | 0.0497 | 0.0314 | 0.6326 |
| cluster2937 | -4595.66379 | 0.0276 | 0.0156 | 0.5663 | 0.0281 | 0.0121 | 0.4307 |
| cluster2938 | -7240.55010 | 0.0177 | 0.0149 | 0.8448 | 0.0128 | 0.0070 | 0.5457 |

| | | | | | | | |
|---|---|---|---|---|---|---|---|
| cluster2939 | -2817.41286 | 0.0277 | 0.0104 | 0.3733 | 0.0190 | 0.0058 | 0.3039 |
| cluster2940 | -11375.80355 | 0.0583 | 0.0954 | 1.6361 | 0.0785 | 0.0337 | 0.4299 |
| cluster2941 | -3029.46391 | 0.0001 | 0.0000 | 0.2025 | 0.0037 | 0.0006 | 0.1614 |
| cluster2942 | -11359.32770 | 0.0516 | 0.0966 | 1.8705 | 0.0570 | 0.0316 | 0.5541 |
| cluster2943 | -4040.38368 | 0.0114 | 0.0115 | 1.0037 | 0.0101 | 0.0073 | 0.7215 |
| cluster2944 | -8491.05443 | 0.0817 | 0.0747 | 0.9146 | 0.1284 | 0.0910 | 0.7082 |
| cluster2945 | -3084.20151 | 0.0017 | 0.0073 | 4.2897 | 0.0036 | 0.0023 | 0.6516 |
| cluster2946 | -6740.48881 | 0.0061 | 0.0021 | 0.3410 | 0.0128 | 0.0039 | 0.3071 |
| cluster2947 | -9659.94851 | 0.0362 | 0.0315 | 0.8694 | 0.0386 | 0.0118 | 0.3071 |
| cluster2948 | -9376.63723 | 0.0500 | 0.0443 | 0.8861 | 0.0566 | 0.0372 | 0.6569 |
| cluster2949 | -6815.81660 | 0.0158 | 0.0100 | 0.6376 | 0.0189 | 0.0097 | 0.5120 |
| cluster2950 | -7085.57463 | 0.0042 | 0.0029 | 0.6795 | 0.0315 | 0.0204 | 0.6491 |
| cluster2951 | -9094.13038 | 0.0683 | 0.1135 | 1.6635 | 0.0575 | 0.0417 | 0.7259 |
| cluster2952 | -10893.21002 | 0.0363 | 0.0701 | 1.9339 | 0.0510 | 0.0245 | 0.4812 |
| cluster2953 | -13285.88099 | 0.0560 | 0.1222 | 2.1834 | 0.0848 | 0.0576 | 0.6796 |
| cluster2954 | -7273.73604 | 0.0178 | 0.0234 | 1.3157 | 0.0216 | 0.0234 | 1.0829 |
| cluster2955 | -6029.65422 | 0.0270 | 0.0374 | 1.3856 | 0.0207 | 0.0072 | 0.3476 |
| cluster2956 | -10134.29992 | 0.0144 | 0.0164 | 1.1365 | 0.0123 | 0.0374 | 3.0361 |
| cluster2957 | -8385.40344 | 0.0778 | 0.1509 | 1.9388 | 0.1136 | 0.0519 | 0.4575 |
| cluster2958 | -9691.17231 | 0.0113 | 0.0183 | 1.6149 | 0.0129 | 0.0066 | 0.5128 |
| cluster2959 | -5169.46503 | 0.0537 | 0.0302 | 0.5625 | 0.0519 | 0.0303 | 0.5839 |
| cluster2960 | -9441.51617 | 0.0210 | 0.0191 | 0.9091 | 0.0263 | 0.0208 | 0.7916 |
| cluster2961 | -8397.96983 | 0.0104 | 0.0299 | 2.8666 | 0.0087 | 0.0127 | 1.4549 |
| cluster2962 | -7495.41991 | 0.0207 | 0.0350 | 1.6927 | 0.0237 | 0.0348 | 1.4665 |
| cluster2963 | -7223.90128 | 0.0361 | 0.0159 | 0.4414 | 0.0355 | 0.0111 | 0.3118 |
| cluster2964 | -5247.15403 | 0.0250 | 0.0125 | 0.5021 | 0.0389 | 0.0168 | 0.4312 |
| cluster2965 | -7390.99417 | 0.0126 | 0.0152 | 1.2106 | 0.0091 | 0.0039 | 0.4296 |
| cluster2966 | -5587.44209 | 0.0206 | 0.0489 | 2.3805 | 0.0253 | 0.0187 | 0.7385 |
| cluster2967 | -9578.73464 | 0.0191 | 0.0118 | 0.6159 | 0.0151 | 0.0084 | 0.5580 |
| cluster2968 | -5456.41994 | 0.0164 | 0.0170 | 1.0396 | 0.0210 | 0.0108 | 0.5129 |
| cluster2969 | -7910.44485 | 0.0443 | 0.0631 | 1.4230 | 0.0400 | 0.0137 | 0.3432 |
| cluster2970 | -4235.92651 | 0.0031 | 0.0072 | 2.2851 | 0.0048 | 0.0022 | 0.4692 |
| cluster2971 | -3608.78171 | 0.0294 | 0.0110 | 0.3741 | 0.0339 | 0.0052 | 0.1532 |
| cluster2972 | -2669.98014 | 0.0054 | 0.0035 | 0.6552 | 0.0070 | 0.0049 | 0.7059 |
| cluster2973 | -6352.89607 | 0.0650 | 0.1297 | 1.9947 | 0.1333 | 0.0482 | 0.3617 |
| cluster2974 | -4388.61493 | 0.0080 | 0.0057 | 0.7105 | 0.0727 | 0.0241 | 0.3320 |
| cluster2975 | -10301.69145 | 0.0668 | 0.1096 | 1.6401 | 0.0771 | 0.0381 | 0.4943 |
| cluster2976 | -1468.182266 | 0.1481 | 0.0409 | 0.2759 | 0.2695 | 0.1084 | 0.4024 |
| cluster2977 | -3392.83520 | 0.0043 | 0.0155 | 3.5861 | 0.0055 | 0.0052 | 0.9421 |
| cluster2978 | -8210.46880 | 0.0025 | 0.0018 | 0.6897 | 0.0003 | 0.0002 | 0.6729 |
| cluster2979 | -8838.63586 | 0.0355 | 0.0418 | 1.1779 | 0.0459 | 0.0177 | 0.3869 |
| cluster2980 | -1553.72411 | 0.0180 | 0.0673 | 3.7401 | 0.0268 | 0.0331 | 1.2372 |
| cluster2981 | -6207.41963 | 0.0423 | 0.0697 | 1.6501 | 0.0783 | 0.0330 | 0.4221 |
| cluster2982 | -8044.42799 | 0.0722 | 0.0517 | 0.7165 | 0.1553 | 0.0441 | 0.2841 |

| | | | | | | | |
|---|---|---|---|---|---|---|---|
| cluster2983 | -7517.86134 | 0.0194 | 0.0519 | 2.6781 | 0.0294 | 0.0172 | 0.5852 |
| cluster2984 | -4831.84088 | 0.0328 | 0.0145 | 0.4427 | 0.0310 | 0.0134 | 0.4325 |
| cluster2985 | -3971.478666 | 0.2794 | 0.1067 | 0.3819 | 0.2032 | 0.0656 | 0.3228 |
| cluster2986 | -3495.66795 | 0.0920 | 0.0938 | 1.0192 | 0.1384 | 0.0258 | 0.1867 |
| cluster2987 | -6077.19005 | 0.0264 | 0.0189 | 0.7151 | 0.0346 | 0.0142 | 0.4102 |
| cluster2988 | -5702.97980 | 0.0222 | 0.0430 | 1.9357 | 0.0274 | 0.0130 | 0.4729 |
| cluster2989 | -6588.32343 | 0.0458 | 0.0188 | 0.4103 | 0.0327 | 0.0143 | 0.4373 |
| cluster2990 | -7511.83869 | 0.0425 | 0.0651 | 1.5302 | 0.0468 | 0.0260 | 0.5546 |
| cluster2991 | -6301.48926 | 0.0390 | 0.1118 | 2.8661 | 0.0853 | 0.0404 | 0.4735 |
| cluster2992 | -4177.93860 | 0.0078 | 0.0037 | 0.4681 | 0.0141 | 0.0039 | 0.2760 |
| cluster2993 | -5420.98681 | 0.0420 | 0.0269 | 0.6394 | 0.0255 | 0.0222 | 0.8734 |
| cluster2994 | -2259.79342 | 0.0028 | 0.0022 | 0.7900 | 0.0057 | 0.0015 | 0.2566 |
| cluster2995 | -740.116173 | 0.0230 | 0.0423 | 1.8355 | 0.0156 | 0.0056 | 0.3617 |
| cluster2996 | -7310.92308 | 0.0425 | 0.0263 | 0.6179 | 0.0439 | 0.0279 | 0.6354 |
| cluster2997 | -8170.65906 | 0.0360 | 0.0494 | 1.3723 | 0.0468 | 0.0186 | 0.3973 |
| cluster2998 | -7504.17728 | 0.0186 | 0.0349 | 1.8796 | 0.0191 | 0.0091 | 0.4762 |
| cluster2999 | -6764.13473 | 0.0297 | 0.0708 | 2.3845 | 0.0320 | 0.0181 | 0.5664 |
| cluster3000 | -6334.38409 | 0.0110 | 0.0095 | 0.8622 | 0.0137 | 0.0027 | 0.1944 |
| cluster3001 | -6053.26815 | 0.0149 | 0.0155 | 1.0403 | 0.0323 | 0.0116 | 0.3593 |
| cluster3002 | -7669.41678 | 0.0260 | 0.0253 | 0.9755 | 0.0273 | 0.0103 | 0.3769 |
| cluster3003 | -7808.98207 | 0.0787 | 0.0668 | 0.8489 | 0.1070 | 0.0274 | 0.2562 |
| cluster3004 | -5965.15977 | 0.0991 | 0.0280 | 0.2829 | 0.0900 | 0.0392 | 0.4358 |
| cluster3005 | -4102.90399 | 0.0586 | 0.0304 | 0.5189 | 0.0587 | 0.0076 | 0.1297 |
| cluster3006 | -8671.98360 | 0.0584 | 0.0542 | 0.9278 | 0.0558 | 0.0388 | 0.6962 |
| cluster3007 | -8243.99526 | 0.0260 | 0.0756 | 2.9056 | 0.0277 | 0.0186 | 0.6701 |
| cluster3008 | -5950.18731 | 0.0251 | 0.0361 | 1.4410 | 0.0442 | 0.0251 | 0.5675 |
| cluster3009 | -5913.05732 | 0.0406 | 0.0347 | 0.8546 | 0.0476 | 0.0134 | 0.2826 |
| cluster3010 | -8052.03560 | 0.1199 | 0.0995 | 0.8304 | 0.1924 | 0.0464 | 0.2409 |
| cluster3011 | -3371.72568 | 0.0079 | 0.0132 | 1.6732 | 0.0112 | 0.0156 | 1.3948 |
| cluster3012 | -5210.02012 | 0.0624 | 0.1216 | 1.9490 | 0.0545 | 0.0353 | 0.6488 |
| cluster3013 | -6735.16817 | 0.0815 | 0.0376 | 0.4608 | 0.0836 | 0.0254 | 0.3033 |
| cluster3014 | -3557.15835 | 0.0115 | 0.0221 | 1.9171 | 0.0032 | 0.0018 | 0.5753 |
| cluster3015 | -2410.35004 | 0.0383 | 0.0239 | 0.6257 | 0.0472 | 0.0102 | 0.2159 |
| cluster3016 | -4556.14556 | 0.0366 | 0.0249 | 0.6813 | 0.0436 | 0.0115 | 0.2649 |
| cluster3017 | -5968.77168 | 0.0185 | 0.0698 | 3.7781 | 0.0367 | 0.0203 | 0.5533 |
| cluster3018 | -5679.15398 | 0.0068 | 0.0098 | 1.4489 | 0.0062 | 0.0031 | 0.5016 |
| cluster3019 | -7569.33485 | 0.0435 | 0.0753 | 1.7302 | 0.0613 | 0.0445 | 0.7259 |
| cluster3020 | -6186.88545 | 0.0240 | 0.0344 | 1.4358 | 0.0547 | 0.0170 | 0.3103 |
| cluster3021 | -3024.77350 | 0.0180 | 0.0086 | 0.4763 | 0.0153 | 0.0070 | 0.4570 |
| cluster3022 | -3430.12440 | 0.0440 | 0.1377 | 3.1278 | 0.0738 | 0.0348 | 0.4717 |
| cluster3023 | -4204.56735 | 0.0108 | 0.0060 | 0.5545 | 0.0093 | 0.0052 | 0.5624 |
| cluster3024 | -1957.94559 | 0.0688 | 0.0403 | 0.5861 | 0.1278 | 0.0193 | 0.1507 |
| cluster3025 | -4691.58964 | 0.0089 | 0.0047 | 0.5258 | 0.0118 | 0.0028 | 0.2350 |
| cluster3026 | -4110.746711 | 0.1571 | 0.1678 | 1.0684 | 0.1939 | 0.0473 | 0.2437 |

| | | | | | | | |
|---|---|---|---|---|---|---|---|
| cluster3027 | -3436.61804 | 0.0022 | 0.0021 | 0.9516 | 0.0024 | 0.0020 | 0.8325 |
| cluster3028 | -6800.11352 | 0.0305 | 0.0868 | 2.8433 | 0.0546 | 0.0287 | 0.5260 |
| cluster3029 | -6675.71398 | 0.0288 | 0.0265 | 0.9211 | 0.0288 | 0.0096 | 0.3338 |
| cluster3030 | -5268.16191 | 0.0580 | 0.0238 | 0.4101 | 0.0493 | 0.0162 | 0.3279 |
| cluster3031 | -8794.56184 | 0.1202 | 0.0733 | 0.6104 | 0.0821 | 0.0632 | 0.7698 |
| cluster3032 | -5914.79441 | 0.0189 | 0.0171 | 0.9033 | 0.0280 | 0.0192 | 0.6848 |
| cluster3033 | -3821.651802 | 0.1830 | 0.0586 | 0.3202 | 0.1009 | 0.0069 | 0.0688 |
| cluster3034 | -4967.04172 | 0.0347 | 0.0666 | 1.9207 | 0.0339 | 0.3805 | 11.2201 |
| cluster3035 | -7642.87647 | 0.0182 | 0.0174 | 0.9531 | 0.0262 | 0.0133 | 0.5081 |
| cluster3036 | -8981.09531 | 0.0637 | 0.1050 | 1.6480 | 0.0805 | 0.0407 | 0.5052 |
| cluster3037 | -2547.50489 | 0.0045 | 0.0035 | 0.7853 | 0.0543 | 0.0099 | 0.1824 |
| cluster3038 | -4471.43426 | 0.0598 | 0.0881 | 1.4737 | 0.0590 | 0.0220 | 0.3731 |
| cluster3039 | -6524.93235 | 0.0436 | 0.0155 | 0.3556 | 0.0235 | 0.0071 | 0.3031 |
| cluster3040 | -4843.83042 | 0.0170 | 0.0078 | 0.4577 | 0.0216 | 0.0067 | 0.3128 |
| cluster3041 | -8441.09451 | 0.0348 | 0.0663 | 1.9073 | 0.0624 | 0.0343 | 0.5490 |
| cluster3042 | -4400.37745 | 0.0254 | 0.0472 | 1.8596 | 0.0492 | 0.0176 | 0.3584 |
| cluster3043 | -2912.40733 | 0.0111 | 0.0730 | 6.5534 | 0.0423 | 0.0276 | 0.6507 |
| cluster3044 | -6427.98536 | 0.0327 | 0.0518 | 1.5839 | 0.0275 | 0.0133 | 0.4839 |
| cluster3045 | -6128.46639 | 0.0489 | 0.0368 | 0.7519 | 0.0578 | 0.0174 | 0.3012 |
| cluster3046 | -7266.49014 | 0.1132 | 0.0558 | 0.4927 | 0.2525 | 0.1288 | 0.5101 |
| cluster3047 | -2123.64377 | 0.0030 | 0.0027 | 0.8996 | 0.0040 | 0.0019 | 0.4756 |
| cluster3048 | -5407.12198 | 0.0200 | 0.0215 | 1.0785 | 0.0255 | 0.0087 | 0.3430 |
| cluster3049 | -7062.02866 | 0.0120 | 0.0068 | 0.5705 | 0.0203 | 0.0143 | 0.7047 |
| cluster3050 | -7562.08278 | 0.0375 | 0.0450 | 1.1995 | 0.0440 | 0.0129 | 0.2941 |
| cluster3051 | -7511.91296 | 0.0336 | 0.0753 | 2.2410 | 0.0461 | 0.0295 | 0.6410 |
| cluster3052 | -6821.76197 | 0.0366 | 0.0744 | 2.0307 | 0.0655 | 0.0309 | 0.4719 |
| cluster3053 | -9371.09384 | 0.0907 | 0.0664 | 0.7317 | 0.0940 | 0.0591 | 0.6286 |
| cluster3054 | -6875.88218 | 0.0065 | 0.0080 | 1.2320 | 0.0146 | 0.0061 | 0.4201 |
| cluster3055 | -3961.19987 | 0.0099 | 0.0104 | 1.0503 | 0.0837 | 0.0260 | 0.3104 |
| cluster3056 | -5995.04159 | 0.1059 | 0.0330 | 0.3112 | 0.0956 | 0.0237 | 0.2476 |
| cluster3057 | -919.81782 | 0.0570 | 0.0204 | 0.3582 | 0.0473 | 0.0112 | 0.2366 |
| cluster3058 | -2530.53071 | 0.0111 | 0.0035 | 0.3160 | 0.0429 | 0.0125 | 0.2907 |
| cluster3059 | -8712.73963 | 0.0613 | 0.1827 | 2.9819 | 0.0821 | 0.0794 | 0.9667 |
| cluster3060 | -5325.45481 | 0.0282 | 0.0473 | 1.6773 | 0.0405 | 0.0214 | 0.5298 |
| cluster3061 | -5740.70457 | 0.0176 | 0.0064 | 0.3645 | 0.0145 | 0.0062 | 0.4276 |
| cluster3062 | -5690.75046 | 0.0337 | 0.0205 | 0.6079 | 0.0412 | 0.0104 | 0.2528 |
| cluster3063 | -6147.23519 | 0.0283 | 0.0338 | 1.1969 | 0.0368 | 0.0155 | 0.4214 |
| cluster3064 | -1912.73331 | 0.0033 | 0.0018 | 0.5471 | 0.0274 | 0.0054 | 0.1978 |
| cluster3065 | -2296.40485 | 0.0641 | 0.0941 | 1.4670 | 0.1366 | 0.0476 | 0.3485 |
| cluster3066 | -6187.60797 | 0.0685 | 0.0353 | 0.5151 | 0.0700 | 0.0372 | 0.5310 |
| cluster3067 | -4919.45828 | 0.0487 | 0.0314 | 0.6455 | 0.0345 | 0.0217 | 0.6311 |
| cluster3068 | -5143.14856 | 0.1010 | 0.0322 | 0.3186 | 0.1186 | 0.0274 | 0.2308 |
| cluster3069 | -1632.07958 | 0.0020 | 0.0007 | 0.3256 | 0.0001 | 0.0000 | 0.0907 |
| cluster3070 | -5192.46608 | 0.0356 | 0.0304 | 0.8535 | 0.0376 | 0.0242 | 0.6421 |

| | | | | | | | |
|---|---|---|---|---|---|---|---|
| cluster3071 | -5857.30150 | 0.0200 | 0.0090 | 0.4500 | 0.0246 | 0.0104 | 0.4214 |
| cluster3072 | -7368.06231 | 0.0662 | 0.0887 | 1.3403 | 0.1088 | 0.0518 | 0.4761 |
| cluster3073 | -3570.58642 | 0.0072 | 0.0076 | 1.0498 | 0.0084 | 0.0021 | 0.2455 |
| cluster3074 | -7650.13283 | 0.0258 | 0.0656 | 2.5403 | 0.0345 | 0.0226 | 0.6549 |
| cluster3075 | -5428.08474 | 0.0369 | 0.0291 | 0.7894 | 0.0370 | 0.0218 | 0.5896 |
| cluster3076 | -1812.27411 | 0.0397 | 0.0297 | 0.7485 | 0.0142 | 0.0461 | 3.2538 |
| cluster3077 | -4384.35891 | 0.0269 | 0.0240 | 0.8901 | 0.0198 | 0.0124 | 0.6250 |
| cluster3078 | -5021.99816 | 0.0144 | 0.0167 | 1.1603 | 0.0317 | 0.0159 | 0.5022 |
| cluster3079 | -7619.52663 | 0.0415 | 0.0234 | 0.5642 | 0.0271 | 0.0207 | 0.7660 |
| cluster3080 | -5650.06733 | 0.0159 | 0.0197 | 1.2391 | 0.0168 | 0.0064 | 0.3836 |
| cluster3081 | -3875.19760 | 0.0124 | 0.0071 | 0.5749 | 0.0078 | 0.0044 | 0.5584 |
| cluster3082 | -7186.46630 | 0.0517 | 0.0707 | 1.3682 | 0.0673 | 0.0304 | 0.4515 |
| cluster3083 | -4157.40104 | 0.0380 | 0.0138 | 0.3630 | 0.0186 | 0.0096 | 0.5156 |
| cluster3084 | -3197.62172 | 0.0433 | 0.0089 | 0.2064 | 0.0309 | 0.0044 | 0.1421 |
| cluster3085 | -7339.50271 | 0.0162 | 0.0616 | 3.7983 | 0.0337 | 0.0277 | 0.8201 |
| cluster3086 | -3385.24489 | 0.0382 | 0.0181 | 0.4742 | 0.1937 | 0.0685 | 0.3537 |
| cluster3087 | -6567.54040 | 0.0206 | 0.0172 | 0.8335 | 0.0165 | 0.0052 | 0.3119 |
| cluster3088 | -6390.34547 | 0.0535 | 0.0475 | 0.8890 | 0.0802 | 0.0284 | 0.3545 |
| cluster3089 | -3685.38943 | 0.0652 | 0.0310 | 0.4762 | 0.0784 | 0.0416 | 0.5316 |
| cluster3090 | -4959.10604 | 0.0109 | 0.0147 | 1.3452 | 0.0056 | 0.0031 | 0.5426 |
| cluster3091 | -5909.67809 | 0.0449 | 0.0187 | 0.4173 | 0.0656 | 0.0204 | 0.3118 |
| cluster3092 | -4264.05671 | 0.0139 | 0.0159 | 1.1439 | 0.0226 | 0.0088 | 0.3874 |
| cluster3093 | -4540.76203 | 0.0042 | 0.0108 | 2.5845 | 0.0140 | 0.0037 | 0.2652 |
| cluster3094 | -7861.81317 | 0.0231 | 0.0718 | 3.1111 | 0.0388 | 0.0297 | 0.7638 |
| cluster3095 | -4869.08617 | 0.0042 | 0.0024 | 0.5881 | 0.0082 | 0.0068 | 0.8348 |
| cluster3096 | -6132.20650 | 0.0309 | 0.0426 | 1.3790 | 0.0371 | 0.0136 | 0.3668 |
| cluster3097 | -7077.60788 | 0.0216 | 0.0108 | 0.4992 | 0.0203 | 0.0068 | 0.3358 |
| cluster3098 | -1418.18477 | 0.0054 | 0.0121 | 2.2530 | 0.0105 | 0.0059 | 0.5622 |
| cluster3099 | -4267.30144 | 0.0234 | 0.0149 | 0.6387 | 0.0553 | 0.0348 | 0.6293 |
| cluster3100 | -4393.80661 | 0.0464 | 0.0555 | 1.1971 | 0.0370 | 0.0152 | 0.4114 |
| cluster3101 | -3436.11176 | 0.0498 | 0.0315 | 0.6328 | 0.0460 | 0.0283 | 0.6142 |
| cluster3102 | -2231.68448 | 0.0140 | 0.0106 | 0.7579 | 0.0151 | 0.0122 | 0.8106 |
| cluster3103 | -5215.21687 | 0.0291 | 0.0882 | 3.0307 | 0.0451 | 0.0326 | 0.7229 |
| cluster3104 | -4325.94619 | 0.0167 | 0.0191 | 1.1406 | 0.0272 | 0.0139 | 0.5115 |
| cluster3105 | -7969.93437 | 0.0217 | 0.0694 | 3.1985 | 0.0351 | 0.0255 | 0.7275 |
| cluster3106 | -5132.24742 | 0.0029 | 0.0014 | 0.4946 | 0.0027 | 0.0012 | 0.4398 |
| cluster3107 | -3137.74707 | 0.0389 | 0.0265 | 0.6807 | 0.0454 | 0.0091 | 0.1997 |
| cluster3108 | -4328.04526 | 0.1069 | 0.1082 | 1.0126 | 0.1859 | 0.0536 | 0.2881 |
| cluster3109 | -3458.43665 | 0.0196 | 0.0351 | 1.7954 | 0.0412 | 0.0190 | 0.4609 |
| cluster3110 | -6380.31188 | 0.0988 | 0.0801 | 0.8110 | 0.1232 | 0.0287 | 0.2331 |
| cluster3111 | -4889.11079 | 0.0350 | 0.0341 | 0.9756 | 0.0290 | 0.0097 | 0.3351 |
| cluster3112 | -6143.47219 | 0.0293 | 0.0894 | 3.0480 | 0.0493 | 0.0334 | 0.6779 |
| cluster3113 | -3067.57692 | 0.0295 | 0.0402 | 1.3619 | 0.0190 | 0.0093 | 0.4903 |
| cluster3114 | -4682.93666 | 0.0205 | 0.0150 | 0.7299 | 0.0482 | 0.0166 | 0.3454 |

| | | | | | | | |
|---|---|---|---|---|---|---|---|
| cluster3115 | -3777.29780 | 0.0164 | 0.0185 | 1.1282 | 0.0208 | 0.0083 | 0.4016 |
| cluster3116 | -2723.86717 | 0.0427 | 0.0482 | 1.1297 | 0.0313 | 0.0109 | 0.3480 |
| cluster3117 | -7494.39582 | 0.0339 | 0.1129 | 3.3281 | 0.0628 | 0.0378 | 0.6029 |
| cluster3118 | -1567.42321 | 0.0095 | 0.0032 | 0.3325 | 0.0907 | 0.0086 | 0.0951 |
| cluster3119 | -2981.73819 | 0.0498 | 0.0378 | 0.7582 | 0.0308 | 0.0351 | 1.1398 |
| cluster3120 | -4909.34217 | 0.0192 | 0.0224 | 1.1665 | 0.0289 | 0.0092 | 0.3177 |
| cluster3121 | -6820.35062 | 0.0226 | 0.0199 | 0.8806 | 0.0318 | 0.0219 | 0.6868 |
| cluster3122 | -3558.19631 | 0.0187 | 0.0089 | 0.4756 | 0.0236 | 0.0089 | 0.3792 |
| cluster3123 | -3830.43265 | 0.0181 | 0.0102 | 0.5651 | 0.0104 | 0.0044 | 0.4267 |
| cluster3124 | -6375.64354 | 0.0334 | 0.0400 | 1.1993 | 0.0478 | 0.0203 | 0.4240 |
| cluster3125 | -4801.84394 | 0.0766 | 0.0630 | 0.8227 | 0.0547 | 0.0177 | 0.3232 |
| cluster3126 | -7551.88292 | 0.0481 | 0.0354 | 0.7363 | 0.0366 | 0.0229 | 0.6258 |
| cluster3127 | -2604.84844 | 0.0052 | 0.0104 | 2.0169 | 0.0142 | 0.0051 | 0.3577 |
| cluster3128 | -2238.18709 | 0.0016 | 0.0010 | 0.6525 | 0.0030 | 0.0008 | 0.2557 |
| cluster3129 | -3261.94400 | 0.0214 | 0.1122 | 5.2318 | 0.0481 | 0.0457 | 0.9501 |
| cluster3130 | -5798.93411 | 0.0326 | 0.0628 | 1.9282 | 0.0404 | 0.0203 | 0.5018 |
| cluster3131 | -1762.33349 | 0.0048 | 0.0107 | 2.2061 | 0.0135 | 0.0087 | 0.6431 |
| cluster3132 | -5289.90993 | 0.0269 | 0.0152 | 0.5662 | 0.0228 | 0.0131 | 0.5750 |
| cluster3133 | -8921.42569 | 0.0481 | 0.1401 | 2.9118 | 0.0778 | 0.0624 | 0.8019 |
| cluster3134 | -3291.00937 | 0.0564 | 0.0280 | 0.4959 | 0.0606 | 0.0172 | 0.2836 |
| cluster3135 | -4395.67720 | 0.0073 | 0.0092 | 1.2584 | 0.0192 | 0.0054 | 0.2829 |
| cluster3136 | -3954.05889 | 0.0370 | 0.0605 | 1.6362 | 0.0392 | 0.0216 | 0.5514 |
| cluster3137 | -3086.94624 | 0.0168 | 0.0335 | 1.9984 | 0.0372 | 0.0201 | 0.5414 |
| cluster3138 | -4743.04326 | 0.0533 | 0.0531 | 0.9977 | 0.0529 | 0.0399 | 0.7530 |
| cluster3139 | -5753.50936 | 0.0625 | 0.1391 | 2.2257 | 0.0739 | 0.0631 | 0.8540 |
| cluster3140 | -3559.21841 | 0.0361 | 0.0192 | 0.5322 | 0.0560 | 0.0089 | 0.1591 |
| cluster3141 | -2414.43073 | 0.0143 | 0.0053 | 0.3722 | 0.0060 | 0.0018 | 0.3052 |
| cluster3142 | -1635.452986 | 0.0428 | 0.0091 | 0.2127 | 0.2498 | 0.0124 | 0.0497 |
| cluster3143 | -3221.87497 | 0.0062 | 0.0034 | 0.5501 | 0.0253 | 0.0095 | 0.3756 |
| cluster3144 | -2088.76585 | 0.0049 | 0.0537 | 10.9659 | 0.0046 | 0.0107 | 2.3287 |
| cluster3145 | -6530.48213 | 0.0049 | 0.0096 | 1.9625 | 0.0061 | 0.0047 | 0.7734 |
| cluster3146 | -1742.11965 | 0.0029 | 0.0029 | 1.0294 | 0.0021 | 0.0137 | 6.4076 |
| cluster3147 | -4472.89127 | 0.0057 | 0.0084 | 1.4688 | 0.0075 | 0.0027 | 0.3637 |
| cluster3148 | -6021.28470 | 0.0245 | 0.0216 | 0.8843 | 0.0371 | 0.0106 | 0.2861 |
| cluster3149 | -2369.25931 | 0.0060 | 0.0028 | 0.4697 | 0.0001 | 0.0000 | 0.4056 |
| cluster3150 | -4524.54657 | 0.0686 | 0.0354 | 0.5161 | 0.0638 | 0.0096 | 0.1497 |
| cluster3151 | -3975.22868 | 0.0535 | 0.0424 | 0.7922 | 0.0471 | 0.0146 | 0.3101 |
| cluster3152 | -8744.54981 | 0.0420 | 0.1140 | 2.7141 | 0.0561 | 0.0428 | 0.7640 |
| cluster3153 | -4517.03905 | 0.0418 | 0.0334 | 0.8002 | 0.0449 | 0.0118 | 0.2635 |
| cluster3154 | -5341.47387 | 0.0111 | 0.0162 | 1.4594 | 0.0200 | 0.0113 | 0.5668 |
| cluster3155 | -3014.36664 | 0.0072 | 0.0112 | 1.5652 | 0.0116 | 0.0047 | 0.4010 |
| cluster3156 | -6784.01914 | 0.0180 | 0.0551 | 3.0541 | 0.0363 | 0.0178 | 0.4922 |
| cluster3157 | -3281.00069 | 0.1000 | 0.0414 | 0.4141 | 0.0963 | 0.0121 | 0.1258 |
| cluster3158 | -3373.10772 | 0.0149 | 0.0123 | 0.8258 | 0.0155 | 0.0151 | 0.9775 |

| | | | | | | | |
|---|---|---|---|---|---|---|---|
| cluster3159 | -5810.85865 | 0.0368 | 0.0934 | 2.5370 | 0.0597 | 0.0381 | 0.6380 |
| cluster3160 | -1098.384689 | 0.0747 | 0.0178 | 0.2388 | 0.0220 | 0.0022 | 0.0989 |
| cluster3161 | -4981.92854 | 0.0286 | 0.0714 | 2.4979 | 0.0367 | 0.0169 | 0.4608 |
| cluster3162 | -3639.24551 | 0.0080 | 0.0058 | 0.7236 | 0.0130 | 0.0018 | 0.1360 |
| cluster3163 | -3429.71459 | 0.0309 | 0.0706 | 2.2871 | 0.0422 | 0.0194 | 0.4604 |
| cluster3164 | -6178.27847 | 0.0479 | 0.0808 | 1.6867 | 0.0717 | 0.0379 | 0.5286 |
| cluster3165 | -8045.69485 | 0.0433 | 0.0908 | 2.0957 | 0.0707 | 0.0468 | 0.6620 |
| cluster3166 | -4631.30621 | 0.0186 | 0.0122 | 0.6563 | 0.0320 | 0.0177 | 0.5514 |
| cluster3167 | -5313.33750 | 0.0984 | 0.0683 | 0.6945 | 0.0839 | 0.0152 | 0.1810 |
| cluster3168 | -7090.15349 | 0.0202 | 0.0223 | 1.1063 | 0.0159 | 0.0199 | 1.2538 |
| cluster3169 | -5027.88042 | 0.0229 | 0.0580 | 2.5267 | 0.0373 | 0.0215 | 0.5756 |
| cluster3170 | -1623.07695 | 0.0153 | 0.0036 | 0.2317 | 0.0095 | 0.0008 | 0.0838 |
| cluster3171 | -6149.87687 | 0.0948 | 0.0693 | 0.7308 | 0.0393 | 0.0205 | 0.5214 |
| cluster3172 | -6262.15786 | 0.0291 | 0.0222 | 0.7624 | 0.0267 | 0.0166 | 0.6212 |
| cluster3173 | -4457.22460 | 0.0134 | 0.0208 | 1.5538 | 0.0177 | 0.0083 | 0.4667 |
| cluster3174 | -1300.73983 | 0.0011 | 0.0011 | 1.0865 | 0.0038 | 0.0015 | 0.3931 |
| cluster3175 | -1938.14077 | 0.0169 | 0.0160 | 0.9419 | 0.0141 | 0.0037 | 0.2592 |
| cluster3176 | -3887.40405 | 0.0278 | 0.0182 | 0.6562 | 0.0199 | 0.0057 | 0.2858 |
| cluster3177 | -3504.19209 | 0.0392 | 0.0236 | 0.6017 | 0.0710 | 0.0238 | 0.3348 |
| cluster3178 | -6743.67227 | 0.0084 | 0.0087 | 1.0340 | 0.0141 | 0.0147 | 1.0426 |
| cluster3179 | -4628.08849 | 0.0461 | 0.1705 | 3.6980 | 0.0828 | 0.0502 | 0.6069 |
| cluster3180 | -1070.72322 | 0.0018 | 0.0124 | 6.9880 | 0.0060 | 0.0018 | 0.2934 |
| cluster3181 | -6693.35303 | 0.0204 | 0.0198 | 0.9704 | 0.0226 | 0.0188 | 0.8339 |
| cluster3182 | -2530.16874 | 0.0050 | 0.0020 | 0.3998 | 0.0084 | 0.0040 | 0.4779 |
| cluster3183 | -4041.32753 | 0.0671 | 0.0776 | 1.1576 | 0.0962 | 0.0527 | 0.5476 |
| cluster3184 | -4206.14969 | 0.0372 | 0.0260 | 0.6997 | 0.0237 | 0.0059 | 0.2484 |
| cluster3185 | -3546.67555 | 0.0160 | 0.0331 | 2.0682 | 0.0180 | 0.0125 | 0.6976 |
| cluster3186 | -2289.81315 | 0.0229 | 0.0296 | 1.2896 | 0.0374 | 0.0096 | 0.2574 |
| cluster3187 | -5392.88884 | 0.0235 | 0.0293 | 1.2463 | 0.0175 | 0.0076 | 0.4347 |
| cluster3188 | -4445.34050 | 0.0924 | 0.0390 | 0.4226 | 0.0881 | 0.0426 | 0.4838 |
| cluster3189 | -6228.06133 | 0.0351 | 0.0836 | 2.3840 | 0.0500 | 0.0287 | 0.5733 |
| cluster3190 | -5396.79986 | 0.0256 | 0.0747 | 2.9234 | 0.0378 | 0.0262 | 0.6926 |
| cluster3191 | -2831.35728 | 0.0518 | 0.0345 | 0.6659 | 0.0419 | 0.0251 | 0.6005 |
| cluster3192 | -5887.00165 | 0.0552 | 0.0274 | 0.4967 | 0.0483 | 0.0243 | 0.5021 |
| cluster3193 | -3444.61237 | 0.0172 | 0.0102 | 0.5896 | 0.0662 | 0.0176 | 0.2665 |
| cluster3194 | -6975.34463 | 0.0368 | 0.0950 | 2.5807 | 0.0804 | 0.0463 | 0.5765 |
| cluster3195 | -3959.45754 | 0.0225 | 0.0244 | 1.0808 | 0.0134 | 0.0105 | 0.7853 |
| cluster3196 | -2244.51818 | 0.0057 | 0.0013 | 0.2348 | 0.0028 | 0.0005 | 0.1631 |
| cluster3197 | -5445.80939 | 0.0426 | 0.0189 | 0.4430 | 0.0614 | 0.0244 | 0.3970 |
| cluster3198 | -6431.16898 | 0.0278 | 0.0638 | 2.2976 | 0.0210 | 0.0151 | 0.7177 |
| cluster3199 | -2975.54875 | 0.0062 | 0.0063 | 1.0108 | 0.0206 | 0.0058 | 0.2838 |
| cluster3200 | -4262.83790 | 0.0024 | 0.0140 | 5.8487 | 0.0281 | 0.0135 | 0.4823 |
| cluster3201 | -4476.70273 | 0.1104 | 0.0458 | 0.4151 | 0.1118 | 0.0161 | 0.1443 |
| cluster3202 | -6071.47952 | 0.0428 | 0.0398 | 0.9296 | 0.0425 | 0.0130 | 0.3067 |

| | | | | | | | |
|---|---|---|---|---|---|---|---|
| cluster3203 | -3017.04464 | 0.0434 | 0.1110 | 2.5562 | 0.0564 | 0.0301 | 0.5337 |
| cluster3204 | -1878.16329 | 0.0344 | 0.0079 | 0.2294 | 0.0427 | 0.0092 | 0.2152 |
| cluster3205 | -2015.80078 | 0.0062 | 0.0017 | 0.2660 | 0.0059 | 0.0010 | 0.1644 |
| cluster3206 | -3914.70479 | 0.0363 | 0.0507 | 1.3971 | 0.0318 | 0.0172 | 0.5400 |
| cluster3207 | -5500.72942 | 0.0143 | 0.0164 | 1.1437 | 0.0134 | 0.0111 | 0.8234 |
| cluster3208 | -3285.75862 | 0.0066 | 0.0037 | 0.5675 | 0.0031 | 0.0012 | 0.3978 |
| cluster3209 | -5553.31439 | 0.0209 | 0.0416 | 1.9948 | 0.0255 | 0.0150 | 0.5889 |
| cluster3210 | -3377.95371 | 0.0033 | 0.0217 | 6.6878 | 0.0027 | 0.0016 | 0.6011 |
| cluster3211 | -3434.98954 | 0.0336 | 0.0539 | 1.6063 | 0.0291 | 0.0148 | 0.5090 |
| cluster3212 | -2809.63302 | 0.0047 | 0.0014 | 0.2969 | 0.0007 | 0.0003 | 0.5317 |
| cluster3213 | -4613.55847 | 0.1164 | 0.0720 | 0.6184 | 0.0764 | 0.0644 | 0.8429 |
| cluster3214 | -6050.70033 | 0.0408 | 0.0302 | 0.7411 | 0.0555 | 0.0267 | 0.4821 |
| cluster3215 | -2279.12936 | 0.0016 | 0.0005 | 0.2890 | 0.0041 | 0.0003 | 0.0760 |
| cluster3216 | -4536.01005 | 0.0209 | 0.0971 | 4.6547 | 0.0385 | 0.0343 | 0.8916 |
| cluster3217 | -3824.69694 | 0.0464 | 0.0232 | 0.5000 | 0.0384 | 0.0182 | 0.4740 |
| cluster3218 | -5083.93102 | 0.0220 | 0.0540 | 2.4519 | 0.0281 | 0.0177 | 0.6297 |
| cluster3219 | -5449.15186 | 0.0103 | 0.0191 | 1.8526 | 0.0146 | 0.0068 | 0.4660 |
| cluster3220 | -2923.99523 | 0.0249 | 0.0106 | 0.4268 | 0.0361 | 0.0150 | 0.4153 |
| cluster3221 | -5732.34955 | 0.0234 | 0.0707 | 3.0179 | 0.0447 | 0.0289 | 0.6464 |
| cluster3222 | -2621.78624 | 0.0448 | 0.0287 | 0.6404 | 0.0358 | 0.0135 | 0.3781 |
| cluster3223 | -3457.47441 | 0.1304 | 0.1423 | 1.0913 | 0.0401 | 0.0091 | 0.2273 |
| cluster3224 | -1597.24353 | 0.0001 | 0.0000 | 0.4757 | 0.0001 | 0.0000 | 0.1182 |
| cluster3225 | -6105.04787 | 0.0285 | 0.0532 | 1.8657 | 0.0370 | 0.0165 | 0.4448 |
| cluster3226 | -3719.85176 | 0.0236 | 0.0381 | 1.6143 | 0.0616 | 0.0239 | 0.3890 |
| cluster3227 | -4398.67648 | 0.0699 | 0.1110 | 1.5881 | 0.0682 | 0.0432 | 0.6331 |
| cluster3228 | -3621.53825 | 0.0215 | 0.0629 | 2.9201 | 0.0476 | 0.0248 | 0.5211 |
| cluster3229 | -3857.86139 | 0.0121 | 0.0084 | 0.6902 | 0.0541 | 0.0281 | 0.5192 |
| cluster3230 | -3482.28727 | 0.0122 | 0.0059 | 0.4849 | 0.0098 | 0.0049 | 0.5018 |
| cluster3231 | -4528.28769 | 0.0213 | 0.0248 | 1.1649 | 0.0287 | 0.0116 | 0.4045 |
| cluster3232 | -5052.18779 | 0.0180 | 0.0349 | 1.9329 | 0.0292 | 0.0181 | 0.6197 |
| cluster3233 | -3436.45268 | 0.0388 | 0.0374 | 0.9640 | 0.0437 | 0.0403 | 0.9233 |
| cluster3234 | -3821.29820 | 0.0527 | 0.0356 | 0.6758 | 0.0491 | 0.0305 | 0.6211 |
| cluster3235 | -7271.36630 | 0.0334 | 0.1057 | 3.1642 | 0.0419 | 0.0339 | 0.8096 |
| cluster3236 | -3956.70997 | 0.0327 | 0.0563 | 1.7223 | 0.0334 | 0.0208 | 0.6211 |
| cluster3237 | -3243.94352 | 0.0505 | 0.0676 | 1.3396 | 0.0688 | 0.0334 | 0.4857 |
| cluster3238 | -2743.43805 | 0.0027 | 0.0010 | 0.3698 | 0.1536 | 0.0297 | 0.1935 |
| cluster3239 | -2738.17502 | 0.0157 | 0.0101 | 0.6471 | 0.0344 | 0.0077 | 0.2227 |
| cluster3240 | -6961.66899 | 0.0353 | 0.0280 | 0.7956 | 0.0289 | 0.0246 | 0.8504 |
| cluster3241 | -5756.22983 | 0.0602 | 0.0867 | 1.4416 | 0.0694 | 0.0314 | 0.4526 |
| cluster3242 | -7159.42806 | 0.0325 | 0.1026 | 3.1549 | 0.0589 | 0.0413 | 0.7006 |
| cluster3243 | -5984.09652 | 0.1436 | 0.0469 | 0.3269 | 0.1615 | 0.0470 | 0.2909 |
| cluster3244 | -6473.16345 | 0.0554 | 0.0396 | 0.7149 | 0.0648 | 0.0374 | 0.5769 |
| cluster3245 | -2584.54745 | 0.0019 | 0.0032 | 1.6477 | 0.0053 | 0.0022 | 0.4230 |
| cluster3246 | -5134.91182 | 0.0451 | 0.0418 | 0.9261 | 0.0402 | 0.0465 | 1.1574 |

| | | | | | | | |
|---|---|---|---|---|---|---|---|
| cluster3247 | -848.89256 | 0.0525 | 0.0590 | 1.1239 | 0.0801 | 0.0198 | 0.2473 |
| cluster3248 | -3484.17390 | 0.0176 | 0.0250 | 1.4224 | 0.0267 | 0.0135 | 0.5061 |
| cluster3249 | -3479.85474 | 0.0119 | 0.0197 | 1.6563 | 0.0257 | 0.0106 | 0.4124 |
| cluster3250 | -3551.46699 | 0.0939 | 0.0421 | 0.4484 | 0.1351 | 0.0355 | 0.2629 |
| cluster3251 | -4466.97248 | 0.0518 | 0.0520 | 1.0037 | 0.0664 | 0.0233 | 0.3512 |
| cluster3252 | -7422.97968 | 0.0640 | 0.1308 | 2.0444 | 0.0634 | 0.0546 | 0.8615 |
| cluster3253 | -4340.21730 | 0.0632 | 0.0796 | 1.2605 | 0.0619 | 0.0247 | 0.3998 |
| cluster3254 | -3108.94960 | 0.0279 | 0.0411 | 1.4728 | 0.0457 | 0.0177 | 0.3866 |
| cluster3255 | -5568.40375 | 0.0132 | 0.0444 | 3.3624 | 0.0225 | 0.0197 | 0.8772 |
| cluster3256 | -6098.53128 | 0.0334 | 0.0455 | 1.3628 | 0.0512 | 0.0251 | 0.4895 |
| cluster3257 | -5144.74136 | 0.0070 | 0.0051 | 0.7311 | 0.0060 | 0.0051 | 0.8517 |
| cluster3258 | -1918.58951 | 0.0020 | 0.0014 | 0.7293 | 0.0001 | 0.0000 | 0.2047 |
| cluster3259 | -3556.52491 | 0.0357 | 0.0087 | 0.2443 | 0.0363 | 0.0096 | 0.2648 |
| cluster3260 | -4482.35940 | 0.0151 | 0.0083 | 0.5484 | 0.0126 | 0.0079 | 0.6290 |
| cluster3261 | -4941.49893 | 0.0464 | 0.1346 | 2.9035 | 0.1004 | 0.0510 | 0.5083 |
| cluster3262 | -5790.57311 | 0.0161 | 0.0421 | 2.6221 | 0.0221 | 0.0132 | 0.5963 |
| cluster3263 | -2031.76492 | 0.0331 | 0.0181 | 0.5470 | 0.2934 | 0.0827 | 0.2818 |
| cluster3264 | -1832.56585 | 0.0055 | 0.0033 | 0.5925 | 0.0238 | 0.0032 | 0.1343 |
| cluster3265 | -4128.75602 | 0.0033 | 0.0040 | 1.2201 | 0.0024 | 0.0011 | 0.4848 |
| cluster3266 | -2849.84064 | 0.0145 | 0.0065 | 0.4480 | 0.0523 | 0.0103 | 0.1965 |
| cluster3267 | -4120.23838 | 0.0204 | 0.0143 | 0.7016 | 0.0464 | 0.0257 | 0.5532 |
| cluster3268 | -2802.29770 | 0.0242 | 0.0293 | 1.2111 | 0.0584 | 0.0215 | 0.3675 |
| cluster3269 | -3537.71429 | 0.0659 | 0.0758 | 1.1491 | 0.0493 | 0.0188 | 0.3817 |
| cluster3270 | -2181.60476 | 0.0161 | 0.0414 | 2.5783 | 0.0459 | 0.0189 | 0.4125 |
| cluster3271 | -4112.49462 | 0.0107 | 0.0081 | 0.7502 | 0.0144 | 0.0057 | 0.3968 |
| cluster3272 | -938.361518 | 0.4222 | 0.0267 | 0.0632 | 0.1731 | 0.0135 | 0.0780 |
| cluster3273 | -4005.11545 | 0.1018 | 0.1520 | 1.4929 | 0.1288 | 0.0576 | 0.4474 |
| cluster3274 | -2029.03806 | 0.0174 | 0.0522 | 3.0000 | 0.0496 | 0.0314 | 0.6341 |
| cluster3275 | -4551.15201 | 0.0468 | 0.0691 | 1.4762 | 0.0507 | 0.0313 | 0.6176 |
| cluster3276 | -2485.27608 | 0.0764 | 0.0131 | 0.1715 | 0.1017 | 0.0195 | 0.1917 |
| cluster3277 | -3226.82452 | 0.0391 | 0.0961 | 2.4560 | 0.0610 | 0.0259 | 0.4251 |
| cluster3278 | -4576.57251 | 0.0021 | 0.0013 | 0.5970 | 0.0053 | 0.0011 | 0.1988 |
| cluster3279 | -3732.04283 | 0.0193 | 0.0318 | 1.6507 | 0.0130 | 0.0069 | 0.5270 |
| cluster3280 | -3788.56260 | 0.0232 | 0.0201 | 0.8695 | 0.0206 | 0.0060 | 0.2917 |
| cluster3281 | -1962.63516 | 0.0107 | 0.0239 | 2.2459 | 0.0372 | 0.0199 | 0.5357 |
| cluster3282 | -2048.72661 | 0.0043 | 0.0046 | 1.0686 | 0.0082 | 0.0025 | 0.3039 |
| cluster3283 | -5464.79992 | 0.0612 | 0.0386 | 0.6317 | 0.0676 | 0.0404 | 0.5979 |
| cluster3284 | -5980.28960 | 0.0335 | 0.0475 | 1.4203 | 0.0362 | 0.0149 | 0.4121 |
| cluster3285 | -2362.67566 | 0.0084 | 0.0054 | 0.6372 | 0.0053 | 0.0011 | 0.2053 |
| cluster3286 | -2205.44023 | 0.0174 | 0.0082 | 0.4715 | 0.0493 | 0.0106 | 0.2145 |
| cluster3287 | -1375.92558 | 0.0080 | 0.0083 | 1.0461 | 0.0220 | 0.0068 | 0.3110 |
| cluster3288 | -1238.44073 | 0.0030 | 0.0111 | 3.6750 | 0.0137 | 0.0084 | 0.6110 |
| cluster3289 | -5797.90272 | 0.0415 | 0.0168 | 0.4051 | 0.0556 | 0.0168 | 0.3025 |
| cluster3290 | -1250.54561 | 0.0134 | 0.0162 | 1.2106 | 0.0522 | 0.0100 | 0.1906 |

| | | | | | | | |
|---|---|---|---|---|---|---|---|
| cluster3291 | -4787.80685 | 0.0410 | 0.0315 | 0.7698 | 0.0388 | 0.0281 | 0.7245 |
| cluster3292 | -2768.50263 | 0.0162 | 0.0159 | 0.9845 | 0.0458 | 0.0123 | 0.2685 |
| cluster3293 | -3184.82953 | 0.0094 | 0.0108 | 1.1466 | 0.0097 | 0.0064 | 0.6547 |
| cluster3294 | -2883.07723 | 0.0120 | 0.0097 | 0.8059 | 0.0154 | 0.0110 | 0.7148 |
| cluster3295 | -2241.21355 | 0.0015 | 0.0011 | 0.7188 | 0.0018 | 0.0004 | 0.2014 |
| cluster3296 | -1799.32567 | 0.0054 | 0.0089 | 1.6643 | 0.0027 | 0.0019 | 0.7070 |
| cluster3297 | -2354.611709 | 0.2041 | 0.1159 | 0.5677 | 0.1314 | 0.0471 | 0.3586 |
| cluster3298 | -2943.17949 | 0.0059 | 0.0050 | 0.8511 | 0.0023 | 0.0019 | 0.8000 |
| cluster3299 | -7903.99754 | 0.0072 | 0.1697 | 23.7323 | 0.0645 | 0.0709 | 1.0993 |
| cluster3300 | -2245.68158 | 0.0012 | 0.0019 | 1.4946 | 0.0113 | 0.0059 | 0.5244 |
| cluster3301 | -1804.30980 | 0.0446 | 0.0158 | 0.3534 | 0.0527 | 0.0207 | 0.3935 |
| cluster3302 | -4544.81828 | 0.0463 | 0.0513 | 1.1070 | 0.1409 | 0.0503 | 0.3566 |
| cluster3303 | -4479.76102 | 0.0440 | 0.0434 | 0.9847 | 0.0300 | 0.0380 | 1.2678 |
| cluster3304 | -2852.56045 | 0.0416 | 0.0175 | 0.4217 | 0.0088 | 0.0032 | 0.3651 |
| cluster3305 | -3804.19788 | 0.0534 | 0.0800 | 1.4998 | 0.0756 | 0.0305 | 0.4029 |
| cluster3306 | -5013.99272 | 0.0497 | 0.0862 | 1.7351 | 0.1051 | 0.0453 | 0.4309 |
| cluster3307 | -1670.29091 | 0.0028 | 0.0049 | 1.7806 | 0.0082 | 0.0033 | 0.4007 |
| cluster3308 | -4387.49135 | 0.0215 | 0.0248 | 1.1568 | 0.0160 | 0.0146 | 0.9150 |
| cluster3309 | -3384.14835 | 0.0993 | 0.0378 | 0.3808 | 0.1419 | 0.0438 | 0.3088 |
| cluster3310 | -4174.15888 | 0.0123 | 0.0125 | 1.0111 | 0.0197 | 0.0094 | 0.4742 |
| cluster3311 | -6156.89097 | 0.0466 | 0.0381 | 0.8190 | 0.0638 | 0.0434 | 0.6799 |
| cluster3312 | -1931.84276 | 0.0279 | 0.0568 | 2.0352 | 0.0253 | 0.0193 | 0.7634 |
| cluster3313 | -2505.10479 | 0.0053 | 0.0023 | 0.4281 | 0.0214 | 0.0060 | 0.2820 |
| cluster3314 | -7138.00563 | 0.0299 | 0.1278 | 4.2760 | 0.0558 | 0.0472 | 0.8461 |
| cluster3315 | -3722.96085 | 0.0067 | 0.0041 | 0.6062 | 0.0051 | 0.0037 | 0.7158 |
| cluster3316 | -5974.07040 | 0.0781 | 0.1236 | 1.5837 | 0.0933 | 0.0410 | 0.4397 |
| cluster3317 | -1139.90637 | 0.0009 | 0.0010 | 1.1479 | 0.0088 | 0.0027 | 0.3005 |
| cluster3318 | -3441.78846 | 0.1086 | 0.1201 | 1.1065 | 0.1073 | 0.0982 | 0.9158 |
| cluster3319 | -3769.58275 | 0.0343 | 0.0615 | 1.7930 | 0.0371 | 0.0260 | 0.7004 |
| cluster3320 | -2814.00034 | 0.0534 | 0.0834 | 1.5622 | 0.0571 | 0.0240 | 0.4205 |
| cluster3321 | -2914.93599 | 0.0006 | 0.0006 | 1.1024 | 0.0073 | 0.0017 | 0.2303 |
| cluster3322 | -3279.03283 | 0.0385 | 0.0770 | 2.0008 | 0.0445 | 0.0302 | 0.6777 |
| cluster3323 | -5518.43353 | 0.0418 | 0.0597 | 1.4265 | 0.0631 | 0.0278 | 0.4409 |
| cluster3324 | -1399.21985 | 0.0330 | 0.0141 | 0.4261 | 0.0110 | 0.0415 | 3.7649 |
| cluster3325 | -2061.15856 | 0.0189 | 0.0324 | 1.7164 | 0.0185 | 0.0080 | 0.4317 |
| cluster3326 | -1700.06957 | 0.0502 | 0.0194 | 0.3872 | 0.1552 | 0.0127 | 0.0817 |
| cluster3327 | -3863.91675 | 0.0201 | 0.0155 | 0.7703 | 0.0303 | 0.0188 | 0.6204 |
| cluster3328 | -2602.26947 | 0.0311 | 0.0334 | 1.0737 | 0.0053 | 0.0015 | 0.2810 |
| cluster3329 | -3145.28275 | 0.0317 | 0.1004 | 3.1644 | 0.1020 | 0.0710 | 0.6961 |
| cluster3330 | -4708.52579 | 0.0363 | 0.0884 | 2.4341 | 0.0646 | 0.0277 | 0.4291 |
| cluster3331 | -2862.78475 | 0.0485 | 0.0313 | 0.6444 | 0.1176 | 0.0244 | 0.2073 |
| cluster3332 | -6186.85934 | 0.0466 | 0.1144 | 2.4565 | 0.0837 | 0.0472 | 0.5636 |
| cluster3333 | -3575.05899 | 0.0183 | 0.0388 | 2.1245 | 0.0288 | 0.0160 | 0.5564 |
| cluster3334 | -1377.93026 | 0.0465 | 0.0360 | 0.7739 | 0.1189 | 0.0254 | 0.2138 |

| | | | | | | | |
|---|---|---|---|---|---|---|---|
| cluster3335 | -2858.35614 | 0.0270 | 0.0183 | 0.6778 | 0.0310 | 0.0339 | 1.0947 |
| cluster3336 | -3028.07686 | 0.0037 | 0.0016 | 0.4443 | 0.0039 | 0.0015 | 0.3737 |
| cluster3337 | -2620.42367 | 0.0304 | 0.0097 | 0.3209 | 0.0314 | 0.0074 | 0.2352 |
| cluster3338 | -6300.33958 | 0.0302 | 0.0934 | 3.0929 | 0.0497 | 0.0340 | 0.6826 |
| cluster3339 | -1907.97145 | 0.0708 | 0.0467 | 0.6592 | 0.0294 | 0.0364 | 1.2359 |
| cluster3340 | -1623.085447 | 0.0656 | 0.0076 | 0.1152 | 0.2325 | 0.0383 | 0.1646 |
| cluster3341 | -1577.00534 | 0.0113 | 0.0271 | 2.4070 | 0.0106 | 0.0062 | 0.5869 |
| cluster3342 | -2554.39119 | 0.0256 | 0.0490 | 1.9180 | 0.0040 | 0.0111 | 2.7641 |
| cluster3343 | -2925.93633 | 0.0006 | 0.0009 | 1.4539 | 0.0001 | 0.0000 | 0.4433 |
| cluster3344 | -3149.30791 | 0.0253 | 0.0116 | 0.4586 | 0.0282 | 0.0086 | 0.3049 |
| cluster3345 | -2792.26743 | 0.0080 | 0.0272 | 3.3905 | 0.0217 | 0.0104 | 0.4780 |
| cluster3346 | -3610.28822 | 0.0034 | 0.0030 | 0.8675 | 0.0037 | 0.0022 | 0.5976 |
| cluster3347 | -4174.24768 | 0.0585 | 0.0264 | 0.4511 | 0.0371 | 0.0154 | 0.4151 |
| cluster3348 | -4559.91177 | 0.0480 | 0.0342 | 0.7131 | 0.0529 | 0.0128 | 0.2418 |
| cluster3349 | -4215.00995 | 0.0100 | 0.0105 | 1.0505 | 0.0081 | 0.0059 | 0.7275 |
| cluster3350 | -4778.90911 | 0.0494 | 0.0760 | 1.5393 | 0.0509 | 0.0282 | 0.5539 |
| cluster3351 | -3134.21908 | 0.0158 | 0.0251 | 1.5877 | 0.0166 | 0.0053 | 0.3169 |
| cluster3352 | -4085.14942 | 0.0539 | 0.0529 | 0.9820 | 0.0606 | 0.0156 | 0.2578 |
| cluster3353 | -1585.12549 | 0.0133 | 0.0109 | 0.8193 | 0.0037 | 0.0006 | 0.1651 |
| cluster3354 | -2120.14051 | 0.0034 | 0.0021 | 0.6159 | 0.0102 | 0.0016 | 0.1613 |
| cluster3355 | -4279.20099 | 0.0263 | 0.0358 | 1.3639 | 0.0640 | 0.0346 | 0.5409 |
| cluster3356 | -4595.31866 | 0.0466 | 0.0260 | 0.5573 | 0.0353 | 0.0166 | 0.4721 |
| cluster3357 | -1289.93239 | 0.0247 | 0.0236 | 0.9529 | 0.0333 | 0.0099 | 0.2963 |
| cluster3358 | -4169.99937 | 0.0005 | 0.0002 | 0.4645 | 0.0028 | 0.0014 | 0.4780 |
| cluster3359 | -6041.42957 | 0.0290 | 0.0777 | 2.6806 | 0.0463 | 0.0265 | 0.5728 |
| cluster3360 | -2471.60075 | 0.0357 | 0.0290 | 0.8117 | 0.0402 | 0.0320 | 0.7976 |
| cluster3361 | -5034.47694 | 0.0307 | 0.0811 | 2.6412 | 0.0433 | 0.0304 | 0.7010 |
| cluster3362 | -3692.63783 | 0.0177 | 0.0137 | 0.7773 | 0.0147 | 0.0089 | 0.6103 |
| cluster3363 | -3721.11771 | 0.0022 | 0.0118 | 5.3396 | 0.0026 | 0.0016 | 0.6231 |
| cluster3364 | -4356.02854 | 0.0213 | 0.0517 | 2.4320 | 0.0263 | 0.0208 | 0.7913 |
| cluster3365 | -5565.94115 | 0.0641 | 0.0846 | 1.3205 | 0.0598 | 0.0252 | 0.4210 |
| cluster3366 | -3148.10292 | 0.0224 | 0.1025 | 4.5799 | 0.0403 | 0.0231 | 0.5738 |
| cluster3367 | -3693.75774 | 0.1470 | 0.0665 | 0.4527 | 0.1250 | 0.0771 | 0.6165 |
| cluster3368 | -1651.95797 | 0.0384 | 0.0109 | 0.2849 | 0.0076 | 0.0011 | 0.1468 |
| cluster3369 | -3766.19553 | 0.0143 | 0.0261 | 1.8193 | 0.0166 | 0.0107 | 0.6457 |
| cluster3370 | -6387.91283 | 0.0985 | 0.1183 | 1.2009 | 0.1003 | 0.0391 | 0.3901 |
| cluster3371 | -2934.33942 | 0.0055 | 0.0037 | 0.6788 | 0.0078 | 0.0016 | 0.2030 |
| cluster3372 | -3841.52549 | 0.0086 | 0.0097 | 1.1176 | 0.0087 | 0.0089 | 1.0290 |
| cluster3373 | -5277.24550 | 0.0094 | 0.0213 | 2.2757 | 0.0074 | 0.0048 | 0.6448 |
| cluster3374 | -5125.25492 | 0.0663 | 0.1797 | 2.7099 | 0.0770 | 0.0570 | 0.7400 |
| cluster3375 | -3052.20271 | 0.0057 | 0.0026 | 0.4647 | 0.0049 | 0.0015 | 0.3053 |
| cluster3376 | -1524.20721 | 0.0094 | 0.0027 | 0.2924 | 0.0713 | 0.0153 | 0.2147 |
| cluster3377 | -2272.62711 | 0.0087 | 0.0226 | 2.5893 | 0.0176 | 0.0087 | 0.4946 |
| cluster3378 | -6506.63214 | 0.0396 | 0.0378 | 0.9536 | 0.0585 | 0.0485 | 0.8302 |

| | | | | | | | |
|---|---|---|---|---|---|---|---|
| cluster3379 | -1566.85971 | 0.0249 | 0.0250 | 1.0059 | 0.0502 | 0.0146 | 0.2900 |
| cluster3380 | -2653.27155 | 0.0085 | 0.0338 | 3.9579 | 0.0139 | 0.0041 | 0.2944 |
| cluster3381 | -2267.93805 | 0.0365 | 0.0197 | 0.5385 | 0.0310 | 0.0212 | 0.6837 |
| cluster3382 | -885.65465 | 0.0054 | 0.0038 | 0.6939 | 0.0087 | 0.0017 | 0.1937 |
| cluster3383 | -5137.61458 | 0.0225 | 0.0960 | 4.2758 | 0.0864 | 0.0517 | 0.5983 |
| cluster3384 | -1965.04175 | 0.0139 | 0.0098 | 0.7079 | 0.0594 | 0.0269 | 0.4533 |
| cluster3385 | -3771.68364 | 0.0223 | 0.0412 | 1.8434 | 0.0216 | 0.0120 | 0.5565 |
| cluster3386 | -4154.23335 | 0.0112 | 0.0239 | 2.1279 | 0.0193 | 0.0100 | 0.5207 |
| cluster3387 | -3020.45953 | 0.0215 | 0.0262 | 1.2240 | 0.0745 | 0.0278 | 0.3735 |
| cluster3388 | -2807.65351 | 0.0191 | 0.0200 | 1.0439 | 0.0338 | 0.0145 | 0.4286 |
| cluster3389 | -1788.91307 | 0.0094 | 0.0228 | 2.4321 | 0.0108 | 0.0041 | 0.3832 |
| cluster3390 | -5117.08365 | 0.0609 | 0.0423 | 0.6944 | 0.0673 | 0.0320 | 0.4760 |
| cluster3391 | -2416.11028 | 0.0072 | 0.0061 | 0.8534 | 0.0109 | 0.0027 | 0.2489 |
| cluster3392 | -2525.39063 | 0.0154 | 0.0253 | 1.6399 | 0.0053 | 0.0035 | 0.6617 |
| cluster3393 | -3179.48629 | 0.0603 | 0.0738 | 1.2236 | 0.0551 | 0.0200 | 0.3627 |
| cluster3394 | -712.73628 | 0.0001 | 0.0001 | 0.5969 | 0.0060 | 0.0052 | 0.8689 |
| cluster3395 | -2713.46108 | 0.0102 | 0.0057 | 0.5591 | 0.0185 | 0.0134 | 0.7220 |
| cluster3396 | -4982.71751 | 0.0468 | 0.0251 | 0.5353 | 0.0272 | 0.0148 | 0.5437 |
| cluster3397 | -4641.23784 | 0.0241 | 0.0518 | 2.1516 | 0.0332 | 0.0207 | 0.6220 |
| cluster3398 | -3833.67754 | 0.0576 | 0.0507 | 0.8809 | 0.0523 | 0.0114 | 0.2187 |
| cluster3399 | -3229.21879 | 0.0328 | 0.0386 | 1.1746 | 0.0588 | 0.0341 | 0.5800 |
| cluster3400 | -1943.288274 | 0.0245 | 0.0039 | 0.1601 | 0.1816 | 0.0345 | 0.1898 |
| cluster3401 | -2868.99905 | 0.0120 | 0.0218 | 1.8061 | 0.0237 | 0.0075 | 0.3179 |
| cluster3402 | -5111.01880 | 0.0315 | 0.0240 | 0.7634 | 0.0157 | 0.0091 | 0.5779 |
| cluster3403 | -3668.59706 | 0.0185 | 0.0203 | 1.0985 | 0.0274 | 0.0118 | 0.4292 |
| cluster3404 | -716.696126 | 0.0001 | 0.0000 | 0.1596 | 0.0001 | 0.0000 | 0.0540 |
| cluster3405 | -1537.216503 | 0.1701 | 0.0889 | 0.5226 | 0.0643 | 0.0162 | 0.2524 |
| cluster3406 | -3800.13888 | 0.0225 | 0.0177 | 0.7868 | 0.0163 | 0.0168 | 1.0332 |
| cluster3407 | -6244.24146 | 0.0564 | 0.0560 | 0.9917 | 0.0648 | 0.0586 | 0.9034 |
| cluster3408 | -1896.69506 | 0.0063 | 0.0021 | 0.3399 | 0.0061 | 0.0014 | 0.2341 |
| cluster3409 | -4909.34237 | 0.0253 | 0.0539 | 2.1287 | 0.0215 | 0.0152 | 0.7046 |
| cluster3410 | -1442.77299 | 0.0293 | 0.0103 | 0.3510 | 0.0182 | 0.0049 | 0.2678 |
| cluster3411 | -2582.73902 | 0.0004 | 0.0005 | 1.2243 | 0.0001 | 0.0000 | 0.2379 |
| cluster3412 | -3389.45318 | 0.1083 | 0.1188 | 1.0972 | 0.0594 | 0.0236 | 0.3966 |
| cluster3413 | -2551.79985 | 0.0232 | 0.0397 | 1.7129 | 0.0818 | 0.0334 | 0.4087 |
| cluster3414 | -2522.913637 | 0.1421 | 0.0334 | 0.2348 | 0.1301 | 0.0277 | 0.2129 |
| cluster3415 | -3489.07947 | 0.0469 | 0.0372 | 0.7934 | 0.0914 | 0.0242 | 0.2649 |
| cluster3416 | -598.05622 | 0.0001 | 0.0002 | 1.7473 | 0.2662 | 0.5401 | 2.0290 |
| cluster3417 | -1215.22065 | 0.0035 | 0.0210 | 5.9838 | 0.0010 | 0.0117 | 11.7520 |
| cluster3418 | -2534.20389 | 0.0517 | 0.0643 | 1.2432 | 0.0584 | 0.0278 | 0.4755 |
| cluster3419 | -2670.153676 | 0.0756 | 0.0402 | 0.5314 | 0.1481 | 0.0199 | 0.1345 |
| cluster3420 | -1780.847549 | 0.1639 | 0.1394 | 0.8506 | 0.4137 | 0.0775 | 0.1873 |
| cluster3421 | -4207.39267 | 0.0095 | 0.0219 | 2.3093 | 0.0129 | 0.0077 | 0.5957 |
| cluster3422 | -5398.67425 | 0.0399 | 0.0307 | 0.7710 | 0.0297 | 0.0146 | 0.4932 |

| | | | | | | | |
|---|---|---|---|---|---|---|---|
| cluster3423 | -4241.25649 | 0.0261 | 0.1037 | 3.9682 | 0.0785 | 0.0505 | 0.6438 |
| cluster3424 | -2854.05861 | 0.0763 | 0.0702 | 0.9208 | 0.1496 | 0.0523 | 0.3498 |
| cluster3425 | -2213.17927 | 0.0106 | 0.0055 | 0.5220 | 0.1082 | 0.0205 | 0.1896 |
| cluster3426 | -3409.06874 | 0.0264 | 0.0266 | 1.0083 | 0.0131 | 0.0148 | 1.1344 |
| cluster3427 | -3796.49366 | 0.0860 | 0.0704 | 0.8178 | 0.1010 | 0.0309 | 0.3058 |
| cluster3428 | -2685.53127 | 0.0740 | 0.0269 | 0.3643 | 0.0562 | 0.0253 | 0.4500 |
| cluster3429 | -4143.81674 | 0.0368 | 0.0298 | 0.8095 | 0.0223 | 0.0167 | 0.7480 |
| cluster3430 | -4824.72273 | 0.0444 | 0.1012 | 2.2783 | 0.0429 | 0.0394 | 0.9167 |
| cluster3431 | -4738.02156 | 0.0334 | 0.0835 | 2.4989 | 0.0495 | 0.0323 | 0.6532 |
| cluster3432 | -3782.77619 | 0.0370 | 0.0184 | 0.4977 | 0.0367 | 0.0150 | 0.4092 |
| cluster3433 | -5390.50723 | 0.1366 | 0.0754 | 0.5518 | 0.2094 | 0.0536 | 0.2561 |
| cluster3434 | -3295.16156 | 0.0358 | 0.0143 | 0.3979 | 0.0372 | 0.0132 | 0.3542 |
| cluster3435 | -4093.33712 | 0.0221 | 0.0318 | 1.4437 | 0.0202 | 0.0104 | 0.5156 |
| cluster3436 | -3135.33072 | 0.0238 | 0.0541 | 2.2744 | 0.0229 | 0.0115 | 0.5027 |
| cluster3437 | -2816.28211 | 0.0109 | 0.0121 | 1.1096 | 0.0151 | 0.0122 | 0.8095 |
| cluster3438 | -3823.95503 | 0.0327 | 0.0381 | 1.1656 | 0.0463 | 0.0182 | 0.3923 |
| cluster3439 | -2056.30004 | 0.0479 | 0.0732 | 1.5300 | 0.0592 | 0.0166 | 0.2805 |
| cluster3440 | -2314.33224 | 0.0279 | 0.0166 | 0.5934 | 0.0227 | 0.0058 | 0.2558 |
| cluster3441 | -2754.76712 | 0.0278 | 0.0591 | 2.1275 | 0.0542 | 0.0316 | 0.5823 |
| cluster3442 | -2920.01616 | 0.0338 | 0.0907 | 2.6804 | 0.0102 | 0.0187 | 1.8450 |
| cluster3443 | -3531.12120 | 0.0117 | 0.0253 | 2.1663 | 0.0206 | 0.0100 | 0.4861 |
| cluster3444 | -3386.69545 | 0.0108 | 0.0129 | 1.1973 | 0.0086 | 0.0024 | 0.2824 |
| cluster3445 | -2961.69552 | 0.0218 | 0.0096 | 0.4403 | 0.0067 | 0.0027 | 0.4063 |
| cluster3446 | -4061.67512 | 0.0276 | 0.0560 | 2.0301 | 0.0352 | 0.0205 | 0.5819 |
| cluster3447 | -5921.42339 | 0.0230 | 0.1013 | 4.3993 | 0.0528 | 0.0403 | 0.7628 |
| cluster3448 | -2378.62461 | 0.0259 | 0.0598 | 2.3129 | 0.0196 | 0.0088 | 0.4480 |
| cluster3449 | -613.12828 | 0.0061 | 0.0009 | 0.1564 | 0.0001 | 0.0000 | 0.1423 |
| cluster3450 | -2516.89814 | 0.0142 | 0.0442 | 3.1103 | 0.0284 | 0.0160 | 0.5627 |
| cluster3451 | -3594.17637 | 0.0290 | 0.0549 | 1.8978 | 0.0260 | 0.0127 | 0.4875 |
| cluster3452 | -3963.45707 | 0.0556 | 0.0842 | 1.5145 | 0.0370 | 0.0207 | 0.5592 |
| cluster3453 | -1845.35409 | 0.0671 | 0.0229 | 0.3414 | 0.0214 | 0.0128 | 0.5965 |
| cluster3454 | -3695.73880 | 0.0092 | 0.0305 | 3.3303 | 0.0118 | 0.0106 | 0.8991 |
| cluster3455 | -3649.31611 | 0.0427 | 0.0126 | 0.2961 | 0.0445 | 0.0098 | 0.2213 |
| cluster3456 | -4229.67716 | 0.0436 | 0.0277 | 0.6348 | 0.0267 | 0.0069 | 0.2594 |
| cluster3457 | -3649.26905 | 0.0333 | 0.0509 | 1.5258 | 0.0280 | 0.0156 | 0.5555 |
| cluster3458 | -1805.96274 | 0.0274 | 0.0150 | 0.5472 | 0.0420 | 0.0183 | 0.4364 |
| cluster3459 | -2638.01903 | 0.0291 | 0.1045 | 3.5924 | 0.0645 | 0.0297 | 0.4609 |
| cluster3460 | -5579.56446 | 0.0268 | 0.0779 | 2.9039 | 0.0380 | 0.0341 | 0.8964 |
| cluster3461 | -3102.92604 | 0.0664 | 0.0987 | 1.4855 | 0.0657 | 0.0316 | 0.4816 |
| cluster3462 | -823.848522 | 0.0217 | 0.0027 | 0.1265 | 0.0533 | 0.0039 | 0.0739 |
| cluster3463 | -4771.28951 | 0.0506 | 0.0719 | 1.4204 | 0.0447 | 0.0251 | 0.5610 |
| cluster3464 | -5602.85421 | 0.0321 | 0.1056 | 3.2897 | 0.0633 | 0.0387 | 0.6112 |
| cluster3465 | -3954.89010 | 0.0130 | 0.0270 | 2.0791 | 0.0077 | 0.0119 | 1.5497 |
| cluster3466 | -2644.06745 | 0.0104 | 0.0084 | 0.8091 | 0.0143 | 0.0133 | 0.9245 |

| | | | | | | | |
|---|---|---|---|---|---|---|---|
| cluster3467 | -4058.03648 | 0.0248 | 0.0320 | 1.2880 | 0.0450 | 0.0191 | 0.4247 |
| cluster3468 | -1849.82352 | 0.0068 | 0.0262 | 3.8725 | 0.0088 | 0.0050 | 0.5676 |
| cluster3469 | -2105.80693 | 0.0211 | 0.0294 | 1.3979 | 0.0248 | 0.0112 | 0.4523 |
| cluster3470 | -3884.29172 | 0.0064 | 0.0243 | 3.7666 | 0.0154 | 0.0067 | 0.4315 |
| cluster3471 | -4173.29060 | 0.0410 | 0.0587 | 1.4315 | 0.0452 | 0.0213 | 0.4718 |
| cluster3472 | -3721.69023 | 0.0205 | 0.0131 | 0.6399 | 0.0154 | 0.0091 | 0.5906 |
| cluster3473 | -3470.37924 | 0.0230 | 0.0143 | 0.6223 | 0.0314 | 0.0237 | 0.7566 |
| cluster3474 | -5365.33334 | 0.0254 | 0.0923 | 3.6339 | 0.0406 | 0.0392 | 0.9660 |
| cluster3475 | -2339.13756 | 0.0074 | 0.0220 | 2.9900 | 0.0172 | 0.0090 | 0.5220 |
| cluster3476 | -5108.80522 | 0.0212 | 0.0902 | 4.2516 | 0.0395 | 0.0405 | 1.0241 |
| cluster3477 | -3277.08867 | 0.0179 | 0.0053 | 0.2956 | 0.0129 | 0.0029 | 0.2262 |
| cluster3478 | -3582.10716 | 0.0189 | 0.0120 | 0.6352 | 0.0200 | 0.0127 | 0.6341 |
| cluster3479 | -2915.63467 | 0.0389 | 0.1092 | 2.8117 | 0.0591 | 0.0519 | 0.8793 |
| cluster3480 | -4508.59869 | 0.0297 | 0.0517 | 1.7382 | 0.0383 | 0.0233 | 0.6082 |
| cluster3481 | -1069.88718 | 0.0149 | 0.0054 | 0.3617 | 0.0015 | 0.0388 | 25.6613 |
| cluster3482 | -1960.08783 | 0.0403 | 0.0159 | 0.3943 | 0.1484 | 0.0220 | 0.1485 |
| cluster3483 | -3656.54545 | 0.0065 | 0.0112 | 1.7343 | 0.0167 | 0.0069 | 0.4134 |
| cluster3484 | -2765.20790 | 0.0187 | 0.0430 | 2.2978 | 0.0476 | 0.0242 | 0.5071 |
| cluster3485 | -3489.34841 | 0.0272 | 0.0539 | 1.9784 | 0.0699 | 0.0167 | 0.2383 |
| cluster3486 | -3174.90592 | 0.0461 | 0.0177 | 0.3844 | 0.0555 | 0.0175 | 0.3144 |
| cluster3487 | -2347.86587 | 0.0202 | 0.0265 | 1.3121 | 0.0285 | 0.0134 | 0.4710 |
| cluster3488 | -1483.40842 | 0.0271 | 0.0251 | 0.9251 | 0.0290 | 0.0078 | 0.2690 |
| cluster3489 | -4058.86175 | 0.0323 | 0.0720 | 2.2293 | 0.0610 | 0.0335 | 0.5484 |
| cluster3490 | -3432.61184 | 0.0183 | 0.0524 | 2.8690 | 0.0257 | 0.0201 | 0.7827 |
| cluster3491 | -3686.63162 | 0.0273 | 0.0410 | 1.5010 | 0.0368 | 0.0149 | 0.4033 |
| cluster3492 | -3371.44078 | 0.0453 | 0.0827 | 1.8271 | 0.0295 | 0.0196 | 0.6647 |
| cluster3493 | -4433.27650 | 0.0329 | 0.0436 | 1.3254 | 0.0367 | 0.0122 | 0.3321 |
| cluster3494 | -2865.76349 | 0.0106 | 0.0275 | 2.6088 | 0.0155 | 0.0269 | 1.7373 |
| cluster3495 | -2840.93680 | 0.0217 | 0.0325 | 1.5010 | 0.0421 | 0.0138 | 0.3272 |
| cluster3496 | -3230.97215 | 0.0269 | 0.0158 | 0.5895 | 0.0182 | 0.0100 | 0.5473 |
| cluster3497 | -1194.87154 | 0.0134 | 0.0025 | 0.1840 | 0.0136 | 0.0022 | 0.1614 |
| cluster3498 | -4134.13304 | 0.0468 | 0.1317 | 2.8164 | 0.0496 | 0.0354 | 0.7130 |
| cluster3499 | -3715.70328 | 0.0461 | 0.1341 | 2.9106 | 0.1602 | 0.0667 | 0.4162 |
| cluster3500 | -3416.95998 | 0.0096 | 0.0302 | 3.1430 | 0.0360 | 0.0143 | 0.3970 |
| cluster3501 | -2358.31923 | 0.0373 | 0.0199 | 0.5339 | 0.0199 | 0.0171 | 0.8600 |
| cluster3502 | -2167.78725 | 0.0100 | 0.0207 | 2.0612 | 0.0203 | 0.0122 | 0.6001 |
| cluster3503 | -1951.06464 | 0.0316 | 0.0886 | 2.8033 | 0.0583 | 0.0237 | 0.4073 |
| cluster3504 | -1141.94590 | 0.0095 | 0.0180 | 1.9001 | 0.0062 | 0.0096 | 1.5536 |
| cluster3505 | -2413.90045 | 0.0315 | 0.0955 | 3.0293 | 0.0838 | 0.0550 | 0.6561 |
| cluster3506 | -2549.97053 | 0.0220 | 0.0422 | 1.9185 | 0.0258 | 0.0115 | 0.4448 |
| cluster3507 | -5016.78743 | 0.0414 | 0.0739 | 1.7840 | 0.0715 | 0.0379 | 0.5307 |
| cluster3508 | -5630.88536 | 0.0211 | 0.1138 | 5.4052 | 0.0405 | 0.0459 | 1.1332 |
| cluster3509 | -2407.43721 | 0.0433 | 0.0543 | 1.2533 | 0.0001 | 0.0000 | 0.1228 |
| cluster3510 | -834.629659 | 9.0000 | 0.0585 | 0.0001 | 0.0413 | 0.0312 | 0.7572 |

| | | | | | | | |
|---|---|---|---|---|---|---|---|
| cluster3511 | -1016.55126 | 0.0184 | 0.0227 | 1.2352 | 0.0602 | 0.0182 | 0.3027 |
| cluster3512 | -2356.43545 | 0.0375 | 0.0244 | 0.6522 | 0.0262 | 0.0070 | 0.2681 |
| cluster3513 | -959.354097 | 0.0686 | 0.0089 | 0.1301 | 0.0751 | 0.0037 | 0.0490 |
| cluster3514 | -5158.11903 | 0.0425 | 0.0281 | 0.6609 | 0.0313 | 0.0203 | 0.6479 |
| cluster3515 | -2602.95335 | 0.0185 | 0.0222 | 1.1999 | 0.0071 | 0.0031 | 0.4410 |
| cluster3516 | -4129.71610 | 0.0237 | 0.0344 | 1.4515 | 0.0327 | 0.0166 | 0.5096 |
| cluster3517 | -2410.32150 | 0.0027 | 0.0022 | 0.8287 | 0.0021 | 0.0004 | 0.1786 |
| cluster3518 | -4783.04394 | 0.0624 | 0.1137 | 1.8219 | 0.0907 | 0.0433 | 0.4775 |
| cluster3519 | -2127.10279 | 0.0383 | 0.0491 | 1.2818 | 0.0451 | 0.0231 | 0.5122 |
| cluster3520 | -1193.429552 | 0.1793 | 0.1174 | 0.6548 | 0.1172 | 0.0247 | 0.2106 |
| cluster3521 | -4221.98999 | 0.0289 | 0.0530 | 1.8355 | 0.0493 | 0.0394 | 0.8008 |
| cluster3522 | -4753.64986 | 0.0294 | 0.0836 | 2.8427 | 0.0370 | 0.0257 | 0.6932 |
| cluster3523 | -1654.60429 | 0.0190 | 0.0410 | 2.1623 | 0.0474 | 0.0262 | 0.5529 |
| cluster3524 | -1960.91412 | 0.0068 | 0.0041 | 0.6007 | 0.0243 | 0.0045 | 0.1865 |
| cluster3525 | -2817.12775 | 0.0475 | 0.0408 | 0.8582 | 0.0431 | 0.0109 | 0.2535 |
| cluster3526 | -2995.84764 | 0.0009 | 0.0049 | 5.5115 | 0.0055 | 0.0025 | 0.4584 |
| cluster3527 | -4936.25634 | 0.0388 | 0.1064 | 2.7459 | 0.0577 | 0.0455 | 0.7878 |
| cluster3528 | -1963.24725 | 0.0031 | 0.0016 | 0.5297 | 0.0201 | 0.0085 | 0.4232 |
| cluster3529 | -3740.56486 | 0.0233 | 0.0783 | 3.3634 | 0.0415 | 0.0214 | 0.5157 |
| cluster3530 | -1954.24261 | 0.0322 | 0.0749 | 2.3255 | 0.0284 | 0.0156 | 0.5485 |
| cluster3531 | -2915.87830 | 0.0349 | 0.0529 | 1.5169 | 0.0797 | 0.0208 | 0.2613 |
| cluster3532 | -5557.44427 | 0.0705 | 0.0519 | 0.7364 | 0.0565 | 0.0581 | 1.0274 |
| cluster3533 | -2655.21747 | 0.0259 | 0.0122 | 0.4702 | 0.0233 | 0.0087 | 0.3717 |
| cluster3534 | -2559.50310 | 0.0801 | 0.0207 | 0.2586 | 0.0508 | 0.0129 | 0.2535 |
| cluster3535 | -3289.69741 | 0.0248 | 0.0893 | 3.5992 | 0.0193 | 0.0139 | 0.7192 |
| cluster3536 | -3556.36039 | 0.0440 | 0.0229 | 0.5191 | 0.0297 | 0.0200 | 0.6731 |
| cluster3537 | -3665.57258 | 0.0067 | 0.0141 | 2.1081 | 0.0070 | 0.0034 | 0.4861 |
| cluster3538 | -1384.71735 | 0.0081 | 0.0018 | 0.2238 | 0.0531 | 0.0199 | 0.3736 |
| cluster3539 | -2945.33001 | 0.0297 | 0.1458 | 4.9037 | 0.0408 | 0.0408 | 0.9990 |
| cluster3540 | -5364.16316 | 0.0935 | 0.1879 | 2.0091 | 0.0883 | 0.0624 | 0.7073 |
| cluster3541 | -3877.47752 | 0.0306 | 0.0210 | 0.6841 | 0.0243 | 0.0155 | 0.6377 |
| cluster3542 | -5426.75169 | 0.0520 | 0.1538 | 2.9582 | 0.0693 | 0.0541 | 0.7814 |
| cluster3543 | -4242.07313 | 0.0214 | 0.0967 | 4.5283 | 0.0533 | 0.0327 | 0.6132 |
| cluster3544 | -1087.83205 | 0.0083 | 0.0043 | 0.5193 | 0.0245 | 0.0050 | 0.2054 |
| cluster3545 | -5146.63081 | 0.0414 | 0.0895 | 2.1629 | 0.0526 | 0.0364 | 0.6923 |
| cluster3546 | -2676.80675 | 0.0143 | 0.0550 | 3.8460 | 0.0334 | 0.0214 | 0.6407 |
| cluster3547 | -1116.47701 | 0.0170 | 0.0025 | 0.1475 | 0.0049 | 0.0014 | 0.2927 |
| cluster3548 | -1295.86349 | 0.0039 | 0.0007 | 0.1820 | 0.1394 | 0.0172 | 0.1231 |
| cluster3549 | -3232.71033 | 0.0451 | 0.0287 | 0.6364 | 0.0327 | 0.0207 | 0.6353 |
| cluster3550 | -1223.54681 | 0.0051 | 0.0019 | 0.3646 | 0.0140 | 0.0019 | 0.1322 |
| cluster3551 | -2798.86413 | 0.0492 | 0.0139 | 0.2816 | 0.0518 | 0.0163 | 0.3138 |
| cluster3552 | -2385.97942 | 0.0258 | 0.0369 | 1.4286 | 0.0203 | 0.0238 | 1.1710 |
| cluster3553 | -2404.58138 | 0.0273 | 0.0294 | 1.0750 | 0.0412 | 0.0164 | 0.3991 |
| cluster3554 | -3824.30523 | 0.0276 | 0.0298 | 1.0774 | 0.0244 | 0.0062 | 0.2544 |

| | | | | | | | |
|---|---|---|---|---|---|---|---|
| cluster3555 | -1628.35836 | 0.0080 | 0.0222 | 2.7695 | 0.0479 | 0.0251 | 0.5237 |
| cluster3556 | -4203.80576 | 0.0193 | 0.0465 | 2.4119 | 0.0345 | 0.0206 | 0.5975 |
| cluster3557 | -1366.880566 | 0.0781 | 0.1647 | 2.1089 | 0.0241 | 0.0889 | 3.6879 |
| cluster3558 | -4776.35523 | 0.0582 | 0.0901 | 1.5497 | 0.0755 | 0.0471 | 0.6244 |
| cluster3559 | -2456.48203 | 0.0087 | 0.0329 | 3.7859 | 0.0238 | 0.0091 | 0.3822 |
| cluster3560 | -5094.00766 | 0.0458 | 0.1132 | 2.4730 | 0.0743 | 0.0486 | 0.6537 |
| cluster3561 | -1798.05664 | 0.0185 | 0.0114 | 0.6192 | 0.0608 | 0.0110 | 0.1803 |
| cluster3562 | -4275.57915 | 0.0290 | 0.0585 | 2.0196 | 0.0501 | 0.0268 | 0.5349 |
| cluster3563 | -1981.72854 | 0.0208 | 0.0126 | 0.6042 | 0.0103 | 0.0070 | 0.6766 |
| cluster3564 | -5023.92269 | 0.0759 | 0.0368 | 0.4849 | 0.0656 | 0.0350 | 0.5338 |
| cluster3565 | -3341.11306 | 0.0149 | 0.0574 | 3.8431 | 0.0633 | 0.0335 | 0.5303 |
| cluster3566 | -1884.89428 | 0.0228 | 0.0393 | 1.7266 | 0.0744 | 0.0253 | 0.3406 |
| cluster3567 | -2403.90768 | 0.0706 | 0.0935 | 1.3232 | 0.0635 | 0.0258 | 0.4069 |
| cluster3568 | -3168.27080 | 0.0278 | 0.0680 | 2.4452 | 0.0291 | 0.0185 | 0.6365 |
| cluster3569 | -3551.63320 | 0.0403 | 0.1050 | 2.6051 | 0.0407 | 0.0279 | 0.6854 |
| cluster3570 | -2400.10573 | 0.0148 | 0.0732 | 4.9591 | 0.0247 | 0.0252 | 1.0197 |
| cluster3571 | -2034.81854 | 0.0124 | 0.0124 | 1.0007 | 0.0324 | 0.0263 | 0.8101 |
| cluster3572 | -3453.56956 | 0.0448 | 0.0831 | 1.8532 | 0.0556 | 0.0335 | 0.6019 |
| cluster3573 | -3244.16933 | 0.0098 | 0.0276 | 2.8292 | 0.0138 | 0.0120 | 0.8668 |
| cluster3574 | -5131.78846 | 0.0779 | 0.1193 | 1.5312 | 0.0746 | 0.0498 | 0.6681 |
| cluster3575 | -3831.18385 | 0.0392 | 0.0654 | 1.6717 | 0.0404 | 0.0204 | 0.5049 |
| cluster3576 | -3274.17902 | 0.0305 | 0.0612 | 2.0093 | 0.0453 | 0.0275 | 0.6082 |
| cluster3577 | -2731.39519 | 0.0380 | 0.0221 | 0.5817 | 0.0410 | 0.0161 | 0.3916 |
| cluster3578 | -3387.52857 | 0.0200 | 0.0405 | 2.0295 | 0.0466 | 0.0234 | 0.5024 |
| cluster3579 | -2924.07653 | 0.0235 | 0.0234 | 0.9929 | 0.0284 | 0.0309 | 1.0897 |
| cluster3580 | -2417.92659 | 0.0107 | 0.0311 | 2.9154 | 0.0099 | 0.0095 | 0.9605 |
| cluster3581 | -1247.07799 | 0.0080 | 0.0245 | 3.0699 | 0.0073 | 0.0030 | 0.4062 |
| cluster3582 | -1104.98877 | 0.0048 | 0.0009 | 0.1875 | 0.0419 | 0.0068 | 0.1627 |
| cluster3583 | -2533.43616 | 0.0001 | 0.0002 | 1.7219 | 0.0001 | 0.0000 | 0.3718 |
| cluster3584 | -4869.86751 | 0.0304 | 0.1063 | 3.4938 | 0.0312 | 0.0290 | 0.9300 |
| cluster3585 | -2093.61821 | 0.0497 | 0.0188 | 0.3773 | 0.0782 | 0.0235 | 0.3003 |
| cluster3586 | -5223.29750 | 0.0590 | 0.1057 | 1.7933 | 0.0870 | 0.0632 | 0.7265 |
| cluster3587 | -3463.03615 | 0.0244 | 0.1156 | 4.7368 | 0.0874 | 0.0609 | 0.6961 |
| cluster3588 | -3358.44821 | 0.0178 | 0.0092 | 0.5155 | 0.0379 | 0.0108 | 0.2849 |
| cluster3589 | -3011.26222 | 0.0257 | 0.0421 | 1.6387 | 0.0770 | 0.0272 | 0.3527 |
| cluster3590 | -4180.48765 | 0.0451 | 0.1503 | 3.3309 | 0.0735 | 0.0622 | 0.8451 |
| cluster3591 | -2874.40101 | 0.0125 | 0.0055 | 0.4386 | 0.0115 | 0.0055 | 0.4754 |
| cluster3592 | -4705.69447 | 0.0348 | 0.0886 | 2.5482 | 0.0438 | 0.0306 | 0.6983 |
| cluster3593 | -3415.22070 | 0.0288 | 0.0865 | 3.0059 | 0.0710 | 0.0487 | 0.6854 |
| cluster3594 | -3170.78793 | 0.0348 | 0.0568 | 1.6303 | 0.0964 | 0.0250 | 0.2588 |
| cluster3595 | -2123.57502 | 0.0106 | 0.0053 | 0.4994 | 0.0134 | 0.0062 | 0.4633 |
| cluster3596 | -2316.26217 | 0.0323 | 0.0407 | 1.2588 | 0.0339 | 0.0151 | 0.4462 |
| cluster3597 | -4193.27922 | 0.0238 | 0.0228 | 0.9566 | 0.0370 | 0.0218 | 0.5908 |
| cluster3598 | -4253.68880 | 0.0265 | 0.1138 | 4.2999 | 0.0203 | 0.0854 | 4.2093 |

| | | | | | | | |
|---|---|---|---|---|---|---|---|
| cluster3599 | -2252.31510 | 0.0185 | 0.0314 | 1.7010 | 0.0344 | 0.0171 | 0.4958 |
| cluster3600 | -2860.35219 | 0.0209 | 0.0247 | 1.1834 | 0.0333 | 0.0083 | 0.2489 |
| cluster3601 | -3280.30497 | 0.0019 | 0.0165 | 8.4981 | 0.0106 | 0.0081 | 0.7633 |
| cluster3602 | -3065.51986 | 0.0045 | 0.0018 | 0.4098 | 0.0023 | 0.0010 | 0.4242 |
| cluster3603 | -4224.35849 | 0.0565 | 0.0875 | 1.5475 | 0.0476 | 0.0251 | 0.5259 |
| cluster3604 | -1058.11484 | 0.0736 | 0.0102 | 0.1388 | 0.0505 | 0.0039 | 0.0775 |
| cluster3605 | -3057.86491 | 0.0370 | 0.0521 | 1.4061 | 0.0391 | 0.0220 | 0.5638 |
| cluster3606 | -4323.61015 | 0.0269 | 0.0671 | 2.4985 | 0.0286 | 0.0186 | 0.6506 |
| cluster3607 | -2594.69192 | 0.0214 | 0.0578 | 2.7070 | 0.0507 | 0.0298 | 0.5870 |
| cluster3608 | -2888.08117 | 0.0097 | 0.0069 | 0.7146 | 0.0117 | 0.0072 | 0.6140 |
| cluster3609 | -4173.00001 | 0.0148 | 0.0720 | 4.8733 | 0.0504 | 0.0362 | 0.7185 |
| cluster3610 | -1393.50280 | 0.0113 | 0.0099 | 0.8764 | 0.0011 | 0.0292 | 25.8007 |
| cluster3611 | -2340.58829 | 0.0228 | 0.1142 | 5.0117 | 0.0303 | 0.0298 | 0.9831 |
| cluster3612 | -2828.60633 | 0.0188 | 0.0885 | 4.6998 | 0.0497 | 0.0353 | 0.7103 |
| cluster3613 | -4059.04629 | 0.0206 | 0.0116 | 0.5616 | 0.0165 | 0.0119 | 0.7205 |
| cluster3614 | -4308.14678 | 0.0235 | 0.0999 | 4.2485 | 0.0265 | 0.0300 | 1.1334 |
| cluster3615 | -1663.11705 | 0.0403 | 0.0071 | 0.1760 | 0.0535 | 0.0113 | 0.2116 |
| cluster3616 | -770.27387 | 0.0020 | 0.0124 | 6.1494 | 0.0375 | 0.0097 | 0.2600 |
| cluster3617 | -1925.22662 | 0.0209 | 0.0498 | 2.3848 | 0.0742 | 0.0384 | 0.5177 |
| cluster3618 | -1898.959729 | 0.1423 | 0.1609 | 1.1303 | 0.2000 | 0.0565 | 0.2823 |
| cluster3619 | -2551.18763 | 0.0013 | 0.0082 | 6.2659 | 0.0061 | 0.0021 | 0.3426 |
| cluster3620 | -3603.77129 | 0.0399 | 0.0525 | 1.3155 | 0.0634 | 0.0282 | 0.4444 |
| cluster3621 | -4764.31818 | 0.0276 | 0.0908 | 3.2909 | 0.0399 | 0.0346 | 0.8680 |
| cluster3622 | -2715.06940 | 0.0037 | 0.0027 | 0.7410 | 0.0268 | 0.0080 | 0.2969 |
| cluster3623 | -2271.74869 | 0.0094 | 0.0461 | 4.8835 | 0.0087 | 0.0107 | 1.2320 |
| cluster3624 | -2909.87685 | 0.0124 | 0.0527 | 4.2524 | 0.0149 | 0.0084 | 0.5621 |
| cluster3625 | -3307.30031 | 0.0027 | 0.0216 | 7.8989 | 0.0086 | 0.0048 | 0.5511 |
| cluster3626 | -1980.65010 | 0.0451 | 0.0212 | 0.4695 | 0.0186 | 0.0155 | 0.8359 |
| cluster3627 | -3001.00939 | 0.0068 | 0.0113 | 1.6683 | 0.0080 | 0.0042 | 0.5220 |
| cluster3628 | -3023.09466 | 0.0270 | 0.0808 | 2.9923 | 0.0286 | 0.0266 | 0.9298 |
| cluster3629 | -4163.49701 | 0.0085 | 0.0408 | 4.8003 | 0.0167 | 0.0162 | 0.9664 |
| cluster3630 | -1208.99095 | 0.0083 | 0.0039 | 0.4689 | 0.0244 | 0.0053 | 0.2166 |
| cluster3631 | -3592.33665 | 0.0101 | 0.0139 | 1.3793 | 0.0280 | 0.0249 | 0.8918 |
| cluster3632 | -2460.00620 | 0.0318 | 0.0484 | 1.5233 | 0.0300 | 0.0437 | 1.4558 |
| cluster3633 | -2502.32498 | 0.0282 | 0.0258 | 0.9138 | 0.0036 | 0.0039 | 1.0980 |
| cluster3634 | -936.24467 | 0.0213 | 0.0173 | 0.8137 | 0.0001 | 0.0000 | 0.1531 |
| cluster3635 | -1620.14107 | 0.0300 | 0.0061 | 0.2024 | 0.0597 | 0.0085 | 0.1424 |
| cluster3636 | -2531.27835 | 0.0195 | 0.0158 | 0.8096 | 0.0112 | 0.0118 | 1.0572 |
| cluster3637 | -1057.22944 | 0.0023 | 0.0012 | 0.5027 | 0.0509 | 0.0053 | 0.1036 |
| cluster3638 | -2731.13258 | 0.0248 | 0.0277 | 1.1149 | 0.0438 | 0.0107 | 0.2452 |
| cluster3639 | -1272.48533 | 0.0318 | 0.0197 | 0.6209 | 0.0388 | 0.0154 | 0.3956 |
| cluster3640 | -4764.16477 | 0.0557 | 0.1373 | 2.4657 | 0.0934 | 0.0619 | 0.6624 |
| cluster3641 | -1769.09223 | 0.0274 | 0.0079 | 0.2886 | 0.0253 | 0.0058 | 0.2288 |
| cluster3642 | -3356.37743 | 0.0253 | 0.0694 | 2.7498 | 0.0410 | 0.0246 | 0.6005 |

| | | | | | | | |
|---|---|---|---|---|---|---|---|
| cluster3643 | -3537.57202 | 0.0875 | 0.1626 | 1.8580 | 0.1164 | 0.0478 | 0.4110 |
| cluster3644 | -1911.92525 | 0.0214 | 0.0117 | 0.5475 | 0.0397 | 0.0093 | 0.2353 |
| cluster3645 | -1807.05572 | 0.0236 | 0.0489 | 2.0741 | 0.0385 | 0.0229 | 0.5967 |
| cluster3646 | -2634.75095 | 0.0143 | 0.0797 | 5.5632 | 0.0289 | 0.0324 | 1.1201 |
| cluster3647 | -3428.61978 | 0.0060 | 0.0337 | 5.6304 | 0.0025 | 0.0044 | 1.7595 |
| cluster3648 | -1758.71849 | 0.0022 | 0.0019 | 0.8818 | 0.0005 | 0.0006 | 1.1581 |
| cluster3649 | -1640.39301 | 0.0086 | 0.0105 | 1.2211 | 0.0129 | 0.0032 | 0.2448 |
| cluster3650 | -1254.31679 | 0.0024 | 0.0658 | 27.7952 | 0.0300 | 0.0344 | 1.1445 |
| cluster3651 | -3971.05866 | 0.0526 | 0.0403 | 0.7657 | 0.0430 | 0.0319 | 0.7430 |
| cluster3652 | -3309.21548 | 0.0183 | 0.0131 | 0.7163 | 0.0318 | 0.0222 | 0.6961 |
| cluster3653 | -1425.58190 | 0.0013 | 0.0009 | 0.7001 | 0.0065 | 0.0031 | 0.4724 |
| cluster3654 | -2838.01614 | 0.0098 | 0.0227 | 2.3274 | 0.0092 | 0.0056 | 0.6105 |
| cluster3655 | -2366.65624 | 0.0666 | 0.0977 | 1.4673 | 0.0657 | 0.0275 | 0.4178 |
| cluster3656 | -3108.53386 | 0.0284 | 0.0239 | 0.8413 | 0.0227 | 0.0184 | 0.8104 |
| cluster3657 | -3194.44716 | 0.0152 | 0.0658 | 4.3320 | 0.0149 | 0.0086 | 0.5785 |
| cluster3658 | -747.212307 | 0.0187 | 0.0039 | 0.2071 | 0.1762 | 0.0348 | 0.1977 |
| cluster3659 | -2269.78952 | 0.0369 | 0.0301 | 0.8147 | 0.0341 | 0.0117 | 0.3421 |
| cluster3660 | -3014.97746 | 0.0110 | 0.0754 | 6.8538 | 0.0292 | 0.0249 | 0.8550 |
| cluster3661 | -1818.54570 | 0.0034 | 0.0005 | 0.1616 | 0.0001 | 0.0000 | 0.0819 |
| cluster3662 | -2803.16039 | 0.0291 | 0.0594 | 2.0428 | 0.0465 | 0.0263 | 0.5661 |
| cluster3663 | -3134.14922 | 0.0486 | 0.0630 | 1.2947 | 0.0489 | 0.0228 | 0.4666 |
| cluster3664 | -2445.22156 | 0.0235 | 0.0863 | 3.6679 | 0.0614 | 0.0435 | 0.7093 |
| cluster3665 | -3603.20584 | 0.0186 | 0.0438 | 2.3586 | 0.0227 | 0.0186 | 0.8168 |
| cluster3666 | -2711.09979 | 0.0303 | 0.0556 | 1.8369 | 0.0410 | 0.0165 | 0.4028 |
| cluster3667 | -1570.44336 | 0.0177 | 0.0091 | 0.5124 | 0.0280 | 0.0043 | 0.1546 |
| cluster3668 | -2983.09264 | 0.0176 | 0.0320 | 1.8144 | 0.0254 | 0.0139 | 0.5458 |
| cluster3669 | -2601.60696 | 0.0406 | 0.0470 | 1.1585 | 0.0306 | 0.0095 | 0.3107 |
| cluster3670 | -4034.91157 | 0.0448 | 0.0403 | 0.8988 | 0.0471 | 0.0394 | 0.8359 |
| cluster3671 | -3995.99566 | 0.0249 | 0.0608 | 2.4403 | 0.0342 | 0.0229 | 0.6689 |
| cluster3672 | -2500.07346 | 0.0373 | 0.0642 | 1.7216 | 0.0538 | 0.0234 | 0.4348 |
| cluster3673 | -4408.74634 | 0.0315 | 0.1808 | 5.7393 | 0.0510 | 0.0628 | 1.2317 |
| cluster3674 | -2025.35751 | 0.0298 | 0.0641 | 2.1496 | 0.0248 | 0.0257 | 1.0343 |
| cluster3675 | -660.30852 | 0.0001 | 0.0000 | 0.1317 | 0.0001 | 0.0000 | 0.1494 |
| cluster3676 | -3519.32269 | 0.0326 | 0.0735 | 2.2553 | 0.0832 | 0.0402 | 0.4835 |
| cluster3677 | -812.08913 | 0.0227 | 0.0142 | 0.6258 | 0.0217 | 0.0088 | 0.4078 |
| cluster3678 | -3307.19957 | 0.0307 | 0.0439 | 1.4299 | 0.0391 | 0.0205 | 0.5252 |
| cluster3679 | -944.53537 | 0.0067 | 0.0029 | 0.4369 | 0.0104 | 0.0037 | 0.3606 |
| cluster3680 | -2314.16536 | 0.0111 | 0.0413 | 3.7254 | 0.0077 | 0.0075 | 0.9736 |
| cluster3681 | -2959.25881 | 0.0196 | 0.0524 | 2.6751 | 0.0195 | 0.0177 | 0.9037 |
| cluster3682 | -4355.10161 | 0.0182 | 0.0713 | 3.9075 | 0.0597 | 0.0337 | 0.5648 |
| cluster3683 | -2804.88177 | 0.0883 | 0.0774 | 0.8770 | 0.1267 | 0.1158 | 0.9142 |
| cluster3684 | -2991.25532 | 0.0253 | 0.0498 | 1.9682 | 0.0452 | 0.0249 | 0.5510 |
| cluster3685 | -3672.54835 | 0.0663 | 0.1608 | 2.4262 | 0.1118 | 0.0653 | 0.5842 |
| cluster3686 | -1622.31749 | 0.0763 | 0.1531 | 2.0069 | 0.0016 | 0.0276 | 16.8088 |

| | | | | | | | |
|---|---|---|---|---|---|---|---|
| cluster3687 | -2877.54338 | 0.0753 | 0.2614 | 3.4690 | 0.0901 | 0.0521 | 0.5777 |
| cluster3688 | -2048.39585 | 0.0032 | 0.0265 | 8.1852 | 0.0133 | 0.0072 | 0.5396 |
| cluster3689 | -2642.10455 | 0.0397 | 0.0300 | 0.7558 | 0.0382 | 0.0161 | 0.4207 |
| cluster3690 | -1454.90639 | 0.0220 | 0.0139 | 0.6325 | 0.0244 | 0.0196 | 0.8048 |
| cluster3691 | -670.03487 | 0.0020 | 0.0020 | 1.0327 | 0.0080 | 0.0040 | 0.5025 |
| cluster3692 | -1434.87199 | 0.0091 | 0.0191 | 2.1069 | 0.0148 | 0.0083 | 0.5625 |
| cluster3693 | -2464.67432 | 0.0078 | 0.0317 | 4.0851 | 0.0112 | 0.0100 | 0.8928 |
| cluster3694 | -2090.40361 | 0.0193 | 0.0059 | 0.3088 | 0.0413 | 0.0138 | 0.3329 |
| cluster3695 | -1737.01807 | 0.0231 | 0.0389 | 1.6810 | 0.0272 | 0.0130 | 0.4786 |
| cluster3696 | -813.66271 | 0.0179 | 0.0576 | 3.2118 | 0.0004 | 0.0134 | 30.4555 |
| cluster3697 | -3584.13897 | 0.0748 | 0.0385 | 0.5141 | 0.0700 | 0.0422 | 0.6030 |
| cluster3698 | -1099.57239 | 0.0001 | 0.0000 | 0.3100 | 0.0001 | 0.0000 | 0.2646 |
| cluster3699 | -4440.58183 | 0.0395 | 0.1086 | 2.7526 | 0.0558 | 0.0374 | 0.6711 |
| cluster3700 | -2819.00265 | 0.0301 | 0.0449 | 1.4901 | 0.0347 | 0.0209 | 0.6034 |
| cluster3701 | -1127.62172 | 0.0031 | 0.0010 | 0.3129 | 0.0001 | 0.0000 | 0.4429 |
| cluster3702 | -3862.40062 | 0.0172 | 0.0858 | 4.9911 | 0.0348 | 0.0250 | 0.7178 |
| cluster3703 | -1511.61703 | 0.0104 | 0.0050 | 0.4819 | 0.0264 | 0.0070 | 0.2633 |
| cluster3704 | -2229.00616 | 0.0102 | 0.0182 | 1.7896 | 0.0039 | 0.0068 | 1.7720 |
| cluster3705 | -1942.44417 | 0.0067 | 0.0057 | 0.8593 | 0.0253 | 0.0109 | 0.4320 |
| cluster3706 | -2094.65573 | 0.0209 | 0.0247 | 1.1825 | 0.0211 | 0.0053 | 0.2496 |
| cluster3707 | -1921.43820 | 0.0393 | 0.1009 | 2.5719 | 0.0445 | 0.0322 | 0.7237 |
| cluster3708 | -2632.34906 | 0.0233 | 0.0544 | 2.3338 | 0.0706 | 0.0226 | 0.3195 |
| cluster3709 | -1579.74860 | 0.0924 | 0.0369 | 0.3991 | 0.0570 | 0.0081 | 0.1425 |
| cluster3710 | -1922.04587 | 0.0070 | 0.0359 | 5.1148 | 0.0221 | 0.0156 | 0.7076 |
| cluster3711 | -1431.87069 | 0.0108 | 0.0048 | 0.4455 | 0.0861 | 0.0082 | 0.0949 |
| cluster3712 | -3402.33824 | 0.0302 | 0.0275 | 0.9095 | 0.0229 | 0.0105 | 0.4611 |
| cluster3713 | -4025.69722 | 0.0347 | 0.1193 | 3.4358 | 0.0724 | 0.0464 | 0.6413 |
| cluster3714 | -1775.53374 | 0.0395 | 0.0157 | 0.3988 | 0.0540 | 0.0183 | 0.3394 |
| cluster3715 | -1285.42457 | 0.0076 | 0.0631 | 8.3294 | 0.0248 | 0.0275 | 1.1068 |
| cluster3716 | -3657.58815 | 0.0404 | 0.0870 | 2.1554 | 0.0535 | 0.0339 | 0.6332 |
| cluster3717 | -1403.84010 | 0.0117 | 0.0033 | 0.2804 | 0.0165 | 0.0040 | 0.2429 |
| cluster3718 | -2989.76659 | 0.0153 | 0.0577 | 3.7665 | 0.0260 | 0.0145 | 0.5567 |
| cluster3719 | -3292.71722 | 0.0321 | 0.0486 | 1.5112 | 0.0621 | 0.0233 | 0.3748 |
| cluster3720 | -2755.92580 | 0.0182 | 0.0280 | 1.5340 | 0.0427 | 0.0119 | 0.2794 |
| cluster3721 | -1098.04369 | 0.0045 | 0.0009 | 0.1888 | 0.0057 | 0.0007 | 0.1274 |
| cluster3722 | -1224.36502 | 0.0318 | 0.1393 | 4.3871 | 0.0667 | 0.0588 | 0.8809 |
| cluster3723 | -989.94266 | 0.0001 | 0.0001 | 0.6368 | 0.0001 | 0.0000 | 0.2590 |
| cluster3724 | -3082.21881 | 0.0266 | 0.0615 | 2.3150 | 0.0426 | 0.0234 | 0.5495 |
| cluster3725 | -1839.85633 | 0.0114 | 0.0308 | 2.7009 | 0.0092 | 0.0059 | 0.6456 |
| cluster3726 | -1935.59399 | 0.0387 | 0.0327 | 0.8453 | 0.0615 | 0.0235 | 0.3820 |
| cluster3727 | -2467.95306 | 0.0413 | 0.1271 | 3.0754 | 0.0666 | 0.0311 | 0.4660 |
| cluster3728 | -2157.06679 | 0.0219 | 0.0191 | 0.8741 | 0.0202 | 0.0105 | 0.5203 |
| cluster3729 | -2947.33120 | 0.0130 | 0.0329 | 2.5404 | 0.0135 | 0.0083 | 0.6169 |
| cluster3730 | -1777.73744 | 0.0434 | 0.0147 | 0.3400 | 0.1441 | 0.0282 | 0.1955 |

| | | | | | | | |
|---|---|---|---|---|---|---|---|
| cluster3731 | -2865.30216 | 0.0077 | 0.0056 | 0.7186 | 0.0105 | 0.0056 | 0.5365 |
| cluster3732 | -2822.89362 | 0.0216 | 0.0292 | 1.3536 | 0.0243 | 0.0100 | 0.4127 |
| cluster3733 | -3069.54833 | 0.0561 | 0.0295 | 0.5269 | 0.1063 | 0.0382 | 0.3595 |
| cluster3734 | -3043.44553 | 0.0424 | 0.0662 | 1.5608 | 0.0443 | 0.0270 | 0.6097 |
| cluster3735 | -3250.36926 | 0.0812 | 0.0363 | 0.4468 | 0.0803 | 0.0374 | 0.4661 |
| cluster3736 | -3537.11041 | 0.0537 | 0.0448 | 0.8354 | 0.0371 | 0.0149 | 0.4009 |
| cluster3737 | -1209.47833 | 0.0023 | 0.0010 | 0.4363 | 0.0043 | 0.0007 | 0.1582 |
| cluster3738 | -3257.53961 | 0.0137 | 0.0102 | 0.7446 | 0.0057 | 0.0046 | 0.7995 |
| cluster3739 | -1156.25352 | 0.0001 | 0.0000 | 0.4279 | 0.0001 | 0.0000 | 0.1404 |
| cluster3740 | -3447.88238 | 0.0224 | 0.0461 | 2.0586 | 0.0406 | 0.0223 | 0.5483 |
| cluster3741 | -1312.77908 | 0.0159 | 0.0107 | 0.6751 | 0.0177 | 0.0046 | 0.2589 |
| cluster3742 | -1615.45067 | 0.0474 | 0.0715 | 1.5081 | 0.0662 | 0.0322 | 0.4871 |
| cluster3743 | -2980.01467 | 0.0212 | 0.0464 | 2.1884 | 0.0280 | 0.0125 | 0.4477 |
| cluster3744 | -1353.835801 | 0.0691 | 0.0777 | 1.1245 | 0.1603 | 0.0284 | 0.1773 |
| cluster3745 | -2584.96547 | 0.0211 | 0.0463 | 2.1962 | 0.0304 | 0.0207 | 0.6810 |
| cluster3746 | -3783.79913 | 0.0275 | 0.0365 | 1.3249 | 0.0559 | 0.0357 | 0.6382 |
| cluster3747 | -1120.84000 | 0.0318 | 0.0352 | 1.1094 | 0.0199 | 0.0076 | 0.3840 |
| cluster3748 | -2165.78889 | 0.0099 | 0.0045 | 0.4558 | 0.0183 | 0.0096 | 0.5285 |
| cluster3749 | -3069.55063 | 0.0113 | 0.0344 | 3.0368 | 0.0242 | 0.0127 | 0.5255 |
| cluster3750 | -1354.76695 | 0.0183 | 0.0752 | 4.1139 | 0.0339 | 0.0255 | 0.7529 |
| cluster3751 | -3011.98700 | 0.0045 | 0.0074 | 1.6638 | 0.0049 | 0.0023 | 0.4602 |
| cluster3752 | -2662.59436 | 0.0059 | 0.0226 | 3.8476 | 0.0113 | 0.0068 | 0.6037 |
| cluster3753 | -1023.374497 | 0.0195 | 0.0011 | 0.0548 | 0.0195 | 0.0028 | 0.1454 |
| cluster3754 | -881.97724 | 0.0001 | 0.0000 | 0.2745 | 0.0001 | 0.0000 | 0.0896 |
| cluster3755 | -2499.78479 | 0.0164 | 0.0153 | 0.9316 | 0.0149 | 0.0109 | 0.7342 |
| cluster3756 | -1689.78263 | 0.0348 | 0.0450 | 1.2956 | 0.0366 | 0.0204 | 0.5567 |
| cluster3757 | -2091.38241 | 0.0456 | 0.0328 | 0.7197 | 0.0453 | 0.0112 | 0.2468 |
| cluster3758 | -1511.28576 | 0.0353 | 0.0236 | 0.6689 | 0.0223 | 0.0093 | 0.4157 |
| cluster3759 | -3062.83066 | 0.0463 | 0.0249 | 0.5389 | 0.0970 | 0.0495 | 0.5100 |
| cluster3760 | -2600.72362 | 0.0255 | 0.0312 | 1.2246 | 0.0203 | 0.0187 | 0.9216 |
| cluster3761 | -767.74720 | 0.0073 | 0.0492 | 6.7197 | 0.0031 | 0.0065 | 2.1370 |
| cluster3762 | -1966.49649 | 0.0085 | 0.0024 | 0.2835 | 0.0225 | 0.0066 | 0.2935 |
| cluster3763 | -2003.28254 | 0.0357 | 0.0124 | 0.3483 | 0.0822 | 0.0166 | 0.2021 |
| cluster3764 | -1629.52159 | 0.0253 | 0.0651 | 2.5713 | 0.0358 | 0.0275 | 0.7690 |
| cluster3765 | -2668.46726 | 0.0301 | 0.0256 | 0.8508 | 0.0119 | 0.0082 | 0.6888 |
| cluster3766 | -1794.69342 | 0.0068 | 0.0206 | 3.0167 | 0.0160 | 0.0065 | 0.4096 |
| cluster3767 | -861.47223 | 0.0085 | 0.0464 | 5.4489 | 0.0365 | 0.0252 | 0.6902 |
| cluster3768 | -2218.21235 | 0.0853 | 0.1269 | 1.4880 | 0.0376 | 0.0128 | 0.3407 |
| cluster3769 | -1764.87369 | 0.0050 | 0.0022 | 0.4410 | 0.0105 | 0.0020 | 0.1894 |
| cluster3770 | -1774.96630 | 0.0477 | 0.0660 | 1.3838 | 0.0588 | 0.0213 | 0.3632 |
| cluster3771 | -2138.50037 | 0.0859 | 0.1197 | 1.3935 | 0.0611 | 0.0239 | 0.3911 |
| cluster3772 | -725.44836 | 0.0080 | 0.0439 | 5.5247 | 0.0001 | 0.0000 | 0.1720 |
| cluster3773 | -1114.46689 | 0.2929 | 0.0918 | 0.3135 | 0.0368 | 0.0018 | 0.0501 |
| cluster3774 | -1409.06723 | 0.0121 | 0.0043 | 0.3573 | 0.0169 | 0.0018 | 0.1042 |

| | | | | | | | |
|---|---|---|---|---|---|---|---|
| cluster3775 | -906.72299 | 0.0122 | 0.0402 | 3.3022 | 0.0135 | 0.0066 | 0.4896 |
| cluster3776 | -2518.35922 | 0.0882 | 0.0330 | 0.3748 | 0.0671 | 0.0278 | 0.4141 |
| cluster3777 | -3064.24328 | 0.0341 | 0.0870 | 2.5552 | 0.0785 | 0.0476 | 0.6061 |
| cluster3778 | -980.71862 | 0.0015 | 0.0010 | 0.6523 | 0.0010 | 0.0009 | 0.8952 |
| cluster3779 | -2995.04813 | 0.0245 | 0.0465 | 1.9004 | 0.0202 | 0.0211 | 1.0482 |
| cluster3780 | -1683.84635 | 0.0069 | 0.0353 | 5.1232 | 0.0180 | 0.0134 | 0.7424 |
| cluster3781 | -2686.31482 | 0.0612 | 0.0416 | 0.6802 | 0.0538 | 0.0318 | 0.5925 |
| cluster3782 | -2290.00340 | 0.0377 | 0.0364 | 0.9647 | 0.0495 | 0.0193 | 0.3893 |
| cluster3783 | -2107.89479 | 0.0055 | 0.0549 | 9.9864 | 0.0088 | 0.0106 | 1.2051 |
| cluster3784 | -2951.78987 | 0.0469 | 0.0360 | 0.7685 | 0.0498 | 0.0507 | 1.0190 |
| cluster3785 | -2687.77252 | 0.0125 | 0.0129 | 1.0328 | 0.0240 | 0.0080 | 0.3352 |
| cluster3786 | -2262.61676 | 0.0032 | 0.0087 | 2.7427 | 0.0750 | 0.0354 | 0.4714 |
| cluster3787 | -1519.68553 | 0.0167 | 0.0236 | 1.4131 | 0.0434 | 0.0189 | 0.4351 |
| cluster3788 | -1160.401549 | 0.0001 | 0.0000 | 0.1768 | 0.0001 | 0.0000 | 0.0195 |
| cluster3789 | -3039.25662 | 0.0048 | 0.0053 | 1.1223 | 0.0292 | 0.0053 | 0.1821 |
| cluster3790 | -2174.02907 | 0.0058 | 0.0479 | 8.2068 | 0.0223 | 0.0118 | 0.5279 |
| cluster3791 | -419.57167 | 0.0062 | 0.0030 | 0.4954 | 0.0205 | 0.0060 | 0.2918 |
| cluster3792 | -1863.28824 | 0.0053 | 0.0040 | 0.7654 | 0.0071 | 0.0023 | 0.3185 |
| cluster3793 | -1850.41442 | 0.0151 | 0.0259 | 1.7076 | 0.0445 | 0.0151 | 0.3398 |
| cluster3794 | -2188.25833 | 0.0119 | 0.0283 | 2.3864 | 0.0226 | 0.0126 | 0.5590 |
| cluster3795 | -1700.96874 | 0.0143 | 0.0313 | 2.1821 | 0.0260 | 0.0084 | 0.3218 |
| cluster3796 | -1249.343531 | 0.0862 | 0.1005 | 1.1657 | 0.1988 | 0.0534 | 0.2687 |
| cluster3797 | -3909.93777 | 0.0377 | 0.2115 | 5.6037 | 0.0739 | 0.0540 | 0.7299 |
| cluster3798 | -3442.93444 | 0.0155 | 0.0344 | 2.2270 | 0.0355 | 0.0215 | 0.6069 |
| cluster3799 | -2340.89021 | 0.0384 | 0.0325 | 0.8470 | 0.0581 | 0.0333 | 0.5742 |
| cluster3800 | -429.878172 | 0.0528 | 0.0022 | 0.0412 | 0.0001 | 0.0000 | 0.1126 |
| cluster3801 | -2689.046772 | 0.2508 | 0.1580 | 0.6299 | 0.1568 | 0.0371 | 0.2365 |
| cluster3802 | -2649.93749 | 0.0348 | 0.0483 | 1.3879 | 0.0333 | 0.0373 | 1.1210 |
| cluster3803 | -2239.57604 | 0.0310 | 0.0427 | 1.3784 | 0.0428 | 0.0162 | 0.3784 |
| cluster3804 | -2635.14891 | 0.0281 | 0.0429 | 1.5259 | 0.0376 | 0.0161 | 0.4285 |
| cluster3805 | -2654.58620 | 0.0174 | 0.0394 | 2.2635 | 0.0309 | 0.0169 | 0.5458 |
| cluster3806 | -3566.82576 | 0.0362 | 0.0637 | 1.7607 | 0.0422 | 0.0267 | 0.6323 |
| cluster3807 | -1441.72020 | 0.0053 | 0.0014 | 0.2632 | 0.0090 | 0.0031 | 0.3430 |
| cluster3808 | -1825.71402 | 0.0357 | 0.0696 | 1.9471 | 0.0266 | 0.0158 | 0.5944 |
| cluster3809 | -2361.19578 | 0.0083 | 0.0138 | 1.6539 | 0.0112 | 0.0039 | 0.3495 |
| cluster3810 | -2464.59329 | 0.0095 | 0.0111 | 1.1673 | 0.0098 | 0.0088 | 0.8958 |
| cluster3811 | -761.377364 | 0.0338 | 0.0099 | 0.2913 | 0.0131 | 0.0015 | 0.1112 |
| cluster3812 | -2999.76080 | 0.0221 | 0.0346 | 1.5665 | 0.0389 | 0.0097 | 0.2500 |
| cluster3813 | -3413.72749 | 0.0342 | 0.1261 | 3.6855 | 0.0358 | 0.0465 | 1.3012 |
| cluster3814 | -1608.071036 | 0.0663 | 0.0135 | 0.2038 | 0.1121 | 0.0214 | 0.1907 |
| cluster3815 | -1707.12017 | 0.0125 | 0.0611 | 4.8988 | 0.0278 | 0.0154 | 0.5539 |
| cluster3816 | -2525.45531 | 0.0205 | 0.0628 | 3.0656 | 0.0353 | 0.0257 | 0.7294 |
| cluster3817 | -990.61991 | 0.0147 | 0.0031 | 0.2085 | 0.0023 | 0.0008 | 0.3673 |
| cluster3818 | -1978.65491 | 0.0524 | 0.0560 | 1.0676 | 0.0733 | 0.0736 | 1.0044 |

| | | | | | | | |
|---|---|---|---|---|---|---|---|
| cluster3819 | -3346.94705 | 0.0247 | 0.0404 | 1.6394 | 0.0312 | 0.0175 | 0.5613 |
| cluster3820 | -2222.54506 | 0.0172 | 0.0386 | 2.2502 | 0.0454 | 0.0171 | 0.3755 |
| cluster3821 | -2243.65311 | 0.0177 | 0.0157 | 0.8861 | 0.0146 | 0.0097 | 0.6657 |
| cluster3822 | -3395.28842 | 0.0166 | 0.0584 | 3.5210 | 0.0419 | 0.0231 | 0.5512 |
| cluster3823 | -3407.75658 | 0.0151 | 0.0271 | 1.7923 | 0.0253 | 0.0260 | 1.0247 |
| cluster3824 | -2341.91570 | 0.0054 | 0.0467 | 8.7124 | 0.0489 | 0.0354 | 0.7243 |
| cluster3825 | -1388.82875 | 0.0058 | 0.0079 | 1.3559 | 0.0077 | 0.0030 | 0.3942 |
| cluster3826 | -3155.18174 | 0.0732 | 0.1426 | 1.9495 | 0.0746 | 0.0491 | 0.6578 |
| cluster3827 | -2403.53963 | 0.0242 | 0.0255 | 1.0537 | 0.0314 | 0.0253 | 0.8074 |
| cluster3828 | -2365.19295 | 0.0460 | 0.0527 | 1.1461 | 0.0510 | 0.0259 | 0.5069 |
| cluster3829 | -2256.70274 | 0.0199 | 0.0466 | 2.3429 | 0.0295 | 0.0160 | 0.5426 |
| cluster3830 | -1622.95609 | 0.0098 | 0.0074 | 0.7599 | 0.0618 | 0.0066 | 0.1073 |
| cluster3831 | -2348.38792 | 0.0289 | 0.0855 | 2.9588 | 0.0348 | 0.0222 | 0.6375 |
| cluster3832 | -1766.81268 | 0.0024 | 0.0032 | 1.3282 | 0.0088 | 0.0032 | 0.3604 |
| cluster3833 | -2130.47629 | 0.0223 | 0.0343 | 1.5386 | 0.0242 | 0.0128 | 0.5294 |
| cluster3834 | -2049.43077 | 0.0011 | 0.0307 | 27.4321 | 0.0251 | 0.0137 | 0.5469 |
| cluster3835 | -2117.89704 | 0.0297 | 0.0414 | 1.3923 | 0.0413 | 0.0166 | 0.4022 |
| cluster3836 | -1895.02033 | 0.0180 | 0.0238 | 1.3238 | 0.0163 | 0.0081 | 0.4957 |
| cluster3837 | -2637.86264 | 0.0324 | 0.0520 | 1.6082 | 0.0549 | 0.0262 | 0.4780 |
| cluster3838 | -2294.23628 | 0.0118 | 0.0176 | 1.4975 | 0.0831 | 0.0320 | 0.3855 |
| cluster3839 | -1492.86541 | 0.0057 | 0.0249 | 4.3740 | 0.0419 | 0.0187 | 0.4470 |
| cluster3840 | -2024.37941 | 0.0030 | 0.0181 | 6.1013 | 0.0054 | 0.0942 | 17.3121 |
| cluster3841 | -2291.94411 | 0.0595 | 0.1490 | 2.5049 | 0.0958 | 0.0547 | 0.5705 |
| cluster3842 | -1604.85197 | 0.0001 | 0.0000 | 0.2473 | 0.0074 | 0.0016 | 0.2130 |
| cluster3843 | -1440.15437 | 0.0147 | 0.0146 | 0.9957 | 0.0049 | 0.0013 | 0.2659 |
| cluster3844 | -1073.431622 | 0.0120 | 0.0477 | 3.9658 | 0.1119 | 0.0214 | 0.1908 |
| cluster3845 | -604.91636 | 0.0001 | 0.0000 | 0.0047 | 0.0001 | 0.0000 | 0.3093 |
| cluster3846 | -1272.90263 | 0.0250 | 0.0204 | 0.8167 | 0.0097 | 0.0065 | 0.6642 |
| cluster3847 | -1156.38813 | 0.0009 | 0.0024 | 2.6490 | 0.0015 | 0.0008 | 0.5488 |
| cluster3848 | -2512.36612 | 0.0185 | 0.0528 | 2.8567 | 0.0536 | 0.0300 | 0.5590 |
| cluster3849 | -2663.41345 | 0.0113 | 0.0687 | 6.0734 | 0.0288 | 0.0383 | 1.3324 |
| cluster3850 | -1677.18818 | 0.0154 | 0.0201 | 1.3040 | 0.0249 | 0.0075 | 0.3018 |
| cluster3851 | -1618.16441 | 0.0103 | 0.0189 | 1.8321 | 0.0087 | 0.0036 | 0.4182 |
| cluster3852 | -1856.04016 | 0.0095 | 0.0132 | 1.3867 | 0.0048 | 0.0094 | 1.9618 |
| cluster3853 | -1101.73961 | 0.0045 | 0.0186 | 4.1225 | 0.0053 | 0.0044 | 0.8240 |
| cluster3854 | -1356.883037 | 0.0683 | 0.0856 | 1.2533 | 0.1414 | 0.0228 | 0.1613 |
| cluster3855 | -2930.40829 | 0.0085 | 0.0473 | 5.6050 | 0.0131 | 0.0183 | 1.3972 |
| cluster3856 | -1694.27134 | 0.0122 | 0.0295 | 2.4197 | 0.0277 | 0.0161 | 0.5804 |
| cluster3857 | -3037.98812 | 0.0157 | 0.0318 | 2.0173 | 0.0178 | 0.0117 | 0.6601 |
| cluster3858 | -2416.70533 | 0.0506 | 0.0416 | 0.8211 | 0.0470 | 0.0334 | 0.7107 |
| cluster3859 | -1976.54087 | 0.0098 | 0.0153 | 1.5690 | 0.0189 | 0.0068 | 0.3599 |
| cluster3860 | -2622.42977 | 0.0562 | 0.0301 | 0.5361 | 0.1174 | 0.0434 | 0.3696 |
| cluster3861 | -1300.57292 | 0.0136 | 0.0195 | 1.4368 | 0.0214 | 0.0164 | 0.7669 |
| cluster3862 | -922.33901 | 0.0001 | 0.0001 | 1.1204 | 0.0001 | 0.0000 | 0.3185 |

| | | | | | | | |
|---|---|---|---|---|---|---|---|
| cluster3863 | -1482.43810 | 0.0001 | 0.0001 | 0.6721 | 0.0018 | 0.0012 | 0.6666 |
| cluster3864 | -2220.07359 | 0.0041 | 0.0145 | 3.5143 | 0.0036 | 0.0024 | 0.6767 |
| cluster3865 | -3004.31564 | 0.0461 | 0.0997 | 2.1622 | 0.0443 | 0.0372 | 0.8411 |
| cluster3866 | -1633.11923 | 0.0147 | 0.0454 | 3.0820 | 0.0224 | 0.0165 | 0.7343 |
| cluster3867 | -608.471137 | 0.0001 | 0.0000 | 0.0943 | 0.0001 | 0.0000 | 0.0420 |
| cluster3868 | -1392.61508 | 0.0204 | 0.0150 | 0.7369 | 0.0418 | 0.0109 | 0.2604 |
| cluster3869 | -2973.29263 | 0.0209 | 0.1204 | 5.7680 | 0.0441 | 0.0439 | 0.9959 |
| cluster3870 | -2461.66441 | 0.0316 | 0.0539 | 1.7031 | 0.0495 | 0.0227 | 0.4594 |
| cluster3871 | -2151.60953 | 0.0096 | 0.0158 | 1.6467 | 0.0040 | 0.0022 | 0.5557 |
| cluster3872 | -2653.99887 | 0.0100 | 0.0448 | 4.4883 | 0.0082 | 0.0063 | 0.7753 |
| cluster3873 | -1785.46810 | 0.0159 | 0.0164 | 1.0314 | 0.0422 | 0.0225 | 0.5334 |
| cluster3874 | -2100.79239 | 0.0175 | 0.0076 | 0.4362 | 0.0064 | 0.0012 | 0.1794 |
| cluster3875 | -769.60663 | 0.0046 | 0.0036 | 0.7808 | 0.0001 | 0.0000 | 0.3286 |
| cluster3876 | -744.42200 | 0.0195 | 0.0101 | 0.5170 | 0.0032 | 0.0012 | 0.3659 |
| cluster3877 | -1433.02123 | 0.0237 | 0.0122 | 0.5160 | 0.0047 | 0.0019 | 0.4039 |
| cluster3879 | -1881.44900 | 0.0171 | 0.0239 | 1.4009 | 0.0189 | 0.0088 | 0.4670 |
| cluster3880 | -1357.09653 | 0.0255 | 0.0416 | 1.6318 | 0.0145 | 0.0109 | 0.7489 |
| cluster3881 | -3016.52667 | 0.0361 | 0.1176 | 3.2565 | 0.0606 | 0.0381 | 0.6291 |
| cluster3882 | -2689.55948 | 0.0200 | 0.0439 | 2.1994 | 0.0314 | 0.0167 | 0.5316 |
| cluster3883 | -422.49962 | 0.0060 | 0.1999 | 33.1093 | 0.0008 | 0.0376 | 48.0854 |
| cluster3884 | -1615.17414 | 0.0003 | 0.0182 | 71.5177 | 2.2637 | 0.0068 | 0.0030 |
| cluster3885 | -2856.30182 | 0.0083 | 0.0191 | 2.3000 | 0.0190 | 0.0117 | 0.6165 |
| cluster3886 | -1393.32244 | 0.0018 | 0.0181 | 10.1316 | 0.0075 | 0.0063 | 0.8305 |
| cluster3887 | -1622.79376 | 0.0010 | 0.0467 | 45.7712 | 0.0162 | 0.0126 | 0.7797 |
| cluster3888 | -1182.20801 | 0.0065 | 0.0619 | 9.5093 | 0.0139 | 0.0223 | 1.6129 |
| cluster3889 | -1004.67821 | 0.0232 | 0.0684 | 2.9462 | 0.0246 | 0.0148 | 0.6001 |
| cluster3890 | -1871.61675 | 0.0139 | 0.0298 | 2.1451 | 0.0209 | 0.0229 | 1.0990 |
| cluster3891 | -2183.90491 | 0.0193 | 0.0136 | 0.7058 | 0.0103 | 0.0098 | 0.9525 |
| cluster3892 | -1374.18394 | 0.0172 | 0.0116 | 0.6719 | 0.0114 | 0.0035 | 0.3094 |
| cluster3893 | -1458.95497 | 0.0243 | 0.0126 | 0.5185 | 0.0257 | 0.0151 | 0.5865 |
| cluster3894 | -965.69181 | 0.0028 | 0.0342 | 12.2437 | 0.0124 | 0.0041 | 0.3330 |
| cluster3895 | -1980.42435 | 0.0010 | 0.0250 | 26.2140 | 0.0149 | 0.0059 | 0.3978 |
| cluster3896 | -818.93448 | 0.0003 | 0.0092 | 32.0625 | 0.0108 | 0.0020 | 0.1889 |
| cluster3897 | -1454.73213 | 0.0042 | 0.0078 | 1.8570 | 0.1155 | 0.0729 | 0.6311 |
| cluster3898 | -2390.23632 | 0.0279 | 0.0089 | 0.3175 | 0.0216 | 0.0064 | 0.2960 |
| cluster3899 | -3001.62580 | 0.0200 | 0.0735 | 3.6783 | 0.0364 | 0.0312 | 0.8557 |
| cluster3900 | -2363.12197 | 0.0409 | 0.1667 | 4.0735 | 0.0945 | 0.0770 | 0.8151 |
| cluster3901 | -1542.34176 | 0.0009 | 0.0021 | 2.4674 | 0.0051 | 0.0034 | 0.6727 |
| cluster3902 | -2493.70709 | 0.0075 | 0.0247 | 3.2837 | 0.0205 | 0.0120 | 0.5884 |
| cluster3903 | -1623.45504 | 0.0083 | 0.0642 | 7.7404 | 0.0115 | 0.0077 | 0.6657 |
| cluster3904 | -905.20283 | 0.0062 | 0.0343 | 5.5762 | 0.0035 | 0.0011 | 0.3255 |
| cluster3905 | -1594.44978 | 0.1080 | 0.1243 | 1.1511 | 0.1528 | 0.0502 | 0.3286 |
| cluster3906 | -1452.39731 | 0.0284 | 0.0343 | 1.2077 | 0.0112 | 0.0036 | 0.3183 |
| cluster3907 | -1944.18144 | 0.0336 | 0.0470 | 1.3981 | 0.0526 | 0.0220 | 0.4185 |

| | | | | | | | |
|---|---|---|---|---|---|---|---|
| cluster3908 | -1848.14091 | 0.0130 | 0.0272 | 2.0910 | 0.0155 | 0.0078 | 0.5021 |
| cluster3909 | -2458.50099 | 0.0110 | 0.0594 | 5.4198 | 0.0462 | 0.0273 | 0.5906 |
| cluster3910 | -1193.43866 | 0.0378 | 0.0833 | 2.2047 | 0.0343 | 0.0252 | 0.7339 |
| cluster3911 | -1748.30604 | 0.0041 | 0.0223 | 5.4168 | 0.0128 | 0.0063 | 0.4925 |
| cluster3912 | -951.76626 | 0.0001 | 0.0000 | 0.3069 | 0.0001 | 0.0000 | 0.3108 |
| cluster3913 | -2544.69609 | 0.0431 | 0.0787 | 1.8253 | 0.0338 | 0.0183 | 0.5397 |
| cluster3914 | -2239.07118 | 0.0603 | 0.1279 | 2.1226 | 0.0527 | 0.0418 | 0.7924 |
| cluster3915 | -2227.40734 | 0.0359 | 0.0638 | 1.7768 | 0.0328 | 0.0210 | 0.6402 |
| cluster3916 | -902.575357 | 0.0721 | 0.0740 | 1.0272 | 0.0898 | 0.0369 | 0.4112 |
| cluster3917 | -2582.19748 | 0.0162 | 0.0360 | 2.2293 | 0.0083 | 0.0031 | 0.3744 |
| cluster3918 | -2755.44108 | 0.0099 | 0.0847 | 8.5364 | 0.0286 | 0.0315 | 1.1027 |
| cluster3919 | -569.756043 | 0.0001 | 0.0000 | 0.2478 | 0.0027 | 0.0011 | 0.3905 |
| cluster3920 | -656.44190 | 0.0052 | 0.1675 | 32.0826 | 0.0319 | 0.0485 | 1.5201 |
| cluster3921 | -2189.91794 | 0.0201 | 0.0386 | 1.9229 | 0.0320 | 0.0164 | 0.5120 |
| cluster3922 | -1240.27666 | 0.0500 | 0.0645 | 1.2894 | 0.0521 | 0.0207 | 0.3983 |
| cluster3923 | -1150.12288 | 0.0005 | 0.0023 | 4.4851 | 0.0001 | 0.0001 | 0.8479 |
| cluster3924 | -1696.29554 | 0.0195 | 0.0352 | 1.8048 | 0.0200 | 0.0105 | 0.5253 |
| cluster3925 | -1086.33302 | 0.0181 | 0.0349 | 1.9273 | 0.1134 | 0.0359 | 0.3169 |
| cluster3926 | -2437.34429 | 0.0523 | 0.1474 | 2.8177 | 0.0428 | 0.0405 | 0.9469 |
| cluster3927 | -1187.72237 | 0.0076 | 0.0217 | 2.8499 | 0.0140 | 0.0072 | 0.5101 |
| cluster3928 | -1159.09630 | 0.0364 | 0.0308 | 0.8461 | 0.0164 | 0.0033 | 0.2026 |
| cluster3929 | -1608.15803 | 0.0066 | 0.0162 | 2.4649 | 0.0053 | 0.0037 | 0.6991 |
| cluster3930 | -2016.85510 | 0.0114 | 0.0081 | 0.7068 | 0.0215 | 0.0090 | 0.4189 |
| cluster3931 | -1795.384051 | 0.0033 | 0.0008 | 0.2506 | 0.3412 | 0.0603 | 0.1769 |
| cluster3932 | -1117.89854 | 0.0113 | 0.0035 | 0.3108 | 0.0107 | 0.0023 | 0.2098 |
| cluster3933 | -1157.74369 | 0.1076 | 0.0578 | 0.5371 | 0.0628 | 0.0115 | 0.1824 |
| cluster3934 | -717.10761 | 0.0001 | 0.0001 | 0.9944 | 0.0001 | 0.0000 | 0.1244 |
| cluster3935 | -3101.79832 | 0.0288 | 0.1209 | 4.2050 | 0.0896 | 0.0556 | 0.6211 |
| cluster3936 | -2125.55760 | 0.0060 | 0.0311 | 5.1846 | 0.0367 | 0.0216 | 0.5891 |
| cluster3937 | -1938.91416 | 0.0176 | 0.0117 | 0.6641 | 0.0224 | 0.0113 | 0.5059 |
| cluster3938 | -1782.25404 | 0.0344 | 0.0342 | 0.9939 | 0.0295 | 0.0093 | 0.3167 |
| cluster3939 | -2055.91869 | 0.0166 | 0.0337 | 2.0306 | 0.0140 | 0.0067 | 0.4819 |
| cluster3940 | -1338.47669 | 0.0198 | 0.0088 | 0.4456 | 0.0157 | 0.0065 | 0.4140 |
| cluster3941 | -968.37343 | 0.0406 | 0.0220 | 0.5402 | 0.0075 | 0.0008 | 0.1120 |
| cluster3942 | -2009.55483 | 0.0297 | 0.0270 | 0.9079 | 0.0337 | 0.0112 | 0.3316 |
| cluster3943 | -2267.69231 | 0.0114 | 0.0626 | 5.5138 | 0.0202 | 0.0137 | 0.6787 |
| cluster3944 | -1588.09383 | 0.0252 | 0.0557 | 2.2117 | 0.0812 | 0.0311 | 0.3825 |
| cluster3945 | -1676.58422 | 0.0415 | 0.0764 | 1.8396 | 0.0344 | 0.0167 | 0.4861 |
| cluster3946 | -1805.98983 | 0.0234 | 0.0476 | 2.0329 | 0.0293 | 0.0124 | 0.4245 |
| cluster3947 | -2285.98484 | 0.0268 | 0.0582 | 2.1683 | 0.0305 | 0.0199 | 0.6528 |
| cluster3948 | -1015.48582 | 0.0244 | 0.0221 | 0.9064 | 0.0398 | 0.0144 | 0.3617 |
| cluster3949 | -878.23321 | 0.0069 | 0.0408 | 5.8983 | 0.0097 | 0.0035 | 0.3645 |
| cluster3950 | -1739.93603 | 0.0058 | 0.0272 | 4.6562 | 0.0175 | 0.0214 | 1.2251 |
| cluster3951 | -2832.95951 | 0.0312 | 0.0642 | 2.0553 | 0.0503 | 0.0302 | 0.6008 |

| | | | | | | | |
|---|---|---|---|---|---|---|---|
| cluster3952 | -1474.67175 | 0.0024 | 0.0088 | 3.7054 | 0.0044 | 0.0033 | 0.7398 |
| cluster3953 | -1362.68331 | 0.0039 | 0.0235 | 6.0115 | 0.0091 | 0.0025 | 0.2746 |
| cluster3954 | -1805.44602 | 0.0217 | 0.0303 | 1.3975 | 0.0388 | 0.0137 | 0.3530 |
| cluster3955 | -1192.14158 | 0.0273 | 0.0580 | 2.1210 | 0.0221 | 0.0145 | 0.6539 |
| cluster3956 | -1995.32775 | 0.0122 | 0.0372 | 3.0534 | 0.0182 | 0.0124 | 0.6813 |
| cluster3957 | -1958.20839 | 0.0083 | 0.0296 | 3.5695 | 0.0179 | 0.0130 | 0.7303 |
| cluster3958 | -1104.33948 | 0.0300 | 0.0460 | 1.5359 | 0.0164 | 0.0061 | 0.3739 |
| cluster3959 | -2686.14195 | 0.0323 | 0.0222 | 0.6867 | 0.0357 | 0.0224 | 0.6261 |
| cluster3960 | -2418.04329 | 0.0184 | 0.1058 | 5.7425 | 0.1045 | 0.0302 | 0.2891 |
| cluster3961 | -1593.76440 | 0.0476 | 0.0785 | 1.6486 | 0.0241 | 0.0146 | 0.6048 |
| cluster3962 | -1532.75306 | 0.0344 | 0.0589 | 1.7139 | 0.0528 | 0.0196 | 0.3706 |
| cluster3963 | -2076.87312 | 0.0310 | 0.0527 | 1.6994 | 0.0534 | 0.0296 | 0.5537 |
| cluster3964 | -2988.38510 | 0.0080 | 0.2560 | 31.8909 | 0.1334 | 0.1031 | 0.7728 |
| cluster3965 | -1515.62907 | 0.0231 | 0.0812 | 3.5181 | 0.0532 | 0.0350 | 0.6593 |
| cluster3966 | -1432.69749 | 0.0204 | 0.0223 | 1.0973 | 0.0300 | 0.0101 | 0.3380 |
| cluster3967 | -1753.77832 | 0.0806 | 0.0375 | 0.4650 | 0.0952 | 0.0303 | 0.3179 |
| cluster3968 | -1167.58351 | 0.0156 | 0.0239 | 1.5347 | 0.0206 | 0.0091 | 0.4423 |
| cluster3969 | -899.56818 | 0.0001 | 0.0000 | 0.4811 | 0.0001 | 0.0000 | 0.1348 |
| cluster3970 | -2482.76106 | 0.0344 | 0.0832 | 2.4178 | 0.0605 | 0.0332 | 0.5490 |
| cluster3971 | -1686.82755 | 0.0153 | 0.0600 | 3.9329 | 0.0538 | 0.0229 | 0.4260 |
| cluster3972 | -990.19221 | 0.0131 | 0.0252 | 1.9216 | 0.0154 | 0.0079 | 0.5103 |
| cluster3973 | -1420.10089 | 0.0092 | 0.0150 | 1.6350 | 0.0100 | 0.0012 | 0.1215 |
| cluster3974 | -1832.45074 | 0.0152 | 0.0226 | 1.4905 | 0.0220 | 0.0115 | 0.5242 |
| cluster3975 | -1269.55199 | 0.0390 | 0.0462 | 1.1833 | 0.0147 | 0.0069 | 0.4674 |
| cluster3976 | -1579.45662 | 0.0057 | 0.0156 | 2.7262 | 0.0319 | 0.0103 | 0.3231 |
| cluster3977 | -2230.15921 | 0.0977 | 0.1196 | 1.2242 | 0.0971 | 0.0443 | 0.4565 |
| cluster3978 | -885.02767 | 0.0014 | 0.0015 | 1.0758 | 0.0179 | 0.0040 | 0.2259 |
| cluster3979 | -1621.10949 | 0.0025 | 0.0025 | 1.0296 | 0.0018 | 0.0013 | 0.7134 |
| cluster3980 | -922.24867 | 0.0001 | 0.0000 | 0.1606 | 0.0001 | 0.0000 | 0.2004 |
| cluster3981 | -1980.31481 | 0.0141 | 0.0338 | 2.3972 | 0.0269 | 0.0127 | 0.4712 |
| cluster3982 | -1829.76196 | 0.0454 | 0.0685 | 1.5108 | 0.0550 | 0.0254 | 0.4619 |
| cluster3983 | -2677.25124 | 0.0305 | 0.0973 | 3.1884 | 0.0369 | 0.0330 | 0.8942 |
| cluster3984 | -1379.33073 | 0.0169 | 0.1010 | 5.9818 | 0.0744 | 0.0467 | 0.6282 |
| cluster3985 | -978.74979 | 0.0164 | 0.0076 | 0.4664 | 0.0076 | 0.0029 | 0.3844 |
| cluster3986 | -1732.46561 | 0.0073 | 0.0453 | 6.1906 | 0.0273 | 0.0202 | 0.7391 |
| cluster3987 | -2085.97565 | 0.0183 | 0.0221 | 1.2064 | 0.0082 | 0.0071 | 0.8691 |
| cluster3988 | -1231.92303 | 0.0001 | 0.0001 | 1.3777 | 0.0309 | 0.0095 | 0.3060 |
| cluster3989 | -1835.03929 | 0.0530 | 0.0418 | 0.7887 | 0.0567 | 0.0549 | 0.9676 |
| cluster3990 | -2207.13261 | 0.0165 | 0.0678 | 4.1164 | 0.0326 | 0.0221 | 0.6778 |
| cluster3991 | -1521.61263 | 0.0243 | 0.0465 | 1.9157 | 0.0157 | 0.0130 | 0.8288 |
| cluster3992 | -1887.96095 | 0.0263 | 0.0122 | 0.4654 | 0.0490 | 0.0227 | 0.4640 |
| cluster3993 | -1674.43869 | 0.0284 | 0.0632 | 2.2208 | 0.0441 | 0.0217 | 0.4922 |
| cluster3994 | -2393.35295 | 0.0426 | 0.0589 | 1.3826 | 0.0793 | 0.0540 | 0.6810 |
| cluster3995 | -1618.10611 | 0.0248 | 0.0412 | 1.6625 | 0.0497 | 0.0259 | 0.5215 |

| | | | | | | | |
|---|---|---|---|---|---|---|---|
| cluster3996 | -1707.30307 | 0.0087 | 0.0238 | 2.7302 | 0.0030 | 0.0070 | 2.3194 |
| cluster3997 | -2516.39728 | 0.0387 | 0.0723 | 1.8668 | 0.0767 | 0.0422 | 0.5500 |
| cluster3998 | -1414.29365 | 0.0359 | 0.0454 | 1.2661 | 0.0827 | 0.0314 | 0.3794 |
| cluster3999 | -2523.69612 | 0.0252 | 0.0740 | 2.9369 | 0.0451 | 0.0257 | 0.5687 |
| cluster4000 | -2288.36362 | 0.0343 | 0.0487 | 1.4173 | 0.0632 | 0.0291 | 0.4599 |
| cluster4001 | -1736.00255 | 0.0070 | 0.0215 | 3.0986 | 0.0361 | 0.0187 | 0.5183 |
| cluster4002 | -1801.66706 | 0.0036 | 0.0226 | 6.2160 | 0.0077 | 0.0078 | 1.0235 |
| cluster4003 | -1417.91027 | 0.0279 | 0.0813 | 2.9139 | 0.1129 | 0.0482 | 0.4266 |
| cluster4004 | -1442.80788 | 0.0088 | 0.0117 | 1.3365 | 0.0066 | 0.0027 | 0.4026 |
| cluster4005 | -2060.79930 | 0.0172 | 0.0965 | 5.6115 | 0.0874 | 0.0488 | 0.5584 |
| cluster4006 | -1089.41657 | 0.0349 | 0.0408 | 1.1717 | 0.0263 | 0.0077 | 0.2919 |
| cluster4007 | -1424.86417 | 0.0014 | 0.0019 | 1.4065 | 0.0056 | 0.0019 | 0.3490 |
| cluster4008 | -1353.51845 | 0.0144 | 0.0348 | 2.4091 | 0.0031 | 0.0034 | 1.0872 |
| cluster4009 | -1542.87849 | 0.0017 | 0.0449 | 26.5201 | 0.0187 | 0.0167 | 0.8932 |
| cluster4010 | -1626.12943 | 0.0419 | 0.1215 | 2.8986 | 0.0868 | 0.0523 | 0.6024 |
| cluster4011 | -1059.870157 | 0.1138 | 0.0177 | 0.1557 | 0.0431 | 0.0076 | 0.1764 |
| cluster4012 | -2440.90761 | 0.0082 | 0.0722 | 8.7852 | 0.0468 | 0.1299 | 2.7736 |
| cluster4013 | -846.34596 | 0.0185 | 0.0136 | 0.7351 | 0.0312 | 0.0082 | 0.2637 |
| cluster4014 | -1138.05442 | 0.0066 | 0.0145 | 2.1911 | 0.0057 | 0.0022 | 0.3821 |
| cluster4015 | -615.974998 | 0.1186 | 0.0035 | 0.0298 | 0.0001 | 0.0000 | 0.0664 |
| cluster4016 | -1810.91150 | 0.0095 | 0.0131 | 1.3793 | 0.0333 | 0.0208 | 0.6254 |
| cluster4017 | -1339.26258 | 0.0230 | 0.0529 | 2.3013 | 0.0766 | 0.0371 | 0.4845 |
| cluster4018 | -1322.914892 | 0.0975 | 0.1295 | 1.3286 | 0.2699 | 0.1235 | 0.4577 |
| cluster4019 | -1825.05177 | 0.0493 | 0.1155 | 2.3429 | 0.0879 | 0.0450 | 0.5123 |
| cluster4020 | -1005.38282 | 0.0377 | 0.0169 | 0.4482 | 0.0345 | 0.0131 | 0.3788 |
| cluster4021 | -1593.14254 | 0.0241 | 0.0203 | 0.8423 | 0.0313 | 0.0080 | 0.2563 |
| cluster4022 | -970.14324 | 0.0108 | 0.0127 | 1.1688 | 0.0057 | 0.0009 | 0.1554 |
| cluster4023 | -1844.27072 | 0.0719 | 0.0765 | 1.0642 | 0.0569 | 0.0272 | 0.4780 |
| cluster4024 | -2359.39954 | 0.0341 | 0.0925 | 2.7165 | 0.0246 | 0.0320 | 1.3012 |
| cluster4025 | -624.30922 | 0.0007 | 0.0037 | 5.2524 | 0.0001 | 0.0000 | 0.2154 |
| cluster4026 | -1087.93354 | 0.0045 | 0.0173 | 3.8397 | 0.0001 | 0.0000 | 0.3354 |
| cluster4027 | -978.42822 | 0.0022 | 0.0266 | 12.2770 | 0.0121 | 0.0039 | 0.3202 |
| cluster4028 | -1847.01553 | 0.0050 | 0.0250 | 4.9904 | 0.0082 | 0.0068 | 0.8352 |
| cluster4029 | -2248.53227 | 0.0457 | 0.1025 | 2.2417 | 0.0973 | 0.0601 | 0.6173 |
| cluster4030 | -1123.65764 | 0.0246 | 0.0375 | 1.5223 | 0.0606 | 0.0121 | 0.1996 |
| cluster4031 | -1097.50673 | 0.0637 | 0.0832 | 1.3067 | 0.0172 | 0.0118 | 0.6851 |
| cluster4032 | -1475.05536 | 0.0017 | 0.0010 | 0.5815 | 0.0001 | 0.0000 | 0.1820 |
| cluster4033 | -2212.65697 | 0.0475 | 0.1205 | 2.5389 | 0.0991 | 0.0587 | 0.5918 |
| cluster4034 | -493.67260 | 0.0001 | 0.0000 | 0.0089 | 0.0001 | 0.0001 | 1.3059 |
| cluster4035 | -2110.71307 | 0.0348 | 0.0257 | 0.7375 | 0.1058 | 0.0393 | 0.3715 |
| cluster4036 | -1076.83359 | 0.0123 | 0.0223 | 1.8157 | 0.0055 | 0.0029 | 0.5213 |
| cluster4037 | -1625.98866 | 0.0202 | 0.0467 | 2.3112 | 0.0180 | 0.0146 | 0.8081 |
| cluster4038 | -902.56872 | 0.0163 | 0.0588 | 3.6137 | 0.0174 | 0.0144 | 0.8294 |
| cluster4039 | -1507.95725 | 0.0681 | 0.0426 | 0.6250 | 0.0454 | 0.0053 | 0.1169 |

| | | | | | | | |
|---|---|---|---|---|---|---|---|
| cluster4040 | -1122.85631 | 0.0652 | 0.0479 | 0.7351 | 0.0145 | 0.0111 | 0.7660 |
| cluster4041 | -1475.34256 | 0.0156 | 0.0348 | 2.2351 | 0.0116 | 0.0101 | 0.8704 |
| cluster4042 | -899.30706 | 0.0256 | 0.0370 | 1.4462 | 0.0439 | 0.0110 | 0.2513 |
| cluster4043 | -1138.59362 | 0.0420 | 0.1612 | 3.8371 | 0.0465 | 0.0608 | 1.3085 |
| cluster4044 | -956.80116 | 0.0021 | 0.0054 | 2.5272 | 0.0001 | 0.0001 | 0.7751 |
| cluster4045 | -2023.77981 | 0.0616 | 0.1168 | 1.8951 | 0.0697 | 0.0359 | 0.5144 |
| cluster4046 | -889.47494 | 0.0035 | 0.0028 | 0.7922 | 0.0001 | 0.0000 | 0.1610 |
| cluster4047 | -2067.05469 | 0.0114 | 0.0469 | 4.0974 | 0.0270 | 0.0194 | 0.7206 |
| cluster4048 | -1469.99396 | 0.0065 | 0.0052 | 0.7884 | 0.0123 | 0.0028 | 0.2296 |
| cluster4049 | -573.230829 | 0.0005 | 0.0016 | 3.1324 | 0.0015 | 0.0052 | 3.4625 |
| cluster4050 | -1216.67972 | 0.0119 | 0.0124 | 1.0479 | 0.0047 | 0.0022 | 0.4704 |
| cluster4051 | -1403.33503 | 0.0123 | 0.0054 | 0.4430 | 0.0031 | 0.0007 | 0.2110 |
| cluster4052 | -1288.17630 | 0.0183 | 0.0093 | 0.5099 | 0.0200 | 0.0107 | 0.5378 |
| cluster4053 | -1641.64590 | 0.0051 | 0.0529 | 10.4177 | 0.0208 | 0.0146 | 0.7015 |
| cluster4054 | -1860.63608 | 0.0157 | 0.0471 | 2.9991 | 0.0219 | 0.0235 | 1.0706 |
| cluster4055 | -719.69343 | 0.0018 | 0.0265 | 14.5525 | 0.0035 | 0.0062 | 1.7618 |
| cluster4056 | -932.84067 | 0.0212 | 0.0399 | 1.8796 | 0.0597 | 0.0181 | 0.3034 |
| cluster4057 | -1563.96055 | 0.0253 | 0.0554 | 2.1881 | 0.0461 | 0.0245 | 0.5313 |
| cluster4058 | -1679.85909 | 0.0268 | 0.0699 | 2.6127 | 0.0574 | 0.0280 | 0.4884 |
| cluster4059 | -718.75195 | 0.0012 | 0.0442 | 38.3173 | 0.0360 | 0.0075 | 0.2087 |
| cluster4060 | -1875.11766 | 0.0221 | 0.0370 | 1.6792 | 0.0348 | 0.0164 | 0.4712 |
| cluster4061 | -1183.86697 | 0.0020 | 0.0110 | 5.5966 | 0.0151 | 0.0117 | 0.7723 |
| cluster4062 | -1201.11495 | 0.0003 | 0.0123 | 45.5147 | 0.0377 | 0.0131 | 0.3469 |
| cluster4063 | -1722.06765 | 0.0185 | 0.0901 | 4.8780 | 0.0370 | 0.0304 | 0.8228 |
| cluster4064 | -783.11597 | 0.0132 | 0.0249 | 1.8936 | 0.0197 | 0.0089 | 0.4529 |
| cluster4065 | -989.34037 | 0.0103 | 0.0086 | 0.8391 | 0.0057 | 0.0023 | 0.3976 |
| cluster4066 | -1616.92237 | 0.0446 | 0.0765 | 1.7155 | 0.0481 | 0.0199 | 0.4133 |
| cluster4067 | -418.206511 | 0.0757 | 0.1165 | 1.5377 | 0.0118 | 0.0366 | 3.1070 |
| cluster4068 | -1354.01446 | 0.0187 | 0.1052 | 5.6383 | 0.0181 | 0.0179 | 0.9902 |
| cluster4069 | -1065.11441 | 0.0645 | 0.0467 | 0.7247 | 0.0590 | 0.0217 | 0.3680 |
| cluster4070 | -695.03541 | 0.0071 | 0.0431 | 6.1023 | 0.0009 | 0.0150 | 15.9144 |
| cluster4071 | -826.53036 | 0.0028 | 0.0085 | 2.9881 | 0.0008 | 0.0011 | 1.4071 |
| cluster4072 | -987.81345 | 0.0205 | 0.0349 | 1.7056 | 0.0324 | 0.0133 | 0.4122 |
| cluster4073 | -1889.06910 | 0.0250 | 0.0638 | 2.5523 | 0.0402 | 0.0257 | 0.6397 |
| cluster4074 | -1256.89712 | 0.0389 | 0.0993 | 2.5533 | 0.0159 | 0.0299 | 1.8833 |
| cluster4075 | -709.176463 | 1.8754 | 0.0234 | 0.0125 | 0.3583 | 3.3482 | 9.3450 |
| cluster4076 | -1650.04796 | 0.0781 | 0.0468 | 0.5989 | 0.1020 | 0.0454 | 0.4454 |
| cluster4077 | -1679.84120 | 0.0394 | 0.0725 | 1.8399 | 0.0515 | 0.0192 | 0.3739 |
| cluster4078 | -1227.48037 | 0.0495 | 0.0322 | 0.6516 | 0.0088 | 0.0159 | 1.8076 |
| cluster4079 | -1206.00982 | 0.0024 | 0.0022 | 0.9127 | 0.0001 | 0.0001 | 0.9717 |
| cluster4080 | -737.50372 | 0.0181 | 0.0018 | 0.0996 | 0.0182 | 0.0306 | 1.6785 |
| cluster4081 | -1131.23972 | 0.0170 | 0.0214 | 1.2595 | 0.0212 | 0.0082 | 0.3839 |
| cluster4082 | -742.82389 | 0.0041 | 0.0016 | 0.3930 | 0.0135 | 0.1039 | 7.7138 |
| cluster4083 | -1700.79221 | 0.0195 | 0.1001 | 5.1389 | 0.0960 | 0.0644 | 0.6709 |

| | | | | | | | |
|---|---|---|---|---|---|---|---|
| cluster4084 | -716.05498 | 0.0175 | 0.0113 | 0.6483 | 0.0132 | 0.0050 | 0.3777 |
| cluster4085 | -858.264996 | 0.0096 | 0.0026 | 0.2725 | 0.0401 | 0.0136 | 0.3402 |
| cluster4086 | -1313.84016 | 0.0210 | 0.0661 | 3.1503 | 0.0411 | 0.0262 | 0.6369 |
| cluster4087 | -1205.49685 | 0.0158 | 0.0491 | 3.1007 | 0.0078 | 0.0113 | 1.4490 |
| cluster4088 | -1313.37046 | 0.0710 | 0.0409 | 0.5763 | 0.0365 | 0.0062 | 0.1689 |
| cluster4089 | -1509.98027 | 0.0034 | 0.0230 | 6.8067 | 0.0096 | 0.0110 | 1.1378 |
| cluster4090 | -755.27553 | 0.0283 | 0.0269 | 0.9524 | 0.0483 | 0.0165 | 0.3423 |
| cluster4091 | -954.05768 | 0.0338 | 0.0169 | 0.4996 | 0.0282 | 0.0063 | 0.2223 |
| cluster4092 | -1422.21451 | 0.0142 | 0.0653 | 4.5871 | 0.0453 | 0.0251 | 0.5547 |
| cluster4093 | -1006.45250 | 0.0129 | 0.0334 | 2.5863 | 0.0234 | 0.0151 | 0.6452 |
| cluster4094 | -374.968891 | 0.0151 | 0.0031 | 0.2049 | 0.0001 | 0.0000 | 0.1003 |
| cluster4095 | -1404.29321 | 0.0535 | 0.0792 | 1.4805 | 0.0362 | 0.0489 | 1.3497 |
| cluster4096 | -1212.32726 | 0.0148 | 0.0582 | 3.9246 | 0.0310 | 0.0196 | 0.6336 |
| cluster4097 | -704.658754 | 0.0001 | 0.0000 | 0.0352 | 0.1225 | 0.0194 | 0.1581 |
| cluster4098 | -1868.59871 | 0.0298 | 0.0290 | 0.9759 | 0.0364 | 0.0456 | 1.2514 |
| cluster4099 | -811.01258 | 0.0026 | 0.0063 | 2.4476 | 0.0028 | 0.0011 | 0.3775 |
| cluster4100 | -1048.78302 | 0.0080 | 0.0148 | 1.8435 | 0.0079 | 0.0049 | 0.6122 |
| cluster4101 | -2089.328288 | 0.2290 | 0.1490 | 0.6510 | 0.2555 | 0.1062 | 0.4158 |
| cluster4102 | -1856.88163 | 0.0441 | 0.2214 | 5.0156 | 0.0497 | 0.0869 | 1.7477 |
| cluster4103 | -1107.34138 | 0.0446 | 0.0147 | 0.3285 | 0.0391 | 0.0074 | 0.1904 |
| cluster4104 | -1018.03737 | 0.0070 | 0.0220 | 3.1694 | 0.0081 | 0.0039 | 0.4799 |
| cluster4105 | -733.40819 | 0.0058 | 0.0136 | 2.3515 | 0.0468 | 0.0745 | 1.5926 |
| cluster4106 | -1000.60740 | 0.0096 | 0.0245 | 2.5559 | 0.0050 | 0.0038 | 0.7469 |
| cluster4107 | -786.72797 | 0.0019 | 0.0036 | 1.9154 | 0.0032 | 0.0024 | 0.7525 |
| cluster4108 | -783.26388 | 0.0090 | 0.0225 | 2.4869 | 0.0174 | 0.0129 | 0.7426 |
| cluster4109 | -1651.88942 | 0.0661 | 0.1637 | 2.4770 | 0.0687 | 0.0397 | 0.5785 |
| cluster4110 | -1243.71317 | 0.0302 | 0.1100 | 3.6375 | 0.1036 | 0.0297 | 0.2865 |
| cluster4111 | -925.16404 | 0.0432 | 0.1177 | 2.7274 | 0.0519 | 0.0240 | 0.4631 |
| cluster4112 | -1113.30444 | 0.0141 | 0.0167 | 1.1871 | 0.0185 | 0.0086 | 0.4644 |
| cluster4113 | -772.58419 | 0.0046 | 0.0135 | 2.9675 | 0.0281 | 0.0011 | 0.0391 |
| cluster4114 | -1074.98134 | 0.0453 | 0.0436 | 0.9635 | 0.0393 | 0.0114 | 0.2895 |
| cluster4115 | -1152.97128 | 0.0033 | 0.0214 | 6.4443 | 0.0205 | 0.0123 | 0.5978 |
| cluster4116 | -635.581584 | 0.0108 | 0.0263 | 2.4347 | 0.1463 | 0.0764 | 0.5223 |
| cluster4117 | -660.63188 | 0.0129 | 0.0146 | 1.1277 | 0.0128 | 0.0014 | 0.1110 |
| cluster4118 | -1288.26318 | 0.0254 | 0.0666 | 2.6241 | 0.0363 | 0.0241 | 0.6649 |
| cluster4119 | -924.47143 | 0.0089 | 0.0440 | 4.9648 | 0.0315 | 0.0152 | 0.4825 |
| cluster4120 | -965.23128 | 0.0001 | 0.0001 | 1.2414 | 0.0122 | 0.0043 | 0.3534 |
| cluster4121 | -1217.36452 | 0.0031 | 0.0387 | 12.3450 | 0.0059 | 0.0156 | 2.6609 |
| cluster4122 | -527.410036 | 0.0267 | 0.0036 | 0.1344 | 0.0760 | 0.0012 | 0.0157 |
| cluster4123 | -467.18166 | 0.0113 | 0.0117 | 1.0370 | 0.0001 | 0.0000 | 0.2888 |
| cluster4124 | -905.64501 | 0.0045 | 0.0307 | 6.7707 | 0.0042 | 0.0024 | 0.5620 |
| cluster4125 | -1065.590635 | 0.0969 | 0.0749 | 0.7723 | 0.0584 | 0.0591 | 1.0119 |
| cluster4126 | -762.587517 | 0.0993 | 0.0497 | 0.5006 | 0.0556 | 0.0077 | 0.1392 |
| cluster4127 | -766.44990 | 0.0552 | 0.0624 | 1.1319 | 0.0605 | 0.0164 | 0.2711 |

| | | | | | | | |
|---|---|---|---|---|---|---|---|
| cluster4128 | -989.65507 | 0.0346 | 0.1600 | 4.6236 | 0.0360 | 0.0408 | 1.1356 |

| Adjusted p value |
| ---: |
| 0.04 |
| 7.43E-65 |
| 0.0005 |
| 1.35E-92 |
| 2.78E-07 |
| 7.21E-06 |
| 5.07E-54 |
| 3.26E-09 |
| 5.26E-124 |
| 9.13E-42 |
| 4.28E-88 |
| 6.57E-44 |
| 2.32E-107 |
| 2.19E-12 |
| 1.42E-48 |
| 3.32E-28 |
| 2.09E-08 |
| 7.05E-06 |
| 3.57E-06 |
| 1.07E-10 |
| 1.69E-59 |
| 4.75E-22 |
| 3.04E-64 |
| 6.87E-11 |
| 8.19E-21 |
| 5.15E-12 |
| 0.005 |
| 1.65E-17 |
| 6.98E-19 |
| 1.89E-27 |
| 1.90E-07 |
| 1.19E-33 |
| 1.10E-32 |
| 7.67E-25 |
| 4.45E-55 |
| 6.39E-08 |
| 1.28E-46 |
| 0.07 |
| 1.29E-24 |
| 1.82E-34 |
| 2.48E-12 |
| 3.92E-17 |

| |
|---|
| 5.08E-19 |
| 4.76E-66 |
| 1.94E-22 |
| 2.18E-14 |
| 1.67E-29 |
| 1.54E-39 |
| 1.73E-20 |
| 1.88E-13 |
| 1.36E-45 |
| 3.79E-05 |
| 2.84E-19 |
| 5.17E-16 |
| 3.00E-11 |
| 5.75E-14 |
| 6.51E-05 |
| 2.42E-20 |
| 4.00E-10 |
| 3.70E-24 |
| 5.28E-41 |
| 5.23E-25 |
| 4.56E-19 |
| 1.64E-26 |
| 7.03E-30 |
| 0.0001 |
| 3.63E-13 |
| 4.24E-30 |
| 6.10E-27 |
| 1.24E-09 |
| 0.04 |
| 4.92E-33 |
| 1.52E-14 |
| 5.28E-41 |
| 2.11E-11 |
| 1.03E-24 |
| 3.62E-52 |
| 1.05E-09 |
| 2.74E-10 |
| 1.21E-12 |
| 2.36E-14 |
| 6.47E-12 |
| 4.39E-14 |
| 3.70E-24 |
| 1.29E-12 |
| 4.04E-29 |

| |
|---|
| 1.43E-06 |
| 7.18E-15 |
| 1.27E-06 |
| 1.03E-16 |
| 1.66E-23 |
| 8.32E-06 |
| 3.63E-21 |
| 2.29E-15 |
| 5.03E-21 |
| 4.95E-16 |
| 2.98E-23 |
| 1.58E-27 |
| 1.47E-10 |
| 4.58E-22 |
| 1.04E-14 |
| 2.73E-18 |
| 1.57E-15 |
| 4.81E-38 |
| 3.23E-06 |
| 1.00E-47 |
| 2.92E-09 |
| 4.84E-36 |
| 9.41E-47 |
| 7.30E-21 |
| 2.22E-14 |
| 9.57E-14 |
| 4.53E-32 |
| 5.00E-18 |
| 5.69E-40 |
| 1.03E-21 |
| 1.13E-15 |
| 3.27E-30 |
| 2.18E-05 |
| 7.08E-17 |
| 1.09E-05 |
| 2.88E-09 |
| 5.71E-11 |
| 0.47 |
| 4.63E-34 |
| 3.05E-57 |
| 8.81E-19 |
| 2.46E-05 |
| 4.68E-11 |
| 4.02E-05 |

| |
|---|
| 2.81E-26 |
| 1.50E-12 |
| 0.23 |
| 0.01 |
| 1.04E-13 |
| 1.02E-19 |
| 3.64E-13 |
| 4.87E-14 |
| 6.90E-08 |
| 5.77E-12 |
| 6.98E-17 |
| 2.85E-10 |
| 0.10 |
| 4.93E-16 |
| 5.67E-11 |
| 5.96E-27 |
| 1.01E-15 |
| 5.51E-18 |
| 3.70E-16 |
| 5.59E-22 |
| 1.75E-06 |
| 3.65E-05 |
| 4.43E-05 |
| 6.42E-10 |
| 2.00E-23 |
| 7.88E-17 |
| 1.08E-09 |
| 0.005 |
| 1.03E-16 |
| 2.34E-07 |
| 1.41E-06 |
| 7.92E-25 |
| 0.0002 |
| 7.72E-09 |
| 2.58E-20 |
| 4.81E-32 |
| 1.06E-11 |
| 2.56E-15 |
| 2.32E-10 |
| 0.0002 |
| 1.46E-19 |
| 0.02 |
| 1.26E-12 |
| 0.14 |

| |
|---|
| 1.39E-30 |
| 2.82E-12 |
| 9.67E-15 |
| 3.45E-08 |
| 5.74E-07 |
| 4.33E-22 |
| 0.14 |
| 0.005 |
| 6.42E-25 |
| 4.18E-08 |
| 2.91E-07 |
| 4.91E-07 |
| 2.75E-11 |
| 5.66E-14 |
| 2.65E-15 |
| 4.21E-12 |
| 5.46E-09 |
| 1.49E-16 |
| 7.60E-11 |
| 0.02 |
| 1.03E-08 |
| 3.59E-17 |
| 3.55E-31 |
| 2.18E-14 |
| 5.40E-14 |
| 1.03E-13 |
| 1.36E-07 |
| 7.88E-33 |
| 5.79E-09 |
| 9.93E-05 |
| 0.05 |
| 1.39E-05 |
| 2.21E-06 |
| 1.17E-11 |
| 2.35E-17 |
| 7.35E-10 |
| 1.15E-16 |
| 2.21E-05 |
| 0.02 |
| 4.99E-08 |
| 6.94E-11 |
| 0.003 |
| 0.001 |
| 2.98E-13 |

| |
|---|
| 2.35E-17 |
| 1.80E-07 |
| 1.00E-14 |
| 1.32E-13 |
| 8.55E-13 |
| 0.001 |
| 2.58E-16 |
| 1.94E-16 |
| 6.40E-20 |
| 9.38E-21 |
| 4.02E-18 |
| 5.03E-10 |
| 3.24E-11 |
| 0.002 |
| 1.73E-20 |
| 0.001 |
| 2.48E-20 |
| 7.08E-09 |
| 0.0001 |
| 6.15E-28 |
| 2.99E-07 |
| 3.47E-14 |
| 4.80E-16 |
| 5.65E-21 |
| 7.72E-27 |
| 2.07E-14 |
| 9.55E-19 |
| 3.09E-11 |
| 3.60E-09 |
| 1.62E-09 |
| 2.82E-08 |
| 2.18E-12 |
| 1.88E-09 |
| 3.35E-15 |
| 9.92E-17 |
| 2.80E-23 |
| 2.82E-05 |
| 0.01 |
| 1.70E-11 |
| 0.0003 |
| 1.75E-11 |
| 5.80E-10 |
| 2.16E-10 |
| 2.45E-10 |

| |
|---|
| 4.28E-14 |
| 1.49E-06 |
| 1.06E-12 |
| 0.01 |
| 5.93E-06 |
| 5.86E-16 |
| 2.74E-18 |
| 2.42E-10 |
| 1.07E-14 |
| 4.21E-14 |
| 4.18E-05 |
| 1.11E-15 |
| 3.01E-09 |
| 5.58E-10 |
| 8.19E-06 |
| 3.86E-14 |
| 2.21E-10 |
| 1.48E-15 |
| 2.08E-08 |
| 2.50E-05 |
| 3.64E-12 |
| 3.01E-09 |
| 1.05E-11 |
| 1.01E-08 |
| 4.02E-06 |
| 1.53E-05 |
| 3.62E-08 |
| 0.16 |
| 5.71E-12 |
| 6.43E-12 |
| 9.37E-51 |
| 2.92E-13 |
| 6.68E-27 |
| 3.34E-14 |
| 7.95E-10 |
| 9.01E-05 |
| 5.15E-06 |
| 1.91E-13 |
| 1.01E-08 |
| 1.69E-27 |
| 3.92E-17 |
| 1.98E-16 |
| 0.02 |
| 8.85E-16 |

| |
|---|
| 2.51E-10 |
| 0.11 |
| 6.19E-11 |
| 9.33E-11 |
| 2.33E-08 |
| 3.22E-09 |
| 5.63E-11 |
| 4.46E-16 |
| 8.58E-05 |
| 7.05E-30 |
| 2.95E-14 |
| 0.02 |
| 1.21E-07 |
| 5.80E-17 |
| 1.31E-20 |
| 4.73E-09 |
| 9.55E-07 |
| 2.51E-07 |
| 0.0004 |
| 3.83E-41 |
| 2.64E-05 |
| 3.67E-07 |
| 1.03E-24 |
| 4.01E-09 |
| 0.0001 |
| 7.98E-09 |
| 7.33E-18 |
| 3.86E-06 |
| 1.13E-17 |
| 0.0005 |
| 1.05E-10 |
| 6.37E-14 |
| 8.55E-05 |
| 4.37E-10 |
| 5.03E-07 |
| 3.02E-09 |
| 8.57E-18 |
| 6.44E-06 |
| 1.21E-07 |
| 1.38E-20 |
| 1.08E-11 |
| 7.49E-20 |
| 0.003 |
| 8.33E-11 |

| |
|---|
| 0.0001 |
| 0.003 |
| 4.30E-05 |
| 1.02E-09 |
| 8.50E-26 |
| 1.53E-09 |
| 2.78E-19 |
| 3.38E-21 |
| 1.08E-09 |
| 1.22E-12 |
| 0.0001 |
| 8.81E-12 |
| 0.0003 |
| 1.27E-10 |
| 1.06E-07 |
| 4.39E-14 |
| 1.22E-23 |
| 4.05E-06 |
| 4.73E-14 |
| 7.98E-06 |
| 0.002 |
| 8.58E-06 |
| 5.56E-17 |
| 1.39E-08 |
| 4.17E-06 |
| 5.14E-10 |
| 1.87E-09 |
| 1.62E-14 |
| 1.74E-11 |
| 1.43E-18 |
| 4.45E-08 |
| 3.46E-07 |
| 5.50E-12 |
| 3.93E-12 |
| 2.95E-05 |
| 0.0005 |
| 2.34E-05 |
| 2.96E-19 |
| 1.24E-07 |
| 3.19E-09 |
| 0.01 |
| 2.15E-10 |
| 1.22E-21 |
| 3.18E-11 |

| |
|---|
| 0.04 |
| 9.55E-13 |
| 5.75E-14 |
| 0.01 |
| 1.30E-07 |
| 5.93E-08 |
| 5.98E-05 |
| 6.15E-09 |
| 2.89E-20 |
| 3.41E-13 |
| 1.67E-29 |
| 3.50E-08 |
| 1.37E-05 |
| 2.12E-18 |
| 1.76E-05 |
| 2.91E-22 |
| 0.04 |
| 2.19E-05 |
| 8.25E-31 |
| 8.55E-05 |
| 6.93E-11 |
| 4.24E-07 |
| 1.45E-05 |
| 4.49E-06 |
| 5.35E-16 |
| 0.71 |
| 0.004 |
| 1.45E-05 |
| 1.36E-08 |
| 0.02 |
| 5.17E-05 |
| 3.20E-17 |
| 1.11E-13 |
| 3.34E-11 |
| 4.99E-08 |
| 9.95E-12 |
| 1.57E-07 |
| 8.57E-15 |
| 3.35E-07 |
| 0.001 |
| 1.89E-06 |
| 8.82E-09 |
| 1.02E-09 |
| 0.001 |

| |
|---|
| 6.33E-12 |
| 1.02E-06 |
| 1.95E-19 |
| 1.46E-10 |
| 1.64E-10 |
| 2.43E-10 |
| 0.16 |
| 7.87E-22 |
| 1.16E-06 |
| 1.82E-15 |
| 0.001 |
| 1.62E-05 |
| 1.66E-11 |
| 5.57E-07 |
| 3.33E-05 |
| 6.31E-06 |
| 3.67E-12 |
| 1.53E-18 |
| 0.02 |
| 2.72E-15 |
| 6.51E-10 |
| 1.87E-08 |
| 1.67E-07 |
| 8.47E-11 |
| 1.90E-21 |
| 1.78E-05 |
| 1.16E-06 |
| 0.03 |
| 4.48E-10 |
| 0.0001 |
| 1.72E-09 |
| 1.94E-07 |
| 8.63E-10 |
| 0.001 |
| 4.47E-08 |
| 0.06 |
| 6.24E-13 |
| 8.79E-07 |
| 2.32E-05 |
| 8.25E-06 |
| 0.001 |
| 4.14E-09 |
| 2.75E-11 |
| 0.04 |

| |
|---|
| 5.75E-09 |
| 3.62E-13 |
| 1.46E-05 |
| 2.94E-12 |
| 6.63E-05 |
| 0.43 |
| 1.99E-08 |
| 3.19E-09 |
| 6.10E-24 |
| 4.23E-08 |
| 1.00E-15 |
| 5.83E-23 |
| 1.84E-07 |
| 1.68E-06 |
| 3.04E-17 |
| 9.63E-09 |
| 3.54E-11 |
| 1.62E-05 |
| 1.53E-05 |
| 8.32E-07 |
| 7.83E-09 |
| 2.83E-09 |
| 0.0002 |
| 7.82E-26 |
| 2.52E-13 |
| 7.98E-06 |
| 6.86E-09 |
| 1.66E-15 |
| 8.24E-35 |
| 1.08E-10 |
| 9.89E-09 |
| 4.32E-07 |
| 0.01 |
| 0.001 |
| 1.55E-16 |
| 7.03E-05 |
| 1.56E-11 |
| 8.26E-18 |
| 4.25E-34 |
| 9.63E-06 |
| 3.00E-09 |
| 0.01 |
| 1.75E-15 |
| 6.47E-12 |

| |
|---|
| 0.0002 |
| 1.06E-06 |
| 9.63E-06 |
| 0.01 |
| 1.91E-07 |
| 4.18E-05 |
| 3.42E-13 |
| 2.40E-20 |
| 2.71E-07 |
| 1.74E-09 |
| 3.97E-09 |
| 1.38E-06 |
| 2.81E-11 |
| 1.58E-12 |
| 0.0002 |
| 1.72E-06 |
| 6.35E-13 |
| 1.09E-12 |
| 5.00E-15 |
| 6.88E-06 |
| 2.79E-07 |
| 0.12 |
| 1.00E-08 |
| 8.04E-15 |
| 2.88E-11 |
| 0.11 |
| 0.06 |
| 5.78E-15 |
| 3.02E-08 |
| 3.31E-06 |
| 2.99E-14 |
| 0.0005 |
| 5.13E-15 |
| 0.001 |
| 1.09E-05 |
| 0.13 |
| 2.16E-05 |
| 0.03 |
| 3.46E-10 |
| 0.0002 |
| 0.06 |
| 0.35 |
| 1.60E-20 |
| 6.70E-14 |

| |
|---|
| 2.44E-08 |
| 0.01 |
| 0.001 |
| 1.48E-15 |
| 0.001 |
| 0.0004 |
| 6.48E-13 |
| 7.73E-09 |
| 4.37E-10 |
| 3.40E-08 |
| 0.02 |
| 4.74E-07 |
| 3.13E-15 |
| 5.77E-12 |
| 7.18E-15 |
| 1.06E-08 |
| 0.002 |
| 6.67E-06 |
| 7.51E-06 |
| 6.39E-08 |
| 9.48E-13 |
| 2.99E-14 |
| 2.51E-11 |
| 4.94E-12 |
| 2.23E-11 |
| 9.52E-08 |
| 0.04 |
| 1.44E-09 |
| 2.72E-10 |
| 6.01E-07 |
| 0.0003 |
| 1.24E-15 |
| 4.30E-05 |
| 2.19E-08 |
| 6.01E-10 |
| 0.0003 |
| 1.50E-05 |
| 1.81E-12 |
| 0.0004 |
| 0.21 |
| 1.84E-08 |
| 1.24E-09 |
| 2.85E-18 |
| 2.52E-13 |

| |
|---|
| 6.41E-10 |
| 2.93E-11 |
| 3.22E-08 |
| 4.39E-16 |
| 1.07E-08 |
| 2.77E-15 |
| 5.63E-11 |
| 7.19E-11 |
| 9.31E-09 |
| 6.92E-12 |
| 9.30E-08 |
| 4.15E-13 |
| 0.0002 |
| 1.01E-10 |
| 0.001 |
| 0.03 |
| 7.94E-19 |
| 6.61E-09 |
| 5.66E-06 |
| 8.30E-05 |
| 6.93E-07 |
| 0.0003 |
| 6.10E-08 |
| 9.30E-12 |
| 3.97E-09 |
| 2.04E-08 |
| 1.07E-10 |
| 1.25E-10 |
| 2.63E-08 |
| 5.97E-11 |
| 0.02 |
| 5.99E-07 |
| 0.03 |
| 1.12E-08 |
| 2.73E-06 |
| 8.00E-10 |
| 0.001 |
| 7.75E-08 |
| 3.56E-05 |
| 8.68E-11 |
| 7.87E-09 |
| 7.72E-11 |
| 0.0002 |
| 0.24 |

| |
|---|
| 0.02 |
| 3.85E-05 |
| 0.41 |
| 2.24E-12 |
| 7.17E-12 |
| 4.72E-12 |
| 5.20E-10 |
| 4.16E-05 |
| 2.90E-05 |
| 0.26 |
| 3.40E-09 |
| 2.14E-10 |
| 2.99E-05 |
| 3.49E-06 |
| 8.01E-05 |
| 3.87E-24 |
| 1.73E-08 |
| 8.30E-08 |
| 6.54E-10 |
| 2.82E-05 |
| 5.05E-06 |
| 1.91E-07 |
| 4.99E-07 |
| 0.01 |
| 3.39E-10 |
| 6.09E-09 |
| 1.72E-24 |
| 0.004 |
| 4.38E-06 |
| 0.0002 |
| 4.74E-11 |
| 1.35E-06 |
| 2.43E-09 |
| 0.005 |
| 6.50E-10 |
| 8.41E-08 |
| 0.005 |
| 0.04 |
| 2.00E-08 |
| 0.003 |
| 0.001 |
| 6.92E-08 |
| 1.53E-07 |
| 5.08E-10 |

| |
|---|
| 6.84E-07 |
| 5.68E-14 |
| 0.08 |
| 5.18E-07 |
| 3.02E-08 |
| 9.96E-08 |
| 0.0005 |
| 1.30E-10 |
| 1.81E-07 |
| 5.36E-07 |
| 9.05E-09 |
| 0.0002 |
| 0.005 |
| 1.31E-07 |
| 1.91E-06 |
| 4.93E-07 |
| 3.91E-07 |
| 3.97E-06 |
| 1.83E-11 |
| 4.85E-06 |
| 8.55E-09 |
| 4.35E-07 |
| 8.63E-06 |
| 5.85E-08 |
| 4.19E-08 |
| 6.91E-12 |
| 2.99E-07 |
| 0.01 |
| 2.80E-18 |
| 6.88E-10 |
| 0.004 |
| 3.70E-06 |
| 2.53E-05 |
| 1.44E-07 |
| 6.44E-07 |
| 0.0001 |
| 1.46E-10 |
| 2.43E-10 |
| 1.03E-07 |
| 0.0001 |
| 1.64E-08 |
| 1.07E-06 |
| 1.30E-11 |
| 8.41E-08 |

| |
|---|
| 5.21E-08 |
| 6.93E-08 |
| 7.77E-19 |
| 6.70E-14 |
| 1.13E-08 |
| 0.24 |
| 4.96E-07 |
| 5.56E-13 |
| 1.77E-06 |
| 3.78E-12 |
| 9.56E-10 |
| 7.33E-05 |
| 2.54E-05 |
| 2.46E-07 |
| 2.00E-16 |
| 2.69E-09 |
| 0.01 |
| 0.003 |
| 0.0004 |
| 0.32 |
| 4.48E-16 |
| 1.55E-07 |
| 1.40E-12 |
| 2.43E-10 |
| 3.60E-09 |
| 2.08E-10 |
| 2.42E-17 |
| 9.83E-07 |
| 1.37E-19 |
| 3.02E-08 |
| 1.09E-22 |
| 0.001 |
| 1.64E-14 |
| 3.95E-07 |
| 2.19E-06 |
| 0.002 |
| 0.02 |
| 3.14E-10 |
| 0.01 |
| 3.77E-06 |
| 0.002 |
| 8.87E-05 |
| 0.0002 |
| 9.14E-08 |

| |
|---|
| 0.003 |
| 8.63E-05 |
| 3.37E-06 |
| 5.25E-08 |
| 7.67E-25 |
| 3.25E-16 |
| 2.13E-06 |
| 0.0005 |
| 3.70E-07 |
| 9.77E-12 |
| 4.73E-09 |
| 2.66E-16 |
| 0.002 |
| 1.12E-10 |
| 1.99E-11 |
| 0.20 |
| 3.00E-05 |
| 2.84E-11 |
| 0.01 |
| 2.99E-07 |
| 1.40E-05 |
| 1.71E-05 |
| 1.42E-14 |
| 1.01E-06 |
| 0.001 |
| 1.09E-11 |
| 4.57E-06 |
| 1.06E-08 |
| 2.42E-10 |
| 1.77E-05 |
| 2.87E-05 |
| 4.64E-05 |
| 0.01 |
| 3.54E-06 |
| 0.01 |
| 2.50E-08 |
| 0.0001 |
| 9.57E-14 |
| 1.52E-10 |
| 2.29E-12 |
| 0.02 |
| 4.26E-09 |
| 0.001 |
| 0.004 |

| |
|---|
| 0.04 |
| 5.34E-11 |
| 2.66E-06 |
| 2.34E-05 |
| 0.0002 |
| 7.61E-09 |
| 3.46E-17 |
| 0.002 |
| 2.10E-06 |
| 0.002 |
| 0.0002 |
| 7.37E-05 |
| 1.03E-09 |
| 9.83E-08 |
| 2.46E-06 |
| 2.18E-10 |
| 5.60E-05 |
| 2.03E-15 |
| 5.32E-09 |
| 2.45E-07 |
| 0.0002 |
| 0.0004 |
| 0.04 |
| 1.87E-08 |
| 0.02 |
| 2.55E-10 |
| 5.19E-07 |
| 0.003 |
| 9.49E-06 |
| 1.86E-07 |
| 0.001 |
| 2.49E-11 |
| 2.14E-06 |
| 1.70E-07 |
| 0.01 |
| 0.02 |
| 2.08E-05 |
| 3.79E-09 |
| 9.74E-11 |
| 0.0002 |
| 0.0003 |
| 1.98E-05 |
| 6.72E-12 |
| 0.02 |

| |
|---|
| 1.87E-13 |
| 2.57E-09 |
| 0.0004 |
| 6.03E-07 |
| 0.001 |
| 4.42E-07 |
| 0.0001 |
| 2.78E-19 |
| 0.001 |
| 3.39E-10 |
| 7.55E-05 |
| 0.001 |
| 1.19E-09 |
| 0.15 |
| 1.16E-08 |
| 1.69E-08 |
| 0.0002 |
| 1.16E-08 |
| 1.88E-14 |
| 4.63E-05 |
| 1.30E-19 |
| 1.16E-13 |
| 0.13 |
| 0.01 |
| 9.52E-12 |
| 2.83E-05 |
| 0.0001 |
| 0.0005 |
| 0.0003 |
| 4.81E-08 |
| 1.38E-05 |
| 5.19E-05 |
| 0.005 |
| 5.19E-05 |
| 4.12E-09 |
| 9.69E-08 |
| 0.0005 |
| 0.01 |
| 4.38E-05 |
| 0.002 |
| 0.005 |
| 0.01 |
| 0.54 |
| 0.003 |

| |
|---|
| 0.001 |
| 0.001 |
| 1.45E-05 |
| 1.12E-06 |
| 9.10E-10 |
| 1.18E-09 |
| 0.0001 |
| 1.38E-05 |
| 8.32E-14 |
| 3.72E-06 |
| 1.32E-08 |
| 5.76E-10 |
| 3.90E-05 |
| 0.11 |
| 1.07E-10 |
| 1.17E-08 |
| 0.02 |
| 1.92E-08 |
| 3.59E-06 |
| 2.13E-09 |
| 5.44E-06 |
| 0.14 |
| 9.02E-05 |
| 1.53E-13 |
| 0.0001 |
| 0.51 |
| 0.33 |
| 1.98E-06 |
| 6.48E-06 |
| 1.72E-06 |
| 1.82E-09 |
| 8.01E-09 |
| 1.46E-05 |
| 1.09E-05 |
| 9.32E-12 |
| 3.23E-13 |
| 0.07 |
| 2.52E-07 |
| 1.84E-06 |
| 0.11 |
| 2.53E-07 |
| 6.72E-09 |
| 0.001 |
| 0.001 |

| |
|---|
| 1.84E-10 |
| 6.21E-07 |
| 1.19E-07 |
| 5.04E-10 |
| 2.99E-14 |
| 1.64E-08 |
| 2.45E-08 |
| 0.002 |
| 6.41E-08 |
| 2.34E-05 |
| 6.06E-07 |
| 0.002 |
| 1.70E-06 |
| 8.89E-11 |
| 2.14E-09 |
| 1.75E-09 |
| 1.06E-05 |
| 1.58E-07 |
| 4.97E-07 |
| 1.54E-07 |
| 3.54E-07 |
| 1.45E-05 |
| 0.01 |
| 4.14E-05 |
| 7.76E-07 |
| 0.11 |
| 0.01 |
| 1.31E-05 |
| 1.26E-12 |
| 1.38E-07 |
| 5.33E-07 |
| 0.0003 |
| 5.24E-09 |
| 1.62E-13 |
| 2.65E-08 |
| 0.08 |
| 1.34E-16 |
| 2.10E-08 |
| 1.71E-14 |
| 4.00E-05 |
| 1.74E-08 |
| 0.003 |
| 6.17E-06 |
| 6.55E-07 |

| |
|---|
| 5.72E-10 |
| 4.30E-19 |
| 9.59E-06 |
| 1.31E-06 |
| 0.29 |
| 0.0005 |
| 1.73E-06 |
| 5.12E-07 |
| 3.01E-13 |
| 1.22E-12 |
| 1.51E-05 |
| 7.26E-06 |
| 0.0002 |
| 0.002 |
| 1.91E-05 |
| 9.38E-08 |
| 0.0002 |
| 3.18E-05 |
| 7.67E-09 |
| 0.06 |
| 6.99E-09 |
| 1.47E-14 |
| 2.78E-10 |
| 6.10E-08 |
| 3.37E-06 |
| 0.001 |
| 3.79E-07 |
| 8.33E-11 |
| 0.0005 |
| 5.32E-05 |
| 7.11E-05 |
| 8.58E-09 |
| 7.07E-06 |
| 7.43E-06 |
| 6.48E-06 |
| 9.31E-09 |
| 0.05 |
| 1.05E-14 |
| 8.20E-14 |
| 3.69E-07 |
| 0.0004 |
| 0.11 |
| 5.05E-05 |
| 7.13E-06 |

| |
|---|
| 1.67E-06 |
| 8.96E-08 |
| 0.002 |
| 3.14E-10 |
| 0.0001 |
| 6.06E-06 |
| 0.004 |
| 1.80E-06 |
| 0.08 |
| 7.57E-10 |
| 3.03E-08 |
| 0.10 |
| 0.02 |
| 3.90E-06 |
| 0.01 |
| 1.64E-13 |
| 2.43E-06 |
| 0.01 |
| 0.004 |
| 9.62E-07 |
| 0.002 |
| 2.61E-11 |
| 1.01E-08 |
| 0.09 |
| 0.01 |
| 0.12 |
| 0.001 |
| 0.0002 |
| 1.91E-08 |
| 2.25E-08 |
| 3.32E-05 |
| 3.01E-09 |
| 0.0001 |
| 0.04 |
| 1.32E-05 |
| 7.44E-10 |
| 0.0002 |
| 0.0002 |
| 4.83E-06 |
| 1.36E-07 |
| 0.003 |
| 0.001 |
| 8.01E-09 |
| 1.45E-07 |

| |
|---|
| 1.22E-11 |
| 0.0001 |
| 4.11E-05 |
| 0.0004 |
| 1.02E-09 |
| 2.49E-10 |
| 0.0004 |
| 2.71E-07 |
| 0.002 |
| 0.0005 |
| 0.001 |
| 1.01E-05 |
| 0.02 |
| 0.0002 |
| 7.26E-05 |
| 0.03 |
| 4.30E-05 |
| 0.0005 |
| 0.0001 |
| 4.81E-05 |
| 3.86E-05 |
| 0.0004 |
| 5.09E-07 |
| 0.001 |
| 1.22E-06 |
| 0.02 |
| 0.0002 |
| 9.36E-13 |
| 0.01 |
| 1.13E-06 |
| 0.0004 |
| 2.78E-06 |
| 0.02 |
| 0.0001 |
| 1.30E-07 |
| 5.83E-10 |
| 2.11E-06 |
| 0.002 |
| 9.47E-08 |
| 1.72E-06 |
| 0.0002 |
| 0.0002 |
| 0.0004 |
| 0.0002 |

| |
|---|
| 5.30E-11 |
| 0.0001 |
| 0.003 |
| 4.12E-06 |
| 8.73E-06 |
| 5.62E-07 |
| 1.44E-12 |
| 0.01 |
| 1.06E-08 |
| 0.0002 |
| 0.001 |
| 6.62E-11 |
| 6.52E-09 |
| 7.56E-06 |
| 0.01 |
| 0.15 |
| 3.67E-06 |
| 0.002 |
| 3.85E-06 |
| 0.42 |
| 1.73E-08 |
| 0.002 |
| 0.25 |
| 0.01 |
| 0.01 |
| 4.08E-05 |
| 0.002 |
| 0.003 |
| 1.66E-05 |
| 3.73E-05 |
| 5.81E-07 |
| 0.0001 |
| 2.09E-12 |
| 0.002 |
| 0.003 |
| 0.005 |
| 1.29E-06 |
| 0.003 |
| 0.01 |
| 0.0004 |
| 1.39E-05 |
| 6.99E-06 |
| 8.86E-05 |
| 0.02 |

| |
|---|
| 0.004 |
| 9.72E-07 |
| 0.01 |
| 0.03 |
| 3.92E-11 |
| 0.01 |
| 0.0003 |
| 7.96E-10 |
| 3.31E-06 |
| 0.12 |
| 5.69E-06 |
| 4.54E-05 |
| 8.50E-06 |
| 3.31E-06 |
| 9.90E-08 |
| 7.68E-06 |
| 0.002 |
| 9.87E-05 |
| 0.002 |
| 0.02 |
| 4.57E-06 |
| 2.68E-09 |
| 4.21E-07 |
| 0.0002 |
| 0.0004 |
| 0.0001 |
| 0.001 |
| 0.14 |
| 0.0003 |
| 0.05 |
| 0.0001 |
| 2.59E-06 |
| 0.003 |
| 4.67E-06 |
| 0.0005 |
| 0.97 |
| 0.02 |
| 0.003 |
| 3.92E-05 |
| 1.65E-10 |
| 0.01 |
| 0.0001 |
| 0.02 |
| 0.001 |

| |
|---|
| 0.02 |
| 0.14 |
| 0.0002 |
| 7.13E-06 |
| 0.02 |
| 9.23E-08 |
| 0.02 |
| 0.01 |
| 0.0002 |
| 0.0001 |
| 0.22 |
| 0.02 |
| 0.0001 |
| 0.49 |
| 1.67E-05 |
| 1.27E-06 |
| 1.78E-07 |
| 0.46 |
| 0.02 |
| 0.02 |
| 1.88E-07 |
| 0.004 |
| 3.82E-06 |
| 7.26E-07 |
| 1.37E-05 |
| 0.01 |
| 0.02 |
| 0.04 |
| 2.28E-05 |
| 0.03 |
| 0.01 |
| 1.41E-06 |
| 0.11 |
| 0.01 |
| 0.0001 |
| 0.0002 |
| 1.40E-06 |
| 4.63E-09 |
| 0.56 |
| 0.01 |
| 3.61E-07 |
| 0.07 |
| 0.20 |
| 0.03 |

0.02

**Supplementary Table 23.** codeml results - w for the parasites/other species

| ID | Likelihood | ϖ saprophagousother species | dN | dS | ϖ parasites |
|---|---|---|---|---|---|
| cluster1127 | -1490.348354 | 0.0567 | 0.0655 | 1.1547 | 1.0000 |
| cluster2853 | -23496.58764 | 0.0064 | 0.0115 | 1.7779 | 1.0000 |
| cluster2854 | -1780.556145 | 0.0303 | 0.0109 | 0.3602 | 1.0000 |
| cluster2856 | -15425.64679 | 0.1006 | 0.1266 | 1.2581 | 0.1205 |
| cluster2857 | -6019.551964 | 0.0216 | 0.0021 | 0.0980 | 0.0114 |
| cluster2858 | -7754.717043 | 0.0026 | 0.0028 | 1.1092 | 0.0045 |
| cluster2859 | -5667.898296 | 0.0116 | 0.0050 | 0.4354 | 0.0516 |
| cluster2860 | -16044.72396 | 0.0019 | 0.0012 | 0.6446 | 0.0017 |
| cluster2861 | -8361.464536 | 0.0020 | 0.0019 | 0.9206 | 0.0138 |
| cluster2862 | -18855.25576 | 0.0115 | 0.0070 | 0.6102 | 0.0168 |
| cluster2863 | -15064.89947 | 0.0197 | 0.0278 | 1.4106 | 0.0372 |
| cluster2864 | -12550.96769 | 0.0028 | 0.0021 | 0.7477 | 0.0041 |
| cluster2865 | -14997.66143 | 0.0636 | 0.0222 | 0.3499 | 0.1139 |
| cluster2866 | -17305.29864 | 0.0195 | 0.0252 | 1.2913 | 0.0336 |
| cluster2867 | -14859.89907 | 0.0371 | 0.0594 | 1.6006 | 0.0559 |
| cluster2869 | -8916.751785 | 0.0841 | 0.0496 | 0.5899 | 0.0947 |
| cluster2870 | -3880.711345 | 0.0643 | 0.0343 | 0.5338 | 0.0510 |
| cluster2871 | -3137.296668 | 0.0074 | 0.0019 | 0.2516 | 0.0305 |
| cluster2872 | -12260.1214 | 0.0741 | 0.0746 | 1.0061 | 0.0879 |
| cluster2873 | -13845.99774 | 0.0084 | 0.0038 | 0.4533 | 0.0186 |
| cluster2874 | -10383.34712 | 0.0246 | 0.0174 | 0.7096 | 0.0430 |
| cluster2875 | -13632.07206 | 0.0102 | 0.0098 | 0.9587 | 0.0194 |
| cluster2876 | -3971.175413 | 0.0274 | 0.0120 | 0.4391 | 0.0387 |
| cluster2877 | -10633.20601 | 0.0331 | 0.0175 | 0.5278 | 0.0589 |
| cluster2878 | -7059.311809 | 0.0326 | 0.0088 | 0.2687 | 0.0417 |
| cluster2879 | -10034.88967 | 0.2336 | 0.0487 | 0.2084 | 0.1774 |
| cluster2880 | -10641.39086 | 0.0427 | 0.0534 | 1.2517 | 0.0527 |
| cluster2881 | -11521.07836 | 0.0392 | 0.0172 | 0.4398 | 0.0579 |
| cluster2882 | -10648.97149 | 0.0252 | 0.0192 | 0.7630 | 0.0328 |
| cluster2883 | -6764.998811 | 0.0738 | 0.0245 | 0.3322 | 0.0691 |
| cluster2884 | -12071.19145 | 0.0234 | 0.0175 | 0.7468 | 0.0269 |
| cluster2885 | -15738.25998 | 0.0351 | 0.0478 | 1.3600 | 0.0491 |
| cluster2886 | -12003.58931 | 0.0326 | 0.0130 | 0.3999 | 0.0431 |
| cluster2887 | -8367.276007 | 0.0042 | 0.0050 | 1.1959 | 0.0077 |
| cluster2888 | -7786.254836 | 0.0586 | 0.0993 | 1.6946 | 0.0684 |
| cluster2889 | -6926.895499 | 0.0044 | 0.0068 | 1.5698 | 0.0105 |
| cluster2890 | -2677.549826 | 0.2759 | 0.0009 | 0.0031 | 0.0740 |
| cluster2891 | -12095.72028 | 0.0302 | 0.0306 | 1.0119 | 0.0436 |
| cluster2892 | -7908.664528 | 0.0068 | 0.0033 | 0.4855 | 0.0178 |
| cluster2893 | -8467.462904 | 0.0271 | 0.0487 | 1.7954 | 0.0411 |
| cluster2894 | -5507.241395 | 0.0063 | 0.0041 | 0.6502 | 0.0219 |
| cluster2895 | -8446.542115 | 0.0278 | 0.0366 | 1.3200 | 0.0411 |

| | | | | | |
|---|---|---|---|---|---|
| cluster2896 | -10286.5694 | 0.0079 | 0.0112 | 1.4243 | 0.0112 |
| cluster2897 | -9153.428508 | 0.0212 | 0.0177 | 0.8360 | 0.0399 |
| cluster2898 | -4500.198665 | 0.0338 | 0.0141 | 0.4172 | 0.0229 |
| cluster2899 | -8004.124437 | 0.0115 | 0.0077 | 0.6647 | 0.0227 |
| cluster2900 | -9924.15163 | 0.0124 | 0.0059 | 0.4732 | 0.0132 |
| cluster2901 | -9240.753861 | 0.0336 | 0.0442 | 1.3172 | 0.0390 |
| cluster2902 | -3696.707288 | 0.0268 | 0.0392 | 1.4603 | 0.0262 |
| cluster2903 | -8863.859297 | 0.0091 | 0.0065 | 0.7121 | 0.0111 |
| cluster2904 | -6412.675234 | 0.0601 | 0.0167 | 0.2779 | 0.1023 |
| cluster2905 | -8874.851805 | 0.0149 | 0.0052 | 0.3513 | 0.0349 |
| cluster2906 | -13664.15733 | 0.0541 | 0.0202 | 0.3741 | 0.0825 |
| cluster2907 | -12738.44979 | 0.0599 | 0.0873 | 1.4568 | 0.0803 |
| cluster2908 | -5642.501158 | 0.0046 | 0.0017 | 0.3825 | 0.0148 |
| cluster2909 | -2924.649235 | 0.0269 | 0.0104 | 0.3876 | 0.0420 |
| cluster2910 | -9270.300896 | 0.0363 | 0.0386 | 1.0615 | 0.0464 |
| cluster2911 | -4004.904608 | 0.0134 | 0.0014 | 0.1058 | 0.0021 |
| cluster2912 | -8768.618137 | 0.0230 | 0.0308 | 1.3403 | 0.0276 |
| cluster2913 | -6297.243511 | 0.0014 | 0.0011 | 0.8028 | 0.0030 |
| cluster2914 | -8063.662563 | 0.0135 | 0.0041 | 0.3067 | 0.0168 |
| cluster2915 | -4203.527631 | 0.0078 | 0.0081 | 1.0308 | 0.0205 |
| cluster2916 | -9768.545182 | 0.0233 | 0.0125 | 0.5366 | 0.0352 |
| cluster2917 | -9021.482005 | 0.0116 | 0.0116 | 0.9980 | 0.0242 |
| cluster2918 | -4475.227766 | 0.0618 | 0.0263 | 0.4252 | 0.0768 |
| cluster2919 | -9565.042304 | 0.0626 | 0.0238 | 0.3811 | 0.0688 |
| cluster2920 | -6957.419061 | 0.0156 | 0.0302 | 1.9372 | 0.0156 |
| cluster2921 | -3809.182828 | 0.0035 | 0.0038 | 1.0876 | 0.0042 |
| cluster2922 | -6244.365595 | 0.0503 | 0.0518 | 1.0287 | 0.0595 |
| cluster2923 | -5843.755798 | 0.0349 | 0.0095 | 0.2725 | 0.2322 |
| cluster2924 | -10975.13632 | 0.0172 | 0.0243 | 1.4161 | 0.0247 |
| cluster2925 | -5771.585495 | 0.0345 | 0.0359 | 1.0413 | 0.0328 |
| cluster2926 | -11219.39698 | 0.0178 | 0.0121 | 0.6759 | 0.0226 |
| cluster2927 | -4904.174485 | 0.0354 | 0.0445 | 1.2572 | 0.0292 |
| cluster2928 | -7178.159059 | 0.0148 | 0.0206 | 1.3870 | 0.0248 |
| cluster2929 | -8009.362729 | 0.0033 | 0.0032 | 0.9685 | 0.0065 |
| cluster2930 | -10988.59155 | 0.0771 | 0.0525 | 0.6809 | 0.0874 |
| cluster2931 | -6866.073182 | 0.0460 | 0.0693 | 1.5075 | 0.0560 |
| cluster2932 | -9692.683913 | 0.0506 | 0.0573 | 1.1322 | 0.0659 |
| cluster2933 | -8734.233021 | 0.0664 | 0.0170 | 0.2565 | 0.0412 |
| cluster2934 | -10531.82554 | 0.0646 | 0.0360 | 0.5573 | 0.0631 |
| cluster2935 | -4951.423234 | 0.0196 | 0.0073 | 0.3735 | 0.0321 |
| cluster2936 | -9510.040836 | 0.0295 | 0.0584 | 1.9789 | 0.0404 |
| cluster2937 | -4595.580846 | 0.0257 | 0.0121 | 0.4715 | 0.0289 |
| cluster2938 | -7241.421547 | 0.0147 | 0.0071 | 0.4853 | 0.0156 |
| cluster2939 | -2815.163079 | 0.0075 | 0.0017 | 0.2278 | 0.0288 |

| | | | | | |
|---|---|---|---|---|---|
| cluster2940 | -11377.06931 | 0.0567 | 0.0534 | 0.9406 | 0.0718 |
| cluster2941 | -3030.371562 | 0.0001 | 0.0000 | 0.1129 | 0.0023 |
| cluster2942 | -11359.56887 | 0.0528 | 0.0583 | 1.1047 | 0.0546 |
| cluster2943 | -4040.364417 | 0.0096 | 0.0064 | 0.6688 | 0.0112 |
| cluster2944 | -8489.921754 | 0.0637 | 0.0320 | 0.5013 | 0.1180 |
| cluster2945 | -3084.836818 | 0.0016 | 0.0023 | 1.4632 | 0.0025 |
| cluster2946 | -6740.260388 | 0.0035 | 0.0006 | 0.1707 | 0.0112 |
| cluster2947 | -9651.772767 | 0.0191 | 0.0115 | 0.5998 | 0.0438 |
| cluster2948 | -9373.614707 | 0.0384 | 0.0218 | 0.5674 | 0.0629 |
| cluster2949 | -6811.852451 | 0.0091 | 0.0044 | 0.4837 | 0.0209 |
| cluster2950 | -7099.783009 | 0.0035 | 0.0020 | 0.5579 | 0.0226 |
| cluster2951 | -9094.266932 | 0.0688 | 0.0542 | 0.7880 | 0.0591 |
| cluster2952 | -10895.41521 | 0.0404 | 0.0456 | 1.1306 | 0.0446 |
| cluster2953 | -13289.80445 | 0.0604 | 0.0744 | 1.2306 | 0.0752 |
| cluster2954 | -7271.571383 | 0.0115 | 0.0068 | 0.5927 | 0.0239 |
| cluster2955 | -6023.35876 | 0.0120 | 0.0105 | 0.8753 | 0.0307 |
| cluster2956 | -10129.87169 | 0.0081 | 0.0080 | 0.9833 | 0.0177 |
| cluster2957 | -8382.461686 | 0.0639 | 0.0902 | 1.4133 | 0.1121 |
| cluster2958 | -9689.47781 | 0.0086 | 0.0106 | 1.2382 | 0.0132 |
| cluster2959 | -5169.247966 | 0.0604 | 0.0176 | 0.2916 | 0.0503 |
| cluster2960 | -9437.546991 | 0.0147 | 0.0116 | 0.7909 | 0.0280 |
| cluster2961 | -8397.357134 | 0.0070 | 0.0094 | 1.3395 | 0.0110 |
| cluster2962 | -7494.86441 | 0.0174 | 0.0204 | 1.1721 | 0.0256 |
| cluster2963 | -7222.520362 | 0.0266 | 0.0074 | 0.2772 | 0.0400 |
| cluster2964 | -5242.614124 | 0.0146 | 0.0047 | 0.3207 | 0.0408 |
| cluster2965 | -7389.386373 | 0.0167 | 0.0129 | 0.7765 | 0.0097 |
| cluster2966 | -5587.633677 | 0.0204 | 0.0302 | 1.4813 | 0.0239 |
| cluster2967 | -9577.612766 | 0.0126 | 0.0063 | 0.5012 | 0.0195 |
| cluster2968 | -5455.734027 | 0.0131 | 0.0094 | 0.7140 | 0.0206 |
| cluster2969 | -7910.619386 | 0.0422 | 0.0361 | 0.8548 | 0.0427 |
| cluster2970 | -4235.383492 | 0.0020 | 0.0021 | 1.0587 | 0.0044 |
| cluster2971 | -3607.738319 | 0.0165 | 0.0033 | 0.2008 | 0.0345 |
| cluster2972 | -2669.63607 | 0.0035 | 0.0011 | 0.3001 | 0.0072 |
| cluster2973 | -6354.783635 | 0.0577 | 0.0763 | 1.3210 | 0.1120 |
| cluster2974 | -4418.100287 | 0.0109 | 0.0046 | 0.4203 | 0.0360 |
| cluster2975 | -10297.92373 | 0.0523 | 0.0467 | 0.8939 | 0.0823 |
| cluster2976 | -1468.562151 | 0.1129 | 0.0193 | 0.1710 | 0.2204 |
| cluster2977 | -3392.830598 | 0.0039 | 0.0094 | 2.4181 | 0.0051 |
| cluster2978 | -8215.51891 | 0.0017 | 0.0008 | 0.4606 | 0.0013 |
| cluster2979 | -8833.730596 | 0.0233 | 0.0170 | 0.7298 | 0.0454 |
| cluster2980 | -1553.810938 | 0.0290 | 0.0243 | 0.8374 | 0.0197 |
| cluster2981 | -6212.499155 | 0.0462 | 0.0491 | 1.0612 | 0.0610 |
| cluster2982 | -8045.275533 | 0.0466 | 0.0212 | 0.4551 | 0.1278 |
| cluster2983 | -7514.084847 | 0.0135 | 0.0201 | 1.4910 | 0.0294 |

| | | | | | |
|---|---|---|---|---|---|
| cluster2984 | -4830.697732 | 0.0219 | 0.0063 | 0.2898 | 0.0356 |
| cluster2985 | -3971.516608 | 0.3189 | 0.0777 | 0.2436 | 0.2154 |
| cluster2986 | -3494.709913 | 0.0704 | 0.0380 | 0.5407 | 0.1247 |
| cluster2987 | -6075.86229 | 0.0185 | 0.0085 | 0.4569 | 0.0334 |
| cluster2988 | -5703.438212 | 0.0238 | 0.0318 | 1.3358 | 0.0246 |
| cluster2989 | -6589.518742 | 0.0356 | 0.0118 | 0.3304 | 0.0409 |
| cluster2990 | -7509.986767 | 0.0337 | 0.0323 | 0.9593 | 0.0494 |
| cluster2991 | -6306.106094 | 0.0434 | 0.0431 | 0.9923 | 0.0716 |
| cluster2992 | -4178.949065 | 0.0100 | 0.0030 | 0.2966 | 0.0108 |
| cluster2993 | -5423.563803 | 0.0322 | 0.0216 | 0.6702 | 0.0325 |
| cluster2994 | -2258.622387 | 0.0001 | 0.0001 | 0.5409 | 0.0049 |
| cluster2995 | -740.225074 | 0.0214 | 0.0163 | 0.7601 | 0.0199 |
| cluster2996 | -7309.093365 | 0.0316 | 0.0174 | 0.5515 | 0.0485 |
| cluster2997 | -8170.839826 | 0.0329 | 0.0291 | 0.8867 | 0.0433 |
| cluster2998 | -7497.896758 | 0.0099 | 0.0105 | 1.0642 | 0.0232 |
| cluster2999 | -6763.774456 | 0.0267 | 0.0245 | 0.9158 | 0.0327 |
| cluster3000 | -6328.539481 | 0.0263 | 0.0088 | 0.3362 | 0.0089 |
| cluster3001 | -6058.351017 | 0.0277 | 0.0179 | 0.6481 | 0.0191 |
| cluster3002 | -7660.52322 | 0.0102 | 0.0062 | 0.6106 | 0.0316 |
| cluster3003 | -7809.957407 | 0.0730 | 0.0355 | 0.4859 | 0.0955 |
| cluster3004 | -5965.207873 | 0.0870 | 0.0138 | 0.1591 | 0.0964 |
| cluster3005 | -4101.330238 | 0.0821 | 0.0235 | 0.2867 | 0.0518 |
| cluster3006 | -8671.926149 | 0.0607 | 0.0325 | 0.5363 | 0.0558 |
| cluster3007 | -8244.006384 | 0.0279 | 0.0410 | 1.4726 | 0.0263 |
| cluster3008 | -5951.644955 | 0.0218 | 0.0241 | 1.1082 | 0.0364 |
| cluster3009 | -5912.039967 | 0.0317 | 0.0145 | 0.4570 | 0.0470 |
| cluster3010 | -8049.261046 | 0.0825 | 0.0330 | 0.3998 | 0.1738 |
| cluster3011 | -3370.636674 | 0.0054 | 0.0035 | 0.6469 | 0.0146 |
| cluster3012 | -5209.470104 | 0.0706 | 0.1095 | 1.5510 | 0.0536 |
| cluster3013 | -6732.279397 | 0.0578 | 0.0187 | 0.3227 | 0.0947 |
| cluster3014 | -3559.707764 | 0.0126 | 0.0172 | 1.3647 | 0.0056 |
| cluster3015 | -2410.178332 | 0.0317 | 0.0132 | 0.4154 | 0.0449 |
| cluster3016 | -4554.999114 | 0.0270 | 0.0134 | 0.4953 | 0.0434 |
| cluster3017 | -5966.546006 | 0.0106 | 0.0637 | 6.0190 | 0.0334 |
| cluster3018 | -5678.00011 | 0.0040 | 0.0036 | 0.9010 | 0.0074 |
| cluster3019 | -7565.762779 | 0.0329 | 0.0385 | 1.1706 | 0.0624 |
| cluster3020 | -6187.338489 | 0.0174 | 0.0187 | 1.0793 | 0.0427 |
| cluster3021 | -3023.867569 | 0.0080 | 0.0023 | 0.2929 | 0.0188 |
| cluster3022 | -3429.972554 | 0.0437 | 0.0535 | 1.2245 | 0.0733 |
| cluster3023 | -4201.267099 | 0.0042 | 0.0018 | 0.4417 | 0.0128 |
| cluster3024 | -1958.148839 | 0.0503 | 0.0191 | 0.3791 | 0.1015 |
| cluster3025 | -4691.177719 | 0.0057 | 0.0017 | 0.2936 | 0.0111 |
| cluster3026 | -4110.887863 | 0.1502 | 0.1174 | 0.7818 | 0.1809 |
| cluster3027 | -3436.590998 | 0.0019 | 0.0014 | 0.7026 | 0.0024 |

| | | | | | |
|---|---|---|---|---|---|
| cluster3028 | -6803.588661 | 0.0358 | 0.0418 | 1.1660 | 0.0438 |
| cluster3029 | -6674.017291 | 0.0201 | 0.0123 | 0.6100 | 0.0315 |
| cluster3030 | -5267.921094 | 0.0648 | 0.0160 | 0.2472 | 0.0499 |
| cluster3031 | -8797.255384 | 0.0965 | 0.0508 | 0.5263 | 0.0998 |
| cluster3032 | -5902.622116 | 0.0075 | 0.0060 | 0.7976 | 0.0338 |
| cluster3033 | -3824.034381 | 0.1420 | 0.0189 | 0.1331 | 0.1621 |
| cluster3034 | -4963.70237 | 0.0989 | 0.0262 | 0.2651 | 0.0324 |
| cluster3035 | -7642.49603 | 0.0143 | 0.0072 | 0.5002 | 0.0243 |
| cluster3036 | -8982.381097 | 0.0697 | 0.0706 | 1.0123 | 0.0716 |
| cluster3037 | -2558.372168 | 0.0001 | 0.0000 | 0.4531 | 0.0242 |
| cluster3038 | -4466.471113 | 0.0325 | 0.0245 | 0.7556 | 0.0741 |
| cluster3039 | -6528.980553 | 0.0323 | 0.0074 | 0.2301 | 0.0333 |
| cluster3040 | -4842.409113 | 0.0115 | 0.0035 | 0.3059 | 0.0223 |
| cluster3041 | -8446.298431 | 0.0355 | 0.0497 | 1.3979 | 0.0490 |
| cluster3042 | -4401.867864 | 0.0223 | 0.0264 | 1.1835 | 0.0399 |
| cluster3043 | -2917.647251 | 0.0205 | 0.0686 | 3.3477 | 0.0257 |
| cluster3044 | -6428.298541 | 0.0287 | 0.0244 | 0.8487 | 0.0305 |
| cluster3045 | -6128.84052 | 0.0548 | 0.0240 | 0.4377 | 0.0523 |
| cluster3046 | -7272.017695 | 0.0822 | 0.0180 | 0.2185 | 0.1991 |
| cluster3047 | -2120.945949 | 0.0001 | 0.0000 | 0.3472 | 0.0046 |
| cluster3048 | -5405.537595 | 0.0144 | 0.0115 | 0.8015 | 0.0251 |
| cluster3049 | -7061.079649 | 0.0077 | 0.0027 | 0.3489 | 0.0195 |
| cluster3050 | -7559.165181 | 0.0259 | 0.0147 | 0.5664 | 0.0454 |
| cluster3051 | -7512.583725 | 0.0329 | 0.0482 | 1.4633 | 0.0419 |
| cluster3052 | -6824.734961 | 0.0356 | 0.0537 | 1.5065 | 0.0544 |
| cluster3053 | -9367.268187 | 0.0663 | 0.0423 | 0.6387 | 0.1078 |
| cluster3054 | -6875.337109 | 0.0034 | 0.0028 | 0.8366 | 0.0114 |
| cluster3055 | -3977.602645 | 0.0032 | 0.0021 | 0.6540 | 0.0408 |
| cluster3056 | -5995.082693 | 0.0926 | 0.0178 | 0.1917 | 0.1034 |
| cluster3057 | -919.777936 | 0.0422 | 0.0064 | 0.1512 | 0.0563 |
| cluster3058 | -2534.206844 | 0.0094 | 0.0016 | 0.1757 | 0.0323 |
| cluster3059 | -8713.597649 | 0.0711 | 0.1169 | 1.6446 | 0.0669 |
| cluster3060 | -5326.294506 | 0.0287 | 0.0249 | 0.8688 | 0.0365 |
| cluster3061 | -5740.61354 | 0.0188 | 0.0060 | 0.3213 | 0.0147 |
| cluster3062 | -5688.832627 | 0.0222 | 0.0085 | 0.3814 | 0.0412 |
| cluster3063 | -6142.307356 | 0.0159 | 0.0108 | 0.6795 | 0.0388 |
| cluster3064 | -1916.690283 | 0.0001 | 0.0000 | 0.2515 | 0.0142 |
| cluster3065 | -2300.341732 | 0.0784 | 0.0553 | 0.7049 | 0.0989 |
| cluster3066 | -6187.590911 | 0.0721 | 0.0196 | 0.2722 | 0.0684 |
| cluster3067 | -4915.378505 | 0.0205 | 0.0121 | 0.5893 | 0.0510 |
| cluster3068 | -5143.443057 | 0.1026 | 0.0168 | 0.1640 | 0.1117 |
| cluster3069 | -1632.273996 | 0.0001 | 0.0000 | 0.1214 | 0.0016 |
| cluster3070 | -5191.891446 | 0.0298 | 0.0178 | 0.5971 | 0.0395 |
| cluster3071 | -5850.922767 | 0.0069 | 0.0019 | 0.2777 | 0.0285 |

| | | | | | |
|---|---|---|---|---|---|
| cluster3072 | -7369.142133 | 0.0575 | 0.0636 | 1.1061 | 0.0959 |
| cluster3073 | -3569.312448 | 0.0039 | 0.0030 | 0.7663 | 0.0090 |
| cluster3074 | -7651.117609 | 0.0313 | 0.0374 | 1.1957 | 0.0286 |
| cluster3075 | -5426.189273 | 0.0268 | 0.0182 | 0.6788 | 0.0427 |
| cluster3076 | -1810.390031 | 0.0074 | 0.0048 | 0.6452 | 0.0561 |
| cluster3077 | -4385.079628 | 0.0241 | 0.0135 | 0.5576 | 0.0230 |
| cluster3078 | -5024.7252 | 0.0117 | 0.0078 | 0.6648 | 0.0256 |
| cluster3079 | -7621.965543 | 0.0355 | 0.0154 | 0.4343 | 0.0326 |
| cluster3080 | -5649.336424 | 0.0125 | 0.0105 | 0.8457 | 0.0175 |
| cluster3081 | -3875.536311 | 0.0072 | 0.0022 | 0.3076 | 0.0111 |
| cluster3082 | -7183.812733 | 0.0401 | 0.0370 | 0.9230 | 0.0667 |
| cluster3083 | -4151.925551 | 0.0601 | 0.0214 | 0.3556 | 0.0172 |
| cluster3084 | -3197.951209 | 0.0437 | 0.0041 | 0.0935 | 0.0360 |
| cluster3085 | -7344.446583 | 0.0210 | 0.0364 | 1.7378 | 0.0233 |
| cluster3086 | -3397.315351 | 0.0257 | 0.0073 | 0.2842 | 0.1239 |
| cluster3087 | -6564.010685 | 0.0093 | 0.0052 | 0.5657 | 0.0221 |
| cluster3088 | -6387.86234 | 0.0389 | 0.0261 | 0.6722 | 0.0750 |
| cluster3089 | -3685.653237 | 0.0697 | 0.0108 | 0.1550 | 0.0733 |
| cluster3090 | -4957.050133 | 0.0028 | 0.0024 | 0.8775 | 0.0104 |
| cluster3091 | -5910.283837 | 0.0415 | 0.0120 | 0.2886 | 0.0598 |
| cluster3092 | -4265.525202 | 0.0167 | 0.0106 | 0.6348 | 0.0176 |
| cluster3093 | -4546.688361 | 0.0075 | 0.0043 | 0.5731 | 0.0062 |
| cluster3094 | -7865.299986 | 0.0253 | 0.0386 | 1.5233 | 0.0323 |
| cluster3095 | -4869.271253 | 0.0035 | 0.0019 | 0.5347 | 0.0075 |
| cluster3096 | -6131.893881 | 0.0273 | 0.0200 | 0.7333 | 0.0359 |
| cluster3097 | -7077.26845 | 0.0174 | 0.0063 | 0.3649 | 0.0222 |
| cluster3098 | -1418.75125 | 0.0062 | 0.0079 | 1.2782 | 0.0075 |
| cluster3099 | -4269.89774 | 0.0183 | 0.0068 | 0.3729 | 0.0469 |
| cluster3100 | -4394.139006 | 0.0378 | 0.0292 | 0.7714 | 0.0442 |
| cluster3101 | -3432.404388 | 0.0235 | 0.0130 | 0.5521 | 0.0586 |
| cluster3102 | -2229.426447 | 0.0067 | 0.0059 | 0.8758 | 0.0198 |
| cluster3103 | -5213.678099 | 0.0244 | 0.0376 | 1.5433 | 0.0451 |
| cluster3104 | -4323.97733 | 0.0094 | 0.0063 | 0.6699 | 0.0257 |
| cluster3105 | -7972.65329 | 0.0295 | 0.0452 | 1.5333 | 0.0270 |
| cluster3106 | -5132.242171 | 0.0030 | 0.0011 | 0.3564 | 0.0027 |
| cluster3107 | -3137.431434 | 0.0302 | 0.0089 | 0.2956 | 0.0437 |
| cluster3108 | -4329.495389 | 0.1000 | 0.0687 | 0.6872 | 0.1605 |
| cluster3109 | -3460.605718 | 0.0180 | 0.0263 | 1.4654 | 0.0327 |
| cluster3110 | -6379.584807 | 0.0838 | 0.0381 | 0.4544 | 0.1175 |
| cluster3111 | -4889.428057 | 0.0312 | 0.0155 | 0.4949 | 0.0330 |
| cluster3112 | -6144.195381 | 0.0326 | 0.0387 | 1.1864 | 0.0480 |
| cluster3113 | -3063.924679 | 0.0472 | 0.0384 | 0.8140 | 0.0187 |
| cluster3114 | -4683.198151 | 0.0098 | 0.0044 | 0.4464 | 0.0383 |
| cluster3115 | -3776.397577 | 0.0112 | 0.0075 | 0.6703 | 0.0201 |

| | | | | | |
|---|---|---|---|---|---|
| cluster3116 | -2722.960175 | 0.0256 | 0.0225 | 0.8791 | 0.0437 |
| cluster3117 | -7492.733614 | 0.0285 | 0.0666 | 2.3349 | 0.0587 |
| cluster3118 | -1574.383876 | 0.0080 | 0.0016 | 0.1966 | 0.0402 |
| cluster3119 | -2982.630853 | 0.0337 | 0.0117 | 0.3461 | 0.0423 |
| cluster3120 | -4909.318416 | 0.0152 | 0.0104 | 0.6840 | 0.0253 |
| cluster3121 | -6821.153153 | 0.0242 | 0.0105 | 0.4351 | 0.0283 |
| cluster3122 | -3554.133727 | 0.0083 | 0.0028 | 0.3323 | 0.0276 |
| cluster3123 | -3827.339272 | 0.0050 | 0.0019 | 0.3887 | 0.0190 |
| cluster3124 | -6373.688558 | 0.0245 | 0.0201 | 0.8214 | 0.0448 |
| cluster3125 | -4802.144737 | 0.0498 | 0.0219 | 0.4406 | 0.0714 |
| cluster3126 | -7551.642144 | 0.0334 | 0.0203 | 0.6074 | 0.0471 |
| cluster3127 | -2606.430436 | 0.0039 | 0.0033 | 0.8331 | 0.0096 |
| cluster3128 | -2238.311211 | 0.0028 | 0.0011 | 0.3724 | 0.0019 |
| cluster3129 | -3263.053577 | 0.0240 | 0.0679 | 2.8345 | 0.0424 |
| cluster3130 | -5799.277627 | 0.0330 | 0.0433 | 1.3109 | 0.0374 |
| cluster3131 | -1763.489411 | 0.0036 | 0.0052 | 1.4290 | 0.0096 |
| cluster3132 | -5289.770016 | 0.0201 | 0.0061 | 0.3057 | 0.0263 |
| cluster3133 | -8924.620562 | 0.0521 | 0.0943 | 1.8104 | 0.0661 |
| cluster3134 | -3290.527925 | 0.0451 | 0.0137 | 0.3034 | 0.0631 |
| cluster3135 | -4399.939771 | 0.0148 | 0.0082 | 0.5525 | 0.0101 |
| cluster3136 | -3954.079995 | 0.0369 | 0.0436 | 1.1821 | 0.0383 |
| cluster3137 | -3086.489819 | 0.0113 | 0.0127 | 1.1271 | 0.0310 |
| cluster3138 | -4742.462421 | 0.0421 | 0.0336 | 0.7974 | 0.0586 |
| cluster3139 | -5753.552854 | 0.0770 | 0.1220 | 1.5839 | 0.0651 |
| cluster3140 | -3555.043956 | 0.0173 | 0.0079 | 0.4536 | 0.0536 |
| cluster3141 | -2414.347152 | 0.0220 | 0.0028 | 0.1265 | 0.0078 |
| cluster3142 | -1641.604663 | 0.0516 | 0.0073 | 0.1422 | 0.1183 |
| cluster3143 | -3226.259089 | 0.0071 | 0.0030 | 0.4249 | 0.0177 |
| cluster3144 | -2088.305038 | 0.0034 | 0.0141 | 4.1876 | 0.0063 |
| cluster3145 | -6530.199047 | 0.0070 | 0.0078 | 1.1164 | 0.0049 |
| cluster3146 | -1742.577095 | 0.0004 | 0.0016 | 3.8681 | 0.0025 |
| cluster3147 | -4471.454739 | 0.0032 | 0.0032 | 0.9902 | 0.0075 |
| cluster3148 | -6019.836172 | 0.0173 | 0.0116 | 0.6725 | 0.0334 |
| cluster3149 | -2372.122306 | 0.0001 | 0.0000 | 0.2200 | 0.0033 |
| cluster3150 | -4523.682155 | 0.0507 | 0.0163 | 0.3205 | 0.0724 |
| cluster3151 | -3975.103787 | 0.0586 | 0.0284 | 0.4853 | 0.0486 |
| cluster3152 | -8745.482862 | 0.0427 | 0.0861 | 2.0185 | 0.0515 |
| cluster3153 | -4516.929648 | 0.0385 | 0.0201 | 0.5206 | 0.0445 |
| cluster3154 | -5343.880062 | 0.0125 | 0.0118 | 0.9505 | 0.0158 |
| cluster3155 | -3007.687698 | 0.0001 | 0.0001 | 0.9577 | 0.0118 |
| cluster3156 | -6786.929986 | 0.0187 | 0.0249 | 1.3310 | 0.0305 |
| cluster3157 | -3279.102891 | 0.1493 | 0.0309 | 0.2073 | 0.0869 |
| cluster3158 | -3370.348925 | 0.0068 | 0.0046 | 0.6821 | 0.0194 |
| cluster3159 | -5812.448668 | 0.0384 | 0.0589 | 1.5351 | 0.0529 |

| | | | | | |
|---|---|---|---|---|---|
| cluster3160 | -1094.240564 | 0.1736 | 0.0311 | 0.1794 | 0.0222 |
| cluster3161 | -4978.976542 | 0.0174 | 0.0134 | 0.7686 | 0.0385 |
| cluster3162 | -3637.122133 | 0.0019 | 0.0009 | 0.4445 | 0.0117 |
| cluster3163 | -3428.807898 | 0.0252 | 0.0280 | 1.1109 | 0.0431 |
| cluster3164 | -6180.604267 | 0.0544 | 0.0577 | 1.0620 | 0.0591 |
| cluster3165 | -8047.657981 | 0.0421 | 0.0486 | 1.1542 | 0.0625 |
| cluster3166 | -4628.031927 | 0.0112 | 0.0066 | 0.5914 | 0.0321 |
| cluster3167 | -5313.663551 | 0.0947 | 0.0388 | 0.4095 | 0.0928 |
| cluster3168 | -7089.762512 | 0.0136 | 0.0118 | 0.8666 | 0.0198 |
| cluster3169 | -5029.533577 | 0.0264 | 0.0328 | 1.2421 | 0.0323 |
| cluster3170 | -1622.185559 | 0.0001 | 0.0000 | 0.0864 | 0.0153 |
| cluster3171 | -6150.947786 | 0.0264 | 0.0088 | 0.3321 | 0.0824 |
| cluster3172 | -6262.233385 | 0.0279 | 0.0149 | 0.5336 | 0.0280 |
| cluster3173 | -4457.558643 | 0.0129 | 0.0125 | 0.9755 | 0.0157 |
| cluster3174 | -1300.677192 | 0.0001 | 0.0001 | 0.6713 | 0.0026 |
| cluster3175 | -1938.122771 | 0.0189 | 0.0093 | 0.4915 | 0.0151 |
| cluster3176 | -3887.565836 | 0.0192 | 0.0109 | 0.5674 | 0.0264 |
| cluster3177 | -3506.540621 | 0.0411 | 0.0065 | 0.1576 | 0.0565 |
| cluster3178 | -6737.768013 | 0.0038 | 0.0032 | 0.8359 | 0.0175 |
| cluster3179 | -4626.5728 | 0.0374 | 0.1072 | 2.8714 | 0.0817 |
| cluster3180 | -1071.249014 | 0.0035 | 0.0025 | 0.7183 | 0.0032 |
| cluster3181 | -6693.341426 | 0.0196 | 0.0147 | 0.7476 | 0.0223 |
| cluster3182 | -2530.616274 | 0.0062 | 0.0014 | 0.2226 | 0.0069 |
| cluster3183 | -4042.079879 | 0.0673 | 0.0527 | 0.7837 | 0.0874 |
| cluster3184 | -4206.243473 | 0.0209 | 0.0091 | 0.4361 | 0.0353 |
| cluster3185 | -3546.602209 | 0.0191 | 0.0205 | 1.0764 | 0.0160 |
| cluster3186 | -2287.535817 | 0.0094 | 0.0051 | 0.5485 | 0.0355 |
| cluster3187 | -5392.600838 | 0.0150 | 0.0099 | 0.6607 | 0.0227 |
| cluster3188 | -4445.178279 | 0.1008 | 0.0290 | 0.2874 | 0.0865 |
| cluster3189 | -6227.352109 | 0.0311 | 0.0327 | 1.0503 | 0.0490 |
| cluster3190 | -5397.584093 | 0.0259 | 0.0514 | 1.9841 | 0.0339 |
| cluster3191 | -2831.570368 | 0.0471 | 0.0228 | 0.4845 | 0.0465 |
| cluster3192 | -5887.211709 | 0.0516 | 0.0178 | 0.3448 | 0.0519 |
| cluster3193 | -3450.33576 | 0.0088 | 0.0034 | 0.3795 | 0.0439 |
| cluster3194 | -6978.917025 | 0.0420 | 0.0469 | 1.1165 | 0.0705 |
| cluster3195 | -3951.655489 | 0.0561 | 0.0280 | 0.4997 | 0.0096 |
| cluster3196 | -2244.479812 | 0.0070 | 0.0011 | 0.1527 | 0.0034 |
| cluster3197 | -5446.255287 | 0.0404 | 0.0138 | 0.3415 | 0.0568 |
| cluster3198 | -6430.184241 | 0.0321 | 0.0365 | 1.1367 | 0.0209 |
| cluster3199 | -2977.226056 | 0.0033 | 0.0022 | 0.6681 | 0.0136 |
| cluster3200 | -4262.978941 | 0.0013 | 0.0008 | 0.5970 | 0.0168 |
| cluster3201 | -4476.690466 | 0.1074 | 0.0239 | 0.2227 | 0.1119 |
| cluster3202 | -6070.915745 | 0.0505 | 0.0260 | 0.5157 | 0.0405 |
| cluster3203 | -3017.331956 | 0.0452 | 0.0452 | 1.0005 | 0.0521 |

| | | | | | |
|---|---|---|---|---|---|
| cluster3204 | -1878.268875 | 0.0425 | 0.0059 | 0.1377 | 0.0379 |
| cluster3205 | -2015.722975 | 0.0044 | 0.0007 | 0.1609 | 0.0067 |
| cluster3206 | -3914.829607 | 0.0354 | 0.0324 | 0.9171 | 0.0338 |
| cluster3207 | -5498.177419 | 0.0081 | 0.0057 | 0.7043 | 0.0173 |
| cluster3208 | -3285.631245 | 0.0016 | 0.0004 | 0.2488 | 0.0060 |
| cluster3209 | -5548.110328 | 0.0144 | 0.0215 | 1.4933 | 0.0285 |
| cluster3210 | -3373.784402 | 0.0001 | 0.0001 | 1.0683 | 0.0042 |
| cluster3211 | -3434.974501 | 0.0279 | 0.0217 | 0.7783 | 0.0330 |
| cluster3212 | -2811.639502 | 0.0015 | 0.0002 | 0.1095 | 0.0026 |
| cluster3213 | -4614.870608 | 0.0800 | 0.0413 | 0.5165 | 0.1004 |
| cluster3214 | -6049.068319 | 0.0328 | 0.0231 | 0.7060 | 0.0558 |
| cluster3215 | -2279.0588 | 0.0001 | 0.0000 | 0.1041 | 0.0027 |
| cluster3216 | -4536.572345 | 0.0194 | 0.0593 | 3.0545 | 0.0349 |
| cluster3217 | -3824.872844 | 0.0478 | 0.0106 | 0.2215 | 0.0407 |
| cluster3218 | -5084.211379 | 0.0265 | 0.0285 | 1.0754 | 0.0234 |
| cluster3219 | -5447.37769 | 0.0064 | 0.0052 | 0.8128 | 0.0144 |
| cluster3220 | -2923.092162 | 0.0161 | 0.0036 | 0.2257 | 0.0362 |
| cluster3221 | -5731.136001 | 0.0191 | 0.0427 | 2.2412 | 0.0415 |
| cluster3222 | -2621.667689 | 0.0539 | 0.0147 | 0.2720 | 0.0374 |
| cluster3223 | -3463.663105 | 0.1460 | 0.1630 | 1.1164 | 0.0757 |
| cluster3224 | -1597.243525 | 0.0001 | 0.0000 | 0.1594 | 0.0001 |
| cluster3225 | -6103.011073 | 0.0220 | 0.0275 | 1.2501 | 0.0369 |
| cluster3226 | -3719.49898 | 0.0128 | 0.0096 | 0.7478 | 0.0501 |
| cluster3227 | -4398.212167 | 0.0595 | 0.0732 | 1.2299 | 0.0740 |
| cluster3228 | -3624.799271 | 0.0242 | 0.0434 | 1.7915 | 0.0364 |
| cluster3229 | -3862.810962 | 0.0076 | 0.0029 | 0.3855 | 0.0392 |
| cluster3230 | -3482.398035 | 0.0102 | 0.0032 | 0.3185 | 0.0113 |
| cluster3231 | -4527.019153 | 0.0152 | 0.0111 | 0.7280 | 0.0277 |
| cluster3232 | -5053.89461 | 0.0206 | 0.0238 | 1.1523 | 0.0234 |
| cluster3233 | -3434.943979 | 0.0275 | 0.0257 | 0.9357 | 0.0519 |
| cluster3234 | -3820.357415 | 0.0365 | 0.0162 | 0.4442 | 0.0563 |
| cluster3235 | -7271.955522 | 0.0368 | 0.0561 | 1.5256 | 0.0382 |
| cluster3236 | -3956.614024 | 0.0305 | 0.0353 | 1.1582 | 0.0341 |
| cluster3237 | -3244.691907 | 0.0535 | 0.0513 | 0.9602 | 0.0602 |
| cluster3238 | -2777.454964 | 0.0050 | 0.0010 | 0.1918 | 0.0714 |
| cluster3239 | -2740.488997 | 0.0176 | 0.0083 | 0.4693 | 0.0242 |
| cluster3240 | -6960.445972 | 0.0231 | 0.0119 | 0.5163 | 0.0363 |
| cluster3241 | -5756.176062 | 0.0570 | 0.0445 | 0.7800 | 0.0677 |
| cluster3242 | -7161.334488 | 0.0315 | 0.0675 | 2.1433 | 0.0510 |
| cluster3243 | -5983.773094 | 0.1305 | 0.0271 | 0.2076 | 0.1627 |
| cluster3244 | -6470.947456 | 0.0432 | 0.0283 | 0.6541 | 0.0692 |
| cluster3245 | -2584.577809 | 0.0009 | 0.0010 | 1.1269 | 0.0037 |
| cluster3246 | -5133.304344 | 0.0288 | 0.0211 | 0.7313 | 0.0497 |
| cluster3247 | -848.241699 | 0.0342 | 0.0208 | 0.6098 | 0.0766 |

| | | | | | |
|---|---|---|---|---|---|
| cluster3248 | -3484.145445 | 0.0146 | 0.0137 | 0.9330 | 0.0239 |
| cluster3249 | -3482.721678 | 0.0128 | 0.0113 | 0.8858 | 0.0181 |
| cluster3250 | -3550.249449 | 0.1709 | 0.0300 | 0.1758 | 0.0939 |
| cluster3251 | -4467.195393 | 0.0488 | 0.0331 | 0.6780 | 0.0611 |
| cluster3252 | -7422.980618 | 0.0638 | 0.0968 | 1.5178 | 0.0637 |
| cluster3253 | -4316.179653 | 0.0534 | 0.0480 | 0.8983 | 0.0789 |
| cluster3254 | -3109.775024 | 0.0265 | 0.0272 | 1.0276 | 0.0383 |
| cluster3255 | -5570.75476 | 0.0164 | 0.0215 | 1.3145 | 0.0183 |
| cluster3256 | -6095.248937 | 0.0236 | 0.0284 | 1.2070 | 0.0492 |
| cluster3257 | -5141.279171 | 0.0018 | 0.0009 | 0.5217 | 0.0080 |
| cluster3258 | -1918.82522 | 0.0001 | 0.0000 | 0.4156 | 0.0017 |
| cluster3259 | -3556.436844 | 0.0400 | 0.0071 | 0.1777 | 0.0345 |
| cluster3260 | -4482.406024 | 0.0156 | 0.0056 | 0.3573 | 0.0132 |
| cluster3261 | -4943.252058 | 0.0386 | 0.1001 | 2.5929 | 0.0831 |
| cluster3262 | -5790.446384 | 0.0143 | 0.0224 | 1.5619 | 0.0205 |
| cluster3263 | -2048.206626 | 0.0134 | 0.0035 | 0.2591 | 0.1550 |
| cluster3264 | -1834.614617 | 0.0039 | 0.0012 | 0.3090 | 0.0121 |
| cluster3265 | -4128.745855 | 0.0022 | 0.0019 | 0.8859 | 0.0032 |
| cluster3266 | -2849.390647 | 0.0001 | 0.0000 | 0.2254 | 0.0360 |
| cluster3267 | -4123.969326 | 0.0234 | 0.0137 | 0.5880 | 0.0360 |
| cluster3268 | -2807.423634 | 0.0380 | 0.0263 | 0.6918 | 0.0372 |
| cluster3269 | -3533.72913 | 0.0288 | 0.0197 | 0.6837 | 0.0730 |
| cluster3270 | -2180.730587 | 0.0088 | 0.0143 | 1.6269 | 0.0346 |
| cluster3271 | -4109.105119 | 0.0050 | 0.0040 | 0.7906 | 0.0161 |
| cluster3272 | -938.361462 | 0.4222 | 0.0534 | 0.1264 | 0.1731 |
| cluster3273 | -4005.525758 | 0.1041 | 0.0988 | 0.9493 | 0.1209 |
| cluster3274 | -2033.094073 | 0.0266 | 0.0319 | 1.2020 | 0.0347 |
| cluster3275 | -4550.343853 | 0.0392 | 0.0419 | 1.0682 | 0.0543 |
| cluster3276 | -2485.014512 | 0.0626 | 0.0061 | 0.0978 | 0.0994 |
| cluster3277 | -3227.749345 | 0.0435 | 0.0401 | 0.9227 | 0.0540 |
| cluster3278 | -4576.983259 | 0.0014 | 0.0006 | 0.4377 | 0.0037 |
| cluster3279 | -3732.816887 | 0.0194 | 0.0186 | 0.9596 | 0.0158 |
| cluster3280 | -3787.172697 | 0.0128 | 0.0062 | 0.4858 | 0.0248 |
| cluster3281 | -1967.23845 | 0.0172 | 0.0224 | 1.2998 | 0.0222 |
| cluster3282 | -2049.200846 | 0.0059 | 0.0049 | 0.8223 | 0.0055 |
| cluster3283 | -5464.87002 | 0.0675 | 0.0265 | 0.3929 | 0.0633 |
| cluster3284 | -5979.730858 | 0.0403 | 0.0388 | 0.9624 | 0.0322 |
| cluster3285 | -2359.641516 | 0.0001 | 0.0000 | 0.4184 | 0.0097 |
| cluster3286 | -2209.11753 | 0.0260 | 0.0061 | 0.2360 | 0.0310 |
| cluster3287 | -1376.194923 | 0.0039 | 0.0025 | 0.6473 | 0.0159 |
| cluster3288 | -1237.677991 | 0.0009 | 0.0024 | 2.7915 | 0.0100 |
| cluster3289 | -5796.738597 | 0.0335 | 0.0103 | 0.3061 | 0.0550 |
| cluster3290 | -1252.876965 | 0.0162 | 0.0149 | 0.9253 | 0.0256 |
| cluster3291 | -4784.990875 | 0.0228 | 0.0130 | 0.5700 | 0.0470 |

| | | | | | |
|---|---|---|---|---|---|
| cluster3292 | -2774.405067 | 0.0249 | 0.0130 | 0.5236 | 0.0268 |
| cluster3293 | -3183.75255 | 0.0060 | 0.0049 | 0.8114 | 0.0119 |
| cluster3294 | -2882.220944 | 0.0078 | 0.0035 | 0.4448 | 0.0168 |
| cluster3295 | -2240.628508 | 0.0001 | 0.0000 | 0.3861 | 0.0020 |
| cluster3296 | -1797.166949 | 0.0120 | 0.0098 | 0.8187 | 0.0027 |
| cluster3297 | -2352.417967 | 0.0798 | 0.0170 | 0.2129 | 0.2167 |
| cluster3298 | -2944.773577 | 0.0040 | 0.0020 | 0.4969 | 0.0040 |
| cluster3299 | -7907.955394 | 0.0584 | 0.1535 | 2.6283 | 0.0443 |
| cluster3300 | -2248.451102 | 0.0001 | 0.0001 | 1.0616 | 0.0065 |
| cluster3301 | -1804.162433 | 0.0366 | 0.0064 | 0.1737 | 0.0526 |
| cluster3302 | -4556.116574 | 0.0467 | 0.0302 | 0.6471 | 0.1003 |
| cluster3303 | -4480.460915 | 0.0297 | 0.0170 | 0.5714 | 0.0379 |
| cluster3304 | -2856.80487 | 0.0063 | 0.0013 | 0.2156 | 0.0310 |
| cluster3305 | -3802.795422 | 0.0412 | 0.0379 | 0.9215 | 0.0741 |
| cluster3306 | -5017.241552 | 0.0452 | 0.0485 | 1.0722 | 0.0842 |
| cluster3307 | -1671.372584 | 0.0047 | 0.0034 | 0.7221 | 0.0045 |
| cluster3308 | -4385.456236 | 0.0085 | 0.0034 | 0.4051 | 0.0229 |
| cluster3309 | -3383.540472 | 0.0843 | 0.0275 | 0.3258 | 0.1371 |
| cluster3310 | -4173.117051 | 0.0074 | 0.0035 | 0.4798 | 0.0183 |
| cluster3311 | -6155.989154 | 0.0383 | 0.0168 | 0.4398 | 0.0632 |
| cluster3312 | -1931.708784 | 0.0308 | 0.0535 | 1.7347 | 0.0247 |
| cluster3313 | -2508.66018 | 0.0085 | 0.0013 | 0.1504 | 0.0125 |
| cluster3314 | -7137.935864 | 0.0295 | 0.0611 | 2.0719 | 0.0546 |
| cluster3315 | -3721.282914 | 0.0020 | 0.0006 | 0.3186 | 0.0073 |
| cluster3316 | -5974.483513 | 0.0875 | 0.0843 | 0.9632 | 0.0840 |
| cluster3317 | -1141.483717 | 0.0001 | 0.0001 | 0.6638 | 0.0041 |
| cluster3318 | -3441.692737 | 0.0989 | 0.0936 | 0.9468 | 0.1147 |
| cluster3319 | -3769.466677 | 0.0394 | 0.0447 | 1.1363 | 0.0338 |
| cluster3320 | -2813.253338 | 0.0426 | 0.0461 | 1.0838 | 0.0608 |
| cluster3321 | -2917.397968 | 0.0014 | 0.0005 | 0.3426 | 0.0028 |
| cluster3322 | -3279.131764 | 0.0390 | 0.0477 | 1.2227 | 0.0428 |
| cluster3323 | -5520.626135 | 0.0493 | 0.0432 | 0.8774 | 0.0508 |
| cluster3324 | -1400.116621 | 0.0395 | 0.0075 | 0.1893 | 0.0180 |
| cluster3325 | -2060.490448 | 0.0134 | 0.0192 | 1.4299 | 0.0214 |
| cluster3326 | -1703.45422 | 0.0427 | 0.0075 | 0.1750 | 0.0835 |
| cluster3327 | -3863.922517 | 0.0173 | 0.0125 | 0.7205 | 0.0286 |
| cluster3328 | -2607.726753 | 0.0149 | 0.0061 | 0.4078 | 0.0093 |
| cluster3329 | -3151.532346 | 0.0330 | 0.0489 | 1.4804 | 0.0744 |
| cluster3330 | -4710.974561 | 0.0479 | 0.0757 | 1.5807 | 0.0473 |
| cluster3331 | -2860.1994 | 0.0161 | 0.0058 | 0.3596 | 0.0909 |
| cluster3332 | -6191.774219 | 0.0577 | 0.0832 | 1.4420 | 0.0648 |
| cluster3333 | -3575.864114 | 0.0207 | 0.0167 | 0.8054 | 0.0254 |
| cluster3334 | -1380.745331 | 0.0598 | 0.0266 | 0.4448 | 0.0778 |
| cluster3335 | -2857.795347 | 0.0209 | 0.0097 | 0.4626 | 0.0343 |

| | | | | | |
|---|---|---|---|---|---|
| cluster3336 | -3027.678211 | 0.0061 | 0.0020 | 0.3352 | 0.0029 |
| cluster3337 | -2619.362334 | 0.0436 | 0.0118 | 0.2707 | 0.0252 |
| cluster3338 | -6302.828105 | 0.0337 | 0.0599 | 1.7765 | 0.0414 |
| cluster3339 | -1910.254582 | 0.0600 | 0.0334 | 0.5568 | 0.0414 |
| cluster3340 | -1624.291472 | 0.0001 | 0.0000 | 0.0222 | 0.1822 |
| cluster3341 | -1576.25304 | 0.0166 | 0.0184 | 1.1089 | 0.0084 |
| cluster3342 | -2556.40873 | 0.0222 | 0.0249 | 1.1174 | 0.0209 |
| cluster3343 | -2926.171756 | 0.0001 | 0.0001 | 0.8753 | 0.0006 |
| cluster3344 | -3149.000199 | 0.0205 | 0.0064 | 0.3128 | 0.0290 |
| cluster3345 | -2792.807986 | 0.0063 | 0.0112 | 1.7984 | 0.0169 |
| cluster3346 | -3610.200651 | 0.0028 | 0.0011 | 0.4102 | 0.0038 |
| cluster3347 | -4176.133151 | 0.0470 | 0.0139 | 0.2963 | 0.0467 |
| cluster3348 | -4557.855029 | 0.0299 | 0.0112 | 0.3742 | 0.0553 |
| cluster3349 | -4213.375305 | 0.0042 | 0.0008 | 0.1988 | 0.0121 |
| cluster3350 | -4778.773099 | 0.0458 | 0.0477 | 1.0425 | 0.0519 |
| cluster3351 | -3132.810881 | 0.0090 | 0.0066 | 0.7393 | 0.0190 |
| cluster3352 | -4079.628238 | 0.0264 | 0.0150 | 0.5680 | 0.0690 |
| cluster3353 | -1584.760481 | 0.0032 | 0.0018 | 0.5593 | 0.0139 |
| cluster3354 | -2120.815096 | 0.0087 | 0.0021 | 0.2379 | 0.0052 |
| cluster3355 | -4281.043 | 0.0189 | 0.0205 | 1.0839 | 0.0519 |
| cluster3356 | -4593.870398 | 0.0237 | 0.0071 | 0.3002 | 0.0465 |
| cluster3357 | -1289.318627 | 0.0164 | 0.0120 | 0.7299 | 0.0325 |
| cluster3358 | -4169.857055 | 0.0001 | 0.0000 | 0.3902 | 0.0024 |
| cluster3359 | -6037.313171 | 0.0217 | 0.0307 | 1.4187 | 0.0480 |
| cluster3360 | -2471.523532 | 0.0436 | 0.0242 | 0.5561 | 0.0355 |
| cluster3361 | -5035.351525 | 0.0339 | 0.0368 | 1.0859 | 0.0389 |
| cluster3362 | -3690.153951 | 0.0079 | 0.0042 | 0.5283 | 0.0203 |
| cluster3363 | -3720.286466 | 0.0013 | 0.0022 | 1.7455 | 0.0029 |
| cluster3364 | -4356.293757 | 0.0248 | 0.0278 | 1.1201 | 0.0232 |
| cluster3365 | -5565.523118 | 0.0709 | 0.0627 | 0.8849 | 0.0588 |
| cluster3366 | -3149.556891 | 0.0302 | 0.0455 | 1.5073 | 0.0316 |
| cluster3367 | -3694.017918 | 0.1289 | 0.0376 | 0.2917 | 0.1354 |
| cluster3368 | -1650.847921 | 0.0041 | 0.0007 | 0.1712 | 0.0396 |
| cluster3369 | -3761.478472 | 0.0067 | 0.0098 | 1.4565 | 0.0192 |
| cluster3370 | -6386.547734 | 0.0799 | 0.0618 | 0.7731 | 0.1079 |
| cluster3371 | -2934.135711 | 0.0037 | 0.0018 | 0.4777 | 0.0070 |
| cluster3372 | -3840.990601 | 0.0055 | 0.0027 | 0.4897 | 0.0097 |
| cluster3373 | -5276.665553 | 0.0063 | 0.0094 | 1.5010 | 0.0097 |
| cluster3374 | -5125.071115 | 0.0612 | 0.1312 | 2.1444 | 0.0756 |
| cluster3375 | -3051.413251 | 0.0018 | 0.0001 | 0.0815 | 0.0066 |
| cluster3376 | -1530.808056 | 0.0107 | 0.0006 | 0.0539 | 0.0416 |
| cluster3377 | -2273.553512 | 0.0084 | 0.0132 | 1.5691 | 0.0133 |
| cluster3378 | -6507.734665 | 0.0417 | 0.0344 | 0.8253 | 0.0516 |
| cluster3379 | -1568.334487 | 0.0268 | 0.0154 | 0.5753 | 0.0361 |

| | | | | | |
|---|---|---|---|---|---|
| cluster3380 | -2652.637299 | 0.0095 | 0.0058 | 0.6104 | 0.0217 |
| cluster3381 | -2267.984093 | 0.0311 | 0.0067 | 0.2140 | 0.0346 |
| cluster3382 | -885.471804 | 0.0150 | 0.0025 | 0.1681 | 0.0056 |
| cluster3383 | -5142.599319 | 0.0281 | 0.0447 | 1.5912 | 0.0777 |
| cluster3384 | -1969.380523 | 0.0106 | 0.0036 | 0.3369 | 0.0426 |
| cluster3385 | -3770.875709 | 0.0283 | 0.0304 | 1.0769 | 0.0193 |
| cluster3386 | -4154.767735 | 0.0102 | 0.0133 | 1.3044 | 0.0165 |
| cluster3387 | -3031.298676 | 0.0417 | 0.0283 | 0.6795 | 0.0431 |
| cluster3388 | -2808.979324 | 0.0194 | 0.0147 | 0.7595 | 0.0272 |
| cluster3389 | -1788.697593 | 0.0076 | 0.0089 | 1.1801 | 0.0117 |
| cluster3390 | -5117.146173 | 0.0601 | 0.0239 | 0.3980 | 0.0654 |
| cluster3391 | -2415.756793 | 0.0043 | 0.0025 | 0.5827 | 0.0097 |
| cluster3392 | -2528.393075 | 0.0092 | 0.0131 | 1.4180 | 0.0108 |
| cluster3393 | -3178.225745 | 0.0400 | 0.0265 | 0.6625 | 0.0653 |
| cluster3394 | -713.243035 | 0.0092 | 0.0038 | 0.4157 | 0.0023 |
| cluster3395 | -2714.177427 | 0.0097 | 0.0033 | 0.3394 | 0.0165 |
| cluster3396 | -4984.804523 | 0.0462 | 0.0198 | 0.4286 | 0.0327 |
| cluster3397 | -4641.694914 | 0.0328 | 0.0316 | 0.9627 | 0.0260 |
| cluster3398 | -3830.45029 | 0.0315 | 0.0169 | 0.5373 | 0.0657 |
| cluster3399 | -3227.50657 | 0.0237 | 0.0161 | 0.6784 | 0.0585 |
| cluster3400 | -1951.535689 | 0.0304 | 0.0030 | 0.0993 | 0.1239 |
| cluster3401 | -2867.713938 | 0.0072 | 0.0057 | 0.7958 | 0.0205 |
| cluster3402 | -5114.76111 | 0.0292 | 0.0124 | 0.4246 | 0.0213 |
| cluster3403 | -3669.342313 | 0.0182 | 0.0140 | 0.7671 | 0.0235 |
| cluster3404 | -716.696126 | 0.0001 | 0.0000 | 0.1104 | 0.0001 |
| cluster3405 | -1537.834881 | 0.2201 | 0.0937 | 0.4259 | 0.0913 |
| cluster3406 | -3799.718461 | 0.0124 | 0.0056 | 0.4520 | 0.0217 |
| cluster3407 | -6244.104657 | 0.0534 | 0.0397 | 0.7427 | 0.0647 |
| cluster3408 | -1895.470002 | 0.0001 | 0.0000 | 0.2615 | 0.0077 |
| cluster3409 | -4909.496159 | 0.0221 | 0.0201 | 0.9103 | 0.0238 |
| cluster3410 | -1443.019507 | 0.0307 | 0.0056 | 0.1820 | 0.0220 |
| cluster3411 | -2582.793791 | 0.0001 | 0.0001 | 0.5269 | 0.0004 |
| cluster3412 | -3390.775517 | 0.1161 | 0.1200 | 1.0331 | 0.0734 |
| cluster3413 | -2560.129014 | 0.0330 | 0.0304 | 0.9191 | 0.0508 |
| cluster3414 | -2522.766319 | 0.1130 | 0.0172 | 0.1525 | 0.1430 |
| cluster3415 | -3491.031986 | 0.0384 | 0.0195 | 0.5071 | 0.0714 |
| cluster3416 | -599.340235 | 0.0001 | 0.0027 | 19.3027 | 0.2459 |
| cluster3417 | -1213.520003 | 0.0003 | 0.0003 | 1.4180 | 0.0055 |
| cluster3418 | -2532.87713 | 0.0355 | 0.0300 | 0.8450 | 0.0632 |
| cluster3419 | -2672.94644 | 0.0826 | 0.0251 | 0.3035 | 0.1033 |
| cluster3420 | -1785.415517 | 0.1930 | 0.1695 | 0.8785 | 0.2815 |
| cluster3421 | -4207.452349 | 0.0091 | 0.0096 | 1.0563 | 0.0122 |
| cluster3422 | -5399.283662 | 0.0377 | 0.0226 | 0.5999 | 0.0337 |
| cluster3423 | -4248.413786 | 0.0295 | 0.0697 | 2.3594 | 0.0561 |

| | | | | | |
|---|---|---|---|---|---|
| cluster3424 | -2854.233334 | 0.0667 | 0.0438 | 0.6573 | 0.1277 |
| cluster3425 | -2228.852034 | 0.0076 | 0.0018 | 0.2335 | 0.0515 |
| cluster3426 | -3410.497644 | 0.0120 | 0.0087 | 0.7250 | 0.0209 |
| cluster3427 | -3796.671633 | 0.1009 | 0.0542 | 0.5365 | 0.0898 |
| cluster3428 | -2685.959668 | 0.0606 | 0.0172 | 0.2837 | 0.0659 |
| cluster3429 | -4144.689316 | 0.0192 | 0.0103 | 0.5371 | 0.0322 |
| cluster3430 | -4824.685965 | 0.0414 | 0.0774 | 1.8687 | 0.0448 |
| cluster3431 | -4738.071204 | 0.0320 | 0.0408 | 1.2781 | 0.0478 |
| cluster3432 | -3781.642668 | 0.0246 | 0.0057 | 0.2334 | 0.0418 |
| cluster3433 | -5392.852631 | 0.1395 | 0.0489 | 0.3506 | 0.1790 |
| cluster3434 | -3294.398515 | 0.0247 | 0.0058 | 0.2332 | 0.0410 |
| cluster3435 | -4093.248198 | 0.0191 | 0.0225 | 1.1780 | 0.0222 |
| cluster3436 | -3133.412683 | 0.0341 | 0.0385 | 1.1293 | 0.0191 |
| cluster3437 | -2816.417333 | 0.0118 | 0.0062 | 0.5289 | 0.0135 |
| cluster3438 | -3824.947756 | 0.0411 | 0.0328 | 0.7963 | 0.0372 |
| cluster3439 | -2055.381373 | 0.0756 | 0.0398 | 0.5270 | 0.0438 |
| cluster3440 | -2313.929682 | 0.0180 | 0.0081 | 0.4514 | 0.0285 |
| cluster3441 | -2756.887451 | 0.0372 | 0.0392 | 1.0540 | 0.0379 |
| cluster3442 | -2920.960987 | 0.0325 | 0.0639 | 1.9675 | 0.0116 |
| cluster3443 | -3531.057006 | 0.0088 | 0.0110 | 1.2473 | 0.0173 |
| cluster3444 | -3384.87604 | 0.0040 | 0.0024 | 0.6041 | 0.0121 |
| cluster3445 | -2963.867809 | 0.0073 | 0.0019 | 0.2559 | 0.0171 |
| cluster3446 | -4061.575773 | 0.0254 | 0.0420 | 1.6515 | 0.0336 |
| cluster3447 | -5926.258671 | 0.0272 | 0.0653 | 2.4010 | 0.0388 |
| cluster3448 | -2378.755083 | 0.0260 | 0.0359 | 1.3796 | 0.0211 |
| cluster3449 | -613.276106 | 0.0001 | 0.0000 | 0.0849 | 0.0052 |
| cluster3450 | -2518.383043 | 0.0161 | 0.0159 | 0.9918 | 0.0218 |
| cluster3451 | -3594.230743 | 0.0264 | 0.0315 | 1.1910 | 0.0280 |
| cluster3452 | -3964.594092 | 0.0538 | 0.0627 | 1.1654 | 0.0445 |
| cluster3453 | -1849.691389 | 0.0377 | 0.0075 | 0.1986 | 0.0388 |
| cluster3454 | -3695.932596 | 0.0099 | 0.0149 | 1.4935 | 0.0113 |
| cluster3455 | -3649.325195 | 0.0435 | 0.0044 | 0.1021 | 0.0436 |
| cluster3456 | -4231.554101 | 0.0361 | 0.0165 | 0.4558 | 0.0368 |
| cluster3457 | -3648.968605 | 0.0244 | 0.0214 | 0.8793 | 0.0334 |
| cluster3458 | -1806.611085 | 0.0346 | 0.0095 | 0.2755 | 0.0340 |
| cluster3459 | -2638.960498 | 0.0287 | 0.0552 | 1.9264 | 0.0557 |
| cluster3460 | -5580.50216 | 0.0299 | 0.0551 | 1.8450 | 0.0336 |
| cluster3461 | -3102.87623 | 0.0705 | 0.0497 | 0.7056 | 0.0644 |
| cluster3462 | -822.914415 | 0.0001 | 0.0000 | 0.0929 | 0.0467 |
| cluster3463 | -4771.344968 | 0.0445 | 0.0338 | 0.7598 | 0.0492 |
| cluster3464 | -5603.725621 | 0.0279 | 0.0553 | 1.9815 | 0.0551 |
| cluster3465 | -3954.430985 | 0.0057 | 0.0059 | 1.0404 | 0.0131 |
| cluster3466 | -2643.003706 | 0.0066 | 0.0037 | 0.5563 | 0.0153 |
| cluster3467 | -4061.009448 | 0.0293 | 0.0207 | 0.7076 | 0.0342 |

| | | | | | |
|---|---|---|---|---|---|
| cluster3468 | -1848.387133 | 0.0029 | 0.0037 | 1.2795 | 0.0101 |
| cluster3469 | -2105.580284 | 0.0178 | 0.0190 | 1.0695 | 0.0247 |
| cluster3470 | -3884.562808 | 0.0112 | 0.0088 | 0.7855 | 0.0148 |
| cluster3471 | -4173.398464 | 0.0430 | 0.0384 | 0.8938 | 0.0428 |
| cluster3472 | -3720.759858 | 0.0109 | 0.0036 | 0.3303 | 0.0208 |
| cluster3473 | -3470.788924 | 0.0239 | 0.0078 | 0.3248 | 0.0287 |
| cluster3474 | -5365.284356 | 0.0220 | 0.0624 | 2.8345 | 0.0366 |
| cluster3475 | -2340.394347 | 0.0088 | 0.0113 | 1.2967 | 0.0142 |
| cluster3476 | -5109.934691 | 0.0205 | 0.0509 | 2.4895 | 0.0346 |
| cluster3477 | -3274.186131 | 0.0040 | 0.0007 | 0.1791 | 0.0203 |
| cluster3478 | -3579.220944 | 0.0092 | 0.0041 | 0.4507 | 0.0239 |
| cluster3479 | -2916.329603 | 0.0532 | 0.0662 | 1.2446 | 0.0450 |
| cluster3480 | -4509.147655 | 0.0341 | 0.0322 | 0.9441 | 0.0329 |
| cluster3481 | -1071.238242 | 0.0223 | 0.0041 | 0.1840 | 0.0038 |
| cluster3482 | -1967.191465 | 0.1031 | 0.0155 | 0.1506 | 0.0723 |
| cluster3483 | -3657.504796 | 0.0033 | 0.0026 | 0.7809 | 0.0120 |
| cluster3484 | -2766.960657 | 0.0176 | 0.0217 | 1.2330 | 0.0375 |
| cluster3485 | -3494.526985 | 0.0307 | 0.0197 | 0.6413 | 0.0454 |
| cluster3486 | -3175.106763 | 0.0485 | 0.0129 | 0.2652 | 0.0514 |
| cluster3487 | -2348.288126 | 0.0219 | 0.0129 | 0.5869 | 0.0250 |
| cluster3488 | -1482.609404 | 0.0143 | 0.0058 | 0.4066 | 0.0310 |
| cluster3489 | -4061.605279 | 0.0412 | 0.0588 | 1.4266 | 0.0464 |
| cluster3490 | -3432.545925 | 0.0169 | 0.0224 | 1.3299 | 0.0247 |
| cluster3491 | -3686.664869 | 0.0250 | 0.0183 | 0.7307 | 0.0343 |
| cluster3492 | -3372.365309 | 0.0410 | 0.0540 | 1.3183 | 0.0354 |
| cluster3493 | -4431.04338 | 0.0220 | 0.0167 | 0.7598 | 0.0402 |
| cluster3494 | -2865.229373 | 0.0085 | 0.0090 | 1.0596 | 0.0169 |
| cluster3495 | -2842.837642 | 0.0246 | 0.0211 | 0.8549 | 0.0314 |
| cluster3496 | -3231.774316 | 0.0211 | 0.0071 | 0.3355 | 0.0225 |
| cluster3497 | -1194.463348 | 0.0269 | 0.0018 | 0.0674 | 0.0106 |
| cluster3498 | -4133.379952 | 0.0628 | 0.1230 | 1.9572 | 0.0443 |
| cluster3499 | -3722.447648 | 0.0464 | 0.0766 | 1.6504 | 0.1051 |
| cluster3500 | -3424.169609 | 0.0175 | 0.0201 | 1.1482 | 0.0225 |
| cluster3501 | -2358.03023 | 0.0922 | 0.0037 | 0.0405 | 0.0216 |
| cluster3502 | -2166.37275 | 0.0049 | 0.0057 | 1.1799 | 0.0182 |
| cluster3503 | -1950.843056 | 0.0260 | 0.0178 | 0.6835 | 0.0551 |
| cluster3504 | -1140.767257 | 0.0170 | 0.0104 | 0.6119 | 0.0046 |
| cluster3505 | -2418.176206 | 0.0466 | 0.0854 | 1.8341 | 0.0555 |
| cluster3506 | -2548.437713 | 0.0151 | 0.0230 | 1.5221 | 0.0284 |
| cluster3507 | -5019.971647 | 0.0580 | 0.0579 | 0.9995 | 0.0525 |
| cluster3508 | -5633.9689 | 0.0263 | 0.0768 | 2.9250 | 0.0314 |
| cluster3509 | -2401.711985 | 0.0001 | 0.0000 | 0.2957 | 0.0012 |
| cluster3510 | -843.813626 | 0.1554 | 0.0686 | 0.4415 | 0.0940 |
| cluster3511 | -1018.351294 | 0.0150 | 0.0069 | 0.4620 | 0.0389 |

| | | | | | |
|---|---|---|---|---|---|
| cluster3512 | -2356.672712 | 0.0251 | 0.0099 | 0.3950 | 0.0350 |
| cluster3513 | -959.363349 | 0.0001 | 0.0000 | 0.0000 | 0.0711 |
| cluster3514 | -5158.885805 | 0.0348 | 0.0181 | 0.5188 | 0.0373 |
| cluster3515 | -2601.596822 | 0.0033 | 0.0028 | 0.8393 | 0.0171 |
| cluster3516 | -4129.34574 | 0.0197 | 0.0202 | 1.0259 | 0.0304 |
| cluster3517 | -2409.47051 | 0.0001 | 0.0000 | 0.3689 | 0.0030 |
| cluster3518 | -4782.103527 | 0.0523 | 0.0564 | 1.0772 | 0.0872 |
| cluster3519 | -2127.160605 | 0.0382 | 0.0267 | 0.7000 | 0.0431 |
| cluster3520 | -1193.985497 | 0.1423 | 0.0438 | 0.3076 | 0.1515 |
| cluster3521 | -4223.603121 | 0.0296 | 0.0359 | 1.2116 | 0.0411 |
| cluster3522 | -4754.017169 | 0.0349 | 0.0490 | 1.4059 | 0.0322 |
| cluster3523 | -1657.2976 | 0.0272 | 0.0327 | 1.2024 | 0.0309 |
| cluster3524 | -1963.536092 | 0.0118 | 0.0037 | 0.3172 | 0.0127 |
| cluster3525 | -2816.186019 | 0.0307 | 0.0143 | 0.4669 | 0.0501 |
| cluster3526 | -2996.833194 | 0.0001 | 0.0002 | 1.6101 | 0.0040 |
| cluster3527 | -4937.353968 | 0.0425 | 0.0762 | 1.7960 | 0.0503 |
| cluster3528 | -1967.272883 | 0.0042 | 0.0013 | 0.3088 | 0.0132 |
| cluster3529 | -3742.444864 | 0.0263 | 0.0459 | 1.7471 | 0.0338 |
| cluster3530 | -1951.230683 | 0.0117 | 0.0102 | 0.8738 | 0.0384 |
| cluster3531 | -2917.894679 | 0.0305 | 0.0264 | 0.8663 | 0.0594 |
| cluster3532 | -5557.76588 | 0.0597 | 0.0313 | 0.5248 | 0.0632 |
| cluster3533 | -2652.624162 | 0.0109 | 0.0047 | 0.4288 | 0.0309 |
| cluster3534 | -2560.438659 | 0.0746 | 0.0072 | 0.0958 | 0.0619 |
| cluster3535 | -3289.418627 | 0.0270 | 0.0352 | 1.3045 | 0.0187 |
| cluster3536 | -3556.904591 | 0.0435 | 0.0156 | 0.3576 | 0.0335 |
| cluster3537 | -3664.460119 | 0.0034 | 0.0032 | 0.9306 | 0.0079 |
| cluster3538 | -1386.76564 | 0.0001 | 0.0000 | 0.0854 | 0.0424 |
| cluster3539 | -2944.947612 | 0.0252 | 0.0468 | 1.8560 | 0.0409 |
| cluster3540 | -5362.920446 | 0.1177 | 0.1628 | 1.3837 | 0.0814 |
| cluster3541 | -3877.847454 | 0.0259 | 0.0103 | 0.3977 | 0.0278 |
| cluster3542 | -5427.613617 | 0.0601 | 0.0902 | 1.4998 | 0.0603 |
| cluster3543 | -4243.333648 | 0.0200 | 0.0488 | 2.4461 | 0.0453 |
| cluster3544 | -1088.624111 | 0.0080 | 0.0002 | 0.0191 | 0.0153 |
| cluster3545 | -5147.19654 | 0.0449 | 0.0609 | 1.3570 | 0.0463 |
| cluster3546 | -2678.231085 | 0.0128 | 0.0204 | 1.5907 | 0.0268 |
| cluster3547 | -1113.667396 | 0.0655 | 0.0039 | 0.0591 | 0.0036 |
| cluster3548 | -1302.956165 | 0.0001 | 0.0000 | 0.0681 | 0.0802 |
| cluster3549 | -3231.854339 | 0.0227 | 0.0087 | 0.3838 | 0.0441 |
| cluster3550 | -1223.294987 | 0.0001 | 0.0000 | 0.1777 | 0.0099 |
| cluster3551 | -2798.534144 | 0.0595 | 0.0126 | 0.2119 | 0.0460 |
| cluster3552 | -2385.943533 | 0.0276 | 0.0150 | 0.5442 | 0.0207 |
| cluster3553 | -2403.453674 | 0.0183 | 0.0149 | 0.8136 | 0.0385 |
| cluster3554 | -3822.133705 | 0.0153 | 0.0095 | 0.6213 | 0.0308 |
| cluster3555 | -1632.448337 | 0.0099 | 0.0141 | 1.4189 | 0.0273 |

| | | | | | |
|---|---|---|---|---|---|
| cluster3556 | -4204.195855 | 0.0169 | 0.0222 | 1.3141 | 0.0296 |
| cluster3557 | -1366.621812 | 0.0900 | 0.0526 | 0.5844 | 0.0292 |
| cluster3558 | -4775.478987 | 0.0487 | 0.0619 | 1.2720 | 0.0734 |
| cluster3559 | -2455.442131 | 0.0070 | 0.0049 | 0.6964 | 0.0229 |
| cluster3560 | -5093.798243 | 0.0381 | 0.0729 | 1.9148 | 0.0678 |
| cluster3561 | -1800.399153 | 0.0122 | 0.0041 | 0.3380 | 0.0372 |
| cluster3562 | -4275.909875 | 0.0256 | 0.0349 | 1.3632 | 0.0444 |
| cluster3563 | -1982.62498 | 0.0110 | 0.0032 | 0.2929 | 0.0159 |
| cluster3564 | -5023.069709 | 0.0547 | 0.0238 | 0.4353 | 0.0783 |
| cluster3565 | -3345.579061 | 0.0147 | 0.0243 | 1.6536 | 0.0493 |
| cluster3566 | -1889.706236 | 0.0302 | 0.0202 | 0.6700 | 0.0488 |
| cluster3567 | -2403.972764 | 0.0657 | 0.0395 | 0.6011 | 0.0673 |
| cluster3568 | -3168.075708 | 0.0320 | 0.0430 | 1.3432 | 0.0260 |
| cluster3569 | -3551.537639 | 0.0437 | 0.0603 | 1.3808 | 0.0381 |
| cluster3570 | -2401.038226 | 0.0187 | 0.0283 | 1.5134 | 0.0201 |
| cluster3571 | -2032.993374 | 0.0066 | 0.0062 | 0.9400 | 0.0323 |
| cluster3572 | -3453.449823 | 0.0416 | 0.0357 | 0.8587 | 0.0550 |
| cluster3573 | -3241.014745 | 0.0179 | 0.0140 | 0.7827 | 0.0040 |
| cluster3574 | -5131.776858 | 0.0794 | 0.0966 | 1.2175 | 0.0752 |
| cluster3575 | -3830.853834 | 0.0344 | 0.0423 | 1.2299 | 0.0424 |
| cluster3576 | -3274.906715 | 0.0416 | 0.0295 | 0.7089 | 0.0344 |
| cluster3577 | -2729.219491 | 0.0255 | 0.0125 | 0.4904 | 0.0497 |
| cluster3578 | -3390.230281 | 0.0221 | 0.0153 | 0.6931 | 0.0376 |
| cluster3579 | -2923.411182 | 0.0393 | 0.0254 | 0.6459 | 0.0225 |
| cluster3580 | -2417.926217 | 0.0098 | 0.0145 | 1.4689 | 0.0107 |
| cluster3581 | -1247.01822 | 0.0093 | 0.0065 | 0.6983 | 0.0070 |
| cluster3582 | -1106.170042 | 0.0001 | 0.0000 | 0.0772 | 0.0296 |
| cluster3583 | -2533.436155 | 0.0001 | 0.0001 | 0.5708 | 0.0001 |
| cluster3584 | -4869.860727 | 0.0316 | 0.0671 | 2.1232 | 0.0303 |
| cluster3585 | -2092.760052 | 0.0337 | 0.0079 | 0.2355 | 0.0755 |
| cluster3586 | -5224.82473 | 0.0661 | 0.0883 | 1.3360 | 0.0746 |
| cluster3587 | -3468.128652 | 0.0306 | 0.0877 | 2.8634 | 0.0627 |
| cluster3588 | -3355.890929 | 0.0031 | 0.0009 | 0.3031 | 0.0322 |
| cluster3589 | -3016.951397 | 0.0308 | 0.0361 | 1.1725 | 0.0493 |
| cluster3590 | -4181.130824 | 0.0467 | 0.0701 | 1.5014 | 0.0688 |
| cluster3591 | -2874.387524 | 0.0134 | 0.0029 | 0.2153 | 0.0115 |
| cluster3592 | -4705.228938 | 0.0310 | 0.0496 | 1.5974 | 0.0436 |
| cluster3593 | -3417.287075 | 0.0311 | 0.0591 | 1.9004 | 0.0593 |
| cluster3594 | -3177.628185 | 0.0518 | 0.0274 | 0.5298 | 0.0641 |
| cluster3595 | -2123.325663 | 0.0064 | 0.0016 | 0.2561 | 0.0141 |
| cluster3596 | -2316.232587 | 0.0303 | 0.0169 | 0.5575 | 0.0339 |
| cluster3597 | -4194.771113 | 0.0303 | 0.0174 | 0.5760 | 0.0293 |
| cluster3598 | -4253.675277 | 0.0293 | 0.0981 | 3.3502 | 0.0213 |
| cluster3599 | -2253.9302 | 0.0213 | 0.0202 | 0.9479 | 0.0259 |

| | | | | | |
|---|---|---|---|---|---|
| cluster3600 | -2858.255938 | 0.0095 | 0.0053 | 0.5566 | 0.0306 |
| cluster3601 | -3281.741698 | 0.0019 | 0.0029 | 1.4908 | 0.0089 |
| cluster3602 | -3065.722283 | 0.0018 | 0.0005 | 0.2514 | 0.0037 |
| cluster3603 | -4224.207416 | 0.0607 | 0.0535 | 0.8813 | 0.0488 |
| cluster3604 | -1058.291995 | 0.0632 | 0.0050 | 0.0798 | 0.0630 |
| cluster3605 | -3057.768769 | 0.0340 | 0.0314 | 0.9219 | 0.0397 |
| cluster3606 | -4323.592568 | 0.0262 | 0.0385 | 1.4723 | 0.0285 |
| cluster3607 | -2596.62466 | 0.0282 | 0.0300 | 1.0651 | 0.0413 |
| cluster3608 | -2887.843735 | 0.0145 | 0.0058 | 0.4025 | 0.0097 |
| cluster3609 | -4181.916609 | 0.0254 | 0.0440 | 1.7321 | 0.0324 |
| cluster3610 | -1394.1214 | 0.0176 | 0.0086 | 0.4884 | 0.0046 |
| cluster3611 | -2340.720966 | 0.0230 | 0.0457 | 1.9898 | 0.0286 |
| cluster3612 | -2828.35876 | 0.0122 | 0.0595 | 4.8699 | 0.0446 |
| cluster3613 | -4059.335259 | 0.0181 | 0.0083 | 0.4562 | 0.0184 |
| cluster3614 | -4306.710176 | 0.0179 | 0.0472 | 2.6360 | 0.0300 |
| cluster3615 | -1661.681755 | 0.0227 | 0.0024 | 0.1060 | 0.0583 |
| cluster3616 | -773.18706 | 0.0035 | 0.0013 | 0.3726 | 0.0202 |
| cluster3617 | -1931.169126 | 0.0346 | 0.0426 | 1.2335 | 0.0479 |
| cluster3618 | -1898.67718 | 0.1245 | 0.1020 | 0.8195 | 0.1962 |
| cluster3619 | -2552.076005 | 0.0026 | 0.0024 | 0.9489 | 0.0056 |
| cluster3620 | -3600.013918 | 0.0221 | 0.0152 | 0.6904 | 0.0640 |
| cluster3621 | -4764.650761 | 0.0273 | 0.0482 | 1.7645 | 0.0377 |
| cluster3622 | -2718.288546 | 0.0001 | 0.0001 | 0.5218 | 0.0163 |
| cluster3623 | -2271.731117 | 0.0084 | 0.0085 | 1.0109 | 0.0096 |
| cluster3624 | -2909.96204 | 0.0136 | 0.0137 | 1.0086 | 0.0139 |
| cluster3625 | -3305.08671 | 0.0021 | 0.0030 | 1.4323 | 0.0108 |
| cluster3626 | -1982.71187 | 0.0403 | 0.0140 | 0.3489 | 0.0255 |
| cluster3627 | -3000.322685 | 0.0043 | 0.0045 | 1.0466 | 0.0082 |
| cluster3628 | -3023.110106 | 0.0280 | 0.0469 | 1.6748 | 0.0277 |
| cluster3629 | -4163.819677 | 0.0077 | 0.0209 | 2.7205 | 0.0145 |
| cluster3630 | -1205.657277 | 0.0001 | 0.0001 | 0.5589 | 0.0231 |
| cluster3631 | -3589.765879 | 0.0055 | 0.0055 | 0.9969 | 0.0290 |
| cluster3632 | -2459.757552 | 0.0240 | 0.0266 | 1.1097 | 0.0331 |
| cluster3633 | -2503.275008 | 0.0024 | 0.0003 | 0.1095 | 0.0145 |
| cluster3634 | -936.245193 | 0.0001 | 0.0000 | 0.4611 | 0.0214 |
| cluster3635 | -1620.641063 | 0.0265 | 0.0011 | 0.0403 | 0.0479 |
| cluster3636 | -2528.313076 | 0.0494 | 0.0143 | 0.2903 | 0.0093 |
| cluster3637 | -1061.256668 | 0.0001 | 0.0000 | 0.3649 | 0.0199 |
| cluster3638 | -2728.629194 | 0.0099 | 0.0049 | 0.4934 | 0.0385 |
| cluster3639 | -1271.905064 | 0.0176 | 0.0043 | 0.2445 | 0.0402 |
| cluster3640 | -4762.263548 | 0.0448 | 0.0664 | 1.4818 | 0.0916 |
| cluster3641 | -1769.106553 | 0.0263 | 0.0059 | 0.2260 | 0.0265 |
| cluster3642 | -3356.555059 | 0.0223 | 0.0308 | 1.3825 | 0.0362 |
| cluster3643 | -3538.05634 | 0.0890 | 0.0725 | 0.8152 | 0.1076 |

| | | | | | |
|---|---|---|---|---|---|
| cluster3644 | -1913.053809 | 0.0243 | 0.0042 | 0.1725 | 0.0297 |
| cluster3645 | -1807.895416 | 0.0315 | 0.0276 | 0.8782 | 0.0304 |
| cluster3646 | -2633.811393 | 0.0050 | 0.0666 | 13.2700 | 0.0269 |
| cluster3647 | -3429.242617 | 0.0057 | 0.0150 | 2.6394 | 0.0029 |
| cluster3648 | -1759.490074 | 0.0009 | 0.0002 | 0.2550 | 0.0016 |
| cluster3649 | -1640.094008 | 0.0055 | 0.0038 | 0.6983 | 0.0114 |
| cluster3650 | -1256.862533 | 0.0031 | 0.0285 | 9.1185 | 0.0187 |
| cluster3651 | -3970.923419 | 0.0390 | 0.0212 | 0.5428 | 0.0523 |
| cluster3652 | -3309.51998 | 0.0164 | 0.0098 | 0.5973 | 0.0291 |
| cluster3653 | -1425.36361 | 0.0001 | 0.0001 | 0.6694 | 0.0052 |
| cluster3654 | -2837.591473 | 0.0073 | 0.0107 | 1.4577 | 0.0108 |
| cluster3655 | -2366.656358 | 0.0656 | 0.0656 | 1.0007 | 0.0664 |
| cluster3656 | -3108.259206 | 0.0197 | 0.0133 | 0.6728 | 0.0287 |
| cluster3657 | -3194.44669 | 0.0153 | 0.0152 | 0.9958 | 0.0149 |
| cluster3658 | -748.908668 | 0.0095 | 0.0015 | 0.1625 | 0.1350 |
| cluster3659 | -2269.517511 | 0.0445 | 0.0223 | 0.5012 | 0.0332 |
| cluster3660 | -3016.463276 | 0.0100 | 0.0510 | 5.0861 | 0.0235 |
| cluster3661 | -1818.945345 | 0.0001 | 0.0000 | 0.0767 | 0.0022 |
| cluster3662 | -2802.903929 | 0.0242 | 0.0333 | 1.3751 | 0.0424 |
| cluster3663 | -3132.766183 | 0.0343 | 0.0409 | 1.1929 | 0.0559 |
| cluster3664 | -2449.494188 | 0.0387 | 0.0899 | 2.3213 | 0.0400 |
| cluster3665 | -3602.537 | 0.0153 | 0.0227 | 1.4844 | 0.0236 |
| cluster3666 | -2710.951247 | 0.0260 | 0.0156 | 0.5997 | 0.0387 |
| cluster3667 | -1566.942432 | 0.0001 | 0.0000 | 0.2958 | 0.0269 |
| cluster3668 | -2980.454039 | 0.0104 | 0.0138 | 1.3209 | 0.0259 |
| cluster3669 | -2601.776577 | 0.0320 | 0.0277 | 0.8663 | 0.0393 |
| cluster3670 | -4034.141579 | 0.0349 | 0.0260 | 0.7466 | 0.0518 |
| cluster3671 | -3996.667338 | 0.0309 | 0.0350 | 1.1355 | 0.0294 |
| cluster3672 | -2500.319151 | 0.0357 | 0.0259 | 0.7265 | 0.0491 |
| cluster3673 | -4410.084623 | 0.0360 | 0.0930 | 2.5835 | 0.0443 |
| cluster3674 | -2025.4169 | 0.0290 | 0.0376 | 1.2972 | 0.0264 |
| cluster3675 | -660.30852 | 0.0001 | 0.0000 | 0.0032 | 0.0001 |
| cluster3676 | -3522.196037 | 0.0281 | 0.0392 | 1.3950 | 0.0651 |
| cluster3677 | -811.299445 | 0.0415 | 0.0053 | 0.1271 | 0.0169 |
| cluster3678 | -3305.276363 | 0.0212 | 0.0212 | 0.9990 | 0.0409 |
| cluster3679 | -944.669922 | 0.0089 | 0.0025 | 0.2843 | 0.0081 |
| cluster3680 | -2314.433772 | 0.0088 | 0.0127 | 1.4468 | 0.0097 |
| cluster3681 | -2959.188763 | 0.0211 | 0.0352 | 1.6705 | 0.0186 |
| cluster3682 | -4359.365849 | 0.0167 | 0.0338 | 2.0176 | 0.0456 |
| cluster3683 | -2805.270009 | 0.0907 | 0.0446 | 0.4919 | 0.1083 |
| cluster3684 | -2987.450213 | 0.0141 | 0.0162 | 1.1539 | 0.0449 |
| cluster3685 | -3674.610951 | 0.0866 | 0.1132 | 1.3072 | 0.0818 |
| cluster3686 | -1627.697236 | 0.0761 | 0.1195 | 1.5692 | 0.0206 |
| cluster3687 | -2876.842964 | 0.1078 | 0.1380 | 1.2802 | 0.0681 |

| | | | | | |
|---|---|---|---|---|---|
| cluster3688 | -2051.092454 | 0.0060 | 0.0059 | 0.9769 | 0.0094 |
| cluster3689 | -2640.92489 | 0.0256 | 0.0162 | 0.6341 | 0.0485 |
| cluster3690 | -1453.769756 | 0.0118 | 0.0065 | 0.5496 | 0.0293 |
| cluster3691 | -670.692271 | 0.0042 | 0.0044 | 1.0578 | 0.0043 |
| cluster3692 | -1434.974638 | 0.0078 | 0.0133 | 1.7154 | 0.0126 |
| cluster3693 | -2464.093175 | 0.0129 | 0.0231 | 1.7905 | 0.0071 |
| cluster3694 | -2090.597284 | 0.0136 | 0.0013 | 0.0963 | 0.0366 |
| cluster3695 | -1735.860115 | 0.0148 | 0.0173 | 1.1689 | 0.0297 |
| cluster3696 | -813.366248 | 0.0223 | 0.0232 | 1.0400 | 0.0010 |
| cluster3697 | -3582.621084 | 0.0494 | 0.0157 | 0.3184 | 0.0846 |
| cluster3698 | -1099.572393 | 0.0001 | 0.0000 | 0.2241 | 0.0001 |
| cluster3699 | -4441.482378 | 0.0539 | 0.0882 | 1.6359 | 0.0443 |
| cluster3700 | -2818.737632 | 0.0254 | 0.0167 | 0.6587 | 0.0349 |
| cluster3701 | -1128.320963 | 0.0001 | 0.0000 | 0.4103 | 0.0015 |
| cluster3702 | -3863.22081 | 0.0142 | 0.0598 | 4.2014 | 0.0303 |
| cluster3703 | -1512.029157 | 0.0064 | 0.0013 | 0.2028 | 0.0209 |
| cluster3704 | -2229.294532 | 0.0024 | 0.0015 | 0.6469 | 0.0063 |
| cluster3705 | -1943.745294 | 0.0060 | 0.0038 | 0.6337 | 0.0181 |
| cluster3706 | -2093.471294 | 0.0121 | 0.0091 | 0.7557 | 0.0246 |
| cluster3707 | -1921.302231 | 0.0488 | 0.0825 | 1.6919 | 0.0383 |
| cluster3708 | -2635.068053 | 0.0278 | 0.0138 | 0.4972 | 0.0554 |
| cluster3709 | -1579.996799 | 0.0507 | 0.0098 | 0.1925 | 0.0851 |
| cluster3710 | -1922.384666 | 0.0061 | 0.0118 | 1.9364 | 0.0187 |
| cluster3711 | -1436.236433 | 0.0001 | 0.0000 | 0.1953 | 0.0357 |
| cluster3712 | -3401.875896 | 0.0348 | 0.0240 | 0.6910 | 0.0229 |
| cluster3713 | -4023.388075 | 0.0257 | 0.0554 | 2.1558 | 0.0713 |
| cluster3714 | -1775.470696 | 0.0354 | 0.0117 | 0.3302 | 0.0518 |
| cluster3715 | -1284.231328 | 0.0039 | 0.0489 | 12.6386 | 0.0212 |
| cluster3716 | -3657.942341 | 0.0410 | 0.0674 | 1.6435 | 0.0490 |
| cluster3717 | -1403.828242 | 0.0084 | 0.0007 | 0.0883 | 0.0153 |
| cluster3718 | -2988.762883 | 0.0132 | 0.0249 | 1.8922 | 0.0277 |
| cluster3719 | -3294.078339 | 0.0319 | 0.0221 | 0.6929 | 0.0505 |
| cluster3720 | -2757.913685 | 0.0161 | 0.0100 | 0.6227 | 0.0302 |
| cluster3721 | -1097.492957 | 0.0001 | 0.0000 | 0.1096 | 0.0068 |
| cluster3722 | -1224.600599 | 0.0276 | 0.1004 | 3.6341 | 0.0450 |
| cluster3723 | -989.942659 | 0.0001 | 0.0000 | 0.4878 | 0.0001 |
| cluster3724 | -3081.555003 | 0.0219 | 0.0333 | 1.5223 | 0.0400 |
| cluster3725 | -1837.76965 | 0.0196 | 0.0227 | 1.1586 | 0.0068 |
| cluster3726 | -1936.486915 | 0.0411 | 0.0191 | 0.4646 | 0.0506 |
| cluster3727 | -2467.635166 | 0.0376 | 0.0578 | 1.5361 | 0.0650 |
| cluster3728 | -2156.542835 | 0.0133 | 0.0052 | 0.3867 | 0.0231 |
| cluster3729 | -2946.308056 | 0.0194 | 0.0177 | 0.9126 | 0.0112 |
| cluster3730 | -1781.203913 | 0.0521 | 0.0107 | 0.2055 | 0.0954 |
| cluster3731 | -2864.733899 | 0.0056 | 0.0028 | 0.4934 | 0.0106 |

| | | | | | |
|---|---|---|---|---|---|
| cluster3732 | -2822.68689 | 0.0185 | 0.0139 | 0.7500 | 0.0243 |
| cluster3733 | -3072.518017 | 0.0659 | 0.0217 | 0.3289 | 0.0843 |
| cluster3734 | -3043.162709 | 0.0364 | 0.0329 | 0.9032 | 0.0463 |
| cluster3735 | -3249.310982 | 0.0573 | 0.0173 | 0.3018 | 0.0903 |
| cluster3736 | -3538.191601 | 0.0504 | 0.0315 | 0.6259 | 0.0447 |
| cluster3737 | -1209.19177 | 0.0070 | 0.0018 | 0.2642 | 0.0019 |
| cluster3738 | -3261.152401 | 0.0068 | 0.0031 | 0.4542 | 0.0088 |
| cluster3739 | -1156.253518 | 0.0001 | 0.0000 | 0.1245 | 0.0001 |
| cluster3740 | -3446.939816 | 0.0169 | 0.0264 | 1.5642 | 0.0364 |
| cluster3741 | -1309.637958 | 0.0001 | 0.0000 | 0.4154 | 0.0206 |
| cluster3742 | -1613.426453 | 0.0267 | 0.0381 | 1.4253 | 0.0736 |
| cluster3743 | -2980.34673 | 0.0254 | 0.0309 | 1.2172 | 0.0229 |
| cluster3744 | -1356.105459 | 0.1006 | 0.0618 | 0.6140 | 0.1088 |
| cluster3745 | -2584.757972 | 0.0187 | 0.0325 | 1.7349 | 0.0293 |
| cluster3746 | -3786.223326 | 0.0380 | 0.0439 | 1.1547 | 0.0382 |
| cluster3747 | -1121.259569 | 0.0248 | 0.0252 | 1.0169 | 0.0286 |
| cluster3748 | -2165.782978 | 0.0080 | 0.0023 | 0.2889 | 0.0169 |
| cluster3749 | -3067.573536 | 0.0079 | 0.0144 | 1.8306 | 0.0234 |
| cluster3750 | -1355.056636 | 0.0182 | 0.0395 | 2.1697 | 0.0290 |
| cluster3751 | -3009.941964 | 0.0011 | 0.0010 | 0.9121 | 0.0057 |
| cluster3752 | -2663.352477 | 0.0064 | 0.0076 | 1.1760 | 0.0086 |
| cluster3753 | -1022.847579 | 0.0001 | 0.0000 | 0.0581 | 0.0220 |
| cluster3754 | -881.977235 | 0.0001 | 0.0000 | 0.1371 | 0.0001 |
| cluster3755 | -2499.738586 | 0.0175 | 0.0130 | 0.7435 | 0.0147 |
| cluster3756 | -1688.666529 | 0.0216 | 0.0190 | 0.8815 | 0.0413 |
| cluster3757 | -2090.930437 | 0.0586 | 0.0270 | 0.4612 | 0.0417 |
| cluster3758 | -1510.284419 | 0.0558 | 0.0167 | 0.2999 | 0.0206 |
| cluster3759 | -3063.809848 | 0.0399 | 0.0182 | 0.4550 | 0.0842 |
| cluster3760 | -2600.110934 | 0.0146 | 0.0109 | 0.7454 | 0.0293 |
| cluster3761 | -767.990045 | 0.0041 | 0.0109 | 2.6807 | 0.0045 |
| cluster3762 | -1965.256036 | 0.0001 | 0.0000 | 0.1632 | 0.0199 |
| cluster3763 | -2004.236843 | 0.0322 | 0.0108 | 0.3354 | 0.0677 |
| cluster3764 | -1630.596539 | 0.0272 | 0.0387 | 1.4233 | 0.0305 |
| cluster3765 | -2670.307621 | 0.0122 | 0.0036 | 0.2975 | 0.0218 |
| cluster3766 | -1796.560284 | 0.0138 | 0.0081 | 0.5901 | 0.0091 |
| cluster3767 | -860.913521 | 0.0075 | 0.0194 | 2.5930 | 0.0330 |
| cluster3768 | -2213.411523 | 0.1807 | 0.1626 | 0.8998 | 0.0362 |
| cluster3769 | -1763.747465 | 0.0001 | 0.0000 | 0.2892 | 0.0091 |
| cluster3770 | -1774.995322 | 0.0453 | 0.0347 | 0.7664 | 0.0563 |
| cluster3771 | -2138.873223 | 0.0586 | 0.0232 | 0.3949 | 0.0745 |
| cluster3772 | -725.227827 | 0.0001 | 0.0000 | 0.3230 | 0.0009 |
| cluster3773 | -1117.62438 | 0.1138 | 0.0060 | 0.0524 | 0.2189 |
| cluster3774 | -1406.83076 | 0.0001 | 0.0000 | 0.2631 | 0.0185 |
| cluster3775 | -904.528086 | 0.0309 | 0.0266 | 0.8611 | 0.0071 |

| | | | | | |
|---|---|---|---|---|---|
| cluster3776 | -2518.639796 | 0.0743 | 0.0205 | 0.2766 | 0.0801 |
| cluster3777 | -3061.968879 | 0.0271 | 0.0457 | 1.6874 | 0.0784 |
| cluster3778 | -980.349947 | 0.0001 | 0.0000 | 0.2903 | 0.0016 |
| cluster3779 | -2995.129688 | 0.0235 | 0.0316 | 1.3414 | 0.0212 |
| cluster3780 | -1683.673565 | 0.0052 | 0.0117 | 2.2631 | 0.0166 |
| cluster3781 | -2686.131305 | 0.0482 | 0.0221 | 0.4582 | 0.0617 |
| cluster3782 | -2290.202181 | 0.0501 | 0.0362 | 0.7224 | 0.0408 |
| cluster3783 | -2107.051133 | 0.0023 | 0.0224 | 9.7243 | 0.0092 |
| cluster3784 | -2951.138685 | 0.0667 | 0.0329 | 0.4941 | 0.0435 |
| cluster3785 | -2688.066923 | 0.0268 | 0.0151 | 0.5646 | 0.0139 |
| cluster3786 | -2265.964594 | 0.0039 | 0.0024 | 0.6233 | 0.0112 |
| cluster3787 | -1522.310385 | 0.0230 | 0.0206 | 0.8932 | 0.0291 |
| cluster3788 | -1160.401549 | 0.0001 | 0.0000 | 0.0376 | 0.0001 |
| cluster3789 | -3037.801617 | 0.0017 | 0.0004 | 0.2129 | 0.0148 |
| cluster3790 | -2173.664285 | 0.0057 | 0.0117 | 2.0368 | 0.0228 |
| cluster3791 | -419.938863 | 0.0247 | 0.0061 | 0.2467 | 0.0114 |
| cluster3792 | -1863.345042 | 0.0045 | 0.0013 | 0.2804 | 0.0063 |
| cluster3793 | -1852.152002 | 0.0138 | 0.0094 | 0.6775 | 0.0328 |
| cluster3794 | -2189.509144 | 0.0179 | 0.0158 | 0.8818 | 0.0153 |
| cluster3795 | -1701.057853 | 0.0133 | 0.0115 | 0.8662 | 0.0231 |
| cluster3796 | -1249.044506 | 0.0576 | 0.0732 | 1.2708 | 0.1594 |
| cluster3797 | -3911.049659 | 0.0365 | 0.1006 | 2.7566 | 0.0655 |
| cluster3798 | -3447.124473 | 0.0237 | 0.0204 | 0.8605 | 0.0243 |
| cluster3799 | -2341.295192 | 0.0393 | 0.0282 | 0.7187 | 0.0513 |
| cluster3800 | -431.301283 | 0.0001 | 0.0000 | 0.0224 | 0.0117 |
| cluster3801 | -2686.728967 | 0.3852 | 0.0871 | 0.2262 | 0.1660 |
| cluster3802 | -2649.120306 | 0.0239 | 0.0249 | 1.0416 | 0.0398 |
| cluster3803 | -2239.959766 | 0.0344 | 0.0341 | 0.9927 | 0.0359 |
| cluster3804 | -2634.772025 | 0.0223 | 0.0163 | 0.7311 | 0.0355 |
| cluster3805 | -2655.680239 | 0.0195 | 0.0235 | 1.2093 | 0.0256 |
| cluster3806 | -3566.950748 | 0.0411 | 0.0510 | 1.2417 | 0.0375 |
| cluster3807 | -1441.826747 | 0.0051 | 0.0009 | 0.1871 | 0.0081 |
| cluster3808 | -1825.913247 | 0.0361 | 0.0483 | 1.3383 | 0.0303 |
| cluster3809 | -2361.188557 | 0.0116 | 0.0087 | 0.7556 | 0.0085 |
| cluster3810 | -2464.593625 | 0.0094 | 0.0075 | 0.7992 | 0.0097 |
| cluster3811 | -761.00903 | 0.0001 | 0.0000 | 0.1224 | 0.0283 |
| cluster3812 | -3000.780832 | 0.0275 | 0.0167 | 0.6070 | 0.0315 |
| cluster3813 | -3413.663408 | 0.0320 | 0.0803 | 2.5111 | 0.0365 |
| cluster3814 | -1607.186149 | 0.0396 | 0.0036 | 0.0918 | 0.1080 |
| cluster3815 | -1707.54743 | 0.0117 | 0.0250 | 2.1409 | 0.0240 |
| cluster3816 | -2526.665949 | 0.0243 | 0.0425 | 1.7455 | 0.0280 |
| cluster3817 | -990.361444 | 0.0285 | 0.0027 | 0.0962 | 0.0035 |
| cluster3818 | -1978.998971 | 0.0545 | 0.0429 | 0.7862 | 0.0664 |
| cluster3819 | -3347.055898 | 0.0242 | 0.0248 | 1.0264 | 0.0291 |

| | | | | | |
|---|---|---|---|---|---|
| cluster3820 | -2222.195346 | 0.0130 | 0.0129 | 0.9932 | 0.0384 |
| cluster3821 | -2242.483661 | 0.0093 | 0.0046 | 0.4956 | 0.0213 |
| cluster3822 | -3396.217925 | 0.0151 | 0.0287 | 1.9011 | 0.0357 |
| cluster3823 | -3406.704698 | 0.0115 | 0.0175 | 1.5243 | 0.0247 |
| cluster3824 | -2350.317361 | 0.0083 | 0.0210 | 2.5247 | 0.0330 |
| cluster3825 | -1386.643951 | 0.0001 | 0.0001 | 0.7211 | 0.0085 |
| cluster3826 | -3154.848789 | 0.0634 | 0.0899 | 1.4181 | 0.0812 |
| cluster3827 | -2403.090292 | 0.0199 | 0.0124 | 0.6256 | 0.0320 |
| cluster3828 | -2365.224407 | 0.0503 | 0.0339 | 0.6741 | 0.0468 |
| cluster3829 | -2256.814649 | 0.0316 | 0.0279 | 0.8834 | 0.0213 |
| cluster3830 | -1628.909611 | 0.0114 | 0.0034 | 0.3036 | 0.0226 |
| cluster3831 | -2348.387956 | 0.0355 | 0.0412 | 1.1604 | 0.0293 |
| cluster3832 | -1766.748567 | 0.0001 | 0.0001 | 0.6231 | 0.0055 |
| cluster3833 | -2130.20773 | 0.0274 | 0.0357 | 1.3033 | 0.0209 |
| cluster3834 | -2054.85602 | 0.0011 | 0.0019 | 1.6636 | 0.0164 |
| cluster3835 | -2117.145736 | 0.0215 | 0.0179 | 0.8329 | 0.0400 |
| cluster3836 | -1894.912275 | 0.0209 | 0.0172 | 0.8261 | 0.0161 |
| cluster3837 | -2639.469093 | 0.0354 | 0.0390 | 1.1011 | 0.0433 |
| cluster3838 | -2305.843591 | 0.0098 | 0.0102 | 1.0365 | 0.0501 |
| cluster3839 | -1498.687486 | 0.0086 | 0.0148 | 1.7245 | 0.0250 |
| cluster3840 | -2024.641652 | 0.0032 | 0.0022 | 0.6877 | 0.0123 |
| cluster3841 | -2293.169013 | 0.0764 | 0.0907 | 1.1861 | 0.0751 |
| cluster3842 | -1606.299856 | 0.0001 | 0.0000 | 0.0813 | 0.0045 |
| cluster3843 | -1440.331851 | 0.0051 | 0.0036 | 0.7169 | 0.0143 |
| cluster3844 | -1076.38588 | 0.0183 | 0.0058 | 0.3161 | 0.0805 |
| cluster3845 | -604.916359 | 0.0001 | 0.0000 | 0.0000 | 0.0001 |
| cluster3846 | -1268.709937 | 0.0021 | 0.0010 | 0.4690 | 0.0329 |
| cluster3847 | -1155.876249 | 0.0001 | 0.0001 | 1.2992 | 0.0014 |
| cluster3848 | -2516.993037 | 0.0228 | 0.0438 | 1.9236 | 0.0353 |
| cluster3849 | -2664.114187 | 0.0211 | 0.0329 | 1.5642 | 0.0102 |
| cluster3850 | -1676.140969 | 0.0498 | 0.0135 | 0.2708 | 0.0178 |
| cluster3851 | -1618.182098 | 0.0093 | 0.0079 | 0.8415 | 0.0099 |
| cluster3852 | -1856.83096 | 0.0074 | 0.0040 | 0.5436 | 0.0065 |
| cluster3853 | -1101.730577 | 0.0042 | 0.0076 | 1.8161 | 0.0051 |
| cluster3854 | -1356.231582 | 0.0523 | 0.0337 | 0.6438 | 0.1277 |
| cluster3855 | -2930.765931 | 0.0084 | 0.0272 | 3.2287 | 0.0112 |
| cluster3856 | -1693.430459 | 0.0074 | 0.0177 | 2.4107 | 0.0242 |
| cluster3857 | -3037.030709 | 0.0109 | 0.0124 | 1.1396 | 0.0190 |
| cluster3858 | -2416.372771 | 0.0381 | 0.0138 | 0.3637 | 0.0534 |
| cluster3859 | -1974.771658 | 0.0037 | 0.0031 | 0.8489 | 0.0170 |
| cluster3860 | -2618.401705 | 0.0310 | 0.0126 | 0.4046 | 0.1160 |
| cluster3861 | -1300.22313 | 0.0085 | 0.0066 | 0.7767 | 0.0196 |
| cluster3862 | -922.33901 | 0.0001 | 0.0000 | 0.4903 | 0.0001 |
| cluster3863 | -1483.126909 | 0.0001 | 0.0000 | 0.4468 | 0.0013 |

| | | | | | |
|---|---|---|---|---|---|
| cluster3864 | -2219.389089 | 0.0020 | 0.0030 | 1.4560 | 0.0052 |
| cluster3865 | -3003.458786 | 0.0333 | 0.0597 | 1.7903 | 0.0505 |
| cluster3866 | -1633.311468 | 0.0150 | 0.0191 | 1.2753 | 0.0199 |
| cluster3867 | -608.471137 | 0.0001 | 0.0000 | 0.0246 | 0.0001 |
| cluster3868 | -1388.885176 | 0.0001 | 0.0000 | 0.4810 | 0.0371 |
| cluster3869 | -2974.00478 | 0.0181 | 0.0849 | 4.6812 | 0.0373 |
| cluster3870 | -2461.634354 | 0.0279 | 0.0287 | 1.0261 | 0.0458 |
| cluster3871 | -2148.485749 | 0.0174 | 0.0178 | 1.0196 | 0.0045 |
| cluster3872 | -2648.903963 | 0.0182 | 0.0470 | 2.5886 | 0.0106 |
| cluster3873 | -1788.160957 | 0.0285 | 0.0087 | 0.3034 | 0.0289 |
| cluster3874 | -2097.361314 | 0.0001 | 0.0000 | 0.2137 | 0.0182 |
| cluster3875 | -770.173788 | 0.0001 | 0.0000 | 0.4199 | 0.0033 |
| cluster3876 | -744.64899 | 0.0001 | 0.0000 | 0.1906 | 0.0142 |
| cluster3877 | -1433.557693 | 0.0026 | 0.0003 | 0.1195 | 0.0192 |
| cluster3879 | -1881.019652 | 0.0116 | 0.0080 | 0.6858 | 0.0196 |
| cluster3880 | -1357.438158 | 0.0276 | 0.0182 | 0.6606 | 0.0183 |
| cluster3881 | -3017.712011 | 0.0428 | 0.0729 | 1.7027 | 0.0503 |
| cluster3882 | -2686.522988 | 0.0110 | 0.0265 | 2.3987 | 0.0316 |
| cluster3883 | -421.863316 | 0.0001 | 0.0013 | 13.0388 | 0.0395 |
| cluster3884 | -1563.376049 | 0.0002 | 0.0146 | 70.5690 | 0.0004 |
| cluster3885 | -2857.695423 | 0.0100 | 0.0130 | 1.3033 | 0.0156 |
| cluster3886 | -1392.620239 | 0.0009 | 0.0112 | 12.5239 | 0.0063 |
| cluster3887 | -1628.245691 | 0.0124 | 0.0327 | 2.6390 | 0.0082 |
| cluster3888 | -1182.494316 | 0.0074 | 0.0193 | 2.6143 | 0.0122 |
| cluster3889 | -1004.560618 | 0.0193 | 0.0288 | 1.4963 | 0.0259 |
| cluster3890 | -1871.092045 | 0.0104 | 0.0124 | 1.1940 | 0.0212 |
| cluster3891 | -2184.694914 | 0.0160 | 0.0093 | 0.5788 | 0.0132 |
| cluster3892 | -1373.349896 | 0.0313 | 0.0088 | 0.2801 | 0.0122 |
| cluster3893 | -1457.112277 | 0.0074 | 0.0016 | 0.2109 | 0.0333 |
| cluster3894 | -964.931863 | 0.0001 | 0.0001 | 0.5813 | 0.0114 |
| cluster3895 | -1978.118782 | 0.0001 | 0.0001 | 0.9639 | 0.0143 |
| cluster3896 | -818.847875 | 0.0001 | 0.0001 | 1.1488 | 0.0055 |
| cluster3897 | -1448.984104 | 0.0001 | 0.0001 | 0.7018 | 0.0117 |
| cluster3898 | -2390.40173 | 0.0221 | 0.0044 | 0.2009 | 0.0253 |
| cluster3899 | -3001.702079 | 0.0168 | 0.0463 | 2.7549 | 0.0326 |
| cluster3900 | -2359.922823 | 0.0124 | 0.0763 | 6.1484 | 0.0953 |
| cluster3901 | -1543.160296 | 0.0001 | 0.0001 | 1.2862 | 0.0029 |
| cluster3902 | -2496.296944 | 0.0099 | 0.0214 | 2.1659 | 0.0122 |
| cluster3903 | -1622.95685 | 0.0067 | 0.0096 | 1.4363 | 0.0135 |
| cluster3904 | -904.817825 | 0.0109 | 0.0069 | 0.6317 | 0.0039 |
| cluster3905 | -1591.587148 | 0.0591 | 0.0618 | 1.0450 | 0.1688 |
| cluster3906 | -1451.589178 | 0.0093 | 0.0091 | 0.9779 | 0.0287 |
| cluster3907 | -1944.694358 | 0.0322 | 0.0320 | 0.9935 | 0.0449 |
| cluster3908 | -1847.959893 | 0.0108 | 0.0102 | 0.9429 | 0.0155 |

| | | | | | |
|---|---|---|---|---|---|
| cluster3909 | -2461.501404 | 0.0089 | 0.0135 | 1.5090 | 0.0332 |
| cluster3910 | -1193.281251 | 0.0293 | 0.0313 | 1.0689 | 0.0404 |
| cluster3911 | -1748.862312 | 0.0037 | 0.0038 | 1.0113 | 0.0109 |
| cluster3912 | -951.766263 | 0.0001 | 0.0000 | 0.2971 | 0.0001 |
| cluster3913 | -2544.922642 | 0.0354 | 0.0348 | 0.9845 | 0.0403 |
| cluster3914 | -2239.068989 | 0.0536 | 0.0808 | 1.5075 | 0.0612 |
| cluster3915 | -2227.20086 | 0.0411 | 0.0467 | 1.1351 | 0.0323 |
| cluster3916 | -902.465738 | 0.0631 | 0.0490 | 0.7763 | 0.0873 |
| cluster3917 | -2582.878164 | 0.0158 | 0.0128 | 0.8091 | 0.0109 |
| cluster3918 | -2758.924417 | 0.0153 | 0.0446 | 2.9220 | 0.0193 |
| cluster3919 | -569.923191 | 0.0001 | 0.0000 | 0.2700 | 0.0022 |
| cluster3920 | -657.65968 | 0.0071 | 0.1081 | 15.2973 | 0.0158 |
| cluster3921 | -2190.57862 | 0.0299 | 0.0220 | 0.7371 | 0.0239 |
| cluster3922 | -1240.206842 | 0.0577 | 0.0365 | 0.6321 | 0.0483 |
| cluster3923 | -1149.926241 | 0.0011 | 0.0022 | 1.9936 | 0.0002 |
| cluster3924 | -1696.289154 | 0.0189 | 0.0162 | 0.8563 | 0.0201 |
| cluster3925 | -1091.662377 | 0.0249 | 0.0423 | 1.7003 | 0.0648 |
| cluster3926 | -2435.266919 | 0.0190 | 0.1122 | 5.8891 | 0.0607 |
| cluster3927 | -1186.206404 | 0.0025 | 0.0026 | 1.0383 | 0.0147 |
| cluster3928 | -1158.569109 | 0.0111 | 0.0057 | 0.5119 | 0.0361 |
| cluster3929 | -1607.885335 | 0.0042 | 0.0064 | 1.5476 | 0.0071 |
| cluster3930 | -2017.826272 | 0.0153 | 0.0087 | 0.5693 | 0.0157 |
| cluster3931 | -1823.594875 | 0.0145 | 0.0023 | 0.1599 | 0.1682 |
| cluster3932 | -1117.870724 | 0.0088 | 0.0006 | 0.0727 | 0.0116 |
| cluster3933 | -1157.654377 | 0.0465 | 0.0112 | 0.2413 | 0.0985 |
| cluster3934 | -717.107608 | 0.0001 | 0.0000 | 0.3399 | 0.0001 |
| cluster3935 | -3102.72817 | 0.0236 | 0.0481 | 2.0336 | 0.0769 |
| cluster3936 | -2131.839537 | 0.0069 | 0.0162 | 2.3449 | 0.0235 |
| cluster3937 | -1937.297274 | 0.0090 | 0.0054 | 0.6046 | 0.0250 |
| cluster3938 | -1781.80766 | 0.0225 | 0.0151 | 0.6702 | 0.0357 |
| cluster3939 | -2056.003695 | 0.0157 | 0.0177 | 1.1292 | 0.0155 |
| cluster3940 | -1338.53153 | 0.0157 | 0.0046 | 0.2910 | 0.0187 |
| cluster3941 | -968.785752 | 0.0001 | 0.0000 | 0.1218 | 0.0338 |
| cluster3942 | -2009.205125 | 0.0228 | 0.0129 | 0.5680 | 0.0342 |
| cluster3943 | -2268.191831 | 0.0125 | 0.0242 | 1.9370 | 0.0182 |
| cluster3944 | -1592.713101 | 0.0326 | 0.0493 | 1.5122 | 0.0521 |
| cluster3945 | -1676.626733 | 0.0347 | 0.0350 | 1.0105 | 0.0400 |
| cluster3946 | -1805.256504 | 0.0159 | 0.0135 | 0.8490 | 0.0301 |
| cluster3947 | -2283.791498 | 0.0466 | 0.0555 | 1.1907 | 0.0223 |
| cluster3948 | -1012.4142 | 0.0046 | 0.0028 | 0.6071 | 0.0406 |
| cluster3949 | -876.250494 | 0.0001 | 0.0001 | 0.5610 | 0.0162 |
| cluster3950 | -1737.378505 | 0.0005 | 0.0142 | 28.9846 | 0.0259 |
| cluster3951 | -2833.810244 | 0.0318 | 0.0349 | 1.0985 | 0.0430 |
| cluster3952 | -1474.784299 | 0.0019 | 0.0027 | 1.4159 | 0.0035 |

| | | | | | |
|---|---|---|---|---|---|
| cluster3953 | -1361.955053 | 0.0029 | 0.0018 | 0.6445 | 0.0192 |
| cluster3954 | -1803.47987 | 0.0119 | 0.0111 | 0.9297 | 0.0364 |
| cluster3955 | -1192.220381 | 0.0253 | 0.0264 | 1.0428 | 0.0251 |
| cluster3956 | -1995.550232 | 0.0114 | 0.0165 | 1.4416 | 0.0157 |
| cluster3957 | -1959.017862 | 0.0085 | 0.0110 | 1.3036 | 0.0143 |
| cluster3958 | -1104.863039 | 0.0231 | 0.0115 | 0.4961 | 0.0229 |
| cluster3959 | -2684.99595 | 0.0239 | 0.0139 | 0.5842 | 0.0401 |
| cluster3960 | -2422.912029 | 0.0850 | 0.0789 | 0.9276 | 0.0453 |
| cluster3961 | -1592.985436 | 0.0675 | 0.0557 | 0.8258 | 0.0281 |
| cluster3962 | -1532.956009 | 0.0317 | 0.0367 | 1.1597 | 0.0459 |
| cluster3963 | -2074.470911 | 0.0172 | 0.0241 | 1.4033 | 0.0520 |
| cluster3964 | -2992.552775 | 0.0054 | 0.1671 | 30.8113 | 0.0935 |
| cluster3965 | -1516.841574 | 0.0329 | 0.0429 | 1.3024 | 0.0355 |
| cluster3966 | -1431.259411 | 0.0097 | 0.0062 | 0.6343 | 0.0297 |
| cluster3967 | -1753.721993 | 0.0723 | 0.0195 | 0.2692 | 0.0938 |
| cluster3968 | -1167.507437 | 0.0133 | 0.0183 | 1.3747 | 0.0193 |
| cluster3969 | -899.568179 | 0.0001 | 0.0000 | 0.2948 | 0.0001 |
| cluster3970 | -2484.154536 | 0.0375 | 0.0401 | 1.0697 | 0.0490 |
| cluster3971 | -1686.351508 | 0.0137 | 0.0162 | 1.1796 | 0.0502 |
| cluster3972 | -988.603724 | 0.0276 | 0.0265 | 0.9572 | 0.0096 |
| cluster3973 | -1419.068546 | 0.0280 | 0.0035 | 0.1253 | 0.0042 |
| cluster3974 | -1831.567939 | 0.0091 | 0.0091 | 0.9951 | 0.0211 |
| cluster3975 | -1266.911531 | 0.0681 | 0.0628 | 0.9223 | 0.0162 |
| cluster3976 | -1578.908982 | 0.0056 | 0.0074 | 1.3218 | 0.0205 |
| cluster3977 | -2230.143198 | 0.0934 | 0.0809 | 0.8668 | 0.0991 |
| cluster3978 | -887.285188 | 0.0001 | 0.0000 | 0.2874 | 0.0069 |
| cluster3979 | -1620.014713 | 0.0070 | 0.0023 | 0.3327 | 0.0014 |
| cluster3980 | -922.248674 | 0.0001 | 0.0000 | 0.0281 | 0.0001 |
| cluster3981 | -1979.457902 | 0.0100 | 0.0078 | 0.7823 | 0.0274 |
| cluster3982 | -1829.900743 | 0.0470 | 0.0410 | 0.8709 | 0.0503 |
| cluster3983 | -2676.129745 | 0.0236 | 0.0468 | 1.9854 | 0.0405 |
| cluster3984 | -1379.596962 | 0.0160 | 0.0437 | 2.7313 | 0.0706 |
| cluster3985 | -979.187872 | 0.0159 | 0.0039 | 0.2433 | 0.0110 |
| cluster3986 | -1732.923985 | 0.0086 | 0.0141 | 1.6314 | 0.0232 |
| cluster3987 | -2087.223978 | 0.0115 | 0.0031 | 0.2716 | 0.0124 |
| cluster3988 | -1243.635505 | 0.0001 | 0.0001 | 0.8562 | 0.0124 |
| cluster3989 | -1835.009475 | 0.0614 | 0.0178 | 0.2903 | 0.0531 |
| cluster3990 | -2207.669555 | 0.0167 | 0.0292 | 1.7477 | 0.0290 |
| cluster3991 | -1522.092251 | 0.0202 | 0.0302 | 1.4943 | 0.0209 |
| cluster3992 | -1887.404934 | 0.0203 | 0.0073 | 0.3603 | 0.0466 |
| cluster3993 | -1674.954846 | 0.0382 | 0.0396 | 1.0382 | 0.0327 |
| cluster3994 | -2394.952787 | 0.0497 | 0.0385 | 0.7746 | 0.0617 |
| cluster3995 | -1618.154416 | 0.0199 | 0.0162 | 0.8105 | 0.0436 |
| cluster3996 | -1708.748614 | 0.0043 | 0.0076 | 1.7769 | 0.0071 |

| | | | | | |
|---|---|---|---|---|---|
| cluster3997 | -2518.917693 | 0.0512 | 0.0532 | 1.0394 | 0.0549 |
| cluster3998 | -1416.521039 | 0.0424 | 0.0423 | 0.9994 | 0.0587 |
| cluster3999 | -2524.788957 | 0.0265 | 0.0407 | 1.5334 | 0.0345 |
| cluster4000 | -2289.398117 | 0.0315 | 0.0265 | 0.8404 | 0.0529 |
| cluster4001 | -1739.568055 | 0.0208 | 0.0167 | 0.8002 | 0.0185 |
| cluster4002 | -1801.841941 | 0.0050 | 0.0070 | 1.4057 | 0.0025 |
| cluster4003 | -1418.800317 | 0.0306 | 0.0311 | 1.0162 | 0.0963 |
| cluster4004 | -1442.052569 | 0.0145 | 0.0092 | 0.6297 | 0.0063 |
| cluster4005 | -2067.22439 | 0.0275 | 0.0754 | 2.7412 | 0.0547 |
| cluster4006 | -1088.963393 | 0.0210 | 0.0182 | 0.8691 | 0.0373 |
| cluster4007 | -1424.567732 | 0.0001 | 0.0001 | 0.8950 | 0.0035 |
| cluster4008 | -1355.67501 | 0.0097 | 0.0115 | 1.1853 | 0.0067 |
| cluster4009 | -1546.481916 | 0.0145 | 0.0255 | 1.7544 | 0.0114 |
| cluster4010 | -1627.725309 | 0.0585 | 0.0973 | 1.6654 | 0.0539 |
| cluster4011 | -1061.44901 | 0.0804 | 0.0133 | 0.1660 | 0.0665 |
| cluster4012 | -2442.798918 | 0.0053 | 0.0471 | 8.8694 | 0.0237 |
| cluster4013 | -843.621104 | 0.0001 | 0.0001 | 0.5842 | 0.0317 |
| cluster4014 | -1137.241725 | 0.0117 | 0.0086 | 0.7381 | 0.0043 |
| cluster4015 | -617.98354 | 0.0970 | 0.0035 | 0.0364 | 0.0159 |
| cluster4016 | -1815.670474 | 0.0167 | 0.0131 | 0.7851 | 0.0197 |
| cluster4017 | -1341.595099 | 0.0284 | 0.0334 | 1.1745 | 0.0494 |
| cluster4018 | -1325.636479 | 0.1202 | 0.0771 | 0.6415 | 0.1585 |
| cluster4019 | -1826.171259 | 0.0615 | 0.0635 | 1.0324 | 0.0695 |
| cluster4020 | -1004.604694 | 0.0169 | 0.0037 | 0.2195 | 0.0439 |
| cluster4021 | -1591.029391 | 0.0080 | 0.0037 | 0.4648 | 0.0325 |
| cluster4022 | -969.50095 | 0.0001 | 0.0000 | 0.2264 | 0.0108 |
| cluster4023 | -1843.216811 | 0.0955 | 0.0534 | 0.5599 | 0.0548 |
| cluster4024 | -2359.145949 | 0.0386 | 0.0591 | 1.5316 | 0.0255 |
| cluster4025 | -624.335272 | 0.0001 | 0.0000 | 0.4858 | 0.0007 |
| cluster4026 | -1088.994697 | 0.0026 | 0.0030 | 1.1389 | 0.0026 |
| cluster4027 | -980.13209 | 0.0042 | 0.0020 | 0.4705 | 0.0056 |
| cluster4028 | -1844.514942 | 0.0014 | 0.0093 | 6.6848 | 0.0093 |
| cluster4029 | -2250.78546 | 0.0536 | 0.0681 | 1.2721 | 0.0775 |
| cluster4030 | -1123.502069 | 0.0158 | 0.0097 | 0.6135 | 0.0470 |
| cluster4031 | -1096.766563 | 0.0115 | 0.0040 | 0.3428 | 0.0367 |
| cluster4032 | -1475.107328 | 0.0001 | 0.0000 | 0.4613 | 0.0016 |
| cluster4033 | -2213.72445 | 0.0442 | 0.0769 | 1.7395 | 0.0825 |
| cluster4034 | -493.672603 | 0.0001 | 0.0000 | 0.0177 | 0.0001 |
| cluster4035 | -2113.948325 | 0.0372 | 0.0223 | 0.5997 | 0.0757 |
| cluster4036 | -1077.296292 | 0.0073 | 0.0057 | 0.7851 | 0.0096 |
| cluster4037 | -1625.236236 | 0.0273 | 0.0290 | 1.0634 | 0.0156 |
| cluster4038 | -901.963409 | 0.0235 | 0.0610 | 2.5961 | 0.0121 |
| cluster4039 | -1508.285523 | 0.0557 | 0.0196 | 0.3519 | 0.0634 |
| cluster4040 | -1122.346005 | 0.1320 | 0.0277 | 0.2097 | 0.0097 |

| | | | | | |
|---|---|---|---|---|---|
| cluster4041 | -1474.29775 | 0.0242 | 0.0288 | 1.1880 | 0.0108 |
| cluster4042 | -899.663205 | 0.0361 | 0.0325 | 0.9022 | 0.0304 |
| cluster4043 | -1138.588515 | 0.0367 | 0.1111 | 3.0281 | 0.0452 |
| cluster4044 | -958.068597 | 0.0015 | 0.0024 | 1.6672 | 0.0014 |
| cluster4045 | -2022.111922 | 0.0380 | 0.0543 | 1.4279 | 0.0780 |
| cluster4046 | -889.479228 | 0.0001 | 0.0000 | 0.4310 | 0.0035 |
| cluster4047 | -2068.867764 | 0.0126 | 0.0437 | 3.4708 | 0.0201 |
| cluster4048 | -1468.45791 | 0.0001 | 0.0000 | 0.4018 | 0.0103 |
| cluster4049 | -573.688456 | 0.0013 | 0.0034 | 2.7236 | 0.0011 |
| cluster4050 | -1216.688339 | 0.0030 | 0.0022 | 0.7195 | 0.0107 |
| cluster4051 | -1404.153632 | 0.0050 | 0.0016 | 0.3263 | 0.0098 |
| cluster4052 | -1287.90854 | 0.0142 | 0.0068 | 0.4792 | 0.0218 |
| cluster4053 | -1643.423076 | 0.0052 | 0.0306 | 5.8299 | 0.0152 |
| cluster4054 | -1856.879012 | 0.0059 | 0.0192 | 3.2252 | 0.0268 |
| cluster4055 | -719.970774 | 0.0025 | 0.0108 | 4.3385 | 0.0022 |
| cluster4056 | -931.631787 | 0.0104 | 0.0093 | 0.8910 | 0.0519 |
| cluster4057 | -1562.938384 | 0.0164 | 0.0191 | 1.1677 | 0.0429 |
| cluster4058 | -1681.465415 | 0.0397 | 0.0314 | 0.7913 | 0.0430 |
| cluster4059 | -721.69248 | 0.0542 | 0.0533 | 0.9842 | 0.0128 |
| cluster4060 | -1875.574962 | 0.0218 | 0.0192 | 0.8797 | 0.0305 |
| cluster4061 | -1187.965356 | 0.0032 | 0.0082 | 2.5582 | 0.0071 |
| cluster4062 | -1202.287968 | 0.0001 | 0.0003 | 2.5168 | 0.0231 |
| cluster4063 | -1721.94084 | 0.0164 | 0.0448 | 2.7291 | 0.0333 |
| cluster4064 | -783.008783 | 0.0108 | 0.0116 | 1.0725 | 0.0188 |
| cluster4065 | -988.415542 | 0.0001 | 0.0000 | 0.4451 | 0.0102 |
| cluster4066 | -1616.651987 | 0.0551 | 0.0512 | 0.9292 | 0.0414 |
| cluster4067 | -418.465838 | 0.0271 | 0.0489 | 1.8095 | 0.0116 |
| cluster4068 | -1353.889416 | 0.0152 | 0.0410 | 2.6934 | 0.0204 |
| cluster4069 | -1064.48628 | 0.0908 | 0.0424 | 0.4668 | 0.0536 |
| cluster4070 | -693.476069 | 0.0135 | 0.0165 | 1.2291 | 0.0013 |
| cluster4071 | -826.20544 | 0.0001 | 0.0001 | 0.9423 | 0.0026 |
| cluster4072 | -987.934457 | 0.0342 | 0.0231 | 0.6759 | 0.0228 |
| cluster4073 | -1889.615845 | 0.0254 | 0.0373 | 1.4693 | 0.0333 |
| cluster4074 | -1256.779507 | 0.0448 | 0.0593 | 1.3236 | 0.0266 |
| cluster4075 | -708.460845 | 9.0000 | 0.0000 | 0.0000 | 0.4954 |
| cluster4076 | -1648.914549 | 0.0546 | 0.0254 | 0.4659 | 0.1093 |
| cluster4077 | -1679.925711 | 0.0375 | 0.0382 | 1.0179 | 0.0466 |
| cluster4078 | -1230.174601 | 0.0581 | 0.0199 | 0.3433 | 0.0146 |
| cluster4079 | -1207.249307 | 0.0001 | 0.0000 | 0.4732 | 0.0012 |
| cluster4080 | -736.760834 | 0.0001 | 0.0000 | 0.1497 | 0.0208 |
| cluster4081 | -1128.016885 | 0.0031 | 0.0026 | 0.8475 | 0.0249 |
| cluster4082 | -742.820363 | 0.0001 | 0.0000 | 0.0000 | 0.0107 |
| cluster4083 | -1702.016099 | 0.0206 | 0.0395 | 1.9195 | 0.0801 |
| cluster4084 | -715.616885 | 9.0000 | 0.0000 | 0.0000 | 0.0102 |

| | | | | | |
|---|---|---|---|---|---|
| cluster4085 | -859.031317 | 0.0130 | 0.0047 | 0.3606 | 0.0324 |
| cluster4086 | -1314.791851 | 0.0324 | 0.0310 | 0.9560 | 0.0290 |
| cluster4087 | -1205.93067 | 0.0150 | 0.0237 | 1.5836 | 0.0111 |
| cluster4088 | -1313.61374 | 0.0351 | 0.0125 | 0.3571 | 0.0663 |
| cluster4089 | -1510.750964 | 0.0035 | 0.0154 | 4.3866 | 0.0073 |
| cluster4090 | -755.196255 | 0.0201 | 0.0123 | 0.6091 | 0.0420 |
| cluster4091 | -951.289898 | 0.0001 | 0.0000 | 0.2539 | 0.0389 |
| cluster4092 | -1421.731668 | 0.0092 | 0.0220 | 2.4037 | 0.0391 |
| cluster4093 | -1006.493579 | 0.0107 | 0.0101 | 0.9466 | 0.0208 |
| cluster4094 | -375.462812 | 0.0001 | 0.0000 | 0.0284 | 0.0090 |
| cluster4095 | -1404.345128 | 0.0640 | 0.0524 | 0.8180 | 0.0408 |
| cluster4096 | -1212.049759 | 0.0110 | 0.0253 | 2.3101 | 0.0291 |
| cluster4097 | -705.32302 | 0.1246 | 0.0025 | 0.0199 | 0.0945 |
| cluster4098 | -1868.691484 | 0.0379 | 0.0144 | 0.3799 | 0.0318 |
| cluster4099 | -809.58465 | 0.0070 | 0.0094 | 1.3436 | 0.0011 |
| cluster4100 | -1045.629994 | 0.0001 | 0.0001 | 1.0523 | 0.0110 |
| cluster4101 | -2087.430033 | 0.1489 | 0.0694 | 0.4659 | 0.3196 |
| cluster4102 | -1856.694425 | 0.0575 | 0.1892 | 3.2927 | 0.0428 |
| cluster4103 | -1103.372758 | 0.0009 | 0.0002 | 0.1999 | 0.0617 |
| cluster4104 | -1018.043033 | 0.0082 | 0.0114 | 1.3888 | 0.0071 |
| cluster4105 | -735.522172 | 0.0060 | 0.0090 | 1.4988 | 0.0114 |
| cluster4106 | -1000.105492 | 0.0138 | 0.0197 | 1.4247 | 0.0054 |
| cluster4107 | -785.650977 | 0.0001 | 0.0002 | 1.6968 | 0.0033 |
| cluster4108 | -780.331348 | 0.0040 | 0.0189 | 4.6674 | 0.0277 |
| cluster4109 | -1651.867539 | 0.0717 | 0.0962 | 1.3415 | 0.0655 |
| cluster4110 | -1243.851369 | 0.0263 | 0.0255 | 0.9716 | 0.0880 |
| cluster4111 | -925.196321 | 0.0508 | 0.0422 | 0.8311 | 0.0461 |
| cluster4112 | -1113.287198 | 0.0129 | 0.0051 | 0.3986 | 0.0177 |
| cluster4113 | -773.195973 | 0.0001 | 0.0000 | 0.0702 | 0.0064 |
| cluster4114 | -1074.129137 | 0.0246 | 0.0180 | 0.7297 | 0.0516 |
| cluster4115 | -1154.94232 | 0.0029 | 0.0025 | 0.8886 | 0.0151 |
| cluster4116 | -636.643958 | 0.0445 | 0.0144 | 0.3247 | 0.0256 |
| cluster4117 | -660.549106 | 0.0091 | 0.0034 | 0.3748 | 0.0145 |
| cluster4118 | -1288.275254 | 0.0235 | 0.0532 | 2.2592 | 0.0335 |
| cluster4119 | -924.970505 | 0.0067 | 0.0073 | 1.0882 | 0.0245 |
| cluster4120 | -968.240795 | 0.0023 | 0.0020 | 0.8668 | 0.0051 |
| cluster4121 | -1216.982213 | 0.0020 | 0.0179 | 8.9169 | 0.0057 |
| cluster4122 | -527.42162 | 0.0746 | 0.0036 | 0.0480 | 0.0268 |
| cluster4123 | -468.668086 | 0.0073 | 0.0035 | 0.4786 | 0.0071 |
| cluster4124 | -905.02019 | 0.0014 | 0.0068 | 4.6910 | 0.0062 |
| cluster4125 | -1065.596318 | 0.0523 | 0.0315 | 0.6022 | 0.0966 |
| cluster4126 | -762.147499 | 0.0427 | 0.0159 | 0.3737 | 0.1013 |
| cluster4127 | -763.036516 | 0.0161 | 0.0146 | 0.9056 | 0.0840 |
| cluster4128 | -989.452098 | 0.0252 | 0.0907 | 3.6010 | 0.0410 |

| dN | dS | Adjusted p value |
|---|---|---|
| 0.1643 | 0.1643 | 0.54 |
| 0.0749 | 0.0749 | 1.72E-34 |
| 0.0599 | 0.0599 | 0.48 |
| 0.1304 | 1.0818 | 3.20E-07 |
| 0.0046 | 0.4040 | 3.77E-05 |
| 0.0033 | 0.7369 | 5.60E-54 |
| 0.0173 | 0.3353 | 2.41E-09 |
| 0.0014 | 0.8013 | 9.20E-124 |
| 0.0107 | 0.7746 | 4.00E-42 |
| 0.0152 | 0.9039 | 4.49E-88 |
| 0.0332 | 0.8942 | 3.64E-44 |
| 0.0042 | 1.0388 | 3.25E-107 |
| 0.0631 | 0.5538 | 1.62E-12 |
| 0.0347 | 1.0337 | 1.03E-48 |
| 0.0519 | 0.9289 | 6.64E-28 |
| 0.0524 | 0.5529 | 2.20E-08 |
| 0.0348 | 0.6818 | 7.10E-06 |
| 0.0095 | 0.3102 | 3.14E-06 |
| 0.0817 | 0.9292 | 1.03E-10 |
| 0.0138 | 0.7435 | 1.30E-59 |
| 0.0225 | 0.5225 | 5.55E-22 |
| 0.0233 | 1.2006 | 2.71E-64 |
| 0.0278 | 0.7190 | 7.27E-11 |
| 0.0426 | 0.7237 | 6.58E-21 |
| 0.0157 | 0.3761 | 6.02E-12 |
| 0.0623 | 0.3514 | 0.005 |
| 0.0542 | 1.0273 | 1.57E-17 |
| 0.0454 | 0.7851 | 6.20E-19 |
| 0.0302 | 0.9207 | 2.23E-27 |
| 0.0422 | 0.6099 | 1.99E-07 |
| 0.0142 | 0.5269 | 1.30E-33 |
| 0.0569 | 1.1592 | 1.73E-32 |
| 0.0290 | 0.6727 | 7.78E-25 |
| 0.0084 | 1.0915 | 9.76E-55 |
| 0.0852 | 1.2458 | 7.24E-08 |
| 0.0100 | 0.9591 | 9.35E-47 |
| 0.0144 | 0.1945 | 0.06 |
| 0.0492 | 1.1296 | 1.38E-24 |
| 0.0122 | 0.6857 | 1.30E-34 |
| 0.0606 | 1.4768 | 4.48E-15 |
| 0.0161 | 0.7368 | 3.30E-17 |
| 0.0393 | 0.9580 | 4.43E-19 |

| | | |
|---|---|---|
| 0.0126 | 1.1280 | 6.15E-66 |
| 0.0249 | 0.6251 | 1.44E-22 |
| 0.0137 | 0.6015 | 2.03E-14 |
| 0.0286 | 1.2571 | 1.34E-29 |
| 0.0093 | 0.7079 | 1.72E-39 |
| 0.0413 | 1.0613 | 1.71E-20 |
| 0.0279 | 1.0652 | 1.89E-13 |
| 0.0108 | 0.9717 | 1.49E-45 |
| 0.0560 | 0.5475 | 5.36E-05 |
| 0.0169 | 0.4836 | 2.90E-19 |
| 0.0395 | 0.4788 | 4.38E-16 |
| 0.0956 | 1.1918 | 3.70E-11 |
| 0.0050 | 0.3360 | 4.63E-14 |
| 0.0130 | 0.3107 | 6.45E-05 |
| 0.0337 | 0.7249 | 2.29E-20 |
| 0.0006 | 0.2798 | 3.61E-10 |
| 0.0290 | 1.0512 | 5.94E-24 |
| 0.0022 | 0.7458 | 5.66E-41 |
| 0.0094 | 0.5609 | 5.42E-25 |
| 0.0158 | 0.7733 | 3.35E-19 |
| 0.0239 | 0.6805 | 1.44E-26 |
| 0.0212 | 0.8790 | 5.09E-30 |
| 0.0334 | 0.4346 | 0.0001 |
| 0.0346 | 0.5029 | 3.68E-13 |
| 0.0251 | 1.6053 | 4.58E-30 |
| 0.0044 | 1.0476 | 6.82E-27 |
| 0.0606 | 1.0184 | 1.28E-09 |
| 0.1451 | 0.6248 | 0.04 |
| 0.0293 | 1.1856 | 7.36E-33 |
| 0.0342 | 1.0441 | 1.52E-14 |
| 0.0198 | 0.8778 | 5.66E-41 |
| 0.0395 | 1.3506 | 2.30E-11 |
| 0.0227 | 0.9146 | 8.53E-25 |
| 0.0058 | 0.8824 | 3.44E-52 |
| 0.0539 | 0.6170 | 1.03E-09 |
| 0.0688 | 1.2290 | 3.03E-10 |
| 0.0596 | 0.9054 | 1.57E-12 |
| 0.0201 | 0.4864 | 2.45E-14 |
| 0.0306 | 0.4848 | 6.52E-12 |
| 0.0235 | 0.7341 | 4.10E-14 |
| 0.0497 | 1.2311 | 5.10E-24 |
| 0.0140 | 0.4840 | 1.29E-12 |
| 0.0111 | 0.7095 | 4.49E-29 |
| 0.0096 | 0.3319 | 1.24E-06 |

| | | |
|---|---|---|
| 0.0600 | 0.8351 | 7.68E-15 |
| 0.0005 | 0.1941 | 1.34E-06 |
| 0.0575 | 1.0524 | 1.03E-16 |
| 0.0097 | 0.8614 | 1.67E-23 |
| 0.0982 | 0.8320 | 8.03E-06 |
| 0.0049 | 1.9602 | 3.93E-21 |
| 0.0040 | 0.3539 | 2.28E-15 |
| 0.0218 | 0.4976 | 2.89E-21 |
| 0.0452 | 0.7198 | 4.22E-16 |
| 0.0117 | 0.5593 | 2.15E-23 |
| 0.0162 | 0.7163 | 5.73E-27 |
| 0.0742 | 1.2552 | 1.48E-10 |
| 0.0422 | 0.9458 | 5.31E-22 |
| 0.0860 | 1.1435 | 1.24E-14 |
| 0.0278 | 1.1641 | 2.34E-18 |
| 0.0216 | 0.7025 | 1.01E-15 |
| 0.0350 | 1.9791 | 3.45E-38 |
| 0.0927 | 0.8275 | 2.89E-06 |
| 0.0114 | 0.8654 | 9.65E-48 |
| 0.0332 | 0.6597 | 2.89E-09 |
| 0.0227 | 0.8095 | 3.60E-36 |
| 0.0222 | 2.0219 | 9.35E-47 |
| 0.0385 | 1.5063 | 7.05E-21 |
| 0.0148 | 0.3717 | 2.03E-14 |
| 0.0189 | 0.4625 | 6.90E-14 |
| 0.0073 | 0.7535 | 3.97E-32 |
| 0.0311 | 1.3018 | 5.05E-18 |
| 0.0112 | 0.5732 | 5.56E-40 |
| 0.0144 | 0.7013 | 9.79E-22 |
| 0.0327 | 0.7671 | 1.15E-15 |
| 0.0047 | 1.0753 | 3.13E-30 |
| 0.0086 | 0.2501 | 2.07E-05 |
| 0.0055 | 0.7655 | 6.88E-17 |
| 0.0833 | 0.7438 | 1.18E-05 |
| 0.0191 | 0.5312 | 1.70E-08 |
| 0.0724 | 0.8795 | 4.86E-11 |
| 0.0954 | 0.4327 | 0.47 |
| 0.0093 | 1.8351 | 4.78E-34 |
| 0.0010 | 0.7249 | 7.42E-57 |
| 0.0300 | 0.6603 | 6.42E-19 |
| 0.0522 | 2.6515 | 2.49E-05 |
| 0.0477 | 0.7824 | 6.26E-11 |
| 0.0541 | 0.4231 | 4.13E-05 |
| 0.0339 | 1.1530 | 2.04E-26 |

| | | |
|---|---|---|
| 0.0162 | 0.4566 | 1.39E-12 |
| 0.0826 | 0.3835 | 0.23 |
| 0.0571 | 0.4584 | 0.01 |
| 0.0180 | 0.5394 | 9.58E-14 |
| 0.0233 | 0.9461 | 1.06E-19 |
| 0.0178 | 0.4367 | 3.92E-13 |
| 0.0440 | 0.8900 | 4.33E-14 |
| 0.0762 | 1.0641 | 8.65E-08 |
| 0.0040 | 0.3666 | 6.22E-12 |
| 0.0252 | 0.7773 | 8.59E-17 |
| 0.0022 | 0.4472 | 2.56E-10 |
| 0.0212 | 1.0653 | 0.10 |
| 0.0303 | 0.6253 | 4.38E-16 |
| 0.0315 | 0.7273 | 5.75E-11 |
| 0.0217 | 0.9369 | 3.50E-27 |
| 0.0429 | 1.3124 | 9.97E-16 |
| 0.0045 | 0.5057 | 3.52E-18 |
| 0.0118 | 0.6215 | 5.28E-16 |
| 0.0189 | 0.5991 | 2.84E-22 |
| 0.0454 | 0.4757 | 1.83E-06 |
| 0.0405 | 0.4197 | 3.65E-05 |
| 0.0150 | 0.2886 | 4.15E-05 |
| 0.0480 | 0.8610 | 6.44E-10 |
| 0.0415 | 1.5781 | 2.03E-23 |
| 0.0310 | 0.8499 | 8.84E-17 |
| 0.0240 | 0.5098 | 1.03E-09 |
| 0.0776 | 0.4465 | 0.004 |
| 0.0176 | 1.2063 | 9.30E-17 |
| 0.0598 | 1.1163 | 2.28E-07 |
| 0.0340 | 0.3597 | 1.26E-06 |
| 0.0081 | 1.4331 | 1.07E-24 |
| 0.0163 | 0.3634 | 0.0002 |
| 0.0178 | 0.4107 | 7.25E-09 |
| 0.0345 | 1.0316 | 2.20E-20 |
| 0.0064 | 0.8640 | 4.34E-32 |
| 0.0620 | 0.9927 | 8.76E-12 |
| 0.0255 | 0.5970 | 2.63E-15 |
| 0.0092 | 0.4890 | 2.15E-10 |
| 0.0820 | 1.1194 | 0.0002 |
| 0.0071 | 0.5556 | 1.06E-19 |
| 0.0301 | 0.2968 | 0.02 |
| 0.0040 | 0.3638 | 1.24E-12 |
| 0.0905 | 0.5000 | 0.14 |
| 0.0022 | 0.9201 | 1.42E-30 |

| | | |
|---|---|---|
| 0.0549 | 1.2553 | 3.51E-12 |
| 0.0174 | 0.5514 | 8.58E-15 |
| 0.0198 | 0.3972 | 3.42E-08 |
| 0.0722 | 0.7230 | 6.38E-07 |
| 0.0227 | 0.6730 | 1.44E-22 |
| 0.0296 | 0.1824 | 0.15 |
| 0.3028 | 9.3374 | 0.005 |
| 0.0171 | 0.7032 | 6.31E-25 |
| 0.0656 | 0.9165 | 4.39E-08 |
| 0.0092 | 0.3805 | 6.25E-07 |
| 0.0546 | 0.7368 | 3.77E-07 |
| 0.0114 | 0.3419 | 3.43E-11 |
| 0.0084 | 0.3767 | 5.05E-14 |
| 0.0468 | 0.9552 | 3.62E-15 |
| 0.0304 | 0.7631 | 4.74E-12 |
| 0.0409 | 1.5886 | 8.08E-09 |
| 0.0297 | 0.9744 | 1.52E-16 |
| 0.0254 | 0.4859 | 7.77E-11 |
| 0.1197 | 0.6010 | 0.02 |
| 0.0031 | 0.6768 | 8.21E-09 |
| 0.0145 | 0.5760 | 3.18E-17 |
| 0.0137 | 0.7038 | 3.31E-31 |
| 0.0286 | 0.6302 | 1.81E-14 |
| 0.0479 | 1.1434 | 5.59E-14 |
| 0.0475 | 0.8725 | 1.25E-13 |
| 0.0681 | 0.6315 | 1.18E-07 |
| 0.0079 | 0.6924 | 7.67E-33 |
| 0.0237 | 0.5800 | 1.63E-08 |
| 0.0300 | 0.2906 | 9.94E-05 |
| 0.0175 | 0.3110 | 0.05 |
| 0.0107 | 0.3316 | 1.74E-05 |
| 0.1216 | 1.8165 | 2.29E-06 |
| 0.0336 | 0.9221 | 1.24E-11 |
| 0.0063 | 0.4253 | 2.31E-17 |
| 0.0161 | 0.3902 | 6.61E-10 |
| 0.0265 | 0.6815 | 8.15E-17 |
| 0.0049 | 0.3484 | 2.91E-05 |
| 0.0695 | 0.7024 | 0.02 |
| 0.0405 | 0.5918 | 4.99E-08 |
| 0.0301 | 0.5903 | 5.37E-11 |
| 0.0324 | 0.2903 | 0.003 |
| 0.0003 | 0.2007 | 0.001 |
| 0.0292 | 0.7399 | 2.84E-13 |
| 0.0125 | 0.4394 | 1.42E-17 |

| | | |
|---|---|---|
| 0.0678 | 0.7077 | 1.90E-07 |
| 0.0046 | 0.5121 | 8.86E-15 |
| 0.0404 | 1.4148 | 1.40E-13 |
| 0.0271 | 0.6340 | 7.44E-13 |
| 0.0500 | 0.8914 | 0.001 |
| 0.0180 | 0.7838 | 2.75E-16 |
| 0.0185 | 0.7227 | 2.44E-16 |
| 0.0239 | 0.7310 | 7.71E-20 |
| 0.0120 | 0.6850 | 8.86E-21 |
| 0.0067 | 0.6048 | 4.13E-18 |
| 0.0491 | 0.7360 | 4.36E-10 |
| 0.0085 | 0.4959 | 2.23E-11 |
| 0.0067 | 0.1870 | 0.002 |
| 0.0428 | 1.8375 | 2.49E-20 |
| 0.0581 | 0.4694 | 0.002 |
| 0.0112 | 0.5051 | 1.92E-20 |
| 0.0389 | 0.5186 | 6.30E-09 |
| 0.0439 | 0.5994 | 0.0001 |
| 0.0090 | 0.8715 | 5.07E-28 |
| 0.0218 | 0.3653 | 3.10E-07 |
| 0.0120 | 0.6826 | 3.91E-14 |
| 0.0071 | 1.1369 | 7.87E-16 |
| 0.0489 | 1.5138 | 7.33E-21 |
| 0.0059 | 0.7887 | 8.02E-27 |
| 0.0266 | 0.7407 | 2.01E-14 |
| 0.0090 | 0.4075 | 9.37E-19 |
| 0.0086 | 1.1511 | 3.31E-11 |
| 0.0320 | 0.6831 | 4.24E-09 |
| 0.0319 | 0.7228 | 1.66E-09 |
| 0.0355 | 0.6053 | 2.20E-08 |
| 0.0137 | 0.6931 | 1.70E-12 |
| 0.0594 | 1.3173 | 1.73E-09 |
| 0.0187 | 0.7298 | 2.85E-15 |
| 0.0428 | 1.5849 | 1.17E-16 |
| 0.0013 | 0.4887 | 2.79E-23 |
| 0.0178 | 0.4077 | 2.78E-05 |
| 0.0782 | 0.4871 | 0.01 |
| 0.0254 | 0.7763 | 2.02E-11 |
| 0.0524 | 0.4455 | 0.0003 |
| 0.0205 | 0.6208 | 1.79E-11 |
| 0.0604 | 1.2567 | 6.08E-10 |
| 0.0172 | 0.9185 | 1.60E-10 |
| 0.0190 | 0.4950 | 2.49E-10 |
| 0.0137 | 0.6787 | 3.94E-14 |

| | | |
|---|---|---|
| 0.0266 | 0.6089 | 1.41E-06 |
| 0.0688 | 1.1731 | 9.52E-13 |
| 0.0075 | 0.1860 | 0.01 |
| 0.0433 | 1.0246 | 6.25E-06 |
| 0.0155 | 0.6138 | 5.91E-16 |
| 0.0236 | 0.8340 | 2.89E-18 |
| 0.0114 | 0.4113 | 1.80E-10 |
| 0.0089 | 0.4684 | 8.24E-15 |
| 0.0303 | 0.6763 | 3.70E-14 |
| 0.0392 | 0.5490 | 4.25E-05 |
| 0.0307 | 0.6519 | 1.10E-15 |
| 0.0082 | 0.8618 | 3.44E-09 |
| 0.0008 | 0.4247 | 5.65E-10 |
| 0.0741 | 1.7454 | 8.70E-06 |
| 0.0359 | 0.9613 | 3.94E-14 |
| 0.0105 | 1.0967 | 2.48E-10 |
| 0.0163 | 0.6189 | 1.45E-15 |
| 0.0937 | 1.4172 | 2.38E-08 |
| 0.0238 | 0.3771 | 2.45E-05 |
| 0.0068 | 0.6752 | 4.96E-12 |
| 0.0356 | 0.9298 | 3.03E-09 |
| 0.0288 | 0.9285 | 1.01E-11 |
| 0.0484 | 0.8249 | 9.82E-09 |
| 0.0866 | 1.3304 | 4.05E-06 |
| 0.0143 | 0.2675 | 1.25E-05 |
| 0.0031 | 0.3947 | 3.61E-08 |
| 0.0123 | 0.1040 | 0.19 |
| 0.0080 | 0.4495 | 8.18E-12 |
| 0.0314 | 4.9876 | 6.16E-12 |
| 0.0064 | 1.2946 | 1.07E-50 |
| 0.0113 | 4.5789 | 3.06E-13 |
| 0.0055 | 0.7301 | 5.86E-27 |
| 0.0159 | 0.4760 | 3.02E-14 |
| 0.0016 | 0.4750 | 1.03E-09 |
| 0.0208 | 0.2875 | 8.68E-05 |
| 0.0250 | 0.5137 | 5.13E-06 |
| 0.0679 | 1.3190 | 2.01E-13 |
| 0.0206 | 0.4625 | 1.01E-08 |
| 0.0136 | 0.8589 | 2.24E-27 |
| 0.0090 | 0.7559 | 1.93E-17 |
| 0.0351 | 1.1487 | 2.45E-16 |
| 0.0220 | 0.2529 | 0.02 |
| 0.0170 | 0.8782 | 6.75E-16 |
| 0.0609 | 1.1519 | 2.74E-10 |

| | | |
|---|---|---|
| 0.0033 | 0.1467 | 0.09 |
| 0.0453 | 1.1772 | 5.14E-11 |
| 0.0040 | 0.3464 | 7.98E-11 |
| 0.0431 | 0.9992 | 2.20E-08 |
| 0.0547 | 0.9269 | 3.65E-09 |
| 0.0689 | 1.1017 | 6.22E-11 |
| 0.0183 | 0.5695 | 3.42E-16 |
| 0.0357 | 0.3850 | 8.68E-05 |
| 0.0236 | 1.1894 | 6.94E-30 |
| 0.0371 | 1.1496 | 3.30E-14 |
| 0.0024 | 0.1566 | 0.02 |
| 0.0502 | 0.6085 | 1.27E-07 |
| 0.0199 | 0.7100 | 5.87E-17 |
| 0.0135 | 0.8593 | 1.35E-20 |
| 0.0017 | 0.6621 | 4.72E-09 |
| 0.0084 | 0.5563 | 9.55E-07 |
| 0.0106 | 0.4029 | 2.53E-07 |
| 0.0278 | 0.4926 | 0.0005 |
| 0.0151 | 0.8636 | 2.10E-41 |
| 0.0968 | 1.1844 | 2.48E-05 |
| 0.0070 | 2.1981 | 3.89E-07 |
| 0.0204 | 0.9144 | 1.06E-24 |
| 0.0035 | 0.5091 | 4.18E-09 |
| 0.0655 | 0.7498 | 0.0001 |
| 0.0151 | 0.4286 | 8.08E-09 |
| 0.0209 | 1.3017 | 7.26E-18 |
| 0.0207 | 0.5843 | 3.32E-06 |
| 0.0179 | 0.7874 | 1.11E-17 |
| 0.0440 | 0.5087 | 0.0005 |
| 0.0554 | 1.1298 | 1.01E-10 |
| 0.0444 | 1.3090 | 6.78E-14 |
| 0.0305 | 0.6574 | 8.64E-05 |
| 0.0277 | 0.5330 | 4.40E-10 |
| 0.0173 | 0.3938 | 7.09E-07 |
| 0.0710 | 1.0080 | 3.63E-09 |
| 0.0139 | 1.4436 | 4.05E-18 |
| 0.0007 | 0.2025 | 6.41E-06 |
| 0.0244 | 0.4303 | 1.23E-07 |
| 0.0340 | 1.6270 | 1.29E-20 |
| 0.0071 | 0.5193 | 1.25E-11 |
| 0.0176 | 1.0493 | 7.71E-20 |
| 0.0290 | 0.2595 | 0.003 |
| 0.0232 | 0.5722 | 8.04E-11 |
| 0.0670 | 1.2863 | 0.0001 |

| | | |
|---|---|---|
| 0.0092 | 0.2434 | 0.003 |
| 0.0014 | 0.2153 | 4.29E-05 |
| 0.0300 | 0.8880 | 1.03E-09 |
| 0.0158 | 0.9124 | 6.75E-26 |
| 0.0030 | 0.5001 | 1.52E-09 |
| 0.0273 | 0.9556 | 1.78E-19 |
| 0.0122 | 2.8770 | 2.08E-21 |
| 0.0327 | 0.9898 | 1.08E-09 |
| 0.0013 | 0.4967 | 1.47E-12 |
| 0.0752 | 0.7490 | 0.0001 |
| 0.0297 | 0.5326 | 7.97E-12 |
| 0.0005 | 0.1750 | 0.0003 |
| 0.0599 | 1.7184 | 1.32E-10 |
| 0.0226 | 0.5542 | 1.08E-07 |
| 0.0331 | 1.4194 | 4.44E-14 |
| 0.0133 | 0.9257 | 1.03E-23 |
| 0.0162 | 0.4480 | 3.87E-06 |
| 0.0467 | 1.1249 | 4.33E-14 |
| 0.0207 | 0.5544 | 7.95E-06 |
| 0.0367 | 0.4848 | 0.002 |
| 0.0000 | 0.2866 | 8.60E-06 |
| 0.0322 | 0.8725 | 4.76E-17 |
| 0.0353 | 0.7049 | 1.36E-08 |
| 0.0695 | 0.9385 | 4.10E-06 |
| 0.0398 | 1.0935 | 6.48E-10 |
| 0.0250 | 0.6389 | 2.62E-09 |
| 0.0060 | 0.5342 | 1.62E-14 |
| 0.0185 | 0.6670 | 1.60E-11 |
| 0.0255 | 1.0875 | 1.66E-18 |
| 0.0432 | 0.8325 | 4.02E-08 |
| 0.0373 | 0.6613 | 3.29E-07 |
| 0.0646 | 1.6907 | 5.69E-12 |
| 0.0349 | 1.0230 | 3.94E-12 |
| 0.0463 | 0.7687 | 3.08E-05 |
| 0.0215 | 0.3017 | 0.002 |
| 0.0088 | 0.3645 | 2.68E-05 |
| 0.0303 | 0.8336 | 2.76E-19 |
| 0.0559 | 0.8257 | 1.23E-07 |
| 0.0663 | 1.3010 | 3.53E-09 |
| 0.0523 | 0.3215 | 0.01 |
| 0.0416 | 0.6013 | 1.89E-10 |
| 0.0031 | 0.8222 | 1.23E-21 |
| 0.0515 | 1.0377 | 2.90E-11 |
| 0.0393 | 0.5137 | 0.04 |

| | | |
|---|---|---|
| 0.0193 | 0.8082 | 9.49E-13 |
| 0.0151 | 0.8341 | 7.36E-14 |
| 0.0372 | 0.3967 | 0.01 |
| 0.0354 | 0.5789 | 1.31E-07 |
| 0.0821 | 1.2897 | 5.97E-08 |
| 0.0466 | 0.5905 | 2.04E-05 |
| 0.0272 | 0.7114 | 6.55E-09 |
| 0.0318 | 1.7382 | 3.61E-20 |
| 0.0348 | 0.7079 | 2.75E-13 |
| 0.0065 | 0.8090 | 1.18E-29 |
| 0.0007 | 0.4148 | 3.59E-08 |
| 0.0096 | 0.2789 | 1.35E-05 |
| 0.0087 | 0.6597 | 2.14E-18 |
| 0.0821 | 0.9874 | 1.91E-05 |
| 0.0254 | 1.2387 | 2.84E-22 |
| 0.0676 | 0.4360 | 0.08 |
| 0.0037 | 0.3096 | 2.54E-05 |
| 0.0024 | 0.7523 | 8.45E-31 |
| 0.0110 | 0.3062 | 8.34E-05 |
| 0.0225 | 0.6260 | 8.92E-11 |
| 0.0245 | 0.6575 | 5.98E-07 |
| 0.0469 | 0.6429 | 1.18E-05 |
| 0.0316 | 0.9118 | 4.22E-06 |
| 0.0075 | 0.4662 | 4.08E-16 |
| 0.0101 | 0.0585 | 0.71 |
| 0.0948 | 0.7842 | 0.004 |
| 0.0418 | 1.2058 | 1.91E-05 |
| 0.0475 | 0.8751 | 1.30E-08 |
| 0.0200 | 0.2015 | 0.02 |
| 0.0578 | 1.0696 | 5.42E-05 |
| 0.0013 | 0.3372 | 3.28E-17 |
| 0.0164 | 1.0382 | 1.19E-13 |
| 0.0130 | 0.5249 | 3.03E-11 |
| 0.0221 | 0.9937 | 7.42E-08 |
| 0.0030 | 0.5533 | 1.04E-11 |
| 0.0427 | 0.6750 | 1.57E-07 |
| 0.0253 | 0.7879 | 8.24E-15 |
| 0.0035 | 0.3659 | 2.67E-07 |
| 0.0104 | 0.3362 | 0.001 |
| 0.0087 | 0.5484 | 1.94E-06 |
| 0.0113 | 1.1330 | 8.14E-09 |
| 0.0189 | 0.3439 | 9.60E-10 |
| 0.0122 | 0.4768 | 0.001 |
| 0.0344 | 0.7319 | 5.22E-12 |

| | | |
|---|---|---|
| 0.0140 | 0.5203 | 1.49E-06 |
| 0.0095 | 0.7951 | 1.73E-19 |
| 0.0124 | 0.7388 | 1.35E-10 |
| 0.0008 | 0.4122 | 1.54E-10 |
| 0.0035 | 1.2710 | 1.95E-10 |
| 0.0934 | 0.4311 | 0.15 |
| 0.0035 | 0.8949 | 9.77E-22 |
| 0.1007 | 2.2731 | 1.36E-06 |
| 0.0053 | 0.8215 | 2.53E-15 |
| 0.0222 | 0.4227 | 0.001 |
| 0.0564 | 0.5621 | 2.71E-05 |
| 0.0465 | 1.2250 | 1.74E-11 |
| 0.0121 | 0.3908 | 7.38E-07 |
| 0.0539 | 0.7271 | 3.12E-05 |
| 0.0656 | 0.7786 | 7.29E-06 |
| 0.0042 | 0.9334 | 4.20E-12 |
| 0.0233 | 1.0179 | 1.28E-18 |
| 0.0455 | 0.3320 | 0.02 |
| 0.0126 | 0.6888 | 2.53E-15 |
| 0.0482 | 0.7616 | 6.26E-10 |
| 0.0295 | 1.1927 | 1.86E-08 |
| 0.0050 | 0.3974 | 2.17E-07 |
| 0.0848 | 1.5532 | 8.43E-11 |
| 0.0052 | 0.7106 | 1.59E-21 |
| 0.0716 | 0.8520 | 1.83E-05 |
| 0.0025 | 0.6028 | 1.37E-06 |
| 0.1111 | 0.9687 | 0.03 |
| 0.0391 | 1.1555 | 4.42E-10 |
| 0.0483 | 0.7933 | 0.0001 |
| 0.0016 | 0.5616 | 2.10E-09 |
| 0.0493 | 1.1533 | 1.97E-07 |
| 0.0397 | 0.7822 | 9.84E-10 |
| 0.0362 | 2.0095 | 0.001 |
| 0.0174 | 0.8126 | 4.22E-08 |
| 0.0171 | 0.2052 | 0.06 |
| 0.0188 | 0.6565 | 6.24E-13 |
| 0.0156 | 1.6790 | 1.28E-06 |
| 0.0918 | 1.2346 | 3.24E-05 |
| 0.0464 | 0.9807 | 9.23E-06 |
| 0.0327 | 0.3597 | 0.001 |
| 0.0722 | 1.1149 | 5.39E-09 |
| 0.0274 | 1.0800 | 2.92E-11 |
| 0.0309 | 0.3967 | 0.04 |
| 0.0326 | 0.9518 | 5.52E-09 |

| | | |
|---|---|---|
| 0.0012 | 0.4229 | 3.47E-13 |
| 0.0070 | 0.2780 | 1.38E-05 |
| 0.0575 | 1.3904 | 3.48E-12 |
| 0.0431 | 1.0424 | 7.70E-05 |
| 0.0323 | 0.1771 | 0.44 |
| 0.0136 | 1.6200 | 1.86E-08 |
| 0.0257 | 1.2294 | 3.77E-09 |
| 0.0005 | 0.8425 | 6.38E-24 |
| 0.0108 | 0.3741 | 4.14E-08 |
| 0.0187 | 1.1082 | 1.06E-15 |
| 0.0029 | 0.7699 | 5.84E-23 |
| 0.0214 | 0.4583 | 2.05E-07 |
| 0.0240 | 0.4337 | 1.51E-06 |
| 0.0099 | 0.8134 | 2.60E-17 |
| 0.0472 | 0.9107 | 9.60E-09 |
| 0.0150 | 0.7867 | 3.16E-11 |
| 0.0344 | 0.4988 | 1.25E-05 |
| 0.0055 | 0.3947 | 1.49E-05 |
| 0.0019 | 0.3677 | 8.80E-07 |
| 0.0390 | 0.7510 | 8.76E-09 |
| 0.0245 | 0.5275 | 2.58E-09 |
| 0.0163 | 0.5005 | 0.0002 |
| 0.0012 | 0.4926 | 7.69E-26 |
| 0.0512 | 1.0677 | 1.91E-13 |
| 0.0321 | 0.9039 | 7.97E-06 |
| 0.0544 | 1.3976 | 7.25E-09 |
| 0.0131 | 0.6417 | 1.33E-15 |
| 0.0066 | 2.2806 | 7.86E-35 |
| 0.0347 | 1.4973 | 1.10E-10 |
| 0.0454 | 0.7723 | 9.70E-09 |
| 0.0569 | 1.8016 | 4.76E-07 |
| 0.0820 | 0.6061 | 0.01 |
| 0.0074 | 0.1874 | 0.001 |
| 0.0185 | 0.9673 | 9.98E-17 |
| 0.0733 | 0.6790 | 6.67E-05 |
| 0.0026 | 0.3707 | 1.53E-11 |
| 0.0109 | 1.1264 | 7.81E-18 |
| 0.0118 | 1.2185 | 4.09E-34 |
| 0.1001 | 1.3249 | 9.56E-06 |
| 0.0027 | 0.4077 | 2.83E-09 |
| 0.0124 | 0.2979 | 0.01 |
| 0.0146 | 1.0990 | 1.97E-15 |
| 0.0466 | 0.9035 | 6.92E-12 |
| 0.0197 | 0.5472 | 0.0002 |

| | | |
|---|---|---|
| 0.0185 | 0.8543 | 1.02E-06 |
| 0.0244 | 0.7064 | 9.65E-06 |
| 0.0025 | 0.4502 | 0.01 |
| 0.0774 | 0.9969 | 2.48E-07 |
| 0.0244 | 0.5736 | 5.56E-05 |
| 0.0220 | 1.1367 | 3.19E-13 |
| 0.0162 | 0.9795 | 2.49E-20 |
| 0.0268 | 0.6214 | 5.50E-07 |
| 0.0173 | 0.6352 | 1.95E-09 |
| 0.0124 | 1.0594 | 3.90E-09 |
| 0.0392 | 0.6001 | 1.39E-06 |
| 0.0045 | 0.4613 | 2.73E-11 |
| 0.0119 | 1.0987 | 2.12E-12 |
| 0.0452 | 0.6921 | 0.0001 |
| 0.0020 | 0.8960 | 1.83E-06 |
| 0.0120 | 0.7318 | 6.82E-13 |
| 0.0187 | 0.5725 | 1.25E-12 |
| 0.0338 | 1.2982 | 5.17E-15 |
| 0.0297 | 0.4522 | 5.82E-06 |
| 0.0416 | 0.7108 | 2.50E-07 |
| 0.0262 | 0.2114 | 0.15 |
| 0.0152 | 0.7402 | 9.19E-09 |
| 0.0157 | 0.7367 | 1.06E-14 |
| 0.0155 | 0.6594 | 3.04E-11 |
| 0.0000 | 0.0927 | 0.11 |
| 0.0329 | 0.3605 | 0.06 |
| 0.0208 | 0.9615 | 5.55E-15 |
| 0.0622 | 0.9608 | 3.02E-08 |
| 0.0022 | 0.2801 | 3.01E-06 |
| 0.0333 | 1.4013 | 3.02E-14 |
| 0.0074 | 0.3370 | 0.0005 |
| 0.0003 | 0.6597 | 5.17E-15 |
| 0.0466 | 0.6347 | 0.001 |
| 0.0384 | 0.7559 | 1.82E-05 |
| 0.0335 | 0.2343 | 0.13 |
| 0.0320 | 0.4486 | 2.39E-05 |
| 0.4217 | 1.7154 | 0.03 |
| 0.0188 | 3.4404 | 2.79E-10 |
| 0.0458 | 0.7252 | 0.0002 |
| 0.0290 | 0.2809 | 0.07 |
| 0.0865 | 0.3072 | 0.39 |
| 0.0144 | 1.1725 | 1.62E-20 |
| 0.0207 | 0.6142 | 7.00E-14 |
| 0.0736 | 1.3129 | 3.75E-08 |

| | | |
|---|---|---|
| 0.0650 | 0.5092 | 0.01 |
| 0.0174 | 0.3388 | 0.002 |
| 0.0224 | 1.0722 | 1.69E-15 |
| 0.0449 | 0.5001 | 0.001 |
| 0.0285 | 0.4320 | 0.0004 |
| 0.0252 | 0.7818 | 6.96E-13 |
| 0.0608 | 1.3555 | 7.75E-09 |
| 0.0561 | 1.1748 | 4.36E-10 |
| 0.0197 | 0.4725 | 3.18E-08 |
| 0.0658 | 0.3674 | 0.03 |
| 0.0162 | 0.3950 | 4.55E-07 |
| 0.0181 | 0.8137 | 3.12E-15 |
| 0.0259 | 1.3545 | 4.92E-12 |
| 0.0137 | 1.0146 | 7.28E-15 |
| 0.0247 | 0.6645 | 1.14E-08 |
| 0.0393 | 0.8985 | 0.002 |
| 0.0107 | 0.3737 | 6.50E-06 |
| 0.0437 | 1.1547 | 8.56E-06 |
| 0.0429 | 3.6984 | 6.89E-08 |
| 0.0176 | 1.0134 | 9.38E-13 |
| 0.0078 | 0.6445 | 2.52E-14 |
| 0.0075 | 0.4413 | 3.00E-11 |
| 0.0330 | 0.9811 | 4.92E-12 |
| 0.0663 | 1.7101 | 2.98E-11 |
| 0.0274 | 1.2976 | 9.69E-08 |
| 0.0008 | 0.1601 | 0.04 |
| 0.0304 | 1.3902 | 1.64E-09 |
| 0.0290 | 1.0371 | 2.73E-10 |
| 0.0417 | 0.9368 | 6.43E-07 |
| 0.0206 | 0.5312 | 0.0003 |
| 0.0195 | 1.7337 | 1.27E-15 |
| 0.0126 | 0.2885 | 4.31E-05 |
| 0.0149 | 0.4044 | 2.46E-08 |
| 0.0316 | 0.9482 | 5.93E-10 |
| 0.0184 | 0.5418 | 0.0003 |
| 0.0612 | 1.0993 | 1.59E-05 |
| 0.0510 | 1.5198 | 1.92E-12 |
| 0.0605 | 0.9396 | 0.0004 |
| 0.0044 | 0.0947 | 0.20 |
| 0.0463 | 0.9409 | 1.86E-08 |
| 0.0686 | 1.2467 | 1.31E-09 |
| 0.0213 | 1.6242 | 2.78E-18 |
| 0.0137 | 0.8947 | 2.24E-13 |
| 0.0253 | 0.7387 | 7.79E-10 |

| | | |
|---|---|---|
| 0.0159 | 1.5804 | 2.54E-11 |
| 0.0184 | 0.7462 | 3.18E-08 |
| 0.0151 | 1.0163 | 4.43E-16 |
| 0.0358 | 0.8355 | 1.09E-08 |
| 0.0133 | 0.6401 | 2.57E-15 |
| 0.0229 | 0.7984 | 5.80E-11 |
| 0.0603 | 1.6477 | 7.20E-11 |
| 0.0150 | 1.0579 | 1.03E-08 |
| 0.0632 | 1.8272 | 7.52E-12 |
| 0.0053 | 0.2618 | 7.69E-08 |
| 0.0151 | 0.6314 | 3.24E-13 |
| 0.0770 | 1.7106 | 0.0002 |
| 0.0354 | 1.0747 | 1.04E-10 |
| 0.0308 | 8.0134 | 0.001 |
| 0.0202 | 0.2787 | 0.04 |
| 0.0102 | 0.8508 | 8.87E-19 |
| 0.0346 | 0.9222 | 7.58E-09 |
| 0.0349 | 0.7696 | 7.46E-06 |
| 0.0187 | 0.3628 | 8.37E-05 |
| 0.0201 | 0.8067 | 7.19E-07 |
| 0.0169 | 0.5462 | 0.0003 |
| 0.0476 | 1.0273 | 7.21E-08 |
| 0.0358 | 1.4508 | 9.21E-12 |
| 0.0272 | 0.7936 | 3.99E-09 |
| 0.0424 | 1.1985 | 2.19E-08 |
| 0.0269 | 0.6685 | 9.21E-11 |
| 0.0321 | 1.8977 | 1.19E-10 |
| 0.0217 | 0.6918 | 3.04E-08 |
| 0.0138 | 0.6137 | 6.36E-11 |
| 0.0021 | 0.2021 | 0.02 |
| 0.0618 | 1.3944 | 5.80E-07 |
| 0.0998 | 0.9492 | 0.04 |
| 0.0212 | 0.9414 | 1.85E-08 |
| 0.0213 | 0.9877 | 2.68E-06 |
| 0.0183 | 1.0078 | 7.07E-10 |
| 0.0582 | 1.0558 | 0.001 |
| 0.0130 | 2.8023 | 6.84E-08 |
| 0.0700 | 1.2623 | 4.59E-05 |
| 0.0240 | 0.8445 | 7.62E-11 |
| 0.0513 | 0.9777 | 9.43E-09 |
| 0.0725 | 2.3076 | 9.21E-11 |
| 0.0269 | 22.5285 | 0.0002 |
| 0.0356 | 0.3790 | 0.34 |
| 0.0235 | 0.6041 | 0.02 |

| | | |
|---|---|---|
| 0.0150 | 0.4290 | 3.90E-05 |
| 0.0072 | 0.1017 | 0.41 |
| 0.0251 | 0.6734 | 2.37E-12 |
| 0.0123 | 0.7165 | 6.38E-12 |
| 0.0247 | 0.8131 | 4.62E-12 |
| 0.0014 | 0.4535 | 4.82E-10 |
| 0.0758 | 0.8684 | 4.01E-05 |
| 0.0353 | 0.8188 | 2.93E-05 |
| 0.0657 | 0.4334 | 0.27 |
| 0.0473 | 1.1498 | 3.78E-09 |
| 0.0489 | 1.5179 | 2.17E-10 |
| 0.0329 | 1.0647 | 3.65E-05 |
| 0.0045 | 0.3566 | 4.21E-06 |
| 0.0250 | 0.4987 | 7.61E-05 |
| 0.0043 | 1.0877 | 4.60E-24 |
| 0.0686 | 1.3637 | 1.86E-08 |
| 0.0066 | 0.5052 | 1.16E-07 |
| 0.0442 | 1.3061 | 7.51E-10 |
| 0.0464 | 1.2081 | 2.30E-05 |
| 0.0360 | 0.6066 | 5.72E-06 |
| 0.0622 | 0.9836 | 1.96E-07 |
| 0.0120 | 0.3882 | 4.20E-07 |
| 0.0183 | 0.2961 | 0.01 |
| 0.0461 | 2.4643 | 3.31E-10 |
| 0.0226 | 0.6735 | 6.34E-09 |
| 0.0087 | 1.0991 | 1.52E-24 |
| 0.0157 | 0.3711 | 0.004 |
| 0.0922 | 2.2532 | 4.29E-06 |
| 0.0999 | 1.2280 | 0.0002 |
| 0.0197 | 0.7096 | 4.88E-11 |
| 0.0954 | 1.5835 | 1.40E-06 |
| 0.0613 | 1.3542 | 2.63E-09 |
| 0.0060 | 0.3894 | 0.005 |
| 0.0569 | 1.2295 | 6.73E-10 |
| 0.0390 | 1.4535 | 9.36E-08 |
| 0.0011 | 0.2969 | 0.004 |
| 0.0131 | 0.1637 | 0.06 |
| 0.0286 | 0.6497 | 1.90E-08 |
| 0.0023 | 0.2352 | 0.003 |
| 0.0153 | 0.3333 | 0.001 |
| 0.0323 | 1.5617 | 6.95E-08 |
| 0.0234 | 0.6093 | 1.40E-07 |
| 0.0172 | 0.5571 | 4.36E-10 |
| 0.0284 | 1.0406 | 9.85E-07 |

| | | |
|---|---|---|
| 0.0335 | 1.1305 | 5.85E-14 |
| 0.1342 | 4.6046 | 0.08 |
| 0.0652 | 0.8887 | 5.00E-07 |
| 0.0223 | 0.9733 | 2.79E-08 |
| 0.0756 | 1.1159 | 9.93E-08 |
| 0.0129 | 0.3468 | 0.001 |
| 0.0408 | 0.9182 | 1.33E-10 |
| 0.0112 | 0.7064 | 1.96E-07 |
| 0.0398 | 0.5082 | 5.19E-07 |
| 0.0485 | 0.9842 | 1.22E-08 |
| 0.0337 | 0.6904 | 0.0002 |
| 0.0561 | 0.8333 | 0.005 |
| 0.0371 | 1.4267 | 1.29E-07 |
| 0.0583 | 1.5299 | 1.91E-06 |
| 0.0486 | 2.4177 | 5.28E-07 |
| 0.0251 | 0.7765 | 3.40E-07 |
| 0.0578 | 1.0509 | 3.97E-06 |
| 0.0192 | 4.7759 | 1.41E-11 |
| 0.0728 | 0.9681 | 4.86E-06 |
| 0.0376 | 0.8865 | 8.38E-09 |
| 0.0439 | 1.2767 | 4.59E-07 |
| 0.0213 | 0.4291 | 7.61E-06 |
| 0.0344 | 0.9148 | 7.00E-08 |
| 0.0281 | 1.2489 | 3.99E-08 |
| 0.0190 | 1.7813 | 6.93E-12 |
| 0.0129 | 1.8357 | 2.99E-07 |
| 0.0057 | 0.1922 | 0.02 |
| 0.0001 | 0.9971 | 2.82E-18 |
| 0.0581 | 1.9142 | 6.90E-10 |
| 0.0256 | 0.3391 | 0.004 |
| 0.0781 | 1.0481 | 3.97E-06 |
| 0.0831 | 1.3250 | 3.29E-05 |
| 0.0125 | 0.3869 | 1.23E-07 |
| 0.0338 | 0.6865 | 9.37E-07 |
| 0.1048 | 1.5237 | 0.0002 |
| 0.0061 | 0.5275 | 1.45E-10 |
| 0.0550 | 1.2616 | 2.37E-10 |
| 0.0657 | 1.1085 | 1.18E-07 |
| 0.0412 | 0.6424 | 0.0002 |
| 0.0074 | 0.5234 | 1.62E-08 |
| 0.0275 | 0.8100 | 1.07E-06 |
| 0.0230 | 0.7841 | 1.46E-11 |
| 0.0963 | 4.5249 | 8.44E-08 |
| 0.0236 | 0.9142 | 5.97E-08 |

| | | |
|---|---|---|
| 0.0172 | 0.5632 | 6.01E-08 |
| 0.0137 | 1.5340 | 9.07E-19 |
| 0.0017 | 0.4531 | 6.86E-14 |
| 0.0490 | 1.0046 | 1.13E-08 |
| 0.0068 | 0.1078 | 0.24 |
| 0.0347 | 0.8752 | 4.95E-07 |
| 0.0379 | 1.3293 | 5.55E-13 |
| 0.0446 | 1.0787 | 2.01E-06 |
| 0.0072 | 0.7391 | 3.73E-12 |
| 0.0529 | 1.6347 | 1.73E-09 |
| 0.0246 | 5.3752 | 7.69E-05 |
| 0.0684 | 2.3960 | 2.57E-05 |
| 0.0564 | 1.2642 | 2.43E-07 |
| 0.0128 | 0.6974 | 2.10E-16 |
| 0.0610 | 2.0324 | 2.46E-09 |
| 0.0124 | 0.2120 | 0.01 |
| 0.0126 | 0.6209 | 0.004 |
| 0.0436 | 0.9095 | 0.001 |
| 0.0980 | 0.4995 | 0.32 |
| 0.0051 | 0.9098 | 4.92E-16 |
| 0.0449 | 0.7006 | 1.23E-07 |
| 0.0595 | 1.5781 | 1.44E-12 |
| 0.0073 | 0.4498 | 3.15E-10 |
| 0.0290 | 3.0294 | 3.63E-09 |
| 0.0292 | 2.0982 | 2.07E-10 |
| 0.0138 | 1.2703 | 1.93E-17 |
| 0.0192 | 0.7526 | 1.16E-06 |
| 0.0076 | 0.9303 | 1.26E-19 |
| 0.0486 | 1.7576 | 3.04E-08 |
| 0.0276 | 1.9042 | 1.14E-22 |
| 0.0059 | 0.2570 | 0.001 |
| 0.0249 | 0.8603 | 1.31E-14 |
| 0.0505 | 1.5275 | 3.89E-07 |
| 0.0162 | 1.1241 | 2.35E-06 |
| 0.0086 | 0.4046 | 0.002 |
| 0.0093 | 0.1941 | 0.02 |
| 0.0131 | 1.4105 | 2.45E-10 |
| 0.0046 | 0.2289 | 0.01 |
| 0.0208 | 0.5391 | 3.23E-06 |
| 0.0208 | 0.5158 | 0.002 |
| 0.0991 | 1.0820 | 8.22E-05 |
| 0.0068 | 0.2585 | 0.0002 |
| 0.0459 | 1.2665 | 9.27E-08 |
| 0.0996 | 0.9261 | 0.003 |

| | | |
|---|---|---|
| 0.0118 | 0.3975 | 9.27E-05 |
| 0.0348 | 1.1434 | 3.59E-06 |
| 0.0480 | 1.7832 | 4.91E-08 |
| 0.0164 | 5.5880 | 8.44E-25 |
| 0.0017 | 1.1106 | 3.62E-16 |
| 0.0067 | 0.5853 | 2.08E-06 |
| 0.0516 | 2.7624 | 0.001 |
| 0.0394 | 0.7527 | 3.69E-07 |
| 0.0209 | 0.7169 | 9.99E-12 |
| 0.0027 | 0.5319 | 4.65E-09 |
| 0.0129 | 1.1954 | 2.57E-16 |
| 0.0530 | 0.7980 | 0.002 |
| 0.0230 | 0.8010 | 1.09E-10 |
| 0.0355 | 2.3858 | 2.00E-11 |
| 0.0278 | 0.2060 | 0.22 |
| 0.0182 | 0.5487 | 2.97E-05 |
| 0.0439 | 1.8683 | 3.20E-11 |
| 0.0003 | 0.1232 | 0.01 |
| 0.0414 | 0.9749 | 2.96E-07 |
| 0.0386 | 0.6897 | 1.29E-05 |
| 0.0534 | 1.3360 | 2.24E-05 |
| 0.0302 | 1.2805 | 1.33E-14 |
| 0.0365 | 0.9441 | 9.99E-07 |
| 0.0078 | 0.2892 | 0.001 |
| 0.0231 | 0.8912 | 8.76E-12 |
| 0.0237 | 0.6038 | 4.62E-06 |
| 0.0436 | 0.8410 | 1.02E-08 |
| 0.0390 | 1.3294 | 2.51E-10 |
| 0.0433 | 0.8808 | 1.81E-05 |
| 0.1150 | 2.5930 | 3.06E-05 |
| 0.0419 | 1.5849 | 4.68E-05 |
| 0.0000 | 0.1771 | 0.01 |
| 0.0578 | 0.8881 | 4.13E-06 |
| 0.0110 | 0.6484 | 0.01 |
| 0.0324 | 0.7940 | 2.20E-08 |
| 0.0035 | 0.4294 | 0.0001 |
| 0.0231 | 2.3658 | 9.86E-14 |
| 0.0306 | 1.6486 | 1.51E-10 |
| 0.0531 | 1.1645 | 3.20E-12 |
| 0.1135 | 1.0478 | 0.02 |
| 0.0395 | 0.8807 | 3.24E-09 |
| 0.1013 | 1.2386 | 0.001 |
| 0.0668 | 3.2459 | 0.01 |
| 0.1340 | 1.9679 | 0.04 |

| | | |
|---|---|---|
| 0.0173 | 1.8426 | 6.96E-11 |
| 0.0234 | 0.4829 | 2.47E-06 |
| 0.0210 | 0.7157 | 2.14E-05 |
| 0.0030 | 0.7064 | 0.0002 |
| 0.0125 | 0.9938 | 7.71E-09 |
| 0.0175 | 2.4526 | 3.22E-17 |
| 0.0134 | 0.3648 | 0.002 |
| 0.0251 | 0.8448 | 1.91E-06 |
| 0.0328 | 33.4098 | 0.002 |
| 0.0485 | 0.5738 | 0.0002 |
| 0.0000 | 0.2974 | 7.38E-05 |
| 0.0604 | 1.3646 | 1.09E-09 |
| 0.0340 | 0.9727 | 9.73E-08 |
| 0.0006 | 0.3828 | 2.64E-06 |
| 0.0470 | 1.5502 | 2.30E-10 |
| 0.0078 | 0.3721 | 5.78E-05 |
| 0.0140 | 2.2145 | 2.13E-15 |
| 0.0103 | 0.5703 | 6.04E-09 |
| 0.0140 | 0.5672 | 2.22E-07 |
| 0.0539 | 1.4098 | 0.0002 |
| 0.0409 | 0.7388 | 0.0004 |
| 0.0220 | 0.2583 | 0.04 |
| 0.0268 | 1.4382 | 1.94E-08 |
| 0.0084 | 0.2346 | 0.03 |
| 0.0152 | 0.6627 | 2.48E-10 |
| 0.0814 | 1.1414 | 4.62E-07 |
| 0.0189 | 0.3652 | 0.003 |
| 0.0406 | 1.9138 | 8.56E-06 |
| 0.0524 | 1.0704 | 1.92E-07 |
| 0.0046 | 0.2978 | 0.001 |
| 0.0335 | 1.2098 | 2.28E-11 |
| 0.0367 | 0.7257 | 2.33E-06 |
| 0.0205 | 0.6807 | 1.96E-07 |
| 0.0011 | 0.1607 | 0.01 |
| 0.0894 | 1.9850 | 0.02 |
| 0.0000 | 0.3907 | 2.09E-05 |
| 0.0402 | 1.0035 | 3.63E-09 |
| 0.0142 | 2.1017 | 7.85E-11 |
| 0.0293 | 0.5782 | 0.0002 |
| 0.0730 | 1.1225 | 0.0003 |
| 0.0163 | 0.7069 | 1.92E-05 |
| 0.0183 | 1.6347 | 6.20E-12 |
| 0.0255 | 0.2674 | 0.02 |
| 0.0066 | 0.6215 | 1.77E-13 |

| | | |
|---|---|---|
| 0.0187 | 0.7694 | 2.53E-09 |
| 0.0377 | 0.4472 | 0.0005 |
| 0.0452 | 0.9767 | 5.99E-07 |
| 0.0429 | 0.4746 | 0.001 |
| 0.0256 | 0.5725 | 4.75E-07 |
| 0.0005 | 0.2715 | 0.0001 |
| 0.0080 | 0.9023 | 4.07E-19 |
| 0.0000 | 0.2881 | 0.001 |
| 0.0334 | 0.9174 | 3.15E-10 |
| 0.0087 | 0.4201 | 5.85E-05 |
| 0.0513 | 0.6969 | 0.001 |
| 0.0249 | 1.0847 | 1.23E-09 |
| 0.0456 | 0.4193 | 0.16 |
| 0.0308 | 1.0520 | 1.15E-08 |
| 0.0334 | 0.8751 | 1.96E-08 |
| 0.0169 | 0.5923 | 0.0002 |
| 0.0091 | 0.5414 | 1.17E-08 |
| 0.0232 | 0.9922 | 1.56E-14 |
| 0.0471 | 1.6275 | 4.74E-05 |
| 0.0052 | 0.9025 | 1.03E-19 |
| 0.0146 | 1.7075 | 1.26E-13 |
| 0.0027 | 0.1211 | 0.13 |
| 0.0000 | 0.1702 | 0.01 |
| 0.0125 | 0.8508 | 9.44E-12 |
| 0.0331 | 0.8006 | 2.61E-05 |
| 0.0181 | 0.4346 | 0.0001 |
| 0.0144 | 0.6990 | 0.0004 |
| 0.0451 | 0.5359 | 0.0003 |
| 0.0271 | 0.9250 | 4.58E-08 |
| 0.0267 | 6.0088 | 1.41E-05 |
| 0.0063 | 0.3144 | 4.79E-05 |
| 0.0162 | 0.2389 | 0.01 |
| 0.0427 | 1.4009 | 5.61E-05 |
| 0.0186 | 0.8553 | 4.78E-09 |
| 0.0128 | 1.4030 | 1.15E-07 |
| 0.0372 | 1.1280 | 0.0005 |
| 0.0338 | 0.9340 | 0.01 |
| 0.0026 | 0.2875 | 4.07E-05 |
| 0.0404 | 0.7182 | 0.002 |
| 0.0718 | 0.9639 | 0.005 |
| 0.0221 | 23.9176 | 0.01 |
| 0.0449 | 0.2052 | 0.59 |
| 0.0035 | 0.1889 | 0.003 |
| 0.0183 | 2.5889 | 0.001 |

| | | |
|---|---|---|
| 0.0330 | 0.4122 | 0.001 |
| 0.0692 | 0.8825 | 1.28E-05 |
| 0.0014 | 0.8977 | 1.08E-06 |
| 0.0311 | 1.4722 | 9.22E-10 |
| 0.0250 | 1.5045 | 1.16E-09 |
| 0.0399 | 0.6467 | 0.0001 |
| 0.0240 | 0.5875 | 1.40E-05 |
| 0.0303 | 3.2831 | 7.52E-14 |
| 0.0469 | 1.0797 | 3.59E-06 |
| 0.0087 | 0.6243 | 1.36E-08 |
| 0.0307 | 2.7367 | 7.76E-10 |
| 0.0207 | 0.7112 | 4.73E-05 |
| 0.0000 | 0.0936 | 0.11 |
| 0.0074 | 0.5006 | 9.45E-11 |
| 0.0301 | 1.3241 | 1.14E-08 |
| 0.0045 | 0.3970 | 0.02 |
| 0.0034 | 0.5470 | 1.94E-08 |
| 0.0223 | 0.6796 | 4.10E-06 |
| 0.0197 | 1.2907 | 2.40E-09 |
| 0.0191 | 0.8294 | 5.49E-06 |
| 0.0737 | 0.4622 | 0.13 |
| 0.1229 | 1.8766 | 9.49E-05 |
| 0.0280 | 1.1513 | 2.19E-13 |
| 0.0342 | 0.6659 | 0.0001 |
| 0.0011 | 0.0976 | 0.54 |
| 0.0836 | 0.5036 | 0.31 |
| 0.0468 | 1.1767 | 1.89E-06 |
| 0.0252 | 0.7026 | 6.64E-06 |
| 0.0296 | 0.8357 | 1.68E-06 |
| 0.0266 | 1.0400 | 2.00E-09 |
| 0.0392 | 1.0451 | 8.14E-09 |
| 0.0028 | 0.3418 | 1.48E-05 |
| 0.0345 | 1.1363 | 1.10E-05 |
| 0.0076 | 0.8933 | 9.28E-12 |
| 0.0102 | 1.0516 | 3.20E-13 |
| 0.0055 | 0.1952 | 0.07 |
| 0.0206 | 0.6534 | 2.70E-07 |
| 0.0780 | 2.1343 | 1.83E-06 |
| 0.0229 | 0.2117 | 0.10 |
| 0.0362 | 1.5096 | 2.63E-07 |
| 0.0401 | 1.4331 | 7.43E-09 |
| 0.0013 | 0.3585 | 0.001 |
| 0.0722 | 1.0877 | 0.001 |
| 0.0273 | 0.9364 | 1.85E-10 |

| | | |
|---|---|---|
| 0.0290 | 0.7565 | 6.12E-07 |
| 0.0147 | 0.6874 | 1.09E-07 |
| 0.0402 | 1.1260 | 5.40E-10 |
| 0.0291 | 1.1766 | 2.72E-14 |
| 0.0447 | 1.3554 | 3.21E-08 |
| 0.0062 | 0.7352 | 1.96E-08 |
| 0.0858 | 1.0559 | 0.002 |
| 0.0294 | 0.9180 | 6.23E-08 |
| 0.0373 | 0.7965 | 2.36E-05 |
| 0.0286 | 1.3460 | 6.17E-07 |
| 0.0078 | 0.3463 | 0.002 |
| 0.0491 | 1.6734 | 1.71E-06 |
| 0.0040 | 0.7262 | 8.85E-11 |
| 0.0177 | 0.8487 | 2.10E-09 |
| 0.0257 | 1.5687 | 2.90E-09 |
| 0.0289 | 0.7242 | 1.00E-05 |
| 0.0136 | 0.8444 | 1.57E-07 |
| 0.0360 | 0.8320 | 5.57E-07 |
| 0.0306 | 0.6099 | 3.73E-07 |
| 0.0232 | 0.9261 | 6.02E-07 |
| 0.0798 | 6.4660 | 1.48E-05 |
| 0.0932 | 1.2413 | 0.01 |
| 0.0012 | 0.2625 | 4.61E-05 |
| 0.0073 | 0.5141 | 7.92E-07 |
| 0.0398 | 0.4946 | 0.13 |
| 0.0000 | 0.2343 | 0.01 |
| 0.0165 | 0.5027 | 9.00E-06 |
| 0.0018 | 1.3320 | 1.18E-12 |
| 0.0392 | 1.1108 | 1.95E-07 |
| 0.0544 | 5.3586 | 5.63E-07 |
| 0.0132 | 0.7438 | 0.0003 |
| 0.0102 | 1.0337 | 5.27E-09 |
| 0.0127 | 1.9519 | 1.78E-13 |
| 0.0107 | 2.0813 | 2.67E-08 |
| 0.0526 | 0.4118 | 0.08 |
| 0.0307 | 2.7448 | 1.40E-16 |
| 0.0225 | 0.9299 | 1.95E-08 |
| 0.0215 | 1.1305 | 1.56E-14 |
| 0.0432 | 0.8088 | 3.92E-05 |
| 0.0121 | 0.7136 | 1.49E-08 |
| 0.0467 | 0.4028 | 0.003 |
| 0.0204 | 1.0415 | 5.98E-06 |
| 0.0001 | 0.6765 | 6.58E-07 |
| 0.0009 | 0.7243 | 6.19E-10 |

| | | |
|---|---|---|
| 0.0083 | 1.6067 | 3.94E-19 |
| 0.0629 | 1.2471 | 9.13E-06 |
| 0.0305 | 1.5331 | 1.33E-06 |
| 0.0000 | 0.0725 | 0.29 |
| 0.0154 | 0.4166 | 0.0004 |
| 0.0723 | 1.9394 | 1.82E-06 |
| 0.0370 | 0.8078 | 5.14E-07 |
| 0.0050 | 1.1303 | 2.12E-13 |
| 0.0163 | 1.5485 | 7.44E-13 |
| 0.0223 | 0.7740 | 1.85E-05 |
| 0.0052 | 0.2876 | 5.74E-06 |
| 0.0018 | 0.5336 | 0.0003 |
| 0.0063 | 0.4461 | 0.002 |
| 0.0089 | 0.4655 | 2.01E-05 |
| 0.0166 | 0.8443 | 9.10E-08 |
| 0.0241 | 1.3174 | 0.0002 |
| 0.0697 | 1.3868 | 3.39E-05 |
| 0.0278 | 0.8813 | 6.09E-09 |
| 0.1325 | 3.3573 | 0.06 |
| 0.0138 | 32.3566 | 2.62E-11 |
| 0.0154 | 0.9936 | 1.66E-14 |
| 0.0110 | 1.7523 | 2.56E-10 |
| 0.0242 | 2.9571 | 1.02E-07 |
| 0.0431 | 3.5480 | 3.47E-06 |
| 0.0382 | 1.4767 | 0.001 |
| 0.0294 | 1.3862 | 3.65E-07 |
| 0.0120 | 0.9036 | 8.92E-11 |
| 0.0062 | 0.5106 | 0.0005 |
| 0.0180 | 0.5395 | 4.65E-05 |
| 0.0202 | 1.7781 | 6.61E-05 |
| 0.0168 | 1.1738 | 7.08E-09 |
| 0.0063 | 1.1474 | 7.01E-06 |
| 0.0578 | 4.9650 | 4.60E-06 |
| 0.0083 | 0.3261 | 6.55E-06 |
| 0.0491 | 1.5061 | 9.42E-09 |
| 0.1234 | 1.2949 | 0.04 |
| 0.0036 | 1.2459 | 1.18E-14 |
| 0.0164 | 1.3464 | 1.07E-13 |
| 0.0354 | 2.6286 | 3.54E-07 |
| 0.0162 | 4.1408 | 0.0004 |
| 0.0848 | 0.5021 | 0.10 |
| 0.0173 | 0.6037 | 4.74E-05 |
| 0.0323 | 0.7191 | 7.42E-06 |
| 0.0169 | 1.0915 | 1.65E-06 |

| | | |
|---|---|---|
| 0.0474 | 1.4287 | 1.12E-07 |
| 0.0527 | 1.3050 | 0.002 |
| 0.0150 | 1.3680 | 3.31E-10 |
| 0.0000 | 0.3123 | 0.0001 |
| 0.0442 | 1.0985 | 6.15E-06 |
| 0.0752 | 1.2283 | 0.004 |
| 0.0359 | 1.1126 | 1.79E-06 |
| 0.0527 | 0.6042 | 0.08 |
| 0.0171 | 1.5684 | 8.02E-10 |
| 0.0548 | 2.8361 | 3.95E-08 |
| 0.0008 | 0.3533 | 0.10 |
| 0.0938 | 5.9353 | 0.02 |
| 0.0263 | 1.0971 | 4.10E-06 |
| 0.0387 | 0.8023 | 0.01 |
| 0.0006 | 2.9355 | 1.60E-13 |
| 0.0215 | 1.0695 | 2.43E-06 |
| 0.0355 | 0.5470 | 0.01 |
| 0.0751 | 1.2372 | 0.004 |
| 0.0156 | 1.0668 | 8.25E-07 |
| 0.0162 | 0.4492 | 0.001 |
| 0.0094 | 1.3349 | 2.53E-11 |
| 0.0084 | 0.5342 | 1.11E-08 |
| 0.0424 | 0.2521 | 0.22 |
| 0.0034 | 0.2917 | 0.01 |
| 0.0346 | 0.3513 | 0.12 |
| 0.0001 | 0.5055 | 0.001 |
| 0.0919 | 1.1952 | 0.0002 |
| 0.0284 | 1.2076 | 3.25E-08 |
| 0.0135 | 0.5386 | 1.96E-08 |
| 0.0202 | 0.5664 | 3.22E-05 |
| 0.0174 | 1.1221 | 3.03E-09 |
| 0.0083 | 0.4453 | 0.0001 |
| 0.0116 | 0.3428 | 0.04 |
| 0.0186 | 0.5448 | 1.28E-05 |
| 0.0357 | 1.9564 | 7.79E-10 |
| 0.0395 | 0.7582 | 0.0003 |
| 0.0420 | 1.0492 | 0.0002 |
| 0.0299 | 0.9961 | 4.58E-06 |
| 0.0302 | 1.3532 | 1.15E-07 |
| 0.0215 | 0.5314 | 0.002 |
| 0.0231 | 1.4226 | 0.001 |
| 0.0281 | 1.0844 | 6.35E-09 |
| 0.0462 | 1.0760 | 1.54E-07 |
| 0.0063 | 1.8151 | 1.24E-11 |

| | | |
|---|---|---|
| 0.0131 | 0.6812 | 0.0001 |
| 0.0231 | 0.6360 | 3.60E-05 |
| 0.0332 | 1.3222 | 0.0004 |
| 0.0239 | 1.5224 | 1.03E-09 |
| 0.0223 | 1.5568 | 2.69E-10 |
| 0.0248 | 1.0849 | 0.0004 |
| 0.0253 | 0.6313 | 2.50E-07 |
| 0.0556 | 1.2271 | 0.003 |
| 0.0360 | 1.2810 | 0.0005 |
| 0.0351 | 0.7657 | 0.001 |
| 0.0431 | 0.8283 | 8.59E-06 |
| 0.1717 | 1.8371 | 0.03 |
| 0.0570 | 1.6065 | 0.0002 |
| 0.0174 | 0.5874 | 6.49E-05 |
| 0.0367 | 0.3916 | 0.03 |
| 0.0143 | 0.7387 | 4.29E-05 |
| 0.0000 | 0.2680 | 0.0005 |
| 0.0568 | 1.1592 | 0.0002 |
| 0.0439 | 0.8742 | 4.67E-05 |
| 0.0119 | 1.2342 | 3.33E-05 |
| 0.0074 | 1.7671 | 0.0004 |
| 0.0178 | 0.8455 | 4.77E-07 |
| 0.0124 | 0.7646 | 0.001 |
| 0.0145 | 0.7059 | 1.17E-06 |
| 0.0728 | 0.7351 | 0.02 |
| 0.0038 | 0.5473 | 0.0003 |
| 0.0014 | 1.0067 | 8.08E-13 |
| 0.0000 | 0.2236 | 0.01 |
| 0.0252 | 0.9204 | 1.06E-06 |
| 0.0432 | 0.8596 | 0.0004 |
| 0.0621 | 1.5315 | 2.59E-06 |
| 0.0761 | 1.0793 | 0.02 |
| 0.0051 | 0.4606 | 0.0001 |
| 0.0344 | 1.4839 | 1.35E-07 |
| 0.0156 | 1.2558 | 6.57E-10 |
| 0.0071 | 0.5735 | 6.30E-06 |
| 0.0571 | 1.0747 | 0.002 |
| 0.0435 | 1.4978 | 9.93E-08 |
| 0.0253 | 1.2088 | 1.81E-06 |
| 0.0220 | 0.4722 | 0.0002 |
| 0.0382 | 1.1686 | 0.0002 |
| 0.0598 | 0.9683 | 0.0004 |
| 0.0362 | 0.8306 | 0.0002 |
| 0.0156 | 2.2039 | 6.20E-11 |

| | | |
|---|---|---|
| 0.0555 | 1.0097 | 0.0001 |
| 0.0358 | 0.6097 | 0.003 |
| 0.0465 | 1.3454 | 4.43E-06 |
| 0.0400 | 0.7563 | 9.37E-06 |
| 0.0195 | 1.0560 | 7.62E-07 |
| 0.0155 | 6.2854 | 1.47E-12 |
| 0.0707 | 0.7341 | 0.01 |
| 0.0055 | 0.8683 | 9.88E-09 |
| 0.0677 | 1.2382 | 0.0003 |
| 0.0219 | 0.5872 | 0.001 |
| 0.0024 | 0.6911 | 6.36E-11 |
| 0.0170 | 2.5474 | 8.20E-09 |
| 0.0279 | 2.4493 | 1.01E-05 |
| 0.0757 | 1.4045 | 0.01 |
| 0.0112 | 0.1678 | 0.16 |
| 0.1201 | 5.0745 | 4.15E-06 |
| 0.0125 | 0.3956 | 0.002 |
| 0.0067 | 1.5587 | 3.59E-06 |
| 0.0009 | 0.0554 | 0.46 |
| 0.0188 | 0.9520 | 2.67E-08 |
| 0.0471 | 0.9543 | 0.002 |
| 0.1380 | 0.8704 | 0.27 |
| 0.0762 | 1.0959 | 0.01 |
| 0.0186 | 0.4233 | 0.005 |
| 0.0153 | 0.4708 | 3.50E-05 |
| 0.0070 | 0.6464 | 0.002 |
| 0.0450 | 0.8214 | 0.003 |
| 0.0552 | 2.1655 | 1.64E-05 |
| 0.0019 | 2.7088 | 3.73E-05 |
| 0.0079 | 2.9893 | 6.53E-07 |
| 0.0157 | 2.8140 | 0.0001 |
| 0.0153 | 1.6571 | 1.56E-12 |
| 0.0815 | 1.0518 | 0.002 |
| 0.0257 | 0.5464 | 0.003 |
| 0.0513 | 1.3953 | 0.005 |
| 0.0005 | 0.3127 | 1.30E-06 |
| 0.0867 | 1.0507 | 0.003 |
| 0.0001 | 0.9794 | 0.01 |
| 0.0364 | 0.4806 | 0.0004 |
| 0.0118 | 1.2272 | 1.46E-05 |
| 0.0268 | 1.7202 | 6.58E-06 |
| 0.0246 | 2.0267 | 8.37E-05 |
| 0.0203 | 0.3198 | 0.02 |
| 0.0223 | 2.3059 | 0.004 |

| | | |
|---|---|---|
| 0.0177 | 1.6419 | 8.85E-07 |
| 0.0186 | 0.6144 | 0.01 |
| 0.0985 | 2.1791 | 0.03 |
| 0.0021 | 1.5115 | 4.77E-11 |
| 0.0727 | 0.9316 | 0.01 |
| 0.0014 | 0.4084 | 0.0003 |
| 0.0275 | 1.3693 | 9.51E-10 |
| 0.0047 | 0.4517 | 2.89E-06 |
| 0.0039 | 3.6313 | 0.12 |
| 0.0073 | 0.6873 | 5.71E-06 |
| 0.0029 | 0.2981 | 4.83E-05 |
| 0.0114 | 0.5241 | 8.34E-06 |
| 0.0300 | 1.9787 | 3.82E-06 |
| 0.0366 | 1.3656 | 7.20E-08 |
| 0.0156 | 7.0565 | 7.96E-06 |
| 0.0322 | 0.6199 | 0.002 |
| 0.0417 | 0.9704 | 9.17E-05 |
| 0.0486 | 1.1309 | 0.002 |
| 0.0140 | 1.0879 | 0.02 |
| 0.0262 | 0.8588 | 4.73E-06 |
| 0.0121 | 1.6956 | 4.32E-09 |
| 0.0161 | 0.7003 | 4.76E-07 |
| 0.0574 | 1.7237 | 0.0002 |
| 0.0163 | 0.8684 | 0.0004 |
| 0.0060 | 0.5890 | 0.0001 |
| 0.0401 | 0.9684 | 0.001 |
| 0.0721 | 6.2393 | 0.15 |
| 0.0557 | 2.7340 | 0.0003 |
| 0.0290 | 0.5410 | 0.05 |
| 0.0277 | 21.2490 | 0.0001 |
| 0.0049 | 1.8903 | 2.49E-06 |
| 0.0221 | 0.9715 | 0.003 |
| 0.0421 | 1.2672 | 4.86E-06 |
| 0.0552 | 2.0751 | 0.0005 |
| 2.8561 | 5.7650 | 0.96 |
| 0.0525 | 0.4804 | 0.02 |
| 0.0414 | 0.8880 | 0.003 |
| 0.0229 | 1.5652 | 4.87E-05 |
| 0.0012 | 1.0217 | 1.93E-10 |
| 0.0239 | 1.1519 | 0.01 |
| 0.0162 | 0.6523 | 9.18E-05 |
| 0.0788 | 7.3647 | 0.02 |
| 0.0900 | 1.1233 | 0.001 |
| 0.0088 | 0.8614 | 0.02 |

| | | |
|---|---|---|
| 0.0103 | 0.3178 | 0.15 |
| 0.0453 | 1.5655 | 0.0002 |
| 0.0269 | 2.4184 | 7.43E-06 |
| 0.0219 | 0.3310 | 0.03 |
| 0.0159 | 2.1653 | 9.93E-08 |
| 0.0229 | 0.5448 | 0.02 |
| 0.0134 | 0.3440 | 0.01 |
| 0.0470 | 1.2016 | 0.0002 |
| 0.0256 | 1.2332 | 0.0001 |
| 0.0015 | 0.1718 | 0.22 |
| 0.0626 | 1.5326 | 0.02 |
| 0.0380 | 1.3059 | 0.0001 |
| 0.0131 | 0.1386 | 0.50 |
| 0.0446 | 1.3998 | 1.70E-05 |
| 0.0016 | 1.3818 | 1.07E-06 |
| 0.0111 | 1.0100 | 1.24E-07 |
| 0.1432 | 0.4481 | 0.44 |
| 0.1287 | 3.0070 | 0.02 |
| 0.0147 | 0.2386 | 0.02 |
| 0.0111 | 1.5671 | 1.90E-07 |
| 0.0575 | 5.0364 | 0.005 |
| 0.0101 | 1.8657 | 3.64E-06 |
| 0.0037 | 1.0974 | 6.39E-07 |
| 0.0180 | 0.6488 | 9.81E-06 |
| 0.0875 | 1.3360 | 0.01 |
| 0.0735 | 0.8359 | 0.02 |
| 0.0664 | 1.4415 | 0.05 |
| 0.0136 | 0.7688 | 2.29E-05 |
| 0.0076 | 1.1733 | 0.03 |
| 0.0259 | 0.5013 | 0.01 |
| 0.0194 | 1.2878 | 1.73E-06 |
| 0.0680 | 2.6535 | 0.12 |
| 0.0076 | 0.5228 | 0.01 |
| 0.0385 | 1.1471 | 0.0001 |
| 0.0320 | 1.3053 | 0.0002 |
| 0.0032 | 0.6276 | 1.93E-06 |
| 0.0267 | 4.6612 | 4.44E-09 |
| 0.0018 | 0.0669 | 0.56 |
| 0.0049 | 0.6905 | 0.02 |
| 0.0157 | 2.5095 | 3.36E-07 |
| 0.0743 | 0.7696 | 0.07 |
| 0.0262 | 0.2588 | 0.20 |
| 0.0420 | 0.5007 | 0.02 |
| 0.0879 | 2.1477 | 0.02 |

**Supplementary Table 24.** codeml results - w for the D. hominis/other species

| ID | Likelihood | ϖ other species | dN | dS | ϖ *D. hominis* | dN |
|---|---|---|---|---|---|---|
| cluster1127 | -1409.1082 | 0.05308 | 0.1025 | 1.9304 | 0.1001 | 0.0012450 |
| cluster2853 | -22452.115 | 0.0236 | 0.0234 | 0.9889 | 0.0642 | 0.0646730 |
| cluster2854 | -1613.6079 | 0.0380 | 0.0150 | 0.3936 | 0.0001 | 0.0000100 |
| cluster2855 | -23548.962 | 0.0193 | 0.0368 | 1.9088 | 0.0764 | 0.0044090 |
| cluster2856 | -15426.043 | 0.1101 | 0.1585 | 1.4399 | 0.1651 | 0.0114250 |
| cluster2857 | -6009.464 | 0.0095 | 0.0040 | 0.4168 | 0.0901 | 0.0046310 |
| cluster2858 | -7755.334 | 0.0038 | 0.0038 | 1.0000 | 0.0093 | 0.0007120 |
| cluster2859 | -5677.7355 | 0.0391 | 0.0174 | 0.4449 | 0.1060 | 0.0043460 |
| cluster2860 | -16041.541 | 0.0020 | 0.0016 | 0.7690 | 0.0005 | 0.0003590 |
| cluster2861 | -8373.0215 | 0.0094 | 0.0087 | 0.9235 | 0.0182 | 0.0084330 |
| cluster2862 | -18851.924 | 0.0135 | 0.0106 | 0.7842 | 0.0226 | 0.0248510 |
| cluster2863 | -15068.936 | 0.0290 | 0.0374 | 1.2887 | 0.0800 | 0.0099770 |
| cluster2864 | -12550.761 | 0.0040 | 0.0037 | 0.9307 | 0.0025 | 0.0033530 |
| cluster2865 | -15003.672 | 0.0935 | 0.0489 | 0.5231 | 0.1173 | 0.0735920 |
| cluster2866 | -17309.508 | 0.0264 | 0.0393 | 1.4863 | 0.0574 | 0.0061140 |
| cluster2867 | -14837.022 | 0.0415 | 0.0569 | 1.3711 | 0.2901 | 0.0274240 |
| cluster2869 | -8923.1711 | 0.0742 | 0.0581 | 0.7830 | 57.4615 | 0.0211300 |
| cluster2870 | -3881.1101 | 0.0551 | 0.0430 | 0.7818 | 0.0461 | 0.0015960 |
| cluster2871 | -3135.398 | 0.0210 | 0.0074 | 0.3499 | 0.0820 | 0.0103250 |
| cluster2872 | -12260.323 | 0.0811 | 0.0972 | 1.1986 | 0.1191 | 0.0112530 |
| cluster2873 | -13847.541 | 0.0132 | 0.0092 | 0.6960 | 0.0249 | 0.0202100 |
| cluster2874 | -10375.77 | 0.0332 | 0.0231 | 0.6964 | 0.1454 | 0.0127740 |
| cluster2875 | -13634.052 | 0.0144 | 0.0188 | 1.3051 | 0.0249 | 0.0252350 |
| cluster2876 | -3971.4858 | 0.0330 | 0.0201 | 0.6101 | 0.0412 | 0.0407060 |
| cluster2877 | -10632.978 | 0.0465 | 0.0359 | 0.7708 | 0.0845 | 0.0416610 |
| cluster2878 | -7059.7588 | 0.0391 | 0.0138 | 0.3519 | 0.0412 | 0.0157120 |
| cluster2879 | -10032.185 | 0.1744 | 0.0603 | 0.3455 | 0.2935 | 0.0609470 |
| cluster2880 | -10635.097 | 0.0456 | 0.0632 | 1.3856 | 0.1356 | 0.0149570 |
| cluster2881 | -11522.907 | 0.0508 | 0.0329 | 0.6473 | 0.0588 | 0.0652110 |
| cluster2882 | -10635.775 | 0.0255 | 0.0283 | 1.1114 | 0.0853 | 0.0253310 |
| cluster2883 | -6764.3935 | 0.0669 | 0.0320 | 0.4788 | 0.0847 | 0.0661440 |
| cluster2884 | -12070.075 | 0.0251 | 0.0176 | 0.7011 | 0.0501 | 0.0034280 |
| cluster2885 | -15731.366 | 0.0399 | 0.0648 | 1.6239 | 0.1583 | 0.0127880 |
| cluster2886 | -12004.755 | 0.0393 | 0.0217 | 0.5527 | 0.0411 | 0.0403310 |
| cluster2887 | -8367.5042 | 0.0060 | 0.0091 | 1.5015 | 0.0180 | 0.0022470 |
| cluster2888 | -7782.0777 | 0.0586 | 0.1016 | 1.7342 | 0.1634 | 0.0275390 |
| cluster2889 | -6929.3331 | 0.0080 | 0.0111 | 1.3886 | 0.0361 | 0.0021850 |
| cluster2890 | -2673.3964 | 0.1183 | 0.0147 | 0.1245 | 0.0175 | 0.00418 |
| cluster2891 | -12094.453 | 0.0364 | 0.0548 | 1.5072 | 0.1302 | 0.0064350 |
| cluster2892 | -7904.3318 | 0.0112 | 0.0068 | 0.6087 | 0.0296 | 0.0238660 |
| cluster2893 | -8462.9441 | 0.0318 | 0.0688 | 2.1660 | 0.1192 | 0.0128920 |
| cluster2894 | -5512.8952 | 0.0164 | 0.0164 | 0.9971 | 0.0217 | 0.0029000 |

| | | | | | | |
|---|---|---|---|---|---|---|
| cluster2895 | -8444.0585 | 0.0331 | 0.0450 | 1.3610 | 0.1097 | 0.0114180 |
| cluster2896 | -10285.416 | 0.0097 | 0.0146 | 1.5055 | 0.0400 | 0.0025570 |
| cluster2897 | -9158.3466 | 0.0332 | 0.0283 | 0.8511 | 0.0782 | 0.0036200 |
| cluster2898 | -4499.189 | 0.0230 | 0.0119 | 0.5175 | 0.0408 | 0.0254310 |
| cluster2899 | -8003.6445 | 0.0170 | 0.0211 | 1.2432 | 0.0334 | 0.0347580 |
| cluster2900 | -9922.0311 | 0.0146 | 0.0077 | 0.5243 | 0.0093 | 0.0119760 |
| cluster2901 | -9240.5702 | 0.0361 | 0.0508 | 1.4056 | 0.0586 | 0.0056590 |
| cluster2902 | -3694.1438 | 0.0241 | 0.0348 | 1.4437 | 0.1015 | 0.0089340 |
| cluster2903 | -8863.9926 | 0.0108 | 0.0082 | 0.7646 | 0.0092 | 0.0155430 |
| cluster2904 | -6411.2928 | 0.1029 | 0.0499 | 0.4848 | 0.0559 | 0.0359380 |
| cluster2905 | -8879.8559 | 0.0273 | 0.0110 | 0.4016 | 0.0348 | 0.0267900 |
| cluster2906 | -13665.042 | 0.0678 | 0.0301 | 0.4445 | 0.0975 | 0.0551410 |
| cluster2907 | -12718.922 | 0.0642 | 0.1093 | 1.7035 | 0.5586 | 0.0231710 |
| cluster2908 | -5642.4778 | 0.0134 | 0.0054 | 0.4031 | 0.0001 | 0.0000120 |
| cluster2909 | -2924.6982 | 0.0363 | 0.0145 | 0.4010 | 0.0860 | 0.0040880 |
| cluster2910 | -9265.063 | 0.0402 | 0.0404 | 1.0072 | 0.2023 | 0.0088210 |
| cluster2911 | -4007.9777 | 0.0056 | 0.0012 | 0.2096 | 0.0017 | 0.0005880 |
| cluster2912 | -8764.4261 | 0.0246 | 0.0350 | 1.4210 | 0.1374 | 0.0056720 |
| cluster2913 | -6297.9235 | 0.0027 | 0.0024 | 0.8894 | 0.0019 | 0.0004420 |
| cluster2914 | -8063.9178 | 0.0161 | 0.0071 | 0.4423 | 0.0155 | 0.0126740 |
| cluster2915 | -4206.9758 | 0.0163 | 0.0170 | 1.0398 | 0.0340 | 0.0033540 |
| cluster2916 | -9770.5278 | 0.0318 | 0.0202 | 0.6360 | 0.0312 | 0.0255520 |
| cluster2917 | -9027.0649 | 0.0204 | 0.0236 | 1.1576 | 0.0146 | 0.0024140 |
| cluster2918 | -4475.2853 | 0.0712 | 0.0370 | 0.5201 | 0.1026 | 0.0108340 |
| cluster2919 | -9562.7949 | 0.0615 | 0.0273 | 0.4445 | 0.0900 | 0.0530260 |
| cluster2920 | -6955.8639 | 0.0149 | 0.0311 | 2.0906 | 0.0524 | 0.0045160 |
| cluster2921 | -3809.1558 | 0.0041 | 0.0053 | 1.2757 | 0.0020 | 0.0003920 |
| cluster2922 | -6244.1406 | 0.0548 | 0.0703 | 1.2820 | 0.0813 | 0.0119430 |
| cluster2923 | -5863.3047 | 0.1878 | 0.1444 | 0.7690 | 0.0001 | 0.00000 |
| cluster2924 | -10971.951 | 0.0197 | 0.0332 | 1.6864 | 0.0740 | 0.0074580 |
| cluster2925 | -5770.4968 | 0.0313 | 0.0402 | 1.2838 | 0.0649 | 0.0095330 |
| cluster2926 | -11217.193 | 0.0192 | 0.0152 | 0.7916 | 0.0303 | 0.0303070 |
| cluster2927 | -4903.7474 | 0.0295 | 0.0478 | 1.6222 | 0.0552 | 0.0096400 |
| cluster2928 | -7180.5505 | 0.0210 | 0.0266 | 1.2701 | 0.0350 | 0.0044620 |
| cluster2929 | -8010.4725 | 0.0055 | 0.0062 | 1.1281 | 0.0138 | 0.0012560 |
| cluster2930 | -10986.826 | 0.0810 | 0.0640 | 0.7895 | 0.1620 | 0.0105450 |
| cluster2931 | -6865.0032 | 0.0498 | 0.0820 | 1.6461 | 0.1074 | 0.0139120 |
| cluster2932 | -9693.6413 | 0.0597 | 0.0687 | 1.1508 | 0.0782 | 0.0196270 |
| cluster2933 | -8737.1674 | 0.0467 | 0.0164 | 0.3519 | 0.0476 | 0.0342920 |
| cluster2934 | -10531.145 | 0.0622 | 0.0379 | 0.6102 | 0.0973 | 0.0060930 |
| cluster2935 | -4952.2148 | 0.0295 | 0.0172 | 0.5815 | 0.0239 | 0.0310730 |
| cluster2936 | -9498.7827 | 0.0316 | 0.0588 | 1.8612 | 0.3831 | 0.0162390 |
| cluster2937 | -4594.6502 | 0.0253 | 0.0118 | 0.4644 | 0.0391 | 0.0209950 |
| cluster2938 | -7241.4398 | 0.0155 | 0.0103 | 0.6638 | 0.0149 | 0.0101880 |

| | | | | | | |
|---|---|---|---|---|---|---|
| cluster2939 | -2817.9494 | 0.0239 | 0.0062 | 0.2594 | 0.0220 | 0.0133720 |
| cluster2940 | -11373.287 | 0.0625 | 0.0695 | 1.1123 | 0.1970 | 0.0125190 |
| cluster2941 | -3030.3689 | 0.0023 | 0.0004 | 0.1529 | 0.0001 | 0.0000280 |
| cluster2942 | -11359.591 | 0.0542 | 0.0698 | 1.2877 | 0.0522 | 0.0092080 |
| cluster2943 | -4037.3495 | 0.0129 | 0.0102 | 0.7931 | 0.0038 | 0.0041550 |
| cluster2944 | -8493.1165 | 0.1086 | 0.0957 | 0.8812 | 0.0827 | 0.0395020 |
| cluster2945 | -3084.9557 | 0.0022 | 0.0053 | 2.3978 | 0.0071 | 0.0005480 |
| cluster2946 | -6842.7134 | 0.0001 | 0.0000 | 0.3250 | 0.0206 | 0.0083260 |
| cluster2947 | -9658.2713 | 0.0353 | 0.0226 | 0.6413 | 0.0696 | 0.0070750 |
| cluster2948 | -9371.0598 | 0.0462 | 0.0299 | 0.6483 | 0.0904 | 0.0816870 |
| cluster2949 | -6811.3435 | 0.0142 | 0.0077 | 0.5406 | 0.0307 | 0.0190440 |
| cluster2950 | -7118.1451 | 0.0048 | 0.0160 | 3.3055 | 0.0064 | 0.0032970 |
| cluster2951 | -9093.7064 | 0.0662 | 0.0857 | 1.2945 | 0.0398 | 0.0103600 |
| cluster2952 | -10891.451 | 0.0402 | 0.0508 | 1.2627 | 0.1093 | 0.0097640 |
| cluster2953 | -13284.964 | 0.0636 | 0.0948 | 1.4911 | 0.1386 | 0.0364680 |
| cluster2954 | -7273.7221 | 0.0207 | 0.0193 | 0.9335 | 0.0165 | 0.0397930 |
| cluster2955 | -6026.791 | 0.0233 | 0.0232 | 1.0001 | 999.0000 | 0.0024210 |
| cluster2956 | -10134.488 | 0.0137 | 0.0324 | 2.3714 | 0.0131 | 0.0156590 |
| cluster2957 | -8387.9623 | 0.0919 | 0.1123 | 1.2225 | 0.1300 | 0.0086130 |
| cluster2958 | -9688.5796 | 0.0111 | 0.0130 | 1.1673 | 0.0317 | 0.0037930 |
| cluster2959 | -5168.5469 | 0.0491 | 0.0253 | 0.5157 | 0.0682 | 0.0509770 |
| cluster2960 | -9440.9224 | 0.0220 | 0.0178 | 0.8083 | 0.0313 | 0.0290330 |
| cluster2961 | -8398.0246 | 0.0092 | 0.0182 | 1.9903 | 0.0110 | 0.0249180 |
| cluster2962 | -7495.1388 | 0.0214 | 0.0283 | 1.3262 | 0.0282 | 0.0609070 |
| cluster2963 | -7223.021 | 0.0333 | 0.0109 | 0.3264 | 0.0448 | 0.0220220 |
| cluster2964 | -5244.4677 | 0.0260 | 0.0118 | 0.4548 | 0.0593 | 0.0279190 |
| cluster2965 | -7391.7607 | 0.0111 | 0.0100 | 0.9013 | 0.0151 | 0.0020980 |
| cluster2966 | -5586.48 | 0.0213 | 0.0358 | 1.6860 | 0.0475 | 0.0092530 |
| cluster2967 | -9579.3326 | 0.0167 | 0.0090 | 0.5389 | 0.0188 | 0.0134480 |
| cluster2968 | -5454.9994 | 0.0166 | 0.0136 | 0.8195 | 0.0346 | 0.0117130 |
| cluster2969 | -7910.6214 | 0.0426 | 0.0408 | 0.9589 | 0.0422 | 0.0038470 |
| cluster2970 | -4236.3195 | 0.0038 | 0.0051 | 1.3723 | 0.0038 | 0.0005300 |
| cluster2971 | -3607.8637 | 0.0336 | 0.0088 | 0.2619 | 0.0149 | 0.0022750 |
| cluster2972 | -2669.0522 | 0.0052 | 0.0037 | 0.7166 | 0.0131 | 0.0070040 |
| cluster2973 | -6343.2744 | 0.0761 | 0.0924 | 1.2131 | 2.1912 | 0.0294630 |
| cluster2974 | -4421.3353 | 0.0313 | 0.0202 | 0.6456 | 0.0001 | 0.0000050 |
| cluster2975 | -10297.6 | 0.0652 | 0.0766 | 1.1757 | 0.1417 | 0.0254900 |
| cluster2976 | -1469.1965 | 0.1876 | 0.0941 | 0.5015 | 0.2864 | 0.02061 |
| cluster2977 | -3392.5245 | 0.0046 | 0.0113 | 2.4879 | 0.0185 | 0.0013070 |
| cluster2978 | -8212.9013 | 0.0019 | 0.0010 | 0.5483 | 0.0003 | 0.0003480 |
| cluster2979 | -8834.4253 | 0.0367 | 0.0318 | 0.8666 | 0.1275 | 0.0085660 |
| cluster2980 | -1554.0646 | 0.0214 | 0.0570 | 2.6642 | 0.0278 | 0.0054010 |
| cluster2981 | -6207.8269 | 0.0518 | 0.0558 | 1.0773 | 0.2201 | 0.0136850 |
| cluster2982 | -8047.1533 | 0.0931 | 0.0497 | 0.5335 | 0.2914 | 0.0361840 |

| | | | | | | |
|---|---|---|---|---|---|---|
| cluster2983 | -7509.5812 | 0.0203 | 0.0361 | 1.7754 | 0.2464 | 0.0094570 |
| cluster2984 | -4826.6678 | 0.0254 | 0.0105 | 0.4127 | 0.0613 | 0.0283350 |
| cluster2985 | -3972.5724 | 0.2339 | 0.0755 | 0.3230 | 0.2510 | 0.10899 |
| cluster2986 | -3495.1625 | 0.1019 | 0.0645 | 0.6328 | 758.1971 | 0.0060900 |
| cluster2987 | -6076.011 | 0.0328 | 0.0190 | 0.5791 | 0.0155 | 0.0048850 |
| cluster2988 | -5702.6411 | 0.0234 | 0.0301 | 1.2857 | 0.0488 | 0.0040910 |
| cluster2989 | -6589.7063 | 0.0394 | 0.0147 | 0.3736 | 0.0384 | 0.0232840 |
| cluster2990 | -7511.6557 | 0.0435 | 0.0493 | 1.1322 | 0.0583 | 0.0107640 |
| cluster2991 | -6307.9922 | 0.0572 | 0.0843 | 1.4747 | 0.1239 | 0.0082990 |
| cluster2992 | -4177.6095 | 0.0119 | 0.0045 | 0.3828 | 0.0027 | 0.0006240 |
| cluster2993 | -5422.7473 | 0.0303 | 0.0203 | 0.6711 | 0.0413 | 0.0417660 |
| cluster2994 | -2259.9174 | 0.0039 | 0.0022 | 0.5589 | 0.0001 | 0.0000100 |
| cluster2995 | -740.22899 | 0.0203 | 0.0253 | 1.2455 | 6.3580 | 0.00000 |
| cluster2996 | -7310.092 | 0.0409 | 0.0239 | 0.5837 | 0.0536 | 0.0413120 |
| cluster2997 | -8171.3782 | 0.0416 | 0.0378 | 0.9106 | 0.0266 | 0.0045230 |
| cluster2998 | -7504.1195 | 0.0186 | 0.0237 | 1.2718 | 0.0237 | 0.0023100 |
| cluster2999 | -6764.1459 | 0.0312 | 0.0480 | 1.5405 | 0.0265 | 0.0040740 |
| cluster3000 | -6334.2206 | 0.0115 | 0.0065 | 0.5678 | 0.0272 | 0.0008050 |
| cluster3001 | -6058.2258 | 0.0197 | 0.0140 | 0.7097 | 0.0341 | 0.0095930 |
| cluster3002 | -7668.0942 | 0.0282 | 0.0191 | 0.6783 | 0.0165 | 0.0052380 |
| cluster3003 | -7810.0954 | 0.0868 | 0.0511 | 0.5886 | 0.1444 | 0.0116690 |
| cluster3004 | -5964.2504 | 0.0868 | 0.0233 | 0.2683 | 0.1132 | 0.0829430 |
| cluster3005 | -4101.9438 | 0.0565 | 0.0199 | 0.3532 | 0.1494 | 0.0036190 |
| cluster3006 | -8671.6346 | 0.0551 | 0.0378 | 0.6865 | 0.0649 | 0.0742600 |
| cluster3007 | -8243.1 | 0.0279 | 0.0514 | 1.8429 | 0.0122 | 0.0017740 |
| cluster3008 | -5954.3421 | 0.0317 | 0.0361 | 1.1380 | 0.0330 | 0.0037420 |
| cluster3009 | -5909.8807 | 0.0394 | 0.0241 | 0.6133 | 0.0973 | 0.0127200 |
| cluster3010 | -8048.2947 | 0.1357 | 0.0790 | 0.5820 | 0.7325 | 0.0222610 |
| cluster3011 | -3371.8828 | 0.0090 | 0.0092 | 1.0155 | 0.0127 | 0.0357350 |
| cluster3012 | -5207.6851 | 0.0633 | 0.0867 | 1.3707 | 0.0195 | 0.0059270 |
| cluster3013 | -6734.9966 | 0.0846 | 0.0290 | 0.3426 | 0.0746 | 0.0349990 |
| cluster3014 | -3561.3869 | 0.0083 | 0.0124 | 1.4930 | 0.0001 | 0.0000060 |
| cluster3015 | -2410.5228 | 0.0420 | 0.0188 | 0.4489 | 0.0381 | 0.0032730 |
| cluster3016 | -4556.0773 | 0.0383 | 0.0200 | 0.5229 | 0.0631 | 0.0041830 |
| cluster3017 | -5972.1963 | 0.0267 | 0.0495 | 1.8550 | 0.0266 | 0.0032070 |
| cluster3018 | -5678.9264 | 0.0064 | 0.0070 | 1.0886 | 0.0126 | 0.0010160 |
| cluster3019 | -7570.3206 | 0.0560 | 0.0652 | 1.1648 | 0.0418 | 0.0240460 |
| cluster3020 | -6188.2061 | 0.0308 | 0.0274 | 0.8908 | 0.2106 | 0.0087180 |
| cluster3021 | -3024.8541 | 0.0161 | 0.0065 | 0.4012 | 0.0175 | 0.0123920 |
| cluster3022 | -3431.4246 | 0.0613 | 0.0942 | 1.5352 | 0.0427 | 0.0041660 |
| cluster3023 | -4202.5332 | 0.0080 | 0.0037 | 0.4704 | 0.0165 | 0.0138110 |
| cluster3024 | -1958.6756 | 0.0814 | 0.0317 | 0.3891 | 0.2207 | 0.0115980 |
| cluster3025 | -4691.8608 | 0.0101 | 0.0042 | 0.4124 | 0.0099 | 0.0010110 |
| cluster3026 | -4111.0837 | 0.1742 | 0.1163 | 0.6675 | 0.1364 | 0.01524 |

| | | | | | | |
|---|---|---|---|---|---|---|
| cluster3027 | -3436.6336 | 0.0023 | 0.0019 | 0.8108 | 0.0023 | 0.0026380 |
| cluster3028 | -6800.3267 | 0.0373 | 0.0620 | 1.6655 | 0.1548 | 0.0102380 |
| cluster3029 | -6675.2428 | 0.0279 | 0.0192 | 0.6887 | 0.0440 | 0.0045460 |
| cluster3030 | -5268.4371 | 0.0541 | 0.0173 | 0.3194 | 0.0529 | 0.0274440 |
| cluster3031 | -8796.7884 | 0.1021 | 0.0628 | 0.6145 | 0.0838 | 0.0876750 |
| cluster3032 | -5911.8083 | 0.0195 | 0.0147 | 0.7523 | 0.0426 | 0.0330860 |
| cluster3033 | -3824.1382 | 0.1584 | 0.0337 | 0.2126 | 0.1557 | 0.00237 |
| cluster3034 | -4966.993 | 0.0340 | 0.3160 | 9.2992 | 0.0278 | 0.0123000 |
| cluster3035 | -7637.5981 | 0.0179 | 0.0129 | 0.7196 | 0.0433 | 0.0230990 |
| cluster3036 | -8980.9494 | 0.0679 | 0.0784 | 1.1547 | 0.1114 | 0.0176060 |
| cluster3037 | -2563.7029 | 0.0158 | 0.0077 | 0.4866 | 0.0702 | 0.0058870 |
| cluster3038 | -4469.1988 | 0.0554 | 0.0584 | 1.0536 | 0.3873 | 0.0066820 |
| cluster3039 | -6525.3833 | 0.0384 | 0.0108 | 0.2806 | 0.0190 | 0.0095940 |
| cluster3040 | -4837.408 | 0.0139 | 0.0050 | 0.3631 | 0.0444 | 0.0163340 |
| cluster3041 | -8448.1087 | 0.0437 | 0.0575 | 1.3156 | 0.0559 | 0.0058390 |
| cluster3042 | -4402.5395 | 0.0320 | 0.0357 | 1.1159 | 0.2467 | 0.0040810 |
| cluster3043 | -2914.9611 | 0.0195 | 0.0540 | 2.7667 | 0.1396 | 0.0116620 |
| cluster3044 | -6428.2868 | 0.0303 | 0.0346 | 1.1426 | 0.0263 | 0.0049540 |
| cluster3045 | -6125.4362 | 0.0469 | 0.0253 | 0.5402 | 0.0902 | 0.0245210 |
| cluster3046 | -7266.5172 | 0.1992 | 0.1217 | 0.6107 | 0.0478 | 0.0107580 |
| cluster3047 | -2123.6959 | 0.0033 | 0.0019 | 0.5855 | 0.0039 | 0.0034090 |
| cluster3048 | -5405.8835 | 0.0228 | 0.0174 | 0.7644 | 0.0001 | 0.0000080 |
| cluster3049 | -7064.0624 | 0.0157 | 0.0092 | 0.5884 | 0.0204 | 0.0188310 |
| cluster3050 | -7561.7135 | 0.0389 | 0.0306 | 0.7865 | 0.0629 | 0.0061690 |
| cluster3051 | -7509.6043 | 0.0354 | 0.0565 | 1.5970 | 0.1052 | 0.0121130 |
| cluster3052 | -6822.6371 | 0.0440 | 0.0567 | 1.2884 | 0.1735 | 0.0134650 |
| cluster3053 | -9370.2834 | 0.0882 | 0.0538 | 0.6104 | 0.1113 | 0.0952290 |
| cluster3054 | -6880.5576 | 0.0093 | 0.0084 | 0.8986 | 0.0112 | 0.0009490 |
| cluster3055 | -3993.4093 | 0.0305 | 0.0237 | 0.7766 | 0.0001 | 0.0000050 |
| cluster3056 | -5995.1868 | 0.1007 | 0.0253 | 0.2511 | 0.1007 | 0.0360970 |
| cluster3057 | -918.77815 | 0.0652 | 0.0151 | 0.2323 | 0.0246 | 0.0130170 |
| cluster3058 | -2525.241 | 0.0099 | 0.0021 | 0.2122 | 0.0617 | 0.0366390 |
| cluster3059 | -8712.4407 | 0.0628 | 0.1465 | 2.3331 | 0.1124 | 0.0156770 |
| cluster3060 | -5326.7539 | 0.0340 | 0.0379 | 1.1142 | 0.0363 | 0.0076710 |
| cluster3061 | -5740.719 | 0.0152 | 0.0053 | 0.3478 | 0.0186 | 0.0111260 |
| cluster3062 | -5690.6482 | 0.0379 | 0.0174 | 0.4592 | 0.0223 | 0.0027300 |
| cluster3063 | -6139.6654 | 0.0263 | 0.0230 | 0.8750 | 0.0874 | 0.0218360 |
| cluster3064 | -1918.646 | 0.0124 | 0.0049 | 0.3947 | 0.0001 | 0.0000060 |
| cluster3065 | -2297.1856 | 0.0824 | 0.0735 | 0.8921 | 0.2986 | 0.0370550 |
| cluster3066 | -6187.0931 | 0.0659 | 0.0294 | 0.4457 | 0.0808 | 0.0650450 |
| cluster3067 | -4920.6831 | 0.0405 | 0.0219 | 0.5395 | 0.0421 | 0.0413450 |
| cluster3068 | -5141.3856 | 0.0982 | 0.0252 | 0.2567 | 0.1539 | 0.0459820 |
| cluster3069 | -1632.3953 | 0.0014 | 0.0003 | 0.2288 | 0.0001 | 0.0000010 |
| cluster3070 | -5192.4946 | 0.0365 | 0.0258 | 0.7061 | 0.0373 | 0.0301910 |

| | | | | | | |
|---|---|---|---|---|---|---|
| cluster3071 | -5850.3646 | 0.0159 | 0.0058 | 0.3637 | 0.0428 | 0.0266850 |
| cluster3072 | -7366.3773 | 0.0760 | 0.0757 | 0.9970 | 0.2519 | 0.0279730 |
| cluster3073 | -3570.6027 | 0.0075 | 0.0052 | 0.6896 | 0.0104 | 0.0007690 |
| cluster3074 | -7650.0727 | 0.0283 | 0.0483 | 1.7104 | 0.0743 | 0.0049660 |
| cluster3075 | -5425.9011 | 0.0326 | 0.0195 | 0.5961 | 0.0544 | 0.0470260 |
| cluster3076 | -1803.0368 | 0.0289 | 0.0204 | 0.7075 | 0.2055 | 0.1305530 |
| cluster3077 | -4385.0051 | 0.0227 | 0.0164 | 0.7243 | 0.0257 | 0.0199410 |
| cluster3078 | -5021.6597 | 0.0185 | 0.0159 | 0.8634 | 0.0557 | 0.0174510 |
| cluster3079 | -7622.042 | 0.0334 | 0.0196 | 0.5853 | 0.0337 | 0.0332140 |
| cluster3080 | -5649.8165 | 0.0157 | 0.0137 | 0.8722 | 0.0239 | 0.0034550 |
| cluster3081 | -3875.7124 | 0.0093 | 0.0043 | 0.4641 | 0.0121 | 0.0110060 |
| cluster3082 | -7187.7348 | 0.0594 | 0.0562 | 0.9470 | 0.0514 | 0.0086740 |
| cluster3083 | -4157.7674 | 0.0312 | 0.0115 | 0.3668 | 0.0140 | 0.0113110 |
| cluster3084 | -3197.9276 | 0.0354 | 0.0053 | 0.1507 | 0.0423 | 0.0098830 |
| cluster3085 | -7344.5022 | 0.0219 | 0.0505 | 2.3030 | 0.0253 | 0.0055280 |
| cluster3086 | -3352.3137 | 0.0370 | 0.0135 | 0.3650 | 0.4830 | 0.2022120 |
| cluster3087 | -6567.7828 | 0.0187 | 0.0118 | 0.6313 | 0.0264 | 0.0024570 |
| cluster3088 | -6393.174 | 0.0667 | 0.0436 | 0.6540 | 0.0495 | 0.0060580 |
| cluster3089 | -3685.0626 | 0.0676 | 0.0290 | 0.4293 | 0.0899 | 0.0704700 |
| cluster3090 | -4960.5466 | 0.0090 | 0.0093 | 1.0396 | 0.0040 | 0.0011610 |
| cluster3091 | -5908.6464 | 0.0472 | 0.0149 | 0.3159 | 0.0800 | 0.0388190 |
| cluster3092 | -4264.5663 | 0.0164 | 0.0135 | 0.8220 | 0.0391 | 0.0041590 |
| cluster3093 | -4546.778 | 0.0064 | 0.0082 | 1.2656 | 0.8096 | 0.0000010 |
| cluster3094 | -7861.8898 | 0.0273 | 0.0552 | 2.0185 | 0.0958 | 0.0115510 |
| cluster3095 | -4863.6148 | 0.0036 | 0.0019 | 0.5394 | 0.0139 | 0.0182640 |
| cluster3096 | -6131.3029 | 0.0319 | 0.0298 | 0.9346 | 0.0683 | 0.0061990 |
| cluster3097 | -7075.7746 | 0.0189 | 0.0081 | 0.4265 | 0.0338 | 0.0099500 |
| cluster3098 | -1418.6373 | 0.0068 | 0.0100 | 1.4762 | 0.0232 | 0.0015820 |
| cluster3099 | -4272.9576 | 0.0427 | 0.0284 | 0.6651 | 0.0308 | 0.0186930 |
| cluster3100 | -4393.3871 | 0.0440 | 0.0387 | 0.8817 | 0.0173 | 0.0020980 |
| cluster3101 | -3429.2866 | 0.0368 | 0.0253 | 0.6864 | 0.1325 | 0.0500230 |
| cluster3102 | -2230.6356 | 0.0124 | 0.0083 | 0.6675 | 0.0233 | 0.0257040 |
| cluster3103 | -5216.4064 | 0.0380 | 0.0674 | 1.7750 | 0.0256 | 0.0055800 |
| cluster3104 | -4326.4084 | 0.0194 | 0.0156 | 0.8037 | 0.0336 | 0.0171460 |
| cluster3105 | -7971.3 | 0.0261 | 0.0515 | 1.9766 | 0.0535 | 0.0095530 |
| cluster3106 | -5132.2231 | 0.0029 | 0.0011 | 0.3981 | 0.0024 | 0.0017400 |
| cluster3107 | -3137.8512 | 0.0417 | 0.0195 | 0.4671 | 0.0332 | 0.0023640 |
| cluster3108 | -4324.0056 | 0.1178 | 0.0825 | 0.7007 | 0.7216 | 0.0423250 |
| cluster3109 | -3461.6809 | 0.0263 | 0.0299 | 1.1397 | 0.0651 | 0.0071920 |
| cluster3110 | -6381.1278 | 0.1074 | 0.0599 | 0.5574 | 0.1288 | 0.0072500 |
| cluster3111 | -4889.3833 | 0.0322 | 0.0234 | 0.7278 | 0.0412 | 0.0034260 |
| cluster3112 | -6141.7496 | 0.0358 | 0.0643 | 1.7964 | 0.0898 | 0.0212410 |
| cluster3113 | -3068.3025 | 0.0244 | 0.0255 | 1.0441 | 0.0431 | 0.0059750 |
| cluster3114 | -4690.0546 | 0.0335 | 0.0192 | 0.5741 | 0.0224 | 0.0030450 |

| | | | | | | |
|---|---|---|---|---|---|---|
| cluster3115 | -3777.6004 | 0.0179 | 0.0147 | 0.8200 | 0.0216 | 0.0033770 |
| cluster3116 | -2723.2541 | 0.0398 | 0.0319 | 0.8033 | 0.0101 | 0.0012720 |
| cluster3117 | -7495.8244 | 0.0431 | 0.0812 | 1.8845 | 0.1431 | 0.0135680 |
| cluster3118 | -1575.8839 | 0.0336 | 0.0078 | 0.2330 | 0.0001 | 0.0000010 |
| cluster3119 | -2982.5758 | 0.0412 | 0.0277 | 0.6724 | 0.0319 | 0.0721800 |
| cluster3120 | -4909.8876 | 0.0216 | 0.0167 | 0.7733 | 0.0410 | 0.0053830 |
| cluster3121 | -6820.9854 | 0.0257 | 0.0164 | 0.6387 | 0.0328 | 0.0389580 |
| cluster3122 | -3558.4274 | 0.0207 | 0.0075 | 0.3607 | 0.0215 | 0.0141430 |
| cluster3123 | -3825.5626 | 0.0182 | 0.0082 | 0.4486 | 0.0017 | 0.0011190 |
| cluster3124 | -6376.6125 | 0.0376 | 0.0330 | 0.8776 | 0.0659 | 0.0084820 |
| cluster3125 | -4802.932 | 0.0677 | 0.0437 | 0.6454 | 0.0449 | 0.0040410 |
| cluster3126 | -7552.7753 | 0.0411 | 0.0268 | 0.6526 | 0.0478 | 0.0334110 |
| cluster3127 | -2606.1922 | 0.0073 | 0.0082 | 1.1215 | 0.0272 | 0.0031960 |
| cluster3128 | -2238.3148 | 0.0022 | 0.0011 | 0.5067 | 0.0001 | 0.0000040 |
| cluster3129 | -3260.2664 | 0.0288 | 0.0862 | 2.9958 | 0.2327 | 0.0166100 |
| cluster3130 | -5794.377 | 0.0319 | 0.0424 | 1.3302 | 0.2156 | 0.0127050 |
| cluster3131 | -1762.1327 | 0.0065 | 0.0102 | 1.5596 | 0.0577 | 0.0062750 |
| cluster3132 | -5289.6429 | 0.0232 | 0.0122 | 0.5244 | 0.0305 | 0.0217570 |
| cluster3133 | -8922.9858 | 0.0564 | 0.1128 | 2.0012 | 0.1347 | 0.0152450 |
| cluster3134 | -3290.9697 | 0.0598 | 0.0214 | 0.3583 | 0.0527 | 0.0214960 |
| cluster3135 | -4395.4879 | 0.0096 | 0.0078 | 0.8068 | 0.0820 | 0.0035590 |
| cluster3136 | -3954.0913 | 0.0378 | 0.0457 | 1.2071 | 0.0387 | 0.0032600 |
| cluster3137 | -3089.3169 | 0.0259 | 0.0319 | 1.2300 | 0.0001 | 0.0000100 |
| cluster3138 | -4742.2016 | 0.0496 | 0.0447 | 0.9003 | 0.0768 | 0.0466970 |
| cluster3139 | -5753.6533 | 0.0662 | 0.1094 | 1.6543 | 0.0816 | 0.0297950 |
| cluster3140 | -3557.054 | 0.0393 | 0.0150 | 0.3816 | 0.4460 | 0.0041860 |
| cluster3141 | -2414.5272 | 0.0119 | 0.0036 | 0.2988 | 0.0048 | 0.0023120 |
| cluster3142 | -1638.1161 | 0.0767 | 0.0106 | 0.1384 | 0.7914 | 0.01276 |
| cluster3143 | -3225.2417 | 0.0171 | 0.0074 | 0.4325 | 0.0039 | 0.0022900 |
| cluster3144 | -2088.7622 | 0.0049 | 0.0349 | 7.1703 | 0.0001 | 0.0000020 |
| cluster3145 | -6529.9487 | 0.0051 | 0.0076 | 1.4879 | 0.0107 | 0.0028440 |
| cluster3146 | -1739.5848 | 0.0007 | 0.0035 | 4.7227 | 0.0128 | 0.0332590 |
| cluster3147 | -4473.0776 | 0.0062 | 0.0060 | 0.9806 | 0.0100 | 0.0006920 |
| cluster3148 | -6023.3584 | 0.0290 | 0.0182 | 0.6264 | 0.0289 | 0.0027520 |
| cluster3149 | -2371.6156 | 0.0037 | 0.0015 | 0.4016 | 0.0001 | 0.0000550 |
| cluster3150 | -4524.2153 | 0.0654 | 0.0240 | 0.3666 | 0.1452 | 0.0027840 |
| cluster3151 | -3975.4891 | 0.0509 | 0.0321 | 0.6301 | 3.6814 | 0.0000010 |
| cluster3152 | -8742.5828 | 0.0446 | 0.0852 | 1.9087 | 0.1317 | 0.0142740 |
| cluster3153 | -4516.9772 | 0.0423 | 0.0245 | 0.5793 | 0.0563 | 0.0041240 |
| cluster3154 | -5338.3532 | 0.0120 | 0.0125 | 1.0368 | 0.0378 | 0.0160070 |
| cluster3155 | -3013.0974 | 0.0078 | 0.0083 | 1.0633 | 0.0586 | 0.0028360 |
| cluster3156 | -6782.7403 | 0.0232 | 0.0380 | 1.6419 | 0.0930 | 0.0113410 |
| cluster3157 | -3280.9876 | 0.0993 | 0.0291 | 0.2930 | 0.0838 | 0.0028100 |
| cluster3158 | -3370.8123 | 0.0126 | 0.0107 | 0.8520 | 0.0285 | 0.0279860 |

| | | | | | | |
|---|---|---|---|---|---|---|
| cluster3159 | -5807.6114 | 0.0407 | 0.0712 | 1.7497 | 0.3902 | 0.0149800 |
| cluster3160 | -1099.5349 | 0.0573 | 0.0101 | 0.1758 | 0.0177 | 0.00141 |
| cluster3161 | -4981.509 | 0.0308 | 0.0469 | 1.5227 | 0.0650 | 0.0057610 |
| cluster3162 | -3637.9885 | 0.0083 | 0.0038 | 0.4550 | 0.0530 | 0.0017610 |
| cluster3163 | -3429.9614 | 0.0352 | 0.0483 | 1.3707 | 0.0780 | 0.0056110 |
| cluster3164 | -6179.6654 | 0.0552 | 0.0658 | 1.1915 | 0.1042 | 0.0117530 |
| cluster3165 | -8049.1587 | 0.0535 | 0.0762 | 1.4241 | 0.0821 | 0.0174910 |
| cluster3166 | -4633.2183 | 0.0237 | 0.0145 | 0.6110 | 0.0321 | 0.0183110 |
| cluster3167 | -5313.4143 | 0.0916 | 0.0439 | 0.4792 | 0.1484 | 0.0050490 |
| cluster3168 | -7087.5253 | 0.0156 | 0.0166 | 1.0639 | 0.0301 | 0.0390280 |
| cluster3169 | -5029.6043 | 0.0292 | 0.0434 | 1.4860 | 0.0434 | 0.0067070 |
| cluster3170 | -1623.243 | 0.0133 | 0.0024 | 0.1789 | 0.0001 | 0.0000000 |
| cluster3171 | -6159.0255 | 0.0661 | 0.0404 | 0.6107 | 0.0501 | 0.0398740 |
| cluster3172 | -6261.9499 | 0.0268 | 0.0156 | 0.5822 | 0.0322 | 0.0322560 |
| cluster3173 | -4456.3486 | 0.0157 | 0.0168 | 1.0734 | 0.0001 | 0.0000130 |
| cluster3174 | -1301.2588 | 0.0021 | 0.0017 | 0.8168 | 0.0001 | 0.0000070 |
| cluster3175 | -1937.5836 | 0.0165 | 0.0107 | 0.6493 | 0.0001 | 0.0000070 |
| cluster3176 | -3886.5109 | 0.0264 | 0.0130 | 0.4920 | 0.0086 | 0.0017740 |
| cluster3177 | -3502.5805 | 0.0428 | 0.0195 | 0.4544 | 0.1179 | 0.0383760 |
| cluster3178 | -6736.0633 | 0.0086 | 0.0079 | 0.9197 | 0.0299 | 0.0296690 |
| cluster3179 | -4627.8566 | 0.0572 | 0.1176 | 2.0564 | 0.1937 | 0.0196410 |
| cluster3180 | -1071.2516 | 0.0033 | 0.0076 | 2.3294 | 0.0001 | 0.0000000 |
| cluster3181 | -6688.4199 | 0.0180 | 0.0160 | 0.8903 | 0.0412 | 0.0328980 |
| cluster3182 | -2530.385 | 0.0059 | 0.0019 | 0.3238 | 0.0086 | 0.0080540 |
| cluster3183 | -4034.1215 | 0.0644 | 0.0594 | 0.9225 | 0.2184 | 0.0793800 |
| cluster3184 | -4207.3955 | 0.0314 | 0.0168 | 0.5358 | 0.0539 | 0.0020350 |
| cluster3185 | -3544.7394 | 0.0154 | 0.0240 | 1.5560 | 0.0597 | 0.0062610 |
| cluster3186 | -2290.2778 | 0.0272 | 0.0210 | 0.7722 | 0.0572 | 0.0036770 |
| cluster3187 | -5393.5753 | 0.0210 | 0.0198 | 0.9425 | 0.0167 | 0.0023030 |
| cluster3188 | -4445.3035 | 0.0918 | 0.0372 | 0.4049 | 0.0843 | 0.0574340 |
| cluster3189 | -6229.1864 | 0.0416 | 0.0615 | 1.4780 | 0.0620 | 0.0072900 |
| cluster3190 | -5398.152 | 0.0315 | 0.0562 | 1.7813 | 0.0254 | 0.0045350 |
| cluster3191 | -2827.8247 | 0.0579 | 0.0286 | 0.4945 | 0.0194 | 0.0279750 |
| cluster3192 | -5887.1708 | 0.0511 | 0.0219 | 0.4283 | 0.0544 | 0.0410930 |
| cluster3193 | -3453.6141 | 0.0384 | 0.0181 | 0.4698 | 0.0001 | 0.0000080 |
| cluster3194 | -6980.4455 | 0.0544 | 0.0731 | 1.3427 | 0.0914 | 0.0353620 |
| cluster3195 | -3958.5135 | 0.0206 | 0.0173 | 0.8378 | 0.0083 | 0.0124140 |
| cluster3196 | -2243.5941 | 0.0061 | 0.0010 | 0.1636 | 0.0001 | 0.0000310 |
| cluster3197 | -5446.3218 | 0.0550 | 0.0225 | 0.4094 | 0.0385 | 0.0187970 |
| cluster3198 | -6431.4288 | 0.0236 | 0.0414 | 1.7596 | 0.0433 | 0.0060130 |
| cluster3199 | -2975.4161 | 0.0089 | 0.0062 | 0.6922 | 0.0674 | 0.0053780 |
| cluster3200 | -4262.2542 | 0.0170 | 0.0169 | 0.9955 | 0.0013 | 0.0022010 |
| cluster3201 | -4476.4468 | 0.1135 | 0.0338 | 0.2980 | 0.0771 | 0.0051450 |
| cluster3202 | -6071.4515 | 0.0429 | 0.0289 | 0.6729 | 0.0374 | 0.0032340 |

| | | | | | | |
|---|---|---|---|---|---|---|
| cluster3203 | -3017.4301 | 0.0491 | 0.0764 | 1.5573 | 0.0543 | 0.0071470 |
| cluster3204 | -1878.0714 | 0.0356 | 0.0071 | 0.1992 | 0.0484 | 0.0149020 |
| cluster3205 | -2014.7219 | 0.0076 | 0.0015 | 0.1971 | 0.0001 | 0.0000240 |
| cluster3206 | -3914.8318 | 0.0340 | 0.0366 | 1.0754 | 0.0374 | 0.0060230 |
| cluster3207 | -5496.7142 | 0.0171 | 0.0131 | 0.7708 | 0.0064 | 0.0132750 |
| cluster3208 | -3286.7042 | 0.0048 | 0.0020 | 0.4202 | 0.0054 | 0.0033470 |
| cluster3209 | -5550.2657 | 0.0220 | 0.0308 | 1.3978 | 0.0849 | 0.0058110 |
| cluster3210 | -3378.0114 | 0.0030 | 0.0121 | 3.9834 | 4.4760 | 0.0000020 |
| cluster3211 | -3434.1571 | 0.0301 | 0.0370 | 1.2284 | 0.2500 | 0.0034510 |
| cluster3212 | -2810.8345 | 0.0034 | 0.0008 | 0.2446 | 0.0009 | 0.0010480 |
| cluster3213 | -4614.1005 | 0.0862 | 0.0586 | 0.6804 | 0.1321 | 0.1082840 |
| cluster3214 | -6046.6803 | 0.0403 | 0.0220 | 0.5453 | 0.0797 | 0.0533120 |
| cluster3215 | -2279.293 | 0.0024 | 0.0005 | 0.1957 | 0.0001 | 0.0000020 |
| cluster3216 | -4536.2345 | 0.0275 | 0.0724 | 2.6314 | 0.1972 | 0.0071640 |
| cluster3217 | -3822.3714 | 0.0355 | 0.0149 | 0.4204 | 0.0649 | 0.0436090 |
| cluster3218 | -5082.4311 | 0.0225 | 0.0384 | 1.7060 | 0.0737 | 0.0063740 |
| cluster3219 | -5449.5024 | 0.0119 | 0.0141 | 1.1861 | 0.0246 | 0.0018880 |
| cluster3220 | -2923.7147 | 0.0265 | 0.0090 | 0.3397 | 0.0422 | 0.0297320 |
| cluster3221 | -5733.4042 | 0.0287 | 0.0541 | 1.8873 | 0.1466 | 0.0100430 |
| cluster3222 | -2619.8216 | 0.0332 | 0.0175 | 0.5252 | 0.0726 | 0.0281500 |
| cluster3223 | -3467.4742 | 0.0953 | 0.0757 | 0.7950 | 0.0957 | 0.0036950 |
| cluster3224 | -1597.2435 | 0.0001 | 0.0000 | 0.3157 | 0.0001 | 0.0000040 |
| cluster3225 | -6104.0647 | 0.0300 | 0.0366 | 1.2202 | 0.0681 | 0.0085960 |
| cluster3226 | -3722.234 | 0.0348 | 0.0328 | 0.9438 | 0.1309 | 0.0176890 |
| cluster3227 | -4397.0033 | 0.0643 | 0.0824 | 1.2808 | 0.1807 | 0.0170950 |
| cluster3228 | -3619.0135 | 0.0267 | 0.0451 | 1.6908 | 0.2064 | 0.0177070 |
| cluster3229 | -3870.2528 | 0.0313 | 0.0201 | 0.6399 | 0.0229 | 0.0163880 |
| cluster3230 | -3482.3036 | 0.0116 | 0.0049 | 0.4195 | 0.0092 | 0.0073840 |
| cluster3231 | -4527.6765 | 0.0260 | 0.0215 | 0.8295 | 0.0011 | 0.0001410 |
| cluster3232 | -5048.7816 | 0.0195 | 0.0278 | 1.4230 | 0.1230 | 0.0105350 |
| cluster3233 | -3436.5207 | 0.0411 | 0.0318 | 0.7724 | 0.0415 | 0.0683290 |
| cluster3234 | -3817.6802 | 0.0426 | 0.0243 | 0.5706 | 0.0866 | 0.0682980 |
| cluster3235 | -7271.9599 | 0.0379 | 0.0767 | 2.0239 | 0.0354 | 0.0077050 |
| cluster3236 | -3956.3144 | 0.0318 | 0.0419 | 1.3184 | 0.0559 | 0.0064530 |
| cluster3237 | -3244.7833 | 0.0582 | 0.0555 | 0.9535 | 0.0607 | 0.0144100 |
| cluster3238 | -2782.4673 | 0.0620 | 0.0215 | 0.3472 | 0.0001 | 0.0000060 |
| cluster3239 | -2738.6478 | 0.0197 | 0.0091 | 0.4619 | 0.0618 | 0.0069080 |
| cluster3240 | -6961.3917 | 0.0301 | 0.0210 | 0.6988 | 0.0400 | 0.0470960 |
| cluster3241 | -5756.4821 | 0.0653 | 0.0639 | 0.9776 | 0.0588 | 0.0131560 |
| cluster3242 | -7158.3774 | 0.0385 | 0.0772 | 2.0033 | 0.1231 | 0.0202460 |
| cluster3243 | -5984.1536 | 0.1562 | 0.0377 | 0.2413 | 0.1402 | 0.0827720 |
| cluster3244 | -6473.2028 | 0.0578 | 0.0340 | 0.5887 | 0.0684 | 0.0548730 |
| cluster3245 | -2585.832 | 0.0029 | 0.0033 | 1.1137 | 0.0001 | 0.0000030 |
| cluster3246 | -5134.3199 | 0.0402 | 0.0364 | 0.9049 | 0.0553 | 0.0790620 |

| | | | | | | |
|---|---|---|---|---|---|---|
| cluster3247 | -848.77847 | 0.0654 | 0.0444 | 0.6789 | 0.0001 | 0.0000040 |
| cluster3248 | -3482.1791 | 0.0186 | 0.0197 | 1.0582 | 0.0649 | 0.0109900 |
| cluster3249 | -3482.546 | 0.0159 | 0.0168 | 1.0572 | 0.0367 | 0.0039850 |
| cluster3250 | -3544.7875 | 0.0868 | 0.0323 | 0.3722 | 0.2972 | 0.0639980 |
| cluster3251 | -4467.0115 | 0.0554 | 0.0413 | 0.7463 | 0.1282 | 0.0071320 |
| cluster3252 | -7421.7064 | 0.0600 | 0.0972 | 1.6202 | 0.1094 | 0.0282330 |
| cluster3253 | -4317.6844 | 0.0701 | 0.0565 | 0.8051 | 0.0862 | 0.0078750 |
| cluster3254 | -3108.4504 | 0.0319 | 0.0315 | 0.9861 | 0.1233 | 0.0089940 |
| cluster3255 | -5568.1541 | 0.0156 | 0.0345 | 2.2198 | 0.0481 | 0.0099280 |
| cluster3256 | -6100.1098 | 0.0387 | 0.0371 | 0.9583 | 0.0642 | 0.0174270 |
| cluster3257 | -5143.9376 | 0.0056 | 0.0039 | 0.7066 | 0.0092 | 0.0099060 |
| cluster3258 | -1919.2438 | 0.0014 | 0.0007 | 0.5196 | 0.1747 | 0.0000010 |
| cluster3259 | -3556.5053 | 0.0353 | 0.0072 | 0.2027 | 0.0377 | 0.0177200 |
| cluster3260 | -4482.5159 | 0.0139 | 0.0070 | 0.5068 | 0.0134 | 0.0126430 |
| cluster3261 | -4945.3287 | 0.0630 | 0.1023 | 1.6243 | 0.3411 | 0.0143380 |
| cluster3262 | -5791.4375 | 0.0182 | 0.0300 | 1.6464 | 0.0234 | 0.0039390 |
| cluster3263 | -2055.3621 | 0.1316 | 0.0675 | 0.5130 | 0.0001 | 0.00000 |
| cluster3264 | -1835.1655 | 0.0104 | 0.0040 | 0.3828 | 0.0001 | 0.0000020 |
| cluster3265 | -4128.5792 | 0.0028 | 0.0027 | 0.9592 | 0.0086 | 0.0006950 |
| cluster3266 | -2855.1819 | 0.0311 | 0.0109 | 0.3524 | 0.0001 | 0.0000060 |
| cluster3267 | -4124.5745 | 0.0298 | 0.0196 | 0.6568 | 0.0414 | 0.0237580 |
| cluster3268 | -2803.0124 | 0.0316 | 0.0256 | 0.8097 | 0.1205 | 0.0205780 |
| cluster3269 | -3538.3937 | 0.0571 | 0.0501 | 0.8778 | 0.0619 | 0.0065460 |
| cluster3270 | -2185.3898 | 0.0262 | 0.0345 | 1.3173 | 0.0242 | 0.0020970 |
| cluster3271 | -4111.1439 | 0.0105 | 0.0059 | 0.5595 | 0.0223 | 0.0096670 |
| cluster3272 | -939.22097 | 0.3182 | 0.0264 | 0.0829 | 0.0001 | 0.00000 |
| cluster3273 | -4003.6964 | 0.1069 | 0.1122 | 1.0504 | 0.2634 | 0.0259520 |
| cluster3274 | -2030.5037 | 0.0276 | 0.0460 | 1.6658 | 0.1440 | 0.0146450 |
| cluster3275 | -4551.2003 | 0.0482 | 0.0565 | 1.1726 | 0.0522 | 0.0059900 |
| cluster3276 | -2485.4348 | 0.0952 | 0.0155 | 0.1623 | 0.0741 | 0.0208990 |
| cluster3277 | -3233.8507 | 0.0464 | 0.0661 | 1.4231 | 124.6987 | 0.0064400 |
| cluster3278 | -4576.2479 | 0.0029 | 0.0013 | 0.4423 | 0.0269 | 0.0006260 |
| cluster3279 | -3730.929 | 0.0175 | 0.0211 | 1.2058 | 0.0001 | 0.0000130 |
| cluster3280 | -3787.095 | 0.0231 | 0.0146 | 0.6334 | 0.0001 | 0.0000070 |
| cluster3281 | -1963.969 | 0.0174 | 0.0235 | 1.3553 | 0.0911 | 0.0137200 |
| cluster3282 | -2049.1933 | 0.0057 | 0.0042 | 0.7358 | 0.0042 | 0.0003880 |
| cluster3283 | -5464.8796 | 0.0651 | 0.0358 | 0.5498 | 0.0617 | 0.0545150 |
| cluster3284 | -5978.4681 | 0.0327 | 0.0331 | 1.0127 | 0.1071 | 0.0057890 |
| cluster3285 | -2362.5459 | 0.0076 | 0.0035 | 0.4587 | 0.0001 | 0.0000060 |
| cluster3286 | -2209.0265 | 0.0294 | 0.0114 | 0.3874 | 0.0514 | 0.0020620 |
| cluster3287 | -1377.3891 | 0.0126 | 0.0089 | 0.7068 | 0.0187 | 0.0018610 |
| cluster3288 | -1238.2688 | 0.0051 | 0.0100 | 1.9582 | 0.0491 | 0.0076550 |
| cluster3289 | -5798.8425 | 0.0480 | 0.0158 | 0.3286 | 0.0468 | 0.0198350 |
| cluster3290 | -1251.4819 | 0.0169 | 0.0104 | 0.6136 | 0.0662 | 0.0195410 |

| | | | | | | |
|---|---|---|---|---|---|---|
| cluster3291 | -4787.7697 | 0.0390 | 0.0273 | 0.7012 | 0.0429 | 0.0381300 |
| cluster3292 | -2756.7683 | 0.0176 | 0.0121 | 0.6903 | 0.3575 | 0.0203700 |
| cluster3293 | -3183.4426 | 0.0080 | 0.0071 | 0.8802 | 0.0195 | 0.0131390 |
| cluster3294 | -2880.4499 | 0.0103 | 0.0061 | 0.5929 | 0.0292 | 0.0279200 |
| cluster3295 | -2241.1094 | 0.0017 | 0.0008 | 0.4892 | 0.0001 | 0.0000080 |
| cluster3296 | -1799.5658 | 0.0041 | 0.0055 | 1.3334 | 0.0097 | 0.0014700 |
| cluster3297 | -2354.5038 | 0.1463 | 0.0679 | 0.4638 | 0.2472 | 0.10511 |
| cluster3298 | -2941.7989 | 0.0053 | 0.0037 | 0.7019 | 0.0007 | 0.0009430 |
| cluster3299 | -7905.0756 | 0.0251 | 0.1351 | 5.3902 | 999.0000 | 0.0116560 |
| cluster3300 | -2254.7486 | 0.0045 | 0.0052 | 1.1456 | 0.4282 | 0.0000010 |
| cluster3301 | -1801.7864 | 0.0390 | 0.0143 | 0.3656 | 0.0971 | 0.0384480 |
| cluster3302 | -4560.5995 | 0.0860 | 0.0582 | 0.6762 | 0.0995 | 0.0208400 |
| cluster3303 | -4480.5639 | 0.0346 | 0.0349 | 1.0089 | 0.0411 | 0.0624400 |
| cluster3304 | -2855.1735 | 0.0318 | 0.0105 | 0.3304 | 0.0054 | 0.0033380 |
| cluster3305 | -3805.2847 | 0.0627 | 0.0604 | 0.9635 | 0.0753 | 0.0107180 |
| cluster3306 | -5020.6581 | 0.0681 | 0.0735 | 1.0790 | 0.1228 | 0.0145500 |
| cluster3307 | -1671.302 | 0.0047 | 0.0051 | 1.0900 | 0.0001 | 0.0000070 |
| cluster3308 | -4387.625 | 0.0171 | 0.0164 | 0.9625 | 0.0225 | 0.0282590 |
| cluster3309 | -3384.1874 | 0.1286 | 0.0419 | 0.3260 | 0.0842 | 0.0371730 |
| cluster3310 | -4174.1512 | 0.0177 | 0.0116 | 0.6546 | 0.0095 | 0.0066510 |
| cluster3311 | -6153.519 | 0.0472 | 0.0328 | 0.6941 | 0.0977 | 0.0757290 |
| cluster3312 | -1926.3644 | 0.0241 | 0.0386 | 1.5982 | 999.0000 | 0.0115040 |
| cluster3313 | -2505.1616 | 0.0066 | 0.0018 | 0.2792 | 0.0257 | 0.0149430 |
| cluster3314 | -7138.7351 | 0.0363 | 0.0955 | 2.6319 | 0.1177 | 0.0142430 |
| cluster3315 | -3722.4938 | 0.0067 | 0.0036 | 0.5404 | 0.0037 | 0.0047010 |
| cluster3316 | -5969.4495 | 0.0763 | 0.0876 | 1.1480 | 0.3799 | 0.0174210 |
| cluster3317 | -1143.1797 | 0.0030 | 0.0024 | 0.7937 | 0.5637 | 0.0000010 |
| cluster3318 | -3441.3424 | 0.1151 | 0.0935 | 0.8128 | 0.0768 | 0.1579460 |
| cluster3319 | -3765.9331 | 0.0317 | 0.0470 | 1.4794 | 0.2127 | 0.0103320 |
| cluster3320 | -2813.8777 | 0.0538 | 0.0584 | 1.0848 | 0.0937 | 0.0047760 |
| cluster3321 | -2913.084 | 0.0014 | 0.0009 | 0.6813 | 0.0844 | 0.0026490 |
| cluster3322 | -3277.2483 | 0.0372 | 0.0562 | 1.5110 | 0.0971 | 0.0186450 |
| cluster3323 | -5518.7582 | 0.0464 | 0.0458 | 0.9887 | 0.0998 | 0.0181330 |
| cluster3324 | -1399.7753 | 0.0418 | 0.0345 | 0.8244 | 0.0138 | 0.0125290 |
| cluster3325 | -2061.1587 | 0.0188 | 0.0208 | 1.1097 | 0.0181 | 0.0053190 |
| cluster3326 | -1704.4543 | 0.0736 | 0.0186 | 0.2532 | 0.0001 | 0.00000 |
| cluster3327 | -3863.0404 | 0.0216 | 0.0135 | 0.6240 | 0.0390 | 0.0336170 |
| cluster3328 | -2608.094 | 0.0114 | 0.0173 | 1.5161 | 3.1567 | 0.0000010 |
| cluster3329 | -3130.3686 | 0.0394 | 0.0752 | 1.9086 | 0.4379 | 0.1150640 |
| cluster3330 | -4709.8278 | 0.0443 | 0.0633 | 1.4315 | 0.1317 | 0.0070890 |
| cluster3331 | -2866.0844 | 0.0668 | 0.0303 | 0.4532 | 0.2085 | 0.0139470 |
| cluster3332 | -6185.2943 | 0.0556 | 0.0857 | 1.5424 | 0.2428 | 0.0244920 |
| cluster3333 | -3572.7414 | 0.0203 | 0.0284 | 1.3985 | 0.0746 | 0.0112320 |
| cluster3334 | -1377.8498 | 0.0608 | 0.0315 | 0.5178 | 0.3071 | 0.0218970 |

| | | | | | | |
|---|---|---|---|---|---|---|
| cluster3335 | -2854.4916 | 0.0233 | 0.0144 | 0.6176 | 0.0711 | 0.0838060 |
| cluster3336 | -3027.9687 | 0.0034 | 0.0013 | 0.3712 | 0.0052 | 0.0026800 |
| cluster3337 | -2619.1111 | 0.0351 | 0.0088 | 0.2495 | 0.0160 | 0.0058240 |
| cluster3338 | -6296.6007 | 0.0330 | 0.0674 | 2.0433 | 0.2656 | 0.0163950 |
| cluster3339 | -1909.6697 | 0.0407 | 0.0318 | 0.7821 | 0.0770 | 0.0835660 |
| cluster3340 | -1627.4506 | 0.1695 | 0.0307 | 0.1813 | 0.1235 | 0.00345 |
| cluster3341 | -1576.959 | 0.0107 | 0.0177 | 1.6630 | 0.0161 | 0.0020110 |
| cluster3342 | -2554.7677 | 0.0193 | 0.0302 | 1.5683 | 0.2456 | 0.0052270 |
| cluster3343 | -2926.5134 | 0.0005 | 0.0005 | 1.0462 | 0.0001 | 0.0000070 |
| cluster3344 | -3149.1046 | 0.0251 | 0.0089 | 0.3555 | 0.0331 | 0.0133120 |
| cluster3345 | -2794.6965 | 0.0119 | 0.0202 | 1.6881 | 0.0506 | 0.0044470 |
| cluster3346 | -3608.5599 | 0.0027 | 0.0019 | 0.7091 | 0.0084 | 0.0052240 |
| cluster3347 | -4176.129 | 0.0465 | 0.0182 | 0.3920 | 0.0477 | 0.0266750 |
| cluster3348 | -4559.4806 | 0.0520 | 0.0256 | 0.4932 | 0.0319 | 0.0048890 |
| cluster3349 | -4211.0927 | 0.0119 | 0.0080 | 0.6767 | 0.0024 | 0.0087810 |
| cluster3350 | -4772.9048 | 0.0437 | 0.0532 | 1.2154 | 0.1852 | 0.0202770 |
| cluster3351 | -3133.472 | 0.0151 | 0.0156 | 1.0332 | 0.0412 | 0.0032210 |
| cluster3352 | -4084.9312 | 0.0549 | 0.0366 | 0.6676 | 0.0923 | 0.0057920 |
| cluster3353 | -1585.7928 | 0.0112 | 0.0059 | 0.5275 | 0.0001 | 0.0000060 |
| cluster3354 | -2117.9497 | 0.0038 | 0.0016 | 0.4207 | 0.0760 | 0.0024750 |
| cluster3355 | -4280.0183 | 0.0350 | 0.0349 | 0.9993 | 0.1186 | 0.0350310 |
| cluster3356 | -4595.6028 | 0.0380 | 0.0168 | 0.4417 | 0.0489 | 0.0359130 |
| cluster3357 | -1289.9447 | 0.0267 | 0.0180 | 0.6732 | 0.0518 | 0.0045260 |
| cluster3358 | -4168.4654 | 0.0006 | 0.0003 | 0.4248 | 0.0051 | 0.0033900 |
| cluster3359 | -6041.9676 | 0.0351 | 0.0555 | 1.5793 | 0.0750 | 0.0125370 |
| cluster3360 | -2470.8802 | 0.0341 | 0.0244 | 0.7152 | 0.0556 | 0.0569960 |
| cluster3361 | -5035.3956 | 0.0364 | 0.0620 | 1.7039 | 0.0477 | 0.0058270 |
| cluster3362 | -3692.7218 | 0.0167 | 0.0109 | 0.6566 | 0.0140 | 0.0107600 |
| cluster3363 | -3721.1671 | 0.0023 | 0.0071 | 3.0372 | 1.6664 | 0.0000010 |
| cluster3364 | -4356.0572 | 0.0229 | 0.0401 | 1.7504 | 0.0374 | 0.0053230 |
| cluster3365 | -5561.5704 | 0.0576 | 0.0574 | 0.9952 | 0.2779 | 0.0117880 |
| cluster3366 | -3140.1788 | 0.0244 | 0.0630 | 2.5839 | 0.5978 | 0.0209530 |
| cluster3367 | -3690.0802 | 0.1553 | 0.0797 | 0.5130 | 0.0681 | 0.0469400 |
| cluster3368 | -1654.4337 | 0.0292 | 0.0056 | 0.1931 | 0.0153 | 0.0034400 |
| cluster3369 | -3764.3506 | 0.0140 | 0.0193 | 1.3756 | 0.0422 | 0.0070030 |
| cluster3370 | -6384.9217 | 0.0918 | 0.0822 | 0.8945 | 0.2095 | 0.0221050 |
| cluster3371 | -2934.3639 | 0.0063 | 0.0030 | 0.4848 | 0.0001 | 0.0000020 |
| cluster3372 | -3840.8422 | 0.0077 | 0.0062 | 0.8045 | 0.0117 | 0.0214740 |
| cluster3373 | -5277.2232 | 0.0087 | 0.0142 | 1.6336 | 0.0023 | 0.0003540 |
| cluster3374 | -5122.4187 | 0.0647 | 0.1267 | 1.9580 | 0.2002 | 0.0209130 |
| cluster3375 | -3051.5935 | 0.0067 | 0.0017 | 0.2578 | 0.0031 | 0.0026730 |
| cluster3376 | -1519.0529 | 0.0078 | 0.0014 | 0.1808 | 0.1023 | 0.0471450 |
| cluster3377 | -2273.1177 | 0.0108 | 0.0166 | 1.5425 | 0.0567 | 0.0038920 |
| cluster3378 | -6508.0468 | 0.0469 | 0.0388 | 0.8273 | 0.0528 | 0.0637910 |

| | | | | | | |
|---|---|---|---|---|---|---|
| cluster3379 | -1568.3418 | 0.0349 | 0.0238 | 0.6811 | 0.0001 | 0.0000050 |
| cluster3380 | -2653.744 | 0.0137 | 0.0198 | 1.4457 | 6.2250 | 0.0003560 |
| cluster3381 | -2267.9889 | 0.0330 | 0.0172 | 0.5231 | 0.0361 | 0.0346760 |
| cluster3382 | -885.72899 | 0.0065 | 0.0031 | 0.4822 | 0.0001 | 0.0000010 |
| cluster3383 | -5145.4422 | 0.0459 | 0.0814 | 1.7728 | 0.5216 | 0.0229050 |
| cluster3384 | -1972.2526 | 0.0326 | 0.0187 | 0.5726 | 0.0384 | 0.0234360 |
| cluster3385 | -3771.0771 | 0.0209 | 0.0278 | 1.3339 | 0.0420 | 0.0065870 |
| cluster3386 | -4155.5925 | 0.0139 | 0.0189 | 1.3548 | 0.0309 | 0.0022600 |
| cluster3387 | -3029.9032 | 0.0387 | 0.0289 | 0.7477 | 0.0767 | 0.0197210 |
| cluster3388 | -2807.5333 | 0.0225 | 0.0180 | 0.8017 | 0.0591 | 0.0111260 |
| cluster3389 | -1788.791 | 0.0105 | 0.0147 | 1.4040 | 0.0001 | 0.0000120 |
| cluster3390 | -5115.642 | 0.0586 | 0.0327 | 0.5579 | 0.0890 | 0.0504450 |
| cluster3391 | -2414.0498 | 0.0070 | 0.0042 | 0.5973 | 0.0453 | 0.0034570 |
| cluster3392 | -2528.4239 | 0.0103 | 0.0147 | 1.4230 | 0.0082 | 0.0017130 |
| cluster3393 | -3179.4013 | 0.0560 | 0.0497 | 0.8870 | 0.0765 | 0.0084880 |
| cluster3394 | -713.50683 | 0.0044 | 0.0023 | 0.5380 | 0.0022 | 0.0039380 |
| cluster3395 | -2714.0407 | 0.0132 | 0.0069 | 0.5212 | 0.0197 | 0.0231720 |
| cluster3396 | -4984.6917 | 0.0391 | 0.0182 | 0.4653 | 0.0271 | 0.0225850 |
| cluster3397 | -4641.0957 | 0.0266 | 0.0395 | 1.4856 | 0.0639 | 0.0068350 |
| cluster3398 | -3833.2922 | 0.0539 | 0.0323 | 0.5998 | 0.0959 | 0.0056700 |
| cluster3399 | -3230.6229 | 0.0428 | 0.0295 | 0.6885 | 0.0498 | 0.0601610 |
| cluster3400 | -1944.629 | 0.1379 | 0.0256 | 0.1855 | 0.0060 | 0.00144 |
| cluster3401 | -2870.6045 | 0.0166 | 0.0166 | 0.9968 | 0.0001 | 0.0000030 |
| cluster3402 | -5114.9705 | 0.0241 | 0.0155 | 0.6428 | 0.0187 | 0.0138990 |
| cluster3403 | -3669.0974 | 0.0210 | 0.0167 | 0.7953 | 0.0331 | 0.0086930 |
| cluster3404 | -716.69613 | 0.0001 | 0.0000 | 0.0932 | 0.0001 | 0.00001 |
| cluster3405 | -1541.0534 | 0.1245 | 0.0554 | 0.4447 | 0.0592 | 0.00359 |
| cluster3406 | -3800.5688 | 0.0182 | 0.0149 | 0.8170 | 0.0211 | 0.0271730 |
| cluster3407 | -6244.045 | 0.0630 | 0.0506 | 0.8026 | 0.0502 | 0.0837260 |
| cluster3408 | -1896.6784 | 0.0060 | 0.0018 | 0.2976 | 0.0075 | 0.0014310 |
| cluster3409 | -4909.1042 | 0.0221 | 0.0363 | 1.6457 | 0.0363 | 0.0074800 |
| cluster3410 | -1442.9568 | 0.0263 | 0.0065 | 0.2472 | 0.0185 | 0.0094970 |
| cluster3411 | -2582.9183 | 0.0003 | 0.0003 | 0.7837 | 0.0001 | 0.0000040 |
| cluster3412 | -3391.7819 | 0.0808 | 0.0734 | 0.9080 | 0.1522 | 0.0089240 |
| cluster3413 | -2548.0391 | 0.0330 | 0.0373 | 1.1328 | 0.9072 | 0.0308930 |
| cluster3414 | -2522.9665 | 0.1357 | 0.0256 | 0.1890 | 0.1361 | 0.04755 |
| cluster3415 | -3492.479 | 0.0605 | 0.0346 | 0.5724 | 0.1240 | 0.0083190 |
| cluster3416 | -600.79495 | 0.0010 | 0.0181 | 18.0245 | 0.1626 | 1.6097910 |
| cluster3417 | -1214.9697 | 0.0018 | 0.0150 | 8.4922 | 0.0003 | 0.0177830 |
| cluster3418 | -2534.1558 | 0.0534 | 0.0482 | 0.9036 | 0.0675 | 0.0189780 |
| cluster3419 | -2669.6058 | 0.0862 | 0.0304 | 0.3523 | 0.3325 | 0.01823 |
| cluster3420 | -1781.5122 | 0.2052 | 0.0988 | 0.4814 | 0.6383 | 0.11881 |
| cluster3421 | -4207.7173 | 0.0111 | 0.0162 | 1.4692 | 0.0148 | 0.0019890 |
| cluster3422 | -5398.885 | 0.0369 | 0.0223 | 0.6047 | 0.0267 | 0.0166470 |

| | | | | | | |
|---|---|---|---|---|---|---|
| cluster3423 | -4242.8275 | 0.0393 | 0.0843 | 2.1473 | 0.5953 | 0.0203550 |
| cluster3424 | -2855.6287 | 0.0929 | 0.0533 | 0.5735 | 0.1479 | 0.0837750 |
| cluster3425 | -2232.4189 | 0.0452 | 0.0181 | 0.4006 | 0.0001 | 0.0000020 |
| cluster3426 | -3411.0645 | 0.0192 | 0.0195 | 1.0181 | 0.0142 | 0.0196670 |
| cluster3427 | -3796.7359 | 0.0936 | 0.0567 | 0.6059 | 0.0771 | 0.0071860 |
| cluster3428 | -2685.5869 | 0.0689 | 0.0213 | 0.3098 | 0.0527 | 0.0446540 |
| cluster3429 | -4145.3803 | 0.0269 | 0.0201 | 0.7462 | 0.0353 | 0.0298560 |
| cluster3430 | -4824.2996 | 0.0417 | 0.0764 | 1.8318 | 0.0639 | 0.0140570 |
| cluster3431 | -4739.1827 | 0.0418 | 0.0628 | 1.5024 | 0.0480 | 0.0130510 |
| cluster3432 | -3782.5917 | 0.0353 | 0.0143 | 0.4065 | 0.0429 | 0.0247460 |
| cluster3433 | -5393.2811 | 0.1649 | 0.0658 | 0.3987 | 0.1977 | 0.0477340 |
| cluster3434 | -3292.0516 | 0.0435 | 0.0145 | 0.3326 | 0.0162 | 0.0095480 |
| cluster3435 | -4091.1558 | 0.0193 | 0.0215 | 1.1161 | 0.0719 | 0.0075550 |
| cluster3436 | -3134.4288 | 0.0225 | 0.0345 | 1.5313 | 0.0721 | 0.0042210 |
| cluster3437 | -2816.2145 | 0.0120 | 0.0096 | 0.7975 | 0.0175 | 0.0225300 |
| cluster3438 | -3824.2207 | 0.0362 | 0.0300 | 0.8290 | 0.0651 | 0.0104770 |
| cluster3439 | -2056.4363 | 0.0517 | 0.0480 | 0.9284 | 0.0742 | 0.0044280 |
| cluster3440 | -2313.7781 | 0.0238 | 0.0116 | 0.4849 | 999.0000 | 0.0021660 |
| cluster3441 | -2756.4059 | 0.0362 | 0.0517 | 1.4270 | 0.1266 | 0.0048710 |
| cluster3442 | -2923.6185 | 0.0182 | 0.0584 | 3.2113 | 0.0510 | 0.0032100 |
| cluster3443 | -3528.2926 | 0.0132 | 0.0185 | 1.4073 | 0.1760 | 0.0057100 |
| cluster3444 | -3384.1074 | 0.0088 | 0.0073 | 0.8320 | 0.0539 | 0.0037060 |
| cluster3445 | -2964.8956 | 0.0152 | 0.0056 | 0.3670 | 0.0110 | 0.0067300 |
| cluster3446 | -4060.066 | 0.0279 | 0.0413 | 1.4789 | 0.6150 | 0.0063340 |
| cluster3447 | -5903.7381 | 0.0249 | 0.0724 | 2.9131 | 4.5036 | 0.0293000 |
| cluster3448 | -2378.1881 | 0.0239 | 0.0365 | 1.5257 | 0.0001 | 0.0000070 |
| cluster3449 | -613.4533 | 0.0043 | 0.0006 | 0.1325 | 0.0001 | 0.0000210 |
| cluster3450 | -2517.7165 | 0.0186 | 0.0328 | 1.7615 | 0.0764 | 0.0050180 |
| cluster3451 | -3594.1384 | 0.0268 | 0.0356 | 1.3261 | 0.0365 | 0.0049950 |
| cluster3452 | -3964.5425 | 0.0462 | 0.0559 | 1.2120 | 0.0875 | 0.0050690 |
| cluster3453 | -1848.0377 | 0.0458 | 0.0183 | 0.3993 | 0.0192 | 0.0157870 |
| cluster3454 | -3695.8427 | 0.0103 | 0.0220 | 2.1297 | 0.0141 | 0.0051120 |
| cluster3455 | -3648.9256 | 0.0400 | 0.0085 | 0.2118 | 0.0524 | 0.0209140 |
| cluster3456 | -4231.5564 | 0.0366 | 0.0172 | 0.4686 | 0.0367 | 0.0073210 |
| cluster3457 | -3649.0761 | 0.0320 | 0.0361 | 1.1292 | 0.0183 | 0.0042580 |
| cluster3458 | -1806.4083 | 0.0320 | 0.0136 | 0.4253 | 0.0417 | 0.0294440 |
| cluster3459 | -2636.1634 | 0.0366 | 0.0702 | 1.9213 | 464.4545 | 0.0132800 |
| cluster3460 | -5573.5775 | 0.0260 | 0.0592 | 2.2810 | 0.1422 | 0.0180360 |
| cluster3461 | -3098.5429 | 0.0560 | 0.0668 | 1.1912 | 0.2947 | 0.0226980 |
| cluster3462 | -821.47962 | 0.0194 | 0.0021 | 0.1092 | 0.2090 | 0.00906 |
| cluster3463 | -4771.3766 | 0.0481 | 0.0526 | 1.0923 | 0.0417 | 0.0088960 |
| cluster3464 | -5603.184 | 0.0405 | 0.0767 | 1.8926 | 0.1597 | 0.0181070 |
| cluster3465 | -3955.1365 | 0.0084 | 0.0161 | 1.9139 | 0.0158 | 0.0250460 |
| cluster3466 | -2643.9757 | 0.0134 | 0.0094 | 0.7020 | 0.0087 | 0.0186040 |

| | | | | | | |
|---|---|---|---|---|---|---|
| cluster3467 | -4059.9368 | 0.0312 | 0.0285 | 0.9116 | 0.0686 | 0.0072960 |
| cluster3468 | -1849.9112 | 0.0078 | 0.0167 | 2.1359 | 0.0050 | 0.0008600 |
| cluster3469 | -2102.2856 | 0.0251 | 0.0233 | 0.9277 | 0.0001 | 0.0000290 |
| cluster3470 | -3883.7309 | 0.0123 | 0.0161 | 1.3034 | 0.0433 | 0.0038050 |
| cluster3471 | -4169.823 | 0.0385 | 0.0410 | 1.0647 | 0.1081 | 0.0162460 |
| cluster3472 | -3722.0417 | 0.0182 | 0.0107 | 0.5857 | 0.0163 | 0.0114690 |
| cluster3473 | -3470.0881 | 0.0248 | 0.0141 | 0.5662 | 0.0375 | 0.0432560 |
| cluster3474 | -5365.9869 | 0.0286 | 0.0736 | 2.5733 | 0.0689 | 0.0069040 |
| cluster3475 | -2340.2615 | 0.0135 | 0.0175 | 1.2990 | 0.0049 | 0.0020170 |
| cluster3476 | -5103.9351 | 0.0230 | 0.0715 | 3.1058 | 999.0000 | 0.0125720 |
| cluster3477 | -3274.0653 | 0.0202 | 0.0043 | 0.2128 | 0.0040 | 0.0017220 |
| cluster3478 | -3581.7083 | 0.0180 | 0.0095 | 0.5272 | 0.0239 | 0.0242790 |
| cluster3479 | -2914.1471 | 0.0393 | 0.0891 | 2.2677 | 0.2639 | 0.0154620 |
| cluster3480 | -4508.0099 | 0.0315 | 0.0412 | 1.3057 | 0.0719 | 0.0077770 |
| cluster3481 | -1071.5868 | 0.0086 | 0.0028 | 0.3300 | 0.0015 | 0.1163640 |
| cluster3482 | -1966.8791 | 0.0795 | 0.0243 | 0.3057 | 0.0001 | 0.0000030 |
| cluster3483 | -3654.4462 | 0.0083 | 0.0091 | 1.0967 | 0.0536 | 0.0068140 |
| cluster3484 | -2769.2117 | 0.0294 | 0.0387 | 1.3136 | 0.0664 | 0.0043280 |
| cluster3485 | -3494.0858 | 0.0394 | 0.0385 | 0.9772 | 0.1237 | 0.0050170 |
| cluster3486 | -3175.0971 | 0.0497 | 0.0159 | 0.3202 | 0.0537 | 0.0235420 |
| cluster3487 | -2348.178 | 0.0232 | 0.0205 | 0.8814 | 0.0307 | 0.0113650 |
| cluster3488 | -1482.8891 | 0.0255 | 0.0161 | 0.6310 | 0.0493 | 0.0088240 |
| cluster3489 | -4054.4899 | 0.0376 | 0.0536 | 1.4257 | 0.2517 | 0.0259310 |
| cluster3490 | -3430.4879 | 0.0193 | 0.0393 | 2.0328 | 0.1066 | 0.0074290 |
| cluster3491 | -3687.1712 | 0.0319 | 0.0312 | 0.9780 | 0.0252 | 0.0024500 |
| cluster3492 | -3371.3595 | 0.0388 | 0.0563 | 1.4502 | 0.0001 | 0.0000100 |
| cluster3493 | -4433.3656 | 0.0347 | 0.0304 | 0.8762 | 0.0308 | 0.0025030 |
| cluster3494 | -2865.944 | 0.0135 | 0.0218 | 1.6125 | 0.0162 | 0.0476870 |
| cluster3495 | -2839.3571 | 0.0253 | 0.0237 | 0.9375 | 0.1375 | 0.0104190 |
| cluster3496 | -3231.6312 | 0.0232 | 0.0120 | 0.5174 | 0.0189 | 0.0140750 |
| cluster3497 | -1194.6737 | 0.0161 | 0.0021 | 0.1288 | 0.0086 | 0.0029580 |
| cluster3498 | -4133.8721 | 0.0470 | 0.0900 | 1.9145 | 0.0766 | 0.0081120 |
| cluster3499 | -3718.7189 | 0.0749 | 0.1102 | 1.4716 | 0.4625 | 0.0262260 |
| cluster3500 | -3421.7195 | 0.0179 | 0.0242 | 1.3499 | 0.1120 | 0.0062460 |
| cluster3501 | -2359.576 | 0.0254 | 0.0130 | 0.5109 | 0.0315 | 0.0410730 |
| cluster3502 | -2165.8572 | 0.0117 | 0.0166 | 1.4252 | 0.0675 | 0.0114300 |
| cluster3503 | -1952.2781 | 0.0451 | 0.0597 | 1.3235 | 0.0563 | 0.0114230 |
| cluster3504 | -1141.7751 | 0.0092 | 0.0107 | 1.1690 | 0.0048 | 0.0220300 |
| cluster3505 | -2413.2986 | 0.0390 | 0.0814 | 2.0871 | 0.3308 | 0.0321050 |
| cluster3506 | -2548.0948 | 0.0216 | 0.0276 | 1.2795 | 0.0919 | 0.0071790 |
| cluster3507 | -5015.2324 | 0.0480 | 0.0605 | 1.2603 | 0.2245 | 0.0169040 |
| cluster3508 | -5634.0562 | 0.0286 | 0.0883 | 3.0877 | 0.0379 | 0.0120470 |
| cluster3509 | -2402.0276 | 0.0012 | 0.0268 | 22.5950 | 29.5195 | 0.0000020 |
| cluster3510 | -843.75645 | 0.1172 | 0.0510 | 0.4351 | 0.0433 | 0.00776 |

| | | | | | | |
|---|---|---|---|---|---|---|
| cluster3511 | -1018.2814 | 0.0296 | 0.0239 | 0.8063 | 0.2149 | 0.0056070 |
| cluster3512 | -2356.2959 | 0.0315 | 0.0167 | 0.5291 | 0.0997 | 0.0029040 |
| cluster3513 | -959.09104 | 0.0616 | 0.0050 | 0.0812 | 0.1058 | 0.00890 |
| cluster3514 | -5158.5389 | 0.0348 | 0.0215 | 0.6176 | 0.0448 | 0.0325980 |
| cluster3515 | -2605.3311 | 0.0135 | 0.0127 | 0.9399 | 0.0115 | 0.0018980 |
| cluster3516 | -4129.8459 | 0.0262 | 0.0282 | 1.0788 | 0.0551 | 0.0054490 |
| cluster3517 | -2412.9843 | 0.0024 | 0.0014 | 0.5668 | 4.5089 | 0.0000020 |
| cluster3518 | -4783.6424 | 0.0719 | 0.0849 | 1.1811 | 0.1376 | 0.0179570 |
| cluster3519 | -2127.2029 | 0.0422 | 0.0399 | 0.9452 | 0.0383 | 0.0079790 |
| cluster3520 | -1193.9729 | 0.1515 | 0.0736 | 0.4862 | 0.1278 | 0.01197 |
| cluster3521 | -4219.9066 | 0.0325 | 0.0514 | 1.5809 | 0.1185 | 0.0174230 |
| cluster3522 | -4752.7947 | 0.0311 | 0.0587 | 1.8847 | 0.0773 | 0.0085810 |
| cluster3523 | -1653.9448 | 0.0241 | 0.0351 | 1.4572 | 0.1377 | 0.0195680 |
| cluster3524 | -1963.4128 | 0.0121 | 0.0051 | 0.4208 | 0.0227 | 0.0014050 |
| cluster3525 | -2816.8575 | 0.0440 | 0.0267 | 0.6074 | 0.0700 | 0.0069520 |
| cluster3526 | -2998.5473 | 0.0018 | 0.0039 | 2.2437 | 0.9075 | 0.0013410 |
| cluster3527 | -4935.1893 | 0.0426 | 0.0828 | 1.9437 | 0.1267 | 0.0166660 |
| cluster3528 | -1962.6714 | 0.0046 | 0.0018 | 0.3836 | 0.0285 | 0.0212230 |
| cluster3529 | -3742.8532 | 0.0310 | 0.0546 | 1.7643 | 0.0346 | 0.0041110 |
| cluster3530 | -1954.1592 | 0.0289 | 0.0470 | 1.6284 | 0.0433 | 0.0072280 |
| cluster3531 | -2917.8839 | 0.0463 | 0.0397 | 0.8585 | 0.3025 | 0.0082480 |
| cluster3532 | -5557.4442 | 0.0589 | 0.0491 | 0.8331 | 0.0752 | 0.0839610 |
| cluster3533 | -2655.1962 | 0.0237 | 0.0084 | 0.3544 | 0.0272 | 0.0170320 |
| cluster3534 | -2560.0861 | 0.0703 | 0.0148 | 0.2104 | 0.0497 | 0.0216520 |
| cluster3535 | -3287.969 | 0.0189 | 0.0526 | 2.7818 | 0.2047 | 0.0062710 |
| cluster3536 | -3555.4521 | 0.0410 | 0.0196 | 0.4789 | 0.0217 | 0.0273590 |
| cluster3537 | -3664.5373 | 0.0062 | 0.0089 | 1.4242 | 0.0270 | 0.0022000 |
| cluster3538 | -1379.5361 | 0.0569 | 0.0156 | 0.2735 | 0.0001 | 0.0000530 |
| cluster3539 | -2945.292 | 0.0407 | 0.0971 | 2.3847 | 0.0278 | 0.0264920 |
| cluster3540 | -5363.0187 | 0.0845 | 0.1342 | 1.5889 | 0.1834 | 0.0212480 |
| cluster3541 | -3877.8378 | 0.0269 | 0.0162 | 0.6037 | 0.0289 | 0.0240890 |
| cluster3542 | -5427.0013 | 0.0572 | 0.1145 | 2.0013 | 0.0994 | 0.0120510 |
| cluster3543 | -4245.8338 | 0.0327 | 0.0698 | 2.1347 | 0.1051 | 0.0112400 |
| cluster3544 | -1088.7313 | 0.0152 | 0.0042 | 0.2742 | 0.0107 | 0.0056410 |
| cluster3545 | -5146.9636 | 0.0447 | 0.0701 | 1.5676 | 0.0723 | 0.0067290 |
| cluster3546 | -2679.7022 | 0.0216 | 0.0426 | 1.9731 | 0.0444 | 0.0049900 |
| cluster3547 | -1116.9149 | 0.0134 | 0.0013 | 0.0982 | 0.0055 | 0.0042650 |
| cluster3548 | -1295.6727 | 0.0101 | 0.0012 | 0.1171 | 0.1842 | 0.0480170 |
| cluster3549 | -3232.3461 | 0.0417 | 0.0238 | 0.5699 | 0.0256 | 0.0252580 |
| cluster3550 | -1223.9291 | 0.0087 | 0.0023 | 0.2670 | 0.0001 | 0.0000050 |
| cluster3551 | -2798.3319 | 0.0542 | 0.0161 | 0.2969 | 0.0337 | 0.0111690 |
| cluster3552 | -2382.8192 | 0.0195 | 0.0233 | 1.1977 | 0.0743 | 0.0562590 |
| cluster3553 | -2405.0562 | 0.0313 | 0.0252 | 0.8028 | 0.0596 | 0.0070720 |
| cluster3554 | -3824.3539 | 0.0263 | 0.0191 | 0.7259 | 0.0349 | 0.0015520 |

| | | | | | | |
|---|---|---|---|---|---|---|
| cluster3555 | -1630.3964 | 0.0170 | 0.0262 | 1.5403 | 0.1668 | 0.0172080 |
| cluster3556 | -4199.8068 | 0.0213 | 0.0349 | 1.6364 | 0.1418 | 0.0129110 |
| cluster3557 | -1366.2234 | 0.0717 | 0.1502 | 2.0948 | 0.0001 | 0.00001 |
| cluster3558 | -4777.0924 | 0.0649 | 0.0730 | 1.1244 | 0.0769 | 0.0311640 |
| cluster3559 | -2457.5823 | 0.0184 | 0.0234 | 1.2742 | 0.0001 | 0.0000050 |
| cluster3560 | -5091.7846 | 0.0520 | 0.0870 | 1.6751 | 0.2254 | 0.0217610 |
| cluster3561 | -1801.1772 | 0.0327 | 0.0138 | 0.4211 | 0.0001 | 0.0000060 |
| cluster3562 | -4278.2159 | 0.0372 | 0.0479 | 1.2899 | 0.0569 | 0.0057340 |
| cluster3563 | -1982.7829 | 0.0149 | 0.0076 | 0.5128 | 0.0147 | 0.0166450 |
| cluster3564 | -5024.1248 | 0.0695 | 0.0296 | 0.4257 | 0.0734 | 0.0613990 |
| cluster3565 | -3327.7557 | 0.0236 | 0.0451 | 1.9072 | 999.0000 | 0.0328510 |
| cluster3566 | -1871.5453 | 0.0253 | 0.0277 | 1.0982 | 0.4189 | 0.0457630 |
| cluster3567 | -2403.913 | 0.0653 | 0.0607 | 0.9288 | 0.0762 | 0.0206090 |
| cluster3568 | -3167.9343 | 0.0270 | 0.0466 | 1.7249 | 0.0634 | 0.0044430 |
| cluster3569 | -3548.3503 | 0.0359 | 0.0698 | 1.9427 | 0.5599 | 0.0113350 |
| cluster3570 | -2401.0514 | 0.0195 | 0.0548 | 2.8085 | 0.0215 | 0.0034420 |
| cluster3571 | -2030.7218 | 0.0139 | 0.0114 | 0.8235 | 0.0842 | 0.0620510 |
| cluster3572 | -3452.9723 | 0.0469 | 0.0614 | 1.3107 | 0.0848 | 0.0204190 |
| cluster3573 | -3244.1002 | 0.0124 | 0.0226 | 1.8240 | 0.0057 | 0.0012410 |
| cluster3574 | -5128.9415 | 0.0831 | 0.0974 | 1.1718 | 0.0208 | 0.0044210 |
| cluster3575 | -3831.1917 | 0.0397 | 0.0462 | 1.1646 | 0.0396 | 0.0073200 |
| cluster3576 | -3274.2404 | 0.0388 | 0.0516 | 1.3327 | 0.0001 | 0.0000110 |
| cluster3577 | -2731.2323 | 0.0375 | 0.0166 | 0.4428 | 0.0475 | 0.0261120 |
| cluster3578 | -3391.4794 | 0.0336 | 0.0357 | 1.0623 | 0.0351 | 0.0095430 |
| cluster3579 | -2924.1608 | 0.0252 | 0.0219 | 0.8698 | 0.0290 | 0.0512270 |
| cluster3580 | -2413.9795 | 0.0079 | 0.0205 | 2.6063 | 999.0000 | 0.0066010 |
| cluster3581 | -1247.0857 | 0.0077 | 0.0145 | 1.8910 | 5.3436 | 0.0000020 |
| cluster3582 | -1103.4153 | 0.0067 | 0.0010 | 0.1441 | 0.0662 | 0.0183890 |
| cluster3583 | -2533.4362 | 0.0001 | 0.0001 | 1.1132 | 0.0001 | 0.0000110 |
| cluster3584 | -4866.8111 | 0.0270 | 0.0722 | 2.6723 | 0.1734 | 0.0099400 |
| cluster3585 | -2090.082 | 0.0458 | 0.0135 | 0.2946 | 0.1312 | 0.0555480 |
| cluster3586 | -5223.2864 | 0.0632 | 0.0913 | 1.4452 | 0.1435 | 0.0340540 |
| cluster3587 | -3466.494 | 0.0444 | 0.0988 | 2.2250 | 0.8544 | 0.0174920 |
| cluster3588 | -3359.1793 | 0.0299 | 0.0124 | 0.4158 | 0.0055 | 0.0009520 |
| cluster3589 | -3005.5498 | 0.0327 | 0.0345 | 1.0578 | 0.5684 | 0.0255510 |
| cluster3590 | -4182.0547 | 0.0631 | 0.1198 | 1.8998 | 0.0460 | 0.0098680 |
| cluster3591 | -2874.3662 | 0.0126 | 0.0041 | 0.3241 | 0.0109 | 0.0111050 |
| cluster3592 | -4702.508 | 0.0343 | 0.0634 | 1.8447 | 0.1936 | 0.0127610 |
| cluster3593 | -3415.184 | 0.0395 | 0.0737 | 1.8687 | 0.2208 | 0.0205660 |
| cluster3594 | -3176.337 | 0.0564 | 0.0455 | 0.8057 | 0.1907 | 0.0084170 |
| cluster3595 | -2123.6489 | 0.0121 | 0.0053 | 0.4420 | 0.0118 | 0.0074070 |
| cluster3596 | -2315.0236 | 0.0302 | 0.0294 | 0.9723 | 0.0987 | 0.0080380 |
| cluster3597 | -4191.3156 | 0.0254 | 0.0201 | 0.7915 | 0.0607 | 0.0298930 |
| cluster3598 | -4253.7418 | 0.0214 | 0.1186 | 5.5354 | 0.0291 | 0.0085090 |

| | | | | | | |
|---|---|---|---|---|---|---|
| cluster3599 | -2242.9564 | 0.0193 | 0.0249 | 1.2953 | 999.0000 | 0.0139760 |
| cluster3600 | -2861.2903 | 0.0255 | 0.0181 | 0.7118 | 0.0224 | 0.0022790 |
| cluster3601 | -3285.3119 | 0.0026 | 0.0141 | 5.4079 | 7.2879 | 0.0006960 |
| cluster3602 | -3065.93 | 0.0031 | 0.0010 | 0.3254 | 0.0037 | 0.0028570 |
| cluster3603 | -4221.428 | 0.0477 | 0.0589 | 1.2333 | 0.3712 | 0.0108040 |
| cluster3604 | -1057.946 | 0.0579 | 0.0071 | 0.1221 | 0.1721 | 0.00389 |
| cluster3605 | -3057.8711 | 0.0377 | 0.0405 | 1.0726 | 0.0415 | 0.0082680 |
| cluster3606 | -4316.8059 | 0.0226 | 0.0431 | 1.9121 | 0.1494 | 0.0163560 |
| cluster3607 | -2596.6708 | 0.0343 | 0.0490 | 1.4289 | 0.0738 | 0.0111430 |
| cluster3608 | -2888.1005 | 0.0113 | 0.0067 | 0.5991 | 0.0091 | 0.0082350 |
| cluster3609 | -4181.7381 | 0.0279 | 0.0603 | 2.1632 | 0.0470 | 0.0131380 |
| cluster3610 | -1395.8048 | 0.0070 | 0.0269 | 3.8350 | 7.5703 | 0.0000020 |
| cluster3611 | -2340.5299 | 0.0248 | 0.0778 | 3.1335 | 0.2356 | 0.0055590 |
| cluster3612 | -2831.5953 | 0.0335 | 0.0689 | 2.0550 | 0.0440 | 0.0061720 |
| cluster3613 | -4058.7587 | 0.0168 | 0.0092 | 0.5489 | 0.0233 | 0.0231300 |
| cluster3614 | -4307.2417 | 0.0234 | 0.0714 | 3.0583 | 0.1290 | 0.0036640 |
| cluster3615 | -1662.7414 | 0.0531 | 0.0090 | 0.1703 | 0.0296 | 0.0097570 |
| cluster3616 | -773.2269 | 0.0099 | 0.0091 | 0.9262 | 999.0000 | 0.0106960 |
| cluster3617 | -1928.8317 | 0.0377 | 0.0493 | 1.3085 | 0.1777 | 0.0183610 |
| cluster3618 | -1898.8036 | 0.1573 | 0.1187 | 0.7546 | 0.4248 | 0.01464 |
| cluster3619 | -2552.1726 | 0.0040 | 0.0053 | 1.3261 | 0.0135 | 0.0012510 |
| cluster3620 | -3597.816 | 0.0398 | 0.0370 | 0.9286 | 0.1608 | 0.0412520 |
| cluster3621 | -4760.1591 | 0.0294 | 0.0674 | 2.2932 | 0.1507 | 0.0139210 |
| cluster3622 | -2722.3699 | 0.0110 | 0.0063 | 0.5695 | 0.0457 | 0.0040830 |
| cluster3623 | -2271.6932 | 0.0094 | 0.0307 | 3.2531 | 0.0065 | 0.0014020 |
| cluster3624 | -2909.3795 | 0.0127 | 0.0319 | 2.5139 | 0.0479 | 0.0026740 |
| cluster3625 | -3308.2436 | 0.0058 | 0.0138 | 2.3941 | 0.0157 | 0.0026270 |
| cluster3626 | -1982.478 | 0.0332 | 0.0170 | 0.5108 | 0.0192 | 0.0248860 |
| cluster3627 | -3000.9981 | 0.0070 | 0.0084 | 1.1910 | 0.0122 | 0.0014120 |
| cluster3628 | -3022.9283 | 0.0268 | 0.0580 | 2.1673 | 0.0404 | 0.0087270 |
| cluster3629 | -4162.4953 | 0.0103 | 0.0304 | 2.9633 | 0.1458 | 0.0072840 |
| cluster3630 | -1208.8941 | 0.0173 | 0.0059 | 0.3419 | 0.0001 | 0.0000200 |
| cluster3631 | -3583.5465 | 0.0111 | 0.0098 | 0.8824 | 0.0676 | 0.0661220 |
| cluster3632 | -2458.7847 | 0.0354 | 0.0409 | 1.1558 | 0.0188 | 0.0638950 |
| cluster3633 | -2496.9684 | 0.0266 | 0.0145 | 0.5463 | 0.0011 | 0.0063000 |
| cluster3634 | -939.30444 | 0.0155 | 0.0085 | 0.5486 | 0.0001 | 0.0000010 |
| cluster3635 | -1620.9926 | 0.0389 | 0.0053 | 0.1374 | 0.0508 | 0.0148030 |
| cluster3636 | -2529.0962 | 0.0178 | 0.0142 | 0.7986 | 0.0052 | 0.0119490 |
| cluster3637 | -1060.2274 | 0.0107 | 0.0035 | 0.3317 | 999.0000 | 0.0041200 |
| cluster3638 | -2732.5105 | 0.0310 | 0.0210 | 0.6758 | 0.0496 | 0.0040660 |
| cluster3639 | -1272.049 | 0.0310 | 0.0149 | 0.4795 | 0.0521 | 0.0263680 |
| cluster3640 | -4764.8345 | 0.0674 | 0.1096 | 1.6260 | 0.1519 | 0.0222790 |
| cluster3641 | -1766.8974 | 0.0334 | 0.0071 | 0.2117 | 0.0084 | 0.0037900 |
| cluster3642 | -3357.5704 | 0.0306 | 0.0510 | 1.6672 | 0.0459 | 0.0086580 |

| | | | | | | |
|---|---|---|---|---|---|---|
| cluster3643 | -3535.8298 | 0.0930 | 0.1121 | 1.2065 | 0.3095 | 0.0183390 |
| cluster3644 | -1911.4575 | 0.0243 | 0.0097 | 0.3973 | 0.0644 | 0.0124740 |
| cluster3645 | -1807.4474 | 0.0285 | 0.0395 | 1.3880 | 0.0685 | 0.0084700 |
| cluster3646 | -2634.7218 | 0.0192 | 0.0626 | 3.2537 | 999.0000 | 0.0043040 |
| cluster3647 | -3429.1376 | 0.0040 | 0.0193 | 4.7972 | 8.2358 | 0.0023570 |
| cluster3648 | -1759.176 | 0.0018 | 0.0011 | 0.5935 | 0.0006 | 0.0017470 |
| cluster3649 | -1640.2347 | 0.0092 | 0.0072 | 0.7806 | 0.0464 | 0.0014130 |
| cluster3650 | -1253.4316 | 0.0072 | 0.0540 | 7.4661 | 999.0000 | 0.0161590 |
| cluster3651 | -3971.1277 | 0.0459 | 0.0292 | 0.6359 | 0.0567 | 0.0610470 |
| cluster3652 | -3310.7341 | 0.0250 | 0.0152 | 0.6088 | 0.0253 | 0.0296470 |
| cluster3653 | -1426.933 | 0.0041 | 0.0023 | 0.5469 | 0.0029 | 0.0018240 |
| cluster3654 | -2837.9411 | 0.0092 | 0.0153 | 1.6638 | 0.0298 | 0.0007020 |
| cluster3655 | -2366.6099 | 0.0654 | 0.0682 | 1.0420 | 0.0931 | 0.0044750 |
| cluster3656 | -3108.7595 | 0.0250 | 0.0178 | 0.7127 | 0.0274 | 0.0324050 |
| cluster3657 | -3194.4488 | 0.0150 | 0.0390 | 2.5983 | 0.0154 | 0.0014200 |
| cluster3658 | -753.69075 | 0.0920 | 0.0147 | 0.1597 | 0.0776 | 0.04367 |
| cluster3659 | -2269.489 | 0.0340 | 0.0219 | 0.6440 | 0.0620 | 0.0067230 |
| cluster3660 | -3017.5517 | 0.0177 | 0.0547 | 3.0971 | 0.0352 | 0.0058010 |
| cluster3661 | -1818.9915 | 0.0021 | 0.0003 | 0.1292 | 0.0001 | 0.0000050 |
| cluster3662 | -2804.273 | 0.0359 | 0.0485 | 1.3525 | 0.0443 | 0.0047540 |
| cluster3663 | -3133.1564 | 0.0456 | 0.0456 | 0.9994 | 0.1052 | 0.0108570 |
| cluster3664 | -2443.0495 | 0.0305 | 0.0675 | 2.2148 | 0.1684 | 0.0322910 |
| cluster3665 | -3603.2124 | 0.0201 | 0.0347 | 1.7316 | 0.0324 | 0.0040620 |
| cluster3666 | -2712.0325 | 0.0333 | 0.0402 | 1.2055 | 4.6227 | 0.0000020 |
| cluster3667 | -1570.8146 | 0.0213 | 0.0075 | 0.3501 | 0.0178 | 0.0013660 |
| cluster3668 | -2980.7622 | 0.0186 | 0.0239 | 1.2850 | 0.0758 | 0.0094380 |
| cluster3669 | -2595.8302 | 0.0305 | 0.0268 | 0.8785 | 0.5199 | 0.0130440 |
| cluster3670 | -4034.202 | 0.0430 | 0.0328 | 0.7630 | 0.0610 | 0.0677770 |
| cluster3671 | -3994.5551 | 0.0260 | 0.0435 | 1.6751 | 0.0683 | 0.0150300 |
| cluster3672 | -2500.7308 | 0.0444 | 0.0489 | 1.1013 | 0.0615 | 0.0029020 |
| cluster3673 | -4407.4962 | 0.0356 | 0.1314 | 3.6921 | 0.1831 | 0.0214470 |
| cluster3674 | -2025.3342 | 0.0268 | 0.0504 | 1.8835 | 0.0491 | 0.0031490 |
| cluster3675 | -660.30852 | 0.0001 | 0.0000 | 0.1120 | 0.0001 | 0.0000260 |
| cluster3676 | -3510.8249 | 0.0383 | 0.0555 | 1.4501 | 0.2875 | 0.0453610 |
| cluster3677 | -812.07325 | 0.0234 | 0.0099 | 0.4248 | 0.0189 | 0.0149430 |
| cluster3678 | -3306.5885 | 0.0315 | 0.0327 | 1.0394 | 0.0594 | 0.0174600 |
| cluster3679 | -944.37913 | 0.0067 | 0.0022 | 0.3215 | 0.0131 | 0.0083880 |
| cluster3680 | -2314.1926 | 0.0091 | 0.0255 | 2.8138 | 0.0805 | 0.0017740 |
| cluster3681 | -2959.2458 | 0.0194 | 0.0389 | 2.0086 | 0.0241 | 0.0021500 |
| cluster3682 | -4361.7951 | 0.0299 | 0.0573 | 1.9191 | 0.1402 | 0.0141330 |
| cluster3683 | -2796.1546 | 0.0868 | 0.0672 | 0.7748 | 0.4116 | 0.2750700 |
| cluster3684 | -2990.0921 | 0.0304 | 0.0397 | 1.3061 | 0.1447 | 0.0138460 |
| cluster3685 | -3667.9827 | 0.0660 | 0.1224 | 1.8559 | 1.5993 | 0.0252960 |
| cluster3686 | -1626.9282 | 0.0457 | 0.0973 | 2.1294 | 0.0001 | 0.0000300 |

| | | | | | | |
|---|---|---|---|---|---|---|
| cluster3687 | -2874.3405 | 0.0738 | 0.1667 | 2.2578 | 999.0000 | 0.0103620 |
| cluster3688 | -2049.9172 | 0.0069 | 0.0175 | 2.5248 | 0.0338 | 0.0042380 |
| cluster3689 | -2638.0984 | 0.0309 | 0.0176 | 0.5716 | 0.1005 | 0.0384580 |
| cluster3690 | -1454.813 | 0.0218 | 0.0129 | 0.5923 | 0.0279 | 0.0350770 |
| cluster3691 | -667.98374 | 0.0014 | 0.0033 | 2.3583 | 999.0000 | 0.0042640 |
| cluster3692 | -1435.0363 | 0.0105 | 0.0152 | 1.4427 | 0.0534 | 0.0020210 |
| cluster3693 | -2463.7331 | 0.0084 | 0.0221 | 2.6370 | 85.2628 | 0.0036650 |
| cluster3694 | -2092.1122 | 0.0328 | 0.0084 | 0.2569 | 0.0251 | 0.0161710 |
| cluster3695 | -1735.7121 | 0.0220 | 0.0273 | 1.2415 | 0.1091 | 0.0072250 |
| cluster3696 | -817.96763 | 0.0105 | 0.0385 | 3.6828 | 0.0001 | 0.0000000 |
| cluster3697 | -3583.8695 | 0.0688 | 0.0324 | 0.4701 | 0.0860 | 0.0756150 |
| cluster3698 | -1099.5724 | 0.0001 | 0.0000 | 0.2917 | 0.0001 | 0.0000250 |
| cluster3699 | -4437.0639 | 0.0418 | 0.0774 | 1.8550 | 0.3002 | 0.0158230 |
| cluster3700 | -2818.2616 | 0.0300 | 0.0358 | 1.1951 | 0.0709 | 0.0081470 |
| cluster3701 | -1128.2553 | 0.0016 | 0.0005 | 0.3321 | 0.0001 | 0.0000620 |
| cluster3702 | -3860.3094 | 0.0203 | 0.0583 | 2.8723 | 258.3070 | 0.0117690 |
| cluster3703 | -1511.3765 | 0.0119 | 0.0038 | 0.3172 | 0.0322 | 0.0151840 |
| cluster3704 | -2229.5472 | 0.0043 | 0.0099 | 2.3209 | 0.0109 | 0.0192190 |
| cluster3705 | -1945.2249 | 0.0136 | 0.0075 | 0.5535 | 0.0134 | 0.0112680 |
| cluster3706 | -2094.4118 | 0.0202 | 0.0155 | 0.7705 | 0.0413 | 0.0026560 |
| cluster3707 | -1920.0011 | 0.0450 | 0.0750 | 1.6659 | 0.0001 | 0.0000140 |
| cluster3708 | -2635.7257 | 0.0440 | 0.0422 | 0.9587 | 0.1673 | 0.0062640 |
| cluster3709 | -1580.3022 | 0.0803 | 0.0246 | 0.3065 | 0.0001 | 0.0000010 |
| cluster3710 | -1919.0697 | 0.0103 | 0.0244 | 2.3708 | 0.0698 | 0.0212290 |
| cluster3711 | -1439.5585 | 0.0285 | 0.0080 | 0.2818 | 0.0507 | 0.0013160 |
| cluster3712 | -3401.1812 | 0.0304 | 0.0192 | 0.6326 | 0.0128 | 0.0091230 |
| cluster3713 | -4020.4339 | 0.0446 | 0.0897 | 2.0094 | 999.0000 | 0.0175200 |
| cluster3714 | -1775.8554 | 0.0458 | 0.0148 | 0.3240 | 0.0477 | 0.0260170 |
| cluster3715 | -1284.5073 | 0.0104 | 0.0483 | 4.6604 | 0.1169 | 0.0145130 |
| cluster3716 | -3657.8051 | 0.0446 | 0.0642 | 1.4383 | 0.0673 | 0.0180570 |
| cluster3717 | -1403.8669 | 0.0154 | 0.0035 | 0.2297 | 0.0108 | 0.0040490 |
| cluster3718 | -2986.6714 | 0.0172 | 0.0374 | 2.1782 | 0.3363 | 0.0079820 |
| cluster3719 | -3294.8846 | 0.0458 | 0.0410 | 0.8953 | 0.0317 | 0.0049680 |
| cluster3720 | -2758.1546 | 0.0251 | 0.0220 | 0.8799 | 0.0952 | 0.0031900 |
| cluster3721 | -1097.4231 | 0.0070 | 0.0009 | 0.1302 | 0.0001 | 0.0000240 |
| cluster3722 | -1223.7491 | 0.0485 | 0.1146 | 2.3630 | 0.0106 | 0.0029650 |
| cluster3723 | -989.94266 | 0.0001 | 0.0001 | 0.5102 | 0.0001 | 0.0000010 |
| cluster3724 | -3080.5748 | 0.0295 | 0.0455 | 1.5430 | 0.2094 | 0.0096340 |
| cluster3725 | -1839.8593 | 0.0107 | 0.0196 | 1.8270 | 0.0055 | 0.0012900 |
| cluster3726 | -1935.4088 | 0.0522 | 0.0337 | 0.6468 | 0.0136 | 0.0021800 |
| cluster3727 | -2466.6598 | 0.0477 | 0.0848 | 1.7792 | 0.6213 | 0.0080190 |
| cluster3728 | -2157.0913 | 0.0210 | 0.0141 | 0.6734 | 0.0210 | 0.0132090 |
| cluster3729 | -2946.8382 | 0.0125 | 0.0215 | 1.7266 | 0.0235 | 0.0046900 |
| cluster3730 | -1782.0557 | 0.0820 | 0.0215 | 0.2616 | 0.0830 | 0.0227560 |

| | | | | | | |
|---|---|---|---|---|---|---|
| cluster3731 | -2864.2062 | 0.0105 | 0.0058 | 0.5553 | 0.0041 | 0.0037180 |
| cluster3732 | -2822.2552 | 0.0213 | 0.0207 | 0.9723 | 0.0609 | 0.0047080 |
| cluster3733 | -3072.6737 | 0.0769 | 0.0340 | 0.4424 | 0.0938 | 0.0347610 |
| cluster3734 | -3043.4313 | 0.0428 | 0.0509 | 1.1882 | 0.0488 | 0.0099400 |
| cluster3735 | -3249.4592 | 0.0882 | 0.0356 | 0.4033 | 0.0590 | 0.0426180 |
| cluster3736 | -3538.2938 | 0.0463 | 0.0316 | 0.6822 | 0.0439 | 0.0077290 |
| cluster3737 | -1208.3728 | 0.0016 | 0.0005 | 0.3106 | 0.0197 | 0.0020560 |
| cluster3738 | -3259.4362 | 0.0099 | 0.0070 | 0.7040 | 0.0062 | 0.0067760 |
| cluster3739 | -1158.3637 | 0.0001 | 0.0000 | 0.3318 | 0.5101 | 0.0000010 |
| cluster3740 | -3446.5185 | 0.0265 | 0.0361 | 1.3595 | 0.1107 | 0.0138950 |
| cluster3741 | -1310.6038 | 0.0136 | 0.0072 | 0.5279 | 0.1011 | 0.0060280 |
| cluster3742 | -1613.8034 | 0.0481 | 0.0549 | 1.1396 | 0.2956 | 0.0186860 |
| cluster3743 | -2977.892 | 0.0211 | 0.0308 | 1.4638 | 0.2139 | 0.0061370 |
| cluster3744 | -1350.6734 | 0.0792 | 0.0551 | 0.6960 | 1.9996 | 0.01977 |
| cluster3745 | -2583.6467 | 0.0211 | 0.0346 | 1.6407 | 0.1082 | 0.0134040 |
| cluster3746 | -3782.1527 | 0.0447 | 0.0379 | 0.8489 | 0.0162 | 0.0229100 |
| cluster3747 | -1120.2437 | 0.0256 | 0.0216 | 0.8461 | 0.1472 | 0.0056920 |
| cluster3748 | -2162.1118 | 0.0092 | 0.0046 | 0.4938 | 0.0400 | 0.0205480 |
| cluster3749 | -3066.1666 | 0.0146 | 0.0246 | 1.6918 | 0.3075 | 0.0078350 |
| cluster3750 | -1355.4029 | 0.0247 | 0.0571 | 2.3160 | 0.0124 | 0.0016160 |
| cluster3751 | -3011.9671 | 0.0046 | 0.0052 | 1.1384 | 0.0064 | 0.0009060 |
| cluster3752 | -2663.829 | 0.0076 | 0.0165 | 2.1570 | 3.4764 | 0.0000020 |
| cluster3753 | -1021.9505 | 0.0264 | 0.0027 | 0.1011 | 0.0001 | 0.00001 |
| cluster3754 | -881.97724 | 0.0001 | 0.0000 | 0.2015 | 0.0001 | 0.0000010 |
| cluster3755 | -2499.4647 | 0.0147 | 0.0115 | 0.7811 | 0.0228 | 0.0189030 |
| cluster3756 | -1686.0491 | 0.0300 | 0.0333 | 1.1106 | 0.1986 | 0.0167950 |
| cluster3757 | -2090.8086 | 0.0429 | 0.0225 | 0.5256 | 0.0879 | 0.0079340 |
| cluster3758 | -1511.707 | 0.0277 | 0.0138 | 0.4995 | 0.0345 | 0.0201620 |
| cluster3759 | -3063.3635 | 0.0578 | 0.0254 | 0.4396 | 0.1159 | 0.0961620 |
| cluster3760 | -2599.6918 | 0.0203 | 0.0204 | 1.0049 | 0.0434 | 0.0358100 |
| cluster3761 | -767.79542 | 0.0054 | 0.0295 | 5.4672 | 0.0001 | 0.0000220 |
| cluster3762 | -1963.5196 | 0.0071 | 0.0015 | 0.2178 | 0.0339 | 0.0187670 |
| cluster3763 | -2001.5972 | 0.0698 | 0.0169 | 0.2418 | 0.0140 | 0.0053140 |
| cluster3764 | -1628.8335 | 0.0265 | 0.0502 | 1.8899 | 0.0770 | 0.0114810 |
| cluster3765 | -2669.8429 | 0.0139 | 0.0137 | 0.9789 | 0.0394 | 0.0207950 |
| cluster3766 | -1795.4746 | 0.0117 | 0.0153 | 1.3042 | 0.0001 | 0.0000040 |
| cluster3767 | -862.69362 | 0.0244 | 0.0422 | 1.7308 | 0.0167 | 0.0008710 |
| cluster3768 | -2219.2415 | 0.0376 | 0.0708 | 1.8847 | 0.0620 | 0.0082450 |
| cluster3769 | -1764.87 | 0.0064 | 0.0022 | 0.3459 | 0.0220 | 0.0014750 |
| cluster3770 | -1775.1439 | 0.0528 | 0.0463 | 0.8775 | 0.0555 | 0.0110710 |
| cluster3771 | -2138.5572 | 0.0759 | 0.0751 | 0.9906 | 0.0429 | 0.0100610 |
| cluster3772 | -725.30225 | 0.0009 | 0.0221 | 24.0025 | 0.0001 | 0.0000020 |
| cluster3773 | -1117.2365 | 0.2249 | 0.0462 | 0.2054 | 0.0001 | 0.00000 |
| cluster3774 | -1408.8883 | 0.0141 | 0.0035 | 0.2476 | 0.0001 | 0.0000040 |

| | | | | | | |
|---|---|---|---|---|---|---|
| cluster3775 | -906.37489 | 0.0118 | 0.0239 | 2.0333 | 0.0388 | 0.0039390 |
| cluster3776 | -2518.6436 | 0.0790 | 0.0297 | 0.3761 | 0.0729 | 0.0325170 |
| cluster3777 | -3059.3394 | 0.0433 | 0.0641 | 1.4827 | 0.2272 | 0.0570530 |
| cluster3778 | -980.50837 | 0.0008 | 0.0006 | 0.7053 | 0.0024 | 0.0028140 |
| cluster3779 | -2995.0266 | 0.0228 | 0.0320 | 1.3995 | 0.0160 | 0.0284760 |
| cluster3780 | -1679.955 | 0.0080 | 0.0234 | 2.9300 | 361.7500 | 0.0146200 |
| cluster3781 | -2686.1562 | 0.0601 | 0.0371 | 0.6168 | 0.0455 | 0.0312980 |
| cluster3782 | -2289.5656 | 0.0395 | 0.0293 | 0.7406 | 0.0861 | 0.0126020 |
| cluster3783 | -2105.7623 | 0.0064 | 0.0345 | 5.4007 | 999.0000 | 0.0034040 |
| cluster3784 | -2951.2645 | 0.0450 | 0.0380 | 0.8438 | 0.0637 | 0.0732920 |
| cluster3785 | -2684.2411 | 0.0139 | 0.0105 | 0.7552 | 0.0961 | 0.0077170 |
| cluster3786 | -2266.6429 | 0.0099 | 0.0290 | 2.9398 | 0.0053 | 0.0067710 |
| cluster3787 | -1522.0568 | 0.0258 | 0.0240 | 0.9337 | 0.0546 | 0.0066790 |
| cluster3788 | -1160.4015 | 0.0001 | 0.0000 | 0.1007 | 0.0001 | 0.00000 |
| cluster3789 | -3038.3016 | 0.0144 | 0.0062 | 0.4351 | 0.0020 | 0.0013220 |
| cluster3790 | -2177.4301 | 0.0131 | 0.0322 | 2.4533 | 0.0343 | 0.0020830 |
| cluster3791 | -420.00572 | 0.0117 | 0.0045 | 0.3890 | 0.0211 | 0.0059760 |
| cluster3792 | -1863.2583 | 0.0057 | 0.0034 | 0.5905 | 0.0110 | 0.0013170 |
| cluster3793 | -1851.7916 | 0.0238 | 0.0216 | 0.9092 | 0.1023 | 0.0105080 |
| cluster3794 | -2189.2818 | 0.0155 | 0.0225 | 1.4527 | 0.0320 | 0.0041420 |
| cluster3795 | -1701.656 | 0.0197 | 0.0219 | 1.1130 | 0.0001 | 0.0000010 |
| cluster3796 | -1251.5025 | 0.1185 | 0.0859 | 0.7245 | 0.2379 | 0.02254 |
| cluster3797 | -3912.5783 | 0.0524 | 0.1450 | 2.7654 | 0.1366 | 0.0073830 |
| cluster3798 | -3443.5201 | 0.0207 | 0.0294 | 1.4170 | 0.0798 | 0.0146310 |
| cluster3799 | -2341.5359 | 0.0472 | 0.0259 | 0.5482 | 0.0446 | 0.0599520 |
| cluster3800 | -431.25192 | 0.0123 | 0.0011 | 0.0918 | 0.0001 | 0.00001 |
| cluster3801 | -2690.5652 | 0.2037 | 0.1039 | 0.5103 | 0.2805 | 0.00725 |
| cluster3802 | -2649.196 | 0.0315 | 0.0395 | 1.2532 | 0.0541 | 0.0515340 |
| cluster3803 | -2236.4914 | 0.0307 | 0.0293 | 0.9538 | 0.1988 | 0.0127030 |
| cluster3804 | -2634.7446 | 0.0303 | 0.0318 | 1.0494 | 0.0761 | 0.0065200 |
| cluster3805 | -2655.9579 | 0.0239 | 0.0319 | 1.3352 | 0.0201 | 0.0027790 |
| cluster3806 | -3563.8746 | 0.0346 | 0.0471 | 1.3600 | 0.1333 | 0.0165810 |
| cluster3807 | -1441.8671 | 0.0082 | 0.0020 | 0.2400 | 0.0061 | 0.0037180 |
| cluster3808 | -1825.8667 | 0.0327 | 0.0467 | 1.4266 | 0.0001 | 0.0000050 |
| cluster3809 | -2360.2965 | 0.0099 | 0.0098 | 0.9914 | 0.0001 | 0.0000180 |
| cluster3810 | -2462.8888 | 0.0114 | 0.0101 | 0.8919 | 0.0035 | 0.0078290 |
| cluster3811 | -761.4375 | 0.0221 | 0.0049 | 0.2205 | 0.0719 | 0.00288 |
| cluster3812 | -3000.7916 | 0.0307 | 0.0244 | 0.7967 | 0.0214 | 0.0013610 |
| cluster3813 | -3413.6604 | 0.0358 | 0.0958 | 2.6742 | 0.0283 | 0.0095160 |
| cluster3814 | -1607.1362 | 0.0720 | 0.0134 | 0.1864 | 0.1649 | 0.03749 |
| cluster3815 | -1707.5406 | 0.0170 | 0.0399 | 2.3499 | 0.0728 | 0.0083560 |
| cluster3816 | -2526.6164 | 0.0261 | 0.0499 | 1.9127 | 0.0457 | 0.0030470 |
| cluster3817 | -989.71726 | 0.0116 | 0.0022 | 0.1910 | 0.0001 | 0.0000770 |
| cluster3818 | -1978.5222 | 0.0577 | 0.0624 | 1.0808 | 0.0980 | 0.0815040 |

| | | | | | | |
|---|---|---|---|---|---|---|
| cluster3819 | -3344.6516 | 0.0246 | 0.0306 | 1.2444 | 0.1393 | 0.0087180 |
| cluster3820 | -2225.8016 | 0.0300 | 0.0316 | 1.0549 | 0.0281 | 0.0028560 |
| cluster3821 | -2243.7444 | 0.0164 | 0.0110 | 0.6754 | 0.0154 | 0.0165940 |
| cluster3822 | -3396.1771 | 0.0247 | 0.0427 | 1.7319 | 0.1015 | 0.0152660 |
| cluster3823 | -3407.8304 | 0.0180 | 0.0258 | 1.4333 | 0.0335 | 0.0279140 |
| cluster3824 | -2344.162 | 0.0171 | 0.0439 | 2.5618 | 999.0000 | 0.0209660 |
| cluster3825 | -1388.6865 | 0.0067 | 0.0062 | 0.9212 | 0.0001 | 0.0000080 |
| cluster3826 | -3155.1311 | 0.0754 | 0.1063 | 1.4093 | 0.0602 | 0.0078360 |
| cluster3827 | -2403.5245 | 0.0296 | 0.0249 | 0.8399 | 0.0216 | 0.0258790 |
| cluster3828 | -2365.0744 | 0.0490 | 0.0462 | 0.9428 | 0.0001 | 0.0000040 |
| cluster3829 | -2252.3641 | 0.0199 | 0.0321 | 1.6147 | 0.5656 | 0.0110910 |
| cluster3830 | -1629.3648 | 0.0204 | 0.0085 | 0.4145 | 0.0187 | 0.0010140 |
| cluster3831 | -2347.4872 | 0.0298 | 0.0572 | 1.9212 | 0.1854 | 0.0066980 |
| cluster3832 | -1768.6053 | 0.0045 | 0.0040 | 0.8864 | 0.0001 | 0.0000030 |
| cluster3833 | -2129.6444 | 0.0215 | 0.0256 | 1.1905 | 999.0000 | 0.0031760 |
| cluster3834 | -2056.4315 | 0.0078 | 0.0247 | 3.1765 | 513.1948 | 0.0051250 |
| cluster3835 | -2117.9497 | 0.0327 | 0.0314 | 0.9604 | 0.0656 | 0.0067070 |
| cluster3836 | -1894.9967 | 0.0170 | 0.0171 | 1.0081 | 0.0226 | 0.0032810 |
| cluster3837 | -2635.9363 | 0.0376 | 0.0430 | 1.1446 | 999.0000 | 0.0084880 |
| cluster3838 | -2291.2555 | 0.0244 | 0.0242 | 0.9908 | 1.8984 | 0.0360470 |
| cluster3839 | -1497.5333 | 0.0158 | 0.0233 | 1.4704 | 0.1078 | 0.0126520 |
| cluster3840 | -2023.8127 | 0.0032 | 0.0763 | 23.6929 | 0.0074 | 0.0115690 |
| cluster3841 | -2293.1704 | 0.0755 | 0.1150 | 1.5233 | 0.0763 | 0.0036360 |
| cluster3842 | -1606.0556 | 0.0048 | 0.0009 | 0.1929 | 0.0001 | 0.0000380 |
| cluster3843 | -1440.9317 | 0.0121 | 0.0083 | 0.6870 | 0.0001 | 0.0000080 |
| cluster3844 | -1067.7154 | 0.0237 | 0.0283 | 1.1910 | 2.0658 | 0.05556 |
| cluster3845 | -604.91636 | 0.0001 | 0.0000 | 0.1561 | 0.0001 | 0.0000310 |
| cluster3846 | -1273.8196 | 0.0184 | 0.0115 | 0.6254 | 0.0132 | 0.0144490 |
| cluster3847 | -1157.4642 | 0.0009 | 0.0018 | 1.9374 | 0.5187 | 0.0000010 |
| cluster3848 | -2511.6515 | 0.0255 | 0.0428 | 1.6824 | 0.2377 | 0.0219640 |
| cluster3849 | -2658.8188 | 0.0116 | 0.0607 | 5.2312 | 999.0000 | 0.0082260 |
| cluster3850 | -1676.2136 | 0.0188 | 0.0149 | 0.7924 | 0.0852 | 0.0065670 |
| cluster3851 | -1618.0646 | 0.0094 | 0.0117 | 1.2456 | 0.0173 | 0.0018980 |
| cluster3852 | -1851.6097 | 0.0099 | 0.0105 | 1.0573 | 0.0012 | 0.0124060 |
| cluster3853 | -1101.5484 | 0.0045 | 0.0119 | 2.6537 | 0.0107 | 0.0025520 |
| cluster3854 | -1357.2727 | 0.0889 | 0.0556 | 0.6252 | 0.2502 | 0.01860 |
| cluster3855 | -2930.1814 | 0.0096 | 0.0366 | 3.8331 | 0.0521 | 0.0027450 |
| cluster3856 | -1691.3932 | 0.0147 | 0.0231 | 1.5680 | 0.1668 | 0.0145690 |
| cluster3857 | -3038.0343 | 0.0165 | 0.0235 | 1.4256 | 0.0188 | 0.0047640 |
| cluster3858 | -2416.699 | 0.0496 | 0.0395 | 0.7977 | 0.0438 | 0.0258420 |
| cluster3859 | -1977.2128 | 0.0124 | 0.0121 | 0.9780 | 0.0357 | 0.0025320 |
| cluster3860 | -2624.7157 | 0.0755 | 0.0357 | 0.4724 | 0.1192 | 0.0430730 |
| cluster3861 | -1300.2379 | 0.0156 | 0.0208 | 1.3320 | 0.0666 | 0.0049000 |
| cluster3862 | -922.33901 | 0.0001 | 0.0001 | 0.7991 | 0.0001 | 0.0000000 |

| | | | | | | |
|---|---|---|---|---|---|---|
| cluster3863 | -1483.259 | 0.0008 | 0.0005 | 0.5627 | 0.0017 | 0.0018210 |
| cluster3864 | -2218.6993 | 0.0029 | 0.0082 | 2.8224 | 0.0479 | 0.0026740 |
| cluster3865 | -3003.4103 | 0.0408 | 0.0738 | 1.8085 | 0.1004 | 0.0141210 |
| cluster3866 | -1627.265 | 0.0124 | 0.0319 | 2.5705 | 999.0000 | 0.0126560 |
| cluster3867 | -608.47114 | 0.0001 | 0.0000 | 0.0708 | 0.0001 | 0.00000 |
| cluster3868 | -1392.5319 | 0.0297 | 0.0158 | 0.5340 | 0.0001 | 0.0000110 |
| cluster3869 | -2971.4636 | 0.0272 | 0.0897 | 3.2986 | 1.4132 | 0.0115500 |
| cluster3870 | -2460.2896 | 0.0356 | 0.0412 | 1.1590 | 0.3058 | 0.0102610 |
| cluster3871 | -2151.3221 | 0.0067 | 0.0087 | 1.2956 | 0.2567 | 0.0024860 |
| cluster3872 | -2647.3061 | 0.0112 | 0.0266 | 2.3754 | 999.0000 | 0.0043400 |
| cluster3873 | -1788.1583 | 0.0286 | 0.0201 | 0.7035 | 0.0297 | 0.0174880 |
| cluster3874 | -2101.8859 | 0.0125 | 0.0039 | 0.3098 | 0.0204 | 0.0034980 |
| cluster3875 | -770.52123 | 0.0028 | 0.0018 | 0.6377 | 0.0001 | 0.0000010 |
| cluster3876 | -744.76712 | 0.0135 | 0.0060 | 0.4483 | 0.0001 | 0.0000420 |
| cluster3877 | -1435.206 | 0.0169 | 0.0068 | 0.4028 | 0.0059 | 0.0038790 |
| cluster3879 | -1881.2661 | 0.0172 | 0.0176 | 1.0280 | 0.0325 | 0.0035230 |
| cluster3880 | -1357.6313 | 0.0200 | 0.0281 | 1.4013 | 0.0333 | 0.0026550 |
| cluster3881 | -3016.2153 | 0.0418 | 0.0832 | 1.9898 | 0.2061 | 0.0148110 |
| cluster3882 | -2686.3226 | 0.0207 | 0.0308 | 1.4873 | 0.1963 | 0.0117430 |
| cluster3883 | -426.40467 | 0.0160 | 0.1122 | 7.0236 | 999.0000 | 0.02499 |
| cluster3884 | -1563.6984 | 0.0004 | 0.0169 | 41.3837 | 293.2992 | 0.0006500 |
| cluster3885 | -2847.9362 | 0.0102 | 0.0151 | 1.4878 | 0.1880 | 0.0122540 |
| cluster3886 | -1394.281 | 0.0028 | 0.0131 | 4.7752 | 0.0222 | 0.0027160 |
| cluster3887 | -1627.7429 | 0.0078 | 0.0318 | 4.0879 | 0.0528 | 0.0025690 |
| cluster3888 | -1182.6856 | 0.0105 | 0.0459 | 4.3821 | 0.0121 | 0.0077850 |
| cluster3889 | -1003.6327 | 0.0265 | 0.0456 | 1.7232 | 0.0001 | 0.0000230 |
| cluster3890 | -1871.7922 | 0.0184 | 0.0249 | 1.3510 | 0.0136 | 0.0285620 |
| cluster3891 | -2184.7592 | 0.0143 | 0.0095 | 0.6647 | 0.0134 | 0.0197060 |
| cluster3892 | -1380.7901 | 0.0124 | 0.0077 | 0.6216 | 5.2832 | 0.0000020 |
| cluster3893 | -1458.1253 | 0.0208 | 0.0084 | 0.4069 | 0.0390 | 0.0365300 |
| cluster3894 | -966.53063 | 0.0080 | 0.0202 | 2.5208 | 0.0001 | 0.0000030 |
| cluster3895 | -1982.7211 | 0.0090 | 0.0165 | 1.8476 | 0.0156 | 0.0021100 |
| cluster3896 | -820.69173 | 0.0032 | 0.0059 | 1.8714 | 1.0166 | 0.0000010 |
| cluster3897 | -1450.6884 | 0.0081 | 0.0575 | 7.1395 | 0.0001 | 0.0000890 |
| cluster3898 | -2390.3308 | 0.0232 | 0.0060 | 0.2571 | 0.0283 | 0.0135750 |
| cluster3899 | -2998.3208 | 0.0218 | 0.0551 | 2.5282 | 0.2498 | 0.0173460 |
| cluster3900 | -2364.6174 | 0.0615 | 0.1333 | 2.1665 | 0.1114 | 0.0338270 |
| cluster3901 | -1545.5579 | 0.0023 | 0.0041 | 1.8084 | 0.8964 | 0.0000010 |
| cluster3902 | -2495.0068 | 0.0105 | 0.0201 | 1.9158 | 0.6649 | 0.0042030 |
| cluster3903 | -1622.858 | 0.0091 | 0.0366 | 4.0303 | 0.0324 | 0.0049330 |
| cluster3904 | -905.26245 | 0.0053 | 0.0179 | 3.3898 | 0.0001 | 0.0000050 |
| cluster3905 | -1593.9442 | 0.1466 | 0.0966 | 0.6585 | 0.0649 | 0.01971 |
| cluster3906 | -1452.5628 | 0.0226 | 0.0199 | 0.8810 | 0.0001 | 0.0000180 |
| cluster3907 | -1942.6291 | 0.0364 | 0.0363 | 0.9988 | 0.1663 | 0.0136210 |

| | | | | | | |
|---|---|---|---|---|---|---|
| cluster3908 | -1847.5315 | 0.0146 | 0.0196 | 1.3410 | 0.0001 | 0.0000120 |
| cluster3909 | -2462.6668 | 0.0204 | 0.0474 | 2.3301 | 0.4258 | 0.0091330 |
| cluster3910 | -1189.8669 | 0.0303 | 0.0537 | 1.7724 | 999.0000 | 0.0147470 |
| cluster3911 | -1749.4346 | 0.0075 | 0.0149 | 1.9925 | 0.0193 | 0.0037740 |
| cluster3912 | -951.76626 | 0.0001 | 0.0000 | 0.3201 | 0.0001 | 0.0000270 |
| cluster3913 | -2544.902 | 0.0394 | 0.0516 | 1.3102 | 0.0294 | 0.0053830 |
| cluster3914 | -2239.1047 | 0.0560 | 0.0924 | 1.6506 | 0.0621 | 0.0110210 |
| cluster3915 | -2226.3236 | 0.0320 | 0.0441 | 1.3787 | 0.0829 | 0.0134980 |
| cluster3916 | -902.07702 | 0.0732 | 0.0605 | 0.8265 | 1.6766 | 0.01124 |
| cluster3917 | -2582.3042 | 0.0132 | 0.0204 | 1.5418 | 0.0001 | 0.0000110 |
| cluster3918 | -2753.6549 | 0.0128 | 0.0630 | 4.9062 | 999.0000 | 0.0097010 |
| cluster3919 | -569.98635 | 0.0021 | 0.0008 | 0.3810 | 0.0001 | 0.00002 |
| cluster3920 | -657.87738 | 0.0099 | 0.1195 | 12.0566 | 474.2962 | 0.0023750 |
| cluster3921 | -2189.1762 | 0.0228 | 0.0291 | 1.2739 | 0.0868 | 0.0090530 |
| cluster3922 | -1239.7003 | 0.0480 | 0.0448 | 0.9330 | 0.2364 | 0.0080190 |
| cluster3923 | -1150.4886 | 0.0004 | 0.0012 | 3.0369 | 1.0234 | 0.0000010 |
| cluster3924 | -1695.9527 | 0.0190 | 0.0246 | 1.2971 | 0.0962 | 0.0024430 |
| cluster3925 | -1092.042 | 0.0434 | 0.0408 | 0.9416 | 0.2158 | 0.0180840 |
| cluster3926 | -2435.7759 | 0.0370 | 0.0984 | 2.6586 | 0.1511 | 0.0185870 |
| cluster3927 | -1186.3018 | 0.0083 | 0.0142 | 1.7041 | 0.0464 | 0.0080450 |
| cluster3928 | -1159.9359 | 0.0295 | 0.0172 | 0.5834 | 0.0241 | 0.0021440 |
| cluster3929 | -1607.786 | 0.0057 | 0.0104 | 1.8443 | 0.0204 | 0.0020590 |
| cluster3930 | -2016.085 | 0.0119 | 0.0057 | 0.4788 | 0.0289 | 0.0206230 |
| cluster3931 | -1829.0835 | 0.1431 | 0.0428 | 0.2989 | 0.0001 | 0.00000 |
| cluster3932 | -1117.6361 | 0.0129 | 0.0028 | 0.2199 | 0.0060 | 0.0022520 |
| cluster3933 | -1158.0962 | 0.0886 | 0.0373 | 0.4212 | 0.0001 | 0.00000 |
| cluster3934 | -717.10761 | 0.0001 | 0.0001 | 0.5905 | 0.0001 | 0.0000000 |
| cluster3935 | -3103.7346 | 0.0462 | 0.0954 | 2.0664 | 0.7036 | 0.0268220 |
| cluster3936 | -2124.1946 | 0.0115 | 0.0271 | 2.3564 | 999.0000 | 0.0188690 |
| cluster3937 | -1937.8539 | 0.0170 | 0.0097 | 0.5694 | 0.0358 | 0.0186450 |
| cluster3938 | -1782.2945 | 0.0318 | 0.0225 | 0.7066 | 0.0390 | 0.0062000 |
| cluster3939 | -2055.6456 | 0.0159 | 0.0219 | 1.3796 | 0.0001 | 0.0000050 |
| cluster3940 | -1338.556 | 0.0182 | 0.0061 | 0.3360 | 0.0167 | 0.0129520 |
| cluster3941 | -971.69228 | 0.0280 | 0.0114 | 0.4070 | 1.9424 | 0.0000010 |
| cluster3942 | -2009.6112 | 0.0314 | 0.0203 | 0.6459 | 0.0319 | 0.0064600 |
| cluster3943 | -2267.9919 | 0.0143 | 0.0401 | 2.8036 | 0.0377 | 0.0058220 |
| cluster3944 | -1590.6573 | 0.0383 | 0.0449 | 1.1726 | 0.1462 | 0.0254400 |
| cluster3945 | -1676.4953 | 0.0397 | 0.0501 | 1.2636 | 0.0190 | 0.0028940 |
| cluster3946 | -1805.9303 | 0.0253 | 0.0325 | 1.2848 | 999.0000 | 0.0020330 |
| cluster3947 | -2285.9949 | 0.0277 | 0.0431 | 1.5574 | 0.0545 | 0.0031700 |
| cluster3948 | -1016.9566 | 0.0284 | 0.0213 | 0.7485 | 1.1873 | 0.0000010 |
| cluster3949 | -878.24927 | 0.0093 | 0.0230 | 2.4762 | 3.4907 | 0.0000020 |
| cluster3950 | -1740.6701 | 0.0104 | 0.0202 | 1.9523 | 0.0230 | 0.0355970 |
| cluster3951 | -2834.1635 | 0.0401 | 0.0539 | 1.3423 | 0.0275 | 0.0057730 |

| | | | | | | |
|---|---|---|---|---|---|---|
| cluster3952 | -1474.835 | 0.0029 | 0.0065 | 2.2636 | 0.0071 | 0.0015880 |
| cluster3953 | -1363.1342 | 0.0061 | 0.0136 | 2.2363 | 0.0001 | 0.0000040 |
| cluster3954 | -1806.3118 | 0.0266 | 0.0245 | 0.9228 | 0.2094 | 0.0029000 |
| cluster3955 | -1192.2205 | 0.0252 | 0.0398 | 1.5805 | 0.3116 | 0.0000010 |
| cluster3956 | -1995.0784 | 0.0132 | 0.0267 | 2.0176 | 0.0432 | 0.0045760 |
| cluster3957 | -1955.7467 | 0.0099 | 0.0218 | 2.2020 | 0.0889 | 0.0100050 |
| cluster3958 | -1104.3836 | 0.0203 | 0.0268 | 1.3190 | 3.1458 | 0.0029910 |
| cluster3959 | -2686.0149 | 0.0354 | 0.0218 | 0.6154 | 0.0278 | 0.0233750 |
| cluster3960 | -2422.5772 | 0.0542 | 0.0738 | 1.3621 | 12.6306 | 0.0059210 |
| cluster3961 | -1594.3782 | 0.0393 | 0.0501 | 1.2746 | 0.0001 | 0.0000060 |
| cluster3962 | -1532.2647 | 0.0377 | 0.0419 | 1.1103 | 0.1418 | 0.0086310 |
| cluster3963 | -2073.1053 | 0.0328 | 0.0433 | 1.3227 | 0.3166 | 0.0200900 |
| cluster3964 | -2990.4727 | 0.0369 | 0.1981 | 5.3758 | 999.0000 | 0.0232720 |
| cluster3965 | -1514.5038 | 0.0270 | 0.0610 | 2.2557 | 0.8612 | 0.0194330 |
| cluster3966 | -1433.0606 | 0.0239 | 0.0187 | 0.7828 | 0.0001 | 0.0000000 |
| cluster3967 | -1752.9438 | 0.0768 | 0.0264 | 0.3441 | 0.1316 | 0.0603100 |
| cluster3968 | -1166.7897 | 0.0160 | 0.0171 | 1.0711 | 0.0884 | 0.0054930 |
| cluster3969 | -901.29213 | 0.0001 | 0.0000 | 0.3542 | 1.7073 | 0.0000010 |
| cluster3970 | -2484.0023 | 0.0471 | 0.0672 | 1.4254 | 0.0001 | 0.0000080 |
| cluster3971 | -1689.5985 | 0.0325 | 0.0439 | 1.3485 | 0.0674 | 0.0147740 |
| cluster3972 | -990.23326 | 0.0139 | 0.0185 | 1.3280 | 0.0001 | 0.0000000 |
| cluster3973 | -1419.8038 | 0.0099 | 0.0084 | 0.8524 | 0.0001 | 0.0000060 |
| cluster3974 | -1832.7845 | 0.0176 | 0.0192 | 1.0921 | 0.0279 | 0.0028470 |
| cluster3975 | -1269.7168 | 0.0300 | 0.0284 | 0.9473 | 0.0001 | 0.0000150 |
| cluster3976 | -1581.3422 | 0.0105 | 0.0106 | 1.0117 | 0.0047 | 0.0201080 |
| cluster3977 | -2227.0089 | 0.0837 | 0.0856 | 1.0227 | 0.7471 | 0.0245240 |
| cluster3978 | -889.04061 | 0.0055 | 0.0037 | 0.6643 | 0.6483 | 0.0000010 |
| cluster3979 | -1620.0079 | 0.0027 | 0.0022 | 0.7994 | 0.0001 | 0.0000990 |
| cluster3980 | -922.24867 | 0.0001 | 0.0000 | 0.1122 | 0.0001 | 0.0000470 |
| cluster3981 | -1978.7909 | 0.0160 | 0.0229 | 1.4286 | 1.8042 | 0.0115740 |
| cluster3982 | -1829.7161 | 0.0478 | 0.0520 | 1.0895 | 0.2030 | 0.0041700 |
| cluster3983 | -2677.4001 | 0.0335 | 0.0727 | 2.1684 | 0.0448 | 0.0026530 |
| cluster3984 | -1381.4342 | 0.0326 | 0.0848 | 2.6034 | 999.0000 | 0.0072380 |
| cluster3985 | -979.25641 | 0.0124 | 0.0039 | 0.3148 | 0.0112 | 0.0090310 |
| cluster3986 | -1733.0238 | 0.0136 | 0.0361 | 2.6498 | 0.1161 | 0.0076400 |
| cluster3987 | -2084.9046 | 0.0159 | 0.0146 | 0.9217 | 0.0037 | 0.0072540 |
| cluster3988 | -1234.7682 | 0.0041 | 0.0039 | 0.9440 | 0.6314 | 0.0126290 |
| cluster3989 | -1834.2869 | 0.0619 | 0.0475 | 0.7684 | 0.0361 | 0.0573620 |
| cluster3990 | -2206.7434 | 0.0284 | 0.0506 | 1.7782 | 0.0051 | 0.0017310 |
| cluster3991 | -1522.1499 | 0.0222 | 0.0335 | 1.5104 | 0.0231 | 0.0014770 |
| cluster3992 | -1889.115 | 0.0382 | 0.0150 | 0.3931 | 0.0327 | 0.0282200 |
| cluster3993 | -1674.8906 | 0.0358 | 0.0488 | 1.3608 | 0.0038 | 0.0004640 |
| cluster3994 | -2390.5579 | 0.0463 | 0.0455 | 0.9835 | 0.1432 | 0.1001810 |
| cluster3995 | -1618.995 | 0.0336 | 0.0368 | 1.0962 | 0.0627 | 0.0130310 |

| | | | | | | |
|---|---|---|---|---|---|---|
| cluster3996 | -1708.9928 | 0.0064 | 0.0174 | 2.7150 | 0.0001 | 0.0000000 |
| cluster3997 | -2516.5717 | 0.0480 | 0.0621 | 1.2928 | 0.1791 | 0.0202630 |
| cluster3998 | -1414.6457 | 0.0465 | 0.0406 | 0.8747 | 0.1881 | 0.0225130 |
| cluster3999 | -2524.2797 | 0.0297 | 0.0546 | 1.8380 | 999.0000 | 0.0052190 |
| cluster4000 | -2289.645 | 0.0434 | 0.0415 | 0.9563 | 0.0839 | 0.0184190 |
| cluster4001 | -1736.2571 | 0.0229 | 0.0240 | 1.0492 | 0.0001 | 0.0000620 |
| cluster4002 | -1801.3215 | 0.0056 | 0.0173 | 3.1198 | 0.0001 | 0.0000330 |
| cluster4003 | -1414.0545 | 0.0508 | 0.0670 | 1.3208 | 999.0000 | 0.0402600 |
| cluster4004 | -1442.7323 | 0.0077 | 0.0074 | 0.9615 | 0.0159 | 0.0019850 |
| cluster4005 | -2064.3959 | 0.0363 | 0.0789 | 2.1742 | 0.2896 | 0.0247070 |
| cluster4006 | -1089.5732 | 0.0320 | 0.0262 | 0.8171 | 0.0001 | 0.0000000 |
| cluster4007 | -1425.8071 | 0.0027 | 0.0024 | 0.8930 | 0.0001 | 0.0000120 |
| cluster4008 | -1355.5051 | 0.0072 | 0.0190 | 2.6414 | 0.0204 | 0.0032060 |
| cluster4009 | -1540.8125 | 0.0018 | 0.0319 | 18.0784 | 15.4676 | 0.0124430 |
| cluster4010 | -1626.5025 | 0.0486 | 0.0942 | 1.9394 | 0.1529 | 0.0206080 |
| cluster4011 | -1061.3988 | 0.0717 | 0.0140 | 0.1960 | 0.0396 | 0.00199 |
| cluster4012 | -2445.3946 | 0.0126 | 0.1281 | 10.1495 | 0.1055 | 0.0069330 |
| cluster4013 | -844.87902 | 0.0190 | 0.0113 | 0.5941 | 999.0000 | 0.0057130 |
| cluster4014 | -1137.9303 | 0.0063 | 0.0089 | 1.4104 | 0.0001 | 0.0000060 |
| cluster4015 | -618.38127 | 0.0311 | 0.0018 | 0.0565 | 0.0001 | 0.00000 |
| cluster4016 | -1814.0212 | 0.0158 | 0.0143 | 0.9100 | 0.0369 | 0.0304670 |
| cluster4017 | -1339.4212 | 0.0368 | 0.0496 | 1.3460 | 999.0000 | 0.0150660 |
| cluster4018 | -1325.475 | 0.1293 | 0.1491 | 1.1533 | 0.2235 | 0.03013 |
| cluster4019 | -1825.5613 | 0.0615 | 0.0867 | 1.4095 | 0.1475 | 0.0190150 |
| cluster4020 | -1005.3808 | 0.0351 | 0.0134 | 0.3816 | 0.0394 | 0.0197670 |
| cluster4021 | -1592.1283 | 0.0235 | 0.0138 | 0.5857 | 0.0687 | 0.0090600 |
| cluster4022 | -970.14394 | 0.0096 | 0.0070 | 0.7279 | 0.0001 | 0.0000050 |
| cluster4023 | -1843.5533 | 0.0626 | 0.0565 | 0.9023 | 0.3445 | 0.0066650 |
| cluster4024 | -2359.7194 | 0.0284 | 0.0697 | 2.4553 | 11.6218 | 0.0020660 |
| cluster4025 | -624.40744 | 0.0006 | 0.0019 | 2.9638 | 0.0001 | 0.0000000 |
| cluster4026 | -1088.9385 | 0.0027 | 0.0086 | 3.1657 | 0.0001 | 0.0000070 |
| cluster4027 | -979.9204 | 0.0056 | 0.0163 | 2.8947 | 0.0001 | 0.0000140 |
| cluster4028 | -1844.002 | 0.0052 | 0.0162 | 3.1218 | 89.4528 | 0.0052200 |
| cluster4029 | -2242.6815 | 0.0506 | 0.0830 | 1.6422 | 0.6058 | 0.0528700 |
| cluster4030 | -1125.0665 | 0.0380 | 0.0282 | 0.7435 | 0.0106 | 0.0004540 |
| cluster4031 | -1097.383 | 0.0325 | 0.0454 | 1.3966 | 0.0155 | 0.0215520 |
| cluster4032 | -1475.3667 | 0.0012 | 0.0005 | 0.4088 | 0.0001 | 0.0000080 |
| cluster4033 | -2213.1899 | 0.0594 | 0.0962 | 1.6185 | 0.2303 | 0.0332260 |
| cluster4034 | -493.6726 | 0.0001 | 0.0000 | 0.1080 | 0.0001 | 0.0003500 |
| cluster4035 | -2113.6352 | 0.0514 | 0.0263 | 0.5122 | 0.1096 | 0.0594150 |
| cluster4036 | -1076.9139 | 0.0080 | 0.0125 | 1.5657 | 0.0290 | 0.0028710 |
| cluster4037 | -1625.0274 | 0.0174 | 0.0329 | 1.8942 | 999.0000 | 0.0045790 |
| cluster4038 | -902.54419 | 0.0160 | 0.0369 | 2.3116 | 0.1953 | 0.0068320 |
| cluster4039 | -1508.2006 | 0.0595 | 0.0242 | 0.4071 | 0.1067 | 0.0034030 |

| | | | | | | |
|---|---|---|---|---|---|---|
| cluster4040 | -1126.3941 | 0.0390 | 0.0326 | 0.8340 | 0.0114 | 0.0042530 |
| cluster4041 | -1475.3952 | 0.0132 | 0.0236 | 1.7834 | 0.0210 | 0.0051250 |
| cluster4042 | -899.09489 | 0.0308 | 0.0256 | 0.8314 | 999.0000 | 0.0035900 |
| cluster4043 | -1138.5106 | 0.0484 | 0.1256 | 2.5976 | 0.0271 | 0.0045280 |
| cluster4044 | -958.06892 | 0.0014 | 0.0027 | 1.9265 | 0.0001 | 0.0000000 |
| cluster4045 | -2022.0265 | 0.0565 | 0.0803 | 1.4222 | 0.2188 | 0.0202010 |
| cluster4046 | -889.94517 | 0.0027 | 0.0014 | 0.5155 | 0.0001 | 0.0000020 |
| cluster4047 | -2066.3996 | 0.0147 | 0.0349 | 2.3759 | 0.2125 | 0.0113640 |
| cluster4048 | -1470.1346 | 0.0078 | 0.0043 | 0.5447 | 0.0230 | 0.0016230 |
| cluster4049 | -571.98909 | 0.0009 | 0.0039 | 4.3615 | 999.0000 | 0.00304 |
| cluster4050 | -1217.6531 | 0.0089 | 0.0078 | 0.8831 | 0.0001 | 0.0000000 |
| cluster4051 | -1404.1556 | 0.0089 | 0.0033 | 0.3670 | 0.0001 | 0.0000050 |
| cluster4052 | -1287.9158 | 0.0171 | 0.0081 | 0.4738 | 0.0266 | 0.0189430 |
| cluster4053 | -1638.9977 | 0.0075 | 0.0338 | 4.4899 | 0.2152 | 0.0132530 |
| cluster4054 | -1860.4834 | 0.0170 | 0.0396 | 2.3283 | 0.0474 | 0.0055920 |
| cluster4055 | -719.56954 | 0.0021 | 0.0163 | 7.7161 | 0.0091 | 0.0052360 |
| cluster4056 | -934.46282 | 0.0332 | 0.0305 | 0.9193 | 0.1536 | 0.0101030 |
| cluster4057 | -1565.0707 | 0.0336 | 0.0442 | 1.3158 | 0.0474 | 0.0078870 |
| cluster4058 | -1678.3287 | 0.0333 | 0.0529 | 1.5860 | 999.0000 | 0.0122240 |
| cluster4059 | -723.11767 | 0.0124 | 0.0278 | 2.2349 | 0.0001 | 0.0000000 |
| cluster4060 | -1875.2557 | 0.0257 | 0.0270 | 1.0476 | 0.0458 | 0.0155480 |
| cluster4061 | -1188.8132 | 0.0058 | 0.0139 | 2.4234 | 7.6272 | 0.0000020 |
| cluster4062 | -1215.6433 | 0.0176 | 0.0156 | 0.8897 | 0.0030 | 0.0024550 |
| cluster4063 | -1720.5739 | 0.0224 | 0.0640 | 2.8625 | 999.0000 | 0.0101680 |
| cluster4064 | -781.25398 | 0.0121 | 0.0158 | 1.3045 | 0.1228 | 0.0126760 |
| cluster4065 | -991.43446 | 0.0075 | 0.0059 | 0.7863 | 1.0194 | 0.0000010 |
| cluster4066 | -1616.9391 | 0.0457 | 0.0522 | 1.1426 | 0.0527 | 0.0037390 |
| cluster4067 | -418.64637 | 0.0201 | 0.0847 | 4.2059 | 0.0001 | 0.00000 |
| cluster4068 | -1353.0169 | 0.0213 | 0.0671 | 3.1452 | 0.0001 | 0.0000380 |
| cluster4069 | -1065.1345 | 0.0618 | 0.0361 | 0.5850 | 0.0639 | 0.0136660 |
| cluster4070 | -692.2985 | 0.0021 | 0.0300 | 14.1693 | 0.0566 | 0.0167340 |
| cluster4071 | -826.73717 | 0.0022 | 0.0049 | 2.2171 | 0.0001 | 0.0000180 |
| cluster4072 | -987.5946 | 0.0271 | 0.0284 | 1.0499 | 0.0001 | 0.0000170 |
| cluster4073 | -1884.9788 | 0.0266 | 0.0474 | 1.7830 | 1.6050 | 0.0122410 |
| cluster4074 | -1256.756 | 0.0331 | 0.0727 | 2.1978 | 0.0001 | 0.0000390 |
| cluster4075 | -710.42792 | 0.6801 | 3.4403 | 5.0583 | 0.8987 | 0.06112 |
| cluster4076 | -1650.344 | 0.0892 | 0.0406 | 0.4552 | 0.0883 | 0.0658410 |
| cluster4077 | -1679.9932 | 0.0430 | 0.0495 | 1.1508 | 0.0606 | 0.0050620 |
| cluster4078 | -1230.4251 | 0.0272 | 0.0211 | 0.7757 | 0.0048 | 0.0292920 |
| cluster4079 | -1206.4491 | 0.0018 | 0.0011 | 0.6017 | 0.0001 | 0.0002390 |
| cluster4080 | -734.91877 | 0.0209 | 0.0121 | 0.5778 | 0.0013 | 0.0458140 |
| cluster4081 | -1132.2501 | 0.0174 | 0.0166 | 0.9508 | 1.3587 | 0.0000010 |
| cluster4082 | -745.24847 | 0.0074 | 0.0774 | 10.4192 | 4.5682 | 0.0000020 |
| cluster4083 | -1694.1452 | 0.0359 | 0.0821 | 2.2852 | 1.3467 | 0.0625480 |

| | | | | | | |
|---|---|---|---|---|---|---|
| cluster4084 | -713.97297 | 0.0291 | 0.0080 | 0.2749 | 0.0028 | 0.0046800 |
| cluster4085 | -859.28584 | 0.0307 | 0.0108 | 0.3526 | 0.0125 | 0.00255 |
| cluster4086 | -1314.0939 | 0.0283 | 0.0509 | 1.8005 | 0.2341 | 0.0063590 |
| cluster4087 | -1201.2516 | 0.0091 | 0.0295 | 3.2405 | 999.0000 | 0.0151610 |
| cluster4088 | -1314.2655 | 0.0563 | 0.0243 | 0.4308 | 0.0922 | 0.0025770 |
| cluster4089 | -1508.0897 | 0.0052 | 0.0185 | 3.5395 | 999.0000 | 0.0045380 |
| cluster4090 | -754.04246 | 0.0298 | 0.0223 | 0.7488 | 0.1925 | 0.0132620 |
| cluster4091 | -953.99832 | 0.0319 | 0.0132 | 0.4125 | 0.0001 | 0.0000020 |
| cluster4092 | -1423.9751 | 0.0235 | 0.0479 | 2.0387 | 0.0912 | 0.0131180 |
| cluster4093 | -1004.9575 | 0.0145 | 0.0253 | 1.7387 | 0.1724 | 0.0106850 |
| cluster4094 | -375.44687 | 0.0091 | 0.0015 | 0.1689 | 0.0001 | 0.00000 |
| cluster4095 | -1403.8144 | 0.0369 | 0.0606 | 1.6412 | 0.0991 | 0.0679500 |
| cluster4096 | -1210.4822 | 0.0181 | 0.0402 | 2.2177 | 0.2523 | 0.0139600 |
| cluster4097 | -705.33004 | 0.0956 | 0.0118 | 0.1238 | 0.1022 | 0.00764 |
| cluster4098 | -1868.2825 | 0.0291 | 0.0363 | 1.2472 | 0.0575 | 0.0482430 |
| cluster4099 | -811.01553 | 0.0026 | 0.0040 | 1.5004 | 8.4638 | 0.0000020 |
| cluster4100 | -1048.5218 | 0.0074 | 0.0103 | 1.3880 | 0.0188 | 0.0028830 |
| cluster4101 | -2089.1469 | 0.2301 | 0.1105 | 0.4805 | 0.3127 | 0.17649 |
| cluster4102 | -1855.3465 | 0.0385 | 0.1682 | 4.3664 | 0.1483 | 0.0224880 |
| cluster4103 | -1107.0942 | 0.0466 | 0.0104 | 0.2221 | 0.0292 | 0.0100700 |
| cluster4104 | -1017.4099 | 0.0079 | 0.0140 | 1.7674 | 0.0001 | 0.0000260 |
| cluster4105 | -734.04163 | 0.0058 | 0.0575 | 9.8720 | 0.0441 | 0.0090740 |
| cluster4106 | -1000.5643 | 0.0083 | 0.0150 | 1.8016 | 0.0001 | 0.0000160 |
| cluster4107 | -786.70815 | 0.0025 | 0.0036 | 1.4555 | 0.0001 | 0.0000230 |
| cluster4108 | -783.34142 | 0.0100 | 0.0161 | 1.6019 | 0.0232 | 0.0213110 |
| cluster4109 | -1651.848 | 0.0688 | 0.1108 | 1.6102 | 0.0430 | 0.0046850 |
| cluster4110 | -1246.3732 | 0.0612 | 0.0773 | 1.2633 | 0.1328 | 0.0050610 |
| cluster4111 | -924.01349 | 0.0405 | 0.0740 | 1.8257 | 0.2914 | 0.0092960 |
| cluster4112 | -1112.3805 | 0.0125 | 0.0098 | 0.7776 | 0.0381 | 0.0196170 |
| cluster4113 | -773.37694 | 0.0061 | 0.0076 | 1.2318 | 0.0001 | 0.0000030 |
| cluster4114 | -1074.5579 | 0.0452 | 0.0307 | 0.6796 | 0.0001 | 0.0000090 |
| cluster4115 | -1155.47 | 0.0092 | 0.0183 | 1.9967 | 999.0000 | 0.0049260 |
| cluster4116 | -636.33055 | 0.0221 | 0.0529 | 2.3923 | 0.0612 | 0.07431 |
| cluster4117 | -660.63231 | 0.0129 | 0.0084 | 0.6479 | 7.2383 | 0.0000020 |
| cluster4118 | -1286.026 | 0.0249 | 0.0435 | 1.7482 | 0.1605 | 0.0263140 |
| cluster4119 | -925.98536 | 0.0172 | 0.0316 | 1.8360 | 0.0442 | 0.0073180 |
| cluster4120 | -964.81161 | 0.0019 | 0.0017 | 0.8654 | 0.1148 | 0.0063500 |
| cluster4121 | -1215.8556 | 0.0036 | 0.0300 | 8.4403 | 999.0000 | 0.0038100 |
| cluster4122 | -525.18541 | 0.0226 | 0.0018 | 0.0794 | 999.0000 | 0.00358 |
| cluster4123 | -468.58494 | 0.0075 | 0.0058 | 0.7816 | 0.0001 | 0.0000070 |
| cluster4124 | -905.64803 | 0.0044 | 0.0171 | 3.9138 | 4.1864 | 0.0000020 |
| cluster4125 | -1064.1639 | 0.0600 | 0.0454 | 0.7563 | 0.2726 | 0.15140 |
| cluster4126 | -762.7028 | 0.0779 | 0.0279 | 0.3587 | 0.2944 | 0.00674 |
| cluster4127 | -766.29365 | 0.0546 | 0.0423 | 0.7742 | 999.0000 | 0.0035080 |

| | | | | | | |
|---|---|---|---|---|---|---|
| cluster4128 | -989.3 | 0.0314 | 0.1066 | 3.3893 | 0.0804 | 0.0137530 |

| dS | Adjusted p value |
| --- | --- |
| 0.0124290 | 0.0354 |
| 1.0070340 | 1.27E-65 |
| 0.0980840 | 0.0004 |
| 0.0577360 | 1.02E-92 |
| 0.0692040 | 3.23E-07 |
| 0.0514280 | 2.56E-05 |
| 0.0761410 | 4.96E-54 |
| 0.0410070 | 4.09E-09 |
| 0.7849730 | 6.02E-124 |
| 0.4645810 | 1.25E-41 |
| 1.0995730 | 2.39E-88 |
| 0.1246950 | 4.55E-44 |
| 1.3340400 | 3.16E-107 |
| 0.6273850 | 2.08E-12 |
| 0.1064750 | 1.24E-48 |
| 0.0945480 | 1.47E-28 |
| 0.0003680 | 2.87E-08 |
| 0.0346040 | 7.24E-06 |
| 0.1259750 | 2.80E-06 |
| 0.0945060 | 1.04E-10 |
| 0.8129670 | 1.14E-59 |
| 0.0878390 | 3.32E-22 |
| 1.0155390 | 2.38E-64 |
| 0.9876900 | 7.43E-11 |
| 0.4931410 | 6.35E-21 |
| 0.3813310 | 6.14E-12 |
| 0.2076260 | 0.0047 |
| 0.1102990 | 1.08E-17 |
| 1.1092330 | 6.77E-19 |
| 0.2971480 | 7.90E-28 |
| 0.7805460 | 1.93E-07 |
| 0.0684350 | 1.16E-33 |
| 0.0807650 | 1.08E-32 |
| 0.9820220 | 8.07E-25 |
| 0.1245250 | 8.04E-55 |
| 0.1685790 | 6.03E-08 |
| 0.0604880 | 1.09E-46 |
| 0.23905 | 0.0600 |
| 0.0494240 | 1.26E-24 |
| 0.8066060 | 8.02E-35 |
| 0.1081340 | 3.42E-15 |
| 0.1335910 | 5.21E-17 |

| | |
|---|---|
| 0.1040780 | 3.75E-19 |
| 0.0639070 | 4.28E-66 |
| 0.0463100 | 2.07E-22 |
| 0.6237580 | 1.89E-14 |
| 1.0421720 | 1.28E-29 |
| 1.2862230 | 1.29E-39 |
| 0.0965660 | 1.65E-20 |
| 0.0880670 | 1.52E-13 |
| 1.6958630 | 1.35E-45 |
| 0.6429130 | 5.09E-05 |
| 0.7703010 | 3.98E-19 |
| 0.5657990 | 4.59E-16 |
| 0.0414780 | 1.67E-11 |
| 0.1191470 | 4.62E-14 |
| 0.0475120 | 6.47E-05 |
| 0.0436080 | 1.63E-20 |
| 0.3542990 | 4.48E-10 |
| 0.0412810 | 4.27E-24 |
| 0.2393670 | 5.62E-41 |
| 0.8176310 | 5.37E-25 |
| 0.0987310 | 4.60E-19 |
| 0.8190910 | 1.65E-26 |
| 0.1654010 | 7.91E-30 |
| 0.1056170 | 0.0001 |
| 0.5895260 | 3.28E-13 |
| 0.0862170 | 3.83E-30 |
| 0.1920130 | 6.80E-27 |
| 0.1469270 | 1.26E-09 |
| 0.03791 | 0.0600 |
| 0.1007920 | 5.48E-33 |
| 0.1469620 | 1.40E-14 |
| 1.0001600 | 4.28E-41 |
| 0.1746030 | 2.22E-11 |
| 0.1273450 | 1.02E-24 |
| 0.0908870 | 3.29E-52 |
| 0.0650960 | 9.56E-10 |
| 0.1295690 | 2.87E-10 |
| 0.2510980 | 1.65E-12 |
| 0.7200580 | 2.90E-14 |
| 0.0626480 | 6.31E-12 |
| 1.2983790 | 4.34E-14 |
| 0.0423920 | 2.16E-24 |
| 0.5377110 | 1.20E-12 |
| 0.6822250 | 4.40E-29 |

| | |
|---|---|
| 0.6078680 | 1.48E-06 |
| 0.0635650 | 6.27E-15 |
| 0.2818800 | 1.34E-06 |
| 0.1763670 | 1.05E-16 |
| 1.0930790 | 1.18E-23 |
| 0.4777800 | 8.89E-06 |
| 0.0776450 | 3.99E-21 |
| 0.4044730 | 1.46E-12 |
| 0.1016480 | 4.47E-21 |
| 0.9033330 | 3.63E-16 |
| 0.6200270 | 2.03E-23 |
| 0.5125990 | 2.81E-26 |
| 0.2600760 | 1.44E-10 |
| 0.0893420 | 4.02E-22 |
| 0.2630760 | 9.85E-15 |
| 2.4071260 | 2.73E-18 |
| 0.0000020 | 1.28E-15 |
| 1.1998620 | 4.89E-38 |
| 0.0662480 | 3.52E-06 |
| 0.1197370 | 7.61E-48 |
| 0.7474200 | 2.79E-09 |
| 0.9287140 | 4.58E-36 |
| 2.2649380 | 9.46E-47 |
| 2.1570110 | 7.06E-21 |
| 0.4917650 | 2.11E-14 |
| 0.4704610 | 7.85E-14 |
| 0.1388040 | 4.76E-32 |
| 0.1947540 | 4.61E-18 |
| 0.7137800 | 6.04E-40 |
| 0.3383830 | 9.04E-22 |
| 0.0912040 | 1.16E-15 |
| 0.1404990 | 3.44E-30 |
| 0.1523900 | 2.09E-05 |
| 0.5342170 | 6.38E-17 |
| 0.0134460 | 7.47E-06 |
| 0.0516510 | 2.04E-08 |
| 0.1798440 | 4.77E-11 |
| 0.07193 | 0.4772 |
| 0.0708460 | 4.42E-34 |
| 1.1899140 | 4.15E-57 |
| 0.0672070 | 6.69E-19 |
| 0.1939670 | 2.54E-05 |
| 0.0621640 | 4.78E-11 |
| 0.1241680 | 4.38E-05 |

| | |
|---|---|
| 0.0383780 | 1.38E-26 |
| 0.4624370 | 1.04E-12 |
| 0.43427 | 0.2363 |
| 0.0000080 | 0.0092 |
| 0.3143280 | 9.57E-14 |
| 0.0839230 | 9.88E-20 |
| 0.6069930 | 3.97E-13 |
| 0.1846850 | 4.75E-14 |
| 0.0669800 | 9.40E-08 |
| 0.2334160 | 5.64E-12 |
| 1.0119860 | 7.98E-17 |
| 0.0989630 | 2.89E-10 |
| 0.00000 | 0.1000 |
| 0.7708580 | 4.70E-16 |
| 0.1698190 | 5.91E-11 |
| 0.0976260 | 5.93E-27 |
| 0.1538940 | 1.02E-15 |
| 0.0295800 | 5.43E-18 |
| 0.2814450 | 5.20E-16 |
| 0.3185160 | 5.02E-22 |
| 0.0808240 | 1.83E-06 |
| 0.7330010 | 3.51E-05 |
| 0.0242280 | 4.28E-05 |
| 1.1448080 | 6.31E-10 |
| 0.1452720 | 1.86E-23 |
| 0.1133710 | 1.07E-16 |
| 0.1306900 | 9.04E-10 |
| 0.0303910 | 0.0042 |
| 2.8193580 | 1.05E-16 |
| 0.3034560 | 2.07E-07 |
| 0.4691700 | 1.41E-06 |
| 0.0582710 | 1.29E-24 |
| 0.0859580 | 0.0002 |
| 0.0662690 | 7.69E-09 |
| 0.1204080 | 3.45E-20 |
| 0.0808900 | 4.76E-32 |
| 0.5757300 | 1.11E-11 |
| 0.0413910 | 2.82E-15 |
| 0.7071710 | 2.31E-10 |
| 0.0976330 | 0.0002 |
| 0.8362340 | 1.21E-19 |
| 0.0525660 | 0.0217 |
| 0.1024570 | 1.29E-12 |
| 0.11171 | 0.1418 |

| | |
|---|---|
| 1.1649560 | 1.39E-30 |
| 0.0661560 | 2.86E-12 |
| 0.1034160 | 9.27E-15 |
| 0.5186560 | 3.50E-08 |
| 1.0464940 | 6.26E-07 |
| 0.7774280 | 3.29E-22 |
| 0.01520 | 0.1509 |
| 0.4427080 | 0.0051 |
| 0.5336690 | 4.13E-25 |
| 0.1580500 | 4.15E-08 |
| 0.0838190 | 9.03E-07 |
| 0.0172550 | 4.36E-07 |
| 0.5063300 | 2.81E-11 |
| 0.3681670 | 3.52E-14 |
| 0.1044960 | 4.06E-15 |
| 0.0165440 | 4.93E-12 |
| 0.0835190 | 6.64E-09 |
| 0.1882560 | 1.52E-16 |
| 0.2717700 | 6.39E-11 |
| 0.2250750 | 0.0174 |
| 0.8813770 | 1.03E-08 |
| 0.0815270 | 3.24E-17 |
| 0.9212300 | 4.35E-31 |
| 0.0980890 | 2.11E-14 |
| 0.1151580 | 4.64E-14 |
| 0.0776150 | 1.08E-13 |
| 0.8553080 | 1.31E-07 |
| 0.0849800 | 1.25E-32 |
| 0.0497350 | 4.28E-08 |
| 0.3583790 | 9.98E-05 |
| 0.5288900 | 0.0500 |
| 0.5942430 | 9.89E-06 |
| 0.1394960 | 2.18E-06 |
| 0.2114190 | 1.27E-11 |
| 0.5987470 | 2.33E-17 |
| 0.1227050 | 7.30E-10 |
| 0.2499350 | 6.59E-17 |
| 0.0643590 | 3.31E-05 |
| 0.1240950 | 0.0167 |
| 0.8053810 | 4.85E-08 |
| 0.9829020 | 7.49E-11 |
| 0.2987620 | 0.0025 |
| 0.0090420 | 0.0009 |
| 0.8094950 | 2.96E-13 |

| | |
|---|---|
| 0.6238510 | 1.34E-17 |
| 0.1110480 | 1.68E-07 |
| 0.0739690 | 9.89E-15 |
| 0.0668540 | 1.31E-13 |
| 0.8648600 | 7.31E-13 |
| 0.6352870 | 0.0008 |
| 0.7761750 | 2.72E-16 |
| 0.3135970 | 1.87E-16 |
| 0.9854400 | 7.69E-20 |
| 0.1446470 | 9.16E-21 |
| 0.9110570 | 4.24E-18 |
| 0.1686800 | 5.33E-10 |
| 0.8056800 | 3.32E-11 |
| 0.2339360 | 0.0023 |
| 0.2184270 | 2.55E-20 |
| 0.4186790 | 0.0003 |
| 0.0932010 | 2.56E-20 |
| 0.1223100 | 8.13E-09 |
| 0.7837900 | 0.0001 |
| 0.2931270 | 7.24E-28 |
| 0.4852290 | 2.86E-07 |
| 0.1063990 | 3.61E-14 |
| 0.0000020 | 7.90E-16 |
| 0.1205610 | 5.69E-21 |
| 1.3160760 | 4.50E-27 |
| 0.0907610 | 1.93E-14 |
| 0.2946740 | 8.35E-19 |
| 0.0682420 | 3.26E-11 |
| 0.6073800 | 5.13E-09 |
| 0.1211250 | 1.57E-09 |
| 0.3775840 | 1.80E-08 |
| 1.1031170 | 1.94E-12 |
| 0.2181520 | 2.03E-09 |
| 0.5107430 | 3.52E-15 |
| 0.1787130 | 1.07E-16 |
| 0.7176900 | 2.75E-23 |
| 0.0711310 | 2.84E-05 |
| 0.0586550 | 0.0105 |
| 0.1104020 | 2.18E-11 |
| 0.0562750 | 0.0003 |
| 0.0832240 | 1.78E-11 |
| 0.2366120 | 5.30E-10 |
| 0.1385480 | 2.26E-10 |
| 0.1356670 | 3.85E-10 |

| | |
|---|---|
| 0.1564820 | 4.34E-14 |
| 0.1266110 | 1.44E-06 |
| 0.0948340 | 1.14E-12 |
| 0.0148520 | 0.0122 |
| 2.2603850 | 6.21E-06 |
| 0.1312810 | 6.17E-16 |
| 1.1882880 | 2.86E-18 |
| 0.6590870 | 2.44E-10 |
| 0.6503750 | 7.03E-15 |
| 0.1286350 | 4.41E-14 |
| 0.0899290 | 4.38E-05 |
| 0.6990350 | 1.18E-15 |
| 0.1173260 | 3.36E-09 |
| 0.0426940 | 5.61E-10 |
| 0.0713800 | 7.47E-06 |
| 0.0589270 | 2.84E-14 |
| 0.1086970 | 2.15E-10 |
| 0.7140550 | 1.44E-15 |
| 0.1131830 | 2.22E-08 |
| 0.4082040 | 2.51E-05 |
| 0.0434060 | 3.60E-12 |
| 0.0841780 | 3.04E-09 |
| 0.1028720 | 1.28E-11 |
| 0.6078880 | 9.62E-09 |
| 0.3653340 | 4.04E-06 |
| 0.0093860 | 1.38E-05 |
| 0.4827380 | 3.66E-08 |
| 0.01612 | 0.1723 |
| 0.5929770 | 7.48E-12 |
| 0.0243250 | 6.38E-12 |
| 0.2663080 | 8.72E-51 |
| 2.5946640 | 2.12E-13 |
| 0.0692710 | 6.83E-27 |
| 0.0950930 | 3.83E-14 |
| 0.5476380 | 9.71E-10 |
| 0.0191700 | 8.89E-05 |
| 0.0000000 | 5.23E-06 |
| 0.1084240 | 1.71E-13 |
| 0.0733030 | 1.01E-08 |
| 0.4240180 | 1.21E-27 |
| 0.0483690 | 3.42E-17 |
| 0.1219360 | 1.81E-16 |
| 0.0335360 | 0.0167 |
| 0.9823750 | 7.04E-16 |

| | |
|---|---|
| 0.0383870 | 2.10E-10 |
| 0.07971 | 0.1157 |
| 0.0885710 | 6.06E-11 |
| 0.0332360 | 8.53E-11 |
| 0.0719210 | 2.38E-08 |
| 0.1127740 | 3.48E-09 |
| 0.2130770 | 6.68E-11 |
| 0.5710580 | 5.15E-16 |
| 0.0340140 | 8.61E-05 |
| 1.2952910 | 5.57E-30 |
| 0.1545960 | 3.32E-14 |
| 0.0000090 | 0.0208 |
| 0.7959960 | 1.85E-07 |
| 1.0014660 | 5.71E-17 |
| 0.1268790 | 1.20E-20 |
| 0.0695410 | 5.04E-09 |
| 0.0721240 | 9.11E-07 |
| 0.2074000 | 2.39E-07 |
| 0.3254120 | 0.0004 |
| 0.9911930 | 1.55E-41 |
| 0.1013830 | 2.62E-05 |
| 0.0000070 | 3.88E-07 |
| 0.7987330 | 6.98E-25 |
| 0.9380840 | 4.09E-09 |
| 0.3634640 | 0.0001 |
| 0.0377790 | 8.59E-09 |
| 0.1048030 | 5.96E-18 |
| 0.0642490 | 3.98E-06 |
| 0.1376530 | 1.19E-17 |
| 0.6810160 | 0.0005 |
| 0.1175490 | 1.11E-10 |
| 0.1783660 | 7.00E-14 |
| 1.4391620 | 7.06E-05 |
| 0.7555010 | 4.42E-10 |
| 0.0847840 | 8.61E-07 |
| 0.3868820 | 3.90E-09 |
| 1.4890180 | 7.80E-18 |
| 0.3062800 | 6.04E-06 |
| 0.4888680 | 1.24E-07 |
| 0.1387420 | 1.41E-20 |
| 0.0798400 | 1.07E-11 |
| 1.7611870 | 7.12E-20 |
| 0.0667350 | 0.0029 |
| 0.0863870 | 8.34E-11 |

| | |
|---|---|
| 0.1316120 | 0.0001 |
| 0.3077260 | 0.0026 |
| 0.2394880 | 4.03E-05 |
| 0.1609840 | 1.02E-09 |
| 2.0708970 | 5.70E-26 |
| 0.6210270 | 1.64E-09 |
| 0.0684300 | 2.14E-19 |
| 0.0000000 | 3.40E-21 |
| 0.0138060 | 1.01E-09 |
| 1.1912990 | 1.35E-12 |
| 0.8194880 | 0.0001 |
| 0.6690570 | 6.85E-12 |
| 0.0220270 | 0.0003 |
| 0.0363320 | 1.28E-10 |
| 0.6718920 | 9.27E-08 |
| 0.0865200 | 3.93E-14 |
| 0.0768190 | 1.24E-23 |
| 0.7049530 | 3.98E-06 |
| 0.0684920 | 5.00E-14 |
| 0.3877720 | 7.08E-06 |
| 0.0385930 | 0.0023 |
| 0.0431110 | 8.56E-06 |
| 0.1262930 | 5.18E-17 |
| 0.1351070 | 1.63E-08 |
| 0.0945910 | 3.85E-06 |
| 0.0857720 | 4.29E-10 |
| 0.7154850 | 4.25E-09 |
| 0.8022460 | 1.61E-14 |
| 0.1320810 | 1.67E-11 |
| 0.0856840 | 1.05E-18 |
| 1.6483450 | 4.45E-08 |
| 0.7889780 | 2.81E-07 |
| 0.2179460 | 5.64E-12 |
| 0.1154560 | 3.82E-12 |
| 0.2374570 | 3.08E-05 |
| 0.0556350 | 0.0029 |
| 0.1117440 | 2.41E-05 |
| 1.1779540 | 2.95E-19 |
| 0.2239100 | 1.25E-07 |
| 0.1644560 | 3.05E-09 |
| 0.5903830 | 0.0133 |
| 0.8018790 | 2.13E-10 |
| 0.0295460 | 1.45E-21 |
| 1.4301600 | 3.06E-11 |

| | |
|---|---|
| 0.0376160 | 0.0361 |
| 0.1692420 | 7.99E-13 |
| 0.1084610 | 7.24E-14 |
| 0.2153520 | 0.0073 |
| 0.0556480 | 1.30E-07 |
| 0.2580030 | 5.58E-08 |
| 0.0914010 | 2.18E-05 |
| 0.0729470 | 5.94E-09 |
| 0.2064360 | 2.86E-20 |
| 0.2715330 | 3.74E-13 |
| 1.0815350 | 1.53E-29 |
| 0.0000040 | 3.72E-08 |
| 0.4704270 | 1.36E-05 |
| 0.9469820 | 2.15E-18 |
| 0.0420410 | 2.09E-05 |
| 0.1683110 | 3.15E-22 |
| 0.04266 | 0.1000 |
| 0.0232660 | 2.63E-05 |
| 0.0808710 | 8.07E-31 |
| 0.0558340 | 0.0001 |
| 0.5732610 | 9.34E-11 |
| 0.1707380 | 4.44E-07 |
| 0.1058070 | 1.49E-05 |
| 0.0867970 | 5.84E-06 |
| 0.4328750 | 4.83E-16 |
| 0.00686 | 0.7227 |
| 0.0985250 | 0.0035 |
| 0.1017340 | 1.60E-05 |
| 0.1146850 | 1.37E-08 |
| 0.2822400 | 0.0203 |
| 0.0000520 | 7.43E-05 |
| 0.0232910 | 3.09E-17 |
| 0.1303230 | 1.01E-13 |
| 0.0729000 | 3.01E-11 |
| 0.1505350 | 5.58E-08 |
| 0.0917260 | 1.05E-11 |
| 0.8833760 | 1.58E-07 |
| 0.0540300 | 7.41E-15 |
| 0.0582560 | 3.31E-07 |
| 0.0401320 | 0.0010 |
| 0.0994250 | 2.16E-06 |
| 0.1557720 | 8.62E-09 |
| 0.4238250 | 1.06E-09 |
| 0.2950860 | 0.0011 |

| | |
|---|---|
| 0.8883720 | 6.29E-12 |
| 0.0569730 | 4.64E-07 |
| 0.6722580 | 1.66E-19 |
| 0.9559280 | 1.16E-10 |
| 0.0819340 | 1.61E-10 |
| 0.1509290 | 2.47E-10 |
| 0.42518 | 0.1617 |
| 1.3317990 | 6.58E-22 |
| 0.0000120 | 1.22E-06 |
| 0.0000030 | 5.24E-15 |
| 0.3959240 | 0.0009 |
| 0.2095400 | 3.31E-05 |
| 1.5211590 | 1.74E-11 |
| 0.6159340 | 6.62E-07 |
| 0.1424260 | 3.51E-05 |
| 0.1185200 | 8.46E-06 |
| 0.0735920 | 4.14E-12 |
| 1.2537880 | 1.55E-18 |
| 0.4413140 | 0.0174 |
| 0.6993200 | 2.76E-15 |
| 0.7751650 | 5.43E-10 |
| 0.0000120 | 1.14E-08 |
| 0.5815400 | 1.68E-07 |
| 0.1210110 | 8.83E-11 |
| 1.2574080 | 1.80E-21 |
| 0.0458550 | 1.49E-05 |
| 0.0000020 | 1.62E-06 |
| 2.0563630 | 0.0249 |
| 0.0485800 | 3.44E-10 |
| 0.0509620 | 0.0001 |
| 0.0313830 | 1.49E-09 |
| 0.1920450 | 1.74E-07 |
| 0.1816390 | 8.76E-10 |
| 0.9090630 | 0.0011 |
| 0.2940660 | 4.45E-08 |
| 0.00001 | 0.0700 |
| 0.8626160 | 5.79E-13 |
| 0.0000000 | 1.31E-06 |
| 0.2627590 | 1.02E-05 |
| 0.0538430 | 8.71E-06 |
| 0.0668850 | 0.0017 |
| 0.1008640 | 3.86E-09 |
| 0.1505940 | 2.28E-11 |
| 0.0713080 | 0.0362 |

| | |
|---|---|
| 1.1786320 | 4.26E-09 |
| 0.5201390 | 3.57E-13 |
| 0.3635680 | 1.36E-05 |
| 0.0617240 | 2.32E-12 |
| 1.0853380 | 7.42E-05 |
| 0.02789 | 0.4744 |
| 0.1253020 | 1.98E-08 |
| 0.0212810 | 3.32E-09 |
| 0.0711210 | 6.60E-24 |
| 0.4027580 | 4.18E-08 |
| 0.0878900 | 1.28E-15 |
| 0.6191670 | 4.74E-23 |
| 0.5590180 | 2.04E-07 |
| 0.1534650 | 1.64E-06 |
| 3.7232250 | 2.12E-17 |
| 0.1095080 | 6.88E-09 |
| 0.0782850 | 3.32E-11 |
| 0.0627690 | 1.61E-05 |
| 0.0602540 | 1.61E-05 |
| 0.0325850 | 7.05E-07 |
| 0.2954980 | 8.21E-09 |
| 0.7340190 | 2.88E-09 |
| 0.0874570 | 0.0002 |
| 0.6621910 | 6.41E-26 |
| 0.1671620 | 2.60E-13 |
| 1.0252770 | 7.63E-06 |
| 0.1220490 | 7.26E-09 |
| 0.7708930 | 1.68E-15 |
| 0.0000010 | 8.02E-35 |
| 0.1422900 | 1.08E-10 |
| 0.0424260 | 7.87E-09 |
| 0.0350510 | 2.61E-07 |
| 0.6894250 | 0.0104 |
| 0.2253050 | 0.0014 |
| 0.1660300 | 1.30E-16 |
| 0.1055390 | 6.31E-05 |
| 0.0232270 | 1.56E-11 |
| 1.8294250 | 7.70E-18 |
| 0.1509940 | 4.24E-34 |
| 0.1044500 | 8.52E-06 |
| 0.8630140 | 2.88E-09 |
| 0.4608570 | 0.0078 |
| 0.0686610 | 1.85E-15 |
| 1.2092250 | 6.99E-12 |

| | |
|---|---|
| 0.0493540 | 0.0002 |
| 0.0000570 | 1.10E-06 |
| 0.9612380 | 9.63E-06 |
| 0.0147720 | 0.0086 |
| 0.0439160 | 2.86E-07 |
| 0.6102940 | 6.72E-05 |
| 0.1567710 | 3.25E-13 |
| 0.0732100 | 2.72E-20 |
| 0.2571030 | 4.99E-07 |
| 0.1882740 | 1.72E-09 |
| 0.1178550 | 3.93E-09 |
| 0.5671450 | 1.30E-06 |
| 0.0763480 | 2.30E-11 |
| 0.2094360 | 2.13E-12 |
| 0.1109300 | 0.0002 |
| 1.8067540 | 1.88E-06 |
| 1.1759060 | 6.71E-13 |
| 0.8329230 | 1.23E-12 |
| 0.1069030 | 4.97E-15 |
| 0.0591300 | 6.71E-06 |
| 1.2085290 | 3.05E-07 |
| 0.23856 | 0.1255 |
| 0.0256130 | 1.12E-08 |
| 0.7454140 | 1.07E-14 |
| 0.2624180 | 3.00E-11 |
| 0.10854 | 0.1113 |
| 0.06072 | 0.0700 |
| 1.2855210 | 6.01E-15 |
| 1.6665640 | 2.97E-08 |
| 0.1920560 | 3.31E-06 |
| 0.2062490 | 2.93E-14 |
| 0.5133720 | 0.0005 |
| 0.0375520 | 5.24E-15 |
| 0.0586240 | 0.0009 |
| 0.0340550 | 8.56E-06 |
| 0.34947 | 0.1297 |
| 0.0671150 | 2.56E-05 |
| 9.8976470 | 0.0348 |
| 54.6922040 | 3.36E-10 |
| 0.2813300 | 0.0002 |
| 0.05482 | 0.0600 |
| 0.18612 | 0.3567 |
| 0.1341670 | 1.64E-20 |
| 0.6233580 | 6.86E-14 |

| | |
|---|---|
| 0.0341900 | 2.67E-08 |
| 0.5664770 | 0.0080 |
| 0.0243000 | 0.0023 |
| 1.3826680 | 1.78E-15 |
| 0.0931850 | 0.0010 |
| 0.8474700 | 0.0004 |
| 0.8466680 | 7.32E-13 |
| 0.2201660 | 7.56E-09 |
| 0.2720240 | 4.68E-10 |
| 0.5763450 | 3.36E-08 |
| 0.2414740 | 0.0256 |
| 0.5899220 | 3.90E-07 |
| 0.1050840 | 2.65E-15 |
| 0.0585780 | 5.33E-12 |
| 1.2841560 | 7.03E-15 |
| 0.1610000 | 1.07E-08 |
| 0.0597070 | 0.0022 |
| 0.0000020 | 6.41E-06 |
| 0.0384900 | 8.31E-06 |
| 0.0629840 | 8.27E-08 |
| 0.0324420 | 7.37E-13 |
| 0.0687070 | 2.35E-14 |
| 0.6105340 | 3.26E-11 |
| 0.0103000 | 4.37E-12 |
| 0.0065060 | 7.51E-12 |
| 0.0700920 | 9.25E-08 |
| 0.2072330 | 0.0415 |
| 0.0656440 | 1.54E-09 |
| 0.1368500 | 2.72E-10 |
| 0.0579100 | 6.42E-07 |
| 0.8240160 | 0.0003 |
| 0.3619160 | 1.26E-15 |
| 0.3988030 | 4.25E-05 |
| 0.1995940 | 2.44E-08 |
| 0.2325310 | 5.92E-10 |
| 0.7063250 | 0.0003 |
| 0.0000290 | 1.35E-05 |
| 0.1268440 | 1.22E-12 |
| 0.0770180 | 0.0004 |
| 0.04334 | 0.1900 |
| 0.2132830 | 1.85E-08 |
| 0.1133550 | 1.26E-09 |
| 1.5828600 | 2.93E-18 |
| 2.1316590 | 2.49E-13 |

| | |
|---|---|
| 0.1063170 | 7.27E-10 |
| 0.1726340 | 2.97E-11 |
| 0.2917320 | 2.39E-08 |
| 0.0878740 | 4.16E-16 |
| 0.1503170 | 8.69E-09 |
| 0.7019400 | 2.89E-15 |
| 1.1541140 | 5.50E-11 |
| 0.1001840 | 7.49E-11 |
| 0.4138440 | 1.01E-08 |
| 0.0000130 | 5.05E-12 |
| 0.4307560 | 7.64E-08 |
| 1.0164870 | 3.99E-13 |
| 0.0585990 | 0.0002 |
| 0.1082480 | 9.68E-11 |
| 76.5802430 | 0.0008 |
| 0.0264610 | 0.0422 |
| 0.1270970 | 6.44E-19 |
| 0.0651780 | 8.84E-09 |
| 0.0405720 | 7.28E-06 |
| 0.4385520 | 8.38E-05 |
| 0.3703070 | 7.13E-07 |
| 0.1789670 | 0.0003 |
| 0.1030090 | 4.66E-08 |
| 0.0696850 | 7.74E-12 |
| 0.0971970 | 4.11E-09 |
| 0.0976160 | 2.04E-08 |
| 0.0812060 | 1.08E-10 |
| 2.9446040 | 1.27E-10 |
| 0.0757490 | 2.35E-08 |
| 0.7465860 | 6.31E-11 |
| 0.3447810 | 0.0172 |
| 0.1059160 | 5.93E-07 |
| 0.0567030 | 0.0346 |
| 0.0557760 | 1.56E-08 |
| 1.3059080 | 2.98E-06 |
| 0.1694020 | 6.73E-10 |
| 0.2030730 | 0.0013 |
| 4.6275270 | 7.64E-08 |
| 0.0970420 | 3.43E-05 |
| 0.0780850 | 7.40E-11 |
| 0.0753020 | 7.26E-09 |
| 0.3179010 | 9.34E-11 |
| 0.0000000 | 0.0002 |
| 0.17919 | 0.3354 |

| | |
|---|---|
| 0.0260860 | 0.0181 |
| 0.0291310 | 3.81E-05 |
| 0.08408 | 0.4027 |
| 0.7270470 | 2.31E-12 |
| 0.1649460 | 8.94E-12 |
| 0.0988270 | 4.76E-12 |
| 0.0000000 | 6.53E-10 |
| 0.1304610 | 4.28E-05 |
| 0.2085120 | 2.93E-05 |
| 0.09370 | 0.2681 |
| 0.1470540 | 3.02E-09 |
| 0.1110700 | 2.00E-10 |
| 0.1421610 | 2.86E-05 |
| 0.0619550 | 4.16E-06 |
| 0.0993030 | 7.88E-05 |
| 0.0014770 | 5.56E-24 |
| 0.1315130 | 1.63E-08 |
| 0.7442350 | 7.89E-08 |
| 0.1189820 | 7.66E-10 |
| 0.1669030 | 2.80E-05 |
| 0.0272690 | 5.71E-06 |
| 1.1172250 | 1.92E-07 |
| 0.6263280 | 4.97E-07 |
| 0.4357230 | 0.0104 |
| 0.0306420 | 2.98E-10 |
| 1.2590580 | 5.72E-09 |
| 0.0815350 | 1.51E-24 |
| 0.5273950 | 0.0029 |
| 0.9516460 | 4.36E-06 |
| 0.1158460 | 0.0002 |
| 0.8322770 | 4.85E-11 |
| 0.1211910 | 1.37E-06 |
| 0.1069420 | 3.09E-09 |
| 0.5281580 | 0.0048 |
| 0.0930100 | 6.62E-10 |
| 0.1123590 | 1.03E-07 |
| 0.7741190 | 0.0055 |
| 0.2607290 | 0.0401 |
| 0.9877070 | 1.96E-08 |
| 0.0472120 | 0.0028 |
| 0.3319370 | 0.0009 |
| 0.7574320 | 5.40E-08 |
| 0.1187160 | 1.59E-07 |
| 0.0444270 | 5.09E-10 |

| | |
|---|---|
| 0.1031760 | 8.23E-07 |
| 0.0910500 | 4.14E-14 |
| 0.14112 | 0.0800 |
| 0.4054720 | 5.39E-07 |
| 0.0468690 | 3.29E-08 |
| 0.0965460 | 8.95E-08 |
| 0.0621320 | 0.0006 |
| 0.1008190 | 1.55E-10 |
| 1.1303110 | 1.98E-07 |
| 0.8362530 | 5.42E-07 |
| 0.0000330 | 3.60E-09 |
| 0.1092480 | 6.49E-05 |
| 0.2703400 | 0.0054 |
| 0.0700380 | 1.29E-07 |
| 0.0202440 | 1.60E-06 |
| 0.1603970 | 5.25E-07 |
| 0.7367680 | 2.83E-07 |
| 0.2407860 | 3.85E-06 |
| 0.2185930 | 1.81E-11 |
| 0.2131100 | 4.26E-06 |
| 0.1847170 | 8.57E-09 |
| 0.1139330 | 4.37E-07 |
| 0.5494060 | 8.55E-06 |
| 0.2719820 | 7.61E-08 |
| 1.7655780 | 4.20E-08 |
| 0.0000070 | 4.76E-12 |
| 0.0000000 | 3.00E-07 |
| 0.2776400 | 0.0133 |
| 0.1063710 | 2.82E-18 |
| 0.0573160 | 5.72E-10 |
| 0.4235220 | 0.0033 |
| 0.2372470 | 3.69E-06 |
| 0.0204740 | 3.02E-05 |
| 0.1723580 | 1.51E-07 |
| 0.0449500 | 4.50E-07 |
| 0.2145530 | 0.0002 |
| 1.0234680 | 1.44E-10 |
| 0.0659080 | 1.98E-10 |
| 0.0931410 | 1.03E-07 |
| 0.0441380 | 0.0002 |
| 0.6284950 | 1.65E-08 |
| 0.0814280 | 9.78E-07 |
| 0.4927520 | 1.12E-11 |
| 0.2924180 | 8.46E-08 |

| | |
|---|---|
| 0.0000140 | 2.41E-08 |
| 0.1017680 | 7.48E-08 |
| 0.0000960 | 1.33E-18 |
| 0.7755860 | 6.98E-14 |
| 0.0291090 | 9.52E-09 |
| 0.02259 | 0.2365 |
| 0.1990920 | 4.95E-07 |
| 0.1095070 | 3.26E-13 |
| 0.1509140 | 2.01E-06 |
| 0.9071190 | 3.80E-12 |
| 0.2798560 | 1.70E-09 |
| 0.0000000 | 8.74E-05 |
| 0.0235930 | 2.54E-05 |
| 0.1402600 | 3.01E-07 |
| 0.9941680 | 1.95E-16 |
| 0.0284030 | 2.55E-09 |
| 0.3292250 | 0.0126 |
| 0.0000110 | 0.0042 |
| 0.1033480 | 0.0005 |
| 0.03446 | 0.3177 |
| 0.0929470 | 4.98E-16 |
| 0.2565370 | 1.08E-07 |
| 0.0923610 | 1.04E-12 |
| 0.0893710 | 4.48E-10 |
| 0.2170830 | 3.60E-09 |
| 0.0558710 | 1.97E-10 |
| 0.1674540 | 2.66E-17 |
| 1.2968590 | 1.13E-06 |
| 0.1161670 | 1.35E-19 |
| 0.2159020 | 2.97E-08 |
| 0.0499650 | 9.74E-23 |
| 0.1973270 | 0.0008 |
| 0.9787900 | 7.40E-15 |
| 3.3947560 | 3.62E-07 |
| 5.7905060 | 1.54E-06 |
| 0.0066440 | 0.0031 |
| 0.2912990 | 0.0203 |
| 2.2847940 | 2.60E-10 |
| 0.0000040 | 0.0089 |
| 0.0820360 | 4.09E-06 |
| 0.5058610 | 0.0017 |
| 0.1466560 | 9.12E-05 |
| 0.4507460 | 0.0002 |
| 0.1886940 | 9.81E-08 |

| | |
|---|---|
| 0.0592540 | 0.0029 |
| 0.1936270 | 8.38E-05 |
| 0.1236230 | 3.47E-06 |
| 0.0000040 | 5.24E-08 |
| 0.0002860 | 8.07E-25 |
| 2.7942150 | 3.47E-16 |
| 0.0304630 | 2.10E-06 |
| 0.0000160 | 0.0005 |
| 1.0765820 | 3.72E-07 |
| 1.1705200 | 1.10E-11 |
| 0.6291660 | 5.49E-09 |
| 0.0235470 | 2.65E-16 |
| 0.0480890 | 0.0016 |
| 1.1810500 | 1.14E-10 |
| 0.0923810 | 1.98E-11 |
| 0.56261 | 0.2697 |
| 0.1084890 | 2.96E-05 |
| 0.1647350 | 3.49E-11 |
| 0.0525340 | 0.0081 |
| 0.1073550 | 3.24E-07 |
| 0.1032200 | 1.33E-05 |
| 0.1917860 | 1.49E-05 |
| 0.1253470 | 1.40E-14 |
| 0.0000000 | 1.07E-06 |
| 0.0767040 | 0.0007 |
| 0.1245970 | 8.94E-12 |
| 0.0250900 | 3.17E-06 |
| 1.1105020 | 1.01E-08 |
| 0.2202080 | 2.17E-10 |
| 0.0472030 | 1.84E-05 |
| 0.1171490 | 2.71E-05 |
| 0.0641610 | 4.65E-05 |
| 0.2634510 | 0.0118 |
| 0.1578010 | 2.18E-06 |
| 0.7916960 | 0.0078 |
| 0.2938490 | 2.41E-08 |
| 0.6427990 | 0.0001 |
| 0.0220370 | 9.56E-14 |
| 0.0891390 | 1.52E-10 |
| 0.1008250 | 3.80E-12 |
| 0.6682690 | 0.0153 |
| 0.0957050 | 3.93E-09 |
| 0.0158170 | 0.0009 |
| 0.2961480 | 0.0054 |

| | |
|---|---|
| 0.0000100 | 0.0374 |
| 0.1253440 | 6.21E-11 |
| 0.3826010 | 2.04E-06 |
| 1.2562790 | 2.33E-05 |
| 0.0000040 | 0.0001 |
| 0.0378280 | 7.71E-09 |
| 0.0000430 | 3.09E-17 |
| 0.6443390 | 0.0018 |
| 0.0662140 | 1.88E-06 |
| 0.0000080 | 0.0025 |
| 0.8788840 | 0.0002 |
| 0.2470690 | 7.38E-05 |
| 0.0527130 | 8.23E-10 |
| 0.1148650 | 9.36E-08 |
| 0.6208110 | 2.62E-06 |
| 0.0000460 | 1.87E-10 |
| 0.4721870 | 5.51E-05 |
| 1.7576620 | 2.18E-15 |
| 0.8434640 | 6.88E-09 |
| 0.0643420 | 2.40E-07 |
| 0.1445230 | 0.0002 |
| 0.0374420 | 0.0005 |
| 0.0138670 | 0.0371 |
| 0.3041150 | 1.44E-08 |
| 0.0259910 | 0.0319 |
| 0.7107720 | 2.34E-10 |
| 0.0000180 | 3.89E-07 |
| 0.5451830 | 0.0030 |
| 0.1241620 | 8.71E-06 |
| 0.2682150 | 1.89E-07 |
| 0.3767030 | 0.0006 |
| 0.0237360 | 1.89E-11 |
| 0.1566950 | 2.43E-06 |
| 0.0335170 | 1.98E-07 |
| 0.2420390 | 0.0070 |
| 0.2800240 | 0.0185 |
| 0.0098140 | 2.09E-05 |
| 0.0460060 | 3.38E-09 |
| 0.2352840 | 9.72E-11 |
| 0.1604270 | 0.0002 |
| 0.0129080 | 0.0002 |
| 0.6281020 | 1.99E-05 |
| 0.1992220 | 6.42E-12 |
| 0.2740960 | 0.0204 |

| | |
|---|---|
| 0.9104590 | 1.67E-13 |
| 0.0773560 | 2.45E-09 |
| 0.3705040 | 0.0005 |
| 0.2035230 | 6.06E-07 |
| 0.7218070 | 0.0011 |
| 0.1759730 | 4.72E-07 |
| 0.1041590 | 9.60E-05 |
| 1.0857210 | 3.40E-19 |
| 0.0000020 | 0.0016 |
| 0.1255000 | 3.05E-10 |
| 0.0596050 | 6.35E-05 |
| 0.0632130 | 0.0012 |
| 0.0286860 | 1.01E-09 |
| 0.00989 | 0.1319 |
| 0.1239210 | 1.05E-08 |
| 1.4104870 | 1.52E-08 |
| 0.0386600 | 0.0002 |
| 0.5136290 | 8.59E-09 |
| 0.0254820 | 1.36E-14 |
| 0.1303520 | 4.88E-05 |
| 0.1415910 | 1.28E-19 |
| 0.0000000 | 1.31E-13 |
| 0.14333 | 0.1258 |
| 0.0119880 | 0.0101 |
| 0.8282550 | 9.16E-12 |
| 0.0845680 | 2.15E-05 |
| 0.0903140 | 0.0001 |
| 0.5844260 | 0.0005 |
| 0.8295670 | 0.0003 |
| 0.8248300 | 4.44E-08 |
| 0.2161580 | 1.38E-05 |
| 0.5536670 | 4.29E-05 |
| 0.3810630 | 0.0051 |
| 0.1491430 | 4.95E-05 |
| 0.5275080 | 4.60E-09 |
| 0.0356830 | 1.04E-07 |
| 0.0521980 | 0.0006 |
| 0.1330870 | 0.0103 |
| 0.0671210 | 4.39E-05 |
| 0.1994260 | 0.0022 |
| 0.2347200 | 0.0050 |
| 0.0180130 | 0.0109 |
| 0.03695 | 0.5800 |
| 0.0367600 | 0.0034 |

| | |
|---|---|
| 0.1014640 | 0.0014 |
| 0.4459080 | 0.0010 |
| 0.2511560 | 1.10E-05 |
| 1.1700760 | 1.10E-06 |
| 1.7780030 | 9.04E-10 |
| 0.0000400 | 7.86E-10 |
| 0.6882730 | 0.0001 |
| 0.1464330 | 1.34E-05 |
| 0.0000030 | 6.58E-14 |
| 1.1498970 | 3.60E-06 |
| 0.0803400 | 1.01E-08 |
| 1.2845830 | 8.17E-10 |
| 0.1223030 | 4.64E-05 |
| 0.00916 | 0.1074 |
| 0.6724350 | 9.88E-11 |
| 0.0607030 | 1.58E-08 |
| 0.2834920 | 0.0221 |
| 0.1201590 | 1.92E-08 |
| 0.1027420 | 3.97E-06 |
| 0.1295420 | 2.34E-09 |
| 0.0062210 | 5.74E-06 |
| 0.09472 | 0.1472 |
| 0.0540610 | 0.0001 |
| 0.1833300 | 1.60E-13 |
| 1.3435060 | 0.0001 |
| 0.04598 | 0.5362 |
| 0.02585 | 0.3387 |
| 0.9534590 | 1.88E-06 |
| 0.0638940 | 5.25E-06 |
| 0.0856520 | 1.67E-06 |
| 0.1382790 | 2.04E-09 |
| 0.1243630 | 6.65E-09 |
| 0.6082130 | 1.48E-05 |
| 0.0544540 | 1.10E-05 |
| 0.1777350 | 8.49E-12 |
| 2.2157800 | 2.70E-13 |
| 0.04004 | 0.0700 |
| 0.0634890 | 2.70E-07 |
| 0.3364530 | 1.83E-06 |
| 0.22738 | 0.1020 |
| 0.1148110 | 2.63E-07 |
| 0.0666620 | 7.37E-09 |
| 0.7695210 | 0.0006 |
| 0.8317420 | 0.0015 |

| | |
|---|---|
| 0.0625940 | 1.53E-10 |
| 0.1016340 | 7.98E-07 |
| 1.0796020 | 1.20E-07 |
| 0.1504680 | 5.35E-10 |
| 0.8321610 | 3.01E-14 |
| 0.0000210 | 1.96E-08 |
| 0.0800600 | 2.41E-08 |
| 0.1301170 | 0.0023 |
| 1.2004040 | 6.42E-08 |
| 0.0409530 | 2.34E-05 |
| 0.0196110 | 4.43E-07 |
| 0.0543020 | 0.0024 |
| 0.0361300 | 1.60E-06 |
| 0.0294700 | 1.07E-10 |
| 0.0000030 | 2.00E-09 |
| 0.0000100 | 3.35E-09 |
| 0.1022150 | 1.06E-05 |
| 0.1454350 | 1.58E-07 |
| 0.0000080 | 4.36E-07 |
| 0.0189880 | 1.23E-07 |
| 0.1174040 | 5.39E-07 |
| 1.5610590 | 1.39E-05 |
| 0.0476340 | 0.0112 |
| 0.3778300 | 4.51E-05 |
| 0.0844230 | 8.36E-07 |
| 0.02690 | 0.0900 |
| 0.3130790 | 0.0081 |
| 1.0944160 | 1.40E-05 |
| 0.0000020 | 1.46E-12 |
| 0.0924040 | 1.31E-07 |
| 0.0000080 | 3.90E-07 |
| 0.0770840 | 0.0003 |
| 0.1099380 | 5.17E-09 |
| 10.5943360 | 9.50E-14 |
| 0.2382600 | 2.59E-08 |
| 0.07432 | 0.0800 |
| 0.0526960 | 1.30E-16 |
| 0.0873660 | 1.61E-08 |
| 0.2531920 | 1.71E-14 |
| 0.5898700 | 4.00E-05 |
| 0.0708440 | 1.85E-08 |
| 0.3614610 | 0.0038 |
| 0.0735860 | 5.99E-06 |
| 0.0000070 | 6.58E-07 |

| | |
|---|---|
| 1.0892750 | 6.23E-10 |
| 0.0558090 | 3.58E-19 |
| 0.1407250 | 9.05E-06 |
| 0.0000130 | 7.92E-07 |
| 0.03152 | 0.2899 |
| 0.1096800 | 0.0005 |
| 0.0081730 | 1.55E-06 |
| 0.0335550 | 4.64E-07 |
| 0.0096850 | 2.91E-13 |
| 0.0000040 | 6.39E-13 |
| 0.5885730 | 1.84E-05 |
| 0.1712180 | 7.82E-06 |
| 0.0136590 | 0.0003 |
| 0.4236200 | 0.0020 |
| 0.6555100 | 2.29E-05 |
| 0.1084280 | 9.25E-08 |
| 0.0798460 | 0.0002 |
| 0.0718510 | 3.12E-05 |
| 0.0598180 | 5.96E-09 |
| 0.00003 | 0.0800 |
| 0.0000020 | 2.71E-11 |
| 0.0651800 | 6.27E-15 |
| 0.1222070 | 3.11E-10 |
| 0.0486860 | 9.69E-08 |
| 0.6437810 | 3.52E-06 |
| 0.2325690 | 0.0010 |
| 2.1006590 | 3.86E-07 |
| 1.4726410 | 9.04E-11 |
| 0.0000000 | 0.0009 |
| 0.9365650 | 5.00E-05 |
| 0.0291240 | 7.67E-05 |
| 0.1352030 | 1.04E-08 |
| 0.0000010 | 8.54E-06 |
| 0.8903780 | 5.31E-06 |
| 0.4789410 | 6.49E-06 |
| 0.0694410 | 7.37E-09 |
| 0.3035910 | 0.0500 |
| 0.0000010 | 1.61E-14 |
| 0.0063210 | 9.39E-14 |
| 0.1522440 | 3.50E-07 |
| 0.0456190 | 0.0004 |
| 0.30362 | 0.1070 |
| 0.1760600 | 5.11E-05 |
| 0.0819240 | 6.39E-06 |

| | |
|---|---|
| 0.1197570 | 1.59E-06 |
| 0.0214480 | 1.22E-07 |
| 0.0000150 | 0.0017 |
| 0.1955580 | 3.53E-10 |
| 0.2656110 | 0.0001 |
| 0.1833780 | 6.13E-06 |
| 0.1774630 | 0.0036 |
| 0.1627340 | 1.67E-06 |
| 0.00671 | 0.0700 |
| 0.1142480 | 7.62E-10 |
| 0.0000100 | 2.65E-08 |
| 0.17077 | 0.1000 |
| 0.0000050 | 0.0243 |
| 0.1042690 | 3.72E-06 |
| 0.0339170 | 0.0082 |
| 0.0000010 | 1.72E-13 |
| 0.0254050 | 2.36E-06 |
| 0.0838040 | 0.0130 |
| 0.1230400 | 0.0039 |
| 0.1732430 | 8.32E-07 |
| 0.0888980 | 0.0016 |
| 0.1010220 | 2.48E-11 |
| 0.7148140 | 9.50E-09 |
| 0.02331 | 0.2535 |
| 0.3778290 | 0.0063 |
| 0.01575 | 0.1261 |
| 0.0000070 | 0.0015 |
| 0.0381210 | 0.0003 |
| 0.0000190 | 1.71E-08 |
| 0.5206640 | 2.05E-08 |
| 0.1589220 | 3.32E-05 |
| 0.0468230 | 2.95E-09 |
| 0.7743020 | 0.0001 |
| 0.0000010 | 0.0500 |
| 0.2023080 | 1.32E-05 |
| 0.1545610 | 7.62E-10 |
| 0.1740420 | 0.0003 |
| 0.1524420 | 0.0002 |
| 0.0000020 | 4.81E-06 |
| 0.0581810 | 1.35E-07 |
| 0.0000010 | 0.0030 |
| 0.0000000 | 0.0010 |
| 1.5463890 | 8.59E-09 |
| 0.2097320 | 1.58E-07 |

| | |
|---|---|
| 0.2232190 | 1.25E-11 |
| 0.0353700 | 0.0001 |
| 0.0138540 | 4.36E-05 |
| 0.0000030 | 0.0004 |
| 0.1058440 | 9.84E-10 |
| 0.1124840 | 1.97E-10 |
| 0.0009510 | 0.0004 |
| 0.8415910 | 2.69E-07 |
| 0.0004690 | 0.0025 |
| 0.0640470 | 0.0005 |
| 0.0608530 | 0.0008 |
| 0.0634580 | 7.81E-06 |
| 0.0000230 | 0.0259 |
| 0.0225650 | 0.0002 |
| 0.0000060 | 7.45E-05 |
| 0.4583200 | 0.0267 |
| 0.0621600 | 4.04E-05 |
| 0.0000010 | 0.0005 |
| 0.0785760 | 0.0002 |
| 0.2190670 | 5.89E-05 |
| 0.0000070 | 3.87E-05 |
| 0.0590350 | 0.0004 |
| 0.1018860 | 5.24E-07 |
| 0.1531510 | 0.0007 |
| 4.3256340 | 1.44E-06 |
| 0.0328240 | 0.0182 |
| 0.0000020 | 0.0004 |
| 0.9882380 | 8.02E-13 |
| 0.4735810 | 0.0077 |
| 0.0064150 | 1.00E-06 |
| 0.0205430 | 0.0004 |
| 0.0592680 | 2.81E-06 |
| 0.0000070 | 0.0213 |
| 0.8065430 | 0.0001 |
| 0.0658000 | 1.36E-07 |
| 1.9558580 | 5.30E-10 |
| 0.0200000 | 2.78E-06 |
| 1.5901880 | 0.0018 |
| 0.3390310 | 9.24E-08 |
| 0.0638340 | 1.81E-06 |
| 0.8628880 | 0.0002 |
| 0.1217380 | 0.0002 |
| 0.6994030 | 0.0003 |
| 0.2078340 | 0.0002 |

| | |
|---|---|
| 0.0000080 | 6.33E-11 |
| 0.1131170 | 0.0001 |
| 0.1197180 | 0.0027 |
| 0.0000050 | 4.27E-06 |
| 0.2195360 | 9.49E-06 |
| 0.6212420 | 5.76E-07 |
| 0.3314210 | 1.38E-12 |
| 0.0000400 | 0.0087 |
| 0.1250740 | 1.05E-08 |
| 0.0853310 | 0.0002 |
| 0.0000060 | 0.0014 |
| 0.1178210 | 7.40E-11 |
| 0.1572630 | 8.05E-09 |
| 0.0008040 | 6.41E-06 |
| 0.1348200 | 0.0071 |
| 0.05041 | 0.1580 |
| 0.0657450 | 4.91E-06 |
| 0.0000060 | 0.0019 |
| 0.0601600 | 3.81E-06 |
| 0.03131 | 0.4639 |
| 0.8250270 | 2.29E-08 |
| 0.0000150 | 0.0016 |
| 0.13479 | 0.2668 |
| 0.1289010 | 0.0075 |
| 0.5019300 | 0.0056 |
| 0.1319680 | 3.80E-05 |
| 0.0513790 | 0.0024 |
| 0.0193490 | 0.0028 |
| 0.0001780 | 1.69E-05 |
| 0.0000080 | 3.77E-05 |
| 0.0674500 | 6.49E-07 |
| 0.1404530 | 0.0001 |
| 0.0000580 | 1.46E-12 |
| 0.0872790 | 0.0015 |
| 0.0427510 | 0.0033 |
| 1.3923630 | 0.0051 |
| 0.0764900 | 1.33E-06 |
| 0.1443090 | 0.0028 |
| 3.5035010 | 0.0092 |
| 0.5419150 | 0.0004 |
| 0.0991320 | 1.40E-05 |
| 0.0000050 | 6.45E-06 |
| 0.0349890 | 8.84E-05 |
| 0.0318920 | 0.0175 |

| | |
|---|---|
| 0.3732550 | 0.0052 |
| 0.2436100 | 9.76E-07 |
| 0.0000040 | 0.0056 |
| 0.1672630 | 0.0311 |
| 0.0000080 | 4.77E-11 |
| 0.0923350 | 0.0069 |
| 0.0185670 | 0.0003 |
| 0.0534920 | 7.51E-10 |
| 0.0706700 | 3.34E-06 |
| 0.00000 | 0.1074 |
| 0.0000080 | 6.21E-06 |
| 0.0518400 | 4.85E-05 |
| 0.7109930 | 8.33E-06 |
| 0.0615720 | 2.65E-06 |
| 0.1179390 | 9.80E-08 |
| 0.5731730 | 7.58E-06 |
| 0.0657930 | 0.0025 |
| 0.1663140 | 0.0001 |
| 0.0000120 | 0.0015 |
| 0.0000080 | 0.0223 |
| 0.3396440 | 4.62E-06 |
| 0.0000000 | 4.76E-09 |
| 0.8088410 | 1.79E-06 |
| 0.0000100 | 0.0002 |
| 0.1032460 | 0.0004 |
| 0.0000010 | 0.0002 |
| 0.0710130 | 0.0014 |
| 0.00001 | 0.1473 |
| 0.3839590 | 0.0003 |
| 0.2138080 | 0.0500 |
| 0.2956170 | 0.0001 |
| 0.1753870 | 2.65E-06 |
| 0.1678090 | 0.0027 |
| 0.0076270 | 3.46E-06 |
| 0.3899460 | 0.0005 |
| 0.06801 | 0.9985 |
| 0.7457940 | 0.0212 |
| 0.0835570 | 0.0027 |
| 6.0697040 | 4.98E-05 |
| 2.3922940 | 1.74E-10 |
| 36.0337060 | 0.0110 |
| 0.0000010 | 0.0001 |
| 0.0000000 | 0.0254 |
| 0.0464450 | 0.0008 |

| | |
|---|---|
| 1.6961300 | 0.0183 |
| 0.20431 | 0.1519 |
| 0.0271600 | 0.0002 |
| 0.0000150 | 4.82E-06 |
| 0.0279460 | 0.0260 |
| 0.0000050 | 7.69E-08 |
| 0.0688950 | 0.0154 |
| 0.0194090 | 0.0087 |
| 0.1439190 | 0.0003 |
| 0.0619650 | 0.0001 |
| 0.03913 | 0.2237 |
| 0.6859190 | 0.0236 |
| 0.0553360 | 9.70E-05 |
| 0.07479 | 0.5007 |
| 0.8395940 | 1.64E-05 |
| 0.0000000 | 1.27E-06 |
| 0.1537920 | 1.72E-07 |
| 0.56442 | 0.4590 |
| 0.1516850 | 0.0185 |
| 0.3444480 | 0.0233 |
| 0.2623970 | 1.75E-07 |
| 0.2056660 | 0.0040 |
| 0.1596310 | 3.81E-06 |
| 0.2268330 | 7.27E-07 |
| 0.9175890 | 1.38E-05 |
| 0.1089870 | 0.0140 |
| 0.0381160 | 0.0265 |
| 0.0318980 | 0.0400 |
| 0.5152520 | 2.10E-05 |
| 0.0348470 | 0.0319 |
| 0.0854440 | 0.0080 |
| 0.0000050 | 1.81E-06 |
| 1.21349 | 0.1185 |
| 0.0000000 | 0.0078 |
| 0.1639750 | 9.20E-05 |
| 0.1656400 | 0.0003 |
| 0.0553240 | 1.34E-06 |
| 0.0000040 | 3.90E-09 |
| 0.00000 | 0.5125 |
| 0.0735040 | 0.0161 |
| 0.0000000 | 3.61E-07 |
| 0.55547 | 0.0600 |
| 0.02290 | 0.2000 |
| 0.0000040 | 0.0302 |

| | |
|---|---|
| 0.1710300 | 0.0203 |

**Supplementary Table 25.** codeml results - w for the O. ovis/other species

| ID | Likelihood | ϖ other species | dN | dS | ϖ *O. ovis* | dN | dS |
|---|---|---|---|---|---|---|---|
| cluster1127 | -1407.8957 | 0.0640 | 0.051 | 0.797 | 0.0213 | 0.2075 | 9.7622 |
| cluster2853 | -22469.879 | 0.0259 | 0.027 | 1.036 | 0.0412 | 0.0497 | 1.2064 |
| cluster2854 | -1615.3622 | 0.0359 | 0.008 | 0.234 | 0.0347 | 0.0262 | 0.7552 |
| cluster2855 | -23554.802 | 0.0196 | 0.018 | 0.903 | 0.0235 | 0.0822 | 3.4938 |
| cluster2856 | -15422.869 | 0.1209 | 0.086 | 0.712 | 0.0726 | 0.2975 | 4.0981 |
| cluster2857 | -5969.8149 | 0.0483 | 0.005 | 0.101 | 0.0012 | 0.0021 | 1.7937 |
| cluster2858 | -7753.8927 | 0.0029 | 0.002 | 0.577 | 0.0055 | 0.0093 | 1.7030 |
| cluster2859 | -5679.1307 | 0.0394 | 0.009 | 0.237 | 0.0440 | 0.0377 | 0.8585 |
| cluster2860 | -16041.379 | 0.0012 | 0.001 | 0.649 | 0.0029 | 0.0036 | 1.2139 |
| cluster2861 | -8375.1097 | 0.0100 | 0.007 | 0.657 | 0.0118 | 0.0172 | 1.4588 |
| cluster2862 | -18856.54 | 0.0145 | 0.012 | 0.813 | 0.0184 | 0.0198 | 1.0730 |
| cluster2863 | -15073.285 | 0.0289 | 0.021 | 0.741 | 0.0385 | 0.0758 | 1.9687 |
| cluster2864 | -12544.293 | 0.0027 | 0.003 | 0.932 | 0.0080 | 0.0088 | 1.0979 |
| cluster2865 | -14997.672 | 0.0879 | 0.042 | 0.478 | 0.1446 | 0.1026 | 0.7095 |
| cluster2866 | -17308.964 | 0.0259 | 0.018 | 0.677 | 0.0394 | 0.0946 | 2.4004 |
| cluster2867 | -14862.209 | 0.0534 | 0.042 | 0.792 | 0.0392 | 0.0979 | 2.4989 |
| cluster2869 | -8916.4524 | 0.0962 | 0.034 | 0.357 | 0.0822 | 0.1211 | 1.4734 |
| cluster2870 | -3878.6903 | 0.0634 | 0.025 | 0.390 | 0.0317 | 0.0737 | 2.3242 |
| cluster2871 | -3139.858 | 0.0247 | 0.005 | 0.217 | 0.0291 | 0.0182 | 0.6262 |
| cluster2872 | -12259.81 | 0.0797 | 0.053 | 0.662 | 0.0992 | 0.1916 | 1.9317 |
| cluster2873 | -13841.724 | 0.0122 | 0.008 | 0.633 | 0.0287 | 0.0264 | 0.9225 |
| cluster2874 | -10388.038 | 0.0399 | 0.016 | 0.402 | 0.0353 | 0.0439 | 1.2412 |
| cluster2875 | -13628.633 | 0.0137 | 0.014 | 1.011 | 0.0282 | 0.0468 | 1.6625 |
| cluster2876 | -3967.617 | 0.0288 | 0.018 | 0.640 | 0.0728 | 0.0495 | 0.6806 |
| cluster2877 | -10637.493 | 0.0502 | 0.027 | 0.539 | 0.0587 | 0.0767 | 1.3076 |
| cluster2878 | -7054.4173 | 0.0314 | 0.009 | 0.280 | 0.0599 | 0.0369 | 0.6149 |
| cluster2879 | -10032.054 | 0.2124 | 0.048 | 0.225 | 0.1437 | 0.1071 | 0.7451 |
| cluster2880 | -10642.154 | 0.0482 | 0.035 | 0.726 | 0.0534 | 0.1304 | 2.4442 |
| cluster2881 | -11522.348 | 0.0498 | 0.037 | 0.734 | 0.0641 | 0.0501 | 0.7804 |
| cluster2882 | -10649.568 | 0.0321 | 0.020 | 0.626 | 0.0272 | 0.0585 | 2.1502 |
| cluster2883 | -6764.0384 | 0.0662 | 0.034 | 0.514 | 0.0900 | 0.0577 | 0.6406 |
| cluster2884 | -12071.352 | 0.0269 | 0.011 | 0.414 | 0.0244 | 0.0296 | 1.2142 |
| cluster2885 | -15738.647 | 0.0468 | 0.037 | 0.783 | 0.0305 | 0.1280 | 4.1931 |
| cluster2886 | -12000.18 | 0.0357 | 0.021 | 0.589 | 0.0601 | 0.0440 | 0.7320 |
| cluster2887 | -8364.7279 | 0.0073 | 0.005 | 0.682 | 0.0018 | 0.0189 | 10.5803 |
| cluster2888 | -7784.1859 | 0.0715 | 0.072 | 1.002 | 0.0424 | 0.1514 | 3.5751 |
| cluster2889 | -6929.8504 | 0.0073 | 0.006 | 0.796 | 0.0110 | 0.0237 | 2.1669 |
| cluster2890 | -2675.7997 | 0.1174 | 0.014 | 0.119 | 0.0308 | 0.0084 | 0.2715 |
| cluster2891 | -12098.448 | 0.0394 | 0.025 | 0.645 | 0.0355 | 0.1251 | 3.5223 |
| cluster2892 | -7912.4716 | 0.0138 | 0.009 | 0.641 | 0.0192 | 0.0150 | 0.7781 |
| cluster2893 | -8461.1297 | 0.0411 | 0.037 | 0.905 | 0.0099 | 0.1387 | 14.0629 |
| cluster2894 | -5510.3237 | 0.0140 | 0.006 | 0.406 | 0.0259 | 0.0467 | 1.8011 |

| | | | | | | | |
|---|---|---|---|---|---|---|---|
| cluster2895 | -8448.0915 | 0.0344 | 0.025 | 0.715 | 0.0438 | 0.0964 | 2.1987 |
| cluster2896 | -10287.856 | 0.0104 | 0.009 | 0.853 | 0.0099 | 0.0261 | 2.6533 |
| cluster2897 | -9158.799 | 0.0321 | 0.015 | 0.473 | 0.0385 | 0.0570 | 1.4811 |
| cluster2898 | -4499.4345 | 0.0291 | 0.014 | 0.477 | 0.0166 | 0.0149 | 0.8980 |
| cluster2899 | -8006.3949 | 0.0184 | 0.021 | 1.119 | 0.0240 | 0.0369 | 1.5395 |
| cluster2900 | -9923.4607 | 0.0122 | 0.008 | 0.645 | 0.0162 | 0.0117 | 0.7197 |
| cluster2901 | -9240.1245 | 0.0350 | 0.025 | 0.721 | 0.0453 | 0.1091 | 2.4095 |
| cluster2902 | -3696.6863 | 0.0260 | 0.021 | 0.812 | 0.0279 | 0.0664 | 2.3806 |
| cluster2903 | -8864.0178 | 0.0101 | 0.010 | 0.980 | 0.0125 | 0.0093 | 0.7427 |
| cluster2904 | -6409.4988 | 0.1049 | 0.051 | 0.488 | 0.0500 | 0.0315 | 0.6298 |
| cluster2905 | -8880.0209 | 0.0308 | 0.014 | 0.446 | 0.0240 | 0.0155 | 0.6469 |
| cluster2906 | -13664.807 | 0.0675 | 0.029 | 0.434 | 0.0980 | 0.0587 | 0.5985 |
| cluster2907 | -12738.318 | 0.0793 | 0.064 | 0.810 | 0.0569 | 0.2097 | 3.6845 |
| cluster2908 | -5643.3725 | 0.0090 | 0.002 | 0.250 | 0.0175 | 0.0127 | 0.7239 |
| cluster2909 | -2925.4119 | 0.0384 | 0.009 | 0.233 | 0.0381 | 0.0268 | 0.7051 |
| cluster2910 | -9271.2419 | 0.0418 | 0.025 | 0.606 | 0.0465 | 0.0721 | 1.5528 |
| cluster2911 | -4008.64 | 0.0053 | 0.001 | 0.167 | 0.0034 | 0.0018 | 0.5240 |
| cluster2912 | -8765.4408 | 0.0301 | 0.022 | 0.723 | 0.0184 | 0.0593 | 3.2178 |
| cluster2913 | -6295.9542 | 0.0016 | 0.001 | 0.504 | 0.0040 | 0.0068 | 1.7173 |
| cluster2914 | -8063.818 | 0.0164 | 0.007 | 0.449 | 0.0146 | 0.0116 | 0.7981 |
| cluster2915 | -4206.554 | 0.0150 | 0.010 | 0.647 | 0.0213 | 0.0334 | 1.5701 |
| cluster2916 | -9769.6436 | 0.0299 | 0.019 | 0.632 | 0.0389 | 0.0311 | 0.7998 |
| cluster2917 | -9014.0653 | 0.0147 | 0.009 | 0.609 | 0.0364 | 0.0620 | 1.7028 |
| cluster2918 | -4475.319 | 0.0773 | 0.026 | 0.334 | 0.0663 | 0.0564 | 0.8514 |
| cluster2919 | -9563.8287 | 0.0627 | 0.027 | 0.431 | 0.0831 | 0.0541 | 0.6512 |
| cluster2920 | -6955.8131 | 0.0166 | 0.018 | 1.082 | 0.0068 | 0.0584 | 8.5662 |
| cluster2921 | -3808.6709 | 0.0047 | 0.003 | 0.729 | 0.0029 | 0.0076 | 2.6201 |
| cluster2922 | -6244.5831 | 0.0582 | 0.037 | 0.644 | 0.0516 | 0.1433 | 2.7751 |
| cluster2923 | -5867.5848 | 0.1853 | 0.071 | 0.385 | 0.1691 | 0.2885 | 1.7059 |
| cluster2924 | -10974.114 | 0.0235 | 0.019 | 0.799 | 0.0110 | 0.0655 | 5.9558 |
| cluster2925 | -5771.6089 | 0.0334 | 0.024 | 0.705 | 0.0329 | 0.0786 | 2.3889 |
| cluster2926 | -11218.956 | 0.0199 | 0.016 | 0.799 | 0.0266 | 0.0270 | 1.0161 |
| cluster2927 | -4900.572 | 0.0370 | 0.031 | 0.844 | 0.0168 | 0.0760 | 4.5224 |
| cluster2928 | -7179.9921 | 0.0197 | 0.014 | 0.723 | 0.0261 | 0.0548 | 2.1004 |
| cluster2929 | -8010.2895 | 0.0049 | 0.003 | 0.619 | 0.0071 | 0.0142 | 1.9915 |
| cluster2930 | -10988.592 | 0.0814 | 0.033 | 0.409 | 0.0915 | 0.1355 | 1.4803 |
| cluster2931 | -6865.3848 | 0.0558 | 0.049 | 0.884 | 0.0378 | 0.1446 | 3.8226 |
| cluster2932 | -9691.2992 | 0.0669 | 0.047 | 0.698 | 0.0462 | 0.1064 | 2.3061 |
| cluster2933 | -8736.8945 | 0.0454 | 0.018 | 0.388 | 0.0523 | 0.0298 | 0.5701 |
| cluster2934 | -10531.791 | 0.0626 | 0.021 | 0.341 | 0.0651 | 0.0730 | 1.1219 |
| cluster2935 | -4950.688 | 0.0249 | 0.015 | 0.619 | 0.0429 | 0.0384 | 0.8947 |
| cluster2936 | -9510.0927 | 0.0392 | 0.037 | 0.954 | 0.0276 | 0.1062 | 3.8523 |
| cluster2937 | -4595.6016 | 0.0272 | 0.012 | 0.442 | 0.0302 | 0.0196 | 0.6482 |
| cluster2938 | -7240.2952 | 0.0138 | 0.007 | 0.521 | 0.0198 | 0.0231 | 1.1682 |

| | | | | | | | |
|---|---|---|---|---|---|---|---|
| cluster2939 | -2814.0059 | 0.0160 | 0.004 | 0.280 | 0.0451 | 0.0212 | 0.4687 |
| cluster2940 | -11378.217 | 0.0691 | 0.039 | 0.558 | 0.0613 | 0.1387 | 2.2619 |
| cluster2941 | -3030.3381 | 0.0024 | 0.000 | 0.150 | 0.0001 | 0.0000 | 0.2925 |
| cluster2942 | -11359.428 | 0.0551 | 0.038 | 0.692 | 0.0504 | 0.1351 | 2.6829 |
| cluster2943 | -4040.1776 | 0.0101 | 0.007 | 0.703 | 0.0131 | 0.0168 | 1.2852 |
| cluster2944 | -8493.9703 | 0.1035 | 0.076 | 0.733 | 0.1042 | 0.1182 | 1.1349 |
| cluster2945 | -3084.8219 | 0.0026 | 0.002 | 0.862 | 0.0018 | 0.0127 | 6.8922 |
| cluster2946 | -6742.5059 | 0.0099 | 0.003 | 0.278 | 0.0082 | 0.0041 | 0.5059 |
| cluster2947 | -9655.711 | 0.0312 | 0.012 | 0.380 | 0.0492 | 0.0517 | 1.0523 |
| cluster2948 | -9375.193 | 0.0491 | 0.033 | 0.670 | 0.0721 | 0.0688 | 0.9542 |
| cluster2949 | -6814.4755 | 0.0154 | 0.008 | 0.513 | 0.0241 | 0.0177 | 0.7340 |
| cluster2950 | -7118.2306 | 0.0050 | 0.016 | 3.186 | 0.0059 | 0.0042 | 0.7164 |
| cluster2951 | -9094.5491 | 0.0619 | 0.045 | 0.726 | 0.0654 | 0.1722 | 2.6333 |
| cluster2952 | -10892.914 | 0.0460 | 0.030 | 0.645 | 0.0239 | 0.0947 | 3.9644 |
| cluster2953 | -13288.435 | 0.0752 | 0.061 | 0.816 | 0.0505 | 0.1703 | 3.3751 |
| cluster2954 | -7273.7228 | 0.0188 | 0.019 | 1.013 | 0.0235 | 0.0409 | 1.7397 |
| cluster2955 | -6020.4296 | 0.0169 | 0.008 | 0.480 | 0.0417 | 0.0653 | 1.5654 |
| cluster2956 | -10128.708 | 0.0110 | 0.030 | 2.719 | 0.0240 | 0.0277 | 1.1539 |
| cluster2957 | -8387.704 | 0.0898 | 0.061 | 0.683 | 0.1085 | 0.2148 | 1.9802 |
| cluster2958 | -9690.7419 | 0.0110 | 0.008 | 0.694 | 0.0136 | 0.0261 | 1.9166 |
| cluster2959 | -5169.3859 | 0.0540 | 0.027 | 0.506 | 0.0487 | 0.0421 | 0.8652 |
| cluster2960 | -9441.2679 | 0.0222 | 0.018 | 0.812 | 0.0303 | 0.0279 | 0.9233 |
| cluster2961 | -8396.2976 | 0.0087 | 0.012 | 1.370 | 0.0151 | 0.0507 | 3.3559 |
| cluster2962 | -7495.5482 | 0.0218 | 0.031 | 1.426 | 0.0242 | 0.0497 | 2.0564 |
| cluster2963 | -7222.7232 | 0.0327 | 0.010 | 0.304 | 0.0452 | 0.0260 | 0.5748 |
| cluster2964 | -5248.0118 | 0.0289 | 0.012 | 0.429 | 0.0412 | 0.0245 | 0.5953 |
| cluster2965 | -7391.6537 | 0.0120 | 0.006 | 0.515 | 0.0101 | 0.0173 | 1.7090 |
| cluster2966 | -5587.7981 | 0.0231 | 0.021 | 0.930 | 0.0213 | 0.0687 | 3.2307 |
| cluster2967 | -9573.9678 | 0.0143 | 0.008 | 0.541 | 0.0305 | 0.0191 | 0.6269 |
| cluster2968 | -5456.8959 | 0.0181 | 0.010 | 0.566 | 0.0196 | 0.0253 | 1.2917 |
| cluster2969 | -7910.3902 | 0.0410 | 0.019 | 0.471 | 0.0463 | 0.0899 | 1.9442 |
| cluster2970 | -4236.277 | 0.0036 | 0.002 | 0.617 | 0.0041 | 0.0123 | 3.0206 |
| cluster2971 | -3608.6975 | 0.0286 | 0.005 | 0.165 | 0.0346 | 0.0187 | 0.5403 |
| cluster2972 | -2670.0647 | 0.0060 | 0.004 | 0.616 | 0.0070 | 0.0067 | 0.9642 |
| cluster2973 | -6360.2479 | 0.0958 | 0.054 | 0.569 | 0.0824 | 0.1902 | 2.3082 |
| cluster2974 | -4402.626 | 0.0483 | 0.019 | 0.388 | 0.0068 | 0.0068 | 1.0046 |
| cluster2975 | -10300.306 | 0.0683 | 0.040 | 0.591 | 0.0923 | 0.1748 | 1.8940 |
| cluster2976 | -1469.2392 | 0.2131 | 0.084 | 0.395 | 0.1716 | 0.0636 | 0.3707 |
| cluster2977 | -3393.0742 | 0.0048 | 0.006 | 1.318 | 0.0047 | 0.0217 | 4.6294 |
| cluster2978 | -8211.575 | 0.0008 | 0.000 | 0.606 | 0.0036 | 0.0031 | 0.8644 |
| cluster2979 | -8839.1817 | 0.0370 | 0.018 | 0.475 | 0.0450 | 0.0673 | 1.4941 |
| cluster2980 | -1552.4572 | 0.0274 | 0.031 | 1.130 | 0.0092 | 0.1097 | 11.9468 |
| cluster2981 | -6210.7605 | 0.0635 | 0.037 | 0.578 | 0.0397 | 0.0919 | 2.3135 |
| cluster2982 | -8058.1196 | 0.1163 | 0.038 | 0.327 | 0.0934 | 0.0835 | 0.8939 |

| | | | | | | | |
|---|---|---|---|---|---|---|---|
| cluster2983 | -7519.2419 | 0.0223 | 0.018 | 0.805 | 0.0286 | 0.0845 | 2.9521 |
| cluster2984 | -4830.559 | 0.0285 | 0.011 | 0.400 | 0.0443 | 0.0245 | 0.5515 |
| cluster2985 | -3972.5604 | 0.2333 | 0.069 | 0.297 | 0.2519 | 0.1345 | 0.5341 |
| cluster2986 | -3497.2106 | 0.1057 | 0.029 | 0.273 | 0.1126 | 0.1514 | 1.3445 |
| cluster2987 | -6078.0943 | 0.0299 | 0.013 | 0.425 | 0.0313 | 0.0295 | 0.9440 |
| cluster2988 | -5702.9494 | 0.0258 | 0.018 | 0.686 | 0.0194 | 0.0541 | 2.7901 |
| cluster2989 | -6586.3346 | 0.0343 | 0.014 | 0.404 | 0.0631 | 0.0280 | 0.4440 |
| cluster2990 | -7511.0098 | 0.0419 | 0.028 | 0.659 | 0.0541 | 0.0986 | 1.8210 |
| cluster2991 | -6306.087 | 0.0649 | 0.041 | 0.624 | 0.0319 | 0.1829 | 5.7401 |
| cluster2992 | -4177.9729 | 0.0127 | 0.004 | 0.283 | 0.0067 | 0.0043 | 0.6424 |
| cluster2993 | -5419.2914 | 0.0278 | 0.022 | 0.808 | 0.0656 | 0.0335 | 0.5116 |
| cluster2994 | -2260.1203 | 0.0033 | 0.001 | 0.328 | 0.0043 | 0.0044 | 1.0236 |
| cluster2995 | -740.15748 | 0.0180 | 0.008 | 0.462 | 0.0242 | 0.0684 | 2.8333 |
| cluster2996 | -7309.4621 | 0.0401 | 0.025 | 0.621 | 0.0589 | 0.0368 | 0.6245 |
| cluster2997 | -8171.9711 | 0.0410 | 0.021 | 0.517 | 0.0391 | 0.0702 | 1.7967 |
| cluster2998 | -7499.4772 | 0.0153 | 0.010 | 0.623 | 0.0280 | 0.0598 | 2.1373 |
| cluster2999 | -6764.0215 | 0.0300 | 0.020 | 0.657 | 0.0346 | 0.1176 | 3.4001 |
| cluster3000 | -6328.092 | 0.0184 | 0.004 | 0.229 | 0.0066 | 0.0101 | 1.5235 |
| cluster3001 | -6045.9498 | 0.0296 | 0.013 | 0.440 | 0.0082 | 0.0127 | 1.5498 |
| cluster3002 | -7665.0302 | 0.0215 | 0.009 | 0.432 | 0.0372 | 0.0448 | 1.2027 |
| cluster3003 | -7810.9036 | 0.0934 | 0.029 | 0.314 | 0.0838 | 0.0993 | 1.1853 |
| cluster3004 | -5964.7058 | 0.0894 | 0.033 | 0.367 | 0.1144 | 0.0438 | 0.3826 |
| cluster3005 | -4101.6821 | 0.0685 | 0.012 | 0.169 | 0.0476 | 0.0370 | 0.7770 |
| cluster3006 | -8672.0147 | 0.0574 | 0.037 | 0.651 | 0.0563 | 0.0755 | 1.3423 |
| cluster3007 | -8243.4669 | 0.0277 | 0.024 | 0.872 | 0.0183 | 0.1098 | 6.0149 |
| cluster3008 | -5954.0766 | 0.0334 | 0.025 | 0.738 | 0.0288 | 0.0495 | 1.7193 |
| cluster3009 | -5913.2643 | 0.0419 | 0.014 | 0.326 | 0.0462 | 0.0553 | 1.1970 |
| cluster3010 | -8057.4728 | 0.1505 | 0.043 | 0.284 | 0.1492 | 0.1680 | 1.1264 |
| cluster3011 | -3371.9832 | 0.0093 | 0.012 | 1.323 | 0.0117 | 0.0235 | 2.0075 |
| cluster3012 | -5210.0384 | 0.0604 | 0.054 | 0.890 | 0.0511 | 0.1325 | 2.5920 |
| cluster3013 | -6733.1919 | 0.0750 | 0.023 | 0.310 | 0.1076 | 0.0595 | 0.5524 |
| cluster3014 | -3561.4061 | 0.0073 | 0.006 | 0.783 | 0.0104 | 0.0269 | 2.5951 |
| cluster3015 | -2410.5221 | 0.0410 | 0.011 | 0.266 | 0.0429 | 0.0348 | 0.8126 |
| cluster3016 | -4556.1885 | 0.0371 | 0.012 | 0.324 | 0.0434 | 0.0368 | 0.8486 |
| cluster3017 | -5972.13 | 0.0261 | 0.031 | 1.193 | 0.0286 | 0.0768 | 2.6903 |
| cluster3018 | -5678.0243 | 0.0054 | 0.003 | 0.602 | 0.0085 | 0.0161 | 1.9001 |
| cluster3019 | -7570.0049 | 0.0507 | 0.043 | 0.844 | 0.0662 | 0.1140 | 1.7220 |
| cluster3020 | -6195.1844 | 0.0350 | 0.017 | 0.492 | 0.0329 | 0.0531 | 1.6125 |
| cluster3021 | -3023.7905 | 0.0139 | 0.006 | 0.414 | 0.0245 | 0.0153 | 0.6241 |
| cluster3022 | -3431.4875 | 0.0595 | 0.039 | 0.661 | 0.0707 | 0.2235 | 3.1626 |
| cluster3023 | -4199.9473 | 0.0073 | 0.004 | 0.544 | 0.0237 | 0.0128 | 0.5393 |
| cluster3024 | -1959.8125 | 0.0912 | 0.019 | 0.209 | 0.0828 | 0.0632 | 0.7638 |
| cluster3025 | -4691.8565 | 0.0100 | 0.002 | 0.250 | 0.0103 | 0.0078 | 0.7522 |
| cluster3026 | -4111.1689 | 0.1739 | 0.065 | 0.373 | 0.1652 | 0.2193 | 1.3280 |

| | | | | | | | |
|---|---|---|---|---|---|---|---|
| cluster3027 | -3436.6312 | 0.0023 | 0.002 | 0.804 | 0.0024 | 0.0028 | 1.1881 |
| cluster3028 | -6800.4202 | 0.0450 | 0.031 | 0.699 | 0.0158 | 0.1332 | 8.4194 |
| cluster3029 | -6674.5605 | 0.0256 | 0.010 | 0.403 | 0.0344 | 0.0407 | 1.1824 |
| cluster3030 | -5268.4339 | 0.0542 | 0.016 | 0.302 | 0.0527 | 0.0310 | 0.5877 |
| cluster3031 | -8792.2678 | 0.0882 | 0.061 | 0.689 | 0.1772 | 0.0984 | 0.5551 |
| cluster3032 | -5912.2176 | 0.0196 | 0.015 | 0.765 | 0.0418 | 0.0317 | 0.7597 |
| cluster3033 | -3821.6283 | 0.1185 | 0.010 | 0.085 | 0.1956 | 0.0981 | 0.5013 |
| cluster3034 | -4964.525 | 0.0346 | 0.297 | 8.599 | 0.0040 | 0.1018 | 25.5438 |
| cluster3035 | -7644.6024 | 0.0222 | 0.011 | 0.516 | 0.0210 | 0.0282 | 1.3434 |
| cluster3036 | -8981.0513 | 0.0758 | 0.048 | 0.633 | 0.0575 | 0.1396 | 2.4282 |
| cluster3037 | -2560.229 | 0.0294 | 0.007 | 0.250 | 0.0067 | 0.0073 | 1.0812 |
| cluster3038 | -4466.6359 | 0.0484 | 0.023 | 0.474 | 0.0964 | 0.1528 | 1.5852 |
| cluster3039 | -6524.2308 | 0.0265 | 0.007 | 0.277 | 0.0529 | 0.0240 | 0.4538 |
| cluster3040 | -4843.8244 | 0.0177 | 0.006 | 0.315 | 0.0228 | 0.0134 | 0.5867 |
| cluster3041 | -8446.3397 | 0.0484 | 0.038 | 0.778 | 0.0350 | 0.0853 | 2.4399 |
| cluster3042 | -4404.2455 | 0.0359 | 0.020 | 0.551 | 0.0295 | 0.0683 | 2.3154 |
| cluster3043 | -2912.9727 | 0.0330 | 0.037 | 1.127 | 0.0075 | 0.0807 | 10.7192 |
| cluster3044 | -6427.4866 | 0.0281 | 0.016 | 0.573 | 0.0381 | 0.0792 | 2.0779 |
| cluster3045 | -6128.2807 | 0.0564 | 0.019 | 0.337 | 0.0457 | 0.0495 | 1.0835 |
| cluster3046 | -7275.2647 | 0.1994 | 0.099 | 0.499 | 0.1337 | 0.0951 | 0.7116 |
| cluster3047 | -2123.1173 | 0.0025 | 0.001 | 0.455 | 0.0051 | 0.0068 | 1.3358 |
| cluster3048 | -5407.3218 | 0.0205 | 0.009 | 0.459 | 0.0248 | 0.0318 | 1.2825 |
| cluster3049 | -7064.4954 | 0.0173 | 0.011 | 0.633 | 0.0148 | 0.0118 | 0.7951 |
| cluster3050 | -7561.6091 | 0.0370 | 0.013 | 0.362 | 0.0467 | 0.0758 | 1.6216 |
| cluster3051 | -7513.1259 | 0.0400 | 0.034 | 0.849 | 0.0353 | 0.1035 | 2.9353 |
| cluster3052 | -6826.5569 | 0.0511 | 0.036 | 0.711 | 0.0396 | 0.0977 | 2.4651 |
| cluster3053 | -9367.7123 | 0.0846 | 0.054 | 0.642 | 0.1377 | 0.0937 | 0.6802 |
| cluster3054 | -6880.4734 | 0.0098 | 0.005 | 0.534 | 0.0087 | 0.0135 | 1.5548 |
| cluster3055 | -3987.8395 | 0.0408 | 0.019 | 0.463 | 0.0150 | 0.0195 | 1.2963 |
| cluster3056 | -5994.7484 | 0.0958 | 0.022 | 0.233 | 0.1152 | 0.0486 | 0.4217 |
| cluster3057 | -919.61379 | 0.0461 | 0.010 | 0.215 | 0.0687 | 0.0364 | 0.5296 |
| cluster3058 | -2532.8503 | 0.0335 | 0.009 | 0.282 | 0.0088 | 0.0044 | 0.4972 |
| cluster3059 | -8711.2202 | 0.0752 | 0.088 | 1.173 | 0.0343 | 0.2461 | 7.1802 |
| cluster3060 | -5326.3702 | 0.0359 | 0.022 | 0.620 | 0.0280 | 0.0698 | 2.4972 |
| cluster3061 | -5740.9505 | 0.0160 | 0.006 | 0.396 | 0.0156 | 0.0065 | 0.4159 |
| cluster3062 | -5690.835 | 0.0344 | 0.010 | 0.287 | 0.0410 | 0.0329 | 0.8038 |
| cluster3063 | -6146.9144 | 0.0290 | 0.014 | 0.485 | 0.0395 | 0.0579 | 1.4672 |
| cluster3064 | -1916.3081 | 0.0189 | 0.004 | 0.212 | 0.0044 | 0.0036 | 0.8301 |
| cluster3065 | -2297.4051 | 0.1146 | 0.049 | 0.429 | 0.0528 | 0.1339 | 2.5372 |
| cluster3066 | -6187.5588 | 0.0704 | 0.033 | 0.466 | 0.0656 | 0.0506 | 0.7717 |
| cluster3067 | -4910.5045 | 0.0313 | 0.019 | 0.610 | 0.0930 | 0.0538 | 0.5785 |
| cluster3068 | -5143.2861 | 0.1142 | 0.025 | 0.215 | 0.1001 | 0.0474 | 0.4737 |
| cluster3069 | -1631.8848 | 0.0001 | 0.000 | 0.098 | 0.0025 | 0.0013 | 0.5289 |
| cluster3070 | -5192.2323 | 0.0352 | 0.022 | 0.635 | 0.0418 | 0.0441 | 1.0548 |

| | | | | | | | |
|---|---|---|---|---|---|---|---|
| cluster3071 | -5855.6765 | 0.0192 | 0.008 | 0.396 | 0.0334 | 0.0185 | 0.5543 |
| cluster3072 | -7373.2883 | 0.0876 | 0.054 | 0.620 | 0.0779 | 0.1169 | 1.5014 |
| cluster3073 | -3570.0775 | 0.0061 | 0.002 | 0.375 | 0.0094 | 0.0123 | 1.3045 |
| cluster3074 | -7646.6489 | 0.0323 | 0.026 | 0.806 | 0.0091 | 0.0927 | 10.1879 |
| cluster3075 | -5425.9334 | 0.0328 | 0.021 | 0.627 | 0.0560 | 0.0423 | 0.7562 |
| cluster3076 | -1808.3028 | 0.0130 | 0.036 | 2.793 | 0.0689 | 0.0563 | 0.8167 |
| cluster3077 | -4384.4591 | 0.0215 | 0.013 | 0.599 | 0.0288 | 0.0345 | 1.1996 |
| cluster3078 | -5025.9807 | 0.0254 | 0.014 | 0.543 | 0.0166 | 0.0257 | 1.5485 |
| cluster3079 | -7619.3481 | 0.0302 | 0.020 | 0.660 | 0.0525 | 0.0328 | 0.6255 |
| cluster3080 | -5649.7227 | 0.0150 | 0.007 | 0.499 | 0.0184 | 0.0289 | 1.5751 |
| cluster3081 | -3873.7706 | 0.0078 | 0.004 | 0.492 | 0.0180 | 0.0134 | 0.7441 |
| cluster3082 | -7187.3526 | 0.0561 | 0.032 | 0.571 | 0.0660 | 0.1056 | 1.6005 |
| cluster3083 | -4159.5153 | 0.0287 | 0.013 | 0.458 | 0.0139 | 0.0056 | 0.4015 |
| cluster3084 | -3197.8321 | 0.0341 | 0.004 | 0.129 | 0.0429 | 0.0136 | 0.3180 |
| cluster3085 | -7337.1771 | 0.0266 | 0.029 | 1.104 | 0.0043 | 0.0894 | 20.6137 |
| cluster3086 | -3398.1846 | 0.1281 | 0.052 | 0.403 | 0.0423 | 0.0273 | 0.6447 |
| cluster3087 | -6563.1174 | 0.0139 | 0.005 | 0.376 | 0.0277 | 0.0293 | 1.0558 |
| cluster3088 | -6393.2964 | 0.0640 | 0.028 | 0.433 | 0.0685 | 0.0695 | 1.0149 |
| cluster3089 | -3685.4085 | 0.0757 | 0.034 | 0.446 | 0.0625 | 0.0510 | 0.8167 |
| cluster3090 | -4952.1815 | 0.0047 | 0.003 | 0.625 | 0.0169 | 0.0272 | 1.6035 |
| cluster3091 | -5911.141 | 0.0567 | 0.018 | 0.315 | 0.0480 | 0.0259 | 0.5399 |
| cluster3092 | -4264.3194 | 0.0202 | 0.009 | 0.451 | 0.0130 | 0.0220 | 1.6993 |
| cluster3093 | -4540.4016 | 0.0112 | 0.004 | 0.337 | 0.0031 | 0.0173 | 5.6391 |
| cluster3094 | -7863.9743 | 0.0329 | 0.032 | 0.960 | 0.0207 | 0.1066 | 5.1618 |
| cluster3095 | -4870.6567 | 0.0066 | 0.005 | 0.800 | 0.0049 | 0.0032 | 0.6499 |
| cluster3096 | -6132.6089 | 0.0331 | 0.015 | 0.459 | 0.0344 | 0.0655 | 1.9074 |
| cluster3097 | -7077.1097 | 0.0195 | 0.007 | 0.343 | 0.0250 | 0.0156 | 0.6244 |
| cluster3098 | -1418.4659 | 0.0084 | 0.006 | 0.744 | 0.0052 | 0.0171 | 3.2763 |
| cluster3099 | -4271.8664 | 0.0446 | 0.027 | 0.606 | 0.0273 | 0.0239 | 0.8769 |
| cluster3100 | -4392.934 | 0.0375 | 0.019 | 0.500 | 0.0541 | 0.0823 | 1.5203 |
| cluster3101 | -3432.2309 | 0.0395 | 0.024 | 0.600 | 0.0903 | 0.0550 | 0.6099 |
| cluster3102 | -2229.5144 | 0.0115 | 0.010 | 0.862 | 0.0380 | 0.0182 | 0.4792 |
| cluster3103 | -5216.1985 | 0.0347 | 0.034 | 0.974 | 0.0459 | 0.1405 | 3.0612 |
| cluster3104 | -4327.8574 | 0.0213 | 0.012 | 0.555 | 0.0221 | 0.0322 | 1.4593 |
| cluster3105 | -7964.9655 | 0.0322 | 0.030 | 0.935 | 0.0094 | 0.0941 | 10.0366 |
| cluster3106 | -5132.2566 | 0.0028 | 0.001 | 0.419 | 0.0027 | 0.0017 | 0.6333 |
| cluster3107 | -3137.8786 | 0.0403 | 0.009 | 0.224 | 0.0424 | 0.0441 | 1.0405 |
| cluster3108 | -4331.54 | 0.1465 | 0.057 | 0.387 | 0.1221 | 0.1505 | 1.2324 |
| cluster3109 | -3462.1379 | 0.0305 | 0.021 | 0.678 | 0.0222 | 0.0447 | 2.0125 |
| cluster3110 | -6381.1757 | 0.1068 | 0.031 | 0.290 | 0.1115 | 0.1235 | 1.1075 |
| cluster3111 | -4888.9738 | 0.0301 | 0.011 | 0.374 | 0.0371 | 0.0529 | 1.4265 |
| cluster3112 | -6143.2089 | 0.0418 | 0.035 | 0.830 | 0.0124 | 0.1391 | 11.2372 |
| cluster3113 | -3067.7451 | 0.0287 | 0.016 | 0.574 | 0.0192 | 0.0415 | 2.1642 |
| cluster3114 | -4689.4919 | 0.0360 | 0.013 | 0.373 | 0.0267 | 0.0258 | 0.9665 |

| | | | | | | | |
|---|---|---|---|---|---|---|---|
| cluster3115 | -3777.5603 | 0.0172 | 0.008 | 0.471 | 0.0195 | 0.0296 | 1.5189 |
| cluster3116 | -2720.7617 | 0.0289 | 0.014 | 0.479 | 0.0608 | 0.0744 | 1.2236 |
| cluster3117 | -7499.8739 | 0.0469 | 0.045 | 0.959 | 0.0514 | 0.1628 | 3.1689 |
| cluster3118 | -1571.9275 | 0.0553 | 0.007 | 0.122 | 0.0102 | 0.0048 | 0.4684 |
| cluster3119 | -2980.5052 | 0.0334 | 0.030 | 0.885 | 0.0781 | 0.0661 | 0.8453 |
| cluster3120 | -4910.784 | 0.0233 | 0.009 | 0.407 | 0.0220 | 0.0346 | 1.5725 |
| cluster3121 | -6820.713 | 0.0286 | 0.019 | 0.656 | 0.0203 | 0.0288 | 1.4233 |
| cluster3122 | -3556.3596 | 0.0172 | 0.007 | 0.379 | 0.0353 | 0.0189 | 0.5342 |
| cluster3123 | -3823.4766 | 0.0086 | 0.004 | 0.414 | 0.0352 | 0.0222 | 0.6301 |
| cluster3124 | -6377.4986 | 0.0381 | 0.020 | 0.528 | 0.0413 | 0.0608 | 1.4725 |
| cluster3125 | -4798.8322 | 0.0532 | 0.019 | 0.354 | 0.0970 | 0.1051 | 1.0840 |
| cluster3126 | -7547.9669 | 0.0368 | 0.022 | 0.606 | 0.0690 | 0.0532 | 0.7711 |
| cluster3127 | -2606.9446 | 0.0098 | 0.005 | 0.471 | 0.0059 | 0.0176 | 2.9901 |
| cluster3128 | -2238.015 | 0.0029 | 0.001 | 0.285 | 0.0012 | 0.0011 | 0.9358 |
| cluster3129 | -3263.5922 | 0.0384 | 0.051 | 1.331 | 0.0067 | 0.1566 | 23.3853 |
| cluster3130 | -5799.3255 | 0.0368 | 0.026 | 0.703 | 0.0328 | 0.0835 | 2.5446 |
| cluster3131 | -1764.3773 | 0.0094 | 0.008 | 0.821 | 0.0061 | 0.0168 | 2.7617 |
| cluster3132 | -5289.0851 | 0.0225 | 0.011 | 0.503 | 0.0329 | 0.0255 | 0.7753 |
| cluster3133 | -8922.8803 | 0.0659 | 0.070 | 1.060 | 0.0124 | 0.1843 | 14.8962 |
| cluster3134 | -3290.8696 | 0.0555 | 0.016 | 0.290 | 0.0654 | 0.0433 | 0.6620 |
| cluster3135 | -4393.0146 | 0.0174 | 0.006 | 0.344 | 0.0051 | 0.0110 | 2.1521 |
| cluster3136 | -3954.0853 | 0.0382 | 0.027 | 0.710 | 0.0372 | 0.0775 | 2.0870 |
| cluster3137 | -3090.9665 | 0.0253 | 0.018 | 0.715 | 0.0236 | 0.0550 | 2.3326 |
| cluster3138 | -4742.5427 | 0.0505 | 0.038 | 0.757 | 0.0665 | 0.0735 | 1.1057 |
| cluster3139 | -5752.0077 | 0.0739 | 0.078 | 1.049 | 0.0446 | 0.1558 | 3.4953 |
| cluster3140 | -3559.5209 | 0.0372 | 0.009 | 0.233 | 0.0529 | 0.0316 | 0.5967 |
| cluster3141 | -2415.3104 | 0.0096 | 0.002 | 0.256 | 0.0110 | 0.0070 | 0.6352 |
| cluster3142 | -1639.6525 | 0.1474 | 0.011 | 0.073 | 0.0467 | 0.0132 | 0.2828 |
| cluster3143 | -3225.9444 | 0.0169 | 0.007 | 0.416 | 0.0056 | 0.0037 | 0.6662 |
| cluster3144 | -2087.8835 | 0.0039 | 0.012 | 2.940 | 0.0102 | 0.0939 | 9.2102 |
| cluster3145 | -6529.0763 | 0.0064 | 0.005 | 0.865 | 0.0035 | 0.0114 | 3.2206 |
| cluster3146 | -1742.0745 | 0.0022 | 0.011 | 4.996 | 0.0041 | 0.0037 | 0.9153 |
| cluster3147 | -4472.7641 | 0.0054 | 0.003 | 0.519 | 0.0075 | 0.0137 | 1.8318 |
| cluster3148 | -6023.3199 | 0.0284 | 0.011 | 0.382 | 0.0300 | 0.0320 | 1.0656 |
| cluster3149 | -2367.3474 | 0.0001 | 0.000 | 0.353 | 0.0092 | 0.0063 | 0.6893 |
| cluster3150 | -4523.6234 | 0.0586 | 0.011 | 0.193 | 0.0789 | 0.0547 | 0.6937 |
| cluster3151 | -3975.377 | 0.0515 | 0.018 | 0.351 | 0.0502 | 0.0561 | 1.1181 |
| cluster3152 | -8745.7337 | 0.0502 | 0.053 | 1.065 | 0.0424 | 0.1428 | 3.3636 |
| cluster3153 | -4517.0727 | 0.0424 | 0.014 | 0.328 | 0.0442 | 0.0470 | 1.0636 |
| cluster3154 | -5342.5355 | 0.0168 | 0.011 | 0.667 | 0.0100 | 0.0204 | 2.0515 |
| cluster3155 | -3013.623 | 0.0064 | 0.003 | 0.544 | 0.0125 | 0.0231 | 1.8478 |
| cluster3156 | -6787.9804 | 0.0283 | 0.019 | 0.687 | 0.0172 | 0.0866 | 5.0403 |
| cluster3157 | -3279.8574 | 0.1147 | 0.017 | 0.147 | 0.0799 | 0.0513 | 0.6424 |
| cluster3158 | -3371.0316 | 0.0128 | 0.012 | 0.930 | 0.0293 | 0.0223 | 0.7607 |

| | | | | | | | |
|---|---|---|---|---|---|---|---|
| cluster3159 | -5812.8217 | 0.0504 | 0.043 | 0.852 | 0.0352 | 0.1287 | 3.6513 |
| cluster3160 | -1098.0644 | 0.0715 | 0.009 | 0.125 | 0.0186 | 0.0055 | 0.2933 |
| cluster3161 | -4981.6129 | 0.0292 | 0.016 | 0.551 | 0.0403 | 0.1313 | 3.2553 |
| cluster3162 | -3639.2597 | 0.0071 | 0.002 | 0.213 | 0.0111 | 0.0109 | 0.9833 |
| cluster3163 | -3430.0267 | 0.0349 | 0.022 | 0.621 | 0.0438 | 0.1143 | 2.6126 |
| cluster3164 | -6178.7255 | 0.0634 | 0.042 | 0.669 | 0.0416 | 0.1057 | 2.5418 |
| cluster3165 | -8049.4939 | 0.0584 | 0.047 | 0.803 | 0.0469 | 0.1352 | 2.8835 |
| cluster3166 | -4633.4653 | 0.0241 | 0.014 | 0.589 | 0.0294 | 0.0196 | 0.6686 |
| cluster3167 | -5313.4519 | 0.0889 | 0.021 | 0.238 | 0.0999 | 0.0974 | 0.9748 |
| cluster3168 | -7086.5889 | 0.0152 | 0.018 | 1.164 | 0.0342 | 0.0344 | 1.0057 |
| cluster3169 | -5027.3939 | 0.0321 | 0.024 | 0.752 | 0.0100 | 0.0836 | 8.3658 |
| cluster3170 | -1622.4211 | 0.0069 | 0.001 | 0.085 | 0.0193 | 0.0072 | 0.3733 |
| cluster3171 | -6131.9886 | 0.0412 | 0.018 | 0.436 | 0.1713 | 0.1376 | 0.8031 |
| cluster3172 | -6262.1311 | 0.0272 | 0.016 | 0.596 | 0.0304 | 0.0297 | 0.9743 |
| cluster3173 | -4457.6863 | 0.0156 | 0.009 | 0.596 | 0.0139 | 0.0293 | 2.1103 |
| cluster3174 | -1301.2501 | 0.0024 | 0.001 | 0.466 | 0.0015 | 0.0023 | 1.4874 |
| cluster3175 | -1938.2118 | 0.0159 | 0.005 | 0.317 | 0.0159 | 0.0226 | 1.4204 |
| cluster3176 | -3886.0633 | 0.0199 | 0.007 | 0.354 | 0.0344 | 0.0255 | 0.7396 |
| cluster3177 | -3505.2032 | 0.0620 | 0.019 | 0.305 | 0.0361 | 0.0397 | 1.1022 |
| cluster3178 | -6744.6144 | 0.0105 | 0.011 | 1.059 | 0.0158 | 0.0162 | 1.0225 |
| cluster3179 | -4630.4575 | 0.0645 | 0.064 | 0.998 | 0.0791 | 0.2372 | 2.9991 |
| cluster3180 | -1070.2742 | 0.0048 | 0.002 | 0.400 | 0.0003 | 0.0220 | 83.4981 |
| cluster3181 | -6693.4655 | 0.0215 | 0.018 | 0.820 | 0.0212 | 0.0250 | 1.1827 |
| cluster3182 | -2529.9485 | 0.0079 | 0.003 | 0.413 | 0.0029 | 0.0018 | 0.6308 |
| cluster3183 | -4042.3026 | 0.0858 | 0.052 | 0.612 | 0.0710 | 0.1021 | 1.4382 |
| cluster3184 | -4203.2805 | 0.0228 | 0.007 | 0.295 | 0.0469 | 0.0431 | 0.9188 |
| cluster3185 | -3546.3324 | 0.0183 | 0.014 | 0.791 | 0.0139 | 0.0459 | 3.2961 |
| cluster3186 | -2290.3133 | 0.0255 | 0.008 | 0.330 | 0.0336 | 0.0545 | 1.6248 |
| cluster3187 | -5390.6748 | 0.0169 | 0.008 | 0.491 | 0.0297 | 0.0488 | 1.6441 |
| cluster3188 | -4445.3044 | 0.0917 | 0.039 | 0.430 | 0.0840 | 0.0483 | 0.5751 |
| cluster3189 | -6229.4631 | 0.0420 | 0.030 | 0.706 | 0.0456 | 0.1361 | 2.9830 |
| cluster3190 | -5397.9414 | 0.0323 | 0.032 | 1.004 | 0.0259 | 0.0987 | 3.8047 |
| cluster3191 | -2831.3223 | 0.0438 | 0.025 | 0.565 | 0.0569 | 0.0468 | 0.8212 |
| cluster3192 | -5886.9225 | 0.0500 | 0.023 | 0.456 | 0.0597 | 0.0377 | 0.6321 |
| cluster3193 | -3453.7321 | 0.0453 | 0.014 | 0.306 | 0.0220 | 0.0171 | 0.7767 |
| cluster3194 | -6978.589 | 0.0625 | 0.046 | 0.735 | 0.0185 | 0.1436 | 7.7530 |
| cluster3195 | -3959.3855 | 0.0201 | 0.014 | 0.691 | 0.0106 | 0.0243 | 2.3032 |
| cluster3196 | -2244.728 | 0.0046 | 0.001 | 0.159 | 0.0043 | 0.0014 | 0.3188 |
| cluster3197 | -5447.1227 | 0.0530 | 0.021 | 0.399 | 0.0469 | 0.0245 | 0.5235 |
| cluster3198 | -6431.8871 | 0.0251 | 0.021 | 0.817 | 0.0210 | 0.0902 | 4.2915 |
| cluster3199 | -2979.3607 | 0.0129 | 0.005 | 0.377 | 0.0085 | 0.0111 | 1.3175 |
| cluster3200 | -4274.0847 | 0.0104 | 0.009 | 0.907 | 0.0169 | 0.0294 | 1.7456 |
| cluster3201 | -4476.7013 | 0.1102 | 0.018 | 0.164 | 0.1120 | 0.0678 | 0.6051 |
| cluster3202 | -6071.0923 | 0.0453 | 0.016 | 0.358 | 0.0386 | 0.0535 | 1.3865 |

| | | | | | | | |
|---|---|---|---|---|---|---|---|
| cluster3203 | -3017.2534 | 0.0512 | 0.034 | 0.659 | 0.0392 | 0.1768 | 4.5135 |
| cluster3204 | -1877.9824 | 0.0425 | 0.008 | 0.196 | 0.0286 | 0.0093 | 0.3257 |
| cluster3205 | -2015.7109 | 0.0053 | 0.001 | 0.164 | 0.0079 | 0.0029 | 0.3681 |
| cluster3206 | -3914.7694 | 0.0333 | 0.021 | 0.633 | 0.0367 | 0.0685 | 1.8657 |
| cluster3207 | -5497.8839 | 0.0119 | 0.010 | 0.807 | 0.0252 | 0.0292 | 1.1598 |
| cluster3208 | -3283.7202 | 0.0027 | 0.001 | 0.356 | 0.0100 | 0.0083 | 0.8328 |
| cluster3209 | -5550.277 | 0.0206 | 0.017 | 0.813 | 0.0318 | 0.0639 | 2.0092 |
| cluster3210 | -3374.9178 | 0.0015 | 0.001 | 0.757 | 0.0073 | 0.0440 | 6.0118 |
| cluster3211 | -3434.4937 | 0.0288 | 0.017 | 0.577 | 0.0393 | 0.0863 | 2.1950 |
| cluster3212 | -2808.0902 | 0.0009 | 0.000 | 0.428 | 0.0109 | 0.0041 | 0.3799 |
| cluster3213 | -4610.6386 | 0.0805 | 0.059 | 0.731 | 0.2072 | 0.1067 | 0.5148 |
| cluster3214 | -6051.8406 | 0.0460 | 0.025 | 0.552 | 0.0522 | 0.0385 | 0.7385 |
| cluster3215 | -2279.3183 | 0.0027 | 0.000 | 0.083 | 0.0020 | 0.0009 | 0.4729 |
| cluster3216 | -4538.4937 | 0.0305 | 0.040 | 1.320 | 0.0250 | 0.1359 | 5.4325 |
| cluster3217 | -3824.909 | 0.0410 | 0.017 | 0.404 | 0.0458 | 0.0357 | 0.7793 |
| cluster3218 | -5083.1621 | 0.0270 | 0.020 | 0.750 | 0.0144 | 0.0793 | 5.5048 |
| cluster3219 | -5449.5645 | 0.0115 | 0.006 | 0.559 | 0.0142 | 0.0332 | 2.3418 |
| cluster3220 | -2924.5718 | 0.0299 | 0.012 | 0.386 | 0.0327 | 0.0190 | 0.5808 |
| cluster3221 | -5736.2677 | 0.0322 | 0.032 | 0.998 | 0.0348 | 0.1002 | 2.8821 |
| cluster3222 | -2622.0523 | 0.0397 | 0.014 | 0.350 | 0.0409 | 0.0425 | 1.0400 |
| cluster3223 | -3466.4252 | 0.0831 | 0.046 | 0.558 | 0.1163 | 0.1215 | 1.0454 |
| cluster3224 | -1597.2435 | 0.0001 | 0.000 | 0.129 | 0.0001 | 0.0001 | 0.7919 |
| cluster3225 | -6105.4048 | 0.0301 | 0.019 | 0.637 | 0.0370 | 0.0797 | 2.1540 |
| cluster3226 | -3727.0457 | 0.0434 | 0.020 | 0.467 | 0.0336 | 0.0666 | 1.9837 |
| cluster3227 | -4398.0173 | 0.0649 | 0.051 | 0.784 | 0.0845 | 0.1493 | 1.7675 |
| cluster3228 | -3624.1336 | 0.0357 | 0.029 | 0.805 | 0.0191 | 0.0856 | 4.4804 |
| cluster3229 | -3867.9395 | 0.0348 | 0.020 | 0.585 | 0.0160 | 0.0147 | 0.9207 |
| cluster3230 | -3482.0505 | 0.0100 | 0.005 | 0.455 | 0.0149 | 0.0093 | 0.6234 |
| cluster3231 | -4528.9023 | 0.0234 | 0.011 | 0.490 | 0.0264 | 0.0392 | 1.4824 |
| cluster3232 | -5052.9365 | 0.0248 | 0.019 | 0.772 | 0.0167 | 0.0488 | 2.9262 |
| cluster3233 | -3435.4738 | 0.0381 | 0.036 | 0.954 | 0.0683 | 0.0501 | 0.7337 |
| cluster3234 | -3820.1209 | 0.0458 | 0.027 | 0.584 | 0.0697 | 0.0565 | 0.8111 |
| cluster3235 | -7270.9712 | 0.0393 | 0.039 | 1.000 | 0.0223 | 0.1547 | 6.9370 |
| cluster3236 | -3956.653 | 0.0323 | 0.024 | 0.755 | 0.0354 | 0.0774 | 2.1865 |
| cluster3237 | -3244.3114 | 0.0628 | 0.038 | 0.602 | 0.0488 | 0.0849 | 1.7393 |
| cluster3238 | -2758.6674 | 0.1010 | 0.022 | 0.214 | 0.0019 | 0.0010 | 0.5445 |
| cluster3239 | -2739.4629 | 0.0272 | 0.008 | 0.285 | 0.0152 | 0.0124 | 0.8191 |
| cluster3240 | -6958.0213 | 0.0276 | 0.021 | 0.767 | 0.0549 | 0.0467 | 0.8507 |
| cluster3241 | -5756.5255 | 0.0647 | 0.035 | 0.536 | 0.0645 | 0.1297 | 2.0100 |
| cluster3242 | -7162.8983 | 0.0466 | 0.048 | 1.019 | 0.0351 | 0.1407 | 4.0154 |
| cluster3243 | -5984.2603 | 0.1502 | 0.042 | 0.277 | 0.1596 | 0.0679 | 0.4255 |
| cluster3244 | -6472.3897 | 0.0559 | 0.034 | 0.616 | 0.0783 | 0.0535 | 0.6827 |
| cluster3245 | -2585.8476 | 0.0031 | 0.002 | 0.601 | 0.0026 | 0.0055 | 2.0927 |
| cluster3246 | -5132.3821 | 0.0373 | 0.040 | 1.068 | 0.0726 | 0.0651 | 0.8974 |

| | | | | | | | |
|---|---|---|---|---|---|---|---|
| cluster3247 | -849.1682 | 0.0592 | 0.020 | 0.336 | 0.0735 | 0.0979 | 1.3329 |
| cluster3248 | -3485.0826 | 0.0216 | 0.013 | 0.622 | 0.0208 | 0.0375 | 1.7995 |
| cluster3249 | -3481.7699 | 0.0198 | 0.011 | 0.534 | 0.0118 | 0.0291 | 2.4628 |
| cluster3250 | -3544.9695 | 0.1428 | 0.035 | 0.246 | 0.0505 | 0.0415 | 0.8214 |
| cluster3251 | -4467.6077 | 0.0589 | 0.026 | 0.436 | 0.0553 | 0.0720 | 1.3020 |
| cluster3252 | -7422.98 | 0.0636 | 0.065 | 1.025 | 0.0641 | 0.1646 | 2.5668 |
| cluster3253 | -4316.6974 | 0.0644 | 0.031 | 0.485 | 0.0864 | 0.1102 | 1.2759 |
| cluster3254 | -3110.2672 | 0.0367 | 0.020 | 0.543 | 0.0303 | 0.0564 | 1.8640 |
| cluster3255 | -5568.3109 | 0.0200 | 0.020 | 0.998 | 0.0096 | 0.0681 | 7.0954 |
| cluster3256 | -6101.1054 | 0.0395 | 0.026 | 0.650 | 0.0455 | 0.0634 | 1.3931 |
| cluster3257 | -5141.3181 | 0.0049 | 0.004 | 0.772 | 0.0135 | 0.0107 | 0.7917 |
| cluster3258 | -1917.9462 | 0.0001 | 0.000 | 0.258 | 0.0028 | 0.0029 | 1.0354 |
| cluster3259 | -3556.3913 | 0.0374 | 0.009 | 0.242 | 0.0308 | 0.0098 | 0.3191 |
| cluster3260 | -4482.5035 | 0.0135 | 0.008 | 0.556 | 0.0145 | 0.0108 | 0.7428 |
| cluster3261 | -4947.9819 | 0.0731 | 0.063 | 0.865 | 0.0587 | 0.1746 | 2.9730 |
| cluster3262 | -5791.5301 | 0.0185 | 0.015 | 0.835 | 0.0186 | 0.0624 | 3.3618 |
| cluster3263 | -2046.4244 | 0.1929 | 0.060 | 0.313 | 0.0421 | 0.0331 | 0.7857 |
| cluster3264 | -1834.3823 | 0.0147 | 0.003 | 0.179 | 0.0062 | 0.0054 | 0.8660 |
| cluster3265 | -4128.3117 | 0.0023 | 0.001 | 0.585 | 0.0041 | 0.0062 | 1.5370 |
| cluster3266 | -2855.4226 | 0.0365 | 0.008 | 0.208 | 0.0201 | 0.0132 | 0.6547 |
| cluster3267 | -4122.6569 | 0.0357 | 0.022 | 0.605 | 0.0171 | 0.0142 | 0.8266 |
| cluster3268 | -2801.73 | 0.0498 | 0.022 | 0.444 | 0.0182 | 0.0347 | 1.9106 |
| cluster3269 | -3530.6067 | 0.0425 | 0.019 | 0.459 | 0.1087 | 0.1332 | 1.2260 |
| cluster3270 | -2185.3885 | 0.0264 | 0.018 | 0.666 | 0.0255 | 0.0696 | 2.7249 |
| cluster3271 | -4111.6368 | 0.0105 | 0.005 | 0.492 | 0.0185 | 0.0127 | 0.6853 |
| cluster3272 | -939.60961 | 0.3125 | 0.026 | 0.085 | 0.0001 | 0.0000 | 0.0000 |
| cluster3273 | -4005.5237 | 0.1192 | 0.068 | 0.569 | 0.0982 | 0.2056 | 2.0942 |
| cluster3274 | -2028.8301 | 0.0398 | 0.031 | 0.786 | 0.0082 | 0.0720 | 8.7389 |
| cluster3275 | -4550.6609 | 0.0455 | 0.034 | 0.746 | 0.0595 | 0.0969 | 1.6294 |
| cluster3276 | -2485.6793 | 0.0896 | 0.015 | 0.172 | 0.0909 | 0.0216 | 0.2371 |
| cluster3277 | -3226.1099 | 0.0530 | 0.029 | 0.552 | 0.0110 | 0.1508 | 13.7010 |
| cluster3278 | -4577.3471 | 0.0037 | 0.001 | 0.259 | 0.0025 | 0.0019 | 0.7551 |
| cluster3279 | -3732.7536 | 0.0154 | 0.010 | 0.636 | 0.0192 | 0.0450 | 2.3480 |
| cluster3280 | -3787.1994 | 0.0178 | 0.006 | 0.342 | 0.0287 | 0.0342 | 1.1915 |
| cluster3281 | -1963.3266 | 0.0277 | 0.020 | 0.715 | 0.0058 | 0.0293 | 5.0552 |
| cluster3282 | -2048.6828 | 0.0069 | 0.003 | 0.434 | 0.0036 | 0.0048 | 1.3287 |
| cluster3283 | -5464.5959 | 0.0670 | 0.037 | 0.550 | 0.0548 | 0.0500 | 0.9133 |
| cluster3284 | -5979.2333 | 0.0378 | 0.021 | 0.550 | 0.0280 | 0.0565 | 2.0181 |
| cluster3285 | -2360.235 | 0.0031 | 0.001 | 0.258 | 0.0131 | 0.0109 | 0.8387 |
| cluster3286 | -2205.943 | 0.0419 | 0.009 | 0.223 | 0.0149 | 0.0107 | 0.7138 |
| cluster3287 | -1377.2998 | 0.0144 | 0.006 | 0.401 | 0.0107 | 0.0146 | 1.3609 |
| cluster3288 | -1241.6018 | 0.0066 | 0.007 | 1.024 | 0.0060 | 0.0200 | 3.3428 |
| cluster3289 | -5798.724 | 0.0465 | 0.015 | 0.313 | 0.0522 | 0.0250 | 0.4789 |
| cluster3290 | -1251.1659 | 0.0270 | 0.012 | 0.429 | 0.0037 | 0.0196 | 5.3701 |

| | | | | | | | |
|---|---|---|---|---|---|---|---|
| cluster3291 | -4785.2893 | 0.0352 | 0.024 | 0.685 | 0.0625 | 0.0523 | 0.8360 |
| cluster3292 | -2768.3307 | 0.0382 | 0.013 | 0.331 | 0.0119 | 0.0182 | 1.5314 |
| cluster3293 | -3184.0302 | 0.0083 | 0.006 | 0.678 | 0.0140 | 0.0187 | 1.3341 |
| cluster3294 | -2883.1278 | 0.0126 | 0.009 | 0.701 | 0.0164 | 0.0166 | 1.0147 |
| cluster3295 | -2241.0851 | 0.0011 | 0.000 | 0.248 | 0.0021 | 0.0022 | 1.0434 |
| cluster3296 | -1799.0481 | 0.0057 | 0.004 | 0.679 | 0.0026 | 0.0078 | 2.9985 |
| cluster3297 | -2350.368 | 0.1255 | 0.039 | 0.313 | 0.3449 | 0.2279 | 0.6606 |
| cluster3298 | -2943.0271 | 0.0028 | 0.002 | 0.706 | 0.0077 | 0.0086 | 1.1160 |
| cluster3299 | -7896.8554 | 0.0601 | 0.091 | 1.510 | 0.0036 | 0.1871 | 51.7239 |
| cluster3300 | -2251.1376 | 0.0066 | 0.004 | 0.666 | 0.0020 | 0.0037 | 1.8591 |
| cluster3301 | -1804.4107 | 0.0488 | 0.017 | 0.343 | 0.0502 | 0.0261 | 0.5197 |
| cluster3302 | -4554.5495 | 0.1052 | 0.045 | 0.427 | 0.0486 | 0.0721 | 1.4823 |
| cluster3303 | -4478.3061 | 0.0308 | 0.033 | 1.063 | 0.0644 | 0.0718 | 1.1145 |
| cluster3304 | -2842.3811 | 0.0085 | 0.003 | 0.312 | 0.0768 | 0.0397 | 0.5169 |
| cluster3305 | -3805.2116 | 0.0615 | 0.032 | 0.526 | 0.0703 | 0.1239 | 1.7627 |
| cluster3306 | -5020.7485 | 0.0768 | 0.046 | 0.594 | 0.0582 | 0.1272 | 2.1848 |
| cluster3307 | -1670.3479 | 0.0066 | 0.003 | 0.485 | 0.0024 | 0.0071 | 3.0106 |
| cluster3308 | -4384.4466 | 0.0151 | 0.011 | 0.758 | 0.0331 | 0.0486 | 1.4669 |
| cluster3309 | -3385.2479 | 0.1176 | 0.039 | 0.330 | 0.1242 | 0.0505 | 0.4067 |
| cluster3310 | -4175.2738 | 0.0157 | 0.008 | 0.491 | 0.0161 | 0.0221 | 1.3733 |
| cluster3311 | -6158.1198 | 0.0546 | 0.036 | 0.659 | 0.0560 | 0.0612 | 1.0926 |
| cluster3312 | -1931.813 | 0.0277 | 0.028 | 1.006 | 0.0237 | 0.0597 | 2.5164 |
| cluster3313 | -2505.0091 | 0.0158 | 0.004 | 0.274 | 0.0018 | 0.0013 | 0.7602 |
| cluster3314 | -7142.1391 | 0.0425 | 0.050 | 1.185 | 0.0456 | 0.1970 | 4.3243 |
| cluster3315 | -3720.9416 | 0.0044 | 0.003 | 0.613 | 0.0120 | 0.0092 | 0.7676 |
| cluster3316 | -5972.9719 | 0.0906 | 0.052 | 0.572 | 0.0468 | 0.1615 | 3.4490 |
| cluster3317 | -1141.7221 | 0.0050 | 0.002 | 0.393 | 0.0012 | 0.0020 | 1.6140 |
| cluster3318 | -3441.5979 | 0.1043 | 0.097 | 0.930 | 0.1374 | 0.1490 | 1.0845 |
| cluster3319 | -3769.2223 | 0.0379 | 0.031 | 0.808 | 0.0296 | 0.0783 | 2.6469 |
| cluster3320 | -2813.5758 | 0.0505 | 0.030 | 0.585 | 0.0654 | 0.1215 | 1.8565 |
| cluster3321 | -2916.1485 | 0.0048 | 0.001 | 0.258 | 0.0009 | 0.0016 | 1.7837 |
| cluster3322 | -3279.1266 | 0.0422 | 0.034 | 0.815 | 0.0376 | 0.1065 | 2.8308 |
| cluster3323 | -5518.3796 | 0.0560 | 0.031 | 0.557 | 0.0349 | 0.0757 | 2.1690 |
| cluster3324 | -1400.474 | 0.0189 | 0.034 | 1.781 | 0.0289 | 0.0203 | 0.7017 |
| cluster3325 | -2060.4462 | 0.0161 | 0.011 | 0.668 | 0.0254 | 0.0461 | 1.8202 |
| cluster3326 | -1703.0956 | 0.0970 | 0.011 | 0.110 | 0.0534 | 0.0317 | 0.5937 |
| cluster3327 | -3864.9561 | 0.0258 | 0.017 | 0.657 | 0.0233 | 0.0188 | 0.8052 |
| cluster3328 | -2603.1074 | 0.0098 | 0.003 | 0.305 | 0.0372 | 0.0605 | 1.6249 |
| cluster3329 | -3152.2577 | 0.0729 | 0.064 | 0.878 | 0.0361 | 0.1536 | 4.2621 |
| cluster3330 | -4708.1042 | 0.0572 | 0.039 | 0.690 | 0.0291 | 0.1021 | 3.5052 |
| cluster3331 | -2868.9679 | 0.0784 | 0.019 | 0.248 | 0.0670 | 0.0576 | 0.8600 |
| cluster3332 | -6187.1015 | 0.0719 | 0.055 | 0.770 | 0.0341 | 0.1475 | 4.3312 |
| cluster3333 | -3575.3736 | 0.0254 | 0.016 | 0.630 | 0.0147 | 0.0615 | 4.1874 |
| cluster3334 | -1379.0659 | 0.0924 | 0.025 | 0.271 | 0.0433 | 0.0484 | 1.1161 |

| | | | | | | | |
|---|---|---|---|---|---|---|---|
| cluster3335 | -2858.1841 | 0.0270 | 0.027 | 1.010 | 0.0382 | 0.0284 | 0.7425 |
| cluster3336 | -3026.9657 | 0.0049 | 0.002 | 0.362 | 0.0001 | 0.0001 | 0.5662 |
| cluster3337 | -2618.7378 | 0.0358 | 0.009 | 0.243 | 0.0149 | 0.0058 | 0.3921 |
| cluster3338 | -6301.5847 | 0.0422 | 0.040 | 0.948 | 0.0247 | 0.1283 | 5.1929 |
| cluster3339 | -1908.5432 | 0.0370 | 0.037 | 0.990 | 0.1195 | 0.0640 | 0.5353 |
| cluster3340 | -1625.2701 | 0.2053 | 0.028 | 0.135 | 0.0813 | 0.0164 | 0.2013 |
| cluster3341 | -1574.5319 | 0.0129 | 0.009 | 0.715 | 0.0009 | 0.0355 | 39.9887 |
| cluster3342 | -2556.1602 | 0.0195 | 0.015 | 0.785 | 0.0261 | 0.0674 | 2.5790 |
| cluster3343 | -2925.3124 | 0.0001 | 0.000 | 0.551 | 0.0009 | 0.0019 | 2.0075 |
| cluster3344 | -3149.2884 | 0.0256 | 0.008 | 0.308 | 0.0293 | 0.0174 | 0.5937 |
| cluster3345 | -2795.817 | 0.0134 | 0.010 | 0.778 | 0.0118 | 0.0448 | 3.7927 |
| cluster3346 | -3610.2765 | 0.0034 | 0.002 | 0.553 | 0.0039 | 0.0050 | 1.2983 |
| cluster3347 | -4173.968 | 0.0405 | 0.015 | 0.376 | 0.0684 | 0.0392 | 0.5736 |
| cluster3348 | -4559.3908 | 0.0453 | 0.012 | 0.275 | 0.0573 | 0.0575 | 1.0038 |
| cluster3349 | -4213.6409 | 0.0079 | 0.004 | 0.567 | 0.0174 | 0.0216 | 1.2431 |
| cluster3350 | -4778.8457 | 0.0490 | 0.033 | 0.677 | 0.0533 | 0.1045 | 1.9599 |
| cluster3351 | -3133.4194 | 0.0136 | 0.006 | 0.421 | 0.0210 | 0.0436 | 2.0804 |
| cluster3352 | -4082.6151 | 0.0470 | 0.016 | 0.333 | 0.0782 | 0.0916 | 1.1712 |
| cluster3353 | -1582.9037 | 0.0034 | 0.001 | 0.263 | 0.0200 | 0.0203 | 1.0115 |
| cluster3354 | -2119.988 | 0.0088 | 0.002 | 0.181 | 0.0029 | 0.0029 | 1.0013 |
| cluster3355 | -4287.2769 | 0.0464 | 0.031 | 0.661 | 0.0348 | 0.0522 | 1.5024 |
| cluster3356 | -4591.5741 | 0.0327 | 0.014 | 0.427 | 0.0672 | 0.0477 | 0.7106 |
| cluster3357 | -1290.0335 | 0.0256 | 0.010 | 0.404 | 0.0317 | 0.0356 | 1.1231 |
| cluster3358 | -4171.7007 | 0.0018 | 0.001 | 0.462 | 0.0006 | 0.0003 | 0.5367 |
| cluster3359 | -6042.2316 | 0.0353 | 0.028 | 0.781 | 0.0513 | 0.1270 | 2.4762 |
| cluster3360 | -2471.1886 | 0.0407 | 0.030 | 0.741 | 0.0270 | 0.0328 | 1.2170 |
| cluster3361 | -5033.816 | 0.0390 | 0.032 | 0.817 | 0.0138 | 0.1243 | 9.0272 |
| cluster3362 | -3689.6446 | 0.0129 | 0.007 | 0.580 | 0.0297 | 0.0251 | 0.8449 |
| cluster3363 | -3720.7294 | 0.0019 | 0.002 | 0.900 | 0.0033 | 0.0216 | 6.5790 |
| cluster3364 | -4353.1398 | 0.0249 | 0.022 | 0.902 | 0.0041 | 0.0747 | 18.4229 |
| cluster3365 | -5565.8397 | 0.0643 | 0.035 | 0.538 | 0.0574 | 0.1058 | 1.8419 |
| cluster3366 | -3146.1149 | 0.0351 | 0.028 | 0.801 | 0.0082 | 0.1577 | 19.3648 |
| cluster3367 | -3693.5185 | 0.1271 | 0.067 | 0.530 | 0.1651 | 0.0968 | 0.5863 |
| cluster3368 | -1645.0316 | 0.0063 | 0.001 | 0.147 | 0.0812 | 0.0268 | 0.3298 |
| cluster3369 | -3763.7903 | 0.0125 | 0.011 | 0.852 | 0.0232 | 0.0430 | 1.8573 |
| cluster3370 | -6386.7462 | 0.0927 | 0.045 | 0.485 | 0.1213 | 0.1767 | 1.4567 |
| cluster3371 | -2934.491 | 0.0059 | 0.002 | 0.272 | 0.0065 | 0.0057 | 0.8731 |
| cluster3372 | -3841.2886 | 0.0080 | 0.007 | 0.913 | 0.0103 | 0.0168 | 1.6283 |
| cluster3373 | -5275.1955 | 0.0070 | 0.006 | 0.856 | 0.0134 | 0.0330 | 2.4616 |
| cluster3374 | -5125.5068 | 0.0704 | 0.075 | 1.072 | 0.0732 | 0.2294 | 3.1320 |
| cluster3375 | -3051.6304 | 0.0040 | 0.001 | 0.248 | 0.0078 | 0.0062 | 0.7884 |
| cluster3376 | -1525.7958 | 0.0579 | 0.011 | 0.193 | 0.0083 | 0.0047 | 0.5648 |
| cluster3377 | -2273.8237 | 0.0129 | 0.010 | 0.753 | 0.0095 | 0.0331 | 3.4810 |
| cluster3378 | -6507.5654 | 0.0503 | 0.044 | 0.881 | 0.0374 | 0.0407 | 1.0888 |

| | | | | | | | |
|---|---|---|---|---|---|---|---|
| cluster3379 | -1567.7767 | 0.0398 | 0.015 | 0.365 | 0.0248 | 0.0354 | 1.4270 |
| cluster3380 | -2650.1969 | 0.0126 | 0.005 | 0.362 | 0.0412 | 0.0635 | 1.5404 |
| cluster3381 | -2267.8114 | 0.0314 | 0.018 | 0.561 | 0.0418 | 0.0341 | 0.8150 |
| cluster3382 | -885.29068 | 0.0102 | 0.002 | 0.186 | 0.0040 | 0.0050 | 1.2678 |
| cluster3383 | -5146.2886 | 0.0615 | 0.049 | 0.804 | 0.0043 | 0.1498 | 34.5627 |
| cluster3384 | -1969.0642 | 0.0421 | 0.021 | 0.488 | 0.0143 | 0.0166 | 1.1647 |
| cluster3385 | -3770.8981 | 0.0238 | 0.017 | 0.694 | 0.0149 | 0.0517 | 3.4763 |
| cluster3386 | -4155.8129 | 0.0150 | 0.011 | 0.718 | 0.0130 | 0.0350 | 2.6853 |
| cluster3387 | -3017.9803 | 0.0614 | 0.028 | 0.450 | 0.0097 | 0.0250 | 2.5698 |
| cluster3388 | -2808.8529 | 0.0280 | 0.014 | 0.513 | 0.0201 | 0.0264 | 1.3152 |
| cluster3389 | -1788.8177 | 0.0092 | 0.005 | 0.579 | 0.0124 | 0.0367 | 2.9574 |
| cluster3390 | -5117.1799 | 0.0649 | 0.030 | 0.460 | 0.0618 | 0.0608 | 0.9838 |
| cluster3391 | -2416.3859 | 0.0080 | 0.003 | 0.332 | 0.0089 | 0.0099 | 1.1090 |
| cluster3392 | -2524.1105 | 0.0069 | 0.006 | 0.866 | 0.0225 | 0.0374 | 1.6616 |
| cluster3393 | -3177.5798 | 0.0496 | 0.022 | 0.438 | 0.0852 | 0.1213 | 1.4236 |
| cluster3394 | -712.98762 | 0.0045 | 0.003 | 0.772 | 0.0001 | 0.0001 | 0.7565 |
| cluster3395 | -2714.0216 | 0.0164 | 0.011 | 0.646 | 0.0097 | 0.0079 | 0.8086 |
| cluster3396 | -4984.6181 | 0.0332 | 0.017 | 0.499 | 0.0483 | 0.0301 | 0.6245 |
| cluster3397 | -4638.3819 | 0.0327 | 0.023 | 0.710 | 0.0121 | 0.0730 | 6.0561 |
| cluster3398 | -3830.4708 | 0.0435 | 0.013 | 0.297 | 0.0785 | 0.0850 | 1.0828 |
| cluster3399 | -3230.5179 | 0.0423 | 0.029 | 0.681 | 0.0530 | 0.0625 | 1.1808 |
| cluster3400 | -1948.4789 | 0.1314 | 0.025 | 0.189 | 0.0164 | 0.0037 | 0.2226 |
| cluster3401 | -2870.6088 | 0.0157 | 0.007 | 0.443 | 0.0178 | 0.0382 | 2.1408 |
| cluster3402 | -5113.0494 | 0.0195 | 0.010 | 0.529 | 0.0326 | 0.0355 | 1.0920 |
| cluster3403 | -3669.313 | 0.0239 | 0.012 | 0.511 | 0.0188 | 0.0268 | 1.4266 |
| cluster3404 | -716.69613 | 0.0001 | 0.000 | 0.068 | 0.0001 | 0.0000 | 0.2088 |
| cluster3405 | -1541.2975 | 0.1174 | 0.035 | 0.301 | 0.1282 | 0.0835 | 0.6515 |
| cluster3406 | -3798.0219 | 0.0161 | 0.014 | 0.863 | 0.0367 | 0.0319 | 0.8694 |
| cluster3407 | -6244.3352 | 0.0598 | 0.053 | 0.895 | 0.0649 | 0.0729 | 1.1229 |
| cluster3408 | -1896.2041 | 0.0045 | 0.001 | 0.241 | 0.0103 | 0.0043 | 0.4167 |
| cluster3409 | -4909.0774 | 0.0219 | 0.017 | 0.755 | 0.0291 | 0.0877 | 3.0187 |
| cluster3410 | -1443.0154 | 0.0222 | 0.005 | 0.239 | 0.0288 | 0.0146 | 0.5065 |
| cluster3411 | -2582.5446 | 0.0001 | 0.000 | 0.310 | 0.0006 | 0.0011 | 1.9094 |
| cluster3412 | -3391.8683 | 0.0779 | 0.047 | 0.606 | 0.0983 | 0.1159 | 1.1790 |
| cluster3413 | -2555.3343 | 0.0607 | 0.032 | 0.521 | 0.0193 | 0.0556 | 2.8737 |
| cluster3414 | -2522.5698 | 0.1263 | 0.025 | 0.198 | 0.1662 | 0.0506 | 0.3043 |
| cluster3415 | -3492.7565 | 0.0694 | 0.023 | 0.330 | 0.0533 | 0.0559 | 1.0495 |
| cluster3416 | -601.09195 | 0.0150 | 0.342 | 22.750 | 0.0001 | 0.0010 | 9.7144 |
| cluster3417 | -1214.6063 | 0.0013 | 0.007 | 5.365 | 0.0130 | 0.0450 | 3.4735 |
| cluster3418 | -2533.6364 | 0.0508 | 0.028 | 0.560 | 0.0723 | 0.0999 | 1.3823 |
| cluster3419 | -2671.6442 | 0.1178 | 0.021 | 0.178 | 0.0743 | 0.0561 | 0.7550 |
| cluster3420 | -1783.3868 | 0.3063 | 0.101 | 0.330 | 0.1455 | 0.1134 | 0.7799 |
| cluster3421 | -4207.7638 | 0.0114 | 0.008 | 0.718 | 0.0107 | 0.0344 | 3.2160 |
| cluster3422 | -5399.0778 | 0.0335 | 0.017 | 0.505 | 0.0407 | 0.0387 | 0.9511 |

| | | | | | | | |
|---|---|---|---|---|---|---|---|
| cluster3423 | -4247.7209 | 0.0549 | 0.054 | 0.992 | 0.0230 | 0.1428 | 6.2086 |
| cluster3424 | -2856.5632 | 0.1031 | 0.049 | 0.471 | 0.0985 | 0.0996 | 1.0121 |
| cluster3425 | -2222.8643 | 0.0713 | 0.015 | 0.213 | 0.0134 | 0.0107 | 0.8044 |
| cluster3426 | -3406.1504 | 0.0141 | 0.014 | 0.969 | 0.0439 | 0.0451 | 1.0288 |
| cluster3427 | -3795.9274 | 0.1006 | 0.037 | 0.364 | 0.0752 | 0.0864 | 1.1496 |
| cluster3428 | -2685.0676 | 0.0584 | 0.023 | 0.398 | 0.0939 | 0.0378 | 0.4024 |
| cluster3429 | -4140.1853 | 0.0223 | 0.015 | 0.671 | 0.0569 | 0.0507 | 0.8919 |
| cluster3430 | -4824.5842 | 0.0425 | 0.049 | 1.152 | 0.0501 | 0.1251 | 2.4982 |
| cluster3431 | -4739.2184 | 0.0427 | 0.034 | 0.802 | 0.0404 | 0.1270 | 3.1474 |
| cluster3432 | -3781.893 | 0.0331 | 0.012 | 0.372 | 0.0492 | 0.0335 | 0.6811 |
| cluster3433 | -5392.3502 | 0.1839 | 0.052 | 0.283 | 0.1390 | 0.1023 | 0.7358 |
| cluster3434 | -3294.7419 | 0.0337 | 0.011 | 0.330 | 0.0458 | 0.0240 | 0.5253 |
| cluster3435 | -4093.0514 | 0.0199 | 0.014 | 0.680 | 0.0248 | 0.0411 | 1.6604 |
| cluster3436 | -3133.3965 | 0.0275 | 0.018 | 0.665 | 0.0138 | 0.0688 | 5.0062 |
| cluster3437 | -2816.4097 | 0.0133 | 0.011 | 0.793 | 0.0111 | 0.0181 | 1.6374 |
| cluster3438 | -3823.281 | 0.0438 | 0.022 | 0.492 | 0.0274 | 0.0445 | 1.6235 |
| cluster3439 | -2052.9726 | 0.0646 | 0.022 | 0.345 | 0.0155 | 0.1065 | 6.8646 |
| cluster3440 | -2313.4576 | 0.0210 | 0.006 | 0.305 | 0.0347 | 0.0251 | 0.7234 |
| cluster3441 | -2753.3636 | 0.0438 | 0.033 | 0.751 | 0.0105 | 0.0829 | 7.8606 |
| cluster3442 | -2923.4127 | 0.0185 | 0.031 | 1.653 | 0.0363 | 0.1169 | 3.2233 |
| cluster3443 | -3532.5213 | 0.0149 | 0.010 | 0.683 | 0.0144 | 0.0409 | 2.8405 |
| cluster3444 | -3384.822 | 0.0066 | 0.002 | 0.363 | 0.0145 | 0.0238 | 1.6358 |
| cluster3445 | -2957.5455 | 0.0077 | 0.003 | 0.361 | 0.0380 | 0.0211 | 0.5548 |
| cluster3446 | -4062.1338 | 0.0306 | 0.026 | 0.842 | 0.0311 | 0.0707 | 2.2718 |
| cluster3447 | -5924.5 | 0.0399 | 0.046 | 1.146 | 0.0193 | 0.1459 | 7.5560 |
| cluster3448 | -2378.8526 | 0.0223 | 0.016 | 0.695 | 0.0254 | 0.0837 | 3.2877 |
| cluster3449 | -612.22624 | 0.0001 | 0.000 | 0.128 | 0.0160 | 0.0033 | 0.2086 |
| cluster3450 | -2517.768 | 0.0229 | 0.016 | 0.686 | 0.0129 | 0.0739 | 5.7139 |
| cluster3451 | -3593.9531 | 0.0262 | 0.017 | 0.664 | 0.0341 | 0.0787 | 2.3097 |
| cluster3452 | -3964.3546 | 0.0441 | 0.031 | 0.708 | 0.0570 | 0.1052 | 1.8463 |
| cluster3453 | -1844.1054 | 0.0269 | 0.013 | 0.478 | 0.1435 | 0.0428 | 0.2982 |
| cluster3454 | -3695.7255 | 0.0111 | 0.012 | 1.051 | 0.0071 | 0.0461 | 6.5139 |
| cluster3455 | -3649.3168 | 0.0442 | 0.008 | 0.192 | 0.0425 | 0.0209 | 0.4910 |
| cluster3456 | -4229.7368 | 0.0307 | 0.009 | 0.306 | 0.0481 | 0.0388 | 0.8069 |
| cluster3457 | -3648.0555 | 0.0270 | 0.017 | 0.634 | 0.0429 | 0.0804 | 1.8730 |
| cluster3458 | -1805.6616 | 0.0390 | 0.016 | 0.408 | 0.0211 | 0.0188 | 0.8884 |
| cluster3459 | -2640.6912 | 0.0459 | 0.036 | 0.782 | 0.0336 | 0.1558 | 4.6441 |
| cluster3460 | -5579.3843 | 0.0336 | 0.039 | 1.164 | 0.0152 | 0.1014 | 6.6727 |
| cluster3461 | -3102.8869 | 0.0671 | 0.036 | 0.538 | 0.0616 | 0.1472 | 2.3889 |
| cluster3462 | -824.34932 | 0.0354 | 0.003 | 0.079 | 0.0374 | 0.0059 | 0.1569 |
| cluster3463 | -4770.8973 | 0.0452 | 0.027 | 0.607 | 0.0596 | 0.1100 | 1.8450 |
| cluster3464 | -5608.1104 | 0.0471 | 0.042 | 0.901 | 0.0408 | 0.1584 | 3.8790 |
| cluster3465 | -3951.3266 | 0.0081 | 0.010 | 1.278 | 0.0246 | 0.0495 | 2.0084 |
| cluster3466 | -2643.816 | 0.0111 | 0.010 | 0.937 | 0.0210 | 0.0146 | 0.6986 |

| | | | | | | | |
|---|---|---|---|---|---|---|---|
| cluster3467 | -4058.8742 | 0.0389 | 0.019 | 0.498 | 0.0221 | 0.0441 | 1.9915 |
| cluster3468 | -1849.2856 | 0.0064 | 0.005 | 0.741 | 0.0127 | 0.0490 | 3.8630 |
| cluster3469 | -2105.8584 | 0.0218 | 0.013 | 0.603 | 0.0246 | 0.0400 | 1.6264 |
| cluster3470 | -3880.8646 | 0.0135 | 0.007 | 0.525 | 0.0019 | 0.0408 | 21.2633 |
| cluster3471 | -4173.2584 | 0.0443 | 0.026 | 0.577 | 0.0391 | 0.0789 | 2.0156 |
| cluster3472 | -3719.2213 | 0.0143 | 0.008 | 0.526 | 0.0328 | 0.0256 | 0.7813 |
| cluster3473 | -3470.6172 | 0.0287 | 0.019 | 0.676 | 0.0216 | 0.0205 | 0.9515 |
| cluster3474 | -5366.9017 | 0.0310 | 0.045 | 1.443 | 0.0311 | 0.1235 | 3.9779 |
| cluster3475 | -2339.5718 | 0.0136 | 0.009 | 0.700 | 0.0039 | 0.0332 | 8.4517 |
| cluster3476 | -5111.5637 | 0.0309 | 0.043 | 1.390 | 0.0240 | 0.1315 | 5.4769 |
| cluster3477 | -3273.3195 | 0.0097 | 0.002 | 0.216 | 0.0332 | 0.0125 | 0.3770 |
| cluster3478 | -3580.4135 | 0.0168 | 0.010 | 0.599 | 0.0309 | 0.0219 | 0.7100 |
| cluster3479 | -2912.3602 | 0.0557 | 0.055 | 0.989 | 0.0034 | 0.1493 | 44.0040 |
| cluster3480 | -4508.3233 | 0.0361 | 0.025 | 0.701 | 0.0254 | 0.0716 | 2.8196 |
| cluster3481 | -1072.0431 | 0.0043 | 0.030 | 7.072 | 0.0175 | 0.0061 | 0.3464 |
| cluster3482 | -1956.9604 | 0.1331 | 0.020 | 0.151 | 0.0252 | 0.0169 | 0.6706 |
| cluster3483 | -3660.0009 | 0.0115 | 0.006 | 0.503 | 0.0082 | 0.0201 | 2.4536 |
| cluster3484 | -2769.1166 | 0.0328 | 0.023 | 0.708 | 0.0237 | 0.0671 | 2.8330 |
| cluster3485 | -3492.3455 | 0.0517 | 0.017 | 0.335 | 0.0245 | 0.0884 | 3.6134 |
| cluster3486 | -3174.966 | 0.0526 | 0.016 | 0.307 | 0.0438 | 0.0222 | 0.5075 |
| cluster3487 | -2348.0344 | 0.0262 | 0.013 | 0.503 | 0.0193 | 0.0403 | 2.0931 |
| cluster3488 | -1483.1405 | 0.0243 | 0.007 | 0.302 | 0.0328 | 0.0445 | 1.3552 |
| cluster3489 | -4058.9314 | 0.0497 | 0.038 | 0.770 | 0.0145 | 0.0937 | 6.4593 |
| cluster3490 | -3433.1783 | 0.0216 | 0.021 | 0.954 | 0.0219 | 0.0830 | 3.7903 |
| cluster3491 | -3687.1989 | 0.0319 | 0.016 | 0.491 | 0.0308 | 0.0644 | 2.0923 |
| cluster3492 | -3371.7446 | 0.0345 | 0.028 | 0.820 | 0.0520 | 0.1112 | 2.1415 |
| cluster3493 | -4432.1848 | 0.0304 | 0.013 | 0.439 | 0.0445 | 0.0711 | 1.5988 |
| cluster3494 | -2865.4564 | 0.0142 | 0.022 | 1.572 | 0.0310 | 0.0474 | 1.5275 |
| cluster3495 | -2842.0907 | 0.0334 | 0.015 | 0.459 | 0.0208 | 0.0461 | 2.2154 |
| cluster3496 | -3230.6337 | 0.0192 | 0.009 | 0.485 | 0.0317 | 0.0254 | 0.8020 |
| cluster3497 | -1194.5059 | 0.0166 | 0.002 | 0.137 | 0.0067 | 0.0021 | 0.3116 |
| cluster3498 | -4133.0284 | 0.0536 | 0.057 | 1.065 | 0.0324 | 0.1397 | 4.3145 |
| cluster3499 | -3721.0084 | 0.1180 | 0.068 | 0.577 | 0.0474 | 0.1932 | 4.0803 |
| cluster3500 | -3414.4982 | 0.0271 | 0.015 | 0.569 | 0.0034 | 0.0430 | 12.8286 |
| cluster3501 | -2359.4363 | 0.0247 | 0.015 | 0.591 | 0.0333 | 0.0347 | 1.0414 |
| cluster3502 | -2169.4982 | 0.0143 | 0.010 | 0.734 | 0.0150 | 0.0362 | 2.4057 |
| cluster3503 | -1952.3445 | 0.0466 | 0.022 | 0.476 | 0.0448 | 0.1615 | 3.6028 |
| cluster3504 | -1141.0795 | 0.0093 | 0.010 | 1.111 | 0.0019 | 0.0238 | 12.5498 |
| cluster3505 | -2414.1356 | 0.0675 | 0.061 | 0.905 | 0.0178 | 0.1156 | 6.5016 |
| cluster3506 | -2549.214 | 0.0206 | 0.014 | 0.698 | 0.0318 | 0.0620 | 1.9468 |
| cluster3507 | -5009.6776 | 0.0639 | 0.042 | 0.660 | 0.0091 | 0.0915 | 10.0657 |
| cluster3508 | -5631.5307 | 0.0328 | 0.053 | 1.604 | 0.0151 | 0.1513 | 10.0073 |
| cluster3509 | -2394.9208 | 0.0001 | 0.000 | 0.158 | 0.0736 | 0.1125 | 1.5291 |
| cluster3510 | -843.81358 | 0.0940 | 0.036 | 0.379 | 0.1554 | 0.0686 | 0.4415 |

| | | | | | | | |
|---|---|---|---|---|---|---|---|
| cluster3511 | -1018.3402 | 0.0424 | 0.015 | 0.359 | 0.0214 | 0.0402 | 1.8810 |
| cluster3512 | -2355.65 | 0.0259 | 0.008 | 0.300 | 0.0443 | 0.0390 | 0.8809 |
| cluster3513 | -959.35411 | 0.0751 | 0.003 | 0.037 | 0.0686 | 0.0178 | 0.2602 |
| cluster3514 | -5157.4539 | 0.0333 | 0.020 | 0.601 | 0.0536 | 0.0390 | 0.7268 |
| cluster3515 | -2597.1997 | 0.0065 | 0.003 | 0.536 | 0.0298 | 0.0398 | 1.3384 |
| cluster3516 | -4130.6185 | 0.0272 | 0.017 | 0.644 | 0.0276 | 0.0491 | 1.7769 |
| cluster3517 | -2409.8435 | 0.0012 | 0.000 | 0.226 | 0.0035 | 0.0044 | 1.2638 |
| cluster3518 | -4784.6691 | 0.0749 | 0.047 | 0.621 | 0.0799 | 0.1737 | 2.1743 |
| cluster3519 | -2127.1954 | 0.0424 | 0.024 | 0.564 | 0.0395 | 0.0717 | 1.8146 |
| cluster3520 | -1193.3228 | 0.1283 | 0.030 | 0.232 | 0.2007 | 0.1912 | 0.9528 |
| cluster3521 | -4223.6211 | 0.0410 | 0.038 | 0.933 | 0.0283 | 0.0714 | 2.5248 |
| cluster3522 | -4752.3689 | 0.0360 | 0.031 | 0.871 | 0.0160 | 0.1176 | 7.3719 |
| cluster3523 | -1654.8987 | 0.0373 | 0.027 | 0.723 | 0.0124 | 0.0537 | 4.3408 |
| cluster3524 | -1961.1808 | 0.0192 | 0.004 | 0.220 | 0.0055 | 0.0050 | 0.9002 |
| cluster3525 | -2816.1263 | 0.0388 | 0.012 | 0.305 | 0.0572 | 0.0675 | 1.1819 |
| cluster3526 | -2998.6794 | 0.0027 | 0.002 | 0.732 | 0.0012 | 0.0095 | 7.6927 |
| cluster3527 | -4936.7492 | 0.0505 | 0.053 | 1.046 | 0.0328 | 0.1374 | 4.1953 |
| cluster3528 | -1964.5744 | 0.0152 | 0.006 | 0.416 | 0.0003 | 0.0003 | 0.8069 |
| cluster3529 | -3741.5859 | 0.0339 | 0.027 | 0.807 | 0.0184 | 0.1124 | 6.1186 |
| cluster3530 | -1951.0989 | 0.0233 | 0.015 | 0.629 | 0.0579 | 0.1419 | 2.4512 |
| cluster3531 | -2920.0081 | 0.0540 | 0.022 | 0.406 | 0.0421 | 0.0826 | 1.9619 |
| cluster3532 | -5557.1475 | 0.0595 | 0.052 | 0.869 | 0.0839 | 0.0738 | 0.8796 |
| cluster3533 | -2652.3524 | 0.0190 | 0.007 | 0.390 | 0.0469 | 0.0215 | 0.4582 |
| cluster3534 | -2559.8799 | 0.0577 | 0.012 | 0.206 | 0.0849 | 0.0346 | 0.4073 |
| cluster3535 | -3289.8067 | 0.0224 | 0.019 | 0.854 | 0.0167 | 0.1422 | 8.5122 |
| cluster3536 | -3557.0334 | 0.0347 | 0.020 | 0.565 | 0.0437 | 0.0301 | 0.6886 |
| cluster3537 | -3665.024 | 0.0057 | 0.003 | 0.599 | 0.0087 | 0.0248 | 2.8609 |
| cluster3538 | -1386.7955 | 0.0462 | 0.015 | 0.316 | 0.0108 | 0.0040 | 0.3717 |
| cluster3539 | -2945.5826 | 0.0366 | 0.042 | 1.152 | 0.0451 | 0.2470 | 5.4772 |
| cluster3540 | -5363.2708 | 0.0979 | 0.087 | 0.890 | 0.0692 | 0.2116 | 3.0586 |
| cluster3541 | -3877.2059 | 0.0252 | 0.014 | 0.565 | 0.0355 | 0.0320 | 0.9024 |
| cluster3542 | -5426.0916 | 0.0653 | 0.063 | 0.966 | 0.0375 | 0.2160 | 5.7542 |
| cluster3543 | -4247.6302 | 0.0360 | 0.036 | 1.000 | 0.0302 | 0.1488 | 4.9228 |
| cluster3544 | -1088.4009 | 0.0176 | 0.003 | 0.184 | 0.0090 | 0.0087 | 0.9653 |
| cluster3545 | -5146.6385 | 0.0487 | 0.042 | 0.869 | 0.0364 | 0.1186 | 3.2555 |
| cluster3546 | -2679.5261 | 0.0247 | 0.021 | 0.846 | 0.0168 | 0.0913 | 5.4447 |
| cluster3547 | -1116.238 | 0.0111 | 0.002 | 0.221 | 0.0001 | 0.0000 | 0.2575 |
| cluster3548 | -1303.4172 | 0.0840 | 0.011 | 0.133 | 0.0062 | 0.0020 | 0.3226 |
| cluster3549 | -3229.9525 | 0.0311 | 0.018 | 0.567 | 0.0729 | 0.0515 | 0.7066 |
| cluster3550 | -1224.0675 | 0.0097 | 0.001 | 0.144 | 0.0068 | 0.0037 | 0.5467 |
| cluster3551 | -2797.9413 | 0.0551 | 0.016 | 0.282 | 0.0298 | 0.0119 | 0.3981 |
| cluster3552 | -2386.1274 | 0.0231 | 0.022 | 0.946 | 0.0228 | 0.0582 | 2.5508 |
| cluster3553 | -2405.522 | 0.0317 | 0.016 | 0.503 | 0.0351 | 0.0444 | 1.2645 |
| cluster3554 | -3821.9722 | 0.0203 | 0.007 | 0.347 | 0.0364 | 0.0503 | 1.3812 |

| | | | | | | | |
|---|---|---|---|---|---|---|---|
| cluster3555 | -1632.9835 | 0.0244 | 0.021 | 0.858 | 0.0134 | 0.0463 | 3.4618 |
| cluster3556 | -4206.0991 | 0.0257 | 0.021 | 0.803 | 0.0239 | 0.0732 | 3.0590 |
| cluster3557 | -1368.2372 | 0.0641 | 0.081 | 1.262 | 0.0422 | 0.2713 | 6.4236 |
| cluster3558 | -4777.1633 | 0.0652 | 0.051 | 0.779 | 0.0707 | 0.1213 | 1.7153 |
| cluster3559 | -2457.9208 | 0.0167 | 0.008 | 0.480 | 0.0226 | 0.0618 | 2.7356 |
| cluster3560 | -5096.7676 | 0.0594 | 0.054 | 0.916 | 0.0571 | 0.1573 | 2.7540 |
| cluster3561 | -1801.3376 | 0.0387 | 0.009 | 0.229 | 0.0224 | 0.0193 | 0.8619 |
| cluster3562 | -4278.3788 | 0.0388 | 0.029 | 0.738 | 0.0357 | 0.0834 | 2.3380 |
| cluster3563 | -1980.8879 | 0.0110 | 0.006 | 0.568 | 0.0287 | 0.0232 | 0.8100 |
| cluster3564 | -5021.4314 | 0.0627 | 0.032 | 0.509 | 0.1144 | 0.0532 | 0.4651 |
| cluster3565 | -3350.205 | 0.0397 | 0.030 | 0.763 | 0.0204 | 0.0986 | 4.8421 |
| cluster3566 | -1886.8601 | 0.0556 | 0.024 | 0.426 | 0.0118 | 0.0599 | 5.0813 |
| cluster3567 | -2403.8508 | 0.0642 | 0.029 | 0.456 | 0.0757 | 0.1476 | 1.9512 |
| cluster3568 | -3165.5625 | 0.0305 | 0.025 | 0.808 | 0.0031 | 0.0925 | 30.2034 |
| cluster3569 | -3549.9926 | 0.0424 | 0.036 | 0.848 | 0.0055 | 0.1465 | 26.8734 |
| cluster3570 | -2397.3121 | 0.0217 | 0.026 | 1.184 | 0.0011 | 0.1178 | 107.5724 |
| cluster3571 | -2036.5825 | 0.0143 | 0.020 | 1.393 | 0.0310 | 0.0192 | 0.6201 |
| cluster3572 | -3453.9086 | 0.0511 | 0.034 | 0.666 | 0.0517 | 0.1311 | 2.5375 |
| cluster3573 | -3238.5145 | 0.0153 | 0.012 | 0.815 | 0.0020 | 0.0415 | 20.9032 |
| cluster3574 | -5131.8053 | 0.0770 | 0.062 | 0.800 | 0.0759 | 0.1416 | 1.8653 |
| cluster3575 | -3830.9531 | 0.0379 | 0.026 | 0.684 | 0.0450 | 0.0892 | 1.9842 |
| cluster3576 | -3271.3794 | 0.0424 | 0.027 | 0.649 | 0.0085 | 0.0920 | 10.8331 |
| cluster3577 | -2729.2542 | 0.0336 | 0.015 | 0.433 | 0.0711 | 0.0361 | 0.5082 |
| cluster3578 | -3389.9651 | 0.0369 | 0.021 | 0.567 | 0.0085 | 0.0669 | 7.8983 |
| cluster3579 | -2922.2151 | 0.0304 | 0.030 | 0.973 | 0.0129 | 0.0207 | 1.6110 |
| cluster3580 | -2417.8752 | 0.0100 | 0.011 | 1.082 | 0.0120 | 0.0475 | 3.9435 |
| cluster3581 | -1247.0514 | 0.0081 | 0.004 | 0.476 | 0.0065 | 0.0425 | 6.5601 |
| cluster3582 | -1106.9685 | 0.0305 | 0.005 | 0.149 | 0.0074 | 0.0023 | 0.3069 |
| cluster3583 | -2533.4362 | 0.0001 | 0.000 | 0.422 | 0.0001 | 0.0003 | 2.8730 |
| cluster3584 | -4869.8223 | 0.0313 | 0.038 | 1.231 | 0.0282 | 0.1452 | 5.1548 |
| cluster3585 | -2094.4304 | 0.0614 | 0.018 | 0.297 | 0.0673 | 0.0325 | 0.4823 |
| cluster3586 | -5223.6249 | 0.0771 | 0.069 | 0.891 | 0.0467 | 0.1233 | 2.6413 |
| cluster3587 | -3467.2967 | 0.0601 | 0.066 | 1.099 | 0.0109 | 0.1484 | 13.5926 |
| cluster3588 | -3361.197 | 0.0286 | 0.008 | 0.289 | 0.0249 | 0.0175 | 0.7023 |
| cluster3589 | -3015.8888 | 0.0517 | 0.028 | 0.545 | 0.0275 | 0.0583 | 2.1199 |
| cluster3590 | -4181.9105 | 0.0623 | 0.064 | 1.020 | 0.0430 | 0.2322 | 5.3937 |
| cluster3591 | -2874.4182 | 0.0120 | 0.005 | 0.408 | 0.0120 | 0.0080 | 0.6649 |
| cluster3592 | -4706.0608 | 0.0380 | 0.035 | 0.927 | 0.0429 | 0.1287 | 3.0011 |
| cluster3593 | -3418.7874 | 0.0502 | 0.050 | 1.001 | 0.0191 | 0.1160 | 6.0597 |
| cluster3594 | -3168.2777 | 0.0773 | 0.025 | 0.327 | 0.0020 | 0.0860 | 43.3520 |
| cluster3595 | -2123.6395 | 0.0118 | 0.005 | 0.420 | 0.0127 | 0.0091 | 0.7119 |
| cluster3596 | -2316.2681 | 0.0328 | 0.016 | 0.476 | 0.0339 | 0.0645 | 1.9051 |
| cluster3597 | -4192.6975 | 0.0336 | 0.020 | 0.609 | 0.0188 | 0.0268 | 1.4275 |
| cluster3598 | -4253.8313 | 0.0228 | 0.089 | 3.890 | 0.0239 | 0.1292 | 5.4099 |

| | | | | | | | |
|---|---|---|---|---|---|---|---|
| cluster3599 | -2252.9745 | 0.0286 | 0.018 | 0.616 | 0.0167 | 0.0437 | 2.6261 |
| cluster3600 | -2861.1244 | 0.0234 | 0.008 | 0.326 | 0.0283 | 0.0444 | 1.5719 |
| cluster3601 | -3284.4637 | 0.0059 | 0.007 | 1.147 | 0.0022 | 0.0303 | 14.1255 |
| cluster3602 | -3064.7212 | 0.0021 | 0.001 | 0.379 | 0.0070 | 0.0038 | 0.5466 |
| cluster3603 | -4224.6278 | 0.0531 | 0.032 | 0.608 | 0.0509 | 0.1207 | 2.3718 |
| cluster3604 | -1058.107 | 0.0537 | 0.004 | 0.078 | 0.0793 | 0.0157 | 0.1975 |
| cluster3605 | -3057.8472 | 0.0373 | 0.024 | 0.655 | 0.0405 | 0.0729 | 1.7991 |
| cluster3606 | -4323.6354 | 0.0277 | 0.024 | 0.852 | 0.0282 | 0.0958 | 3.3961 |
| cluster3607 | -2594.0588 | 0.0392 | 0.029 | 0.746 | 0.0037 | 0.0901 | 24.6846 |
| cluster3608 | -2887.2248 | 0.0123 | 0.007 | 0.562 | 0.0065 | 0.0072 | 1.1125 |
| cluster3609 | -4169.0239 | 0.0383 | 0.037 | 0.972 | 0.0027 | 0.1045 | 39.4857 |
| cluster3610 | -1395.5537 | 0.0081 | 0.024 | 3.035 | 0.0074 | 0.0104 | 1.4064 |
| cluster3611 | -2340.84 | 0.0268 | 0.034 | 1.248 | 0.0229 | 0.1834 | 8.0102 |
| cluster3612 | -2831.6127 | 0.0347 | 0.040 | 1.167 | 0.0319 | 0.1197 | 3.7596 |
| cluster3613 | -4058.8683 | 0.0171 | 0.011 | 0.649 | 0.0240 | 0.0154 | 0.6409 |
| cluster3614 | -4306.9621 | 0.0227 | 0.034 | 1.508 | 0.0376 | 0.1543 | 4.0999 |
| cluster3615 | -1662.679 | 0.0414 | 0.008 | 0.194 | 0.0746 | 0.0157 | 0.2104 |
| cluster3616 | -770.98944 | 0.0211 | 0.007 | 0.350 | 0.0007 | 0.0222 | 30.9723 |
| cluster3617 | -1921.7935 | 0.0550 | 0.039 | 0.703 | 0.0009 | 0.0579 | 63.9626 |
| cluster3618 | -1899.3282 | 0.1675 | 0.068 | 0.405 | 0.1946 | 0.2228 | 1.1446 |
| cluster3619 | -2552.5885 | 0.0044 | 0.002 | 0.493 | 0.0050 | 0.0140 | 2.8085 |
| cluster3620 | -3604.7684 | 0.0481 | 0.024 | 0.506 | 0.0627 | 0.0914 | 1.4593 |
| cluster3621 | -4765.4069 | 0.0345 | 0.038 | 1.099 | 0.0295 | 0.1337 | 4.5377 |
| cluster3622 | -2722.1045 | 0.0166 | 0.006 | 0.358 | 0.0061 | 0.0057 | 0.9379 |
| cluster3623 | -2271.6868 | 0.0085 | 0.010 | 1.188 | 0.0105 | 0.0839 | 8.0215 |
| cluster3624 | -2909.4943 | 0.0143 | 0.010 | 0.674 | 0.0055 | 0.0910 | 16.6096 |
| cluster3625 | -3307.0908 | 0.0058 | 0.004 | 0.752 | 0.0156 | 0.0411 | 2.6320 |
| cluster3626 | -1982.12 | 0.0252 | 0.016 | 0.653 | 0.0537 | 0.0297 | 0.5543 |
| cluster3627 | -3000.8801 | 0.0065 | 0.004 | 0.654 | 0.0084 | 0.0181 | 2.1620 |
| cluster3628 | -3023.0613 | 0.0282 | 0.032 | 1.120 | 0.0241 | 0.1145 | 4.7616 |
| cluster3629 | -4165.4613 | 0.0120 | 0.017 | 1.432 | 0.0113 | 0.0637 | 5.6313 |
| cluster3630 | -1210.1774 | 0.0132 | 0.004 | 0.298 | 0.0205 | 0.0079 | 0.3834 |
| cluster3631 | -3594.8673 | 0.0141 | 0.019 | 1.383 | 0.0189 | 0.0242 | 1.2791 |
| cluster3632 | -2459.6598 | 0.0290 | 0.039 | 1.360 | 0.0405 | 0.0704 | 1.7407 |
| cluster3633 | -2501.0657 | 0.0055 | 0.004 | 0.649 | 0.0484 | 0.0509 | 1.0508 |
| cluster3634 | -931.87974 | 0.0001 | 0.000 | 0.227 | 0.0348 | 0.0359 | 1.0308 |
| cluster3635 | -1620.9616 | 0.0459 | 0.006 | 0.126 | 0.0341 | 0.0122 | 0.3566 |
| cluster3636 | -2532.3436 | 0.0147 | 0.013 | 0.853 | 0.0132 | 0.0178 | 1.3492 |
| cluster3637 | -1061.2182 | 0.0237 | 0.004 | 0.167 | 0.0056 | 0.0035 | 0.6339 |
| cluster3638 | -2732.6534 | 0.0304 | 0.009 | 0.307 | 0.0343 | 0.0510 | 1.4864 |
| cluster3639 | -1272.5097 | 0.0337 | 0.012 | 0.363 | 0.0404 | 0.0366 | 0.9074 |
| cluster3640 | -4766.2667 | 0.0713 | 0.063 | 0.883 | 0.0884 | 0.2125 | 2.4050 |
| cluster3641 | -1769.0836 | 0.0257 | 0.006 | 0.227 | 0.0289 | 0.0101 | 0.3487 |
| cluster3642 | -3357.7853 | 0.0322 | 0.026 | 0.803 | 0.0300 | 0.1094 | 3.6491 |

| | | | | | | | |
|---|---|---|---|---|---|---|---|
| cluster3643 | -3538.1325 | 0.1052 | 0.054 | 0.512 | 0.0853 | 0.2531 | 2.9677 |
| cluster3644 | -1912.2579 | 0.0352 | 0.008 | 0.224 | 0.0207 | 0.0191 | 0.9224 |
| cluster3645 | -1806.54 | 0.0351 | 0.024 | 0.676 | 0.0145 | 0.0710 | 4.8824 |
| cluster3646 | -2636.2101 | 0.0216 | 0.040 | 1.872 | 0.0212 | 0.0930 | 4.3902 |
| cluster3647 | -3429.7537 | 0.0040 | 0.007 | 1.814 | 0.0066 | 0.0523 | 7.8977 |
| cluster3648 | -1757.2905 | 0.0004 | 0.001 | 1.554 | 999.0000 | 0.0053 | 0.0000 |
| cluster3649 | -1640.5882 | 0.0093 | 0.003 | 0.358 | 0.0106 | 0.0175 | 1.6527 |
| cluster3650 | -1256.4377 | 0.0174 | 0.032 | 1.862 | 0.0014 | 0.1057 | 77.6265 |
| cluster3651 | -3969.7526 | 0.0424 | 0.029 | 0.687 | 0.0731 | 0.0614 | 0.8397 |
| cluster3652 | -3310.6171 | 0.0259 | 0.018 | 0.711 | 0.0215 | 0.0169 | 0.7847 |
| cluster3653 | -1426.8535 | 0.0043 | 0.002 | 0.523 | 0.0025 | 0.0018 | 0.7319 |
| cluster3654 | -2837.3801 | 0.0084 | 0.007 | 0.820 | 0.0132 | 0.0348 | 2.6339 |
| cluster3655 | -2366.6522 | 0.0657 | 0.037 | 0.564 | 0.0681 | 0.1297 | 1.9040 |
| cluster3656 | -3106.9421 | 0.0219 | 0.017 | 0.775 | 0.0461 | 0.0363 | 0.7872 |
| cluster3657 | -3194.449 | 0.0150 | 0.010 | 0.682 | 0.0150 | 0.1163 | 7.7624 |
| cluster3658 | -752.15807 | 0.1037 | 0.024 | 0.235 | 0.0254 | 0.0078 | 0.3058 |
| cluster3659 | -2269.7158 | 0.0375 | 0.014 | 0.381 | 0.0321 | 0.0377 | 1.1725 |
| cluster3660 | -3017.1654 | 0.0211 | 0.031 | 1.472 | 0.0120 | 0.1006 | 8.3711 |
| cluster3661 | -1818.2845 | 0.0001 | 0.000 | 0.081 | 0.0044 | 0.0011 | 0.2465 |
| cluster3662 | -2804.2923 | 0.0360 | 0.028 | 0.776 | 0.0368 | 0.0876 | 2.3768 |
| cluster3663 | -3132.7955 | 0.0430 | 0.027 | 0.637 | 0.0681 | 0.0863 | 1.2673 |
| cluster3664 | -2444.9745 | 0.0517 | 0.054 | 1.052 | 0.0171 | 0.0845 | 4.9311 |
| cluster3665 | -3603.1343 | 0.0195 | 0.020 | 1.004 | 0.0252 | 0.0654 | 2.5936 |
| cluster3666 | -2711.494 | 0.0347 | 0.016 | 0.468 | 0.0355 | 0.0963 | 2.7161 |
| cluster3667 | -1570.4955 | 0.0171 | 0.003 | 0.190 | 0.0258 | 0.0183 | 0.7071 |
| cluster3668 | -2983.1902 | 0.0191 | 0.014 | 0.725 | 0.0270 | 0.0512 | 1.8994 |
| cluster3669 | -2600.9649 | 0.0317 | 0.014 | 0.449 | 0.0490 | 0.0661 | 1.3505 |
| cluster3670 | -4034.4773 | 0.0435 | 0.036 | 0.822 | 0.0579 | 0.0556 | 0.9608 |
| cluster3671 | -3994.1631 | 0.0323 | 0.026 | 0.795 | 0.0113 | 0.0861 | 7.6210 |
| cluster3672 | -2500.7029 | 0.0462 | 0.024 | 0.517 | 0.0403 | 0.1031 | 2.5572 |
| cluster3673 | -4408.8869 | 0.0445 | 0.069 | 1.561 | 0.0197 | 0.2652 | 13.4592 |
| cluster3674 | -2025.399 | 0.0270 | 0.029 | 1.065 | 0.0317 | 0.0904 | 2.8511 |
| cluster3675 | -660.30852 | 0.0001 | 0.000 | 0.113 | 0.0001 | 0.0000 | 0.2602 |
| cluster3676 | -3525.9452 | 0.0578 | 0.040 | 0.684 | 0.0411 | 0.1095 | 2.6656 |
| cluster3677 | -807.60405 | 0.0291 | 0.008 | 0.264 | 0.0009 | 0.0172 | 18.6889 |
| cluster3678 | -3306.9773 | 0.0319 | 0.021 | 0.644 | 0.0437 | 0.0678 | 1.5531 |
| cluster3679 | -944.40086 | 0.0097 | 0.003 | 0.352 | 0.0044 | 0.0028 | 0.6315 |
| cluster3680 | -2313.6816 | 0.0082 | 0.009 | 1.074 | 0.0164 | 0.0700 | 4.2643 |
| cluster3681 | -2959.1047 | 0.0201 | 0.022 | 1.094 | 0.0144 | 0.0694 | 4.8156 |
| cluster3682 | -4364.7044 | 0.0373 | 0.033 | 0.884 | 0.0237 | 0.1142 | 4.8218 |
| cluster3683 | -2805.2249 | 0.1087 | 0.097 | 0.893 | 0.0874 | 0.1061 | 1.2138 |
| cluster3684 | -2992.6344 | 0.0321 | 0.023 | 0.713 | 0.0434 | 0.0856 | 1.9737 |
| cluster3685 | -3671.8025 | 0.0991 | 0.076 | 0.769 | 0.0487 | 0.2080 | 4.2714 |
| cluster3686 | -1627.4987 | 0.0249 | 0.050 | 2.026 | 0.0624 | 0.1849 | 2.9638 |

| | | | | | | | |
|---|---|---|---|---|---|---|---|
| cluster3687 | -2875.9708 | 0.0947 | 0.073 | 0.769 | 0.0452 | 0.3753 | 8.2995 |
| cluster3688 | -2048.2275 | 0.0104 | 0.007 | 0.647 | 0.0012 | 0.0474 | 39.9787 |
| cluster3689 | -2640.9793 | 0.0344 | 0.016 | 0.462 | 0.0600 | 0.0452 | 0.7533 |
| cluster3690 | -1454.12 | 0.0204 | 0.016 | 0.766 | 0.0443 | 0.0243 | 0.5494 |
| cluster3691 | -669.56422 | 0.0063 | 0.004 | 0.633 | 0.0001 | 0.0001 | 1.0473 |
| cluster3692 | -1435.3581 | 0.0113 | 0.009 | 0.839 | 0.0104 | 0.0251 | 2.4280 |
| cluster3693 | -2461.3873 | 0.0117 | 0.013 | 1.114 | 0.0023 | 0.0404 | 17.4185 |
| cluster3694 | -2092.0856 | 0.0324 | 0.010 | 0.295 | 0.0236 | 0.0118 | 0.5012 |
| cluster3695 | -1736.5982 | 0.0218 | 0.014 | 0.648 | 0.0319 | 0.0611 | 1.9123 |
| cluster3696 | -817.46452 | 0.0124 | 0.016 | 1.276 | 0.0026 | 0.0909 | 35.3524 |
| cluster3697 | -3582.0314 | 0.0635 | 0.035 | 0.549 | 0.1228 | 0.0661 | 0.5386 |
| cluster3698 | -1099.5724 | 0.0001 | 0.000 | 0.254 | 0.0001 | 0.0000 | 0.3959 |
| cluster3699 | -4438.4776 | 0.0541 | 0.050 | 0.917 | 0.0267 | 0.1293 | 4.8418 |
| cluster3700 | -2819.0365 | 0.0312 | 0.020 | 0.636 | 0.0350 | 0.0734 | 2.0967 |
| cluster3701 | -1126.4168 | 0.0001 | 0.000 | 0.434 | 0.0115 | 0.0023 | 0.1974 |
| cluster3702 | -3864.1159 | 0.0282 | 0.034 | 1.192 | 0.0185 | 0.1124 | 6.0754 |
| cluster3703 | -1512.5315 | 0.0184 | 0.005 | 0.266 | 0.0128 | 0.0093 | 0.7261 |
| cluster3704 | -2226.6249 | 0.0036 | 0.005 | 1.520 | 0.0164 | 0.0359 | 2.1933 |
| cluster3705 | -1944.8294 | 0.0151 | 0.008 | 0.542 | 0.0087 | 0.0082 | 0.9464 |
| cluster3706 | -2093.7998 | 0.0168 | 0.006 | 0.373 | 0.0277 | 0.0403 | 1.4588 |
| cluster3707 | -1920.8903 | 0.0460 | 0.045 | 0.971 | 0.0269 | 0.1186 | 4.4144 |
| cluster3708 | -2634.3929 | 0.0526 | 0.020 | 0.374 | 0.0120 | 0.0958 | 7.9941 |
| cluster3709 | -1578.8049 | 0.0557 | 0.009 | 0.155 | 0.1098 | 0.0638 | 0.5812 |
| cluster3710 | -1924.8089 | 0.0150 | 0.015 | 0.973 | 0.0124 | 0.0613 | 4.9315 |
| cluster3711 | -1436.8783 | 0.0483 | 0.006 | 0.123 | 0.0143 | 0.0098 | 0.6865 |
| cluster3712 | -3402.5092 | 0.0283 | 0.014 | 0.505 | 0.0232 | 0.0293 | 1.2648 |
| cluster3713 | -4029.6364 | 0.0512 | 0.048 | 0.947 | 0.0652 | 0.1859 | 2.8526 |
| cluster3714 | -1775.8601 | 0.0463 | 0.016 | 0.348 | 0.0459 | 0.0208 | 0.4522 |
| cluster3715 | -1286.7638 | 0.0121 | 0.033 | 2.705 | 0.0181 | 0.0825 | 4.5513 |
| cluster3716 | -3658.0177 | 0.0477 | 0.042 | 0.879 | 0.0411 | 0.1089 | 2.6513 |
| cluster3717 | -1403.9483 | 0.0148 | 0.003 | 0.206 | 0.0130 | 0.0061 | 0.4691 |
| cluster3718 | -2990.2996 | 0.0204 | 0.017 | 0.839 | 0.0340 | 0.0911 | 2.6796 |
| cluster3719 | -3294.621 | 0.0478 | 0.023 | 0.474 | 0.0357 | 0.0767 | 2.1488 |
| cluster3720 | -2758.1316 | 0.0309 | 0.011 | 0.367 | 0.0196 | 0.0462 | 2.3538 |
| cluster3721 | -1097.8982 | 0.0036 | 0.000 | 0.123 | 0.0083 | 0.0022 | 0.2635 |
| cluster3722 | -1224.5824 | 0.0398 | 0.070 | 1.752 | 0.0398 | 0.1781 | 4.4700 |
| cluster3723 | -989.94266 | 0.0001 | 0.000 | 0.316 | 0.0001 | 0.0001 | 0.7858 |
| cluster3724 | -3083.5642 | 0.0328 | 0.026 | 0.786 | 0.0360 | 0.0911 | 2.5303 |
| cluster3725 | -1837.6449 | 0.0129 | 0.010 | 0.782 | 0.0020 | 0.0385 | 19.4163 |
| cluster3726 | -1936.0836 | 0.0542 | 0.022 | 0.411 | 0.0381 | 0.0467 | 1.2275 |
| cluster3727 | -2469.0085 | 0.0531 | 0.038 | 0.709 | 0.0601 | 0.1986 | 3.3053 |
| cluster3728 | -2156.5806 | 0.0186 | 0.009 | 0.489 | 0.0272 | 0.0338 | 1.2424 |
| cluster3729 | -2945.9274 | 0.0154 | 0.011 | 0.688 | 0.0073 | 0.0480 | 6.5675 |
| cluster3730 | -1780.3797 | 0.0968 | 0.022 | 0.225 | 0.0442 | 0.0199 | 0.4492 |

| | | | | | | | |
|---|---|---|---|---|---|---|---|
| cluster3731 | -2865.403 | 0.0084 | 0.005 | 0.544 | 0.0107 | 0.0094 | 0.8785 |
| cluster3732 | -2822.909 | 0.0220 | 0.011 | 0.499 | 0.0243 | 0.0450 | 1.8500 |
| cluster3733 | -3070.6204 | 0.0911 | 0.033 | 0.365 | 0.0503 | 0.0367 | 0.7287 |
| cluster3734 | -3043.2604 | 0.0414 | 0.029 | 0.690 | 0.0504 | 0.1002 | 1.9896 |
| cluster3735 | -3249.3767 | 0.0740 | 0.032 | 0.434 | 0.1122 | 0.0578 | 0.5154 |
| cluster3736 | -3537.6725 | 0.0423 | 0.019 | 0.455 | 0.0550 | 0.0575 | 1.0454 |
| cluster3737 | -1208.4594 | 0.0053 | 0.001 | 0.185 | 0.0001 | 0.0001 | 0.6134 |
| cluster3738 | -3256.843 | 0.0065 | 0.004 | 0.683 | 0.0183 | 0.0173 | 0.9452 |
| cluster3739 | -1156.2555 | 0.0001 | 0.000 | 0.136 | 0.0001 | 0.0001 | 0.7290 |
| cluster3740 | -3450.4131 | 0.0296 | 0.023 | 0.787 | 0.0314 | 0.0680 | 2.1698 |
| cluster3741 | -1311.9743 | 0.0120 | 0.004 | 0.298 | 0.0238 | 0.0212 | 0.8912 |
| cluster3742 | -1615.2077 | 0.0508 | 0.034 | 0.673 | 0.0786 | 0.1060 | 1.3488 |
| cluster3743 | -2979.62 | 0.0264 | 0.017 | 0.640 | 0.0167 | 0.0621 | 3.7302 |
| cluster3744 | -1353.7039 | 0.1338 | 0.037 | 0.275 | 0.0507 | 0.0934 | 1.8432 |
| cluster3745 | -2585.3007 | 0.0245 | 0.024 | 0.961 | 0.0274 | 0.0624 | 2.2819 |
| cluster3746 | -3783.1617 | 0.0437 | 0.037 | 0.844 | 0.0189 | 0.0268 | 1.4203 |
| cluster3747 | -1120.5813 | 0.0223 | 0.012 | 0.541 | 0.0388 | 0.0453 | 1.1692 |
| cluster3748 | -2166.6608 | 0.0146 | 0.007 | 0.487 | 0.0124 | 0.0076 | 0.6102 |
| cluster3749 | -3071.2923 | 0.0164 | 0.013 | 0.806 | 0.0224 | 0.0553 | 2.4713 |
| cluster3750 | -1355.3836 | 0.0253 | 0.029 | 1.142 | 0.0201 | 0.1132 | 5.6241 |
| cluster3751 | -3011.1604 | 0.0034 | 0.002 | 0.573 | 0.0064 | 0.0139 | 2.1664 |
| cluster3752 | -2663.0527 | 0.0090 | 0.007 | 0.777 | 0.0050 | 0.0381 | 7.5868 |
| cluster3753 | -1023.151 | 0.0172 | 0.002 | 0.124 | 0.0414 | 0.0021 | 0.0516 |
| cluster3754 | -881.97724 | 0.0001 | 0.000 | 0.101 | 0.0001 | 0.0000 | 0.4120 |
| cluster3755 | -2499.7961 | 0.0160 | 0.012 | 0.727 | 0.0148 | 0.0171 | 1.1566 |
| cluster3756 | -1689.0347 | 0.0312 | 0.020 | 0.645 | 0.0494 | 0.0713 | 1.4417 |
| cluster3757 | -2091.0436 | 0.0500 | 0.015 | 0.301 | 0.0384 | 0.0386 | 1.0059 |
| cluster3758 | -1511.7073 | 0.0303 | 0.011 | 0.375 | 0.0246 | 0.0298 | 1.2119 |
| cluster3759 | -3066.0287 | 0.0727 | 0.040 | 0.552 | 0.0527 | 0.0319 | 0.6064 |
| cluster3760 | -2599.5028 | 0.0204 | 0.017 | 0.816 | 0.0424 | 0.0524 | 1.2363 |
| cluster3761 | -766.76933 | 0.0049 | 0.008 | 1.592 | 0.0255 | 0.0879 | 3.4433 |
| cluster3762 | -1967.8506 | 0.0180 | 0.005 | 0.267 | 0.0129 | 0.0052 | 0.4040 |
| cluster3763 | -2005.1958 | 0.0605 | 0.015 | 0.244 | 0.0400 | 0.0141 | 0.3535 |
| cluster3764 | -1629.4335 | 0.0327 | 0.031 | 0.932 | 0.0185 | 0.0905 | 4.8807 |
| cluster3765 | -2665.82 | 0.0134 | 0.007 | 0.545 | 0.0512 | 0.0506 | 0.9884 |
| cluster3766 | -1791.356 | 0.0156 | 0.007 | 0.444 | 0.0017 | 0.0329 | 19.9203 |
| cluster3767 | -862.49887 | 0.0213 | 0.023 | 1.096 | 0.0324 | 0.0766 | 2.3602 |
| cluster3768 | -2217.5326 | 0.0429 | 0.048 | 1.128 | 0.0013 | 0.0942 | 70.7672 |
| cluster3769 | -1765.311 | 0.0068 | 0.001 | 0.215 | 0.0076 | 0.0045 | 0.5851 |
| cluster3770 | -1775.1381 | 0.0537 | 0.025 | 0.460 | 0.0509 | 0.0975 | 1.9152 |
| cluster3771 | -2137.4054 | 0.0610 | 0.024 | 0.389 | 0.1146 | 0.2208 | 1.9276 |
| cluster3772 | -723.09975 | 0.0001 | 0.000 | 0.211 | 0.0483 | 0.0897 | 1.8562 |
| cluster3773 | -1114.8639 | 0.0823 | 0.004 | 0.044 | 0.3195 | 0.1804 | 0.5647 |
| cluster3774 | -1408.6168 | 0.0090 | 0.001 | 0.145 | 0.0196 | 0.0087 | 0.4439 |

| | | | | | | | |
|---|---|---|---|---|---|---|---|
| cluster3775 | -903.62389 | 0.0201 | 0.011 | 0.571 | 0.0022 | 0.0522 | 23.8124 |
| cluster3776 | -2518.1737 | 0.0728 | 0.026 | 0.362 | 0.1029 | 0.0483 | 0.4692 |
| cluster3777 | -3066.5953 | 0.0561 | 0.046 | 0.816 | 0.0777 | 0.1308 | 1.6842 |
| cluster3778 | -980.35293 | 0.0008 | 0.001 | 0.752 | 0.0032 | 0.0028 | 0.8744 |
| cluster3779 | -2995.126 | 0.0219 | 0.024 | 1.094 | 0.0256 | 0.0614 | 2.4032 |
| cluster3780 | -1685.5793 | 0.0122 | 0.013 | 1.047 | 0.0114 | 0.0597 | 5.2336 |
| cluster3781 | -2685.7455 | 0.0531 | 0.029 | 0.554 | 0.0777 | 0.0635 | 0.8163 |
| cluster3782 | -2289.3063 | 0.0489 | 0.023 | 0.474 | 0.0305 | 0.0381 | 1.2505 |
| cluster3783 | -2108.1914 | 0.0069 | 0.013 | 1.960 | 0.0088 | 0.0874 | 9.9239 |
| cluster3784 | -2950.8152 | 0.0538 | 0.047 | 0.868 | 0.0312 | 0.0361 | 1.1584 |
| cluster3785 | -2683.2441 | 0.0248 | 0.010 | 0.393 | 0.0065 | 0.0108 | 1.6714 |
| cluster3786 | -2267.011 | 0.0087 | 0.026 | 3.032 | 0.0083 | 0.0171 | 2.0743 |
| cluster3787 | -1520.0071 | 0.0355 | 0.019 | 0.542 | 0.0112 | 0.0263 | 2.3500 |
| cluster3788 | -1160.4015 | 0.0001 | 0.000 | 0.024 | 0.0001 | 0.0000 | 0.3159 |
| cluster3789 | -3040.9952 | 0.0106 | 0.003 | 0.315 | 0.0124 | 0.0132 | 1.0667 |
| cluster3790 | -2177.2244 | 0.0136 | 0.012 | 0.855 | 0.0267 | 0.0862 | 3.2307 |
| cluster3791 | -418.31419 | 0.0211 | 0.006 | 0.284 | 0.0001 | 0.0001 | 0.7625 |
| cluster3792 | -1863.3645 | 0.0065 | 0.002 | 0.310 | 0.0055 | 0.0069 | 1.2449 |
| cluster3793 | -1852.7941 | 0.0312 | 0.013 | 0.429 | 0.0189 | 0.0458 | 2.4222 |
| cluster3794 | -2187.1782 | 0.0206 | 0.013 | 0.647 | 0.0064 | 0.0408 | 6.4195 |
| cluster3795 | -1701.6558 | 0.0197 | 0.009 | 0.458 | 0.0185 | 0.0513 | 2.7705 |
| cluster3796 | -1251.9276 | 0.1287 | 0.058 | 0.453 | 0.1229 | 0.1381 | 1.1240 |
| cluster3797 | -3913.0725 | 0.0571 | 0.065 | 1.147 | 0.0427 | 0.3231 | 7.5681 |
| cluster3798 | -3441.8757 | 0.0301 | 0.021 | 0.689 | 0.0086 | 0.0489 | 5.6767 |
| cluster3799 | -2341.3402 | 0.0491 | 0.032 | 0.647 | 0.0366 | 0.0365 | 0.9975 |
| cluster3800 | -429.54977 | 0.0001 | 0.000 | 0.090 | 0.0766 | 0.0046 | 0.0600 |
| cluster3801 | -2690.3441 | 0.2201 | 0.050 | 0.226 | 0.1782 | 0.2229 | 1.2505 |
| cluster3802 | -2648.9273 | 0.0303 | 0.034 | 1.121 | 0.0539 | 0.0740 | 1.3725 |
| cluster3803 | -2239.6734 | 0.0386 | 0.020 | 0.530 | 0.0290 | 0.0527 | 1.8173 |
| cluster3804 | -2635.5876 | 0.0314 | 0.016 | 0.514 | 0.0331 | 0.0699 | 2.1134 |
| cluster3805 | -2655.2673 | 0.0259 | 0.018 | 0.709 | 0.0146 | 0.0562 | 3.8545 |
| cluster3806 | -3566.5706 | 0.0413 | 0.033 | 0.788 | 0.0322 | 0.0769 | 2.3883 |
| cluster3807 | -1441.8582 | 0.0080 | 0.002 | 0.306 | 0.0054 | 0.0018 | 0.3414 |
| cluster3808 | -1825.9664 | 0.0309 | 0.024 | 0.780 | 0.0351 | 0.0901 | 2.5709 |
| cluster3809 | -2359.104 | 0.0109 | 0.005 | 0.464 | 0.0013 | 0.0185 | 14.5716 |
| cluster3810 | -2464.5959 | 0.0097 | 0.008 | 0.876 | 0.0096 | 0.0146 | 1.5210 |
| cluster3811 | -761.00025 | 0.0137 | 0.002 | 0.114 | 0.0381 | 0.0170 | 0.4455 |
| cluster3812 | -2997.2082 | 0.0333 | 0.011 | 0.341 | 0.0032 | 0.0524 | 16.4744 |
| cluster3813 | -3413.696 | 0.0343 | 0.055 | 1.603 | 0.0384 | 0.1724 | 4.4928 |
| cluster3814 | -1609.0304 | 0.0883 | 0.016 | 0.177 | 0.0908 | 0.0263 | 0.2895 |
| cluster3815 | -1708.512 | 0.0198 | 0.018 | 0.887 | 0.0148 | 0.0987 | 6.6790 |
| cluster3816 | -2525.5897 | 0.0300 | 0.030 | 0.993 | 0.0152 | 0.0833 | 5.4914 |
| cluster3817 | -992.05474 | 0.0060 | 0.002 | 0.290 | 0.0074 | 0.0025 | 0.3333 |
| cluster3818 | -1978.8486 | 0.0658 | 0.065 | 0.993 | 0.0484 | 0.0685 | 1.4152 |

| | | | | | | | |
|---|---|---|---|---|---|---|---|
| cluster3819 | -3347.2015 | 0.0279 | 0.019 | 0.689 | 0.0262 | 0.0569 | 2.1698 |
| cluster3820 | -2225.802 | 0.0301 | 0.016 | 0.529 | 0.0294 | 0.0656 | 2.2289 |
| cluster3821 | -2241.9084 | 0.0132 | 0.008 | 0.610 | 0.0330 | 0.0297 | 0.8998 |
| cluster3822 | -3399.8116 | 0.0295 | 0.024 | 0.824 | 0.0247 | 0.0916 | 3.7047 |
| cluster3823 | -3408.7723 | 0.0192 | 0.023 | 1.202 | 0.0227 | 0.0382 | 1.6840 |
| cluster3824 | -2349.759 | 0.0299 | 0.030 | 1.021 | 0.0053 | 0.0741 | 13.9151 |
| cluster3825 | -1388.4368 | 0.0047 | 0.002 | 0.477 | 0.0093 | 0.0158 | 1.7049 |
| cluster3826 | -3154.5024 | 0.0704 | 0.059 | 0.842 | 0.1088 | 0.1978 | 1.8180 |
| cluster3827 | -2403.5221 | 0.0265 | 0.021 | 0.806 | 0.0354 | 0.0414 | 1.1681 |
| cluster3828 | -2365.1011 | 0.0505 | 0.028 | 0.550 | 0.0426 | 0.0715 | 1.6788 |
| cluster3829 | -2254.9158 | 0.0298 | 0.019 | 0.623 | 0.0124 | 0.0668 | 5.3680 |
| cluster3830 | -1625.9479 | 0.0366 | 0.006 | 0.155 | 0.0097 | 0.0119 | 1.2314 |
| cluster3831 | -2347.6497 | 0.0350 | 0.027 | 0.767 | 0.0191 | 0.1288 | 6.7630 |
| cluster3832 | -1768.4101 | 0.0055 | 0.002 | 0.429 | 0.0033 | 0.0064 | 1.9419 |
| cluster3833 | -2130.0166 | 0.0255 | 0.018 | 0.723 | 0.0179 | 0.0330 | 1.8430 |
| cluster3834 | -2057.1838 | 0.0125 | 0.010 | 0.800 | 0.0015 | 0.0635 | 42.5415 |
| cluster3835 | -2118.1865 | 0.0325 | 0.017 | 0.520 | 0.0388 | 0.0660 | 1.7014 |
| cluster3836 | -1895.0229 | 0.0178 | 0.010 | 0.578 | 0.0163 | 0.0306 | 1.8767 |
| cluster3837 | -2638.7365 | 0.0461 | 0.029 | 0.632 | 0.0304 | 0.0663 | 2.1785 |
| cluster3838 | -2308.303 | 0.0493 | 0.026 | 0.527 | 0.0172 | 0.0290 | 1.6862 |
| cluster3839 | -1495.7809 | 0.0252 | 0.017 | 0.687 | 0.0008 | 0.0383 | 50.5479 |
| cluster3840 | -2021.6992 | 0.0056 | 0.071 | 12.701 | 0.0310 | 0.0335 | 1.0823 |
| cluster3841 | -2290.8368 | 0.0878 | 0.063 | 0.723 | 0.0322 | 0.2034 | 6.3120 |
| cluster3842 | -1605.9097 | 0.0051 | 0.001 | 0.184 | 0.0001 | 0.0000 | 0.4153 |
| cluster3843 | -1438.2301 | 0.0051 | 0.002 | 0.376 | 0.0212 | 0.0256 | 1.2067 |
| cluster3844 | -1076.3139 | 0.0781 | 0.017 | 0.220 | 0.0092 | 0.0894 | 9.7650 |
| cluster3845 | -604.91636 | 0.0001 | 0.000 | 0.232 | 0.0001 | 0.0000 | 0.0094 |
| cluster3846 | -1267.1619 | 0.0095 | 0.005 | 0.516 | 0.0831 | 0.0443 | 0.5329 |
| cluster3847 | -1156.3874 | 0.0008 | 0.001 | 0.735 | 0.0013 | 0.0049 | 3.7214 |
| cluster3848 | -2515.4167 | 0.0384 | 0.032 | 0.841 | 0.0179 | 0.0696 | 3.8856 |
| cluster3849 | -2657.8487 | 0.0231 | 0.037 | 1.581 | 0.0015 | 0.1037 | 68.1930 |
| cluster3850 | -1673.0346 | 0.0371 | 0.010 | 0.265 | 0.0082 | 0.0273 | 3.3292 |
| cluster3851 | -1618.1507 | 0.0092 | 0.005 | 0.525 | 0.0107 | 0.0294 | 2.7373 |
| cluster3852 | -1856.1344 | 0.0054 | 0.008 | 1.527 | 0.0104 | 0.0221 | 2.1324 |
| cluster3853 | -1101.7644 | 0.0048 | 0.005 | 1.076 | 0.0049 | 0.0296 | 6.0494 |
| cluster3854 | -1358.096 | 0.0961 | 0.025 | 0.265 | 0.1138 | 0.1451 | 1.2749 |
| cluster3855 | -2930.8015 | 0.0110 | 0.020 | 1.863 | 0.0085 | 0.0678 | 7.9346 |
| cluster3856 | -1696.502 | 0.0185 | 0.017 | 0.893 | 0.0185 | 0.0416 | 2.2515 |
| cluster3857 | -3037.6223 | 0.0152 | 0.012 | 0.788 | 0.0207 | 0.0512 | 2.4795 |
| cluster3858 | -2416.2402 | 0.0453 | 0.028 | 0.622 | 0.0639 | 0.0723 | 1.1304 |
| cluster3859 | -1977.5117 | 0.0122 | 0.006 | 0.480 | 0.0149 | 0.0282 | 1.8890 |
| cluster3860 | -2625.1794 | 0.0778 | 0.033 | 0.419 | 0.0993 | 0.0554 | 0.5581 |
| cluster3861 | -1301.0629 | 0.0172 | 0.014 | 0.809 | 0.0171 | 0.0328 | 1.9216 |
| cluster3862 | -922.33901 | 0.0001 | 0.000 | 0.361 | 0.0001 | 0.0002 | 1.7505 |

| | | | | | | | |
|---|---|---|---|---|---|---|---|
| cluster3863 | -1482.8116 | 0.0015 | 0.001 | 0.612 | 0.0001 | 0.0001 | 0.8993 |
| cluster3864 | -2219.0222 | 0.0030 | 0.003 | 0.872 | 0.0075 | 0.0260 | 3.4837 |
| cluster3865 | -3003.1582 | 0.0402 | 0.043 | 1.072 | 0.0637 | 0.1413 | 2.2162 |
| cluster3866 | -1633.3455 | 0.0189 | 0.017 | 0.897 | 0.0143 | 0.0719 | 5.0310 |
| cluster3867 | -608.47114 | 0.0001 | 0.000 | 0.038 | 0.0001 | 0.0000 | 0.1641 |
| cluster3868 | -1393.5966 | 0.0251 | 0.008 | 0.335 | 0.0325 | 0.0294 | 0.9049 |
| cluster3869 | -2975.4242 | 0.0329 | 0.054 | 1.637 | 0.0258 | 0.1566 | 6.0663 |
| cluster3870 | -2462.7321 | 0.0405 | 0.024 | 0.596 | 0.0389 | 0.0796 | 2.0461 |
| cluster3871 | -2151.7445 | 0.0095 | 0.006 | 0.657 | 0.0046 | 0.0127 | 2.7772 |
| cluster3872 | -2649.5872 | 0.0127 | 0.017 | 1.339 | 0.0109 | 0.0444 | 4.0920 |
| cluster3873 | -1785.1236 | 0.0365 | 0.018 | 0.504 | 0.0114 | 0.0227 | 1.9829 |
| cluster3874 | -2096.033 | 0.0039 | 0.001 | 0.186 | 0.0287 | 0.0175 | 0.6116 |
| cluster3875 | -768.99507 | 0.0001 | 0.000 | 0.353 | 0.0064 | 0.0071 | 1.1124 |
| cluster3876 | -742.59339 | 0.0026 | 0.001 | 0.299 | 0.0325 | 0.0221 | 0.6800 |
| cluster3877 | -1428.3095 | 0.0044 | 0.001 | 0.309 | 0.0454 | 0.0307 | 0.6762 |
| cluster3879 | -1881.3317 | 0.0164 | 0.009 | 0.526 | 0.0205 | 0.0400 | 1.9508 |
| cluster3880 | -1357.6031 | 0.0189 | 0.013 | 0.677 | 0.0240 | 0.0644 | 2.6816 |
| cluster3881 | -3016.492 | 0.0520 | 0.046 | 0.892 | 0.0219 | 0.1626 | 7.4139 |
| cluster3882 | -2689.7806 | 0.0219 | 0.019 | 0.877 | 0.0322 | 0.0629 | 1.9513 |
| cluster3883 | -422.06529 | 0.0007 | 0.027 | 39.253 | 0.2144 | 0.4026 | 1.8782 |
| cluster3884 | -1563.5757 | 0.0003 | 0.008 | 26.864 | 0.0004 | 0.0383 | 88.6011 |
| cluster3885 | -2856.5926 | 0.0150 | 0.012 | 0.791 | 0.0025 | 0.0268 | 10.5624 |
| cluster3886 | -1395.0099 | 0.0031 | 0.007 | 2.425 | 0.0039 | 0.0255 | 6.5514 |
| cluster3887 | -1621.6115 | 0.0142 | 0.017 | 1.218 | 0.0006 | 0.0592 | 106.9742 |
| cluster3888 | -1182.5274 | 0.0115 | 0.021 | 1.870 | 0.0054 | 0.1050 | 19.5282 |
| cluster3889 | -1004.6065 | 0.0224 | 0.018 | 0.816 | 0.0283 | 0.1090 | 3.8481 |
| cluster3890 | -1871.8973 | 0.0166 | 0.020 | 1.196 | 0.0198 | 0.0491 | 2.4838 |
| cluster3891 | -2184.1791 | 0.0128 | 0.010 | 0.791 | 0.0243 | 0.0189 | 0.7803 |
| cluster3892 | -1374.3905 | 0.0160 | 0.005 | 0.303 | 0.0128 | 0.0143 | 1.1135 |
| cluster3893 | -1458.32 | 0.0218 | 0.011 | 0.502 | 0.0395 | 0.0261 | 0.6617 |
| cluster3894 | -966.59869 | 0.0077 | 0.003 | 0.400 | 0.0087 | 0.0685 | 7.9098 |
| cluster3895 | -1982.5126 | 0.0082 | 0.004 | 0.541 | 0.0138 | 0.0507 | 3.6842 |
| cluster3896 | -820.68933 | 0.0031 | 0.001 | 0.419 | 0.0033 | 0.0187 | 5.6380 |
| cluster3897 | -1445.532 | 0.0020 | 0.051 | 25.802 | 0.0237 | 0.0228 | 0.9600 |
| cluster3898 | -2390.0323 | 0.0220 | 0.006 | 0.270 | 0.0324 | 0.0136 | 0.4200 |
| cluster3899 | -3003.2458 | 0.0279 | 0.035 | 1.247 | 0.0261 | 0.1034 | 3.9653 |
| cluster3900 | -2364.7398 | 0.0668 | 0.076 | 1.143 | 0.0890 | 0.2660 | 2.9879 |
| cluster3901 | -1543.8442 | 0.0030 | 0.003 | 0.863 | 0.0014 | 0.0046 | 3.3875 |
| cluster3902 | -2494.5927 | 0.0145 | 0.014 | 0.962 | 0.0054 | 0.0307 | 5.6540 |
| cluster3903 | -1622.709 | 0.0095 | 0.008 | 0.856 | 0.0237 | 0.1203 | 5.0835 |
| cluster3904 | -905.22436 | 0.0064 | 0.003 | 0.399 | 0.0042 | 0.0615 | 14.8015 |
| cluster3905 | -1594.1045 | 0.1162 | 0.054 | 0.463 | 0.1842 | 0.1903 | 1.0331 |
| cluster3906 | -1448.656 | 0.0112 | 0.005 | 0.463 | 0.0489 | 0.0595 | 1.2167 |
| cluster3907 | -1944.9642 | 0.0436 | 0.024 | 0.558 | 0.0360 | 0.0636 | 1.7642 |

| | | | | | | | |
|---|---|---|---|---|---|---|---|
| cluster3908 | -1848.1677 | 0.0135 | 0.008 | 0.621 | 0.0155 | 0.0444 | 2.8575 |
| cluster3909 | -2464.8093 | 0.0273 | 0.023 | 0.849 | 0.0148 | 0.1076 | 7.2593 |
| cluster3910 | -1193.0944 | 0.0326 | 0.027 | 0.825 | 0.0512 | 0.1349 | 2.6343 |
| cluster3911 | -1749.7095 | 0.0092 | 0.006 | 0.620 | 0.0050 | 0.0408 | 8.2143 |
| cluster3912 | -951.76626 | 0.0001 | 0.000 | 0.307 | 0.0001 | 0.0000 | 0.3168 |
| cluster3913 | -2544.1475 | 0.0347 | 0.023 | 0.650 | 0.0532 | 0.1230 | 2.3130 |
| cluster3914 | -2238.7516 | 0.0544 | 0.052 | 0.952 | 0.0833 | 0.1763 | 2.1177 |
| cluster3915 | -2227.3831 | 0.0356 | 0.027 | 0.771 | 0.0315 | 0.0804 | 2.5562 |
| cluster3916 | -902.68444 | 0.0779 | 0.040 | 0.508 | 0.0839 | 0.1004 | 1.1971 |
| cluster3917 | -2582.8624 | 0.0113 | 0.005 | 0.474 | 0.0188 | 0.0592 | 3.1590 |
| cluster3918 | -2754.8756 | 0.0216 | 0.034 | 1.592 | 0.0039 | 0.1252 | 31.7378 |
| cluster3919 | -569.95031 | 0.0022 | 0.001 | 0.365 | 0.0001 | 0.0000 | 0.2281 |
| cluster3920 | -657.20718 | 0.0155 | 0.063 | 4.085 | 0.0026 | 0.2254 | 85.8655 |
| cluster3921 | -2184.4977 | 0.0309 | 0.018 | 0.569 | 0.0006 | 0.0555 | 99.8358 |
| cluster3922 | -1240.1631 | 0.0543 | 0.025 | 0.454 | 0.0429 | 0.0925 | 2.1551 |
| cluster3923 | -1150.4267 | 0.0005 | 0.001 | 1.138 | 0.0003 | 0.0024 | 8.2872 |
| cluster3924 | -1696.2953 | 0.0196 | 0.012 | 0.610 | 0.0201 | 0.0543 | 2.7054 |
| cluster3925 | -1091.6989 | 0.0616 | 0.036 | 0.582 | 0.0201 | 0.0441 | 2.1960 |
| cluster3926 | -2433.6355 | 0.0389 | 0.059 | 1.517 | 0.1147 | 0.1855 | 1.6166 |
| cluster3927 | -1187.8828 | 0.0094 | 0.006 | 0.637 | 0.0162 | 0.0412 | 2.5449 |
| cluster3928 | -1156.832 | 0.0148 | 0.004 | 0.279 | 0.0541 | 0.0552 | 1.0198 |
| cluster3929 | -1607.5321 | 0.0048 | 0.004 | 0.910 | 0.0095 | 0.0264 | 2.7700 |
| cluster3930 | -2016.4799 | 0.0187 | 0.009 | 0.473 | 0.0072 | 0.0064 | 0.8980 |
| cluster3931 | -1808.1318 | 0.2244 | 0.044 | 0.195 | 0.0001 | 0.0000 | 0.3297 |
| cluster3932 | -1117.8711 | 0.0102 | 0.002 | 0.176 | 0.0125 | 0.0068 | 0.5422 |
| cluster3933 | -1155.9655 | 0.0571 | 0.011 | 0.199 | 0.1499 | 0.1068 | 0.7125 |
| cluster3934 | -717.10761 | 0.0001 | 0.000 | 0.178 | 0.0001 | 0.0002 | 1.6488 |
| cluster3935 | -3108.7091 | 0.0615 | 0.053 | 0.859 | 0.0464 | 0.1970 | 4.2498 |
| cluster3936 | -2132.4571 | 0.0210 | 0.019 | 0.923 | 0.0030 | 0.0504 | 17.1094 |
| cluster3937 | -1938.557 | 0.0179 | 0.010 | 0.537 | 0.0286 | 0.0192 | 0.6692 |
| cluster3938 | -1781.528 | 0.0270 | 0.011 | 0.403 | 0.0432 | 0.0532 | 1.2298 |
| cluster3939 | -2055.9257 | 0.0148 | 0.010 | 0.643 | 0.0174 | 0.0497 | 2.8585 |
| cluster3940 | -1338.2916 | 0.0158 | 0.006 | 0.378 | 0.0254 | 0.0141 | 0.5546 |
| cluster3941 | -966.79098 | 0.0055 | 0.001 | 0.114 | 0.0488 | 0.0441 | 0.9040 |
| cluster3942 | -2009.5149 | 0.0297 | 0.012 | 0.392 | 0.0350 | 0.0411 | 1.1729 |
| cluster3943 | -2268.0356 | 0.0168 | 0.016 | 0.962 | 0.0062 | 0.1006 | 16.2333 |
| cluster3944 | -1590.0694 | 0.0575 | 0.035 | 0.613 | 0.0163 | 0.0651 | 4.0072 |
| cluster3945 | -1676.2538 | 0.0349 | 0.021 | 0.617 | 0.0521 | 0.1185 | 2.2768 |
| cluster3946 | -1805.9734 | 0.0237 | 0.013 | 0.534 | 0.0301 | 0.0824 | 2.7408 |
| cluster3947 | -2282.5506 | 0.0367 | 0.029 | 0.777 | 0.0122 | 0.0596 | 4.8862 |
| cluster3948 | -1015.358 | 0.0248 | 0.011 | 0.451 | 0.0423 | 0.0431 | 1.0174 |
| cluster3949 | -876.26308 | 0.0060 | 0.003 | 0.431 | 0.0373 | 0.0810 | 2.1696 |
| cluster3950 | -1740.4659 | 0.0133 | 0.019 | 1.415 | 0.0042 | 0.0415 | 9.8639 |
| cluster3951 | -2833.9963 | 0.0414 | 0.031 | 0.749 | 0.0322 | 0.0956 | 2.9671 |

| | | | | | | | |
|---|---|---|---|---|---|---|---|
| cluster3952 | -1474.9259 | 0.0033 | 0.003 | 0.929 | 0.0028 | 0.0155 | 5.5794 |
| cluster3953 | -1359.9906 | 0.0062 | 0.002 | 0.371 | 0.0342 | 0.0454 | 1.3279 |
| cluster3954 | -1806.4621 | 0.0257 | 0.013 | 0.499 | 0.0331 | 0.0513 | 1.5498 |
| cluster3955 | -1192.1348 | 0.0237 | 0.018 | 0.739 | 0.0299 | 0.0897 | 3.0028 |
| cluster3956 | -1995.7417 | 0.0150 | 0.013 | 0.889 | 0.0130 | 0.0585 | 4.4812 |
| cluster3957 | -1959.2288 | 0.0133 | 0.012 | 0.920 | 0.0093 | 0.0512 | 5.5334 |
| cluster3958 | -1104.2839 | 0.0192 | 0.008 | 0.399 | 0.0357 | 0.0813 | 2.2799 |
| cluster3959 | -2685.1681 | 0.0306 | 0.020 | 0.644 | 0.0539 | 0.0338 | 0.6261 |
| cluster3960 | -2405.7745 | 0.0924 | 0.042 | 0.452 | 0.0033 | 0.1312 | 39.9148 |
| cluster3961 | -1594.9233 | 0.0397 | 0.025 | 0.620 | 0.0340 | 0.1005 | 2.9526 |
| cluster3962 | -1533.3698 | 0.0426 | 0.024 | 0.558 | 0.0389 | 0.0815 | 2.0986 |
| cluster3963 | -2077.9979 | 0.0375 | 0.029 | 0.766 | 0.0479 | 0.0824 | 1.7199 |
| cluster3964 | -2997.6195 | 0.0739 | 0.118 | 1.593 | 0.0173 | 0.3491 | 20.1593 |
| cluster3965 | -1513.5018 | 0.0402 | 0.035 | 0.878 | 0.0013 | 0.1229 | 92.7279 |
| cluster3966 | -1432.7757 | 0.0209 | 0.009 | 0.426 | 0.0295 | 0.0394 | 1.3367 |
| cluster3967 | -1753.9034 | 0.0876 | 0.027 | 0.311 | 0.0860 | 0.0561 | 0.6522 |
| cluster3968 | -1167.7101 | 0.0171 | 0.011 | 0.666 | 0.0182 | 0.0302 | 1.6640 |
| cluster3969 | -899.56818 | 0.0001 | 0.000 | 0.175 | 0.0001 | 0.0001 | 0.6675 |
| cluster3970 | -2483.5123 | 0.0508 | 0.035 | 0.684 | 0.0259 | 0.1263 | 4.8683 |
| cluster3971 | -1690.2232 | 0.0356 | 0.021 | 0.592 | 0.0432 | 0.1075 | 2.4922 |
| cluster3972 | -986.62807 | 0.0210 | 0.013 | 0.605 | 0.0016 | 0.0238 | 15.2152 |
| cluster3973 | -1418.5947 | 0.0143 | 0.002 | 0.125 | 0.0021 | 0.0259 | 12.5775 |
| cluster3974 | -1832.7548 | 0.0168 | 0.011 | 0.642 | 0.0202 | 0.0370 | 1.8287 |
| cluster3975 | -1270.801 | 0.0315 | 0.021 | 0.658 | 0.0199 | 0.0299 | 1.5072 |
| cluster3976 | -1581.0975 | 0.0098 | 0.009 | 0.916 | 0.0192 | 0.0266 | 1.3884 |
| cluster3977 | -2230.1391 | 0.0957 | 0.054 | 0.559 | 0.1023 | 0.1584 | 1.5489 |
| cluster3978 | -886.18612 | 0.0119 | 0.003 | 0.253 | 0.0017 | 0.0030 | 1.7510 |
| cluster3979 | -1620.7161 | 0.0028 | 0.002 | 0.612 | 0.0010 | 0.0019 | 1.9127 |
| cluster3980 | -922.24867 | 0.0001 | 0.000 | 0.157 | 0.0001 | 0.0000 | 0.2931 |
| cluster3981 | -1980.8899 | 0.0194 | 0.011 | 0.575 | 0.0250 | 0.0608 | 2.4337 |
| cluster3982 | -1829.827 | 0.0515 | 0.029 | 0.566 | 0.0436 | 0.0964 | 2.2133 |
| cluster3983 | -2676.6869 | 0.0314 | 0.036 | 1.161 | 0.0482 | 0.1501 | 3.1133 |
| cluster3984 | -1382.2901 | 0.0434 | 0.045 | 1.046 | 0.0268 | 0.1610 | 6.0077 |
| cluster3985 | -978.98239 | 0.0097 | 0.003 | 0.342 | 0.0174 | 0.0117 | 0.6749 |
| cluster3986 | -1734.303 | 0.0168 | 0.018 | 1.091 | 0.0171 | 0.0800 | 4.6795 |
| cluster3987 | -2086.0542 | 0.0099 | 0.007 | 0.659 | 0.0213 | 0.0410 | 1.9237 |
| cluster3988 | -1239.6289 | 0.0163 | 0.007 | 0.431 | 0.0001 | 0.0002 | 1.7581 |
| cluster3989 | -1834.938 | 0.0576 | 0.046 | 0.793 | 0.0467 | 0.0641 | 1.3746 |
| cluster3990 | -2208.4612 | 0.0257 | 0.024 | 0.919 | 0.0178 | 0.1071 | 6.0228 |
| cluster3991 | -1521.5405 | 0.0179 | 0.017 | 0.972 | 0.0283 | 0.0623 | 2.2037 |
| cluster3992 | -1889.167 | 0.0362 | 0.017 | 0.476 | 0.0398 | 0.0199 | 0.5004 |
| cluster3993 | -1673.785 | 0.0403 | 0.026 | 0.639 | 0.0184 | 0.0871 | 4.7229 |
| cluster3994 | -2393.873 | 0.0662 | 0.049 | 0.747 | 0.0355 | 0.0772 | 2.1774 |
| cluster3995 | -1619.4484 | 0.0375 | 0.023 | 0.620 | 0.0335 | 0.0682 | 2.0379 |

| | | | | | | | |
|---|---|---|---|---|---|---|---|
| cluster3996 | -1706.6574 | 0.0040 | 0.008 | 1.952 | 0.0113 | 0.0389 | 3.4268 |
| cluster3997 | -2516.1961 | 0.0635 | 0.044 | 0.691 | 0.0205 | 0.0947 | 4.6250 |
| cluster3998 | -1415.5445 | 0.0636 | 0.034 | 0.527 | 0.0328 | 0.0509 | 1.5544 |
| cluster3999 | -2524.5947 | 0.0349 | 0.029 | 0.828 | 0.0244 | 0.1083 | 4.4415 |
| cluster4000 | -2290.0452 | 0.0505 | 0.028 | 0.556 | 0.0383 | 0.0721 | 1.8850 |
| cluster4001 | -1733.8657 | 0.0248 | 0.018 | 0.713 | 0.0023 | 0.0236 | 10.0996 |
| cluster4002 | -1801.3379 | 0.0060 | 0.008 | 1.259 | 0.0016 | 0.0385 | 24.6089 |
| cluster4003 | -1421.6473 | 0.0789 | 0.044 | 0.554 | 0.0537 | 0.1371 | 2.5507 |
| cluster4004 | -1442.6386 | 0.0095 | 0.004 | 0.458 | 0.0061 | 0.0140 | 2.2919 |
| cluster4005 | -2064.2857 | 0.0608 | 0.054 | 0.888 | 0.0136 | 0.1235 | 9.0518 |
| cluster4006 | -1088.4005 | 0.0236 | 0.010 | 0.436 | 0.0479 | 0.0654 | 1.3667 |
| cluster4007 | -1425.9151 | 0.0029 | 0.001 | 0.488 | 0.0022 | 0.0039 | 1.7849 |
| cluster4008 | -1353.8888 | 0.0060 | 0.006 | 0.929 | 0.0223 | 0.0580 | 2.6047 |
| cluster4009 | -1541.7148 | 0.0151 | 0.018 | 1.218 | 0.0005 | 0.0623 | 123.6060 |
| cluster4010 | -1625.5785 | 0.0696 | 0.063 | 0.902 | 0.0249 | 0.1460 | 5.8631 |
| cluster4011 | -1060.1549 | 0.0541 | 0.009 | 0.170 | 0.1345 | 0.0214 | 0.1591 |
| cluster4012 | -2446.865 | 0.0160 | 0.106 | 6.617 | 0.0133 | 0.0977 | 7.3698 |
| cluster4013 | -846.28473 | 0.0191 | 0.007 | 0.347 | 0.0325 | 0.0265 | 0.8159 |
| cluster4014 | -1137.4544 | 0.0079 | 0.004 | 0.478 | 0.0028 | 0.0201 | 7.0938 |
| cluster4015 | -617.71293 | 0.0151 | 0.001 | 0.058 | 0.1416 | 0.0035 | 0.0250 |
| cluster4016 | -1815.8581 | 0.0261 | 0.019 | 0.711 | 0.0003 | 0.0117 | 43.6185 |
| cluster4017 | -1341.3786 | 0.0492 | 0.035 | 0.708 | 0.0126 | 0.0819 | 6.5133 |
| cluster4018 | -1324.0335 | 0.1853 | 0.111 | 0.597 | 0.0615 | 0.1821 | 2.9613 |
| cluster4019 | -1824.4416 | 0.0761 | 0.049 | 0.650 | 0.0159 | 0.1677 | 10.5242 |
| cluster4020 | -1004.6939 | 0.0299 | 0.010 | 0.343 | 0.0589 | 0.0337 | 0.5714 |
| cluster4021 | -1592.815 | 0.0225 | 0.007 | 0.310 | 0.0339 | 0.0375 | 1.1060 |
| cluster4022 | -969.5245 | 0.0038 | 0.001 | 0.176 | 0.0138 | 0.0256 | 1.8563 |
| cluster4023 | -1844.1617 | 0.0709 | 0.034 | 0.477 | 0.0522 | 0.0982 | 1.8824 |
| cluster4024 | -2359.788 | 0.0306 | 0.039 | 1.272 | 0.0274 | 0.1249 | 4.5617 |
| cluster4025 | -624.22565 | 0.0001 | 0.000 | 0.282 | 0.0008 | 0.0075 | 9.4721 |
| cluster4026 | -1088.1743 | 0.0014 | 0.001 | 0.545 | 0.0073 | 0.0319 | 4.3523 |
| cluster4027 | -978.63952 | 0.0087 | 0.003 | 0.365 | 0.0016 | 0.0508 | 32.7856 |
| cluster4028 | -1846.6197 | 0.0051 | 0.008 | 1.481 | 0.0101 | 0.0409 | 4.0401 |
| cluster4029 | -2250.2936 | 0.0774 | 0.061 | 0.787 | 0.0439 | 0.1497 | 3.4095 |
| cluster4030 | -1125.11 | 0.0376 | 0.011 | 0.303 | 0.0365 | 0.0668 | 1.8319 |
| cluster4031 | -1093.2949 | 0.0194 | 0.012 | 0.600 | 0.1559 | 0.1733 | 1.1114 |
| cluster4032 | -1474.5576 | 0.0001 | 0.000 | 0.252 | 0.0028 | 0.0020 | 0.6997 |
| cluster4033 | -2214.999 | 0.0748 | 0.063 | 0.838 | 0.0569 | 0.1714 | 3.0151 |
| cluster4034 | -493.6726 | 0.0001 | 0.000 | 0.984 | 0.0001 | 0.0000 | 0.0000 |
| cluster4035 | -2114.2962 | 0.0678 | 0.033 | 0.481 | 0.0354 | 0.0309 | 0.8741 |
| cluster4036 | -1076.2279 | 0.0063 | 0.004 | 0.560 | 0.0209 | 0.0389 | 1.8646 |
| cluster4037 | -1625.1667 | 0.0214 | 0.018 | 0.849 | 0.0080 | 0.0629 | 7.9018 |
| cluster4038 | -901.53938 | 0.0210 | 0.026 | 1.257 | 0.0071 | 0.0549 | 7.7158 |
| cluster4039 | -1507.8143 | 0.0511 | 0.009 | 0.176 | 0.0767 | 0.0653 | 0.8509 |

| | | | | | | | |
|---|---|---|---|---|---|---|---|
| cluster4040 | -1126.401 | 0.0248 | 0.017 | 0.678 | 0.0387 | 0.0561 | 1.4482 |
| cluster4041 | -1474.792 | 0.0174 | 0.015 | 0.855 | 0.0023 | 0.0393 | 17.0749 |
| cluster4042 | -896.85015 | 0.0396 | 0.016 | 0.408 | 0.0006 | 0.0419 | 70.2534 |
| cluster4043 | -1138.6042 | 0.0450 | 0.073 | 1.631 | 0.0439 | 0.2115 | 4.8222 |
| cluster4044 | -957.37375 | 0.0007 | 0.001 | 0.992 | 0.0026 | 0.0085 | 3.2724 |
| cluster4045 | -2022.9279 | 0.0573 | 0.040 | 0.707 | 0.0911 | 0.1822 | 2.0002 |
| cluster4046 | -891.83568 | 0.0001 | 0.000 | 0.295 | 0.0038 | 0.0053 | 1.3829 |
| cluster4047 | -2068.4668 | 0.0207 | 0.025 | 1.212 | 0.0103 | 0.0521 | 5.0869 |
| cluster4048 | -1470.4043 | 0.0078 | 0.002 | 0.273 | 0.0091 | 0.0103 | 1.1355 |
| cluster4049 | -572.58299 | 0.0015 | 0.005 | 3.178 | 0.0001 | 0.0003 | 3.3499 |
| cluster4050 | -1214.7895 | 0.0041 | 0.002 | 0.536 | 0.0180 | 0.0231 | 1.2823 |
| cluster4051 | -1402.6209 | 0.0038 | 0.001 | 0.239 | 0.0176 | 0.0097 | 0.5514 |
| cluster4052 | -1287.9608 | 0.0177 | 0.009 | 0.532 | 0.0280 | 0.0133 | 0.4758 |
| cluster4053 | -1642.7949 | 0.0166 | 0.018 | 1.113 | 0.0009 | 0.0747 | 83.2007 |
| cluster4054 | -1858.8635 | 0.0145 | 0.022 | 1.545 | 0.0357 | 0.0765 | 2.1419 |
| cluster4055 | -719.65086 | 0.0029 | 0.007 | 2.393 | 0.0014 | 0.0445 | 31.4732 |
| cluster4056 | -934.72234 | 0.0344 | 0.016 | 0.455 | 0.0371 | 0.0726 | 1.9562 |
| cluster4057 | -1565.1128 | 0.0335 | 0.023 | 0.687 | 0.0379 | 0.0928 | 2.4500 |
| cluster4058 | -1678.7145 | 0.0495 | 0.029 | 0.579 | 0.0084 | 0.1083 | 12.9214 |
| cluster4059 | -719.26364 | 0.0384 | 0.019 | 0.487 | 0.0016 | 0.0328 | 20.8874 |
| cluster4060 | -1875.6807 | 0.0300 | 0.017 | 0.567 | 0.0227 | 0.0556 | 2.4462 |
| cluster4061 | -1185.704 | 0.0089 | 0.011 | 1.197 | 0.0012 | 0.0139 | 11.9369 |
| cluster4062 | -1212.5377 | 0.0146 | 0.010 | 0.680 | 0.0104 | 0.0246 | 2.3660 |
| cluster4063 | -1723.0021 | 0.0247 | 0.033 | 1.352 | 0.0273 | 0.1398 | 5.1215 |
| cluster4064 | -783.03635 | 0.0144 | 0.009 | 0.608 | 0.0217 | 0.0420 | 1.9337 |
| cluster4065 | -988.40498 | 0.0041 | 0.002 | 0.417 | 0.0154 | 0.0174 | 1.1341 |
| cluster4066 | -1616.3544 | 0.0506 | 0.028 | 0.545 | 0.0302 | 0.1010 | 3.3490 |
| cluster4067 | -418.08865 | 0.0254 | 0.040 | 1.570 | 0.1250 | 0.1887 | 1.5093 |
| cluster4068 | -1353.5568 | 0.0173 | 0.024 | 1.380 | 0.0370 | 0.1719 | 4.6511 |
| cluster4069 | -1064.763 | 0.0690 | 0.027 | 0.388 | 0.0472 | 0.0502 | 1.0630 |
| cluster4070 | -702.3702 | 0.0014 | 0.013 | 9.604 | 0.0017 | 0.0682 | 39.9535 |
| cluster4071 | -825.89014 | 0.0006 | 0.001 | 1.323 | 0.0041 | 0.0171 | 4.2138 |
| cluster4072 | -986.76139 | 0.0332 | 0.016 | 0.469 | 0.0113 | 0.0486 | 4.3133 |
| cluster4073 | -1889.6722 | 0.0328 | 0.028 | 0.858 | 0.0259 | 0.0926 | 3.5760 |
| cluster4074 | -1257.2057 | 0.0339 | 0.038 | 1.132 | 0.0290 | 0.1234 | 4.2632 |
| cluster4075 | -710.4342 | 0.7224 | 3.630 | 5.025 | 0.9278 | 0.0393 | 0.0424 |
| cluster4076 | -1650.0561 | 0.0834 | 0.040 | 0.474 | 0.1131 | 0.0712 | 0.6293 |
| cluster4077 | -1680.0288 | 0.0448 | 0.024 | 0.533 | 0.0415 | 0.1077 | 2.5952 |
| cluster4078 | -1231.5289 | 0.0187 | 0.018 | 0.977 | 0.0486 | 0.0456 | 0.9369 |
| cluster4079 | -1205.1706 | 0.0001 | 0.000 | 0.846 | 0.0038 | 0.0049 | 1.3153 |
| cluster4080 | -736.9408 | 0.0155 | 0.023 | 1.478 | 0.0780 | 0.0036 | 0.0467 |
| cluster4081 | -1130.0669 | 0.0135 | 0.007 | 0.495 | 0.0304 | 0.0405 | 1.3337 |
| cluster4082 | -743.23569 | 0.0111 | 0.077 | 6.973 | 0.0057 | 0.0043 | 0.7599 |
| cluster4083 | -1706.3291 | 0.0612 | 0.057 | 0.928 | 0.0436 | 0.1631 | 3.7377 |

| | | | | | | | |
|---|---|---|---|---|---|---|---|
| cluster4084 | -715.98369 | 0.0173 | 0.004 | 0.235 | 0.0113 | 0.0199 | 1.7539 |
| cluster4085 | -859.2586 | 0.0311 | 0.011 | 0.352 | 0.0110 | 0.0021 | 0.1932 |
| cluster4086 | -1312.3485 | 0.0365 | 0.027 | 0.740 | 0.0082 | 0.1023 | 12.4257 |
| cluster4087 | -1205.8278 | 0.0106 | 0.015 | 1.404 | 0.0160 | 0.0731 | 4.5560 |
| cluster4088 | -1311.6997 | 0.0365 | 0.008 | 0.214 | 0.0993 | 0.0705 | 0.7098 |
| cluster4089 | -1510.9473 | 0.0073 | 0.012 | 1.642 | 0.0029 | 0.0308 | 10.5277 |
| cluster4090 | -755.72798 | 0.0370 | 0.015 | 0.418 | 0.0352 | 0.0427 | 1.2115 |
| cluster4091 | -952.73061 | 0.0195 | 0.005 | 0.235 | 0.0505 | 0.0347 | 0.6859 |
| cluster4092 | -1425.0194 | 0.0265 | 0.024 | 0.892 | 0.0275 | 0.1141 | 4.1517 |
| cluster4093 | -1006.9426 | 0.0186 | 0.014 | 0.743 | 0.0166 | 0.0578 | 3.4863 |
| cluster4094 | -374.90135 | 0.0001 | 0.000 | 0.082 | 0.0163 | 0.0062 | 0.3820 |
| cluster4095 | -1404.6555 | 0.0411 | 0.051 | 1.234 | 0.0496 | 0.1052 | 2.1216 |
| cluster4096 | -1213.113 | 0.0232 | 0.021 | 0.906 | 0.0245 | 0.0929 | 3.7864 |
| cluster4097 | -705.06892 | 0.1056 | 0.015 | 0.138 | 0.0001 | 0.0000 | 0.0309 |
| cluster4098 | -1867.348 | 0.0349 | 0.039 | 1.112 | 0.0006 | 0.0406 | 63.1852 |
| cluster4099 | -809.33941 | 0.0049 | 0.003 | 0.641 | 0.0005 | 0.0031 | 6.6578 |
| cluster4100 | -1047.2822 | 0.0050 | 0.004 | 0.721 | 0.0154 | 0.0302 | 1.9606 |
| cluster4101 | -2088.092 | 0.2119 | 0.095 | 0.451 | 0.4018 | 0.2459 | 0.6120 |
| cluster4102 | -1856.0767 | 0.0515 | 0.112 | 2.176 | 0.0177 | 0.2511 | 14.1624 |
| cluster4103 | -1104.8894 | 0.0266 | 0.005 | 0.196 | 0.0836 | 0.0328 | 0.3928 |
| cluster4104 | -1017.9907 | 0.0081 | 0.006 | 0.714 | 0.0062 | 0.0326 | 5.2955 |
| cluster4105 | -735.23365 | 0.0146 | 0.055 | 3.783 | 0.0057 | 0.0182 | 3.1795 |
| cluster4106 | -1000.8644 | 0.0087 | 0.008 | 0.893 | 0.0060 | 0.0290 | 4.8759 |
| cluster4107 | -786.67551 | 0.0018 | 0.002 | 0.989 | 0.0035 | 0.0073 | 2.1272 |
| cluster4108 | -780.85104 | 0.0073 | 0.012 | 1.619 | 999.0000 | 0.0336 | 0.0000 |
| cluster4109 | -1651.8309 | 0.0698 | 0.054 | 0.771 | 0.0604 | 0.2299 | 3.8089 |
| cluster4110 | -1246.4817 | 0.0637 | 0.028 | 0.437 | 0.0592 | 0.2041 | 3.4504 |
| cluster4111 | -923.41889 | 0.0525 | 0.029 | 0.546 | 0.0024 | 0.1861 | 76.5358 |
| cluster4112 | -1113.3942 | 0.0162 | 0.007 | 0.455 | 0.0163 | 0.0288 | 1.7730 |
| cluster4113 | -773.0448 | 0.0147 | 0.001 | 0.047 | 0.0048 | 0.0271 | 5.6256 |
| cluster4114 | -1073.9734 | 0.0332 | 0.013 | 0.398 | 0.0650 | 0.0699 | 1.0764 |
| cluster4115 | -1155.3081 | 0.0136 | 0.010 | 0.705 | 0.0032 | 0.0402 | 12.6715 |
| cluster4116 | -635.26696 | 0.0654 | 0.060 | 0.915 | 0.0079 | 0.0441 | 5.6098 |
| cluster4117 | -660.56395 | 0.0107 | 0.002 | 0.177 | 0.0153 | 0.0261 | 1.7084 |
| cluster4118 | -1288.4938 | 0.0301 | 0.031 | 1.032 | 0.0297 | 0.0829 | 2.7892 |
| cluster4119 | -926.26572 | 0.0209 | 0.013 | 0.626 | 0.0134 | 0.0812 | 6.0428 |
| cluster4120 | -967.39765 | 0.0067 | 0.003 | 0.477 | 0.0004 | 0.0006 | 1.6222 |
| cluster4121 | -1217.5636 | 0.0040 | 0.016 | 4.062 | 0.0056 | 0.0599 | 10.6717 |
| cluster4122 | -526.93116 | 0.0759 | 0.002 | 0.024 | 0.0160 | 0.0036 | 0.2246 |
| cluster4123 | -467.91826 | 0.0033 | 0.001 | 0.320 | 0.0134 | 0.0203 | 1.5135 |
| cluster4124 | -905.08209 | 0.0030 | 0.003 | 1.146 | 0.0086 | 0.0557 | 6.4852 |
| cluster4125 | -1064.6267 | 0.0611 | 0.053 | 0.871 | 0.2085 | 0.1175 | 0.5635 |
| cluster4126 | -761.09182 | 0.0522 | 0.010 | 0.195 | 0.1427 | 0.0839 | 0.5876 |
| cluster4127 | -764.23799 | 0.0382 | 0.016 | 0.413 | 0.1081 | 0.1185 | 1.0956 |

| | | | | | | | |
|---|---|---|---|---|---|---|---|
| cluster4128 | -989.41963 | 0.0320 | 0.054 | 1.674 | 0.0540 | 0.2317 | 4.2893 |

| Adjusted p value |
| --- |
| 0.03 |
| 5.47E-65 |
| 0.0005 |
| 1.78E-92 |
| 2.96E-07 |
| 5.19E-06 |
| 4.13E-54 |
| 4.41E-09 |
| 5.89E-124 |
| 1.56E-41 |
| 3.85E-88 |
| 6.50E-44 |
| 1.28E-107 |
| 1.62E-12 |
| 1.19E-48 |
| 7.57E-28 |
| 2.20E-08 |
| 6.40E-06 |
| 3.65E-06 |
| 1.01E-10 |
| 6.38E-60 |
| 7.60E-22 |
| 1.35E-64 |
| 5.64E-11 |
| 8.49E-21 |
| 4.55E-12 |
| 0.005 |
| 1.65E-17 |
| 6.65E-19 |
| 2.24E-27 |
| 1.91E-07 |
| 1.28E-33 |
| 1.73E-32 |
| 6.16E-25 |
| 5.71E-55 |
| 6.62E-08 |
| 1.16E-46 |
| 0.06 |
| 1.61E-24 |
| 1.78E-34 |
| 3.07E-15 |
| 4.24E-17 |

| |
|---|
| 4.91E-19 |
| 5.76E-66 |
| 2.14E-22 |
| 1.93E-14 |
| 1.62E-29 |
| 1.48E-39 |
| 1.62E-20 |
| 1.87E-13 |
| 1.35E-45 |
| 4.78E-05 |
| 4.06E-19 |
| 4.51E-16 |
| 3.68E-11 |
| 4.94E-14 |
| 6.73E-05 |
| 2.42E-20 |
| 4.70E-10 |
| 4.69E-24 |
| 4.62E-41 |
| 5.41E-25 |
| 4.44E-19 |
| 1.57E-26 |
| 2.73E-30 |
| 0.0001 |
| 3.45E-13 |
| 3.72E-30 |
| 6.52E-27 |
| 1.30E-09 |
| 0.07 |
| 6.56E-33 |
| 1.52E-14 |
| 4.81E-41 |
| 1.80E-11 |
| 9.89E-25 |
| 3.21E-52 |
| 1.03E-09 |
| 2.94E-10 |
| 1.46E-12 |
| 2.84E-14 |
| 6.49E-12 |
| 3.91E-14 |
| 5.05E-24 |
| 1.29E-12 |
| 3.95E-29 |

| |
|---|
| 1.15E-06 |
| 8.06E-15 |
| 1.33E-06 |
| 1.04E-16 |
| 1.62E-23 |
| 9.20E-06 |
| 3.88E-21 |
| 2.65E-15 |
| 3.77E-21 |
| 4.58E-16 |
| 2.68E-23 |
| 2.88E-26 |
| 1.50E-10 |
| 4.44E-22 |
| 1.16E-14 |
| 2.71E-18 |
| 8.08E-16 |
| 2.86E-38 |
| 3.51E-06 |
| 9.58E-48 |
| 2.93E-09 |
| 4.73E-36 |
| 7.78E-47 |
| 7.37E-21 |
| 2.08E-14 |
| 1.02E-13 |
| 4.71E-32 |
| 5.12E-18 |
| 3.57E-40 |
| 1.08E-21 |
| 1.12E-15 |
| 3.34E-30 |
| 2.18E-05 |
| 7.10E-17 |
| 1.45E-05 |
| 6.77E-09 |
| 5.37E-11 |
| 0.48 |
| 4.80E-34 |
| 3.51E-57 |
| 9.30E-19 |
| 2.24E-05 |
| 5.65E-11 |
| 6.27E-05 |

| |
|---|
| 3.18E-26 |
| 1.37E-12 |
| 0.24 |
| 0.01 |
| 1.10E-13 |
| 1.01E-19 |
| 3.21E-13 |
| 4.63E-14 |
| 8.59E-08 |
| 5.77E-12 |
| 6.06E-17 |
| 2.96E-10 |
| 0.10 |
| 4.51E-16 |
| 6.02E-11 |
| 3.97E-27 |
| 9.99E-16 |
| 3.38E-18 |
| 2.16E-16 |
| 4.01E-22 |
| 1.89E-06 |
| 3.58E-05 |
| 4.22E-05 |
| 6.46E-10 |
| 1.92E-23 |
| 1.06E-16 |
| 1.09E-09 |
| 0.01 |
| 1.06E-16 |
| 2.33E-07 |
| 1.31E-06 |
| 1.31E-24 |
| 0.0002 |
| 7.77E-09 |
| 3.43E-20 |
| 4.35E-32 |
| 1.10E-11 |
| 4.55E-15 |
| 2.13E-10 |
| 0.0002 |
| 9.42E-20 |
| 0.02 |
| 1.29E-12 |
| 0.14 |

| |
|---|
| 1.39E-30 |
| 2.89E-12 |
| 8.88E-15 |
| 3.50E-08 |
| 5.28E-07 |
| 3.41E-22 |
| 0.14 |
| 0.005 |
| 7.33E-25 |
| 4.16E-08 |
| 7.10E-07 |
| 3.80E-07 |
| 2.65E-11 |
| 5.66E-14 |
| 3.63E-15 |
| 5.55E-12 |
| 5.70E-09 |
| 1.43E-16 |
| 7.54E-11 |
| 0.02 |
| 9.88E-09 |
| 3.63E-17 |
| 4.54E-31 |
| 2.10E-14 |
| 5.79E-14 |
| 1.39E-13 |
| 1.20E-07 |
| 1.27E-32 |
| 3.05E-08 |
| 9.83E-05 |
| 0.05 |
| 1.59E-05 |
| 2.09E-06 |
| 1.25E-11 |
| 2.36E-17 |
| 7.35E-10 |
| 1.12E-16 |
| 2.83E-05 |
| 0.02 |
| 4.98E-08 |
| 3.86E-11 |
| 0.003 |
| 0.001 |
| 2.92E-13 |

| |
|---|
| 2.06E-17 |
| 2.25E-07 |
| 9.45E-15 |
| 1.07E-13 |
| 7.30E-13 |
| 0.001 |
| 2.60E-16 |
| 2.63E-16 |
| 6.31E-20 |
| 9.09E-21 |
| 3.45E-18 |
| 5.24E-10 |
| 3.78E-11 |
| 0.002 |
| 1.47E-20 |
| 0.002 |
| 1.75E-20 |
| 8.22E-09 |
| 0.0001 |
| 2.86E-28 |
| 3.22E-07 |
| 3.55E-14 |
| 4.63E-16 |
| 6.69E-21 |
| 9.37E-27 |
| 2.10E-14 |
| 9.33E-19 |
| 3.20E-11 |
| 4.80E-09 |
| 1.53E-09 |
| 2.20E-08 |
| 1.72E-12 |
| 2.00E-09 |
| 3.94E-15 |
| 7.10E-17 |
| 2.76E-23 |
| 2.84E-05 |
| 0.01 |
| 2.28E-11 |
| 0.0003 |
| 1.74E-11 |
| 5.74E-10 |
| 2.17E-10 |
| 3.73E-10 |

| |
|---|
| 4.35E-14 |
| 1.22E-06 |
| 1.42E-12 |
| 0.01 |
| 5.49E-06 |
| 6.62E-16 |
| 2.78E-18 |
| 2.12E-10 |
| 5.82E-15 |
| 4.69E-14 |
| 3.69E-05 |
| 8.58E-16 |
| 3.59E-09 |
| 5.50E-10 |
| 8.96E-06 |
| 3.94E-14 |
| 2.73E-10 |
| 1.37E-15 |
| 2.22E-08 |
| 2.50E-05 |
| 2.98E-12 |
| 3.04E-09 |
| 1.49E-11 |
| 9.88E-09 |
| 3.78E-06 |
| 1.56E-05 |
| 3.89E-08 |
| 0.18 |
| 7.96E-12 |
| 5.83E-12 |
| 7.76E-51 |
| 2.89E-13 |
| 6.70E-27 |
| 3.83E-14 |
| 6.76E-10 |
| 8.68E-05 |
| 5.20E-06 |
| 2.04E-13 |
| 1.02E-08 |
| 1.94E-27 |
| 3.63E-17 |
| 2.60E-16 |
| 0.02 |
| 7.16E-16 |

| |
|---|
| 2.83E-10 |
| 0.11 |
| 6.02E-11 |
| 9.32E-11 |
| 2.40E-08 |
| 3.35E-09 |
| 6.78E-11 |
| 5.20E-16 |
| 8.62E-05 |
| 4.97E-30 |
| 2.83E-14 |
| 0.02 |
| 5.11E-08 |
| 5.79E-17 |
| 1.37E-20 |
| 5.04E-09 |
| 9.60E-07 |
| 2.32E-07 |
| 0.0004 |
| 4.28E-41 |
| 2.93E-05 |
| 3.48E-07 |
| 1.06E-24 |
| 3.96E-09 |
| 0.0001 |
| 6.75E-09 |
| 7.06E-18 |
| 4.01E-06 |
| 9.34E-18 |
| 0.001 |
| 1.13E-10 |
| 6.87E-14 |
| 8.54E-05 |
| 4.36E-10 |
| 8.71E-07 |
| 3.59E-09 |
| 8.51E-18 |
| 6.49E-06 |
| 1.29E-07 |
| 1.47E-20 |
| 1.50E-11 |
| 2.23E-19 |
| 0.003 |
| 8.11E-11 |

| |
|---|
| 0.0001 |
| 0.003 |
| 4.29E-05 |
| 1.03E-09 |
| 6.44E-26 |
| 1.31E-09 |
| 2.16E-19 |
| 2.35E-21 |
| 1.04E-09 |
| 1.06E-12 |
| 0.0001 |
| 9.46E-12 |
| 0.0003 |
| 1.50E-10 |
| 1.08E-07 |
| 4.15E-14 |
| 1.27E-23 |
| 4.21E-06 |
| 6.17E-14 |
| 8.10E-06 |
| 0.002 |
| 8.58E-06 |
| 5.71E-17 |
| 2.22E-08 |
| 4.07E-06 |
| 6.18E-10 |
| 3.68E-09 |
| 1.58E-14 |
| 1.81E-11 |
| 1.51E-18 |
| 4.16E-08 |
| 3.23E-07 |
| 5.33E-12 |
| 3.93E-12 |
| 3.00E-05 |
| 0.001 |
| 2.53E-05 |
| 2.26E-19 |
| 1.26E-07 |
| 3.78E-09 |
| 0.01 |
| 2.05E-10 |
| 1.45E-21 |
| 2.72E-11 |

| |
|---|
| 0.04 |
| 1.03E-12 |
| 6.80E-14 |
| 0.01 |
| 1.35E-07 |
| 5.92E-08 |
| 2.08E-05 |
| 6.77E-09 |
| 2.90E-20 |
| 3.98E-13 |
| 1.16E-29 |
| 3.32E-08 |
| 1.34E-05 |
| 2.13E-18 |
| 2.34E-05 |
| 3.18E-22 |
| 0.07 |
| 2.50E-05 |
| 7.79E-31 |
| 0.0001 |
| 8.13E-11 |
| 4.07E-07 |
| 9.96E-06 |
| 5.83E-06 |
| 4.99E-16 |
| 0.73 |
| 0.004 |
| 1.43E-05 |
| 1.32E-08 |
| 0.02 |
| 4.99E-05 |
| 3.41E-17 |
| 1.18E-13 |
| 3.03E-11 |
| 5.29E-08 |
| 9.94E-12 |
| 1.56E-07 |
| 7.92E-15 |
| 2.80E-07 |
| 0.001 |
| 2.14E-06 |
| 1.26E-08 |
| 1.06E-09 |
| 0.001 |

| |
|---|
| 5.31E-12 |
| 1.01E-06 |
| 1.79E-19 |
| 1.47E-10 |
| 1.61E-10 |
| 2.37E-10 |
| 0.14 |
| 7.72E-22 |
| 8.83E-07 |
| 3.45E-15 |
| 0.001 |
| 2.53E-05 |
| 1.49E-11 |
| 2.85E-07 |
| 3.50E-05 |
| 8.48E-06 |
| 3.70E-12 |
| 1.16E-18 |
| 0.02 |
| 3.04E-15 |
| 6.97E-10 |
| 1.88E-08 |
| 1.66E-07 |
| 1.04E-10 |
| 1.51E-21 |
| 1.71E-05 |
| 1.40E-06 |
| 0.02 |
| 4.37E-10 |
| 0.0001 |
| 1.90E-09 |
| 1.96E-07 |
| 8.60E-10 |
| 0.001 |
| 4.20E-08 |
| 0.06 |
| 6.75E-13 |
| 9.32E-07 |
| 3.36E-05 |
| 8.10E-06 |
| 0.002 |
| 4.21E-09 |
| 2.79E-11 |
| 0.04 |

| |
|---|
| 5.68E-09 |
| 3.24E-13 |
| 1.32E-05 |
| 3.19E-12 |
| 6.89E-05 |
| 0.45 |
| 1.57E-08 |
| 3.70E-09 |
| 5.60E-24 |
| 4.21E-08 |
| 1.43E-15 |
| 5.82E-23 |
| 1.82E-07 |
| 1.64E-06 |
| 2.65E-17 |
| 9.69E-09 |
| 3.32E-11 |
| 1.44E-05 |
| 1.30E-05 |
| 8.27E-07 |
| 1.27E-08 |
| 2.24E-09 |
| 0.0002 |
| 9.41E-26 |
| 2.65E-13 |
| 7.78E-06 |
| 6.66E-09 |
| 1.26E-15 |
| 7.79E-35 |
| 8.90E-11 |
| 9.88E-09 |
| 3.80E-07 |
| 0.01 |
| 0.001 |
| 1.23E-16 |
| 6.74E-05 |
| 1.57E-11 |
| 8.06E-18 |
| 3.31E-34 |
| 9.76E-06 |
| 2.90E-09 |
| 0.01 |
| 2.01E-15 |
| 6.83E-12 |

| |
|---|
| 0.0002 |
| 8.68E-07 |
| 9.55E-06 |
| 0.01 |
| 2.99E-07 |
| 5.45E-05 |
| 3.20E-13 |
| 2.78E-20 |
| 2.30E-07 |
| 1.92E-09 |
| 3.95E-09 |
| 1.39E-06 |
| 2.87E-11 |
| 1.39E-12 |
| 0.0001 |
| 1.77E-06 |
| 6.69E-13 |
| 1.24E-12 |
| 3.94E-15 |
| 5.82E-06 |
| 3.03E-07 |
| 0.14 |
| 1.13E-08 |
| 9.27E-15 |
| 3.03E-11 |
| 0.11 |
| 0.07 |
| 4.74E-15 |
| 3.04E-08 |
| 3.20E-06 |
| 2.91E-14 |
| 0.0005 |
| 4.97E-15 |
| 0.001 |
| 1.35E-05 |
| 0.13 |
| 2.60E-05 |
| 0.04 |
| 3.23E-10 |
| 0.0002 |
| 0.07 |
| 0.37 |
| 1.63E-20 |
| 6.87E-14 |

| |
|---|
| 3.59E-08 |
| 0.01 |
| 0.001 |
| 1.12E-15 |
| 0.001 |
| 0.0004 |
| 4.83E-13 |
| 7.73E-09 |
| 4.70E-10 |
| 3.23E-08 |
| 0.02 |
| 4.60E-07 |
| 3.08E-15 |
| 4.90E-12 |
| 7.21E-15 |
| 1.02E-08 |
| 0.002 |
| 6.30E-06 |
| 6.89E-06 |
| 8.13E-08 |
| 1.06E-12 |
| 2.50E-14 |
| 1.75E-11 |
| 5.09E-12 |
| 2.68E-11 |
| 9.73E-08 |
| 0.04 |
| 1.55E-09 |
| 2.70E-10 |
| 6.35E-07 |
| 0.0002 |
| 1.23E-15 |
| 4.31E-05 |
| 2.22E-08 |
| 5.54E-10 |
| 0.0003 |
| 1.75E-05 |
| 1.77E-12 |
| 0.0004 |
| 0.21 |
| 1.81E-08 |
| 1.68E-09 |
| 2.05E-18 |
| 2.45E-13 |

| |
|---|
| 6.78E-10 |
| 2.77E-11 |
| 3.25E-08 |
| 3.16E-16 |
| 1.07E-08 |
| 2.24E-15 |
| 5.68E-11 |
| 7.93E-11 |
| 9.73E-09 |
| 8.37E-12 |
| 7.23E-08 |
| 3.56E-13 |
| 0.0002 |
| 9.87E-11 |
| 0.001 |
| 0.03 |
| 1.14E-18 |
| 8.89E-09 |
| 6.61E-06 |
| 8.33E-05 |
| 7.07E-07 |
| 0.0003 |
| 6.12E-08 |
| 9.70E-12 |
| 4.15E-09 |
| 2.11E-08 |
| 9.93E-11 |
| 1.22E-10 |
| 2.89E-08 |
| 5.78E-11 |
| 0.02 |
| 5.65E-07 |
| 0.04 |
| 9.65E-09 |
| 2.96E-06 |
| 9.35E-10 |
| 0.001 |
| 7.02E-08 |
| 3.61E-05 |
| 8.11E-11 |
| 5.29E-09 |
| 8.03E-11 |
| 0.0001 |
| 0.34 |

| |
|---|
| 0.02 |
| 3.66E-05 |
| 0.41 |
| 2.13E-12 |
| 4.25E-12 |
| 5.03E-12 |
| 4.99E-10 |
| 4.46E-05 |
| 2.93E-05 |
| 0.26 |
| 3.78E-09 |
| 1.97E-10 |
| 3.06E-05 |
| 3.56E-06 |
| 7.59E-05 |
| 5.60E-24 |
| 1.79E-08 |
| 9.27E-08 |
| 7.03E-10 |
| 2.28E-05 |
| 6.45E-06 |
| 1.89E-07 |
| 4.08E-07 |
| 0.01 |
| 3.45E-10 |
| 6.41E-09 |
| 1.61E-24 |
| 0.004 |
| 4.45E-06 |
| 0.0002 |
| 4.63E-11 |
| 1.31E-06 |
| 3.48E-09 |
| 0.005 |
| 6.53E-10 |
| 1.02E-07 |
| 0.01 |
| 0.06 |
| 1.67E-08 |
| 0.003 |
| 0.001 |
| 7.02E-08 |
| 1.65E-07 |
| 4.32E-10 |

| |
|---|
| 1.03E-06 |
| 6.80E-14 |
| 0.08 |
| 5.40E-07 |
| 3.38E-08 |
| 1.15E-07 |
| 0.001 |
| 1.56E-10 |
| 1.70E-07 |
| 4.74E-07 |
| 1.69E-08 |
| 0.0002 |
| 0.01 |
| 1.10E-07 |
| 1.76E-06 |
| 4.01E-07 |
| 4.49E-07 |
| 4.07E-06 |
| 1.15E-11 |
| 4.87E-06 |
| 8.47E-09 |
| 3.65E-07 |
| 7.61E-06 |
| 6.86E-08 |
| 3.65E-08 |
| 6.87E-12 |
| 3.00E-07 |
| 0.02 |
| 2.78E-18 |
| 6.89E-10 |
| 0.004 |
| 3.75E-06 |
| 3.14E-05 |
| 1.72E-07 |
| 8.74E-07 |
| 0.0002 |
| 1.46E-10 |
| 2.49E-10 |
| 1.30E-07 |
| 0.0001 |
| 1.66E-08 |
| 1.07E-06 |
| 1.25E-11 |
| 8.49E-08 |

| |
|---|
| 5.49E-08 |
| 7.31E-08 |
| 1.20E-18 |
| 6.24E-14 |
| 1.16E-08 |
| 0.24 |
| 4.93E-07 |
| 5.53E-13 |
| 1.71E-06 |
| 3.51E-12 |
| 7.34E-10 |
| 8.59E-05 |
| 2.59E-05 |
| 3.02E-07 |
| 1.99E-16 |
| 2.50E-09 |
| 0.01 |
| 0.004 |
| 0.0003 |
| 0.32 |
| 5.17E-16 |
| 1.66E-07 |
| 1.50E-12 |
| 4.37E-10 |
| 3.62E-09 |
| 2.00E-10 |
| 2.36E-17 |
| 1.10E-06 |
| 1.34E-19 |
| 3.03E-08 |
| 1.35E-22 |
| 0.001 |
| 2.05E-14 |
| 3.85E-07 |
| 2.02E-06 |
| 0.002 |
| 0.02 |
| 3.47E-10 |
| 0.01 |
| 4.15E-06 |
| 0.002 |
| 9.70E-05 |
| 0.0002 |
| 9.98E-08 |

| |
|---|
| 0.003 |
| 8.82E-05 |
| 3.24E-06 |
| 5.88E-08 |
| 8.87E-25 |
| 2.63E-16 |
| 2.16E-06 |
| 0.001 |
| 3.44E-07 |
| 1.10E-11 |
| 5.46E-09 |
| 2.51E-16 |
| 0.002 |
| 9.87E-11 |
| 1.99E-11 |
| 0.25 |
| 3.00E-05 |
| 3.39E-11 |
| 0.01 |
| 3.24E-07 |
| 1.30E-05 |
| 1.69E-05 |
| 1.41E-14 |
| 1.03E-06 |
| 0.001 |
| 1.11E-11 |
| 4.39E-06 |
| 1.04E-08 |
| 2.13E-10 |
| 1.84E-05 |
| 2.89E-05 |
| 4.67E-05 |
| 0.01 |
| 5.06E-06 |
| 0.01 |
| 2.49E-08 |
| 0.0001 |
| 9.05E-14 |
| 1.50E-10 |
| 4.70E-12 |
| 0.02 |
| 4.76E-09 |
| 0.001 |
| 0.01 |

| |
|---|
| 0.04 |
| 5.23E-11 |
| 2.47E-06 |
| 2.20E-05 |
| 0.0002 |
| 8.00E-09 |
| 2.36E-17 |
| 0.002 |
| 2.03E-06 |
| 0.002 |
| 0.0002 |
| 7.38E-05 |
| 9.06E-10 |
| 9.91E-08 |
| 2.17E-06 |
| 2.45E-10 |
| 6.03E-05 |
| 1.53E-15 |
| 6.66E-09 |
| 2.28E-07 |
| 0.0002 |
| 0.0004 |
| 0.03 |
| 2.40E-08 |
| 0.03 |
| 2.61E-10 |
| 6.48E-07 |
| 0.003 |
| 1.07E-05 |
| 1.92E-07 |
| 0.001 |
| 2.60E-11 |
| 2.40E-06 |
| 1.98E-07 |
| 0.01 |
| 0.02 |
| 2.09E-05 |
| 4.20E-09 |
| 7.77E-11 |
| 0.0002 |
| 0.0003 |
| 1.92E-05 |
| 5.93E-12 |
| 0.02 |

| |
|---|
| 1.87E-13 |
| 2.58E-09 |
| 0.0004 |
| 5.99E-07 |
| 0.001 |
| 4.56E-07 |
| 9.70E-05 |
| 2.52E-19 |
| 0.001 |
| 4.12E-10 |
| 7.09E-05 |
| 0.001 |
| 1.15E-09 |
| 0.15 |
| 1.20E-08 |
| 1.63E-08 |
| 0.0002 |
| 1.26E-08 |
| 2.19E-14 |
| 4.88E-05 |
| 1.18E-19 |
| 1.22E-13 |
| 0.13 |
| 0.01 |
| 9.56E-12 |
| 2.68E-05 |
| 0.0001 |
| 0.0005 |
| 0.0004 |
| 4.36E-08 |
| 1.23E-05 |
| 5.70E-05 |
| 0.01 |
| 5.17E-05 |
| 3.39E-09 |
| 7.22E-08 |
| 0.001 |
| 0.01 |
| 4.52E-05 |
| 0.002 |
| 0.005 |
| 0.01 |
| 0.55 |
| 0.003 |

| |
|---|
| 0.001 |
| 0.001 |
| 1.65E-05 |
| 1.08E-06 |
| 9.12E-10 |
| 1.41E-09 |
| 0.0001 |
| 1.31E-05 |
| 8.62E-14 |
| 3.52E-06 |
| 9.49E-09 |
| 8.48E-10 |
| 3.98E-05 |
| 0.11 |
| 1.22E-10 |
| 1.56E-08 |
| 0.02 |
| 1.95E-08 |
| 4.30E-06 |
| 1.93E-09 |
| 5.74E-06 |
| 0.15 |
| 0.0001 |
| 1.38E-13 |
| 0.0001 |
| 0.50 |
| 0.34 |
| 1.86E-06 |
| 6.49E-06 |
| 1.77E-06 |
| 1.93E-09 |
| 7.96E-09 |
| 1.47E-05 |
| 1.10E-05 |
| 7.59E-12 |
| 3.20E-13 |
| 0.07 |
| 2.13E-07 |
| 1.83E-06 |
| 0.11 |
| 2.87E-07 |
| 6.77E-09 |
| 0.001 |
| 0.001 |

| |
|---|
| 1.88E-10 |
| 8.01E-07 |
| 1.04E-07 |
| 6.97E-10 |
| 3.28E-14 |
| 3.06E-08 |
| 2.38E-08 |
| 0.002 |
| 6.42E-08 |
| 2.34E-05 |
| 5.34E-07 |
| 0.002 |
| 1.62E-06 |
| 1.04E-10 |
| 2.06E-09 |
| 3.59E-09 |
| 1.08E-05 |
| 1.59E-07 |
| 5.27E-07 |
| 4.46E-07 |
| 4.60E-07 |
| 1.21E-05 |
| 0.01 |
| 4.46E-05 |
| 6.51E-07 |
| 0.13 |
| 0.01 |
| 7.84E-06 |
| 1.27E-12 |
| 1.73E-07 |
| 3.65E-07 |
| 0.0003 |
| 5.24E-09 |
| 1.63E-13 |
| 2.68E-08 |
| 0.09 |
| 1.39E-16 |
| 2.62E-08 |
| 1.65E-14 |
| 3.88E-05 |
| 1.90E-08 |
| 0.004 |
| 6.42E-06 |
| 6.58E-07 |

| |
|---|
| 5.95E-10 |
| 3.75E-19 |
| 8.96E-06 |
| 1.33E-06 |
| 0.29 |
| 0.001 |
| 1.97E-06 |
| 5.51E-07 |
| 3.04E-13 |
| 7.96E-13 |
| 1.47E-05 |
| 5.21E-06 |
| 0.0002 |
| 0.002 |
| 1.31E-05 |
| 9.29E-08 |
| 0.0002 |
| 3.16E-05 |
| 7.81E-09 |
| 0.06 |
| 2.68E-11 |
| 1.51E-14 |
| 3.43E-10 |
| 5.45E-08 |
| 3.49E-06 |
| 0.001 |
| 3.87E-07 |
| 8.49E-11 |
| 0.001 |
| 5.08E-05 |
| 7.73E-05 |
| 1.03E-08 |
| 8.54E-06 |
| 3.45E-06 |
| 6.40E-06 |
| 1.04E-08 |
| 0.05 |
| 1.29E-14 |
| 8.97E-14 |
| 3.44E-07 |
| 0.0004 |
| 0.11 |
| 3.80E-05 |
| 7.55E-06 |

| |
|---|
| 1.67E-06 |
| 1.43E-07 |
| 0.002 |
| 3.65E-10 |
| 0.0001 |
| 5.83E-06 |
| 0.004 |
| 1.80E-06 |
| 0.08 |
| 7.99E-10 |
| 2.94E-08 |
| 0.10 |
| 0.02 |
| 2.67E-06 |
| 0.01 |
| 1.71E-13 |
| 2.43E-06 |
| 0.01 |
| 0.004 |
| 9.77E-07 |
| 0.001 |
| 2.43E-11 |
| 9.88E-09 |
| 0.14 |
| 0.01 |
| 0.12 |
| 0.001 |
| 0.0003 |
| 3.42E-08 |
| 2.21E-08 |
| 3.14E-05 |
| 3.03E-09 |
| 0.0001 |
| 0.03 |
| 1.31E-05 |
| 7.65E-10 |
| 0.0003 |
| 0.0002 |
| 4.84E-06 |
| 1.04E-07 |
| 0.003 |
| 0.001 |
| 8.49E-09 |
| 1.56E-07 |

| |
|---|
| 1.26E-11 |
| 9.77E-05 |
| 4.40E-05 |
| 0.0004 |
| 1.05E-09 |
| 2.76E-10 |
| 0.0004 |
| 2.53E-07 |
| 0.001 |
| 0.001 |
| 0.001 |
| 1.09E-05 |
| 0.03 |
| 0.0002 |
| 7.31E-05 |
| 0.03 |
| 4.36E-05 |
| 0.0005 |
| 0.0002 |
| 6.16E-05 |
| 2.76E-05 |
| 0.0004 |
| 5.23E-07 |
| 0.001 |
| 1.40E-06 |
| 0.02 |
| 0.0003 |
| 8.83E-13 |
| 0.01 |
| 1.18E-06 |
| 0.0004 |
| 2.68E-06 |
| 0.02 |
| 0.0001 |
| 1.54E-07 |
| 5.90E-10 |
| 4.37E-06 |
| 0.002 |
| 1.05E-07 |
| 1.72E-06 |
| 0.0002 |
| 0.0002 |
| 0.0004 |
| 0.0002 |

| |
|---|
| 4.90E-11 |
| 0.0001 |
| 0.003 |
| 4.38E-06 |
| 9.77E-06 |
| 4.71E-07 |
| 1.39E-12 |
| 0.01 |
| 1.04E-08 |
| 0.0002 |
| 0.001 |
| 7.50E-11 |
| 6.77E-09 |
| 6.88E-06 |
| 0.01 |
| 0.15 |
| 5.40E-06 |
| 0.002 |
| 3.65E-06 |
| 0.45 |
| 2.72E-08 |
| 0.002 |
| 0.25 |
| 0.01 |
| 0.01 |
| 3.98E-05 |
| 0.002 |
| 0.003 |
| 1.70E-05 |
| 3.68E-05 |
| 5.99E-07 |
| 0.0001 |
| 1.97E-12 |
| 0.002 |
| 0.003 |
| 0.004 |
| 1.23E-06 |
| 0.003 |
| 0.01 |
| 0.0005 |
| 1.31E-05 |
| 6.51E-06 |
| 8.05E-05 |
| 0.02 |

| |
|---|
| 0.01 |
| 9.26E-07 |
| 0.005 |
| 0.03 |
| 4.26E-11 |
| 0.01 |
| 0.0003 |
| 9.09E-10 |
| 3.43E-06 |
| 0.11 |
| 4.78E-06 |
| 4.31E-05 |
| 8.35E-06 |
| 3.63E-06 |
| 8.53E-08 |
| 7.64E-06 |
| 0.003 |
| 0.0001 |
| 0.002 |
| 0.02 |
| 4.78E-06 |
| 3.35E-09 |
| 1.32E-06 |
| 0.0002 |
| 0.0004 |
| 0.0001 |
| 0.001 |
| 0.14 |
| 0.0003 |
| 0.05 |
| 0.0003 |
| 2.40E-06 |
| 0.003 |
| 4.89E-06 |
| 0.001 |
| 1.00 |
| 0.02 |
| 0.003 |
| 5.47E-05 |
| 1.49E-10 |
| 0.01 |
| 0.0001 |
| 0.02 |
| 0.002 |

| |
|---|
| 0.02 |
| 0.15 |
| 0.0002 |
| 7.34E-06 |
| 0.02 |
| 1.01E-07 |
| 0.02 |
| 0.01 |
| 0.0003 |
| 0.0001 |
| 0.22 |
| 0.02 |
| 0.0001 |
| 0.50 |
| 1.53E-05 |
| 1.03E-06 |
| 1.51E-07 |
| 0.45 |
| 0.02 |
| 0.02 |
| 1.88E-07 |
| 0.004 |
| 3.93E-06 |
| 7.25E-07 |
| 1.04E-05 |
| 0.01 |
| 0.03 |
| 0.04 |
| 2.30E-05 |
| 0.03 |
| 0.01 |
| 1.78E-06 |
| 0.11 |
| 0.01 |
| 0.0001 |
| 0.0003 |
| 1.77E-06 |
| 4.76E-09 |
| 0.55 |
| 0.02 |
| 3.37E-07 |
| 0.06 |
| 0.19 |
| 0.03 |

0.02
